# Fibrillar Aβ causes profound microglial metabolic perturbations in a novel APP knock-in mouse model

**DOI:** 10.1101/2021.01.19.426731

**Authors:** Dan Xia, Steve Lianoglou, Thomas Sandmann, Meredith Calvert, Jung H. Suh, Elliot Thomsen, Jason Dugas, Michelle E. Pizzo, Sarah L. DeVos, Timothy K. Earr, Chia-Ching Lin, Sonnet Davis, Connie Ha, Hoang Nguyen, Roni Chau, Ernie Yulyaningsih, Hilda Solanoy, Shababa T. Masoud, Richard Liang, Karin Lin, Robert G. Thorne, Dylan Garceau, Jennifer D. Whitesell, Michael Sasner, Julie A. Harris, Kimberly Scearce-Levie, Joseph W. Lewcock, Gilbert Di Paolo, Pascal E. Sanchez

## Abstract

Microglial dysfunction is believed to play a pathogenic role in Alzheimer’s disease (AD). Here, we characterize the amyloid-β related pathology and microglial responses in an engineered APP knock-in mouse model of familial AD. This model recapitulates key pathological features of AD such as a progressive accumulation of parenchymal amyloid plaques and vascular amyloid deposits, altered glial responses and neurodegeneration. Leveraging multi-omics approaches, we found lipid accumulation and an exacerbated disease-associated transcriptomic response in methoxy-X04-positive, phagocytic microglia. Together, these findings highlight the potential of this novel, open-access mouse model to investigate AD pathogenesis and demonstrate that fibrillar Aβ triggers lipid dysregulation and immuno-metabolic perturbations in phagocytic microglia.

**Highlights:**

- Novel open-access APP KI mouse model shows salient AD pathological features
- Deep phenotyping of sorted microglia reveals profound lipidomic perturbations in line with Alois Alzheimer’s original descriptions of glial adipose inclusions
- Immunometabolic perturbations are exacerbated in microglia accumulating fibrillar Aβ

---

Alzheimer’s disease (AD) is a devastating and complex neurological condition. The production and detailed evaluation of new, optimized AD mouse models has been instrumental in gaining insight into the neurobiology of AD^1^. Over the last decade, emerging human genetics and preclinical research revealed that microglia, the resident immune cells of the central nervous system (CNS), are likely contributing to AD pathophysiology^2-4^. Microglia respond to various pathogenic drivers of AD, including amyloid-β (Aβ) and tau, but the mechanisms by which microglia may become dysfunctional and contribute to disease remain unclear. Microglia expression profiles from AD and preclinical mouse models thereof have revealed that microglia undergo profound changes in their gene expression profiles^2,5-7^. However, the functional implications of this molecular remodeling are poorly defined. It is hypothesized that these alterations in gene expression may enable microglia to contain and/or resorb AD pathologies such as amyloid deposits and associated downstream events. Notably, the phagocytic activity of microglia in this context is critical and the metabolic demand and burden on microglial degradation pathways are likely high, potentially leading to immunometabolic failure and increasing cellular dysfunction over time. Indeed, alterations of lipid metabolism have been observed in AD brain, and several genes associated with late onset AD (LOAD) risk that control lipid metabolism (*e.g., TREM2, APOE)^3,8^* are highly expressed in microglia. Here, we sought to determine if the pathogenic fibrillar form of Aβ causes molecular and metabolic dysregulation in microglia as a result of phagocytic uptake. To do so, we engineered and characterized an amyloid-precursor protein (APP) knock-in (KI) mouse model carrying familial AD (FAD) mutations and applied multi-omics approaches to deeply characterize molecular and cellular microglial responses.

To circumvent the numerous limitations inherent to transgenesis^9^, we used a knock-in strategy to humanize the Aβ sequence of the murine *App* gene and introduced three FAD mutations - Swedish (KM670/671NL), Arctic (E693G) and Austrian (T712I) - using homologous recombination (**Supplementary Fig. 1a**). This model, which we have named *App*^SAA^, differs from the *App*^NL-G-F^ knock-in mouse model^10^ in the FAD mutation present at the gamma secretase cleavage site (Austrian instead of Iberian/Beyreuther). Our biochemical analysis showed that levels of brain *App* mRNA and full-length protein were similar across the three genotypes (**Supplementary Fig. 1b, c** and **Fig. 1a**), but as anticipated, the levels of human APP and C-terminal APP fragments were elevated in a gene dose-dependent manner (**Supplementary Fig. 1c, d** and **Fig. 1b**). In the *App*^SAA^ homozygous mice (KI/KI), we observed a dramatic increase in the Aβ_42/40_ ratio in brain extracts (soluble and insoluble fractions), CSF, and plasma at both 2 and 4 months of age (**Fig. 1c, d** and **Supplementary Fig. 1e-j**). At 2 months of age, when amyloid plaques were not detectable yet, the increased Aβ_42/40_ ratio was driven by a reduction of Aβ_40_ levels. By contrast, at 4 months of age, when amyloid plaques were present, the increased ratio in the guanidine fraction of the brain was driven by an increase of Aβ_42_ levels in the *App*^SAA^ homozygous mice, which is consistent with the reported effects of the Austrian mutation on APP cleavage^11^.

**Figure 1.**
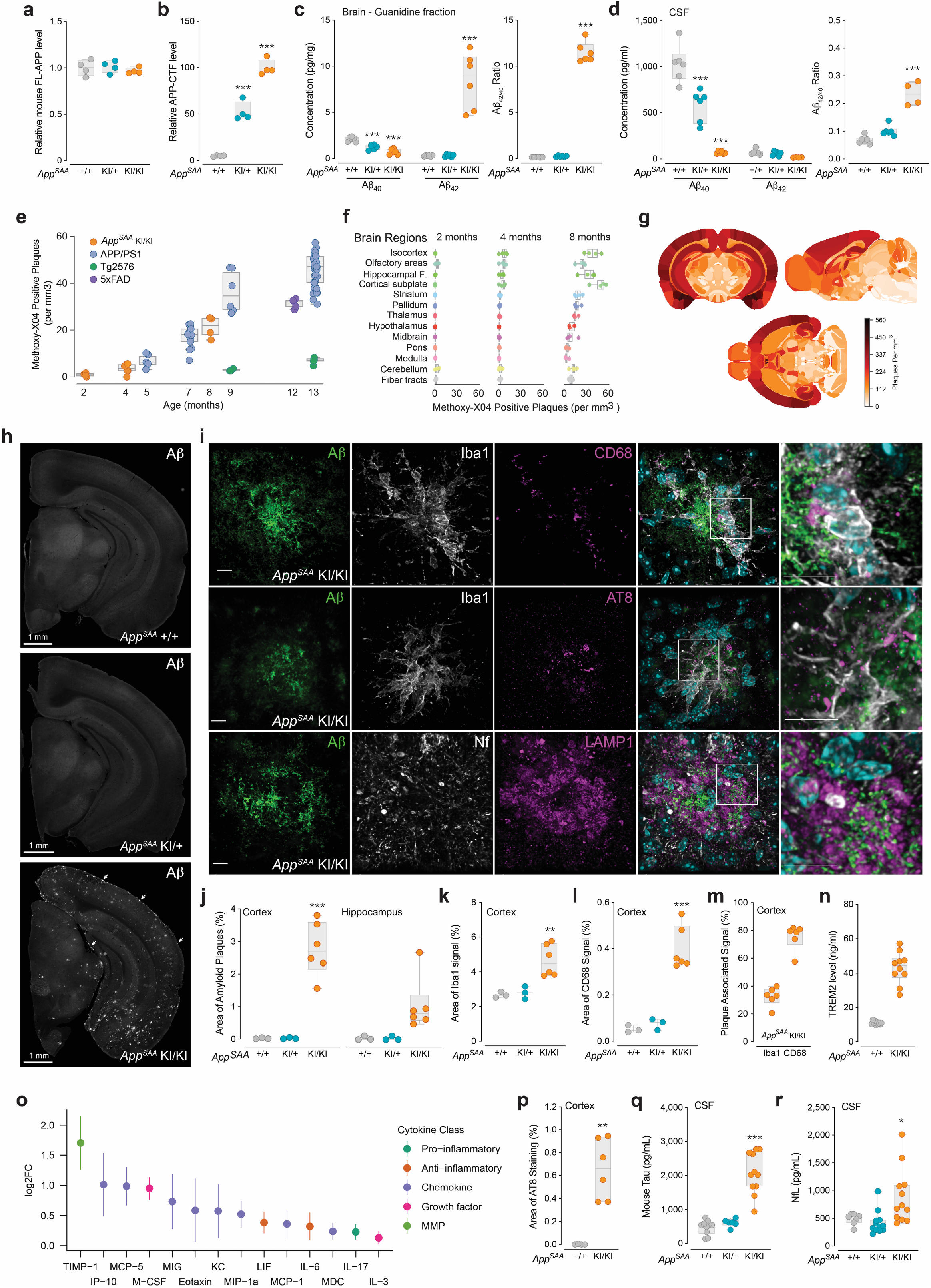
*App*^SAA^ knock-in mice display key features of amyloid-β associated pathologies. **(a-d)** Brain and CSF samples were analyzed in 2- and 4-month-old cohorts for the three genotypes of *App^SAA^* mice. Quantification of full-length (FL) murine APP **(a)** and the C-terminal fragment of APP **(b)** in brain lysates at 2 months of age by western blot analysis. **(c-d)** Aβ_40_ and Aβ42 concentrations were measured and the ratio of Aβ_42/40_ calculated in the guanidine fraction of the brain and in the CSF. **(e)** Brain-wide plaque density was measured by automated segmentation and registration to the CCFv3 atlas in four mouse lines at various ages. **(f)** Plaque density in major brain divisions in the *App*^SAA^ KI/KI mice at 2, 4 and 8 months of age. Box plots show the median and IQR, with whiskers extending to 1.5 times the IQR. Hippocampal F. = Hippocampal formation. Images and data from panels **(g)** to **(r)** are all from 8-month-old mice. **(g)** Whole brain 3D heatmaps showing the anatomical distribution of plaques in *App*^SAA^ KI/KI mice. **(h)** Representative images of brain sections showing staining for Aβ for the three genotypes of *App*^SAA^ mice (leptomeningeal CAA indicated by arrows). **(i)** Super-resolution confocal images showing Aβ plaques with microglia markers Iba1 and CD68, phospho-tau (AT8), neurofilament (Nf) and/or the lysosomal marker LAMP1; scale bars = 10 μm. Quantification of areas covered by Aβ plaques **(j)**, Iba1 **(k)**, CD68 **(l)**, percentage of the Iba1 and CD68 signals overlapping with plaques in *App*^SAA^ KI/KI mice **(m)**. **(n)** TREM2 levels measured in brain homogenates. **(o)** Differential abundance (log2) of cytokines extracted from the brains of *App*^SAA^ and WT mice. Bars represent 95% confidence intervals of fold change; positive values indicate higher abundance in *App*^SAA^ KI/KI. **(p)** Quantification of the area of phospho-tau (AT8) signal on sections. Total mouse tau **(q)** and Nf-L **(r)** measured in CSF. Graphs from panels a-d and j-n and p-r display means ± SEM and P values: one-way ANOVA with Dunnett’s multiple comparison test, each group compared to the *App*^SAA^ +/+ control group (n=4-6 per group); *P < 0.05, **P < 0.001, and ***P < 0.0001.

To analyze the brain-wide spatial distribution of Aβ deposition in *App*^SAA^ mice, we quantified methoxy-X04 labeled Aβ plaques using serial two-photon tomography imaging at 2, 4 and 8 months of age^12^. We found that the total brain density of Aβ plaques increased in an age-dependent manner in *App*^SAA^ homozygous mice. The amount of Aβ deposition resembled what has been reported for commonly used transgenic models, such as the APP/PS1 and the 5×FAD lines (**Fig. 1e**). At 8 months of age, Aβ plaques were detected in multiple brain regions, but the highest density of pathology was found in cortical and hippocampal regions (**Fig. 1f, g**). Comparing the temporal and spatial plaque deposition patterns in *App*^SAA^ homozygous mice with amyloid-based phases of human AD^12,13^ revealed remarkable similarities in plaque distribution (**Supplementary Fig. 2a-c**). We further confirmed the presence of amyloid plaques by Aβ immunolabeling in several brain regions of the *App*^SAA^ homozygous mice at 8 and 16 months of age (**Fig. 1h-j. Supplementary Fig. 3a**), and in the heterozygous mice at 16 months of age (**Supplementary Fig. 3a**). We also identified vascular Aβ deposition in *App*^SAA^ homozygous mice at 8 and 16 months of age, particularly in leptomeningeal (pial) vessels at the brain surface, demonstrating the presence of cerebral amyloid angiopathy (CAA) in this model (**Supplementary Fig. 3b-e**). Microglia density was drastically increased in the vicinity of amyloid plaques with an enrichment in CD68 positive microglia (**Fig. 1i, k-m** and **Supplementary Fig. 4a-d**). The levels of TREM2 and various cytokines were also elevated in brain lysates (**Fig. 1n, o**), showing profound immune responses in 8-month-old *App*^SAA^ homozygous mice. In addition, we found evidence of dystrophic neurites as shown by the accumulation of phosphorylated tau (AT8), neurofilament-light (Nf-L), and the lysosomal marker LAMP1 in aberrant (bulbous) shapes at the site of amyloid deposition (**Fig. 1i, p**). These characteristics resemble the Aβ pathology of human AD and were associated with an increase of total tau and Nf-L in the CSF (**Fig. 1q, r**), indicating presence of neurodegenerative processes in the *App*^SAA^ mouse model. Taken together with the observation that humanized wild-type APP knock-in rodent models do not develop amyloid pathology even with the combination of Presenilin 1 mutation^14^, these results suggest that the AD-relevant pathologies observed in our *App*^SAA^ mouse model are mediated by FAD mutations on *App*.

The presence of microglia clusters around amyloid plaques and the elevation of cytokines indicated that microglia were responding to the amyloid deposits in this model. To further characterize the glial response, we leveraged both transcriptomics and metabolomics approaches to profile microglia in 8-month-old *App*^SAA^ mice. First, we performed an RNA sequencing (RNA-seq) analysis of FACS-isolated microglia from wildtype (WT), *App*^SAA^ heterozygous, and *App*^SAA^ homozygous mice. We confirmed the enrichment and purity of isolated microglia by comparing the expression levels of previously reported marker genes from different cell types in the mouse brain^15^ (**Supplementary Fig. 5a**). We identified more than six hundred differentially expressed genes (DEGs) between cells from *App*^SAA^ homozygous and WT mice (absolute log2 fold change [FC] > 1.2, false discovery rate [FDR] < 10%), with an up regulation of disease associated microglia (DAM) genes observed in the *App*^SAA^ homozygous mice (**Fig. 2a**). Minimal expression changes were found in microglia from heterozygous animals at this age (**Fig. 2a**). Increased activity of other gene sets related to microglial state and function, such as cholesterol metabolism, glycolysis, phagocytic and lysosomal function resembled those observed in microglia from the 5×FAD transgenic mouse model^16,17^ (**Fig. 2b** and **Supplementary Fig. 5b**), demonstrating that some aspects of the microglia signature are conserved across diverse APP mouse models, including the *App*^SAA^ mouse model.

**Figure 2.**
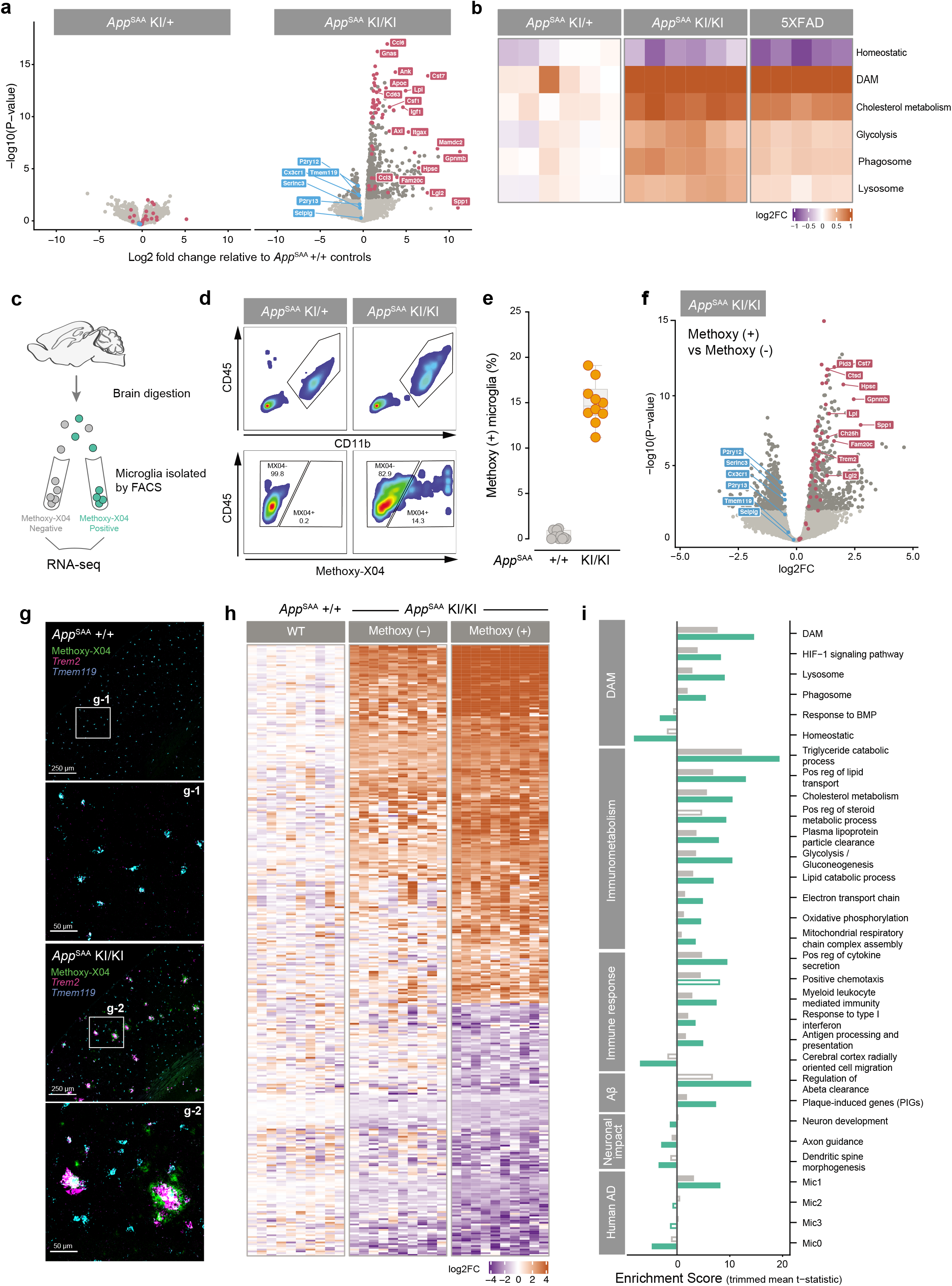
Transcriptome analysis of isolated microglia from *App*^SAA^ knock-in mice reveals shifts in immunometabolism driven by fibrillar Aβ. **(a)** Volcano plots showing log2 fold changes between *App^SAA^* heterozygous (left) and homozygous (right) knock-in compared to WT; dark grey, fuchsia, and blue indicate DEGs, DAM, and homeostatic genes, respectively. **(b)** Single sample activity scores for curated gene sets in *App*^SAA^ heterozygous, *App*^SAA^ homozygous, and 5×FAD. Intensities indicate log2 fold change of aggregated gene set score (row) per mouse (column) with respect to mean gene set score from matched WT mice. **(c, d)** Schematic of FACS experiment and gating strategy used to isolate pure populations of microglia that are negative (-) or positive (+) for methoxy-X04. **(e)** Fraction of methoxy (+) microglia purified from WT and *App*^SAA^ KI/KI mouse brains. **(f)** Volcano plot showing log2 fold changes between methoxy (+) and methoxy (-) expression profiles. Genes expressed higher in methoxy (+) microglia have log2 fold changes > 0; colors same as in (a). **(g)** Images of brain sections showing Methoxy-X04 labeling and detection of *trem2* and *tmem119* transcripts by *in situ* hybridization. **(h)** Gene expression profiles from a representative subset of differentially expressed genes in methoxy (-) and (+) *App*^SAA^ KI/KI microglia when compared to WT. Intensities correspond to the log2 foldchange for each gene (row) per sample (column) as compared to the mean expression of the gene in the methoxy (-) WT group. **(i)** Gene set enrichment analysis from methoxy (-) (grey) or methoxy (+) (light green) *App*^SAA^ KI/KI vs methoxy (-) WT mice. Enrichment scores are calculated as the mean t-statistic of genes in the leading edge of the gene set. Filled bars indicate gene set enrichment results with FDR <= 10%.

Next, we wondered if microglia at the vicinity of amyloid plaques might exhibit a different gene expression profile as compared to those distal from the plaques. To address this question, we hypothesized that most microglia close to amyloid plaques may have more phagocytic activity and therefore show intracellular accumulation of the fibrillar form of Aβ. We therefore labelled fibrillar Aβ in *App*^SAA^ mice by peripheral injection of methoxy-X04 and then sorted methoxy-X04-negative (-) and positive (+) microglia by FACS, prior to conducting RNA-seq analysis (**Fig. 2c-e**). Direct comparison of methoxy-X04 (+) vs (-) microglia from *App*^SAA^ homozygous mice identified more than 800 DEGs, with the methoxy-X04 (+) microglia showing increased DAM activation (**Fig. 2f**), as well as an induction of the plaque-induced gene (PIG) signature^7^ (**Supplementary Fig. 5c**). *In situ* analysis of the *Trem2* transcript, one of the DAM genes, shows high level of *Trem2* signal in methoxy-X04 positive amyloid plaques (**Fig. 2g**), which is consistent with an enrichment of a responsive microglial population clustering around amyloid plaques. Methoxy-X04 (-) microglia from *App*^SAA^ homozygous mice also showed significant changes compared to methoxy-X04 (-) microglia from WT control mice, but the effects were not as strong as the ones measured in *App*^SAA^ homozygous methoxy-X04 (+) microglia (**Fig. 2h**; **Supplementary Fig. 5c**). Gene set enrichment analysis (GSEA) revealed about twice as many significant pathways (FDR <= 10%) in the methoxy-X04 (+) as compared to the methoxy-X04 (-) microglia when compared against WT. Most notably, methoxy-X04 (+) cells exhibited higher enrichment for genes regulating innate immune function and metabolic functions that are critical for supporting lipid clearance and metabolism (**Fig 2i**). As anticipated, those cells also showed an enrichment in genes associated with regulation of Aβ clearance and amyloid plaques (PIGs). The reduced expression of genes involved in neuronal development, axon guidance and spine morphogenesis in the methoxy-X04 (+) population supports the notion that phagocytic microglia may not be able to sustain their neurotrophic roles. Generally, our data show that uptake of fibrillar Aβ by microglia may not only induce a generic DAM response, but also results in additional gene expression changes that reflect impact of fibrillar Aβ on microglial physiology. These data are also consistent with the evidence that fibrillar Aβ triggers major proteomic alterations in microglia that are associated with amyloid plaques^18^. Finally, the signature genes from the “Mic1” microglia sub-cluster in human AD^6^ were enriched in the phagocytic microglia from the *App*^SAA^ homozygous mice (**Fig 2i**), suggesting that this mouse model successfully recapitulates a subset of the microglia response found in human disease.

Next, we subjected FACS-isolated *App*^SAA^ microglia as well as microglia from wildtype animals to lipidomic and metabolomic analyses, as described before in non-amyloid models^17^. Using this approach, we identified significant alterations of the *App*^SAA^ microglia lipidome, including upregulation of the ganglioside GM3 (d36:1), various species of triglyceride (TG) (including species of arachidonate-containing TG) as well as a variety of phospholipid species (*i.e.*, PG, PS and PI) (**Fig. 3a,b**). Conversely, lysolipid lysophosphatidylinositol LPI 18:1 and coenzyme Q10 were downregulated in *App*^SAA^ microglia (**Fig. 3a,b**). Since phagocytic methoxy-X04 (+) microglia showed an increase in genes involved in metabolic functions compared to methoxy-X04 (-) microglia in *App*^SAA^ homozygous mice, we sought to directly measure levels of lipids and metabolites from those two microglia populations. Mirroring our transcriptomic analyses, lipid alterations were exacerbated in *App*^SAA^ methoxy-X04 (+) microglia relative to methoxy-X04 (-) and WT microglia, as shown for instance for GM3(d36:1) and TG 20:4/36:3 species (**Fig. 3c-f**). However, specific lipid changes appeared to be unique to methoxy-X04 (+) microglia, including an increase in cholesteryl ester CE 22:6 and a decrease in other CE species, such as CE 20:4 (**Fig. 3c-f**). Accumulation of GM3 is consistent with lysosomal storage defects and has been previously described in the brain of AD patients and mouse models^19^. Accumulation of TG and CE typically reflects metabolic imbalance and has been associated with pro-inflammatory states and neurotoxicity in other models^3,20^. Remarkably, these biochemical findings are consistent with Dr. Alzheimer’s original observations of adipose inclusions in glial cells neighboring amyloid plaques^21^. When taken along with a growing amount of human genetic data linking various genes involved in lipid metabolism to AD (*e.g., APOE4, ABCA7, CLU, TREM2*) and recent studies showing storage of neutral lipids in microglia from AD and aging models^3^, our data argues that dysregulation of lipid metabolism in microglia contributes to AD pathogenesis. Finally, metabolomic analyses of sorted microglia generally revealed subtle changes in several polar metabolites, with accumulation of the polyamine spermine standing out as the most dramatic change observed in methoxy-X04 (+) microglia (**Fig. 3e,f**). Spermine has been shown to modulate various cellular functions, including gene expression, ion channel function, response to oxidative stress, autophagy, metabolism and immune response. Spermine is critical as genetic ablation of spermine synthase (SMS) has been shown to cause oxidative stress and lysosomal impairment^22^. Additionally, spermine binds and neutralizes Aβ^23^ and is upregulated in PSAPP mouse^24^, suggesting that increase in spermine levels in methoxy-X04 (+) microglia may be a compensatory response to prevent the deleterious effects of Aβ. Therefore, our data lends further support to the notion that phagocytic microglia undergo profound cellular alterations, including lysosomal dysfunction, lipid dyshomeostasis, and other metabolic changes.

**Figure 3.**
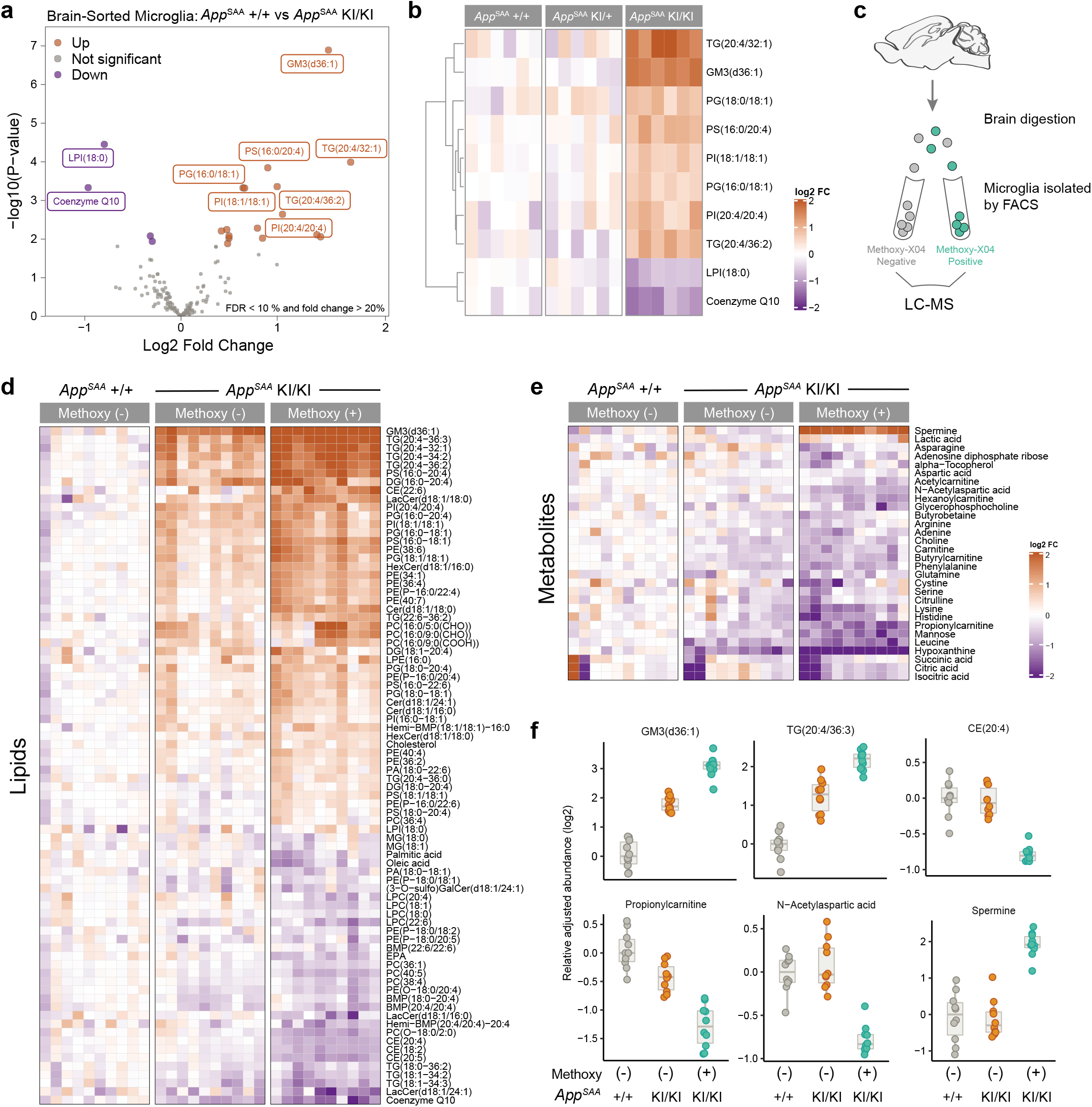
Microglia isolated from *App^SAA^* knock-in mice have profound metabolic dysregulation. **(a, b)** Lipidomics and metabolomics analysis was conducted in FACS-isolated microglia from brains of *App*^SAA^ KI/KI, *App*^SAA^ KI/+ and *App*^SAA^ +/+ control mice at 8 months of age (n=6 per group). **(a)** Volcano plot showing differential levels of lipids. FDR < 10% and absolute fold change > 20%. **(b)** Heatmap of top 10 lipids significantly altered by genotype in isolated microglia; columns represent individual mice. **(c)** Schematic of FACS experiment used to isolate pure populations of microglia that are negative (-) or positive (+) for methoxy-X04. Heatmap of **(d)** lipids and **(e)** metabolites significantly altered by genotype and/or presence of methoxy-X04 dye in microglia isolated from *App*^SAA^ KI/KI and *App*^SAA^ +/+ mouse brain (n=10 per group); columns represent individual mice. **(f)** Representative plots showing genotype and methoxy-X04 effects on key lipids and metabolites from FACS-isolated microglia. Statistical analysis was performed by fitting linear models using limma; FDR was calculated according to Benjamini’s and Hochberg’s method.

Altogether, our in-depth analysis of the *App*^SAA^ knock-in mouse model confirms emergence of disease-relevant biology and progressive accumulation of pathological hallmarks of AD. Since this new mouse model, together with control humanized rodent models^14^, can be used to investigate multiple relevant aspects of AD biology, we have made it open access to enable unrestricted pre-clinical research in the scientific community.

## Materials and Methods

### Mouse model generation

The *App*^SAA^ knock-in mouse model was engineered by insertion of 6 mutations into the genomic *App* locus via homologous recombination. Three amino acids APP G676R, APP F681Y and APP R684H were substituted to humanize the mouse Aβ_1-42_ region and the following three FAD-linked APP mutations were inserted: KM670/671NL (Swedish), E693G (Arctic) and T714I (Austrian). This mouse model was created on a C57BL/6J background by Ozgene (Australia) using goGermline technology^21^.

To generate the targeting vector for the *App*^SAA^ KI mouse model, six individual fragments (A-F) that cover the targeting area were engineered and introduced into six cloning vectors respectively. Fragment A encoding a 2828bp right homology arm fragment within intron 17 (mouse genomic DNA chr16:84,959,292-84,962,119 GRCm38/mm10) was amplified by PCR from BAC genomic DNA (clones RP23-126H12 and RP23-99P18) with primers P1915_41 and P1915_51. Fragment B encoding a portion of intron 17, exon 17 with the T714I and E693G mutations, a portion of intron 16 (corresponding to mouse genomic DNA region chr16:84,962,120-84,963,051 GRCm38/mm10) and a portion of the hygromycin cassette was synthesized as a gBlock by Integrated DNA Technologies. Fragment C encoding the remainder of the hygromycin cassette was amplified by PCR from an Ozgene in-house cloning vector using primers P1915_74 and P1915_53. Fragment D encoding a 2032bp portion of intron 16 (mouse genomic DNA region chr16:84,963,052-84,965,083 GRCm38/mm10) was synthesized as a gBlock by Integrated DNA Technologies. Fragment E encoding a portion of intron 16, exon 16 with the R684H, F681Y, G676R, and KM670/671NL mutations, and a portion of intron 15 (corresponding to mouse genomic DNA chr16:84,965,084-84,965,993 GRCm38/mm10) was synthesized by GENEWIZ and was housed within a vector that contained the neo cassette on the vector backbone. Fragment F encoding a 5870bp left homology arm within intron 15 (mouse genomic DNA chr16:84,965,994-84,971,863 GRCm38/mm10) was amplified by PCR from BAC genomic DNA (clones RP23-126H12 and RP23-99P18) with primers P1915_46 and P1915_56.

The final targeting vector was then generated by the sequential assembly of fragments A-F. Fragment A digested with AatII was ligated into vector B digested with the same enzyme to generate vector AB. Fragment AB digested with AgeI was ligated into fragment C digested with the same enzyme to generate vector ABC. Fragment ABC digested with AscI was ligated into vector D digested with MluI to generate vector ABCD. Fragment ABCD digested with AscI was ligated into vector E digested with the same enzymes to generate vector ABCDE. Fragment ABCDE digested with enzyme MluI was ligated into vector F digested with the same enzyme to generate vector ABCDEF.

The targeting vector containing Fragment ABCDEF was linearized with PmeI and electroporated into Bruce ES cells which were derived from C57BL/6-Thy1.1 mice. ES cells were maintained in the medium supplemented with G418 drug at 200ug/ml, and surviving clones were picked after 8 days of drug selection. ES cells were then screened for correct targeting events by qPCR and then confirmed by Southern analysis. For southern analysis, genomic DNA was purified from two ES clones (II_1C11 and II_1G11), digested with SpeI and then detected with 5’probe, 3’ probe, Hygro probe, enP probe, and neoP probe. Both clones were tested positive for proper homologous recombination in left and right arms. The ES cell clones (clone II_1C11 and clone II_1G11) that were confirmed to carry the correct homologous recombination events were injected into goGermline blastocysts, and the resulting chimeric mice were crossed to Flp transgenic mice (OzFlp) to excise the neo and hygro selection cassettes. The resulting heterozygous KI mice were backcrossed to C57BL/6J for more than two generations and then intercrossed to obtain homozygous KI mice for further characterization. The *App*^SAA^ model will be available from the Jackson Laboratory as B6(Cg)-Apptm1.1Dnli/J (https://www.jax.org/strain/034711). Primer sequences and assays for mouse model generation are listed in the **Supplementary Table 1**.

### Animal breeding and sample collection

Mice were maintained on the C57BL/6J genetic background and *App*^SAA^ heterozygous KI mice were intercrossed to obtain three genotypes of interest. Mice were bred either at the Jackson Laboratory (USA, Bar Harbor or Sacramento) or Ozgene (Australia, Perth) and then were shipped to Denali for an acclimation period of at least 2 weeks. Housing conditions included standard pellet food and water *ad libitum*, a 12-h light-dark cycle at 22°C with a maximum of 5 mice per cage. Housing conditions were similar between Denali Therapeutics, the Jackson Laboratory and Ozgene. Animals used for related studies are listed in the **Supplementary Table 2**. All mouse husbandry and experimental procedures were reviewed and approved by Denali Institutional Animal Care and Use Committee and were conducted in full compliance with regulatory statutes, Institutional Animal Care and Use Committee policies, and National Institutes of Health guidelines.

### Tissue collection

Prior to tissue harvest, animals were deeply anesthetized via intraperitoneal (i.p.) injection of 2.5% Avertin. All collected tissues were snap-frozen on dry ice and stored at −80°C. Blood samples were collected by cardiac puncture into an EDTA tube (Sarstedt Microvette 500 K3E, Ref# 201341102), then slowly inverted 10 times prior to centrifugation at 12,700 rpm for 7 minutes at 4°C to collect plasma. CSF samples were collected by pre-pulled glass capillary tube from the cisterna magna and then transferred to 0.5mL Protein LoBind Eppendorf tubes (Eppendorf Cat #022431064) for centrifugation at 12,700 rpm for 7 minutes at 4°C, supernatant was then collected. The mice were then perfused intracardially with cold PBS, and brain tissues were subdissected to separate the cortical and hippocampal regions.

### SDS-PAGE and Western blotting

Brains were homogenized using Qiagen Tissue-Lyser II (28 Hz, 3 min, 3 times) in lysis buffer (Cell Signaling #9803) containing protease inhibitor cocktail (Roche #4693159001) and PhosSTOP (Roche #4906837001) (1 ml lysis buffer for 100 mg tissue). Lysates were then incubated on ice for 10 min followed by centrifugation at 18,660 g for 20 min at 4°. Supernatants were transferred to new tubes for protein concentration determination and loading sample preparation. Protein lysate samples were boiled at 95°C and SDS-PAGE was performed using standard BioRad reagents. For Western blotting, PDVF membranes were incubated overnight at 4°C with the following primary antibodies diluted blocking buffer (Rockland): Mouse anti APP A4 clone 22C11 (Millipore MAB348, 1:1,000), Mouse anti APP clone 6E10 (Biolegend 803001, 1:2,000), Rabbit anti APP CTF (Sigma A8717, 1:4,000), α-β-Actin (Sigma A2228, 1:4,000). Membranes were then incubated with the appropriate fluorescently conjugated secondary antibody (1:10,000, Li-Cor) and imaged using a Li-Cor Odyssey CLx system.

### Aβ extraction and measurement by MSD

Cortical and hippocampal tissues were weighed and homogenized in TBS (140 mM NaCl, 3 mM KCl, 25 mM Tris-HCl, pH 7.4, 5 mM EDTA, 2 mM 1,10-phenanthroline) containing protease inhibitor (Roche,#4693159001). The homogenates were centrifuged at 100,000 g, 4°C for 1 hour, and then the supernatant was collected (TBS-soluble fraction). Pellets were resuspended in 5 M guanidine, 50 m M Tris, pH 8.0, further homogenized, and incubated at room temperature for 3 hours. Samples were then centrifuged at 20,800g, 4°C, for 20 minutes, and the supernatant was collected (TBS-insoluble GuHCl fraction). MSD V-PLEX Aβ Peptide Panel 1 (4G8) Kit (cat#K15199E) were used for Aβ measurement. Samples were diluted by Diluent 35 provided in the kit (GuHCl fractions 1:20; TBS fractions, no dilution; plasma 1:4; CSF 1:26). Plates were read using the SECTOR Imager 2400A.

### Tau measurement by MSD

Total mouse tau in CSF was measured using an ECL-immunoassay MSD 96-well mouse total tau assay (Meso Scale Discovery) according to the manufacturer’s instructions. Briefly, CSF samples (5 μL) were diluted 1 to 5 in 10% blocking solution (10% BSA in tris wash buffer) and incubated at room temperature for 1 hour on a shaker plate. Standard samples were also diluted in 10% blocking solution. Plates were washed 4 times with 1XTris washing buffer and then incubated at room temperature with sulfo-tag total tau detection antibody for 1 hour. After a few additional washes, plates were read on the MSD SECTOR 600 Reader. Samples were fit against a 9-point standard curve.

### TREM2 measurement by MSD

TREM2 levels in brain homogenates prepared by Cell Signaling lysis buffer (#9803) and plasma were quantified by using MSD technology. The sandwich layout of this TREM2 MSD assay from the bottom to the top is (1) Capture-antibody (1 μg/mL; BAF1729, R&D Systems), (2) Samples, (3) Primary-antibody (10 μg/mL; anti-TREM2 antibody - clone 4D9) ^25^, and (4) Detection-antibody (0.5 μg/mL; SULFO-TAG-labeled anti-human antibody, Meso Scale Discovery). MSD 96-well plates coated with streptavidin were first incubated with 150μL MSD-Blocker A. For building each layer for the assay, 25μL of the reagent was added to each well, and the plates were incubated at R.T on a shaker at 800-rpm for 1hr. Between every incubation step, each well was washed with TBST buffer (0.05% Tween 20 in TBS) using ELx406 plate washer (BioTek). Diluted samples (Brain lysates = 1:5; plasma = 1:20) and TREM2 protein standard (Denali, CA) in assay buffer (25% MSD-Blocker A and 75% TBST) were prepared and aliquoted for running duplicates in the assay. A four-fold serial-dilution standard curve of recombinant murine TREM2 extracellular domain was prepared to include the linear detection range from 62.5 ng/ml to 15.25 pg/mL. For acquiring the MSD units on the MSD Sector Imager S600 reader (MSD), 150μL of 2X MSD read buffer was added to each well after the final TBST wash. TREM2 levels were calculated using the MSD Discovery Workbench software and normalized by each lysate’s concentration (brain samples only).

### Cytokines

Brain homogenates prepared by Cell Signaling lysis buffer (#9803) were diluted to 5 μg/μl with PBS and plasma samples were prepared at a 2-fold dilution with PBS. Diluted samples were sent to Eve Technologies (Canada) for cytokines and chemokines measurement by using the Mouse Cytokine Array / Chemokine Array 44-Plex (MD44).

### Immunohistochemistry

Fresh brains were fixed by immersion in 4% paraformaldehyde at 4°C for 24 hours then transferred to a phosphate buffered saline (PBS) solution with 0.1% sodium azide for storage until ready for processing. Brains were initially transferred to a 30% sucrose solution in PBS for two days before sectioning and then subsequently sectioned coronally at a thickness of 40 μm on a freezing microtome. Sections were stored in cryoprotectant buffer (30% glycerol, 30% ethoxyethanol, and 40% PBS) at −20°C prior to staining. Seven to twelve coronal brain sections (from approximately bregma to 4.8 mm posterior to bregma) were selected for immunostaining for Aβ and microglia markers, IBA1 and CD68. Sections were incubated for 1 hour at room temperature in 1× Tris-buffered saline solution containing 0.05% Tween (TBST) and 5% donkey serum, followed by primary antibodies overnight at 4°C. Sections were then washed in TBST and a solution of secondary antibodies was then applied for 1 hour at room temperature. Sections were washed in TBST prior to mounting and cover slipping with Prolong Glass Antifade Mountant solution (Thermo Fisher, P36984). Immunofluorescence was performed using the following primary antibodies: rabbit anti-human amyloid beta (IBL America, #18584; 1:500), rat anti-CD68 (BioRad, MCA1957A; 1:500), goat anti-Iba1 (Novus, NB100-1028; 1:500), mouse anti-pTau (Invitrogen, MN1020; 1:500), goat-anti-CD31 (R&D Systems, AF3628, 1:500), mouse-anti-alpha smooth muscle actin Cy3 (Sigma, C6198; 1:500), rat anti-LAMP1 (DSHB, 1D4B; 1:250) and mouse anti-Neurofilament (BioLegend, 837801; 1:500); and the following secondary antibodies: Alexa Fluor 488 Donkey anti-rabbit IgG (Invitrogen, #A-21206; 1:200), DyLight 550 Donkey anti rat-IgG (Invitrogen, #SA5-10027, 1:200), Alexa Fluor Plus 555 Donkey anti-mouse IgG (Invitrogen, #A32773; 1:200), Alexa Fluor 647 Donkey anti-goat IgG (Invitrogen, #A-21447; 1:200) and Alexa Fluor Plus 647 Donkey anti-mouse IgG (Invitrogen, #A32787; 1:200).

### Microscopy

Fluorescence microscopy of brain sections was performed using a Zeiss Axioscan.Z1 slide scanner with a 10x / 0.45 NA objective. A custom macro in Zeiss Zen software was used to quantify signal for Aβ, CD68, and Iba1 following median smoothing, channel extraction, and local background subtraction to create binary masks of fluorescent signal for each channel. Objects smaller than 10 pixels were excluded as staining artifacts / debris. Detected Aβ signal was divided into three size bins for plaques 125-250 μm^2^, 250-500 μm^2^, or > 500 μm^2^ in area; individual plaques were also dilated to create a region surrounding the plaques to detect microglia adjacent to plaques. The cortex and hippocampus regions of interest (ROIs) were manually outlined for each section and analyzed using the macro to quantify ROI area and measure the number and area of all plaques and microglia, including overlapping area between microglia and the dilated region surrounding and including plaques.

For 3D assessment of plaque architecture and the associated cellular changes, super resolution confocal microscopy of fluorescently stained sections was performed using a scanning confocal microscope (Leica SP8, Leica Microsystems, Inc.) operated in super resolution lightning mode. Images were acquired using a 63× / 1.4 NA oil immersion objective at a pixel size of 50 nm and processed using the Adaptive processing algorithm. Confocal z-stacks of 18-25 μm were acquired for each channel using sequential scan settings. A minimum of four representative fields were captured from within the cortical brain region of *App*^SAA^ KI/KI (n=3-4) and *App*^SAA^ +/+ control (n=2) mice.

Microscopic evaluation of cerebral amyloid angiopathy (CAA) pathology was performed on larger fields using a 10× / 0.45 NA objective at a pixel size of 150 nm and z-stacks of 30-40 μm, and at higher resolution using a 20× / 0.75 NA oil immersion objective at a pixel size of 67 μm and z-stacks of 25-35 μm. In both cases, images were processed using the lightning Adaptive processing algorithm.

### Whole brain imaging and quantification of plaque distribution

To label amyloid plaques, mice were administered 3.3 mg/kg methoxy-X04 (R&D Systems) by intraperitoneal (i.p.) injection. Twenty-four hours later, mice were perfused with PBS and 4% paraformaldehyde (PFA) sequentially and then intact brains were dissected and postfixed in 4% PFA at room temperature for 6 hours, followed by overnight at 4°C. Whole brain fluorescence imaging was performed as described in Whitesell et al^12^ with serial two-photon (STP) tomography (TissueCyte 1,000, TissueVision Inc., Somerville, MA), using 925 nm excitation, a 500 nm dichroic mirror, and a 447/60 bandpass emission filter on the blue channel. One hundred and forty serial block-face images were acquired from each brain at 0.35 μm/pixel lateral resolution with a 100 μm sectioning interval. Automated segmentation of the fluorescent signal from methoxy-X04 labeled plaques and registration to the 3D Allen Mouse Brain Common Coordinate Framework, v3 (CCFv3)^26^ were performed as previously described^12^. Briefly, segmented fluorescence output is a full resolution mask that classifies each 0.35 μm × 0.35 μm pixel as either signal or background. An isotropic 3D summary of each brain is constructed by dividing each image series into 10 μm × 10 μm × 10 μm grid voxels. Total signal is computed for each voxel by summing the number of signal positive pixels in that voxel. Each image stack is registered to the CCFv3 in a multi-step process using both global affine and local deformable registration. Plaque density for each structure in the reference atlas ontology was calculated by summing voxels from the same structure. To obtain plaque counts within each structure, we used a standard feature labeling algorithm. Adjacent and orthogonally adjacent voxels in the segmentation signal were grouped together as one plaque object. Due to the 100 μm z-sampling interval, our resolution limit for detecting separate plaques in the z-axis was 100 μm. Plaque quantification is reported for one hemisphere per brain, chosen based on image and tissue quality (**Supplementary Table 3**). All image series were subjected to manual QC checks for completeness and uniformity of raw fluorescence images, minimum fluorescence intensity, and artifacts. Automatic segmentation results were checked for overall quality and false positive signals by overlaying segmentation results for 3-5 single coronal sections with raw fluorescent images from STP imaging. Four brains were failed for low signal strength resulting in segmentation failure (n=1 two-month HOM, n=1 two-month WILD, n=2 eight-month HOM).

### Neurofilament Light detection

CSF neurofilament light (Nf-L) concentrations were measured using Simoa NF-Light^®^ (SR-X version, Quanterix 103400) bead-based digital ELISA kits and read on the Quanterix SR-X instrument. CSF samples were diluted 100× with Sample Diluent (Quanterix 102252) before being added to Simoa 96-well microplates (Quanterix 101457). Following kit instructions, Simoa Detector Reagent and Bead Reagent (Quanterix 103159, 102246) were added to the samples before incubating and shaking for 30 mins, 30°C at 800rpm. After incubation, the sample plate was washed with Simoa Wash Buffer A (Quanterix 103078) on a Simoa Microplate Washer according to Quanterix’s two step protocol. After initial washes, SBG Reagent (Quanterix 102250) was added and samples were again incubated at 30°C, 800rpm for 10 min. The two-step washer protocol was continued, with the sample beads being twice resuspended in Simoa Wash Buffer B (Quanterix 103079) before final aspiration of buffer. Sample Nf-L levels were measured using the NF Light analysis protocol on the Quanterix SR-X instrument and interpolated against a calibration curve provided with the Quanterix assay kit.

### Fluorescence activated cell sorting (FACS)

Mice were perfused with cold PBS and cortical and hippocampal tissues were dissected and dissociated into a single cell suspension using the Adult Brain Dissociation Kit (Miltenyi Biotec 130-107-677), according to the manufacturer’s protocol. Cell suspensions were stained with the following antibodies in FACS buffer (1% fatty acid-free BSA + 1mM EDTA in PBS) for 25 minutes on ice: Fixable Viability Stain BV510 (BD Biosciences, 564406), CD11b-BV421 (BioLegend 101251), CD45-APC (BD Biosciences, 559864), ACSA-2-PE (Miltenyi Biotec, 130-102-365), Mouse Fc blocker (anti-mouse CD16/32, BioLegend 101320). Cells were washed twice with FACS buffer and strained through a 100 μm filter before sorting CD11b^+^ microglia and ACSA-2^+^ astrocytes on a FACS Aria III (BD Biosciences) with a 100 μm nozzle. For bulk RNAseq analysis, cells were sorted directly into freshly prepared RLT-plus buffer (Qiagen) containing betamercaptoethanol; for lipid extraction, cells were sorted directly into LC/MS grade Methanol with internal standards.

To label Aβ-containing microglia, mice were injected i.p. with 10 mg/kg methoxy-X04 (R&D Systems). Mice were perfused with PBS 24 hours after injection and cortical and hippocampal tissues were dissected and processed into single-cell suspension for staining as described above except the antibodies used for FACS were: Fixable/viability Dye 780 (Thermo fisher 65-0865-14), Cd11b-PE (BioLegend, 101208), CD45-APC (BD Biosciences, 559864), Mouse FC block 1:50 (BioLegend, 101320). Methoxy-X04 positive and negative microglia (CD11b+) were collected for RNAseq and LC-MS analysis.

### Multiplexed Fluorescence in situ hybridization and imaging

Fresh-frozen mouse brains embedded in OCT were sectioned at 15 μm onto Superfrost Plus glass slides (Fisher Scientific). Sections were stored at −80°C until RNAscope treatment. The RNAscope multiplex fluorescent v2 kit was used following the manufacturer’s protocol for fresh-frozen tissue sections (ACD 323100) except that protease treatment was performed for 15 minutes. Probe sets for *Trem2* (ACD 404111-C2) and *Tmem119* (ACD 472901-C3) were used with Opal 520 and Opal 570 dyes (Akoya Biosciences FP1487001KT), respectively. Methoxy-X04 (Tocris 4920) was applied to the tissue sections at 100 μM in 1×PBS for 30min at room temperature immediately following in situ hybridization. Sections were then washed three times for 10min in 1×PBS before mounting in ProLong Gold Antifade Mountant (ThermoFisher P36930) followed by a coverslip. Sections were imaged using a confocal microscope (Leica SP8; Leica Microsystems, Inc.) with a 20×/.75IMM oil objective.

### RNA isolation, qPCR, and SMARTSeq library preparation

RNA from bulk mouse brain tissue was extracted using the RNeasy Plus Mini Kit (Qiagen) and resuspended in nuclease-free water. For qPCR, 3μl of RNA was transcribed into cDNA using SuperScript IV (Invitrogen). *App* gene expression level was assessed using Taqman probe (FAM-Mm01344172_m1) on a QuantStudio 6 Flex (Applied Biosystems) and normalized to *Gapdh*.

For RNA-seq analysis of FACS sorted cells, 200 live cells were sorted directly into 11.5 microliters of Clontech SMARTseq 10X reaction buffer for reverse transcription. cDNA synthesis was performed with the Clontech SMARTSeq v4 3’ DE kit (Takara Bio USA, Inc. 635040) kit. Reverse transcription was followed by 14 cycles of cDNA amplification. cDNA was purified with 0.8X volume of SPRIselect beads (Beckman Coulter B23318), quantified on a Bioanalyzer 2100 System with a High Sensitivity DNA chip (Agilent 5067-4626). 300 pg of cDNA from each sample was used as input for library preparation with the Nextera XT DNA Library Prep Kit (Illumina FC-131-1096). Fragmentation and adaptors insertion were performed by tagmentation, followed by 12 cycles of PCR amplification. The final libraries were purified using 0.8X SPRIselect beads. Library quantity and quality were assessed with Qubit 4 Fluorometer with the 1X dsDNA HS Assay Kit (Invitrogen Q33231) and on a Bioanalyzer 2100 System with a High Sensitivity DNA chip. Sequencing reads were generated on an Illumina NovaSeq 6000 instrument (100 bp single end) by SeqMatic (Fremont, CA, USA).

Sequencing adapters were trimmed from the raw reads with skewer (version 0.2.2) ^27^Reads were aligned to the mouse genome version GRCm38_p6. A STAR index (version 2.7.1a) ^28^ and built with the --sjdbOverhang=50 argument. Splice junctions from Gencode gene models (release M17) were provided via the --sjdbGTFfile argument. STAR alignments were generated with the following parameters: --outFilterType BySJout, --quantMode TranscriptomeSAM, -- outFilterIntronMotifs RemoveNoncanonicalUnannotated, --outSAMstrandField intronMotif, -- outSAMattributes NH HI AS nM MD XS and --outSAMunmapped Within. Alignments were obtained with the following parameters: --readFilesCommand zcat --outFilterType BySJout -- outFilterMultimapNmax 20 --alignSJoverhangMin 8 --alignSJDBoverhangMin 1 -- outFilterMismatchNmax 999 --outFilterMismatchNoverLmax 0.6 --alignIntronMin 20 -- alignIntronMax 1000000 --alignMatesGapMax 1000000 --quantMode GeneCounts -- outSAMunmapped Within --outSAMattributes NH HI AS nM MD XS --outSAMstrandField intronMotif --outSAMtype BAM SortedByCoordinate --outBAMcompression 6. Gene level counts were obtained using featureCounts from the subread package (version 1.6.2)^29^. Gene symbols and biotype information were extracted from the Gencode GTF file.

### RNA-seq data analysis

Following alignment and expression quantitation, lowly expressed genes were removed, and differential expression analysis was performed using the limma/voom with sample weighting framework^30,31^. Lowly expressed genes are those that did not have more than ten reads assigned to them in at least as many samples as the minimum replicate size per experiment (six samples in GSE158152 and ten in GSE158153). Technical replicate libraries from GSE158152 (identified by the *animal_id* column) were summed together prior to analysis. Linear models were fit against the covariate(s) of interest (genotype and/or methoxy-X04-status) with “take-down day” and “sex” encoded as batch covariates. Because the direct comparison of the methoxy-X04-positive vs methoxy-X04-negative profiles in *App*^SAA^ KI/KI mice utilized repeated measures from the same animal, “subject_id” was fit as a random effect using the duplicateCorrelation function from limma^32^. Empirical Bayes moderated t-statistics and *p-*values were computed relative to a 1.2 fold-change cutoff using treat^33^. Significantly differentially expressed genes were then defined as those having an FDR <= 10%.

Single sample gene set activity scores were calculated in **Fig. 2b** by taking a weighted average of the expression of the genes in each gene set. Weights correspond to the loadings of each gene on the first principal component of the mean-centered gene-by-sample matrix for each gene set.

Gene set enrichment analyses were performed via the fgseaMultiLevel function in the fgsea R package using the moderated t-statistic as the gene ranking statistic^34^. Gene sets used for testing were taken from the Biological Process collection of the Gene Ontology database, the KEGG database, as well as a custom set of genes compiled from the literature that are enumerated in **Supplemental Table 4**^35–37^. Gene set enrichment scores shown **Fig. 2i** and **Supplemental Fig. 5b** were calculated by averaging the moderated t-statistics of the genes in the leading edge of the gene set. Because the leading edge can consist of different genes for each comparison, we defined the genes used in this calculation as the universe of the genes that appeared in the leading edge of the gene set across all comparisons shown in these figures.

All software versions for the RNA-seq analysis correspond to Bioconductor release version 3.11^38^.

### FACS lipid extraction

Lipid extraction was performed using Matyash liquid-liquid extraction protocol with the following modifications. Briefly, 50,000 cells were sorted directly into 400 μl of methanol containing surrogate internal standards and kept on ice. To each tube, 200-400 μl of water was added to achieve final volume of 800 μl. These samples were vortexed for 5 min at room temperature. To these samples, 800 μl of tert-Butyl methyl ether (MTBE) was added and vortexed for an additional 5 min at room temperature. Samples were then centrifuged at 21,000 x g for 10 min at 4°C. Following centrifugation, 700 μl of upper organic layer was collected and dried under constant stream of N_2_ gas. Dried samples were reconstituted in 100 μl of MS-grade methanol for further analysis by LC-MS/MS.

### LCMS analysis of lipids

Lipid analyses were performed by liquid chromatography UHPLC Nexera X2, coupled to electrospray mass spectrometry (QTRAP 6500+, Sciex). For each analysis, 5 μL of sample was injected on a BEH C18 1.7 μm, 2.1×100 mm column (Waters) using a flow rate of 0.25 mL/min at 55°C. Electrospray ionization was performed in positive and negative ion modes.For positive ionization mode, mobile phase A consisted of 60:40 acetonitrile/water (v/v) with 10 mM ammonium formate + 0.1% formic acid; mobile phase B consisted of 90:10 isopropyl alcohol/acetonitrile (v/v) with 10 mM ammonium formate + 0.1% formic acid. For negative ionization mode, mobile phase A consisted of 60:40 acetonitrile/water (v/v) with 10 mM ammonium acetate + 0.1% acetic acid; mobile phase B consisted of 90:10 isopropyl alcohol/acetonitrile (v/v) with 10 mM ammonium acetate + 0.1% acetic acid. The gradient was programmed as follows: 0.0-8.0 min from 45% B to 99% B, 8.0-9.0 min at 99% B, 9.0-9.1 min to 45% B, and 9.1-10.0 min at 45% B.

Electrospray ionization was performed using the following settings: curtain gas at 30 psi; collision gas at 8 psi; ion spray voltage at 5500 V (positive mode) or −4500 V (negative mode); temperature at 250°C (positive mode) or 600°C (negative mode); ion source Gas 1 at 55 psi; ion source Gas 2 at 60 psi; entrance potential at 10 V (positive mode) or −10 V (negative mode); and collision cell exit potential at 12.5 V (positive mode) or −15.0 V (negative mode). Data acquisition was performed in multiple reaction monitoring mode (MRM) with the collision energy (CE) values reported in **Supplementary Table 5**. Lipids were quantified as area normalized to specific non-endogenous internal standards as reported in **Supplementary Table 5**. Quantification was performed using MultiQuant 3.02 (Sciex). Lipids were normalized to protein amount. Protein concentration was measured using the bicinchoninic acid (BCA) assay (Pierce, Rockford, IL, USA).

### LCMS analysis of polar metabolites

#### Positive mode

Metabolites analyses were performed liquid chromatography (UHPLC Nexera X2) coupled to electrospray mass spectrometry (QTRAP 6500+, Sciex). For each analysis, 5 μL of sample was injected on a BEH amide 1.7 μm, 2.1×150 mm column (Waters Corporation, Milford, Massachusetts, USA) using a flow rate of 0.40 mL/min at 40°C. Mobile phase A consisted of water with 10 mM ammonium formate + 0.1% formic acid. Mobile phase B consisted of acetonitrile with 0.1% formic acid. The gradient was programmed as follows: 0.0–1.0 min at 95% B; 1.0–7.0 min to 50% B; 7.0–7.1 min to 95% B; and 7.1–10.0 min at 95% B. The following source settings were applied: curtain gas at 30 psi; collision gas was set at at 8 psi; ion spray voltage at 5500 V; temperature at 600°C; ion source Gas 1 at 50 psi; ion source Gas 2 at 60 psi; entrance potential at 10 V; and collision cell exit potential at 12.5 V. Data acquisition was performed in multiple reaction monitoring mode (MRM) with the collision energy (CE) values reported in **Supplementary Table 6**. Quantification was performed using MultiQuant 3.02 (Sciex).

#### Negative mode

Metabolites analyses were performed liquid chromatography (ACQUITY I-Class Plus UPLC FL, Waters Corp) coupled to electrospray mass spectrometry (XEVO TQ-S Micro, Waters Corp). For each analysis, 5 μL of sample was injected on an Agilent InfinityLab Poroshell 120 HILIC-Z P 2.7 μm, 2.1×50 mm (Agilent Technologies Inc., Santa Clara, CA USA); using a flow rate of 0.5 mL/min at 25°C. Mobile phase A consisted of water with 10 mM ammonium acetate + 5 μM medronic acid (Agilent), pH 9. Mobile phase B consisted of acetonitrile:water 9:1 with 10 mM ammonium acetate + 5 μM medronic acid (Agilent), pH 9. The gradient was programmed as follows: 0.0–1.0 min at 95% B; 1.0–6.0 min to 50% B; 6.0–6.5 min to 95% B; and 6.5–10.0 min at 95% B. Electrospray ionization was performed in negative ion mode. The following source settings were applied: capillary voltage at 1.9 kV; source temperature at 150°C; desolvation temperature at 600°C; desolvation gas flow at 1000 L/hr; cone gas flow at 50 L/hr; cone voltage at 20 V; nebulizer gas at 7 bar. Data acquisition was performed in multiple reaction monitoring mode (MRM) with the collision energy (CE) values reported in **Supplementary Table 6**. Quantification was performed using Skyline (v19.1;University of Washington).

### Statistical analysis

Data have either been expressed as means ± SEM or as indicated in graphs. Statistical analysis of data was performed in GraphPad Prism 8 or as indicated otherwise for whole brain plaque analysis, RNAseq and LC-MS data. Analysis was performed using *t* test or with one-way analysis of variance (ANOVA) with Dunnett multiple comparison as indicated in figure legends. Criterion for differences to be considered significant was *P* < 0.05.

Comparison of LC/MS results from FACS-isolated microglia from *App*^SAA^ +/+, *App*^SAA^ KI/+ and *App*^SAA^ KI/KI mice: Integrated peak areas were divided by the area of pre-assigned internallimma standards and the resulting ratios were log2 transformed. Analytes detected in less than 75% of the samples were removed. To identify and account for unwanted sources of variation, two surrogate variables (SV) were identified with the sva R package^39^. Genotypes were compared by fitting the following linear model using the limma R package^40^: log_2_(abundance) ~ genotype + batch + sex + SV. Sample weights were incorporated via limma’s arrayWeights function. Analytes with an estimated absolute fold difference of more than 20% at a false discovery rate of less than 10% were deemed significant. To visualize adjusted abundances in the heatmap, the batch, sex and SV covariates were regressed out with limma’s removeBatchEffect function. LC/MS results from methoxy-positive and −negative microglia were analyzed in a similar way, but as no significant surrogate variables were identified, median scaling was applied to the log_2_ transformed ratios of each sample instead and the following linear model was fit using limma: log_2_(abundance) ~ genotype + batch + sex.

## Data availability

RNA-seq datasets have been deposited online to the Gene Expression Omnibus database under accession GSE158156.

## Author contributions

Conceptualization: D.X., M.S., J.W.L., G.D.P., P.E.S. Experiments: D.X., M.C., E.T., J.D., M.P., S. L.D., T.E., C.C.L., S.D., C.H., H.N., R.C., E.Y., H.S., S.T.M., R.C., K.L., D.G., J.D.W. Data analysis: D.X., S.L., T.S., M.C., J.H.S., M.P., S.L.D., R.G.T., T.K.E., R.C., J.D.W., J.A.H, P.E.S. Writing original draft: D.X., S.L., T.S., J.S., G.D.P., P.E.S. Writing – review and editing: D.X., S.L., T. S., M.C., J.S., M.P., S.L.D., R.G.T., J.D.W., M.S., J.A.H., K.S.L., J.W.L., P.E.S. Supervision: P.E.S

## Acknowledgements

5×FAD brains were a generous gift from Drs. Sandro Da Mesquita and Jonathan Kipnis to J.A.H. J.D.W. and J.A.H thank the Allen Institute founder, Paul G. Allen, for his vision, encouragement, and support.

