## Supplementary Figures for "Fibrillar Aβ causes profound microglial metabolic perturbations in a novel APP knock-in mouse model"

Supplementary Figure 1 (Extended Figure 1)

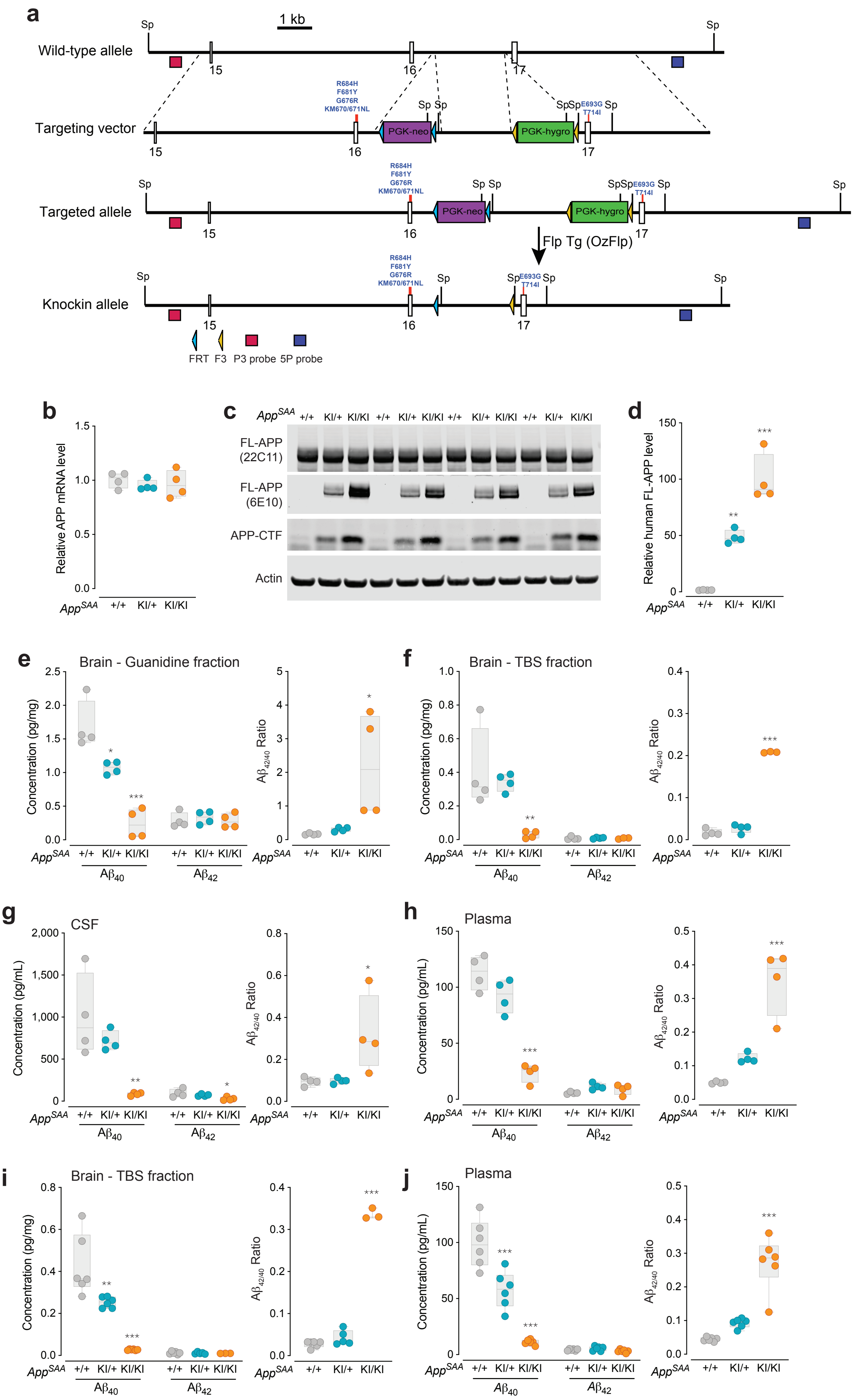

**Supplementary Figure 1 (Extended Figure 1). Levels of APP and Aβ in brain, CSF and plasma from *App*$^{SAA}$ KI mice. (a)** Schematic illustrating the genetic engineering approach of the *App*$^{SAA}$ mouse line. **(b)** *App* mRNA levels were measured in brain by RT-qPCR. **(c)** Western blotting analysis of brain lysates at 2 months of age and corresponding **(d)** quantification of human full-length APP. **(e-j)** $Aβ_{40}$ and $Aβ_{42}$ concentrations were measured and the ratio of $Aβ_{42/40}$ calculated from biofluids and tissues from the 3 genotypes of the *App*$^{SAA}$ mouse line, at 2-month-old (e-h) and 4-month-old (i-j). Graphs display means ± SEM and P values: one-way ANOVA with Dunnett's multiple comparison test, each group compared to the *App*$^{SAA}$ +/+ wild-type control group; *P < 0.05, **P < 0.001, and ***P < 0.0001.

Supplementary Figure 2 (Extended Figure 1)

**a**

| Human region (Thal *et al.* 2002) | Mouse Region (Allen CCFv3) |
|---|---|
| Neocortex | Isocortex |
| CA1 | CA1 |
| Entorhinal Region | ENTl, ENTm |
| Gyrus Cinguli | ACA, RSP |
| Amygdala | LA,BLA,BMA,PA,CEA,MEA,COA |
| Fascia Dentata | DG |
| Presubiculum | PRE, POST |
| Thalamus | TH |
| Striatum | CP, ACB |
| Hypothalamus | HY |
| Basal Forebrain Nuclei (Meynert) | SI, MA, NDB |
| CA4 | Not annotated |
| Central Gray | PAG |
| Superior Collicle | SCs, SCm |
| Red Nucleus | RN |
| Inferior Olivary Nucleus | IO |
| Substantia Nigra | SNr, SNc |
| Ret. Form. of the Med. Oblongata | GRN, IRN, PARN |
| Cerebellar Molecular Layer | CBX |
| Reticular Formation of the Pons | PRNr, PRNc |
| Ant. and Central Raphe nuclei | RAMb |
| Locus coeruleus | LC |
| Parabrachial Nuclei | PB |
| Reticulo Tegmental Nucleus | TRN |
| Dorsal Tegmental Nucleus | DTN |
| Nuclei Pontis | PG |
| Cerebellar Granule Cell Layer | Not annotated |
| Dentate Nucleus | DN |

age -> *App*SAA KI/KI: 2, 4, 8; APP/PS1: 5, 7, 9, 13, 19; Tg2576: 9, 13, 19; 5xFAD: 12

Phase 1

Phase 2

Phase 3

Phase 4

Phase 5

No Deposits

**b**

| | *App*SAA KI/KI | APP/PS1 | Tg2576 | 5xFAD |
|---|---|---|---|---|
| 90% | 51 | 109 | 45 | 61 |
| 50% | 16 | 41 | 9 | 30 |
| 10% | 0 | 0 | 0 | 0 |

-0.4 0 0.4 0.8

Pearson Correlation

**c**

| | *App*SAA KI/KI 2 | 4 | 8 | APP/PS1 5 | 7 | 9 | 13 | 19 | Tg2576 9 | 13 | 19 | 5xFAD 12 |
|---|---|---|---|---|---|---|---|---|---|---|---|---|
| Phase 1 | 0.03 | 0.60 | 0.34 | 0.26 | 0.42 | 0.42 | 0.38 | 0.34 | -0.08 | 0.06 | 0.22 | 0.12 |
| Phase 2 | -0.30 | 0.88 | 0.80 | 0.54 | 0.81 | 0.82 | 0.80 | 0.75 | 0.01 | 0.46 | 0.70 | 0.20 |
| Phase 3 | -0.43 | 0.69 | 0.83 | 0.42 | 0.63 | 0.73 | 0.77 | 0.77 | -0.07 | 0.28 | 0.56 | 0.68 |
| Phase 4 | -0.31 | 0.49 | 0.62 | 0.37 | 0.47 | 0.52 | 0.56 | 0.54 | -0.13 | 0.08 | 0.34 | 0.83 |
| Phase 5 | -0.21 | 0.35 | 0.50 | 0.30 | 0.39 | 0.46 | 0.48 | 0.49 | 0.16 | 0.40 | 0.45 | 0.62 |

**d**

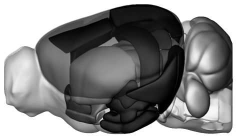

APPSAA (8 mo)

**Supplementary Figure 2 (Extended Figure 1). Brain-wide spatio–temporal patterns of plaque deposition in *App*[SAA] KI/KI mice recapitulates the pattern seen in human patients with AD. (a)** Comparison of relative plaque density in similar structures between human autopsy tissue and four APP mouse models. Human brain regions where Aβ pathology was quantified by Thal *et al*. (2002) are listed in the left column and the corresponding region(s) from the Allen CCFv3 reference atlas are listed in the second column. The median plaque density (plaques per $mm^3$) for each age group and mouse line is indicated by the heatmap in the columns to the right. Cartoons on the right show the anatomical location of the structures that were included in each Phase. The colormap for the plaque density spans from the 10th percentile to the 90th percentile of the plaque density for all structures at the oldest age in each mouse line. **(b)** 10, 50, and 90% values and corresponding colormaps are shown for each mouse line. **(c)** Similarity measured using the Pearson correlation coefficient for comparisons between plaque density in each mouse line and age group with the fraction of patients showing plaques in each region during the five phases of Aβ deposition. **(d)** Anatomical location of the structures used to compare mouse plaque deposition with Thal phases, colored by the measured plaque density in 8-month-old *App*[SAA] KI/KI mice. Abbreviations: ENTl = entorhinal area, lateral part; ENTm = entorhinal area, medial part, dorsal zone; ACA = anterior cingulate area; RSP = retrosplenial area; LA = lateral amygdalar nucleus; BLA = basolateral amygdalar nucleus; BMA = basomedial amygdalar nucleus; PA = posterior amygdalar nucleus; CEA = central amygdalar nucleus; MEA = medial amygdalar nucleus; COA = cortical amygdalar area; DG = dentate gyrus; PRE = presubiculum; POST = postsubiculum; TH = thalamus; CP = caudoputamen; ACB = nucleus accumbens; HY = hypothalamus; SI = substantia innominata; MA = magnocellular nucleus; NDB = diagonal band nucleus; PAG = periaqueductal gray; SCs = Superior colliculus, sensory related; SCm = superior colliculus, motor related; RN = red nucleus; IO = inferior olivary complex; SNr = substantia nigra, reticular part; SNc = substantia nigra, compact part; GRN = gigantocellular reticular nucleus; IRN = intermediate reticular nucleus; PARN = parvicellular reticular nucleus; CBX = cerebellar cortex; PRNr = pontine reticular nucleus; PRNc = pontine reticular nucleus, caudal part; RAmb = midbrain raphe nuclei; LC = locus ceruleus; PB = parabrachial nucleus; DN = dentate nucleus.

Supplementary Figure 3 (Extension of Figure 1)

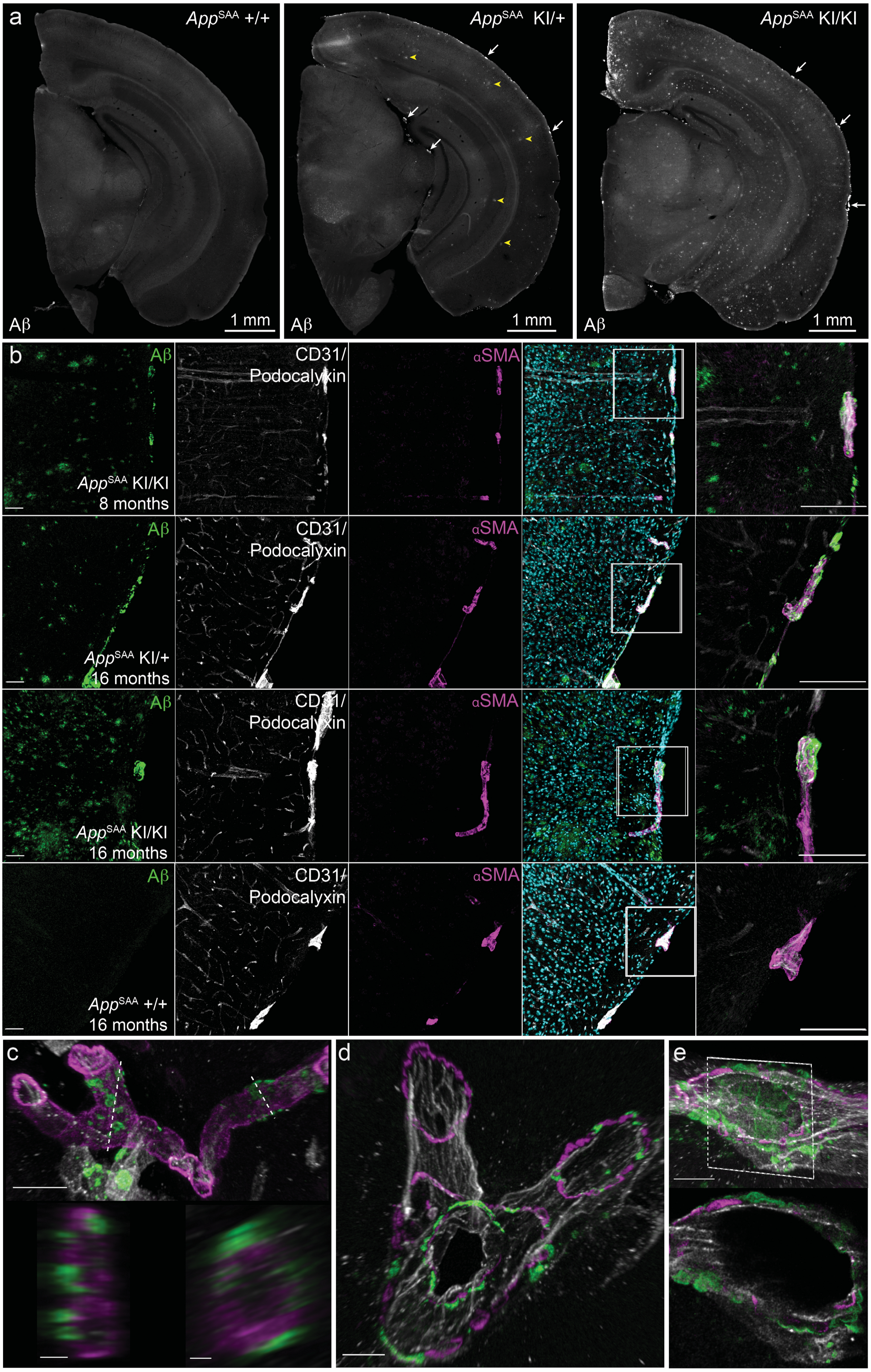

**Supplementary Figure 3 (Extended Figure 1). Evidence of CAA pathology in *App*$^{SAA}$ mice at 8 and 16 months of age** (a) Representative images of Aβ deposition in *App*$^{SAA}$ +/+, *App*$^{SAA}$ KI/+, and *App*$^{SAA}$ KI/KI mice at 16 months of age; *App*$^{SAA}$ KI/+ and *App*$^{SAA}$ KI/KI exhibit parenchymal Aβ plaque deposition (examples indicated with yellow arrowheads for *App*$^{SAA}$ KI/+) and Aβ deposition associated with leptomeningeal vessels (CAA; examples indicated with white arrows for both *App*$^{SAA}$ KI/+ and *App*$^{SAA}$ KI/KI). (b) Representative confocal images showing accumulation of Aβ surrounding leptomeningeal (pial) vessels (endothelial cells labeled by CD31 and smooth muscle cells labeled by alpha-smooth muscle actin, consistent with putative arteries/arterioles). Super resolution confocal images at 16 months of age showing Aβ accumulation in a penetrating parenchymal vessel **(c)**, branching leptomeningeal vessel in the ambient cistern **(d)** and a leptomeningeal vessel **(e).** Scale bars = 20 μm.

### Supplementary Figure 4 (Extended Figure 1)

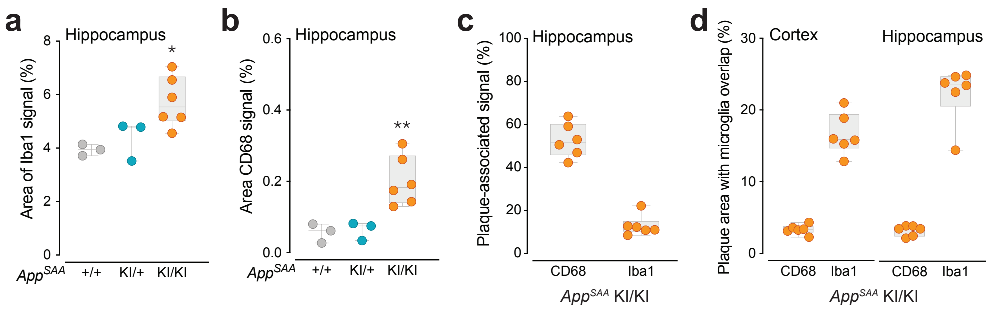

**Supplementary Figure 4 (Extended Figure 1). Histological analysis of microglia and amyloid-β plaques.** Analysis was conducted in brain sections from 8-month-old mice. Quantification of areas covered by Iba1 **(a)**, CD68 **(b)** and the percentage of the CD68 and Iba1 signals overlapping with amyloid-β plaques **(c)** in the hippocampus from *App*$^{SAA}$ KI/KI mice. **(d)** Percentage of plaque overlapping with CD68 or Iba1 in the hippocampus or cortex. Graphs display means ± SEM and P values: one-way ANOVA with Dunnett's multiple comparison test, each group compared to the *App*$^{SAA}$ +/+ control group (n=4-6 per group); *P < 0.05 and **P < 0.001.

### Supplementary Figure 5

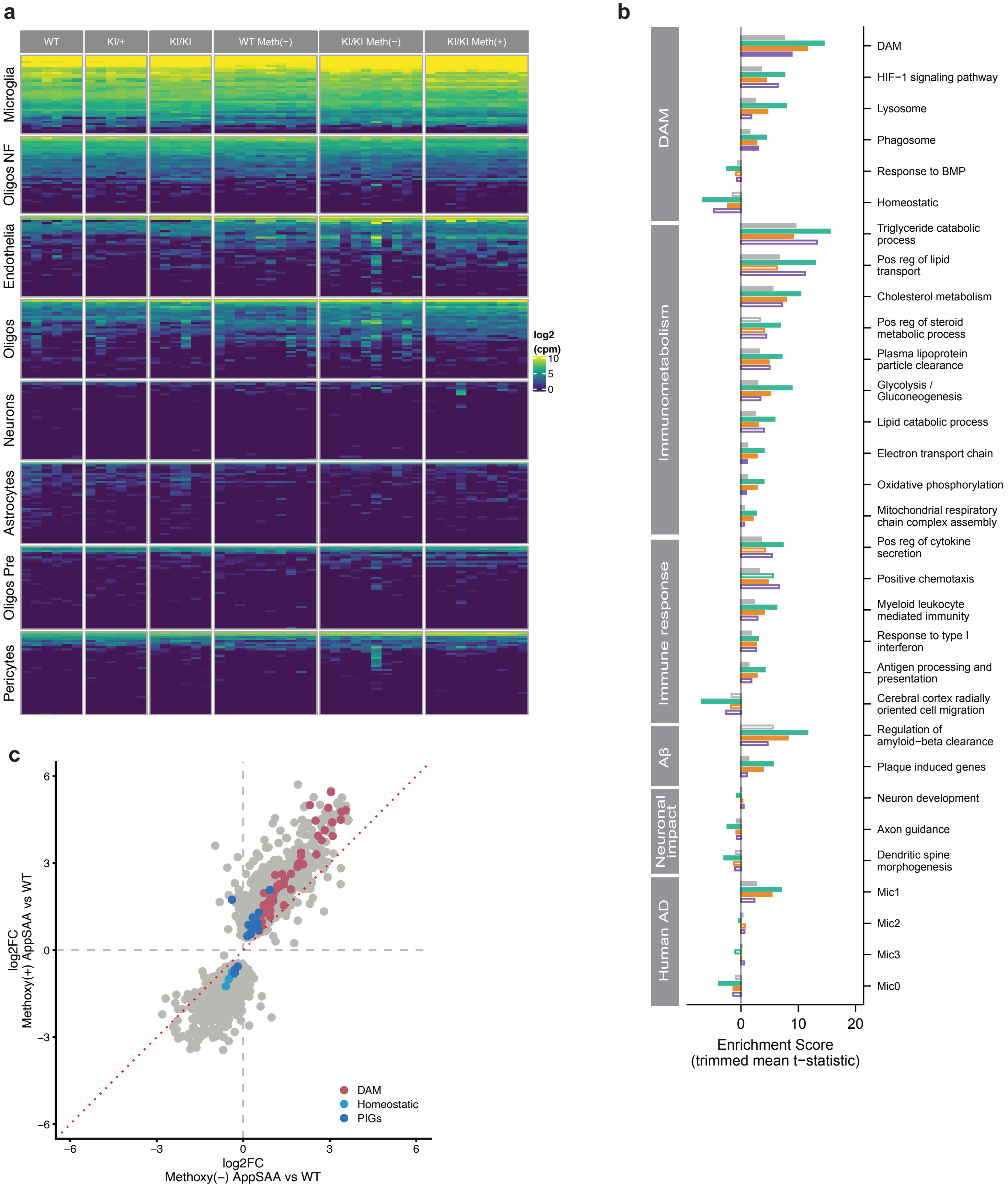

**Supplementary Figure 5 (Extended Figure 2). (a)** Absolute expression (log2 counts per million) of cell type specific markers shows strong enrichment and purity of microglia cell population analyzed in this study. **(b)** Extended results of GSEA analysis from Fig 2i with statistics for non-methoxy sorted microglia in *App*$^{SAA}$ vs WT (orange), as well as 5XFAD vs WT (purple). **(c)** Log2 fold change of methoxy-X04 (-) *App*$^{SAA}$ +/+ microglia vs methoxy-X04 (-) WT microglia (*x*-axis) vs log2 fold change of methoxy-X04 (+) *App*$^{SAA}$ +/+ microglia vs methoxy-X04 (-) WT (*y*-axis). Only genes with log2 fold change >= 1.2 and FDR <= 10% in either comparison are shown. Genes from the homeostatic (light blue), DAM (fuscia), and PIGs (dark blue) signatures are highlighted.
