## Supplementary Table 1 for "Fibrillar Aβ causes profound microglial metabolic perturbations in a novel APP knock-in mouse model"

### Supplementary Table 1. Primer sequence and assays for mouse model generation

| | |
|---|---|
| P1915_41 | CTAATACTGACATGAATGAGGTCTGCTC |
| P1915_51 | TAAGCATTGGTAAGACGTCTATATGCTGGACTTCTTTCTGCCC |
| P1915_74 | TAAGCATTGGTAAACCGGTAGGCGCCAACCGGCTCCGTTCTTTGGTGGCCCCTTCGCGCCACCTTCTACTCCT<br>CCCCTAGTCAG |
| P1915_53 | CTAAGGCGCGCCGAAGTTCCTATACTATTTGAAGAATAGGAACTTCGATCTATAGATCATGAGTGGGAGGAATG<br>AGCTGGCCCTTAATTTG |
| P1915_46 | TAAGCATTGGTAAACGCGTCATGCCATAATTAAAAGGGGAGGG |
| P1915_56 | CTAAATTGAGTAGCTGTAGGAGGAGGTA |
| 5'probe for southern blotting | GGCATCTCAATTCCCTGTCTTTTCTTGGTTTGCTTTGCGTGCTAATCCTCCATGCCCTTTGCTGCTGGGAAGC<br>AGAAGGGGTTCACGATTCCTGCCATGGGGACAGCTTAGCCCTCAGTGTTCACAGGGCTCTGGGCTTCCGAA<br>TAGATTGCTTTTAAAGGCACCCGAGGAGAAGAGTCCAAATGGCATCTTTTAATCCCCAATCGTCTTGAATTCT<br>GGAAGTGAATATTCATGTCTGGGGAGAAACCTCCGGCTAAACAAAGAACAAGGATGAGCCTTCCTCCTAGCA<br>AGGGCCCTCAAAGCTACCTCGAGGACTGTGCCAATCCCTCTGTGACCACTGTGTGCCTCTAGCAACAGACTG<br>AAATTGTGGG |
| 3'probe for southern blotting | CCTTGTCAGAAAACAACTTCCCTCAGCTCAGACTGGATTTTGAAGAAATAAGGAAGGGAGAGAGAGGGCAAC<br>CAAGCTTTAAGGATTAACAAGTGGCTGAAGGGCAGATGAAGGCAAGGGAGGAGGAAGTCCCCTGGGTCAGG<br>TCTGGCAAGTGTAAAAGAGAGAAAGAAATCTGAGCGTAGTGGTGGAGGGAAGCCAGCGGGAGTTTGATGGG<br>AGCCTTGGGGACCCACAGTTTCAACCATTTAGCTTACAGAGGCTCACAGGCCAAGTGGTCGAAAGTGACCAC<br>AACCCCAAATCCGTTCTGAATGGATGGCACAGTTGATAAGAATGG |
| Hygro1 probe for southern blotting | CTGTCGAGAAGTTTCTGATCGAAAAGTTCGACAGCGTCTCCGACCTGATGCAGCTCTCGGAGGGCGAAGAAT<br>CTCGTGCTTTCAGCTTCGATGTAGGAGGGCGTGGATATGTCCTGCGGGTAAATAGCTGCGCCGATGGTTTCT<br>ACAAAGATCGTTATGTTTATCGGCACTTTGCATCGGCCGCGCTCCCGATTCCGGAAGTGCTTGACATTGGGGA<br>ATTCAGCGAGAGCCTGACCTATTGCATCTCCCGCCGTGCACAGGGTGTCACGTTGCAAGACCTGCCTGAAAC<br>CGAACTGCCCGCTGTTCTGCAGCCGGTCGCGGAGGCCATGGATGCGATCGCTGCGGCCGATCTTAGCCAGA<br>CGAGCGGGTTCGGCCCATTCGGACCGCAAGGAATCGGTCAATACACTACATGGCGTGATTTCATATGCGCGA<br>TTGCTGATCCCCATGTGTATCACTGGCAAACTGTGATGGACGACACCGTCAGTGCGTCCGTCGCGCAGGCTC<br>TCGATGAGCTGATGCTTTGGGCCGAGGACTGCCCCGAAGTCCGGCACCTCGTGCACGCGGATTTCGGCTCCA<br>ACAATGTCCTGACGGACAATGGCCGCATAACAGCGGTCATTGACTGGAGCGAGGCGATGTTCGGGGATTCCC<br>AATACGAGGTCGCCAACATCTTCTTCTGGAGGCCGTGGTTGGCTTGTATGGAGCAGCAGACGCGCTACTTCGA<br>GCGGAGGCATCCGGAGCTTGCAGGATCGCCGCGGCTCCGGGCGTATATGCTCCGCATTGGTCTTGACCAACT<br>CTATCAGAGCTTGGTTGACGGCAATTTCGATGATGCAGCTTGGGCGCAGGGTCGATGCGACGCAATCGTCCGA<br>TCCGGAGCCGGGACTGTCGGGCGTACACAAATCGCCCGCAGAAGCGCGGCCGTCTGGACCGATGGCTGTGT<br>AGAAGTACTCG |
| enP probe for southern blotting | TAAGCATTGGTAAAGAGCACATTTGTTATGTAAGTTAGTGCCAACAGCTCCCTAAATAATTCTTTCCAGGCGGCT<br>TCAAGATAGGTGAATGACCGATTTCCATCGCTAAATCCATCCCTGCTGCAGTTTGCAAAGGCGAGGTAAACTAG<br>AGCAGATAAAATTTTTCTAAGTGGATGTTACTCACAGCGCATCTGCGGCAAAATTAGAAATGGCTCGTAATTAAT<br>CCCCTCGCCAACCAAAAATGCTAGCTCAACACATTTATTAAAGGAGATCATTAATTAACTAATGATACTACAGTG<br>AGGTGATCGGGAATAAATTTAG |
| neo probe for southern blotting | TAAGCATTGGTAAGACTGGGCACAACAGACAATCGGCTGCTCTGATGCCGCCGTGTTCCGGCTGTCAGCGCAG<br>GGGCGCCCGGTTCTTTTTGTCAAGACCGACCTGTCCGGTGCCCTGAATGAACTGCAGGACGAGGCAGCGCGGC<br>TATCGTGGCTGGCCACGACGGGCGTTCCTTGCGCAGCTGTGCTCGACGTTGTCACTGAAGCGGGAAGGGACTG<br>GCTGCTATTGGGCGAAGTGCCGGGGCAGGATCTCCTGTCATCTCACCTTGCTCCTGCCGAGAAAGTATCCATCA<br>TGGCTGATGCAATGCGGCGGCTGCATACGCTTGATCCGGCTACCTGCCCATTCGACCACCAAGCGAAACATCGC<br>ATCGAGCGAGCACGTACTCGGATGGAAGCCGGTCTTGTCGATCAGGATGATCTGGACGAAGAGCATCAGGGGCT<br>CGCGCCAGCCGAACTGTTCGCCAGGCTCAAGGCGCGCATGCCCGACGGCGATGATCTCGTCGTGACCCATGGC<br>GATGCCTGCTTGCCGAATATCATGGTGGAAAATGGCCGCTTTTCTGGATTCATCGACTGTGGCCGGCTGGGTGTG<br>GCGGACCGCTATCAGGACATAGCGTTGGCTACCCGTGATATTGCTGAAGAGCTTGGTTAG |
| qPCR for ES cell screening | 1915_Lo5WT assay (Ozgene) |
| qPCR to confirm 5'targeting | 1915_Lo5WT assay (Ozgene) |
| qPCR to confirm 3'targeting | 1915_LoWT3 assay (Ozgene) |
| qPCR to confirm absence of random integration: | 1638_goNoz assay (Ozgene) |
| qPCR to confirm the copy numbers of the Y Chromosome in the ES cells | 1638_LoChrY assay (Ozgene) |
| qPCR to confirms the copy numbers of the Chromosome 8 in the ES cells | 1638_goChr8 assay (Ozgene) |
