## Supplementary Table 2 for "Fibrillar Aβ causes profound microglial metabolic perturbations in a novel APP knock-in mouse model"

**Supplementary Table 2. Animal cohorts used for each figure panel**

| Mouse Cohort | Age (months) | Sample size and sex by genotype (M: males, F: females) | | | Corresponding figure panels |
|---|---|---|---|---|---|
| | | ***App*$^{SAA}$ +/+** | ***App*$^{SAA}$ KI/+** | ***App*$^{SAA}$ KI/KI** | |
| 1 | 2 | 2MMM/ 2F | 3M/ 1F | 1M/ 3F | • Fig. 1a-b<br>• Suppl. Fig. 1b-h |
| 2 | 4 | 4M/ 2F | 4M/ 2F | 4M/ 2F | • Fig. 1c-d<br>• Suppl. Fig. 1i-j |
| 3 | 8 | 3M/ 3F | 2M/ 4F | 4M/ 2F | • Fig. 1h-m, p-r<br>• Fig. 2a-b<br>• Fig. 3a-b<br>• Suppl. Fig. 3b<br>• Suppl. Fig. 4<br>• Suppl. Fig. 5a |
| 4 | 2 | 1M/ 1F | 1M | 4M/ 1F | • Fig. 1e-g, q-r<br>• Suppl. Fig. 2a-d |
| | 4 | 2F | 6M | 3M/ 3F | |
| | 8 | 1M/ 1F | 3M/ 3F | 3M/ 3F | |
| 5 | 8 | 3M/ 7F | - | 8M/ 2F | • Fig. 1n-o<br>• Fig. 2 c-i<br>• Fig. 3c-f<br>• Suppl. Fig. 5a-c |
| 5 | 16 | 1M/3F | 1M/1F | 3M | • Suppl. Fig. 3a-e |
| ***Total*** | | ***15M/ 21F*** | ***20M/ 11F*** | ***30M/ 16F*** | |
