## Supplementary Table 3 for "Fibrillar Aβ causes profound microglial metabolic perturbations in a novel APP knock-in mouse model"

Supplementary Table 3 - Plaque Count Whole-Brain

| image_series_id | hemisphere_id | structure_acronym | structure_id | structure_volume | plaque_volume | plaque_count | plaque_density | count_per_mm_3 | age_group | genotype |
|---|---|---|---|---|---|---|---|---|---|---|
| 1013978665 | 1 | TMv | 1 | 0.111060313 | 0 | 0 | 0 | 0 | 2 mo | HOM |
| 1013445609 | 1 | TMv | 1 | 0.111060313 | 0 | 0 | 0 | 0 | 4 mo | HOM |
| 1013609015 | 1 | TMv | 1 | 0.111060313 | 5.81E-06 | 1 | 5.23E-05 | 9.004116529 | 4 mo | HOM |
| 1013608106 | 2 | TMv | 1 | 0.111060313 | 0 | 0 | 0 | 0 | 4 mo | HOM |
| 1011937543 | 1 | TMv | 1 | 0.111060313 | 0.000208916 | 5 | 0.001881105 | 45.02058265 | 8 mo | HOM |
| 1010419538 | 2 | TMv | 1 | 0.111060313 | 6.59E-22 | 0 | 5.94E-21 | 0 | 4 mo | WILD |
| 1010419187 | 1 | TMv | 1 | 0.111060313 | 9.7E-05 | 2 | 0.000873196 | 18.00823306 | 8 mo | HOM |
| 1013978366 | 1 | TMv | 1 | 0.111060313 | 8.84E-07 | 0 | 7.96E-06 | 0 | 2 mo | HOM |
| 1014416155 | 2 | TMv | 1 | 0.111060313 | 0 | 0 | 0 | 0 | 2 mo | HOM |
| 1013978065 | 2 | TMv | 1 | 0.111060313 | 0 | 0 | 0 | 0 | 2 mo | HOM |
| 1009934102 | 2 | TMv | 1 | 0.111060313 | 1.03E-07 | 0 | 9.27E-07 | 0 | 4 mo | WILD |
| 1009322937 | 2 | TMv | 1 | 0.111060313 | 0 | 0 | 0 | 0 | 8 mo | WILD |
| 1011080637 | 2 | TMv | 1 | 0.111060313 | 2.3E-06 | 0 | 2.07E-05 | 0 | 8 mo | HOM |
| 1013609713 | 2 | TMv | 1 | 0.111060313 | 0 | 0 | 0 | 0 | 2 mo | WILD |
| 1010418842 | 1 | TMv | 1 | 0.111060313 | 1.77E-05 | 2 | 0.000159614 | 18.00823306 | 8 mo | HOM |
| 1013288213 | 2 | TMv | 1 | 0.111060313 | 0 | 0 | 0 | 0 | 4 mo | HOM |
| 1008976060 | 2 | TMv | 1 | 0.111060313 | 0 | 0 | 0 | 0 | 8 mo | WILD |
| 1013446243 | 1 | TMv | 1 | 0.111060313 | 0 | 0 | 0 | 0 | 4 mo | HOM |
| 1013288951 | 2 | TMv | 1 | 0.111060313 | 1.25E-06 | 0 | 1.13E-05 | 0 | 4 mo | HOM |
| 1013978665 | 1 | SSp-m6b | 2 | 0.11770733 | 0 | 0 | 0 | 0 | 2 mo | HOM |
| 1013445609 | 1 | SSp-m6b | 2 | 0.11770733 | 0 | 0 | 0 | 0 | 4 mo | HOM |
| 1013609015 | 1 | SSp-m6b | 2 | 0.11770733 | 0 | 0 | 0 | 0 | 4 mo | HOM |
| 1013608106 | 2 | SSp-m6b | 2 | 0.11770733 | 0 | 0 | 0 | 0 | 4 mo | HOM |
| 1011937543 | 1 | SSp-m6b | 2 | 0.11770733 | 0.000108446 | 3 | 0.000921317 | 25.486943 | 8 mo | HOM |
| 1010419538 | 2 | SSp-m6b | 2 | 0.11770733 | 0 | 0 | 0 | 0 | 4 mo | WILD |
| 1010419187 | 1 | SSp-m6b | 2 | 0.11770733 | 1.39E-05 | 1 | 0.000118371 | 8.495647666 | 8 mo | HOM |
| 1013978366 | 1 | SSp-m6b | 2 | 0.11770733 | 0 | 0 | 0 | 0 | 2 mo | HOM |
| 1014416155 | 2 | SSp-m6b | 2 | 0.11770733 | 0 | 0 | 0 | 0 | 2 mo | HOM |
| 1013978065 | 2 | SSp-m6b | 2 | 0.11770733 | 0 | 0 | 0 | 0 | 2 mo | HOM |
| 1009934102 | 2 | SSp-m6b | 2 | 0.11770733 | 0 | 0 | 0 | 0 | 4 mo | WILD |
| 1009322937 | 2 | SSp-m6b | 2 | 0.11770733 | 0 | 0 | 0 | 0 | 8 mo | WILD |
| 1011080637 | 2 | SSp-m6b | 2 | 0.11770733 | 1.48E-05 | 1 | 0.000125898 | 8.495647666 | 8 mo | HOM |
| 1013609713 | 2 | SSp-m6b | 2 | 0.11770733 | 0 | 0 | 0 | 0 | 2 mo | WILD |
| 1010418842 | 1 | SSp-m6b | 2 | 0.11770733 | 9.11E-05 | 6 | 0.000774211 | 50.973886 | 8 mo | HOM |
| 1013288213 | 2 | SSp-m6b | 2 | 0.11770733 | 3.39E-06 | 1 | 2.88E-05 | 8.495647666 | 4 mo | HOM |
| 1008976060 | 2 | SSp-m6b | 2 | 0.11770733 | 0 | 0 | 0 | 0 | 8 mo | WILD |
| 1013446243 | 1 | SSp-m6b | 2 | 0.11770733 | 6.64E-06 | 1 | 5.64E-05 | 8.495647666 | 4 mo | HOM |
| 1013288951 | 2 | SSp-m6b | 2 | 0.11770733 | 0 | 0 | 0 | 0 | 4 mo | HOM |
| 1013978665 | 1 | IC | 4 | 4.579933254 | 6.81E-05 | 6 | 1.49E-05 | 1.31006276 | 2 mo | HOM |
| 1013445609 | 1 | IC | 4 | 4.579933254 | 0 | 0 | 0 | 0 | 4 mo | HOM |
| 1013609015 | 1 | IC | 4 | 4.579933254 | 0.000121693 | 5 | 2.66E-05 | 1.091718967 | 4 mo | HOM |
| 1013608106 | 2 | IC | 4 | 4.579933254 | 0 | 0 | 0 | 0 | 4 mo | HOM |
| 1011937543 | 1 | IC | 4 | 4.579933254 | 0.011545672 | 59 | 0.002520926 | 12.88228381 | 8 mo | HOM |
| 1010419538 | 2 | IC | 4 | 4.579933254 | 0 | 0 | 0 | 0 | 4 mo | WILD |
| 1010419187 | 1 | IC | 4 | 4.579933254 | 0.001674213 | 41 | 0.000365554 | 8.952095527 | 8 mo | HOM |
| 1013978366 | 1 | IC | 4 | 4.579933254 | 3.95E-05 | 2 | 8.62E-06 | 0.436687587 | 2 mo | HOM |
| 1014416155 | 2 | IC | 4 | 4.579933254 | 0 | 0 | 0 | 0 | 2 mo | HOM |
| 1013978065 | 2 | IC | 4 | 4.579933254 | 0 | 0 | 0 | 0 | 2 mo | HOM |
| 1009934102 | 2 | IC | 4 | 4.579933254 | 0 | 0 | 0 | 0 | 4 mo | WILD |
| 1009322937 | 2 | IC | 4 | 4.579933254 | 2.55E-07 | 0 | 5.56E-08 | 0 | 8 mo | WILD |
| 1011080637 | 2 | IC | 4 | 4.579933254 | 0.00061267 | 27 | 0.000133773 | 5.89528242 | 8 mo | HOM |
| 1013609713 | 2 | IC | 4 | 4.579933254 | 6.94E-07 | 0 | 1.52E-07 | 0 | 2 mo | WILD |
| 1010418842 | 1 | IC | 4 | 4.579933254 | 0.002044139 | 62 | 0.000446325 | 13.53731519 | 8 mo | HOM |
| 1013288213 | 2 | IC | 4 | 4.579933254 | 0 | 0 | 0 | 0 | 4 mo | HOM |
| 1008976060 | 2 | IC | 4 | 4.579933254 | 0.000225556 | 18 | 4.92E-05 | 3.93018828 | 8 mo | WILD |
| 1013446243 | 1 | IC | 4 | 4.579933254 | 9.31E-06 | 1 | 2.03E-06 | 0.218343793 | 4 mo | HOM |
| 1013288951 | 2 | IC | 4 | 4.579933254 | 3.35E-05 | 2 | 7.32E-06 | 0.436687587 | 4 mo | HOM |

| | | | | | | | | | | |
|---|---|---|---|---|---|---|---|---|---|---|
| **1013978665** | 1 | int | 6 | 2.160672326 | 0 | 0 | 0 | 0 | 2 mo | HOM |
| **1013445609** | 1 | int | 6 | 2.160672326 | 0 | 0 | 0 | 0 | 4 mo | HOM |
| **1013609015** | 1 | int | 6 | 2.160672326 | 0 | 0 | 0 | 0 | 4 mo | HOM |
| **1013608106** | 2 | int | 6 | 2.160672326 | 0 | 0 | 0 | 0 | 4 mo | HOM |
| **1011937543** | 1 | int | 6 | 2.160672326 | 0.001794354 | 31 | 0.000830461 | 14.34738605 | 8 mo | HOM |
| **1010419538** | 2 | int | 6 | 2.160672326 | 0 | 0 | 0 | 0 | 4 mo | WILD |
| **1010419187** | 1 | int | 6 | 2.160672326 | 1.18E-05 | 3 | 5.44E-06 | 1.388456715 | 8 mo | HOM |
| **1013978366** | 1 | int | 6 | 2.160672326 | 0 | 0 | 0 | 0 | 2 mo | HOM |
| **1014416155** | 2 | int | 6 | 2.160672326 | 0 | 0 | 0 | 0 | 2 mo | HOM |
| **1013978065** | 2 | int | 6 | 2.160672326 | 0 | 0 | 0 | 0 | 2 mo | HOM |
| **1009934102** | 2 | int | 6 | 2.160672326 | 0 | 0 | 0 | 0 | 4 mo | WILD |
| **1009322937** | 2 | int | 6 | 2.160672326 | 0 | 0 | 0 | 0 | 8 mo | WILD |
| **1011080637** | 2 | int | 6 | 2.160672326 | 4.16E-05 | 5 | 1.93E-05 | 2.314094525 | 8 mo | HOM |
| **1013609713** | 2 | int | 6 | 2.160672326 | 0 | 0 | 0 | 0 | 2 mo | WILD |
| **1010418842** | 1 | int | 6 | 2.160672326 | 0.000146006 | 10 | 6.76E-05 | 4.628189049 | 8 mo | HOM |
| **1013288213** | 2 | int | 6 | 2.160672326 | 0 | 0 | 0 | 0 | 4 mo | HOM |
| **1008976060** | 2 | int | 6 | 2.160672326 | 2E-05 | 3 | 9.26E-06 | 1.388456715 | 8 mo | WILD |
| **1013446243** | 1 | int | 6 | 2.160672326 | 1.79E-05 | 3 | 8.28E-06 | 1.388456715 | 4 mo | HOM |
| **1013288951** | 2 | int | 6 | 2.160672326 | 3.09E-05 | 1 | 1.43E-05 | 0.462818905 | 4 mo | HOM |
| **1013978665** | 1 | PSV | 7 | 1.139507152 | 4.98E-06 | 1 | 4.37E-06 | 0.877572377 | 2 mo | HOM |
| **1013445609** | 1 | PSV | 7 | 1.139507152 | 0 | 0 | 0 | 0 | 4 mo | HOM |
| **1013609015** | 1 | PSV | 7 | 1.139507152 | 0 | 0 | 0 | 0 | 4 mo | HOM |
| **1013608106** | 2 | PSV | 7 | 1.139507152 | 0 | 0 | 0 | 0 | 4 mo | HOM |
| **1011937543** | 1 | PSV | 7 | 1.139507152 | 0.001256449 | 5 | 0.001102625 | 4.387861885 | 8 mo | HOM |
| **1010419538** | 2 | PSV | 7 | 1.139507152 | 0 | 0 | 0 | 0 | 4 mo | WILD |
| **1010419187** | 1 | PSV | 7 | 1.139507152 | 9.2E-06 | 1 | 8.07E-06 | 0.877572377 | 8 mo | HOM |
| **1013978366** | 1 | PSV | 7 | 1.139507152 | 0 | 0 | 0 | 0 | 2 mo | HOM |
| **1014416155** | 2 | PSV | 7 | 1.139507152 | 0 | 0 | 0 | 0 | 2 mo | HOM |
| **1013978065** | 2 | PSV | 7 | 1.139507152 | 0 | 0 | 0 | 0 | 2 mo | HOM |
| **1009934102** | 2 | PSV | 7 | 1.139507152 | 0 | 0 | 0 | 0 | 4 mo | WILD |
| **1009322937** | 2 | PSV | 7 | 1.139507152 | 0 | 0 | 0 | 0 | 8 mo | WILD |
| **1011080637** | 2 | PSV | 7 | 1.139507152 | 0 | 0 | 0 | 0 | 8 mo | HOM |
| **1013609713** | 2 | PSV | 7 | 1.139507152 | 0 | 0 | 0 | 0 | 2 mo | WILD |
| **1010418842** | 1 | PSV | 7 | 1.139507152 | 0 | 0 | 0 | 0 | 8 mo | HOM |
| **1013288213** | 2 | PSV | 7 | 1.139507152 | 0 | 0 | 0 | 0 | 4 mo | HOM |
| **1008976060** | 2 | PSV | 7 | 1.139507152 | 1.5E-05 | 2 | 1.32E-05 | 1.755144754 | 8 mo | WILD |
| **1013446243** | 1 | PSV | 7 | 1.139507152 | 0 | 0 | 0 | 0 | 4 mo | HOM |
| **1013288951** | 2 | PSV | 7 | 1.139507152 | 0 | 0 | 0 | 0 | 4 mo | HOM |
| **1013978665** | 1 | grey | 8 | 459.5709405 | 0.012515766 | 536 | 2.72E-05 | 1.166305249 | 2 mo | HOM |
| **1013445609** | 1 | grey | 8 | 459.8042894 | 0.018187101 | 1216 | 3.96E-05 | 2.644603428 | 4 mo | HOM |
| **1013609015** | 1 | grey | 8 | 457.5716822 | 0.107002434 | 2481 | 0.000233848 | 5.422101272 | 4 mo | HOM |
| **1013608106** | 2 | grey | 8 | 459.4682352 | 8.41E-06 | 1 | 1.83E-08 | 0.002176429 | 4 mo | HOM |
| **1011937543** | 1 | grey | 8 | 460.2129654 | 0.63739478 | 7584 | 0.001385 | 16.47932712 | 8 mo | HOM |
| **1010419538** | 2 | grey | 8 | 460.5067235 | 0.004426211 | 253 | 9.61E-06 | 0.549394802 | 4 mo | WILD |
| **1010419187** | 1 | grey | 8 | 456.7454773 | 0.494820557 | 12092 | 0.001083362 | 26.47426324 | 8 mo | HOM |
| **1013978366** | 1 | grey | 8 | 461.9075522 | 0.029915234 | 734 | 6.48E-05 | 1.589062566 | 2 mo | HOM |
| **1014416155** | 2 | grey | 8 | 460.7324971 | 0.000107661 | 11 | 2.34E-07 | 0.023875025 | 2 mo | HOM |
| **1013978065** | 2 | grey | 8 | 462.3636266 | 0.004052296 | 204 | 8.76E-06 | 0.441211177 | 2 mo | HOM |
| **1009934102** | 2 | grey | 8 | 458.8259693 | 0.006937623 | 373 | 1.51E-05 | 0.812944395 | 4 mo | WILD |
| **1009322937** | 2 | grey | 8 | 456.4024717 | 0.008798231 | 456 | 1.93E-05 | 0.999118165 | 8 mo | WILD |
| **1011080637** | 2 | grey | 8 | 456.015054 | 0.298703692 | 9178 | 0.00065503 | 20.12652854 | 8 mo | HOM |
| **1013609713** | 2 | grey | 8 | 463.0745877 | 0.002492172 | 132 | 5.38E-06 | 0.285051271 | 2 mo | WILD |
| **1010418842** | 1 | grey | 8 | 458.605479 | 0.462122802 | 12220 | 0.00100767 | 26.64599652 | 8 mo | HOM |
| **1013288213** | 2 | grey | 8 | 460.2699978 | 0.06259359 | 1239 | 0.000135993 | 2.691898246 | 4 mo | HOM |
| **1008976060** | 2 | grey | 8 | 461.4537169 | 0.053953091 | 1773 | 0.00011692 | 3.842205481 | 8 mo | WILD |
| **1013446243** | 1 | grey | 8 | 459.2074847 | 0.048017947 | 2117 | 0.000104567 | 4.610116495 | 4 mo | HOM |
| **1013288951** | 2 | grey | 8 | 459.0154477 | 0.10536934 | 2626 | 0.000229555 | 5.720940359 | 4 mo | HOM |
| **1013978665** | 1 | SSp-tr6a | 9 | 0.239945591 | 0 | 0 | 0 | 0 | 2 mo | HOM |
| **1013445609** | 1 | SSp-tr6a | 9 | 0.239945591 | 3.71E-06 | 1 | 1.55E-05 | 4.167611478 | 4 mo | HOM |

| | | | | | | | | | | |
|---|---|---|---|---|---|---|---|---|---|---|
| 1013609015 | 1 | SSp-tr6a | 9 | 0.239945591 | 0.000121308 | 3 | 0.000505565 | 12.50283443 | 4 mo | HOM |
| 1013608106 | 2 | SSp-tr6a | 9 | 0.239945591 | 0 | 0 | 0 | 0 | 4 mo | HOM |
| 1011937543 | 1 | SSp-tr6a | 9 | 0.239945591 | 0.000311758 | 6 | 0.001299285 | 25.00566887 | 8 mo | HOM |
| 1010419538 | 2 | SSp-tr6a | 9 | 0.239945591 | 0 | 0 | 0 | 0 | 4 mo | WILD |
| 1010419187 | 1 | SSp-tr6a | 9 | 0.239945591 | 0.00065845 | 12 | 0.002744165 | 50.01133774 | 8 mo | HOM |
| 1013978366 | 1 | SSp-tr6a | 9 | 0.239945591 | 0 | 0 | 0 | 0 | 2 mo | HOM |
| 1014416155 | 2 | SSp-tr6a | 9 | 0.239945591 | 0 | 0 | 0 | 0 | 2 mo | HOM |
| 1013978065 | 2 | SSp-tr6a | 9 | 0.239945591 | 0 | 0 | 0 | 0 | 2 mo | HOM |
| 1009934102 | 2 | SSp-tr6a | 9 | 0.239945591 | 0 | 0 | 0 | 0 | 4 mo | WILD |
| 1009322937 | 2 | SSp-tr6a | 9 | 0.239945591 | 0 | 0 | 0 | 0 | 8 mo | WILD |
| 1011080637 | 2 | SSp-tr6a | 9 | 0.239945591 | 0.000229414 | 6 | 0.000956108 | 25.00566887 | 8 mo | HOM |
| 1013609713 | 2 | SSp-tr6a | 9 | 0.239945591 | 0 | 0 | 0 | 0 | 2 mo | WILD |
| 1010418842 | 1 | SSp-tr6a | 9 | 0.239945591 | 0.000195465 | 10 | 0.000814621 | 41.67611478 | 8 mo | HOM |
| 1013288213 | 2 | SSp-tr6a | 9 | 0.239945591 | 6.82E-05 | 2 | 0.000284218 | 8.335222956 | 4 mo | HOM |
| 1008976060 | 2 | SSp-tr6a | 9 | 0.239945591 | 0 | 0 | 0 | 0 | 8 mo | WILD |
| 1013446243 | 1 | SSp-tr6a | 9 | 0.239945591 | 7.57E-05 | 5 | 0.000315471 | 20.83805739 | 4 mo | HOM |
| 1013288951 | 2 | SSp-tr6a | 9 | 0.239945591 | 8.64E-05 | 2 | 0.000359987 | 8.335222956 | 4 mo | HOM |
| 1013978665 | 1 | SCig | 10 | 2.099087558 | 8.02E-06 | 1 | 3.82E-06 | 0.476397469 | 2 mo | HOM |
| 1013445609 | 1 | SCig | 10 | 2.099087558 | 0 | 0 | 0 | 0 | 4 mo | HOM |
| 1013609015 | 1 | SCig | 10 | 2.099087558 | 1.36E-07 | 0 | 6.47E-08 | 0 | 4 mo | HOM |
| 1013608106 | 2 | SCig | 10 | 2.099087558 | 0 | 0 | 0 | 0 | 4 mo | HOM |
| 1011937543 | 1 | SCig | 10 | 2.099087558 | 0.003097938 | 52 | 0.00147585 | 24.77266839 | 8 mo | HOM |
| 1010419538 | 2 | SCig | 10 | 2.099087558 | 0 | 0 | 0 | 0 | 4 mo | WILD |
| 1010419187 | 1 | SCig | 10 | 2.099087558 | 0.000291862 | 13 | 0.000139042 | 6.193167097 | 8 mo | HOM |
| 1013978366 | 1 | SCig | 10 | 2.099087558 | 0 | 0 | 0 | 0 | 2 mo | HOM |
| 1014416155 | 2 | SCig | 10 | 2.099087558 | 0 | 0 | 0 | 0 | 2 mo | HOM |
| 1013978065 | 2 | SCig | 10 | 2.099087558 | 0 | 0 | 0 | 0 | 2 mo | HOM |
| 1009934102 | 2 | SCig | 10 | 2.099087558 | 0 | 0 | 0 | 0 | 4 mo | WILD |
| 1009322937 | 2 | SCig | 10 | 2.099087558 | 1.54E-06 | 0 | 7.36E-07 | 0 | 8 mo | WILD |
| 1011080637 | 2 | SCig | 10 | 2.099087558 | 0.000129235 | 4 | 6.16E-05 | 1.905589876 | 8 mo | HOM |
| 1013609713 | 2 | SCig | 10 | 2.099087558 | 0 | 0 | 0 | 0 | 2 mo | WILD |
| 1010418842 | 1 | SCig | 10 | 2.099087558 | 0.000186236 | 6 | 8.87E-05 | 2.858384814 | 8 mo | HOM |
| 1013288213 | 2 | SCig | 10 | 2.099087558 | 0 | 0 | 0 | 0 | 4 mo | HOM |
| 1008976060 | 2 | SCig | 10 | 2.099087558 | 5.83E-05 | 6 | 2.78E-05 | 2.858384814 | 8 mo | WILD |
| 1013446243 | 1 | SCig | 10 | 2.099087558 | 7.56E-06 | 1 | 3.6E-06 | 0.476397469 | 4 mo | HOM |
| 1013288951 | 2 | SCig | 10 | 2.099087558 | 0 | 0 | 0 | 0 | 4 mo | HOM |
| 1013978665 | 1 | IF | 12 | 0.089419414 | 0 | 0 | 0 | 0 | 2 mo | HOM |
| 1013445609 | 1 | IF | 12 | 0.089419414 | 0 | 0 | 0 | 0 | 4 mo | HOM |
| 1013609015 | 1 | IF | 12 | 0.089419414 | 4.73E-06 | 1 | 5.3E-05 | 11.18325378 | 4 mo | HOM |
| 1013608106 | 2 | IF | 12 | 0.089419414 | 0 | 0 | 0 | 0 | 4 mo | HOM |
| 1011937543 | 1 | IF | 12 | 0.089419414 | 0 | 0 | 0 | 0 | 8 mo | HOM |
| 1010419538 | 2 | IF | 12 | 0.089419414 | 0 | 0 | 0 | 0 | 4 mo | WILD |
| 1010419187 | 1 | IF | 12 | 0.089419414 | 0 | 0 | 0 | 0 | 8 mo | HOM |
| 1013978366 | 1 | IF | 12 | 0.089419414 | 0 | 0 | 0 | 0 | 2 mo | HOM |
| 1014416155 | 2 | IF | 12 | 0.089419414 | 0 | 0 | 0 | 0 | 2 mo | HOM |
| 1013978065 | 2 | IF | 12 | 0.089419414 | 0 | 0 | 0 | 0 | 2 mo | HOM |
| 1009934102 | 2 | IF | 12 | 0.089419414 | 0 | 0 | 0 | 0 | 4 mo | WILD |
| 1009322937 | 2 | IF | 12 | 0.089419414 | 0 | 0 | 0 | 0 | 8 mo | WILD |
| 1011080637 | 2 | IF | 12 | 0.089419414 | 6.75E-10 | 0 | 7.54E-09 | 0 | 8 mo | HOM |
| 1013609713 | 2 | IF | 12 | 0.089419414 | 0 | 0 | 0 | 0 | 2 mo | WILD |
| 1010418842 | 1 | IF | 12 | 0.089419414 | 1.19E-06 | 0 | 1.33E-05 | 0 | 8 mo | HOM |
| 1013288213 | 2 | IF | 12 | 0.089419414 | 0 | 0 | 0 | 0 | 4 mo | HOM |
| 1008976060 | 2 | IF | 12 | 0.089419414 | 4.02E-06 | 0 | 4.5E-05 | 0 | 8 mo | WILD |
| 1013446243 | 1 | IF | 12 | 0.089419414 | 0 | 0 | 0 | 0 | 4 mo | HOM |
| 1013288951 | 2 | IF | 12 | 0.089419414 | 0 | 0 | 0 | 0 | 4 mo | HOM |
| 1013978665 | 1 | PT | 15 | 0.236902299 | 0 | 0 | 0 | 0 | 2 mo | HOM |
| 1013445609 | 1 | PT | 15 | 0.236902299 | 2.25E-07 | 0 | 9.48E-07 | 0 | 4 mo | HOM |
| 1013609015 | 1 | PT | 15 | 0.236902299 | 7.05E-06 | 1 | 2.97E-05 | 4.221149408 | 4 mo | HOM |
| 1013608106 | 2 | PT | 15 | 0.236902299 | 0 | 0 | 0 | 0 | 4 mo | HOM |

| | | | | | | | | | | |
|---|---|---|---|---|---|---|---|---|---|---|
| **1011937543** | 1 | PT | 15 | 0.236902299 | 2.66E-05 | 1 | 0.000112159 | 4.221149408 | 8 mo | HOM |
| **1010419538** | 2 | PT | 15 | 0.236902299 | 0 | 0 | 0 | 0 | 4 mo | WILD |
| **1010419187** | 1 | PT | 15 | 0.236902299 | 0.000120737 | 2 | 0.000509648 | 8.442298816 | 8 mo | HOM |
| **1013978366** | 1 | PT | 15 | 0.236902299 | 1.12E-06 | 0 | 4.74E-06 | 0 | 2 mo | HOM |
| **1014416155** | 2 | PT | 15 | 0.236902299 | 0 | 0 | 0 | 0 | 2 mo | HOM |
| **1013978065** | 2 | PT | 15 | 0.236902299 | 0 | 0 | 0 | 0 | 2 mo | HOM |
| **1009934102** | 2 | PT | 15 | 0.236902299 | 0 | 0 | 0 | 0 | 4 mo | WILD |
| **1009322937** | 2 | PT | 15 | 0.236902299 | 0 | 0 | 0 | 0 | 8 mo | WILD |
| **1011080637** | 2 | PT | 15 | 0.236902299 | 0.00010513 | 3 | 0.00044377 | 12.66344822 | 8 mo | HOM |
| **1013609713** | 2 | PT | 15 | 0.236902299 | 0 | 0 | 0 | 0 | 2 mo | WILD |
| **1010418842** | 1 | PT | 15 | 0.236902299 | 7.14E-05 | 1 | 0.000301445 | 4.221149408 | 8 mo | HOM |
| **1013288213** | 2 | PT | 15 | 0.236902299 | 9.91E-08 | 0 | 4.18E-07 | 0 | 4 mo | HOM |
| **1008976060** | 2 | PT | 15 | 0.236902299 | 1.48E-05 | 1 | 6.23E-05 | 4.221149408 | 8 mo | WILD |
| **1013446243** | 1 | PT | 15 | 0.236902299 | 0 | 0 | 0 | 0 | 4 mo | HOM |
| **1013288951** | 2 | PT | 15 | 0.236902299 | 0 | 0 | 0 | 0 | 4 mo | HOM |
| **1013978665** | 1 | SCiw | 17 | 2.152974544 | 0 | 0 | 0 | 0 | 2 mo | HOM |
| **1013445609** | 1 | SCiw | 17 | 2.152974544 | 0 | 0 | 0 | 0 | 4 mo | HOM |
| **1013609015** | 1 | SCiw | 17 | 2.152974544 | 5.77E-07 | 1 | 2.68E-07 | 0.464473676 | 4 mo | HOM |
| **1013608106** | 2 | SCiw | 17 | 2.152974544 | 0 | 0 | 0 | 0 | 4 mo | HOM |
| **1011937543** | 1 | SCiw | 17 | 2.152974544 | 0.002384174 | 46 | 0.001107386 | 21.36578908 | 8 mo | HOM |
| **1010419538** | 2 | SCiw | 17 | 2.152974544 | 0 | 0 | 0 | 0 | 4 mo | WILD |
| **1010419187** | 1 | SCiw | 17 | 2.152974544 | 0.000289755 | 10 | 0.000134584 | 4.644736756 | 8 mo | HOM |
| **1013978366** | 1 | SCiw | 17 | 2.152974544 | 0 | 0 | 0 | 0 | 2 mo | HOM |
| **1014416155** | 2 | SCiw | 17 | 2.152974544 | 0 | 0 | 0 | 0 | 2 mo | HOM |
| **1013978065** | 2 | SCiw | 17 | 2.152974544 | 0 | 0 | 0 | 0 | 2 mo | HOM |
| **1009934102** | 2 | SCiw | 17 | 2.152974544 | 0 | 0 | 0 | 0 | 4 mo | WILD |
| **1009322937** | 2 | SCiw | 17 | 2.152974544 | 0 | 0 | 0 | 0 | 8 mo | WILD |
| **1011080637** | 2 | SCiw | 17 | 2.152974544 | 5.71E-05 | 3 | 2.65E-05 | 1.393421027 | 8 mo | HOM |
| **1013609713** | 2 | SCiw | 17 | 2.152974544 | 0 | 0 | 0 | 0 | 2 mo | WILD |
| **1010418842** | 1 | SCiw | 17 | 2.152974544 | 0.000277696 | 12 | 0.000128982 | 5.573684108 | 8 mo | HOM |
| **1013288213** | 2 | SCiw | 17 | 2.152974544 | 0 | 0 | 0 | 0 | 4 mo | HOM |
| **1008976060** | 2 | SCiw | 17 | 2.152974544 | 4.37E-05 | 5 | 2.03E-05 | 2.322368378 | 8 mo | WILD |
| **1013446243** | 1 | SCiw | 17 | 2.152974544 | 1.42E-06 | 0 | 6.6E-07 | 0 | 4 mo | HOM |
| **1013288951** | 2 | SCiw | 17 | 2.152974544 | 0 | 0 | 0 | 0 | 4 mo | HOM |
| **1013978665** | 1 | IG | 19 | 0.129488978 | 0 | 0 | 0 | 0 | 2 mo | HOM |
| **1013445609** | 1 | IG | 19 | 0.129488978 | 9.42E-06 | 1 | 7.27E-05 | 7.722665025 | 4 mo | HOM |
| **1013609015** | 1 | IG | 19 | 0.129488978 | 4.13E-05 | 1 | 0.000319141 | 7.722665025 | 4 mo | HOM |
| **1013608106** | 2 | IG | 19 | 0.129488978 | 0 | 0 | 0 | 0 | 4 mo | HOM |
| **1011937543** | 1 | IG | 19 | 0.129488978 | 8.3E-06 | 0 | 6.41E-05 | 0 | 8 mo | HOM |
| **1010419538** | 2 | IG | 19 | 0.129488978 | 0 | 0 | 0 | 0 | 4 mo | WILD |
| **1010419187** | 1 | IG | 19 | 0.129488978 | 0.000250019 | 7 | 0.001930812 | 54.05865518 | 8 mo | HOM |
| **1013978366** | 1 | IG | 19 | 0.129488978 | 2.02E-05 | 1 | 0.00015581 | 7.722665025 | 2 mo | HOM |
| **1014416155** | 2 | IG | 19 | 0.129488978 | 0 | 0 | 0 | 0 | 2 mo | HOM |
| **1013978065** | 2 | IG | 19 | 0.129488978 | 0 | 0 | 0 | 0 | 2 mo | HOM |
| **1009934102** | 2 | IG | 19 | 0.129488978 | 0.000119135 | 2 | 0.00092004 | 15.44533005 | 4 mo | WILD |
| **1009322937** | 2 | IG | 19 | 0.129488978 | 0.000323417 | 1 | 0.002497645 | 7.722665025 | 8 mo | WILD |
| **1011080637** | 2 | IG | 19 | 0.129488978 | 0.00095164 | 11 | 0.007349199 | 84.94931528 | 8 mo | HOM |
| **1013609713** | 2 | IG | 19 | 0.129488978 | 0 | 0 | 0 | 0 | 2 mo | WILD |
| **1010418842** | 1 | IG | 19 | 0.129488978 | 0.001332357 | 2 | 0.01028935 | 15.44533005 | 8 mo | HOM |
| **1013288213** | 2 | IG | 19 | 0.129488978 | 0.002943961 | 4 | 0.022735225 | 30.8906601 | 4 mo | HOM |
| **1008976060** | 2 | IG | 19 | 0.129488978 | 1.05E-05 | 1 | 8.09E-05 | 7.722665025 | 8 mo | WILD |
| **1013446243** | 1 | IG | 19 | 0.129488978 | 3.82E-05 | 2 | 0.000295337 | 15.44533005 | 4 mo | HOM |
| **1013288951** | 2 | IG | 19 | 0.129488978 | 0.000270894 | 17 | 0.002092025 | 131.2853054 | 4 mo | HOM |
| **1013978665** | 1 | ENTl2 | 20 | 1.652909395 | 0 | 0 | 0 | 0 | 2 mo | HOM |
| **1013445609** | 1 | ENTl2 | 20 | 1.652909395 | 1.06E-05 | 1 | 6.42E-06 | 0.604993839 | 4 mo | HOM |
| **1013609015** | 1 | ENTl2 | 20 | 1.652909395 | 2.4E-05 | 2 | 1.45E-05 | 1.209987677 | 4 mo | HOM |
| **1013608106** | 2 | ENTl2 | 20 | 1.652909395 | 0 | 0 | 0 | 0 | 4 mo | HOM |
| **1011937543** | 1 | ENTl2 | 20 | 1.652909395 | 0.001290681 | 33 | 0.000780854 | 19.96479668 | 8 mo | HOM |
| **1010419538** | 2 | ENTl2 | 20 | 1.652909395 | 0 | 0 | 0 | 0 | 4 mo | WILD |

| | | | | | | | | | | |
|---|---|---|---|---|---|---|---|---|---|---|
| **1010419187** | 1 | ENTl2 | 20 | 1.652909395 | 0.000632602 | 25 | 0.000382721 | 15.12484597 | 8 mo | HOM |
| **1013978366** | 1 | ENTl2 | 20 | 1.652909395 | 0 | 0 | 0 | 0 | 2 mo | HOM |
| **1014416155** | 2 | ENTl2 | 20 | 1.652909395 | 0 | 0 | 0 | 0 | 2 mo | HOM |
| **1013978065** | 2 | ENTl2 | 20 | 1.652909395 | 0 | 0 | 0 | 0 | 2 mo | HOM |
| **1009934102** | 2 | ENTl2 | 20 | 1.652909395 | 0 | 0 | 0 | 0 | 4 mo | WILD |
| **1009322937** | 2 | ENTl2 | 20 | 1.652909395 | 0 | 0 | 0 | 0 | 8 mo | WILD |
| **1011080637** | 2 | ENTl2 | 20 | 1.652909395 | 0.00049501 | 25 | 0.000299478 | 15.12484597 | 8 mo | HOM |
| **1013609713** | 2 | ENTl2 | 20 | 1.652909395 | 0 | 0 | 0 | 0 | 2 mo | WILD |
| **1010418842** | 1 | ENTl2 | 20 | 1.652909395 | 0.000447902 | 17 | 0.000270978 | 10.28489526 | 8 mo | HOM |
| **1013288213** | 2 | ENTl2 | 20 | 1.652909395 | 4.81E-06 | 1 | 2.91E-06 | 0.604993839 | 4 mo | HOM |
| **1008976060** | 2 | ENTl2 | 20 | 1.652909395 | 0 | 0 | 0 | 0 | 8 mo | WILD |
| **1013446243** | 1 | ENTl2 | 20 | 1.652909395 | 8.58E-05 | 6 | 5.19E-05 | 3.629963032 | 4 mo | HOM |
| **1013288951** | 2 | ENTl2 | 20 | 1.652909395 | 7.27E-05 | 4 | 4.4E-05 | 2.419975355 | 4 mo | HOM |
| **1013978665** | 1 | lotg | 21 | 1.094051577 | 8.51E-07 | 0 | 7.78E-07 | 0 | 2 mo | HOM |
| **1013445609** | 1 | lotg | 21 | 1.094051577 | 3.89E-06 | 0 | 3.56E-06 | 0 | 4 mo | HOM |
| **1013609015** | 1 | lotg | 21 | 1.094051577 | 3.14E-06 | 0 | 2.87E-06 | 0 | 4 mo | HOM |
| **1013608106** | 2 | lotg | 21 | 1.094051577 | 0 | 0 | 0 | 0 | 4 mo | HOM |
| **1011937543** | 1 | lotg | 21 | 1.094051577 | 0.000242193 | 3 | 0.000221373 | 2.742101071 | 8 mo | HOM |
| **1010419538** | 2 | lotg | 21 | 1.094051577 | 0 | 0 | 0 | 0 | 4 mo | WILD |
| **1010419187** | 1 | lotg | 21 | 1.094051577 | 0.000197826 | 4 | 0.000180819 | 3.656134761 | 8 mo | HOM |
| **1013978366** | 1 | lotg | 21 | 1.094051577 | 0 | 0 | 0 | 0 | 2 mo | HOM |
| **1014416155** | 2 | lotg | 21 | 1.094051577 | 0 | 0 | 0 | 0 | 2 mo | HOM |
| **1013978065** | 2 | lotg | 21 | 1.094051577 | 0 | 0 | 0 | 0 | 2 mo | HOM |
| **1009934102** | 2 | lotg | 21 | 1.094051577 | 2.16E-06 | 0 | 1.97E-06 | 0 | 4 mo | WILD |
| **1009322937** | 2 | lotg | 21 | 1.094051577 | 2.27E-05 | 3 | 2.08E-05 | 2.742101071 | 8 mo | WILD |
| **1011080637** | 2 | lotg | 21 | 1.094051577 | 0.000246846 | 4 | 0.000225626 | 3.656134761 | 8 mo | HOM |
| **1013609713** | 2 | lotg | 21 | 1.094051577 | 0 | 0 | 0 | 0 | 2 mo | WILD |
| **1010418842** | 1 | lotg | 21 | 1.094051577 | 0.0002084 | 3 | 0.000190484 | 2.742101071 | 8 mo | HOM |
| **1013288213** | 2 | lotg | 21 | 1.094051577 | 0.000536716 | 2 | 0.000490576 | 1.82806738 | 4 mo | HOM |
| **1008976060** | 2 | lotg | 21 | 1.094051577 | 5.61E-06 | 1 | 5.13E-06 | 0.91403369 | 8 mo | WILD |
| **1013446243** | 1 | lotg | 21 | 1.094051577 | 2.62E-06 | 0 | 2.39E-06 | 0 | 4 mo | HOM |
| **1013288951** | 2 | lotg | 21 | 1.094051577 | 3.21E-05 | 2 | 2.93E-05 | 1.82806738 | 4 mo | HOM |
| **1013978665** | 1 | PTLp | 22 | 2.531085609 | 1.22E-05 | 1 | 4.83E-06 | 0.395087387 | 2 mo | HOM |
| **1013445609** | 1 | PTLp | 22 | 2.531085609 | 0.000343615 | 24 | 0.000135758 | 9.482097292 | 4 mo | HOM |
| **1013609015** | 1 | PTLp | 22 | 2.531085609 | 0.001136382 | 38 | 0.00044897 | 15.01332071 | 4 mo | HOM |
| **1013608106** | 2 | PTLp | 22 | 2.531085609 | 0 | 0 | 0 | 0 | 4 mo | HOM |
| **1011937543** | 1 | PTLp | 22 | 2.531085609 | 0.005273596 | 58 | 0.002083531 | 22.91506846 | 8 mo | HOM |
| **1010419538** | 2 | PTLp | 22 | 2.531085609 | 0 | 0 | 0 | 0 | 4 mo | WILD |
| **1010419187** | 1 | PTLp | 22 | 2.531085609 | 0.004756183 | 144 | 0.001879108 | 56.89258375 | 8 mo | HOM |
| **1013978366** | 1 | PTLp | 22 | 2.531085609 | 3.71E-06 | 1 | 1.47E-06 | 0.395087387 | 2 mo | HOM |
| **1014416155** | 2 | PTLp | 22 | 2.531085609 | 0 | 0 | 0 | 0 | 2 mo | HOM |
| **1013978065** | 2 | PTLp | 22 | 2.531085609 | 1.46E-06 | 0 | 5.78E-07 | 0 | 2 mo | HOM |
| **1009934102** | 2 | PTLp | 22 | 2.531085609 | 6.67E-08 | 0 | 2.64E-08 | 0 | 4 mo | WILD |
| **1009322937** | 2 | PTLp | 22 | 2.531085609 | 5.18E-05 | 2 | 2.05E-05 | 0.790174774 | 8 mo | WILD |
| **1011080637** | 2 | PTLp | 22 | 2.531085609 | 0.00204383 | 79 | 0.000807492 | 31.21190359 | 8 mo | HOM |
| **1013609713** | 2 | PTLp | 22 | 2.531085609 | 1.12E-05 | 1 | 4.41E-06 | 0.395087387 | 2 mo | WILD |
| **1010418842** | 1 | PTLp | 22 | 2.531085609 | 0.004367202 | 120 | 0.001725426 | 47.41048646 | 8 mo | HOM |
| **1013288213** | 2 | PTLp | 22 | 2.531085609 | 0.000221291 | 21 | 8.74E-05 | 8.29683513 | 4 mo | HOM |
| **1008976060** | 2 | PTLp | 22 | 2.531085609 | 6.45E-05 | 1 | 2.55E-05 | 0.395087387 | 8 mo | WILD |
| **1013446243** | 1 | PTLp | 22 | 2.531085609 | 0.000773566 | 35 | 0.000305626 | 13.82805855 | 4 mo | HOM |
| **1013288951** | 2 | PTLp | 22 | 2.531085609 | 0.001111247 | 31 | 0.00043904 | 12.247709 | 4 mo | HOM |
| **1013978665** | 1 | AAA | 23 | 0.473452386 | 8.08E-07 | 0 | 1.71E-06 | 0 | 2 mo | HOM |
| **1013445609** | 1 | AAA | 23 | 0.50931235 | 0 | 0 | 0 | 0 | 4 mo | HOM |
| **1013609015** | 1 | AAA | 23 | 0.50931235 | 0 | 0 | 0 | 0 | 4 mo | HOM |
| **1013608106** | 2 | AAA | 23 | 0.50931235 | 0 | 0 | 0 | 0 | 4 mo | HOM |
| **1011937543** | 1 | AAA | 23 | 0.50931235 | 0.000464993 | 4 | 0.000912982 | 7.85372669 | 8 mo | HOM |
| **1010419538** | 2 | AAA | 23 | 0.50931235 | 0 | 0 | 0 | 0 | 4 mo | WILD |
| **1010419187** | 1 | AAA | 23 | 0.50931235 | 0.00010304 | 7 | 0.000202312 | 13.74402171 | 8 mo | HOM |
| **1013978366** | 1 | AAA | 23 | 0.50931235 | 0 | 0 | 0 | 0 | 2 mo | HOM |

| | | | | | | | | | | |
|---|---|---|---|---|---|---|---|---|---|---|
| **1014416155** | 2 | AAA | 23 | 0.50931235 | 0 | 0 | 0 | 0 | 2 mo | HOM |
| **1013978065** | 2 | AAA | 23 | 0.50931235 | 0 | 0 | 0 | 0 | 2 mo | HOM |
| **1009934102** | 2 | AAA | 23 | 0.50931235 | 0 | 0 | 0 | 0 | 4 mo | WILD |
| **1009322937** | 2 | AAA | 23 | 0.50931235 | 0 | 0 | 0 | 0 | 8 mo | WILD |
| **1011080637** | 2 | AAA | 23 | 0.50931235 | 0.000124528 | 4 | 0.000244502 | 7.85372669 | 8 mo | HOM |
| **1013609713** | 2 | AAA | 23 | 0.50931235 | 0 | 0 | 0 | 0 | 2 mo | WILD |
| **1010418842** | 1 | AAA | 23 | 0.50931235 | 0.000295347 | 10 | 0.000579893 | 19.63431673 | 8 mo | HOM |
| **1013288213** | 2 | AAA | 23 | 0.50931235 | 3.78E-06 | 0 | 7.43E-06 | 0 | 4 mo | HOM |
| **1008976060** | 2 | AAA | 23 | 0.50931235 | 1.41E-05 | 1 | 2.77E-05 | 1.963431673 | 8 mo | WILD |
| **1013446243** | 1 | AAA | 23 | 0.50931235 | 2.79E-06 | 0 | 5.47E-06 | 0 | 4 mo | HOM |
| **1013288951** | 2 | AAA | 23 | 0.50931235 | 1.41E-07 | 1 | 2.77E-07 | 1.963431673 | 4 mo | HOM |
| **1013978665** | 1 | SCdg | 26 | 1.230941701 | 0 | 0 | 0 | 0 | 2 mo | HOM |
| **1013445609** | 1 | SCdg | 26 | 1.230941701 | 0 | 0 | 0 | 0 | 4 mo | HOM |
| **1013609015** | 1 | SCdg | 26 | 1.230941701 | 0 | 0 | 0 | 0 | 4 mo | HOM |
| **1013608106** | 2 | SCdg | 26 | 1.230941701 | 0 | 0 | 0 | 0 | 4 mo | HOM |
| **1011937543** | 1 | SCdg | 26 | 1.230941701 | 0.001097196 | 27 | 0.000891347 | 21.93442629 | 8 mo | HOM |
| **1010419538** | 2 | SCdg | 26 | 1.230941701 | 0 | 0 | 0 | 0 | 4 mo | WILD |
| **1010419187** | 1 | SCdg | 26 | 1.230941701 | 3.1E-05 | 3 | 2.52E-05 | 2.437158476 | 8 mo | HOM |
| **1013978366** | 1 | SCdg | 26 | 1.230941701 | 0 | 0 | 0 | 0 | 2 mo | HOM |
| **1014416155** | 2 | SCdg | 26 | 1.230941701 | 0 | 0 | 0 | 0 | 2 mo | HOM |
| **1013978065** | 2 | SCdg | 26 | 1.230941701 | 0 | 0 | 0 | 0 | 2 mo | HOM |
| **1009934102** | 2 | SCdg | 26 | 1.230941701 | 0 | 0 | 0 | 0 | 4 mo | WILD |
| **1009322937** | 2 | SCdg | 26 | 1.230941701 | 0 | 0 | 0 | 0 | 8 mo | WILD |
| **1011080637** | 2 | SCdg | 26 | 1.230941701 | 1.03E-05 | 1 | 8.35E-06 | 0.812386159 | 8 mo | HOM |
| **1013609713** | 2 | SCdg | 26 | 1.230941701 | 0 | 0 | 0 | 0 | 2 mo | WILD |
| **1010418842** | 1 | SCdg | 26 | 1.230941701 | 3.42E-05 | 3 | 2.78E-05 | 2.437158476 | 8 mo | HOM |
| **1013288213** | 2 | SCdg | 26 | 1.230941701 | 0 | 0 | 0 | 0 | 4 mo | HOM |
| **1008976060** | 2 | SCdg | 26 | 1.230941701 | 0 | 0 | 0 | 0 | 8 mo | WILD |
| **1013446243** | 1 | SCdg | 26 | 1.230941701 | 0 | 0 | 0 | 0 | 4 mo | HOM |
| **1013288951** | 2 | SCdg | 26 | 1.230941701 | 0 | 0 | 0 | 0 | 4 mo | HOM |
| **1013978665** | 1 | IGL | 27 | 0.070487994 | 0 | 0 | 0 | 0 | 2 mo | HOM |
| **1013445609** | 1 | IGL | 27 | 0.070487994 | 0 | 0 | 0 | 0 | 4 mo | HOM |
| **1013609015** | 1 | IGL | 27 | 0.070487994 | 0 | 0 | 0 | 0 | 4 mo | HOM |
| **1013608106** | 2 | IGL | 27 | 0.070487994 | 0 | 0 | 0 | 0 | 4 mo | HOM |
| **1011937543** | 1 | IGL | 27 | 0.070487994 | 0.000340909 | 1 | 0.004836407 | 14.18681305 | 8 mo | HOM |
| **1010419538** | 2 | IGL | 27 | 0.070487994 | 0 | 0 | 0 | 0 | 4 mo | WILD |
| **1010419187** | 1 | IGL | 27 | 0.070487994 | 6.47E-06 | 0 | 9.17E-05 | 0 | 8 mo | HOM |
| **1013978366** | 1 | IGL | 27 | 0.070487994 | 0 | 0 | 0 | 0 | 2 mo | HOM |
| **1014416155** | 2 | IGL | 27 | 0.070487994 | 0 | 0 | 0 | 0 | 2 mo | HOM |
| **1013978065** | 2 | IGL | 27 | 0.070487994 | 0 | 0 | 0 | 0 | 2 mo | HOM |
| **1009934102** | 2 | IGL | 27 | 0.070487994 | 0 | 0 | 0 | 0 | 4 mo | WILD |
| **1009322937** | 2 | IGL | 27 | 0.070487994 | 0 | 0 | 0 | 0 | 8 mo | WILD |
| **1011080637** | 2 | IGL | 27 | 0.070487994 | 9.92E-06 | 2 | 0.000140686 | 28.37362609 | 8 mo | HOM |
| **1013609713** | 2 | IGL | 27 | 0.070487994 | 0 | 0 | 0 | 0 | 2 mo | WILD |
| **1010418842** | 1 | IGL | 27 | 0.070487994 | 8.22E-06 | 0 | 0.000116662 | 0 | 8 mo | HOM |
| **1013288213** | 2 | IGL | 27 | 0.070487994 | 0 | 0 | 0 | 0 | 4 mo | HOM |
| **1008976060** | 2 | IGL | 27 | 0.070487994 | 2.38E-07 | 0 | 3.38E-06 | 0 | 8 mo | WILD |
| **1013446243** | 1 | IGL | 27 | 0.070487994 | 0 | 0 | 0 | 0 | 4 mo | HOM |
| **1013288951** | 2 | IGL | 27 | 0.070487994 | 0 | 0 | 0 | 0 | 4 mo | HOM |
| **1013978665** | 1 | ENTl6a | 28 | 1.080372227 | 0 | 0 | 0 | 0 | 2 mo | HOM |
| **1013445609** | 1 | ENTl6a | 28 | 1.080372227 | 3.18E-05 | 1 | 2.94E-05 | 0.925606911 | 4 mo | HOM |
| **1013609015** | 1 | ENTl6a | 28 | 1.080372227 | 0.000100652 | 5 | 9.32E-05 | 4.628034555 | 4 mo | HOM |
| **1013608106** | 2 | ENTl6a | 28 | 1.080372227 | 0 | 0 | 0 | 0 | 4 mo | HOM |
| **1011937543** | 1 | ENTl6a | 28 | 1.080372227 | 0.001893444 | 31 | 0.001752585 | 28.69381424 | 8 mo | HOM |
| **1010419538** | 2 | ENTl6a | 28 | 1.080372227 | 0 | 0 | 0 | 0 | 4 mo | WILD |
| **1010419187** | 1 | ENTl6a | 28 | 1.080372227 | 0.001183308 | 45 | 0.001095278 | 41.652311 | 8 mo | HOM |
| **1013978366** | 1 | ENTl6a | 28 | 1.080372227 | 0 | 0 | 0 | 0 | 2 mo | HOM |
| **1014416155** | 2 | ENTl6a | 28 | 1.080372227 | 0 | 0 | 0 | 0 | 2 mo | HOM |
| **1013978065** | 2 | ENTl6a | 28 | 1.080372227 | 0 | 0 | 0 | 0 | 2 mo | HOM |

| | | | | | | | | | | |
|---|---|---|---|---|---|---|---|---|---|---|
| **1009934102** | 2 | ENTl6a | 28 | 1.080372227 | 0 | 0 | 0 | 0 | 4 mo | WILD |
| **1009322937** | 2 | ENTl6a | 28 | 1.080372227 | 0 | 0 | 0 | 0 | 8 mo | WILD |
| **1011080637** | 2 | ENTl6a | 28 | 1.080372227 | 0.000978662 | 34 | 0.000905857 | 31.47063498 | 8 mo | HOM |
| **1013609713** | 2 | ENTl6a | 28 | 1.080372227 | 0 | 0 | 0 | 0 | 2 mo | WILD |
| **1010418842** | 1 | ENTl6a | 28 | 1.080372227 | 0.001518058 | 49 | 0.001405125 | 45.35473864 | 8 mo | HOM |
| **1013288213** | 2 | ENTl6a | 28 | 1.080372227 | 1.07E-05 | 1 | 9.93E-06 | 0.925606911 | 4 mo | HOM |
| **1008976060** | 2 | ENTl6a | 28 | 1.080372227 | 8.82E-07 | 0 | 8.17E-07 | 0 | 8 mo | WILD |
| **1013446243** | 1 | ENTl6a | 28 | 1.080372227 | 0.000117645 | 7 | 0.000108893 | 6.479248377 | 4 mo | HOM |
| **1013288951** | 2 | ENTl6a | 28 | 1.080372227 | 2.33E-05 | 2 | 2.16E-05 | 1.851213822 | 4 mo | HOM |
| **1013978665** | 1 | PVa | 30 | 0.052316888 | 0 | 0 | 0 | 0 | 2 mo | HOM |
| **1013445609** | 1 | PVa | 30 | 0.052316888 | 0 | 0 | 0 | 0 | 4 mo | HOM |
| **1013609015** | 1 | PVa | 30 | 0.052316888 | 0 | 0 | 0 | 0 | 4 mo | HOM |
| **1013608106** | 2 | PVa | 30 | 0.052316888 | 0 | 0 | 0 | 0 | 4 mo | HOM |
| **1011937543** | 1 | PVa | 30 | 0.052316888 | 0 | 0 | 0 | 0 | 8 mo | HOM |
| **1010419538** | 2 | PVa | 30 | 0.052316888 | 0 | 0 | 0 | 0 | 4 mo | WILD |
| **1010419187** | 1 | PVa | 30 | 0.052316888 | 0 | 0 | 0 | 0 | 8 mo | HOM |
| **1013978366** | 1 | PVa | 30 | 0.052316888 | 0 | 0 | 0 | 0 | 2 mo | HOM |
| **1014416155** | 2 | PVa | 30 | 0.052316888 | 0 | 0 | 0 | 0 | 2 mo | HOM |
| **1013978065** | 2 | PVa | 30 | 0.052316888 | 0 | 0 | 0 | 0 | 2 mo | HOM |
| **1009934102** | 2 | PVa | 30 | 0.052316888 | 0 | 0 | 0 | 0 | 4 mo | WILD |
| **1009322937** | 2 | PVa | 30 | 0.052316888 | 0 | 0 | 0 | 0 | 8 mo | WILD |
| **1011080637** | 2 | PVa | 30 | 0.052316888 | 0 | 0 | 0 | 0 | 8 mo | HOM |
| **1013609713** | 2 | PVa | 30 | 0.052316888 | 0 | 0 | 0 | 0 | 2 mo | WILD |
| **1010418842** | 1 | PVa | 30 | 0.052316888 | 8.7E-05 | 2 | 0.001663756 | 38.22857324 | 8 mo | HOM |
| **1013288213** | 2 | PVa | 30 | 0.052316888 | 0 | 0 | 0 | 0 | 4 mo | HOM |
| **1008976060** | 2 | PVa | 30 | 0.052316888 | 1.03E-05 | 2 | 0.000196646 | 38.22857324 | 8 mo | WILD |
| **1013446243** | 1 | PVa | 30 | 0.052316888 | 0 | 0 | 0 | 0 | 4 mo | HOM |
| **1013288951** | 2 | PVa | 30 | 0.052316888 | 0 | 0 | 0 | 0 | 4 mo | HOM |
| **1013978665** | 1 | ACA | 31 | 5.707304376 | 4.77E-05 | 1 | 8.35E-06 | 0.175214065 | 2 mo | HOM |
| **1013445609** | 1 | ACA | 31 | 5.707304376 | 0.000354826 | 24 | 6.22E-05 | 4.205137561 | 4 mo | HOM |
| **1013609015** | 1 | ACA | 31 | 5.707304376 | 0.001701712 | 58 | 0.000298164 | 10.16241577 | 4 mo | HOM |
| **1013608106** | 2 | ACA | 31 | 5.707304376 | 0 | 0 | 0 | 0 | 4 mo | HOM |
| **1011937543** | 1 | ACA | 31 | 5.707304376 | 0.005240953 | 91 | 0.000918289 | 15.94447992 | 8 mo | HOM |
| **1010419538** | 2 | ACA | 31 | 5.707304376 | 4.21E-06 | 0 | 7.38E-07 | 0 | 4 mo | WILD |
| **1010419187** | 1 | ACA | 31 | 5.707304376 | 0.008738468 | 252 | 0.001531102 | 44.15394439 | 8 mo | HOM |
| **1013978366** | 1 | ACA | 31 | 5.707304376 | 0.000485136 | 7 | 8.5E-05 | 1.226498455 | 2 mo | HOM |
| **1014416155** | 2 | ACA | 31 | 5.707304376 | 0 | 0 | 0 | 0 | 2 mo | HOM |
| **1013978065** | 2 | ACA | 31 | 5.707304376 | 0 | 0 | 0 | 0 | 2 mo | HOM |
| **1009934102** | 2 | ACA | 31 | 5.707304376 | 0.000165184 | 5 | 2.89E-05 | 0.876070325 | 4 mo | WILD |
| **1009322937** | 2 | ACA | 31 | 5.707304376 | 3.66E-05 | 3 | 6.42E-06 | 0.525642195 | 8 mo | WILD |
| **1011080637** | 2 | ACA | 31 | 5.707304376 | 0.004292109 | 204 | 0.000752038 | 35.74366927 | 8 mo | HOM |
| **1013609713** | 2 | ACA | 31 | 5.707304376 | 4.12E-05 | 1 | 7.22E-06 | 0.175214065 | 2 mo | WILD |
| **1010418842** | 1 | ACA | 31 | 5.707304376 | 0.006463023 | 220 | 0.001132413 | 38.54709431 | 8 mo | HOM |
| **1013288213** | 2 | ACA | 31 | 5.707304376 | 0.00080106 | 22 | 0.000140357 | 3.854709431 | 4 mo | HOM |
| **1008976060** | 2 | ACA | 31 | 5.707304376 | 6.04E-05 | 4 | 1.06E-05 | 0.70085626 | 8 mo | WILD |
| **1013446243** | 1 | ACA | 31 | 5.707304376 | 0.001050983 | 32 | 0.000184147 | 5.606850081 | 4 mo | HOM |
| **1013288951** | 2 | ACA | 31 | 5.707304376 | 0.001625295 | 54 | 0.000284775 | 9.461559512 | 4 mo | HOM |
| **1013978665** | 1 | VISp6a | 33 | 1.103230845 | 0 | 0 | 0 | 0 | 2 mo | HOM |
| **1013445609** | 1 | VISp6a | 33 | 1.103230845 | 5.74E-05 | 4 | 5.2E-05 | 3.625714436 | 4 mo | HOM |
| **1013609015** | 1 | VISp6a | 33 | 1.103230845 | 0.000609425 | 19 | 0.0005524 | 17.22214357 | 4 mo | HOM |
| **1013608106** | 2 | VISp6a | 33 | 1.103230845 | 0 | 0 | 0 | 0 | 4 mo | HOM |
| **1011937543** | 1 | VISp6a | 33 | 1.103230845 | 0.001516089 | 27 | 0.001374227 | 24.47357244 | 8 mo | HOM |
| **1010419538** | 2 | VISp6a | 33 | 1.103230845 | 0 | 0 | 0 | 0 | 4 mo | WILD |
| **1010419187** | 1 | VISp6a | 33 | 1.103230845 | 0.002860038 | 75 | 0.00259242 | 67.98214567 | 8 mo | HOM |
| **1013978366** | 1 | VISp6a | 33 | 1.103230845 | 0 | 0 | 0 | 0 | 2 mo | HOM |
| **1014416155** | 2 | VISp6a | 33 | 1.103230845 | 0 | 0 | 0 | 0 | 2 mo | HOM |
| **1013978065** | 2 | VISp6a | 33 | 1.103230845 | 0 | 0 | 0 | 0 | 2 mo | HOM |
| **1009934102** | 2 | VISp6a | 33 | 1.103230845 | 0 | 0 | 0 | 0 | 4 mo | WILD |
| **1009322937** | 2 | VISp6a | 33 | 1.103230845 | 0 | 0 | 0 | 0 | 8 mo | WILD |

| | | | | | | | | | | |
|---|---|---|---|---|---|---|---|---|---|---|
| 1011080637 | 2 | VISp6a | 33 | 1.103230845 | 0.001164409 | 43 | 0.001055453 | 38.97643018 | 8 mo | HOM |
| 1013609713 | 2 | VISp6a | 33 | 1.103230845 | 0 | 0 | 0 | 0 | 2 mo | WILD |
| 1010418842 | 1 | VISp6a | 33 | 1.103230845 | 0.001532752 | 50 | 0.00138933 | 45.32143045 | 8 mo | HOM |
| 1013288213 | 2 | VISp6a | 33 | 1.103230845 | 0.000136599 | 10 | 0.000123817 | 9.064286089 | 4 mo | HOM |
| 1008976060 | 2 | VISp6a | 33 | 1.103230845 | 6E-06 | 1 | 5.43E-06 | 0.906428609 | 8 mo | WILD |
| 1013446243 | 1 | VISp6a | 33 | 1.103230845 | 0.000302353 | 12 | 0.000274061 | 10.87714331 | 4 mo | HOM |
| 1013288951 | 2 | VISp6a | 33 | 1.103230845 | 0.000316082 | 14 | 0.000286506 | 12.69000053 | 4 mo | HOM |
| 1013978665 | 1 | III | 35 | 0.03351528 | 0 | 0 | 0 | 0 | 2 mo | HOM |
| 1013445609 | 1 | III | 35 | 0.03351528 | 0 | 0 | 0 | 0 | 4 mo | HOM |
| 1013609015 | 1 | III | 35 | 0.03351528 | 0 | 0 | 0 | 0 | 4 mo | HOM |
| 1013608106 | 2 | III | 35 | 0.03351528 | 0 | 0 | 0 | 0 | 4 mo | HOM |
| 1011937543 | 1 | III | 35 | 0.03351528 | 1.98E-07 | 0 | 5.9E-06 | 0 | 8 mo | HOM |
| 1010419538 | 2 | III | 35 | 0.03351528 | 0 | 0 | 0 | 0 | 4 mo | WILD |
| 1010419187 | 1 | III | 35 | 0.03351528 | 0 | 0 | 0 | 0 | 8 mo | HOM |
| 1013978366 | 1 | III | 35 | 0.03351528 | 0 | 0 | 0 | 0 | 2 mo | HOM |
| 1014416155 | 2 | III | 35 | 0.03351528 | 0 | 0 | 0 | 0 | 2 mo | HOM |
| 1013978065 | 2 | III | 35 | 0.03351528 | 0 | 0 | 0 | 0 | 2 mo | HOM |
| 1009934102 | 2 | III | 35 | 0.03351528 | 0 | 0 | 0 | 0 | 4 mo | WILD |
| 1009322937 | 2 | III | 35 | 0.03351528 | 0 | 0 | 0 | 0 | 8 mo | WILD |
| 1011080637 | 2 | III | 35 | 0.03351528 | 0 | 0 | 0 | 0 | 8 mo | HOM |
| 1013609713 | 2 | III | 35 | 0.03351528 | 0 | 0 | 0 | 0 | 2 mo | WILD |
| 1010418842 | 1 | III | 35 | 0.03351528 | 0 | 0 | 0 | 0 | 8 mo | HOM |
| 1013288213 | 2 | III | 35 | 0.03351528 | 0 | 0 | 0 | 0 | 4 mo | HOM |
| 1008976060 | 2 | III | 35 | 0.03351528 | 0 | 0 | 0 | 0 | 8 mo | WILD |
| 1013446243 | 1 | III | 35 | 0.03351528 | 0 | 0 | 0 | 0 | 4 mo | HOM |
| 1013288951 | 2 | III | 35 | 0.03351528 | 0 | 0 | 0 | 0 | 4 mo | HOM |
| 1013978665 | 1 | GU1 | 36 | 0.202177528 | 4.23E-06 | 1 | 2.09E-05 | 4.94614812 | 2 mo | HOM |
| 1013445609 | 1 | GU1 | 36 | 0.202177528 | 2.09E-05 | 1 | 0.000103611 | 4.94614812 | 4 mo | HOM |
| 1013609015 | 1 | GU1 | 36 | 0.202177528 | 7.14E-05 | 3 | 0.0003532 | 14.83844436 | 4 mo | HOM |
| 1013608106 | 2 | GU1 | 36 | 0.202177528 | 0 | 0 | 0 | 0 | 4 mo | HOM |
| 1011937543 | 1 | GU1 | 36 | 0.202177528 | 0.000109817 | 3 | 0.00054317 | 14.83844436 | 8 mo | HOM |
| 1010419538 | 2 | GU1 | 36 | 0.202177528 | 2.47E-06 | 0 | 1.22E-05 | 0 | 4 mo | WILD |
| 1010419187 | 1 | GU1 | 36 | 0.202177528 | 0.00030584 | 16 | 0.00151273 | 79.13836992 | 8 mo | HOM |
| 1013978366 | 1 | GU1 | 36 | 0.202177528 | 0 | 0 | 0 | 0 | 2 mo | HOM |
| 1014416155 | 2 | GU1 | 36 | 0.202177528 | 0 | 0 | 0 | 0 | 2 mo | HOM |
| 1013978065 | 2 | GU1 | 36 | 0.202177528 | 0 | 0 | 0 | 0 | 2 mo | HOM |
| 1009934102 | 2 | GU1 | 36 | 0.202177528 | 0 | 0 | 0 | 0 | 4 mo | WILD |
| 1009322937 | 2 | GU1 | 36 | 0.202177528 | 0 | 0 | 0 | 0 | 8 mo | WILD |
| 1011080637 | 2 | GU1 | 36 | 0.202177528 | 0.000178696 | 11 | 0.000883858 | 54.40762932 | 8 mo | HOM |
| 1013609713 | 2 | GU1 | 36 | 0.202177528 | 0 | 0 | 0 | 0 | 2 mo | WILD |
| 1010418842 | 1 | GU1 | 36 | 0.202177528 | 0.000160508 | 10 | 0.000793898 | 49.4614812 | 8 mo | HOM |
| 1013288213 | 2 | GU1 | 36 | 0.202177528 | 5.4E-06 | 0 | 2.67E-05 | 0 | 4 mo | HOM |
| 1008976060 | 2 | GU1 | 36 | 0.202177528 | 1.02E-06 | 0 | 5.05E-06 | 0 | 8 mo | WILD |
| 1013446243 | 1 | GU1 | 36 | 0.202177528 | 9.38E-06 | 1 | 4.64E-05 | 4.94614812 | 4 mo | HOM |
| 1013288951 | 2 | GU1 | 36 | 0.202177528 | 8.41E-05 | 3 | 0.00041589 | 14.83844436 | 4 mo | HOM |
| 1013978665 | 1 | PVH | 38 | 0.194454971 | 0 | 0 | 0 | 0 | 2 mo | HOM |
| 1013445609 | 1 | PVH | 38 | 0.194454971 | 0 | 0 | 0 | 0 | 4 mo | HOM |
| 1013609015 | 1 | PVH | 38 | 0.194454971 | 0 | 0 | 0 | 0 | 4 mo | HOM |
| 1013608106 | 2 | PVH | 38 | 0.194454971 | 0 | 0 | 0 | 0 | 4 mo | HOM |
| 1011937543 | 1 | PVH | 38 | 0.194454971 | 8.26E-07 | 0 | 4.25E-06 | 0 | 8 mo | HOM |
| 1010419538 | 2 | PVH | 38 | 0.194454971 | 0 | 0 | 0 | 0 | 4 mo | WILD |
| 1010419187 | 1 | PVH | 38 | 0.194454971 | 1.92E-05 | 1 | 9.86E-05 | 5.142578736 | 8 mo | HOM |
| 1013978366 | 1 | PVH | 38 | 0.194454971 | 0 | 0 | 0 | 0 | 2 mo | HOM |
| 1014416155 | 2 | PVH | 38 | 0.194454971 | 0 | 0 | 0 | 0 | 2 mo | HOM |
| 1013978065 | 2 | PVH | 38 | 0.194454971 | 0 | 0 | 0 | 0 | 2 mo | HOM |
| 1009934102 | 2 | PVH | 38 | 0.194454971 | 0 | 0 | 0 | 0 | 4 mo | WILD |
| 1009322937 | 2 | PVH | 38 | 0.194454971 | 0 | 0 | 0 | 0 | 8 mo | WILD |
| 1011080637 | 2 | PVH | 38 | 0.194454971 | 0 | 0 | 0 | 0 | 8 mo | HOM |
| 1013609713 | 2 | PVH | 38 | 0.194454971 | 0 | 0 | 0 | 0 | 2 mo | WILD |

| | | | | | | | | | | |
|---|---|---|---|---|---|---|---|---|---|---|
| **1010418842** | 1 | PVH | 38 | 0.194454971 | 6.14E-05 | 5 | 0.000315751 | 25.71289368 | 8 mo | HOM |
| **1013288213** | 2 | PVH | 38 | 0.194454971 | 0 | 0 | 0 | 0 | 4 mo | HOM |
| **1008976060** | 2 | PVH | 38 | 0.194454971 | 2.22E-05 | 4 | 0.000114226 | 20.57031494 | 8 mo | WILD |
| **1013446243** | 1 | PVH | 38 | 0.194454971 | 1.23E-06 | 0 | 6.3E-06 | 0 | 4 mo | HOM |
| **1013288951** | 2 | PVH | 38 | 0.194454971 | 0 | 0 | 0 | 0 | 4 mo | HOM |
| **1013978665** | 1 | ACAd | 39 | 3.225978597 | 4.77E-05 | 1 | 1.48E-05 | 0.309983458 | 2 mo | HOM |
| **1013445609** | 1 | ACAd | 39 | 3.225978597 | 0.000225942 | 13 | 7E-05 | 4.02978495 | 4 mo | HOM |
| **1013609015** | 1 | ACAd | 39 | 3.225978597 | 0.001209706 | 38 | 0.000374989 | 11.77937139 | 4 mo | HOM |
| **1013608106** | 2 | ACAd | 39 | 3.225978597 | 0 | 0 | 0 | 0 | 4 mo | HOM |
| **1011937543** | 1 | ACAd | 39 | 3.225978597 | 0.003326369 | 53 | 0.001031119 | 16.42912326 | 8 mo | HOM |
| **1010419538** | 2 | ACAd | 39 | 3.225978597 | 4.21E-06 | 0 | 1.31E-06 | 0 | 4 mo | WILD |
| **1010419187** | 1 | ACAd | 39 | 3.225978597 | 0.005076861 | 147 | 0.001573743 | 45.56756828 | 8 mo | HOM |
| **1013978366** | 1 | ACAd | 39 | 3.225978597 | 0.000132034 | 3 | 4.09E-05 | 0.929950373 | 2 mo | HOM |
| **1014416155** | 2 | ACAd | 39 | 3.225978597 | 0 | 0 | 0 | 0 | 2 mo | HOM |
| **1013978065** | 2 | ACAd | 39 | 3.225978597 | 0 | 0 | 0 | 0 | 2 mo | HOM |
| **1009934102** | 2 | ACAd | 39 | 3.225978597 | 7.15E-05 | 2 | 2.22E-05 | 0.619966915 | 4 mo | WILD |
| **1009322937** | 2 | ACAd | 39 | 3.225978597 | 1.9E-05 | 2 | 5.88E-06 | 0.619966915 | 8 mo | WILD |
| **1011080637** | 2 | ACAd | 39 | 3.225978597 | 0.001949468 | 110 | 0.000604303 | 34.09818035 | 8 mo | HOM |
| **1013609713** | 2 | ACAd | 39 | 3.225978597 | 3.98E-05 | 1 | 1.23E-05 | 0.309983458 | 2 mo | WILD |
| **1010418842** | 1 | ACAd | 39 | 3.225978597 | 0.002884577 | 131 | 0.000894171 | 40.60783296 | 8 mo | HOM |
| **1013288213** | 2 | ACAd | 39 | 3.225978597 | 0.000261638 | 9 | 8.11E-05 | 2.789851119 | 4 mo | HOM |
| **1008976060** | 2 | ACAd | 39 | 3.225978597 | 4.5E-05 | 2 | 1.4E-05 | 0.619966915 | 8 mo | WILD |
| **1013446243** | 1 | ACAd | 39 | 3.225978597 | 0.000674719 | 23 | 0.000209152 | 7.129619527 | 4 mo | HOM |
| **1013288951** | 2 | ACAd | 39 | 3.225978597 | 0.001146084 | 34 | 0.000355267 | 10.53943756 | 4 mo | HOM |
| **1013978665** | 1 | VISpm2/3 | 41 | 0.301037929 | 0 | 0 | 0 | 0 | 2 mo | HOM |
| **1013445609** | 1 | VISpm2/3 | 41 | 0.301037929 | 1.51E-06 | 0 | 5.01E-06 | 0 | 4 mo | HOM |
| **1013609015** | 1 | VISpm2/3 | 41 | 0.301037929 | 7.15E-05 | 3 | 0.00023757 | 9.96552166 | 4 mo | HOM |
| **1013608106** | 2 | VISpm2/3 | 41 | 0.301037929 | 0 | 0 | 0 | 0 | 4 mo | HOM |
| **1011937543** | 1 | VISpm2/3 | 41 | 0.301037929 | 0.000693572 | 5 | 0.002303934 | 16.60920277 | 8 mo | HOM |
| **1010419538** | 2 | VISpm2/3 | 41 | 0.301037929 | 0 | 0 | 0 | 0 | 4 mo | WILD |
| **1010419187** | 1 | VISpm2/3 | 41 | 0.301037929 | 0.000400046 | 13 | 0.001328888 | 43.18392719 | 8 mo | HOM |
| **1013978366** | 1 | VISpm2/3 | 41 | 0.301037929 | 0 | 0 | 0 | 0 | 2 mo | HOM |
| **1014416155** | 2 | VISpm2/3 | 41 | 0.301037929 | 0 | 0 | 0 | 0 | 2 mo | HOM |
| **1013978065** | 2 | VISpm2/3 | 41 | 0.301037929 | 0 | 0 | 0 | 0 | 2 mo | HOM |
| **1009934102** | 2 | VISpm2/3 | 41 | 0.301037929 | 0 | 0 | 0 | 0 | 4 mo | WILD |
| **1009322937** | 2 | VISpm2/3 | 41 | 0.301037929 | 0 | 0 | 0 | 0 | 8 mo | WILD |
| **1011080637** | 2 | VISpm2/3 | 41 | 0.301037929 | 0.000140996 | 6 | 0.000468365 | 19.93104332 | 8 mo | HOM |
| **1013609713** | 2 | VISpm2/3 | 41 | 0.301037929 | 0 | 0 | 0 | 0 | 2 mo | WILD |
| **1010418842** | 1 | VISpm2/3 | 41 | 0.301037929 | 0.000499591 | 11 | 0.001659562 | 36.54024609 | 8 mo | HOM |
| **1013288213** | 2 | VISpm2/3 | 41 | 0.301037929 | 7.9E-06 | 1 | 2.62E-05 | 3.321840553 | 4 mo | HOM |
| **1008976060** | 2 | VISpm2/3 | 41 | 0.301037929 | 4.05E-06 | 0 | 1.35E-05 | 0 | 8 mo | WILD |
| **1013446243** | 1 | VISpm2/3 | 41 | 0.301037929 | 0.000155559 | 3 | 0.000516742 | 9.96552166 | 4 mo | HOM |
| **1013288951** | 2 | VISpm2/3 | 41 | 0.301037929 | 8.2E-07 | 0 | 2.72E-06 | 0 | 4 mo | HOM |
| **1013978665** | 1 | SCdw | 42 | 0.34027509 | 0 | 0 | 0 | 0 | 2 mo | HOM |
| **1013445609** | 1 | SCdw | 42 | 0.34027509 | 0 | 0 | 0 | 0 | 4 mo | HOM |
| **1013609015** | 1 | SCdw | 42 | 0.34027509 | 0 | 0 | 0 | 0 | 4 mo | HOM |
| **1013608106** | 2 | SCdw | 42 | 0.34027509 | 0 | 0 | 0 | 0 | 4 mo | HOM |
| **1011937543** | 1 | SCdw | 42 | 0.34027509 | 0.000337282 | 9 | 0.000991203 | 26.44918848 | 8 mo | HOM |
| **1010419538** | 2 | SCdw | 42 | 0.34027509 | 0 | 0 | 0 | 0 | 4 mo | WILD |
| **1010419187** | 1 | SCdw | 42 | 0.34027509 | 5.97E-06 | 1 | 1.76E-05 | 2.93879872 | 8 mo | HOM |
| **1013978366** | 1 | SCdw | 42 | 0.34027509 | 0 | 0 | 0 | 0 | 2 mo | HOM |
| **1014416155** | 2 | SCdw | 42 | 0.34027509 | 0 | 0 | 0 | 0 | 2 mo | HOM |
| **1013978065** | 2 | SCdw | 42 | 0.34027509 | 0 | 0 | 0 | 0 | 2 mo | HOM |
| **1009934102** | 2 | SCdw | 42 | 0.34027509 | 0 | 0 | 0 | 0 | 4 mo | WILD |
| **1009322937** | 2 | SCdw | 42 | 0.34027509 | 0 | 0 | 0 | 0 | 8 mo | WILD |
| **1011080637** | 2 | SCdw | 42 | 0.34027509 | 0 | 0 | 0 | 0 | 8 mo | HOM |
| **1013609713** | 2 | SCdw | 42 | 0.34027509 | 0 | 0 | 0 | 0 | 2 mo | WILD |
| **1010418842** | 1 | SCdw | 42 | 0.34027509 | 3.97E-05 | 2 | 0.00011666 | 5.877597439 | 8 mo | HOM |
| **1013288213** | 2 | SCdw | 42 | 0.34027509 | 0 | 0 | 0 | 0 | 4 mo | HOM |

| | | | | | | | | | | |
|---|---|---|---|---|---|---|---|---|---|---|
| 1008976060 | 2 | SCdw | 42 | 0.34027509 | 3.19E-08 | 0 | 9.38E-08 | 0 | 8 mo | WILD |
| 1013446243 | 1 | SCdw | 42 | 0.34027509 | 0 | 0 | 0 | 0 | 4 mo | HOM |
| 1013288951 | 2 | SCdw | 42 | 0.34027509 | 0 | 0 | 0 | 0 | 4 mo | HOM |
| 1013978665 | 1 | ILA | 44 | 0.869841625 | 0 | 0 | 0 | 0 | 2 mo | HOM |
| 1013445609 | 1 | ILA | 44 | 0.869841625 | 4.32E-05 | 1 | 4.97E-05 | 1.149634567 | 4 mo | HOM |
| 1013609015 | 1 | ILA | 44 | 0.869841625 | 0.000496751 | 11 | 0.000571082 | 12.64598024 | 4 mo | HOM |
| 1013608106 | 2 | ILA | 44 | 0.869841625 | 0 | 0 | 0 | 0 | 4 mo | HOM |
| 1011937543 | 1 | ILA | 44 | 0.869841625 | 0.000716137 | 14 | 0.000823296 | 16.09488394 | 8 mo | HOM |
| 1010419538 | 2 | ILA | 44 | 0.869841625 | 0 | 0 | 0 | 0 | 4 mo | WILD |
| 1010419187 | 1 | ILA | 44 | 0.869841625 | 0.00146797 | 29 | 0.001687629 | 33.33940244 | 8 mo | HOM |
| 1013978366 | 1 | ILA | 44 | 0.869841625 | 0 | 0 | 0 | 0 | 2 mo | HOM |
| 1014416155 | 2 | ILA | 44 | 0.869841625 | 0 | 0 | 0 | 0 | 2 mo | HOM |
| 1013978065 | 2 | ILA | 44 | 0.869841625 | 0 | 0 | 0 | 0 | 2 mo | HOM |
| 1009934102 | 2 | ILA | 44 | 0.869841625 | 0 | 0 | 0 | 0 | 4 mo | WILD |
| 1009322937 | 2 | ILA | 44 | 0.869841625 | 0.000223563 | 4 | 0.000257016 | 4.598538268 | 8 mo | WILD |
| 1011080637 | 2 | ILA | 44 | 0.869841625 | 0.000938329 | 32 | 0.001078735 | 36.78830614 | 8 mo | HOM |
| 1013609713 | 2 | ILA | 44 | 0.869841625 | 0 | 0 | 0 | 0 | 2 mo | WILD |
| 1010418842 | 1 | ILA | 44 | 0.869841625 | 0.002430496 | 21 | 0.002794182 | 24.14232591 | 8 mo | HOM |
| 1013288213 | 2 | ILA | 44 | 0.869841625 | 0.001373851 | 5 | 0.001579427 | 5.748172835 | 4 mo | HOM |
| 1008976060 | 2 | ILA | 44 | 0.869841625 | 7.9E-05 | 2 | 9.08E-05 | 2.299269134 | 8 mo | WILD |
| 1013446243 | 1 | ILA | 44 | 0.869841625 | 0.000111304 | 4 | 0.000127959 | 4.598538268 | 4 mo | HOM |
| 1013288951 | 2 | ILA | 44 | 0.869841625 | 0.000464682 | 14 | 0.000534214 | 16.09488394 | 4 mo | HOM |
| 1013978665 | 1 | mfbsma | 46 | 0.268268522 | 0 | 0 | 0 | 0 | 2 mo | HOM |
| 1013445609 | 1 | mfbsma | 46 | 0.268268522 | 0 | 0 | 0 | 0 | 4 mo | HOM |
| 1013609015 | 1 | mfbsma | 46 | 0.268268522 | 0 | 0 | 0 | 0 | 4 mo | HOM |
| 1013608106 | 2 | mfbsma | 46 | 0.268268522 | 0 | 0 | 0 | 0 | 4 mo | HOM |
| 1011937543 | 1 | mfbsma | 46 | 0.268268522 | 2.86E-06 | 0 | 1.07E-05 | 0 | 8 mo | HOM |
| 1010419538 | 2 | mfbsma | 46 | 0.268268522 | 0 | 0 | 0 | 0 | 4 mo | WILD |
| 1010419187 | 1 | mfbsma | 46 | 0.268268522 | 1.29E-10 | 0 | 4.8E-10 | 0 | 8 mo | HOM |
| 1013978366 | 1 | mfbsma | 46 | 0.268268522 | 0 | 0 | 0 | 0 | 2 mo | HOM |
| 1014416155 | 2 | mfbsma | 46 | 0.268268522 | 0 | 0 | 0 | 0 | 2 mo | HOM |
| 1013978065 | 2 | mfbsma | 46 | 0.268268522 | 0 | 0 | 0 | 0 | 2 mo | HOM |
| 1009934102 | 2 | mfbsma | 46 | 0.268268522 | 0 | 0 | 0 | 0 | 4 mo | WILD |
| 1009322937 | 2 | mfbsma | 46 | 0.268268522 | 0 | 0 | 0 | 0 | 8 mo | WILD |
| 1011080637 | 2 | mfbsma | 46 | 0.268268522 | 1.32E-07 | 0 | 4.93E-07 | 0 | 8 mo | HOM |
| 1013609713 | 2 | mfbsma | 46 | 0.268268522 | 0 | 0 | 0 | 0 | 2 mo | WILD |
| 1010418842 | 1 | mfbsma | 46 | 0.268268522 | 1.22E-05 | 0 | 4.56E-05 | 0 | 8 mo | HOM |
| 1013288213 | 2 | mfbsma | 46 | 0.268268522 | 0 | 0 | 0 | 0 | 4 mo | HOM |
| 1008976060 | 2 | mfbsma | 46 | 0.268268522 | 0 | 0 | 0 | 0 | 8 mo | WILD |
| 1013446243 | 1 | mfbsma | 46 | 0.268268522 | 0 | 0 | 0 | 0 | 4 mo | HOM |
| 1013288951 | 2 | mfbsma | 46 | 0.268268522 | 0 | 0 | 0 | 0 | 4 mo | HOM |
| 1013978665 | 1 | ACAv | 48 | 2.481325778 | 0 | 0 | 0 | 0 | 2 mo | HOM |
| 1013445609 | 1 | ACAv | 48 | 2.481325778 | 0.000128885 | 11 | 5.19E-05 | 4.433113981 | 4 mo | HOM |
| 1013609015 | 1 | ACAv | 48 | 2.481325778 | 0.000492007 | 20 | 0.000198284 | 8.060207238 | 4 mo | HOM |
| 1013608106 | 2 | ACAv | 48 | 2.481325778 | 0 | 0 | 0 | 0 | 4 mo | HOM |
| 1011937543 | 1 | ACAv | 48 | 2.481325778 | 0.001914584 | 38 | 0.000771597 | 15.31439375 | 8 mo | HOM |
| 1010419538 | 2 | ACAv | 48 | 2.481325778 | 0 | 0 | 0 | 0 | 4 mo | WILD |
| 1010419187 | 1 | ACAv | 48 | 2.481325778 | 0.003661607 | 105 | 0.001475666 | 42.316088 | 8 mo | HOM |
| 1013978366 | 1 | ACAv | 48 | 2.481325778 | 0.000353102 | 4 | 0.000142304 | 1.612041448 | 2 mo | HOM |
| 1014416155 | 2 | ACAv | 48 | 2.481325778 | 0 | 0 | 0 | 0 | 2 mo | HOM |
| 1013978065 | 2 | ACAv | 48 | 2.481325778 | 0 | 0 | 0 | 0 | 2 mo | HOM |
| 1009934102 | 2 | ACAv | 48 | 2.481325778 | 9.37E-05 | 3 | 3.78E-05 | 1.209031086 | 4 mo | WILD |
| 1009322937 | 2 | ACAv | 48 | 2.481325778 | 1.77E-05 | 1 | 7.12E-06 | 0.403010362 | 8 mo | WILD |
| 1011080637 | 2 | ACAv | 48 | 2.481325778 | 0.002342641 | 94 | 0.000944109 | 37.88297402 | 8 mo | HOM |
| 1013609713 | 2 | ACAv | 48 | 2.481325778 | 1.44E-06 | 0 | 5.79E-07 | 0 | 2 mo | WILD |
| 1010418842 | 1 | ACAv | 48 | 2.481325778 | 0.003578446 | 89 | 0.001442151 | 35.86792221 | 8 mo | HOM |
| 1013288213 | 2 | ACAv | 48 | 2.481325778 | 0.000539422 | 13 | 0.000217393 | 5.239134705 | 4 mo | HOM |
| 1008976060 | 2 | ACAv | 48 | 2.481325778 | 1.53E-05 | 2 | 6.17E-06 | 0.806020724 | 8 mo | WILD |
| 1013446243 | 1 | ACAv | 48 | 2.481325778 | 0.000376265 | 9 | 0.000151638 | 3.627093257 | 4 mo | HOM |

| | | | | | | | | | | |
|---|---|---|---|---|---|---|---|---|---|---|
| **1013288951** | 2 | ACAv | 48 | 2.481325778 | 0.000479212 | 20 | 0.000193127 | 8.060207238 | 4 mo | HOM |
| **1013978665** | 1 | PRC | 50 | 0.175865586 | 0 | 0 | 0 | 0 | 2 mo | HOM |
| **1013445609** | 1 | PRC | 50 | 0.175865586 | 0 | 0 | 0 | 0 | 4 mo | HOM |
| **1013609015** | 1 | PRC | 50 | 0.175865586 | 4.74E-05 | 3 | 0.000269367 | 17.05848234 | 4 mo | HOM |
| **1013608106** | 2 | PRC | 50 | 0.175865586 | 0 | 0 | 0 | 0 | 4 mo | HOM |
| **1011937543** | 1 | PRC | 50 | 0.175865586 | 0.000218752 | 3 | 0.001243859 | 17.05848234 | 8 mo | HOM |
| **1010419538** | 2 | PRC | 50 | 0.175865586 | 0 | 0 | 0 | 0 | 4 mo | WILD |
| **1010419187** | 1 | PRC | 50 | 0.175865586 | 1.21E-05 | 1 | 6.88E-05 | 5.68616078 | 8 mo | HOM |
| **1013978366** | 1 | PRC | 50 | 0.175865586 | 0 | 0 | 0 | 0 | 2 mo | HOM |
| **1014416155** | 2 | PRC | 50 | 0.175865586 | 0 | 0 | 0 | 0 | 2 mo | HOM |
| **1013978065** | 2 | PRC | 50 | 0.175865586 | 0 | 0 | 0 | 0 | 2 mo | HOM |
| **1009934102** | 2 | PRC | 50 | 0.175865586 | 0 | 0 | 0 | 0 | 4 mo | WILD |
| **1009322937** | 2 | PRC | 50 | 0.175865586 | 0 | 0 | 0 | 0 | 8 mo | WILD |
| **1011080637** | 2 | PRC | 50 | 0.175865586 | 4.77E-06 | 0 | 2.71E-05 | 0 | 8 mo | HOM |
| **1013609713** | 2 | PRC | 50 | 0.175865586 | 0 | 0 | 0 | 0 | 2 mo | WILD |
| **1010418842** | 1 | PRC | 50 | 0.175865586 | 5.78E-05 | 2 | 0.000328909 | 11.37232156 | 8 mo | HOM |
| **1013288213** | 2 | PRC | 50 | 0.175865586 | 0 | 0 | 0 | 0 | 4 mo | HOM |
| **1008976060** | 2 | PRC | 50 | 0.175865586 | 2.4E-05 | 3 | 0.000136672 | 17.05848234 | 8 mo | WILD |
| **1013446243** | 1 | PRC | 50 | 0.175865586 | 6.89E-06 | 1 | 3.92E-05 | 5.68616078 | 4 mo | HOM |
| **1013288951** | 2 | PRC | 50 | 0.175865586 | 3.77E-09 | 0 | 2.14E-08 | 0 | 4 mo | HOM |
| **1013978665** | 1 | ILM | 51 | 1.649338716 | 0 | 0 | 0 | 0 | 2 mo | HOM |
| **1013445609** | 1 | ILM | 51 | 1.649338716 | 0 | 0 | 0 | 0 | 4 mo | HOM |
| **1013609015** | 1 | ILM | 51 | 1.649338716 | 4.31E-06 | 1 | 2.61E-06 | 0.606303599 | 4 mo | HOM |
| **1013608106** | 2 | ILM | 51 | 1.649338716 | 0 | 0 | 0 | 0 | 4 mo | HOM |
| **1011937543** | 1 | ILM | 51 | 1.649338716 | 0.001014569 | 22 | 0.000615137 | 13.33867918 | 8 mo | HOM |
| **1010419538** | 2 | ILM | 51 | 1.649338716 | 0 | 0 | 0 | 0 | 4 mo | WILD |
| **1010419187** | 1 | ILM | 51 | 1.649338716 | 0.000211387 | 9 | 0.000128165 | 5.456732393 | 8 mo | HOM |
| **1013978366** | 1 | ILM | 51 | 1.649338716 | 0 | 0 | 0 | 0 | 2 mo | HOM |
| **1014416155** | 2 | ILM | 51 | 1.649338716 | 0 | 0 | 0 | 0 | 2 mo | HOM |
| **1013978065** | 2 | ILM | 51 | 1.649338716 | 0 | 0 | 0 | 0 | 2 mo | HOM |
| **1009934102** | 2 | ILM | 51 | 1.649338716 | 0 | 0 | 0 | 0 | 4 mo | WILD |
| **1009322937** | 2 | ILM | 51 | 1.649338716 | 7.39E-06 | 2 | 4.48E-06 | 1.212607198 | 8 mo | WILD |
| **1011080637** | 2 | ILM | 51 | 1.649338716 | 0.000307842 | 12 | 0.000186646 | 7.275643191 | 8 mo | HOM |
| **1013609713** | 2 | ILM | 51 | 1.649338716 | 0 | 0 | 0 | 0 | 2 mo | WILD |
| **1010418842** | 1 | ILM | 51 | 1.649338716 | 0.000606114 | 17 | 0.000367489 | 10.30716119 | 8 mo | HOM |
| **1013288213** | 2 | ILM | 51 | 1.649338716 | 0 | 0 | 0 | 0 | 4 mo | HOM |
| **1008976060** | 2 | ILM | 51 | 1.649338716 | 3.74E-05 | 4 | 2.27E-05 | 2.425214397 | 8 mo | WILD |
| **1013446243** | 1 | ILM | 51 | 1.649338716 | 2.86E-05 | 2 | 1.73E-05 | 1.212607198 | 4 mo | HOM |
| **1013288951** | 2 | ILM | 51 | 1.649338716 | 0 | 0 | 0 | 0 | 4 mo | HOM |
| **1013978665** | 1 | ENTl3 | 52 | 1.356888714 | 0 | 0 | 0 | 0 | 2 mo | HOM |
| **1013445609** | 1 | ENTl3 | 52 | 1.356888714 | 6.67E-05 | 7 | 4.91E-05 | 5.158860804 | 4 mo | HOM |
| **1013609015** | 1 | ENTl3 | 52 | 1.356888714 | 0.000127507 | 5 | 9.4E-05 | 3.684900574 | 4 mo | HOM |
| **1013608106** | 2 | ENTl3 | 52 | 1.356888714 | 0 | 0 | 0 | 0 | 4 mo | HOM |
| **1011937543** | 1 | ENTl3 | 52 | 1.356888714 | 0.000953629 | 27 | 0.000702805 | 19.8984631 | 8 mo | HOM |
| **1010419538** | 2 | ENTl3 | 52 | 1.356888714 | 0 | 0 | 0 | 0 | 4 mo | WILD |
| **1010419187** | 1 | ENTl3 | 52 | 1.356888714 | 0.001257129 | 50 | 0.000926479 | 36.84900574 | 8 mo | HOM |
| **1013978366** | 1 | ENTl3 | 52 | 1.356888714 | 0 | 0 | 0 | 0 | 2 mo | HOM |
| **1014416155** | 2 | ENTl3 | 52 | 1.356888714 | 0 | 0 | 0 | 0 | 2 mo | HOM |
| **1013978065** | 2 | ENTl3 | 52 | 1.356888714 | 0 | 0 | 0 | 0 | 2 mo | HOM |
| **1009934102** | 2 | ENTl3 | 52 | 1.356888714 | 0 | 0 | 0 | 0 | 4 mo | WILD |
| **1009322937** | 2 | ENTl3 | 52 | 1.356888714 | 0 | 0 | 0 | 0 | 8 mo | WILD |
| **1011080637** | 2 | ENTl3 | 52 | 1.356888714 | 0.000843653 | 39 | 0.000621756 | 28.74222448 | 8 mo | HOM |
| **1013609713** | 2 | ENTl3 | 52 | 1.356888714 | 0 | 0 | 0 | 0 | 2 mo | WILD |
| **1010418842** | 1 | ENTl3 | 52 | 1.356888714 | 0.001138681 | 47 | 0.000839185 | 34.6380654 | 8 mo | HOM |
| **1013288213** | 2 | ENTl3 | 52 | 1.356888714 | 0.000160091 | 8 | 0.000117984 | 5.895840919 | 4 mo | HOM |
| **1008976060** | 2 | ENTl3 | 52 | 1.356888714 | 1.68E-06 | 0 | 1.24E-06 | 0 | 8 mo | WILD |
| **1013446243** | 1 | ENTl3 | 52 | 1.356888714 | 0.000171173 | 6 | 0.000126151 | 4.421880689 | 4 mo | HOM |
| **1013288951** | 2 | ENTl3 | 52 | 1.356888714 | 0.0001515 | 11 | 0.000111652 | 8.106781263 | 4 mo | HOM |
| **1013978665** | 1 | mfb | 54 | 0.050835422 | 0 | 0 | 0 | 0 | 2 mo | HOM |

| | | | | | | | | | | |
|---|---|---|---|---|---|---|---|---|---|---|
| **1013445609** | 1 | mfb | 54 | 0.050835422 | 0 | 0 | 0 | 0 | 4 mo | HOM |
| **1013609015** | 1 | mfb | 54 | 0.050835422 | 0 | 0 | 0 | 0 | 4 mo | HOM |
| **1013608106** | 2 | mfb | 54 | 0.050835422 | 0 | 0 | 0 | 0 | 4 mo | HOM |
| **1011937543** | 1 | mfb | 54 | 0.050835422 | 6.89E-05 | 2 | 0.001355778 | 39.34264545 | 8 mo | HOM |
| **1010419538** | 2 | mfb | 54 | 0.050835422 | 0 | 0 | 0 | 0 | 4 mo | WILD |
| **1010419187** | 1 | mfb | 54 | 0.050835422 | 3.92E-08 | 0 | 7.71E-07 | 0 | 8 mo | HOM |
| **1013978366** | 1 | mfb | 54 | 0.050835422 | 0 | 0 | 0 | 0 | 2 mo | HOM |
| **1014416155** | 2 | mfb | 54 | 0.050835422 | 0 | 0 | 0 | 0 | 2 mo | HOM |
| **1013978065** | 2 | mfb | 54 | 0.050835422 | 0 | 0 | 0 | 0 | 2 mo | HOM |
| **1009934102** | 2 | mfb | 54 | 0.050835422 | 0 | 0 | 0 | 0 | 4 mo | WILD |
| **1009322937** | 2 | mfb | 54 | 0.050835422 | 0 | 0 | 0 | 0 | 8 mo | WILD |
| **1011080637** | 2 | mfb | 54 | 0.050835422 | 0 | 0 | 0 | 0 | 8 mo | HOM |
| **1013609713** | 2 | mfb | 54 | 0.050835422 | 0 | 0 | 0 | 0 | 2 mo | WILD |
| **1010418842** | 1 | mfb | 54 | 0.050835422 | 6.39E-09 | 0 | 1.26E-07 | 0 | 8 mo | HOM |
| **1013288213** | 2 | mfb | 54 | 0.050835422 | 0 | 0 | 0 | 0 | 4 mo | HOM |
| **1008976060** | 2 | mfb | 54 | 0.050835422 | 0 | 0 | 0 | 0 | 8 mo | WILD |
| **1013446243** | 1 | mfb | 54 | 0.050835422 | 0 | 0 | 0 | 0 | 4 mo | HOM |
| **1013288951** | 2 | mfb | 54 | 0.050835422 | 0 | 0 | 0 | 0 | 4 mo | HOM |
| **1013978665** | 1 | ACB | 56 | 4.53439728 | 0 | 0 | 0 | 0 | 2 mo | HOM |
| **1013445609** | 1 | ACB | 56 | 4.53439728 | 1.42E-05 | 2 | 3.13E-06 | 0.441072953 | 4 mo | HOM |
| **1013609015** | 1 | ACB | 56 | 4.53439728 | 7.99E-05 | 4 | 1.76E-05 | 0.882145907 | 4 mo | HOM |
| **1013608106** | 2 | ACB | 56 | 4.53439728 | 0 | 0 | 0 | 0 | 4 mo | HOM |
| **1011937543** | 1 | ACB | 56 | 4.53439728 | 0.003204542 | 78 | 0.000706718 | 17.20184518 | 8 mo | HOM |
| **1010419538** | 2 | ACB | 56 | 4.53439728 | 0 | 0 | 0 | 0 | 4 mo | WILD |
| **1010419187** | 1 | ACB | 56 | 4.53439728 | 0.002599333 | 127 | 0.000573248 | 28.00813254 | 8 mo | HOM |
| **1013978366** | 1 | ACB | 56 | 4.53439728 | 0 | 0 | 0 | 0 | 2 mo | HOM |
| **1014416155** | 2 | ACB | 56 | 4.53439728 | 0 | 0 | 0 | 0 | 2 mo | HOM |
| **1013978065** | 2 | ACB | 56 | 4.53439728 | 0 | 0 | 0 | 0 | 2 mo | HOM |
| **1009934102** | 2 | ACB | 56 | 4.53439728 | 0 | 0 | 0 | 0 | 4 mo | WILD |
| **1009322937** | 2 | ACB | 56 | 4.53439728 | 0 | 0 | 0 | 0 | 8 mo | WILD |
| **1011080637** | 2 | ACB | 56 | 4.53439728 | 0.001339156 | 77 | 0.000295333 | 16.9813087 | 8 mo | HOM |
| **1013609713** | 2 | ACB | 56 | 4.53439728 | 0 | 0 | 0 | 0 | 2 mo | WILD |
| **1010418842** | 1 | ACB | 56 | 4.53439728 | 0.002268319 | 104 | 0.000500247 | 22.93579357 | 8 mo | HOM |
| **1013288213** | 2 | ACB | 56 | 4.53439728 | 1.84E-05 | 1 | 4.06E-06 | 0.220536477 | 4 mo | HOM |
| **1008976060** | 2 | ACB | 56 | 4.53439728 | 7.08E-06 | 2 | 1.56E-06 | 0.441072953 | 8 mo | WILD |
| **1013446243** | 1 | ACB | 56 | 4.53439728 | 6.9E-05 | 8 | 1.52E-05 | 1.764291813 | 4 mo | HOM |
| **1013288951** | 2 | ACB | 56 | 4.53439728 | 1.75E-05 | 2 | 3.86E-06 | 0.441072953 | 4 mo | HOM |
| **1013978665** | 1 | MT | 58 | 0.044668498 | 0 | 0 | 0 | 0 | 2 mo | HOM |
| **1013445609** | 1 | MT | 58 | 0.044668498 | 0 | 0 | 0 | 0 | 4 mo | HOM |
| **1013609015** | 1 | MT | 58 | 0.044668498 | 4.18E-06 | 1 | 9.35E-05 | 22.38714183 | 4 mo | HOM |
| **1013608106** | 2 | MT | 58 | 0.044668498 | 0 | 0 | 0 | 0 | 4 mo | HOM |
| **1011937543** | 1 | MT | 58 | 0.044668498 | 4.72E-05 | 2 | 0.001055633 | 44.77428366 | 8 mo | HOM |
| **1010419538** | 2 | MT | 58 | 0.044668498 | 0 | 0 | 0 | 0 | 4 mo | WILD |
| **1010419187** | 1 | MT | 58 | 0.044668498 | 0 | 0 | 0 | 0 | 8 mo | HOM |
| **1013978366** | 1 | MT | 58 | 0.044668498 | 0 | 0 | 0 | 0 | 2 mo | HOM |
| **1014416155** | 2 | MT | 58 | 0.044668498 | 0 | 0 | 0 | 0 | 2 mo | HOM |
| **1013978065** | 2 | MT | 58 | 0.044668498 | 0 | 0 | 0 | 0 | 2 mo | HOM |
| **1009934102** | 2 | MT | 58 | 0.044668498 | 0 | 0 | 0 | 0 | 4 mo | WILD |
| **1009322937** | 2 | MT | 58 | 0.044668498 | 0 | 0 | 0 | 0 | 8 mo | WILD |
| **1011080637** | 2 | MT | 58 | 0.044668498 | 0 | 0 | 0 | 0 | 8 mo | HOM |
| **1013609713** | 2 | MT | 58 | 0.044668498 | 0 | 0 | 0 | 0 | 2 mo | WILD |
| **1010418842** | 1 | MT | 58 | 0.044668498 | 7.78E-07 | 0 | 1.74E-05 | 0 | 8 mo | HOM |
| **1013288213** | 2 | MT | 58 | 0.044668498 | 0 | 0 | 0 | 0 | 4 mo | HOM |
| **1008976060** | 2 | MT | 58 | 0.044668498 | 0 | 0 | 0 | 0 | 8 mo | WILD |
| **1013446243** | 1 | MT | 58 | 0.044668498 | 0 | 0 | 0 | 0 | 4 mo | HOM |
| **1013288951** | 2 | MT | 58 | 0.044668498 | 0 | 0 | 0 | 0 | 4 mo | HOM |
| **1013978665** | 1 | IMD | 59 | 0.181742 | 0 | 0 | 0 | 0 | 2 mo | HOM |
| **1013445609** | 1 | IMD | 59 | 0.181742 | 0 | 0 | 0 | 0 | 4 mo | HOM |
| **1013609015** | 1 | IMD | 59 | 0.181742 | 0 | 0 | 0 | 0 | 4 mo | HOM |

| | | | | | | | | | | |
|---|---|---|---|---|---|---|---|---|---|---|
| 1013608106 | 2 | IMD | 59 | 0.181742 | 0 | 0 | 0 | 0 | 4 mo | HOM |
| 1011937543 | 1 | IMD | 59 | 0.181742 | 0.000150992 | 2 | 0.000830805 | 11.00461096 | 8 mo | HOM |
| 1010419538 | 2 | IMD | 59 | 0.181742 | 0 | 0 | 0 | 0 | 4 mo | WILD |
| 1010419187 | 1 | IMD | 59 | 0.181742 | 6.2E-08 | 0 | 3.41E-07 | 0 | 8 mo | HOM |
| 1013978366 | 1 | IMD | 59 | 0.181742 | 0 | 0 | 0 | 0 | 2 mo | HOM |
| 1014416155 | 2 | IMD | 59 | 0.181742 | 0 | 0 | 0 | 0 | 2 mo | HOM |
| 1013978065 | 2 | IMD | 59 | 0.181742 | 0 | 0 | 0 | 0 | 2 mo | HOM |
| 1009934102 | 2 | IMD | 59 | 0.181742 | 0 | 0 | 0 | 0 | 4 mo | WILD |
| 1009322937 | 2 | IMD | 59 | 0.181742 | 0 | 0 | 0 | 0 | 8 mo | WILD |
| 1011080637 | 2 | IMD | 59 | 0.181742 | 0.00012468 | 5 | 0.000686025 | 27.51152739 | 8 mo | HOM |
| 1013609713 | 2 | IMD | 59 | 0.181742 | 0 | 0 | 0 | 0 | 2 mo | WILD |
| 1010418842 | 1 | IMD | 59 | 0.181742 | 0.000102524 | 3 | 0.000564117 | 16.50691643 | 8 mo | HOM |
| 1013288213 | 2 | IMD | 59 | 0.181742 | 0 | 0 | 0 | 0 | 4 mo | HOM |
| 1008976060 | 2 | IMD | 59 | 0.181742 | 4.43E-06 | 1 | 2.44E-05 | 5.502305478 | 8 mo | WILD |
| 1013446243 | 1 | IMD | 59 | 0.181742 | 0 | 0 | 0 | 0 | 4 mo | HOM |
| 1013288951 | 2 | IMD | 59 | 0.181742 | 0 | 0 | 0 | 0 | 4 mo | HOM |
| 1013978665 | 1 | mlf | 62 | 0.153604485 | 0 | 0 | 0 | 0 | 2 mo | HOM |
| 1013445609 | 1 | mlf | 62 | 0.153604485 | 0 | 0 | 0 | 0 | 4 mo | HOM |
| 1013609015 | 1 | mlf | 62 | 0.153604485 | 0 | 0 | 0 | 0 | 4 mo | HOM |
| 1013608106 | 2 | mlf | 62 | 0.153604485 | 0 | 0 | 0 | 0 | 4 mo | HOM |
| 1011937543 | 1 | mlf | 62 | 0.153604485 | 1.58E-06 | 0 | 1.03E-05 | 0 | 8 mo | HOM |
| 1010419538 | 2 | mlf | 62 | 0.153604485 | 0 | 0 | 0 | 0 | 4 mo | WILD |
| 1010419187 | 1 | mlf | 62 | 0.153604485 | 0 | 0 | 0 | 0 | 8 mo | HOM |
| 1013978366 | 1 | mlf | 62 | 0.153604485 | 0 | 0 | 0 | 0 | 2 mo | HOM |
| 1014416155 | 2 | mlf | 62 | 0.153604485 | 0 | 0 | 0 | 0 | 2 mo | HOM |
| 1013978065 | 2 | mlf | 62 | 0.153604485 | 0 | 0 | 0 | 0 | 2 mo | HOM |
| 1009934102 | 2 | mlf | 62 | 0.153604485 | 0 | 0 | 0 | 0 | 4 mo | WILD |
| 1009322937 | 2 | mlf | 62 | 0.153604485 | 0 | 0 | 0 | 0 | 8 mo | WILD |
| 1011080637 | 2 | mlf | 62 | 0.153604485 | 0 | 0 | 0 | 0 | 8 mo | HOM |
| 1013609713 | 2 | mlf | 62 | 0.153604485 | 0 | 0 | 0 | 0 | 2 mo | WILD |
| 1010418842 | 1 | mlf | 62 | 0.153604485 | 0 | 0 | 0 | 0 | 8 mo | HOM |
| 1013288213 | 2 | mlf | 62 | 0.153604485 | 0 | 0 | 0 | 0 | 4 mo | HOM |
| 1008976060 | 2 | mlf | 62 | 0.153604485 | 4.85E-08 | 0 | 3.16E-07 | 0 | 8 mo | WILD |
| 1013446243 | 1 | mlf | 62 | 0.153604485 | 0 | 0 | 0 | 0 | 4 mo | HOM |
| 1013288951 | 2 | mlf | 62 | 0.153604485 | 0 | 0 | 0 | 0 | 4 mo | HOM |
| 1013978665 | 1 | PVHd | 63 | 0.134590652 | 0 | 0 | 0 | 0 | 2 mo | HOM |
| 1013445609 | 1 | PVHd | 63 | 0.134590652 | 0 | 0 | 0 | 0 | 4 mo | HOM |
| 1013609015 | 1 | PVHd | 63 | 0.134590652 | 0 | 0 | 0 | 0 | 4 mo | HOM |
| 1013608106 | 2 | PVHd | 63 | 0.134590652 | 0 | 0 | 0 | 0 | 4 mo | HOM |
| 1011937543 | 1 | PVHd | 63 | 0.134590652 | 0 | 0 | 0 | 0 | 8 mo | HOM |
| 1010419538 | 2 | PVHd | 63 | 0.134590652 | 0 | 0 | 0 | 0 | 4 mo | WILD |
| 1010419187 | 1 | PVHd | 63 | 0.134590652 | 7.76E-05 | 2 | 0.000576493 | 14.85987303 | 8 mo | HOM |
| 1013978366 | 1 | PVHd | 63 | 0.134590652 | 0 | 0 | 0 | 0 | 2 mo | HOM |
| 1014416155 | 2 | PVHd | 63 | 0.134590652 | 0 | 0 | 0 | 0 | 2 mo | HOM |
| 1013978065 | 2 | PVHd | 63 | 0.134590652 | 0 | 0 | 0 | 0 | 2 mo | HOM |
| 1009934102 | 2 | PVHd | 63 | 0.134590652 | 0 | 0 | 0 | 0 | 4 mo | WILD |
| 1009322937 | 2 | PVHd | 63 | 0.134590652 | 0 | 0 | 0 | 0 | 8 mo | WILD |
| 1011080637 | 2 | PVHd | 63 | 0.134590652 | 8.61E-06 | 1 | 6.4E-05 | 7.429936516 | 8 mo | HOM |
| 1013609713 | 2 | PVHd | 63 | 0.134590652 | 0 | 0 | 0 | 0 | 2 mo | WILD |
| 1010418842 | 1 | PVHd | 63 | 0.134590652 | 8.43E-06 | 0 | 6.26E-05 | 0 | 8 mo | HOM |
| 1013288213 | 2 | PVHd | 63 | 0.134590652 | 0 | 0 | 0 | 0 | 4 mo | HOM |
| 1008976060 | 2 | PVHd | 63 | 0.134590652 | 0 | 0 | 0 | 0 | 8 mo | WILD |
| 1013446243 | 1 | PVHd | 63 | 0.134590652 | 0 | 0 | 0 | 0 | 4 mo | HOM |
| 1013288951 | 2 | PVHd | 63 | 0.134590652 | 0 | 0 | 0 | 0 | 4 mo | HOM |
| 1013978665 | 1 | AD | 64 | 0.165983678 | 0 | 0 | 0 | 0 | 2 mo | HOM |
| 1013445609 | 1 | AD | 64 | 0.165983678 | 1.61E-06 | 0 | 9.69E-06 | 0 | 4 mo | HOM |
| 1013609015 | 1 | AD | 64 | 0.165983678 | 9.44E-05 | 6 | 0.000568735 | 36.14813259 | 4 mo | HOM |
| 1013608106 | 2 | AD | 64 | 0.165983678 | 0 | 0 | 0 | 0 | 4 mo | HOM |
| 1011937543 | 1 | AD | 64 | 0.165983678 | 0.000114266 | 5 | 0.000688416 | 30.12344382 | 8 mo | HOM |

| | | | | | | | | | | |
|---|---|---|---|---|---|---|---|---|---|---|
| **1010419538** | 2 | AD | 64 | 0.165983678 | 0 | 0 | 0 | 0 | 4 mo | WILD |
| **1010419187** | 1 | AD | 64 | 0.165983678 | 0.000545696 | 7 | 0.003287646 | 42.17282135 | 8 mo | HOM |
| **1013978366** | 1 | AD | 64 | 0.165983678 | 1.08E-05 | 1 | 6.51E-05 | 6.024688765 | 2 mo | HOM |
| **1014416155** | 2 | AD | 64 | 0.165983678 | 0 | 0 | 0 | 0 | 2 mo | HOM |
| **1013978065** | 2 | AD | 64 | 0.165983678 | 0 | 0 | 0 | 0 | 2 mo | HOM |
| **1009934102** | 2 | AD | 64 | 0.165983678 | 1.07E-07 | 0 | 6.43E-07 | 0 | 4 mo | WILD |
| **1009322937** | 2 | AD | 64 | 0.165983678 | 9.05E-07 | 0 | 5.45E-06 | 0 | 8 mo | WILD |
| **1011080637** | 2 | AD | 64 | 0.165983678 | 0.00034465 | 5 | 0.00207641 | 30.12344382 | 8 mo | HOM |
| **1013609713** | 2 | AD | 64 | 0.165983678 | 0 | 0 | 0 | 0 | 2 mo | WILD |
| **1010418842** | 1 | AD | 64 | 0.165983678 | 0.000365198 | 7 | 0.002200207 | 42.17282135 | 8 mo | HOM |
| **1013288213** | 2 | AD | 64 | 0.165983678 | 4.32E-06 | 1 | 2.6E-05 | 6.024688765 | 4 mo | HOM |
| **1008976060** | 2 | AD | 64 | 0.165983678 | 2.69E-05 | 3 | 0.000162299 | 18.07406629 | 8 mo | WILD |
| **1013446243** | 1 | AD | 64 | 0.165983678 | 1.62E-05 | 2 | 9.73E-05 | 12.04937753 | 4 mo | HOM |
| **1013288951** | 2 | AD | 64 | 0.165983678 | 0 | 0 | 0 | 0 | 4 mo | HOM |
| **1013978665** | 1 | LT | 66 | 0.018424544 | 0 | 0 | 0 | 0 | 2 mo | HOM |
| **1013445609** | 1 | LT | 66 | 0.018424544 | 0 | 0 | 0 | 0 | 4 mo | HOM |
| **1013609015** | 1 | LT | 66 | 0.018424544 | 0 | 0 | 0 | 0 | 4 mo | HOM |
| **1013608106** | 2 | LT | 66 | 0.018424544 | 0 | 0 | 0 | 0 | 4 mo | HOM |
| **1011937543** | 1 | LT | 66 | 0.018424544 | 3.86E-05 | 1 | 0.002092835 | 54.27542768 | 8 mo | HOM |
| **1010419538** | 2 | LT | 66 | 0.018424544 | 0 | 0 | 0 | 0 | 4 mo | WILD |
| **1010419187** | 1 | LT | 66 | 0.018424544 | 0 | 0 | 0 | 0 | 8 mo | HOM |
| **1013978366** | 1 | LT | 66 | 0.018424544 | 0 | 0 | 0 | 0 | 2 mo | HOM |
| **1014416155** | 2 | LT | 66 | 0.018424544 | 0 | 0 | 0 | 0 | 2 mo | HOM |
| **1013978065** | 2 | LT | 66 | 0.018424544 | 0 | 0 | 0 | 0 | 2 mo | HOM |
| **1009934102** | 2 | LT | 66 | 0.018424544 | 0 | 0 | 0 | 0 | 4 mo | WILD |
| **1009322937** | 2 | LT | 66 | 0.018424544 | 0 | 0 | 0 | 0 | 8 mo | WILD |
| **1011080637** | 2 | LT | 66 | 0.018424544 | 0 | 0 | 0 | 0 | 8 mo | HOM |
| **1013609713** | 2 | LT | 66 | 0.018424544 | 0 | 0 | 0 | 0 | 2 mo | WILD |
| **1010418842** | 1 | LT | 66 | 0.018424544 | 0 | 0 | 0 | 0 | 8 mo | HOM |
| **1013288213** | 2 | LT | 66 | 0.018424544 | 0 | 0 | 0 | 0 | 4 mo | HOM |
| **1008976060** | 2 | LT | 66 | 0.018424544 | 2.75E-06 | 1 | 0.000149165 | 54.27542768 | 8 mo | WILD |
| **1013446243** | 1 | LT | 66 | 0.018424544 | 0 | 0 | 0 | 0 | 4 mo | HOM |
| **1013288951** | 2 | LT | 66 | 0.018424544 | 0 | 0 | 0 | 0 | 4 mo | HOM |
| **1013978665** | 1 | INC | 67 | 0.082137779 | 0 | 0 | 0 | 0 | 2 mo | HOM |
| **1013445609** | 1 | INC | 67 | 0.082137779 | 0 | 0 | 0 | 0 | 4 mo | HOM |
| **1013609015** | 1 | INC | 67 | 0.082137779 | 0 | 0 | 0 | 0 | 4 mo | HOM |
| **1013608106** | 2 | INC | 67 | 0.082137779 | 0 | 0 | 0 | 0 | 4 mo | HOM |
| **1011937543** | 1 | INC | 67 | 0.082137779 | 4.23E-10 | 0 | 5.15E-09 | 0 | 8 mo | HOM |
| **1010419538** | 2 | INC | 67 | 0.082137779 | 0 | 0 | 0 | 0 | 4 mo | WILD |
| **1010419187** | 1 | INC | 67 | 0.082137779 | 0 | 0 | 0 | 0 | 8 mo | HOM |
| **1013978366** | 1 | INC | 67 | 0.082137779 | 0 | 0 | 0 | 0 | 2 mo | HOM |
| **1014416155** | 2 | INC | 67 | 0.082137779 | 0 | 0 | 0 | 0 | 2 mo | HOM |
| **1013978065** | 2 | INC | 67 | 0.082137779 | 0 | 0 | 0 | 0 | 2 mo | HOM |
| **1009934102** | 2 | INC | 67 | 0.082137779 | 0 | 0 | 0 | 0 | 4 mo | WILD |
| **1009322937** | 2 | INC | 67 | 0.082137779 | 0 | 0 | 0 | 0 | 8 mo | WILD |
| **1011080637** | 2 | INC | 67 | 0.082137779 | 0 | 0 | 0 | 0 | 8 mo | HOM |
| **1013609713** | 2 | INC | 67 | 0.082137779 | 0 | 0 | 0 | 0 | 2 mo | WILD |
| **1010418842** | 1 | INC | 67 | 0.082137779 | 0 | 0 | 0 | 0 | 8 mo | HOM |
| **1013288213** | 2 | INC | 67 | 0.082137779 | 0 | 0 | 0 | 0 | 4 mo | HOM |
| **1008976060** | 2 | INC | 67 | 0.082137779 | 0 | 0 | 0 | 0 | 8 mo | WILD |
| **1013446243** | 1 | INC | 67 | 0.082137779 | 0 | 0 | 0 | 0 | 4 mo | HOM |
| **1013288951** | 2 | INC | 67 | 0.082137779 | 0 | 0 | 0 | 0 | 4 mo | HOM |
| **1013978665** | 1 | FRP1 | 68 | 0.264273307 | 0 | 0 | 0 | 0 | 2 mo | HOM |
| **1013445609** | 1 | FRP1 | 68 | 0.264273307 | 0 | 0 | 0 | 0 | 4 mo | HOM |
| **1013609015** | 1 | FRP1 | 68 | 0.264273307 | 9.14E-07 | 0 | 3.46E-06 | 0 | 4 mo | HOM |
| **1013608106** | 2 | FRP1 | 68 | 0.264273307 | 0 | 0 | 0 | 0 | 4 mo | HOM |
| **1011937543** | 1 | FRP1 | 68 | 0.264273307 | 0.000142331 | 3 | 0.000538574 | 11.35188427 | 8 mo | HOM |
| **1010419538** | 2 | FRP1 | 68 | 0.264273307 | 0 | 0 | 0 | 0 | 4 mo | WILD |
| **1010419187** | 1 | FRP1 | 68 | 0.264273307 | 5.02E-05 | 6 | 0.000190054 | 22.70376855 | 8 mo | HOM |

| | | | | | | | | | | |
|---|---|---|---|---|---|---|---|---|---|---|
| **1013978366** | 1 | FRP1 | 68 | 0.264273307 | 1.04E-05 | 0 | 3.93E-05 | 0 | 2 mo | HOM |
| **1014416155** | 2 | FRP1 | 68 | 0.264273307 | 0 | 0 | 0 | 0 | 2 mo | HOM |
| **1013978065** | 2 | FRP1 | 68 | 0.264273307 | 0 | 0 | 0 | 0 | 2 mo | HOM |
| **1009934102** | 2 | FRP1 | 68 | 0.264273307 | 0 | 0 | 0 | 0 | 4 mo | WILD |
| **1009322937** | 2 | FRP1 | 68 | 0.264273307 | 1.69E-05 | 2 | 6.38E-05 | 7.567922849 | 8 mo | WILD |
| **1011080637** | 2 | FRP1 | 68 | 0.264273307 | 7.11E-06 | 1 | 2.69E-05 | 3.783961425 | 8 mo | HOM |
| **1013609713** | 2 | FRP1 | 68 | 0.264273307 | 0 | 0 | 0 | 0 | 2 mo | WILD |
| **1010418842** | 1 | FRP1 | 68 | 0.264273307 | 5.52E-05 | 3 | 0.000208857 | 11.35188427 | 8 mo | HOM |
| **1013288213** | 2 | FRP1 | 68 | 0.264273307 | 0.000822509 | 3 | 0.003112343 | 11.35188427 | 4 mo | HOM |
| **1008976060** | 2 | FRP1 | 68 | 0.264273307 | 1.35E-05 | 1 | 5.12E-05 | 3.783961425 | 8 mo | WILD |
| **1013446243** | 1 | FRP1 | 68 | 0.264273307 | 2.17E-07 | 0 | 8.2E-07 | 0 | 4 mo | HOM |
| **1013288951** | 2 | FRP1 | 68 | 0.264273307 | 4E-05 | 3 | 0.00015147 | 11.35188427 | 4 mo | HOM |
| **1013978665** | 1 | ADP | 72 | 0.102538292 | 0 | 0 | 0 | 0 | 2 mo | HOM |
| **1013445609** | 1 | ADP | 72 | 0.102538292 | 0 | 0 | 0 | 0 | 4 mo | HOM |
| **1013609015** | 1 | ADP | 72 | 0.102538292 | 0 | 0 | 0 | 0 | 4 mo | HOM |
| **1013608106** | 2 | ADP | 72 | 0.102538292 | 0 | 0 | 0 | 0 | 4 mo | HOM |
| **1011937543** | 1 | ADP | 72 | 0.102538292 | 9.15E-05 | 4 | 0.000892087 | 39.00981701 | 8 mo | HOM |
| **1010419538** | 2 | ADP | 72 | 0.102538292 | 0 | 0 | 0 | 0 | 4 mo | WILD |
| **1010419187** | 1 | ADP | 72 | 0.102538292 | 0.000151105 | 1 | 0.001473645 | 9.752454253 | 8 mo | HOM |
| **1013978366** | 1 | ADP | 72 | 0.102538292 | 0 | 0 | 0 | 0 | 2 mo | HOM |
| **1014416155** | 2 | ADP | 72 | 0.102538292 | 0 | 0 | 0 | 0 | 2 mo | HOM |
| **1013978065** | 2 | ADP | 72 | 0.102538292 | 0 | 0 | 0 | 0 | 2 mo | HOM |
| **1009934102** | 2 | ADP | 72 | 0.102538292 | 0 | 0 | 0 | 0 | 4 mo | WILD |
| **1009322937** | 2 | ADP | 72 | 0.102538292 | 0 | 0 | 0 | 0 | 8 mo | WILD |
| **1011080637** | 2 | ADP | 72 | 0.102538292 | 0.000102055 | 2 | 0.000995283 | 19.50490851 | 8 mo | HOM |
| **1013609713** | 2 | ADP | 72 | 0.102538292 | 0 | 0 | 0 | 0 | 2 mo | WILD |
| **1010418842** | 1 | ADP | 72 | 0.102538292 | 0.000180655 | 2 | 0.001761834 | 19.50490851 | 8 mo | HOM |
| **1013288213** | 2 | ADP | 72 | 0.102538292 | 0 | 0 | 0 | 0 | 4 mo | HOM |
| **1008976060** | 2 | ADP | 72 | 0.102538292 | 0 | 0 | 0 | 0 | 8 mo | WILD |
| **1013446243** | 1 | ADP | 72 | 0.102538292 | 0 | 0 | 0 | 0 | 4 mo | HOM |
| **1013288951** | 2 | ADP | 72 | 0.102538292 | 0 | 0 | 0 | 0 | 4 mo | HOM |
| **1013978665** | 1 | VS | 73 | 6.139504255 | 0.001276114 | 30 | 0.000207853 | 4.886388014 | 2 mo | HOM |
| **1013445609** | 1 | VS | 73 | 6.139504255 | 0.002186239 | 80 | 0.000356094 | 13.03036804 | 4 mo | HOM |
| **1013609015** | 1 | VS | 73 | 6.139504255 | 0.029727474 | 119 | 0.004841999 | 19.38267245 | 4 mo | HOM |
| **1013608106** | 2 | VS | 73 | 6.139504255 | 0 | 0 | 0 | 0 | 4 mo | HOM |
| **1011937543** | 1 | VS | 73 | 6.139504255 | 0.008558436 | 55 | 0.001393995 | 8.958378025 | 8 mo | HOM |
| **1010419538** | 2 | VS | 73 | 6.139504255 | 0.000286852 | 15 | 4.67E-05 | 2.443194007 | 4 mo | WILD |
| **1010419187** | 1 | VS | 73 | 6.139504255 | 0.039003393 | 177 | 0.006352857 | 28.82968928 | 8 mo | HOM |
| **1013978366** | 1 | VS | 73 | 6.139504255 | 0.002413244 | 59 | 0.000393068 | 9.609896427 | 2 mo | HOM |
| **1014416155** | 2 | VS | 73 | 6.139504255 | 0 | 0 | 0 | 0 | 2 mo | HOM |
| **1013978065** | 2 | VS | 73 | 6.139504255 | 0.000680994 | 26 | 0.00011092 | 4.234869612 | 2 mo | HOM |
| **1009934102** | 2 | VS | 73 | 6.139504255 | 0.000161205 | 6 | 2.63E-05 | 0.977277603 | 4 mo | WILD |
| **1009322937** | 2 | VS | 73 | 6.139504255 | 0.001080608 | 36 | 0.000176009 | 5.863665616 | 8 mo | WILD |
| **1011080637** | 2 | VS | 73 | 6.139504255 | 0.022315707 | 171 | 0.003634773 | 27.85241168 | 8 mo | HOM |
| **1013609713** | 2 | VS | 73 | 6.139504255 | 0.00016279 | 4 | 2.65E-05 | 0.651518402 | 2 mo | WILD |
| **1010418842** | 1 | VS | 73 | 6.139504255 | 0.045628767 | 153 | 0.007431995 | 24.92057887 | 8 mo | HOM |
| **1013288213** | 2 | VS | 73 | 6.139504255 | 0.001606881 | 58 | 0.000261728 | 9.447016826 | 4 mo | HOM |
| **1008976060** | 2 | VS | 73 | 6.139504255 | 0.013024881 | 121 | 0.002121487 | 19.70843165 | 8 mo | WILD |
| **1013446243** | 1 | VS | 73 | 6.139504255 | 0.010472154 | 140 | 0.0017057 | 22.80314406 | 4 mo | HOM |
| **1013288951** | 2 | VS | 73 | 6.139504255 | 0.023792055 | 146 | 0.00387524 | 23.78042167 | 4 mo | HOM |
| **1013978665** | 1 | VISl6a | 74 | 0.213299842 | 6.53E-06 | 1 | 3.06E-05 | 4.688236009 | 2 mo | HOM |
| **1013445609** | 1 | VISl6a | 74 | 0.213299842 | 5.34E-06 | 1 | 2.5E-05 | 4.688236009 | 4 mo | HOM |
| **1013609015** | 1 | VISl6a | 74 | 0.213299842 | 3.19E-05 | 2 | 0.000149642 | 9.376472018 | 4 mo | HOM |
| **1013608106** | 2 | VISl6a | 74 | 0.213299842 | 0 | 0 | 0 | 0 | 4 mo | HOM |
| **1011937543** | 1 | VISl6a | 74 | 0.213299842 | 0.000252598 | 5 | 0.001184237 | 23.44118005 | 8 mo | HOM |
| **1010419538** | 2 | VISl6a | 74 | 0.213299842 | 0 | 0 | 0 | 0 | 4 mo | WILD |
| **1010419187** | 1 | VISl6a | 74 | 0.213299842 | 0.000409014 | 15 | 0.001917554 | 70.32354014 | 8 mo | HOM |
| **1013978366** | 1 | VISl6a | 74 | 0.213299842 | 0 | 0 | 0 | 0 | 2 mo | HOM |
| **1014416155** | 2 | VISl6a | 74 | 0.213299842 | 0 | 0 | 0 | 0 | 2 mo | HOM |

| | | | | | | | | | | |
|---|---|---|---|---|---|---|---|---|---|---|
| **1013978065** | 2 | VISl6a | 74 | 0.213299842 | 0 | 0 | 0 | 0 | 2 mo | HOM |
| **1009934102** | 2 | VISl6a | 74 | 0.213299842 | 0 | 0 | 0 | 0 | 4 mo | WILD |
| **1009322937** | 2 | VISl6a | 74 | 0.213299842 | 0 | 0 | 0 | 0 | 8 mo | WILD |
| **1011080637** | 2 | VISl6a | 74 | 0.213299842 | 7.63E-05 | 9 | 0.000357725 | 42.19412408 | 8 mo | HOM |
| **1013609713** | 2 | VISl6a | 74 | 0.213299842 | 0 | 0 | 0 | 0 | 2 mo | WILD |
| **1010418842** | 1 | VISl6a | 74 | 0.213299842 | 0.00029413 | 13 | 0.001378949 | 60.94706812 | 8 mo | HOM |
| **1013288213** | 2 | VISl6a | 74 | 0.213299842 | 1.06E-05 | 2 | 4.95E-05 | 9.376472018 | 4 mo | HOM |
| **1008976060** | 2 | VISl6a | 74 | 0.213299842 | 0 | 0 | 0 | 0 | 8 mo | WILD |
| **1013446243** | 1 | VISl6a | 74 | 0.213299842 | 4.79E-05 | 1 | 0.000224659 | 4.688236009 | 4 mo | HOM |
| **1013288951** | 2 | VISl6a | 74 | 0.213299842 | 3.25E-05 | 2 | 0.000152526 | 9.376472018 | 4 mo | HOM |
| **1013978665** | 1 | DT | 75 | 0.011798137 | 0 | 0 | 0 | 0 | 2 mo | HOM |
| **1013445609** | 1 | DT | 75 | 0.011798137 | 0 | 0 | 0 | 0 | 4 mo | HOM |
| **1013609015** | 1 | DT | 75 | 0.011798137 | 0 | 0 | 0 | 0 | 4 mo | HOM |
| **1013608106** | 2 | DT | 75 | 0.011798137 | 0 | 0 | 0 | 0 | 4 mo | HOM |
| **1011937543** | 1 | DT | 75 | 0.011798137 | 9.31E-06 | 1 | 0.000788771 | 84.75914676 | 8 mo | HOM |
| **1010419538** | 2 | DT | 75 | 0.011798137 | 0 | 0 | 0 | 0 | 4 mo | WILD |
| **1010419187** | 1 | DT | 75 | 0.011798137 | 4.55E-06 | 1 | 0.000385525 | 84.75914676 | 8 mo | HOM |
| **1013978366** | 1 | DT | 75 | 0.011798137 | 0 | 0 | 0 | 0 | 2 mo | HOM |
| **1014416155** | 2 | DT | 75 | 0.011798137 | 0 | 0 | 0 | 0 | 2 mo | HOM |
| **1013978065** | 2 | DT | 75 | 0.011798137 | 0 | 0 | 0 | 0 | 2 mo | HOM |
| **1009934102** | 2 | DT | 75 | 0.011798137 | 0 | 0 | 0 | 0 | 4 mo | WILD |
| **1009322937** | 2 | DT | 75 | 0.011798137 | 0 | 0 | 0 | 0 | 8 mo | WILD |
| **1011080637** | 2 | DT | 75 | 0.011798137 | 0 | 0 | 0 | 0 | 8 mo | HOM |
| **1013609713** | 2 | DT | 75 | 0.011798137 | 0 | 0 | 0 | 0 | 2 mo | WILD |
| **1010418842** | 1 | DT | 75 | 0.011798137 | 0 | 0 | 0 | 0 | 8 mo | HOM |
| **1013288213** | 2 | DT | 75 | 0.011798137 | 0 | 0 | 0 | 0 | 4 mo | HOM |
| **1008976060** | 2 | DT | 75 | 0.011798137 | 0 | 0 | 0 | 0 | 8 mo | WILD |
| **1013446243** | 1 | DT | 75 | 0.011798137 | 0 | 0 | 0 | 0 | 4 mo | HOM |
| **1013288951** | 2 | DT | 75 | 0.011798137 | 0 | 0 | 0 | 0 | 4 mo | HOM |
| **1013978665** | 1 | mcp | 78 | 1.127150606 | 0 | 0 | 0 | 0 | 2 mo | HOM |
| **1013445609** | 1 | mcp | 78 | 1.127150606 | 0 | 0 | 0 | 0 | 4 mo | HOM |
| **1013609015** | 1 | mcp | 78 | 1.127150606 | 0 | 0 | 0 | 0 | 4 mo | HOM |
| **1013608106** | 2 | mcp | 78 | 1.127150606 | 0 | 0 | 0 | 0 | 4 mo | HOM |
| **1011937543** | 1 | mcp | 78 | 1.127150606 | 0.001313085 | 12 | 0.00116496 | 10.64631464 | 8 mo | HOM |
| **1010419538** | 2 | mcp | 78 | 1.127150606 | 0 | 0 | 0 | 0 | 4 mo | WILD |
| **1010419187** | 1 | mcp | 78 | 1.127150606 | 2.6E-05 | 1 | 2.3E-05 | 0.887192886 | 8 mo | HOM |
| **1013978366** | 1 | mcp | 78 | 1.127150606 | 1.71E-06 | 0 | 1.51E-06 | 0 | 2 mo | HOM |
| **1014416155** | 2 | mcp | 78 | 1.127150606 | 0 | 0 | 0 | 0 | 2 mo | HOM |
| **1013978065** | 2 | mcp | 78 | 1.127150606 | 0 | 0 | 0 | 0 | 2 mo | HOM |
| **1009934102** | 2 | mcp | 78 | 1.127150606 | 6.01E-06 | 0 | 5.33E-06 | 0 | 4 mo | WILD |
| **1009322937** | 2 | mcp | 78 | 1.127150606 | 0 | 0 | 0 | 0 | 8 mo | WILD |
| **1011080637** | 2 | mcp | 78 | 1.127150606 | 4.09E-06 | 1 | 3.63E-06 | 0.887192886 | 8 mo | HOM |
| **1013609713** | 2 | mcp | 78 | 1.127150606 | 0 | 0 | 0 | 0 | 2 mo | WILD |
| **1010418842** | 1 | mcp | 78 | 1.127150606 | 5.88E-08 | 0 | 5.22E-08 | 0 | 8 mo | HOM |
| **1013288213** | 2 | mcp | 78 | 1.127150606 | 0 | 0 | 0 | 0 | 4 mo | HOM |
| **1008976060** | 2 | mcp | 78 | 1.127150606 | 4.55E-06 | 0 | 4.04E-06 | 0 | 8 mo | WILD |
| **1013446243** | 1 | mcp | 78 | 1.127150606 | 0 | 0 | 0 | 0 | 4 mo | HOM |
| **1013288951** | 2 | mcp | 78 | 1.127150606 | 0 | 0 | 0 | 0 | 4 mo | HOM |
| **1013978665** | 1 | VL | 81 | 3.813130451 | 5.33E-05 | 2 | 1.4E-05 | 0.52450343 | 2 mo | HOM |
| **1013445609** | 1 | VL | 81 | 3.813130451 | 0.001099194 | 44 | 0.000288266 | 11.53907546 | 4 mo | HOM |
| **1013609015** | 1 | VL | 81 | 3.813130451 | 0.009887166 | 98 | 0.002592926 | 25.70066806 | 4 mo | HOM |
| **1013608106** | 2 | VL | 81 | 3.813130765 | 0 | 0 | 0 | 0 | 4 mo | HOM |
| **1011937543** | 1 | VL | 81 | 3.813130451 | 0.003137044 | 33 | 0.000822695 | 8.654306592 | 8 mo | HOM |
| **1010419538** | 2 | VL | 81 | 3.813130765 | 4.52E-05 | 4 | 1.18E-05 | 1.049006773 | 4 mo | WILD |
| **1010419187** | 1 | VL | 81 | 3.813130451 | 0.025497682 | 146 | 0.006686811 | 38.28875038 | 8 mo | HOM |
| **1013978366** | 1 | VL | 81 | 3.813130451 | 0.001299087 | 34 | 0.000340688 | 8.916558307 | 2 mo | HOM |
| **1014416155** | 2 | VL | 81 | 3.813130765 | 0 | 0 | 0 | 0 | 2 mo | HOM |
| **1013978065** | 2 | VL | 81 | 3.813130765 | 0.000384207 | 13 | 0.000100759 | 3.409272013 | 2 mo | HOM |
| **1009934102** | 2 | VL | 81 | 3.813130765 | 9.18E-06 | 1 | 2.41E-06 | 0.262251693 | 4 mo | WILD |

| | | | | | | | | | | |
|---|---|---|---|---|---|---|---|---|---|---|
| 1009322937 | 2 | VL | 81 | 3.813130765 | 3.71E-05 | 5 | 9.73E-06 | 1.311258467 | 8 mo | WILD |
| 1011080637 | 2 | VL | 81 | 3.813130765 | 0.007071945 | 136 | 0.001854629 | 35.66623029 | 8 mo | HOM |
| 1013609713 | 2 | VL | 81 | 3.813130765 | 1.92E-05 | 0 | 5.03E-06 | 0 | 2 mo | WILD |
| 1010418842 | 1 | VL | 81 | 3.813130451 | 0.020996772 | 112 | 0.00550644 | 29.37219207 | 8 mo | HOM |
| 1013288213 | 2 | VL | 81 | 3.813130765 | 0.001082004 | 27 | 0.000283757 | 7.08079572 | 4 mo | HOM |
| 1008976060 | 2 | VL | 81 | 3.813130765 | 0.003695062 | 79 | 0.000969036 | 20.71788377 | 8 mo | WILD |
| 1013446243 | 1 | VL | 81 | 3.813130451 | 0.003482394 | 108 | 0.000913264 | 28.32318521 | 4 mo | HOM |
| 1013288951 | 2 | VL | 81 | 3.813130765 | 0.016057479 | 127 | 0.004211101 | 33.30596505 | 4 mo | HOM |
| 1013978665 | 1 | IO | 83 | 0.527225888 | 0 | 0 | 0 | 0 | 2 mo | HOM |
| 1013445609 | 1 | IO | 83 | 0.527225888 | 0 | 0 | 0 | 0 | 4 mo | HOM |
| 1013609015 | 1 | IO | 83 | 0.527225888 | 0 | 0 | 0 | 0 | 4 mo | HOM |
| 1013608106 | 2 | IO | 83 | 0.527187315 | 0 | 0 | 0 | 0 | 4 mo | HOM |
| 1011937543 | 1 | IO | 83 | 0.527225888 | 0 | 0 | 0 | 0 | 8 mo | HOM |
| 1010419538 | 2 | IO | 83 | 0.527225888 | 0 | 0 | 0 | 0 | 4 mo | WILD |
| 1010419187 | 1 | IO | 83 | 0.527225888 | 0 | 0 | 0 | 0 | 8 mo | HOM |
| 1013978366 | 1 | IO | 83 | 0.527225888 | 0 | 0 | 0 | 0 | 2 mo | HOM |
| 1014416155 | 2 | IO | 83 | 0.527225888 | 0 | 0 | 0 | 0 | 2 mo | HOM |
| 1013978065 | 2 | IO | 83 | 0.527225888 | 0 | 0 | 0 | 0 | 2 mo | HOM |
| 1009934102 | 2 | IO | 83 | 0.527225888 | 0 | 0 | 0 | 0 | 4 mo | WILD |
| 1009322937 | 2 | IO | 83 | 0.527225888 | 0 | 0 | 0 | 0 | 8 mo | WILD |
| 1011080637 | 2 | IO | 83 | 0.527225888 | 0 | 0 | 0 | 0 | 8 mo | HOM |
| 1013609713 | 2 | IO | 83 | 0.527225888 | 0 | 0 | 0 | 0 | 2 mo | WILD |
| 1010418842 | 1 | IO | 83 | 0.527225888 | 0 | 0 | 0 | 0 | 8 mo | HOM |
| 1013288213 | 2 | IO | 83 | 0.527225888 | 0 | 0 | 0 | 0 | 4 mo | HOM |
| 1008976060 | 2 | IO | 83 | 0.527225888 | 0 | 0 | 0 | 0 | 8 mo | WILD |
| 1013446243 | 1 | IO | 83 | 0.527225888 | 0 | 0 | 0 | 0 | 4 mo | HOM |
| 1013288951 | 2 | IO | 83 | 0.527225888 | 0 | 0 | 0 | 0 | 4 mo | HOM |
| 1013978665 | 1 | PL6a | 84 | 0.479194755 | 0 | 0 | 0 | 0 | 2 mo | HOM |
| 1013445609 | 1 | PL6a | 84 | 0.479194755 | 7.92E-06 | 1 | 1.65E-05 | 2.086834192 | 4 mo | HOM |
| 1013609015 | 1 | PL6a | 84 | 0.479194755 | 9.99E-05 | 3 | 0.000208463 | 6.260502577 | 4 mo | HOM |
| 1013608106 | 2 | PL6a | 84 | 0.479194755 | 0 | 0 | 0 | 0 | 4 mo | HOM |
| 1011937543 | 1 | PL6a | 84 | 0.479194755 | 0.00023741 | 2 | 0.000495435 | 4.173668385 | 8 mo | HOM |
| 1010419538 | 2 | PL6a | 84 | 0.479194755 | 0 | 0 | 0 | 0 | 4 mo | WILD |
| 1010419187 | 1 | PL6a | 84 | 0.479194755 | 0.000595773 | 20 | 0.001243279 | 41.73668385 | 8 mo | HOM |
| 1013978366 | 1 | PL6a | 84 | 0.479194755 | 0 | 0 | 0 | 0 | 2 mo | HOM |
| 1014416155 | 2 | PL6a | 84 | 0.479194755 | 0 | 0 | 0 | 0 | 2 mo | HOM |
| 1013978065 | 2 | PL6a | 84 | 0.479194755 | 0 | 0 | 0 | 0 | 2 mo | HOM |
| 1009934102 | 2 | PL6a | 84 | 0.479194755 | 0 | 0 | 0 | 0 | 4 mo | WILD |
| 1009322937 | 2 | PL6a | 84 | 0.479194755 | 0 | 0 | 0 | 0 | 8 mo | WILD |
| 1011080637 | 2 | PL6a | 84 | 0.479194755 | 0.00036061 | 19 | 0.000752534 | 39.64984966 | 8 mo | HOM |
| 1013609713 | 2 | PL6a | 84 | 0.479194755 | 0 | 0 | 0 | 0 | 2 mo | WILD |
| 1010418842 | 1 | PL6a | 84 | 0.479194755 | 0.000546102 | 17 | 0.001139624 | 35.47618127 | 8 mo | HOM |
| 1013288213 | 2 | PL6a | 84 | 0.479194755 | 1.46E-05 | 2 | 3.04E-05 | 4.173668385 | 4 mo | HOM |
| 1008976060 | 2 | PL6a | 84 | 0.479194755 | 6.35E-06 | 1 | 1.32E-05 | 2.086834192 | 8 mo | WILD |
| 1013446243 | 1 | PL6a | 84 | 0.479194755 | 3.44E-05 | 2 | 7.18E-05 | 4.173668385 | 4 mo | HOM |
| 1013288951 | 2 | PL6a | 84 | 0.479194755 | 5.99E-05 | 1 | 0.000124951 | 2.086834192 | 4 mo | HOM |
| 1013978665 | 1 | AHN | 88 | 0.732071682 | 0 | 0 | 0 | 0 | 2 mo | HOM |
| 1013445609 | 1 | AHN | 88 | 0.732071682 | 0 | 0 | 0 | 0 | 4 mo | HOM |
| 1013609015 | 1 | AHN | 88 | 0.732071682 | 0 | 0 | 0 | 0 | 4 mo | HOM |
| 1013608106 | 2 | AHN | 88 | 0.732071682 | 0 | 0 | 0 | 0 | 4 mo | HOM |
| 1011937543 | 1 | AHN | 88 | 0.732071682 | 0.00012676 | 6 | 0.000173153 | 8.195918721 | 8 mo | HOM |
| 1010419538 | 2 | AHN | 88 | 0.732071682 | 0 | 0 | 0 | 0 | 4 mo | WILD |
| 1010419187 | 1 | AHN | 88 | 0.732071682 | 0.000112279 | 5 | 0.000153372 | 6.829932267 | 8 mo | HOM |
| 1013978366 | 1 | AHN | 88 | 0.732071682 | 0 | 0 | 0 | 0 | 2 mo | HOM |
| 1014416155 | 2 | AHN | 88 | 0.732071682 | 0 | 0 | 0 | 0 | 2 mo | HOM |
| 1013978065 | 2 | AHN | 88 | 0.732071682 | 0 | 0 | 0 | 0 | 2 mo | HOM |
| 1009934102 | 2 | AHN | 88 | 0.732071682 | 0 | 0 | 0 | 0 | 4 mo | WILD |
| 1009322937 | 2 | AHN | 88 | 0.732071682 | 0 | 0 | 0 | 0 | 8 mo | WILD |
| 1011080637 | 2 | AHN | 88 | 0.732071682 | 0.000316896 | 10 | 0.000432876 | 13.65986453 | 8 mo | HOM |

| | | | | | | | | | | |
|---|---|---|---|---|---|---|---|---|---|---|
| 1013609713 | 2 | AHN | 88 | 0.732071682 | 0 | 0 | 0 | 0 | 2 mo | WILD |
| 1010418842 | 1 | AHN | 88 | 0.732071682 | 0.000361617 | 12 | 0.000493964 | 16.39183744 | 8 mo | HOM |
| 1013288213 | 2 | AHN | 88 | 0.732071682 | 0 | 0 | 0 | 0 | 4 mo | HOM |
| 1008976060 | 2 | AHN | 88 | 0.732071682 | 0 | 0 | 0 | 0 | 8 mo | WILD |
| 1013446243 | 1 | AHN | 88 | 0.732071682 | 0 | 0 | 0 | 0 | 4 mo | HOM |
| 1013288951 | 2 | AHN | 88 | 0.732071682 | 0 | 0 | 0 | 0 | 4 mo | HOM |
| 1013978665 | 1 | IP | 91 | 0.885863709 | 0 | 0 | 0 | 0 | 2 mo | HOM |
| 1013445609 | 1 | IP | 91 | 0.885863709 | 0 | 0 | 0 | 0 | 4 mo | HOM |
| 1013609015 | 1 | IP | 91 | 0.885863709 | 0 | 0 | 0 | 0 | 4 mo | HOM |
| 1013608106 | 2 | IP | 91 | 0.885863709 | 0 | 0 | 0 | 0 | 4 mo | HOM |
| 1011937543 | 1 | IP | 91 | 0.885863709 | 0.000692567 | 8 | 0.000781799 | 9.030734548 | 8 mo | HOM |
| 1010419538 | 2 | IP | 91 | 0.885863709 | 0 | 0 | 0 | 0 | 4 mo | WILD |
| 1010419187 | 1 | IP | 91 | 0.885863709 | 2.55E-07 | 0 | 2.88E-07 | 0 | 8 mo | HOM |
| 1013978366 | 1 | IP | 91 | 0.885863709 | 0 | 0 | 0 | 0 | 2 mo | HOM |
| 1014416155 | 2 | IP | 91 | 0.885863709 | 0 | 0 | 0 | 0 | 2 mo | HOM |
| 1013978065 | 2 | IP | 91 | 0.885863709 | 0 | 0 | 0 | 0 | 2 mo | HOM |
| 1009934102 | 2 | IP | 91 | 0.885863709 | 0 | 0 | 0 | 0 | 4 mo | WILD |
| 1009322937 | 2 | IP | 91 | 0.885863709 | 0 | 0 | 0 | 0 | 8 mo | WILD |
| 1011080637 | 2 | IP | 91 | 0.885863709 | 0 | 0 | 0 | 0 | 8 mo | HOM |
| 1013609713 | 2 | IP | 91 | 0.885863709 | 0 | 0 | 0 | 0 | 2 mo | WILD |
| 1010418842 | 1 | IP | 91 | 0.885863709 | 0 | 0 | 0 | 0 | 8 mo | HOM |
| 1013288213 | 2 | IP | 91 | 0.885863709 | 0 | 0 | 0 | 0 | 4 mo | HOM |
| 1008976060 | 2 | IP | 91 | 0.885863709 | 1.84E-05 | 2 | 2.08E-05 | 2.257683637 | 8 mo | WILD |
| 1013446243 | 1 | IP | 91 | 0.885863709 | 0 | 0 | 0 | 0 | 4 mo | HOM |
| 1013288951 | 2 | IP | 91 | 0.885863709 | 0 | 0 | 0 | 0 | 4 mo | HOM |
| 1013978665 | 1 | moV | 93 | 0.061349901 | 0 | 0 | 0 | 0 | 2 mo | HOM |
| 1013445609 | 1 | moV | 93 | 0.061349901 | 0 | 0 | 0 | 0 | 4 mo | HOM |
| 1013609015 | 1 | moV | 93 | 0.061349901 | 0 | 0 | 0 | 0 | 4 mo | HOM |
| 1013608106 | 2 | moV | 93 | 0.061349901 | 0 | 0 | 0 | 0 | 4 mo | HOM |
| 1011937543 | 1 | moV | 93 | 0.061349901 | 3.41E-05 | 1 | 0.000555463 | 16.29994475 | 8 mo | HOM |
| 1010419538 | 2 | moV | 93 | 0.061349901 | 0 | 0 | 0 | 0 | 4 mo | WILD |
| 1010419187 | 1 | moV | 93 | 0.061349901 | 0 | 0 | 0 | 0 | 8 mo | HOM |
| 1013978366 | 1 | moV | 93 | 0.061349901 | 0 | 0 | 0 | 0 | 2 mo | HOM |
| 1014416155 | 2 | moV | 93 | 0.061349901 | 0 | 0 | 0 | 0 | 2 mo | HOM |
| 1013978065 | 2 | moV | 93 | 0.061349901 | 0 | 0 | 0 | 0 | 2 mo | HOM |
| 1009934102 | 2 | moV | 93 | 0.061349901 | 0 | 0 | 0 | 0 | 4 mo | WILD |
| 1009322937 | 2 | moV | 93 | 0.061349901 | 0 | 0 | 0 | 0 | 8 mo | WILD |
| 1011080637 | 2 | moV | 93 | 0.061349901 | 0 | 0 | 0 | 0 | 8 mo | HOM |
| 1013609713 | 2 | moV | 93 | 0.061349901 | 0 | 0 | 0 | 0 | 2 mo | WILD |
| 1010418842 | 1 | moV | 93 | 0.061349901 | 0 | 0 | 0 | 0 | 8 mo | HOM |
| 1013288213 | 2 | moV | 93 | 0.061349901 | 0 | 0 | 0 | 0 | 4 mo | HOM |
| 1008976060 | 2 | moV | 93 | 0.061349901 | 0 | 0 | 0 | 0 | 8 mo | WILD |
| 1013446243 | 1 | moV | 93 | 0.061349901 | 0 | 0 | 0 | 0 | 4 mo | HOM |
| 1013288951 | 2 | moV | 93 | 0.061349901 | 0 | 0 | 0 | 0 | 4 mo | HOM |
| 1013978665 | 1 | AI | 95 | 8.132705364 | 0 | 0 | 0 | 0 | 2 mo | HOM |
| 1013445609 | 1 | AI | 95 | 8.132705364 | 0.000595068 | 40 | 7.32E-05 | 4.918412535 | 4 mo | HOM |
| 1013609015 | 1 | AI | 95 | 8.132705364 | 0.001260676 | 60 | 0.000155013 | 7.377618802 | 4 mo | HOM |
| 1013608106 | 2 | AI | 95 | 8.132705364 | 2.26E-07 | 0 | 2.78E-08 | 0 | 4 mo | HOM |
| 1011937543 | 1 | AI | 95 | 8.132705364 | 0.008606941 | 233 | 0.001058312 | 28.64975301 | 8 mo | HOM |
| 1010419538 | 2 | AI | 95 | 8.132705364 | 1.68E-05 | 0 | 2.06E-06 | 0 | 4 mo | WILD |
| 1010419187 | 1 | AI | 95 | 8.132705364 | 0.010227167 | 398 | 0.001257536 | 48.93820472 | 8 mo | HOM |
| 1013978366 | 1 | AI | 95 | 8.132705364 | 2.82E-05 | 3 | 3.46E-06 | 0.36888094 | 2 mo | HOM |
| 1014416155 | 2 | AI | 95 | 8.132705364 | 0 | 0 | 0 | 0 | 2 mo | HOM |
| 1013978065 | 2 | AI | 95 | 8.132705364 | 0 | 0 | 0 | 0 | 2 mo | HOM |
| 1009934102 | 2 | AI | 95 | 8.132705364 | 0 | 0 | 0 | 0 | 4 mo | WILD |
| 1009322937 | 2 | AI | 95 | 8.132705364 | 9.69E-05 | 10 | 1.19E-05 | 1.229603134 | 8 mo | WILD |
| 1011080637 | 2 | AI | 95 | 8.132705364 | 0.00473267 | 250 | 0.000581931 | 30.74007834 | 8 mo | HOM |
| 1013609713 | 2 | AI | 95 | 8.132705364 | 0 | 0 | 0 | 0 | 2 mo | WILD |
| 1010418842 | 1 | AI | 95 | 8.132705364 | 0.007525944 | 326 | 0.000925392 | 40.08506216 | 8 mo | HOM |

| | | | | | | | | | | |
|---|---|---|---|---|---|---|---|---|---|---|
| **1013288213** | 2 | AI | 95 | 8.132705364 | 0.000311591 | 27 | 3.83E-05 | 3.319928461 | 4 mo | HOM |
| **1008976060** | 2 | AI | 95 | 8.132705364 | 0.000177189 | 12 | 2.18E-05 | 1.47552376 | 8 mo | WILD |
| **1013446243** | 1 | AI | 95 | 8.132705364 | 0.001106555 | 74 | 0.000136062 | 9.099063189 | 4 mo | HOM |
| **1013288951** | 2 | AI | 95 | 8.132705364 | 0.001538169 | 66 | 0.000189134 | 8.115380682 | 4 mo | HOM |
| **1013978665** | 1 | DCO | 96 | 0.630699619 | 2.26E-05 | 3 | 3.58E-05 | 4.756622501 | 2 mo | HOM |
| **1013445609** | 1 | DCO | 96 | 0.630699619 | 1.33E-07 | 0 | 2.1E-07 | 0 | 4 mo | HOM |
| **1013609015** | 1 | DCO | 96 | 0.630699619 | 0.00288367 | 6 | 0.004572176 | 9.513245002 | 4 mo | HOM |
| **1013608106** | 2 | DCO | 96 | 0.630699619 | 0 | 0 | 0 | 0 | 4 mo | HOM |
| **1011937543** | 1 | DCO | 96 | 0.630699619 | 0.0033537 | 4 | 0.005317429 | 6.342163335 | 8 mo | HOM |
| **1010419538** | 2 | DCO | 96 | 0.630699619 | 1.95E-05 | 4 | 3.09E-05 | 6.342163335 | 4 mo | WILD |
| **1010419187** | 1 | DCO | 96 | 0.630699619 | 0.00016533 | 6 | 0.000262137 | 9.513245002 | 8 mo | HOM |
| **1013978366** | 1 | DCO | 96 | 0.630699619 | 0.00023254 | 4 | 0.000368702 | 6.342163335 | 2 mo | HOM |
| **1014416155** | 2 | DCO | 96 | 0.630699619 | 0 | 0 | 0 | 0 | 2 mo | HOM |
| **1013978065** | 2 | DCO | 96 | 0.630699619 | 3.82E-06 | 0 | 6.05E-06 | 0 | 2 mo | HOM |
| **1009934102** | 2 | DCO | 96 | 0.630699619 | 1.91E-05 | 2 | 3.03E-05 | 3.171081667 | 4 mo | WILD |
| **1009322937** | 2 | DCO | 96 | 0.630699619 | 0.000139066 | 6 | 0.000220495 | 9.513245002 | 8 mo | WILD |
| **1011080637** | 2 | DCO | 96 | 0.630699619 | 0.000463069 | 12 | 0.000734215 | 19.02649 | 8 mo | HOM |
| **1013609713** | 2 | DCO | 96 | 0.630699619 | 0 | 0 | 0 | 0 | 2 mo | WILD |
| **1010418842** | 1 | DCO | 96 | 0.630699619 | 0.000146627 | 9 | 0.000232483 | 14.2698675 | 8 mo | HOM |
| **1013288213** | 2 | DCO | 96 | 0.630699619 | 2.66E-06 | 0 | 4.22E-06 | 0 | 4 mo | HOM |
| **1008976060** | 2 | DCO | 96 | 0.630699619 | 0.001185884 | 14 | 0.001880267 | 22.19757167 | 8 mo | WILD |
| **1013446243** | 1 | DCO | 96 | 0.630699619 | 1.26E-05 | 1 | 2E-05 | 1.585540834 | 4 mo | HOM |
| **1013288951** | 2 | DCO | 96 | 0.630699619 | 0.000244867 | 4 | 0.000388246 | 6.342163335 | 4 mo | HOM |
| **1013978665** | 1 | TEa1 | 97 | 0.541286379 | 2.41E-07 | 0 | 4.46E-07 | 0 | 2 mo | HOM |
| **1013445609** | 1 | TEa1 | 97 | 0.541286379 | 2.17E-05 | 3 | 4.01E-05 | 5.542352653 | 4 mo | HOM |
| **1013609015** | 1 | TEa1 | 97 | 0.541286379 | 0.000283114 | 6 | 0.000523039 | 11.08470531 | 4 mo | HOM |
| **1013608106** | 2 | TEa1 | 97 | 0.541286379 | 0 | 0 | 0 | 0 | 4 mo | HOM |
| **1011937543** | 1 | TEa1 | 97 | 0.541286379 | 0.004887176 | 16 | 0.009028818 | 29.55921415 | 8 mo | HOM |
| **1010419538** | 2 | TEa1 | 97 | 0.541286379 | 8.25E-06 | 1 | 1.52E-05 | 1.847450884 | 4 mo | WILD |
| **1010419187** | 1 | TEa1 | 97 | 0.541286379 | 0.002126178 | 30 | 0.003928008 | 55.42352653 | 8 mo | HOM |
| **1013978366** | 1 | TEa1 | 97 | 0.541286379 | 0 | 0 | 0 | 0 | 2 mo | HOM |
| **1014416155** | 2 | TEa1 | 97 | 0.541286379 | 0 | 0 | 0 | 0 | 2 mo | HOM |
| **1013978065** | 2 | TEa1 | 97 | 0.541286379 | 0 | 0 | 0 | 0 | 2 mo | HOM |
| **1009934102** | 2 | TEa1 | 97 | 0.541286379 | 0 | 0 | 0 | 0 | 4 mo | WILD |
| **1009322937** | 2 | TEa1 | 97 | 0.541286379 | 0 | 0 | 0 | 0 | 8 mo | WILD |
| **1011080637** | 2 | TEa1 | 97 | 0.541286379 | 0.000774887 | 28 | 0.001431567 | 51.72862476 | 8 mo | HOM |
| **1013609713** | 2 | TEa1 | 97 | 0.541286379 | 9.31E-06 | 2 | 1.72E-05 | 3.694901769 | 2 mo | WILD |
| **1010418842** | 1 | TEa1 | 97 | 0.541286379 | 0.001446558 | 37 | 0.002672445 | 68.35568273 | 8 mo | HOM |
| **1013288213** | 2 | TEa1 | 97 | 0.541286379 | 7.89E-05 | 7 | 0.000145742 | 12.93215619 | 4 mo | HOM |
| **1008976060** | 2 | TEa1 | 97 | 0.541286379 | 4.91E-05 | 7 | 9.08E-05 | 12.93215619 | 8 mo | WILD |
| **1013446243** | 1 | TEa1 | 97 | 0.541286379 | 0.000148893 | 9 | 0.000275073 | 16.62705796 | 4 mo | HOM |
| **1013288951** | 2 | TEa1 | 97 | 0.541286379 | 0.000209966 | 10 | 0.000387901 | 18.47450884 | 4 mo | HOM |
| **1013978665** | 1 | SEZ | 98 | 0.140609224 | 0 | 0 | 0 | 0 | 2 mo | HOM |
| **1013445609** | 1 | SEZ | 98 | 0.140609224 | 0.000342445 | 17 | 0.002435436 | 120.9024523 | 4 mo | HOM |
| **1013609015** | 1 | SEZ | 98 | 0.140609224 | 0.000404518 | 3 | 0.002876896 | 21.33572688 | 4 mo | HOM |
| **1013608106** | 2 | SEZ | 98 | 0.140609224 | 0 | 0 | 0 | 0 | 4 mo | HOM |
| **1011937543** | 1 | SEZ | 98 | 0.140609224 | 6.77E-06 | 0 | 4.81E-05 | 0 | 8 mo | HOM |
| **1010419538** | 2 | SEZ | 98 | 0.140609224 | 0 | 0 | 0 | 0 | 4 mo | WILD |
| **1010419187** | 1 | SEZ | 98 | 0.140609224 | 0.002832608 | 9 | 0.020145247 | 64.00718064 | 8 mo | HOM |
| **1013978366** | 1 | SEZ | 98 | 0.140609224 | 2.49E-05 | 0 | 0.000177126 | 0 | 2 mo | HOM |
| **1014416155** | 2 | SEZ | 98 | 0.140609224 | 0 | 0 | 0 | 0 | 2 mo | HOM |
| **1013978065** | 2 | SEZ | 98 | 0.140609224 | 3.99E-06 | 0 | 2.84E-05 | 0 | 2 mo | HOM |
| **1009934102** | 2 | SEZ | 98 | 0.140609224 | 0 | 0 | 0 | 0 | 4 mo | WILD |
| **1009322937** | 2 | SEZ | 98 | 0.140609224 | 1.08E-06 | 0 | 7.7E-06 | 0 | 8 mo | WILD |
| **1011080637** | 2 | SEZ | 98 | 0.140609224 | 0.000211229 | 8 | 0.001502243 | 56.89527168 | 8 mo | HOM |
| **1013609713** | 2 | SEZ | 98 | 0.140609224 | 0 | 0 | 0 | 0 | 2 mo | WILD |
| **1010418842** | 1 | SEZ | 98 | 0.140609224 | 0.000141412 | 5 | 0.001005712 | 35.5595448 | 8 mo | HOM |
| **1013288213** | 2 | SEZ | 98 | 0.140609224 | 0.000288525 | 6 | 0.002051966 | 42.67145376 | 4 mo | HOM |
| **1008976060** | 2 | SEZ | 98 | 0.140609224 | 0.00012086 | 3 | 0.000859546 | 21.33572688 | 8 mo | WILD |

| | | | | | | | | | | |
|---|---|---|---|---|---|---|---|---|---|---|
| **1013446243** | 1 | SEZ | 98 | 0.140609224 | 0.000163961 | 14 | 0.001166079 | 99.56672544 | 4 mo | HOM |
| **1013288951** | 2 | SEZ | 98 | 0.140609224 | 0.000411636 | 2 | 0.002927521 | 14.22381792 | 4 mo | HOM |
| **1013978665** | 1 | IPN | 100 | 0.365700994 | 0 | 0 | 0 | 0 | 2 mo | HOM |
| **1013445609** | 1 | IPN | 100 | 0.365700994 | 0 | 0 | 0 | 0 | 4 mo | HOM |
| **1013609015** | 1 | IPN | 100 | 0.365700994 | 0 | 0 | 0 | 0 | 4 mo | HOM |
| **1013608106** | 2 | IPN | 100 | 0.365701034 | 0 | 0 | 0 | 0 | 4 mo | HOM |
| **1011937543** | 1 | IPN | 100 | 0.365700994 | 0.000223015 | 1 | 0.000609829 | 2.73447438 | 8 mo | HOM |
| **1010419538** | 2 | IPN | 100 | 0.365701034 | 1.83E-07 | 0 | 5.01E-07 | 0 | 4 mo | WILD |
| **1010419187** | 1 | IPN | 100 | 0.365700994 | 0.000150324 | 3 | 0.000411057 | 8.203423141 | 8 mo | HOM |
| **1013978366** | 1 | IPN | 100 | 0.365700994 | 0 | 0 | 0 | 0 | 2 mo | HOM |
| **1014416155** | 2 | IPN | 100 | 0.365701034 | 0 | 0 | 0 | 0 | 2 mo | HOM |
| **1013978065** | 2 | IPN | 100 | 0.365701034 | 0 | 0 | 0 | 0 | 2 mo | HOM |
| **1009934102** | 2 | IPN | 100 | 0.365701034 | 1.1E-05 | 1 | 3E-05 | 2.734474087 | 4 mo | WILD |
| **1009322937** | 2 | IPN | 100 | 0.365701034 | 1.71E-05 | 2 | 4.68E-05 | 5.468948174 | 8 mo | WILD |
| **1011080637** | 2 | IPN | 100 | 0.365701034 | 5.45E-06 | 1 | 1.49E-05 | 2.734474087 | 8 mo | HOM |
| **1013609713** | 2 | IPN | 100 | 0.365701034 | 0 | 0 | 0 | 0 | 2 mo | WILD |
| **1010418842** | 1 | IPN | 100 | 0.365700994 | 6.1E-05 | 7 | 0.000166897 | 19.14132066 | 8 mo | HOM |
| **1013288213** | 2 | IPN | 100 | 0.365701034 | 0 | 0 | 0 | 0 | 4 mo | HOM |
| **1008976060** | 2 | IPN | 100 | 0.365701034 | 0.0001181 | 8 | 0.00032294 | 21.8757927 | 8 mo | WILD |
| **1013446243** | 1 | IPN | 100 | 0.365700994 | 1.11E-06 | 0 | 3.03E-06 | 0 | 4 mo | HOM |
| **1013288951** | 2 | IPN | 100 | 0.365701034 | 1.1E-05 | 1 | 3E-05 | 2.734474087 | 4 mo | HOM |
| **1013978665** | 1 | VCO | 101 | 1.047827446 | 1.09E-05 | 1 | 1.04E-05 | 0.954355608 | 2 mo | HOM |
| **1013445609** | 1 | VCO | 101 | 1.047827446 | 0.000160803 | 4 | 0.000153463 | 3.817422433 | 4 mo | HOM |
| **1013609015** | 1 | VCO | 101 | 1.047827446 | 2.94E-05 | 2 | 2.8E-05 | 1.908711217 | 4 mo | HOM |
| **1013608106** | 2 | VCO | 101 | 1.047827446 | 0 | 0 | 0 | 0 | 4 mo | HOM |
| **1011937543** | 1 | VCO | 101 | 1.047827446 | 0.001293781 | 2 | 0.001234727 | 1.908711217 | 8 mo | HOM |
| **1010419538** | 2 | VCO | 101 | 1.047827446 | 6.26E-06 | 1 | 5.97E-06 | 0.954355608 | 4 mo | WILD |
| **1010419187** | 1 | VCO | 101 | 1.047827446 | 9.87E-05 | 2 | 9.42E-05 | 1.908711217 | 8 mo | HOM |
| **1013978366** | 1 | VCO | 101 | 1.047827446 | 0.000829742 | 1 | 0.000791869 | 0.954355608 | 2 mo | HOM |
| **1014416155** | 2 | VCO | 101 | 1.047827446 | 0 | 0 | 0 | 0 | 2 mo | HOM |
| **1013978065** | 2 | VCO | 101 | 1.047827446 | 6.88E-05 | 3 | 6.57E-05 | 2.863066825 | 2 mo | HOM |
| **1009934102** | 2 | VCO | 101 | 1.047827446 | 9.39E-10 | 0 | 8.96E-10 | 0 | 4 mo | WILD |
| **1009322937** | 2 | VCO | 101 | 1.047827446 | 1.26E-05 | 0 | 1.21E-05 | 0 | 8 mo | WILD |
| **1011080637** | 2 | VCO | 101 | 1.047827446 | 0.000154824 | 8 | 0.000147757 | 7.634844866 | 8 mo | HOM |
| **1013609713** | 2 | VCO | 101 | 1.047827446 | 3.69E-07 | 0 | 3.52E-07 | 0 | 2 mo | WILD |
| **1010418842** | 1 | VCO | 101 | 1.047827446 | 2.22E-05 | 2 | 2.12E-05 | 1.908711217 | 8 mo | HOM |
| **1013288213** | 2 | VCO | 101 | 1.047827446 | 8.08E-11 | 0 | 7.71E-11 | 0 | 4 mo | HOM |
| **1008976060** | 2 | VCO | 101 | 1.047827446 | 0.000374215 | 6 | 0.000357134 | 5.72613365 | 8 mo | WILD |
| **1013446243** | 1 | VCO | 101 | 1.047827446 | 7.28E-05 | 2 | 6.94E-05 | 1.908711217 | 4 mo | HOM |
| **1013288951** | 2 | VCO | 101 | 1.047827446 | 4.32E-06 | 0 | 4.12E-06 | 0 | 4 mo | HOM |
| **1013978665** | 1 | nst | 102 | 0.115760207 | 0 | 0 | 0 | 0 | 2 mo | HOM |
| **1013445609** | 1 | nst | 102 | 0.115760207 | 0 | 0 | 0 | 0 | 4 mo | HOM |
| **1013609015** | 1 | nst | 102 | 0.115760207 | 0 | 0 | 0 | 0 | 4 mo | HOM |
| **1013608106** | 2 | nst | 102 | 0.115760207 | 0 | 0 | 0 | 0 | 4 mo | HOM |
| **1011937543** | 1 | nst | 102 | 0.115760207 | 0.000165372 | 3 | 0.001428578 | 25.91564133 | 8 mo | HOM |
| **1010419538** | 2 | nst | 102 | 0.115760207 | 0 | 0 | 0 | 0 | 4 mo | WILD |
| **1010419187** | 1 | nst | 102 | 0.115760207 | 0 | 0 | 0 | 0 | 8 mo | HOM |
| **1013978366** | 1 | nst | 102 | 0.115760207 | 0 | 0 | 0 | 0 | 2 mo | HOM |
| **1014416155** | 2 | nst | 102 | 0.115760207 | 0 | 0 | 0 | 0 | 2 mo | HOM |
| **1013978065** | 2 | nst | 102 | 0.115760207 | 0 | 0 | 0 | 0 | 2 mo | HOM |
| **1009934102** | 2 | nst | 102 | 0.115760207 | 0 | 0 | 0 | 0 | 4 mo | WILD |
| **1009322937** | 2 | nst | 102 | 0.115760207 | 0 | 0 | 0 | 0 | 8 mo | WILD |
| **1011080637** | 2 | nst | 102 | 0.115760207 | 0 | 0 | 0 | 0 | 8 mo | HOM |
| **1013609713** | 2 | nst | 102 | 0.115760207 | 0 | 0 | 0 | 0 | 2 mo | WILD |
| **1010418842** | 1 | nst | 102 | 0.115760207 | 2.23E-05 | 2 | 0.000192795 | 17.27709422 | 8 mo | HOM |
| **1013288213** | 2 | nst | 102 | 0.115760207 | 0 | 0 | 0 | 0 | 4 mo | HOM |
| **1008976060** | 2 | nst | 102 | 0.115760207 | 0 | 0 | 0 | 0 | 8 mo | WILD |
| **1013446243** | 1 | nst | 102 | 0.115760207 | 0 | 0 | 0 | 0 | 4 mo | HOM |
| **1013288951** | 2 | nst | 102 | 0.115760207 | 0 | 0 | 0 | 0 | 4 mo | HOM |

| | | | | | | | | | | |
|---|---|---|---|---|---|---|---|---|---|---|
| **1013978665** | 1 | Ald | 104 | 3.837227563 | 0 | 0 | 0 | 0 | 2 mo | HOM |
| **1013445609** | 1 | Ald | 104 | 3.837227563 | 0.000264666 | 15 | 6.9E-05 | 3.909072306 | 4 mo | HOM |
| **1013609015** | 1 | Ald | 104 | 3.837227563 | 0.000593055 | 25 | 0.000154553 | 6.51512051 | 4 mo | HOM |
| **1013608106** | 2 | Ald | 104 | 3.837227563 | 0 | 0 | 0 | 0 | 4 mo | HOM |
| **1011937543** | 1 | Ald | 104 | 3.837227563 | 0.004101675 | 122 | 0.001068916 | 31.79378809 | 8 mo | HOM |
| **1010419538** | 2 | Ald | 104 | 3.837227563 | 1.68E-05 | 0 | 4.37E-06 | 0 | 4 mo | WILD |
| **1010419187** | 1 | Ald | 104 | 3.837227563 | 0.004112846 | 172 | 0.001071827 | 44.82402911 | 8 mo | HOM |
| **1013978366** | 1 | Ald | 104 | 3.837227563 | 2.24E-05 | 2 | 5.85E-06 | 0.521209641 | 2 mo | HOM |
| **1014416155** | 2 | Ald | 104 | 3.837227563 | 0 | 0 | 0 | 0 | 2 mo | HOM |
| **1013978065** | 2 | Ald | 104 | 3.837227563 | 0 | 0 | 0 | 0 | 2 mo | HOM |
| **1009934102** | 2 | Ald | 104 | 3.837227563 | 0 | 0 | 0 | 0 | 4 mo | WILD |
| **1009322937** | 2 | Ald | 104 | 3.837227563 | 2.84E-05 | 5 | 7.41E-06 | 1.303024102 | 8 mo | WILD |
| **1011080637** | 2 | Ald | 104 | 3.837227563 | 0.00191847 | 97 | 0.000499962 | 25.27866758 | 8 mo | HOM |
| **1013609713** | 2 | Ald | 104 | 3.837227563 | 0 | 0 | 0 | 0 | 2 mo | WILD |
| **1010418842** | 1 | Ald | 104 | 3.837227563 | 0.002987823 | 132 | 0.000778641 | 34.39983629 | 8 mo | HOM |
| **1013288213** | 2 | Ald | 104 | 3.837227563 | 0.000117361 | 12 | 3.06E-05 | 3.127257845 | 4 mo | HOM |
| **1008976060** | 2 | Ald | 104 | 3.837227563 | 2.42E-05 | 4 | 6.29E-06 | 1.042419282 | 8 mo | WILD |
| **1013446243** | 1 | Ald | 104 | 3.837227563 | 0.00069363 | 36 | 0.000180763 | 9.381773534 | 4 mo | HOM |
| **1013288951** | 2 | Ald | 104 | 3.837227563 | 0.000667345 | 24 | 0.000173913 | 6.25451569 | 4 mo | HOM |
| **1013978665** | 1 | SOCm | 105 | 0.201114346 | 0 | 0 | 0 | 0 | 2 mo | HOM |
| **1013445609** | 1 | SOCm | 105 | 0.201114346 | 0 | 0 | 0 | 0 | 4 mo | HOM |
| **1013609015** | 1 | SOCm | 105 | 0.201114346 | 0 | 0 | 0 | 0 | 4 mo | HOM |
| **1013608106** | 2 | SOCm | 105 | 0.201114346 | 0 | 0 | 0 | 0 | 4 mo | HOM |
| **1011937543** | 1 | SOCm | 105 | 0.201114346 | 0 | 0 | 0 | 0 | 8 mo | HOM |
| **1010419538** | 2 | SOCm | 105 | 0.201114346 | 0 | 0 | 0 | 0 | 4 mo | WILD |
| **1010419187** | 1 | SOCm | 105 | 0.201114346 | 0 | 0 | 0 | 0 | 8 mo | HOM |
| **1013978366** | 1 | SOCm | 105 | 0.201114346 | 0 | 0 | 0 | 0 | 2 mo | HOM |
| **1014416155** | 2 | SOCm | 105 | 0.201114346 | 0 | 0 | 0 | 0 | 2 mo | HOM |
| **1013978065** | 2 | SOCm | 105 | 0.201114346 | 0 | 0 | 0 | 0 | 2 mo | HOM |
| **1009934102** | 2 | SOCm | 105 | 0.201114346 | 0 | 0 | 0 | 0 | 4 mo | WILD |
| **1009322937** | 2 | SOCm | 105 | 0.201114346 | 0 | 0 | 0 | 0 | 8 mo | WILD |
| **1011080637** | 2 | SOCm | 105 | 0.201114346 | 0 | 0 | 0 | 0 | 8 mo | HOM |
| **1013609713** | 2 | SOCm | 105 | 0.201114346 | 0 | 0 | 0 | 0 | 2 mo | WILD |
| **1010418842** | 1 | SOCm | 105 | 0.201114346 | 0 | 0 | 0 | 0 | 8 mo | HOM |
| **1013288213** | 2 | SOCm | 105 | 0.201114346 | 0 | 0 | 0 | 0 | 4 mo | HOM |
| **1008976060** | 2 | SOCm | 105 | 0.201114346 | 0 | 0 | 0 | 0 | 8 mo | WILD |
| **1013446243** | 1 | SOCm | 105 | 0.201114346 | 0 | 0 | 0 | 0 | 4 mo | HOM |
| **1013288951** | 2 | SOCm | 105 | 0.201114346 | 0 | 0 | 0 | 0 | 4 mo | HOM |
| **1013978665** | 1 | ISN | 106 | 0.007442345 | 0 | 0 | 0 | 0 | 2 mo | HOM |
| **1013445609** | 1 | ISN | 106 | 0.007442345 | 0 | 0 | 0 | 0 | 4 mo | HOM |
| **1013609015** | 1 | ISN | 106 | 0.007442345 | 0 | 0 | 0 | 0 | 4 mo | HOM |
| **1013608106** | 2 | ISN | 106 | 0.007442345 | 0 | 0 | 0 | 0 | 4 mo | HOM |
| **1011937543** | 1 | ISN | 106 | 0.007442345 | 1.19E-10 | 0 | 1.6E-08 | 0 | 8 mo | HOM |
| **1010419538** | 2 | ISN | 106 | 0.007442345 | 0 | 0 | 0 | 0 | 4 mo | WILD |
| **1010419187** | 1 | ISN | 106 | 0.007442345 | 0 | 0 | 0 | 0 | 8 mo | HOM |
| **1013978366** | 1 | ISN | 106 | 0.007442345 | 0 | 0 | 0 | 0 | 2 mo | HOM |
| **1014416155** | 2 | ISN | 106 | 0.007442345 | 0 | 0 | 0 | 0 | 2 mo | HOM |
| **1013978065** | 2 | ISN | 106 | 0.007442345 | 0 | 0 | 0 | 0 | 2 mo | HOM |
| **1009934102** | 2 | ISN | 106 | 0.007442345 | 0 | 0 | 0 | 0 | 4 mo | WILD |
| **1009322937** | 2 | ISN | 106 | 0.007442345 | 0 | 0 | 0 | 0 | 8 mo | WILD |
| **1011080637** | 2 | ISN | 106 | 0.007442345 | 0 | 0 | 0 | 0 | 8 mo | HOM |
| **1013609713** | 2 | ISN | 106 | 0.007442345 | 0 | 0 | 0 | 0 | 2 mo | WILD |
| **1010418842** | 1 | ISN | 106 | 0.007442345 | 0 | 0 | 0 | 0 | 8 mo | HOM |
| **1013288213** | 2 | ISN | 106 | 0.007442345 | 0 | 0 | 0 | 0 | 4 mo | HOM |
| **1008976060** | 2 | ISN | 106 | 0.007442345 | 0 | 0 | 0 | 0 | 8 mo | WILD |
| **1013446243** | 1 | ISN | 106 | 0.007442345 | 0 | 0 | 0 | 0 | 4 mo | HOM |
| **1013288951** | 2 | ISN | 106 | 0.007442345 | 0 | 0 | 0 | 0 | 4 mo | HOM |
| **1013978665** | 1 | chpl | 108 | 1.489740336 | 1.07E-05 | 0 | 7.16E-06 | 0 | 2 mo | HOM |
| **1013445609** | 1 | chpl | 108 | 1.489740336 | 0.000119018 | 2 | 7.99E-05 | 1.342515841 | 4 mo | HOM |

| | | | | | | | | | | |
|---|---|---|---|---|---|---|---|---|---|---|
| **1013609015** | 1 | chpl | 108 | 1.489740336 | 0.00504279 | 37 | 0.003385013 | 24.83654306 | 4 mo | HOM |
| **1013608106** | 2 | chpl | 108 | 1.489740336 | 0 | 0 | 0 | 0 | 4 mo | HOM |
| **1011937543** | 1 | chpl | 108 | 1.489740336 | 0.00203511 | 15 | 0.001366084 | 10.06886881 | 8 mo | HOM |
| **1010419538** | 2 | chpl | 108 | 1.489740336 | 4.52E-05 | 4 | 3.03E-05 | 2.685031682 | 4 mo | WILD |
| **1010419187** | 1 | chpl | 108 | 1.489740336 | 0.009483489 | 58 | 0.006365867 | 38.93295939 | 8 mo | HOM |
| **1013978366** | 1 | chpl | 108 | 1.489740336 | 0.000495233 | 11 | 0.000332429 | 7.383837125 | 2 mo | HOM |
| **1014416155** | 2 | chpl | 108 | 1.489740336 | 0 | 0 | 0 | 0 | 2 mo | HOM |
| **1013978065** | 2 | chpl | 108 | 1.489740336 | 0.000284562 | 8 | 0.000191015 | 5.370063364 | 2 mo | HOM |
| **1009934102** | 2 | chpl | 108 | 1.489740336 | 9.18E-06 | 1 | 6.16E-06 | 0.67125792 | 4 mo | WILD |
| **1009322937** | 2 | chpl | 108 | 1.489740336 | 1.86E-05 | 1 | 1.25E-05 | 0.67125792 | 8 mo | WILD |
| **1011080637** | 2 | chpl | 108 | 1.489740336 | 0.004258844 | 58 | 0.002858783 | 38.93295939 | 8 mo | HOM |
| **1013609713** | 2 | chpl | 108 | 1.489740336 | 1.92E-05 | 0 | 1.29E-05 | 0 | 2 mo | WILD |
| **1010418842** | 1 | chpl | 108 | 1.489740336 | 0.010355208 | 37 | 0.006951015 | 24.83654306 | 8 mo | HOM |
| **1013288213** | 2 | chpl | 108 | 1.489740336 | 4.63E-05 | 3 | 3.11E-05 | 2.013773761 | 4 mo | HOM |
| **1008976060** | 2 | chpl | 108 | 1.489740336 | 0.002119164 | 22 | 0.001422505 | 14.76767425 | 8 mo | WILD |
| **1013446243** | 1 | chpl | 108 | 1.489740336 | 0.001878508 | 40 | 0.001260963 | 26.85031682 | 4 mo | HOM |
| **1013288951** | 2 | chpl | 108 | 1.489740336 | 0.00812899 | 52 | 0.005456649 | 34.90541187 | 4 mo | HOM |
| **1013978665** | 1 | AIp | 111 | 2.508212517 | 0 | 0 | 0 | 0 | 2 mo | HOM |
| **1013445609** | 1 | AIp | 111 | 2.508212517 | 0.000272192 | 22 | 0.00010852 | 8.771186593 | 4 mo | HOM |
| **1013609015** | 1 | AIp | 111 | 2.508212517 | 0.000567996 | 28 | 0.000226455 | 11.16332839 | 4 mo | HOM |
| **1013608106** | 2 | AIp | 111 | 2.508212517 | 0 | 0 | 0 | 0 | 4 mo | HOM |
| **1011937543** | 1 | AIp | 111 | 2.508212517 | 0.002733588 | 73 | 0.001089855 | 29.10439188 | 8 mo | HOM |
| **1010419538** | 2 | AIp | 111 | 2.508212517 | 3.9E-09 | 0 | 1.55E-09 | 0 | 4 mo | WILD |
| **1010419187** | 1 | AIp | 111 | 2.508212517 | 0.003640995 | 140 | 0.001451629 | 55.81664195 | 8 mo | HOM |
| **1013978366** | 1 | AIp | 111 | 2.508212517 | 0 | 0 | 0 | 0 | 2 mo | HOM |
| **1014416155** | 2 | AIp | 111 | 2.508212517 | 0 | 0 | 0 | 0 | 2 mo | HOM |
| **1013978065** | 2 | AIp | 111 | 2.508212517 | 0 | 0 | 0 | 0 | 2 mo | HOM |
| **1009934102** | 2 | AIp | 111 | 2.508212517 | 0 | 0 | 0 | 0 | 4 mo | WILD |
| **1009322937** | 2 | AIp | 111 | 2.508212517 | 0 | 0 | 0 | 0 | 8 mo | WILD |
| **1011080637** | 2 | AIp | 111 | 2.508212517 | 0.001811166 | 100 | 0.000722094 | 39.86902997 | 8 mo | HOM |
| **1013609713** | 2 | AIp | 111 | 2.508212517 | 0 | 0 | 0 | 0 | 2 mo | WILD |
| **1010418842** | 1 | AIp | 111 | 2.508212517 | 0.00312457 | 132 | 0.001245736 | 52.62711956 | 8 mo | HOM |
| **1013288213** | 2 | AIp | 111 | 2.508212517 | 0.000100464 | 12 | 4.01E-05 | 4.784283596 | 4 mo | HOM |
| **1008976060** | 2 | AIp | 111 | 2.508212517 | 7.56E-06 | 2 | 3.01E-06 | 0.797380599 | 8 mo | WILD |
| **1013446243** | 1 | AIp | 111 | 2.508212517 | 0.000313777 | 27 | 0.0001251 | 10.76463809 | 4 mo | HOM |
| **1013288951** | 2 | AIp | 111 | 2.508212517 | 0.000670895 | 38 | 0.000267479 | 15.15023139 | 4 mo | HOM |
| **1013978665** | 1 | SSp-ll2/3 | 113 | 0.666114624 | 0 | 0 | 0 | 0 | 2 mo | HOM |
| **1013445609** | 1 | SSp-ll2/3 | 113 | 0.666114624 | 1.93E-05 | 1 | 2.89E-05 | 1.501243125 | 4 mo | HOM |
| **1013609015** | 1 | SSp-ll2/3 | 113 | 0.666114624 | 0.00022883 | 6 | 0.000343529 | 9.007458752 | 4 mo | HOM |
| **1013608106** | 2 | SSp-ll2/3 | 113 | 0.666114624 | 0 | 0 | 0 | 0 | 4 mo | HOM |
| **1011937543** | 1 | SSp-ll2/3 | 113 | 0.666114624 | 0.000640504 | 17 | 0.000961553 | 25.52113313 | 8 mo | HOM |
| **1010419538** | 2 | SSp-ll2/3 | 113 | 0.666114624 | 0 | 0 | 0 | 0 | 4 mo | WILD |
| **1010419187** | 1 | SSp-ll2/3 | 113 | 0.666114624 | 0.001449225 | 34 | 0.002175638 | 51.04226626 | 8 mo | HOM |
| **1013978366** | 1 | SSp-ll2/3 | 113 | 0.666114624 | 0 | 0 | 0 | 0 | 2 mo | HOM |
| **1014416155** | 2 | SSp-ll2/3 | 113 | 0.666114624 | 0 | 0 | 0 | 0 | 2 mo | HOM |
| **1013978065** | 2 | SSp-ll2/3 | 113 | 0.666114624 | 0 | 0 | 0 | 0 | 2 mo | HOM |
| **1009934102** | 2 | SSp-ll2/3 | 113 | 0.666114624 | 0 | 0 | 0 | 0 | 4 mo | WILD |
| **1009322937** | 2 | SSp-ll2/3 | 113 | 0.666114624 | 0 | 0 | 0 | 0 | 8 mo | WILD |
| **1011080637** | 2 | SSp-ll2/3 | 113 | 0.666114624 | 0.000386732 | 19 | 0.000580579 | 28.52361938 | 8 mo | HOM |
| **1013609713** | 2 | SSp-ll2/3 | 113 | 0.666114624 | 0 | 0 | 0 | 0 | 2 mo | WILD |
| **1010418842** | 1 | SSp-ll2/3 | 113 | 0.666114624 | 0.000674647 | 20 | 0.001012809 | 30.02486251 | 8 mo | HOM |
| **1013288213** | 2 | SSp-ll2/3 | 113 | 0.666114624 | 4.79E-05 | 3 | 7.19E-05 | 4.503729376 | 4 mo | HOM |
| **1008976060** | 2 | SSp-ll2/3 | 113 | 0.666114624 | 0 | 0 | 0 | 0 | 8 mo | WILD |
| **1013446243** | 1 | SSp-ll2/3 | 113 | 0.666114624 | 0.000116598 | 5 | 0.000175042 | 7.506215627 | 4 mo | HOM |
| **1013288951** | 2 | SSp-ll2/3 | 113 | 0.666114624 | 0.000109592 | 4 | 0.000164524 | 6.004972502 | 4 mo | HOM |
| **1013978665** | 1 | SOCl | 114 | 0.347694749 | 0 | 0 | 0 | 0 | 2 mo | HOM |
| **1013445609** | 1 | SOCl | 114 | 0.347694749 | 0 | 0 | 0 | 0 | 4 mo | HOM |
| **1013609015** | 1 | SOCl | 114 | 0.347694749 | 0 | 0 | 0 | 0 | 4 mo | HOM |
| **1013608106** | 2 | SOCl | 114 | 0.347694749 | 0 | 0 | 0 | 0 | 4 mo | HOM |

| | | | | | | | | | | |
|---|---|---|---|---|---|---|---|---|---|---|
| **1011937543** | 1 | SOCI | 114 | 0.347694749 | 0.000375573 | 1 | 0.001080181 | 2.876086002 | 8 mo | HOM |
| **1010419538** | 2 | SOCI | 114 | 0.347694749 | 0 | 0 | 0 | 0 | 4 mo | WILD |
| **1010419187** | 1 | SOCI | 114 | 0.347694749 | 0 | 0 | 0 | 0 | 8 mo | HOM |
| **1013978366** | 1 | SOCI | 114 | 0.347694749 | 0 | 0 | 0 | 0 | 2 mo | HOM |
| **1014416155** | 2 | SOCI | 114 | 0.347694749 | 0 | 0 | 0 | 0 | 2 mo | HOM |
| **1013978065** | 2 | SOCI | 114 | 0.347694749 | 0 | 0 | 0 | 0 | 2 mo | HOM |
| **1009934102** | 2 | SOCI | 114 | 0.347694749 | 0 | 0 | 0 | 0 | 4 mo | WILD |
| **1009322937** | 2 | SOCI | 114 | 0.347694749 | 0 | 0 | 0 | 0 | 8 mo | WILD |
| **1011080637** | 2 | SOCI | 114 | 0.347694749 | 0 | 0 | 0 | 0 | 8 mo | HOM |
| **1013609713** | 2 | SOCI | 114 | 0.347694749 | 0 | 0 | 0 | 0 | 2 mo | WILD |
| **1010418842** | 1 | SOCI | 114 | 0.347694749 | 6E-06 | 0 | 1.73E-05 | 0 | 8 mo | HOM |
| **1013288213** | 2 | SOCI | 114 | 0.347694749 | 0 | 0 | 0 | 0 | 4 mo | HOM |
| **1008976060** | 2 | SOCI | 114 | 0.347694749 | 0 | 0 | 0 | 0 | 8 mo | WILD |
| **1013446243** | 1 | SOCI | 114 | 0.347694749 | 0 | 0 | 0 | 0 | 4 mo | HOM |
| **1013288951** | 2 | SOCI | 114 | 0.347694749 | 0 | 0 | 0 | 0 | 4 mo | HOM |
| **1013978665** | 1 | IV | 115 | 0.007477373 | 0 | 0 | 0 | 0 | 2 mo | HOM |
| **1013445609** | 1 | IV | 115 | 0.007477373 | 0 | 0 | 0 | 0 | 4 mo | HOM |
| **1013609015** | 1 | IV | 115 | 0.007477373 | 0 | 0 | 0 | 0 | 4 mo | HOM |
| **1013608106** | 2 | IV | 115 | 0.007477373 | 0 | 0 | 0 | 0 | 4 mo | HOM |
| **1011937543** | 1 | IV | 115 | 0.007477373 | 0 | 0 | 0 | 0 | 8 mo | HOM |
| **1010419538** | 2 | IV | 115 | 0.007477373 | 0 | 0 | 0 | 0 | 4 mo | WILD |
| **1010419187** | 1 | IV | 115 | 0.007477373 | 0 | 0 | 0 | 0 | 8 mo | HOM |
| **1013978366** | 1 | IV | 115 | 0.007477373 | 0 | 0 | 0 | 0 | 2 mo | HOM |
| **1014416155** | 2 | IV | 115 | 0.007477373 | 0 | 0 | 0 | 0 | 2 mo | HOM |
| **1013978065** | 2 | IV | 115 | 0.007477373 | 0 | 0 | 0 | 0 | 2 mo | HOM |
| **1009934102** | 2 | IV | 115 | 0.007477373 | 0 | 0 | 0 | 0 | 4 mo | WILD |
| **1009322937** | 2 | IV | 115 | 0.007477373 | 0 | 0 | 0 | 0 | 8 mo | WILD |
| **1011080637** | 2 | IV | 115 | 0.007477373 | 0 | 0 | 0 | 0 | 8 mo | HOM |
| **1013609713** | 2 | IV | 115 | 0.007477373 | 0 | 0 | 0 | 0 | 2 mo | WILD |
| **1010418842** | 1 | IV | 115 | 0.007477373 | 0 | 0 | 0 | 0 | 8 mo | HOM |
| **1013288213** | 2 | IV | 115 | 0.007477373 | 0 | 0 | 0 | 0 | 4 mo | HOM |
| **1008976060** | 2 | IV | 115 | 0.007477373 | 0 | 0 | 0 | 0 | 8 mo | WILD |
| **1013446243** | 1 | IV | 115 | 0.007477373 | 0 | 0 | 0 | 0 | 4 mo | HOM |
| **1013288951** | 2 | IV | 115 | 0.007477373 | 0 | 0 | 0 | 0 | 4 mo | HOM |
| **1013978665** | 1 | och | 117 | 0.320474151 | 0 | 0 | 0 | 0 | 2 mo | HOM |
| **1013445609** | 1 | och | 117 | 0.320474151 | 0 | 0 | 0 | 0 | 4 mo | HOM |
| **1013609015** | 1 | och | 117 | 0.320474151 | 0 | 0 | 0 | 0 | 4 mo | HOM |
| **1013608106** | 2 | och | 117 | 0.320474151 | 0 | 0 | 0 | 0 | 4 mo | HOM |
| **1011937543** | 1 | och | 117 | 0.320474151 | 0.002031615 | 5 | 0.006339403 | 15.60188234 | 8 mo | HOM |
| **1010419538** | 2 | och | 117 | 0.320474151 | 0 | 0 | 0 | 0 | 4 mo | WILD |
| **1010419187** | 1 | och | 117 | 0.320474151 | 9.6E-05 | 2 | 0.000299681 | 6.240752936 | 8 mo | HOM |
| **1013978366** | 1 | och | 117 | 0.320474151 | 3.75E-05 | 1 | 0.000116936 | 3.120376468 | 2 mo | HOM |
| **1014416155** | 2 | och | 117 | 0.320474151 | 0 | 0 | 0 | 0 | 2 mo | HOM |
| **1013978065** | 2 | och | 117 | 0.320474151 | 0 | 0 | 0 | 0 | 2 mo | HOM |
| **1009934102** | 2 | och | 117 | 0.320474151 | 5.67E-06 | 1 | 1.77E-05 | 3.120376468 | 4 mo | WILD |
| **1009322937** | 2 | och | 117 | 0.320474151 | 0 | 0 | 0 | 0 | 8 mo | WILD |
| **1011080637** | 2 | och | 117 | 0.320474151 | 1.47E-06 | 0 | 4.58E-06 | 0 | 8 mo | HOM |
| **1013609713** | 2 | och | 117 | 0.320474151 | 0 | 0 | 0 | 0 | 2 mo | WILD |
| **1010418842** | 1 | och | 117 | 0.320474151 | 0.000140606 | 4 | 0.000438742 | 12.48150587 | 8 mo | HOM |
| **1013288213** | 2 | och | 117 | 0.320474151 | 0 | 0 | 0 | 0 | 4 mo | HOM |
| **1008976060** | 2 | och | 117 | 0.320474151 | 2.02E-05 | 1 | 6.3E-05 | 3.120376468 | 8 mo | WILD |
| **1013446243** | 1 | och | 117 | 0.320474151 | 0 | 0 | 0 | 0 | 4 mo | HOM |
| **1013288951** | 2 | och | 117 | 0.320474151 | 0 | 0 | 0 | 0 | 4 mo | HOM |
| **1013978665** | 1 | PVi | 118 | 0.211233218 | 0 | 0 | 0 | 0 | 2 mo | HOM |
| **1013445609** | 1 | PVi | 118 | 0.211233218 | 1.96E-06 | 1 | 9.3E-06 | 4.734103895 | 4 mo | HOM |
| **1013609015** | 1 | PVi | 118 | 0.211233218 | 0 | 0 | 0 | 0 | 4 mo | HOM |
| **1013608106** | 2 | PVi | 118 | 0.211233218 | 0 | 0 | 0 | 0 | 4 mo | HOM |
| **1011937543** | 1 | PVi | 118 | 0.211233218 | 2.77E-11 | 0 | 1.31E-10 | 0 | 8 mo | HOM |
| **1010419538** | 2 | PVi | 118 | 0.211233218 | 2.7E-06 | 1 | 1.28E-05 | 4.734103895 | 4 mo | WILD |

| | | | | | | | | | | |
|---|---|---|---|---|---|---|---|---|---|---|
| **1010419187** | 1 | PVi | 118 | 0.211233218 | 0 | 0 | 0 | 0 | 8 mo | HOM |
| **1013978366** | 1 | PVi | 118 | 0.211233218 | 0 | 0 | 0 | 0 | 2 mo | HOM |
| **1014416155** | 2 | PVi | 118 | 0.211233218 | 0 | 0 | 0 | 0 | 2 mo | HOM |
| **1013978065** | 2 | PVi | 118 | 0.211233218 | 0 | 0 | 0 | 0 | 2 mo | HOM |
| **1009934102** | 2 | PVi | 118 | 0.211233218 | 0 | 0 | 0 | 0 | 4 mo | WILD |
| **1009322937** | 2 | PVi | 118 | 0.211233218 | 0 | 0 | 0 | 0 | 8 mo | WILD |
| **1011080637** | 2 | PVi | 118 | 0.211233218 | 2.39E-05 | 0 | 0.000113351 | 0 | 8 mo | HOM |
| **1013609713** | 2 | PVi | 118 | 0.211233218 | 0 | 0 | 0 | 0 | 2 mo | WILD |
| **1010418842** | 1 | PVi | 118 | 0.211233218 | 0.000149048 | 9 | 0.000705607 | 42.60693505 | 8 mo | HOM |
| **1013288213** | 2 | PVi | 118 | 0.211233218 | 1.3E-07 | 0 | 6.13E-07 | 0 | 4 mo | HOM |
| **1008976060** | 2 | PVi | 118 | 0.211233218 | 0.000192232 | 10 | 0.000910047 | 47.34103895 | 8 mo | WILD |
| **1013446243** | 1 | PVi | 118 | 0.211233218 | 1.49E-06 | 1 | 7.07E-06 | 4.734103895 | 4 mo | HOM |
| **1013288951** | 2 | PVi | 118 | 0.211233218 | 0 | 0 | 0 | 0 | 4 mo | HOM |
| **1013978665** | 1 | AIv | 119 | 1.787265283 | 0 | 0 | 0 | 0 | 2 mo | HOM |
| **1013445609** | 1 | AIv | 119 | 1.787265283 | 5.82E-05 | 3 | 3.26E-05 | 1.678542088 | 4 mo | HOM |
| **1013609015** | 1 | AIv | 119 | 1.787265283 | 9.96E-05 | 7 | 5.57E-05 | 3.916598205 | 4 mo | HOM |
| **1013608106** | 2 | AIv | 119 | 1.787265283 | 2.26E-07 | 0 | 1.26E-07 | 0 | 4 mo | HOM |
| **1011937543** | 1 | AIv | 119 | 1.787265283 | 0.001771679 | 38 | 0.000991279 | 21.26153311 | 8 mo | HOM |
| **1010419538** | 2 | AIv | 119 | 1.787265283 | 0 | 0 | 0 | 0 | 4 mo | WILD |
| **1010419187** | 1 | AIv | 119 | 1.787265283 | 0.002473327 | 86 | 0.001383861 | 48.11820652 | 8 mo | HOM |
| **1013978366** | 1 | AIv | 119 | 1.787265283 | 5.71E-06 | 1 | 3.19E-06 | 0.559514029 | 2 mo | HOM |
| **1014416155** | 2 | AIv | 119 | 1.787265283 | 0 | 0 | 0 | 0 | 2 mo | HOM |
| **1013978065** | 2 | AIv | 119 | 1.787265283 | 0 | 0 | 0 | 0 | 2 mo | HOM |
| **1009934102** | 2 | AIv | 119 | 1.787265283 | 0 | 0 | 0 | 0 | 4 mo | WILD |
| **1009322937** | 2 | AIv | 119 | 1.787265283 | 6.85E-05 | 5 | 3.83E-05 | 2.797570146 | 8 mo | WILD |
| **1011080637** | 2 | AIv | 119 | 1.787265283 | 0.001003034 | 53 | 0.000561212 | 29.65424355 | 8 mo | HOM |
| **1013609713** | 2 | AIv | 119 | 1.787265283 | 0 | 0 | 0 | 0 | 2 mo | WILD |
| **1010418842** | 1 | AIv | 119 | 1.787265283 | 0.001413552 | 62 | 0.000790902 | 34.68986981 | 8 mo | HOM |
| **1013288213** | 2 | AIv | 119 | 1.787265283 | 9.38E-05 | 3 | 5.25E-05 | 1.678542088 | 4 mo | HOM |
| **1008976060** | 2 | AIv | 119 | 1.787265283 | 0.000145476 | 6 | 8.14E-05 | 3.357084176 | 8 mo | WILD |
| **1013446243** | 1 | AIv | 119 | 1.787265283 | 9.91E-05 | 11 | 5.55E-05 | 6.154654322 | 4 mo | HOM |
| **1013288951** | 2 | AIv | 119 | 1.787265283 | 0.000199929 | 4 | 0.000111863 | 2.238056117 | 4 mo | HOM |
| **1013978665** | 1 | AIp1 | 120 | 0.434794091 | 0 | 0 | 0 | 0 | 2 mo | HOM |
| **1013445609** | 1 | AIp1 | 120 | 0.434794091 | 5.56E-05 | 3 | 0.000127988 | 6.899817777 | 4 mo | HOM |
| **1013609015** | 1 | AIp1 | 120 | 0.434794091 | 0.000122553 | 5 | 0.000281864 | 11.4996963 | 4 mo | HOM |
| **1013608106** | 2 | AIp1 | 120 | 0.434794091 | 0 | 0 | 0 | 0 | 4 mo | HOM |
| **1011937543** | 1 | AIp1 | 120 | 0.434794091 | 0.000470949 | 13 | 0.001083154 | 29.89921037 | 8 mo | HOM |
| **1010419538** | 2 | AIp1 | 120 | 0.434794091 | 3.9E-09 | 0 | 8.97E-09 | 0 | 4 mo | WILD |
| **1010419187** | 1 | AIp1 | 120 | 0.434794091 | 0.000852147 | 35 | 0.001959887 | 80.49787407 | 8 mo | HOM |
| **1013978366** | 1 | AIp1 | 120 | 0.434794091 | 0 | 0 | 0 | 0 | 2 mo | HOM |
| **1014416155** | 2 | AIp1 | 120 | 0.434794091 | 0 | 0 | 0 | 0 | 2 mo | HOM |
| **1013978065** | 2 | AIp1 | 120 | 0.434794091 | 0 | 0 | 0 | 0 | 2 mo | HOM |
| **1009934102** | 2 | AIp1 | 120 | 0.434794091 | 0 | 0 | 0 | 0 | 4 mo | WILD |
| **1009322937** | 2 | AIp1 | 120 | 0.434794091 | 0 | 0 | 0 | 0 | 8 mo | WILD |
| **1011080637** | 2 | AIp1 | 120 | 0.434794091 | 0.00035499 | 17 | 0.000816457 | 39.09896741 | 8 mo | HOM |
| **1013609713** | 2 | AIp1 | 120 | 0.434794091 | 0 | 0 | 0 | 0 | 2 mo | WILD |
| **1010418842** | 1 | AIp1 | 120 | 0.434794091 | 0.000644839 | 30 | 0.001483091 | 68.99817777 | 8 mo | HOM |
| **1013288213** | 2 | AIp1 | 120 | 0.434794091 | 2.19E-05 | 3 | 5.03E-05 | 6.899817777 | 4 mo | HOM |
| **1008976060** | 2 | AIp1 | 120 | 0.434794091 | 1.93E-07 | 0 | 4.44E-07 | 0 | 8 mo | WILD |
| **1013446243** | 1 | AIp1 | 120 | 0.434794091 | 0.000103137 | 9 | 0.000237208 | 20.69945333 | 4 mo | HOM |
| **1013288951** | 2 | AIp1 | 120 | 0.434794091 | 0.000182488 | 6 | 0.000419712 | 13.79963555 | 4 mo | HOM |
| **1013978665** | 1 | VISl6b | 121 | 0.039395804 | 0 | 0 | 0 | 0 | 2 mo | HOM |
| **1013445609** | 1 | VISl6b | 121 | 0.039395804 | 0 | 0 | 0 | 0 | 4 mo | HOM |
| **1013609015** | 1 | VISl6b | 121 | 0.039395804 | 2.85E-05 | 2 | 0.000723363 | 50.76682786 | 4 mo | HOM |
| **1013608106** | 2 | VISl6b | 121 | 0.039395804 | 0 | 0 | 0 | 0 | 4 mo | HOM |
| **1011937543** | 1 | VISl6b | 121 | 0.039395804 | 2.94E-05 | 1 | 0.000746629 | 25.38341393 | 8 mo | HOM |
| **1010419538** | 2 | VISl6b | 121 | 0.039395804 | 0 | 0 | 0 | 0 | 4 mo | WILD |
| **1010419187** | 1 | VISl6b | 121 | 0.039395804 | 5.52E-05 | 3 | 0.001401618 | 76.15024179 | 8 mo | HOM |
| **1013978366** | 1 | VISl6b | 121 | 0.039395804 | 0 | 0 | 0 | 0 | 2 mo | HOM |

| | | | | | | | | | | |
|---|---|---|---|---|---|---|---|---|---|---|
| 1014416155 | 2 | VISl6b | 121 | 0.039395804 | 0 | 0 | 0 | 0 | 2 mo | HOM |
| 1013978065 | 2 | VISl6b | 121 | 0.039395804 | 0 | 0 | 0 | 0 | 2 mo | HOM |
| 1009934102 | 2 | VISl6b | 121 | 0.039395804 | 0 | 0 | 0 | 0 | 4 mo | WILD |
| 1009322937 | 2 | VISl6b | 121 | 0.039395804 | 0 | 0 | 0 | 0 | 8 mo | WILD |
| 1011080637 | 2 | VISl6b | 121 | 0.039395804 | 1.27E-05 | 0 | 0.000321461 | 0 | 8 mo | HOM |
| 1013609713 | 2 | VISl6b | 121 | 0.039395804 | 0 | 0 | 0 | 0 | 2 mo | WILD |
| 1010418842 | 1 | VISl6b | 121 | 0.039395804 | 3.63E-05 | 3 | 0.000922388 | 76.15024179 | 8 mo | HOM |
| 1013288213 | 2 | VISl6b | 121 | 0.039395804 | 6.97E-07 | 0 | 1.77E-05 | 0 | 4 mo | HOM |
| 1008976060 | 2 | VISl6b | 121 | 0.039395804 | 0 | 0 | 0 | 0 | 8 mo | WILD |
| 1013446243 | 1 | VISl6b | 121 | 0.039395804 | 9.73E-09 | 0 | 2.47E-07 | 0 | 4 mo | HOM |
| 1013288951 | 2 | VISl6b | 121 | 0.039395804 | 8.97E-09 | 0 | 2.28E-07 | 0 | 4 mo | HOM |
| 1013978665 | 1 | POR | 122 | 0.376994358 | 0 | 0 | 0 | 0 | 2 mo | HOM |
| 1013445609 | 1 | POR | 122 | 0.376994358 | 0 | 0 | 0 | 0 | 4 mo | HOM |
| 1013609015 | 1 | POR | 122 | 0.376994358 | 0 | 0 | 0 | 0 | 4 mo | HOM |
| 1013608106 | 2 | POR | 122 | 0.376994358 | 0 | 0 | 0 | 0 | 4 mo | HOM |
| 1011937543 | 1 | POR | 122 | 0.376994358 | 5.82E-06 | 0 | 1.54E-05 | 0 | 8 mo | HOM |
| 1010419538 | 2 | POR | 122 | 0.376994358 | 0 | 0 | 0 | 0 | 4 mo | WILD |
| 1010419187 | 1 | POR | 122 | 0.376994358 | 0 | 0 | 0 | 0 | 8 mo | HOM |
| 1013978366 | 1 | POR | 122 | 0.376994358 | 0 | 0 | 0 | 0 | 2 mo | HOM |
| 1014416155 | 2 | POR | 122 | 0.376994358 | 0 | 0 | 0 | 0 | 2 mo | HOM |
| 1013978065 | 2 | POR | 122 | 0.376994358 | 0 | 0 | 0 | 0 | 2 mo | HOM |
| 1009934102 | 2 | POR | 122 | 0.376994358 | 0 | 0 | 0 | 0 | 4 mo | WILD |
| 1009322937 | 2 | POR | 122 | 0.376994358 | 0 | 0 | 0 | 0 | 8 mo | WILD |
| 1011080637 | 2 | POR | 122 | 0.376994358 | 0 | 0 | 0 | 0 | 8 mo | HOM |
| 1013609713 | 2 | POR | 122 | 0.376994358 | 0 | 0 | 0 | 0 | 2 mo | WILD |
| 1010418842 | 1 | POR | 122 | 0.376994358 | 1.62E-05 | 1 | 4.3E-05 | 2.652559594 | 8 mo | HOM |
| 1013288213 | 2 | POR | 122 | 0.376994358 | 0 | 0 | 0 | 0 | 4 mo | HOM |
| 1008976060 | 2 | POR | 122 | 0.376994358 | 9.92E-06 | 2 | 2.63E-05 | 5.305119187 | 8 mo | WILD |
| 1013446243 | 1 | POR | 122 | 0.376994358 | 0 | 0 | 0 | 0 | 4 mo | HOM |
| 1013288951 | 2 | POR | 122 | 0.376994358 | 0 | 0 | 0 | 0 | 4 mo | HOM |
| 1013978665 | 1 | KF | 123 | 0.180501594 | 0 | 0 | 0 | 0 | 2 mo | HOM |
| 1013445609 | 1 | KF | 123 | 0.180501594 | 0 | 0 | 0 | 0 | 4 mo | HOM |
| 1013609015 | 1 | KF | 123 | 0.180501594 | 0 | 0 | 0 | 0 | 4 mo | HOM |
| 1013608106 | 2 | KF | 123 | 0.180501594 | 0 | 0 | 0 | 0 | 4 mo | HOM |
| 1011937543 | 1 | KF | 123 | 0.180501594 | 4.77E-05 | 1 | 0.000264409 | 5.54011728 | 8 mo | HOM |
| 1010419538 | 2 | KF | 123 | 0.180501594 | 0 | 0 | 0 | 0 | 4 mo | WILD |
| 1010419187 | 1 | KF | 123 | 0.180501594 | 6.14E-06 | 1 | 3.4E-05 | 5.54011728 | 8 mo | HOM |
| 1013978366 | 1 | KF | 123 | 0.180501594 | 0 | 0 | 0 | 0 | 2 mo | HOM |
| 1014416155 | 2 | KF | 123 | 0.180501594 | 0 | 0 | 0 | 0 | 2 mo | HOM |
| 1013978065 | 2 | KF | 123 | 0.180501594 | 0 | 0 | 0 | 0 | 2 mo | HOM |
| 1009934102 | 2 | KF | 123 | 0.180501594 | 0 | 0 | 0 | 0 | 4 mo | WILD |
| 1009322937 | 2 | KF | 123 | 0.180501594 | 0 | 0 | 0 | 0 | 8 mo | WILD |
| 1011080637 | 2 | KF | 123 | 0.180501594 | 4.83E-06 | 1 | 2.67E-05 | 5.54011728 | 8 mo | HOM |
| 1013609713 | 2 | KF | 123 | 0.180501594 | 0 | 0 | 0 | 0 | 2 mo | WILD |
| 1010418842 | 1 | KF | 123 | 0.180501594 | 0 | 0 | 0 | 0 | 8 mo | HOM |
| 1013288213 | 2 | KF | 123 | 0.180501594 | 0 | 0 | 0 | 0 | 4 mo | HOM |
| 1008976060 | 2 | KF | 123 | 0.180501594 | 1.59E-05 | 1 | 8.81E-05 | 5.54011728 | 8 mo | WILD |
| 1013446243 | 1 | KF | 123 | 0.180501594 | 0 | 0 | 0 | 0 | 4 mo | HOM |
| 1013288951 | 2 | KF | 123 | 0.180501594 | 0 | 0 | 0 | 0 | 4 mo | HOM |
| 1013978665 | 1 | opt | 125 | 0.718404758 | 0 | 0 | 0 | 0 | 2 mo | HOM |
| 1013445609 | 1 | opt | 125 | 0.718404758 | 0 | 0 | 0 | 0 | 4 mo | HOM |
| 1013609015 | 1 | opt | 125 | 0.718404758 | 0 | 0 | 0 | 0 | 4 mo | HOM |
| 1013608106 | 2 | opt | 125 | 0.718404758 | 0 | 0 | 0 | 0 | 4 mo | HOM |
| 1011937543 | 1 | opt | 125 | 0.718404758 | 0.001544581 | 7 | 0.002150015 | 9.743810739 | 8 mo | HOM |
| 1010419538 | 2 | opt | 125 | 0.718404758 | 0 | 0 | 0 | 0 | 4 mo | WILD |
| 1010419187 | 1 | opt | 125 | 0.718404758 | 2.74E-05 | 3 | 3.81E-05 | 4.175918888 | 8 mo | HOM |
| 1013978366 | 1 | opt | 125 | 0.718404758 | 0 | 0 | 0 | 0 | 2 mo | HOM |
| 1014416155 | 2 | opt | 125 | 0.718404758 | 0 | 0 | 0 | 0 | 2 mo | HOM |
| 1013978065 | 2 | opt | 125 | 0.718404758 | 0 | 0 | 0 | 0 | 2 mo | HOM |

| | | | | | | | | | | |
|---|---|---|---|---|---|---|---|---|---|---|
| 1009934102 | 2 | opt | 125 | 0.718404758 | 0 | 0 | 0 | 0 | 4 mo | WILD |
| 1009322937 | 2 | opt | 125 | 0.718404758 | 0 | 0 | 0 | 0 | 8 mo | WILD |
| 1011080637 | 2 | opt | 125 | 0.718404758 | 2.22E-05 | 1 | 3.08E-05 | 1.391972963 | 8 mo | HOM |
| 1013609713 | 2 | opt | 125 | 0.718404758 | 0 | 0 | 0 | 0 | 2 mo | WILD |
| 1010418842 | 1 | opt | 125 | 0.718404758 | 1.99E-05 | 0 | 2.77E-05 | 0 | 8 mo | HOM |
| 1013288213 | 2 | opt | 125 | 0.718404758 | 0 | 0 | 0 | 0 | 4 mo | HOM |
| 1008976060 | 2 | opt | 125 | 0.718404758 | 5.32E-08 | 0 | 7.4E-08 | 0 | 8 mo | WILD |
| 1013446243 | 1 | opt | 125 | 0.718404758 | 7.9E-22 | 0 | 1.1E-21 | 0 | 4 mo | HOM |
| 1013288951 | 2 | opt | 125 | 0.718404758 | 0 | 0 | 0 | 0 | 4 mo | HOM |
| 1013978665 | 1 | PVp | 126 | 0.128322768 | 0 | 0 | 0 | 0 | 2 mo | HOM |
| 1013445609 | 1 | PVp | 126 | 0.128322768 | 0 | 0 | 0 | 0 | 4 mo | HOM |
| 1013609015 | 1 | PVp | 126 | 0.128322768 | 3.63E-05 | 3 | 0.000282973 | 23.37854807 | 4 mo | HOM |
| 1013608106 | 2 | PVp | 126 | 0.128322768 | 0 | 0 | 0 | 0 | 4 mo | HOM |
| 1011937543 | 1 | PVp | 126 | 0.128322768 | 0.000378429 | 1 | 0.00294904 | 7.792849356 | 8 mo | HOM |
| 1010419538 | 2 | PVp | 126 | 0.128322768 | 3.69E-08 | 0 | 2.87E-07 | 0 | 4 mo | WILD |
| 1010419187 | 1 | PVp | 126 | 0.128322768 | 0.000776077 | 1 | 0.00604785 | 7.792849356 | 8 mo | HOM |
| 1013978366 | 1 | PVp | 126 | 0.128322768 | 0 | 0 | 0 | 0 | 2 mo | HOM |
| 1014416155 | 2 | PVp | 126 | 0.128322768 | 0 | 0 | 0 | 0 | 2 mo | HOM |
| 1013978065 | 2 | PVp | 126 | 0.128322768 | 0 | 0 | 0 | 0 | 2 mo | HOM |
| 1009934102 | 2 | PVp | 126 | 0.128322768 | 5.52E-06 | 0 | 4.3E-05 | 0 | 4 mo | WILD |
| 1009322937 | 2 | PVp | 126 | 0.128322768 | 2.9E-06 | 1 | 2.26E-05 | 7.792849356 | 8 mo | WILD |
| 1011080637 | 2 | PVp | 126 | 0.128322768 | 0 | 0 | 0 | 0 | 8 mo | HOM |
| 1013609713 | 2 | PVp | 126 | 0.128322768 | 0 | 0 | 0 | 0 | 2 mo | WILD |
| 1010418842 | 1 | PVp | 126 | 0.128322768 | 0 | 0 | 0 | 0 | 8 mo | HOM |
| 1013288213 | 2 | PVp | 126 | 0.128322768 | 0 | 0 | 0 | 0 | 4 mo | HOM |
| 1008976060 | 2 | PVp | 126 | 0.128322768 | 1.65E-07 | 0 | 1.29E-06 | 0 | 8 mo | WILD |
| 1013446243 | 1 | PVp | 126 | 0.128322768 | 2.53E-06 | 1 | 1.97E-05 | 7.792849356 | 4 mo | HOM |
| 1013288951 | 2 | PVp | 126 | 0.128322768 | 0 | 0 | 0 | 0 | 4 mo | HOM |
| 1013978665 | 1 | AM | 127 | 0.423033052 | 0 | 0 | 0 | 0 | 2 mo | HOM |
| 1013445609 | 1 | AM | 127 | 0.423033052 | 0 | 0 | 0 | 0 | 4 mo | HOM |
| 1013609015 | 1 | AM | 127 | 0.423033052 | 0 | 0 | 0 | 0 | 4 mo | HOM |
| 1013608106 | 2 | AM | 127 | 0.423033052 | 0 | 0 | 0 | 0 | 4 mo | HOM |
| 1011937543 | 1 | AM | 127 | 0.423033052 | 0.000165286 | 5 | 0.000390716 | 11.81940742 | 8 mo | HOM |
| 1010419538 | 2 | AM | 127 | 0.423033052 | 0 | 0 | 0 | 0 | 4 mo | WILD |
| 1010419187 | 1 | AM | 127 | 0.423033052 | 0.000245201 | 9 | 0.000579626 | 21.27493336 | 8 mo | HOM |
| 1013978366 | 1 | AM | 127 | 0.423033052 | 0 | 0 | 0 | 0 | 2 mo | HOM |
| 1014416155 | 2 | AM | 127 | 0.423033052 | 0 | 0 | 0 | 0 | 2 mo | HOM |
| 1013978065 | 2 | AM | 127 | 0.423033052 | 0 | 0 | 0 | 0 | 2 mo | HOM |
| 1009934102 | 2 | AM | 127 | 0.423033052 | 0 | 0 | 0 | 0 | 4 mo | WILD |
| 1009322937 | 2 | AM | 127 | 0.423033052 | 1.07E-05 | 1 | 2.52E-05 | 2.363881485 | 8 mo | WILD |
| 1011080637 | 2 | AM | 127 | 0.423033052 | 0.000378054 | 8 | 0.000893675 | 18.91105188 | 8 mo | HOM |
| 1013609713 | 2 | AM | 127 | 0.423033052 | 0 | 0 | 0 | 0 | 2 mo | WILD |
| 1010418842 | 1 | AM | 127 | 0.423033052 | 0.00019595 | 10 | 0.000463202 | 23.63881485 | 8 mo | HOM |
| 1013288213 | 2 | AM | 127 | 0.423033052 | 7.49E-06 | 1 | 1.77E-05 | 2.363881485 | 4 mo | HOM |
| 1008976060 | 2 | AM | 127 | 0.423033052 | 4.72E-06 | 1 | 1.11E-05 | 2.363881485 | 8 mo | WILD |
| 1013446243 | 1 | AM | 127 | 0.423033052 | 7.46E-05 | 1 | 0.000176249 | 2.363881485 | 4 mo | HOM |
| 1013288951 | 2 | AM | 127 | 0.423033052 | 0 | 0 | 0 | 0 | 4 mo | HOM |
| 1013978665 | 1 | MRN | 128 | 5.29871104 | 4.84E-09 | 0 | 9.13E-10 | 0 | 2 mo | HOM |
| 1013445609 | 1 | MRN | 128 | 5.29871104 | 0 | 0 | 0 | 0 | 4 mo | HOM |
| 1013609015 | 1 | MRN | 128 | 5.29871104 | 0 | 0 | 0 | 0 | 4 mo | HOM |
| 1013608106 | 2 | MRN | 128 | 5.29871104 | 0 | 0 | 0 | 0 | 4 mo | HOM |
| 1011937543 | 1 | MRN | 128 | 5.29871104 | 0.017345477 | 82 | 0.003273528 | 15.47546175 | 8 mo | HOM |
| 1010419538 | 2 | MRN | 128 | 5.29871104 | 0 | 0 | 0 | 0 | 4 mo | WILD |
| 1010419187 | 1 | MRN | 128 | 5.29871104 | 4.2E-05 | 3 | 7.94E-06 | 0.56617543 | 8 mo | HOM |
| 1013978366 | 1 | MRN | 128 | 5.29871104 | 0 | 0 | 0 | 0 | 2 mo | HOM |
| 1014416155 | 2 | MRN | 128 | 5.29871104 | 0 | 0 | 0 | 0 | 2 mo | HOM |
| 1013978065 | 2 | MRN | 128 | 5.29871104 | 0 | 0 | 0 | 0 | 2 mo | HOM |
| 1009934102 | 2 | MRN | 128 | 5.29871104 | 0 | 0 | 0 | 0 | 4 mo | WILD |
| 1009322937 | 2 | MRN | 128 | 5.29871104 | 1.53E-10 | 0 | 2.89E-11 | 0 | 8 mo | WILD |

| | | | | | | | | | | |
|---|---|---|---|---|---|---|---|---|---|---|
| **1011080637** | 2 | MRN | 128 | 5.29871104 | 0.000129417 | 7 | 2.44E-05 | 1.321076003 | 8 mo | HOM |
| **1013609713** | 2 | MRN | 128 | 5.29871104 | 0 | 0 | 0 | 0 | 2 mo | WILD |
| **1010418842** | 1 | MRN | 128 | 5.29871104 | 0.000539394 | 17 | 0.000101797 | 3.208327435 | 8 mo | HOM |
| **1013288213** | 2 | MRN | 128 | 5.29871104 | 0 | 0 | 0 | 0 | 4 mo | HOM |
| **1008976060** | 2 | MRN | 128 | 5.29871104 | 0.000125816 | 12 | 2.37E-05 | 2.264701719 | 8 mo | WILD |
| **1013446243** | 1 | MRN | 128 | 5.29871104 | 0 | 0 | 0 | 0 | 4 mo | HOM |
| **1013288951** | 2 | MRN | 128 | 5.29871104 | 8.45E-06 | 1 | 1.59E-06 | 0.188725143 | 4 mo | HOM |
| **1013978665** | 1 | V3 | 129 | 0.918124133 | 0.000860475 | 15 | 0.00093721 | 16.33766009 | 2 mo | HOM |
| **1013445609** | 1 | V3 | 129 | 0.918124133 | 0.00062118 | 14 | 0.000676575 | 15.24848275 | 4 mo | HOM |
| **1013609015** | 1 | V3 | 129 | 0.918124133 | 0.003422693 | 11 | 0.003727919 | 11.98095073 | 4 mo | HOM |
| **1013608106** | 2 | V3 | 129 | 0.918124133 | 0 | 0 | 0 | 0 | 4 mo | HOM |
| **1011937543** | 1 | V3 | 129 | 0.918124133 | 0.000792455 | 10 | 0.000863124 | 10.89177339 | 8 mo | HOM |
| **1010419538** | 2 | V3 | 129 | 0.918124133 | 7.74E-05 | 7 | 8.43E-05 | 7.624241374 | 4 mo | WILD |
| **1010419187** | 1 | V3 | 129 | 0.918124133 | 0.005142682 | 18 | 0.005601293 | 19.60519211 | 8 mo | HOM |
| **1013978366** | 1 | V3 | 129 | 0.918124133 | 0.000503468 | 7 | 0.000548366 | 7.624241374 | 2 mo | HOM |
| **1014416155** | 2 | V3 | 129 | 0.918124133 | 0 | 0 | 0 | 0 | 2 mo | HOM |
| **1013978065** | 2 | V3 | 129 | 0.918124133 | 1.27E-05 | 2 | 1.39E-05 | 2.178354678 | 2 mo | HOM |
| **1009934102** | 2 | V3 | 129 | 0.918124133 | 7.72E-05 | 4 | 8.41E-05 | 4.356709357 | 4 mo | WILD |
| **1009322937** | 2 | V3 | 129 | 0.918124133 | 0.000318278 | 13 | 0.000346662 | 14.15930541 | 8 mo | WILD |
| **1011080637** | 2 | V3 | 129 | 0.918124133 | 0.00692635 | 18 | 0.007544023 | 19.60519211 | 8 mo | HOM |
| **1013609713** | 2 | V3 | 129 | 0.918124133 | 6.51E-05 | 3 | 7.09E-05 | 3.267532018 | 2 mo | WILD |
| **1010418842** | 1 | V3 | 129 | 0.918124133 | 0.009665919 | 21 | 0.0105279 | 22.87272412 | 8 mo | HOM |
| **1013288213** | 2 | V3 | 129 | 0.918124133 | 0.000225768 | 17 | 0.000245901 | 18.51601477 | 4 mo | HOM |
| **1008976060** | 2 | V3 | 129 | 0.918124133 | 0.005031314 | 20 | 0.005479993 | 21.78354678 | 8 mo | WILD |
| **1013446243** | 1 | V3 | 129 | 0.918124133 | 0.002748802 | 13 | 0.002993933 | 14.15930541 | 4 mo | HOM |
| **1013288951** | 2 | V3 | 129 | 0.918124133 | 0.005052254 | 8 | 0.0055028 | 8.713418713 | 4 mo | HOM |
| **1013978665** | 1 | LA | 131 | 0.865685744 | 0 | 0 | 0 | 0 | 2 mo | HOM |
| **1013445609** | 1 | LA | 131 | 0.865685744 | 0.000151375 | 10 | 0.000174862 | 11.55153596 | 4 mo | HOM |
| **1013609015** | 1 | LA | 131 | 0.865685744 | 0.000328795 | 12 | 0.000379809 | 13.86184315 | 4 mo | HOM |
| **1013608106** | 2 | LA | 131 | 0.865685744 | 0 | 0 | 0 | 0 | 4 mo | HOM |
| **1011937543** | 1 | LA | 131 | 0.865685744 | 0.001303416 | 30 | 0.001505646 | 34.65460787 | 8 mo | HOM |
| **1010419538** | 2 | LA | 131 | 0.865685744 | 0 | 0 | 0 | 0 | 4 mo | WILD |
| **1010419187** | 1 | LA | 131 | 0.865685744 | 0.001943734 | 57 | 0.002245311 | 65.84375496 | 8 mo | HOM |
| **1013978366** | 1 | LA | 131 | 0.865685744 | 0 | 0 | 0 | 0 | 2 mo | HOM |
| **1014416155** | 2 | LA | 131 | 0.865685744 | 0 | 0 | 0 | 0 | 2 mo | HOM |
| **1013978065** | 2 | LA | 131 | 0.865685744 | 0 | 0 | 0 | 0 | 2 mo | HOM |
| **1009934102** | 2 | LA | 131 | 0.865685744 | 0 | 0 | 0 | 0 | 4 mo | WILD |
| **1009322937** | 2 | LA | 131 | 0.865685744 | 0 | 0 | 0 | 0 | 8 mo | WILD |
| **1011080637** | 2 | LA | 131 | 0.865685744 | 0.001343048 | 46 | 0.001551426 | 53.13706541 | 8 mo | HOM |
| **1013609713** | 2 | LA | 131 | 0.865685744 | 0 | 0 | 0 | 0 | 2 mo | WILD |
| **1010418842** | 1 | LA | 131 | 0.865685744 | 0.001924895 | 42 | 0.00222355 | 48.51645102 | 8 mo | HOM |
| **1013288213** | 2 | LA | 131 | 0.865685744 | 2.8E-05 | 3 | 3.23E-05 | 3.465460787 | 4 mo | HOM |
| **1008976060** | 2 | LA | 131 | 0.865685744 | 3.84E-07 | 0 | 4.44E-07 | 0 | 8 mo | WILD |
| **1013446243** | 1 | LA | 131 | 0.865685744 | 0.000131677 | 8 | 0.000152107 | 9.241228766 | 4 mo | HOM |
| **1013288951** | 2 | LA | 131 | 0.865685744 | 0.000287414 | 10 | 0.000332007 | 11.55153596 | 4 mo | HOM |
| **1013978665** | 1 | PL6b | 132 | 0.022013848 | 0 | 0 | 0 | 0 | 2 mo | HOM |
| **1013445609** | 1 | PL6b | 132 | 0.022013848 | 0 | 0 | 0 | 0 | 4 mo | HOM |
| **1013609015** | 1 | PL6b | 132 | 0.022013848 | 0 | 0 | 0 | 0 | 4 mo | HOM |
| **1013608106** | 2 | PL6b | 132 | 0.022013848 | 0 | 0 | 0 | 0 | 4 mo | HOM |
| **1011937543** | 1 | PL6b | 132 | 0.022013848 | 0 | 0 | 0 | 0 | 8 mo | HOM |
| **1010419538** | 2 | PL6b | 132 | 0.022013848 | 0 | 0 | 0 | 0 | 4 mo | WILD |
| **1010419187** | 1 | PL6b | 132 | 0.022013848 | 1.95E-05 | 1 | 0.000885767 | 45.4259523 | 8 mo | HOM |
| **1013978366** | 1 | PL6b | 132 | 0.022013848 | 0 | 0 | 0 | 0 | 2 mo | HOM |
| **1014416155** | 2 | PL6b | 132 | 0.022013848 | 0 | 0 | 0 | 0 | 2 mo | HOM |
| **1013978065** | 2 | PL6b | 132 | 0.022013848 | 0 | 0 | 0 | 0 | 2 mo | HOM |
| **1009934102** | 2 | PL6b | 132 | 0.022013848 | 0 | 0 | 0 | 0 | 4 mo | WILD |
| **1009322937** | 2 | PL6b | 132 | 0.022013848 | 0 | 0 | 0 | 0 | 8 mo | WILD |
| **1011080637** | 2 | PL6b | 132 | 0.022013848 | 0 | 0 | 0 | 0 | 8 mo | HOM |
| **1013609713** | 2 | PL6b | 132 | 0.022013848 | 0 | 0 | 0 | 0 | 2 mo | WILD |

| | | | | | | | | | | |
|---|---|---|---|---|---|---|---|---|---|---|
| 1010418842 | 1 | PL6b | 132 | 0.022013848 | 0 | 0 | 0 | 0 | 8 mo | HOM |
| 1013288213 | 2 | PL6b | 132 | 0.022013848 | 0 | 0 | 0 | 0 | 4 mo | HOM |
| 1008976060 | 2 | PL6b | 132 | 0.022013848 | 0 | 0 | 0 | 0 | 8 mo | WILD |
| 1013446243 | 1 | PL6b | 132 | 0.022013848 | 0 | 0 | 0 | 0 | 4 mo | HOM |
| 1013288951 | 2 | PL6b | 132 | 0.022013848 | 0 | 0 | 0 | 0 | 4 mo | HOM |
| 1013978665 | 1 | PVpo | 133 | 0.152110661 | 0 | 0 | 0 | 0 | 2 mo | HOM |
| 1013445609 | 1 | PVpo | 133 | 0.152110661 | 0 | 0 | 0 | 0 | 4 mo | HOM |
| 1013609015 | 1 | PVpo | 133 | 0.152110661 | 9.51E-23 | 0 | 6.25E-22 | 0 | 4 mo | HOM |
| 1013608106 | 2 | PVpo | 133 | 0.152110661 | 0 | 0 | 0 | 0 | 4 mo | HOM |
| 1011937543 | 1 | PVpo | 133 | 0.152110661 | 2.45E-06 | 0 | 1.61E-05 | 0 | 8 mo | HOM |
| 1010419538 | 2 | PVpo | 133 | 0.152110661 | 0 | 0 | 0 | 0 | 4 mo | WILD |
| 1010419187 | 1 | PVpo | 133 | 0.152110661 | 0 | 0 | 0 | 0 | 8 mo | HOM |
| 1013978366 | 1 | PVpo | 133 | 0.152110661 | 0 | 0 | 0 | 0 | 2 mo | HOM |
| 1014416155 | 2 | PVpo | 133 | 0.152110661 | 0 | 0 | 0 | 0 | 2 mo | HOM |
| 1013978065 | 2 | PVpo | 133 | 0.152110661 | 0 | 0 | 0 | 0 | 2 mo | HOM |
| 1009934102 | 2 | PVpo | 133 | 0.152110661 | 0 | 0 | 0 | 0 | 4 mo | WILD |
| 1009322937 | 2 | PVpo | 133 | 0.152110661 | 0 | 0 | 0 | 0 | 8 mo | WILD |
| 1011080637 | 2 | PVpo | 133 | 0.152110661 | 7.62E-05 | 1 | 0.000501035 | 6.574161171 | 8 mo | HOM |
| 1013609713 | 2 | PVpo | 133 | 0.152110661 | 0 | 0 | 0 | 0 | 2 mo | WILD |
| 1010418842 | 1 | PVpo | 133 | 0.152110661 | 0.000738328 | 9 | 0.004853887 | 59.16745054 | 8 mo | HOM |
| 1013288213 | 2 | PVpo | 133 | 0.152110661 | 4.88E-07 | 0 | 3.21E-06 | 0 | 4 mo | HOM |
| 1008976060 | 2 | PVpo | 133 | 0.152110661 | 5.99E-05 | 5 | 0.000393674 | 32.87080586 | 8 mo | WILD |
| 1013446243 | 1 | PVpo | 133 | 0.152110661 | 1.89E-07 | 0 | 1.24E-06 | 0 | 4 mo | HOM |
| 1013288951 | 2 | PVpo | 133 | 0.152110661 | 7.11E-06 | 0 | 4.68E-05 | 0 | 4 mo | HOM |
| 1013978665 | 1 | AMB | 135 | 0.044390335 | 0 | 0 | 0 | 0 | 2 mo | HOM |
| 1013445609 | 1 | AMB | 135 | 0.044390335 | 0 | 0 | 0 | 0 | 4 mo | HOM |
| 1013609015 | 1 | AMB | 135 | 0.044390335 | 0 | 0 | 0 | 0 | 4 mo | HOM |
| 1013608106 | 2 | AMB | 135 | 0.044390335 | 0 | 0 | 0 | 0 | 4 mo | HOM |
| 1011937543 | 1 | AMB | 135 | 0.044390335 | 0 | 0 | 0 | 0 | 8 mo | HOM |
| 1010419538 | 2 | AMB | 135 | 0.044390335 | 0 | 0 | 0 | 0 | 4 mo | WILD |
| 1010419187 | 1 | AMB | 135 | 0.044390335 | 0 | 0 | 0 | 0 | 8 mo | HOM |
| 1013978366 | 1 | AMB | 135 | 0.044390335 | 0 | 0 | 0 | 0 | 2 mo | HOM |
| 1014416155 | 2 | AMB | 135 | 0.044390335 | 0 | 0 | 0 | 0 | 2 mo | HOM |
| 1013978065 | 2 | AMB | 135 | 0.044390335 | 0 | 0 | 0 | 0 | 2 mo | HOM |
| 1009934102 | 2 | AMB | 135 | 0.044390335 | 0 | 0 | 0 | 0 | 4 mo | WILD |
| 1009322937 | 2 | AMB | 135 | 0.044390335 | 0 | 0 | 0 | 0 | 8 mo | WILD |
| 1011080637 | 2 | AMB | 135 | 0.044390335 | 0 | 0 | 0 | 0 | 8 mo | HOM |
| 1013609713 | 2 | AMB | 135 | 0.044390335 | 0 | 0 | 0 | 0 | 2 mo | WILD |
| 1010418842 | 1 | AMB | 135 | 0.044390335 | 0 | 0 | 0 | 0 | 8 mo | HOM |
| 1013288213 | 2 | AMB | 135 | 0.044390335 | 0 | 0 | 0 | 0 | 4 mo | HOM |
| 1008976060 | 2 | AMB | 135 | 0.044390335 | 0 | 0 | 0 | 0 | 8 mo | WILD |
| 1013446243 | 1 | AMB | 135 | 0.044390335 | 0 | 0 | 0 | 0 | 4 mo | HOM |
| 1013288951 | 2 | AMB | 135 | 0.044390335 | 0 | 0 | 0 | 0 | 4 mo | HOM |
| 1013978665 | 1 | IRN | 136 | 2.861427206 | 0 | 0 | 0 | 0 | 2 mo | HOM |
| 1013445609 | 1 | IRN | 136 | 2.861427206 | 0 | 0 | 0 | 0 | 4 mo | HOM |
| 1013609015 | 1 | IRN | 136 | 2.861427206 | 0 | 0 | 0 | 0 | 4 mo | HOM |
| 1013608106 | 2 | IRN | 136 | 2.861427206 | 0 | 0 | 0 | 0 | 4 mo | HOM |
| 1011937543 | 1 | IRN | 136 | 2.861427206 | 0 | 0 | 0 | 0 | 8 mo | HOM |
| 1010419538 | 2 | IRN | 136 | 2.861427206 | 0 | 0 | 0 | 0 | 4 mo | WILD |
| 1010419187 | 1 | IRN | 136 | 2.861427206 | 1.04E-05 | 1 | 3.65E-06 | 0.349475953 | 8 mo | HOM |
| 1013978366 | 1 | IRN | 136 | 2.861427206 | 0 | 0 | 0 | 0 | 2 mo | HOM |
| 1014416155 | 2 | IRN | 136 | 2.861427206 | 0 | 0 | 0 | 0 | 2 mo | HOM |
| 1013978065 | 2 | IRN | 136 | 2.861427206 | 0 | 0 | 0 | 0 | 2 mo | HOM |
| 1009934102 | 2 | IRN | 136 | 2.861427206 | 0 | 0 | 0 | 0 | 4 mo | WILD |
| 1009322937 | 2 | IRN | 136 | 2.861427206 | 0 | 0 | 0 | 0 | 8 mo | WILD |
| 1011080637 | 2 | IRN | 136 | 2.861427206 | 8.8E-06 | 1 | 3.08E-06 | 0.349475953 | 8 mo | HOM |
| 1013609713 | 2 | IRN | 136 | 2.861427206 | 0 | 0 | 0 | 0 | 2 mo | WILD |
| 1010418842 | 1 | IRN | 136 | 2.861427206 | 0.000181235 | 13 | 6.33E-05 | 4.543187389 | 8 mo | HOM |
| 1013288213 | 2 | IRN | 136 | 2.861427206 | 0 | 0 | 0 | 0 | 4 mo | HOM |

| | | | | | | | | | | |
|---|---|---|---|---|---|---|---|---|---|---|
| **1008976060** | 2 | IRN | 136 | 2.861427206 | 1.12E-05 | 2 | 3.93E-06 | 0.698951906 | 8 mo | WILD |
| **1013446243** | 1 | IRN | 136 | 2.861427206 | 0 | 0 | 0 | 0 | 4 mo | HOM |
| **1013288951** | 2 | IRN | 136 | 2.861427206 | 0 | 0 | 0 | 0 | 4 mo | HOM |
| **1013978665** | 1 | LAT | 138 | 3.168351838 | 0 | 0 | 0 | 0 | 2 mo | HOM |
| **1013445609** | 1 | LAT | 138 | 3.168351838 | 8.75E-06 | 1 | 2.76E-06 | 0.315621513 | 4 mo | HOM |
| **1013609015** | 1 | LAT | 138 | 3.168351838 | 4.46E-05 | 3 | 1.41E-05 | 0.946864538 | 4 mo | HOM |
| **1013608106** | 2 | LAT | 138 | 3.168351838 | 0 | 0 | 0 | 0 | 4 mo | HOM |
| **1011937543** | 1 | LAT | 138 | 3.168351838 | 0.004078835 | 65 | 0.001287368 | 20.51539833 | 8 mo | HOM |
| **1010419538** | 2 | LAT | 138 | 3.168351838 | 0 | 0 | 0 | 0 | 4 mo | WILD |
| **1010419187** | 1 | LAT | 138 | 3.168351838 | 0.001991359 | 57 | 0.000628516 | 17.99042622 | 8 mo | HOM |
| **1013978366** | 1 | LAT | 138 | 3.168351838 | 0 | 0 | 0 | 0 | 2 mo | HOM |
| **1014416155** | 2 | LAT | 138 | 3.168351838 | 0 | 0 | 0 | 0 | 2 mo | HOM |
| **1013978065** | 2 | LAT | 138 | 3.168351838 | 0 | 0 | 0 | 0 | 2 mo | HOM |
| **1009934102** | 2 | LAT | 138 | 3.168351838 | 7.75E-09 | 0 | 2.45E-09 | 0 | 4 mo | WILD |
| **1009322937** | 2 | LAT | 138 | 3.168351838 | 1.69E-05 | 3 | 5.34E-06 | 0.946864538 | 8 mo | WILD |
| **1011080637** | 2 | LAT | 138 | 3.168351838 | 0.002049359 | 65 | 0.000646822 | 20.51539833 | 8 mo | HOM |
| **1013609713** | 2 | LAT | 138 | 3.168351838 | 0 | 0 | 0 | 0 | 2 mo | WILD |
| **1010418842** | 1 | LAT | 138 | 3.168351838 | 0.004337667 | 84 | 0.001369061 | 26.51220707 | 8 mo | HOM |
| **1013288213** | 2 | LAT | 138 | 3.168351838 | 4.63E-05 | 3 | 1.46E-05 | 0.946864538 | 4 mo | HOM |
| **1008976060** | 2 | LAT | 138 | 3.168351838 | 8.85E-05 | 11 | 2.79E-05 | 3.47183664 | 8 mo | WILD |
| **1013446243** | 1 | LAT | 138 | 3.168351838 | 3.42E-05 | 2 | 1.08E-05 | 0.631243025 | 4 mo | HOM |
| **1013288951** | 2 | LAT | 138 | 3.168351838 | 2.81E-05 | 2 | 8.87E-06 | 0.631243025 | 4 mo | HOM |
| **1013978665** | 1 | ENTI5 | 139 | 1.471958746 | 0 | 0 | 0 | 0 | 2 mo | HOM |
| **1013445609** | 1 | ENTI5 | 139 | 1.471958746 | 7.11E-05 | 7 | 4.83E-05 | 4.755568063 | 4 mo | HOM |
| **1013609015** | 1 | ENTI5 | 139 | 1.471958746 | 0.000250689 | 12 | 0.00017031 | 8.152402393 | 4 mo | HOM |
| **1013608106** | 2 | ENTI5 | 139 | 1.471958746 | 0 | 0 | 0 | 0 | 4 mo | HOM |
| **1011937543** | 1 | ENTI5 | 139 | 1.471958746 | 0.002029514 | 58 | 0.001378785 | 39.40327823 | 8 mo | HOM |
| **1010419538** | 2 | ENTI5 | 139 | 1.471958746 | 0 | 0 | 0 | 0 | 4 mo | WILD |
| **1010419187** | 1 | ENTI5 | 139 | 1.471958746 | 0.002030638 | 90 | 0.001379548 | 61.14301795 | 8 mo | HOM |
| **1013978366** | 1 | ENTI5 | 139 | 1.471958746 | 0 | 0 | 0 | 0 | 2 mo | HOM |
| **1014416155** | 2 | ENTI5 | 139 | 1.471958746 | 0 | 0 | 0 | 0 | 2 mo | HOM |
| **1013978065** | 2 | ENTI5 | 139 | 1.471958746 | 0 | 0 | 0 | 0 | 2 mo | HOM |
| **1009934102** | 2 | ENTI5 | 139 | 1.471958746 | 0 | 0 | 0 | 0 | 4 mo | WILD |
| **1009322937** | 2 | ENTI5 | 139 | 1.471958746 | 0 | 0 | 0 | 0 | 8 mo | WILD |
| **1011080637** | 2 | ENTI5 | 139 | 1.471958746 | 0.001063682 | 53 | 0.00072263 | 36.0064439 | 8 mo | HOM |
| **1013609713** | 2 | ENTI5 | 139 | 1.471958746 | 0 | 0 | 0 | 0 | 2 mo | WILD |
| **1010418842** | 1 | ENTI5 | 139 | 1.471958746 | 0.002013163 | 70 | 0.001367676 | 47.55568063 | 8 mo | HOM |
| **1013288213** | 2 | ENTI5 | 139 | 1.471958746 | 0.00013219 | 11 | 8.98E-05 | 7.473035527 | 4 mo | HOM |
| **1008976060** | 2 | ENTI5 | 139 | 1.471958746 | 1.12E-05 | 3 | 7.61E-06 | 2.038100598 | 8 mo | WILD |
| **1013446243** | 1 | ENTI5 | 139 | 1.471958746 | 0.000102193 | 4 | 6.94E-05 | 2.717467464 | 4 mo | HOM |
| **1013288951** | 2 | ENTI5 | 139 | 1.471958746 | 0.000129286 | 11 | 8.78E-05 | 7.473035527 | 4 mo | HOM |
| **1013978665** | 1 | AQ | 140 | 0.417504853 | 5.45E-05 | 5 | 0.000130654 | 11.9759087 | 2 mo | HOM |
| **1013445609** | 1 | AQ | 140 | 0.417504853 | 0.000395292 | 18 | 0.000946797 | 43.11327133 | 4 mo | HOM |
| **1013609015** | 1 | AQ | 140 | 0.417504853 | 0.012584118 | 3 | 0.030141249 | 7.185545222 | 4 mo | HOM |
| **1013608106** | 2 | AQ | 140 | 0.417504853 | 0 | 0 | 0 | 0 | 4 mo | HOM |
| **1011937543** | 1 | AQ | 140 | 0.417504853 | 0.003394724 | 9 | 0.00813098 | 21.55663566 | 8 mo | HOM |
| **1010419538** | 2 | AQ | 140 | 0.417504853 | 0.000145735 | 4 | 0.000349063 | 9.580726962 | 4 mo | WILD |
| **1010419187** | 1 | AQ | 140 | 0.417504853 | 0.007073694 | 5 | 0.016942782 | 11.9759087 | 8 mo | HOM |
| **1013978366** | 1 | AQ | 140 | 0.417504853 | 0.000313988 | 9 | 0.000752058 | 21.55663566 | 2 mo | HOM |
| **1014416155** | 2 | AQ | 140 | 0.417504853 | 0 | 0 | 0 | 0 | 2 mo | HOM |
| **1013978065** | 2 | AQ | 140 | 0.417504853 | 0 | 0 | 0 | 0 | 2 mo | HOM |
| **1009934102** | 2 | AQ | 140 | 0.417504853 | 7.91E-06 | 1 | 1.89E-05 | 2.395181741 | 4 mo | WILD |
| **1009322937** | 2 | AQ | 140 | 0.417504853 | 0.000237386 | 6 | 0.000568582 | 14.37109044 | 8 mo | WILD |
| **1011080637** | 2 | AQ | 140 | 0.417504853 | 0.004593124 | 5 | 0.011001367 | 11.9759087 | 8 mo | HOM |
| **1013609713** | 2 | AQ | 140 | 0.417504853 | 4.55E-05 | 0 | 0.00010897 | 0 | 2 mo | WILD |
| **1010418842** | 1 | AQ | 140 | 0.417504853 | 0.011306069 | 6 | 0.027080091 | 14.37109044 | 8 mo | HOM |
| **1013288213** | 2 | AQ | 140 | 0.417504853 | 4.84E-05 | 3 | 0.000115997 | 7.185545222 | 4 mo | HOM |
| **1008976060** | 2 | AQ | 140 | 0.417504853 | 0.001608106 | 4 | 0.003851705 | 9.580726962 | 8 mo | WILD |
| **1013446243** | 1 | AQ | 140 | 0.417504853 | 0.003880682 | 12 | 0.009294939 | 28.74218089 | 4 mo | HOM |

| | | | | | | | | | | |
|---|---|---|---|---|---|---|---|---|---|---|
| **1013288951** | 2 | AQ | 140 | 0.417504853 | 0.001784419 | 7 | 0.004274008 | 16.76627218 | 4 mo | HOM |
| **1013978665** | 1 | PVR | 141 | 2.107607514 | 3.71E-05 | 1 | 1.76E-05 | 0.474471643 | 2 mo | HOM |
| **1013445609** | 1 | PVR | 141 | 2.107607514 | 0.000197628 | 6 | 9.38E-05 | 2.846829858 | 4 mo | HOM |
| **1013609015** | 1 | PVR | 141 | 2.107607514 | 0.000724826 | 7 | 0.000343909 | 3.321301501 | 4 mo | HOM |
| **1013608106** | 2 | PVR | 141 | 2.107607514 | 0 | 0 | 0 | 0 | 4 mo | HOM |
| **1011937543** | 1 | PVR | 141 | 2.107607514 | 0.001311328 | 13 | 0.000622188 | 6.168131358 | 8 mo | HOM |
| **1010419538** | 2 | PVR | 141 | 2.107607514 | 3.69E-08 | 0 | 1.75E-08 | 0 | 4 mo | WILD |
| **1010419187** | 1 | PVR | 141 | 2.107607514 | 0.00199403 | 21 | 0.000946111 | 9.963904502 | 8 mo | HOM |
| **1013978366** | 1 | PVR | 141 | 2.107607514 | 0.000252838 | 2 | 0.000119965 | 0.948943286 | 2 mo | HOM |
| **1014416155** | 2 | PVR | 141 | 2.107607514 | 0 | 0 | 0 | 0 | 2 mo | HOM |
| **1013978065** | 2 | PVR | 141 | 2.107607514 | 5.25E-06 | 1 | 2.49E-06 | 0.474471643 | 2 mo | HOM |
| **1009934102** | 2 | PVR | 141 | 2.107607514 | 1.58E-05 | 2 | 7.5E-06 | 0.948943286 | 4 mo | WILD |
| **1009322937** | 2 | PVR | 141 | 2.107607514 | 4.82E-05 | 4 | 2.29E-05 | 1.897886572 | 8 mo | WILD |
| **1011080637** | 2 | PVR | 141 | 2.107607514 | 0.001066222 | 19 | 0.000505892 | 9.014961216 | 8 mo | HOM |
| **1013609713** | 2 | PVR | 141 | 2.107607514 | 0 | 0 | 0 | 0 | 2 mo | WILD |
| **1010418842** | 1 | PVR | 141 | 2.107607514 | 0.002955328 | 32 | 0.001402219 | 15.18309257 | 8 mo | HOM |
| **1013288213** | 2 | PVR | 141 | 2.107607514 | 3.39E-05 | 3 | 1.61E-05 | 1.423414929 | 4 mo | HOM |
| **1008976060** | 2 | PVR | 141 | 2.107607514 | 0.00025337 | 8 | 0.000120217 | 3.795773144 | 8 mo | WILD |
| **1013446243** | 1 | PVR | 141 | 2.107607514 | 0.000931913 | 2 | 0.000442166 | 0.948943286 | 4 mo | HOM |
| **1013288951** | 2 | PVR | 141 | 2.107607514 | 0.000433468 | 4 | 0.000205668 | 1.897886572 | 4 mo | HOM |
| **1013978665** | 1 | AMBv | 143 | 0.022541323 | 0 | 0 | 0 | 0 | 2 mo | HOM |
| **1013445609** | 1 | AMBv | 143 | 0.022541323 | 0 | 0 | 0 | 0 | 4 mo | HOM |
| **1013609015** | 1 | AMBv | 143 | 0.022541323 | 0 | 0 | 0 | 0 | 4 mo | HOM |
| **1013608106** | 2 | AMBv | 143 | 0.022541323 | 0 | 0 | 0 | 0 | 4 mo | HOM |
| **1011937543** | 1 | AMBv | 143 | 0.022541323 | 0 | 0 | 0 | 0 | 8 mo | HOM |
| **1010419538** | 2 | AMBv | 143 | 0.022541323 | 0 | 0 | 0 | 0 | 4 mo | WILD |
| **1010419187** | 1 | AMBv | 143 | 0.022541323 | 0 | 0 | 0 | 0 | 8 mo | HOM |
| **1013978366** | 1 | AMBv | 143 | 0.022541323 | 0 | 0 | 0 | 0 | 2 mo | HOM |
| **1014416155** | 2 | AMBv | 143 | 0.022541323 | 0 | 0 | 0 | 0 | 2 mo | HOM |
| **1013978065** | 2 | AMBv | 143 | 0.022541323 | 0 | 0 | 0 | 0 | 2 mo | HOM |
| **1009934102** | 2 | AMBv | 143 | 0.022541323 | 0 | 0 | 0 | 0 | 4 mo | WILD |
| **1009322937** | 2 | AMBv | 143 | 0.022541323 | 0 | 0 | 0 | 0 | 8 mo | WILD |
| **1011080637** | 2 | AMBv | 143 | 0.022541323 | 0 | 0 | 0 | 0 | 8 mo | HOM |
| **1013609713** | 2 | AMBv | 143 | 0.022541323 | 0 | 0 | 0 | 0 | 2 mo | WILD |
| **1010418842** | 1 | AMBv | 143 | 0.022541323 | 0 | 0 | 0 | 0 | 8 mo | HOM |
| **1013288213** | 2 | AMBv | 143 | 0.022541323 | 0 | 0 | 0 | 0 | 4 mo | HOM |
| **1008976060** | 2 | AMBv | 143 | 0.022541323 | 0 | 0 | 0 | 0 | 8 mo | WILD |
| **1013446243** | 1 | AMBv | 143 | 0.022541323 | 0 | 0 | 0 | 0 | 4 mo | HOM |
| **1013288951** | 2 | AMBv | 143 | 0.022541323 | 0 | 0 | 0 | 0 | 4 mo | HOM |
| **1013978665** | 1 | V4 | 145 | 0.990299722 | 0.000307832 | 8 | 0.000310847 | 8.078362364 | 2 mo | HOM |
| **1013445609** | 1 | V4 | 145 | 0.990299722 | 7.06E-05 | 4 | 7.13E-05 | 4.039181182 | 4 mo | HOM |
| **1013609015** | 1 | V4 | 145 | 0.990299722 | 0.003833498 | 7 | 0.003871048 | 7.068567068 | 4 mo | HOM |
| **1013608106** | 2 | V4 | 145 | 0.990299722 | 0 | 0 | 0 | 0 | 4 mo | HOM |
| **1011937543** | 1 | V4 | 145 | 0.990299722 | 0.001234212 | 3 | 0.001246302 | 3.029385886 | 8 mo | HOM |
| **1010419538** | 2 | V4 | 145 | 0.990299722 | 1.86E-05 | 0 | 1.88E-05 | 0 | 4 mo | WILD |
| **1010419187** | 1 | V4 | 145 | 0.990299722 | 0.001289189 | 7 | 0.001301817 | 7.068567068 | 8 mo | HOM |
| **1013978366** | 1 | V4 | 145 | 0.990299722 | 0.000296701 | 9 | 0.000299608 | 9.088157659 | 2 mo | HOM |
| **1014416155** | 2 | V4 | 145 | 0.990299722 | 0 | 0 | 0 | 0 | 2 mo | HOM |
| **1013978065** | 2 | V4 | 145 | 0.990299722 | 0.000284039 | 11 | 0.000286821 | 11.10774825 | 2 mo | HOM |
| **1009934102** | 2 | V4 | 145 | 0.990299722 | 6.69E-05 | 0 | 6.76E-05 | 0 | 4 mo | WILD |
| **1009322937** | 2 | V4 | 145 | 0.990299722 | 0.000487825 | 12 | 0.000492603 | 12.11754355 | 8 mo | WILD |
| **1011080637** | 2 | V4 | 145 | 0.990299722 | 0.003724289 | 12 | 0.003760769 | 12.11754355 | 8 mo | HOM |
| **1013609713** | 2 | V4 | 145 | 0.990299722 | 3.3E-05 | 1 | 3.33E-05 | 1.009795295 | 2 mo | WILD |
| **1010418842** | 1 | V4 | 145 | 0.990299722 | 0.003660008 | 14 | 0.003695858 | 14.13713414 | 8 mo | HOM |
| **1013288213** | 2 | V4 | 145 | 0.990299722 | 0.00025068 | 11 | 0.000253135 | 11.10774825 | 4 mo | HOM |
| **1008976060** | 2 | V4 | 145 | 0.990299722 | 0.002690399 | 18 | 0.002716752 | 18.17631532 | 8 mo | WILD |
| **1013446243** | 1 | V4 | 145 | 0.990299722 | 0.000349424 | 6 | 0.000352847 | 6.058771773 | 4 mo | HOM |
| **1013288951** | 2 | V4 | 145 | 0.990299722 | 0.000897904 | 4 | 0.0009067 | 4.039181182 | 4 mo | HOM |
| **1013978665** | 1 | PRNr | 146 | 2.43447727 | 0 | 0 | 0 | 0 | 2 mo | HOM |

| | | | | | | | | | | |
|---|---|---|---|---|---|---|---|---|---|---|
| **1013445609** | 1 | PRNr | 146 | 2.43447727 | 0 | 0 | 0 | 0 | 4 mo | HOM |
| **1013609015** | 1 | PRNr | 146 | 2.43447727 | 0 | 0 | 0 | 0 | 4 mo | HOM |
| **1013608106** | 2 | PRNr | 146 | 2.43447727 | 0 | 0 | 0 | 0 | 4 mo | HOM |
| **1011937543** | 1 | PRNr | 146 | 2.43447727 | 0.007526238 | 14 | 0.003091521 | 5.75072118 | 8 mo | HOM |
| **1010419538** | 2 | PRNr | 146 | 2.43447727 | 0 | 0 | 0 | 0 | 4 mo | WILD |
| **1010419187** | 1 | PRNr | 146 | 2.43447727 | 0 | 0 | 0 | 0 | 8 mo | HOM |
| **1013978366** | 1 | PRNr | 146 | 2.43447727 | 0 | 0 | 0 | 0 | 2 mo | HOM |
| **1014416155** | 2 | PRNr | 146 | 2.43447727 | 0 | 0 | 0 | 0 | 2 mo | HOM |
| **1013978065** | 2 | PRNr | 146 | 2.43447727 | 0 | 0 | 0 | 0 | 2 mo | HOM |
| **1009934102** | 2 | PRNr | 146 | 2.43447727 | 0 | 0 | 0 | 0 | 4 mo | WILD |
| **1009322937** | 2 | PRNr | 146 | 2.43447727 | 7.05E-06 | 1 | 2.89E-06 | 0.410765799 | 8 mo | WILD |
| **1011080637** | 2 | PRNr | 146 | 2.43447727 | 1.46E-05 | 1 | 6E-06 | 0.410765799 | 8 mo | HOM |
| **1013609713** | 2 | PRNr | 146 | 2.43447727 | 0 | 0 | 0 | 0 | 2 mo | WILD |
| **1010418842** | 1 | PRNr | 146 | 2.43447727 | 0.000194347 | 14 | 7.98E-05 | 5.75072118 | 8 mo | HOM |
| **1013288213** | 2 | PRNr | 146 | 2.43447727 | 0 | 0 | 0 | 0 | 4 mo | HOM |
| **1008976060** | 2 | PRNr | 146 | 2.43447727 | 0.000231205 | 16 | 9.5E-05 | 6.572252777 | 8 mo | WILD |
| **1013446243** | 1 | PRNr | 146 | 2.43447727 | 0 | 0 | 0 | 0 | 4 mo | HOM |
| **1013288951** | 2 | PRNr | 146 | 2.43447727 | 0 | 0 | 0 | 0 | 4 mo | HOM |
| **1013978665** | 1 | LC | 147 | 0.014923839 | 0 | 0 | 0 | 0 | 2 mo | HOM |
| **1013445609** | 1 | LC | 147 | 0.014923839 | 0 | 0 | 0 | 0 | 4 mo | HOM |
| **1013609015** | 1 | LC | 147 | 0.014923839 | 8.85E-06 | 0 | 0.000593078 | 0 | 4 mo | HOM |
| **1013608106** | 2 | LC | 147 | 0.014923839 | 0 | 0 | 0 | 0 | 4 mo | HOM |
| **1011937543** | 1 | LC | 147 | 0.014923839 | 0 | 0 | 0 | 0 | 8 mo | HOM |
| **1010419538** | 2 | LC | 147 | 0.014923839 | 0 | 0 | 0 | 0 | 4 mo | WILD |
| **1010419187** | 1 | LC | 147 | 0.014923839 | 9.63E-07 | 0 | 6.45E-05 | 0 | 8 mo | HOM |
| **1013978366** | 1 | LC | 147 | 0.014923839 | 1.92E-06 | 0 | 0.00012865 | 0 | 2 mo | HOM |
| **1014416155** | 2 | LC | 147 | 0.014923839 | 0 | 0 | 0 | 0 | 2 mo | HOM |
| **1013978065** | 2 | LC | 147 | 0.014923839 | 0 | 0 | 0 | 0 | 2 mo | HOM |
| **1009934102** | 2 | LC | 147 | 0.014923839 | 0 | 0 | 0 | 0 | 4 mo | WILD |
| **1009322937** | 2 | LC | 147 | 0.014923839 | 0 | 0 | 0 | 0 | 8 mo | WILD |
| **1011080637** | 2 | LC | 147 | 0.014923839 | 0 | 0 | 0 | 0 | 8 mo | HOM |
| **1013609713** | 2 | LC | 147 | 0.014923839 | 0 | 0 | 0 | 0 | 2 mo | WILD |
| **1010418842** | 1 | LC | 147 | 0.014923839 | 5.09E-06 | 0 | 0.000340951 | 0 | 8 mo | HOM |
| **1013288213** | 2 | LC | 147 | 0.014923839 | 0 | 0 | 0 | 0 | 4 mo | HOM |
| **1008976060** | 2 | LC | 147 | 0.014923839 | 5.35E-06 | 1 | 0.000358462 | 67.00688587 | 8 mo | WILD |
| **1013446243** | 1 | LC | 147 | 0.014923839 | 1.02E-08 | 0 | 6.81E-07 | 0 | 4 mo | HOM |
| **1013288951** | 2 | LC | 147 | 0.014923839 | 3.34E-07 | 0 | 2.24E-05 | 0 | 4 mo | HOM |
| **1013978665** | 1 | GU4 | 148 | 0.151061894 | 0 | 0 | 0 | 0 | 2 mo | HOM |
| **1013445609** | 1 | GU4 | 148 | 0.151061894 | 0 | 0 | 0 | 0 | 4 mo | HOM |
| **1013609015** | 1 | GU4 | 148 | 0.151061894 | 1.49E-05 | 0 | 9.87E-05 | 0 | 4 mo | HOM |
| **1013608106** | 2 | GU4 | 148 | 0.151061894 | 0 | 0 | 0 | 0 | 4 mo | HOM |
| **1011937543** | 1 | GU4 | 148 | 0.151061894 | 9.81E-05 | 4 | 0.000649157 | 26.47921252 | 8 mo | HOM |
| **1010419538** | 2 | GU4 | 148 | 0.151061894 | 0 | 0 | 0 | 0 | 4 mo | WILD |
| **1010419187** | 1 | GU4 | 148 | 0.151061894 | 0.00017763 | 9 | 0.001175873 | 59.57822817 | 8 mo | HOM |
| **1013978366** | 1 | GU4 | 148 | 0.151061894 | 0 | 0 | 0 | 0 | 2 mo | HOM |
| **1014416155** | 2 | GU4 | 148 | 0.151061894 | 0 | 0 | 0 | 0 | 2 mo | HOM |
| **1013978065** | 2 | GU4 | 148 | 0.151061894 | 0 | 0 | 0 | 0 | 2 mo | HOM |
| **1009934102** | 2 | GU4 | 148 | 0.151061894 | 0 | 0 | 0 | 0 | 4 mo | WILD |
| **1009322937** | 2 | GU4 | 148 | 0.151061894 | 0 | 0 | 0 | 0 | 8 mo | WILD |
| **1011080637** | 2 | GU4 | 148 | 0.151061894 | 0.00011691 | 5 | 0.000773921 | 33.09901565 | 8 mo | HOM |
| **1013609713** | 2 | GU4 | 148 | 0.151061894 | 0 | 0 | 0 | 0 | 2 mo | WILD |
| **1010418842** | 1 | GU4 | 148 | 0.151061894 | 9.79E-05 | 6 | 0.000648105 | 39.71881878 | 8 mo | HOM |
| **1013288213** | 2 | GU4 | 148 | 0.151061894 | 0 | 0 | 0 | 0 | 4 mo | HOM |
| **1008976060** | 2 | GU4 | 148 | 0.151061894 | 8.1E-06 | 1 | 5.36E-05 | 6.61980313 | 8 mo | WILD |
| **1013446243** | 1 | GU4 | 148 | 0.151061894 | 0 | 0 | 0 | 0 | 4 mo | HOM |
| **1013288951** | 2 | GU4 | 148 | 0.151061894 | 1.36E-05 | 1 | 8.98E-05 | 6.61980313 | 4 mo | HOM |
| **1013978665** | 1 | PVT | 149 | 0.457918524 | 6.06E-09 | 0 | 1.32E-08 | 0 | 2 mo | HOM |
| **1013445609** | 1 | PVT | 149 | 0.457918524 | 0 | 0 | 0 | 0 | 4 mo | HOM |
| **1013609015** | 1 | PVT | 149 | 0.457918524 | 0.000354894 | 12 | 0.000775015 | 26.2055352 | 4 mo | HOM |

| | | | | | | | | | | |
|---|---|---|---|---|---|---|---|---|---|---|
| 1013608106 | 2 | PVT | 149 | 0.457918524 | 0 | 0 | 0 | 0 | 4 mo | HOM |
| 1011937543 | 1 | PVT | 149 | 0.457918524 | 0.000333711 | 5 | 0.000728756 | 10.918973 | 8 mo | HOM |
| 1010419538 | 2 | PVT | 149 | 0.457918524 | 0 | 0 | 0 | 0 | 4 mo | WILD |
| 1010419187 | 1 | PVT | 149 | 0.457918524 | 0.000342581 | 13 | 0.000748126 | 28.3893298 | 8 mo | HOM |
| 1013978366 | 1 | PVT | 149 | 0.457918524 | 1.41E-07 | 0 | 3.08E-07 | 0 | 2 mo | HOM |
| 1014416155 | 2 | PVT | 149 | 0.457918524 | 0 | 0 | 0 | 0 | 2 mo | HOM |
| 1013978065 | 2 | PVT | 149 | 0.457918524 | 9.78E-07 | 0 | 2.14E-06 | 0 | 2 mo | HOM |
| 1009934102 | 2 | PVT | 149 | 0.457918524 | 0 | 0 | 0 | 0 | 4 mo | WILD |
| 1009322937 | 2 | PVT | 149 | 0.457918524 | 1.11E-05 | 1 | 2.43E-05 | 2.1837946 | 8 mo | WILD |
| 1011080637 | 2 | PVT | 149 | 0.457918524 | 1.94E-05 | 2 | 4.25E-05 | 4.3675892 | 8 mo | HOM |
| 1013609713 | 2 | PVT | 149 | 0.457918524 | 0 | 0 | 0 | 0 | 2 mo | WILD |
| 1010418842 | 1 | PVT | 149 | 0.457918524 | 1.14E-06 | 0 | 2.49E-06 | 0 | 8 mo | HOM |
| 1013288213 | 2 | PVT | 149 | 0.457918524 | 0 | 0 | 0 | 0 | 4 mo | HOM |
| 1008976060 | 2 | PVT | 149 | 0.457918524 | 2.28E-05 | 2 | 4.98E-05 | 4.3675892 | 8 mo | WILD |
| 1013446243 | 1 | PVT | 149 | 0.457918524 | 3.21E-05 | 2 | 7E-05 | 4.3675892 | 4 mo | HOM |
| 1013288951 | 2 | PVT | 149 | 0.457918524 | 0 | 0 | 0 | 0 | 4 mo | HOM |
| 1013978665 | 1 | AOB | 151 | 0.671739677 | 0 | 0 | 0 | 0 | 2 mo | HOM |
| 1013445609 | 1 | AOB | 151 | 0.671739677 | 0 | 0 | 0 | 0 | 4 mo | HOM |
| 1013609015 | 1 | AOB | 151 | 0.671739677 | 2.22E-05 | 1 | 3.3E-05 | 1.488671928 | 4 mo | HOM |
| 1013608106 | 2 | AOB | 151 | 0.671739677 | 0 | 0 | 0 | 0 | 4 mo | HOM |
| 1011937543 | 1 | AOB | 151 | 0.671739677 | 0.001849396 | 6 | 0.002753144 | 8.932031567 | 8 mo | HOM |
| 1010419538 | 2 | AOB | 151 | 0.671739677 | 0 | 0 | 0 | 0 | 4 mo | WILD |
| 1010419187 | 1 | AOB | 151 | 0.671739677 | 0.00060703 | 21 | 0.000903668 | 31.26211049 | 8 mo | HOM |
| 1013978366 | 1 | AOB | 151 | 0.671739677 | 1.38E-07 | 0 | 2.05E-07 | 0 | 2 mo | HOM |
| 1014416155 | 2 | AOB | 151 | 0.671739677 | 0 | 0 | 0 | 0 | 2 mo | HOM |
| 1013978065 | 2 | AOB | 151 | 0.671739677 | 0 | 0 | 0 | 0 | 2 mo | HOM |
| 1009934102 | 2 | AOB | 151 | 0.671739677 | 5.81E-06 | 0 | 8.64E-06 | 0 | 4 mo | WILD |
| 1009322937 | 2 | AOB | 151 | 0.671739677 | 2.37E-05 | 4 | 3.53E-05 | 5.954687712 | 8 mo | WILD |
| 1011080637 | 2 | AOB | 151 | 0.671739677 | 0.000163674 | 13 | 0.000243657 | 19.35273506 | 8 mo | HOM |
| 1013609713 | 2 | AOB | 151 | 0.671739677 | 4.37E-06 | 0 | 6.5E-06 | 0 | 2 mo | WILD |
| 1010418842 | 1 | AOB | 151 | 0.671739677 | 0.000181021 | 15 | 0.000269481 | 22.33007892 | 8 mo | HOM |
| 1013288213 | 2 | AOB | 151 | 0.671739677 | 0.000837092 | 5 | 0.001246156 | 7.443359639 | 4 mo | HOM |
| 1008976060 | 2 | AOB | 151 | 0.671739677 | 8.29E-07 | 0 | 1.23E-06 | 0 | 8 mo | WILD |
| 1013446243 | 1 | AOB | 151 | 0.671739677 | 2.18E-05 | 2 | 3.25E-05 | 2.977343856 | 4 mo | HOM |
| 1013288951 | 2 | AOB | 151 | 0.671739677 | 0.000289505 | 14 | 0.000430977 | 20.84140699 | 4 mo | HOM |
| 1013978665 | 1 | V4r | 153 | 0.475634298 | 7.27E-06 | 1 | 1.53E-05 | 2.102455616 | 2 mo | HOM |
| 1013445609 | 1 | V4r | 153 | 0.475634298 | 2.83E-05 | 1 | 5.95E-05 | 2.102455616 | 4 mo | HOM |
| 1013609015 | 1 | V4r | 153 | 0.475634298 | 0.000646474 | 3 | 0.001359183 | 6.307366847 | 4 mo | HOM |
| 1013608106 | 2 | V4r | 153 | 0.475634298 | 0 | 0 | 0 | 0 | 4 mo | HOM |
| 1011937543 | 1 | V4r | 153 | 0.475634298 | 0.000982626 | 1 | 0.002065927 | 2.102455616 | 8 mo | HOM |
| 1010419538 | 2 | V4r | 153 | 0.475634298 | 1.29E-05 | 0 | 2.72E-05 | 0 | 4 mo | WILD |
| 1010419187 | 1 | V4r | 153 | 0.475634298 | 0.000414322 | 5 | 0.000871094 | 10.51227808 | 8 mo | HOM |
| 1013978366 | 1 | V4r | 153 | 0.475634298 | 0.000110346 | 3 | 0.000231999 | 6.307366847 | 2 mo | HOM |
| 1014416155 | 2 | V4r | 153 | 0.475634298 | 0 | 0 | 0 | 0 | 2 mo | HOM |
| 1013978065 | 2 | V4r | 153 | 0.475634298 | 6E-05 | 3 | 0.000126225 | 6.307366847 | 2 mo | HOM |
| 1009934102 | 2 | V4r | 153 | 0.475634298 | 2.19E-06 | 0 | 4.61E-06 | 0 | 4 mo | WILD |
| 1009322937 | 2 | V4r | 153 | 0.475634298 | 4.54E-05 | 2 | 9.55E-05 | 4.204911231 | 8 mo | WILD |
| 1011080637 | 2 | V4r | 153 | 0.475634298 | 0.000560175 | 8 | 0.001177743 | 16.81964493 | 8 mo | HOM |
| 1013609713 | 2 | V4r | 153 | 0.475634298 | 2.93E-06 | 0 | 6.17E-06 | 0 | 2 mo | WILD |
| 1010418842 | 1 | V4r | 153 | 0.475634298 | 0.000282663 | 5 | 0.000594286 | 10.51227808 | 8 mo | HOM |
| 1013288213 | 2 | V4r | 153 | 0.475634298 | 1.01E-05 | 1 | 2.13E-05 | 2.102455616 | 4 mo | HOM |
| 1008976060 | 2 | V4r | 153 | 0.475634298 | 0.000952811 | 10 | 0.002003244 | 21.02455616 | 8 mo | WILD |
| 1013446243 | 1 | V4r | 153 | 0.475634298 | 2.89E-06 | 0 | 6.08E-06 | 0 | 4 mo | HOM |
| 1013288951 | 2 | V4r | 153 | 0.475634298 | 0.000122488 | 0 | 0.000257526 | 0 | 4 mo | HOM |
| 1013978665 | 1 | PHY | 154 | 0.282312549 | 1.24E-06 | 0 | 4.39E-06 | 0 | 2 mo | HOM |
| 1013445609 | 1 | PHY | 154 | 0.282312549 | 0 | 0 | 0 | 0 | 4 mo | HOM |
| 1013609015 | 1 | PHY | 154 | 0.282312549 | 8.19E-05 | 0 | 0.000290114 | 0 | 4 mo | HOM |
| 1013608106 | 2 | PHY | 154 | 0.282312549 | 0 | 0 | 0 | 0 | 4 mo | HOM |
| 1011937543 | 1 | PHY | 154 | 0.282312549 | 0 | 0 | 0 | 0 | 8 mo | HOM |

| | | | | | | | | | | |
|---|---|---|---|---|---|---|---|---|---|---|
| **1010419538** | 2 | PHY | 154 | 0.282312549 | 0 | 0 | 0 | 0 | 4 mo | WILD |
| **1010419187** | 1 | PHY | 154 | 0.282312549 | 3.58E-10 | 0 | 1.27E-09 | 0 | 8 mo | HOM |
| **1013978366** | 1 | PHY | 154 | 0.282312549 | 7.67E-08 | 0 | 2.72E-07 | 0 | 2 mo | HOM |
| **1014416155** | 2 | PHY | 154 | 0.282312549 | 0 | 0 | 0 | 0 | 2 mo | HOM |
| **1013978065** | 2 | PHY | 154 | 0.282312549 | 0 | 0 | 0 | 0 | 2 mo | HOM |
| **1009934102** | 2 | PHY | 154 | 0.282312549 | 0 | 0 | 0 | 0 | 4 mo | WILD |
| **1009322937** | 2 | PHY | 154 | 0.282312549 | 0 | 0 | 0 | 0 | 8 mo | WILD |
| **1011080637** | 2 | PHY | 154 | 0.282312549 | 0.000270426 | 2 | 0.000957896 | 7.084346779 | 8 mo | HOM |
| **1013609713** | 2 | PHY | 154 | 0.282312549 | 0 | 0 | 0 | 0 | 2 mo | WILD |
| **1010418842** | 1 | PHY | 154 | 0.282312549 | 3.13E-05 | 4 | 0.000111014 | 14.16869356 | 8 mo | HOM |
| **1013288213** | 2 | PHY | 154 | 0.282312549 | 0 | 0 | 0 | 0 | 4 mo | HOM |
| **1008976060** | 2 | PHY | 154 | 0.282312549 | 0.000156133 | 6 | 0.00055305 | 21.25304034 | 8 mo | WILD |
| **1013446243** | 1 | PHY | 154 | 0.282312549 | 0 | 0 | 0 | 0 | 4 mo | HOM |
| **1013288951** | 2 | PHY | 154 | 0.282312549 | 6.29E-06 | 0 | 2.23E-05 | 0 | 4 mo | HOM |
| **1013978665** | 1 | LD | 155 | 1.047207224 | 0 | 0 | 0 | 0 | 2 mo | HOM |
| **1013445609** | 1 | LD | 155 | 1.047207224 | 5.92E-05 | 3 | 5.65E-05 | 2.864762514 | 4 mo | HOM |
| **1013609015** | 1 | LD | 155 | 1.047207224 | 0.000263782 | 9 | 0.00025189 | 8.594287543 | 4 mo | HOM |
| **1013608106** | 2 | LD | 155 | 1.047207224 | 0 | 0 | 0 | 0 | 4 mo | HOM |
| **1011937543** | 1 | LD | 155 | 1.047207224 | 0.003178522 | 16 | 0.003035236 | 15.27873341 | 8 mo | HOM |
| **1010419538** | 2 | LD | 155 | 1.047207224 | 0 | 0 | 0 | 0 | 4 mo | WILD |
| **1010419187** | 1 | LD | 155 | 1.047207224 | 0.002802559 | 49 | 0.002676222 | 46.79112107 | 8 mo | HOM |
| **1013978366** | 1 | LD | 155 | 1.047207224 | 3.02E-06 | 1 | 2.88E-06 | 0.954920838 | 2 mo | HOM |
| **1014416155** | 2 | LD | 155 | 1.047207224 | 0 | 0 | 0 | 0 | 2 mo | HOM |
| **1013978065** | 2 | LD | 155 | 1.047207224 | 0 | 0 | 0 | 0 | 2 mo | HOM |
| **1009934102** | 2 | LD | 155 | 1.047207224 | 5.01E-06 | 1 | 4.78E-06 | 0.954920838 | 4 mo | WILD |
| **1009322937** | 2 | LD | 155 | 1.047207224 | 5.98E-06 | 1 | 5.71E-06 | 0.954920838 | 8 mo | WILD |
| **1011080637** | 2 | LD | 155 | 1.047207224 | 0.00088498 | 29 | 0.000845086 | 27.6927043 | 8 mo | HOM |
| **1013609713** | 2 | LD | 155 | 1.047207224 | 0 | 0 | 0 | 0 | 2 mo | WILD |
| **1010418842** | 1 | LD | 155 | 1.047207224 | 0.002655499 | 44 | 0.002535792 | 42.01651688 | 8 mo | HOM |
| **1013288213** | 2 | LD | 155 | 1.047207224 | 1.12E-05 | 1 | 1.07E-05 | 0.954920838 | 4 mo | HOM |
| **1008976060** | 2 | LD | 155 | 1.047207224 | 9.81E-05 | 9 | 9.37E-05 | 8.594287543 | 8 mo | WILD |
| **1013446243** | 1 | LD | 155 | 1.047207224 | 0.000148774 | 7 | 0.000142067 | 6.684445867 | 4 mo | HOM |
| **1013288951** | 2 | LD | 155 | 1.047207224 | 9.12E-05 | 4 | 8.71E-05 | 3.819683352 | 4 mo | HOM |
| **1013978665** | 1 | AUDd6a | 156 | 0.190115668 | 0 | 0 | 0 | 0 | 2 mo | HOM |
| **1013445609** | 1 | AUDd6a | 156 | 0.190115668 | 5.73E-06 | 1 | 3.02E-05 | 5.259955733 | 4 mo | HOM |
| **1013609015** | 1 | AUDd6a | 156 | 0.190115668 | 5.99E-05 | 4 | 0.000315144 | 21.03982293 | 4 mo | HOM |
| **1013608106** | 2 | AUDd6a | 156 | 0.190115668 | 0 | 0 | 0 | 0 | 4 mo | HOM |
| **1011937543** | 1 | AUDd6a | 156 | 0.190115668 | 0.000259408 | 5 | 0.001364474 | 26.29977867 | 8 mo | HOM |
| **1010419538** | 2 | AUDd6a | 156 | 0.190115668 | 0 | 0 | 0 | 0 | 4 mo | WILD |
| **1010419187** | 1 | AUDd6a | 156 | 0.190115668 | 0.000214679 | 13 | 0.001129201 | 68.37942453 | 8 mo | HOM |
| **1013978366** | 1 | AUDd6a | 156 | 0.190115668 | 0 | 0 | 0 | 0 | 2 mo | HOM |
| **1014416155** | 2 | AUDd6a | 156 | 0.190115668 | 0 | 0 | 0 | 0 | 2 mo | HOM |
| **1013978065** | 2 | AUDd6a | 156 | 0.190115668 | 0 | 0 | 0 | 0 | 2 mo | HOM |
| **1009934102** | 2 | AUDd6a | 156 | 0.190115668 | 0 | 0 | 0 | 0 | 4 mo | WILD |
| **1009322937** | 2 | AUDd6a | 156 | 0.190115668 | 0 | 0 | 0 | 0 | 8 mo | WILD |
| **1011080637** | 2 | AUDd6a | 156 | 0.190115668 | 0.000165093 | 4 | 0.000868381 | 21.03982293 | 8 mo | HOM |
| **1013609713** | 2 | AUDd6a | 156 | 0.190115668 | 0 | 0 | 0 | 0 | 2 mo | WILD |
| **1010418842** | 1 | AUDd6a | 156 | 0.190115668 | 0.000253307 | 8 | 0.001332384 | 42.07964587 | 8 mo | HOM |
| **1013288213** | 2 | AUDd6a | 156 | 0.190115668 | 0 | 0 | 0 | 0 | 4 mo | HOM |
| **1008976060** | 2 | AUDd6a | 156 | 0.190115668 | 0 | 0 | 0 | 0 | 8 mo | WILD |
| **1013446243** | 1 | AUDd6a | 156 | 0.190115668 | 5.67E-05 | 2 | 0.000298446 | 10.51991147 | 4 mo | HOM |
| **1013288951** | 2 | AUDd6a | 156 | 0.190115668 | 3.99E-05 | 1 | 0.00020986 | 5.259955733 | 4 mo | HOM |
| **1013978665** | 1 | PVZ | 157 | 0.806320017 | 0 | 0 | 0 | 0 | 2 mo | HOM |
| **1013445609** | 1 | PVZ | 157 | 0.806320017 | 4.44E-06 | 1 | 5.5E-06 | 1.240202375 | 4 mo | HOM |
| **1013609015** | 1 | PVZ | 157 | 0.806320017 | 0 | 0 | 0 | 0 | 4 mo | HOM |
| **1013608106** | 2 | PVZ | 157 | 0.806320017 | 0 | 0 | 0 | 0 | 4 mo | HOM |
| **1011937543** | 1 | PVZ | 157 | 0.806320017 | 0.001356795 | 6 | 0.0016827 | 7.44121425 | 8 mo | HOM |
| **1010419538** | 2 | PVZ | 157 | 0.806320017 | 1.2E-05 | 3 | 1.49E-05 | 3.720607125 | 4 mo | WILD |
| **1010419187** | 1 | PVZ | 157 | 0.806320017 | 2.78E-05 | 1 | 3.45E-05 | 1.240202375 | 8 mo | HOM |

| | | | | | | | | | | |
|---|---|---|---|---|---|---|---|---|---|---|
| **1013978366** | 1 | PVZ | 157 | 0.806320017 | 0 | 0 | 0 | 0 | 2 mo | HOM |
| **1014416155** | 2 | PVZ | 157 | 0.806320017 | 0 | 0 | 0 | 0 | 2 mo | HOM |
| **1013978065** | 2 | PVZ | 157 | 0.806320017 | 0 | 0 | 0 | 0 | 2 mo | HOM |
| **1009934102** | 2 | PVZ | 157 | 0.806320017 | 0 | 0 | 0 | 0 | 4 mo | WILD |
| **1009322937** | 2 | PVZ | 157 | 0.806320017 | 0 | 0 | 0 | 0 | 8 mo | WILD |
| **1011080637** | 2 | PVZ | 157 | 0.806320017 | 8.48E-05 | 1 | 0.00010519 | 1.240202375 | 8 mo | HOM |
| **1013609713** | 2 | PVZ | 157 | 0.806320017 | 0 | 0 | 0 | 0 | 2 mo | WILD |
| **1010418842** | 1 | PVZ | 157 | 0.806320017 | 0.000348596 | 18 | 0.00043233 | 22.32364275 | 8 mo | HOM |
| **1013288213** | 2 | PVZ | 157 | 0.806320017 | 1.3E-07 | 0 | 1.61E-07 | 0 | 4 mo | HOM |
| **1008976060** | 2 | PVZ | 157 | 0.806320017 | 0.000229705 | 17 | 0.00028488 | 21.08344037 | 8 mo | WILD |
| **1013446243** | 1 | PVZ | 157 | 0.806320017 | 2.72E-06 | 1 | 3.37E-06 | 1.240202375 | 4 mo | HOM |
| **1013288951** | 2 | PVZ | 157 | 0.806320017 | 5.2E-08 | 0 | 6.45E-08 | 0 | 4 mo | HOM |
| **1013978665** | 1 | pc | 158 | 0.054045603 | 0 | 0 | 0 | 0 | 2 mo | HOM |
| **1013445609** | 1 | pc | 158 | 0.054045603 | 0 | 0 | 0 | 0 | 4 mo | HOM |
| **1013609015** | 1 | pc | 158 | 0.054045603 | 0 | 0 | 0 | 0 | 4 mo | HOM |
| **1013608106** | 2 | pc | 158 | 0.054045603 | 0 | 0 | 0 | 0 | 4 mo | HOM |
| **1011937543** | 1 | pc | 158 | 0.054045603 | 1.4E-05 | 0 | 0.000258135 | 0 | 8 mo | HOM |
| **1010419538** | 2 | pc | 158 | 0.054045603 | 0 | 0 | 0 | 0 | 4 mo | WILD |
| **1010419187** | 1 | pc | 158 | 0.054045603 | 0 | 0 | 0 | 0 | 8 mo | HOM |
| **1013978366** | 1 | pc | 158 | 0.054045603 | 0 | 0 | 0 | 0 | 2 mo | HOM |
| **1014416155** | 2 | pc | 158 | 0.054045603 | 0 | 0 | 0 | 0 | 2 mo | HOM |
| **1013978065** | 2 | pc | 158 | 0.054045603 | 0 | 0 | 0 | 0 | 2 mo | HOM |
| **1009934102** | 2 | pc | 158 | 0.054045603 | 0 | 0 | 0 | 0 | 4 mo | WILD |
| **1009322937** | 2 | pc | 158 | 0.054045603 | 0 | 0 | 0 | 0 | 8 mo | WILD |
| **1011080637** | 2 | pc | 158 | 0.054045603 | 0 | 0 | 0 | 0 | 8 mo | HOM |
| **1013609713** | 2 | pc | 158 | 0.054045603 | 0 | 0 | 0 | 0 | 2 mo | WILD |
| **1010418842** | 1 | pc | 158 | 0.054045603 | 0 | 0 | 0 | 0 | 8 mo | HOM |
| **1013288213** | 2 | pc | 158 | 0.054045603 | 3.01E-09 | 0 | 5.57E-08 | 0 | 4 mo | HOM |
| **1008976060** | 2 | pc | 158 | 0.054045603 | 0 | 0 | 0 | 0 | 8 mo | WILD |
| **1013446243** | 1 | pc | 158 | 0.054045603 | 0 | 0 | 0 | 0 | 4 mo | HOM |
| **1013288951** | 2 | pc | 158 | 0.054045603 | 0 | 0 | 0 | 0 | 4 mo | HOM |
| **1013978665** | 1 | AON | 159 | 5.017900429 | 9.39E-08 | 0 | 1.87E-08 | 0 | 2 mo | HOM |
| **1013445609** | 1 | AON | 159 | 5.017900429 | 2.42E-05 | 2 | 4.83E-06 | 0.398573074 | 4 mo | HOM |
| **1013609015** | 1 | AON | 159 | 5.017900429 | 8.71E-05 | 7 | 1.74E-05 | 1.39500576 | 4 mo | HOM |
| **1013608106** | 2 | AON | 159 | 5.017900429 | 0 | 0 | 0 | 0 | 4 mo | HOM |
| **1011937543** | 1 | AON | 159 | 5.017900429 | 0.002502338 | 52 | 0.000498682 | 10.36289993 | 8 mo | HOM |
| **1010419538** | 2 | AON | 159 | 5.017900429 | 0 | 0 | 0 | 0 | 4 mo | WILD |
| **1010419187** | 1 | AON | 159 | 5.017900429 | 0.001891152 | 91 | 0.000376881 | 18.13507488 | 8 mo | HOM |
| **1013978366** | 1 | AON | 159 | 5.017900429 | 0 | 0 | 0 | 0 | 2 mo | HOM |
| **1014416155** | 2 | AON | 159 | 5.017900429 | 2.17E-06 | 0 | 4.32E-07 | 0 | 2 mo | HOM |
| **1013978065** | 2 | AON | 159 | 5.017900429 | 0 | 0 | 0 | 0 | 2 mo | HOM |
| **1009934102** | 2 | AON | 159 | 5.017900429 | 0 | 0 | 0 | 0 | 4 mo | WILD |
| **1009322937** | 2 | AON | 159 | 5.017900429 | 8.31E-05 | 5 | 1.66E-05 | 0.996432686 | 8 mo | WILD |
| **1011080637** | 2 | AON | 159 | 5.017900429 | 0.001542293 | 76 | 0.000307358 | 15.14577682 | 8 mo | HOM |
| **1013609713** | 2 | AON | 159 | 5.017900429 | 0 | 0 | 0 | 0 | 2 mo | WILD |
| **1010418842** | 1 | AON | 159 | 5.017900429 | 0.00348619 | 118 | 0.000694751 | 23.51581138 | 8 mo | HOM |
| **1013288213** | 2 | AON | 159 | 5.017900429 | 0.001723784 | 16 | 0.000343527 | 3.188584594 | 4 mo | HOM |
| **1008976060** | 2 | AON | 159 | 5.017900429 | 0.000731327 | 30 | 0.000145744 | 5.978596113 | 8 mo | WILD |
| **1013446243** | 1 | AON | 159 | 5.017900429 | 7.71E-05 | 6 | 1.54E-05 | 1.195719223 | 4 mo | HOM |
| **1013288951** | 2 | AON | 159 | 5.017900429 | 0.000467633 | 16 | 9.32E-05 | 3.188584594 | 4 mo | HOM |
| **1013978665** | 1 | LDT | 162 | 0.209626057 | 0 | 0 | 0 | 0 | 2 mo | HOM |
| **1013445609** | 1 | LDT | 162 | 0.209626057 | 0 | 0 | 0 | 0 | 4 mo | HOM |
| **1013609015** | 1 | LDT | 162 | 0.209626057 | 0 | 0 | 0 | 0 | 4 mo | HOM |
| **1013608106** | 2 | LDT | 162 | 0.209626057 | 0 | 0 | 0 | 0 | 4 mo | HOM |
| **1011937543** | 1 | LDT | 162 | 0.209626057 | 0.000153139 | 2 | 0.000730536 | 9.540798633 | 8 mo | HOM |
| **1010419538** | 2 | LDT | 162 | 0.209626057 | 0 | 0 | 0 | 0 | 4 mo | WILD |
| **1010419187** | 1 | LDT | 162 | 0.209626057 | 2.35E-05 | 4 | 0.000112114 | 19.08159727 | 8 mo | HOM |
| **1013978366** | 1 | LDT | 162 | 0.209626057 | 0 | 0 | 0 | 0 | 2 mo | HOM |
| **1014416155** | 2 | LDT | 162 | 0.209626057 | 0 | 0 | 0 | 0 | 2 mo | HOM |

| | | | | | | | | | | |
|---|---|---|---|---|---|---|---|---|---|---|
| **1013978065** | 2 | LDT | 162 | 0.209626057 | 0 | 0 | 0 | 0 | 2 mo | HOM |
| **1009934102** | 2 | LDT | 162 | 0.209626057 | 0 | 0 | 0 | 0 | 4 mo | WILD |
| **1009322937** | 2 | LDT | 162 | 0.209626057 | 2.79E-07 | 0 | 1.33E-06 | 0 | 8 mo | WILD |
| **1011080637** | 2 | LDT | 162 | 0.209626057 | 1.31E-07 | 0 | 6.25E-07 | 0 | 8 mo | HOM |
| **1013609713** | 2 | LDT | 162 | 0.209626057 | 0 | 0 | 0 | 0 | 2 mo | WILD |
| **1010418842** | 1 | LDT | 162 | 0.209626057 | 6.99E-05 | 3 | 0.000333442 | 14.31119795 | 8 mo | HOM |
| **1013288213** | 2 | LDT | 162 | 0.209626057 | 0 | 0 | 0 | 0 | 4 mo | HOM |
| **1008976060** | 2 | LDT | 162 | 0.209626057 | 1.09E-05 | 2 | 5.2E-05 | 9.540798633 | 8 mo | WILD |
| **1013446243** | 1 | LDT | 162 | 0.209626057 | 0 | 0 | 0 | 0 | 4 mo | HOM |
| **1013288951** | 2 | LDT | 162 | 0.209626057 | 1.95E-07 | 0 | 9.31E-07 | 0 | 4 mo | HOM |
| **1013978665** | 1 | AIp2/3 | 163 | 0.827934106 | 0 | 0 | 0 | 0 | 2 mo | HOM |
| **1013445609** | 1 | AIp2/3 | 163 | 0.827934106 | 9.32E-05 | 8 | 0.000112629 | 9.662604724 | 4 mo | HOM |
| **1013609015** | 1 | AIp2/3 | 163 | 0.827934106 | 0.00017119 | 7 | 0.000206767 | 8.454779134 | 4 mo | HOM |
| **1013608106** | 2 | AIp2/3 | 163 | 0.827934106 | 0 | 0 | 0 | 0 | 4 mo | HOM |
| **1011937543** | 1 | AIp2/3 | 163 | 0.827934106 | 0.000293072 | 10 | 0.00035398 | 12.07825591 | 8 mo | HOM |
| **1010419538** | 2 | AIp2/3 | 163 | 0.827934106 | 0 | 0 | 0 | 0 | 4 mo | WILD |
| **1010419187** | 1 | AIp2/3 | 163 | 0.827934106 | 0.000659697 | 29 | 0.000796799 | 35.02694212 | 8 mo | HOM |
| **1013978366** | 1 | AIp2/3 | 163 | 0.827934106 | 0 | 0 | 0 | 0 | 2 mo | HOM |
| **1014416155** | 2 | AIp2/3 | 163 | 0.827934106 | 0 | 0 | 0 | 0 | 2 mo | HOM |
| **1013978065** | 2 | AIp2/3 | 163 | 0.827934106 | 0 | 0 | 0 | 0 | 2 mo | HOM |
| **1009934102** | 2 | AIp2/3 | 163 | 0.827934106 | 0 | 0 | 0 | 0 | 4 mo | WILD |
| **1009322937** | 2 | AIp2/3 | 163 | 0.827934106 | 0 | 0 | 0 | 0 | 8 mo | WILD |
| **1011080637** | 2 | AIp2/3 | 163 | 0.827934106 | 0.000391746 | 17 | 0.00047316 | 20.53303504 | 8 mo | HOM |
| **1013609713** | 2 | AIp2/3 | 163 | 0.827934106 | 0 | 0 | 0 | 0 | 2 mo | WILD |
| **1010418842** | 1 | AIp2/3 | 163 | 0.827934106 | 0.000539267 | 26 | 0.000651341 | 31.40346535 | 8 mo | HOM |
| **1013288213** | 2 | AIp2/3 | 163 | 0.827934106 | 2.41E-05 | 3 | 2.91E-05 | 3.623476772 | 4 mo | HOM |
| **1008976060** | 2 | AIp2/3 | 163 | 0.827934106 | 0 | 0 | 0 | 0 | 8 mo | WILD |
| **1013446243** | 1 | AIp2/3 | 163 | 0.827934106 | 6.78E-05 | 6 | 8.19E-05 | 7.246953543 | 4 mo | HOM |
| **1013288951** | 2 | AIp2/3 | 163 | 0.827934106 | 0.000143961 | 8 | 0.00017388 | 9.662604724 | 4 mo | HOM |
| **1013978665** | 1 | c | 164 | 0.000445057 | 0 | 0 | 0 | 0 | 2 mo | HOM |
| **1013445609** | 1 | c | 164 | 0.000445057 | 0 | 0 | 0 | 0 | 4 mo | HOM |
| **1013609015** | 1 | c | 164 | 0.000445057 | 0 | 0 | 0 | 0 | 4 mo | HOM |
| **1013608106** | 2 | c | 164 | 0.000445057 | 0 | 0 | 0 | 0 | 4 mo | HOM |
| **1011937543** | 1 | c | 164 | 0.000445057 | 0 | 0 | 0 | 0 | 8 mo | HOM |
| **1010419538** | 2 | c | 164 | 0.000445057 | 0 | 0 | 0 | 0 | 4 mo | WILD |
| **1010419187** | 1 | c | 164 | 0.000445057 | 1.47E-07 | 1 | 0.000329288 | 2246.902196 | 8 mo | HOM |
| **1013978366** | 1 | c | 164 | 0.000445057 | 0 | 0 | 0 | 0 | 2 mo | HOM |
| **1014416155** | 2 | c | 164 | 0.000445057 | 0 | 0 | 0 | 0 | 2 mo | HOM |
| **1013978065** | 2 | c | 164 | 0.000445057 | 0 | 0 | 0 | 0 | 2 mo | HOM |
| **1009934102** | 2 | c | 164 | 0.000445057 | 0 | 0 | 0 | 0 | 4 mo | WILD |
| **1009322937** | 2 | c | 164 | 0.000445057 | 0 | 0 | 0 | 0 | 8 mo | WILD |
| **1011080637** | 2 | c | 164 | 0.000445057 | 3.07E-21 | 0 | 6.89E-18 | 0 | 8 mo | HOM |
| **1013609713** | 2 | c | 164 | 0.000445057 | 0 | 0 | 0 | 0 | 2 mo | WILD |
| **1010418842** | 1 | c | 164 | 0.000445057 | 0 | 0 | 0 | 0 | 8 mo | HOM |
| **1013288213** | 2 | c | 164 | 0.000445057 | 0 | 0 | 0 | 0 | 4 mo | HOM |
| **1008976060** | 2 | c | 164 | 0.000445057 | 0 | 0 | 0 | 0 | 8 mo | WILD |
| **1013446243** | 1 | c | 164 | 0.000445057 | 1.09E-05 | 1 | 0.024380807 | 2246.902196 | 4 mo | HOM |
| **1013288951** | 2 | c | 164 | 0.000445057 | 0 | 0 | 0 | 0 | 4 mo | HOM |
| **1013978665** | 1 | RAmb | 165 | 0.755670045 | 0 | 0 | 0 | 0 | 2 mo | HOM |
| **1013445609** | 1 | RAmb | 165 | 0.755670045 | 0 | 0 | 0 | 0 | 4 mo | HOM |
| **1013609015** | 1 | RAmb | 165 | 0.755670045 | 1.12E-05 | 2 | 1.49E-05 | 2.646657776 | 4 mo | HOM |
| **1013608106** | 2 | RAmb | 165 | 0.755670045 | 0 | 0 | 0 | 0 | 4 mo | HOM |
| **1011937543** | 1 | RAmb | 165 | 0.755670045 | 0.000549443 | 2 | 0.000727094 | 2.646657776 | 8 mo | HOM |
| **1010419538** | 2 | RAmb | 165 | 0.755670045 | 1.83E-07 | 0 | 2.42E-07 | 0 | 4 mo | WILD |
| **1010419187** | 1 | RAmb | 165 | 0.755670045 | 0.000189262 | 4 | 0.000250455 | 5.293315551 | 8 mo | HOM |
| **1013978366** | 1 | RAmb | 165 | 0.755670045 | 0 | 0 | 0 | 0 | 2 mo | HOM |
| **1014416155** | 2 | RAmb | 165 | 0.755670045 | 0 | 0 | 0 | 0 | 2 mo | HOM |
| **1013978065** | 2 | RAmb | 165 | 0.755670045 | 0 | 0 | 0 | 0 | 2 mo | HOM |
| **1009934102** | 2 | RAmb | 165 | 0.755670045 | 1.1E-05 | 1 | 1.45E-05 | 1.323328888 | 4 mo | WILD |

| | | | | | | | | | | |
|---|---|---|---|---|---|---|---|---|---|---|
| 1009322937 | 2 | RAmb | 165 | 0.755670045 | 2.25E-05 | 3 | 2.98E-05 | 3.969986663 | 8 mo | WILD |
| 1011080637 | 2 | RAmb | 165 | 0.755670045 | 6.93E-06 | 1 | 9.17E-06 | 1.323328888 | 8 mo | HOM |
| 1013609713 | 2 | RAmb | 165 | 0.755670045 | 0 | 0 | 0 | 0 | 2 mo | WILD |
| 1010418842 | 1 | RAmb | 165 | 0.755670045 | 7.87E-05 | 9 | 0.000104211 | 11.90995999 | 8 mo | HOM |
| 1013288213 | 2 | RAmb | 165 | 0.755670045 | 0 | 0 | 0 | 0 | 4 mo | HOM |
| 1008976060 | 2 | RAmb | 165 | 0.755670045 | 0.000122125 | 8 | 0.000161612 | 10.5866311 | 8 mo | WILD |
| 1013446243 | 1 | RAmb | 165 | 0.755670045 | 1.11E-06 | 0 | 1.47E-06 | 0 | 4 mo | HOM |
| 1013288951 | 2 | RAmb | 165 | 0.755670045 | 2.41E-05 | 1 | 3.19E-05 | 1.323328888 | 4 mo | HOM |
| 1013978665 | 1 | PRP | 169 | 0.251414041 | 1.24E-06 | 0 | 4.93E-06 | 0 | 2 mo | HOM |
| 1013445609 | 1 | PRP | 169 | 0.251414041 | 0 | 0 | 0 | 0 | 4 mo | HOM |
| 1013609015 | 1 | PRP | 169 | 0.251414041 | 8.19E-05 | 0 | 0.000325769 | 0 | 4 mo | HOM |
| 1013608106 | 2 | PRP | 169 | 0.251414041 | 0 | 0 | 0 | 0 | 4 mo | HOM |
| 1011937543 | 1 | PRP | 169 | 0.251414041 | 0 | 0 | 0 | 0 | 8 mo | HOM |
| 1010419538 | 2 | PRP | 169 | 0.251414041 | 0 | 0 | 0 | 0 | 4 mo | WILD |
| 1010419187 | 1 | PRP | 169 | 0.251414041 | 3.58E-10 | 0 | 1.42E-09 | 0 | 8 mo | HOM |
| 1013978366 | 1 | PRP | 169 | 0.251414041 | 7.67E-08 | 0 | 3.05E-07 | 0 | 2 mo | HOM |
| 1014416155 | 2 | PRP | 169 | 0.251414041 | 0 | 0 | 0 | 0 | 2 mo | HOM |
| 1013978065 | 2 | PRP | 169 | 0.251414041 | 0 | 0 | 0 | 0 | 2 mo | HOM |
| 1009934102 | 2 | PRP | 169 | 0.251414041 | 0 | 0 | 0 | 0 | 4 mo | WILD |
| 1009322937 | 2 | PRP | 169 | 0.251414041 | 0 | 0 | 0 | 0 | 8 mo | WILD |
| 1011080637 | 2 | PRP | 169 | 0.251414041 | 0.000270426 | 2 | 0.00107562 | 7.95500518 | 8 mo | HOM |
| 1013609713 | 2 | PRP | 169 | 0.251414041 | 0 | 0 | 0 | 0 | 2 mo | WILD |
| 1010418842 | 1 | PRP | 169 | 0.251414041 | 2.36E-05 | 3 | 9.37E-05 | 11.93250777 | 8 mo | HOM |
| 1013288213 | 2 | PRP | 169 | 0.251414041 | 0 | 0 | 0 | 0 | 4 mo | HOM |
| 1008976060 | 2 | PRP | 169 | 0.251414041 | 0.000147903 | 5 | 0.000588286 | 19.88751295 | 8 mo | WILD |
| 1013446243 | 1 | PRP | 169 | 0.251414041 | 0 | 0 | 0 | 0 | 4 mo | HOM |
| 1013288951 | 2 | PRP | 169 | 0.251414041 | 6.29E-06 | 0 | 2.5E-05 | 0 | 4 mo | HOM |
| 1013978665 | 1 | LGd | 170 | 0.744158286 | 0 | 0 | 0 | 0 | 2 mo | HOM |
| 1013445609 | 1 | LGd | 170 | 0.744158286 | 2.33E-05 | 2 | 3.13E-05 | 2.687600255 | 4 mo | HOM |
| 1013609015 | 1 | LGd | 170 | 0.744158286 | 0.00018662 | 4 | 0.000250779 | 5.375200511 | 4 mo | HOM |
| 1013608106 | 2 | LGd | 170 | 0.744158286 | 0 | 0 | 0 | 0 | 4 mo | HOM |
| 1011937543 | 1 | LGd | 170 | 0.744158286 | 0.001172056 | 16 | 0.001575009 | 21.50080204 | 8 mo | HOM |
| 1010419538 | 2 | LGd | 170 | 0.744158286 | 0 | 0 | 0 | 0 | 4 mo | WILD |
| 1010419187 | 1 | LGd | 170 | 0.744158286 | 0.000328557 | 11 | 0.000441515 | 14.7818014 | 8 mo | HOM |
| 1013978366 | 1 | LGd | 170 | 0.744158286 | 0 | 0 | 0 | 0 | 2 mo | HOM |
| 1014416155 | 2 | LGd | 170 | 0.744158286 | 0 | 0 | 0 | 0 | 2 mo | HOM |
| 1013978065 | 2 | LGd | 170 | 0.744158286 | 0 | 0 | 0 | 0 | 2 mo | HOM |
| 1009934102 | 2 | LGd | 170 | 0.744158286 | 0 | 0 | 0 | 0 | 4 mo | WILD |
| 1009322937 | 2 | LGd | 170 | 0.744158286 | 1.48E-05 | 3 | 1.99E-05 | 4.031400383 | 8 mo | WILD |
| 1011080637 | 2 | LGd | 170 | 0.744158286 | 0.000390788 | 10 | 0.000525142 | 13.43800128 | 8 mo | HOM |
| 1013609713 | 2 | LGd | 170 | 0.744158286 | 0 | 0 | 0 | 0 | 2 mo | WILD |
| 1010418842 | 1 | LGd | 170 | 0.744158286 | 0.001105244 | 22 | 0.001485228 | 29.56360281 | 8 mo | HOM |
| 1013288213 | 2 | LGd | 170 | 0.744158286 | 0 | 0 | 0 | 0 | 4 mo | HOM |
| 1008976060 | 2 | LGd | 170 | 0.744158286 | 3.78E-05 | 2 | 5.08E-05 | 2.687600255 | 8 mo | WILD |
| 1013446243 | 1 | LGd | 170 | 0.744158286 | 3.87E-05 | 2 | 5.19E-05 | 2.687600255 | 4 mo | HOM |
| 1013288951 | 2 | LGd | 170 | 0.744158286 | 2.02E-05 | 3 | 2.72E-05 | 4.031400383 | 4 mo | HOM |
| 1013978665 | 1 | PL1 | 171 | 0.53966687 | 5.29E-06 | 0 | 9.8E-06 | 0 | 2 mo | HOM |
| 1013445609 | 1 | PL1 | 171 | 0.53966687 | 0 | 0 | 0 | 0 | 4 mo | HOM |
| 1013609015 | 1 | PL1 | 171 | 0.53966687 | 0.000239783 | 7 | 0.000444316 | 12.97096484 | 4 mo | HOM |
| 1013608106 | 2 | PL1 | 171 | 0.53966687 | 0 | 0 | 0 | 0 | 4 mo | HOM |
| 1011937543 | 1 | PL1 | 171 | 0.53966687 | 0.000137661 | 3 | 0.000255086 | 5.55898493 | 8 mo | HOM |
| 1010419538 | 2 | PL1 | 171 | 0.53966687 | 0 | 0 | 0 | 0 | 4 mo | WILD |
| 1010419187 | 1 | PL1 | 171 | 0.53966687 | 0.00123251 | 22 | 0.002283835 | 40.76588949 | 8 mo | HOM |
| 1013978366 | 1 | PL1 | 171 | 0.53966687 | 7.84E-05 | 2 | 0.000145253 | 3.705989954 | 2 mo | HOM |
| 1014416155 | 2 | PL1 | 171 | 0.53966687 | 0 | 0 | 0 | 0 | 2 mo | HOM |
| 1013978065 | 2 | PL1 | 171 | 0.53966687 | 0 | 0 | 0 | 0 | 2 mo | HOM |
| 1009934102 | 2 | PL1 | 171 | 0.53966687 | 0 | 0 | 0 | 0 | 4 mo | WILD |
| 1009322937 | 2 | PL1 | 171 | 0.53966687 | 1.99E-05 | 3 | 3.69E-05 | 5.55898493 | 8 mo | WILD |
| 1011080637 | 2 | PL1 | 171 | 0.53966687 | 0.000524507 | 18 | 0.000971909 | 33.35390958 | 8 mo | HOM |

| | | | | | | | | | | |
|---|---|---|---|---|---|---|---|---|---|---|
| **1013609713** | 2 | PL1 | 171 | 0.53966687 | 1.09E-05 | 1 | 2.01E-05 | 1.852994977 | 2 mo | WILD |
| **1010418842** | 1 | PL1 | 171 | 0.53966687 | 0.001537917 | 20 | 0.002849753 | 37.05989954 | 8 mo | HOM |
| **1013288213** | 2 | PL1 | 171 | 0.53966687 | 0.001663919 | 12 | 0.003083234 | 22.23593972 | 4 mo | HOM |
| **1008976060** | 2 | PL1 | 171 | 0.53966687 | 1.26E-05 | 2 | 2.34E-05 | 3.705989954 | 8 mo | WILD |
| **1013446243** | 1 | PL1 | 171 | 0.53966687 | 2.8E-05 | 3 | 5.19E-05 | 5.55898493 | 4 mo | HOM |
| **1013288951** | 2 | PL1 | 171 | 0.53966687 | 0.000510349 | 14 | 0.000945673 | 25.94192968 | 4 mo | HOM |
| **1013978665** | 1 | RCH | 173 | 0.154873722 | 0 | 0 | 0 | 0 | 2 mo | HOM |
| **1013445609** | 1 | RCH | 173 | 0.154873722 | 0 | 0 | 0 | 0 | 4 mo | HOM |
| **1013609015** | 1 | RCH | 173 | 0.154873722 | 0 | 0 | 0 | 0 | 4 mo | HOM |
| **1013608106** | 2 | RCH | 173 | 0.154873722 | 0 | 0 | 0 | 0 | 4 mo | HOM |
| **1011937543** | 1 | RCH | 173 | 0.154873722 | 0.00119416 | 1 | 0.007710537 | 6.456873305 | 8 mo | HOM |
| **1010419538** | 2 | RCH | 173 | 0.154873722 | 0 | 0 | 0 | 0 | 4 mo | WILD |
| **1010419187** | 1 | RCH | 173 | 0.154873722 | 2.68E-05 | 2 | 0.000172876 | 12.91374661 | 8 mo | HOM |
| **1013978366** | 1 | RCH | 173 | 0.154873722 | 0 | 0 | 0 | 0 | 2 mo | HOM |
| **1014416155** | 2 | RCH | 173 | 0.154873722 | 0 | 0 | 0 | 0 | 2 mo | HOM |
| **1013978065** | 2 | RCH | 173 | 0.154873722 | 0 | 0 | 0 | 0 | 2 mo | HOM |
| **1009934102** | 2 | RCH | 173 | 0.154873722 | 0 | 0 | 0 | 0 | 4 mo | WILD |
| **1009322937** | 2 | RCH | 173 | 0.154873722 | 0 | 0 | 0 | 0 | 8 mo | WILD |
| **1011080637** | 2 | RCH | 173 | 0.154873722 | 2.47E-05 | 3 | 0.000159336 | 19.37061992 | 8 mo | HOM |
| **1013609713** | 2 | RCH | 173 | 0.154873722 | 0 | 0 | 0 | 0 | 2 mo | WILD |
| **1010418842** | 1 | RCH | 173 | 0.154873722 | 7.58E-05 | 3 | 0.000489486 | 19.37061992 | 8 mo | HOM |
| **1013288213** | 2 | RCH | 173 | 0.154873722 | 0 | 0 | 0 | 0 | 4 mo | HOM |
| **1008976060** | 2 | RCH | 173 | 0.154873722 | 0 | 0 | 0 | 0 | 8 mo | WILD |
| **1013446243** | 1 | RCH | 173 | 0.154873722 | 0 | 0 | 0 | 0 | 4 mo | HOM |
| **1013288951** | 2 | RCH | 173 | 0.154873722 | 0 | 0 | 0 | 0 | 4 mo | HOM |
| **1013978665** | 1 | NR | 177 | 0.030898508 | 0 | 0 | 0 | 0 | 2 mo | HOM |
| **1013445609** | 1 | NR | 177 | 0.030898508 | 0 | 0 | 0 | 0 | 4 mo | HOM |
| **1013609015** | 1 | NR | 177 | 0.030898508 | 0 | 0 | 0 | 0 | 4 mo | HOM |
| **1013608106** | 2 | NR | 177 | 0.030898508 | 0 | 0 | 0 | 0 | 4 mo | HOM |
| **1011937543** | 1 | NR | 177 | 0.030898508 | 0 | 0 | 0 | 0 | 8 mo | HOM |
| **1010419538** | 2 | NR | 177 | 0.030898508 | 0 | 0 | 0 | 0 | 4 mo | WILD |
| **1010419187** | 1 | NR | 177 | 0.030898508 | 0 | 0 | 0 | 0 | 8 mo | HOM |
| **1013978366** | 1 | NR | 177 | 0.030898508 | 0 | 0 | 0 | 0 | 2 mo | HOM |
| **1014416155** | 2 | NR | 177 | 0.030898508 | 0 | 0 | 0 | 0 | 2 mo | HOM |
| **1013978065** | 2 | NR | 177 | 0.030898508 | 0 | 0 | 0 | 0 | 2 mo | HOM |
| **1009934102** | 2 | NR | 177 | 0.030898508 | 0 | 0 | 0 | 0 | 4 mo | WILD |
| **1009322937** | 2 | NR | 177 | 0.030898508 | 0 | 0 | 0 | 0 | 8 mo | WILD |
| **1011080637** | 2 | NR | 177 | 0.030898508 | 0 | 0 | 0 | 0 | 8 mo | HOM |
| **1013609713** | 2 | NR | 177 | 0.030898508 | 0 | 0 | 0 | 0 | 2 mo | WILD |
| **1010418842** | 1 | NR | 177 | 0.030898508 | 7.78E-06 | 1 | 0.000251697 | 32.36402203 | 8 mo | HOM |
| **1013288213** | 2 | NR | 177 | 0.030898508 | 0 | 0 | 0 | 0 | 4 mo | HOM |
| **1008976060** | 2 | NR | 177 | 0.030898508 | 8.23E-06 | 1 | 0.000266347 | 32.36402203 | 8 mo | WILD |
| **1013446243** | 1 | NR | 177 | 0.030898508 | 0 | 0 | 0 | 0 | 4 mo | HOM |
| **1013288951** | 2 | NR | 177 | 0.030898508 | 0 | 0 | 0 | 0 | 4 mo | HOM |
| **1013978665** | 1 | LGv | 178 | 0.418005554 | 0 | 0 | 0 | 0 | 2 mo | HOM |
| **1013445609** | 1 | LGv | 178 | 0.418005554 | 2.82E-06 | 1 | 6.75E-06 | 2.392312708 | 4 mo | HOM |
| **1013609015** | 1 | LGv | 178 | 0.418005554 | 2.48E-05 | 1 | 5.94E-05 | 2.392312708 | 4 mo | HOM |
| **1013608106** | 2 | LGv | 178 | 0.418005554 | 0 | 0 | 0 | 0 | 4 mo | HOM |
| **1011937543** | 1 | LGv | 178 | 0.418005554 | 0.000985473 | 5 | 0.00235756 | 11.96156354 | 8 mo | HOM |
| **1010419538** | 2 | LGv | 178 | 0.418005554 | 0 | 0 | 0 | 0 | 4 mo | WILD |
| **1010419187** | 1 | LGv | 178 | 0.418005554 | 0.000194131 | 11 | 0.000464423 | 26.31543979 | 8 mo | HOM |
| **1013978366** | 1 | LGv | 178 | 0.418005554 | 0 | 0 | 0 | 0 | 2 mo | HOM |
| **1014416155** | 2 | LGv | 178 | 0.418005554 | 0 | 0 | 0 | 0 | 2 mo | HOM |
| **1013978065** | 2 | LGv | 178 | 0.418005554 | 0 | 0 | 0 | 0 | 2 mo | HOM |
| **1009934102** | 2 | LGv | 178 | 0.418005554 | 0 | 0 | 0 | 0 | 4 mo | WILD |
| **1009322937** | 2 | LGv | 178 | 0.418005554 | 0 | 0 | 0 | 0 | 8 mo | WILD |
| **1011080637** | 2 | LGv | 178 | 0.418005554 | 7.82E-05 | 4 | 0.000187025 | 9.569250834 | 8 mo | HOM |
| **1013609713** | 2 | LGv | 178 | 0.418005554 | 0 | 0 | 0 | 0 | 2 mo | WILD |
| **1010418842** | 1 | LGv | 178 | 0.418005554 | 0.000883275 | 17 | 0.00211307 | 40.66931604 | 8 mo | HOM |

| | | | | | | | | | | |
|---|---|---|---|---|---|---|---|---|---|---|
| 1013288213 | 2 | LGv | 178 | 0.418005554 | 0 | 0 | 0 | 0 | 4 mo | HOM |
| 1008976060 | 2 | LGv | 178 | 0.418005554 | 3.92E-05 | 2 | 9.38E-05 | 4.784625417 | 8 mo | WILD |
| 1013446243 | 1 | LGv | 178 | 0.418005554 | 0 | 0 | 0 | 0 | 4 mo | HOM |
| 1013288951 | 2 | LGv | 178 | 0.418005554 | 0 | 0 | 0 | 0 | 4 mo | HOM |
| 1013978665 | 1 | GU2/3 | 180 | 0.414096883 | 8.49E-07 | 0 | 2.05E-06 | 0 | 2 mo | HOM |
| 1013445609 | 1 | GU2/3 | 180 | 0.414096883 | 3.49E-05 | 1 | 8.42E-05 | 2.41489381 | 4 mo | HOM |
| 1013609015 | 1 | GU2/3 | 180 | 0.414096883 | 0.000167801 | 4 | 0.000405222 | 9.65957524 | 4 mo | HOM |
| 1013608106 | 2 | GU2/3 | 180 | 0.414096883 | 0 | 0 | 0 | 0 | 4 mo | HOM |
| 1011937543 | 1 | GU2/3 | 180 | 0.414096883 | 0.000314761 | 9 | 0.000760114 | 21.73404429 | 8 mo | HOM |
| 1010419538 | 2 | GU2/3 | 180 | 0.414096883 | 0 | 0 | 0 | 0 | 4 mo | WILD |
| 1010419187 | 1 | GU2/3 | 180 | 0.414096883 | 0.000254793 | 13 | 0.000615297 | 31.39361953 | 8 mo | HOM |
| 1013978366 | 1 | GU2/3 | 180 | 0.414096883 | 0 | 0 | 0 | 0 | 2 mo | HOM |
| 1014416155 | 2 | GU2/3 | 180 | 0.414096883 | 0 | 0 | 0 | 0 | 2 mo | HOM |
| 1013978065 | 2 | GU2/3 | 180 | 0.414096883 | 0 | 0 | 0 | 0 | 2 mo | HOM |
| 1009934102 | 2 | GU2/3 | 180 | 0.414096883 | 0 | 0 | 0 | 0 | 4 mo | WILD |
| 1009322937 | 2 | GU2/3 | 180 | 0.414096883 | 0 | 0 | 0 | 0 | 8 mo | WILD |
| 1011080637 | 2 | GU2/3 | 180 | 0.414096883 | 0.000246376 | 16 | 0.000594973 | 38.63830096 | 8 mo | HOM |
| 1013609713 | 2 | GU2/3 | 180 | 0.414096883 | 0 | 0 | 0 | 0 | 2 mo | WILD |
| 1010418842 | 1 | GU2/3 | 180 | 0.414096883 | 0.000280222 | 10 | 0.000676705 | 24.1489381 | 8 mo | HOM |
| 1013288213 | 2 | GU2/3 | 180 | 0.414096883 | 2.23E-06 | 0 | 5.38E-06 | 0 | 4 mo | HOM |
| 1008976060 | 2 | GU2/3 | 180 | 0.414096883 | 7.27E-07 | 0 | 1.75E-06 | 0 | 8 mo | WILD |
| 1013446243 | 1 | GU2/3 | 180 | 0.414096883 | 2.12E-05 | 1 | 5.11E-05 | 2.41489381 | 4 mo | HOM |
| 1013288951 | 2 | GU2/3 | 180 | 0.414096883 | 5.35E-05 | 4 | 0.000129171 | 9.65957524 | 4 mo | HOM |
| 1013978665 | 1 | RE | 181 | 0.436768007 | 0 | 0 | 0 | 0 | 2 mo | HOM |
| 1013445609 | 1 | RE | 181 | 0.436768007 | 0 | 0 | 0 | 0 | 4 mo | HOM |
| 1013609015 | 1 | RE | 181 | 0.436768007 | 0 | 0 | 0 | 0 | 4 mo | HOM |
| 1013608106 | 2 | RE | 181 | 0.436768007 | 0 | 0 | 0 | 0 | 4 mo | HOM |
| 1011937543 | 1 | RE | 181 | 0.436768007 | 0.000123879 | 2 | 0.000283626 | 4.579089968 | 8 mo | HOM |
| 1010419538 | 2 | RE | 181 | 0.436768007 | 0 | 0 | 0 | 0 | 4 mo | WILD |
| 1010419187 | 1 | RE | 181 | 0.436768007 | 0.000154773 | 5 | 0.000354359 | 11.44772492 | 8 mo | HOM |
| 1013978366 | 1 | RE | 181 | 0.436768007 | 0 | 0 | 0 | 0 | 2 mo | HOM |
| 1014416155 | 2 | RE | 181 | 0.436768007 | 0 | 0 | 0 | 0 | 2 mo | HOM |
| 1013978065 | 2 | RE | 181 | 0.436768007 | 0 | 0 | 0 | 0 | 2 mo | HOM |
| 1009934102 | 2 | RE | 181 | 0.436768007 | 0 | 0 | 0 | 0 | 4 mo | WILD |
| 1009322937 | 2 | RE | 181 | 0.436768007 | 7.28E-07 | 0 | 1.67E-06 | 0 | 8 mo | WILD |
| 1011080637 | 2 | RE | 181 | 0.436768007 | 8.3E-05 | 4 | 0.000190085 | 9.158179935 | 8 mo | HOM |
| 1013609713 | 2 | RE | 181 | 0.436768007 | 0 | 0 | 0 | 0 | 2 mo | WILD |
| 1010418842 | 1 | RE | 181 | 0.436768007 | 0.000180837 | 4 | 0.000414035 | 9.158179935 | 8 mo | HOM |
| 1013288213 | 2 | RE | 181 | 0.436768007 | 0 | 0 | 0 | 0 | 4 mo | HOM |
| 1008976060 | 2 | RE | 181 | 0.436768007 | 0 | 0 | 0 | 0 | 8 mo | WILD |
| 1013446243 | 1 | RE | 181 | 0.436768007 | 0 | 0 | 0 | 0 | 4 mo | HOM |
| 1013288951 | 2 | RE | 181 | 0.436768007 | 0 | 0 | 0 | 0 | 4 mo | HOM |
| 1013978665 | 1 | FRP | 184 | 1.003311395 | 0 | 0 | 0 | 0 | 2 mo | HOM |
| 1013445609 | 1 | FRP | 184 | 1.003311395 | 0 | 0 | 0 | 0 | 4 mo | HOM |
| 1013609015 | 1 | FRP | 184 | 1.003311395 | 5.97E-05 | 2 | 5.95E-05 | 1.993399069 | 4 mo | HOM |
| 1013608106 | 2 | FRP | 184 | 1.003311395 | 0 | 0 | 0 | 0 | 4 mo | HOM |
| 1011937543 | 1 | FRP | 184 | 1.003311395 | 0.000557795 | 14 | 0.000555954 | 13.95379348 | 8 mo | HOM |
| 1010419538 | 2 | FRP | 184 | 1.003311395 | 0 | 0 | 0 | 0 | 4 mo | WILD |
| 1010419187 | 1 | FRP | 184 | 1.003311395 | 0.000775698 | 31 | 0.000773138 | 30.89768557 | 8 mo | HOM |
| 1013978366 | 1 | FRP | 184 | 1.003311395 | 1.04E-05 | 0 | 1.03E-05 | 0 | 2 mo | HOM |
| 1014416155 | 2 | FRP | 184 | 1.003311395 | 0 | 0 | 0 | 0 | 2 mo | HOM |
| 1013978065 | 2 | FRP | 184 | 1.003311395 | 0 | 0 | 0 | 0 | 2 mo | HOM |
| 1009934102 | 2 | FRP | 184 | 1.003311395 | 0 | 0 | 0 | 0 | 4 mo | WILD |
| 1009322937 | 2 | FRP | 184 | 1.003311395 | 2.42E-05 | 3 | 2.41E-05 | 2.990098603 | 8 mo | WILD |
| 1011080637 | 2 | FRP | 184 | 1.003311395 | 0.000211561 | 7 | 0.000210863 | 6.976896741 | 8 mo | HOM |
| 1013609713 | 2 | FRP | 184 | 1.003311395 | 0 | 0 | 0 | 0 | 2 mo | WILD |
| 1010418842 | 1 | FRP | 184 | 1.003311395 | 0.000446327 | 25 | 0.000444854 | 24.91748836 | 8 mo | HOM |
| 1013288213 | 2 | FRP | 184 | 1.003311395 | 0.000869782 | 6 | 0.000866911 | 5.980197206 | 4 mo | HOM |
| 1008976060 | 2 | FRP | 184 | 1.003311395 | 2.17E-05 | 2 | 2.16E-05 | 1.993399069 | 8 mo | WILD |

| | | | | | | | | | | |
|---|---|---|---|---|---|---|---|---|---|---|
| 1013446243 | 1 | FRP | 184 | 1.003311395 | 5.94E-05 | 4 | 5.92E-05 | 3.986798138 | 4 mo | HOM |
| 1013288951 | 2 | FRP | 184 | 1.003311395 | 0.000144501 | 11 | 0.000144024 | 10.96369488 | 4 mo | HOM |
| 1013978665 | 1 | LH | 186 | 0.355137104 | 0 | 0 | 0 | 0 | 2 mo | HOM |
| 1013445609 | 1 | LH | 186 | 0.355137104 | 0 | 0 | 0 | 0 | 4 mo | HOM |
| 1013609015 | 1 | LH | 186 | 0.355137104 | 5.08E-07 | 0 | 1.43E-06 | 0 | 4 mo | HOM |
| 1013608106 | 2 | LH | 186 | 0.355137104 | 0 | 0 | 0 | 0 | 4 mo | HOM |
| 1011937543 | 1 | LH | 186 | 0.355137104 | 0.001007591 | 7 | 0.002837189 | 19.71069742 | 8 mo | HOM |
| 1010419538 | 2 | LH | 186 | 0.355137104 | 2.14E-06 | 0 | 6.02E-06 | 0 | 4 mo | WILD |
| 1010419187 | 1 | LH | 186 | 0.355137104 | 1.27E-20 | 0 | 3.59E-20 | 0 | 8 mo | HOM |
| 1013978366 | 1 | LH | 186 | 0.355137104 | 0 | 0 | 0 | 0 | 2 mo | HOM |
| 1014416155 | 2 | LH | 186 | 0.355137104 | 0 | 0 | 0 | 0 | 2 mo | HOM |
| 1013978065 | 2 | LH | 186 | 0.355137104 | 0 | 0 | 0 | 0 | 2 mo | HOM |
| 1009934102 | 2 | LH | 186 | 0.355137104 | 0 | 0 | 0 | 0 | 4 mo | WILD |
| 1009322937 | 2 | LH | 186 | 0.355137104 | 1.07E-07 | 0 | 3E-07 | 0 | 8 mo | WILD |
| 1011080637 | 2 | LH | 186 | 0.355137104 | 9.2E-05 | 3 | 0.00025911 | 8.447441752 | 8 mo | HOM |
| 1013609713 | 2 | LH | 186 | 0.355137104 | 0 | 0 | 0 | 0 | 2 mo | WILD |
| 1010418842 | 1 | LH | 186 | 0.355137104 | 0 | 0 | 0 | 0 | 8 mo | HOM |
| 1013288213 | 2 | LH | 186 | 0.355137104 | 0 | 0 | 0 | 0 | 4 mo | HOM |
| 1008976060 | 2 | LH | 186 | 0.355137104 | 0 | 0 | 0 | 0 | 8 mo | WILD |
| 1013446243 | 1 | LH | 186 | 0.355137104 | 7.4E-06 | 1 | 2.08E-05 | 2.815813917 | 4 mo | HOM |
| 1013288951 | 2 | LH | 186 | 0.355137104 | 5.58E-05 | 2 | 0.000157256 | 5.631627835 | 4 mo | HOM |
| 1013978665 | 1 | GU5 | 187 | 0.561800249 | 0 | 0 | 0 | 0 | 2 mo | HOM |
| 1013445609 | 1 | GU5 | 187 | 0.561800249 | 2.12E-05 | 2 | 3.78E-05 | 3.559984183 | 4 mo | HOM |
| 1013609015 | 1 | GU5 | 187 | 0.561800249 | 8.48E-05 | 3 | 0.000150874 | 5.339976275 | 4 mo | HOM |
| 1013608106 | 2 | GU5 | 187 | 0.561800249 | 0 | 0 | 0 | 0 | 4 mo | HOM |
| 1011937543 | 1 | GU5 | 187 | 0.561800249 | 0.000850801 | 19 | 0.001514419 | 33.81984974 | 8 mo | HOM |
| 1010419538 | 2 | GU5 | 187 | 0.561800249 | 0 | 0 | 0 | 0 | 4 mo | WILD |
| 1010419187 | 1 | GU5 | 187 | 0.561800249 | 0.000681245 | 27 | 0.001212611 | 48.05978648 | 8 mo | HOM |
| 1013978366 | 1 | GU5 | 187 | 0.561800249 | 8.24E-21 | 0 | 1.47E-20 | 0 | 2 mo | HOM |
| 1014416155 | 2 | GU5 | 187 | 0.561800249 | 0 | 0 | 0 | 0 | 2 mo | HOM |
| 1013978065 | 2 | GU5 | 187 | 0.561800249 | 0 | 0 | 0 | 0 | 2 mo | HOM |
| 1009934102 | 2 | GU5 | 187 | 0.561800249 | 0 | 0 | 0 | 0 | 4 mo | WILD |
| 1009322937 | 2 | GU5 | 187 | 0.561800249 | 1.12E-06 | 0 | 2E-06 | 0 | 8 mo | WILD |
| 1011080637 | 2 | GU5 | 187 | 0.561800249 | 0.000328015 | 26 | 0.000583864 | 46.27979439 | 8 mo | HOM |
| 1013609713 | 2 | GU5 | 187 | 0.561800249 | 0 | 0 | 0 | 0 | 2 mo | WILD |
| 1010418842 | 1 | GU5 | 187 | 0.561800249 | 0.000473445 | 28 | 0.000842728 | 49.83977857 | 8 mo | HOM |
| 1013288213 | 2 | GU5 | 187 | 0.561800249 | 2.44E-05 | 3 | 4.35E-05 | 5.339976275 | 4 mo | HOM |
| 1008976060 | 2 | GU5 | 187 | 0.561800249 | 0 | 0 | 0 | 0 | 8 mo | WILD |
| 1013446243 | 1 | GU5 | 187 | 0.561800249 | 0.000106402 | 4 | 0.000189395 | 7.119968367 | 4 mo | HOM |
| 1013288951 | 2 | GU5 | 187 | 0.561800249 | 5.23E-05 | 5 | 9.31E-05 | 8.899960459 | 4 mo | HOM |
| 1013978665 | 1 | AOBgl | 188 | 0.159122375 | 0 | 0 | 0 | 0 | 2 mo | HOM |
| 1013445609 | 1 | AOBgl | 188 | 0.159122375 | 0 | 0 | 0 | 0 | 4 mo | HOM |
| 1013609015 | 1 | AOBgl | 188 | 0.159122375 | 1.48E-06 | 0 | 9.28E-06 | 0 | 4 mo | HOM |
| 1013608106 | 2 | AOBgl | 188 | 0.159122375 | 0 | 0 | 0 | 0 | 4 mo | HOM |
| 1011937543 | 1 | AOBgl | 188 | 0.159122375 | 3.2E-05 | 0 | 0.000201283 | 0 | 8 mo | HOM |
| 1010419538 | 2 | AOBgl | 188 | 0.159122375 | 0 | 0 | 0 | 0 | 4 mo | WILD |
| 1010419187 | 1 | AOBgl | 188 | 0.159122375 | 0.000419638 | 14 | 0.002637202 | 87.98259852 | 8 mo | HOM |
| 1013978366 | 1 | AOBgl | 188 | 0.159122375 | 0 | 0 | 0 | 0 | 2 mo | HOM |
| 1014416155 | 2 | AOBgl | 188 | 0.159122375 | 0 | 0 | 0 | 0 | 2 mo | HOM |
| 1013978065 | 2 | AOBgl | 188 | 0.159122375 | 0 | 0 | 0 | 0 | 2 mo | HOM |
| 1009934102 | 2 | AOBgl | 188 | 0.159122375 | 1.67E-06 | 0 | 1.05E-05 | 0 | 4 mo | WILD |
| 1009322937 | 2 | AOBgl | 188 | 0.159122375 | 9.96E-06 | 2 | 6.26E-05 | 12.56894265 | 8 mo | WILD |
| 1011080637 | 2 | AOBgl | 188 | 0.159122375 | 2.4E-05 | 3 | 0.00015058 | 18.85341397 | 8 mo | HOM |
| 1013609713 | 2 | AOBgl | 188 | 0.159122375 | 4.37E-06 | 0 | 2.75E-05 | 0 | 2 mo | WILD |
| 1010418842 | 1 | AOBgl | 188 | 0.159122375 | 2.6E-05 | 4 | 0.000163666 | 25.13788529 | 8 mo | HOM |
| 1013288213 | 2 | AOBgl | 188 | 0.159122375 | 0.000827291 | 5 | 0.005199087 | 31.42235661 | 4 mo | HOM |
| 1008976060 | 2 | AOBgl | 188 | 0.159122375 | 0 | 0 | 0 | 0 | 8 mo | WILD |
| 1013446243 | 1 | AOBgl | 188 | 0.159122375 | 9.68E-06 | 1 | 6.09E-05 | 6.284471323 | 4 mo | HOM |
| 1013288951 | 2 | AOBgl | 188 | 0.159122375 | 0.000227038 | 10 | 0.001426815 | 62.84471323 | 4 mo | HOM |

| | | | | | | | | | | |
|---|---|---|---|---|---|---|---|---|---|---|
| 1013978665 | 1 | RH | 189 | 0.093435229 | 0 | 0 | 0 | 0 | 2 mo | HOM |
| 1013445609 | 1 | RH | 189 | 0.093435229 | 0 | 0 | 0 | 0 | 4 mo | HOM |
| 1013609015 | 1 | RH | 189 | 0.093435229 | 0 | 0 | 0 | 0 | 4 mo | HOM |
| 1013608106 | 2 | RH | 189 | 0.093435229 | 0 | 0 | 0 | 0 | 4 mo | HOM |
| 1011937543 | 1 | RH | 189 | 0.093435229 | 3.61E-06 | 0 | 3.86E-05 | 0 | 8 mo | HOM |
| 1010419538 | 2 | RH | 189 | 0.093435229 | 0 | 0 | 0 | 0 | 4 mo | WILD |
| 1010419187 | 1 | RH | 189 | 0.093435229 | 5.5E-06 | 1 | 5.88E-05 | 10.70260131 | 8 mo | HOM |
| 1013978366 | 1 | RH | 189 | 0.093435229 | 0 | 0 | 0 | 0 | 2 mo | HOM |
| 1014416155 | 2 | RH | 189 | 0.093435229 | 0 | 0 | 0 | 0 | 2 mo | HOM |
| 1013978065 | 2 | RH | 189 | 0.093435229 | 0 | 0 | 0 | 0 | 2 mo | HOM |
| 1009934102 | 2 | RH | 189 | 0.093435229 | 0 | 0 | 0 | 0 | 4 mo | WILD |
| 1009322937 | 2 | RH | 189 | 0.093435229 | 0 | 0 | 0 | 0 | 8 mo | WILD |
| 1011080637 | 2 | RH | 189 | 0.093435229 | 2.21E-05 | 2 | 0.000236771 | 21.40520262 | 8 mo | HOM |
| 1013609713 | 2 | RH | 189 | 0.093435229 | 0 | 0 | 0 | 0 | 2 mo | WILD |
| 1010418842 | 1 | RH | 189 | 0.093435229 | 4.31E-06 | 0 | 4.61E-05 | 0 | 8 mo | HOM |
| 1013288213 | 2 | RH | 189 | 0.093435229 | 0 | 0 | 0 | 0 | 4 mo | HOM |
| 1008976060 | 2 | RH | 189 | 0.093435229 | 0 | 0 | 0 | 0 | 8 mo | WILD |
| 1013446243 | 1 | RH | 189 | 0.093435229 | 0 | 0 | 0 | 0 | 4 mo | HOM |
| 1013288951 | 2 | RH | 189 | 0.093435229 | 0 | 0 | 0 | 0 | 4 mo | HOM |
| 1013978665 | 1 | py | 190 | 0.577678052 | 0 | 0 | 0 | 0 | 2 mo | HOM |
| 1013445609 | 1 | py | 190 | 0.577678052 | 0 | 0 | 0 | 0 | 4 mo | HOM |
| 1013609015 | 1 | py | 190 | 0.577678052 | 0 | 0 | 0 | 0 | 4 mo | HOM |
| 1013608106 | 2 | py | 190 | 0.577678052 | 0 | 0 | 0 | 0 | 4 mo | HOM |
| 1011937543 | 1 | py | 190 | 0.577678052 | 1.95E-05 | 1 | 3.38E-05 | 1.731068017 | 8 mo | HOM |
| 1010419538 | 2 | py | 190 | 0.577678052 | 0 | 0 | 0 | 0 | 4 mo | WILD |
| 1010419187 | 1 | py | 190 | 0.577678052 | 0 | 0 | 0 | 0 | 8 mo | HOM |
| 1013978366 | 1 | py | 190 | 0.577678052 | 0 | 0 | 0 | 0 | 2 mo | HOM |
| 1014416155 | 2 | py | 190 | 0.577678052 | 0 | 0 | 0 | 0 | 2 mo | HOM |
| 1013978065 | 2 | py | 190 | 0.577678052 | 0 | 0 | 0 | 0 | 2 mo | HOM |
| 1009934102 | 2 | py | 190 | 0.577678052 | 0 | 0 | 0 | 0 | 4 mo | WILD |
| 1009322937 | 2 | py | 190 | 0.577678052 | 0 | 0 | 0 | 0 | 8 mo | WILD |
| 1011080637 | 2 | py | 190 | 0.577678052 | 0 | 0 | 0 | 0 | 8 mo | HOM |
| 1013609713 | 2 | py | 190 | 0.577678052 | 0 | 0 | 0 | 0 | 2 mo | WILD |
| 1010418842 | 1 | py | 190 | 0.577678052 | 0 | 0 | 0 | 0 | 8 mo | HOM |
| 1013288213 | 2 | py | 190 | 0.577678052 | 0 | 0 | 0 | 0 | 4 mo | HOM |
| 1008976060 | 2 | py | 190 | 0.577678052 | 0 | 0 | 0 | 0 | 8 mo | WILD |
| 1013446243 | 1 | py | 190 | 0.577678052 | 0 | 0 | 0 | 0 | 4 mo | HOM |
| 1013288951 | 2 | py | 190 | 0.577678052 | 0 | 0 | 0 | 0 | 4 mo | HOM |
| 1013978665 | 1 | LHA | 194 | 2.126907798 | 0 | 0 | 0 | 0 | 2 mo | HOM |
| 1013445609 | 1 | LHA | 194 | 2.126907798 | 2.62E-06 | 0 | 1.23E-06 | 0 | 4 mo | HOM |
| 1013609015 | 1 | LHA | 194 | 2.126907798 | 1E-09 | 0 | 4.71E-10 | 0 | 4 mo | HOM |
| 1013608106 | 2 | LHA | 194 | 2.126907798 | 0 | 0 | 0 | 0 | 4 mo | HOM |
| 1011937543 | 1 | LHA | 194 | 2.126907798 | 0.002809572 | 56 | 0.001320965 | 26.32930306 | 8 mo | HOM |
| 1010419538 | 2 | LHA | 194 | 2.126907798 | 0 | 0 | 0 | 0 | 4 mo | WILD |
| 1010419187 | 1 | LHA | 194 | 2.126907798 | 2.08E-05 | 1 | 9.77E-06 | 0.470166126 | 8 mo | HOM |
| 1013978366 | 1 | LHA | 194 | 2.126907798 | 7.38E-08 | 0 | 3.47E-08 | 0 | 2 mo | HOM |
| 1014416155 | 2 | LHA | 194 | 2.126907798 | 0 | 0 | 0 | 0 | 2 mo | HOM |
| 1013978065 | 2 | LHA | 194 | 2.126907798 | 0 | 0 | 0 | 0 | 2 mo | HOM |
| 1009934102 | 2 | LHA | 194 | 2.126907798 | 0 | 0 | 0 | 0 | 4 mo | WILD |
| 1009322937 | 2 | LHA | 194 | 2.126907798 | 0 | 0 | 0 | 0 | 8 mo | WILD |
| 1011080637 | 2 | LHA | 194 | 2.126907798 | 8.65E-05 | 6 | 4.06E-05 | 2.820996756 | 8 mo | HOM |
| 1013609713 | 2 | LHA | 194 | 2.126907798 | 0 | 0 | 0 | 0 | 2 mo | WILD |
| 1010418842 | 1 | LHA | 194 | 2.126907798 | 0.000234332 | 14 | 0.000110175 | 6.582325764 | 8 mo | HOM |
| 1013288213 | 2 | LHA | 194 | 2.126907798 | 0 | 0 | 0 | 0 | 4 mo | HOM |
| 1008976060 | 2 | LHA | 194 | 2.126907798 | 0 | 0 | 0 | 0 | 8 mo | WILD |
| 1013446243 | 1 | LHA | 194 | 2.126907798 | 0 | 0 | 0 | 0 | 4 mo | HOM |
| 1013288951 | 2 | LHA | 194 | 2.126907798 | 0 | 0 | 0 | 0 | 4 mo | HOM |
| 1013978665 | 1 | AOBgr | 196 | 0.226946205 | 0 | 0 | 0 | 0 | 2 mo | HOM |
| 1013445609 | 1 | AOBgr | 196 | 0.226946205 | 0 | 0 | 0 | 0 | 4 mo | HOM |

| | | | | | | | | | | |
|---|---|---|---|---|---|---|---|---|---|---|
| **1013609015** | 1 | AOBgr | 196 | 0.226946205 | 0 | 0 | 0 | 0 | 4 mo | HOM |
| **1013608106** | 2 | AOBgr | 196 | 0.226946205 | 0 | 0 | 0 | 0 | 4 mo | HOM |
| **1011937543** | 1 | AOBgr | 196 | 0.226946205 | 0.000867509 | 2 | 0.003822532 | 8.81266114 | 8 mo | HOM |
| **1010419538** | 2 | AOBgr | 196 | 0.226946205 | 0 | 0 | 0 | 0 | 4 mo | WILD |
| **1010419187** | 1 | AOBgr | 196 | 0.226946205 | 3.76E-06 | 1 | 1.66E-05 | 4.40633057 | 8 mo | HOM |
| **1013978366** | 1 | AOBgr | 196 | 0.226946205 | 0 | 0 | 0 | 0 | 2 mo | HOM |
| **1014416155** | 2 | AOBgr | 196 | 0.226946205 | 0 | 0 | 0 | 0 | 2 mo | HOM |
| **1013978065** | 2 | AOBgr | 196 | 0.226946205 | 0 | 0 | 0 | 0 | 2 mo | HOM |
| **1009934102** | 2 | AOBgr | 196 | 0.226946205 | 0 | 0 | 0 | 0 | 4 mo | WILD |
| **1009322937** | 2 | AOBgr | 196 | 0.226946205 | 0 | 0 | 0 | 0 | 8 mo | WILD |
| **1011080637** | 2 | AOBgr | 196 | 0.226946205 | 7.79E-06 | 1 | 3.43E-05 | 4.40633057 | 8 mo | HOM |
| **1013609713** | 2 | AOBgr | 196 | 0.226946205 | 0 | 0 | 0 | 0 | 2 mo | WILD |
| **1010418842** | 1 | AOBgr | 196 | 0.226946205 | 5.2E-05 | 4 | 0.000229225 | 17.62532228 | 8 mo | HOM |
| **1013288213** | 2 | AOBgr | 196 | 0.226946205 | 1.62E-07 | 0 | 7.16E-07 | 0 | 4 mo | HOM |
| **1008976060** | 2 | AOBgr | 196 | 0.226946205 | 4.89E-08 | 0 | 2.16E-07 | 0 | 8 mo | WILD |
| **1013446243** | 1 | AOBgr | 196 | 0.226946205 | 0 | 0 | 0 | 0 | 4 mo | HOM |
| **1013288951** | 2 | AOBgr | 196 | 0.226946205 | 0 | 0 | 0 | 0 | 4 mo | HOM |
| **1013978665** | 1 | RL | 197 | 0.057346444 | 0 | 0 | 0 | 0 | 2 mo | HOM |
| **1013445609** | 1 | RL | 197 | 0.057346444 | 0 | 0 | 0 | 0 | 4 mo | HOM |
| **1013609015** | 1 | RL | 197 | 0.057346444 | 0 | 0 | 0 | 0 | 4 mo | HOM |
| **1013608106** | 2 | RL | 197 | 0.057346444 | 0 | 0 | 0 | 0 | 4 mo | HOM |
| **1011937543** | 1 | RL | 197 | 0.057346444 | 0 | 0 | 0 | 0 | 8 mo | HOM |
| **1010419538** | 2 | RL | 197 | 0.057346444 | 0 | 0 | 0 | 0 | 4 mo | WILD |
| **1010419187** | 1 | RL | 197 | 0.057346444 | 0 | 0 | 0 | 0 | 8 mo | HOM |
| **1013978366** | 1 | RL | 197 | 0.057346444 | 0 | 0 | 0 | 0 | 2 mo | HOM |
| **1014416155** | 2 | RL | 197 | 0.057346444 | 0 | 0 | 0 | 0 | 2 mo | HOM |
| **1013978065** | 2 | RL | 197 | 0.057346444 | 0 | 0 | 0 | 0 | 2 mo | HOM |
| **1009934102** | 2 | RL | 197 | 0.057346444 | 0 | 0 | 0 | 0 | 4 mo | WILD |
| **1009322937** | 2 | RL | 197 | 0.057346444 | 0 | 0 | 0 | 0 | 8 mo | WILD |
| **1011080637** | 2 | RL | 197 | 0.057346444 | 0 | 0 | 0 | 0 | 8 mo | HOM |
| **1013609713** | 2 | RL | 197 | 0.057346444 | 0 | 0 | 0 | 0 | 2 mo | WILD |
| **1010418842** | 1 | RL | 197 | 0.057346444 | 6.47E-06 | 1 | 0.000112896 | 17.43787275 | 8 mo | HOM |
| **1013288213** | 2 | RL | 197 | 0.057346444 | 0 | 0 | 0 | 0 | 4 mo | HOM |
| **1008976060** | 2 | RL | 197 | 0.057346444 | 0 | 0 | 0 | 0 | 8 mo | WILD |
| **1013446243** | 1 | RL | 197 | 0.057346444 | 0 | 0 | 0 | 0 | 4 mo | HOM |
| **1013288951** | 2 | RL | 197 | 0.057346444 | 0 | 0 | 0 | 0 | 4 mo | HOM |
| **1013978665** | 1 | pyd | 198 | 0.069385656 | 0 | 0 | 0 | 0 | 2 mo | HOM |
| **1013445609** | 1 | pyd | 198 | 0.069385656 | 0 | 0 | 0 | 0 | 4 mo | HOM |
| **1013609015** | 1 | pyd | 198 | 0.069385656 | 0 | 0 | 0 | 0 | 4 mo | HOM |
| **1013608106** | 2 | pyd | 198 | 0.069369775 | 0 | 0 | 0 | 0 | 4 mo | HOM |
| **1011937543** | 1 | pyd | 198 | 0.069385656 | 0 | 0 | 0 | 0 | 8 mo | HOM |
| **1010419538** | 2 | pyd | 198 | 0.069385656 | 0 | 0 | 0 | 0 | 4 mo | WILD |
| **1010419187** | 1 | pyd | 198 | 0.069385656 | 5.5E-05 | 1 | 0.00079232 | 14.41220066 | 8 mo | HOM |
| **1013978366** | 1 | pyd | 198 | 0.069385656 | 0 | 0 | 0 | 0 | 2 mo | HOM |
| **1014416155** | 2 | pyd | 198 | 0.069385656 | 0 | 0 | 0 | 0 | 2 mo | HOM |
| **1013978065** | 2 | pyd | 198 | 0.069385656 | 0 | 0 | 0 | 0 | 2 mo | HOM |
| **1009934102** | 2 | pyd | 198 | 0.069385656 | 0 | 0 | 0 | 0 | 4 mo | WILD |
| **1009322937** | 2 | pyd | 198 | 0.069385656 | 0 | 0 | 0 | 0 | 8 mo | WILD |
| **1011080637** | 2 | pyd | 198 | 0.069385656 | 0 | 0 | 0 | 0 | 8 mo | HOM |
| **1013609713** | 2 | pyd | 198 | 0.069385656 | 0 | 0 | 0 | 0 | 2 mo | WILD |
| **1010418842** | 1 | pyd | 198 | 0.069385656 | 0 | 0 | 0 | 0 | 8 mo | HOM |
| **1013288213** | 2 | pyd | 198 | 0.069385656 | 0 | 0 | 0 | 0 | 4 mo | HOM |
| **1008976060** | 2 | pyd | 198 | 0.069385656 | 0 | 0 | 0 | 0 | 8 mo | WILD |
| **1013446243** | 1 | pyd | 198 | 0.069385656 | 0 | 0 | 0 | 0 | 4 mo | HOM |
| **1013288951** | 2 | pyd | 198 | 0.069385656 | 0 | 0 | 0 | 0 | 4 mo | HOM |
| **1013978665** | 1 | SSp-bfd2/3 | 201 | 1.561862378 | 0 | 0 | 0 | 0 | 2 mo | HOM |
| **1013445609** | 1 | SSp-bfd2/3 | 201 | 1.561862378 | 0.000313586 | 15 | 0.000200777 | 9.603919151 | 4 mo | HOM |
| **1013609015** | 1 | SSp-bfd2/3 | 201 | 1.561862378 | 0.000733843 | 15 | 0.000469851 | 9.603919151 | 4 mo | HOM |
| **1013608106** | 2 | SSp-bfd2/3 | 201 | 1.561862378 | 0 | 0 | 0 | 0 | 4 mo | HOM |

| | | | | | | | | | | |
|---|---|---|---|---|---|---|---|---|---|---|
| **1011937543** | 1 | SSp-bfd2/3 | 201 | 1.561862378 | 0.002623786 | 44 | 0.001679908 | 28.17149618 | 8 mo | HOM |
| **1010419538** | 2 | SSp-bfd2/3 | 201 | 1.561862378 | 0 | 0 | 0 | 0 | 4 mo | WILD |
| **1010419187** | 1 | SSp-bfd2/3 | 201 | 1.561862378 | 0.003215404 | 82 | 0.002058698 | 52.50142469 | 8 mo | HOM |
| **1013978366** | 1 | SSp-bfd2/3 | 201 | 1.561862378 | 5.79E-06 | 1 | 3.7E-06 | 0.640261277 | 2 mo | HOM |
| **1014416155** | 2 | SSp-bfd2/3 | 201 | 1.561862378 | 0 | 0 | 0 | 0 | 2 mo | HOM |
| **1013978065** | 2 | SSp-bfd2/3 | 201 | 1.561862378 | 0 | 0 | 0 | 0 | 2 mo | HOM |
| **1009934102** | 2 | SSp-bfd2/3 | 201 | 1.561862378 | 0 | 0 | 0 | 0 | 4 mo | WILD |
| **1009322937** | 2 | SSp-bfd2/3 | 201 | 1.561862378 | 0 | 0 | 0 | 0 | 8 mo | WILD |
| **1011080637** | 2 | SSp-bfd2/3 | 201 | 1.561862378 | 0.00142771 | 53 | 0.000914107 | 33.93384767 | 8 mo | HOM |
| **1013609713** | 2 | SSp-bfd2/3 | 201 | 1.561862378 | 0 | 0 | 0 | 0 | 2 mo | WILD |
| **1010418842** | 1 | SSp-bfd2/3 | 201 | 1.561862378 | 0.00242988 | 64 | 0.001555758 | 40.97672171 | 8 mo | HOM |
| **1013288213** | 2 | SSp-bfd2/3 | 201 | 1.561862378 | 0.000122301 | 11 | 7.83E-05 | 7.042874044 | 4 mo | HOM |
| **1008976060** | 2 | SSp-bfd2/3 | 201 | 1.561862378 | 0 | 0 | 0 | 0 | 8 mo | WILD |
| **1013446243** | 1 | SSp-bfd2/3 | 201 | 1.561862378 | 0.000509602 | 21 | 0.000326279 | 13.44548681 | 4 mo | HOM |
| **1013288951** | 2 | SSp-bfd2/3 | 201 | 1.561862378 | 0.000563538 | 25 | 0.000360812 | 16.00653192 | 4 mo | HOM |
| **1013978665** | 1 | MV | 202 | 1.87206311 | 0.00075852 | 7 | 0.000405179 | 3.739190181 | 2 mo | HOM |
| **1013445609** | 1 | MV | 202 | 1.87206311 | 4.52E-06 | 1 | 2.41E-06 | 0.534170026 | 4 mo | HOM |
| **1013609015** | 1 | MV | 202 | 1.87206311 | 0.001018178 | 7 | 0.00054388 | 3.739190181 | 4 mo | HOM |
| **1013608106** | 2 | MV | 202 | 1.87206311 | 0 | 0 | 0 | 0 | 4 mo | HOM |
| **1011937543** | 1 | MV | 202 | 1.87206311 | 0.007190297 | 10 | 0.003840841 | 5.341700258 | 8 mo | HOM |
| **1010419538** | 2 | MV | 202 | 1.87206311 | 1.38E-05 | 1 | 7.36E-06 | 0.534170026 | 4 mo | WILD |
| **1010419187** | 1 | MV | 202 | 1.87206311 | 0.001567242 | 9 | 0.000837174 | 4.807530232 | 8 mo | HOM |
| **1013978366** | 1 | MV | 202 | 1.87206311 | 0.000278469 | 3 | 0.00014875 | 1.602510077 | 2 mo | HOM |
| **1014416155** | 2 | MV | 202 | 1.87206311 | 0 | 0 | 0 | 0 | 2 mo | HOM |
| **1013978065** | 2 | MV | 202 | 1.87206311 | 0.000151805 | 1 | 8.11E-05 | 0.534170026 | 2 mo | HOM |
| **1009934102** | 2 | MV | 202 | 1.87206311 | 0 | 0 | 0 | 0 | 4 mo | WILD |
| **1009322937** | 2 | MV | 202 | 1.87206311 | 5.55E-05 | 4 | 2.97E-05 | 2.136680103 | 8 mo | WILD |
| **1011080637** | 2 | MV | 202 | 1.87206311 | 0.001274951 | 9 | 0.000681041 | 4.807530232 | 8 mo | HOM |
| **1013609713** | 2 | MV | 202 | 1.87206311 | 0 | 0 | 0 | 0 | 2 mo | WILD |
| **1010418842** | 1 | MV | 202 | 1.87206311 | 0.001625174 | 32 | 0.000868119 | 17.09344083 | 8 mo | HOM |
| **1013288213** | 2 | MV | 202 | 1.87206311 | 0.000341389 | 6 | 0.00018236 | 3.205020155 | 4 mo | HOM |
| **1008976060** | 2 | MV | 202 | 1.87206311 | 0.002446431 | 41 | 0.00130681 | 21.90097106 | 8 mo | WILD |
| **1013446243** | 1 | MV | 202 | 1.87206311 | 2.88E-05 | 2 | 1.54E-05 | 1.068340052 | 4 mo | HOM |
| **1013288951** | 2 | MV | 202 | 1.87206311 | 0.000340491 | 3 | 0.00018188 | 1.602510077 | 4 mo | HOM |
| **1013978665** | 1 | LIN | 203 | 0.063148672 | 0 | 0 | 0 | 0 | 2 mo | HOM |
| **1013445609** | 1 | LIN | 203 | 0.063148672 | 0 | 0 | 0 | 0 | 4 mo | HOM |
| **1013609015** | 1 | LIN | 203 | 0.063148672 | 0 | 0 | 0 | 0 | 4 mo | HOM |
| **1013608106** | 2 | LIN | 203 | 0.063148672 | 0 | 0 | 0 | 0 | 4 mo | HOM |
| **1011937543** | 1 | LIN | 203 | 0.063148672 | 0 | 0 | 0 | 0 | 8 mo | HOM |
| **1010419538** | 2 | LIN | 203 | 0.063148672 | 0 | 0 | 0 | 0 | 4 mo | WILD |
| **1010419187** | 1 | LIN | 203 | 0.063148672 | 0 | 0 | 0 | 0 | 8 mo | HOM |
| **1013978366** | 1 | LIN | 203 | 0.063148672 | 0 | 0 | 0 | 0 | 2 mo | HOM |
| **1014416155** | 2 | LIN | 203 | 0.063148672 | 0 | 0 | 0 | 0 | 2 mo | HOM |
| **1013978065** | 2 | LIN | 203 | 0.063148672 | 0 | 0 | 0 | 0 | 2 mo | HOM |
| **1009934102** | 2 | LIN | 203 | 0.063148672 | 0 | 0 | 0 | 0 | 4 mo | WILD |
| **1009322937** | 2 | LIN | 203 | 0.063148672 | 0 | 0 | 0 | 0 | 8 mo | WILD |
| **1011080637** | 2 | LIN | 203 | 0.063148672 | 0 | 0 | 0 | 0 | 8 mo | HOM |
| **1013609713** | 2 | LIN | 203 | 0.063148672 | 0 | 0 | 0 | 0 | 2 mo | WILD |
| **1010418842** | 1 | LIN | 203 | 0.063148672 | 4.04E-05 | 2 | 0.000639063 | 31.67129172 | 8 mo | HOM |
| **1013288213** | 2 | LIN | 203 | 0.063148672 | 0 | 0 | 0 | 0 | 4 mo | HOM |
| **1008976060** | 2 | LIN | 203 | 0.063148672 | 1.03E-05 | 1 | 0.000163899 | 15.83564586 | 8 mo | WILD |
| **1013446243** | 1 | LIN | 203 | 0.063148672 | 0 | 0 | 0 | 0 | 4 mo | HOM |
| **1013288951** | 2 | LIN | 203 | 0.063148672 | 0 | 0 | 0 | 0 | 4 mo | HOM |
| **1013978665** | 1 | AOBmi | 204 | 0.285671098 | 0 | 0 | 0 | 0 | 2 mo | HOM |
| **1013445609** | 1 | AOBmi | 204 | 0.285671098 | 0 | 0 | 0 | 0 | 4 mo | HOM |
| **1013609015** | 1 | AOBmi | 204 | 0.285671098 | 2.07E-05 | 1 | 7.24E-05 | 3.500529134 | 4 mo | HOM |
| **1013608106** | 2 | AOBmi | 204 | 0.285671098 | 0 | 0 | 0 | 0 | 4 mo | HOM |
| **1011937543** | 1 | AOBmi | 204 | 0.285671098 | 0.000949858 | 4 | 0.003325006 | 14.00211654 | 8 mo | HOM |
| **1010419538** | 2 | AOBmi | 204 | 0.285671098 | 0 | 0 | 0 | 0 | 4 mo | WILD |

| | | | | | | | | | | |
|---|---|---|---|---|---|---|---|---|---|---|
| **1010419187** | 1 | AOBmi | 204 | 0.285671098 | 0.000183627 | 6 | 0.000642793 | 21.00317481 | 8 mo | HOM |
| **1013978366** | 1 | AOBmi | 204 | 0.285671098 | 1.38E-07 | 0 | 4.82E-07 | 0 | 2 mo | HOM |
| **1014416155** | 2 | AOBmi | 204 | 0.285671098 | 0 | 0 | 0 | 0 | 2 mo | HOM |
| **1013978065** | 2 | AOBmi | 204 | 0.285671098 | 0 | 0 | 0 | 0 | 2 mo | HOM |
| **1009934102** | 2 | AOBmi | 204 | 0.285671098 | 4.14E-06 | 0 | 1.45E-05 | 0 | 4 mo | WILD |
| **1009322937** | 2 | AOBmi | 204 | 0.285671098 | 1.37E-05 | 2 | 4.81E-05 | 7.001058269 | 8 mo | WILD |
| **1011080637** | 2 | AOBmi | 204 | 0.285671098 | 0.000131919 | 9 | 0.000461786 | 31.50476221 | 8 mo | HOM |
| **1013609713** | 2 | AOBmi | 204 | 0.285671098 | 0 | 0 | 0 | 0 | 2 mo | WILD |
| **1010418842** | 1 | AOBmi | 204 | 0.285671098 | 0.000102957 | 7 | 0.000360403 | 24.50370394 | 8 mo | HOM |
| **1013288213** | 2 | AOBmi | 204 | 0.285671098 | 9.64E-06 | 0 | 3.37E-05 | 0 | 4 mo | HOM |
| **1008976060** | 2 | AOBmi | 204 | 0.285671098 | 7.8E-07 | 0 | 2.73E-06 | 0 | 8 mo | WILD |
| **1013446243** | 1 | AOBmi | 204 | 0.285671098 | 1.21E-05 | 1 | 4.24E-05 | 3.500529134 | 4 mo | HOM |
| **1013288951** | 2 | AOBmi | 204 | 0.285671098 | 6.25E-05 | 4 | 0.000218666 | 14.00211654 | 4 mo | HOM |
| **1013978665** | 1 | RM | 206 | 0.120466265 | 0 | 0 | 0 | 0 | 2 mo | HOM |
| **1013445609** | 1 | RM | 206 | 0.120466265 | 0 | 0 | 0 | 0 | 4 mo | HOM |
| **1013609015** | 1 | RM | 206 | 0.120466265 | 0 | 0 | 0 | 0 | 4 mo | HOM |
| **1013608106** | 2 | RM | 206 | 0.117954054 | 0 | 0 | 0 | 0 | 4 mo | HOM |
| **1011937543** | 1 | RM | 206 | 0.120466265 | 0 | 0 | 0 | 0 | 8 mo | HOM |
| **1010419538** | 2 | RM | 206 | 0.120466265 | 0 | 0 | 0 | 0 | 4 mo | WILD |
| **1010419187** | 1 | RM | 206 | 0.120466265 | 0 | 0 | 0 | 0 | 8 mo | HOM |
| **1013978366** | 1 | RM | 206 | 0.120466265 | 0 | 0 | 0 | 0 | 2 mo | HOM |
| **1014416155** | 2 | RM | 206 | 0.120466265 | 0 | 0 | 0 | 0 | 2 mo | HOM |
| **1013978065** | 2 | RM | 206 | 0.120466265 | 0 | 0 | 0 | 0 | 2 mo | HOM |
| **1009934102** | 2 | RM | 206 | 0.120466265 | 0 | 0 | 0 | 0 | 4 mo | WILD |
| **1009322937** | 2 | RM | 206 | 0.120466265 | 0 | 0 | 0 | 0 | 8 mo | WILD |
| **1011080637** | 2 | RM | 206 | 0.120466265 | 1.3E-05 | 1 | 0.000107727 | 8.301079158 | 8 mo | HOM |
| **1013609713** | 2 | RM | 206 | 0.120466265 | 0 | 0 | 0 | 0 | 2 mo | WILD |
| **1010418842** | 1 | RM | 206 | 0.120466265 | 2.93E-05 | 4 | 0.000242969 | 33.20431663 | 8 mo | HOM |
| **1013288213** | 2 | RM | 206 | 0.120466265 | 0 | 0 | 0 | 0 | 4 mo | HOM |
| **1008976060** | 2 | RM | 206 | 0.120466265 | 2.98E-05 | 3 | 0.000247558 | 24.90323747 | 8 mo | WILD |
| **1013446243** | 1 | RM | 206 | 0.120466265 | 0 | 0 | 0 | 0 | 4 mo | HOM |
| **1013288951** | 2 | RM | 206 | 0.120466265 | 0 | 0 | 0 | 0 | 4 mo | HOM |
| **1013978665** | 1 | AP | 207 | 0.054379396 | 0 | 0 | 0 | 0 | 2 mo | HOM |
| **1013445609** | 1 | AP | 207 | 0.054379396 | 7.62E-09 | 0 | 1.4E-07 | 0 | 4 mo | HOM |
| **1013609015** | 1 | AP | 207 | 0.054379396 | 4.71E-07 | 0 | 8.66E-06 | 0 | 4 mo | HOM |
| **1013608106** | 2 | AP | 207 | 0.054379396 | 0 | 0 | 0 | 0 | 4 mo | HOM |
| **1011937543** | 1 | AP | 207 | 0.054379396 | 1.61E-05 | 0 | 0.000296929 | 0 | 8 mo | HOM |
| **1010419538** | 2 | AP | 207 | 0.054379396 | 0 | 0 | 0 | 0 | 4 mo | WILD |
| **1010419187** | 1 | AP | 207 | 0.054379396 | 0 | 0 | 0 | 0 | 8 mo | HOM |
| **1013978366** | 1 | AP | 207 | 0.054379396 | 0 | 0 | 0 | 0 | 2 mo | HOM |
| **1014416155** | 2 | AP | 207 | 0.054379396 | 0 | 0 | 0 | 0 | 2 mo | HOM |
| **1013978065** | 2 | AP | 207 | 0.054379396 | 9.05E-05 | 1 | 0.001664042 | 18.38931776 | 2 mo | HOM |
| **1009934102** | 2 | AP | 207 | 0.054379396 | 0 | 0 | 0 | 0 | 4 mo | WILD |
| **1009322937** | 2 | AP | 207 | 0.054379396 | 0 | 0 | 0 | 0 | 8 mo | WILD |
| **1011080637** | 2 | AP | 207 | 0.054379396 | 0 | 0 | 0 | 0 | 8 mo | HOM |
| **1013609713** | 2 | AP | 207 | 0.054379396 | 0 | 0 | 0 | 0 | 2 mo | WILD |
| **1010418842** | 1 | AP | 207 | 0.054379396 | 8.38E-06 | 1 | 0.000154053 | 18.38931776 | 8 mo | HOM |
| **1013288213** | 2 | AP | 207 | 0.054379396 | 0 | 0 | 0 | 0 | 4 mo | HOM |
| **1008976060** | 2 | AP | 207 | 0.054379396 | 1.06E-09 | 0 | 1.95E-08 | 0 | 8 mo | WILD |
| **1013446243** | 1 | AP | 207 | 0.054379396 | 6.41E-05 | 1 | 0.001179403 | 18.38931776 | 4 mo | HOM |
| **1013288951** | 2 | AP | 207 | 0.054379396 | 7.53E-06 | 1 | 0.000138431 | 18.38931776 | 4 mo | HOM |
| **1013978665** | 1 | LAV | 209 | 0.296257685 | 0 | 0 | 0 | 0 | 2 mo | HOM |
| **1013445609** | 1 | LAV | 209 | 0.296257685 | 0 | 0 | 0 | 0 | 4 mo | HOM |
| **1013609015** | 1 | LAV | 209 | 0.296257685 | 0.000184482 | 2 | 0.000622709 | 6.750879733 | 4 mo | HOM |
| **1013608106** | 2 | LAV | 209 | 0.296257685 | 0 | 0 | 0 | 0 | 4 mo | HOM |
| **1011937543** | 1 | LAV | 209 | 0.296257685 | 0.000105772 | 4 | 0.000357026 | 13.50175947 | 8 mo | HOM |
| **1010419538** | 2 | LAV | 209 | 0.296257685 | 9.15E-07 | 0 | 3.09E-06 | 0 | 4 mo | WILD |
| **1010419187** | 1 | LAV | 209 | 0.296257685 | 0 | 0 | 0 | 0 | 8 mo | HOM |
| **1013978366** | 1 | LAV | 209 | 0.296257685 | 0 | 0 | 0 | 0 | 2 mo | HOM |

| | | | | | | | | | | |
|---|---|---|---|---|---|---|---|---|---|---|
| **1014416155** | 2 | LAV | 209 | 0.296257685 | 0 | 0 | 0 | 0 | 2 mo | HOM |
| **1013978065** | 2 | LAV | 209 | 0.296257685 | 0 | 0 | 0 | 0 | 2 mo | HOM |
| **1009934102** | 2 | LAV | 209 | 0.296257685 | 0 | 0 | 0 | 0 | 4 mo | WILD |
| **1009322937** | 2 | LAV | 209 | 0.296257685 | 4.79E-06 | 1 | 1.62E-05 | 3.375439866 | 8 mo | WILD |
| **1011080637** | 2 | LAV | 209 | 0.296257685 | 9.97E-06 | 1 | 3.36E-05 | 3.375439866 | 8 mo | HOM |
| **1013609713** | 2 | LAV | 209 | 0.296257685 | 0 | 0 | 0 | 0 | 2 mo | WILD |
| **1010418842** | 1 | LAV | 209 | 0.296257685 | 0.000151432 | 13 | 0.00051115 | 43.88071826 | 8 mo | HOM |
| **1013288213** | 2 | LAV | 209 | 0.296257685 | 1.36E-08 | 0 | 4.6E-08 | 0 | 4 mo | HOM |
| **1008976060** | 2 | LAV | 209 | 0.296257685 | 0.000170593 | 10 | 0.000575826 | 33.75439866 | 8 mo | WILD |
| **1013446243** | 1 | LAV | 209 | 0.296257685 | 0 | 0 | 0 | 0 | 4 mo | HOM |
| **1013288951** | 2 | LAV | 209 | 0.296257685 | 0 | 0 | 0 | 0 | 4 mo | HOM |
| **1013978665** | 1 | LM | 210 | 0.079784745 | 0 | 0 | 0 | 0 | 2 mo | HOM |
| **1013445609** | 1 | LM | 210 | 0.079784745 | 2.09E-07 | 0 | 2.62E-06 | 0 | 4 mo | HOM |
| **1013609015** | 1 | LM | 210 | 0.079784745 | 7.66E-06 | 1 | 9.6E-05 | 12.53372435 | 4 mo | HOM |
| **1013608106** | 2 | LM | 210 | 0.079784745 | 0 | 0 | 0 | 0 | 4 mo | HOM |
| **1011937543** | 1 | LM | 210 | 0.079784745 | 0.000243152 | 3 | 0.003047604 | 37.60117305 | 8 mo | HOM |
| **1010419538** | 2 | LM | 210 | 0.079784745 | 4.83E-07 | 0 | 6.05E-06 | 0 | 4 mo | WILD |
| **1010419187** | 1 | LM | 210 | 0.079784745 | 0.00038391 | 0 | 0.004811817 | 0 | 8 mo | HOM |
| **1013978366** | 1 | LM | 210 | 0.079784745 | 7.35E-07 | 0 | 9.21E-06 | 0 | 2 mo | HOM |
| **1014416155** | 2 | LM | 210 | 0.079784745 | 0 | 0 | 0 | 0 | 2 mo | HOM |
| **1013978065** | 2 | LM | 210 | 0.079784745 | 0 | 0 | 0 | 0 | 2 mo | HOM |
| **1009934102** | 2 | LM | 210 | 0.079784745 | 0 | 0 | 0 | 0 | 4 mo | WILD |
| **1009322937** | 2 | LM | 210 | 0.079784745 | 0 | 0 | 0 | 0 | 8 mo | WILD |
| **1011080637** | 2 | LM | 210 | 0.079784745 | 1.42E-08 | 0 | 1.78E-07 | 0 | 8 mo | HOM |
| **1013609713** | 2 | LM | 210 | 0.079784745 | 0 | 0 | 0 | 0 | 2 mo | WILD |
| **1010418842** | 1 | LM | 210 | 0.079784745 | 4.19E-05 | 3 | 0.000525061 | 37.60117305 | 8 mo | HOM |
| **1013288213** | 2 | LM | 210 | 0.079784745 | 2.31E-06 | 0 | 2.89E-05 | 0 | 4 mo | HOM |
| **1008976060** | 2 | LM | 210 | 0.079784745 | 0.000155576 | 1 | 0.001949945 | 12.53372435 | 8 mo | WILD |
| **1013446243** | 1 | LM | 210 | 0.079784745 | 0 | 0 | 0 | 0 | 4 mo | HOM |
| **1013288951** | 2 | LM | 210 | 0.079784745 | 0 | 0 | 0 | 0 | 4 mo | HOM |
| **1013978665** | 1 | ACAd2/3 | 211 | 0.622579378 | 0 | 0 | 0 | 0 | 2 mo | HOM |
| **1013445609** | 1 | ACAd2/3 | 211 | 0.622579378 | 4.28E-05 | 2 | 6.88E-05 | 3.212441769 | 4 mo | HOM |
| **1013609015** | 1 | ACAd2/3 | 211 | 0.622579378 | 0.000219867 | 7 | 0.000353156 | 11.24354619 | 4 mo | HOM |
| **1013608106** | 2 | ACAd2/3 | 211 | 0.622579378 | 0 | 0 | 0 | 0 | 4 mo | HOM |
| **1011937543** | 1 | ACAd2/3 | 211 | 0.622579378 | 0.000963739 | 15 | 0.001547977 | 24.09331327 | 8 mo | HOM |
| **1010419538** | 2 | ACAd2/3 | 211 | 0.622579378 | 0 | 0 | 0 | 0 | 4 mo | WILD |
| **1010419187** | 1 | ACAd2/3 | 211 | 0.622579378 | 0.000532632 | 22 | 0.000855525 | 35.33685946 | 8 mo | HOM |
| **1013978366** | 1 | ACAd2/3 | 211 | 0.622579378 | 0 | 0 | 0 | 0 | 2 mo | HOM |
| **1014416155** | 2 | ACAd2/3 | 211 | 0.622579378 | 0 | 0 | 0 | 0 | 2 mo | HOM |
| **1013978065** | 2 | ACAd2/3 | 211 | 0.622579378 | 0 | 0 | 0 | 0 | 2 mo | HOM |
| **1009934102** | 2 | ACAd2/3 | 211 | 0.622579378 | 0 | 0 | 0 | 0 | 4 mo | WILD |
| **1009322937** | 2 | ACAd2/3 | 211 | 0.622579378 | 0 | 0 | 0 | 0 | 8 mo | WILD |
| **1011080637** | 2 | ACAd2/3 | 211 | 0.622579378 | 0.000211466 | 13 | 0.000339661 | 20.8808715 | 8 mo | HOM |
| **1013609713** | 2 | ACAd2/3 | 211 | 0.622579378 | 0 | 0 | 0 | 0 | 2 mo | WILD |
| **1010418842** | 1 | ACAd2/3 | 211 | 0.622579378 | 0.000561719 | 31 | 0.000902245 | 49.79284743 | 8 mo | HOM |
| **1013288213** | 2 | ACAd2/3 | 211 | 0.622579378 | 1.48E-05 | 1 | 2.38E-05 | 1.606220885 | 4 mo | HOM |
| **1008976060** | 2 | ACAd2/3 | 211 | 0.622579378 | 0 | 0 | 0 | 0 | 8 mo | WILD |
| **1013446243** | 1 | ACAd2/3 | 211 | 0.622579378 | 7.83E-05 | 1 | 0.000125839 | 1.606220885 | 4 mo | HOM |
| **1013288951** | 2 | ACAd2/3 | 211 | 0.622579378 | 0.000113699 | 3 | 0.000182626 | 4.818662654 | 4 mo | HOM |
| **1013978665** | 1 | RN | 214 | 0.811576259 | 0 | 0 | 0 | 0 | 2 mo | HOM |
| **1013445609** | 1 | RN | 214 | 0.811576259 | 0 | 0 | 0 | 0 | 4 mo | HOM |
| **1013609015** | 1 | RN | 214 | 0.811576259 | 0 | 0 | 0 | 0 | 4 mo | HOM |
| **1013608106** | 2 | RN | 214 | 0.811576259 | 0 | 0 | 0 | 0 | 4 mo | HOM |
| **1011937543** | 1 | RN | 214 | 0.811576259 | 0.000734234 | 15 | 0.000904701 | 18.48255149 | 8 mo | HOM |
| **1010419538** | 2 | RN | 214 | 0.811576259 | 0 | 0 | 0 | 0 | 4 mo | WILD |
| **1010419187** | 1 | RN | 214 | 0.811576259 | 3.78E-06 | 1 | 4.65E-06 | 1.232170099 | 8 mo | HOM |
| **1013978366** | 1 | RN | 214 | 0.811576259 | 0 | 0 | 0 | 0 | 2 mo | HOM |
| **1014416155** | 2 | RN | 214 | 0.811576259 | 0 | 0 | 0 | 0 | 2 mo | HOM |
| **1013978065** | 2 | RN | 214 | 0.811576259 | 0 | 0 | 0 | 0 | 2 mo | HOM |

| | | | | | | | | | | |
|---|---|---|---|---|---|---|---|---|---|---|
| 1009934102 | 2 | RN | 214 | 0.811576259 | 0 | 0 | 0 | 0 | 4 mo | WILD |
| 1009322937 | 2 | RN | 214 | 0.811576259 | 0 | 0 | 0 | 0 | 8 mo | WILD |
| 1011080637 | 2 | RN | 214 | 0.811576259 | 1.84E-07 | 0 | 2.27E-07 | 0 | 8 mo | HOM |
| 1013609713 | 2 | RN | 214 | 0.811576259 | 0 | 0 | 0 | 0 | 2 mo | WILD |
| 1010418842 | 1 | RN | 214 | 0.811576259 | 3.65E-05 | 4 | 4.49E-05 | 4.928680398 | 8 mo | HOM |
| 1013288213 | 2 | RN | 214 | 0.811576259 | 0 | 0 | 0 | 0 | 4 mo | HOM |
| 1008976060 | 2 | RN | 214 | 0.811576259 | 0.000111003 | 15 | 0.000136775 | 18.48255149 | 8 mo | WILD |
| 1013446243 | 1 | RN | 214 | 0.811576259 | 0 | 0 | 0 | 0 | 4 mo | HOM |
| 1013288951 | 2 | RN | 214 | 0.811576259 | 0 | 0 | 0 | 0 | 4 mo | HOM |
| 1013978665 | 1 | APN | 215 | 1.314591835 | 0 | 0 | 0 | 0 | 2 mo | HOM |
| 1013445609 | 1 | APN | 215 | 1.314591835 | 0 | 0 | 0 | 0 | 4 mo | HOM |
| 1013609015 | 1 | APN | 215 | 1.314591835 | 0 | 0 | 0 | 0 | 4 mo | HOM |
| 1013608106 | 2 | APN | 215 | 1.314591835 | 0 | 0 | 0 | 0 | 4 mo | HOM |
| 1011937543 | 1 | APN | 215 | 1.314591835 | 0.002303756 | 50 | 0.00175245 | 38.03461931 | 8 mo | HOM |
| 1010419538 | 2 | APN | 215 | 1.314591835 | 0 | 0 | 0 | 0 | 4 mo | WILD |
| 1010419187 | 1 | APN | 215 | 1.314591835 | 0.000184596 | 8 | 0.000140421 | 6.08553909 | 8 mo | HOM |
| 1013978366 | 1 | APN | 215 | 1.314591835 | 5.3E-06 | 1 | 4.03E-06 | 0.760692386 | 2 mo | HOM |
| 1014416155 | 2 | APN | 215 | 1.314591835 | 0 | 0 | 0 | 0 | 2 mo | HOM |
| 1013978065 | 2 | APN | 215 | 1.314591835 | 0 | 0 | 0 | 0 | 2 mo | HOM |
| 1009934102 | 2 | APN | 215 | 1.314591835 | 0 | 0 | 0 | 0 | 4 mo | WILD |
| 1009322937 | 2 | APN | 215 | 1.314591835 | 0 | 0 | 0 | 0 | 8 mo | WILD |
| 1011080637 | 2 | APN | 215 | 1.314591835 | 0.000100094 | 7 | 7.61E-05 | 5.324846703 | 8 mo | HOM |
| 1013609713 | 2 | APN | 215 | 1.314591835 | 1.38E-06 | 0 | 1.05E-06 | 0 | 2 mo | WILD |
| 1010418842 | 1 | APN | 215 | 1.314591835 | 0.000563796 | 22 | 0.000428875 | 16.7352325 | 8 mo | HOM |
| 1013288213 | 2 | APN | 215 | 1.314591835 | 3.88E-07 | 0 | 2.95E-07 | 0 | 4 mo | HOM |
| 1008976060 | 2 | APN | 215 | 1.314591835 | 3E-05 | 4 | 2.28E-05 | 3.042769545 | 8 mo | WILD |
| 1013446243 | 1 | APN | 215 | 1.314591835 | 0 | 0 | 0 | 0 | 4 mo | HOM |
| 1013288951 | 2 | APN | 215 | 1.314591835 | 1.46E-05 | 1 | 1.11E-05 | 0.760692386 | 4 mo | HOM |
| 1013978665 | 1 | SUV | 217 | 0.364384384 | 5.64E-05 | 1 | 0.000154653 | 2.744354709 | 2 mo | HOM |
| 1013445609 | 1 | SUV | 217 | 0.364384384 | 0 | 0 | 0 | 0 | 4 mo | HOM |
| 1013609015 | 1 | SUV | 217 | 0.364384384 | 0.000694407 | 1 | 0.001905699 | 2.744354709 | 4 mo | HOM |
| 1013608106 | 2 | SUV | 217 | 0.364384384 | 0 | 0 | 0 | 0 | 4 mo | HOM |
| 1011937543 | 1 | SUV | 217 | 0.364384384 | 0 | 0 | 0 | 0 | 8 mo | HOM |
| 1010419538 | 2 | SUV | 217 | 0.364384384 | 0 | 0 | 0 | 0 | 4 mo | WILD |
| 1010419187 | 1 | SUV | 217 | 0.364384384 | 2.19E-05 | 0 | 6.02E-05 | 0 | 8 mo | HOM |
| 1013978366 | 1 | SUV | 217 | 0.364384384 | 3.73E-05 | 0 | 0.000102499 | 0 | 2 mo | HOM |
| 1014416155 | 2 | SUV | 217 | 0.364384384 | 0 | 0 | 0 | 0 | 2 mo | HOM |
| 1013978065 | 2 | SUV | 217 | 0.364384384 | 0 | 0 | 0 | 0 | 2 mo | HOM |
| 1009934102 | 2 | SUV | 217 | 0.364384384 | 2.63E-10 | 0 | 7.23E-10 | 0 | 4 mo | WILD |
| 1009322937 | 2 | SUV | 217 | 0.364384384 | 7.89E-07 | 0 | 2.17E-06 | 0 | 8 mo | WILD |
| 1011080637 | 2 | SUV | 217 | 0.364384384 | 0.000297518 | 0 | 0.000816496 | 0 | 8 mo | HOM |
| 1013609713 | 2 | SUV | 217 | 0.364384384 | 0 | 0 | 0 | 0 | 2 mo | WILD |
| 1010418842 | 1 | SUV | 217 | 0.364384384 | 0.000296742 | 7 | 0.000814365 | 19.21048296 | 8 mo | HOM |
| 1013288213 | 2 | SUV | 217 | 0.364384384 | 3.36E-06 | 0 | 9.22E-06 | 0 | 4 mo | HOM |
| 1008976060 | 2 | SUV | 217 | 0.364384384 | 0.000411746 | 12 | 0.001129978 | 32.9322565 | 8 mo | WILD |
| 1013446243 | 1 | SUV | 217 | 0.364384384 | 1.63E-05 | 1 | 4.49E-05 | 2.744354709 | 4 mo | HOM |
| 1013288951 | 2 | SUV | 217 | 0.364384384 | 0.000693982 | 1 | 0.001904532 | 2.744354709 | 4 mo | HOM |
| 1013978665 | 1 | LP | 218 | 1.236976698 | 0 | 0 | 0 | 0 | 2 mo | HOM |
| 1013445609 | 1 | LP | 218 | 1.236976698 | 8.75E-06 | 1 | 7.07E-06 | 0.808422666 | 4 mo | HOM |
| 1013609015 | 1 | LP | 218 | 1.236976698 | 1.88E-05 | 1 | 1.52E-05 | 0.808422666 | 4 mo | HOM |
| 1013608106 | 2 | LP | 218 | 1.236976698 | 0 | 0 | 0 | 0 | 4 mo | HOM |
| 1011937543 | 1 | LP | 218 | 1.236976698 | 0.001877044 | 29 | 0.001517445 | 23.44425732 | 8 mo | HOM |
| 1010419538 | 2 | LP | 218 | 1.236976698 | 0 | 0 | 0 | 0 | 4 mo | WILD |
| 1010419187 | 1 | LP | 218 | 1.236976698 | 0.000710905 | 22 | 0.000574712 | 17.78529866 | 8 mo | HOM |
| 1013978366 | 1 | LP | 218 | 1.236976698 | 0 | 0 | 0 | 0 | 2 mo | HOM |
| 1014416155 | 2 | LP | 218 | 1.236976698 | 0 | 0 | 0 | 0 | 2 mo | HOM |
| 1013978065 | 2 | LP | 218 | 1.236976698 | 0 | 0 | 0 | 0 | 2 mo | HOM |
| 1009934102 | 2 | LP | 218 | 1.236976698 | 7.75E-09 | 0 | 6.27E-09 | 0 | 4 mo | WILD |
| 1009322937 | 2 | LP | 218 | 1.236976698 | 9.01E-06 | 2 | 7.29E-06 | 1.616845333 | 8 mo | WILD |

| | | | | | | | | | | |
|---|---|---|---|---|---|---|---|---|---|---|
| **1011080637** | 2 | LP | 218 | 1.236976698 | 0.000623714 | 23 | 0.000504225 | 18.59372132 | 8 mo | HOM |
| **1013609713** | 2 | LP | 218 | 1.236976698 | 0 | 0 | 0 | 0 | 2 mo | WILD |
| **1010418842** | 1 | LP | 218 | 1.236976698 | 0.000933583 | 28 | 0.00075473 | 22.63583466 | 8 mo | HOM |
| **1013288213** | 2 | LP | 218 | 1.236976698 | 2.4E-05 | 1 | 1.94E-05 | 0.808422666 | 4 mo | HOM |
| **1008976060** | 2 | LP | 218 | 1.236976698 | 3.61E-05 | 5 | 2.92E-05 | 4.042113331 | 8 mo | WILD |
| **1013446243** | 1 | LP | 218 | 1.236976698 | 2.64E-05 | 1 | 2.13E-05 | 0.808422666 | 4 mo | HOM |
| **1013288951** | 2 | LP | 218 | 1.236976698 | 1.39E-05 | 1 | 1.12E-05 | 0.808422666 | 4 mo | HOM |
| **1013978665** | 1 | RO | 222 | 0.0411987 | 0 | 0 | 0 | 0 | 2 mo | HOM |
| **1013445609** | 1 | RO | 222 | 0.0411987 | 0 | 0 | 0 | 0 | 4 mo | HOM |
| **1013609015** | 1 | RO | 222 | 0.0411987 | 0 | 0 | 0 | 0 | 4 mo | HOM |
| **1013608106** | 2 | RO | 222 | 0.037437627 | 0 | 0 | 0 | 0 | 4 mo | HOM |
| **1011937543** | 1 | RO | 222 | 0.0411987 | 0 | 0 | 0 | 0 | 8 mo | HOM |
| **1010419538** | 2 | RO | 222 | 0.0411987 | 0 | 0 | 0 | 0 | 4 mo | WILD |
| **1010419187** | 1 | RO | 222 | 0.0411987 | 0 | 0 | 0 | 0 | 8 mo | HOM |
| **1013978366** | 1 | RO | 222 | 0.0411987 | 0 | 0 | 0 | 0 | 2 mo | HOM |
| **1014416155** | 2 | RO | 222 | 0.0411987 | 0 | 0 | 0 | 0 | 2 mo | HOM |
| **1013978065** | 2 | RO | 222 | 0.0411987 | 0 | 0 | 0 | 0 | 2 mo | HOM |
| **1009934102** | 2 | RO | 222 | 0.0411987 | 0 | 0 | 0 | 0 | 4 mo | WILD |
| **1009322937** | 2 | RO | 222 | 0.0411987 | 0 | 0 | 0 | 0 | 8 mo | WILD |
| **1011080637** | 2 | RO | 222 | 0.0411987 | 0 | 0 | 0 | 0 | 8 mo | HOM |
| **1013609713** | 2 | RO | 222 | 0.0411987 | 0 | 0 | 0 | 0 | 2 mo | WILD |
| **1010418842** | 1 | RO | 222 | 0.0411987 | 9.12E-08 | 0 | 2.21E-06 | 0 | 8 mo | HOM |
| **1013288213** | 2 | RO | 222 | 0.0411987 | 0 | 0 | 0 | 0 | 4 mo | HOM |
| **1008976060** | 2 | RO | 222 | 0.0411987 | 0 | 0 | 0 | 0 | 8 mo | WILD |
| **1013446243** | 1 | RO | 222 | 0.0411987 | 0 | 0 | 0 | 0 | 4 mo | HOM |
| **1013288951** | 2 | RO | 222 | 0.0411987 | 0 | 0 | 0 | 0 | 4 mo | HOM |
| **1013978665** | 1 | ARH | 223 | 0.293548181 | 0 | 0 | 0 | 0 | 2 mo | HOM |
| **1013445609** | 1 | ARH | 223 | 0.293548181 | 2.47E-06 | 0 | 8.43E-06 | 0 | 4 mo | HOM |
| **1013609015** | 1 | ARH | 223 | 0.293548181 | 0 | 0 | 0 | 0 | 4 mo | HOM |
| **1013608106** | 2 | ARH | 223 | 0.293548181 | 0 | 0 | 0 | 0 | 4 mo | HOM |
| **1011937543** | 1 | ARH | 223 | 0.293548181 | 0.001221855 | 5 | 0.004162365 | 17.032979 | 8 mo | HOM |
| **1010419538** | 2 | ARH | 223 | 0.293548181 | 9.34E-06 | 2 | 3.18E-05 | 6.813191601 | 4 mo | WILD |
| **1010419187** | 1 | ARH | 223 | 0.293548181 | 8.65E-06 | 0 | 2.95E-05 | 0 | 8 mo | HOM |
| **1013978366** | 1 | ARH | 223 | 0.293548181 | 0 | 0 | 0 | 0 | 2 mo | HOM |
| **1014416155** | 2 | ARH | 223 | 0.293548181 | 0 | 0 | 0 | 0 | 2 mo | HOM |
| **1013978065** | 2 | ARH | 223 | 0.293548181 | 0 | 0 | 0 | 0 | 2 mo | HOM |
| **1009934102** | 2 | ARH | 223 | 0.293548181 | 0 | 0 | 0 | 0 | 4 mo | WILD |
| **1009322937** | 2 | ARH | 223 | 0.293548181 | 0 | 0 | 0 | 0 | 8 mo | WILD |
| **1011080637** | 2 | ARH | 223 | 0.293548181 | 4.76E-05 | 1 | 0.000162267 | 3.406595801 | 8 mo | HOM |
| **1013609713** | 2 | ARH | 223 | 0.293548181 | 0 | 0 | 0 | 0 | 2 mo | WILD |
| **1010418842** | 1 | ARH | 223 | 0.293548181 | 5.11E-05 | 2 | 0.000174101 | 6.813191601 | 8 mo | HOM |
| **1013288213** | 2 | ARH | 223 | 0.293548181 | 0 | 0 | 0 | 0 | 4 mo | HOM |
| **1008976060** | 2 | ARH | 223 | 0.293548181 | 4.97E-06 | 1 | 1.69E-05 | 3.406595801 | 8 mo | WILD |
| **1013446243** | 1 | ARH | 223 | 0.293548181 | 0 | 0 | 0 | 0 | 4 mo | HOM |
| **1013288951** | 2 | ARH | 223 | 0.293548181 | 5.2E-08 | 0 | 1.77E-07 | 0 | 4 mo | HOM |
| **1013978665** | 1 | SPIV | 225 | 0.771706488 | 0.000188254 | 0 | 0.000243945 | 0 | 2 mo | HOM |
| **1013445609** | 1 | SPIV | 225 | 0.771706488 | 2.93E-05 | 1 | 3.8E-05 | 1.295829458 | 4 mo | HOM |
| **1013609015** | 1 | SPIV | 225 | 0.771706488 | 3.89E-05 | 0 | 5.05E-05 | 0 | 4 mo | HOM |
| **1013608106** | 2 | SPIV | 225 | 0.771706488 | 0 | 0 | 0 | 0 | 4 mo | HOM |
| **1011937543** | 1 | SPIV | 225 | 0.771706488 | 0.001793029 | 4 | 0.002323459 | 5.183317832 | 8 mo | HOM |
| **1010419538** | 2 | SPIV | 225 | 0.771706488 | 3.15E-05 | 2 | 4.08E-05 | 2.591658916 | 4 mo | WILD |
| **1010419187** | 1 | SPIV | 225 | 0.771706488 | 0.000712546 | 2 | 0.000923337 | 2.591658916 | 8 mo | HOM |
| **1013978366** | 1 | SPIV | 225 | 0.771706488 | 7.13E-06 | 1 | 9.24E-06 | 1.295829458 | 2 mo | HOM |
| **1014416155** | 2 | SPIV | 225 | 0.771706488 | 0 | 0 | 0 | 0 | 2 mo | HOM |
| **1013978065** | 2 | SPIV | 225 | 0.771706488 | 3.22E-05 | 1 | 4.17E-05 | 1.295829458 | 2 mo | HOM |
| **1009934102** | 2 | SPIV | 225 | 0.771706488 | 0 | 0 | 0 | 0 | 4 mo | WILD |
| **1009322937** | 2 | SPIV | 225 | 0.771706488 | 2.71E-06 | 0 | 3.51E-06 | 0 | 8 mo | WILD |
| **1011080637** | 2 | SPIV | 225 | 0.771706488 | 0.000221231 | 2 | 0.000286678 | 2.591658916 | 8 mo | HOM |
| **1013609713** | 2 | SPIV | 225 | 0.771706488 | 0 | 0 | 0 | 0 | 2 mo | WILD |

| | | | | | | | | | | |
|---|---|---|---|---|---|---|---|---|---|---|
| **1010418842** | 1 | SPIV | 225 | 0.771706488 | 0.00035613 | 4 | 0.000461484 | 5.183317832 | 8 mo | HOM |
| **1013288213** | 2 | SPIV | 225 | 0.771706488 | 0.00018061 | 1 | 0.00023404 | 1.295829458 | 4 mo | HOM |
| **1008976060** | 2 | SPIV | 225 | 0.771706488 | 0.001438906 | 8 | 0.001864577 | 10.36663566 | 8 mo | WILD |
| **1013446243** | 1 | SPIV | 225 | 0.771706488 | 9.57E-07 | 0 | 1.24E-06 | 0 | 4 mo | HOM |
| **1013288951** | 2 | SPIV | 225 | 0.771706488 | 0.000126887 | 1 | 0.000164425 | 1.295829458 | 4 mo | HOM |
| **1013978665** | 1 | LPO | 226 | 0.576901265 | 0 | 0 | 0 | 0 | 2 mo | HOM |
| **1013445609** | 1 | LPO | 226 | 0.576901265 | 0 | 0 | 0 | 0 | 4 mo | HOM |
| **1013609015** | 1 | LPO | 226 | 0.576901265 | 0 | 0 | 0 | 0 | 4 mo | HOM |
| **1013608106** | 2 | LPO | 226 | 0.576901265 | 0 | 0 | 0 | 0 | 4 mo | HOM |
| **1011937543** | 1 | LPO | 226 | 0.576901265 | 0.000200174 | 5 | 0.000346981 | 8.666994346 | 8 mo | HOM |
| **1010419538** | 2 | LPO | 226 | 0.576901265 | 0 | 0 | 0 | 0 | 4 mo | WILD |
| **1010419187** | 1 | LPO | 226 | 0.576901265 | 5.39E-05 | 5 | 9.35E-05 | 8.666994346 | 8 mo | HOM |
| **1013978366** | 1 | LPO | 226 | 0.576901265 | 0 | 0 | 0 | 0 | 2 mo | HOM |
| **1014416155** | 2 | LPO | 226 | 0.576901265 | 0 | 0 | 0 | 0 | 2 mo | HOM |
| **1013978065** | 2 | LPO | 226 | 0.576901265 | 0 | 0 | 0 | 0 | 2 mo | HOM |
| **1009934102** | 2 | LPO | 226 | 0.576901265 | 0 | 0 | 0 | 0 | 4 mo | WILD |
| **1009322937** | 2 | LPO | 226 | 0.576901265 | 0 | 0 | 0 | 0 | 8 mo | WILD |
| **1011080637** | 2 | LPO | 226 | 0.576901265 | 0.000134384 | 4 | 0.000232941 | 6.933595476 | 8 mo | HOM |
| **1013609713** | 2 | LPO | 226 | 0.576901265 | 0 | 0 | 0 | 0 | 2 mo | WILD |
| **1010418842** | 1 | LPO | 226 | 0.576901265 | 0.000392988 | 13 | 0.000681204 | 22.5341853 | 8 mo | HOM |
| **1013288213** | 2 | LPO | 226 | 0.576901265 | 0 | 0 | 0 | 0 | 4 mo | HOM |
| **1008976060** | 2 | LPO | 226 | 0.576901265 | 0 | 0 | 0 | 0 | 8 mo | WILD |
| **1013446243** | 1 | LPO | 226 | 0.576901265 | 0 | 0 | 0 | 0 | 4 mo | HOM |
| **1013288951** | 2 | LPO | 226 | 0.576901265 | 0 | 0 | 0 | 0 | 4 mo | HOM |
| **1013978665** | 1 | sV | 229 | 2.388106806 | 0 | 0 | 0 | 0 | 2 mo | HOM |
| **1013445609** | 1 | sV | 229 | 2.388106806 | 0 | 0 | 0 | 0 | 4 mo | HOM |
| **1013609015** | 1 | sV | 229 | 2.388106806 | 0 | 0 | 0 | 0 | 4 mo | HOM |
| **1013608106** | 2 | sV | 229 | 2.388106806 | 0 | 0 | 0 | 0 | 4 mo | HOM |
| **1011937543** | 1 | sV | 229 | 2.388106806 | 0.004525818 | 5 | 0.001895149 | 2.093708701 | 8 mo | HOM |
| **1010419538** | 2 | sV | 229 | 2.388106806 | 0 | 0 | 0 | 0 | 4 mo | WILD |
| **1010419187** | 1 | sV | 229 | 2.388106806 | 0.000807922 | 1 | 0.000338311 | 0.41874174 | 8 mo | HOM |
| **1013978366** | 1 | sV | 229 | 2.388106806 | 1.01E-05 | 0 | 4.24E-06 | 0 | 2 mo | HOM |
| **1014416155** | 2 | sV | 229 | 2.388106806 | 0 | 0 | 0 | 0 | 2 mo | HOM |
| **1013978065** | 2 | sV | 229 | 2.388106806 | 0 | 0 | 0 | 0 | 2 mo | HOM |
| **1009934102** | 2 | sV | 229 | 2.388106806 | 1.95E-09 | 0 | 8.16E-10 | 0 | 4 mo | WILD |
| **1009322937** | 2 | sV | 229 | 2.388106806 | 0 | 0 | 0 | 0 | 8 mo | WILD |
| **1011080637** | 2 | sV | 229 | 2.388106806 | 0 | 0 | 0 | 0 | 8 mo | HOM |
| **1013609713** | 2 | sV | 229 | 2.388106806 | 0 | 0 | 0 | 0 | 2 mo | WILD |
| **1010418842** | 1 | sV | 229 | 2.388106806 | 4.1E-07 | 0 | 1.72E-07 | 0 | 8 mo | HOM |
| **1013288213** | 2 | sV | 229 | 2.388106806 | 0 | 0 | 0 | 0 | 4 mo | HOM |
| **1008976060** | 2 | sV | 229 | 2.388106806 | 3.49E-05 | 4 | 1.46E-05 | 1.674966961 | 8 mo | WILD |
| **1013446243** | 1 | sV | 229 | 2.388106806 | 2.71E-07 | 0 | 1.14E-07 | 0 | 4 mo | HOM |
| **1013288951** | 2 | sV | 229 | 2.388106806 | 3.91E-08 | 0 | 1.64E-08 | 0 | 4 mo | HOM |
| **1013978665** | 1 | RPA | 230 | 0.068627411 | 0 | 0 | 0 | 0 | 2 mo | HOM |
| **1013445609** | 1 | RPA | 230 | 0.068627411 | 0 | 0 | 0 | 0 | 4 mo | HOM |
| **1013609015** | 1 | RPA | 230 | 0.068627411 | 0 | 0 | 0 | 0 | 4 mo | HOM |
| **1013608106** | 2 | RPA | 230 | 0.068295798 | 0 | 0 | 0 | 0 | 4 mo | HOM |
| **1011937543** | 1 | RPA | 230 | 0.068627411 | 0 | 0 | 0 | 0 | 8 mo | HOM |
| **1010419538** | 2 | RPA | 230 | 0.068627411 | 0 | 0 | 0 | 0 | 4 mo | WILD |
| **1010419187** | 1 | RPA | 230 | 0.068627411 | 0 | 0 | 0 | 0 | 8 mo | HOM |
| **1013978366** | 1 | RPA | 230 | 0.068627411 | 0 | 0 | 0 | 0 | 2 mo | HOM |
| **1014416155** | 2 | RPA | 230 | 0.068627411 | 0 | 0 | 0 | 0 | 2 mo | HOM |
| **1013978065** | 2 | RPA | 230 | 0.068627411 | 0 | 0 | 0 | 0 | 2 mo | HOM |
| **1009934102** | 2 | RPA | 230 | 0.068627411 | 0 | 0 | 0 | 0 | 4 mo | WILD |
| **1009322937** | 2 | RPA | 230 | 0.068627411 | 0 | 0 | 0 | 0 | 8 mo | WILD |
| **1011080637** | 2 | RPA | 230 | 0.068627411 | 0 | 0 | 0 | 0 | 8 mo | HOM |
| **1013609713** | 2 | RPA | 230 | 0.068627411 | 0 | 0 | 0 | 0 | 2 mo | WILD |
| **1010418842** | 1 | RPA | 230 | 0.068627411 | 0 | 0 | 0 | 0 | 8 mo | HOM |
| **1013288213** | 2 | RPA | 230 | 0.068627411 | 0 | 0 | 0 | 0 | 4 mo | HOM |

| | | | | | | | | | | |
|---|---|---|---|---|---|---|---|---|---|---|
| 1008976060 | 2 | RPA | 230 | 0.068627411 | 0 | 0 | 0 | 0 | 8 mo | WILD |
| 1013446243 | 1 | RPA | 230 | 0.068627411 | 0 | 0 | 0 | 0 | 4 mo | HOM |
| 1013288951 | 2 | RPA | 230 | 0.068627411 | 0 | 0 | 0 | 0 | 4 mo | HOM |
| 1013978665 | 1 | AT | 231 | 0.044365609 | 0 | 0 | 0 | 0 | 2 mo | HOM |
| 1013445609 | 1 | AT | 231 | 0.044365609 | 0 | 0 | 0 | 0 | 4 mo | HOM |
| 1013609015 | 1 | AT | 231 | 0.044365609 | 0 | 0 | 0 | 0 | 4 mo | HOM |
| 1013608106 | 2 | AT | 231 | 0.044365609 | 0 | 0 | 0 | 0 | 4 mo | HOM |
| 1011937543 | 1 | AT | 231 | 0.044365609 | 0 | 0 | 0 | 0 | 8 mo | HOM |
| 1010419538 | 2 | AT | 231 | 0.044365609 | 0 | 0 | 0 | 0 | 4 mo | WILD |
| 1010419187 | 1 | AT | 231 | 0.044365609 | 0 | 0 | 0 | 0 | 8 mo | HOM |
| 1013978366 | 1 | AT | 231 | 0.044365609 | 0 | 0 | 0 | 0 | 2 mo | HOM |
| 1014416155 | 2 | AT | 231 | 0.044365609 | 0 | 0 | 0 | 0 | 2 mo | HOM |
| 1013978065 | 2 | AT | 231 | 0.044365609 | 0 | 0 | 0 | 0 | 2 mo | HOM |
| 1009934102 | 2 | AT | 231 | 0.044365609 | 0 | 0 | 0 | 0 | 4 mo | WILD |
| 1009322937 | 2 | AT | 231 | 0.044365609 | 0 | 0 | 0 | 0 | 8 mo | WILD |
| 1011080637 | 2 | AT | 231 | 0.044365609 | 0 | 0 | 0 | 0 | 8 mo | HOM |
| 1013609713 | 2 | AT | 231 | 0.044365609 | 0 | 0 | 0 | 0 | 2 mo | WILD |
| 1010418842 | 1 | AT | 231 | 0.044365609 | 0 | 0 | 0 | 0 | 8 mo | HOM |
| 1013288213 | 2 | AT | 231 | 0.044365609 | 0 | 0 | 0 | 0 | 4 mo | HOM |
| 1008976060 | 2 | AT | 231 | 0.044365609 | 0 | 0 | 0 | 0 | 8 mo | WILD |
| 1013446243 | 1 | AT | 231 | 0.044365609 | 0 | 0 | 0 | 0 | 4 mo | HOM |
| 1013288951 | 2 | AT | 231 | 0.044365609 | 0 | 0 | 0 | 0 | 4 mo | HOM |
| 1013978665 | 1 | VISal5 | 233 | 0.209075924 | 0 | 0 | 0 | 0 | 2 mo | HOM |
| 1013445609 | 1 | VISal5 | 233 | 0.209075924 | 1.89E-05 | 2 | 9.03E-05 | 9.56590294 | 4 mo | HOM |
| 1013609015 | 1 | VISal5 | 233 | 0.209075924 | 4.48E-05 | 4 | 0.000214414 | 19.13180588 | 4 mo | HOM |
| 1013608106 | 2 | VISal5 | 233 | 0.209075924 | 0 | 0 | 0 | 0 | 4 mo | HOM |
| 1011937543 | 1 | VISal5 | 233 | 0.209075924 | 0.000168266 | 6 | 0.00080481 | 28.69770882 | 8 mo | HOM |
| 1010419538 | 2 | VISal5 | 233 | 0.209075924 | 0 | 0 | 0 | 0 | 4 mo | WILD |
| 1010419187 | 1 | VISal5 | 233 | 0.209075924 | 0.000233994 | 12 | 0.00111918 | 57.39541764 | 8 mo | HOM |
| 1013978366 | 1 | VISal5 | 233 | 0.209075924 | 0 | 0 | 0 | 0 | 2 mo | HOM |
| 1014416155 | 2 | VISal5 | 233 | 0.209075924 | 0 | 0 | 0 | 0 | 2 mo | HOM |
| 1013978065 | 2 | VISal5 | 233 | 0.209075924 | 0 | 0 | 0 | 0 | 2 mo | HOM |
| 1009934102 | 2 | VISal5 | 233 | 0.209075924 | 0 | 0 | 0 | 0 | 4 mo | WILD |
| 1009322937 | 2 | VISal5 | 233 | 0.209075924 | 0 | 0 | 0 | 0 | 8 mo | WILD |
| 1011080637 | 2 | VISal5 | 233 | 0.209075924 | 0.000272371 | 13 | 0.001302739 | 62.17836911 | 8 mo | HOM |
| 1013609713 | 2 | VISal5 | 233 | 0.209075924 | 0 | 0 | 0 | 0 | 2 mo | WILD |
| 1010418842 | 1 | VISal5 | 233 | 0.209075924 | 0.000230063 | 7 | 0.001100381 | 33.48066029 | 8 mo | HOM |
| 1013288213 | 2 | VISal5 | 233 | 0.209075924 | 4.34E-05 | 2 | 0.000207397 | 9.56590294 | 4 mo | HOM |
| 1008976060 | 2 | VISal5 | 233 | 0.209075924 | 0 | 0 | 0 | 0 | 8 mo | WILD |
| 1013446243 | 1 | VISal5 | 233 | 0.209075924 | 3.73E-06 | 0 | 1.79E-05 | 0 | 4 mo | HOM |
| 1013288951 | 2 | VISal5 | 233 | 0.209075924 | 1.45E-05 | 0 | 6.94E-05 | 0 | 4 mo | HOM |
| 1013978665 | 1 | TEa4 | 234 | 0.322272922 | 0 | 0 | 0 | 0 | 2 mo | HOM |
| 1013445609 | 1 | TEa4 | 234 | 0.322272922 | 7.37E-05 | 2 | 0.00022884 | 6.205920094 | 4 mo | HOM |
| 1013609015 | 1 | TEa4 | 234 | 0.322272922 | 0.000164459 | 5 | 0.000510309 | 15.51480024 | 4 mo | HOM |
| 1013608106 | 2 | TEa4 | 234 | 0.322272922 | 0 | 0 | 0 | 0 | 4 mo | HOM |
| 1011937543 | 1 | TEa4 | 234 | 0.322272922 | 0.000202982 | 10 | 0.000629845 | 31.02960047 | 8 mo | HOM |
| 1010419538 | 2 | TEa4 | 234 | 0.322272922 | 0 | 0 | 0 | 0 | 4 mo | WILD |
| 1010419187 | 1 | TEa4 | 234 | 0.322272922 | 0.000590172 | 19 | 0.001831281 | 58.9562409 | 8 mo | HOM |
| 1013978366 | 1 | TEa4 | 234 | 0.322272922 | 0 | 0 | 0 | 0 | 2 mo | HOM |
| 1014416155 | 2 | TEa4 | 234 | 0.322272922 | 0 | 0 | 0 | 0 | 2 mo | HOM |
| 1013978065 | 2 | TEa4 | 234 | 0.322272922 | 0 | 0 | 0 | 0 | 2 mo | HOM |
| 1009934102 | 2 | TEa4 | 234 | 0.322272922 | 0 | 0 | 0 | 0 | 4 mo | WILD |
| 1009322937 | 2 | TEa4 | 234 | 0.322272922 | 0 | 0 | 0 | 0 | 8 mo | WILD |
| 1011080637 | 2 | TEa4 | 234 | 0.322272922 | 0.000289903 | 16 | 0.000899556 | 49.64736075 | 8 mo | HOM |
| 1013609713 | 2 | TEa4 | 234 | 0.322272922 | 0 | 0 | 0 | 0 | 2 mo | WILD |
| 1010418842 | 1 | TEa4 | 234 | 0.322272922 | 0.000495382 | 18 | 0.001537152 | 55.85328085 | 8 mo | HOM |
| 1013288213 | 2 | TEa4 | 234 | 0.322272922 | 7.59E-05 | 3 | 0.000235425 | 9.308880141 | 4 mo | HOM |
| 1008976060 | 2 | TEa4 | 234 | 0.322272922 | 0 | 0 | 0 | 0 | 8 mo | WILD |
| 1013446243 | 1 | TEa4 | 234 | 0.322272922 | 0.000107594 | 5 | 0.000333859 | 15.51480024 | 4 mo | HOM |

| | | | | | | | | | | |
|---|---|---|---|---|---|---|---|---|---|---|
| 1013288951 | 2 | TEa4 | 234 | 0.322272922 | 3.66E-05 | 1 | 0.000113501 | 3.102960047 | 4 mo | HOM |
| 1013978665 | 1 | LRN | 235 | 0.586954198 | 0 | 0 | 0 | 0 | 2 mo | HOM |
| 1013445609 | 1 | LRN | 235 | 0.586954198 | 0 | 0 | 0 | 0 | 4 mo | HOM |
| 1013609015 | 1 | LRN | 235 | 0.586954198 | 0 | 0 | 0 | 0 | 4 mo | HOM |
| 1013608106 | 2 | LRN | 235 | 0.586954198 | 0 | 0 | 0 | 0 | 4 mo | HOM |
| 1011937543 | 1 | LRN | 235 | 0.586954198 | 0.000410433 | 0 | 0.000699259 | 0 | 8 mo | HOM |
| 1010419538 | 2 | LRN | 235 | 0.586954198 | 0 | 0 | 0 | 0 | 4 mo | WILD |
| 1010419187 | 1 | LRN | 235 | 0.586954198 | 9.03E-09 | 0 | 1.54E-08 | 0 | 8 mo | HOM |
| 1013978366 | 1 | LRN | 235 | 0.586954198 | 0 | 0 | 0 | 0 | 2 mo | HOM |
| 1014416155 | 2 | LRN | 235 | 0.586954198 | 0 | 0 | 0 | 0 | 2 mo | HOM |
| 1013978065 | 2 | LRN | 235 | 0.586954198 | 0 | 0 | 0 | 0 | 2 mo | HOM |
| 1009934102 | 2 | LRN | 235 | 0.586954198 | 0 | 0 | 0 | 0 | 4 mo | WILD |
| 1009322937 | 2 | LRN | 235 | 0.586954198 | 0 | 0 | 0 | 0 | 8 mo | WILD |
| 1011080637 | 2 | LRN | 235 | 0.586954198 | 0 | 0 | 0 | 0 | 8 mo | HOM |
| 1013609713 | 2 | LRN | 235 | 0.586954198 | 0 | 0 | 0 | 0 | 2 mo | WILD |
| 1010418842 | 1 | LRN | 235 | 0.586954198 | 2.26E-05 | 4 | 3.84E-05 | 6.814841796 | 8 mo | HOM |
| 1013288213 | 2 | LRN | 235 | 0.586954198 | 0 | 0 | 0 | 0 | 4 mo | HOM |
| 1008976060 | 2 | LRN | 235 | 0.586954198 | 6.95E-05 | 10 | 0.000118389 | 17.03710449 | 8 mo | WILD |
| 1013446243 | 1 | LRN | 235 | 0.586954198 | 0 | 0 | 0 | 0 | 4 mo | HOM |
| 1013288951 | 2 | LRN | 235 | 0.586954198 | 0 | 0 | 0 | 0 | 4 mo | HOM |
| 1013978665 | 1 | ts | 237 | 0.009593455 | 0 | 0 | 0 | 0 | 2 mo | HOM |
| 1013445609 | 1 | ts | 237 | 0.009593455 | 0 | 0 | 0 | 0 | 4 mo | HOM |
| 1013609015 | 1 | ts | 237 | 0.009593455 | 0 | 0 | 0 | 0 | 4 mo | HOM |
| 1013608106 | 2 | ts | 237 | 0.009593455 | 0 | 0 | 0 | 0 | 4 mo | HOM |
| 1011937543 | 1 | ts | 237 | 0.009593455 | 0 | 0 | 0 | 0 | 8 mo | HOM |
| 1010419538 | 2 | ts | 237 | 0.009593455 | 0 | 0 | 0 | 0 | 4 mo | WILD |
| 1010419187 | 1 | ts | 237 | 0.009593455 | 0 | 0 | 0 | 0 | 8 mo | HOM |
| 1013978366 | 1 | ts | 237 | 0.009593455 | 0 | 0 | 0 | 0 | 2 mo | HOM |
| 1014416155 | 2 | ts | 237 | 0.009593455 | 0 | 0 | 0 | 0 | 2 mo | HOM |
| 1013978065 | 2 | ts | 237 | 0.009593455 | 0 | 0 | 0 | 0 | 2 mo | HOM |
| 1009934102 | 2 | ts | 237 | 0.009593455 | 0 | 0 | 0 | 0 | 4 mo | WILD |
| 1009322937 | 2 | ts | 237 | 0.009593455 | 0 | 0 | 0 | 0 | 8 mo | WILD |
| 1011080637 | 2 | ts | 237 | 0.009593455 | 0 | 0 | 0 | 0 | 8 mo | HOM |
| 1013609713 | 2 | ts | 237 | 0.009593455 | 0 | 0 | 0 | 0 | 2 mo | WILD |
| 1010418842 | 1 | ts | 237 | 0.009593455 | 0 | 0 | 0 | 0 | 8 mo | HOM |
| 1013288213 | 2 | ts | 237 | 0.009593455 | 0 | 0 | 0 | 0 | 4 mo | HOM |
| 1008976060 | 2 | ts | 237 | 0.009593455 | 0 | 0 | 0 | 0 | 8 mo | WILD |
| 1013446243 | 1 | ts | 237 | 0.009593455 | 0 | 0 | 0 | 0 | 4 mo | HOM |
| 1013288951 | 2 | ts | 237 | 0.009593455 | 0 | 0 | 0 | 0 | 4 mo | HOM |
| 1013978665 | 1 | RPO | 238 | 0.100106961 | 0 | 0 | 0 | 0 | 2 mo | HOM |
| 1013445609 | 1 | RPO | 238 | 0.100106961 | 0 | 0 | 0 | 0 | 4 mo | HOM |
| 1013609015 | 1 | RPO | 238 | 0.100106961 | 0 | 0 | 0 | 0 | 4 mo | HOM |
| 1013608106 | 2 | RPO | 238 | 0.100106961 | 0 | 0 | 0 | 0 | 4 mo | HOM |
| 1011937543 | 1 | RPO | 238 | 0.100106961 | 0 | 0 | 0 | 0 | 8 mo | HOM |
| 1010419538 | 2 | RPO | 238 | 0.100106961 | 0 | 0 | 0 | 0 | 4 mo | WILD |
| 1010419187 | 1 | RPO | 238 | 0.100106961 | 0 | 0 | 0 | 0 | 8 mo | HOM |
| 1013978366 | 1 | RPO | 238 | 0.100106961 | 0 | 0 | 0 | 0 | 2 mo | HOM |
| 1014416155 | 2 | RPO | 238 | 0.100106961 | 0 | 0 | 0 | 0 | 2 mo | HOM |
| 1013978065 | 2 | RPO | 238 | 0.100106961 | 0 | 0 | 0 | 0 | 2 mo | HOM |
| 1009934102 | 2 | RPO | 238 | 0.100106961 | 0 | 0 | 0 | 0 | 4 mo | WILD |
| 1009322937 | 2 | RPO | 238 | 0.100106961 | 0 | 0 | 0 | 0 | 8 mo | WILD |
| 1011080637 | 2 | RPO | 238 | 0.100106961 | 6.27E-07 | 0 | 6.26E-06 | 0 | 8 mo | HOM |
| 1013609713 | 2 | RPO | 238 | 0.100106961 | 0 | 0 | 0 | 0 | 2 mo | WILD |
| 1010418842 | 1 | RPO | 238 | 0.100106961 | 0 | 0 | 0 | 0 | 8 mo | HOM |
| 1013288213 | 2 | RPO | 238 | 0.100106961 | 0 | 0 | 0 | 0 | 4 mo | HOM |
| 1008976060 | 2 | RPO | 238 | 0.100106961 | 0 | 0 | 0 | 0 | 8 mo | WILD |
| 1013446243 | 1 | RPO | 238 | 0.100106961 | 0 | 0 | 0 | 0 | 4 mo | HOM |
| 1013288951 | 2 | RPO | 238 | 0.100106961 | 0 | 0 | 0 | 0 | 4 mo | HOM |
| 1013978665 | 1 | ATN | 239 | 2.224002362 | 0 | 0 | 0 | 0 | 2 mo | HOM |

| | | | | | | | | | | |
|---|---|---|---|---|---|---|---|---|---|---|
| **1013445609** | 1 | ATN | 239 | 2.224002362 | 8.23E-05 | 5 | 3.7E-05 | 2.248199051 | 4 mo | HOM |
| **1013609015** | 1 | ATN | 239 | 2.224002362 | 0.000518231 | 17 | 0.000233017 | 7.643876775 | 4 mo | HOM |
| **1013608106** | 2 | ATN | 239 | 2.224002362 | 0 | 0 | 0 | 0 | 4 mo | HOM |
| **1011937543** | 1 | ATN | 239 | 2.224002362 | 0.003754886 | 29 | 0.001688346 | 13.0395545 | 8 mo | HOM |
| **1010419538** | 2 | ATN | 239 | 2.224002362 | 0 | 0 | 0 | 0 | 4 mo | WILD |
| **1010419187** | 1 | ATN | 239 | 2.224002362 | 0.005714049 | 85 | 0.002569264 | 38.21938387 | 8 mo | HOM |
| **1013978366** | 1 | ATN | 239 | 2.224002362 | 1.38E-05 | 2 | 6.21E-06 | 0.899279621 | 2 mo | HOM |
| **1014416155** | 2 | ATN | 239 | 2.224002362 | 0 | 0 | 0 | 0 | 2 mo | HOM |
| **1013978065** | 2 | ATN | 239 | 2.224002362 | 0 | 0 | 0 | 0 | 2 mo | HOM |
| **1009934102** | 2 | ATN | 239 | 2.224002362 | 5.12E-06 | 1 | 2.3E-06 | 0.44963981 | 4 mo | WILD |
| **1009322937** | 2 | ATN | 239 | 2.224002362 | 1.93E-05 | 2 | 8.68E-06 | 0.899279621 | 8 mo | WILD |
| **1011080637** | 2 | ATN | 239 | 2.224002362 | 0.002353523 | 58 | 0.001058237 | 26.079109 | 8 mo | HOM |
| **1013609713** | 2 | ATN | 239 | 2.224002362 | 0 | 0 | 0 | 0 | 2 mo | WILD |
| **1010418842** | 1 | ATN | 239 | 2.224002362 | 0.004500136 | 78 | 0.00202344 | 35.0719052 | 8 mo | HOM |
| **1013288213** | 2 | ATN | 239 | 2.224002362 | 2.79E-05 | 4 | 1.25E-05 | 1.798559241 | 4 mo | HOM |
| **1008976060** | 2 | ATN | 239 | 2.224002362 | 0.000147386 | 13 | 6.63E-05 | 5.845317534 | 8 mo | WILD |
| **1013446243** | 1 | ATN | 239 | 2.224002362 | 0.000260719 | 12 | 0.000117229 | 5.395677723 | 4 mo | HOM |
| **1013288951** | 2 | ATN | 239 | 2.224002362 | 9.77E-05 | 5 | 4.4E-05 | 2.248199051 | 4 mo | HOM |
| **1013978665** | 1 | LS | 242 | 3.14502141 | 0 | 0 | 0 | 0 | 2 mo | HOM |
| **1013445609** | 1 | LS | 242 | 3.14502141 | 5.85E-07 | 0 | 1.86E-07 | 0 | 4 mo | HOM |
| **1013609015** | 1 | LS | 242 | 3.14502141 | 0.000146047 | 4 | 4.64E-05 | 1.271851437 | 4 mo | HOM |
| **1013608106** | 2 | LS | 242 | 3.14502141 | 0 | 0 | 0 | 0 | 4 mo | HOM |
| **1011937543** | 1 | LS | 242 | 3.14502141 | 0.001025267 | 48 | 0.000325997 | 15.26221725 | 8 mo | HOM |
| **1010419538** | 2 | LS | 242 | 3.14502141 | 0 | 0 | 0 | 0 | 4 mo | WILD |
| **1010419187** | 1 | LS | 242 | 3.14502141 | 0.002246972 | 69 | 0.000714454 | 21.93943729 | 8 mo | HOM |
| **1013978366** | 1 | LS | 242 | 3.14502141 | 9.44E-07 | 0 | 3E-07 | 0 | 2 mo | HOM |
| **1014416155** | 2 | LS | 242 | 3.14502141 | 0 | 0 | 0 | 0 | 2 mo | HOM |
| **1013978065** | 2 | LS | 242 | 3.14502141 | 0 | 0 | 0 | 0 | 2 mo | HOM |
| **1009934102** | 2 | LS | 242 | 3.14502141 | 0 | 0 | 0 | 0 | 4 mo | WILD |
| **1009322937** | 2 | LS | 242 | 3.14502141 | 9.1E-07 | 0 | 2.89E-07 | 0 | 8 mo | WILD |
| **1011080637** | 2 | LS | 242 | 3.14502141 | 0.001586221 | 52 | 0.000504359 | 16.53406868 | 8 mo | HOM |
| **1013609713** | 2 | LS | 242 | 3.14502141 | 0 | 0 | 0 | 0 | 2 mo | WILD |
| **1010418842** | 1 | LS | 242 | 3.14502141 | 0.005439798 | 109 | 0.001729654 | 34.65795167 | 8 mo | HOM |
| **1013288213** | 2 | LS | 242 | 3.14502141 | 8.14E-05 | 5 | 2.59E-05 | 1.589814297 | 4 mo | HOM |
| **1008976060** | 2 | LS | 242 | 3.14502141 | 0.000169775 | 19 | 5.4E-05 | 6.041294327 | 8 mo | WILD |
| **1013446243** | 1 | LS | 242 | 3.14502141 | 9.44E-05 | 4 | 3E-05 | 1.271851437 | 4 mo | HOM |
| **1013288951** | 2 | LS | 242 | 3.14502141 | 0.000118747 | 4 | 3.78E-05 | 1.271851437 | 4 mo | HOM |
| **1013978665** | 1 | AUDd6b | 243 | 0.031380653 | 0 | 0 | 0 | 0 | 2 mo | HOM |
| **1013445609** | 1 | AUDd6b | 243 | 0.031380653 | 0 | 0 | 0 | 0 | 4 mo | HOM |
| **1013609015** | 1 | AUDd6b | 243 | 0.031380653 | 8.25E-07 | 0 | 2.63E-05 | 0 | 4 mo | HOM |
| **1013608106** | 2 | AUDd6b | 243 | 0.031380653 | 0 | 0 | 0 | 0 | 4 mo | HOM |
| **1011937543** | 1 | AUDd6b | 243 | 0.031380653 | 2.37E-05 | 0 | 0.000755151 | 0 | 8 mo | HOM |
| **1010419538** | 2 | AUDd6b | 243 | 0.031380653 | 0 | 0 | 0 | 0 | 4 mo | WILD |
| **1010419187** | 1 | AUDd6b | 243 | 0.031380653 | 2.71E-05 | 1 | 0.000863717 | 31.86676785 | 8 mo | HOM |
| **1013978366** | 1 | AUDd6b | 243 | 0.031380653 | 0 | 0 | 0 | 0 | 2 mo | HOM |
| **1014416155** | 2 | AUDd6b | 243 | 0.031380653 | 0 | 0 | 0 | 0 | 2 mo | HOM |
| **1013978065** | 2 | AUDd6b | 243 | 0.031380653 | 0 | 0 | 0 | 0 | 2 mo | HOM |
| **1009934102** | 2 | AUDd6b | 243 | 0.031380653 | 0 | 0 | 0 | 0 | 4 mo | WILD |
| **1009322937** | 2 | AUDd6b | 243 | 0.031380653 | 0 | 0 | 0 | 0 | 8 mo | WILD |
| **1011080637** | 2 | AUDd6b | 243 | 0.031380653 | 4.74E-05 | 3 | 0.001511985 | 95.60030354 | 8 mo | HOM |
| **1013609713** | 2 | AUDd6b | 243 | 0.031380653 | 0 | 0 | 0 | 0 | 2 mo | WILD |
| **1010418842** | 1 | AUDd6b | 243 | 0.031380653 | 2.07E-05 | 1 | 0.000659399 | 31.86676785 | 8 mo | HOM |
| **1013288213** | 2 | AUDd6b | 243 | 0.031380653 | 0 | 0 | 0 | 0 | 4 mo | HOM |
| **1008976060** | 2 | AUDd6b | 243 | 0.031380653 | 0 | 0 | 0 | 0 | 8 mo | WILD |
| **1013446243** | 1 | AUDd6b | 243 | 0.031380653 | 8.64E-10 | 0 | 2.75E-08 | 0 | 4 mo | HOM |
| **1013288951** | 2 | AUDd6b | 243 | 0.031380653 | 3.83E-06 | 1 | 0.00012202 | 31.86676785 | 4 mo | HOM |
| **1013978665** | 1 | RR | 246 | 0.14194028 | 0 | 0 | 0 | 0 | 2 mo | HOM |
| **1013445609** | 1 | RR | 246 | 0.14194028 | 0 | 0 | 0 | 0 | 4 mo | HOM |
| **1013609015** | 1 | RR | 246 | 0.14194028 | 0 | 0 | 0 | 0 | 4 mo | HOM |

| | | | | | | | | | | |
|---|---|---|---|---|---|---|---|---|---|---|
| **1013608106** | 2 | RR | 246 | 0.14194028 | 0 | 0 | 0 | 0 | 4 mo | HOM |
| **1011937543** | 1 | RR | 246 | 0.14194028 | 3.75E-05 | 3 | 0.000264541 | 21.1356495 | 8 mo | HOM |
| **1010419538** | 2 | RR | 246 | 0.14194028 | 0 | 0 | 0 | 0 | 4 mo | WILD |
| **1010419187** | 1 | RR | 246 | 0.14194028 | 3.8E-06 | 1 | 2.68E-05 | 7.045216501 | 8 mo | HOM |
| **1013978366** | 1 | RR | 246 | 0.14194028 | 0 | 0 | 0 | 0 | 2 mo | HOM |
| **1014416155** | 2 | RR | 246 | 0.14194028 | 0 | 0 | 0 | 0 | 2 mo | HOM |
| **1013978065** | 2 | RR | 246 | 0.14194028 | 0 | 0 | 0 | 0 | 2 mo | HOM |
| **1009934102** | 2 | RR | 246 | 0.14194028 | 0 | 0 | 0 | 0 | 4 mo | WILD |
| **1009322937** | 2 | RR | 246 | 0.14194028 | 0 | 0 | 0 | 0 | 8 mo | WILD |
| **1011080637** | 2 | RR | 246 | 0.14194028 | 0 | 0 | 0 | 0 | 8 mo | HOM |
| **1013609713** | 2 | RR | 246 | 0.14194028 | 0 | 0 | 0 | 0 | 2 mo | WILD |
| **1010418842** | 1 | RR | 246 | 0.14194028 | 5.8E-06 | 0 | 4.09E-05 | 0 | 8 mo | HOM |
| **1013288213** | 2 | RR | 246 | 0.14194028 | 0 | 0 | 0 | 0 | 4 mo | HOM |
| **1008976060** | 2 | RR | 246 | 0.14194028 | 0 | 0 | 0 | 0 | 8 mo | WILD |
| **1013446243** | 1 | RR | 246 | 0.14194028 | 0 | 0 | 0 | 0 | 4 mo | HOM |
| **1013288951** | 2 | RR | 246 | 0.14194028 | 0 | 0 | 0 | 0 | 4 mo | HOM |
| **1013978665** | 1 | AUD | 247 | 5.969379119 | 1.07E-05 | 2 | 1.8E-06 | 0.33504322 | 2 mo | HOM |
| **1013445609** | 1 | AUD | 247 | 5.969379119 | 0.000529209 | 42 | 8.87E-05 | 7.035907615 | 4 mo | HOM |
| **1013609015** | 1 | AUD | 247 | 5.969379119 | 0.001870037 | 69 | 0.000313272 | 11.55899108 | 4 mo | HOM |
| **1013608106** | 2 | AUD | 247 | 5.969379119 | 0 | 0 | 0 | 0 | 4 mo | HOM |
| **1011937543** | 1 | AUD | 247 | 5.969379119 | 0.009995765 | 202 | 0.001674507 | 33.8393652 | 8 mo | HOM |
| **1010419538** | 2 | AUD | 247 | 5.969379119 | 2.02E-06 | 0 | 3.39E-07 | 0 | 4 mo | WILD |
| **1010419187** | 1 | AUD | 247 | 5.969379119 | 0.009112326 | 299 | 0.001526511 | 50.08896136 | 8 mo | HOM |
| **1013978366** | 1 | AUD | 247 | 5.969379119 | 1.93E-05 | 2 | 3.23E-06 | 0.33504322 | 2 mo | HOM |
| **1014416155** | 2 | AUD | 247 | 5.969379119 | 0 | 0 | 0 | 0 | 2 mo | HOM |
| **1013978065** | 2 | AUD | 247 | 5.969379119 | 0 | 0 | 0 | 0 | 2 mo | HOM |
| **1009934102** | 2 | AUD | 247 | 5.969379119 | 2.32E-06 | 1 | 3.88E-07 | 0.16752161 | 4 mo | WILD |
| **1009322937** | 2 | AUD | 247 | 5.969379119 | 1.12E-05 | 1 | 1.88E-06 | 0.16752161 | 8 mo | WILD |
| **1011080637** | 2 | AUD | 247 | 5.969379119 | 0.004799049 | 193 | 0.000803944 | 32.33167071 | 8 mo | HOM |
| **1013609713** | 2 | AUD | 247 | 5.969379119 | 1.54E-05 | 1 | 2.58E-06 | 0.16752161 | 2 mo | WILD |
| **1010418842** | 1 | AUD | 247 | 5.969379119 | 0.009432945 | 243 | 0.001580222 | 40.7077512 | 8 mo | HOM |
| **1013288213** | 2 | AUD | 247 | 5.969379119 | 0.000535257 | 37 | 8.97E-05 | 6.198299566 | 4 mo | HOM |
| **1008976060** | 2 | AUD | 247 | 5.969379119 | 8.87E-05 | 10 | 1.49E-05 | 1.675216099 | 8 mo | WILD |
| **1013446243** | 1 | AUD | 247 | 5.969379119 | 0.001092527 | 60 | 0.000183022 | 10.05129659 | 4 mo | HOM |
| **1013288951** | 2 | AUD | 247 | 5.969379119 | 0.002751065 | 87 | 0.000460863 | 14.57438006 | 4 mo | HOM |
| **1013978665** | 1 | AUDpo6a | 249 | 0.081783381 | 0 | 0 | 0 | 0 | 2 mo | HOM |
| **1013445609** | 1 | AUDpo6a | 249 | 0.081783381 | 9.59E-06 | 2 | 0.000117227 | 24.45484603 | 4 mo | HOM |
| **1013609015** | 1 | AUDpo6a | 249 | 0.081783381 | 2.48E-05 | 2 | 0.000303046 | 24.45484603 | 4 mo | HOM |
| **1013608106** | 2 | AUDpo6a | 249 | 0.081783381 | 0 | 0 | 0 | 0 | 4 mo | HOM |
| **1011937543** | 1 | AUDpo6a | 249 | 0.081783381 | 0.00030738 | 2 | 0.003758462 | 24.45484603 | 8 mo | HOM |
| **1010419538** | 2 | AUDpo6a | 249 | 0.081783381 | 0 | 0 | 0 | 0 | 4 mo | WILD |
| **1010419187** | 1 | AUDpo6a | 249 | 0.081783381 | 4.27E-05 | 2 | 0.000522213 | 24.45484603 | 8 mo | HOM |
| **1013978366** | 1 | AUDpo6a | 249 | 0.081783381 | 0 | 0 | 0 | 0 | 2 mo | HOM |
| **1014416155** | 2 | AUDpo6a | 249 | 0.081783381 | 0 | 0 | 0 | 0 | 2 mo | HOM |
| **1013978065** | 2 | AUDpo6a | 249 | 0.081783381 | 0 | 0 | 0 | 0 | 2 mo | HOM |
| **1009934102** | 2 | AUDpo6a | 249 | 0.081783381 | 0 | 0 | 0 | 0 | 4 mo | WILD |
| **1009322937** | 2 | AUDpo6a | 249 | 0.081783381 | 0 | 0 | 0 | 0 | 8 mo | WILD |
| **1011080637** | 2 | AUDpo6a | 249 | 0.081783381 | 4.01E-05 | 1 | 0.000490338 | 12.22742302 | 8 mo | HOM |
| **1013609713** | 2 | AUDpo6a | 249 | 0.081783381 | 0 | 0 | 0 | 0 | 2 mo | WILD |
| **1010418842** | 1 | AUDpo6a | 249 | 0.081783381 | 4.79E-05 | 2 | 0.000585774 | 24.45484603 | 8 mo | HOM |
| **1013288213** | 2 | AUDpo6a | 249 | 0.081783381 | 0 | 0 | 0 | 0 | 4 mo | HOM |
| **1008976060** | 2 | AUDpo6a | 249 | 0.081783381 | 0 | 0 | 0 | 0 | 8 mo | WILD |
| **1013446243** | 1 | AUDpo6a | 249 | 0.081783381 | 3.88E-05 | 3 | 0.000474167 | 36.68226905 | 4 mo | HOM |
| **1013288951** | 2 | AUDpo6a | 249 | 0.081783381 | 6.13E-06 | 0 | 7.5E-05 | 0 | 4 mo | HOM |
| **1013978665** | 1 | LSc | 250 | 0.580636868 | 0 | 0 | 0 | 0 | 2 mo | HOM |
| **1013445609** | 1 | LSc | 250 | 0.580636868 | 3.87E-09 | 0 | 6.66E-09 | 0 | 4 mo | HOM |
| **1013609015** | 1 | LSc | 250 | 0.580636868 | 0 | 0 | 0 | 0 | 4 mo | HOM |
| **1013608106** | 2 | LSc | 250 | 0.580636868 | 0 | 0 | 0 | 0 | 4 mo | HOM |
| **1011937543** | 1 | LSc | 250 | 0.580636868 | 4.72E-05 | 3 | 8.12E-05 | 5.166740463 | 8 mo | HOM |

| | | | | | | | | | | |
|---|---|---|---|---|---|---|---|---|---|---|
| 1010419538 | 2 | LSc | 250 | 0.580636868 | 0 | 0 | 0 | 0 | 4 mo | WILD |
| 1010419187 | 1 | LSc | 250 | 0.580636868 | 0.000113846 | 6 | 0.000196071 | 10.33348093 | 8 mo | HOM |
| 1013978366 | 1 | LSc | 250 | 0.580636868 | 9.44E-07 | 0 | 1.63E-06 | 0 | 2 mo | HOM |
| 1014416155 | 2 | LSc | 250 | 0.580636868 | 0 | 0 | 0 | 0 | 2 mo | HOM |
| 1013978065 | 2 | LSc | 250 | 0.580636868 | 0 | 0 | 0 | 0 | 2 mo | HOM |
| 1009934102 | 2 | LSc | 250 | 0.580636868 | 0 | 0 | 0 | 0 | 4 mo | WILD |
| 1009322937 | 2 | LSc | 250 | 0.580636868 | 9.1E-07 | 0 | 1.57E-06 | 0 | 8 mo | WILD |
| 1011080637 | 2 | LSc | 250 | 0.580636868 | 6.04E-05 | 3 | 0.000104007 | 5.166740463 | 8 mo | HOM |
| 1013609713 | 2 | LSc | 250 | 0.580636868 | 0 | 0 | 0 | 0 | 2 mo | WILD |
| 1010418842 | 1 | LSc | 250 | 0.580636868 | 0.000496245 | 14 | 0.000854656 | 24.11145549 | 8 mo | HOM |
| 1013288213 | 2 | LSc | 250 | 0.580636868 | 1.28E-05 | 1 | 2.2E-05 | 1.722246821 | 4 mo | HOM |
| 1008976060 | 2 | LSc | 250 | 0.580636868 | 7.12E-05 | 9 | 0.000122596 | 15.50022139 | 8 mo | WILD |
| 1013446243 | 1 | LSc | 250 | 0.580636868 | 4.43E-11 | 0 | 7.63E-11 | 0 | 4 mo | HOM |
| 1013288951 | 2 | LSc | 250 | 0.580636868 | 0 | 0 | 0 | 0 | 4 mo | HOM |
| 1013978665 | 1 | AUDp2/3 | 251 | 0.486480467 | 0 | 0 | 0 | 0 | 2 mo | HOM |
| 1013445609 | 1 | AUDp2/3 | 251 | 0.486480467 | 6.12E-05 | 4 | 0.000125816 | 8.222323957 | 4 mo | HOM |
| 1013609015 | 1 | AUDp2/3 | 251 | 0.486480467 | 0.000285562 | 7 | 0.000586995 | 14.38906692 | 4 mo | HOM |
| 1013608106 | 2 | AUDp2/3 | 251 | 0.486480467 | 0 | 0 | 0 | 0 | 4 mo | HOM |
| 1011937543 | 1 | AUDp2/3 | 251 | 0.486480467 | 0.000524308 | 14 | 0.001077757 | 28.77813385 | 8 mo | HOM |
| 1010419538 | 2 | AUDp2/3 | 251 | 0.486480467 | 0 | 0 | 0 | 0 | 4 mo | WILD |
| 1010419187 | 1 | AUDp2/3 | 251 | 0.486480467 | 0.000790844 | 18 | 0.001625645 | 37.00045781 | 8 mo | HOM |
| 1013978366 | 1 | AUDp2/3 | 251 | 0.486480467 | 0 | 0 | 0 | 0 | 2 mo | HOM |
| 1014416155 | 2 | AUDp2/3 | 251 | 0.486480467 | 0 | 0 | 0 | 0 | 2 mo | HOM |
| 1013978065 | 2 | AUDp2/3 | 251 | 0.486480467 | 0 | 0 | 0 | 0 | 2 mo | HOM |
| 1009934102 | 2 | AUDp2/3 | 251 | 0.486480467 | 0 | 0 | 0 | 0 | 4 mo | WILD |
| 1009322937 | 2 | AUDp2/3 | 251 | 0.486480467 | 0 | 0 | 0 | 0 | 8 mo | WILD |
| 1011080637 | 2 | AUDp2/3 | 251 | 0.486480467 | 0.000296746 | 10 | 0.000609985 | 20.55580989 | 8 mo | HOM |
| 1013609713 | 2 | AUDp2/3 | 251 | 0.486480467 | 0 | 0 | 0 | 0 | 2 mo | WILD |
| 1010418842 | 1 | AUDp2/3 | 251 | 0.486480467 | 0.000268495 | 12 | 0.000551913 | 24.66697187 | 8 mo | HOM |
| 1013288213 | 2 | AUDp2/3 | 251 | 0.486480467 | 4.15E-05 | 2 | 8.53E-05 | 4.111161978 | 4 mo | HOM |
| 1008976060 | 2 | AUDp2/3 | 251 | 0.486480467 | 0 | 0 | 0 | 0 | 8 mo | WILD |
| 1013446243 | 1 | AUDp2/3 | 251 | 0.486480467 | 2.21E-05 | 2 | 4.55E-05 | 4.111161978 | 4 mo | HOM |
| 1013288951 | 2 | AUDp2/3 | 251 | 0.486480467 | 3.85E-05 | 2 | 7.92E-05 | 4.111161978 | 4 mo | HOM |
| 1013978665 | 1 | AUDd5 | 252 | 0.353435158 | 0 | 0 | 0 | 0 | 2 mo | HOM |
| 1013445609 | 1 | AUDd5 | 252 | 0.353435158 | 4.21E-05 | 4 | 0.000119213 | 11.31749322 | 4 mo | HOM |
| 1013609015 | 1 | AUDd5 | 252 | 0.353435158 | 0.000185214 | 6 | 0.00052404 | 16.97623983 | 4 mo | HOM |
| 1013608106 | 2 | AUDd5 | 252 | 0.353435158 | 0 | 0 | 0 | 0 | 4 mo | HOM |
| 1011937543 | 1 | AUDd5 | 252 | 0.353435158 | 0.000464212 | 15 | 0.001313428 | 42.44059958 | 8 mo | HOM |
| 1010419538 | 2 | AUDd5 | 252 | 0.353435158 | 0 | 0 | 0 | 0 | 4 mo | WILD |
| 1010419187 | 1 | AUDd5 | 252 | 0.353435158 | 0.000575682 | 22 | 0.00162882 | 62.24621271 | 8 mo | HOM |
| 1013978366 | 1 | AUDd5 | 252 | 0.353435158 | 2.35E-08 | 0 | 6.65E-08 | 0 | 2 mo | HOM |
| 1014416155 | 2 | AUDd5 | 252 | 0.353435158 | 0 | 0 | 0 | 0 | 2 mo | HOM |
| 1013978065 | 2 | AUDd5 | 252 | 0.353435158 | 0 | 0 | 0 | 0 | 2 mo | HOM |
| 1009934102 | 2 | AUDd5 | 252 | 0.353435158 | 0 | 0 | 0 | 0 | 4 mo | WILD |
| 1009322937 | 2 | AUDd5 | 252 | 0.353435158 | 0 | 0 | 0 | 0 | 8 mo | WILD |
| 1011080637 | 2 | AUDd5 | 252 | 0.353435158 | 0.000269478 | 10 | 0.000762455 | 28.29373305 | 8 mo | HOM |
| 1013609713 | 2 | AUDd5 | 252 | 0.353435158 | 0 | 0 | 0 | 0 | 2 mo | WILD |
| 1010418842 | 1 | AUDd5 | 252 | 0.353435158 | 0.000240297 | 12 | 0.000679889 | 33.95247966 | 8 mo | HOM |
| 1013288213 | 2 | AUDd5 | 252 | 0.353435158 | 5.89E-05 | 3 | 0.000166546 | 8.488119915 | 4 mo | HOM |
| 1008976060 | 2 | AUDd5 | 252 | 0.353435158 | 1.54E-06 | 0 | 4.37E-06 | 0 | 8 mo | WILD |
| 1013446243 | 1 | AUDd5 | 252 | 0.353435158 | 0.000102256 | 5 | 0.000289321 | 14.14686653 | 4 mo | HOM |
| 1013288951 | 2 | AUDd5 | 252 | 0.353435158 | 6.46E-05 | 4 | 0.000182882 | 11.31749322 | 4 mo | HOM |
| 1013978665 | 1 | RSP | 254 | 10.82110833 | 0.000150014 | 5 | 1.39E-05 | 0.462059879 | 2 mo | HOM |
| 1013445609 | 1 | RSP | 254 | 10.82110833 | 0.000579014 | 36 | 5.35E-05 | 3.326831126 | 4 mo | HOM |
| 1013609015 | 1 | RSP | 254 | 10.82110833 | 0.003109887 | 101 | 0.000287391 | 9.333609548 | 4 mo | HOM |
| 1013608106 | 2 | RSP | 254 | 10.82110833 | 0 | 0 | 0 | 0 | 4 mo | HOM |
| 1011937543 | 1 | RSP | 254 | 10.82110833 | 0.017457331 | 194 | 0.001613266 | 17.92792329 | 8 mo | HOM |
| 1010419538 | 2 | RSP | 254 | 10.72609113 | 0.000554572 | 19 | 5.17E-05 | 1.771381556 | 4 mo | WILD |
| 1010419187 | 1 | RSP | 254 | 10.82110833 | 0.027936871 | 455 | 0.002581701 | 42.04744895 | 8 mo | HOM |

| | | | | | | | | | | |
|---|---|---|---|---|---|---|---|---|---|---|
| **1013978366** | 1 | RSP | 254 | 10.82110833 | 0.002281084 | 30 | 0.0002108 | 2.772359272 | 2 mo | HOM |
| **1014416155** | 2 | RSP | 254 | 10.82110833 | 0 | 0 | 0 | 0 | 2 mo | HOM |
| **1013978065** | 2 | RSP | 254 | 10.82110833 | 8.63E-09 | 0 | 7.98E-10 | 0 | 2 mo | HOM |
| **1009934102** | 2 | RSP | 254 | 10.82110833 | 0.000370058 | 15 | 3.42E-05 | 1.386179636 | 4 mo | WILD |
| **1009322937** | 2 | RSP | 254 | 10.82110833 | 8.49E-05 | 6 | 7.85E-06 | 0.554471854 | 8 mo | WILD |
| **1011080637** | 2 | RSP | 254 | 10.82110833 | 0.013122353 | 356 | 0.001212663 | 32.89866336 | 8 mo | HOM |
| **1013609713** | 2 | RSP | 254 | 10.82110833 | 3.83E-05 | 2 | 3.54E-06 | 0.184823951 | 2 mo | WILD |
| **1010418842** | 1 | RSP | 254 | 10.82110833 | 0.013291661 | 425 | 0.001228309 | 39.27508968 | 8 mo | HOM |
| **1013288213** | 2 | RSP | 254 | 10.82110833 | 0.00060282 | 34 | 5.57E-05 | 3.142007175 | 4 mo | HOM |
| **1008976060** | 2 | RSP | 254 | 10.82110833 | 0.000720331 | 32 | 6.66E-05 | 2.957183223 | 8 mo | WILD |
| **1013446243** | 1 | RSP | 254 | 10.82110833 | 0.001665606 | 81 | 0.000153922 | 7.485370033 | 4 mo | HOM |
| **1013288951** | 2 | RSP | 254 | 10.82110833 | 0.005574777 | 82 | 0.000515176 | 7.577782009 | 4 mo | HOM |
| **1013978665** | 1 | AV | 255 | 0.428165646 | 0 | 0 | 0 | 0 | 2 mo | HOM |
| **1013445609** | 1 | AV | 255 | 0.428165646 | 2.15E-05 | 2 | 5.03E-05 | 4.67108938 | 4 mo | HOM |
| **1013609015** | 1 | AV | 255 | 0.428165646 | 0.000160049 | 2 | 0.000373802 | 4.67108938 | 4 mo | HOM |
| **1013608106** | 2 | AV | 255 | 0.428165646 | 0 | 0 | 0 | 0 | 4 mo | HOM |
| **1011937543** | 1 | AV | 255 | 0.428165646 | 0.000258708 | 2 | 0.000604225 | 4.67108938 | 8 mo | HOM |
| **1010419538** | 2 | AV | 255 | 0.428165646 | 0 | 0 | 0 | 0 | 4 mo | WILD |
| **1010419187** | 1 | AV | 255 | 0.428165646 | 0.002043467 | 18 | 0.004772608 | 42.03980442 | 8 mo | HOM |
| **1013978366** | 1 | AV | 255 | 0.428165646 | 0 | 0 | 0 | 0 | 2 mo | HOM |
| **1014416155** | 2 | AV | 255 | 0.428165646 | 0 | 0 | 0 | 0 | 2 mo | HOM |
| **1013978065** | 2 | AV | 255 | 0.428165646 | 0 | 0 | 0 | 0 | 2 mo | HOM |
| **1009934102** | 2 | AV | 255 | 0.428165646 | 0 | 0 | 0 | 0 | 4 mo | WILD |
| **1009322937** | 2 | AV | 255 | 0.428165646 | 7.98E-08 | 0 | 1.86E-07 | 0 | 8 mo | WILD |
| **1011080637** | 2 | AV | 255 | 0.428165646 | 0.000723344 | 15 | 0.001689402 | 35.03317035 | 8 mo | HOM |
| **1013609713** | 2 | AV | 255 | 0.428165646 | 0 | 0 | 0 | 0 | 2 mo | WILD |
| **1010418842** | 1 | AV | 255 | 0.428165646 | 0.001239773 | 15 | 0.002895546 | 35.03317035 | 8 mo | HOM |
| **1013288213** | 2 | AV | 255 | 0.428165646 | 4.85E-06 | 1 | 1.13E-05 | 2.33554469 | 4 mo | HOM |
| **1008976060** | 2 | AV | 255 | 0.428165646 | 1.76E-05 | 0 | 4.12E-05 | 0 | 8 mo | WILD |
| **1013446243** | 1 | AV | 255 | 0.428165646 | 2.12E-05 | 2 | 4.96E-05 | 4.67108938 | 4 mo | HOM |
| **1013288951** | 2 | AV | 255 | 0.428165646 | 6.52E-06 | 1 | 1.52E-05 | 2.33554469 | 4 mo | HOM |
| **1013978665** | 1 | VISpm6a | 257 | 0.156567417 | 0 | 0 | 0 | 0 | 2 mo | HOM |
| **1013445609** | 1 | VISpm6a | 257 | 0.156567417 | 1.43E-05 | 1 | 9.15E-05 | 6.387024974 | 4 mo | HOM |
| **1013609015** | 1 | VISpm6a | 257 | 0.156567417 | 6.76E-06 | 0 | 4.32E-05 | 0 | 4 mo | HOM |
| **1013608106** | 2 | VISpm6a | 257 | 0.156567417 | 0 | 0 | 0 | 0 | 4 mo | HOM |
| **1011937543** | 1 | VISpm6a | 257 | 0.156567417 | 0.000188346 | 3 | 0.001202968 | 19.16107492 | 8 mo | HOM |
| **1010419538** | 2 | VISpm6a | 257 | 0.156567417 | 0 | 0 | 0 | 0 | 4 mo | WILD |
| **1010419187** | 1 | VISpm6a | 257 | 0.156567417 | 0.000269223 | 11 | 0.001719532 | 70.25727472 | 8 mo | HOM |
| **1013978366** | 1 | VISpm6a | 257 | 0.156567417 | 0 | 0 | 0 | 0 | 2 mo | HOM |
| **1014416155** | 2 | VISpm6a | 257 | 0.156567417 | 0 | 0 | 0 | 0 | 2 mo | HOM |
| **1013978065** | 2 | VISpm6a | 257 | 0.156567417 | 0 | 0 | 0 | 0 | 2 mo | HOM |
| **1009934102** | 2 | VISpm6a | 257 | 0.156567417 | 0 | 0 | 0 | 0 | 4 mo | WILD |
| **1009322937** | 2 | VISpm6a | 257 | 0.156567417 | 0 | 0 | 0 | 0 | 8 mo | WILD |
| **1011080637** | 2 | VISpm6a | 257 | 0.156567417 | 5.52E-05 | 3 | 0.000352396 | 19.16107492 | 8 mo | HOM |
| **1013609713** | 2 | VISpm6a | 257 | 0.156567417 | 0 | 0 | 0 | 0 | 2 mo | WILD |
| **1010418842** | 1 | VISpm6a | 257 | 0.156567417 | 0.000158272 | 6 | 0.00101089 | 38.32214985 | 8 mo | HOM |
| **1013288213** | 2 | VISpm6a | 257 | 0.156567417 | 1.32E-05 | 2 | 8.44E-05 | 12.77404995 | 4 mo | HOM |
| **1008976060** | 2 | VISpm6a | 257 | 0.156567417 | 0 | 0 | 0 | 0 | 8 mo | WILD |
| **1013446243** | 1 | VISpm6a | 257 | 0.156567417 | 4.09E-05 | 2 | 0.000261191 | 12.77404995 | 4 mo | HOM |
| **1013288951** | 2 | VISpm6a | 257 | 0.156567417 | 1.44E-05 | 0 | 9.21E-05 | 0 | 4 mo | HOM |
| **1013978665** | 1 | LSr | 258 | 1.938794838 | 0 | 0 | 0 | 0 | 2 mo | HOM |
| **1013445609** | 1 | LSr | 258 | 1.938794838 | 5.82E-07 | 0 | 3E-07 | 0 | 4 mo | HOM |
| **1013609015** | 1 | LSr | 258 | 1.938794838 | 0.000104233 | 4 | 5.38E-05 | 2.063137327 | 4 mo | HOM |
| **1013608106** | 2 | LSr | 258 | 1.938794838 | 0 | 0 | 0 | 0 | 4 mo | HOM |
| **1011937543** | 1 | LSr | 258 | 1.938794838 | 0.000623472 | 27 | 0.000321577 | 13.92617696 | 8 mo | HOM |
| **1010419538** | 2 | LSr | 258 | 1.938794838 | 0 | 0 | 0 | 0 | 4 mo | WILD |
| **1010419187** | 1 | LSr | 258 | 1.938794838 | 0.001760293 | 54 | 0.000907932 | 27.85235391 | 8 mo | HOM |
| **1013978366** | 1 | LSr | 258 | 1.938794838 | 0 | 0 | 0 | 0 | 2 mo | HOM |
| **1014416155** | 2 | LSr | 258 | 1.938794838 | 0 | 0 | 0 | 0 | 2 mo | HOM |

| | | | | | | | | | | |
|---|---|---|---|---|---|---|---|---|---|---|
| **1013978065** | 2 | LSr | 258 | 1.938794838 | 0 | 0 | 0 | 0 | 2 mo | HOM |
| **1009934102** | 2 | LSr | 258 | 1.938794838 | 0 | 0 | 0 | 0 | 4 mo | WILD |
| **1009322937** | 2 | LSr | 258 | 1.938794838 | 0 | 0 | 0 | 0 | 8 mo | WILD |
| **1011080637** | 2 | LSr | 258 | 1.938794838 | 0.001331353 | 42 | 0.000686691 | 21.66294193 | 8 mo | HOM |
| **1013609713** | 2 | LSr | 258 | 1.938794838 | 0 | 0 | 0 | 0 | 2 mo | WILD |
| **1010418842** | 1 | LSr | 258 | 1.938794838 | 0.003708283 | 80 | 0.001912674 | 41.26274653 | 8 mo | HOM |
| **1013288213** | 2 | LSr | 258 | 1.938794838 | 6.83E-05 | 4 | 3.52E-05 | 2.063137327 | 4 mo | HOM |
| **1008976060** | 2 | LSr | 258 | 1.938794838 | 1.84E-05 | 3 | 9.5E-06 | 1.547352995 | 8 mo | WILD |
| **1013446243** | 1 | LSr | 258 | 1.938794838 | 7.92E-05 | 4 | 4.09E-05 | 2.063137327 | 4 mo | HOM |
| **1013288951** | 2 | LSr | 258 | 1.938794838 | 5.47E-05 | 3 | 2.82E-05 | 1.547352995 | 4 mo | HOM |
| **1013978665** | 1 | NLOT1 | 260 | 0.11487215 | 0 | 0 | 0 | 0 | 2 mo | HOM |
| **1013445609** | 1 | NLOT1 | 260 | 0.11487215 | 0 | 0 | 0 | 0 | 4 mo | HOM |
| **1013609015** | 1 | NLOT1 | 260 | 0.11487215 | 0 | 0 | 0 | 0 | 4 mo | HOM |
| **1013608106** | 2 | NLOT1 | 260 | 0.11487215 | 0 | 0 | 0 | 0 | 4 mo | HOM |
| **1011937543** | 1 | NLOT1 | 260 | 0.11487215 | 5.07E-05 | 0 | 0.000441613 | 0 | 8 mo | HOM |
| **1010419538** | 2 | NLOT1 | 260 | 0.11487215 | 0 | 0 | 0 | 0 | 4 mo | WILD |
| **1010419187** | 1 | NLOT1 | 260 | 0.11487215 | 1.89E-05 | 0 | 0.000164483 | 0 | 8 mo | HOM |
| **1013978366** | 1 | NLOT1 | 260 | 0.11487215 | 0 | 0 | 0 | 0 | 2 mo | HOM |
| **1014416155** | 2 | NLOT1 | 260 | 0.11487215 | 0 | 0 | 0 | 0 | 2 mo | HOM |
| **1013978065** | 2 | NLOT1 | 260 | 0.11487215 | 0 | 0 | 0 | 0 | 2 mo | HOM |
| **1009934102** | 2 | NLOT1 | 260 | 0.11487215 | 0 | 0 | 0 | 0 | 4 mo | WILD |
| **1009322937** | 2 | NLOT1 | 260 | 0.11487215 | 0 | 0 | 0 | 0 | 8 mo | WILD |
| **1011080637** | 2 | NLOT1 | 260 | 0.11487215 | 2.72E-05 | 3 | 0.000236755 | 26.1159906 | 8 mo | HOM |
| **1013609713** | 2 | NLOT1 | 260 | 0.11487215 | 0 | 0 | 0 | 0 | 2 mo | WILD |
| **1010418842** | 1 | NLOT1 | 260 | 0.11487215 | 3.37E-05 | 2 | 0.000293162 | 17.4106604 | 8 mo | HOM |
| **1013288213** | 2 | NLOT1 | 260 | 0.11487215 | 5.2E-07 | 0 | 4.53E-06 | 0 | 4 mo | HOM |
| **1008976060** | 2 | NLOT1 | 260 | 0.11487215 | 0 | 0 | 0 | 0 | 8 mo | WILD |
| **1013446243** | 1 | NLOT1 | 260 | 0.11487215 | 0 | 0 | 0 | 0 | 4 mo | HOM |
| **1013288951** | 2 | NLOT1 | 260 | 0.11487215 | 0 | 0 | 0 | 0 | 4 mo | HOM |
| **1013978665** | 1 | RT | 262 | 1.491106534 | 0 | 0 | 0 | 0 | 2 mo | HOM |
| **1013445609** | 1 | RT | 262 | 1.491106534 | 0 | 0 | 0 | 0 | 4 mo | HOM |
| **1013609015** | 1 | RT | 262 | 1.491106534 | 0 | 0 | 0 | 0 | 4 mo | HOM |
| **1013608106** | 2 | RT | 262 | 1.491106534 | 0 | 0 | 0 | 0 | 4 mo | HOM |
| **1011937543** | 1 | RT | 262 | 1.491106534 | 0.000275377 | 8 | 0.00018468 | 5.365143144 | 8 mo | HOM |
| **1010419538** | 2 | RT | 262 | 1.491106534 | 0 | 0 | 0 | 0 | 4 mo | WILD |
| **1010419187** | 1 | RT | 262 | 1.491106534 | 0.000516646 | 21 | 0.000346485 | 14.08350075 | 8 mo | HOM |
| **1013978366** | 1 | RT | 262 | 1.491106534 | 0 | 0 | 0 | 0 | 2 mo | HOM |
| **1014416155** | 2 | RT | 262 | 1.491106534 | 0 | 0 | 0 | 0 | 2 mo | HOM |
| **1013978065** | 2 | RT | 262 | 1.491106534 | 0 | 0 | 0 | 0 | 2 mo | HOM |
| **1009934102** | 2 | RT | 262 | 1.491106534 | 0 | 0 | 0 | 0 | 4 mo | WILD |
| **1009322937** | 2 | RT | 262 | 1.491106534 | 0 | 0 | 0 | 0 | 8 mo | WILD |
| **1011080637** | 2 | RT | 262 | 1.491106534 | 0.000623422 | 26 | 0.000418094 | 17.43671522 | 8 mo | HOM |
| **1013609713** | 2 | RT | 262 | 1.491106534 | 0 | 0 | 0 | 0 | 2 mo | WILD |
| **1010418842** | 1 | RT | 262 | 1.491106534 | 0.001040362 | 41 | 0.000697711 | 27.49635861 | 8 mo | HOM |
| **1013288213** | 2 | RT | 262 | 1.491106534 | 5.69E-06 | 1 | 3.81E-06 | 0.670642893 | 4 mo | HOM |
| **1008976060** | 2 | RT | 262 | 1.491106534 | 3.91E-05 | 7 | 2.62E-05 | 4.694500251 | 8 mo | WILD |
| **1013446243** | 1 | RT | 262 | 1.491106534 | 1.93E-07 | 0 | 1.3E-07 | 0 | 4 mo | HOM |
| **1013288951** | 2 | RT | 262 | 1.491106534 | 0 | 0 | 0 | 0 | 4 mo | HOM |
| **1013978665** | 1 | AVP | 263 | 0.09535351 | 0 | 0 | 0 | 0 | 2 mo | HOM |
| **1013445609** | 1 | AVP | 263 | 0.09535351 | 0 | 0 | 0 | 0 | 4 mo | HOM |
| **1013609015** | 1 | AVP | 263 | 0.09535351 | 0 | 0 | 0 | 0 | 4 mo | HOM |
| **1013608106** | 2 | AVP | 263 | 0.09535351 | 0 | 0 | 0 | 0 | 4 mo | HOM |
| **1011937543** | 1 | AVP | 263 | 0.09535351 | 0 | 0 | 0 | 0 | 8 mo | HOM |
| **1010419538** | 2 | AVP | 263 | 0.09535351 | 0 | 0 | 0 | 0 | 4 mo | WILD |
| **1010419187** | 1 | AVP | 263 | 0.09535351 | 1.33E-05 | 1 | 0.00013975 | 10.48729092 | 8 mo | HOM |
| **1013978366** | 1 | AVP | 263 | 0.09535351 | 0 | 0 | 0 | 0 | 2 mo | HOM |
| **1014416155** | 2 | AVP | 263 | 0.09535351 | 0 | 0 | 0 | 0 | 2 mo | HOM |
| **1013978065** | 2 | AVP | 263 | 0.09535351 | 0 | 0 | 0 | 0 | 2 mo | HOM |
| **1009934102** | 2 | AVP | 263 | 0.09535351 | 0 | 0 | 0 | 0 | 4 mo | WILD |

| | | | | | | | | | | |
|---|---|---|---|---|---|---|---|---|---|---|
| **1009322937** | 2 | AVP | 263 | 0.09535351 | 0 | 0 | 0 | 0 | 8 mo | WILD |
| **1011080637** | 2 | AVP | 263 | 0.09535351 | 0 | 0 | 0 | 0 | 8 mo | HOM |
| **1013609713** | 2 | AVP | 263 | 0.09535351 | 0 | 0 | 0 | 0 | 2 mo | WILD |
| **1010418842** | 1 | AVP | 263 | 0.09535351 | 2.46E-05 | 1 | 0.000258459 | 10.48729092 | 8 mo | HOM |
| **1013288213** | 2 | AVP | 263 | 0.09535351 | 0 | 0 | 0 | 0 | 4 mo | HOM |
| **1008976060** | 2 | AVP | 263 | 0.09535351 | 0 | 0 | 0 | 0 | 8 mo | WILD |
| **1013446243** | 1 | AVP | 263 | 0.09535351 | 0 | 0 | 0 | 0 | 4 mo | HOM |
| **1013288951** | 2 | AVP | 263 | 0.09535351 | 0 | 0 | 0 | 0 | 4 mo | HOM |
| **1013978665** | 1 | LSv | 266 | 0.625589703 | 0 | 0 | 0 | 0 | 2 mo | HOM |
| **1013445609** | 1 | LSv | 266 | 0.625589703 | 0 | 0 | 0 | 0 | 4 mo | HOM |
| **1013609015** | 1 | LSv | 266 | 0.625589703 | 4.18E-05 | 0 | 6.68E-05 | 0 | 4 mo | HOM |
| **1013608106** | 2 | LSv | 266 | 0.625589703 | 0 | 0 | 0 | 0 | 4 mo | HOM |
| **1011937543** | 1 | LSv | 266 | 0.625589703 | 0.000354638 | 18 | 0.000566887 | 28.77285209 | 8 mo | HOM |
| **1010419538** | 2 | LSv | 266 | 0.625589703 | 0 | 0 | 0 | 0 | 4 mo | WILD |
| **1010419187** | 1 | LSv | 266 | 0.625589703 | 0.000372833 | 9 | 0.000595971 | 14.38642605 | 8 mo | HOM |
| **1013978366** | 1 | LSv | 266 | 0.625589703 | 0 | 0 | 0 | 0 | 2 mo | HOM |
| **1014416155** | 2 | LSv | 266 | 0.625589703 | 0 | 0 | 0 | 0 | 2 mo | HOM |
| **1013978065** | 2 | LSv | 266 | 0.625589703 | 0 | 0 | 0 | 0 | 2 mo | HOM |
| **1009934102** | 2 | LSv | 266 | 0.625589703 | 0 | 0 | 0 | 0 | 4 mo | WILD |
| **1009322937** | 2 | LSv | 266 | 0.625589703 | 0 | 0 | 0 | 0 | 8 mo | WILD |
| **1011080637** | 2 | LSv | 266 | 0.625589703 | 0.000194477 | 7 | 0.00031087 | 11.18944248 | 8 mo | HOM |
| **1013609713** | 2 | LSv | 266 | 0.625589703 | 0 | 0 | 0 | 0 | 2 mo | WILD |
| **1010418842** | 1 | LSv | 266 | 0.625589703 | 0.001235271 | 15 | 0.00197457 | 23.97737674 | 8 mo | HOM |
| **1013288213** | 2 | LSv | 266 | 0.625589703 | 2.93E-07 | 0 | 4.68E-07 | 0 | 4 mo | HOM |
| **1008976060** | 2 | LSv | 266 | 0.625589703 | 8.02E-05 | 7 | 0.000128157 | 11.18944248 | 8 mo | WILD |
| **1013446243** | 1 | LSv | 266 | 0.625589703 | 1.52E-05 | 0 | 2.43E-05 | 0 | 4 mo | HOM |
| **1013288951** | 2 | LSv | 266 | 0.625589703 | 6.41E-05 | 1 | 0.000102425 | 1.598491783 | 4 mo | HOM |
| **1013978665** | 1 | NLOT2 | 268 | 0.154101972 | 0 | 0 | 0 | 0 | 2 mo | HOM |
| **1013445609** | 1 | NLOT2 | 268 | 0.154103118 | 0 | 0 | 0 | 0 | 4 mo | HOM |
| **1013609015** | 1 | NLOT2 | 268 | 0.154103118 | 0 | 0 | 0 | 0 | 4 mo | HOM |
| **1013608106** | 2 | NLOT2 | 268 | 0.154103118 | 0 | 0 | 0 | 0 | 4 mo | HOM |
| **1011937543** | 1 | NLOT2 | 268 | 0.154103118 | 6.11E-05 | 1 | 0.000396414 | 6.489161351 | 8 mo | HOM |
| **1010419538** | 2 | NLOT2 | 268 | 0.154103118 | 0 | 0 | 0 | 0 | 4 mo | WILD |
| **1010419187** | 1 | NLOT2 | 268 | 0.154103118 | 5.24E-05 | 3 | 0.000339904 | 19.46748405 | 8 mo | HOM |
| **1013978366** | 1 | NLOT2 | 268 | 0.154103118 | 0 | 0 | 0 | 0 | 2 mo | HOM |
| **1014416155** | 2 | NLOT2 | 268 | 0.154103118 | 0 | 0 | 0 | 0 | 2 mo | HOM |
| **1013978065** | 2 | NLOT2 | 268 | 0.154103118 | 0 | 0 | 0 | 0 | 2 mo | HOM |
| **1009934102** | 2 | NLOT2 | 268 | 0.154103118 | 0 | 0 | 0 | 0 | 4 mo | WILD |
| **1009322937** | 2 | NLOT2 | 268 | 0.154103118 | 0 | 0 | 0 | 0 | 8 mo | WILD |
| **1011080637** | 2 | NLOT2 | 268 | 0.154103118 | 0.000126125 | 2 | 0.000818444 | 12.9783227 | 8 mo | HOM |
| **1013609713** | 2 | NLOT2 | 268 | 0.154103118 | 0 | 0 | 0 | 0 | 2 mo | WILD |
| **1010418842** | 1 | NLOT2 | 268 | 0.154103118 | 0.000177499 | 5 | 0.001151822 | 32.44580676 | 8 mo | HOM |
| **1013288213** | 2 | NLOT2 | 268 | 0.154103118 | 0 | 0 | 0 | 0 | 4 mo | HOM |
| **1008976060** | 2 | NLOT2 | 268 | 0.154103118 | 0 | 0 | 0 | 0 | 8 mo | WILD |
| **1013446243** | 1 | NLOT2 | 268 | 0.154103118 | 0 | 0 | 0 | 0 | 4 mo | HOM |
| **1013288951** | 2 | NLOT2 | 268 | 0.154103118 | 0 | 0 | 0 | 0 | 4 mo | HOM |
| **1013978665** | 1 | VISpl2/3 | 269 | 0.211350661 | 0 | 0 | 0 | 0 | 2 mo | HOM |
| **1013445609** | 1 | VISpl2/3 | 269 | 0.211350661 | 2.54E-05 | 1 | 0.000120185 | 4.731473251 | 4 mo | HOM |
| **1013609015** | 1 | VISpl2/3 | 269 | 0.211350661 | 7.45E-05 | 3 | 0.000352527 | 14.19441975 | 4 mo | HOM |
| **1013608106** | 2 | VISpl2/3 | 269 | 0.211350661 | 0 | 0 | 0 | 0 | 4 mo | HOM |
| **1011937543** | 1 | VISpl2/3 | 269 | 0.211350661 | 0.000368592 | 2 | 0.001743985 | 9.462946502 | 8 mo | HOM |
| **1010419538** | 2 | VISpl2/3 | 269 | 0.211350661 | 3.29E-06 | 0 | 1.56E-05 | 0 | 4 mo | WILD |
| **1010419187** | 1 | VISpl2/3 | 269 | 0.211350661 | 0.000435171 | 7 | 0.002059002 | 33.12031276 | 8 mo | HOM |
| **1013978366** | 1 | VISpl2/3 | 269 | 0.211350661 | 9.2E-06 | 0 | 4.35E-05 | 0 | 2 mo | HOM |
| **1014416155** | 2 | VISpl2/3 | 269 | 0.211350661 | 0 | 0 | 0 | 0 | 2 mo | HOM |
| **1013978065** | 2 | VISpl2/3 | 269 | 0.211350661 | 0 | 0 | 0 | 0 | 2 mo | HOM |
| **1009934102** | 2 | VISpl2/3 | 269 | 0.211350661 | 4.92E-07 | 0 | 2.33E-06 | 0 | 4 mo | WILD |
| **1009322937** | 2 | VISpl2/3 | 269 | 0.211350661 | 5.51E-24 | 0 | 2.61E-23 | 0 | 8 mo | WILD |
| **1011080637** | 2 | VISpl2/3 | 269 | 0.211350661 | 0.000144048 | 7 | 0.000681561 | 33.12031276 | 8 mo | HOM |

| | | | | | | | | | | |
|---|---|---|---|---|---|---|---|---|---|---|
| 1013609713 | 2 | VISpl2/3 | 269 | 0.211350661 | 2.83E-07 | 0 | 1.34E-06 | 0 | 2 mo | WILD |
| 1010418842 | 1 | VISpl2/3 | 269 | 0.211350661 | 0.000562143 | 12 | 0.002659766 | 56.77767901 | 8 mo | HOM |
| 1013288213 | 2 | VISpl2/3 | 269 | 0.211350661 | 2.45E-09 | 0 | 1.16E-08 | 0 | 4 mo | HOM |
| 1008976060 | 2 | VISpl2/3 | 269 | 0.211350661 | 0 | 0 | 0 | 0 | 8 mo | WILD |
| 1013446243 | 1 | VISpl2/3 | 269 | 0.211350661 | 7E-05 | 1 | 0.000331344 | 4.731473251 | 4 mo | HOM |
| 1013288951 | 2 | VISpl2/3 | 269 | 0.211350661 | 8.76E-05 | 2 | 0.00041432 | 9.462946502 | 4 mo | HOM |
| 1013978665 | 1 | SAG | 271 | 0.101157795 | 0 | 0 | 0 | 0 | 2 mo | HOM |
| 1013445609 | 1 | SAG | 271 | 0.101157795 | 0 | 0 | 0 | 0 | 4 mo | HOM |
| 1013609015 | 1 | SAG | 271 | 0.101157795 | 0 | 0 | 0 | 0 | 4 mo | HOM |
| 1013608106 | 2 | SAG | 271 | 0.101157795 | 0 | 0 | 0 | 0 | 4 mo | HOM |
| 1011937543 | 1 | SAG | 271 | 0.101157795 | 6.57E-07 | 0 | 6.5E-06 | 0 | 8 mo | HOM |
| 1010419538 | 2 | SAG | 271 | 0.101157795 | 0 | 0 | 0 | 0 | 4 mo | WILD |
| 1010419187 | 1 | SAG | 271 | 0.101157795 | 1.14E-05 | 1 | 0.00011245 | 9.885545627 | 8 mo | HOM |
| 1013978366 | 1 | SAG | 271 | 0.101157795 | 0 | 0 | 0 | 0 | 2 mo | HOM |
| 1014416155 | 2 | SAG | 271 | 0.101157795 | 0 | 0 | 0 | 0 | 2 mo | HOM |
| 1013978065 | 2 | SAG | 271 | 0.101157795 | 0 | 0 | 0 | 0 | 2 mo | HOM |
| 1009934102 | 2 | SAG | 271 | 0.101157795 | 0 | 0 | 0 | 0 | 4 mo | WILD |
| 1009322937 | 2 | SAG | 271 | 0.101157795 | 0 | 0 | 0 | 0 | 8 mo | WILD |
| 1011080637 | 2 | SAG | 271 | 0.101157795 | 1.41E-05 | 1 | 0.000139816 | 9.885545627 | 8 mo | HOM |
| 1013609713 | 2 | SAG | 271 | 0.101157795 | 0 | 0 | 0 | 0 | 2 mo | WILD |
| 1010418842 | 1 | SAG | 271 | 0.101157795 | 8.59E-06 | 1 | 8.49E-05 | 9.885545627 | 8 mo | HOM |
| 1013288213 | 2 | SAG | 271 | 0.101157795 | 0 | 0 | 0 | 0 | 4 mo | HOM |
| 1008976060 | 2 | SAG | 271 | 0.101157795 | 0 | 0 | 0 | 0 | 8 mo | WILD |
| 1013446243 | 1 | SAG | 271 | 0.101157795 | 0 | 0 | 0 | 0 | 4 mo | HOM |
| 1013288951 | 2 | SAG | 271 | 0.101157795 | 0 | 0 | 0 | 0 | 4 mo | HOM |
| 1013978665 | 1 | AVPV | 272 | 0.186660286 | 0 | 0 | 0 | 0 | 2 mo | HOM |
| 1013445609 | 1 | AVPV | 272 | 0.186660286 | 0 | 0 | 0 | 0 | 4 mo | HOM |
| 1013609015 | 1 | AVPV | 272 | 0.186660286 | 2.17E-23 | 0 | 1.16E-22 | 0 | 4 mo | HOM |
| 1013608106 | 2 | AVPV | 272 | 0.186660286 | 0 | 0 | 0 | 0 | 4 mo | HOM |
| 1011937543 | 1 | AVPV | 272 | 0.186660286 | 2.93E-05 | 1 | 0.000156709 | 5.357325971 | 8 mo | HOM |
| 1010419538 | 2 | AVPV | 272 | 0.186660286 | 0 | 0 | 0 | 0 | 4 mo | WILD |
| 1010419187 | 1 | AVPV | 272 | 0.186660286 | 0 | 0 | 0 | 0 | 8 mo | HOM |
| 1013978366 | 1 | AVPV | 272 | 0.186660286 | 0 | 0 | 0 | 0 | 2 mo | HOM |
| 1014416155 | 2 | AVPV | 272 | 0.186660286 | 0 | 0 | 0 | 0 | 2 mo | HOM |
| 1013978065 | 2 | AVPV | 272 | 0.186660286 | 4.11E-07 | 0 | 2.2E-06 | 0 | 2 mo | HOM |
| 1009934102 | 2 | AVPV | 272 | 0.186660286 | 1.16E-06 | 0 | 6.23E-06 | 0 | 4 mo | WILD |
| 1009322937 | 2 | AVPV | 272 | 0.186660286 | 0 | 0 | 0 | 0 | 8 mo | WILD |
| 1011080637 | 2 | AVPV | 272 | 0.186660286 | 1.8E-05 | 2 | 9.66E-05 | 10.71465194 | 8 mo | HOM |
| 1013609713 | 2 | AVPV | 272 | 0.186660286 | 0 | 0 | 0 | 0 | 2 mo | WILD |
| 1010418842 | 1 | AVPV | 272 | 0.186660286 | 0.000176184 | 1 | 0.000943873 | 5.357325971 | 8 mo | HOM |
| 1013288213 | 2 | AVPV | 272 | 0.186660286 | 1.51E-06 | 0 | 8.07E-06 | 0 | 4 mo | HOM |
| 1008976060 | 2 | AVPV | 272 | 0.186660286 | 9.65E-06 | 0 | 5.17E-05 | 0 | 8 mo | WILD |
| 1013446243 | 1 | AVPV | 272 | 0.186660286 | 0 | 0 | 0 | 0 | 4 mo | HOM |
| 1013288951 | 2 | AVPV | 272 | 0.186660286 | 0 | 0 | 0 | 0 | 4 mo | HOM |
| 1013978665 | 1 | RSPd6a | 274 | 0.779565069 | 0 | 0 | 0 | 0 | 2 mo | HOM |
| 1013445609 | 1 | RSPd6a | 274 | 0.779565069 | 3.31E-05 | 2 | 4.24E-05 | 2.565533116 | 4 mo | HOM |
| 1013609015 | 1 | RSPd6a | 274 | 0.779565069 | 0.000243685 | 7 | 0.000312591 | 8.979365906 | 4 mo | HOM |
| 1013608106 | 2 | RSPd6a | 274 | 0.779565069 | 0 | 0 | 0 | 0 | 4 mo | HOM |
| 1011937543 | 1 | RSPd6a | 274 | 0.779565069 | 0.000949813 | 13 | 0.001218388 | 16.67596525 | 8 mo | HOM |
| 1010419538 | 2 | RSPd6a | 274 | 0.770448325 | 0 | 0 | 0 | 0 | 4 mo | WILD |
| 1010419187 | 1 | RSPd6a | 274 | 0.779565069 | 0.001122469 | 27 | 0.001439866 | 34.63469706 | 8 mo | HOM |
| 1013978366 | 1 | RSPd6a | 274 | 0.779565069 | 0 | 0 | 0 | 0 | 2 mo | HOM |
| 1014416155 | 2 | RSPd6a | 274 | 0.779565069 | 0 | 0 | 0 | 0 | 2 mo | HOM |
| 1013978065 | 2 | RSPd6a | 274 | 0.779565069 | 0 | 0 | 0 | 0 | 2 mo | HOM |
| 1009934102 | 2 | RSPd6a | 274 | 0.779565069 | 0 | 0 | 0 | 0 | 4 mo | WILD |
| 1009322937 | 2 | RSPd6a | 274 | 0.779565069 | 0 | 0 | 0 | 0 | 8 mo | WILD |
| 1011080637 | 2 | RSPd6a | 274 | 0.779565069 | 0.00052391 | 21 | 0.000672054 | 26.93809772 | 8 mo | HOM |
| 1013609713 | 2 | RSPd6a | 274 | 0.779565069 | 0 | 0 | 0 | 0 | 2 mo | WILD |
| 1010418842 | 1 | RSPd6a | 274 | 0.779565069 | 0.000728004 | 28 | 0.000933859 | 35.91746362 | 8 mo | HOM |

| | | | | | | | | | | |
|---|---|---|---|---|---|---|---|---|---|---|
| **1013288213** | 2 | RSPd6a | 274 | 0.779565069 | 6.99E-06 | 0 | 8.97E-06 | 0 | 4 mo | HOM |
| **1008976060** | 2 | RSPd6a | 274 | 0.779565069 | 0 | 0 | 0 | 0 | 8 mo | WILD |
| **1013446243** | 1 | RSPd6a | 274 | 0.779565069 | 3.07E-05 | 3 | 3.94E-05 | 3.848299674 | 4 mo | HOM |
| **1013288951** | 2 | RSPd6a | 274 | 0.779565069 | 5.82E-05 | 7 | 7.47E-05 | 8.979365906 | 4 mo | HOM |
| **1013978665** | 1 | LSX | 275 | 3.671785762 | 0 | 0 | 0 | 0 | 2 mo | HOM |
| **1013445609** | 1 | LSX | 275 | 3.671785762 | 1.32E-05 | 2 | 3.58E-06 | 0.544694089 | 4 mo | HOM |
| **1013609015** | 1 | LSX | 275 | 3.671785762 | 0.000240759 | 5 | 6.56E-05 | 1.361735222 | 4 mo | HOM |
| **1013608106** | 2 | LSX | 275 | 3.671785762 | 0 | 0 | 0 | 0 | 4 mo | HOM |
| **1011937543** | 1 | LSX | 275 | 3.671785762 | 0.001190541 | 57 | 0.00032424 | 15.52378153 | 8 mo | HOM |
| **1010419538** | 2 | LSX | 275 | 3.671785762 | 0 | 0 | 0 | 0 | 4 mo | WILD |
| **1010419187** | 1 | LSX | 275 | 3.671785762 | 0.002569025 | 74 | 0.000699666 | 20.15368129 | 8 mo | HOM |
| **1013978366** | 1 | LSX | 275 | 3.671785762 | 1.76E-06 | 0 | 4.81E-07 | 0 | 2 mo | HOM |
| **1014416155** | 2 | LSX | 275 | 3.671785762 | 0 | 0 | 0 | 0 | 2 mo | HOM |
| **1013978065** | 2 | LSX | 275 | 3.671785762 | 0 | 0 | 0 | 0 | 2 mo | HOM |
| **1009934102** | 2 | LSX | 275 | 3.671785762 | 0 | 0 | 0 | 0 | 4 mo | WILD |
| **1009322937** | 2 | LSX | 275 | 3.671785762 | 1.75E-06 | 0 | 4.76E-07 | 0 | 8 mo | WILD |
| **1011080637** | 2 | LSX | 275 | 3.671785762 | 0.00193368 | 59 | 0.000526632 | 16.06847562 | 8 mo | HOM |
| **1013609713** | 2 | LSX | 275 | 3.671785762 | 0 | 0 | 0 | 0 | 2 mo | WILD |
| **1010418842** | 1 | LSX | 275 | 3.671785762 | 0.006938407 | 125 | 0.001889655 | 34.04338055 | 8 mo | HOM |
| **1013288213** | 2 | LSX | 275 | 3.671785762 | 8.81E-05 | 5 | 2.4E-05 | 1.361735222 | 4 mo | HOM |
| **1008976060** | 2 | LSX | 275 | 3.671785762 | 0.000270774 | 20 | 7.37E-05 | 5.446940889 | 8 mo | WILD |
| **1013446243** | 1 | LSX | 275 | 3.671785762 | 0.00019358 | 7 | 5.27E-05 | 1.906429311 | 4 mo | HOM |
| **1013288951** | 2 | LSX | 275 | 3.671785762 | 0.000277055 | 8 | 7.55E-05 | 2.178776355 | 4 mo | HOM |
| **1013978665** | 1 | sAMY | 278 | 4.134103518 | 8.08E-07 | 0 | 1.95E-07 | 0 | 2 mo | HOM |
| **1013445609** | 1 | sAMY | 278 | 4.169963482 | 0.000145152 | 11 | 3.48E-05 | 2.637912789 | 4 mo | HOM |
| **1013609015** | 1 | sAMY | 278 | 4.169963482 | 0.000612969 | 19 | 0.000146996 | 4.556394818 | 4 mo | HOM |
| **1013608106** | 2 | sAMY | 278 | 4.169963482 | 0 | 0 | 0 | 0 | 4 mo | HOM |
| **1011937543** | 1 | sAMY | 278 | 4.169963482 | 0.008139668 | 114 | 0.001951976 | 27.33836891 | 8 mo | HOM |
| **1010419538** | 2 | sAMY | 278 | 4.169963482 | 1.47E-08 | 0 | 3.54E-09 | 0 | 4 mo | WILD |
| **1010419187** | 1 | sAMY | 278 | 4.169963482 | 0.00555197 | 180 | 0.001331419 | 43.16584564 | 8 mo | HOM |
| **1013978366** | 1 | sAMY | 278 | 4.169963482 | 1.54E-09 | 0 | 3.7E-10 | 0 | 2 mo | HOM |
| **1014416155** | 2 | sAMY | 278 | 4.169963482 | 0 | 0 | 0 | 0 | 2 mo | HOM |
| **1013978065** | 2 | sAMY | 278 | 4.169963482 | 4.81E-06 | 1 | 1.15E-06 | 0.239810254 | 2 mo | HOM |
| **1009934102** | 2 | sAMY | 278 | 4.169963482 | 2.27E-05 | 1 | 5.45E-06 | 0.239810254 | 4 mo | WILD |
| **1009322937** | 2 | sAMY | 278 | 4.169963482 | 0 | 0 | 0 | 0 | 8 mo | WILD |
| **1011080637** | 2 | sAMY | 278 | 4.169963482 | 0.004587924 | 153 | 0.001100231 | 36.6909688 | 8 mo | HOM |
| **1013609713** | 2 | sAMY | 278 | 4.169963482 | 0 | 0 | 0 | 0 | 2 mo | WILD |
| **1010418842** | 1 | sAMY | 278 | 4.169963482 | 0.006648425 | 179 | 0.001594361 | 42.92603539 | 8 mo | HOM |
| **1013288213** | 2 | sAMY | 278 | 4.169963482 | 0.000143731 | 10 | 3.45E-05 | 2.398102536 | 4 mo | HOM |
| **1008976060** | 2 | sAMY | 278 | 4.169963482 | 4.4E-05 | 3 | 1.06E-05 | 0.719430761 | 8 mo | WILD |
| **1013446243** | 1 | sAMY | 278 | 4.169963482 | 0.000329912 | 19 | 7.91E-05 | 4.556394818 | 4 mo | HOM |
| **1013288951** | 2 | sAMY | 278 | 4.169963482 | 0.000364491 | 17 | 8.74E-05 | 4.076774311 | 4 mo | HOM |
| **1013978665** | 1 | RSPagl6b | 279 | 0.035400591 | 0 | 0 | 0 | 0 | 2 mo | HOM |
| **1013445609** | 1 | RSPagl6b | 279 | 0.035400591 | 0 | 0 | 0 | 0 | 4 mo | HOM |
| **1013609015** | 1 | RSPagl6b | 279 | 0.035400591 | 5.52E-07 | 0 | 1.56E-05 | 0 | 4 mo | HOM |
| **1013608106** | 2 | RSPagl6b | 279 | 0.035400591 | 0 | 0 | 0 | 0 | 4 mo | HOM |
| **1011937543** | 1 | RSPagl6b | 279 | 0.035400591 | 7.61E-05 | 1 | 0.002149485 | 28.24811561 | 8 mo | HOM |
| **1010419538** | 2 | RSPagl6b | 279 | 0.035400591 | 0 | 0 | 0 | 0 | 4 mo | WILD |
| **1010419187** | 1 | RSPagl6b | 279 | 0.035400591 | 5.95E-05 | 3 | 0.00168054 | 84.74434683 | 8 mo | HOM |
| **1013978366** | 1 | RSPagl6b | 279 | 0.035400591 | 0 | 0 | 0 | 0 | 2 mo | HOM |
| **1014416155** | 2 | RSPagl6b | 279 | 0.035400591 | 0 | 0 | 0 | 0 | 2 mo | HOM |
| **1013978065** | 2 | RSPagl6b | 279 | 0.035400591 | 0 | 0 | 0 | 0 | 2 mo | HOM |
| **1009934102** | 2 | RSPagl6b | 279 | 0.035400591 | 0 | 0 | 0 | 0 | 4 mo | WILD |
| **1009322937** | 2 | RSPagl6b | 279 | 0.035400591 | 0 | 0 | 0 | 0 | 8 mo | WILD |
| **1011080637** | 2 | RSPagl6b | 279 | 0.035400591 | 5.57E-06 | 1 | 0.000157366 | 28.24811561 | 8 mo | HOM |
| **1013609713** | 2 | RSPagl6b | 279 | 0.035400591 | 0 | 0 | 0 | 0 | 2 mo | WILD |
| **1010418842** | 1 | RSPagl6b | 279 | 0.035400591 | 0.000102712 | 4 | 0.002901434 | 112.9924624 | 8 mo | HOM |
| **1013288213** | 2 | RSPagl6b | 279 | 0.035400591 | 0 | 0 | 0 | 0 | 4 mo | HOM |
| **1008976060** | 2 | RSPagl6b | 279 | 0.035400591 | 0 | 0 | 0 | 0 | 8 mo | WILD |

| | | | | | | | | | | |
|---|---|---|---|---|---|---|---|---|---|---|
| **1013446243** | 1 | RSPagl6b | 279 | 0.035400591 | 1.82E-05 | 1 | 0.000513272 | 28.24811561 | 4 mo | HOM |
| **1013288951** | 2 | RSPagl6b | 279 | 0.035400591 | 5.37E-06 | 1 | 0.000151775 | 28.24811561 | 4 mo | HOM |
| **1013978665** | 1 | B | 280 | 0.016036482 | 0 | 0 | 0 | 0 | 2 mo | HOM |
| **1013445609** | 1 | B | 280 | 0.016036482 | 0 | 0 | 0 | 0 | 4 mo | HOM |
| **1013609015** | 1 | B | 280 | 0.016036482 | 0 | 0 | 0 | 0 | 4 mo | HOM |
| **1013608106** | 2 | B | 280 | 0.016036482 | 0 | 0 | 0 | 0 | 4 mo | HOM |
| **1011937543** | 1 | B | 280 | 0.016036482 | 0 | 0 | 0 | 0 | 8 mo | HOM |
| **1010419538** | 2 | B | 280 | 0.016036482 | 0 | 0 | 0 | 0 | 4 mo | WILD |
| **1010419187** | 1 | B | 280 | 0.016036482 | 0 | 0 | 0 | 0 | 8 mo | HOM |
| **1013978366** | 1 | B | 280 | 0.016036482 | 0 | 0 | 0 | 0 | 2 mo | HOM |
| **1014416155** | 2 | B | 280 | 0.016036482 | 0 | 0 | 0 | 0 | 2 mo | HOM |
| **1013978065** | 2 | B | 280 | 0.016036482 | 0 | 0 | 0 | 0 | 2 mo | HOM |
| **1009934102** | 2 | B | 280 | 0.016036482 | 0 | 0 | 0 | 0 | 4 mo | WILD |
| **1009322937** | 2 | B | 280 | 0.016036482 | 0 | 0 | 0 | 0 | 8 mo | WILD |
| **1011080637** | 2 | B | 280 | 0.016036482 | 1.16E-08 | 0 | 7.25E-07 | 0 | 8 mo | HOM |
| **1013609713** | 2 | B | 280 | 0.016036482 | 0 | 0 | 0 | 0 | 2 mo | WILD |
| **1010418842** | 1 | B | 280 | 0.016036482 | 0 | 0 | 0 | 0 | 8 mo | HOM |
| **1013288213** | 2 | B | 280 | 0.016036482 | 0 | 0 | 0 | 0 | 4 mo | HOM |
| **1008976060** | 2 | B | 280 | 0.016036482 | 0 | 0 | 0 | 0 | 8 mo | WILD |
| **1013446243** | 1 | B | 280 | 0.016036482 | 0 | 0 | 0 | 0 | 4 mo | HOM |
| **1013288951** | 2 | B | 280 | 0.016036482 | 0 | 0 | 0 | 0 | 4 mo | HOM |
| **1013978665** | 1 | VISam1 | 281 | 0.140149751 | 0 | 0 | 0 | 0 | 2 mo | HOM |
| **1013445609** | 1 | VISam1 | 281 | 0.140149751 | 2.94E-06 | 0 | 2.1E-05 | 0 | 4 mo | HOM |
| **1013609015** | 1 | VISam1 | 281 | 0.140149751 | 4.17E-05 | 5 | 0.000297262 | 35.67612475 | 4 mo | HOM |
| **1013608106** | 2 | VISam1 | 281 | 0.140149751 | 0 | 0 | 0 | 0 | 4 mo | HOM |
| **1011937543** | 1 | VISam1 | 281 | 0.140149751 | 0.001049796 | 3 | 0.007490528 | 21.40567485 | 8 mo | HOM |
| **1010419538** | 2 | VISam1 | 281 | 0.140149751 | 0 | 0 | 0 | 0 | 4 mo | WILD |
| **1010419187** | 1 | VISam1 | 281 | 0.140149751 | 0.000273613 | 8 | 0.001952288 | 57.0817996 | 8 mo | HOM |
| **1013978366** | 1 | VISam1 | 281 | 0.140149751 | 0 | 0 | 0 | 0 | 2 mo | HOM |
| **1014416155** | 2 | VISam1 | 281 | 0.140149751 | 0 | 0 | 0 | 0 | 2 mo | HOM |
| **1013978065** | 2 | VISam1 | 281 | 0.140149751 | 6.08E-06 | 0 | 4.34E-05 | 0 | 2 mo | HOM |
| **1009934102** | 2 | VISam1 | 281 | 0.140149751 | 0 | 0 | 0 | 0 | 4 mo | WILD |
| **1009322937** | 2 | VISam1 | 281 | 0.140149751 | 3.18E-08 | 0 | 2.27E-07 | 0 | 8 mo | WILD |
| **1011080637** | 2 | VISam1 | 281 | 0.140149751 | 0.000154786 | 10 | 0.00110443 | 71.35224949 | 8 mo | HOM |
| **1013609713** | 2 | VISam1 | 281 | 0.140149751 | 0 | 0 | 0 | 0 | 2 mo | WILD |
| **1010418842** | 1 | VISam1 | 281 | 0.140149751 | 0.000500364 | 6 | 0.003570206 | 42.8113497 | 8 mo | HOM |
| **1013288213** | 2 | VISam1 | 281 | 0.140149751 | 2.23E-05 | 3 | 0.000159159 | 21.40567485 | 4 mo | HOM |
| **1008976060** | 2 | VISam1 | 281 | 0.140149751 | 6.55E-05 | 4 | 0.000467027 | 28.5408998 | 8 mo | WILD |
| **1013446243** | 1 | VISam1 | 281 | 0.140149751 | 5.89E-05 | 3 | 0.000420055 | 21.40567485 | 4 mo | HOM |
| **1013288951** | 2 | VISam1 | 281 | 0.140149751 | 1.53E-05 | 2 | 0.000109208 | 14.2704499 | 4 mo | HOM |
| **1013978665** | 1 | SCH | 286 | 0.06776408 | 0 | 0 | 0 | 0 | 2 mo | HOM |
| **1013445609** | 1 | SCH | 286 | 0.06776408 | 0 | 0 | 0 | 0 | 4 mo | HOM |
| **1013609015** | 1 | SCH | 286 | 0.06776408 | 0 | 0 | 0 | 0 | 4 mo | HOM |
| **1013608106** | 2 | SCH | 286 | 0.06776408 | 0 | 0 | 0 | 0 | 4 mo | HOM |
| **1011937543** | 1 | SCH | 286 | 0.06776408 | 0 | 0 | 0 | 0 | 8 mo | HOM |
| **1010419538** | 2 | SCH | 286 | 0.06776408 | 0 | 0 | 0 | 0 | 4 mo | WILD |
| **1010419187** | 1 | SCH | 286 | 0.06776408 | 0 | 0 | 0 | 0 | 8 mo | HOM |
| **1013978366** | 1 | SCH | 286 | 0.06776408 | 0 | 0 | 0 | 0 | 2 mo | HOM |
| **1014416155** | 2 | SCH | 286 | 0.06776408 | 0 | 0 | 0 | 0 | 2 mo | HOM |
| **1013978065** | 2 | SCH | 286 | 0.06776408 | 0 | 0 | 0 | 0 | 2 mo | HOM |
| **1009934102** | 2 | SCH | 286 | 0.06776408 | 0 | 0 | 0 | 0 | 4 mo | WILD |
| **1009322937** | 2 | SCH | 286 | 0.06776408 | 0 | 0 | 0 | 0 | 8 mo | WILD |
| **1011080637** | 2 | SCH | 286 | 0.06776408 | 5.24E-07 | 0 | 7.74E-06 | 0 | 8 mo | HOM |
| **1013609713** | 2 | SCH | 286 | 0.06776408 | 0 | 0 | 0 | 0 | 2 mo | WILD |
| **1010418842** | 1 | SCH | 286 | 0.06776408 | 0 | 0 | 0 | 0 | 8 mo | HOM |
| **1013288213** | 2 | SCH | 286 | 0.06776408 | 0 | 0 | 0 | 0 | 4 mo | HOM |
| **1008976060** | 2 | SCH | 286 | 0.06776408 | 0 | 0 | 0 | 0 | 8 mo | WILD |
| **1013446243** | 1 | SCH | 286 | 0.06776408 | 0 | 0 | 0 | 0 | 4 mo | HOM |
| **1013288951** | 2 | SCH | 286 | 0.06776408 | 0 | 0 | 0 | 0 | 4 mo | HOM |

| | | | | | | | | | | |
|---|---|---|---|---|---|---|---|---|---|---|
| **1013978665** | 1 | BAC | 287 | 0.007990425 | 0 | 0 | 0 | 0 | 2 mo | HOM |
| **1013445609** | 1 | BAC | 287 | 0.007990425 | 0 | 0 | 0 | 0 | 4 mo | HOM |
| **1013609015** | 1 | BAC | 287 | 0.007990425 | 0 | 0 | 0 | 0 | 4 mo | HOM |
| **1013608106** | 2 | BAC | 287 | 0.007990425 | 0 | 0 | 0 | 0 | 4 mo | HOM |
| **1011937543** | 1 | BAC | 287 | 0.007990425 | 0 | 0 | 0 | 0 | 8 mo | HOM |
| **1010419538** | 2 | BAC | 287 | 0.007990425 | 0 | 0 | 0 | 0 | 4 mo | WILD |
| **1010419187** | 1 | BAC | 287 | 0.007990425 | 0 | 0 | 0 | 0 | 8 mo | HOM |
| **1013978366** | 1 | BAC | 287 | 0.007990425 | 0 | 0 | 0 | 0 | 2 mo | HOM |
| **1014416155** | 2 | BAC | 287 | 0.007990425 | 0 | 0 | 0 | 0 | 2 mo | HOM |
| **1013978065** | 2 | BAC | 287 | 0.007990425 | 0 | 0 | 0 | 0 | 2 mo | HOM |
| **1009934102** | 2 | BAC | 287 | 0.007990425 | 0 | 0 | 0 | 0 | 4 mo | WILD |
| **1009322937** | 2 | BAC | 287 | 0.007990425 | 0 | 0 | 0 | 0 | 8 mo | WILD |
| **1011080637** | 2 | BAC | 287 | 0.007990425 | 0 | 0 | 0 | 0 | 8 mo | HOM |
| **1013609713** | 2 | BAC | 287 | 0.007990425 | 0 | 0 | 0 | 0 | 2 mo | WILD |
| **1010418842** | 1 | BAC | 287 | 0.007990425 | 9.18E-06 | 0 | 0.001148984 | 0 | 8 mo | HOM |
| **1013288213** | 2 | BAC | 287 | 0.007990425 | 0 | 0 | 0 | 0 | 4 mo | HOM |
| **1008976060** | 2 | BAC | 287 | 0.007990425 | 0 | 0 | 0 | 0 | 8 mo | WILD |
| **1013446243** | 1 | BAC | 287 | 0.007990425 | 0 | 0 | 0 | 0 | 4 mo | HOM |
| **1013288951** | 2 | BAC | 287 | 0.007990425 | 0 | 0 | 0 | 0 | 4 mo | HOM |
| **1013978665** | 1 | ORBvl2/3 | 288 | 0.539133241 | 0 | 0 | 0 | 0 | 2 mo | HOM |
| **1013445609** | 1 | ORBvl2/3 | 288 | 0.539133241 | 5.17E-05 | 1 | 9.59E-05 | 1.854829056 | 4 mo | HOM |
| **1013609015** | 1 | ORBvl2/3 | 288 | 0.539133241 | 1.26E-05 | 1 | 2.33E-05 | 1.854829056 | 4 mo | HOM |
| **1013608106** | 2 | ORBvl2/3 | 288 | 0.539133241 | 0 | 0 | 0 | 0 | 4 mo | HOM |
| **1011937543** | 1 | ORBvl2/3 | 288 | 0.539133241 | 0.000409926 | 12 | 0.000760342 | 22.25794867 | 8 mo | HOM |
| **1010419538** | 2 | ORBvl2/3 | 288 | 0.539133241 | 0 | 0 | 0 | 0 | 4 mo | WILD |
| **1010419187** | 1 | ORBvl2/3 | 288 | 0.539133241 | 0.000494751 | 17 | 0.000917678 | 31.53209395 | 8 mo | HOM |
| **1013978366** | 1 | ORBvl2/3 | 288 | 0.539133241 | 0 | 0 | 0 | 0 | 2 mo | HOM |
| **1014416155** | 2 | ORBvl2/3 | 288 | 0.539133241 | 0 | 0 | 0 | 0 | 2 mo | HOM |
| **1013978065** | 2 | ORBvl2/3 | 288 | 0.539133241 | 0 | 0 | 0 | 0 | 2 mo | HOM |
| **1009934102** | 2 | ORBvl2/3 | 288 | 0.539133241 | 0 | 0 | 0 | 0 | 4 mo | WILD |
| **1009322937** | 2 | ORBvl2/3 | 288 | 0.539133241 | 3.95E-05 | 2 | 7.33E-05 | 3.709658112 | 8 mo | WILD |
| **1011080637** | 2 | ORBvl2/3 | 288 | 0.539133241 | 0.000176054 | 12 | 0.000326551 | 22.25794867 | 8 mo | HOM |
| **1013609713** | 2 | ORBvl2/3 | 288 | 0.539133241 | 0 | 0 | 0 | 0 | 2 mo | WILD |
| **1010418842** | 1 | ORBvl2/3 | 288 | 0.539133241 | 0.000732839 | 20 | 0.00135929 | 37.09658112 | 8 mo | HOM |
| **1013288213** | 2 | ORBvl2/3 | 288 | 0.539133241 | 0.000232132 | 2 | 0.000430565 | 3.709658112 | 4 mo | HOM |
| **1008976060** | 2 | ORBvl2/3 | 288 | 0.539133241 | 5.36E-07 | 0 | 9.94E-07 | 0 | 8 mo | WILD |
| **1013446243** | 1 | ORBvl2/3 | 288 | 0.539133241 | 4.23E-05 | 1 | 7.85E-05 | 1.854829056 | 4 mo | HOM |
| **1013288951** | 2 | ORBvl2/3 | 288 | 0.539133241 | 5.87E-05 | 4 | 0.000108944 | 7.419316223 | 4 mo | HOM |
| **1013978665** | 1 | TEa5 | 289 | 0.977922557 | 0 | 0 | 0 | 0 | 2 mo | HOM |
| **1013445609** | 1 | TEa5 | 289 | 0.977922557 | 0.000112487 | 6 | 0.000115026 | 6.135455163 | 4 mo | HOM |
| **1013609015** | 1 | TEa5 | 289 | 0.977922557 | 0.000349571 | 9 | 0.000357463 | 9.203182744 | 4 mo | HOM |
| **1013608106** | 2 | TEa5 | 289 | 0.977922557 | 0 | 0 | 0 | 0 | 4 mo | HOM |
| **1011937543** | 1 | TEa5 | 289 | 0.977922557 | 0.001720188 | 47 | 0.001759023 | 48.06106544 | 8 mo | HOM |
| **1010419538** | 2 | TEa5 | 289 | 0.977922557 | 0 | 0 | 0 | 0 | 4 mo | WILD |
| **1010419187** | 1 | TEa5 | 289 | 0.977922557 | 0.001655076 | 65 | 0.00169244 | 66.46743093 | 8 mo | HOM |
| **1013978366** | 1 | TEa5 | 289 | 0.977922557 | 0 | 0 | 0 | 0 | 2 mo | HOM |
| **1014416155** | 2 | TEa5 | 289 | 0.977922557 | 0 | 0 | 0 | 0 | 2 mo | HOM |
| **1013978065** | 2 | TEa5 | 289 | 0.977922557 | 0 | 0 | 0 | 0 | 2 mo | HOM |
| **1009934102** | 2 | TEa5 | 289 | 0.977922557 | 0 | 0 | 0 | 0 | 4 mo | WILD |
| **1009322937** | 2 | TEa5 | 289 | 0.977922557 | 0 | 0 | 0 | 0 | 8 mo | WILD |
| **1011080637** | 2 | TEa5 | 289 | 0.977922557 | 0.000659845 | 35 | 0.000674741 | 35.79015512 | 8 mo | HOM |
| **1013609713** | 2 | TEa5 | 289 | 0.977922557 | 0 | 0 | 0 | 0 | 2 mo | WILD |
| **1010418842** | 1 | TEa5 | 289 | 0.977922557 | 0.001455327 | 44 | 0.001488183 | 44.99333786 | 8 mo | HOM |
| **1013288213** | 2 | TEa5 | 289 | 0.977922557 | 2.54E-05 | 4 | 2.6E-05 | 4.090303442 | 4 mo | HOM |
| **1008976060** | 2 | TEa5 | 289 | 0.977922557 | 0 | 0 | 0 | 0 | 8 mo | WILD |
| **1013446243** | 1 | TEa5 | 289 | 0.977922557 | 0.000214033 | 10 | 0.000218865 | 10.2257586 | 4 mo | HOM |
| **1013288951** | 2 | TEa5 | 289 | 0.977922557 | 0.000181971 | 9 | 0.000186079 | 9.203182744 | 4 mo | HOM |
| **1013978665** | 1 | LZ | 290 | 5.857434892 | 0 | 0 | 0 | 0 | 2 mo | HOM |
| **1013445609** | 1 | LZ | 290 | 5.857434892 | 2.62E-06 | 0 | 4.47E-07 | 0 | 4 mo | HOM |

| | | | | | | | | | | |
|---|---|---|---|---|---|---|---|---|---|---|
| **1013609015** | 1 | LZ | 290 | 5.857434892 | 1E-09 | 0 | 1.71E-10 | 0 | 4 mo | HOM |
| **1013608106** | 2 | LZ | 290 | 5.857434892 | 0 | 0 | 0 | 0 | 4 mo | HOM |
| **1011937543** | 1 | LZ | 290 | 5.857434892 | 0.008059628 | 112 | 0.001375966 | 19.12099785 | 8 mo | HOM |
| **1010419538** | 2 | LZ | 290 | 5.857434892 | 1.71E-07 | 0 | 2.92E-08 | 0 | 4 mo | WILD |
| **1010419187** | 1 | LZ | 290 | 5.857434892 | 0.000324987 | 16 | 5.55E-05 | 2.731571122 | 8 mo | HOM |
| **1013978366** | 1 | LZ | 290 | 5.857434892 | 7.38E-08 | 0 | 1.26E-08 | 0 | 2 mo | HOM |
| **1014416155** | 2 | LZ | 290 | 5.857434892 | 0 | 0 | 0 | 0 | 2 mo | HOM |
| **1013978065** | 2 | LZ | 290 | 5.857434892 | 3.03E-05 | 1 | 5.17E-06 | 0.170723195 | 2 mo | HOM |
| **1009934102** | 2 | LZ | 290 | 5.857434892 | 0 | 0 | 0 | 0 | 4 mo | WILD |
| **1009322937** | 2 | LZ | 290 | 5.857434892 | 0 | 0 | 0 | 0 | 8 mo | WILD |
| **1011080637** | 2 | LZ | 290 | 5.857434892 | 0.00048027 | 28 | 8.2E-05 | 4.780249463 | 8 mo | HOM |
| **1013609713** | 2 | LZ | 290 | 5.857434892 | 0 | 0 | 0 | 0 | 2 mo | WILD |
| **1010418842** | 1 | LZ | 290 | 5.857434892 | 0.002009593 | 74 | 0.000343084 | 12.63351644 | 8 mo | HOM |
| **1013288213** | 2 | LZ | 290 | 5.857434892 | 0 | 0 | 0 | 0 | 4 mo | HOM |
| **1008976060** | 2 | LZ | 290 | 5.857434892 | 4.61E-05 | 3 | 7.88E-06 | 0.512169585 | 8 mo | WILD |
| **1013446243** | 1 | LZ | 290 | 5.857434892 | 9.1E-09 | 0 | 1.55E-09 | 0 | 4 mo | HOM |
| **1013288951** | 2 | LZ | 290 | 5.857434892 | 0 | 0 | 0 | 0 | 4 mo | HOM |
| **1013978665** | 1 | BA | 292 | 0.025712356 | 0 | 0 | 0 | 0 | 2 mo | HOM |
| **1013445609** | 1 | BA | 292 | 0.025712356 | 0 | 0 | 0 | 0 | 4 mo | HOM |
| **1013609015** | 1 | BA | 292 | 0.025712356 | 0 | 0 | 0 | 0 | 4 mo | HOM |
| **1013608106** | 2 | BA | 292 | 0.025712356 | 0 | 0 | 0 | 0 | 4 mo | HOM |
| **1011937543** | 1 | BA | 292 | 0.025712356 | 9.17E-06 | 1 | 0.000356478 | 38.89180818 | 8 mo | HOM |
| **1010419538** | 2 | BA | 292 | 0.025712356 | 0 | 0 | 0 | 0 | 4 mo | WILD |
| **1010419187** | 1 | BA | 292 | 0.025712356 | 8.77E-06 | 1 | 0.00034107 | 38.89180818 | 8 mo | HOM |
| **1013978366** | 1 | BA | 292 | 0.025712356 | 0 | 0 | 0 | 0 | 2 mo | HOM |
| **1014416155** | 2 | BA | 292 | 0.025712356 | 0 | 0 | 0 | 0 | 2 mo | HOM |
| **1013978065** | 2 | BA | 292 | 0.025712356 | 0 | 0 | 0 | 0 | 2 mo | HOM |
| **1009934102** | 2 | BA | 292 | 0.025712356 | 0 | 0 | 0 | 0 | 4 mo | WILD |
| **1009322937** | 2 | BA | 292 | 0.025712356 | 0 | 0 | 0 | 0 | 8 mo | WILD |
| **1011080637** | 2 | BA | 292 | 0.025712356 | 0 | 0 | 0 | 0 | 8 mo | HOM |
| **1013609713** | 2 | BA | 292 | 0.025712356 | 0 | 0 | 0 | 0 | 2 mo | WILD |
| **1010418842** | 1 | BA | 292 | 0.025712356 | 3.51E-05 | 1 | 0.001364698 | 38.89180818 | 8 mo | HOM |
| **1013288213** | 2 | BA | 292 | 0.025712356 | 0 | 0 | 0 | 0 | 4 mo | HOM |
| **1008976060** | 2 | BA | 292 | 0.025712356 | 2.13E-06 | 0 | 8.29E-05 | 0 | 8 mo | WILD |
| **1013446243** | 1 | BA | 292 | 0.025712356 | 0 | 0 | 0 | 0 | 4 mo | HOM |
| **1013288951** | 2 | BA | 292 | 0.025712356 | 0 | 0 | 0 | 0 | 4 mo | HOM |
| **1013978665** | 1 | SCm | 294 | 5.823278894 | 8.02E-06 | 1 | 1.38E-06 | 0.171724559 | 2 mo | HOM |
| **1013445609** | 1 | SCm | 294 | 5.823278894 | 0 | 0 | 0 | 0 | 4 mo | HOM |
| **1013609015** | 1 | SCm | 294 | 5.823278894 | 7.13E-07 | 1 | 1.22E-07 | 0.171724559 | 4 mo | HOM |
| **1013608106** | 2 | SCm | 294 | 5.823278894 | 0 | 0 | 0 | 0 | 4 mo | HOM |
| **1011937543** | 1 | SCm | 294 | 5.823278894 | 0.006916589 | 134 | 0.001187748 | 23.01109091 | 8 mo | HOM |
| **1010419538** | 2 | SCm | 294 | 5.823278894 | 0 | 0 | 0 | 0 | 4 mo | WILD |
| **1010419187** | 1 | SCm | 294 | 5.823278894 | 0.000618583 | 27 | 0.000106226 | 4.636563093 | 8 mo | HOM |
| **1013978366** | 1 | SCm | 294 | 5.823278894 | 0 | 0 | 0 | 0 | 2 mo | HOM |
| **1014416155** | 2 | SCm | 294 | 5.823278894 | 0 | 0 | 0 | 0 | 2 mo | HOM |
| **1013978065** | 2 | SCm | 294 | 5.823278894 | 0 | 0 | 0 | 0 | 2 mo | HOM |
| **1009934102** | 2 | SCm | 294 | 5.823278894 | 0 | 0 | 0 | 0 | 4 mo | WILD |
| **1009322937** | 2 | SCm | 294 | 5.823278894 | 1.54E-06 | 0 | 2.65E-07 | 0 | 8 mo | WILD |
| **1011080637** | 2 | SCm | 294 | 5.823278894 | 0.000196596 | 8 | 3.38E-05 | 1.373796472 | 8 mo | HOM |
| **1013609713** | 2 | SCm | 294 | 5.823278894 | 0 | 0 | 0 | 0 | 2 mo | WILD |
| **1010418842** | 1 | SCm | 294 | 5.823278894 | 0.000537824 | 23 | 9.24E-05 | 3.949664857 | 8 mo | HOM |
| **1013288213** | 2 | SCm | 294 | 5.823278894 | 0 | 0 | 0 | 0 | 4 mo | HOM |
| **1008976060** | 2 | SCm | 294 | 5.823278894 | 0.000102079 | 11 | 1.75E-05 | 1.888970149 | 8 mo | WILD |
| **1013446243** | 1 | SCm | 294 | 5.823278894 | 8.98E-06 | 1 | 1.54E-06 | 0.171724559 | 4 mo | HOM |
| **1013288951** | 2 | SCm | 294 | 5.823278894 | 0 | 0 | 0 | 0 | 4 mo | HOM |
| **1013978665** | 1 | BLA | 295 | 1.957718011 | 0 | 0 | 0 | 0 | 2 mo | HOM |
| **1013445609** | 1 | BLA | 295 | 1.957718011 | 0.000228233 | 15 | 0.000116581 | 7.661981917 | 4 mo | HOM |
| **1013609015** | 1 | BLA | 295 | 1.957718011 | 0.000569117 | 22 | 0.000290704 | 11.23757348 | 4 mo | HOM |
| **1013608106** | 2 | BLA | 295 | 1.957718011 | 0 | 0 | 0 | 0 | 4 mo | HOM |

| | | | | | | | | | | |
|---|---|---|---|---|---|---|---|---|---|---|
| 1011937543 | 1 | BLA | 295 | 1.957718011 | 0.003478556 | 75 | 0.001776842 | 38.30990958 | 8 mo | HOM |
| 1010419538 | 2 | BLA | 295 | 1.957718011 | 0 | 0 | 0 | 0 | 4 mo | WILD |
| 1010419187 | 1 | BLA | 295 | 1.957718011 | 0.002759845 | 124 | 0.001409726 | 63.33905051 | 8 mo | HOM |
| 1013978366 | 1 | BLA | 295 | 1.957718011 | 0 | 0 | 0 | 0 | 2 mo | HOM |
| 1014416155 | 2 | BLA | 295 | 1.957718011 | 0 | 0 | 0 | 0 | 2 mo | HOM |
| 1013978065 | 2 | BLA | 295 | 1.957718011 | 0 | 0 | 0 | 0 | 2 mo | HOM |
| 1009934102 | 2 | BLA | 295 | 1.957718011 | 0 | 0 | 0 | 0 | 4 mo | WILD |
| 1009322937 | 2 | BLA | 295 | 1.957718011 | 8.21E-06 | 2 | 4.2E-06 | 1.021597589 | 8 mo | WILD |
| 1011080637 | 2 | BLA | 295 | 1.957718011 | 0.00200332 | 112 | 0.001023294 | 57.20946498 | 8 mo | HOM |
| 1013609713 | 2 | BLA | 295 | 1.957718011 | 0 | 0 | 0 | 0 | 2 mo | WILD |
| 1010418842 | 1 | BLA | 295 | 1.957718011 | 0.003869921 | 122 | 0.001976751 | 62.31745292 | 8 mo | HOM |
| 1013288213 | 2 | BLA | 295 | 1.957718011 | 0.00012074 | 10 | 6.17E-05 | 5.107987945 | 4 mo | HOM |
| 1008976060 | 2 | BLA | 295 | 1.957718011 | 7.49E-06 | 1 | 3.83E-06 | 0.510798794 | 8 mo | WILD |
| 1013446243 | 1 | BLA | 295 | 1.957718011 | 0.000322152 | 18 | 0.000164555 | 9.1943783 | 4 mo | HOM |
| 1013288951 | 2 | BLA | 295 | 1.957718011 | 0.000512475 | 26 | 0.000261772 | 13.28076866 | 4 mo | HOM |
| 1013978665 | 1 | ACAv2/3 | 296 | 0.517455249 | 0 | 0 | 0 | 0 | 2 mo | HOM |
| 1013445609 | 1 | ACAv2/3 | 296 | 0.517455249 | 2.3E-05 | 2 | 4.45E-05 | 3.865068534 | 4 mo | HOM |
| 1013609015 | 1 | ACAv2/3 | 296 | 0.517455249 | 0.000236864 | 9 | 0.000457748 | 17.3928084 | 4 mo | HOM |
| 1013608106 | 2 | ACAv2/3 | 296 | 0.517455249 | 0 | 0 | 0 | 0 | 4 mo | HOM |
| 1011937543 | 1 | ACAv2/3 | 296 | 0.517455249 | 0.000458996 | 9 | 0.000887026 | 17.3928084 | 8 mo | HOM |
| 1010419538 | 2 | ACAv2/3 | 296 | 0.517455249 | 0 | 0 | 0 | 0 | 4 mo | WILD |
| 1010419187 | 1 | ACAv2/3 | 296 | 0.517455249 | 0.000627849 | 14 | 0.001213339 | 27.05547974 | 8 mo | HOM |
| 1013978366 | 1 | ACAv2/3 | 296 | 0.517455249 | 2.14E-08 | 0 | 4.13E-08 | 0 | 2 mo | HOM |
| 1014416155 | 2 | ACAv2/3 | 296 | 0.517455249 | 0 | 0 | 0 | 0 | 2 mo | HOM |
| 1013978065 | 2 | ACAv2/3 | 296 | 0.517455249 | 0 | 0 | 0 | 0 | 2 mo | HOM |
| 1009934102 | 2 | ACAv2/3 | 296 | 0.517455249 | 0 | 0 | 0 | 0 | 4 mo | WILD |
| 1009322937 | 2 | ACAv2/3 | 296 | 0.517455249 | 1.49E-06 | 0 | 2.88E-06 | 0 | 8 mo | WILD |
| 1011080637 | 2 | ACAv2/3 | 296 | 0.517455249 | 0.000304064 | 20 | 0.000587615 | 38.65068534 | 8 mo | HOM |
| 1013609713 | 2 | ACAv2/3 | 296 | 0.517455249 | 0 | 0 | 0 | 0 | 2 mo | WILD |
| 1010418842 | 1 | ACAv2/3 | 296 | 0.517455249 | 0.000869186 | 19 | 0.001679732 | 36.71815108 | 8 mo | HOM |
| 1013288213 | 2 | ACAv2/3 | 296 | 0.517455249 | 0.000158673 | 3 | 0.000306641 | 5.797602801 | 4 mo | HOM |
| 1008976060 | 2 | ACAv2/3 | 296 | 0.517455249 | 0 | 0 | 0 | 0 | 8 mo | WILD |
| 1013446243 | 1 | ACAv2/3 | 296 | 0.517455249 | 0.000252286 | 5 | 0.000487552 | 9.662671336 | 4 mo | HOM |
| 1013288951 | 2 | ACAv2/3 | 296 | 0.517455249 | 6.82E-05 | 2 | 0.000131787 | 3.865068534 | 4 mo | HOM |
| 1013978665 | 1 | MA | 298 | 0.374277092 | 4.64E-06 | 1 | 1.24E-05 | 2.671817275 | 2 mo | HOM |
| 1013445609 | 1 | MA | 298 | 0.375286257 | 0 | 0 | 0 | 0 | 4 mo | HOM |
| 1013609015 | 1 | MA | 298 | 0.375286257 | 0 | 0 | 0 | 0 | 4 mo | HOM |
| 1013608106 | 2 | MA | 298 | 0.375286257 | 0 | 0 | 0 | 0 | 4 mo | HOM |
| 1011937543 | 1 | MA | 298 | 0.375286257 | 0.000177344 | 4 | 0.000472556 | 10.65853046 | 8 mo | HOM |
| 1010419538 | 2 | MA | 298 | 0.375286257 | 0 | 0 | 0 | 0 | 4 mo | WILD |
| 1010419187 | 1 | MA | 298 | 0.375286257 | 2.71E-05 | 3 | 7.21E-05 | 7.993897846 | 8 mo | HOM |
| 1013978366 | 1 | MA | 298 | 0.375286257 | 0 | 0 | 0 | 0 | 2 mo | HOM |
| 1014416155 | 2 | MA | 298 | 0.375286257 | 0 | 0 | 0 | 0 | 2 mo | HOM |
| 1013978065 | 2 | MA | 298 | 0.375286257 | 0 | 0 | 0 | 0 | 2 mo | HOM |
| 1009934102 | 2 | MA | 298 | 0.375286257 | 0 | 0 | 0 | 0 | 4 mo | WILD |
| 1009322937 | 2 | MA | 298 | 0.375286257 | 0 | 0 | 0 | 0 | 8 mo | WILD |
| 1011080637 | 2 | MA | 298 | 0.375286257 | 4.99E-05 | 4 | 0.000133064 | 10.65853046 | 8 mo | HOM |
| 1013609713 | 2 | MA | 298 | 0.375286257 | 0 | 0 | 0 | 0 | 2 mo | WILD |
| 1010418842 | 1 | MA | 298 | 0.375286257 | 7.22E-05 | 2 | 0.000192437 | 5.329265231 | 8 mo | HOM |
| 1013288213 | 2 | MA | 298 | 0.375286257 | 0 | 0 | 0 | 0 | 4 mo | HOM |
| 1008976060 | 2 | MA | 298 | 0.375286257 | 0 | 0 | 0 | 0 | 8 mo | WILD |
| 1013446243 | 1 | MA | 298 | 0.375286257 | 0 | 0 | 0 | 0 | 4 mo | HOM |
| 1013288951 | 2 | MA | 298 | 0.375286257 | 0 | 0 | 0 | 0 | 4 mo | HOM |
| 1013978665 | 1 | st | 301 | 0.35137265 | 0 | 0 | 0 | 0 | 2 mo | HOM |
| 1013445609 | 1 | st | 301 | 0.35137265 | 0 | 0 | 0 | 0 | 4 mo | HOM |
| 1013609015 | 1 | st | 301 | 0.35137265 | 0 | 0 | 0 | 0 | 4 mo | HOM |
| 1013608106 | 2 | st | 301 | 0.35137265 | 0 | 0 | 0 | 0 | 4 mo | HOM |
| 1011937543 | 1 | st | 301 | 0.35137265 | 0.000147297 | 1 | 0.000419205 | 2.845981328 | 8 mo | HOM |
| 1010419538 | 2 | st | 301 | 0.35137265 | 0 | 0 | 0 | 0 | 4 mo | WILD |

| | | | | | | | | | | |
|---|---|---|---|---|---|---|---|---|---|---|
| 1010419187 | 1 | st | 301 | 0.35137265 | 7.21E-05 | 5 | 0.000205233 | 14.22990664 | 8 mo | HOM |
| 1013978366 | 1 | st | 301 | 0.35137265 | 0 | 0 | 0 | 0 | 2 mo | HOM |
| 1014416155 | 2 | st | 301 | 0.35137265 | 0 | 0 | 0 | 0 | 2 mo | HOM |
| 1013978065 | 2 | st | 301 | 0.35137265 | 0 | 0 | 0 | 0 | 2 mo | HOM |
| 1009934102 | 2 | st | 301 | 0.35137265 | 0 | 0 | 0 | 0 | 4 mo | WILD |
| 1009322937 | 2 | st | 301 | 0.35137265 | 0 | 0 | 0 | 0 | 8 mo | WILD |
| 1011080637 | 2 | st | 301 | 0.35137265 | 5.46E-05 | 3 | 0.000155481 | 8.537943985 | 8 mo | HOM |
| 1013609713 | 2 | st | 301 | 0.35137265 | 0 | 0 | 0 | 0 | 2 mo | WILD |
| 1010418842 | 1 | st | 301 | 0.35137265 | 0.000208499 | 8 | 0.000593384 | 22.76785063 | 8 mo | HOM |
| 1013288213 | 2 | st | 301 | 0.35137265 | 1.13E-07 | 0 | 3.2E-07 | 0 | 4 mo | HOM |
| 1008976060 | 2 | st | 301 | 0.35137265 | 9.74E-06 | 1 | 2.77E-05 | 2.845981328 | 8 mo | WILD |
| 1013446243 | 1 | st | 301 | 0.35137265 | 4E-07 | 0 | 1.14E-06 | 0 | 4 mo | HOM |
| 1013288951 | 2 | st | 301 | 0.35137265 | 7.81E-06 | 0 | 2.22E-05 | 0 | 4 mo | HOM |
| 1013978665 | 1 | SCs | 302 | 2.221632834 | 1.86E-05 | 2 | 8.39E-06 | 0.900238765 | 2 mo | HOM |
| 1013445609 | 1 | SCs | 302 | 2.221632834 | 0 | 0 | 0 | 0 | 4 mo | HOM |
| 1013609015 | 1 | SCs | 302 | 2.221632834 | 2.74E-05 | 1 | 1.23E-05 | 0.450119383 | 4 mo | HOM |
| 1013608106 | 2 | SCs | 302 | 2.221632834 | 0 | 0 | 0 | 0 | 4 mo | HOM |
| 1011937543 | 1 | SCs | 302 | 2.221632834 | 0.00416803 | 75 | 0.001876111 | 33.7589537 | 8 mo | HOM |
| 1010419538 | 2 | SCs | 302 | 2.221632834 | 6.23E-06 | 1 | 2.8E-06 | 0.450119383 | 4 mo | WILD |
| 1010419187 | 1 | SCs | 302 | 2.221632834 | 0.000598141 | 21 | 0.000269235 | 9.452507037 | 8 mo | HOM |
| 1013978366 | 1 | SCs | 302 | 2.221632834 | 5.86E-05 | 1 | 2.64E-05 | 0.450119383 | 2 mo | HOM |
| 1014416155 | 2 | SCs | 302 | 2.221632834 | 0 | 0 | 0 | 0 | 2 mo | HOM |
| 1013978065 | 2 | SCs | 302 | 2.221632834 | 0 | 0 | 0 | 0 | 2 mo | HOM |
| 1009934102 | 2 | SCs | 302 | 2.221632834 | 2.03E-05 | 2 | 9.14E-06 | 0.900238765 | 4 mo | WILD |
| 1009322937 | 2 | SCs | 302 | 2.221632834 | 5.05E-05 | 2 | 2.27E-05 | 0.900238765 | 8 mo | WILD |
| 1011080637 | 2 | SCs | 302 | 2.221632834 | 0.000713777 | 28 | 0.000321285 | 12.60334272 | 8 mo | HOM |
| 1013609713 | 2 | SCs | 302 | 2.221632834 | 1.34E-05 | 2 | 6.02E-06 | 0.900238765 | 2 mo | WILD |
| 1010418842 | 1 | SCs | 302 | 2.221632834 | 0.000473073 | 24 | 0.000212939 | 10.80286518 | 8 mo | HOM |
| 1013288213 | 2 | SCs | 302 | 2.221632834 | 9.37E-06 | 0 | 4.22E-06 | 0 | 4 mo | HOM |
| 1008976060 | 2 | SCs | 302 | 2.221632834 | 0.000149702 | 7 | 6.74E-05 | 3.150835679 | 8 mo | WILD |
| 1013446243 | 1 | SCs | 302 | 2.221632834 | 3.93E-05 | 1 | 1.77E-05 | 0.450119383 | 4 mo | HOM |
| 1013288951 | 2 | SCs | 302 | 2.221632834 | 0.00034145 | 18 | 0.000153693 | 8.102148889 | 4 mo | HOM |
| 1013978665 | 1 | BLAa | 303 | 0.79287143 | 0 | 0 | 0 | 0 | 2 mo | HOM |
| 1013445609 | 1 | BLAa | 303 | 0.79287143 | 4.59E-05 | 4 | 5.79E-05 | 5.044954133 | 4 mo | HOM |
| 1013609015 | 1 | BLAa | 303 | 0.79287143 | 0.000344061 | 11 | 0.000433942 | 13.87362387 | 4 mo | HOM |
| 1013608106 | 2 | BLAa | 303 | 0.79287143 | 0 | 0 | 0 | 0 | 4 mo | HOM |
| 1011937543 | 1 | BLAa | 303 | 0.79287143 | 0.00100526 | 30 | 0.001267873 | 37.837156 | 8 mo | HOM |
| 1010419538 | 2 | BLAa | 303 | 0.79287143 | 0 | 0 | 0 | 0 | 4 mo | WILD |
| 1010419187 | 1 | BLAa | 303 | 0.79287143 | 0.001294349 | 56 | 0.001632483 | 70.62935787 | 8 mo | HOM |
| 1013978366 | 1 | BLAa | 303 | 0.79287143 | 0 | 0 | 0 | 0 | 2 mo | HOM |
| 1014416155 | 2 | BLAa | 303 | 0.79287143 | 0 | 0 | 0 | 0 | 2 mo | HOM |
| 1013978065 | 2 | BLAa | 303 | 0.79287143 | 0 | 0 | 0 | 0 | 2 mo | HOM |
| 1009934102 | 2 | BLAa | 303 | 0.79287143 | 0 | 0 | 0 | 0 | 4 mo | WILD |
| 1009322937 | 2 | BLAa | 303 | 0.79287143 | 8.21E-06 | 2 | 1.04E-05 | 2.522477067 | 8 mo | WILD |
| 1011080637 | 2 | BLAa | 303 | 0.79287143 | 0.000826583 | 53 | 0.001042518 | 66.84564227 | 8 mo | HOM |
| 1013609713 | 2 | BLAa | 303 | 0.79287143 | 0 | 0 | 0 | 0 | 2 mo | WILD |
| 1010418842 | 1 | BLAa | 303 | 0.79287143 | 0.001887253 | 52 | 0.002380277 | 65.58440373 | 8 mo | HOM |
| 1013288213 | 2 | BLAa | 303 | 0.79287143 | 5.71E-05 | 4 | 7.2E-05 | 5.044954133 | 4 mo | HOM |
| 1008976060 | 2 | BLAa | 303 | 0.79287143 | 0 | 0 | 0 | 0 | 8 mo | WILD |
| 1013446243 | 1 | BLAa | 303 | 0.79287143 | 0.000119713 | 10 | 0.000150987 | 12.61238533 | 4 mo | HOM |
| 1013288951 | 2 | BLAa | 303 | 0.79287143 | 0.000229132 | 12 | 0.00028899 | 15.1348624 | 4 mo | HOM |
| 1013978665 | 1 | PL2/3 | 304 | 0.433335302 | 0 | 0 | 0 | 0 | 2 mo | HOM |
| 1013445609 | 1 | PL2/3 | 304 | 0.433335302 | 5.01E-10 | 0 | 1.16E-09 | 0 | 4 mo | HOM |
| 1013609015 | 1 | PL2/3 | 304 | 0.433335302 | 0.000130537 | 6 | 0.000301238 | 13.84609093 | 4 mo | HOM |
| 1013608106 | 2 | PL2/3 | 304 | 0.433335302 | 0 | 0 | 0 | 0 | 4 mo | HOM |
| 1011937543 | 1 | PL2/3 | 304 | 0.433335302 | 0.000209375 | 6 | 0.000483171 | 13.84609093 | 8 mo | HOM |
| 1010419538 | 2 | PL2/3 | 304 | 0.433335302 | 0 | 0 | 0 | 0 | 4 mo | WILD |
| 1010419187 | 1 | PL2/3 | 304 | 0.433335302 | 0.000482096 | 21 | 0.001112524 | 48.46131825 | 8 mo | HOM |
| 1013978366 | 1 | PL2/3 | 304 | 0.433335302 | 0 | 0 | 0 | 0 | 2 mo | HOM |

| | | | | | | | | | | |
|---|---|---|---|---|---|---|---|---|---|---|
| 1014416155 | 2 | PL2/3 | 304 | 0.433335302 | 0 | 0 | 0 | 0 | 2 mo | HOM |
| 1013978065 | 2 | PL2/3 | 304 | 0.433335302 | 0 | 0 | 0 | 0 | 2 mo | HOM |
| 1009934102 | 2 | PL2/3 | 304 | 0.433335302 | 0 | 0 | 0 | 0 | 4 mo | WILD |
| 1009322937 | 2 | PL2/3 | 304 | 0.433335302 | 0 | 0 | 0 | 0 | 8 mo | WILD |
| 1011080637 | 2 | PL2/3 | 304 | 0.433335302 | 0.000265441 | 8 | 0.000612553 | 18.46145457 | 8 mo | HOM |
| 1013609713 | 2 | PL2/3 | 304 | 0.433335302 | 0 | 0 | 0 | 0 | 2 mo | WILD |
| 1010418842 | 1 | PL2/3 | 304 | 0.433335302 | 0.000392813 | 16 | 0.000906487 | 36.92290915 | 8 mo | HOM |
| 1013288213 | 2 | PL2/3 | 304 | 0.433335302 | 2.2E-05 | 2 | 5.08E-05 | 4.615363643 | 4 mo | HOM |
| 1008976060 | 2 | PL2/3 | 304 | 0.433335302 | 0 | 0 | 0 | 0 | 8 mo | WILD |
| 1013446243 | 1 | PL2/3 | 304 | 0.433335302 | 0 | 0 | 0 | 0 | 4 mo | HOM |
| 1013288951 | 2 | PL2/3 | 304 | 0.433335302 | 3.35E-05 | 1 | 7.73E-05 | 2.307681822 | 4 mo | HOM |
| 1013978665 | 1 | VISp6b | 305 | 0.169443176 | 0 | 0 | 0 | 0 | 2 mo | HOM |
| 1013445609 | 1 | VISp6b | 305 | 0.169443176 | 1.06E-05 | 2 | 6.28E-05 | 11.80336705 | 4 mo | HOM |
| 1013609015 | 1 | VISp6b | 305 | 0.169443176 | 2.99E-05 | 3 | 0.000176755 | 17.70505057 | 4 mo | HOM |
| 1013608106 | 2 | VISp6b | 305 | 0.169443176 | 0 | 0 | 0 | 0 | 4 mo | HOM |
| 1011937543 | 1 | VISp6b | 305 | 0.169443176 | 0.000277581 | 5 | 0.001638195 | 29.50841762 | 8 mo | HOM |
| 1010419538 | 2 | VISp6b | 305 | 0.169443176 | 0 | 0 | 0 | 0 | 4 mo | WILD |
| 1010419187 | 1 | VISp6b | 305 | 0.169443176 | 0.000250216 | 6 | 0.001476699 | 35.41010114 | 8 mo | HOM |
| 1013978366 | 1 | VISp6b | 305 | 0.169443176 | 0 | 0 | 0 | 0 | 2 mo | HOM |
| 1014416155 | 2 | VISp6b | 305 | 0.169443176 | 0 | 0 | 0 | 0 | 2 mo | HOM |
| 1013978065 | 2 | VISp6b | 305 | 0.169443176 | 0 | 0 | 0 | 0 | 2 mo | HOM |
| 1009934102 | 2 | VISp6b | 305 | 0.169443176 | 0 | 0 | 0 | 0 | 4 mo | WILD |
| 1009322937 | 2 | VISp6b | 305 | 0.169443176 | 0 | 0 | 0 | 0 | 8 mo | WILD |
| 1011080637 | 2 | VISp6b | 305 | 0.169443176 | 0.00022155 | 11 | 0.001307516 | 64.91851876 | 8 mo | HOM |
| 1013609713 | 2 | VISp6b | 305 | 0.169443176 | 0 | 0 | 0 | 0 | 2 mo | WILD |
| 1010418842 | 1 | VISp6b | 305 | 0.169443176 | 0.000308827 | 6 | 0.001822598 | 35.41010114 | 8 mo | HOM |
| 1013288213 | 2 | VISp6b | 305 | 0.169443176 | 0 | 0 | 0 | 0 | 4 mo | HOM |
| 1008976060 | 2 | VISp6b | 305 | 0.169443176 | 0 | 0 | 0 | 0 | 8 mo | WILD |
| 1013446243 | 1 | VISp6b | 305 | 0.169443176 | 3.6E-05 | 4 | 0.000212721 | 23.60673409 | 4 mo | HOM |
| 1013288951 | 2 | VISp6b | 305 | 0.169443176 | 6.18E-05 | 5 | 0.000364846 | 29.50841762 | 4 mo | HOM |
| 1013978665 | 1 | MARN | 307 | 0.549451974 | 0 | 0 | 0 | 0 | 2 mo | HOM |
| 1013445609 | 1 | MARN | 307 | 0.549451974 | 0 | 0 | 0 | 0 | 4 mo | HOM |
| 1013609015 | 1 | MARN | 307 | 0.549451974 | 0 | 0 | 0 | 0 | 4 mo | HOM |
| 1013608106 | 2 | MARN | 307 | 0.517549446 | 0 | 0 | 0 | 0 | 4 mo | HOM |
| 1011937543 | 1 | MARN | 307 | 0.549451974 | 0 | 0 | 0 | 0 | 8 mo | HOM |
| 1010419538 | 2 | MARN | 307 | 0.549451974 | 0 | 0 | 0 | 0 | 4 mo | WILD |
| 1010419187 | 1 | MARN | 307 | 0.549451974 | 0 | 0 | 0 | 0 | 8 mo | HOM |
| 1013978366 | 1 | MARN | 307 | 0.549451974 | 0 | 0 | 0 | 0 | 2 mo | HOM |
| 1014416155 | 2 | MARN | 307 | 0.549451974 | 0 | 0 | 0 | 0 | 2 mo | HOM |
| 1013978065 | 2 | MARN | 307 | 0.549451974 | 0 | 0 | 0 | 0 | 2 mo | HOM |
| 1009934102 | 2 | MARN | 307 | 0.549451974 | 0 | 0 | 0 | 0 | 4 mo | WILD |
| 1009322937 | 2 | MARN | 307 | 0.549451974 | 4.43E-06 | 1 | 8.06E-06 | 1.81999528 | 8 mo | WILD |
| 1011080637 | 2 | MARN | 307 | 0.549451974 | 0 | 0 | 0 | 0 | 8 mo | HOM |
| 1013609713 | 2 | MARN | 307 | 0.549451974 | 0 | 0 | 0 | 0 | 2 mo | WILD |
| 1010418842 | 1 | MARN | 307 | 0.549451974 | 3.9E-05 | 5 | 7.1E-05 | 9.0999764 | 8 mo | HOM |
| 1013288213 | 2 | MARN | 307 | 0.549451974 | 0 | 0 | 0 | 0 | 4 mo | HOM |
| 1008976060 | 2 | MARN | 307 | 0.549451974 | 8.15E-05 | 10 | 0.000148323 | 18.1999528 | 8 mo | WILD |
| 1013446243 | 1 | MARN | 307 | 0.549451974 | 0 | 0 | 0 | 0 | 4 mo | HOM |
| 1013288951 | 2 | MARN | 307 | 0.549451974 | 0 | 0 | 0 | 0 | 4 mo | HOM |
| 1013978665 | 1 | SF | 310 | 0.488650148 | 0 | 0 | 0 | 0 | 2 mo | HOM |
| 1013445609 | 1 | SF | 310 | 0.488650148 | 4.95E-06 | 1 | 1.01E-05 | 2.046453898 | 4 mo | HOM |
| 1013609015 | 1 | SF | 310 | 0.488650148 | 9.47E-05 | 1 | 0.000193824 | 2.046453898 | 4 mo | HOM |
| 1013608106 | 2 | SF | 310 | 0.488650148 | 0 | 0 | 0 | 0 | 4 mo | HOM |
| 1011937543 | 1 | SF | 310 | 0.488650148 | 0.000117251 | 7 | 0.00023995 | 14.32517728 | 8 mo | HOM |
| 1010419538 | 2 | SF | 310 | 0.488650148 | 0 | 0 | 0 | 0 | 4 mo | WILD |
| 1010419187 | 1 | SF | 310 | 0.488650148 | 0.000267652 | 4 | 0.000547737 | 8.185815591 | 8 mo | HOM |
| 1013978366 | 1 | SF | 310 | 0.488650148 | 8.21E-07 | 0 | 1.68E-06 | 0 | 2 mo | HOM |
| 1014416155 | 2 | SF | 310 | 0.488650148 | 0 | 0 | 0 | 0 | 2 mo | HOM |
| 1013978065 | 2 | SF | 310 | 0.488650148 | 0 | 0 | 0 | 0 | 2 mo | HOM |

| | | | | | | | | | | |
|---|---|---|---|---|---|---|---|---|---|---|
| **1009934102** | 2 | SF | 310 | 0.488650148 | 0 | 0 | 0 | 0 | 4 mo | WILD |
| **1009322937** | 2 | SF | 310 | 0.488650148 | 8.38E-07 | 0 | 1.71E-06 | 0 | 8 mo | WILD |
| **1011080637** | 2 | SF | 310 | 0.488650148 | 0.000325565 | 6 | 0.000666253 | 12.27872339 | 8 mo | HOM |
| **1013609713** | 2 | SF | 310 | 0.488650148 | 0 | 0 | 0 | 0 | 2 mo | WILD |
| **1010418842** | 1 | SF | 310 | 0.488650148 | 0.001340397 | 12 | 0.002743062 | 24.55744677 | 8 mo | HOM |
| **1013288213** | 2 | SF | 310 | 0.488650148 | 6.77E-06 | 0 | 1.39E-05 | 0 | 4 mo | HOM |
| **1008976060** | 2 | SF | 310 | 0.488650148 | 0.000101 | 1 | 0.000206691 | 2.046453898 | 8 mo | WILD |
| **1013446243** | 1 | SF | 310 | 0.488650148 | 9.92E-05 | 3 | 0.000202929 | 6.139361693 | 4 mo | HOM |
| **1013288951** | 2 | SF | 310 | 0.488650148 | 0.000158216 | 4 | 0.000323782 | 8.185815591 | 4 mo | HOM |
| **1013978665** | 1 | BLAp | 311 | 0.732733142 | 0 | 0 | 0 | 0 | 2 mo | HOM |
| **1013445609** | 1 | BLAp | 311 | 0.732733142 | 0.000120199 | 6 | 0.000164042 | 8.188520012 | 4 mo | HOM |
| **1013609015** | 1 | BLAp | 311 | 0.732733142 | 8.58E-05 | 4 | 0.000117147 | 5.459013341 | 4 mo | HOM |
| **1013608106** | 2 | BLAp | 311 | 0.732733142 | 0 | 0 | 0 | 0 | 4 mo | HOM |
| **1011937543** | 1 | BLAp | 311 | 0.732733142 | 0.001338066 | 28 | 0.001826129 | 38.21309339 | 8 mo | HOM |
| **1010419538** | 2 | BLAp | 311 | 0.732733142 | 0 | 0 | 0 | 0 | 4 mo | WILD |
| **1010419187** | 1 | BLAp | 311 | 0.732733142 | 0.000949405 | 44 | 0.001295703 | 60.04914675 | 8 mo | HOM |
| **1013978366** | 1 | BLAp | 311 | 0.732733142 | 0 | 0 | 0 | 0 | 2 mo | HOM |
| **1014416155** | 2 | BLAp | 311 | 0.732733142 | 0 | 0 | 0 | 0 | 2 mo | HOM |
| **1013978065** | 2 | BLAp | 311 | 0.732733142 | 0 | 0 | 0 | 0 | 2 mo | HOM |
| **1009934102** | 2 | BLAp | 311 | 0.732733142 | 0 | 0 | 0 | 0 | 4 mo | WILD |
| **1009322937** | 2 | BLAp | 311 | 0.732733142 | 0 | 0 | 0 | 0 | 8 mo | WILD |
| **1011080637** | 2 | BLAp | 311 | 0.732733142 | 0.000853012 | 37 | 0.001164151 | 50.49587341 | 8 mo | HOM |
| **1013609713** | 2 | BLAp | 311 | 0.732733142 | 0 | 0 | 0 | 0 | 2 mo | WILD |
| **1010418842** | 1 | BLAp | 311 | 0.732733142 | 0.001054373 | 40 | 0.001438958 | 54.59013341 | 8 mo | HOM |
| **1013288213** | 2 | BLAp | 311 | 0.732733142 | 5.99E-05 | 6 | 8.17E-05 | 8.188520012 | 4 mo | HOM |
| **1008976060** | 2 | BLAp | 311 | 0.732733142 | 0 | 0 | 0 | 0 | 8 mo | WILD |
| **1013446243** | 1 | BLAp | 311 | 0.732733142 | 0.000119633 | 4 | 0.000163269 | 5.459013341 | 4 mo | HOM |
| **1013288951** | 2 | BLAp | 311 | 0.732733142 | 9.64E-05 | 6 | 0.000131531 | 8.188520012 | 4 mo | HOM |
| **1013978665** | 1 | MB | 313 | 37.85097057 | 0.000199315 | 17 | 5.27E-06 | 0.449129831 | 2 mo | HOM |
| **1013445609** | 1 | MB | 313 | 37.85097057 | 0.000147886 | 11 | 3.91E-06 | 0.29061342 | 4 mo | HOM |
| **1013609015** | 1 | MB | 313 | 37.85097057 | 0.007424324 | 23 | 0.000196146 | 0.607646241 | 4 mo | HOM |
| **1013608106** | 2 | MB | 313 | 37.85096305 | 0 | 0 | 0 | 0 | 4 mo | HOM |
| **1011937543** | 1 | MB | 313 | 37.85097057 | 0.090947695 | 720 | 0.002402784 | 19.02196929 | 8 mo | HOM |
| **1010419538** | 2 | MB | 313 | 37.85096305 | 5.19E-05 | 6 | 1.37E-06 | 0.158516442 | 4 mo | WILD |
| **1010419187** | 1 | MB | 313 | 37.85097057 | 0.011763594 | 186 | 0.000310787 | 4.914008734 | 8 mo | HOM |
| **1013978366** | 1 | MB | 313 | 37.85097057 | 0.000983884 | 23 | 2.6E-05 | 0.607646241 | 2 mo | HOM |
| **1014416155** | 2 | MB | 313 | 37.85096305 | 0 | 0 | 0 | 0 | 2 mo | HOM |
| **1013978065** | 2 | MB | 313 | 37.85096305 | 1.97E-05 | 1 | 5.21E-07 | 0.026419407 | 2 mo | HOM |
| **1009934102** | 2 | MB | 313 | 37.85096305 | 8.97E-05 | 4 | 2.37E-06 | 0.105677628 | 4 mo | WILD |
| **1009322937** | 2 | MB | 313 | 37.85096305 | 0.000935656 | 31 | 2.47E-05 | 0.819001619 | 8 mo | WILD |
| **1011080637** | 2 | MB | 313 | 37.85096305 | 0.014910973 | 143 | 0.000393939 | 3.777975208 | 8 mo | HOM |
| **1013609713** | 2 | MB | 313 | 37.85096305 | 2.06E-05 | 2 | 5.43E-07 | 0.052838814 | 2 mo | WILD |
| **1010418842** | 1 | MB | 313 | 37.85097057 | 0.013366184 | 280 | 0.000353127 | 7.397432503 | 8 mo | HOM |
| **1013288213** | 2 | MB | 313 | 37.85096305 | 0.000146842 | 7 | 3.88E-06 | 0.184935849 | 4 mo | HOM |
| **1008976060** | 2 | MB | 313 | 37.85096305 | 0.005300527 | 179 | 0.000140037 | 4.729073862 | 8 mo | WILD |
| **1013446243** | 1 | MB | 313 | 37.85097057 | 0.001791168 | 22 | 4.73E-05 | 0.58122684 | 4 mo | HOM |
| **1013288951** | 2 | MB | 313 | 37.85096305 | 0.002861105 | 47 | 7.56E-05 | 1.241712131 | 4 mo | HOM |
| **1013978665** | 1 | Alp6a | 314 | 0.448908169 | 0 | 0 | 0 | 0 | 2 mo | HOM |
| **1013445609** | 1 | Alp6a | 314 | 0.448908169 | 4.97E-05 | 6 | 0.000110814 | 13.36576257 | 4 mo | HOM |
| **1013609015** | 1 | Alp6a | 314 | 0.448908169 | 0.000104729 | 6 | 0.000233298 | 13.36576257 | 4 mo | HOM |
| **1013608106** | 2 | Alp6a | 314 | 0.448908169 | 0 | 0 | 0 | 0 | 4 mo | HOM |
| **1011937543** | 1 | Alp6a | 314 | 0.448908169 | 0.001023518 | 23 | 0.002280016 | 51.23542319 | 8 mo | HOM |
| **1010419538** | 2 | Alp6a | 314 | 0.448908169 | 0 | 0 | 0 | 0 | 4 mo | WILD |
| **1010419187** | 1 | Alp6a | 314 | 0.448908169 | 0.000845121 | 28 | 0.001882615 | 62.37355866 | 8 mo | HOM |
| **1013978366** | 1 | Alp6a | 314 | 0.448908169 | 0 | 0 | 0 | 0 | 2 mo | HOM |
| **1014416155** | 2 | Alp6a | 314 | 0.448908169 | 0 | 0 | 0 | 0 | 2 mo | HOM |
| **1013978065** | 2 | Alp6a | 314 | 0.448908169 | 0 | 0 | 0 | 0 | 2 mo | HOM |
| **1009934102** | 2 | Alp6a | 314 | 0.448908169 | 0 | 0 | 0 | 0 | 4 mo | WILD |
| **1009322937** | 2 | Alp6a | 314 | 0.448908169 | 0 | 0 | 0 | 0 | 8 mo | WILD |

| | | | | | | | | | | |
|---|---|---|---|---|---|---|---|---|---|---|
| **1011080637** | 2 | Alp6a | 314 | 0.448908169 | 0.000436877 | 28 | 0.0009732 | 62.37355866 | 8 mo | HOM |
| **1013609713** | 2 | Alp6a | 314 | 0.448908169 | 0 | 0 | 0 | 0 | 2 mo | WILD |
| **1010418842** | 1 | Alp6a | 314 | 0.448908169 | 0.000998643 | 35 | 0.002224605 | 77.96694833 | 8 mo | HOM |
| **1013288213** | 2 | Alp6a | 314 | 0.448908169 | 1.52E-05 | 1 | 3.38E-05 | 2.227627095 | 4 mo | HOM |
| **1008976060** | 2 | Alp6a | 314 | 0.448908169 | 7.36E-06 | 2 | 1.64E-05 | 4.45525419 | 8 mo | WILD |
| **1013446243** | 1 | Alp6a | 314 | 0.448908169 | 2.19E-05 | 3 | 4.87E-05 | 6.682881285 | 4 mo | HOM |
| **1013288951** | 2 | Alp6a | 314 | 0.448908169 | 0.000106142 | 11 | 0.000236446 | 24.50389805 | 4 mo | HOM |
| **1013978665** | 1 | Isocortex | 315 | 127.0088414 | 0.001399702 | 71 | 1.1E-05 | 0.559016201 | 2 mo | HOM |
| **1013445609** | 1 | Isocortex | 315 | 127.0088414 | 0.011104307 | 695 | 8.74E-05 | 5.472059995 | 4 mo | HOM |
| **1013609015** | 1 | Isocortex | 315 | 127.0088414 | 0.036024452 | 1252 | 0.000283637 | 9.857581458 | 4 mo | HOM |
| **1013608106** | 2 | Isocortex | 315 | 127.0088414 | 2.26E-07 | 0 | 1.78E-09 | 0 | 4 mo | HOM |
| **1011937543** | 1 | Isocortex | 315 | 127.0088414 | 0.205732723 | 3404 | 0.00161983 | 26.80128377 | 8 mo | HOM |
| **1010419538** | 2 | Isocortex | 315 | 126.9138242 | 0.001131577 | 43 | 8.92E-06 | 0.338812578 | 4 mo | WILD |
| **1010419187** | 1 | Isocortex | 315 | 127.0088414 | 0.225159712 | 6342 | 0.001772788 | 49.93353164 | 8 mo | HOM |
| **1013978366** | 1 | Isocortex | 315 | 127.0088414 | 0.004141283 | 86 | 3.26E-05 | 0.677118215 | 2 mo | HOM |
| **1014416155** | 2 | Isocortex | 315 | 127.0088414 | 2.11E-05 | 2 | 1.66E-07 | 0.015746935 | 2 mo | HOM |
| **1013978065** | 2 | Isocortex | 315 | 127.0088414 | 9.31E-05 | 5 | 7.33E-07 | 0.039367338 | 2 mo | HOM |
| **1009934102** | 2 | Isocortex | 315 | 127.0088414 | 0.001140741 | 51 | 8.98E-06 | 0.401546849 | 4 mo | WILD |
| **1009322937** | 2 | Isocortex | 315 | 127.0088414 | 0.00104425 | 74 | 8.22E-06 | 0.582636604 | 8 mo | WILD |
| **1011080637** | 2 | Isocortex | 315 | 127.0088414 | 0.108633345 | 4222 | 0.000855321 | 33.24178029 | 8 mo | HOM |
| **1013609713** | 2 | Isocortex | 315 | 127.0088414 | 0.000293095 | 20 | 2.31E-06 | 0.157469352 | 2 mo | WILD |
| **1010418842** | 1 | Isocortex | 315 | 127.0088414 | 0.164157571 | 5352 | 0.001292489 | 42.13879869 | 8 mo | HOM |
| **1013288213** | 2 | Isocortex | 315 | 127.0088414 | 0.016688135 | 650 | 0.000131393 | 5.117753952 | 4 mo | HOM |
| **1008976060** | 2 | Isocortex | 315 | 127.0088414 | 0.007301651 | 265 | 5.75E-05 | 2.086468919 | 8 mo | WILD |
| **1013446243** | 1 | Isocortex | 315 | 127.0088414 | 0.025543148 | 1219 | 0.000201113 | 9.597757027 | 4 mo | HOM |
| **1013288951** | 2 | Isocortex | 315 | 127.0088414 | 0.043591737 | 1398 | 0.000343218 | 11.00710773 | 4 mo | HOM |
| **1013978665** | 1 | SG | 318 | 0.017884706 | 0 | 0 | 0 | 0 | 2 mo | HOM |
| **1013445609** | 1 | SG | 318 | 0.017884706 | 0 | 0 | 0 | 0 | 4 mo | HOM |
| **1013609015** | 1 | SG | 318 | 0.017884706 | 0 | 0 | 0 | 0 | 4 mo | HOM |
| **1013608106** | 2 | SG | 318 | 0.017884706 | 0 | 0 | 0 | 0 | 4 mo | HOM |
| **1011937543** | 1 | SG | 318 | 0.017884706 | 0 | 0 | 0 | 0 | 8 mo | HOM |
| **1010419538** | 2 | SG | 318 | 0.017884706 | 0 | 0 | 0 | 0 | 4 mo | WILD |
| **1010419187** | 1 | SG | 318 | 0.017884706 | 0 | 0 | 0 | 0 | 8 mo | HOM |
| **1013978366** | 1 | SG | 318 | 0.017884706 | 0 | 0 | 0 | 0 | 2 mo | HOM |
| **1014416155** | 2 | SG | 318 | 0.017884706 | 0 | 0 | 0 | 0 | 2 mo | HOM |
| **1013978065** | 2 | SG | 318 | 0.017884706 | 0 | 0 | 0 | 0 | 2 mo | HOM |
| **1009934102** | 2 | SG | 318 | 0.017884706 | 0 | 0 | 0 | 0 | 4 mo | WILD |
| **1009322937** | 2 | SG | 318 | 0.017884706 | 0 | 0 | 0 | 0 | 8 mo | WILD |
| **1011080637** | 2 | SG | 318 | 0.017884706 | 0 | 0 | 0 | 0 | 8 mo | HOM |
| **1013609713** | 2 | SG | 318 | 0.017884706 | 0 | 0 | 0 | 0 | 2 mo | WILD |
| **1010418842** | 1 | SG | 318 | 0.017884706 | 0 | 0 | 0 | 0 | 8 mo | HOM |
| **1013288213** | 2 | SG | 318 | 0.017884706 | 0 | 0 | 0 | 0 | 4 mo | HOM |
| **1008976060** | 2 | SG | 318 | 0.017884706 | 0 | 0 | 0 | 0 | 8 mo | WILD |
| **1013446243** | 1 | SG | 318 | 0.017884706 | 0 | 0 | 0 | 0 | 4 mo | HOM |
| **1013288951** | 2 | SG | 318 | 0.017884706 | 0 | 0 | 0 | 0 | 4 mo | HOM |
| **1013978665** | 1 | BMA | 319 | 1.528891022 | 0 | 0 | 0 | 0 | 2 mo | HOM |
| **1013445609** | 1 | BMA | 319 | 1.530077842 | 0.000124207 | 7 | 8.12E-05 | 4.574930641 | 4 mo | HOM |
| **1013609015** | 1 | BMA | 319 | 1.530077842 | 0.000381558 | 12 | 0.000249371 | 7.842738241 | 4 mo | HOM |
| **1013608106** | 2 | BMA | 319 | 1.530077842 | 0 | 0 | 0 | 0 | 4 mo | HOM |
| **1011937543** | 1 | BMA | 319 | 1.530077842 | 0.004061586 | 42 | 0.002654496 | 27.44958384 | 8 mo | HOM |
| **1010419538** | 2 | BMA | 319 | 1.530077842 | 0 | 0 | 0 | 0 | 4 mo | WILD |
| **1010419187** | 1 | BMA | 319 | 1.530077842 | 0.002478486 | 85 | 0.001619843 | 55.55272921 | 8 mo | HOM |
| **1013978366** | 1 | BMA | 319 | 1.530077842 | 0 | 0 | 0 | 0 | 2 mo | HOM |
| **1014416155** | 2 | BMA | 319 | 1.530077842 | 0 | 0 | 0 | 0 | 2 mo | HOM |
| **1013978065** | 2 | BMA | 319 | 1.530077842 | 0 | 0 | 0 | 0 | 2 mo | HOM |
| **1009934102** | 2 | BMA | 319 | 1.530077842 | 0 | 0 | 0 | 0 | 4 mo | WILD |
| **1009322937** | 2 | BMA | 319 | 1.530077842 | 0 | 0 | 0 | 0 | 8 mo | WILD |
| **1011080637** | 2 | BMA | 319 | 1.530077842 | 0.001677218 | 74 | 0.001096165 | 48.36355249 | 8 mo | HOM |
| **1013609713** | 2 | BMA | 319 | 1.530077842 | 0 | 0 | 0 | 0 | 2 mo | WILD |

| | | | | | | | | | | |
|---|---|---|---|---|---|---|---|---|---|---|
| **1010418842** | 1 | BMA | 319 | 1.530077842 | 0.002539821 | 67 | 0.001659929 | 43.78862185 | 8 mo | HOM |
| **1013288213** | 2 | BMA | 319 | 1.530077842 | 7.76E-05 | 5 | 5.07E-05 | 3.2678076 | 4 mo | HOM |
| **1008976060** | 2 | BMA | 319 | 1.530077842 | 8.14E-06 | 1 | 5.32E-06 | 0.65356152 | 8 mo | WILD |
| **1013446243** | 1 | BMA | 319 | 1.530077842 | 0.000240849 | 13 | 0.00015741 | 8.496299761 | 4 mo | HOM |
| **1013288951** | 2 | BMA | 319 | 1.530077842 | 0.000267815 | 8 | 0.000175034 | 5.228492161 | 4 mo | HOM |
| **1013978665** | 1 | MOp1 | 320 | 1.369150474 | 3.79E-05 | 6 | 2.77E-05 | 4.382279461 | 2 mo | HOM |
| **1013445609** | 1 | MOp1 | 320 | 1.369150474 | 0.00015257 | 11 | 0.000111434 | 8.034179013 | 4 mo | HOM |
| **1013609015** | 1 | MOp1 | 320 | 1.369150474 | 0.000356204 | 20 | 0.000260164 | 14.6075982 | 4 mo | HOM |
| **1013608106** | 2 | MOp1 | 320 | 1.369150474 | 0 | 0 | 0 | 0 | 4 mo | HOM |
| **1011937543** | 1 | MOp1 | 320 | 1.369150474 | 0.003274125 | 65 | 0.002391355 | 47.47469416 | 8 mo | HOM |
| **1010419538** | 2 | MOp1 | 320 | 1.369150474 | 9.44E-06 | 1 | 6.9E-06 | 0.73037991 | 4 mo | WILD |
| **1010419187** | 1 | MOp1 | 320 | 1.369150474 | 0.003423051 | 112 | 0.002500128 | 81.80254995 | 8 mo | HOM |
| **1013978366** | 1 | MOp1 | 320 | 1.369150474 | 0 | 0 | 0 | 0 | 2 mo | HOM |
| **1014416155** | 2 | MOp1 | 320 | 1.369150474 | 0 | 0 | 0 | 0 | 2 mo | HOM |
| **1013978065** | 2 | MOp1 | 320 | 1.369150474 | 0 | 0 | 0 | 0 | 2 mo | HOM |
| **1009934102** | 2 | MOp1 | 320 | 1.369150474 | 2.69E-06 | 1 | 1.97E-06 | 0.73037991 | 4 mo | WILD |
| **1009322937** | 2 | MOp1 | 320 | 1.369150474 | 2.99E-06 | 1 | 2.19E-06 | 0.73037991 | 8 mo | WILD |
| **1011080637** | 2 | MOp1 | 320 | 1.369150474 | 0.000987433 | 38 | 0.000721201 | 27.75443659 | 8 mo | HOM |
| **1013609713** | 2 | MOp1 | 320 | 1.369150474 | 1.87E-05 | 1 | 1.37E-05 | 0.73037991 | 2 mo | WILD |
| **1010418842** | 1 | MOp1 | 320 | 1.369150474 | 0.002854746 | 96 | 0.002085049 | 70.11647138 | 8 mo | HOM |
| **1013288213** | 2 | MOp1 | 320 | 1.369150474 | 9.84E-05 | 8 | 7.19E-05 | 5.843039282 | 4 mo | HOM |
| **1008976060** | 2 | MOp1 | 320 | 1.369150474 | 0.000245198 | 13 | 0.000179088 | 9.494938833 | 8 mo | WILD |
| **1013446243** | 1 | MOp1 | 320 | 1.369150474 | 0.000488963 | 25 | 0.000357129 | 18.25949776 | 4 mo | HOM |
| **1013288951** | 2 | MOp1 | 320 | 1.369150474 | 0.001008334 | 41 | 0.000736467 | 29.94557632 | 4 mo | HOM |
| **1013978665** | 1 | SubG | 321 | 0.027671843 | 0 | 0 | 0 | 0 | 2 mo | HOM |
| **1013445609** | 1 | SubG | 321 | 0.027671843 | 0 | 0 | 0 | 0 | 4 mo | HOM |
| **1013609015** | 1 | SubG | 321 | 0.027671843 | 0 | 0 | 0 | 0 | 4 mo | HOM |
| **1013608106** | 2 | SubG | 321 | 0.027671843 | 0 | 0 | 0 | 0 | 4 mo | HOM |
| **1011937543** | 1 | SubG | 321 | 0.027671843 | 0 | 0 | 0 | 0 | 8 mo | HOM |
| **1010419538** | 2 | SubG | 321 | 0.027671843 | 0 | 0 | 0 | 0 | 4 mo | WILD |
| **1010419187** | 1 | SubG | 321 | 0.027671843 | 4.94E-06 | 1 | 0.000178357 | 36.13781641 | 8 mo | HOM |
| **1013978366** | 1 | SubG | 321 | 0.027671843 | 0 | 0 | 0 | 0 | 2 mo | HOM |
| **1014416155** | 2 | SubG | 321 | 0.027671843 | 0 | 0 | 0 | 0 | 2 mo | HOM |
| **1013978065** | 2 | SubG | 321 | 0.027671843 | 0 | 0 | 0 | 0 | 2 mo | HOM |
| **1009934102** | 2 | SubG | 321 | 0.027671843 | 0 | 0 | 0 | 0 | 4 mo | WILD |
| **1009322937** | 2 | SubG | 321 | 0.027671843 | 0 | 0 | 0 | 0 | 8 mo | WILD |
| **1011080637** | 2 | SubG | 321 | 0.027671843 | 0 | 0 | 0 | 0 | 8 mo | HOM |
| **1013609713** | 2 | SubG | 321 | 0.027671843 | 0 | 0 | 0 | 0 | 2 mo | WILD |
| **1010418842** | 1 | SubG | 321 | 0.027671843 | 9.66E-06 | 0 | 0.000349006 | 0 | 8 mo | HOM |
| **1013288213** | 2 | SubG | 321 | 0.027671843 | 0 | 0 | 0 | 0 | 4 mo | HOM |
| **1008976060** | 2 | SubG | 321 | 0.027671843 | 0 | 0 | 0 | 0 | 8 mo | WILD |
| **1013446243** | 1 | SubG | 321 | 0.027671843 | 0 | 0 | 0 | 0 | 4 mo | HOM |
| **1013288951** | 2 | SubG | 321 | 0.027671843 | 0 | 0 | 0 | 0 | 4 mo | HOM |
| **1013978665** | 1 | SSp | 322 | 25.03662918 | 5.05E-05 | 3 | 2.02E-06 | 0.119824437 | 2 mo | HOM |
| **1013445609** | 1 | SSp | 322 | 25.03662918 | 0.002995361 | 162 | 0.000119639 | 6.470519606 | 4 mo | HOM |
| **1013609015** | 1 | SSp | 322 | 25.03662918 | 0.008062002 | 281 | 0.000322008 | 11.22355561 | 4 mo | HOM |
| **1013608106** | 2 | SSp | 322 | 25.03662918 | 0 | 0 | 0 | 0 | 4 mo | HOM |
| **1011937543** | 1 | SSp | 322 | 25.03662918 | 0.048045726 | 773 | 0.001919017 | 30.8747633 | 8 mo | HOM |
| **1010419538** | 2 | SSp | 322 | 25.03662918 | 1.72E-05 | 3 | 6.89E-07 | 0.119824437 | 4 mo | WILD |
| **1010419187** | 1 | SSp | 322 | 25.03662918 | 0.045193105 | 1387 | 0.001805079 | 55.39883144 | 8 mo | HOM |
| **1013978366** | 1 | SSp | 322 | 25.03662918 | 9.03E-05 | 8 | 3.61E-06 | 0.319531832 | 2 mo | HOM |
| **1014416155** | 2 | SSp | 322 | 25.03662918 | 4.49E-06 | 1 | 1.79E-07 | 0.039941479 | 2 mo | HOM |
| **1013978065** | 2 | SSp | 322 | 25.03662918 | 4.42E-06 | 1 | 1.77E-07 | 0.039941479 | 2 mo | HOM |
| **1009934102** | 2 | SSp | 322 | 25.03662918 | 0.000106441 | 8 | 4.25E-06 | 0.319531832 | 4 mo | WILD |
| **1009322937** | 2 | SSp | 322 | 25.03662918 | 3.19E-05 | 5 | 1.27E-06 | 0.199707395 | 8 mo | WILD |
| **1011080637** | 2 | SSp | 322 | 25.03662918 | 0.025503709 | 915 | 0.001018656 | 36.54645333 | 8 mo | HOM |
| **1013609713** | 2 | SSp | 322 | 25.03662918 | 5.02E-05 | 5 | 2.01E-06 | 0.199707395 | 2 mo | WILD |
| **1010418842** | 1 | SSp | 322 | 25.03662918 | 0.03874333 | 1199 | 0.001547466 | 47.88983338 | 8 mo | HOM |
| **1013288213** | 2 | SSp | 322 | 25.03662918 | 0.00207001 | 161 | 8.27E-05 | 6.430578127 | 4 mo | HOM |

| | | | | | | | | | | |
|---|---|---|---|---|---|---|---|---|---|---|
| **1008976060** | 2 | SSp | 322 | 25.03662918 | 0.001494977 | 63 | 5.97E-05 | 2.51631318 | 8 mo | WILD |
| **1013446243** | 1 | SSp | 322 | 25.03662918 | 0.006666561 | 325 | 0.000266272 | 12.98098069 | 4 mo | HOM |
| **1013288951** | 2 | SSp | 322 | 25.03662918 | 0.006475598 | 289 | 0.000258645 | 11.54308745 | 4 mo | HOM |
| **1013978665** | 1 | MBmot | 323 | 21.94169514 | 5.73E-05 | 7 | 2.61E-06 | 0.319027311 | 2 mo | HOM |
| **1013445609** | 1 | MBmot | 323 | 21.94169514 | 0.000138956 | 10 | 6.33E-06 | 0.455753301 | 4 mo | HOM |
| **1013609015** | 1 | MBmot | 323 | 21.94169514 | 0.006278416 | 13 | 0.000286141 | 0.592479292 | 4 mo | HOM |
| **1013608106** | 2 | MBmot | 323 | 21.94169514 | 0 | 0 | 0 | 0 | 4 mo | HOM |
| **1011937543** | 1 | MBmot | 323 | 21.94169514 | 0.057432382 | 437 | 0.0026175 | 19.91641927 | 8 mo | HOM |
| **1010419538** | 2 | MBmot | 323 | 21.94169514 | 3.8E-05 | 4 | 1.73E-06 | 0.182301321 | 4 mo | WILD |
| **1010419187** | 1 | MBmot | 323 | 21.94169514 | 0.004866763 | 82 | 0.000221804 | 3.737177072 | 8 mo | HOM |
| **1013978366** | 1 | MBmot | 323 | 21.94169514 | 0.000301576 | 15 | 1.37E-05 | 0.683629952 | 2 mo | HOM |
| **1014416155** | 2 | MBmot | 323 | 21.94169514 | 0 | 0 | 0 | 0 | 2 mo | HOM |
| **1013978065** | 2 | MBmot | 323 | 21.94169514 | 1.28E-05 | 1 | 5.83E-07 | 0.04557533 | 2 mo | HOM |
| **1009934102** | 2 | MBmot | 323 | 21.94169514 | 4.74E-05 | 1 | 2.16E-06 | 0.04557533 | 4 mo | WILD |
| **1009322937** | 2 | MBmot | 323 | 21.94169514 | 0.000694458 | 23 | 3.17E-05 | 1.048232593 | 8 mo | WILD |
| **1011080637** | 2 | MBmot | 323 | 21.94169514 | 0.011982535 | 64 | 0.000546108 | 2.916821129 | 8 mo | HOM |
| **1013609713** | 2 | MBmot | 323 | 21.94169514 | 4.65E-06 | 0 | 2.12E-07 | 0 | 2 mo | WILD |
| **1010418842** | 1 | MBmot | 323 | 21.94169514 | 0.008607949 | 144 | 0.00039231 | 6.56284754 | 8 mo | HOM |
| **1013288213** | 2 | MBmot | 323 | 21.94169514 | 0.0001207 | 5 | 5.5E-06 | 0.227876651 | 4 mo | HOM |
| **1008976060** | 2 | MBmot | 323 | 21.94169514 | 0.003647131 | 109 | 0.000166219 | 4.967710985 | 8 mo | WILD |
| **1013446243** | 1 | MBmot | 323 | 21.94169514 | 0.001593021 | 10 | 7.26E-05 | 0.455753301 | 4 mo | HOM |
| **1013288951** | 2 | MBmot | 323 | 21.94169514 | 0.002223044 | 20 | 0.000101316 | 0.911506603 | 4 mo | HOM |
| **1013978665** | 1 | SGN | 325 | 0.186359466 | 0 | 0 | 0 | 0 | 2 mo | HOM |
| **1013445609** | 1 | SGN | 325 | 0.186359466 | 0 | 0 | 0 | 0 | 4 mo | HOM |
| **1013609015** | 1 | SGN | 325 | 0.186359466 | 0 | 0 | 0 | 0 | 4 mo | HOM |
| **1013608106** | 2 | SGN | 325 | 0.186359466 | 0 | 0 | 0 | 0 | 4 mo | HOM |
| **1011937543** | 1 | SGN | 325 | 0.186359466 | 0.000664173 | 8 | 0.003563937 | 42.92778998 | 8 mo | HOM |
| **1010419538** | 2 | SGN | 325 | 0.186359466 | 0 | 0 | 0 | 0 | 4 mo | WILD |
| **1010419187** | 1 | SGN | 325 | 0.186359466 | 6.56E-05 | 3 | 0.000351915 | 16.09792124 | 8 mo | HOM |
| **1013978366** | 1 | SGN | 325 | 0.186359466 | 0 | 0 | 0 | 0 | 2 mo | HOM |
| **1014416155** | 2 | SGN | 325 | 0.186359466 | 0 | 0 | 0 | 0 | 2 mo | HOM |
| **1013978065** | 2 | SGN | 325 | 0.186359466 | 0 | 0 | 0 | 0 | 2 mo | HOM |
| **1009934102** | 2 | SGN | 325 | 0.186359466 | 0 | 0 | 0 | 0 | 4 mo | WILD |
| **1009322937** | 2 | SGN | 325 | 0.186359466 | 0 | 0 | 0 | 0 | 8 mo | WILD |
| **1011080637** | 2 | SGN | 325 | 0.186359466 | 1.32E-06 | 0 | 7.07E-06 | 0 | 8 mo | HOM |
| **1013609713** | 2 | SGN | 325 | 0.186359466 | 0 | 0 | 0 | 0 | 2 mo | WILD |
| **1010418842** | 1 | SGN | 325 | 0.186359466 | 7.11E-06 | 1 | 3.82E-05 | 5.365973747 | 8 mo | HOM |
| **1013288213** | 2 | SGN | 325 | 0.186359466 | 0 | 0 | 0 | 0 | 4 mo | HOM |
| **1008976060** | 2 | SGN | 325 | 0.186359466 | 2.2E-05 | 1 | 0.000117951 | 5.365973747 | 8 mo | WILD |
| **1013446243** | 1 | SGN | 325 | 0.186359466 | 0 | 0 | 0 | 0 | 4 mo | HOM |
| **1013288951** | 2 | SGN | 325 | 0.186359466 | 0 | 0 | 0 | 0 | 4 mo | HOM |
| **1013978665** | 1 | scp | 326 | 0.882255898 | 0 | 0 | 0 | 0 | 2 mo | HOM |
| **1013445609** | 1 | scp | 326 | 0.882255898 | 2.64E-06 | 0 | 2.99E-06 | 0 | 4 mo | HOM |
| **1013609015** | 1 | scp | 326 | 0.882255898 | 3.08E-06 | 0 | 3.49E-06 | 0 | 4 mo | HOM |
| **1013608106** | 2 | scp | 326 | 0.882255898 | 0 | 0 | 0 | 0 | 4 mo | HOM |
| **1011937543** | 1 | scp | 326 | 0.882255898 | 0.0025147 | 5 | 0.002850306 | 5.667289971 | 8 mo | HOM |
| **1010419538** | 2 | scp | 326 | 0.882255898 | 0 | 0 | 0 | 0 | 4 mo | WILD |
| **1010419187** | 1 | scp | 326 | 0.882255898 | 5.86E-06 | 0 | 6.64E-06 | 0 | 8 mo | HOM |
| **1013978366** | 1 | scp | 326 | 0.882255898 | 0 | 0 | 0 | 0 | 2 mo | HOM |
| **1014416155** | 2 | scp | 326 | 0.882255898 | 0 | 0 | 0 | 0 | 2 mo | HOM |
| **1013978065** | 2 | scp | 326 | 0.882255898 | 0 | 0 | 0 | 0 | 2 mo | HOM |
| **1009934102** | 2 | scp | 326 | 0.882255898 | 0 | 0 | 0 | 0 | 4 mo | WILD |
| **1009322937** | 2 | scp | 326 | 0.882255898 | 8.62E-06 | 0 | 9.77E-06 | 0 | 8 mo | WILD |
| **1011080637** | 2 | scp | 326 | 0.882255898 | 8.46E-06 | 0 | 9.59E-06 | 0 | 8 mo | HOM |
| **1013609713** | 2 | scp | 326 | 0.882255898 | 3.65E-10 | 0 | 4.14E-10 | 0 | 2 mo | WILD |
| **1010418842** | 1 | scp | 326 | 0.882255898 | 0.000244493 | 3 | 0.000277122 | 3.400373982 | 8 mo | HOM |
| **1013288213** | 2 | scp | 326 | 0.882255898 | 0 | 0 | 0 | 0 | 4 mo | HOM |
| **1008976060** | 2 | scp | 326 | 0.882255898 | 3.6E-05 | 2 | 4.08E-05 | 2.266915988 | 8 mo | WILD |
| **1013446243** | 1 | scp | 326 | 0.882255898 | 7.42E-06 | 0 | 8.41E-06 | 0 | 4 mo | HOM |

| | | | | | | | | | | |
|---|---|---|---|---|---|---|---|---|---|---|
| 1013288951 | 2 | scp | 326 | 0.882255898 | 1.15E-06 | 0 | 1.3E-06 | 0 | 4 mo | HOM |
| 1013978665 | 1 | BMAa | 327 | 0.789430533 | 0 | 0 | 0 | 0 | 2 mo | HOM |
| 1013445609 | 1 | BMAa | 327 | 0.790617352 | 3.12E-05 | 1 | 3.94E-05 | 1.26483437 | 4 mo | HOM |
| 1013609015 | 1 | BMAa | 327 | 0.790617352 | 0.000137542 | 4 | 0.000173967 | 5.059337478 | 4 mo | HOM |
| 1013608106 | 2 | BMAa | 327 | 0.790617352 | 0 | 0 | 0 | 0 | 4 mo | HOM |
| 1011937543 | 1 | BMAa | 327 | 0.790617352 | 0.001826424 | 16 | 0.002310124 | 20.23734991 | 8 mo | HOM |
| 1010419538 | 2 | BMAa | 327 | 0.790617352 | 0 | 0 | 0 | 0 | 4 mo | WILD |
| 1010419187 | 1 | BMAa | 327 | 0.790617352 | 0.001137855 | 33 | 0.001439199 | 41.7395342 | 8 mo | HOM |
| 1013978366 | 1 | BMAa | 327 | 0.790617352 | 0 | 0 | 0 | 0 | 2 mo | HOM |
| 1014416155 | 2 | BMAa | 327 | 0.790617352 | 0 | 0 | 0 | 0 | 2 mo | HOM |
| 1013978065 | 2 | BMAa | 327 | 0.790617352 | 0 | 0 | 0 | 0 | 2 mo | HOM |
| 1009934102 | 2 | BMAa | 327 | 0.790617352 | 0 | 0 | 0 | 0 | 4 mo | WILD |
| 1009322937 | 2 | BMAa | 327 | 0.790617352 | 0 | 0 | 0 | 0 | 8 mo | WILD |
| 1011080637 | 2 | BMAa | 327 | 0.790617352 | 0.0007225 | 30 | 0.000913843 | 37.94503109 | 8 mo | HOM |
| 1013609713 | 2 | BMAa | 327 | 0.790617352 | 0 | 0 | 0 | 0 | 2 mo | WILD |
| 1010418842 | 1 | BMAa | 327 | 0.790617352 | 0.001117041 | 23 | 0.001412872 | 29.0911905 | 8 mo | HOM |
| 1013288213 | 2 | BMAa | 327 | 0.790617352 | 1.44E-05 | 1 | 1.82E-05 | 1.26483437 | 4 mo | HOM |
| 1008976060 | 2 | BMAa | 327 | 0.790617352 | 1.64E-06 | 0 | 2.08E-06 | 0 | 8 mo | WILD |
| 1013446243 | 1 | BMAa | 327 | 0.790617352 | 0.000109941 | 6 | 0.000139058 | 7.589006218 | 4 mo | HOM |
| 1013288951 | 2 | BMAa | 327 | 0.790617352 | 6.01E-05 | 3 | 7.6E-05 | 3.794503109 | 4 mo | HOM |
| 1013978665 | 1 | Ald2/3 | 328 | 0.95348559 | 0 | 0 | 0 | 0 | 2 mo | HOM |
| 1013445609 | 1 | Ald2/3 | 328 | 0.95348559 | 5.01E-05 | 3 | 5.25E-05 | 3.146350643 | 4 mo | HOM |
| 1013609015 | 1 | Ald2/3 | 328 | 0.95348559 | 0.000105235 | 4 | 0.000110369 | 4.19513419 | 4 mo | HOM |
| 1013608106 | 2 | Ald2/3 | 328 | 0.95348559 | 0 | 0 | 0 | 0 | 4 mo | HOM |
| 1011937543 | 1 | Ald2/3 | 328 | 0.95348559 | 0.000930078 | 27 | 0.00097545 | 28.31715578 | 8 mo | HOM |
| 1010419538 | 2 | Ald2/3 | 328 | 0.95348559 | 0 | 0 | 0 | 0 | 4 mo | WILD |
| 1010419187 | 1 | Ald2/3 | 328 | 0.95348559 | 0.0010448 | 33 | 0.001095769 | 34.60985707 | 8 mo | HOM |
| 1013978366 | 1 | Ald2/3 | 328 | 0.95348559 | 1E-05 | 1 | 1.05E-05 | 1.048783548 | 2 mo | HOM |
| 1014416155 | 2 | Ald2/3 | 328 | 0.95348559 | 0 | 0 | 0 | 0 | 2 mo | HOM |
| 1013978065 | 2 | Ald2/3 | 328 | 0.95348559 | 0 | 0 | 0 | 0 | 2 mo | HOM |
| 1009934102 | 2 | Ald2/3 | 328 | 0.95348559 | 0 | 0 | 0 | 0 | 4 mo | WILD |
| 1009322937 | 2 | Ald2/3 | 328 | 0.95348559 | 4.77E-06 | 1 | 5E-06 | 1.048783548 | 8 mo | WILD |
| 1011080637 | 2 | Ald2/3 | 328 | 0.95348559 | 0.000356947 | 22 | 0.00037436 | 23.07323805 | 8 mo | HOM |
| 1013609713 | 2 | Ald2/3 | 328 | 0.95348559 | 0 | 0 | 0 | 0 | 2 mo | WILD |
| 1010418842 | 1 | Ald2/3 | 328 | 0.95348559 | 0.000554667 | 27 | 0.000581726 | 28.31715578 | 8 mo | HOM |
| 1013288213 | 2 | Ald2/3 | 328 | 0.95348559 | 1.3E-05 | 2 | 1.36E-05 | 2.097567095 | 4 mo | HOM |
| 1008976060 | 2 | Ald2/3 | 328 | 0.95348559 | 3.25E-09 | 0 | 3.41E-09 | 0 | 8 mo | WILD |
| 1013446243 | 1 | Ald2/3 | 328 | 0.95348559 | 0.00012103 | 6 | 0.000126934 | 6.292701285 | 4 mo | HOM |
| 1013288951 | 2 | Ald2/3 | 328 | 0.95348559 | 4.24E-05 | 1 | 4.44E-05 | 1.048783548 | 4 mo | HOM |
| 1013978665 | 1 | SSp-bfd | 329 | 6.477828234 | 0 | 0 | 0 | 0 | 2 mo | HOM |
| 1013445609 | 1 | SSp-bfd | 329 | 6.477828234 | 0.000842483 | 48 | 0.000130056 | 7.409890826 | 4 mo | HOM |
| 1013609015 | 1 | SSp-bfd | 329 | 6.477828234 | 0.002489914 | 89 | 0.000384375 | 13.73917257 | 4 mo | HOM |
| 1013608106 | 2 | SSp-bfd | 329 | 6.477828234 | 0 | 0 | 0 | 0 | 4 mo | HOM |
| 1011937543 | 1 | SSp-bfd | 329 | 6.477828234 | 0.014928599 | 196 | 0.002304568 | 30.2570542 | 8 mo | HOM |
| 1010419538 | 2 | SSp-bfd | 329 | 6.477828234 | 0 | 0 | 0 | 0 | 4 mo | WILD |
| 1010419187 | 1 | SSp-bfd | 329 | 6.477828234 | 0.014382661 | 391 | 0.002220291 | 60.35973568 | 8 mo | HOM |
| 1013978366 | 1 | SSp-bfd | 329 | 6.477828234 | 3.11E-05 | 5 | 4.8E-06 | 0.771863628 | 2 mo | HOM |
| 1014416155 | 2 | SSp-bfd | 329 | 6.477828234 | 0 | 0 | 0 | 0 | 2 mo | HOM |
| 1013978065 | 2 | SSp-bfd | 329 | 6.477828234 | 0 | 0 | 0 | 0 | 2 mo | HOM |
| 1009934102 | 2 | SSp-bfd | 329 | 6.477828234 | 0 | 0 | 0 | 0 | 4 mo | WILD |
| 1009322937 | 2 | SSp-bfd | 329 | 6.477828234 | 1.54E-05 | 3 | 2.38E-06 | 0.463118177 | 8 mo | WILD |
| 1011080637 | 2 | SSp-bfd | 329 | 6.477828234 | 0.008008774 | 286 | 0.001236336 | 44.1505995 | 8 mo | HOM |
| 1013609713 | 2 | SSp-bfd | 329 | 6.477828234 | 1.88E-05 | 1 | 2.9E-06 | 0.154372726 | 2 mo | WILD |
| 1010418842 | 1 | SSp-bfd | 329 | 6.477828234 | 0.013157983 | 305 | 0.002031234 | 47.08368129 | 8 mo | HOM |
| 1013288213 | 2 | SSp-bfd | 329 | 6.477828234 | 0.000585028 | 51 | 9.03E-05 | 7.873009002 | 4 mo | HOM |
| 1008976060 | 2 | SSp-bfd | 329 | 6.477828234 | 0.000293773 | 18 | 4.54E-05 | 2.77870906 | 8 mo | WILD |
| 1013446243 | 1 | SSp-bfd | 329 | 6.477828234 | 0.001821169 | 79 | 0.000281139 | 12.19544532 | 4 mo | HOM |
| 1013288951 | 2 | SSp-bfd | 329 | 6.477828234 | 0.001967615 | 85 | 0.000303746 | 13.12168167 | 4 mo | HOM |
| 1013978665 | 1 | RSPd6b | 330 | 0.031947277 | 0 | 0 | 0 | 0 | 2 mo | HOM |

| | | | | | | | | | | |
|---|---|---|---|---|---|---|---|---|---|---|
| **1013445609** | 1 | RSPd6b | 330 | 0.031947277 | 0 | 0 | 0 | 0 | 4 mo | HOM |
| **1013609015** | 1 | RSPd6b | 330 | 0.031947277 | 0 | 0 | 0 | 0 | 4 mo | HOM |
| **1013608106** | 2 | RSPd6b | 330 | 0.031947277 | 0 | 0 | 0 | 0 | 4 mo | HOM |
| **1011937543** | 1 | RSPd6b | 330 | 0.031947277 | 2.06E-05 | 0 | 0.000645876 | 0 | 8 mo | HOM |
| **1010419538** | 2 | RSPd6b | 330 | 0.031947277 | 0 | 0 | 0 | 0 | 4 mo | WILD |
| **1010419187** | 1 | RSPd6b | 330 | 0.031947277 | 2.33E-05 | 1 | 0.000728274 | 31.30157203 | 8 mo | HOM |
| **1013978366** | 1 | RSPd6b | 330 | 0.031947277 | 0 | 0 | 0 | 0 | 2 mo | HOM |
| **1014416155** | 2 | RSPd6b | 330 | 0.031947277 | 0 | 0 | 0 | 0 | 2 mo | HOM |
| **1013978065** | 2 | RSPd6b | 330 | 0.031947277 | 0 | 0 | 0 | 0 | 2 mo | HOM |
| **1009934102** | 2 | RSPd6b | 330 | 0.031947277 | 0 | 0 | 0 | 0 | 4 mo | WILD |
| **1009322937** | 2 | RSPd6b | 330 | 0.031947277 | 0 | 0 | 0 | 0 | 8 mo | WILD |
| **1011080637** | 2 | RSPd6b | 330 | 0.031947277 | 1.62E-05 | 1 | 0.000505873 | 31.30157203 | 8 mo | HOM |
| **1013609713** | 2 | RSPd6b | 330 | 0.031947277 | 0 | 0 | 0 | 0 | 2 mo | WILD |
| **1010418842** | 1 | RSPd6b | 330 | 0.031947277 | 3.71E-05 | 0 | 0.001159918 | 0 | 8 mo | HOM |
| **1013288213** | 2 | RSPd6b | 330 | 0.031947277 | 0 | 0 | 0 | 0 | 4 mo | HOM |
| **1008976060** | 2 | RSPd6b | 330 | 0.031947277 | 0 | 0 | 0 | 0 | 8 mo | WILD |
| **1013446243** | 1 | RSPd6b | 330 | 0.031947277 | 0 | 0 | 0 | 0 | 4 mo | HOM |
| **1013288951** | 2 | RSPd6b | 330 | 0.031947277 | 1.78E-06 | 0 | 5.57E-05 | 0 | 4 mo | HOM |
| **1013978665** | 1 | MBO | 331 | 1.04651285 | 1.52E-06 | 0 | 1.45E-06 | 0 | 2 mo | HOM |
| **1013445609** | 1 | MBO | 331 | 1.04651285 | 2.09E-07 | 0 | 2E-07 | 0 | 4 mo | HOM |
| **1013609015** | 1 | MBO | 331 | 1.04651285 | 2.79E-05 | 3 | 2.66E-05 | 2.866663319 | 4 mo | HOM |
| **1013608106** | 2 | MBO | 331 | 1.04651285 | 0 | 0 | 0 | 0 | 4 mo | HOM |
| **1011937543** | 1 | MBO | 331 | 1.04651285 | 0.00324039 | 23 | 0.003096369 | 21.97775211 | 8 mo | HOM |
| **1010419538** | 2 | MBO | 331 | 1.04651285 | 3.06E-06 | 0 | 2.93E-06 | 0 | 4 mo | WILD |
| **1010419187** | 1 | MBO | 331 | 1.04651285 | 0.002571828 | 7 | 0.002457521 | 6.688881078 | 8 mo | HOM |
| **1013978366** | 1 | MBO | 331 | 1.04651285 | 1.37E-05 | 1 | 1.31E-05 | 0.95555444 | 2 mo | HOM |
| **1014416155** | 2 | MBO | 331 | 1.04651285 | 0 | 0 | 0 | 0 | 2 mo | HOM |
| **1013978065** | 2 | MBO | 331 | 1.04651285 | 4.08E-06 | 1 | 3.89E-06 | 0.95555444 | 2 mo | HOM |
| **1009934102** | 2 | MBO | 331 | 1.04651285 | 7.92E-06 | 1 | 7.57E-06 | 0.95555444 | 4 mo | WILD |
| **1009322937** | 2 | MBO | 331 | 1.04651285 | 5.57E-11 | 0 | 5.32E-11 | 0 | 8 mo | WILD |
| **1011080637** | 2 | MBO | 331 | 1.04651285 | 0.000119073 | 5 | 0.00011378 | 4.777772199 | 8 mo | HOM |
| **1013609713** | 2 | MBO | 331 | 1.04651285 | 0 | 0 | 0 | 0 | 2 mo | WILD |
| **1010418842** | 1 | MBO | 331 | 1.04651285 | 0.000182271 | 12 | 0.00017417 | 11.46665328 | 8 mo | HOM |
| **1013288213** | 2 | MBO | 331 | 1.04651285 | 1.16E-05 | 2 | 1.11E-05 | 1.91110888 | 4 mo | HOM |
| **1008976060** | 2 | MBO | 331 | 1.04651285 | 0.000275183 | 10 | 0.000262952 | 9.555544398 | 8 mo | WILD |
| **1013446243** | 1 | MBO | 331 | 1.04651285 | 0 | 0 | 0 | 0 | 4 mo | HOM |
| **1013288951** | 2 | MBO | 331 | 1.04651285 | 5.22E-06 | 1 | 4.99E-06 | 0.95555444 | 4 mo | HOM |
| **1013978665** | 1 | ASO | 332 | 0.006401818 | 0 | 0 | 0 | 0 | 2 mo | HOM |
| **1013445609** | 1 | ASO | 332 | 0.006401818 | 0 | 0 | 0 | 0 | 4 mo | HOM |
| **1013609015** | 1 | ASO | 332 | 0.006401818 | 0 | 0 | 0 | 0 | 4 mo | HOM |
| **1013608106** | 2 | ASO | 332 | 0.006401818 | 0 | 0 | 0 | 0 | 4 mo | HOM |
| **1011937543** | 1 | ASO | 332 | 0.006401818 | 0 | 0 | 0 | 0 | 8 mo | HOM |
| **1010419538** | 2 | ASO | 332 | 0.006401818 | 0 | 0 | 0 | 0 | 4 mo | WILD |
| **1010419187** | 1 | ASO | 332 | 0.006401818 | 0 | 0 | 0 | 0 | 8 mo | HOM |
| **1013978366** | 1 | ASO | 332 | 0.006401818 | 0 | 0 | 0 | 0 | 2 mo | HOM |
| **1014416155** | 2 | ASO | 332 | 0.006401818 | 0 | 0 | 0 | 0 | 2 mo | HOM |
| **1013978065** | 2 | ASO | 332 | 0.006401818 | 0 | 0 | 0 | 0 | 2 mo | HOM |
| **1009934102** | 2 | ASO | 332 | 0.006401818 | 0 | 0 | 0 | 0 | 4 mo | WILD |
| **1009322937** | 2 | ASO | 332 | 0.006401818 | 0 | 0 | 0 | 0 | 8 mo | WILD |
| **1011080637** | 2 | ASO | 332 | 0.006401818 | 0 | 0 | 0 | 0 | 8 mo | HOM |
| **1013609713** | 2 | ASO | 332 | 0.006401818 | 0 | 0 | 0 | 0 | 2 mo | WILD |
| **1010418842** | 1 | ASO | 332 | 0.006401818 | 0 | 0 | 0 | 0 | 8 mo | HOM |
| **1013288213** | 2 | ASO | 332 | 0.006401818 | 0 | 0 | 0 | 0 | 4 mo | HOM |
| **1008976060** | 2 | ASO | 332 | 0.006401818 | 0 | 0 | 0 | 0 | 8 mo | WILD |
| **1013446243** | 1 | ASO | 332 | 0.006401818 | 0 | 0 | 0 | 0 | 4 mo | HOM |
| **1013288951** | 2 | ASO | 332 | 0.006401818 | 0 | 0 | 0 | 0 | 4 mo | HOM |
| **1013978665** | 1 | SH | 333 | 0.038114204 | 0 | 0 | 0 | 0 | 2 mo | HOM |
| **1013445609** | 1 | SH | 333 | 0.038114204 | 7.61E-06 | 1 | 0.00019976 | 26.23693774 | 4 mo | HOM |
| **1013609015** | 1 | SH | 333 | 0.038114204 | 0 | 0 | 0 | 0 | 4 mo | HOM |

| | | | | | | | | | | |
|---|---|---|---|---|---|---|---|---|---|---|
| 1013608106 | 2 | SH | 333 | 0.038114204 | 0 | 0 | 0 | 0 | 4 mo | HOM |
| 1011937543 | 1 | SH | 333 | 0.038114204 | 4.8E-05 | 2 | 0.001259964 | 52.47387548 | 8 mo | HOM |
| 1010419538 | 2 | SH | 333 | 0.038114204 | 0 | 0 | 0 | 0 | 4 mo | WILD |
| 1010419187 | 1 | SH | 333 | 0.038114204 | 5.44E-05 | 1 | 0.001427302 | 26.23693774 | 8 mo | HOM |
| 1013978366 | 1 | SH | 333 | 0.038114204 | 0 | 0 | 0 | 0 | 2 mo | HOM |
| 1014416155 | 2 | SH | 333 | 0.038114204 | 0 | 0 | 0 | 0 | 2 mo | HOM |
| 1013978065 | 2 | SH | 333 | 0.038114204 | 0 | 0 | 0 | 0 | 2 mo | HOM |
| 1009934102 | 2 | SH | 333 | 0.038114204 | 0 | 0 | 0 | 0 | 4 mo | WILD |
| 1009322937 | 2 | SH | 333 | 0.038114204 | 0 | 0 | 0 | 0 | 8 mo | WILD |
| 1011080637 | 2 | SH | 333 | 0.038114204 | 2.19E-05 | 1 | 0.000574445 | 26.23693774 | 8 mo | HOM |
| 1013609713 | 2 | SH | 333 | 0.038114204 | 0 | 0 | 0 | 0 | 2 mo | WILD |
| 1010418842 | 1 | SH | 333 | 0.038114204 | 0.000158211 | 4 | 0.004150979 | 104.947751 | 8 mo | HOM |
| 1013288213 | 2 | SH | 333 | 0.038114204 | 0 | 0 | 0 | 0 | 4 mo | HOM |
| 1008976060 | 2 | SH | 333 | 0.038114204 | 0 | 0 | 0 | 0 | 8 mo | WILD |
| 1013446243 | 1 | SH | 333 | 0.038114204 | 0 | 0 | 0 | 0 | 4 mo | HOM |
| 1013288951 | 2 | SH | 333 | 0.038114204 | 9.22E-08 | 0 | 2.42E-06 | 0 | 4 mo | HOM |
| 1013978665 | 1 | BMAp | 334 | 0.73946049 | 0 | 0 | 0 | 0 | 2 mo | HOM |
| 1013445609 | 1 | BMAp | 334 | 0.73946049 | 9.3E-05 | 6 | 0.000125805 | 8.114023784 | 4 mo | HOM |
| 1013609015 | 1 | BMAp | 334 | 0.73946049 | 0.000244016 | 8 | 0.000329992 | 10.81869838 | 4 mo | HOM |
| 1013608106 | 2 | BMAp | 334 | 0.73946049 | 0 | 0 | 0 | 0 | 4 mo | HOM |
| 1011937543 | 1 | BMAp | 334 | 0.73946049 | 0.002235162 | 26 | 0.003022693 | 35.16076973 | 8 mo | HOM |
| 1010419538 | 2 | BMAp | 334 | 0.73946049 | 0 | 0 | 0 | 0 | 4 mo | WILD |
| 1010419187 | 1 | BMAp | 334 | 0.73946049 | 0.00134063 | 52 | 0.001812984 | 70.32153946 | 8 mo | HOM |
| 1013978366 | 1 | BMAp | 334 | 0.73946049 | 0 | 0 | 0 | 0 | 2 mo | HOM |
| 1014416155 | 2 | BMAp | 334 | 0.73946049 | 0 | 0 | 0 | 0 | 2 mo | HOM |
| 1013978065 | 2 | BMAp | 334 | 0.73946049 | 0 | 0 | 0 | 0 | 2 mo | HOM |
| 1009934102 | 2 | BMAp | 334 | 0.73946049 | 0 | 0 | 0 | 0 | 4 mo | WILD |
| 1009322937 | 2 | BMAp | 334 | 0.73946049 | 0 | 0 | 0 | 0 | 8 mo | WILD |
| 1011080637 | 2 | BMAp | 334 | 0.73946049 | 0.000954718 | 44 | 0.001291101 | 59.50284108 | 8 mo | HOM |
| 1013609713 | 2 | BMAp | 334 | 0.73946049 | 0 | 0 | 0 | 0 | 2 mo | WILD |
| 1010418842 | 1 | BMAp | 334 | 0.73946049 | 0.00142278 | 44 | 0.001924079 | 59.50284108 | 8 mo | HOM |
| 1013288213 | 2 | BMAp | 334 | 0.73946049 | 6.33E-05 | 4 | 8.55E-05 | 5.409349189 | 4 mo | HOM |
| 1008976060 | 2 | BMAp | 334 | 0.73946049 | 6.5E-06 | 1 | 8.79E-06 | 1.352337297 | 8 mo | WILD |
| 1013446243 | 1 | BMAp | 334 | 0.73946049 | 0.000130908 | 7 | 0.000177032 | 9.466361081 | 4 mo | HOM |
| 1013288951 | 2 | BMAp | 334 | 0.73946049 | 0.000207695 | 5 | 0.000280873 | 6.761686487 | 4 mo | HOM |
| 1013978665 | 1 | PERI6a | 335 | 0.055522949 | 0 | 0 | 0 | 0 | 2 mo | HOM |
| 1013445609 | 1 | PERI6a | 335 | 0.055522949 | 1.61E-08 | 0 | 2.9E-07 | 0 | 4 mo | HOM |
| 1013609015 | 1 | PERI6a | 335 | 0.055522949 | 1.21E-06 | 0 | 2.18E-05 | 0 | 4 mo | HOM |
| 1013608106 | 2 | PERI6a | 335 | 0.055522949 | 0 | 0 | 0 | 0 | 4 mo | HOM |
| 1011937543 | 1 | PERI6a | 335 | 0.055522949 | 0.000143675 | 5 | 0.002587667 | 90.05285429 | 8 mo | HOM |
| 1010419538 | 2 | PERI6a | 335 | 0.055522949 | 0 | 0 | 0 | 0 | 4 mo | WILD |
| 1010419187 | 1 | PERI6a | 335 | 0.055522949 | 5.44E-05 | 2 | 0.000980206 | 36.02114172 | 8 mo | HOM |
| 1013978366 | 1 | PERI6a | 335 | 0.055522949 | 0 | 0 | 0 | 0 | 2 mo | HOM |
| 1014416155 | 2 | PERI6a | 335 | 0.055522949 | 0 | 0 | 0 | 0 | 2 mo | HOM |
| 1013978065 | 2 | PERI6a | 335 | 0.055522949 | 0 | 0 | 0 | 0 | 2 mo | HOM |
| 1009934102 | 2 | PERI6a | 335 | 0.055522949 | 0 | 0 | 0 | 0 | 4 mo | WILD |
| 1009322937 | 2 | PERI6a | 335 | 0.055522949 | 0 | 0 | 0 | 0 | 8 mo | WILD |
| 1011080637 | 2 | PERI6a | 335 | 0.055522949 | 5.58E-05 | 2 | 0.001005349 | 36.02114172 | 8 mo | HOM |
| 1013609713 | 2 | PERI6a | 335 | 0.055522949 | 0 | 0 | 0 | 0 | 2 mo | WILD |
| 1010418842 | 1 | PERI6a | 335 | 0.055522949 | 8.9E-05 | 4 | 0.001602807 | 72.04228343 | 8 mo | HOM |
| 1013288213 | 2 | PERI6a | 335 | 0.055522949 | 1.42E-07 | 0 | 2.55E-06 | 0 | 4 mo | HOM |
| 1008976060 | 2 | PERI6a | 335 | 0.055522949 | 0 | 0 | 0 | 0 | 8 mo | WILD |
| 1013446243 | 1 | PERI6a | 335 | 0.055522949 | 1.89E-05 | 0 | 0.0003396 | 0 | 4 mo | HOM |
| 1013288951 | 2 | PERI6a | 335 | 0.055522949 | 2.75E-10 | 0 | 4.95E-09 | 0 | 4 mo | HOM |
| 1013978665 | 1 | csc | 336 | 0.028728854 | 0 | 0 | 0 | 0 | 2 mo | HOM |
| 1013445609 | 1 | csc | 336 | 0.028728854 | 0 | 0 | 0 | 0 | 4 mo | HOM |
| 1013609015 | 1 | csc | 336 | 0.028728854 | 0 | 0 | 0 | 0 | 4 mo | HOM |
| 1013608106 | 2 | csc | 336 | 0.028728854 | 0 | 0 | 0 | 0 | 4 mo | HOM |
| 1011937543 | 1 | csc | 336 | 0.028728854 | 0 | 0 | 0 | 0 | 8 mo | HOM |

| | | | | | | | | | | |
|---|---|---|---|---|---|---|---|---|---|---|
| **1010419538** | 2 | csc | 336 | 0.028728854 | 0 | 0 | 0 | 0 | 4 mo | WILD |
| **1010419187** | 1 | csc | 336 | 0.028728854 | 0 | 0 | 0 | 0 | 8 mo | HOM |
| **1013978366** | 1 | csc | 336 | 0.028728854 | 0 | 0 | 0 | 0 | 2 mo | HOM |
| **1014416155** | 2 | csc | 336 | 0.028728854 | 0 | 0 | 0 | 0 | 2 mo | HOM |
| **1013978065** | 2 | csc | 336 | 0.028728854 | 0 | 0 | 0 | 0 | 2 mo | HOM |
| **1009934102** | 2 | csc | 336 | 0.028728854 | 0 | 0 | 0 | 0 | 4 mo | WILD |
| **1009322937** | 2 | csc | 336 | 0.028728854 | 0 | 0 | 0 | 0 | 8 mo | WILD |
| **1011080637** | 2 | csc | 336 | 0.028728854 | 0 | 0 | 0 | 0 | 8 mo | HOM |
| **1013609713** | 2 | csc | 336 | 0.028728854 | 0 | 0 | 0 | 0 | 2 mo | WILD |
| **1010418842** | 1 | csc | 336 | 0.028728854 | 0 | 0 | 0 | 0 | 8 mo | HOM |
| **1013288213** | 2 | csc | 336 | 0.028728854 | 0 | 0 | 0 | 0 | 4 mo | HOM |
| **1008976060** | 2 | csc | 336 | 0.028728854 | 1.3E-06 | 0 | 4.53E-05 | 0 | 8 mo | WILD |
| **1013446243** | 1 | csc | 336 | 0.028728854 | 0 | 0 | 0 | 0 | 4 mo | HOM |
| **1013288951** | 2 | csc | 336 | 0.028728854 | 0 | 0 | 0 | 0 | 4 mo | HOM |
| **1013978665** | 1 | SSp-II | 337 | 2.423219011 | 2.64E-05 | 1 | 1.09E-05 | 0.412674214 | 2 mo | HOM |
| **1013445609** | 1 | SSp-II | 337 | 2.423219011 | 0.000165631 | 9 | 6.84E-05 | 3.714067924 | 4 mo | HOM |
| **1013609015** | 1 | SSp-II | 337 | 2.423219011 | 0.000988528 | 29 | 0.00040794 | 11.9675522 | 4 mo | HOM |
| **1013608106** | 2 | SSp-II | 337 | 2.423219011 | 0 | 0 | 0 | 0 | 4 mo | HOM |
| **1011937543** | 1 | SSp-II | 337 | 2.423219011 | 0.00506754 | 72 | 0.002091243 | 29.71254339 | 8 mo | HOM |
| **1010419538** | 2 | SSp-II | 337 | 2.423219011 | 0 | 0 | 0 | 0 | 4 mo | WILD |
| **1010419187** | 1 | SSp-II | 337 | 2.423219011 | 0.004898974 | 139 | 0.00202168 | 57.36171571 | 8 mo | HOM |
| **1013978366** | 1 | SSp-II | 337 | 2.423219011 | 0 | 0 | 0 | 0 | 2 mo | HOM |
| **1014416155** | 2 | SSp-II | 337 | 2.423219011 | 0 | 0 | 0 | 0 | 2 mo | HOM |
| **1013978065** | 2 | SSp-II | 337 | 2.423219011 | 4.42E-06 | 1 | 1.83E-06 | 0.412674214 | 2 mo | HOM |
| **1009934102** | 2 | SSp-II | 337 | 2.423219011 | 7.31E-05 | 3 | 3.02E-05 | 1.238022641 | 4 mo | WILD |
| **1009322937** | 2 | SSp-II | 337 | 2.423219011 | 8.24E-06 | 1 | 3.4E-06 | 0.412674214 | 8 mo | WILD |
| **1011080637** | 2 | SSp-II | 337 | 2.423219011 | 0.001879288 | 78 | 0.000775534 | 32.18858867 | 8 mo | HOM |
| **1013609713** | 2 | SSp-II | 337 | 2.423219011 | 3.73E-06 | 1 | 1.54E-06 | 0.412674214 | 2 mo | WILD |
| **1010418842** | 1 | SSp-II | 337 | 2.423219011 | 0.003693566 | 117 | 0.00152424 | 48.28288301 | 8 mo | HOM |
| **1013288213** | 2 | SSp-II | 337 | 2.423219011 | 0.000130206 | 8 | 5.37E-05 | 3.30139371 | 4 mo | HOM |
| **1008976060** | 2 | SSp-II | 337 | 2.423219011 | 0.000150612 | 9 | 6.22E-05 | 3.714067924 | 8 mo | WILD |
| **1013446243** | 1 | SSp-II | 337 | 2.423219011 | 0.000899602 | 28 | 0.000371243 | 11.55487798 | 4 mo | HOM |
| **1013288951** | 2 | SSp-II | 337 | 2.423219011 | 0.000618676 | 27 | 0.000255312 | 11.14220377 | 4 mo | HOM |
| **1013978665** | 1 | SFO | 338 | 0.021342141 | 3.71E-05 | 1 | 0.001738143 | 46.85565498 | 2 mo | HOM |
| **1013445609** | 1 | SFO | 338 | 0.021342141 | 0.000154832 | 3 | 0.007254758 | 140.566965 | 4 mo | HOM |
| **1013609015** | 1 | SFO | 338 | 0.021342141 | 0.000163715 | 2 | 0.007670967 | 93.71130997 | 4 mo | HOM |
| **1013608106** | 2 | SFO | 338 | 0.021342141 | 0 | 0 | 0 | 0 | 4 mo | HOM |
| **1011937543** | 1 | SFO | 338 | 0.021342141 | 7.88E-10 | 0 | 3.69E-08 | 0 | 8 mo | HOM |
| **1010419538** | 2 | SFO | 338 | 0.021342141 | 0 | 0 | 0 | 0 | 4 mo | WILD |
| **1010419187** | 1 | SFO | 338 | 0.021342141 | 0.00070997 | 2 | 0.033266108 | 93.71130997 | 8 mo | HOM |
| **1013978366** | 1 | SFO | 338 | 0.021342141 | 0.000252838 | 2 | 0.011846912 | 93.71130997 | 2 mo | HOM |
| **1014416155** | 2 | SFO | 338 | 0.021342141 | 0 | 0 | 0 | 0 | 2 mo | HOM |
| **1013978065** | 2 | SFO | 338 | 0.021342141 | 4.92E-22 | 0 | 2.3E-20 | 0 | 2 mo | HOM |
| **1009934102** | 2 | SFO | 338 | 0.021342141 | 1.66E-06 | 0 | 7.79E-05 | 0 | 4 mo | WILD |
| **1009322937** | 2 | SFO | 338 | 0.021342141 | 4.03E-05 | 2 | 0.001888864 | 93.71130997 | 8 mo | WILD |
| **1011080637** | 2 | SFO | 338 | 0.021342141 | 0.000367698 | 0 | 0.017228751 | 0 | 8 mo | HOM |
| **1013609713** | 2 | SFO | 338 | 0.021342141 | 0 | 0 | 0 | 0 | 2 mo | WILD |
| **1010418842** | 1 | SFO | 338 | 0.021342141 | 0.000499378 | 0 | 0.023398686 | 0 | 8 mo | HOM |
| **1013288213** | 2 | SFO | 338 | 0.021342141 | 0 | 0 | 0 | 0 | 4 mo | HOM |
| **1008976060** | 2 | SFO | 338 | 0.021342141 | 4.36E-05 | 0 | 0.00204161 | 0 | 8 mo | WILD |
| **1013446243** | 1 | SFO | 338 | 0.021342141 | 0.000690726 | 1 | 0.032364398 | 46.85565498 | 4 mo | HOM |
| **1013288951** | 2 | SFO | 338 | 0.021342141 | 0.000368024 | 0 | 0.017243985 | 0 | 4 mo | HOM |
| **1013978665** | 1 | MBsen | 339 | 7.064110693 | 8.68E-05 | 8 | 1.23E-05 | 1.132485085 | 2 mo | HOM |
| **1013445609** | 1 | MBsen | 339 | 7.064110693 | 0 | 0 | 0 | 0 | 4 mo | HOM |
| **1013609015** | 1 | MBsen | 339 | 7.064110693 | 0.000149136 | 6 | 2.11E-05 | 0.849363814 | 4 mo | HOM |
| **1013608106** | 2 | MBsen | 339 | 7.064110693 | 0 | 0 | 0 | 0 | 4 mo | HOM |
| **1011937543** | 1 | MBsen | 339 | 7.064110693 | 0.01599486 | 138 | 0.002264243 | 19.53536772 | 8 mo | HOM |
| **1010419538** | 2 | MBsen | 339 | 7.064110693 | 6.23E-06 | 1 | 8.82E-07 | 0.141560636 | 4 mo | WILD |
| **1010419187** | 1 | MBsen | 339 | 7.064110693 | 0.002283792 | 63 | 0.000323295 | 8.918320046 | 8 mo | HOM |

| | | | | | | | | | | |
|---|---|---|---|---|---|---|---|---|---|---|
| **1013978366** | 1 | MBsen | 339 | 7.064110693 | 0.000100595 | 3 | 1.42E-05 | 0.424681907 | 2 mo | HOM |
| **1014416155** | 2 | MBsen | 339 | 7.064110693 | 0 | 0 | 0 | 0 | 2 mo | HOM |
| **1013978065** | 2 | MBsen | 339 | 7.064110693 | 0 | 0 | 0 | 0 | 2 mo | HOM |
| **1009934102** | 2 | MBsen | 339 | 7.064110693 | 2.03E-05 | 2 | 2.88E-06 | 0.283121271 | 4 mo | WILD |
| **1009322937** | 2 | MBsen | 339 | 7.064110693 | 5.07E-05 | 2 | 7.18E-06 | 0.283121271 | 8 mo | WILD |
| **1011080637** | 2 | MBsen | 339 | 7.064110693 | 0.001343298 | 57 | 0.000190158 | 8.068956232 | 8 mo | HOM |
| **1013609713** | 2 | MBsen | 339 | 7.064110693 | 1.41E-05 | 2 | 1.99E-06 | 0.283121271 | 2 mo | WILD |
| **1010418842** | 1 | MBsen | 339 | 7.064110693 | 0.002530462 | 87 | 0.000358214 | 12.3157753 | 8 mo | HOM |
| **1013288213** | 2 | MBsen | 339 | 7.064110693 | 1.37E-05 | 0 | 1.94E-06 | 0 | 4 mo | HOM |
| **1008976060** | 2 | MBsen | 339 | 7.064110693 | 0.000375284 | 25 | 5.31E-05 | 3.539015891 | 8 mo | WILD |
| **1013446243** | 1 | MBsen | 339 | 7.064110693 | 5.03E-05 | 2 | 7.12E-06 | 0.283121271 | 4 mo | HOM |
| **1013288951** | 2 | MBsen | 339 | 7.064110693 | 0.000382558 | 21 | 5.42E-05 | 2.972773349 | 4 mo | HOM |
| **1013978665** | 1 | SI | 342 | 3.118454707 | 0 | 0 | 0 | 0 | 2 mo | HOM |
| **1013445609** | 1 | SI | 342 | 3.11860649 | 2.17E-06 | 1 | 6.97E-07 | 0.320656038 | 4 mo | HOM |
| **1013609015** | 1 | SI | 342 | 3.11860649 | 0.000318601 | 4 | 0.000102161 | 1.282624151 | 4 mo | HOM |
| **1013608106** | 2 | SI | 342 | 3.11860649 | 0 | 0 | 0 | 0 | 4 mo | HOM |
| **1011937543** | 1 | SI | 342 | 3.11860649 | 0.00160667 | 33 | 0.000515189 | 10.58164924 | 8 mo | HOM |
| **1010419538** | 2 | SI | 342 | 3.11860649 | 0 | 0 | 0 | 0 | 4 mo | WILD |
| **1010419187** | 1 | SI | 342 | 3.11860649 | 0.001214 | 64 | 0.000389277 | 20.52198641 | 8 mo | HOM |
| **1013978366** | 1 | SI | 342 | 3.11860649 | 0 | 0 | 0 | 0 | 2 mo | HOM |
| **1014416155** | 2 | SI | 342 | 3.11860649 | 0 | 0 | 0 | 0 | 2 mo | HOM |
| **1013978065** | 2 | SI | 342 | 3.11860649 | 0 | 0 | 0 | 0 | 2 mo | HOM |
| **1009934102** | 2 | SI | 342 | 3.11860649 | 0 | 0 | 0 | 0 | 4 mo | WILD |
| **1009322937** | 2 | SI | 342 | 3.11860649 | 0 | 0 | 0 | 0 | 8 mo | WILD |
| **1011080637** | 2 | SI | 342 | 3.11860649 | 0.001132698 | 53 | 0.000363207 | 16.99477 | 8 mo | HOM |
| **1013609713** | 2 | SI | 342 | 3.11860649 | 0 | 0 | 0 | 0 | 2 mo | WILD |
| **1010418842** | 1 | SI | 342 | 3.11860649 | 0.001994549 | 87 | 0.000639564 | 27.89707528 | 8 mo | HOM |
| **1013288213** | 2 | SI | 342 | 3.11860649 | 1.24E-05 | 1 | 3.97E-06 | 0.320656038 | 4 mo | HOM |
| **1008976060** | 2 | SI | 342 | 3.11860649 | 2.06E-05 | 4 | 6.62E-06 | 1.282624151 | 8 mo | WILD |
| **1013446243** | 1 | SI | 342 | 3.11860649 | 7.75E-05 | 4 | 2.49E-05 | 1.282624151 | 4 mo | HOM |
| **1013288951** | 2 | SI | 342 | 3.11860649 | 4.51E-06 | 1 | 1.44E-06 | 0.320656038 | 4 mo | HOM |
| **1013978665** | 1 | BS | 343 | 123.5155117 | 0.001622671 | 46 | 1.31E-05 | 0.372422859 | 2 mo | HOM |
| **1013445609** | 1 | BS | 343 | 123.5155117 | 0.000702373 | 37 | 5.69E-06 | 0.299557517 | 4 mo | HOM |
| **1013609015** | 1 | BS | 343 | 123.5155117 | 0.019448925 | 120 | 0.000157461 | 0.971537893 | 4 mo | HOM |
| **1013608106** | 2 | BS | 343 | 123.4456554 | 0 | 0 | 0 | 0 | 4 mo | HOM |
| **1011937543** | 1 | BS | 343 | 123.5155117 | 0.184119762 | 1343 | 0.001490661 | 10.87312825 | 8 mo | HOM |
| **1010419538** | 2 | BS | 343 | 123.5155117 | 0.000210188 | 20 | 1.7E-06 | 0.161922982 | 4 mo | WILD |
| **1010419187** | 1 | BS | 343 | 123.5155117 | 0.04679675 | 665 | 0.000378873 | 5.383939155 | 8 mo | HOM |
| **1013978366** | 1 | BS | 343 | 123.5155117 | 0.003250172 | 50 | 2.63E-05 | 0.404807455 | 2 mo | HOM |
| **1014416155** | 2 | BS | 343 | 123.5155117 | 0 | 0 | 0 | 0 | 2 mo | HOM |
| **1013978065** | 2 | BS | 343 | 123.5155117 | 0.000494678 | 12 | 4E-06 | 0.097153789 | 2 mo | HOM |
| **1009934102** | 2 | BS | 343 | 123.5155117 | 0.000161843 | 12 | 1.31E-06 | 0.097153789 | 4 mo | WILD |
| **1009322937** | 2 | BS | 343 | 123.5155117 | 0.00163288 | 79 | 1.32E-05 | 0.639595779 | 8 mo | WILD |
| **1011080637** | 2 | BS | 343 | 123.5155117 | 0.035821147 | 620 | 0.000290013 | 5.019612446 | 8 mo | HOM |
| **1013609713** | 2 | BS | 343 | 123.5155117 | 3.6E-05 | 3 | 2.91E-07 | 0.024288447 | 2 mo | WILD |
| **1010418842** | 1 | BS | 343 | 123.5155117 | 0.046502906 | 1203 | 0.000376494 | 9.739667374 | 8 mo | HOM |
| **1013288213** | 2 | BS | 343 | 123.513939 | 0.000932656 | 37 | 7.55E-06 | 0.299561331 | 4 mo | HOM |
| **1008976060** | 2 | BS | 343 | 123.5155117 | 0.019172384 | 596 | 0.000155222 | 4.825304867 | 8 mo | WILD |
| **1013446243** | 1 | BS | 343 | 123.5155117 | 0.004367817 | 77 | 3.54E-05 | 0.623403481 | 4 mo | HOM |
| **1013288951** | 2 | BS | 343 | 123.5155117 | 0.008282866 | 100 | 6.71E-05 | 0.809614911 | 4 mo | HOM |
| **1013978665** | 1 | Alp5 | 344 | 0.77532261 | 0 | 0 | 0 | 0 | 2 mo | HOM |
| **1013445609** | 1 | Alp5 | 344 | 0.77532261 | 7.05E-05 | 5 | 9.1E-05 | 6.448928405 | 4 mo | HOM |
| **1013609015** | 1 | Alp5 | 344 | 0.77532261 | 0.000167148 | 10 | 0.000215586 | 12.89785681 | 4 mo | HOM |
| **1013608106** | 2 | Alp5 | 344 | 0.77532261 | 0 | 0 | 0 | 0 | 4 mo | HOM |
| **1011937543** | 1 | Alp5 | 344 | 0.77532261 | 0.000899535 | 27 | 0.001160208 | 34.82421339 | 8 mo | HOM |
| **1010419538** | 2 | Alp5 | 344 | 0.77532261 | 0 | 0 | 0 | 0 | 4 mo | WILD |
| **1010419187** | 1 | Alp5 | 344 | 0.77532261 | 0.001265279 | 48 | 0.001631939 | 61.90971269 | 8 mo | HOM |
| **1013978366** | 1 | Alp5 | 344 | 0.77532261 | 0 | 0 | 0 | 0 | 2 mo | HOM |
| **1014416155** | 2 | Alp5 | 344 | 0.77532261 | 0 | 0 | 0 | 0 | 2 mo | HOM |

| | | | | | | | | | | |
|---|---|---|---|---|---|---|---|---|---|---|
| 1013978065 | 2 | Alp5 | 344 | 0.77532261 | 0 | 0 | 0 | 0 | 2 mo | HOM |
| 1009934102 | 2 | Alp5 | 344 | 0.77532261 | 0 | 0 | 0 | 0 | 4 mo | WILD |
| 1009322937 | 2 | Alp5 | 344 | 0.77532261 | 0 | 0 | 0 | 0 | 8 mo | WILD |
| 1011080637 | 2 | Alp5 | 344 | 0.77532261 | 0.000614879 | 37 | 0.000793062 | 47.7220702 | 8 mo | HOM |
| 1013609713 | 2 | Alp5 | 344 | 0.77532261 | 0 | 0 | 0 | 0 | 2 mo | WILD |
| 1010418842 | 1 | Alp5 | 344 | 0.77532261 | 0.000939751 | 41 | 0.001212078 | 52.88121292 | 8 mo | HOM |
| 1013288213 | 2 | Alp5 | 344 | 0.77532261 | 3.94E-05 | 5 | 5.08E-05 | 6.448928405 | 4 mo | HOM |
| 1008976060 | 2 | Alp5 | 344 | 0.77532261 | 0 | 0 | 0 | 0 | 8 mo | WILD |
| 1013446243 | 1 | Alp5 | 344 | 0.77532261 | 0.000119025 | 9 | 0.000153517 | 11.60807113 | 4 mo | HOM |
| 1013288951 | 2 | Alp5 | 344 | 0.77532261 | 0.000238303 | 13 | 0.00030736 | 16.76721385 | 4 mo | HOM |
| 1013978665 | 1 | SSp-m | 345 | 6.397305858 | 1.86E-05 | 1 | 2.9E-06 | 0.156315803 | 2 mo | HOM |
| 1013445609 | 1 | SSp-m | 345 | 6.397305858 | 0.000667923 | 35 | 0.000104407 | 5.471053093 | 4 mo | HOM |
| 1013609015 | 1 | SSp-m | 345 | 6.397305858 | 0.001608331 | 60 | 0.000251408 | 9.37894816 | 4 mo | HOM |
| 1013608106 | 2 | SSp-m | 345 | 6.397305858 | 0 | 0 | 0 | 0 | 4 mo | HOM |
| 1011937543 | 1 | SSp-m | 345 | 6.397305858 | 0.00960178 | 198 | 0.00150091 | 30.95052893 | 8 mo | HOM |
| 1010419538 | 2 | SSp-m | 345 | 6.397305858 | 0 | 0 | 0 | 0 | 4 mo | WILD |
| 1010419187 | 1 | SSp-m | 345 | 6.397305858 | 0.00833545 | 302 | 0.001302963 | 47.2073724 | 8 mo | HOM |
| 1013978366 | 1 | SSp-m | 345 | 6.397305858 | 1.72E-05 | 2 | 2.69E-06 | 0.312631605 | 2 mo | HOM |
| 1014416155 | 2 | SSp-m | 345 | 6.397305858 | 4.49E-06 | 1 | 7.02E-07 | 0.156315803 | 2 mo | HOM |
| 1013978065 | 2 | SSp-m | 345 | 6.397305858 | 0 | 0 | 0 | 0 | 2 mo | HOM |
| 1009934102 | 2 | SSp-m | 345 | 6.397305858 | 2.2E-05 | 3 | 3.43E-06 | 0.468947408 | 4 mo | WILD |
| 1009322937 | 2 | SSp-m | 345 | 6.397305858 | 8.26E-06 | 1 | 1.29E-06 | 0.156315803 | 8 mo | WILD |
| 1011080637 | 2 | SSp-m | 345 | 6.397305858 | 0.005262344 | 206 | 0.000822588 | 32.20105535 | 8 mo | HOM |
| 1013609713 | 2 | SSp-m | 345 | 6.397305858 | 1.27E-05 | 2 | 1.99E-06 | 0.312631605 | 2 mo | WILD |
| 1010418842 | 1 | SSp-m | 345 | 6.397305858 | 0.007424913 | 311 | 0.001160631 | 48.61421463 | 8 mo | HOM |
| 1013288213 | 2 | SSp-m | 345 | 6.397305858 | 0.000407936 | 35 | 6.38E-05 | 5.471053093 | 4 mo | HOM |
| 1008976060 | 2 | SSp-m | 345 | 6.397305858 | 0.000401638 | 10 | 6.28E-05 | 1.563158027 | 8 mo | WILD |
| 1013446243 | 1 | SSp-m | 345 | 6.397305858 | 0.001375274 | 71 | 0.000214977 | 11.09842199 | 4 mo | HOM |
| 1013288951 | 2 | SSp-m | 345 | 6.397305858 | 0.001443233 | 64 | 0.0002256 | 10.00421137 | 4 mo | HOM |
| 1013978665 | 1 | SBPV | 347 | 0.112706615 | 0 | 0 | 0 | 0 | 2 mo | HOM |
| 1013445609 | 1 | SBPV | 347 | 0.112706615 | 0 | 0 | 0 | 0 | 4 mo | HOM |
| 1013609015 | 1 | SBPV | 347 | 0.112706615 | 0 | 0 | 0 | 0 | 4 mo | HOM |
| 1013608106 | 2 | SBPV | 347 | 0.112706615 | 0 | 0 | 0 | 0 | 4 mo | HOM |
| 1011937543 | 1 | SBPV | 347 | 0.112706615 | 0 | 0 | 0 | 0 | 8 mo | HOM |
| 1010419538 | 2 | SBPV | 347 | 0.112706615 | 0 | 0 | 0 | 0 | 4 mo | WILD |
| 1010419187 | 1 | SBPV | 347 | 0.112706615 | 0 | 0 | 0 | 0 | 8 mo | HOM |
| 1013978366 | 1 | SBPV | 347 | 0.112706615 | 0 | 0 | 0 | 0 | 2 mo | HOM |
| 1014416155 | 2 | SBPV | 347 | 0.112706615 | 0 | 0 | 0 | 0 | 2 mo | HOM |
| 1013978065 | 2 | SBPV | 347 | 0.112706615 | 0 | 0 | 0 | 0 | 2 mo | HOM |
| 1009934102 | 2 | SBPV | 347 | 0.112706615 | 0 | 0 | 0 | 0 | 4 mo | WILD |
| 1009322937 | 2 | SBPV | 347 | 0.112706615 | 0 | 0 | 0 | 0 | 8 mo | WILD |
| 1011080637 | 2 | SBPV | 347 | 0.112706615 | 0 | 0 | 0 | 0 | 8 mo | HOM |
| 1013609713 | 2 | SBPV | 347 | 0.112706615 | 0 | 0 | 0 | 0 | 2 mo | WILD |
| 1010418842 | 1 | SBPV | 347 | 0.112706615 | 1.22E-07 | 0 | 1.08E-06 | 0 | 8 mo | HOM |
| 1013288213 | 2 | SBPV | 347 | 0.112706615 | 0 | 0 | 0 | 0 | 4 mo | HOM |
| 1008976060 | 2 | SBPV | 347 | 0.112706615 | 0 | 0 | 0 | 0 | 8 mo | WILD |
| 1013446243 | 1 | SBPV | 347 | 0.112706615 | 0 | 0 | 0 | 0 | 4 mo | HOM |
| 1013288951 | 2 | SBPV | 347 | 0.112706615 | 0 | 0 | 0 | 0 | 4 mo | HOM |
| 1013978665 | 1 | MBsta | 348 | 1.877871494 | 0 | 0 | 0 | 0 | 2 mo | HOM |
| 1013445609 | 1 | MBsta | 348 | 1.877871494 | 0 | 0 | 0 | 0 | 4 mo | HOM |
| 1013609015 | 1 | MBsta | 348 | 1.877871494 | 1.14E-05 | 2 | 6.06E-06 | 1.065035603 | 4 mo | HOM |
| 1013608106 | 2 | MBsta | 348 | 1.877871494 | 0 | 0 | 0 | 0 | 4 mo | HOM |
| 1011937543 | 1 | MBsta | 348 | 1.877871494 | 0.006186543 | 35 | 0.003294444 | 18.63812306 | 8 mo | HOM |
| 1010419538 | 2 | MBsta | 348 | 1.877871494 | 1.83E-07 | 0 | 9.75E-08 | 0 | 4 mo | WILD |
| 1010419187 | 1 | MBsta | 348 | 1.877871494 | 0.000236396 | 8 | 0.000125885 | 4.260142413 | 8 mo | HOM |
| 1013978366 | 1 | MBsta | 348 | 1.877871494 | 0 | 0 | 0 | 0 | 2 mo | HOM |
| 1014416155 | 2 | MBsta | 348 | 1.877871494 | 0 | 0 | 0 | 0 | 2 mo | HOM |
| 1013978065 | 2 | MBsta | 348 | 1.877871494 | 0 | 0 | 0 | 0 | 2 mo | HOM |
| 1009934102 | 2 | MBsta | 348 | 1.877871494 | 1.1E-05 | 1 | 5.85E-06 | 0.532517802 | 4 mo | WILD |

| | | | | | | | | | | |
|---|---|---|---|---|---|---|---|---|---|---|
| **1009322937** | 2 | MBsta | 348 | 1.877871494 | 2.25E-05 | 3 | 1.2E-05 | 1.597553405 | 8 mo | WILD |
| **1011080637** | 2 | MBsta | 348 | 1.877871494 | 3.71E-05 | 2 | 1.97E-05 | 1.065035603 | 8 mo | HOM |
| **1013609713** | 2 | MBsta | 348 | 1.877871494 | 0 | 0 | 0 | 0 | 2 mo | WILD |
| **1010418842** | 1 | MBsta | 348 | 1.877871494 | 0.000201729 | 13 | 0.000107424 | 6.922731421 | 8 mo | HOM |
| **1013288213** | 2 | MBsta | 348 | 1.877871494 | 0 | 0 | 0 | 0 | 4 mo | HOM |
| **1008976060** | 2 | MBsta | 348 | 1.877871494 | 0.000124391 | 8 | 6.62E-05 | 4.260142413 | 8 mo | WILD |
| **1013446243** | 1 | MBsta | 348 | 1.877871494 | 1.11E-06 | 0 | 5.93E-07 | 0 | 4 mo | HOM |
| **1013288951** | 2 | MBsta | 348 | 1.877871494 | 2.41E-05 | 1 | 1.28E-05 | 0.532517802 | 4 mo | HOM |
| **1013978665** | 1 | sup | 349 | 0.025784471 | 0 | 0 | 0 | 0 | 2 mo | HOM |
| **1013445609** | 1 | sup | 349 | 0.025784471 | 0 | 0 | 0 | 0 | 4 mo | HOM |
| **1013609015** | 1 | sup | 349 | 0.025784471 | 0 | 0 | 0 | 0 | 4 mo | HOM |
| **1013608106** | 2 | sup | 349 | 0.025784471 | 0 | 0 | 0 | 0 | 4 mo | HOM |
| **1011937543** | 1 | sup | 349 | 0.025784471 | 3.59E-05 | 1 | 0.001391578 | 38.78303295 | 8 mo | HOM |
| **1010419538** | 2 | sup | 349 | 0.025784471 | 0 | 0 | 0 | 0 | 4 mo | WILD |
| **1010419187** | 1 | sup | 349 | 0.025784471 | 3.3E-08 | 0 | 1.28E-06 | 0 | 8 mo | HOM |
| **1013978366** | 1 | sup | 349 | 0.025784471 | 0 | 0 | 0 | 0 | 2 mo | HOM |
| **1014416155** | 2 | sup | 349 | 0.025784471 | 0 | 0 | 0 | 0 | 2 mo | HOM |
| **1013978065** | 2 | sup | 349 | 0.025784471 | 0 | 0 | 0 | 0 | 2 mo | HOM |
| **1009934102** | 2 | sup | 349 | 0.025784471 | 0 | 0 | 0 | 0 | 4 mo | WILD |
| **1009322937** | 2 | sup | 349 | 0.025784471 | 0 | 0 | 0 | 0 | 8 mo | WILD |
| **1011080637** | 2 | sup | 349 | 0.025784471 | 0 | 0 | 0 | 0 | 8 mo | HOM |
| **1013609713** | 2 | sup | 349 | 0.025784471 | 0 | 0 | 0 | 0 | 2 mo | WILD |
| **1010418842** | 1 | sup | 349 | 0.025784471 | 1.74E-06 | 1 | 6.76E-05 | 38.78303295 | 8 mo | HOM |
| **1013288213** | 2 | sup | 349 | 0.025784471 | 0 | 0 | 0 | 0 | 4 mo | HOM |
| **1008976060** | 2 | sup | 349 | 0.025784471 | 0 | 0 | 0 | 0 | 8 mo | WILD |
| **1013446243** | 1 | sup | 349 | 0.025784471 | 0 | 0 | 0 | 0 | 4 mo | HOM |
| **1013288951** | 2 | sup | 349 | 0.025784471 | 0 | 0 | 0 | 0 | 4 mo | HOM |
| **1013978665** | 1 | SLC | 350 | 0.027542035 | 0 | 0 | 0 | 0 | 2 mo | HOM |
| **1013445609** | 1 | SLC | 350 | 0.027542035 | 0 | 0 | 0 | 0 | 4 mo | HOM |
| **1013609015** | 1 | SLC | 350 | 0.027542035 | 0 | 0 | 0 | 0 | 4 mo | HOM |
| **1013608106** | 2 | SLC | 350 | 0.027542035 | 0 | 0 | 0 | 0 | 4 mo | HOM |
| **1011937543** | 1 | SLC | 350 | 0.027542035 | 0 | 0 | 0 | 0 | 8 mo | HOM |
| **1010419538** | 2 | SLC | 350 | 0.027542035 | 0 | 0 | 0 | 0 | 4 mo | WILD |
| **1010419187** | 1 | SLC | 350 | 0.027542035 | 0 | 0 | 0 | 0 | 8 mo | HOM |
| **1013978366** | 1 | SLC | 350 | 0.027542035 | 0 | 0 | 0 | 0 | 2 mo | HOM |
| **1014416155** | 2 | SLC | 350 | 0.027542035 | 0 | 0 | 0 | 0 | 2 mo | HOM |
| **1013978065** | 2 | SLC | 350 | 0.027542035 | 0 | 0 | 0 | 0 | 2 mo | HOM |
| **1009934102** | 2 | SLC | 350 | 0.027542035 | 0 | 0 | 0 | 0 | 4 mo | WILD |
| **1009322937** | 2 | SLC | 350 | 0.027542035 | 0 | 0 | 0 | 0 | 8 mo | WILD |
| **1011080637** | 2 | SLC | 350 | 0.027542035 | 0 | 0 | 0 | 0 | 8 mo | HOM |
| **1013609713** | 2 | SLC | 350 | 0.027542035 | 0 | 0 | 0 | 0 | 2 mo | WILD |
| **1010418842** | 1 | SLC | 350 | 0.027542035 | 1.76E-05 | 3 | 0.000639668 | 108.9244126 | 8 mo | HOM |
| **1013288213** | 2 | SLC | 350 | 0.027542035 | 0 | 0 | 0 | 0 | 4 mo | HOM |
| **1008976060** | 2 | SLC | 350 | 0.027542035 | 4.58E-06 | 1 | 0.000166317 | 36.30813753 | 8 mo | WILD |
| **1013446243** | 1 | SLC | 350 | 0.027542035 | 0 | 0 | 0 | 0 | 4 mo | HOM |
| **1013288951** | 2 | SLC | 350 | 0.027542035 | 0 | 0 | 0 | 0 | 4 mo | HOM |
| **1013978665** | 1 | BST | 351 | 1.381224701 | 0 | 0 | 0 | 0 | 2 mo | HOM |
| **1013445609** | 1 | BST | 351 | 1.381224701 | 3.17E-05 | 3 | 2.29E-05 | 2.171985484 | 4 mo | HOM |
| **1013609015** | 1 | BST | 351 | 1.381224701 | 1.21E-05 | 2 | 8.77E-06 | 1.447990323 | 4 mo | HOM |
| **1013608106** | 2 | BST | 351 | 1.381224701 | 0 | 0 | 0 | 0 | 4 mo | HOM |
| **1011937543** | 1 | BST | 351 | 1.381224701 | 0.000494286 | 21 | 0.000357861 | 15.20389839 | 8 mo | HOM |
| **1010419538** | 2 | BST | 351 | 1.381224701 | 0 | 0 | 0 | 0 | 4 mo | WILD |
| **1010419187** | 1 | BST | 351 | 1.381224701 | 0.001028202 | 39 | 0.000744413 | 28.23581129 | 8 mo | HOM |
| **1013978366** | 1 | BST | 351 | 1.381224701 | 0 | 0 | 0 | 0 | 2 mo | HOM |
| **1014416155** | 2 | BST | 351 | 1.381224701 | 0 | 0 | 0 | 0 | 2 mo | HOM |
| **1013978065** | 2 | BST | 351 | 1.381224701 | 0 | 0 | 0 | 0 | 2 mo | HOM |
| **1009934102** | 2 | BST | 351 | 1.381224701 | 0 | 0 | 0 | 0 | 4 mo | WILD |
| **1009322937** | 2 | BST | 351 | 1.381224701 | 1.56E-06 | 0 | 1.13E-06 | 0 | 8 mo | WILD |
| **1011080637** | 2 | BST | 351 | 1.381224701 | 0.001052575 | 40 | 0.000762059 | 28.95980645 | 8 mo | HOM |

| | | | | | | | | | | |
|---|---|---|---|---|---|---|---|---|---|---|
| 1013609713 | 2 | BST | 351 | 1.381224701 | 0 | 0 | 0 | 0 | 2 mo | WILD |
| 1010418842 | 1 | BST | 351 | 1.381224701 | 0.001446254 | 42 | 0.001047081 | 30.40779677 | 8 mo | HOM |
| 1013288213 | 2 | BST | 351 | 1.381224701 | 4.92E-05 | 4 | 3.56E-05 | 2.895980645 | 4 mo | HOM |
| 1008976060 | 2 | BST | 351 | 1.381224701 | 5E-05 | 3 | 3.62E-05 | 2.171985484 | 8 mo | WILD |
| 1013446243 | 1 | BST | 351 | 1.381224701 | 5.41E-05 | 4 | 3.92E-05 | 2.895980645 | 4 mo | HOM |
| 1013288951 | 2 | BST | 351 | 1.381224701 | 3.56E-05 | 4 | 2.58E-05 | 2.895980645 | 4 mo | HOM |
| 1013978665 | 1 | SSp-n | 353 | 3.110331737 | 5.48E-06 | 1 | 1.76E-06 | 0.321509114 | 2 mo | HOM |
| 1013445609 | 1 | SSp-n | 353 | 3.110331737 | 0.000482283 | 26 | 0.000155058 | 8.359236955 | 4 mo | HOM |
| 1013609015 | 1 | SSp-n | 353 | 3.110331737 | 0.000922619 | 30 | 0.00029663 | 9.64527341 | 4 mo | HOM |
| 1013608106 | 2 | SSp-n | 353 | 3.110331737 | 0 | 0 | 0 | 0 | 4 mo | HOM |
| 1011937543 | 1 | SSp-n | 353 | 3.110331737 | 0.00605922 | 108 | 0.001948094 | 34.72298428 | 8 mo | HOM |
| 1010419538 | 2 | SSp-n | 353 | 3.110331737 | 4.94E-06 | 1 | 1.59E-06 | 0.321509114 | 4 mo | WILD |
| 1010419187 | 1 | SSp-n | 353 | 3.110331737 | 0.004658833 | 190 | 0.001497857 | 61.0867316 | 8 mo | HOM |
| 1013978366 | 1 | SSp-n | 353 | 3.110331737 | 4.2E-05 | 1 | 1.35E-05 | 0.321509114 | 2 mo | HOM |
| 1014416155 | 2 | SSp-n | 353 | 3.110331737 | 0 | 0 | 0 | 0 | 2 mo | HOM |
| 1013978065 | 2 | SSp-n | 353 | 3.110331737 | 0 | 0 | 0 | 0 | 2 mo | HOM |
| 1009934102 | 2 | SSp-n | 353 | 3.110331737 | 3.77E-06 | 1 | 1.21E-06 | 0.321509114 | 4 mo | WILD |
| 1009322937 | 2 | SSp-n | 353 | 3.110331737 | 0 | 0 | 0 | 0 | 8 mo | WILD |
| 1011080637 | 2 | SSp-n | 353 | 3.110331737 | 0.003758584 | 132 | 0.001208419 | 42.439203 | 8 mo | HOM |
| 1013609713 | 2 | SSp-n | 353 | 3.110331737 | 1.05E-05 | 1 | 3.38E-06 | 0.321509114 | 2 mo | WILD |
| 1010418842 | 1 | SSp-n | 353 | 3.110331737 | 0.006168636 | 162 | 0.001983273 | 52.08447641 | 8 mo | HOM |
| 1013288213 | 2 | SSp-n | 353 | 3.110331737 | 0.000235509 | 18 | 7.57E-05 | 5.787164046 | 4 mo | HOM |
| 1008976060 | 2 | SSp-n | 353 | 3.110331737 | 5.34E-05 | 6 | 1.72E-05 | 1.929054682 | 8 mo | WILD |
| 1013446243 | 1 | SSp-n | 353 | 3.110331737 | 0.000851984 | 46 | 0.00027392 | 14.78941923 | 4 mo | HOM |
| 1013288951 | 2 | SSp-n | 353 | 3.110331737 | 0.000806298 | 41 | 0.000259232 | 13.18187366 | 4 mo | HOM |
| 1013978665 | 1 | MY | 354 | 31.90121841 | 0.001270616 | 16 | 3.98E-05 | 0.501548242 | 2 mo | HOM |
| 1013445609 | 1 | MY | 354 | 31.90121841 | 0.000201738 | 7 | 6.32E-06 | 0.219427356 | 4 mo | HOM |
| 1013609015 | 1 | MY | 354 | 31.90121841 | 0.007186878 | 23 | 0.000225285 | 0.720975597 | 4 mo | HOM |
| 1013608106 | 2 | MY | 354 | 31.83135836 | 0 | 0 | 0 | 0 | 4 mo | HOM |
| 1011937543 | 1 | MY | 354 | 31.90121841 | 0.031783634 | 68 | 0.000996314 | 2.131580027 | 8 mo | HOM |
| 1010419538 | 2 | MY | 354 | 31.90121339 | 0.000103332 | 10 | 3.24E-06 | 0.3134677 | 4 mo | WILD |
| 1010419187 | 1 | MY | 354 | 31.90121841 | 0.013967522 | 46 | 0.000437837 | 1.441951195 | 8 mo | HOM |
| 1013978366 | 1 | MY | 354 | 31.90121841 | 0.001856157 | 13 | 5.82E-05 | 0.407507946 | 2 mo | HOM |
| 1014416155 | 2 | MY | 354 | 31.90121339 | 0 | 0 | 0 | 0 | 2 mo | HOM |
| 1013978065 | 2 | MY | 354 | 31.90121339 | 0.000433514 | 8 | 1.36E-05 | 0.25077416 | 2 mo | HOM |
| 1009934102 | 2 | MY | 354 | 31.90121339 | 2.74E-05 | 3 | 8.58E-07 | 0.09404031 | 4 mo | WILD |
| 1009322937 | 2 | MY | 354 | 31.90121339 | 0.000389992 | 17 | 1.22E-05 | 0.532895091 | 8 mo | WILD |
| 1011080637 | 2 | MY | 354 | 31.90121339 | 0.003896686 | 56 | 0.000122149 | 1.755419122 | 8 mo | HOM |
| 1013609713 | 2 | MY | 354 | 31.90121339 | 2.19E-06 | 1 | 6.87E-08 | 0.03134677 | 2 mo | WILD |
| 1010418842 | 1 | MY | 354 | 31.90121841 | 0.005083139 | 205 | 0.00015934 | 6.426086847 | 8 mo | HOM |
| 1013288213 | 2 | MY | 354 | 31.89964037 | 0.000594109 | 9 | 1.86E-05 | 0.282134842 | 4 mo | HOM |
| 1008976060 | 2 | MY | 354 | 31.90121339 | 0.008517735 | 187 | 0.000267003 | 5.861845997 | 8 mo | WILD |
| 1013446243 | 1 | MY | 354 | 31.90121841 | 0.000990599 | 11 | 3.11E-05 | 0.344814416 | 4 mo | HOM |
| 1013288951 | 2 | MY | 354 | 31.90121339 | 0.00388977 | 13 | 0.000121932 | 0.40750801 | 4 mo | HOM |
| 1013978665 | 1 | Alp6b | 355 | 0.021253542 | 0 | 0 | 0 | 0 | 2 mo | HOM |
| 1013445609 | 1 | Alp6b | 355 | 0.021253542 | 3.01E-06 | 0 | 0.000141756 | 0 | 4 mo | HOM |
| 1013609015 | 1 | Alp6b | 355 | 0.021253542 | 2.38E-06 | 0 | 0.000111811 | 0 | 4 mo | HOM |
| 1013608106 | 2 | Alp6b | 355 | 0.021253542 | 0 | 0 | 0 | 0 | 4 mo | HOM |
| 1011937543 | 1 | Alp6b | 355 | 0.021253542 | 4.65E-05 | 0 | 0.002188494 | 0 | 8 mo | HOM |
| 1010419538 | 2 | Alp6b | 355 | 0.021253542 | 0 | 0 | 0 | 0 | 4 mo | WILD |
| 1010419187 | 1 | Alp6b | 355 | 0.021253542 | 1.87E-05 | 0 | 0.000882199 | 0 | 8 mo | HOM |
| 1013978366 | 1 | Alp6b | 355 | 0.021253542 | 0 | 0 | 0 | 0 | 2 mo | HOM |
| 1014416155 | 2 | Alp6b | 355 | 0.021253542 | 0 | 0 | 0 | 0 | 2 mo | HOM |
| 1013978065 | 2 | Alp6b | 355 | 0.021253542 | 0 | 0 | 0 | 0 | 2 mo | HOM |
| 1009934102 | 2 | Alp6b | 355 | 0.021253542 | 0 | 0 | 0 | 0 | 4 mo | WILD |
| 1009322937 | 2 | Alp6b | 355 | 0.021253542 | 0 | 0 | 0 | 0 | 8 mo | WILD |
| 1011080637 | 2 | Alp6b | 355 | 0.021253542 | 1.27E-05 | 1 | 0.000596302 | 47.05098151 | 8 mo | HOM |
| 1013609713 | 2 | Alp6b | 355 | 0.021253542 | 0 | 0 | 0 | 0 | 2 mo | WILD |
| 1010418842 | 1 | Alp6b | 355 | 0.021253542 | 2.07E-06 | 0 | 9.73E-05 | 0 | 8 mo | HOM |

| | | | | | | | | | | |
|---|---|---|---|---|---|---|---|---|---|---|
| **1013288213** | 2 | Alp6b | 355 | 0.021253542 | 0 | 0 | 0 | 0 | 4 mo | HOM |
| **1008976060** | 2 | Alp6b | 355 | 0.021253542 | 8.83E-09 | 0 | 4.15E-07 | 0 | 8 mo | WILD |
| **1013446243** | 1 | Alp6b | 355 | 0.021253542 | 1.89E-06 | 0 | 8.91E-05 | 0 | 4 mo | HOM |
| **1013288951** | 2 | Alp6b | 355 | 0.021253542 | 0 | 0 | 0 | 0 | 4 mo | HOM |
| **1013978665** | 1 | PST | 356 | 0.012832483 | 0 | 0 | 0 | 0 | 2 mo | HOM |
| **1013445609** | 1 | PST | 356 | 0.012832483 | 0 | 0 | 0 | 0 | 4 mo | HOM |
| **1013609015** | 1 | PST | 356 | 0.012832483 | 0 | 0 | 0 | 0 | 4 mo | HOM |
| **1013608106** | 2 | PST | 356 | 0.012832483 | 0 | 0 | 0 | 0 | 4 mo | HOM |
| **1011937543** | 1 | PST | 356 | 0.012832483 | 3.66E-05 | 0 | 0.002851581 | 0 | 8 mo | HOM |
| **1010419538** | 2 | PST | 356 | 0.012832483 | 0 | 0 | 0 | 0 | 4 mo | WILD |
| **1010419187** | 1 | PST | 356 | 0.012832483 | 0 | 0 | 0 | 0 | 8 mo | HOM |
| **1013978366** | 1 | PST | 356 | 0.012832483 | 0 | 0 | 0 | 0 | 2 mo | HOM |
| **1014416155** | 2 | PST | 356 | 0.012832483 | 0 | 0 | 0 | 0 | 2 mo | HOM |
| **1013978065** | 2 | PST | 356 | 0.012832483 | 0 | 0 | 0 | 0 | 2 mo | HOM |
| **1009934102** | 2 | PST | 356 | 0.012832483 | 0 | 0 | 0 | 0 | 4 mo | WILD |
| **1009322937** | 2 | PST | 356 | 0.012832483 | 0 | 0 | 0 | 0 | 8 mo | WILD |
| **1011080637** | 2 | PST | 356 | 0.012832483 | 0 | 0 | 0 | 0 | 8 mo | HOM |
| **1013609713** | 2 | PST | 356 | 0.012832483 | 0 | 0 | 0 | 0 | 2 mo | WILD |
| **1010418842** | 1 | PST | 356 | 0.012832483 | 0 | 0 | 0 | 0 | 8 mo | HOM |
| **1013288213** | 2 | PST | 356 | 0.012832483 | 0 | 0 | 0 | 0 | 4 mo | HOM |
| **1008976060** | 2 | PST | 356 | 0.012832483 | 0 | 0 | 0 | 0 | 8 mo | WILD |
| **1013446243** | 1 | PST | 356 | 0.012832483 | 0 | 0 | 0 | 0 | 4 mo | HOM |
| **1013288951** | 2 | PST | 356 | 0.012832483 | 0 | 0 | 0 | 0 | 4 mo | HOM |
| **1013978665** | 1 | SLD | 358 | 0.048638983 | 0 | 0 | 0 | 0 | 2 mo | HOM |
| **1013445609** | 1 | SLD | 358 | 0.048638983 | 0 | 0 | 0 | 0 | 4 mo | HOM |
| **1013609015** | 1 | SLD | 358 | 0.048638983 | 8.2E-06 | 0 | 0.000168511 | 0 | 4 mo | HOM |
| **1013608106** | 2 | SLD | 358 | 0.048638983 | 0 | 0 | 0 | 0 | 4 mo | HOM |
| **1011937543** | 1 | SLD | 358 | 0.048638983 | 0 | 0 | 0 | 0 | 8 mo | HOM |
| **1010419538** | 2 | SLD | 358 | 0.048638983 | 0 | 0 | 0 | 0 | 4 mo | WILD |
| **1010419187** | 1 | SLD | 358 | 0.048638983 | 5.36E-06 | 0 | 0.000110274 | 0 | 8 mo | HOM |
| **1013978366** | 1 | SLD | 358 | 0.048638983 | 0 | 0 | 0 | 0 | 2 mo | HOM |
| **1014416155** | 2 | SLD | 358 | 0.048638983 | 0 | 0 | 0 | 0 | 2 mo | HOM |
| **1013978065** | 2 | SLD | 358 | 0.048638983 | 0 | 0 | 0 | 0 | 2 mo | HOM |
| **1009934102** | 2 | SLD | 358 | 0.048638983 | 0 | 0 | 0 | 0 | 4 mo | WILD |
| **1009322937** | 2 | SLD | 358 | 0.048638983 | 2.8E-06 | 1 | 5.76E-05 | 20.55964053 | 8 mo | WILD |
| **1011080637** | 2 | SLD | 358 | 0.048638983 | 4.29E-07 | 0 | 8.81E-06 | 0 | 8 mo | HOM |
| **1013609713** | 2 | SLD | 358 | 0.048638983 | 0 | 0 | 0 | 0 | 2 mo | WILD |
| **1010418842** | 1 | SLD | 358 | 0.048638983 | 1.06E-05 | 2 | 0.000217924 | 41.11928106 | 8 mo | HOM |
| **1013288213** | 2 | SLD | 358 | 0.048638983 | 0 | 0 | 0 | 0 | 4 mo | HOM |
| **1008976060** | 2 | SLD | 358 | 0.048638983 | 2.94E-06 | 0 | 6.05E-05 | 0 | 8 mo | WILD |
| **1013446243** | 1 | SLD | 358 | 0.048638983 | 0 | 0 | 0 | 0 | 4 mo | HOM |
| **1013288951** | 2 | SLD | 358 | 0.048638983 | 4.94E-06 | 1 | 0.000101658 | 20.55964053 | 4 mo | HOM |
| **1013978665** | 1 | SSp-tr | 361 | 1.444064312 | 0 | 0 | 0 | 0 | 2 mo | HOM |
| **1013445609** | 1 | SSp-tr | 361 | 1.444064312 | 0.000143296 | 11 | 9.92E-05 | 7.617389272 | 4 mo | HOM |
| **1013609015** | 1 | SSp-tr | 361 | 1.444064312 | 0.00051822 | 17 | 0.000358862 | 11.77232887 | 4 mo | HOM |
| **1013608106** | 2 | SSp-tr | 361 | 1.444064312 | 0 | 0 | 0 | 0 | 4 mo | HOM |
| **1011937543** | 1 | SSp-tr | 361 | 1.444064312 | 0.003475771 | 37 | 0.002406936 | 25.62212755 | 8 mo | HOM |
| **1010419538** | 2 | SSp-tr | 361 | 1.444064312 | 0 | 0 | 0 | 0 | 4 mo | WILD |
| **1010419187** | 1 | SSp-tr | 361 | 1.444064312 | 0.003020324 | 75 | 0.002091544 | 51.93674504 | 8 mo | HOM |
| **1013978366** | 1 | SSp-tr | 361 | 1.444064312 | 0 | 0 | 0 | 0 | 2 mo | HOM |
| **1014416155** | 2 | SSp-tr | 361 | 1.444064312 | 0 | 0 | 0 | 0 | 2 mo | HOM |
| **1013978065** | 2 | SSp-tr | 361 | 1.444064312 | 0 | 0 | 0 | 0 | 2 mo | HOM |
| **1009934102** | 2 | SSp-tr | 361 | 1.444064312 | 0 | 0 | 0 | 0 | 4 mo | WILD |
| **1009322937** | 2 | SSp-tr | 361 | 1.444064312 | 0 | 0 | 0 | 0 | 8 mo | WILD |
| **1011080637** | 2 | SSp-tr | 361 | 1.444064312 | 0.001776293 | 49 | 0.001230065 | 33.93200676 | 8 mo | HOM |
| **1013609713** | 2 | SSp-tr | 361 | 1.444064312 | 0 | 0 | 0 | 0 | 2 mo | WILD |
| **1010418842** | 1 | SSp-tr | 361 | 1.444064312 | 0.001798824 | 72 | 0.001245667 | 49.85927523 | 8 mo | HOM |
| **1013288213** | 2 | SSp-tr | 361 | 1.444064312 | 0.00023712 | 14 | 0.000164203 | 9.694859073 | 4 mo | HOM |
| **1008976060** | 2 | SSp-tr | 361 | 1.444064312 | 2.93E-05 | 4 | 2.03E-05 | 2.769959735 | 8 mo | WILD |

| | | | | | | | | | | |
|---|---|---|---|---|---|---|---|---|---|---|
| **1013446243** | 1 | SSp-tr | 361 | 1.444064312 | 0.00062666 | 30 | 0.000433956 | 20.77469801 | 4 mo | HOM |
| **1013288951** | 2 | SSp-tr | 361 | 1.444064312 | 0.000444441 | 16 | 0.000307771 | 11.07983894 | 4 mo | HOM |
| **1013978665** | 1 | MD | 362 | 1.425203024 | 0 | 0 | 0 | 0 | 2 mo | HOM |
| **1013445609** | 1 | MD | 362 | 1.425203024 | 7.7E-08 | 0 | 5.4E-08 | 0 | 4 mo | HOM |
| **1013609015** | 1 | MD | 362 | 1.425203024 | 3.41E-05 | 0 | 2.39E-05 | 0 | 4 mo | HOM |
| **1013608106** | 2 | MD | 362 | 1.425203024 | 0 | 0 | 0 | 0 | 4 mo | HOM |
| **1011937543** | 1 | MD | 362 | 1.425203024 | 0.000708973 | 18 | 0.000497454 | 12.62977955 | 8 mo | HOM |
| **1010419538** | 2 | MD | 362 | 1.425203024 | 2.31E-05 | 1 | 1.62E-05 | 0.701654419 | 4 mo | WILD |
| **1010419187** | 1 | MD | 362 | 1.425203024 | 0.0005759 | 16 | 0.000404083 | 11.22647071 | 8 mo | HOM |
| **1013978366** | 1 | MD | 362 | 1.425203024 | 0 | 0 | 0 | 0 | 2 mo | HOM |
| **1014416155** | 2 | MD | 362 | 1.425203024 | 0 | 0 | 0 | 0 | 2 mo | HOM |
| **1013978065** | 2 | MD | 362 | 1.425203024 | 0 | 0 | 0 | 0 | 2 mo | HOM |
| **1009934102** | 2 | MD | 362 | 1.425203024 | 0 | 0 | 0 | 0 | 4 mo | WILD |
| **1009322937** | 2 | MD | 362 | 1.425203024 | 2.32E-05 | 2 | 1.63E-05 | 1.403308838 | 8 mo | WILD |
| **1011080637** | 2 | MD | 362 | 1.425203024 | 0.000321442 | 12 | 0.000225541 | 8.41985303 | 8 mo | HOM |
| **1013609713** | 2 | MD | 362 | 1.425203024 | 0 | 0 | 0 | 0 | 2 mo | WILD |
| **1010418842** | 1 | MD | 362 | 1.425203024 | 0.000945142 | 27 | 0.000663163 | 18.94466932 | 8 mo | HOM |
| **1013288213** | 2 | MD | 362 | 1.425203024 | 0 | 0 | 0 | 0 | 4 mo | HOM |
| **1008976060** | 2 | MD | 362 | 1.425203024 | 2.79E-05 | 4 | 1.96E-05 | 2.806617677 | 8 mo | WILD |
| **1013446243** | 1 | MD | 362 | 1.425203024 | 4.73E-06 | 0 | 3.32E-06 | 0 | 4 mo | HOM |
| **1013288951** | 2 | MD | 362 | 1.425203024 | 6.03E-06 | 1 | 4.23E-06 | 0.701654419 | 4 mo | HOM |
| **1013978665** | 1 | PL5 | 363 | 1.009041477 | 0 | 0 | 0 | 0 | 2 mo | HOM |
| **1013445609** | 1 | PL5 | 363 | 1.009041477 | 7.41E-05 | 3 | 7.35E-05 | 2.973118617 | 4 mo | HOM |
| **1013609015** | 1 | PL5 | 363 | 1.009041477 | 0.000272059 | 7 | 0.000269621 | 6.937276773 | 4 mo | HOM |
| **1013608106** | 2 | PL5 | 363 | 1.009041477 | 0 | 0 | 0 | 0 | 4 mo | HOM |
| **1011937543** | 1 | PL5 | 363 | 1.009041477 | 0.000797241 | 23 | 0.000790097 | 22.7939094 | 8 mo | HOM |
| **1010419538** | 2 | PL5 | 363 | 1.009041477 | 0 | 0 | 0 | 0 | 4 mo | WILD |
| **1010419187** | 1 | PL5 | 363 | 1.009041477 | 0.001918787 | 49 | 0.001901594 | 48.56093741 | 8 mo | HOM |
| **1013978366** | 1 | PL5 | 363 | 1.009041477 | 0 | 0 | 0 | 0 | 2 mo | HOM |
| **1014416155** | 2 | PL5 | 363 | 1.009041477 | 0 | 0 | 0 | 0 | 2 mo | HOM |
| **1013978065** | 2 | PL5 | 363 | 1.009041477 | 0 | 0 | 0 | 0 | 2 mo | HOM |
| **1009934102** | 2 | PL5 | 363 | 1.009041477 | 4.9E-09 | 0 | 4.86E-09 | 0 | 4 mo | WILD |
| **1009322937** | 2 | PL5 | 363 | 1.009041477 | 0 | 0 | 0 | 0 | 8 mo | WILD |
| **1011080637** | 2 | PL5 | 363 | 1.009041477 | 0.000743905 | 34 | 0.000737239 | 33.69534433 | 8 mo | HOM |
| **1013609713** | 2 | PL5 | 363 | 1.009041477 | 0 | 0 | 0 | 0 | 2 mo | WILD |
| **1010418842** | 1 | PL5 | 363 | 1.009041477 | 0.001388892 | 48 | 0.001376447 | 47.56989787 | 8 mo | HOM |
| **1013288213** | 2 | PL5 | 363 | 1.009041477 | 0.000104563 | 8 | 0.000103626 | 7.928316312 | 4 mo | HOM |
| **1008976060** | 2 | PL5 | 363 | 1.009041477 | 0 | 0 | 0 | 0 | 8 mo | WILD |
| **1013446243** | 1 | PL5 | 363 | 1.009041477 | 0.000146506 | 6 | 0.000145193 | 5.946237234 | 4 mo | HOM |
| **1013288951** | 2 | PL5 | 363 | 1.009041477 | 0.000222091 | 5 | 0.000220101 | 4.955197695 | 4 mo | HOM |
| **1013978665** | 1 | PSTN | 364 | 0.171538651 | 0 | 0 | 0 | 0 | 2 mo | HOM |
| **1013445609** | 1 | PSTN | 364 | 0.171538651 | 0 | 0 | 0 | 0 | 4 mo | HOM |
| **1013609015** | 1 | PSTN | 364 | 0.171538651 | 0 | 0 | 0 | 0 | 4 mo | HOM |
| **1013608106** | 2 | PSTN | 364 | 0.171538651 | 0 | 0 | 0 | 0 | 4 mo | HOM |
| **1011937543** | 1 | PSTN | 364 | 0.171538651 | 0.00022114 | 5 | 0.001289158 | 29.14794984 | 8 mo | HOM |
| **1010419538** | 2 | PSTN | 364 | 0.171538651 | 0 | 0 | 0 | 0 | 4 mo | WILD |
| **1010419187** | 1 | PSTN | 364 | 0.171538651 | 0 | 0 | 0 | 0 | 8 mo | HOM |
| **1013978366** | 1 | PSTN | 364 | 0.171538651 | 0 | 0 | 0 | 0 | 2 mo | HOM |
| **1014416155** | 2 | PSTN | 364 | 0.171538651 | 0 | 0 | 0 | 0 | 2 mo | HOM |
| **1013978065** | 2 | PSTN | 364 | 0.171538651 | 0 | 0 | 0 | 0 | 2 mo | HOM |
| **1009934102** | 2 | PSTN | 364 | 0.171538651 | 0 | 0 | 0 | 0 | 4 mo | WILD |
| **1009322937** | 2 | PSTN | 364 | 0.171538651 | 0 | 0 | 0 | 0 | 8 mo | WILD |
| **1011080637** | 2 | PSTN | 364 | 0.171538651 | 2.09E-05 | 1 | 0.000121644 | 5.829589967 | 8 mo | HOM |
| **1013609713** | 2 | PSTN | 364 | 0.171538651 | 0 | 0 | 0 | 0 | 2 mo | WILD |
| **1010418842** | 1 | PSTN | 364 | 0.171538651 | 0.00015356 | 2 | 0.000895192 | 11.65917993 | 8 mo | HOM |
| **1013288213** | 2 | PSTN | 364 | 0.171538651 | 0 | 0 | 0 | 0 | 4 mo | HOM |
| **1008976060** | 2 | PSTN | 364 | 0.171538651 | 0 | 0 | 0 | 0 | 8 mo | WILD |
| **1013446243** | 1 | PSTN | 364 | 0.171538651 | 0 | 0 | 0 | 0 | 4 mo | HOM |
| **1013288951** | 2 | PSTN | 364 | 0.171538651 | 0 | 0 | 0 | 0 | 4 mo | HOM |

| | | | | | | | | | | |
|---|---|---|---|---|---|---|---|---|---|---|
| **1013978665** | 1 | SMT | 366 | 0.315595006 | 0 | 0 | 0 | 0 | 2 mo | HOM |
| **1013445609** | 1 | SMT | 366 | 0.315595006 | 0 | 0 | 0 | 0 | 4 mo | HOM |
| **1013609015** | 1 | SMT | 366 | 0.315595006 | 0 | 0 | 0 | 0 | 4 mo | HOM |
| **1013608106** | 2 | SMT | 366 | 0.315595006 | 0 | 0 | 0 | 0 | 4 mo | HOM |
| **1011937543** | 1 | SMT | 366 | 0.315595006 | 3.75E-05 | 3 | 0.000118885 | 9.505853853 | 8 mo | HOM |
| **1010419538** | 2 | SMT | 366 | 0.315595006 | 0 | 0 | 0 | 0 | 4 mo | WILD |
| **1010419187** | 1 | SMT | 366 | 0.315595006 | 0.000148436 | 3 | 0.000470338 | 9.505853853 | 8 mo | HOM |
| **1013978366** | 1 | SMT | 366 | 0.315595006 | 0 | 0 | 0 | 0 | 2 mo | HOM |
| **1014416155** | 2 | SMT | 366 | 0.315595006 | 0 | 0 | 0 | 0 | 2 mo | HOM |
| **1013978065** | 2 | SMT | 366 | 0.315595006 | 0 | 0 | 0 | 0 | 2 mo | HOM |
| **1009934102** | 2 | SMT | 366 | 0.315595006 | 0 | 0 | 0 | 0 | 4 mo | WILD |
| **1009322937** | 2 | SMT | 366 | 0.315595006 | 4.57E-06 | 1 | 1.45E-05 | 3.168617951 | 8 mo | WILD |
| **1011080637** | 2 | SMT | 366 | 0.315595006 | 0.000258064 | 6 | 0.000817707 | 19.01170771 | 8 mo | HOM |
| **1013609713** | 2 | SMT | 366 | 0.315595006 | 0 | 0 | 0 | 0 | 2 mo | WILD |
| **1010418842** | 1 | SMT | 366 | 0.315595006 | 0.000139818 | 5 | 0.000443029 | 15.84308976 | 8 mo | HOM |
| **1013288213** | 2 | SMT | 366 | 0.315595006 | 0 | 0 | 0 | 0 | 4 mo | HOM |
| **1008976060** | 2 | SMT | 366 | 0.315595006 | 1.96E-06 | 0 | 6.2E-06 | 0 | 8 mo | WILD |
| **1013446243** | 1 | SMT | 366 | 0.315595006 | 0 | 0 | 0 | 0 | 4 mo | HOM |
| **1013288951** | 2 | SMT | 366 | 0.315595006 | 0 | 0 | 0 | 0 | 4 mo | HOM |
| **1013978665** | 1 | PERI6b | 368 | 0.013545398 | 0 | 0 | 0 | 0 | 2 mo | HOM |
| **1013445609** | 1 | PERI6b | 368 | 0.013545398 | 0 | 0 | 0 | 0 | 4 mo | HOM |
| **1013609015** | 1 | PERI6b | 368 | 0.013545398 | 0 | 0 | 0 | 0 | 4 mo | HOM |
| **1013608106** | 2 | PERI6b | 368 | 0.013545398 | 0 | 0 | 0 | 0 | 4 mo | HOM |
| **1011937543** | 1 | PERI6b | 368 | 0.013545398 | 8.27E-06 | 0 | 0.000610663 | 0 | 8 mo | HOM |
| **1010419538** | 2 | PERI6b | 368 | 0.013545398 | 0 | 0 | 0 | 0 | 4 mo | WILD |
| **1010419187** | 1 | PERI6b | 368 | 0.013545398 | 1.02E-22 | 0 | 7.51E-21 | 0 | 8 mo | HOM |
| **1013978366** | 1 | PERI6b | 368 | 0.013545398 | 0 | 0 | 0 | 0 | 2 mo | HOM |
| **1014416155** | 2 | PERI6b | 368 | 0.013545398 | 0 | 0 | 0 | 0 | 2 mo | HOM |
| **1013978065** | 2 | PERI6b | 368 | 0.013545398 | 0 | 0 | 0 | 0 | 2 mo | HOM |
| **1009934102** | 2 | PERI6b | 368 | 0.013545398 | 0 | 0 | 0 | 0 | 4 mo | WILD |
| **1009322937** | 2 | PERI6b | 368 | 0.013545398 | 0 | 0 | 0 | 0 | 8 mo | WILD |
| **1011080637** | 2 | PERI6b | 368 | 0.013545398 | 4.18E-06 | 1 | 0.000308955 | 73.82580991 | 8 mo | HOM |
| **1013609713** | 2 | PERI6b | 368 | 0.013545398 | 0 | 0 | 0 | 0 | 2 mo | WILD |
| **1010418842** | 1 | PERI6b | 368 | 0.013545398 | 7.53E-07 | 0 | 5.56E-05 | 0 | 8 mo | HOM |
| **1013288213** | 2 | PERI6b | 368 | 0.013545398 | 0 | 0 | 0 | 0 | 4 mo | HOM |
| **1008976060** | 2 | PERI6b | 368 | 0.013545398 | 0 | 0 | 0 | 0 | 8 mo | WILD |
| **1013446243** | 1 | PERI6b | 368 | 0.013545398 | 0 | 0 | 0 | 0 | 4 mo | HOM |
| **1013288951** | 2 | PERI6b | 368 | 0.013545398 | 0 | 0 | 0 | 0 | 4 mo | HOM |
| **1013978665** | 1 | SSp-ul | 369 | 3.883394016 | 0 | 0 | 0 | 0 | 2 mo | HOM |
| **1013445609** | 1 | SSp-ul | 369 | 3.883394016 | 0.000530268 | 25 | 0.000136548 | 6.437667643 | 4 mo | HOM |
| **1013609015** | 1 | SSp-ul | 369 | 3.883394016 | 0.001107596 | 46 | 0.000285213 | 11.84530846 | 4 mo | HOM |
| **1013608106** | 2 | SSp-ul | 369 | 3.883394016 | 0 | 0 | 0 | 0 | 4 mo | HOM |
| **1011937543** | 1 | SSp-ul | 369 | 3.883394016 | 0.007160972 | 126 | 0.001843998 | 32.44584492 | 8 mo | HOM |
| **1010419538** | 2 | SSp-ul | 369 | 3.883394016 | 6.36E-06 | 1 | 1.64E-06 | 0.257506706 | 4 mo | WILD |
| **1010419187** | 1 | SSp-ul | 369 | 3.883394016 | 0.00695638 | 212 | 0.001791314 | 54.59142161 | 8 mo | HOM |
| **1013978366** | 1 | SSp-ul | 369 | 3.883394016 | 0 | 0 | 0 | 0 | 2 mo | HOM |
| **1014416155** | 2 | SSp-ul | 369 | 3.883394016 | 0 | 0 | 0 | 0 | 2 mo | HOM |
| **1013978065** | 2 | SSp-ul | 369 | 3.883394016 | 0 | 0 | 0 | 0 | 2 mo | HOM |
| **1009934102** | 2 | SSp-ul | 369 | 3.883394016 | 7.57E-06 | 1 | 1.95E-06 | 0.257506706 | 4 mo | WILD |
| **1009322937** | 2 | SSp-ul | 369 | 3.883394016 | 0 | 0 | 0 | 0 | 8 mo | WILD |
| **1011080637** | 2 | SSp-ul | 369 | 3.883394016 | 0.003503561 | 120 | 0.00090219 | 30.90080468 | 8 mo | HOM |
| **1013609713** | 2 | SSp-ul | 369 | 3.883394016 | 4.5E-06 | 0 | 1.16E-06 | 0 | 2 mo | WILD |
| **1010418842** | 1 | SSp-ul | 369 | 3.883394016 | 0.004522066 | 162 | 0.001164462 | 41.71608632 | 8 mo | HOM |
| **1013288213** | 2 | SSp-ul | 369 | 3.883394016 | 0.000257624 | 22 | 6.63E-05 | 5.665147526 | 4 mo | HOM |
| **1008976060** | 2 | SSp-ul | 369 | 3.883394016 | 0.000548313 | 14 | 0.000141194 | 3.60509388 | 8 mo | WILD |
| **1013446243** | 1 | SSp-ul | 369 | 3.883394016 | 0.000819232 | 51 | 0.000210958 | 13.13284199 | 4 mo | HOM |
| **1013288951** | 2 | SSp-ul | 369 | 3.883394016 | 0.000956879 | 44 | 0.000246403 | 11.33029505 | 4 mo | HOM |
| **1013978665** | 1 | MY-mot | 370 | 18.01775335 | 0.001004367 | 8 | 5.57E-05 | 0.444006522 | 2 mo | HOM |
| **1013445609** | 1 | MY-mot | 370 | 18.01775335 | 3.39E-05 | 2 | 1.88E-06 | 0.11100163 | 4 mo | HOM |

| | | | | | | | | | | |
|---|---|---|---|---|---|---|---|---|---|---|
| **1013609015** | 1 | MY-mot | 370 | 18.01775335 | 0.002027611 | 10 | 0.000112534 | 0.555008152 | 4 mo | HOM |
| **1013608106** | 2 | MY-mot | 370 | 17.97069681 | 0 | 0 | 0 | 0 | 4 mo | HOM |
| **1011937543** | 1 | MY-mot | 370 | 18.01775335 | 0.012561385 | 29 | 0.000697167 | 1.609523642 | 8 mo | HOM |
| **1010419538** | 2 | MY-mot | 370 | 18.01775335 | 4.9E-05 | 3 | 2.72E-06 | 0.166502446 | 4 mo | WILD |
| **1010419187** | 1 | MY-mot | 370 | 18.01775335 | 0.002448315 | 22 | 0.000135883 | 1.221017935 | 8 mo | HOM |
| **1013978366** | 1 | MY-mot | 370 | 18.01775335 | 0.000323025 | 4 | 1.79E-05 | 0.222003261 | 2 mo | HOM |
| **1014416155** | 2 | MY-mot | 370 | 18.01775335 | 0 | 0 | 0 | 0 | 2 mo | HOM |
| **1013978065** | 2 | MY-mot | 370 | 18.01775335 | 0.000183994 | 2 | 1.02E-05 | 0.11100163 | 2 mo | HOM |
| **1009934102** | 2 | MY-mot | 370 | 18.01775335 | 2.63E-10 | 0 | 1.46E-11 | 0 | 4 mo | WILD |
| **1009322937** | 2 | MY-mot | 370 | 18.01775335 | 8.14E-05 | 7 | 4.52E-06 | 0.388505707 | 8 mo | WILD |
| **1011080637** | 2 | MY-mot | 370 | 18.01775335 | 0.002189098 | 22 | 0.000121497 | 1.221017935 | 8 mo | HOM |
| **1013609713** | 2 | MY-mot | 370 | 18.01775335 | 0 | 0 | 0 | 0 | 2 mo | WILD |
| **1010418842** | 1 | MY-mot | 370 | 18.01775335 | 0.003853433 | 148 | 0.000213869 | 8.214120657 | 8 mo | HOM |
| **1013288213** | 2 | MY-mot | 370 | 18.01775202 | 0.00053075 | 8 | 2.95E-05 | 0.444006555 | 4 mo | HOM |
| **1008976060** | 2 | MY-mot | 370 | 18.01775335 | 0.005347038 | 142 | 0.000296765 | 7.881115765 | 8 mo | WILD |
| **1013446243** | 1 | MY-mot | 370 | 18.01775335 | 4.61E-05 | 3 | 2.56E-06 | 0.166502446 | 4 mo | HOM |
| **1013288951** | 2 | MY-mot | 370 | 18.01775335 | 0.001167648 | 5 | 6.48E-05 | 0.277504076 | 4 mo | HOM |
| **1013978665** | 1 | ICB | 372 | 0.05508407 | 0 | 0 | 0 | 0 | 2 mo | HOM |
| **1013445609** | 1 | ICB | 372 | 0.05508407 | 0 | 0 | 0 | 0 | 4 mo | HOM |
| **1013609015** | 1 | ICB | 372 | 0.05508407 | 0 | 0 | 0 | 0 | 4 mo | HOM |
| **1013608106** | 2 | ICB | 372 | 0.05508407 | 0 | 0 | 0 | 0 | 4 mo | HOM |
| **1011937543** | 1 | ICB | 372 | 0.05508407 | 0 | 0 | 0 | 0 | 8 mo | HOM |
| **1010419538** | 2 | ICB | 372 | 0.05508407 | 0 | 0 | 0 | 0 | 4 mo | WILD |
| **1010419187** | 1 | ICB | 372 | 0.05508407 | 0 | 0 | 0 | 0 | 8 mo | HOM |
| **1013978366** | 1 | ICB | 372 | 0.05508407 | 0 | 0 | 0 | 0 | 2 mo | HOM |
| **1014416155** | 2 | ICB | 372 | 0.05508407 | 0 | 0 | 0 | 0 | 2 mo | HOM |
| **1013978065** | 2 | ICB | 372 | 0.05508407 | 0 | 0 | 0 | 0 | 2 mo | HOM |
| **1009934102** | 2 | ICB | 372 | 0.05508407 | 0 | 0 | 0 | 0 | 4 mo | WILD |
| **1009322937** | 2 | ICB | 372 | 0.05508407 | 0 | 0 | 0 | 0 | 8 mo | WILD |
| **1011080637** | 2 | ICB | 372 | 0.05508407 | 0 | 0 | 0 | 0 | 8 mo | HOM |
| **1013609713** | 2 | ICB | 372 | 0.05508407 | 0 | 0 | 0 | 0 | 2 mo | WILD |
| **1010418842** | 1 | ICB | 372 | 0.05508407 | 0 | 0 | 0 | 0 | 8 mo | HOM |
| **1013288213** | 2 | ICB | 372 | 0.05508407 | 0 | 0 | 0 | 0 | 4 mo | HOM |
| **1008976060** | 2 | ICB | 372 | 0.05508407 | 0 | 0 | 0 | 0 | 8 mo | WILD |
| **1013446243** | 1 | ICB | 372 | 0.05508407 | 0 | 0 | 0 | 0 | 4 mo | HOM |
| **1013288951** | 2 | ICB | 372 | 0.05508407 | 0 | 0 | 0 | 0 | 4 mo | HOM |
| **1013978665** | 1 | SNc | 374 | 0.202740048 | 0 | 0 | 0 | 0 | 2 mo | HOM |
| **1013445609** | 1 | SNc | 374 | 0.202740048 | 0 | 0 | 0 | 0 | 4 mo | HOM |
| **1013609015** | 1 | SNc | 374 | 0.202740048 | 1.45E-07 | 0 | 7.14E-07 | 0 | 4 mo | HOM |
| **1013608106** | 2 | SNc | 374 | 0.202740048 | 0 | 0 | 0 | 0 | 4 mo | HOM |
| **1011937543** | 1 | SNc | 374 | 0.202740048 | 0.000431761 | 5 | 0.002129626 | 24.662123 | 8 mo | HOM |
| **1010419538** | 2 | SNc | 374 | 0.202740048 | 0 | 0 | 0 | 0 | 4 mo | WILD |
| **1010419187** | 1 | SNc | 374 | 0.202740048 | 4.71E-05 | 4 | 0.000232489 | 19.7296984 | 8 mo | HOM |
| **1013978366** | 1 | SNc | 374 | 0.202740048 | 0 | 0 | 0 | 0 | 2 mo | HOM |
| **1014416155** | 2 | SNc | 374 | 0.202740048 | 0 | 0 | 0 | 0 | 2 mo | HOM |
| **1013978065** | 2 | SNc | 374 | 0.202740048 | 0 | 0 | 0 | 0 | 2 mo | HOM |
| **1009934102** | 2 | SNc | 374 | 0.202740048 | 0 | 0 | 0 | 0 | 4 mo | WILD |
| **1009322937** | 2 | SNc | 374 | 0.202740048 | 0 | 0 | 0 | 0 | 8 mo | WILD |
| **1011080637** | 2 | SNc | 374 | 0.202740048 | 3.01E-05 | 1 | 0.000148609 | 4.932424599 | 8 mo | HOM |
| **1013609713** | 2 | SNc | 374 | 0.202740048 | 0 | 0 | 0 | 0 | 2 mo | WILD |
| **1010418842** | 1 | SNc | 374 | 0.202740048 | 8.87E-05 | 2 | 0.000437363 | 9.864849198 | 8 mo | HOM |
| **1013288213** | 2 | SNc | 374 | 0.202740048 | 0 | 0 | 0 | 0 | 4 mo | HOM |
| **1008976060** | 2 | SNc | 374 | 0.202740048 | 4.28E-08 | 0 | 2.11E-07 | 0 | 8 mo | WILD |
| **1013446243** | 1 | SNc | 374 | 0.202740048 | 0 | 0 | 0 | 0 | 4 mo | HOM |
| **1013288951** | 2 | SNc | 374 | 0.202740048 | 0 | 0 | 0 | 0 | 4 mo | HOM |
| **1013978665** | 1 | CA | 375 | 17.4427683 | 3.57E-05 | 5 | 2.04E-06 | 0.286651747 | 2 mo | HOM |
| **1013445609** | 1 | CA | 375 | 17.4427683 | 0.000524431 | 40 | 3.01E-05 | 2.293213974 | 4 mo | HOM |
| **1013609015** | 1 | CA | 375 | 17.4427683 | 0.002811396 | 124 | 0.000161178 | 7.108963319 | 4 mo | HOM |
| **1013608106** | 2 | CA | 375 | 17.4427683 | 0 | 0 | 0 | 0 | 4 mo | HOM |

| | | | | | | | | | | |
|---|---|---|---|---|---|---|---|---|---|---|
| **1011937543** | 1 | CA | 375 | 17.4427683 | 0.01794156 | 264 | 0.001028596 | 15.13521223 | 8 mo | HOM |
| **1010419538** | 2 | CA | 375 | 17.4427683 | 2.97E-07 | 0 | 1.7E-08 | 0 | 4 mo | WILD |
| **1010419187** | 1 | CA | 375 | 17.4427683 | 0.029106717 | 789 | 0.001668698 | 45.23364563 | 8 mo | HOM |
| **1013978366** | 1 | CA | 375 | 17.4427683 | 5.99E-06 | 1 | 3.43E-07 | 0.057330349 | 2 mo | HOM |
| **1014416155** | 2 | CA | 375 | 17.4427683 | 0 | 0 | 0 | 0 | 2 mo | HOM |
| **1013978065** | 2 | CA | 375 | 17.4427683 | 0 | 0 | 0 | 0 | 2 mo | HOM |
| **1009934102** | 2 | CA | 375 | 17.4427683 | 1.7E-05 | 2 | 9.73E-07 | 0.114660699 | 4 mo | WILD |
| **1009322937** | 2 | CA | 375 | 17.4427683 | 7.49E-05 | 9 | 4.29E-06 | 0.515973144 | 8 mo | WILD |
| **1011080637** | 2 | CA | 375 | 17.4427683 | 0.020460884 | 652 | 0.00117303 | 37.37938777 | 8 mo | HOM |
| **1013609713** | 2 | CA | 375 | 17.4427683 | 0 | 0 | 0 | 0 | 2 mo | WILD |
| **1010418842** | 1 | CA | 375 | 17.4427683 | 0.028799876 | 790 | 0.001651107 | 45.29097598 | 8 mo | HOM |
| **1013288213** | 2 | CA | 375 | 17.4427683 | 0.000845446 | 62 | 4.85E-05 | 3.554481659 | 4 mo | HOM |
| **1008976060** | 2 | CA | 375 | 17.4427683 | 0.000378893 | 52 | 2.17E-05 | 2.981178166 | 8 mo | WILD |
| **1013446243** | 1 | CA | 375 | 17.4427683 | 0.001939647 | 108 | 0.000111201 | 6.191677729 | 4 mo | HOM |
| **1013288951** | 2 | CA | 375 | 17.4427683 | 0.001866981 | 109 | 0.000107035 | 6.249008079 | 4 mo | HOM |
| **1013978665** | 1 | VISpl6a | 377 | 0.110516354 | 0 | 0 | 0 | 0 | 2 mo | HOM |
| **1013445609** | 1 | VISpl6a | 377 | 0.110516354 | 0 | 0 | 0 | 0 | 4 mo | HOM |
| **1013609015** | 1 | VISpl6a | 377 | 0.110516354 | 7.01E-07 | 0 | 6.34E-06 | 0 | 4 mo | HOM |
| **1013608106** | 2 | VISpl6a | 377 | 0.110516354 | 0 | 0 | 0 | 0 | 4 mo | HOM |
| **1011937543** | 1 | VISpl6a | 377 | 0.110516354 | 0.000207849 | 2 | 0.00188071 | 18.09686914 | 8 mo | HOM |
| **1010419538** | 2 | VISpl6a | 377 | 0.110516354 | 0 | 0 | 0 | 0 | 4 mo | WILD |
| **1010419187** | 1 | VISpl6a | 377 | 0.110516354 | 0.000282344 | 10 | 0.002554768 | 90.48434571 | 8 mo | HOM |
| **1013978366** | 1 | VISpl6a | 377 | 0.110516354 | 0 | 0 | 0 | 0 | 2 mo | HOM |
| **1014416155** | 2 | VISpl6a | 377 | 0.110516354 | 0 | 0 | 0 | 0 | 2 mo | HOM |
| **1013978065** | 2 | VISpl6a | 377 | 0.110516354 | 0 | 0 | 0 | 0 | 2 mo | HOM |
| **1009934102** | 2 | VISpl6a | 377 | 0.110516354 | 0 | 0 | 0 | 0 | 4 mo | WILD |
| **1009322937** | 2 | VISpl6a | 377 | 0.110516354 | 0 | 0 | 0 | 0 | 8 mo | WILD |
| **1011080637** | 2 | VISpl6a | 377 | 0.110516354 | 5.24E-05 | 4 | 0.000474374 | 36.19373828 | 8 mo | HOM |
| **1013609713** | 2 | VISpl6a | 377 | 0.110516354 | 0 | 0 | 0 | 0 | 2 mo | WILD |
| **1010418842** | 1 | VISpl6a | 377 | 0.110516354 | 0.00019031 | 9 | 0.001722012 | 81.43591114 | 8 mo | HOM |
| **1013288213** | 2 | VISpl6a | 377 | 0.110516354 | 1.65E-05 | 1 | 0.000149513 | 9.048434571 | 4 mo | HOM |
| **1008976060** | 2 | VISpl6a | 377 | 0.110516354 | 0 | 0 | 0 | 0 | 8 mo | WILD |
| **1013446243** | 1 | VISpl6a | 377 | 0.110516354 | 4.47E-05 | 3 | 0.000404241 | 27.14530371 | 4 mo | HOM |
| **1013288951** | 2 | VISpl6a | 377 | 0.110516354 | 6.14E-06 | 0 | 5.56E-05 | 0 | 4 mo | HOM |
| **1013978665** | 1 | SSs | 378 | 9.290735866 | 3.02E-05 | 2 | 3.25E-06 | 0.215268201 | 2 mo | HOM |
| **1013445609** | 1 | SSs | 378 | 9.290735866 | 0.001105179 | 64 | 0.000118955 | 6.888582447 | 4 mo | HOM |
| **1013609015** | 1 | SSs | 378 | 9.290735866 | 0.002740209 | 94 | 0.00029494 | 10.11760547 | 4 mo | HOM |
| **1013608106** | 2 | SSs | 378 | 9.290735866 | 0 | 0 | 0 | 0 | 4 mo | HOM |
| **1011937543** | 1 | SSs | 378 | 9.290735866 | 0.014726611 | 301 | 0.001585086 | 32.39786432 | 8 mo | HOM |
| **1010419538** | 2 | SSs | 378 | 9.290735866 | 4.22E-05 | 2 | 4.55E-06 | 0.215268201 | 4 mo | WILD |
| **1010419187** | 1 | SSs | 378 | 9.290735866 | 0.011466391 | 430 | 0.001234175 | 46.28266331 | 8 mo | HOM |
| **1013978366** | 1 | SSs | 378 | 9.290735866 | 0.000179109 | 6 | 1.93E-05 | 0.645804604 | 2 mo | HOM |
| **1014416155** | 2 | SSs | 378 | 9.290735866 | 0 | 0 | 0 | 0 | 2 mo | HOM |
| **1013978065** | 2 | SSs | 378 | 9.290735866 | 5.85E-05 | 2 | 6.3E-06 | 0.215268201 | 2 mo | HOM |
| **1009934102** | 2 | SSs | 378 | 9.290735866 | 1.53E-05 | 2 | 1.65E-06 | 0.215268201 | 4 mo | WILD |
| **1009322937** | 2 | SSs | 378 | 9.290735866 | 2.76E-05 | 4 | 2.97E-06 | 0.430536403 | 8 mo | WILD |
| **1011080637** | 2 | SSs | 378 | 9.290735866 | 0.00886582 | 346 | 0.000954265 | 37.24139885 | 8 mo | HOM |
| **1013609713** | 2 | SSs | 378 | 9.290735866 | 1.74E-05 | 0 | 1.87E-06 | 0 | 2 mo | WILD |
| **1010418842** | 1 | SSs | 378 | 9.290735866 | 0.013186763 | 380 | 0.001419345 | 40.90095828 | 8 mo | HOM |
| **1013288213** | 2 | SSs | 378 | 9.290735866 | 0.000453822 | 37 | 4.88E-05 | 3.982461727 | 4 mo | HOM |
| **1008976060** | 2 | SSs | 378 | 9.290735866 | 0.000265585 | 14 | 2.86E-05 | 1.50687741 | 8 mo | WILD |
| **1013446243** | 1 | SSs | 378 | 9.290735866 | 0.001776613 | 85 | 0.000191224 | 9.148898562 | 4 mo | HOM |
| **1013288951** | 2 | SSs | 378 | 9.290735866 | 0.00233716 | 91 | 0.000251558 | 9.794703166 | 4 mo | HOM |
| **1013978665** | 1 | MY-sat | 379 | 0.230292376 | 0 | 0 | 0 | 0 | 2 mo | HOM |
| **1013445609** | 1 | MY-sat | 379 | 0.230292376 | 0 | 0 | 0 | 0 | 4 mo | HOM |
| **1013609015** | 1 | MY-sat | 379 | 0.230292376 | 0 | 0 | 0 | 0 | 4 mo | HOM |
| **1013608106** | 2 | MY-sat | 379 | 0.223687479 | 0 | 0 | 0 | 0 | 4 mo | HOM |
| **1011937543** | 1 | MY-sat | 379 | 0.230292376 | 0 | 0 | 0 | 0 | 8 mo | HOM |
| **1010419538** | 2 | MY-sat | 379 | 0.230292376 | 0 | 0 | 0 | 0 | 4 mo | WILD |

| | | | | | | | | | | |
|---|---|---|---|---|---|---|---|---|---|---|
| **1010419187** | 1 | MY-sat | 379 | 0.230292376 | 0 | 0 | 0 | 0 | 8 mo | HOM |
| **1013978366** | 1 | MY-sat | 379 | 0.230292376 | 0 | 0 | 0 | 0 | 2 mo | HOM |
| **1014416155** | 2 | MY-sat | 379 | 0.230292376 | 0 | 0 | 0 | 0 | 2 mo | HOM |
| **1013978065** | 2 | MY-sat | 379 | 0.230292376 | 0 | 0 | 0 | 0 | 2 mo | HOM |
| **1009934102** | 2 | MY-sat | 379 | 0.230292376 | 0 | 0 | 0 | 0 | 4 mo | WILD |
| **1009322937** | 2 | MY-sat | 379 | 0.230292376 | 0 | 0 | 0 | 0 | 8 mo | WILD |
| **1011080637** | 2 | MY-sat | 379 | 0.230292376 | 1.3E-05 | 1 | 5.64E-05 | 4.342306155 | 8 mo | HOM |
| **1013609713** | 2 | MY-sat | 379 | 0.230292376 | 0 | 0 | 0 | 0 | 2 mo | WILD |
| **1010418842** | 1 | MY-sat | 379 | 0.230292376 | 2.94E-05 | 4 | 0.000127494 | 17.36922462 | 8 mo | HOM |
| **1013288213** | 2 | MY-sat | 379 | 0.230292376 | 0 | 0 | 0 | 0 | 4 mo | HOM |
| **1008976060** | 2 | MY-sat | 379 | 0.230292376 | 2.98E-05 | 3 | 0.000129498 | 13.02691846 | 8 mo | WILD |
| **1013446243** | 1 | MY-sat | 379 | 0.230292376 | 0 | 0 | 0 | 0 | 4 mo | HOM |
| **1013288951** | 2 | MY-sat | 379 | 0.230292376 | 0 | 0 | 0 | 0 | 4 mo | HOM |
| **1013978665** | 1 | cuf | 380 | 0.02298638 | 0 | 0 | 0 | 0 | 2 mo | HOM |
| **1013445609** | 1 | cuf | 380 | 0.02298638 | 0 | 0 | 0 | 0 | 4 mo | HOM |
| **1013609015** | 1 | cuf | 380 | 0.02298638 | 0 | 0 | 0 | 0 | 4 mo | HOM |
| **1013608106** | 2 | cuf | 380 | 0.02298638 | 0 | 0 | 0 | 0 | 4 mo | HOM |
| **1011937543** | 1 | cuf | 380 | 0.02298638 | 0 | 0 | 0 | 0 | 8 mo | HOM |
| **1010419538** | 2 | cuf | 380 | 0.02298638 | 0 | 0 | 0 | 0 | 4 mo | WILD |
| **1010419187** | 1 | cuf | 380 | 0.02298638 | 0 | 0 | 0 | 0 | 8 mo | HOM |
| **1013978366** | 1 | cuf | 380 | 0.02298638 | 0 | 0 | 0 | 0 | 2 mo | HOM |
| **1014416155** | 2 | cuf | 380 | 0.02298638 | 0 | 0 | 0 | 0 | 2 mo | HOM |
| **1013978065** | 2 | cuf | 380 | 0.02298638 | 0 | 0 | 0 | 0 | 2 mo | HOM |
| **1009934102** | 2 | cuf | 380 | 0.02298638 | 0 | 0 | 0 | 0 | 4 mo | WILD |
| **1009322937** | 2 | cuf | 380 | 0.02298638 | 0 | 0 | 0 | 0 | 8 mo | WILD |
| **1011080637** | 2 | cuf | 380 | 0.02298638 | 0 | 0 | 0 | 0 | 8 mo | HOM |
| **1013609713** | 2 | cuf | 380 | 0.02298638 | 0 | 0 | 0 | 0 | 2 mo | WILD |
| **1010418842** | 1 | cuf | 380 | 0.02298638 | 0 | 0 | 0 | 0 | 8 mo | HOM |
| **1013288213** | 2 | cuf | 380 | 0.02298638 | 0 | 0 | 0 | 0 | 4 mo | HOM |
| **1008976060** | 2 | cuf | 380 | 0.02298638 | 0 | 0 | 0 | 0 | 8 mo | WILD |
| **1013446243** | 1 | cuf | 380 | 0.02298638 | 0 | 0 | 0 | 0 | 4 mo | HOM |
| **1013288951** | 2 | cuf | 380 | 0.02298638 | 0 | 0 | 0 | 0 | 4 mo | HOM |
| **1013978665** | 1 | SNr | 381 | 1.596824074 | 0 | 0 | 0 | 0 | 2 mo | HOM |
| **1013445609** | 1 | SNr | 381 | 1.596824074 | 7.94E-06 | 1 | 4.97E-06 | 0.626243064 | 4 mo | HOM |
| **1013609015** | 1 | SNr | 381 | 1.596824074 | 9.08E-06 | 1 | 5.69E-06 | 0.626243064 | 4 mo | HOM |
| **1013608106** | 2 | SNr | 381 | 1.596824074 | 0 | 0 | 0 | 0 | 4 mo | HOM |
| **1011937543** | 1 | SNr | 381 | 1.596824074 | 0.002479031 | 48 | 0.001552476 | 30.05966706 | 8 mo | HOM |
| **1010419538** | 2 | SNr | 381 | 1.596824074 | 0 | 0 | 0 | 0 | 4 mo | WILD |
| **1010419187** | 1 | SNr | 381 | 1.596824074 | 0.000614862 | 9 | 0.000385053 | 5.636187573 | 8 mo | HOM |
| **1013978366** | 1 | SNr | 381 | 1.596824074 | 0 | 0 | 0 | 0 | 2 mo | HOM |
| **1014416155** | 2 | SNr | 381 | 1.596824074 | 0 | 0 | 0 | 0 | 2 mo | HOM |
| **1013978065** | 2 | SNr | 381 | 1.596824074 | 0 | 0 | 0 | 0 | 2 mo | HOM |
| **1009934102** | 2 | SNr | 381 | 1.596824074 | 0 | 0 | 0 | 0 | 4 mo | WILD |
| **1009322937** | 2 | SNr | 381 | 1.596824074 | 0 | 0 | 0 | 0 | 8 mo | WILD |
| **1011080637** | 2 | SNr | 381 | 1.596824074 | 0.000289125 | 12 | 0.000181063 | 7.514916764 | 8 mo | HOM |
| **1013609713** | 2 | SNr | 381 | 1.596824074 | 0 | 0 | 0 | 0 | 2 mo | WILD |
| **1010418842** | 1 | SNr | 381 | 1.596824074 | 0.001738825 | 33 | 0.001088927 | 20.6660211 | 8 mo | HOM |
| **1013288213** | 2 | SNr | 381 | 1.596824074 | 0 | 0 | 0 | 0 | 4 mo | HOM |
| **1008976060** | 2 | SNr | 381 | 1.596824074 | 8.49E-06 | 2 | 5.32E-06 | 1.252486127 | 8 mo | WILD |
| **1013446243** | 1 | SNr | 381 | 1.596824074 | 9.93E-06 | 1 | 6.22E-06 | 0.626243064 | 4 mo | HOM |
| **1013288951** | 2 | SNr | 381 | 1.596824074 | 9.58E-06 | 1 | 6E-06 | 0.626243064 | 4 mo | HOM |
| **1013978665** | 1 | CA1 | 382 | 10.49971498 | 3.57E-05 | 5 | 3.4E-06 | 0.476203403 | 2 mo | HOM |
| **1013445609** | 1 | CA1 | 382 | 10.49971498 | 0.000385884 | 28 | 3.68E-05 | 2.666739054 | 4 mo | HOM |
| **1013609015** | 1 | CA1 | 382 | 10.49971498 | 0.001638966 | 88 | 0.000156096 | 8.381179884 | 4 mo | HOM |
| **1013608106** | 2 | CA1 | 382 | 10.49971498 | 0 | 0 | 0 | 0 | 4 mo | HOM |
| **1011937543** | 1 | CA1 | 382 | 10.49971498 | 0.011762989 | 160 | 0.001120315 | 15.23850888 | 8 mo | HOM |
| **1010419538** | 2 | CA1 | 382 | 10.49971498 | 0 | 0 | 0 | 0 | 4 mo | WILD |
| **1010419187** | 1 | CA1 | 382 | 10.49971498 | 0.019873997 | 529 | 0.001892813 | 50.38231999 | 8 mo | HOM |
| **1013978366** | 1 | CA1 | 382 | 10.49971498 | 5.99E-06 | 1 | 5.7E-07 | 0.095240681 | 2 mo | HOM |

| | | | | | | | | | | |
|---|---|---|---|---|---|---|---|---|---|---|
| **1014416155** | 2 | CA1 | 382 | 10.49971498 | 0 | 0 | 0 | 0 | 2 mo | HOM |
| **1013978065** | 2 | CA1 | 382 | 10.49971498 | 0 | 0 | 0 | 0 | 2 mo | HOM |
| **1009934102** | 2 | CA1 | 382 | 10.49971498 | 1.63E-05 | 2 | 1.55E-06 | 0.190481361 | 4 mo | WILD |
| **1009322937** | 2 | CA1 | 382 | 10.49971498 | 7.38E-05 | 9 | 7.03E-06 | 0.857166125 | 8 mo | WILD |
| **1011080637** | 2 | CA1 | 382 | 10.49971498 | 0.014280255 | 423 | 0.001360061 | 40.28680785 | 8 mo | HOM |
| **1013609713** | 2 | CA1 | 382 | 10.49971498 | 0 | 0 | 0 | 0 | 2 mo | WILD |
| **1010418842** | 1 | CA1 | 382 | 10.49971498 | 0.018634176 | 502 | 0.001774732 | 47.81082161 | 8 mo | HOM |
| **1013288213** | 2 | CA1 | 382 | 10.49971498 | 0.000524742 | 39 | 5E-05 | 3.71438654 | 4 mo | HOM |
| **1008976060** | 2 | CA1 | 382 | 10.49971498 | 0.000209148 | 29 | 1.99E-05 | 2.761979735 | 8 mo | WILD |
| **1013446243** | 1 | CA1 | 382 | 10.49971498 | 0.001462826 | 77 | 0.000139321 | 7.333532399 | 4 mo | HOM |
| **1013288951** | 2 | CA1 | 382 | 10.49971498 | 0.001324654 | 88 | 0.000126161 | 8.381179884 | 4 mo | HOM |
| **1013978665** | 1 | VISp | 385 | 7.287949693 | 0.000668509 | 24 | 9.17E-05 | 3.293107254 | 2 mo | HOM |
| **1013445609** | 1 | VISp | 385 | 7.287949693 | 0.000920653 | 61 | 0.000126325 | 8.369980937 | 4 mo | HOM |
| **1013609015** | 1 | VISp | 385 | 7.287949693 | 0.003535166 | 105 | 0.00048507 | 14.40734424 | 4 mo | HOM |
| **1013608106** | 2 | VISp | 385 | 7.287949693 | 0 | 0 | 0 | 0 | 4 mo | HOM |
| **1011937543** | 1 | VISp | 385 | 7.287949693 | 0.014321318 | 181 | 0.001965068 | 24.83551721 | 8 mo | HOM |
| **1010419538** | 2 | VISp | 385 | 7.287949693 | 0.000305241 | 10 | 4.19E-05 | 1.372128022 | 4 mo | WILD |
| **1010419187** | 1 | VISp | 385 | 7.287949693 | 0.026731615 | 384 | 0.00366792 | 52.68971606 | 8 mo | HOM |
| **1013978366** | 1 | VISp | 385 | 7.287949693 | 0.000628226 | 14 | 8.62E-05 | 1.920979231 | 2 mo | HOM |
| **1014416155** | 2 | VISp | 385 | 7.287949693 | 0 | 0 | 0 | 0 | 2 mo | HOM |
| **1013978065** | 2 | VISp | 385 | 7.287949693 | 1.4E-05 | 2 | 1.92E-06 | 0.274425604 | 2 mo | HOM |
| **1009934102** | 2 | VISp | 385 | 7.287949693 | 0.0002852 | 10 | 3.91E-05 | 1.372128022 | 4 mo | WILD |
| **1009322937** | 2 | VISp | 385 | 7.287949693 | 7.67E-05 | 7 | 1.05E-05 | 0.960489616 | 8 mo | WILD |
| **1011080637** | 2 | VISp | 385 | 7.287949693 | 0.009881237 | 315 | 0.001355832 | 43.22203271 | 8 mo | HOM |
| **1013609713** | 2 | VISp | 385 | 7.287949693 | 5.47E-05 | 4 | 7.51E-06 | 0.548851209 | 2 mo | WILD |
| **1010418842** | 1 | VISp | 385 | 7.287949693 | 0.013172202 | 315 | 0.001807395 | 43.22203271 | 8 mo | HOM |
| **1013288213** | 2 | VISp | 385 | 7.287949693 | 0.000847409 | 66 | 0.000116275 | 9.056044948 | 4 mo | HOM |
| **1008976060** | 2 | VISp | 385 | 7.287949693 | 0.001915354 | 22 | 0.000262811 | 3.018681649 | 8 mo | WILD |
| **1013446243** | 1 | VISp | 385 | 7.287949693 | 0.002039295 | 96 | 0.000279817 | 13.17242902 | 4 mo | HOM |
| **1013288951** | 2 | VISp | 385 | 7.287949693 | 0.005812271 | 124 | 0.000797518 | 17.01438748 | 4 mo | HOM |
| **1013978665** | 1 | MY-sen | 386 | 8.172892368 | 3.34E-05 | 4 | 4.09E-06 | 0.489422816 | 2 mo | HOM |
| **1013445609** | 1 | MY-sen | 386 | 8.172892368 | 0.000160944 | 4 | 1.97E-05 | 0.489422816 | 4 mo | HOM |
| **1013609015** | 1 | MY-sen | 386 | 8.172892368 | 0.002913501 | 8 | 0.000356483 | 0.978845633 | 4 mo | HOM |
| **1013608106** | 2 | MY-sen | 386 | 8.172892368 | 0 | 0 | 0 | 0 | 4 mo | HOM |
| **1011937543** | 1 | MY-sen | 386 | 8.172892368 | 0.014122051 | 26 | 0.001727913 | 3.181248306 | 8 mo | HOM |
| **1010419538** | 2 | MY-sen | 386 | 8.172892368 | 2.68E-05 | 5 | 3.28E-06 | 0.61177852 | 4 mo | WILD |
| **1010419187** | 1 | MY-sen | 386 | 8.172892368 | 0.002673025 | 14 | 0.00032706 | 1.712979857 | 8 mo | HOM |
| **1013978366** | 1 | MY-sen | 386 | 8.172892368 | 0.001065937 | 6 | 0.000130424 | 0.734134224 | 2 mo | HOM |
| **1014416155** | 2 | MY-sen | 386 | 8.172892368 | 0 | 0 | 0 | 0 | 2 mo | HOM |
| **1013978065** | 2 | MY-sen | 386 | 8.172892368 | 0.000165463 | 4 | 2.02E-05 | 0.489422816 | 2 mo | HOM |
| **1009934102** | 2 | MY-sen | 386 | 8.172892368 | 1.91E-05 | 2 | 2.34E-06 | 0.244711408 | 4 mo | WILD |
| **1009322937** | 2 | MY-sen | 386 | 8.172892368 | 0.0002716 | 8 | 3.32E-05 | 0.978845633 | 8 mo | WILD |
| **1011080637** | 2 | MY-sen | 386 | 8.172892368 | 0.000812223 | 28 | 9.94E-05 | 3.425959714 | 8 mo | HOM |
| **1013609713** | 2 | MY-sen | 386 | 8.172892368 | 3.69E-07 | 0 | 4.51E-08 | 0 | 2 mo | WILD |
| **1010418842** | 1 | MY-sen | 386 | 8.172892368 | 0.000840816 | 43 | 0.000102879 | 5.261295275 | 8 mo | HOM |
| **1013288213** | 2 | MY-sen | 386 | 8.171344752 | 2.66E-06 | 0 | 3.25E-07 | 0 | 4 mo | HOM |
| **1008976060** | 2 | MY-sen | 386 | 8.172892368 | 0.001692713 | 35 | 0.000207113 | 4.282449642 | 8 mo | WILD |
| **1013446243** | 1 | MY-sen | 386 | 8.172892368 | 0.000713502 | 7 | 8.73E-05 | 0.856489928 | 4 mo | HOM |
| **1013288951** | 2 | MY-sen | 386 | 8.172892368 | 0.001474238 | 8 | 0.000180381 | 0.978845633 | 4 mo | HOM |
| **1013978665** | 1 | SO | 390 | 0.04836494 | 0 | 0 | 0 | 0 | 2 mo | HOM |
| **1013445609** | 1 | SO | 390 | 0.04836494 | 0 | 0 | 0 | 0 | 4 mo | HOM |
| **1013609015** | 1 | SO | 390 | 0.04836494 | 0 | 0 | 0 | 0 | 4 mo | HOM |
| **1013608106** | 2 | SO | 390 | 0.04836494 | 0 | 0 | 0 | 0 | 4 mo | HOM |
| **1011937543** | 1 | SO | 390 | 0.04836494 | 0.000134114 | 1 | 0.002772965 | 20.67613424 | 8 mo | HOM |
| **1010419538** | 2 | SO | 390 | 0.04836494 | 0 | 0 | 0 | 0 | 4 mo | WILD |
| **1010419187** | 1 | SO | 390 | 0.04836494 | 0 | 0 | 0 | 0 | 8 mo | HOM |
| **1013978366** | 1 | SO | 390 | 0.04836494 | 0 | 0 | 0 | 0 | 2 mo | HOM |
| **1014416155** | 2 | SO | 390 | 0.04836494 | 0 | 0 | 0 | 0 | 2 mo | HOM |
| **1013978065** | 2 | SO | 390 | 0.04836494 | 0 | 0 | 0 | 0 | 2 mo | HOM |

| | | | | | | | | | | |
|---|---|---|---|---|---|---|---|---|---|---|
| **1009934102** | 2 | SO | 390 | 0.04836494 | 0 | 0 | 0 | 0 | 4 mo | WILD |
| **1009322937** | 2 | SO | 390 | 0.04836494 | 0 | 0 | 0 | 0 | 8 mo | WILD |
| **1011080637** | 2 | SO | 390 | 0.04836494 | 1.32E-05 | 0 | 0.000273761 | 0 | 8 mo | HOM |
| **1013609713** | 2 | SO | 390 | 0.04836494 | 0 | 0 | 0 | 0 | 2 mo | WILD |
| **1010418842** | 1 | SO | 390 | 0.04836494 | 0 | 0 | 0 | 0 | 8 mo | HOM |
| **1013288213** | 2 | SO | 390 | 0.04836494 | 0 | 0 | 0 | 0 | 4 mo | HOM |
| **1008976060** | 2 | SO | 390 | 0.04836494 | 0 | 0 | 0 | 0 | 8 mo | WILD |
| **1013446243** | 1 | SO | 390 | 0.04836494 | 0 | 0 | 0 | 0 | 4 mo | HOM |
| **1013288951** | 2 | SO | 390 | 0.04836494 | 0 | 0 | 0 | 0 | 4 mo | HOM |
| **1013978665** | 1 | VISpl6b | 393 | 0.00906804 | 0 | 0 | 0 | 0 | 2 mo | HOM |
| **1013445609** | 1 | VISpl6b | 393 | 0.00906804 | 0 | 0 | 0 | 0 | 4 mo | HOM |
| **1013609015** | 1 | VISpl6b | 393 | 0.00906804 | 1.45E-08 | 0 | 1.6E-06 | 0 | 4 mo | HOM |
| **1013608106** | 2 | VISpl6b | 393 | 0.00906804 | 0 | 0 | 0 | 0 | 4 mo | HOM |
| **1011937543** | 1 | VISpl6b | 393 | 0.00906804 | 0 | 0 | 0 | 0 | 8 mo | HOM |
| **1010419538** | 2 | VISpl6b | 393 | 0.00906804 | 0 | 0 | 0 | 0 | 4 mo | WILD |
| **1010419187** | 1 | VISpl6b | 393 | 0.00906804 | 8.96E-06 | 1 | 0.00098773 | 110.2774084 | 8 mo | HOM |
| **1013978366** | 1 | VISpl6b | 393 | 0.00906804 | 0 | 0 | 0 | 0 | 2 mo | HOM |
| **1014416155** | 2 | VISpl6b | 393 | 0.00906804 | 0 | 0 | 0 | 0 | 2 mo | HOM |
| **1013978065** | 2 | VISpl6b | 393 | 0.00906804 | 0 | 0 | 0 | 0 | 2 mo | HOM |
| **1009934102** | 2 | VISpl6b | 393 | 0.00906804 | 0 | 0 | 0 | 0 | 4 mo | WILD |
| **1009322937** | 2 | VISpl6b | 393 | 0.00906804 | 0 | 0 | 0 | 0 | 8 mo | WILD |
| **1011080637** | 2 | VISpl6b | 393 | 0.00906804 | 2.02E-06 | 0 | 0.000222428 | 0 | 8 mo | HOM |
| **1013609713** | 2 | VISpl6b | 393 | 0.00906804 | 0 | 0 | 0 | 0 | 2 mo | WILD |
| **1010418842** | 1 | VISpl6b | 393 | 0.00906804 | 9.3E-09 | 0 | 1.03E-06 | 0 | 8 mo | HOM |
| **1013288213** | 2 | VISpl6b | 393 | 0.00906804 | 0 | 0 | 0 | 0 | 4 mo | HOM |
| **1008976060** | 2 | VISpl6b | 393 | 0.00906804 | 0 | 0 | 0 | 0 | 8 mo | WILD |
| **1013446243** | 1 | VISpl6b | 393 | 0.00906804 | 0 | 0 | 0 | 0 | 4 mo | HOM |
| **1013288951** | 2 | VISpl6b | 393 | 0.00906804 | 0 | 0 | 0 | 0 | 4 mo | HOM |
| **1013978665** | 1 | VISam | 394 | 0.816513051 | 0 | 0 | 0 | 0 | 2 mo | HOM |
| **1013445609** | 1 | VISam | 394 | 0.816513051 | 6.79E-05 | 3 | 8.32E-05 | 3.674160501 | 4 mo | HOM |
| **1013609015** | 1 | VISam | 394 | 0.816513051 | 0.000214322 | 10 | 0.000262485 | 12.24720167 | 4 mo | HOM |
| **1013608106** | 2 | VISam | 394 | 0.816513051 | 0 | 0 | 0 | 0 | 4 mo | HOM |
| **1011937543** | 1 | VISam | 394 | 0.816513051 | 0.002396751 | 18 | 0.00293535 | 22.04496301 | 8 mo | HOM |
| **1010419538** | 2 | VISam | 394 | 0.816513051 | 0 | 0 | 0 | 0 | 4 mo | WILD |
| **1010419187** | 1 | VISam | 394 | 0.816513051 | 0.001376966 | 48 | 0.001686398 | 58.78656802 | 8 mo | HOM |
| **1013978366** | 1 | VISam | 394 | 0.816513051 | 8.19E-06 | 1 | 1E-05 | 1.224720167 | 2 mo | HOM |
| **1014416155** | 2 | VISam | 394 | 0.816513051 | 0 | 0 | 0 | 0 | 2 mo | HOM |
| **1013978065** | 2 | VISam | 394 | 0.816513051 | 6.08E-06 | 0 | 7.44E-06 | 0 | 2 mo | HOM |
| **1009934102** | 2 | VISam | 394 | 0.816513051 | 0 | 0 | 0 | 0 | 4 mo | WILD |
| **1009322937** | 2 | VISam | 394 | 0.816513051 | 3.18E-08 | 0 | 3.9E-08 | 0 | 8 mo | WILD |
| **1011080637** | 2 | VISam | 394 | 0.816513051 | 0.000549062 | 40 | 0.000672447 | 48.98880668 | 8 mo | HOM |
| **1013609713** | 2 | VISam | 394 | 0.816513051 | 0 | 0 | 0 | 0 | 2 mo | WILD |
| **1010418842** | 1 | VISam | 394 | 0.816513051 | 0.001140636 | 32 | 0.00139696 | 39.19104534 | 8 mo | HOM |
| **1013288213** | 2 | VISam | 394 | 0.816513051 | 0.000106251 | 11 | 0.000130128 | 13.47192184 | 4 mo | HOM |
| **1008976060** | 2 | VISam | 394 | 0.816513051 | 6.55E-05 | 4 | 8.02E-05 | 4.898880668 | 8 mo | WILD |
| **1013446243** | 1 | VISam | 394 | 0.816513051 | 0.000169145 | 11 | 0.000207156 | 13.47192184 | 4 mo | HOM |
| **1013288951** | 2 | VISam | 394 | 0.816513051 | 0.00011835 | 8 | 0.000144946 | 9.797761336 | 4 mo | HOM |
| **1013978665** | 1 | MDRN | 395 | 2.097840998 | 0 | 0 | 0 | 0 | 2 mo | HOM |
| **1013445609** | 1 | MDRN | 395 | 2.097840998 | 0 | 0 | 0 | 0 | 4 mo | HOM |
| **1013609015** | 1 | MDRN | 395 | 2.097840998 | 0 | 0 | 0 | 0 | 4 mo | HOM |
| **1013608106** | 2 | MDRN | 395 | 2.097218894 | 0 | 0 | 0 | 0 | 4 mo | HOM |
| **1011937543** | 1 | MDRN | 395 | 2.097840998 | 4.48E-05 | 1 | 2.14E-05 | 0.47668055 | 8 mo | HOM |
| **1010419538** | 2 | MDRN | 395 | 2.097840998 | 0 | 0 | 0 | 0 | 4 mo | WILD |
| **1010419187** | 1 | MDRN | 395 | 2.097840998 | 6.34E-05 | 6 | 3.02E-05 | 2.860083297 | 8 mo | HOM |
| **1013978366** | 1 | MDRN | 395 | 2.097840998 | 0 | 0 | 0 | 0 | 2 mo | HOM |
| **1014416155** | 2 | MDRN | 395 | 2.097840998 | 0 | 0 | 0 | 0 | 2 mo | HOM |
| **1013978065** | 2 | MDRN | 395 | 2.097840998 | 0 | 0 | 0 | 0 | 2 mo | HOM |
| **1009934102** | 2 | MDRN | 395 | 2.097840998 | 0 | 0 | 0 | 0 | 4 mo | WILD |
| **1009322937** | 2 | MDRN | 395 | 2.097840998 | 0 | 0 | 0 | 0 | 8 mo | WILD |

| | | | | | | | | | | |
|---|---|---|---|---|---|---|---|---|---|---|
| **1011080637** | 2 | MDRN | 395 | 2.097840998 | 4.49E-05 | 3 | 2.14E-05 | 1.430041649 | 8 mo | HOM |
| **1013609713** | 2 | MDRN | 395 | 2.097840998 | 0 | 0 | 0 | 0 | 2 mo | WILD |
| **1010418842** | 1 | MDRN | 395 | 2.097840998 | 0.000320717 | 10 | 0.000152879 | 4.766805496 | 8 mo | HOM |
| **1013288213** | 2 | MDRN | 395 | 2.097839666 | 0 | 0 | 0 | 0 | 4 mo | HOM |
| **1008976060** | 2 | MDRN | 395 | 2.097840998 | 2.04E-05 | 4 | 9.74E-06 | 1.906722198 | 8 mo | WILD |
| **1013446243** | 1 | MDRN | 395 | 2.097840998 | 0 | 0 | 0 | 0 | 4 mo | HOM |
| **1013288951** | 2 | MDRN | 395 | 2.097840998 | 0 | 0 | 0 | 0 | 4 mo | HOM |
| **1013978665** | 1 | vtd | 397 | 0.038489206 | 0 | 0 | 0 | 0 | 2 mo | HOM |
| **1013445609** | 1 | vtd | 397 | 0.038489206 | 0 | 0 | 0 | 0 | 4 mo | HOM |
| **1013609015** | 1 | vtd | 397 | 0.038489206 | 0 | 0 | 0 | 0 | 4 mo | HOM |
| **1013608106** | 2 | vtd | 397 | 0.038489206 | 0 | 0 | 0 | 0 | 4 mo | HOM |
| **1011937543** | 1 | vtd | 397 | 0.038489206 | 0 | 0 | 0 | 0 | 8 mo | HOM |
| **1010419538** | 2 | vtd | 397 | 0.038489206 | 0 | 0 | 0 | 0 | 4 mo | WILD |
| **1010419187** | 1 | vtd | 397 | 0.038489206 | 0 | 0 | 0 | 0 | 8 mo | HOM |
| **1013978366** | 1 | vtd | 397 | 0.038489206 | 0 | 0 | 0 | 0 | 2 mo | HOM |
| **1014416155** | 2 | vtd | 397 | 0.038489206 | 0 | 0 | 0 | 0 | 2 mo | HOM |
| **1013978065** | 2 | vtd | 397 | 0.038489206 | 0 | 0 | 0 | 0 | 2 mo | HOM |
| **1009934102** | 2 | vtd | 397 | 0.038489206 | 0 | 0 | 0 | 0 | 4 mo | WILD |
| **1009322937** | 2 | vtd | 397 | 0.038489206 | 0 | 0 | 0 | 0 | 8 mo | WILD |
| **1011080637** | 2 | vtd | 397 | 0.038489206 | 0 | 0 | 0 | 0 | 8 mo | HOM |
| **1013609713** | 2 | vtd | 397 | 0.038489206 | 0 | 0 | 0 | 0 | 2 mo | WILD |
| **1010418842** | 1 | vtd | 397 | 0.038489206 | 0 | 0 | 0 | 0 | 8 mo | HOM |
| **1013288213** | 2 | vtd | 397 | 0.038489206 | 0 | 0 | 0 | 0 | 4 mo | HOM |
| **1008976060** | 2 | vtd | 397 | 0.038489206 | 0 | 0 | 0 | 0 | 8 mo | WILD |
| **1013446243** | 1 | vtd | 397 | 0.038489206 | 0 | 0 | 0 | 0 | 4 mo | HOM |
| **1013288951** | 2 | vtd | 397 | 0.038489206 | 0 | 0 | 0 | 0 | 4 mo | HOM |
| **1013978665** | 1 | SOC | 398 | 0.925803452 | 0 | 0 | 0 | 0 | 2 mo | HOM |
| **1013445609** | 1 | SOC | 398 | 0.925803452 | 0 | 0 | 0 | 0 | 4 mo | HOM |
| **1013609015** | 1 | SOC | 398 | 0.925803452 | 0 | 0 | 0 | 0 | 4 mo | HOM |
| **1013608106** | 2 | SOC | 398 | 0.925803452 | 0 | 0 | 0 | 0 | 4 mo | HOM |
| **1011937543** | 1 | SOC | 398 | 0.925803452 | 0.000381396 | 1 | 0.000411962 | 1.080142872 | 8 mo | HOM |
| **1010419538** | 2 | SOC | 398 | 0.925803452 | 0 | 0 | 0 | 0 | 4 mo | WILD |
| **1010419187** | 1 | SOC | 398 | 0.925803452 | 0 | 0 | 0 | 0 | 8 mo | HOM |
| **1013978366** | 1 | SOC | 398 | 0.925803452 | 0 | 0 | 0 | 0 | 2 mo | HOM |
| **1014416155** | 2 | SOC | 398 | 0.925803452 | 0 | 0 | 0 | 0 | 2 mo | HOM |
| **1013978065** | 2 | SOC | 398 | 0.925803452 | 0 | 0 | 0 | 0 | 2 mo | HOM |
| **1009934102** | 2 | SOC | 398 | 0.925803452 | 0 | 0 | 0 | 0 | 4 mo | WILD |
| **1009322937** | 2 | SOC | 398 | 0.925803452 | 0 | 0 | 0 | 0 | 8 mo | WILD |
| **1011080637** | 2 | SOC | 398 | 0.925803452 | 0 | 0 | 0 | 0 | 8 mo | HOM |
| **1013609713** | 2 | SOC | 398 | 0.925803452 | 0 | 0 | 0 | 0 | 2 mo | WILD |
| **1010418842** | 1 | SOC | 398 | 0.925803452 | 2.22E-05 | 1 | 2.4E-05 | 1.080142872 | 8 mo | HOM |
| **1013288213** | 2 | SOC | 398 | 0.925803452 | 0 | 0 | 0 | 0 | 4 mo | HOM |
| **1008976060** | 2 | SOC | 398 | 0.925803452 | 9.92E-06 | 2 | 1.07E-05 | 2.160285745 | 8 mo | WILD |
| **1013446243** | 1 | SOC | 398 | 0.925803452 | 0 | 0 | 0 | 0 | 4 mo | HOM |
| **1013288951** | 2 | SOC | 398 | 0.925803452 | 0 | 0 | 0 | 0 | 4 mo | HOM |
| **1013978665** | 1 | VISam4 | 401 | 0.061667206 | 0 | 0 | 0 | 0 | 2 mo | HOM |
| **1013445609** | 1 | VISam4 | 401 | 0.061667206 | 8.99E-07 | 0 | 1.46E-05 | 0 | 4 mo | HOM |
| **1013609015** | 1 | VISam4 | 401 | 0.061667206 | 6.57E-06 | 0 | 0.000106471 | 0 | 4 mo | HOM |
| **1013608106** | 2 | VISam4 | 401 | 0.061667206 | 0 | 0 | 0 | 0 | 4 mo | HOM |
| **1011937543** | 1 | VISam4 | 401 | 0.061667206 | 2.61E-05 | 0 | 0.000423611 | 0 | 8 mo | HOM |
| **1010419538** | 2 | VISam4 | 401 | 0.061667206 | 0 | 0 | 0 | 0 | 4 mo | WILD |
| **1010419187** | 1 | VISam4 | 401 | 0.061667206 | 0.000166178 | 4 | 0.002694749 | 64.86429799 | 8 mo | HOM |
| **1013978366** | 1 | VISam4 | 401 | 0.061667206 | 0 | 0 | 0 | 0 | 2 mo | HOM |
| **1014416155** | 2 | VISam4 | 401 | 0.061667206 | 0 | 0 | 0 | 0 | 2 mo | HOM |
| **1013978065** | 2 | VISam4 | 401 | 0.061667206 | 0 | 0 | 0 | 0 | 2 mo | HOM |
| **1009934102** | 2 | VISam4 | 401 | 0.061667206 | 0 | 0 | 0 | 0 | 4 mo | WILD |
| **1009322937** | 2 | VISam4 | 401 | 0.061667206 | 0 | 0 | 0 | 0 | 8 mo | WILD |
| **1011080637** | 2 | VISam4 | 401 | 0.061667206 | 2.48E-05 | 2 | 0.000402061 | 32.43214899 | 8 mo | HOM |
| **1013609713** | 2 | VISam4 | 401 | 0.061667206 | 0 | 0 | 0 | 0 | 2 mo | WILD |

| | | | | | | | | | | |
|---|---|---|---|---|---|---|---|---|---|---|
| **1010418842** | 1 | VISam4 | 401 | 0.061667206 | 0.000120424 | 5 | 0.001952803 | 81.08037248 | 8 mo | HOM |
| **1013288213** | 2 | VISam4 | 401 | 0.061667206 | 2.58E-05 | 2 | 0.000417703 | 32.43214899 | 4 mo | HOM |
| **1008976060** | 2 | VISam4 | 401 | 0.061667206 | 0 | 0 | 0 | 0 | 8 mo | WILD |
| **1013446243** | 1 | VISam4 | 401 | 0.061667206 | 9.32E-06 | 0 | 0.000151095 | 0 | 4 mo | HOM |
| **1013288951** | 2 | VISam4 | 401 | 0.061667206 | 7.67E-06 | 0 | 0.000124353 | 0 | 4 mo | HOM |
| **1013978665** | 1 | VISal | 402 | 0.781141313 | 3.43E-10 | 0 | 4.4E-10 | 0 | 2 mo | HOM |
| **1013445609** | 1 | VISal | 402 | 0.781141313 | 8.14E-05 | 7 | 0.000104147 | 8.961246682 | 4 mo | HOM |
| **1013609015** | 1 | VISal | 402 | 0.781141313 | 0.000325986 | 15 | 0.00041732 | 19.20267146 | 4 mo | HOM |
| **1013608106** | 2 | VISal | 402 | 0.781141313 | 0 | 0 | 0 | 0 | 4 mo | HOM |
| **1011937543** | 1 | VISal | 402 | 0.781141313 | 0.001467235 | 27 | 0.001878322 | 34.56480863 | 8 mo | HOM |
| **1010419538** | 2 | VISal | 402 | 0.781141313 | 0 | 0 | 0 | 0 | 4 mo | WILD |
| **1010419187** | 1 | VISal | 402 | 0.781141313 | 0.001148485 | 44 | 0.001470265 | 56.32783629 | 8 mo | HOM |
| **1013978366** | 1 | VISal | 402 | 0.781141313 | 8.2E-07 | 0 | 1.05E-06 | 0 | 2 mo | HOM |
| **1014416155** | 2 | VISal | 402 | 0.781141313 | 0 | 0 | 0 | 0 | 2 mo | HOM |
| **1013978065** | 2 | VISal | 402 | 0.781141313 | 0 | 0 | 0 | 0 | 2 mo | HOM |
| **1009934102** | 2 | VISal | 402 | 0.781141313 | 2.38E-06 | 1 | 3.05E-06 | 1.280178097 | 4 mo | WILD |
| **1009322937** | 2 | VISal | 402 | 0.781141313 | 0 | 0 | 0 | 0 | 8 mo | WILD |
| **1011080637** | 2 | VISal | 402 | 0.781141313 | 0.000760968 | 29 | 0.000974175 | 37.12516483 | 8 mo | HOM |
| **1013609713** | 2 | VISal | 402 | 0.781141313 | 0 | 0 | 0 | 0 | 2 mo | WILD |
| **1010418842** | 1 | VISal | 402 | 0.781141313 | 0.001161312 | 35 | 0.001486686 | 44.80623341 | 8 mo | HOM |
| **1013288213** | 2 | VISal | 402 | 0.781141313 | 0.000105809 | 8 | 0.000135455 | 10.24142478 | 4 mo | HOM |
| **1008976060** | 2 | VISal | 402 | 0.781141313 | 5.19E-05 | 5 | 6.64E-05 | 6.400890487 | 8 mo | WILD |
| **1013446243** | 1 | VISal | 402 | 0.781141313 | 0.000177829 | 11 | 0.000227653 | 14.08195907 | 4 mo | HOM |
| **1013288951** | 2 | VISal | 402 | 0.781141313 | 0.00028664 | 10 | 0.00036695 | 12.80178097 | 4 mo | HOM |
| **1013978665** | 1 | MEA | 403 | 2.089895942 | 0 | 0 | 0 | 0 | 2 mo | HOM |
| **1013445609** | 1 | MEA | 403 | 2.089895942 | 2.93E-05 | 6 | 1.4E-05 | 2.870956337 | 4 mo | HOM |
| **1013609015** | 1 | MEA | 403 | 2.089895942 | 0.000329079 | 8 | 0.000157462 | 3.827941783 | 4 mo | HOM |
| **1013608106** | 2 | MEA | 403 | 2.089895942 | 0 | 0 | 0 | 0 | 4 mo | HOM |
| **1011937543** | 1 | MEA | 403 | 2.089895942 | 0.00402411 | 59 | 0.001925507 | 28.23107065 | 8 mo | HOM |
| **1010419538** | 2 | MEA | 403 | 2.089895942 | 1.47E-08 | 0 | 7.06E-09 | 0 | 4 mo | WILD |
| **1010419187** | 1 | MEA | 403 | 2.089895942 | 0.003244854 | 97 | 0.001552639 | 46.41379412 | 8 mo | HOM |
| **1013978366** | 1 | MEA | 403 | 2.089895942 | 1.54E-09 | 0 | 7.38E-10 | 0 | 2 mo | HOM |
| **1014416155** | 2 | MEA | 403 | 2.089895942 | 0 | 0 | 0 | 0 | 2 mo | HOM |
| **1013978065** | 2 | MEA | 403 | 2.089895942 | 0 | 0 | 0 | 0 | 2 mo | HOM |
| **1009934102** | 2 | MEA | 403 | 2.089895942 | 2.27E-05 | 1 | 1.09E-05 | 0.478492723 | 4 mo | WILD |
| **1009322937** | 2 | MEA | 403 | 2.089895942 | 0 | 0 | 0 | 0 | 8 mo | WILD |
| **1011080637** | 2 | MEA | 403 | 2.089895942 | 0.002522246 | 80 | 0.001206876 | 38.27941783 | 8 mo | HOM |
| **1013609713** | 2 | MEA | 403 | 2.089895942 | 0 | 0 | 0 | 0 | 2 mo | WILD |
| **1010418842** | 1 | MEA | 403 | 2.089895942 | 0.003383276 | 89 | 0.001618873 | 42.58585234 | 8 mo | HOM |
| **1013288213** | 2 | MEA | 403 | 2.089895942 | 6.89E-05 | 5 | 3.3E-05 | 2.392463614 | 4 mo | HOM |
| **1008976060** | 2 | MEA | 403 | 2.089895942 | 2.32E-05 | 1 | 1.11E-05 | 0.478492723 | 8 mo | WILD |
| **1013446243** | 1 | MEA | 403 | 2.089895942 | 4.59E-05 | 3 | 2.2E-05 | 1.435478169 | 4 mo | HOM |
| **1013288951** | 2 | MEA | 403 | 2.089895942 | 0.000182374 | 5 | 8.73E-05 | 2.392463614 | 4 mo | HOM |
| **1013978665** | 1 | SPF | 406 | 0.211187883 | 0 | 0 | 0 | 0 | 2 mo | HOM |
| **1013445609** | 1 | SPF | 406 | 0.211187883 | 0 | 0 | 0 | 0 | 4 mo | HOM |
| **1013609015** | 1 | SPF | 406 | 0.211187883 | 0 | 0 | 0 | 0 | 4 mo | HOM |
| **1013608106** | 2 | SPF | 406 | 0.211187883 | 0 | 0 | 0 | 0 | 4 mo | HOM |
| **1011937543** | 1 | SPF | 406 | 0.211187883 | 1.52E-05 | 1 | 7.21E-05 | 4.735120144 | 8 mo | HOM |
| **1010419538** | 2 | SPF | 406 | 0.211187883 | 0 | 0 | 0 | 0 | 4 mo | WILD |
| **1010419187** | 1 | SPF | 406 | 0.211187883 | 4.78E-06 | 0 | 2.26E-05 | 0 | 8 mo | HOM |
| **1013978366** | 1 | SPF | 406 | 0.211187883 | 0 | 0 | 0 | 0 | 2 mo | HOM |
| **1014416155** | 2 | SPF | 406 | 0.211187883 | 0 | 0 | 0 | 0 | 2 mo | HOM |
| **1013978065** | 2 | SPF | 406 | 0.211187883 | 0 | 0 | 0 | 0 | 2 mo | HOM |
| **1009934102** | 2 | SPF | 406 | 0.211187883 | 0 | 0 | 0 | 0 | 4 mo | WILD |
| **1009322937** | 2 | SPF | 406 | 0.211187883 | 0 | 0 | 0 | 0 | 8 mo | WILD |
| **1011080637** | 2 | SPF | 406 | 0.211187883 | 0 | 0 | 0 | 0 | 8 mo | HOM |
| **1013609713** | 2 | SPF | 406 | 0.211187883 | 0 | 0 | 0 | 0 | 2 mo | WILD |
| **1010418842** | 1 | SPF | 406 | 0.211187883 | 5.51E-05 | 2 | 0.00026072 | 9.470240289 | 8 mo | HOM |
| **1013288213** | 2 | SPF | 406 | 0.211187883 | 0 | 0 | 0 | 0 | 4 mo | HOM |

| | | | | | | | | | | |
|---|---|---|---|---|---|---|---|---|---|---|
| **1008976060** | 2 | SPF | 406 | 0.211187883 | 0 | 0 | 0 | 0 | 8 mo | WILD |
| **1013446243** | 1 | SPF | 406 | 0.211187883 | 0 | 0 | 0 | 0 | 4 mo | HOM |
| **1013288951** | 2 | SPF | 406 | 0.211187883 | 0 | 0 | 0 | 0 | 4 mo | HOM |
| **1013978665** | 1 | VISI | 409 | 1.262668495 | 2.87E-05 | 3 | 2.27E-05 | 2.37592053 | 2 mo | HOM |
| **1013445609** | 1 | VISI | 409 | 1.262668495 | 8.87E-05 | 10 | 7.03E-05 | 7.919735099 | 4 mo | HOM |
| **1013609015** | 1 | VISI | 409 | 1.262668495 | 0.000253196 | 13 | 0.000200525 | 10.29565563 | 4 mo | HOM |
| **1013608106** | 2 | VISI | 409 | 1.262668495 | 0 | 0 | 0 | 0 | 4 mo | HOM |
| **1011937543** | 1 | VISI | 409 | 1.262668495 | 0.002115351 | 40 | 0.001675302 | 31.6789404 | 8 mo | HOM |
| **1010419538** | 2 | VISI | 409 | 1.262668495 | 0 | 0 | 0 | 0 | 4 mo | WILD |
| **1010419187** | 1 | VISI | 409 | 1.262668495 | 0.003580134 | 77 | 0.002835371 | 60.98196026 | 8 mo | HOM |
| **1013978366** | 1 | VISI | 409 | 1.262668495 | 1.5E-05 | 1 | 1.18E-05 | 0.79197351 | 2 mo | HOM |
| **1014416155** | 2 | VISI | 409 | 1.262668495 | 0 | 0 | 0 | 0 | 2 mo | HOM |
| **1013978065** | 2 | VISI | 409 | 1.262668495 | 0 | 0 | 0 | 0 | 2 mo | HOM |
| **1009934102** | 2 | VISI | 409 | 1.262668495 | 0 | 0 | 0 | 0 | 4 mo | WILD |
| **1009322937** | 2 | VISI | 409 | 1.262668495 | 0 | 0 | 0 | 0 | 8 mo | WILD |
| **1011080637** | 2 | VISI | 409 | 1.262668495 | 0.001324962 | 58 | 0.001049335 | 45.93446357 | 8 mo | HOM |
| **1013609713** | 2 | VISI | 409 | 1.262668495 | 0 | 0 | 0 | 0 | 2 mo | WILD |
| **1010418842** | 1 | VISI | 409 | 1.262668495 | 0.001772555 | 67 | 0.001403817 | 53.06222516 | 8 mo | HOM |
| **1013288213** | 2 | VISI | 409 | 1.262668495 | 0.000253878 | 12 | 0.000201064 | 9.503682119 | 4 mo | HOM |
| **1008976060** | 2 | VISI | 409 | 1.262668495 | 0.000186691 | 10 | 0.000147854 | 7.919735099 | 8 mo | WILD |
| **1013446243** | 1 | VISI | 409 | 1.262668495 | 0.000503052 | 26 | 0.000398404 | 20.59131126 | 4 mo | HOM |
| **1013288951** | 2 | VISI | 409 | 1.262668495 | 0.000561952 | 20 | 0.000445051 | 15.8394702 | 4 mo | HOM |
| **1013978665** | 1 | ORBI2/3 | 412 | 0.639963402 | 0 | 0 | 0 | 0 | 2 mo | HOM |
| **1013445609** | 1 | ORBI2/3 | 412 | 0.639963402 | 4.34E-05 | 2 | 6.78E-05 | 3.125178709 | 4 mo | HOM |
| **1013609015** | 1 | ORBI2/3 | 412 | 0.639963402 | 7.86E-06 | 1 | 1.23E-05 | 1.562589355 | 4 mo | HOM |
| **1013608106** | 2 | ORBI2/3 | 412 | 0.639963402 | 0 | 0 | 0 | 0 | 4 mo | HOM |
| **1011937543** | 1 | ORBI2/3 | 412 | 0.639963402 | 0.000426367 | 11 | 0.000666237 | 17.1884829 | 8 mo | HOM |
| **1010419538** | 2 | ORBI2/3 | 412 | 0.639963402 | 0 | 0 | 0 | 0 | 4 mo | WILD |
| **1010419187** | 1 | ORBI2/3 | 412 | 0.639963402 | 0.00083518 | 31 | 0.001305044 | 48.44027 | 8 mo | HOM |
| **1013978366** | 1 | ORBI2/3 | 412 | 0.639963402 | 3.98E-08 | 0 | 6.23E-08 | 0 | 2 mo | HOM |
| **1014416155** | 2 | ORBI2/3 | 412 | 0.639963402 | 0 | 0 | 0 | 0 | 2 mo | HOM |
| **1013978065** | 2 | ORBI2/3 | 412 | 0.639963402 | 0 | 0 | 0 | 0 | 2 mo | HOM |
| **1009934102** | 2 | ORBI2/3 | 412 | 0.639963402 | 0 | 0 | 0 | 0 | 4 mo | WILD |
| **1009322937** | 2 | ORBI2/3 | 412 | 0.639963402 | 3.15E-05 | 1 | 4.93E-05 | 1.562589355 | 8 mo | WILD |
| **1011080637** | 2 | ORBI2/3 | 412 | 0.639963402 | 0.00035712 | 17 | 0.000558032 | 26.56401903 | 8 mo | HOM |
| **1013609713** | 2 | ORBI2/3 | 412 | 0.639963402 | 0 | 0 | 0 | 0 | 2 mo | WILD |
| **1010418842** | 1 | ORBI2/3 | 412 | 0.639963402 | 0.000537461 | 24 | 0.000839831 | 37.50214451 | 8 mo | HOM |
| **1013288213** | 2 | ORBI2/3 | 412 | 0.639963402 | 4.2E-05 | 4 | 6.57E-05 | 6.250357419 | 4 mo | HOM |
| **1008976060** | 2 | ORBI2/3 | 412 | 0.639963402 | 3.19E-07 | 0 | 4.98E-07 | 0 | 8 mo | WILD |
| **1013446243** | 1 | ORBI2/3 | 412 | 0.639963402 | 9.02E-05 | 5 | 0.000140892 | 7.812946774 | 4 mo | HOM |
| **1013288951** | 2 | ORBI2/3 | 412 | 0.639963402 | 3.55E-05 | 5 | 5.55E-05 | 7.812946774 | 4 mo | HOM |
| **1013978665** | 1 | vVIIIn | 413 | 0.25771697 | 0 | 0 | 0 | 0 | 2 mo | HOM |
| **1013445609** | 1 | vVIIIn | 413 | 0.25771697 | 0 | 0 | 0 | 0 | 4 mo | HOM |
| **1013609015** | 1 | vVIIIn | 413 | 0.25771697 | 0 | 0 | 0 | 0 | 4 mo | HOM |
| **1013608106** | 2 | vVIIIn | 413 | 0.25771697 | 0 | 0 | 0 | 0 | 4 mo | HOM |
| **1011937543** | 1 | vVIIIn | 413 | 0.25771697 | 0.000220887 | 0 | 0.00085709 | 0 | 8 mo | HOM |
| **1010419538** | 2 | vVIIIn | 413 | 0.25771697 | 0 | 0 | 0 | 0 | 4 mo | WILD |
| **1010419187** | 1 | vVIIIn | 413 | 0.25771697 | 1.1E-05 | 1 | 4.28E-05 | 3.88022566 | 8 mo | HOM |
| **1013978366** | 1 | vVIIIn | 413 | 0.25771697 | 0 | 0 | 0 | 0 | 2 mo | HOM |
| **1014416155** | 2 | vVIIIn | 413 | 0.25771697 | 0 | 0 | 0 | 0 | 2 mo | HOM |
| **1013978065** | 2 | vVIIIn | 413 | 0.25771697 | 0 | 0 | 0 | 0 | 2 mo | HOM |
| **1009934102** | 2 | vVIIIn | 413 | 0.25771697 | 0 | 0 | 0 | 0 | 4 mo | WILD |
| **1009322937** | 2 | vVIIIn | 413 | 0.25771697 | 0 | 0 | 0 | 0 | 8 mo | WILD |
| **1011080637** | 2 | vVIIIn | 413 | 0.25771697 | 0 | 0 | 0 | 0 | 8 mo | HOM |
| **1013609713** | 2 | vVIIIn | 413 | 0.25771697 | 0 | 0 | 0 | 0 | 2 mo | WILD |
| **1010418842** | 1 | vVIIIn | 413 | 0.25771697 | 1.78E-07 | 0 | 6.91E-07 | 0 | 8 mo | HOM |
| **1013288213** | 2 | vVIIIn | 413 | 0.25771697 | 0 | 0 | 0 | 0 | 4 mo | HOM |
| **1008976060** | 2 | vVIIIn | 413 | 0.25771697 | 0 | 0 | 0 | 0 | 8 mo | WILD |
| **1013446243** | 1 | vVIIIn | 413 | 0.25771697 | 0 | 0 | 0 | 0 | 4 mo | HOM |

| | | | | | | | | | | |
|---|---|---|---|---|---|---|---|---|---|---|
| **1013288951** | 2 | vVIIIn | 413 | 0.25771697 | 0 | 0 | 0 | 0 | 4 mo | HOM |
| **1013978665** | 1 | SPFm | 414 | 0.072191984 | 0 | 0 | 0 | 0 | 2 mo | HOM |
| **1013445609** | 1 | SPFm | 414 | 0.072191984 | 0 | 0 | 0 | 0 | 4 mo | HOM |
| **1013609015** | 1 | SPFm | 414 | 0.072191984 | 0 | 0 | 0 | 0 | 4 mo | HOM |
| **1013608106** | 2 | SPFm | 414 | 0.072191984 | 0 | 0 | 0 | 0 | 4 mo | HOM |
| **1011937543** | 1 | SPFm | 414 | 0.072191984 | 5.76E-06 | 0 | 7.97E-05 | 0 | 8 mo | HOM |
| **1010419538** | 2 | SPFm | 414 | 0.072191984 | 0 | 0 | 0 | 0 | 4 mo | WILD |
| **1010419187** | 1 | SPFm | 414 | 0.072191984 | 4.78E-06 | 0 | 6.62E-05 | 0 | 8 mo | HOM |
| **1013978366** | 1 | SPFm | 414 | 0.072191984 | 0 | 0 | 0 | 0 | 2 mo | HOM |
| **1014416155** | 2 | SPFm | 414 | 0.072191984 | 0 | 0 | 0 | 0 | 2 mo | HOM |
| **1013978065** | 2 | SPFm | 414 | 0.072191984 | 0 | 0 | 0 | 0 | 2 mo | HOM |
| **1009934102** | 2 | SPFm | 414 | 0.072191984 | 0 | 0 | 0 | 0 | 4 mo | WILD |
| **1009322937** | 2 | SPFm | 414 | 0.072191984 | 0 | 0 | 0 | 0 | 8 mo | WILD |
| **1011080637** | 2 | SPFm | 414 | 0.072191984 | 0 | 0 | 0 | 0 | 8 mo | HOM |
| **1013609713** | 2 | SPFm | 414 | 0.072191984 | 0 | 0 | 0 | 0 | 2 mo | WILD |
| **1010418842** | 1 | SPFm | 414 | 0.072191984 | 1.37E-05 | 1 | 0.000190112 | 13.85195339 | 8 mo | HOM |
| **1013288213** | 2 | SPFm | 414 | 0.072191984 | 0 | 0 | 0 | 0 | 4 mo | HOM |
| **1008976060** | 2 | SPFm | 414 | 0.072191984 | 0 | 0 | 0 | 0 | 8 mo | WILD |
| **1013446243** | 1 | SPFm | 414 | 0.072191984 | 0 | 0 | 0 | 0 | 4 mo | HOM |
| **1013288951** | 2 | SPFm | 414 | 0.072191984 | 0 | 0 | 0 | 0 | 4 mo | HOM |
| **1013978665** | 1 | VISrl | 417 | 1.048422872 | 1.22E-05 | 1 | 1.17E-05 | 0.953813606 | 2 mo | HOM |
| **1013445609** | 1 | VISrl | 417 | 1.048422872 | 0.000179946 | 15 | 0.000171635 | 14.30720408 | 4 mo | HOM |
| **1013609015** | 1 | VISrl | 417 | 1.048422872 | 0.000529013 | 19 | 0.00050458 | 18.12245851 | 4 mo | HOM |
| **1013608106** | 2 | VISrl | 417 | 1.048422872 | 0 | 0 | 0 | 0 | 4 mo | HOM |
| **1011937543** | 1 | VISrl | 417 | 1.048422872 | 0.002506892 | 32 | 0.002391108 | 30.52203538 | 8 mo | HOM |
| **1010419538** | 2 | VISrl | 417 | 1.048422872 | 0 | 0 | 0 | 0 | 4 mo | WILD |
| **1010419187** | 1 | VISrl | 417 | 1.048422872 | 0.002448348 | 62 | 0.002335268 | 59.13644354 | 8 mo | HOM |
| **1013978366** | 1 | VISrl | 417 | 1.048422872 | 3.71E-06 | 1 | 3.54E-06 | 0.953813606 | 2 mo | HOM |
| **1014416155** | 2 | VISrl | 417 | 1.048422872 | 0 | 0 | 0 | 0 | 2 mo | HOM |
| **1013978065** | 2 | VISrl | 417 | 1.048422872 | 1.46E-06 | 0 | 1.39E-06 | 0 | 2 mo | HOM |
| **1009934102** | 2 | VISrl | 417 | 1.048422872 | 6.67E-08 | 0 | 6.37E-08 | 0 | 4 mo | WILD |
| **1009322937** | 2 | VISrl | 417 | 1.048422872 | 0 | 0 | 0 | 0 | 8 mo | WILD |
| **1011080637** | 2 | VISrl | 417 | 1.048422872 | 0.001069275 | 31 | 0.00101989 | 29.56822177 | 8 mo | HOM |
| **1013609713** | 2 | VISrl | 417 | 1.048422872 | 1.12E-05 | 1 | 1.06E-05 | 0.953813606 | 2 mo | WILD |
| **1010418842** | 1 | VISrl | 417 | 1.048422872 | 0.002056841 | 46 | 0.001961843 | 43.87542585 | 8 mo | HOM |
| **1013288213** | 2 | VISrl | 417 | 1.048422872 | 0.000130758 | 12 | 0.000124719 | 11.44576327 | 4 mo | HOM |
| **1008976060** | 2 | VISrl | 417 | 1.048422872 | 5.1E-05 | 1 | 4.87E-05 | 0.953813606 | 8 mo | WILD |
| **1013446243** | 1 | VISrl | 417 | 1.048422872 | 0.000415999 | 17 | 0.000396786 | 16.21483129 | 4 mo | HOM |
| **1013288951** | 2 | VISrl | 417 | 1.048422872 | 0.000432741 | 9 | 0.000412754 | 8.58432245 | 4 mo | HOM |
| **1013978665** | 1 | VISl1 | 421 | 0.190744103 | 2.21E-05 | 2 | 0.000116105 | 10.48525205 | 2 mo | HOM |
| **1013445609** | 1 | VISl1 | 421 | 0.190744103 | 6.47E-06 | 1 | 3.39E-05 | 5.242626027 | 4 mo | HOM |
| **1013609015** | 1 | VISl1 | 421 | 0.190744103 | 8.06E-06 | 1 | 4.23E-05 | 5.242626027 | 4 mo | HOM |
| **1013608106** | 2 | VISl1 | 421 | 0.190744103 | 0 | 0 | 0 | 0 | 4 mo | HOM |
| **1011937543** | 1 | VISl1 | 421 | 0.190744103 | 0.000643068 | 10 | 0.003371366 | 52.42626027 | 8 mo | HOM |
| **1010419538** | 2 | VISl1 | 421 | 0.190744103 | 0 | 0 | 0 | 0 | 4 mo | WILD |
| **1010419187** | 1 | VISl1 | 421 | 0.190744103 | 0.001629221 | 5 | 0.008541399 | 26.21313014 | 8 mo | HOM |
| **1013978366** | 1 | VISl1 | 421 | 0.190744103 | 0 | 0 | 0 | 0 | 2 mo | HOM |
| **1014416155** | 2 | VISl1 | 421 | 0.190744103 | 0 | 0 | 0 | 0 | 2 mo | HOM |
| **1013978065** | 2 | VISl1 | 421 | 0.190744103 | 0 | 0 | 0 | 0 | 2 mo | HOM |
| **1009934102** | 2 | VISl1 | 421 | 0.190744103 | 0 | 0 | 0 | 0 | 4 mo | WILD |
| **1009322937** | 2 | VISl1 | 421 | 0.190744103 | 0 | 0 | 0 | 0 | 8 mo | WILD |
| **1011080637** | 2 | VISl1 | 421 | 0.190744103 | 0.000368093 | 16 | 0.001929772 | 83.88201644 | 8 mo | HOM |
| **1013609713** | 2 | VISl1 | 421 | 0.190744103 | 0 | 0 | 0 | 0 | 2 mo | WILD |
| **1010418842** | 1 | VISl1 | 421 | 0.190744103 | 0.000374186 | 10 | 0.001961719 | 52.42626027 | 8 mo | HOM |
| **1013288213** | 2 | VISl1 | 421 | 0.190744103 | 2.66E-05 | 2 | 0.000139236 | 10.48525205 | 4 mo | HOM |
| **1008976060** | 2 | VISl1 | 421 | 0.190744103 | 0.000186691 | 10 | 0.000978751 | 52.42626027 | 8 mo | WILD |
| **1013446243** | 1 | VISl1 | 421 | 0.190744103 | 0.000176191 | 6 | 0.000923705 | 31.45575616 | 4 mo | HOM |
| **1013288951** | 2 | VISl1 | 421 | 0.190744103 | 0.000379958 | 9 | 0.001991979 | 47.18363425 | 4 mo | HOM |
| **1013978665** | 1 | SPFp | 422 | 0.138995899 | 0 | 0 | 0 | 0 | 2 mo | HOM |

| | | | | | | | | | | |
|---|---|---|---|---|---|---|---|---|---|---|
| **1013445609** | 1 | SPFp | 422 | 0.138995899 | 0 | 0 | 0 | 0 | 4 mo | HOM |
| **1013609015** | 1 | SPFp | 422 | 0.138995899 | 0 | 0 | 0 | 0 | 4 mo | HOM |
| **1013608106** | 2 | SPFp | 422 | 0.138995899 | 0 | 0 | 0 | 0 | 4 mo | HOM |
| **1011937543** | 1 | SPFp | 422 | 0.138995899 | 9.47E-06 | 1 | 6.81E-05 | 7.194456867 | 8 mo | HOM |
| **1010419538** | 2 | SPFp | 422 | 0.138995899 | 0 | 0 | 0 | 0 | 4 mo | WILD |
| **1010419187** | 1 | SPFp | 422 | 0.138995899 | 0 | 0 | 0 | 0 | 8 mo | HOM |
| **1013978366** | 1 | SPFp | 422 | 0.138995899 | 0 | 0 | 0 | 0 | 2 mo | HOM |
| **1014416155** | 2 | SPFp | 422 | 0.138995899 | 0 | 0 | 0 | 0 | 2 mo | HOM |
| **1013978065** | 2 | SPFp | 422 | 0.138995899 | 0 | 0 | 0 | 0 | 2 mo | HOM |
| **1009934102** | 2 | SPFp | 422 | 0.138995899 | 0 | 0 | 0 | 0 | 4 mo | WILD |
| **1009322937** | 2 | SPFp | 422 | 0.138995899 | 0 | 0 | 0 | 0 | 8 mo | WILD |
| **1011080637** | 2 | SPFp | 422 | 0.138995899 | 0 | 0 | 0 | 0 | 8 mo | HOM |
| **1013609713** | 2 | SPFp | 422 | 0.138995899 | 0 | 0 | 0 | 0 | 2 mo | WILD |
| **1010418842** | 1 | SPFp | 422 | 0.138995899 | 4.13E-05 | 1 | 0.000297393 | 7.194456867 | 8 mo | HOM |
| **1013288213** | 2 | SPFp | 422 | 0.138995899 | 0 | 0 | 0 | 0 | 4 mo | HOM |
| **1008976060** | 2 | SPFp | 422 | 0.138995899 | 0 | 0 | 0 | 0 | 8 mo | WILD |
| **1013446243** | 1 | SPFp | 422 | 0.138995899 | 0 | 0 | 0 | 0 | 4 mo | HOM |
| **1013288951** | 2 | SPFp | 422 | 0.138995899 | 0 | 0 | 0 | 0 | 4 mo | HOM |
| **1013978665** | 1 | CA2 | 423 | 0.490314972 | 0 | 0 | 0 | 0 | 2 mo | HOM |
| **1013445609** | 1 | CA2 | 423 | 0.490314972 | 2.36E-05 | 3 | 4.81E-05 | 6.118515998 | 4 mo | HOM |
| **1013609015** | 1 | CA2 | 423 | 0.490314972 | 9.09E-05 | 3 | 0.000185491 | 6.118515998 | 4 mo | HOM |
| **1013608106** | 2 | CA2 | 423 | 0.490314972 | 0 | 0 | 0 | 0 | 4 mo | HOM |
| **1011937543** | 1 | CA2 | 423 | 0.490314972 | 0.000653953 | 7 | 0.00133374 | 14.27653733 | 8 mo | HOM |
| **1010419538** | 2 | CA2 | 423 | 0.490314972 | 0 | 0 | 0 | 0 | 4 mo | WILD |
| **1010419187** | 1 | CA2 | 423 | 0.490314972 | 0.000806115 | 20 | 0.001644076 | 40.79010665 | 8 mo | HOM |
| **1013978366** | 1 | CA2 | 423 | 0.490314972 | 0 | 0 | 0 | 0 | 2 mo | HOM |
| **1014416155** | 2 | CA2 | 423 | 0.490314972 | 0 | 0 | 0 | 0 | 2 mo | HOM |
| **1013978065** | 2 | CA2 | 423 | 0.490314972 | 0 | 0 | 0 | 0 | 2 mo | HOM |
| **1009934102** | 2 | CA2 | 423 | 0.490314972 | 1.58E-08 | 0 | 3.22E-08 | 0 | 4 mo | WILD |
| **1009322937** | 2 | CA2 | 423 | 0.490314972 | 0 | 0 | 0 | 0 | 8 mo | WILD |
| **1011080637** | 2 | CA2 | 423 | 0.490314972 | 0.000435698 | 13 | 0.000888607 | 26.51356932 | 8 mo | HOM |
| **1013609713** | 2 | CA2 | 423 | 0.490314972 | 0 | 0 | 0 | 0 | 2 mo | WILD |
| **1010418842** | 1 | CA2 | 423 | 0.490314972 | 0.000827317 | 27 | 0.001687318 | 55.06664398 | 8 mo | HOM |
| **1013288213** | 2 | CA2 | 423 | 0.490314972 | 3.05E-05 | 2 | 6.22E-05 | 4.079010665 | 4 mo | HOM |
| **1008976060** | 2 | CA2 | 423 | 0.490314972 | 1.76E-05 | 3 | 3.6E-05 | 6.118515998 | 8 mo | WILD |
| **1013446243** | 1 | CA2 | 423 | 0.490314972 | 7.01E-05 | 6 | 0.00014307 | 12.237032 | 4 mo | HOM |
| **1013288951** | 2 | CA2 | 423 | 0.490314972 | 5.91E-05 | 2 | 0.000120566 | 4.079010665 | 4 mo | HOM |
| **1013978665** | 1 | VISpl | 425 | 0.818478742 | 9.44E-06 | 1 | 1.15E-05 | 1.22177883 | 2 mo | HOM |
| **1013445609** | 1 | VISpl | 425 | 0.818478742 | 8.55E-05 | 6 | 0.00010442 | 7.33067298 | 4 mo | HOM |
| **1013609015** | 1 | VISpl | 425 | 0.818478742 | 0.000210381 | 11 | 0.000257039 | 13.43956713 | 4 mo | HOM |
| **1013608106** | 2 | VISpl | 425 | 0.818478742 | 0 | 0 | 0 | 0 | 4 mo | HOM |
| **1011937543** | 1 | VISpl | 425 | 0.818478742 | 0.004999684 | 15 | 0.006108508 | 18.32668245 | 8 mo | HOM |
| **1010419538** | 2 | VISpl | 425 | 0.818478742 | 5.37E-05 | 2 | 6.56E-05 | 2.44355766 | 4 mo | WILD |
| **1010419187** | 1 | VISpl | 425 | 0.818478742 | 0.004396159 | 38 | 0.005371135 | 46.42759554 | 8 mo | HOM |
| **1013978366** | 1 | VISpl | 425 | 0.818478742 | 0.000240929 | 6 | 0.000294362 | 7.33067298 | 2 mo | HOM |
| **1014416155** | 2 | VISpl | 425 | 0.818478742 | 0 | 0 | 0 | 0 | 2 mo | HOM |
| **1013978065** | 2 | VISpl | 425 | 0.818478742 | 5.36E-06 | 0 | 6.55E-06 | 0 | 2 mo | HOM |
| **1009934102** | 2 | VISpl | 425 | 0.818478742 | 9.46E-06 | 2 | 1.16E-05 | 2.44355766 | 4 mo | WILD |
| **1009322937** | 2 | VISpl | 425 | 0.818478742 | 6.58E-06 | 1 | 8.04E-06 | 1.22177883 | 8 mo | WILD |
| **1011080637** | 2 | VISpl | 425 | 0.818478742 | 0.000675347 | 33 | 0.000825124 | 40.31870139 | 8 mo | HOM |
| **1013609713** | 2 | VISpl | 425 | 0.818478742 | 1.91E-05 | 1 | 2.33E-05 | 1.22177883 | 2 mo | WILD |
| **1010418842** | 1 | VISpl | 425 | 0.818478742 | 0.001488057 | 37 | 0.001818076 | 45.20581671 | 8 mo | HOM |
| **1013288213** | 2 | VISpl | 425 | 0.818478742 | 3.43E-05 | 2 | 4.18E-05 | 2.44355766 | 4 mo | HOM |
| **1008976060** | 2 | VISpl | 425 | 0.818478742 | 0 | 0 | 0 | 0 | 8 mo | WILD |
| **1013446243** | 1 | VISpl | 425 | 0.818478742 | 0.000386119 | 13 | 0.000471752 | 15.88312479 | 4 mo | HOM |
| **1013288951** | 2 | VISpl | 425 | 0.818478742 | 0.002383506 | 6 | 0.002912117 | 7.33067298 | 4 mo | HOM |
| **1013978665** | 1 | ECT2/3 | 427 | 0.490541626 | 0 | 0 | 0 | 0 | 2 mo | HOM |
| **1013445609** | 1 | ECT2/3 | 427 | 0.490541626 | 1.49E-05 | 2 | 3.03E-05 | 4.077125961 | 4 mo | HOM |
| **1013609015** | 1 | ECT2/3 | 427 | 0.490541626 | 0.000155334 | 2 | 0.000316659 | 4.077125961 | 4 mo | HOM |

| | | | | | | | | | | |
|---|---|---|---|---|---|---|---|---|---|---|
| **1013608106** | 2 | ECT2/3 | 427 | 0.490541626 | 0 | 0 | 0 | 0 | 4 mo | HOM |
| **1011937543** | 1 | ECT2/3 | 427 | 0.490541626 | 0.0006974 | 12 | 0.001421694 | 24.46275577 | 8 mo | HOM |
| **1010419538** | 2 | ECT2/3 | 427 | 0.490541626 | 0 | 0 | 0 | 0 | 4 mo | WILD |
| **1010419187** | 1 | ECT2/3 | 427 | 0.490541626 | 0.000651696 | 27 | 0.001328524 | 55.04120047 | 8 mo | HOM |
| **1013978366** | 1 | ECT2/3 | 427 | 0.490541626 | 0 | 0 | 0 | 0 | 2 mo | HOM |
| **1014416155** | 2 | ECT2/3 | 427 | 0.490541626 | 0 | 0 | 0 | 0 | 2 mo | HOM |
| **1013978065** | 2 | ECT2/3 | 427 | 0.490541626 | 0 | 0 | 0 | 0 | 2 mo | HOM |
| **1009934102** | 2 | ECT2/3 | 427 | 0.490541626 | 0 | 0 | 0 | 0 | 4 mo | WILD |
| **1009322937** | 2 | ECT2/3 | 427 | 0.490541626 | 0 | 0 | 0 | 0 | 8 mo | WILD |
| **1011080637** | 2 | ECT2/3 | 427 | 0.490541626 | 0.000241098 | 11 | 0.000491493 | 22.42419279 | 8 mo | HOM |
| **1013609713** | 2 | ECT2/3 | 427 | 0.490541626 | 0 | 0 | 0 | 0 | 2 mo | WILD |
| **1010418842** | 1 | ECT2/3 | 427 | 0.490541626 | 0.000450633 | 25 | 0.000918644 | 50.96407451 | 8 mo | HOM |
| **1013288213** | 2 | ECT2/3 | 427 | 0.490541626 | 2.8E-05 | 2 | 5.71E-05 | 4.077125961 | 4 mo | HOM |
| **1008976060** | 2 | ECT2/3 | 427 | 0.490541626 | 0 | 0 | 0 | 0 | 8 mo | WILD |
| **1013446243** | 1 | ECT2/3 | 427 | 0.490541626 | 8.82E-05 | 4 | 0.000179795 | 8.154251922 | 4 mo | HOM |
| **1013288951** | 2 | ECT2/3 | 427 | 0.490541626 | 9.37E-05 | 4 | 0.000191043 | 8.154251922 | 4 mo | HOM |
| **1013978665** | 1 | mct | 428 | 0.008581774 | 0 | 0 | 0 | 0 | 2 mo | HOM |
| **1013445609** | 1 | mct | 428 | 0.008581774 | 0 | 0 | 0 | 0 | 4 mo | HOM |
| **1013609015** | 1 | mct | 428 | 0.008581774 | 0 | 0 | 0 | 0 | 4 mo | HOM |
| **1013608106** | 2 | mct | 428 | 0.008581774 | 0 | 0 | 0 | 0 | 4 mo | HOM |
| **1011937543** | 1 | mct | 428 | 0.008581774 | 0 | 0 | 0 | 0 | 8 mo | HOM |
| **1010419538** | 2 | mct | 428 | 0.008581774 | 0 | 0 | 0 | 0 | 4 mo | WILD |
| **1010419187** | 1 | mct | 428 | 0.008581774 | 0 | 0 | 0 | 0 | 8 mo | HOM |
| **1013978366** | 1 | mct | 428 | 0.008581774 | 0 | 0 | 0 | 0 | 2 mo | HOM |
| **1014416155** | 2 | mct | 428 | 0.008581774 | 0 | 0 | 0 | 0 | 2 mo | HOM |
| **1013978065** | 2 | mct | 428 | 0.008581774 | 0 | 0 | 0 | 0 | 2 mo | HOM |
| **1009934102** | 2 | mct | 428 | 0.008581774 | 0 | 0 | 0 | 0 | 4 mo | WILD |
| **1009322937** | 2 | mct | 428 | 0.008581774 | 0 | 0 | 0 | 0 | 8 mo | WILD |
| **1011080637** | 2 | mct | 428 | 0.008581774 | 0 | 0 | 0 | 0 | 8 mo | HOM |
| **1013609713** | 2 | mct | 428 | 0.008581774 | 0 | 0 | 0 | 0 | 2 mo | WILD |
| **1010418842** | 1 | mct | 428 | 0.008581774 | 0 | 0 | 0 | 0 | 8 mo | HOM |
| **1013288213** | 2 | mct | 428 | 0.008581774 | 0 | 0 | 0 | 0 | 4 mo | HOM |
| **1008976060** | 2 | mct | 428 | 0.008581774 | 5.13E-09 | 0 | 5.98E-07 | 0 | 8 mo | WILD |
| **1013446243** | 1 | mct | 428 | 0.008581774 | 0 | 0 | 0 | 0 | 4 mo | HOM |
| **1013288951** | 2 | mct | 428 | 0.008581774 | 0 | 0 | 0 | 0 | 4 mo | HOM |
| **1013978665** | 1 | SPVC | 429 | 1.731707197 | 0 | 0 | 0 | 0 | 2 mo | HOM |
| **1013445609** | 1 | SPVC | 429 | 1.731707197 | 0 | 0 | 0 | 0 | 4 mo | HOM |
| **1013609015** | 1 | SPVC | 429 | 1.731707197 | 0 | 0 | 0 | 0 | 4 mo | HOM |
| **1013608106** | 2 | SPVC | 429 | 1.731707197 | 0 | 0 | 0 | 0 | 4 mo | HOM |
| **1011937543** | 1 | SPVC | 429 | 1.731707197 | 0 | 0 | 0 | 0 | 8 mo | HOM |
| **1010419538** | 2 | SPVC | 429 | 1.731707197 | 0 | 0 | 0 | 0 | 4 mo | WILD |
| **1010419187** | 1 | SPVC | 429 | 1.731707197 | 9.71E-05 | 4 | 5.61E-05 | 2.309859315 | 8 mo | HOM |
| **1013978366** | 1 | SPVC | 429 | 1.731707197 | 0 | 0 | 0 | 0 | 2 mo | HOM |
| **1014416155** | 2 | SPVC | 429 | 1.731707197 | 0 | 0 | 0 | 0 | 2 mo | HOM |
| **1013978065** | 2 | SPVC | 429 | 1.731707197 | 0 | 0 | 0 | 0 | 2 mo | HOM |
| **1009934102** | 2 | SPVC | 429 | 1.731707197 | 2.84E-10 | 0 | 1.64E-10 | 0 | 4 mo | WILD |
| **1009322937** | 2 | SPVC | 429 | 1.731707197 | 1.92E-10 | 0 | 1.11E-10 | 0 | 8 mo | WILD |
| **1011080637** | 2 | SPVC | 429 | 1.731707197 | 2.13E-05 | 2 | 1.23E-05 | 1.154929658 | 8 mo | HOM |
| **1013609713** | 2 | SPVC | 429 | 1.731707197 | 0 | 0 | 0 | 0 | 2 mo | WILD |
| **1010418842** | 1 | SPVC | 429 | 1.731707197 | 0.00026367 | 14 | 0.00015226 | 8.084507604 | 8 mo | HOM |
| **1013288213** | 2 | SPVC | 429 | 1.730159581 | 0 | 0 | 0 | 0 | 4 mo | HOM |
| **1008976060** | 2 | SPVC | 429 | 1.731707197 | 1.45E-05 | 2 | 8.36E-06 | 1.154929658 | 8 mo | WILD |
| **1013446243** | 1 | SPVC | 429 | 1.731707197 | 0 | 0 | 0 | 0 | 4 mo | HOM |
| **1013288951** | 2 | SPVC | 429 | 1.731707197 | 0 | 0 | 0 | 0 | 4 mo | HOM |
| **1013978665** | 1 | RSPv2/3 | 430 | 1.043587026 | 0 | 0 | 0 | 0 | 2 mo | HOM |
| **1013445609** | 1 | RSPv2/3 | 430 | 1.043587026 | 3.36E-05 | 2 | 3.22E-05 | 1.916466908 | 4 mo | HOM |
| **1013609015** | 1 | RSPv2/3 | 430 | 1.043587026 | 9.53E-05 | 3 | 9.13E-05 | 2.874700362 | 4 mo | HOM |
| **1013608106** | 2 | RSPv2/3 | 430 | 1.043587026 | 0 | 0 | 0 | 0 | 4 mo | HOM |
| **1011937543** | 1 | RSPv2/3 | 430 | 1.043587026 | 0.001510508 | 21 | 0.001447419 | 20.12290253 | 8 mo | HOM |

| | | | | | | | | | | |
|---|---|---|---|---|---|---|---|---|---|---|
| **1010419538** | 2 | RSPv2/3 | 430 | 1.038010198 | 4.3E-07 | 0 | 4.14E-07 | 0 | 4 mo | WILD |
| **1010419187** | 1 | RSPv2/3 | 430 | 1.043587026 | 0.001359608 | 34 | 0.001302822 | 32.57993743 | 8 mo | HOM |
| **1013978366** | 1 | RSPv2/3 | 430 | 1.043587026 | 0.000312728 | 3 | 0.000299666 | 2.874700362 | 2 mo | HOM |
| **1014416155** | 2 | RSPv2/3 | 430 | 1.043587026 | 0 | 0 | 0 | 0 | 2 mo | HOM |
| **1013978065** | 2 | RSPv2/3 | 430 | 1.043587026 | 0 | 0 | 0 | 0 | 2 mo | HOM |
| **1009934102** | 2 | RSPv2/3 | 430 | 1.043587026 | 6.22E-11 | 0 | 5.96E-11 | 0 | 4 mo | WILD |
| **1009322937** | 2 | RSPv2/3 | 430 | 1.043587026 | 0 | 0 | 0 | 0 | 8 mo | WILD |
| **1011080637** | 2 | RSPv2/3 | 430 | 1.043587026 | 0.000855269 | 32 | 0.000819547 | 30.66347053 | 8 mo | HOM |
| **1013609713** | 2 | RSPv2/3 | 430 | 1.043587026 | 0 | 0 | 0 | 0 | 2 mo | WILD |
| **1010418842** | 1 | RSPv2/3 | 430 | 1.043587026 | 0.001202601 | 36 | 0.001152373 | 34.49640434 | 8 mo | HOM |
| **1013288213** | 2 | RSPv2/3 | 430 | 1.043587026 | 2.1E-06 | 0 | 2.02E-06 | 0 | 4 mo | HOM |
| **1008976060** | 2 | RSPv2/3 | 430 | 1.043587026 | 6.06E-06 | 1 | 5.8E-06 | 0.958233454 | 8 mo | WILD |
| **1013446243** | 1 | RSPv2/3 | 430 | 1.043587026 | 1.54E-05 | 1 | 1.47E-05 | 0.958233454 | 4 mo | HOM |
| **1013288951** | 2 | RSPv2/3 | 430 | 1.043587026 | 3.48E-05 | 2 | 3.34E-05 | 1.916466908 | 4 mo | HOM |
| **1013978665** | 1 | VISam5 | 433 | 0.23433909 | 0 | 0 | 0 | 0 | 2 mo | HOM |
| **1013445609** | 1 | VISam5 | 433 | 0.23433909 | 4.43E-05 | 1 | 0.000188848 | 4.267320496 | 4 mo | HOM |
| **1013609015** | 1 | VISam5 | 433 | 0.23433909 | 8.98E-05 | 2 | 0.000383287 | 8.534640991 | 4 mo | HOM |
| **1013608106** | 2 | VISam5 | 433 | 0.23433909 | 0 | 0 | 0 | 0 | 4 mo | HOM |
| **1011937543** | 1 | VISam5 | 433 | 0.23433909 | 0.000412059 | 7 | 0.00175839 | 29.87124347 | 8 mo | HOM |
| **1010419538** | 2 | VISam5 | 433 | 0.23433909 | 0 | 0 | 0 | 0 | 4 mo | WILD |
| **1010419187** | 1 | VISam5 | 433 | 0.23433909 | 0.000340181 | 16 | 0.001451663 | 68.27712793 | 8 mo | HOM |
| **1013978366** | 1 | VISam5 | 433 | 0.23433909 | 8.19E-06 | 1 | 3.5E-05 | 4.267320496 | 2 mo | HOM |
| **1014416155** | 2 | VISam5 | 433 | 0.23433909 | 0 | 0 | 0 | 0 | 2 mo | HOM |
| **1013978065** | 2 | VISam5 | 433 | 0.23433909 | 0 | 0 | 0 | 0 | 2 mo | HOM |
| **1009934102** | 2 | VISam5 | 433 | 0.23433909 | 0 | 0 | 0 | 0 | 4 mo | WILD |
| **1009322937** | 2 | VISam5 | 433 | 0.23433909 | 0 | 0 | 0 | 0 | 8 mo | WILD |
| **1011080637** | 2 | VISam5 | 433 | 0.23433909 | 0.00014488 | 13 | 0.00061825 | 55.47516644 | 8 mo | HOM |
| **1013609713** | 2 | VISam5 | 433 | 0.23433909 | 0 | 0 | 0 | 0 | 2 mo | WILD |
| **1010418842** | 1 | VISam5 | 433 | 0.23433909 | 0.000211304 | 9 | 0.000901703 | 38.40588446 | 8 mo | HOM |
| **1013288213** | 2 | VISam5 | 433 | 0.23433909 | 2.28E-05 | 2 | 9.71E-05 | 8.534640991 | 4 mo | HOM |
| **1008976060** | 2 | VISam5 | 433 | 0.23433909 | 0 | 0 | 0 | 0 | 8 mo | WILD |
| **1013446243** | 1 | VISam5 | 433 | 0.23433909 | 1.99E-05 | 2 | 8.48E-05 | 8.534640991 | 4 mo | HOM |
| **1013288951** | 2 | VISam5 | 433 | 0.23433909 | 3.24E-05 | 1 | 0.000138189 | 4.267320496 | 4 mo | HOM |
| **1013978665** | 1 | RSPd2/3 | 434 | 1.001842318 | 0 | 0 | 0 | 0 | 2 mo | HOM |
| **1013445609** | 1 | RSPd2/3 | 434 | 1.001842318 | 3.84E-05 | 3 | 3.84E-05 | 2.994483208 | 4 mo | HOM |
| **1013609015** | 1 | RSPd2/3 | 434 | 1.001842318 | 0.000169503 | 10 | 0.000169191 | 9.981610695 | 4 mo | HOM |
| **1013608106** | 2 | RSPd2/3 | 434 | 1.001842318 | 0 | 0 | 0 | 0 | 4 mo | HOM |
| **1011937543** | 1 | RSPd2/3 | 434 | 1.001842318 | 0.002138923 | 16 | 0.00213499 | 15.97057711 | 8 mo | HOM |
| **1010419538** | 2 | RSPd2/3 | 434 | 0.97926547 | 2.89E-06 | 0 | 2.95E-06 | 0 | 4 mo | WILD |
| **1010419187** | 1 | RSPd2/3 | 434 | 1.001842318 | 0.001111818 | 35 | 0.001109774 | 34.93563743 | 8 mo | HOM |
| **1013978366** | 1 | RSPd2/3 | 434 | 1.001842318 | 0 | 0 | 0 | 0 | 2 mo | HOM |
| **1014416155** | 2 | RSPd2/3 | 434 | 1.001842318 | 0 | 0 | 0 | 0 | 2 mo | HOM |
| **1013978065** | 2 | RSPd2/3 | 434 | 1.001842318 | 0 | 0 | 0 | 0 | 2 mo | HOM |
| **1009934102** | 2 | RSPd2/3 | 434 | 1.001842318 | 7.4E-07 | 0 | 7.38E-07 | 0 | 4 mo | WILD |
| **1009322937** | 2 | RSPd2/3 | 434 | 1.001842318 | 6.76E-08 | 0 | 6.75E-08 | 0 | 8 mo | WILD |
| **1011080637** | 2 | RSPd2/3 | 434 | 1.001842318 | 0.000870609 | 25 | 0.000869008 | 24.95402674 | 8 mo | HOM |
| **1013609713** | 2 | RSPd2/3 | 434 | 1.001842318 | 0 | 0 | 0 | 0 | 2 mo | WILD |
| **1010418842** | 1 | RSPd2/3 | 434 | 1.001842318 | 0.000858785 | 37 | 0.000857205 | 36.93195957 | 8 mo | HOM |
| **1013288213** | 2 | RSPd2/3 | 434 | 1.001842318 | 2.15E-05 | 2 | 2.14E-05 | 1.996322139 | 4 mo | HOM |
| **1008976060** | 2 | RSPd2/3 | 434 | 1.001842318 | 1.46E-06 | 0 | 1.46E-06 | 0 | 8 mo | WILD |
| **1013446243** | 1 | RSPd2/3 | 434 | 1.001842318 | 8.13E-05 | 3 | 8.12E-05 | 2.994483208 | 4 mo | HOM |
| **1013288951** | 2 | RSPd2/3 | 434 | 1.001842318 | 0.000178951 | 7 | 0.000178622 | 6.987127486 | 4 mo | HOM |
| **1013978665** | 1 | fx | 436 | 0.277767202 | 4.87E-09 | 0 | 1.75E-08 | 0 | 2 mo | HOM |
| **1013445609** | 1 | fx | 436 | 0.277767202 | 5.59E-08 | 0 | 2.01E-07 | 0 | 4 mo | HOM |
| **1013609015** | 1 | fx | 436 | 0.277767202 | 4.18E-07 | 0 | 1.51E-06 | 0 | 4 mo | HOM |
| **1013608106** | 2 | fx | 436 | 0.277767202 | 0 | 0 | 0 | 0 | 4 mo | HOM |
| **1011937543** | 1 | fx | 436 | 0.277767202 | 4.42E-05 | 1 | 0.000159167 | 3.600137072 | 8 mo | HOM |
| **1010419538** | 2 | fx | 436 | 0.277767202 | 0 | 0 | 0 | 0 | 4 mo | WILD |
| **1010419187** | 1 | fx | 436 | 0.277767202 | 4.29E-06 | 0 | 1.54E-05 | 0 | 8 mo | HOM |

| | | | | | | | | | | |
|---|---|---|---|---|---|---|---|---|---|---|
| **1013978366** | 1 | fx | 436 | 0.277767202 | 4.25E-08 | 0 | 1.53E-07 | 0 | 2 mo | HOM |
| **1014416155** | 2 | fx | 436 | 0.277767202 | 0 | 0 | 0 | 0 | 2 mo | HOM |
| **1013978065** | 2 | fx | 436 | 0.277767202 | 0 | 0 | 0 | 0 | 2 mo | HOM |
| **1009934102** | 2 | fx | 436 | 0.277767202 | 3.23E-06 | 0 | 1.16E-05 | 0 | 4 mo | WILD |
| **1009322937** | 2 | fx | 436 | 0.277767202 | 0 | 0 | 0 | 0 | 8 mo | WILD |
| **1011080637** | 2 | fx | 436 | 0.277767202 | 6.07E-05 | 0 | 0.000218648 | 0 | 8 mo | HOM |
| **1013609713** | 2 | fx | 436 | 0.277767202 | 1.56E-06 | 0 | 5.6E-06 | 0 | 2 mo | WILD |
| **1010418842** | 1 | fx | 436 | 0.277767202 | 0.0001054 | 4 | 0.000379455 | 14.40054829 | 8 mo | HOM |
| **1013288213** | 2 | fx | 436 | 0.277767202 | 1.45E-06 | 0 | 5.23E-06 | 0 | 4 mo | HOM |
| **1008976060** | 2 | fx | 436 | 0.277767202 | 2.22E-05 | 0 | 7.99E-05 | 0 | 8 mo | WILD |
| **1013446243** | 1 | fx | 436 | 0.277767202 | 2.38E-06 | 0 | 8.55E-06 | 0 | 4 mo | HOM |
| **1013288951** | 2 | fx | 436 | 0.277767202 | 4.93E-05 | 0 | 0.000177619 | 0 | 4 mo | HOM |
| **1013978665** | 1 | SPVI | 437 | 1.904253523 | 0 | 0 | 0 | 0 | 2 mo | HOM |
| **1013445609** | 1 | SPVI | 437 | 1.904253523 | 0 | 0 | 0 | 0 | 4 mo | HOM |
| **1013609015** | 1 | SPVI | 437 | 1.904253523 | 0 | 0 | 0 | 0 | 4 mo | HOM |
| **1013608106** | 2 | SPVI | 437 | 1.904253523 | 0 | 0 | 0 | 0 | 4 mo | HOM |
| **1011937543** | 1 | SPVI | 437 | 1.904253523 | 0.001232411 | 3 | 0.000647189 | 1.57542048 | 8 mo | HOM |
| **1010419538** | 2 | SPVI | 437 | 1.904253523 | 0 | 0 | 0 | 0 | 4 mo | WILD |
| **1010419187** | 1 | SPVI | 437 | 1.904253523 | 0 | 0 | 0 | 0 | 8 mo | HOM |
| **1013978366** | 1 | SPVI | 437 | 1.904253523 | 0 | 0 | 0 | 0 | 2 mo | HOM |
| **1014416155** | 2 | SPVI | 437 | 1.904253523 | 0 | 0 | 0 | 0 | 2 mo | HOM |
| **1013978065** | 2 | SPVI | 437 | 1.904253523 | 0 | 0 | 0 | 0 | 2 mo | HOM |
| **1009934102** | 2 | SPVI | 437 | 1.904253523 | 0 | 0 | 0 | 0 | 4 mo | WILD |
| **1009322937** | 2 | SPVI | 437 | 1.904253523 | 0 | 0 | 0 | 0 | 8 mo | WILD |
| **1011080637** | 2 | SPVI | 437 | 1.904253523 | 0 | 0 | 0 | 0 | 8 mo | HOM |
| **1013609713** | 2 | SPVI | 437 | 1.904253523 | 0 | 0 | 0 | 0 | 2 mo | WILD |
| **1010418842** | 1 | SPVI | 437 | 1.904253523 | 8.77E-05 | 6 | 4.61E-05 | 3.15084096 | 8 mo | HOM |
| **1013288213** | 2 | SPVI | 437 | 1.904253523 | 0 | 0 | 0 | 0 | 4 mo | HOM |
| **1008976060** | 2 | SPVI | 437 | 1.904253523 | 2.16E-05 | 3 | 1.13E-05 | 1.57542048 | 8 mo | WILD |
| **1013446243** | 1 | SPVI | 437 | 1.904253523 | 0 | 0 | 0 | 0 | 4 mo | HOM |
| **1013288951** | 2 | SPVI | 437 | 1.904253523 | 0 | 0 | 0 | 0 | 4 mo | HOM |
| **1013978665** | 1 | ORBI6a | 440 | 0.495414618 | 0 | 0 | 0 | 0 | 2 mo | HOM |
| **1013445609** | 1 | ORBI6a | 440 | 0.495414618 | 1.62E-07 | 0 | 3.28E-07 | 0 | 4 mo | HOM |
| **1013609015** | 1 | ORBI6a | 440 | 0.495414618 | 1.97E-05 | 1 | 3.97E-05 | 2.018511292 | 4 mo | HOM |
| **1013608106** | 2 | ORBI6a | 440 | 0.495414618 | 0 | 0 | 0 | 0 | 4 mo | HOM |
| **1011937543** | 1 | ORBI6a | 440 | 0.495414618 | 0.000342395 | 8 | 0.000691129 | 16.14809034 | 8 mo | HOM |
| **1010419538** | 2 | ORBI6a | 440 | 0.495414618 | 0 | 0 | 0 | 0 | 4 mo | WILD |
| **1010419187** | 1 | ORBI6a | 440 | 0.495414618 | 0.000211309 | 14 | 0.000426529 | 28.25915809 | 8 mo | HOM |
| **1013978366** | 1 | ORBI6a | 440 | 0.495414618 | 0 | 0 | 0 | 0 | 2 mo | HOM |
| **1014416155** | 2 | ORBI6a | 440 | 0.495414618 | 0 | 0 | 0 | 0 | 2 mo | HOM |
| **1013978065** | 2 | ORBI6a | 440 | 0.495414618 | 0 | 0 | 0 | 0 | 2 mo | HOM |
| **1009934102** | 2 | ORBI6a | 440 | 0.495414618 | 0 | 0 | 0 | 0 | 4 mo | WILD |
| **1009322937** | 2 | ORBI6a | 440 | 0.495414618 | 0 | 0 | 0 | 0 | 8 mo | WILD |
| **1011080637** | 2 | ORBI6a | 440 | 0.495414618 | 0.000220674 | 10 | 0.000445433 | 20.18511292 | 8 mo | HOM |
| **1013609713** | 2 | ORBI6a | 440 | 0.495414618 | 0 | 0 | 0 | 0 | 2 mo | WILD |
| **1010418842** | 1 | ORBI6a | 440 | 0.495414618 | 0.000289282 | 12 | 0.00058392 | 24.22213551 | 8 mo | HOM |
| **1013288213** | 2 | ORBI6a | 440 | 0.495414618 | 0 | 0 | 0 | 0 | 4 mo | HOM |
| **1008976060** | 2 | ORBI6a | 440 | 0.495414618 | 0 | 0 | 0 | 0 | 8 mo | WILD |
| **1013446243** | 1 | ORBI6a | 440 | 0.495414618 | 9.3E-06 | 1 | 1.88E-05 | 2.018511292 | 4 mo | HOM |
| **1013288951** | 2 | ORBI6a | 440 | 0.495414618 | 8.24E-06 | 0 | 1.66E-05 | 0 | 4 mo | HOM |
| **1013978665** | 1 | VISam6b | 441 | 0.024340096 | 0 | 0 | 0 | 0 | 2 mo | HOM |
| **1013445609** | 1 | VISam6b | 441 | 0.024340096 | 3.66E-09 | 0 | 1.51E-07 | 0 | 4 mo | HOM |
| **1013609015** | 1 | VISam6b | 441 | 0.024340096 | 1.03E-05 | 1 | 0.000424356 | 41.08447256 | 4 mo | HOM |
| **1013608106** | 2 | VISam6b | 441 | 0.024340096 | 0 | 0 | 0 | 0 | 4 mo | HOM |
| **1011937543** | 1 | VISam6b | 441 | 0.024340096 | 5.38E-06 | 0 | 0.000221132 | 0 | 8 mo | HOM |
| **1010419538** | 2 | VISam6b | 441 | 0.024340096 | 0 | 0 | 0 | 0 | 4 mo | WILD |
| **1010419187** | 1 | VISam6b | 441 | 0.024340096 | 2.81E-05 | 1 | 0.00115641 | 41.08447256 | 8 mo | HOM |
| **1013978366** | 1 | VISam6b | 441 | 0.024340096 | 0 | 0 | 0 | 0 | 2 mo | HOM |
| **1014416155** | 2 | VISam6b | 441 | 0.024340096 | 0 | 0 | 0 | 0 | 2 mo | HOM |

| | | | | | | | | | | |
|---|---|---|---|---|---|---|---|---|---|---|
| **1013978065** | 2 | VISam6b | 441 | 0.024340096 | 0 | 0 | 0 | 0 | 2 mo | HOM |
| **1009934102** | 2 | VISam6b | 441 | 0.024340096 | 0 | 0 | 0 | 0 | 4 mo | WILD |
| **1009322937** | 2 | VISam6b | 441 | 0.024340096 | 0 | 0 | 0 | 0 | 8 mo | WILD |
| **1011080637** | 2 | VISam6b | 441 | 0.024340096 | 6.36E-07 | 0 | 2.61E-05 | 0 | 8 mo | HOM |
| **1013609713** | 2 | VISam6b | 441 | 0.024340096 | 0 | 0 | 0 | 0 | 2 mo | WILD |
| **1010418842** | 1 | VISam6b | 441 | 0.024340096 | 2.16E-05 | 0 | 0.00088607 | 0 | 8 mo | HOM |
| **1013288213** | 2 | VISam6b | 441 | 0.024340096 | 0 | 0 | 0 | 0 | 4 mo | HOM |
| **1008976060** | 2 | VISam6b | 441 | 0.024340096 | 0 | 0 | 0 | 0 | 8 mo | WILD |
| **1013446243** | 1 | VISam6b | 441 | 0.024340096 | 0 | 0 | 0 | 0 | 4 mo | HOM |
| **1013288951** | 2 | VISam6b | 441 | 0.024340096 | 0 | 0 | 0 | 0 | 4 mo | HOM |
| **1013978665** | 1 | RSPd1 | 442 | 0.85842465 | 0.000111698 | 2 | 0.00013012 | 2.32984922 | 2 mo | HOM |
| **1013445609** | 1 | RSPd1 | 442 | 0.85842465 | 3.05E-05 | 3 | 3.55E-05 | 3.494773829 | 4 mo | HOM |
| **1013609015** | 1 | RSPd1 | 442 | 0.85842465 | 0.000245303 | 12 | 0.00028576 | 13.97909532 | 4 mo | HOM |
| **1013608106** | 2 | RSPd1 | 442 | 0.85842465 | 0 | 0 | 0 | 0 | 4 mo | HOM |
| **1011937543** | 1 | RSPd1 | 442 | 0.85842465 | 0.001723805 | 14 | 0.002008103 | 16.30894454 | 8 mo | HOM |
| **1010419538** | 2 | RSPd1 | 442 | 0.839070826 | 0.000369134 | 9 | 0.000439931 | 10.72615055 | 4 mo | WILD |
| **1010419187** | 1 | RSPd1 | 442 | 0.85842465 | 0.006108429 | 35 | 0.00711586 | 40.77236134 | 8 mo | HOM |
| **1013978366** | 1 | RSPd1 | 442 | 0.85842465 | 0.000170251 | 2 | 0.000198329 | 2.32984922 | 2 mo | HOM |
| **1014416155** | 2 | RSPd1 | 442 | 0.85842465 | 0 | 0 | 0 | 0 | 2 mo | HOM |
| **1013978065** | 2 | RSPd1 | 442 | 0.85842465 | 0 | 0 | 0 | 0 | 2 mo | HOM |
| **1009934102** | 2 | RSPd1 | 442 | 0.85842465 | 6.34E-05 | 4 | 7.38E-05 | 4.659698439 | 4 mo | WILD |
| **1009322937** | 2 | RSPd1 | 442 | 0.85842465 | 5.08E-05 | 4 | 5.92E-05 | 4.659698439 | 8 mo | WILD |
| **1011080637** | 2 | RSPd1 | 442 | 0.85842465 | 0.001785692 | 36 | 0.002080196 | 41.93728595 | 8 mo | HOM |
| **1013609713** | 2 | RSPd1 | 442 | 0.85842465 | 0 | 0 | 0 | 0 | 2 mo | WILD |
| **1010418842** | 1 | RSPd1 | 442 | 0.85842465 | 0.001016693 | 38 | 0.001184371 | 44.26713517 | 8 mo | HOM |
| **1013288213** | 2 | RSPd1 | 442 | 0.85842465 | 1.9E-05 | 2 | 2.22E-05 | 2.32984922 | 4 mo | HOM |
| **1008976060** | 2 | RSPd1 | 442 | 0.85842465 | 0.000129749 | 9 | 0.000151148 | 10.48432149 | 8 mo | WILD |
| **1013446243** | 1 | RSPd1 | 442 | 0.85842465 | 0.000198072 | 10 | 0.000230739 | 11.6492461 | 4 mo | HOM |
| **1013288951** | 2 | RSPd1 | 442 | 0.85842465 | 0.002242883 | 10 | 0.00261279 | 11.6492461 | 4 mo | HOM |
| **1013978665** | 1 | dhc | 443 | 0.982622402 | 0 | 0 | 0 | 0 | 2 mo | HOM |
| **1013445609** | 1 | dhc | 443 | 0.982622402 | 8.5E-06 | 1 | 8.65E-06 | 1.01768492 | 4 mo | HOM |
| **1013609015** | 1 | dhc | 443 | 0.982622402 | 1.58E-05 | 2 | 1.61E-05 | 2.03536984 | 4 mo | HOM |
| **1013608106** | 2 | dhc | 443 | 0.982622402 | 0 | 0 | 0 | 0 | 4 mo | HOM |
| **1011937543** | 1 | dhc | 443 | 0.982622402 | 0.001203975 | 11 | 0.001225267 | 11.19453412 | 8 mo | HOM |
| **1010419538** | 2 | dhc | 443 | 0.982622402 | 0 | 0 | 0 | 0 | 4 mo | WILD |
| **1010419187** | 1 | dhc | 443 | 0.982622402 | 0.000365113 | 21 | 0.00037157 | 21.37138332 | 8 mo | HOM |
| **1013978366** | 1 | dhc | 443 | 0.982622402 | 0 | 0 | 0 | 0 | 2 mo | HOM |
| **1014416155** | 2 | dhc | 443 | 0.982622402 | 0 | 0 | 0 | 0 | 2 mo | HOM |
| **1013978065** | 2 | dhc | 443 | 0.982622402 | 0 | 0 | 0 | 0 | 2 mo | HOM |
| **1009934102** | 2 | dhc | 443 | 0.982622402 | 0 | 0 | 0 | 0 | 4 mo | WILD |
| **1009322937** | 2 | dhc | 443 | 0.982622402 | 0 | 0 | 0 | 0 | 8 mo | WILD |
| **1011080637** | 2 | dhc | 443 | 0.982622402 | 3E-05 | 2 | 3.05E-05 | 2.03536984 | 8 mo | HOM |
| **1013609713** | 2 | dhc | 443 | 0.982622402 | 0 | 0 | 0 | 0 | 2 mo | WILD |
| **1010418842** | 1 | dhc | 443 | 0.982622402 | 0.001198954 | 28 | 0.001220157 | 28.49517776 | 8 mo | HOM |
| **1013288213** | 2 | dhc | 443 | 0.982622402 | 0 | 0 | 0 | 0 | 4 mo | HOM |
| **1008976060** | 2 | dhc | 443 | 0.982622402 | 3.66E-06 | 1 | 3.72E-06 | 1.01768492 | 8 mo | WILD |
| **1013446243** | 1 | dhc | 443 | 0.982622402 | 4.71E-06 | 1 | 4.79E-06 | 1.01768492 | 4 mo | HOM |
| **1013288951** | 2 | dhc | 443 | 0.982622402 | 1.01E-05 | 1 | 1.03E-05 | 1.01768492 | 4 mo | HOM |
| **1013978665** | 1 | MED | 444 | 2.078437802 | 0 | 0 | 0 | 0 | 2 mo | HOM |
| **1013445609** | 1 | MED | 444 | 2.078437802 | 7.7E-08 | 0 | 3.7E-08 | 0 | 4 mo | HOM |
| **1013609015** | 1 | MED | 444 | 2.078437802 | 3.41E-05 | 0 | 1.64E-05 | 0 | 4 mo | HOM |
| **1013608106** | 2 | MED | 444 | 2.078437802 | 0 | 0 | 0 | 0 | 4 mo | HOM |
| **1011937543** | 1 | MED | 444 | 2.078437802 | 0.000897484 | 23 | 0.000431807 | 11.0660035 | 8 mo | HOM |
| **1010419538** | 2 | MED | 444 | 2.078437802 | 2.31E-05 | 1 | 1.11E-05 | 0.481130587 | 4 mo | WILD |
| **1010419187** | 1 | MED | 444 | 2.078437802 | 0.000767367 | 20 | 0.000369204 | 9.622611743 | 8 mo | HOM |
| **1013978366** | 1 | MED | 444 | 2.078437802 | 0 | 0 | 0 | 0 | 2 mo | HOM |
| **1014416155** | 2 | MED | 444 | 2.078437802 | 0 | 0 | 0 | 0 | 2 mo | HOM |
| **1013978065** | 2 | MED | 444 | 2.078437802 | 0 | 0 | 0 | 0 | 2 mo | HOM |
| **1009934102** | 2 | MED | 444 | 2.078437802 | 0 | 0 | 0 | 0 | 4 mo | WILD |

| | | | | | | | | | | |
|---|---|---|---|---|---|---|---|---|---|---|
| **1009322937** | 2 | MED | 444 | 2.078437802 | 2.78E-05 | 3 | 1.34E-05 | 1.443391761 | 8 mo | WILD |
| **1011080637** | 2 | MED | 444 | 2.078437802 | 0.000769102 | 24 | 0.000370039 | 11.54713409 | 8 mo | HOM |
| **1013609713** | 2 | MED | 444 | 2.078437802 | 0 | 0 | 0 | 0 | 2 mo | WILD |
| **1010418842** | 1 | MED | 444 | 2.078437802 | 0.001230992 | 39 | 0.000592268 | 18.7640929 | 8 mo | HOM |
| **1013288213** | 2 | MED | 444 | 2.078437802 | 0 | 0 | 0 | 0 | 4 mo | HOM |
| **1008976060** | 2 | MED | 444 | 2.078437802 | 3.43E-05 | 5 | 1.65E-05 | 2.405652936 | 8 mo | WILD |
| **1013446243** | 1 | MED | 444 | 2.078437802 | 4.73E-06 | 0 | 2.27E-06 | 0 | 4 mo | HOM |
| **1013288951** | 2 | MED | 444 | 2.078437802 | 6.03E-06 | 1 | 2.9E-06 | 0.481130587 | 4 mo | HOM |
| **1013978665** | 1 | SPVO | 445 | 1.02648532 | 0 | 0 | 0 | 0 | 2 mo | HOM |
| **1013445609** | 1 | SPVO | 445 | 1.02648532 | 0 | 0 | 0 | 0 | 4 mo | HOM |
| **1013609015** | 1 | SPVO | 445 | 1.02648532 | 0 | 0 | 0 | 0 | 4 mo | HOM |
| **1013608106** | 2 | SPVO | 445 | 1.02648532 | 0 | 0 | 0 | 0 | 4 mo | HOM |
| **1011937543** | 1 | SPVO | 445 | 1.02648532 | 0.004097009 | 13 | 0.003991298 | 12.66457469 | 8 mo | HOM |
| **1010419538** | 2 | SPVO | 445 | 1.02648532 | 0 | 0 | 0 | 0 | 4 mo | WILD |
| **1010419187** | 1 | SPVO | 445 | 1.02648532 | 0 | 0 | 0 | 0 | 8 mo | HOM |
| **1013978366** | 1 | SPVO | 445 | 1.02648532 | 0 | 0 | 0 | 0 | 2 mo | HOM |
| **1014416155** | 2 | SPVO | 445 | 1.02648532 | 0 | 0 | 0 | 0 | 2 mo | HOM |
| **1013978065** | 2 | SPVO | 445 | 1.02648532 | 0 | 0 | 0 | 0 | 2 mo | HOM |
| **1009934102** | 2 | SPVO | 445 | 1.02648532 | 0 | 0 | 0 | 0 | 4 mo | WILD |
| **1009322937** | 2 | SPVO | 445 | 1.02648532 | 0 | 0 | 0 | 0 | 8 mo | WILD |
| **1011080637** | 2 | SPVO | 445 | 1.02648532 | 5.99E-06 | 1 | 5.83E-06 | 0.974198053 | 8 mo | HOM |
| **1013609713** | 2 | SPVO | 445 | 1.02648532 | 0 | 0 | 0 | 0 | 2 mo | WILD |
| **1010418842** | 1 | SPVO | 445 | 1.02648532 | 1.86E-05 | 2 | 1.81E-05 | 1.948396106 | 8 mo | HOM |
| **1013288213** | 2 | SPVO | 445 | 1.02648532 | 0 | 0 | 0 | 0 | 4 mo | HOM |
| **1008976060** | 2 | SPVO | 445 | 1.02648532 | 9.1E-06 | 2 | 8.87E-06 | 1.948396106 | 8 mo | WILD |
| **1013446243** | 1 | SPVO | 445 | 1.02648532 | 0 | 0 | 0 | 0 | 4 mo | HOM |
| **1013288951** | 2 | SPVO | 445 | 1.02648532 | 0 | 0 | 0 | 0 | 4 mo | HOM |
| **1013978665** | 1 | ORBI1 | 448 | 0.386764997 | 2.91E-05 | 4 | 7.53E-05 | 10.34219754 | 2 mo | HOM |
| **1013445609** | 1 | ORBI1 | 448 | 0.386764997 | 0 | 0 | 0 | 0 | 4 mo | HOM |
| **1013609015** | 1 | ORBI1 | 448 | 0.386764997 | 3.43E-05 | 2 | 8.87E-05 | 5.171098772 | 4 mo | HOM |
| **1013608106** | 2 | ORBI1 | 448 | 0.386764997 | 0 | 0 | 0 | 0 | 4 mo | HOM |
| **1011937543** | 1 | ORBI1 | 448 | 0.386764997 | 0.000439062 | 7 | 0.001135217 | 18.0988457 | 8 mo | HOM |
| **1010419538** | 2 | ORBI1 | 448 | 0.386764997 | 0 | 0 | 0 | 0 | 4 mo | WILD |
| **1010419187** | 1 | ORBI1 | 448 | 0.386764997 | 0.000363579 | 10 | 0.000940052 | 25.85549386 | 8 mo | HOM |
| **1013978366** | 1 | ORBI1 | 448 | 0.386764997 | 2.11E-05 | 0 | 5.47E-05 | 0 | 2 mo | HOM |
| **1014416155** | 2 | ORBI1 | 448 | 0.386764997 | 0 | 0 | 0 | 0 | 2 mo | HOM |
| **1013978065** | 2 | ORBI1 | 448 | 0.386764997 | 0 | 0 | 0 | 0 | 2 mo | HOM |
| **1009934102** | 2 | ORBI1 | 448 | 0.386764997 | 1.21E-05 | 0 | 3.12E-05 | 0 | 4 mo | WILD |
| **1009322937** | 2 | ORBI1 | 448 | 0.386764997 | 9.51E-05 | 5 | 0.000245938 | 12.92774693 | 8 mo | WILD |
| **1011080637** | 2 | ORBI1 | 448 | 0.386764997 | 7.9E-05 | 5 | 0.000204314 | 12.92774693 | 8 mo | HOM |
| **1013609713** | 2 | ORBI1 | 448 | 0.386764997 | 0 | 0 | 0 | 0 | 2 mo | WILD |
| **1010418842** | 1 | ORBI1 | 448 | 0.386764997 | 0.000239461 | 10 | 0.000619138 | 25.85549386 | 8 mo | HOM |
| **1013288213** | 2 | ORBI1 | 448 | 0.386764997 | 0.000740593 | 3 | 0.001914839 | 7.756648158 | 4 mo | HOM |
| **1008976060** | 2 | ORBI1 | 448 | 0.386764997 | 2.04E-05 | 2 | 5.27E-05 | 5.171098772 | 8 mo | WILD |
| **1013446243** | 1 | ORBI1 | 448 | 0.386764997 | 1.78E-06 | 0 | 4.61E-06 | 0 | 4 mo | HOM |
| **1013288951** | 2 | ORBI1 | 448 | 0.386764997 | 0.000409087 | 6 | 0.001057714 | 15.51329632 | 4 mo | HOM |
| **1013978665** | 1 | vhc | 449 | 0.066659678 | 0 | 0 | 0 | 0 | 2 mo | HOM |
| **1013445609** | 1 | vhc | 449 | 0.066659678 | 5.6E-26 | 0 | 8.4E-25 | 0 | 4 mo | HOM |
| **1013609015** | 1 | vhc | 449 | 0.066659678 | 2.51E-06 | 0 | 3.77E-05 | 0 | 4 mo | HOM |
| **1013608106** | 2 | vhc | 449 | 0.066659678 | 0 | 0 | 0 | 0 | 4 mo | HOM |
| **1011937543** | 1 | vhc | 449 | 0.066659678 | 1.36E-05 | 1 | 0.000204513 | 15.00157252 | 8 mo | HOM |
| **1010419538** | 2 | vhc | 449 | 0.066659678 | 0 | 0 | 0 | 0 | 4 mo | WILD |
| **1010419187** | 1 | vhc | 449 | 0.066659678 | 4.84E-05 | 0 | 0.000726187 | 0 | 8 mo | HOM |
| **1013978366** | 1 | vhc | 449 | 0.066659678 | 7.61E-08 | 0 | 1.14E-06 | 0 | 2 mo | HOM |
| **1014416155** | 2 | vhc | 449 | 0.066659678 | 0 | 0 | 0 | 0 | 2 mo | HOM |
| **1013978065** | 2 | vhc | 449 | 0.066659678 | 0 | 0 | 0 | 0 | 2 mo | HOM |
| **1009934102** | 2 | vhc | 449 | 0.066659678 | 4.93E-06 | 0 | 7.39E-05 | 0 | 4 mo | WILD |
| **1009322937** | 2 | vhc | 449 | 0.066659678 | 3.28E-07 | 0 | 4.92E-06 | 0 | 8 mo | WILD |
| **1011080637** | 2 | vhc | 449 | 0.066659678 | 1.39E-05 | 1 | 0.000209018 | 15.00157252 | 8 mo | HOM |

| | | | | | | | | | | |
|---|---|---|---|---|---|---|---|---|---|---|
| **1013609713** | 2 | vhc | 449 | 0.066659678 | 0 | 0 | 0 | 0 | 2 mo | WILD |
| **1010418842** | 1 | vhc | 449 | 0.066659678 | 0.000100932 | 1 | 0.001514143 | 15.00157252 | 8 mo | HOM |
| **1013288213** | 2 | vhc | 449 | 0.066659678 | 0 | 0 | 0 | 0 | 4 mo | HOM |
| **1008976060** | 2 | vhc | 449 | 0.066659678 | 3.12E-07 | 0 | 4.68E-06 | 0 | 8 mo | WILD |
| **1013446243** | 1 | vhc | 449 | 0.066659678 | 8.04E-06 | 0 | 0.000120659 | 0 | 4 mo | HOM |
| **1013288951** | 2 | vhc | 449 | 0.066659678 | 1.29E-05 | 0 | 0.000193475 | 0 | 4 mo | HOM |
| **1013978665** | 1 | SSp-ul1 | 450 | 0.473718045 | 0 | 0 | 0 | 0 | 2 mo | HOM |
| **1013445609** | 1 | SSp-ul1 | 450 | 0.473718045 | 0.000189898 | 5 | 0.000400867 | 10.55480165 | 4 mo | HOM |
| **1013609015** | 1 | SSp-ul1 | 450 | 0.473718045 | 0.000244273 | 8 | 0.00051565 | 16.88768264 | 4 mo | HOM |
| **1013608106** | 2 | SSp-ul1 | 450 | 0.473718045 | 0 | 0 | 0 | 0 | 4 mo | HOM |
| **1011937543** | 1 | SSp-ul1 | 450 | 0.473718045 | 0.002157969 | 20 | 0.004555388 | 42.21920659 | 8 mo | HOM |
| **1010419538** | 2 | SSp-ul1 | 450 | 0.473718045 | 6.36E-06 | 1 | 1.34E-05 | 2.11096033 | 4 mo | WILD |
| **1010419187** | 1 | SSp-ul1 | 450 | 0.473718045 | 0.001338631 | 30 | 0.002825797 | 63.32880989 | 8 mo | HOM |
| **1013978366** | 1 | SSp-ul1 | 450 | 0.473718045 | 0 | 0 | 0 | 0 | 2 mo | HOM |
| **1014416155** | 2 | SSp-ul1 | 450 | 0.473718045 | 0 | 0 | 0 | 0 | 2 mo | HOM |
| **1013978065** | 2 | SSp-ul1 | 450 | 0.473718045 | 0 | 0 | 0 | 0 | 2 mo | HOM |
| **1009934102** | 2 | SSp-ul1 | 450 | 0.473718045 | 7.57E-06 | 1 | 1.6E-05 | 2.11096033 | 4 mo | WILD |
| **1009322937** | 2 | SSp-ul1 | 450 | 0.473718045 | 0 | 0 | 0 | 0 | 8 mo | WILD |
| **1011080637** | 2 | SSp-ul1 | 450 | 0.473718045 | 0.000370309 | 13 | 0.000781708 | 27.44248429 | 8 mo | HOM |
| **1013609713** | 2 | SSp-ul1 | 450 | 0.473718045 | 4.5E-06 | 0 | 9.49E-06 | 0 | 2 mo | WILD |
| **1010418842** | 1 | SSp-ul1 | 450 | 0.473718045 | 0.001404188 | 34 | 0.002964186 | 71.77265121 | 8 mo | HOM |
| **1013288213** | 2 | SSp-ul1 | 450 | 0.473718045 | 2.45E-05 | 2 | 5.18E-05 | 4.221920659 | 4 mo | HOM |
| **1008976060** | 2 | SSp-ul1 | 450 | 0.473718045 | 0.000531691 | 12 | 0.001122378 | 25.33152396 | 8 mo | WILD |
| **1013446243** | 1 | SSp-ul1 | 450 | 0.473718045 | 0.000291992 | 10 | 0.000616384 | 21.1096033 | 4 mo | HOM |
| **1013288951** | 2 | SSp-ul1 | 450 | 0.473718045 | 0.000220727 | 10 | 0.000465947 | 21.1096033 | 4 mo | HOM |
| **1013978665** | 1 | BLAv | 451 | 0.432113438 | 0 | 0 | 0 | 0 | 2 mo | HOM |
| **1013445609** | 1 | BLAv | 451 | 0.432113438 | 6.21E-05 | 5 | 0.000143779 | 11.57103565 | 4 mo | HOM |
| **1013609015** | 1 | BLAv | 451 | 0.432113438 | 0.000139219 | 7 | 0.000322181 | 16.19944991 | 4 mo | HOM |
| **1013608106** | 2 | BLAv | 451 | 0.432113438 | 0 | 0 | 0 | 0 | 4 mo | HOM |
| **1011937543** | 1 | BLAv | 451 | 0.432113438 | 0.00113523 | 17 | 0.002627159 | 39.34152121 | 8 mo | HOM |
| **1010419538** | 2 | BLAv | 451 | 0.432113438 | 0 | 0 | 0 | 0 | 4 mo | WILD |
| **1010419187** | 1 | BLAv | 451 | 0.432113438 | 0.000516092 | 24 | 0.001194343 | 55.54097111 | 8 mo | HOM |
| **1013978366** | 1 | BLAv | 451 | 0.432113438 | 0 | 0 | 0 | 0 | 2 mo | HOM |
| **1014416155** | 2 | BLAv | 451 | 0.432113438 | 0 | 0 | 0 | 0 | 2 mo | HOM |
| **1013978065** | 2 | BLAv | 451 | 0.432113438 | 0 | 0 | 0 | 0 | 2 mo | HOM |
| **1009934102** | 2 | BLAv | 451 | 0.432113438 | 0 | 0 | 0 | 0 | 4 mo | WILD |
| **1009322937** | 2 | BLAv | 451 | 0.432113438 | 0 | 0 | 0 | 0 | 8 mo | WILD |
| **1011080637** | 2 | BLAv | 451 | 0.432113438 | 0.000323725 | 22 | 0.000749167 | 50.91255685 | 8 mo | HOM |
| **1013609713** | 2 | BLAv | 451 | 0.432113438 | 0 | 0 | 0 | 0 | 2 mo | WILD |
| **1010418842** | 1 | BLAv | 451 | 0.432113438 | 0.000928295 | 30 | 0.002148266 | 69.42621389 | 8 mo | HOM |
| **1013288213** | 2 | BLAv | 451 | 0.432113438 | 3.79E-06 | 0 | 8.76E-06 | 0 | 4 mo | HOM |
| **1008976060** | 2 | BLAv | 451 | 0.432113438 | 7.49E-06 | 1 | 1.73E-05 | 2.31420713 | 8 mo | WILD |
| **1013446243** | 1 | BLAv | 451 | 0.432113438 | 8.28E-05 | 4 | 0.00019163 | 9.256828519 | 4 mo | HOM |
| **1013288951** | 2 | BLAv | 451 | 0.432113438 | 0.000186966 | 8 | 0.000432677 | 18.51365704 | 4 mo | HOM |
| **1013978665** | 1 | MEPO | 452 | 0.041476858 | 4.19E-09 | 0 | 1.01E-07 | 0 | 2 mo | HOM |
| **1013445609** | 1 | MEPO | 452 | 0.041476858 | 3.6E-05 | 2 | 0.000868066 | 48.21965964 | 4 mo | HOM |
| **1013609015** | 1 | MEPO | 452 | 0.041476858 | 0.000506729 | 2 | 0.012217141 | 48.21965964 | 4 mo | HOM |
| **1013608106** | 2 | MEPO | 452 | 0.041476858 | 0 | 0 | 0 | 0 | 4 mo | HOM |
| **1011937543** | 1 | MEPO | 452 | 0.041476858 | 1.02E-05 | 1 | 0.000245081 | 24.10982982 | 8 mo | HOM |
| **1010419538** | 2 | MEPO | 452 | 0.041476858 | 0 | 0 | 0 | 0 | 4 mo | WILD |
| **1010419187** | 1 | MEPO | 452 | 0.041476858 | 3.52E-05 | 1 | 0.000849509 | 24.10982982 | 8 mo | HOM |
| **1013978366** | 1 | MEPO | 452 | 0.041476858 | 0 | 0 | 0 | 0 | 2 mo | HOM |
| **1014416155** | 2 | MEPO | 452 | 0.041476858 | 0 | 0 | 0 | 0 | 2 mo | HOM |
| **1013978065** | 2 | MEPO | 452 | 0.041476858 | 0 | 0 | 0 | 0 | 2 mo | HOM |
| **1009934102** | 2 | MEPO | 452 | 0.041476858 | 0 | 0 | 0 | 0 | 4 mo | WILD |
| **1009322937** | 2 | MEPO | 452 | 0.041476858 | 4.96E-06 | 1 | 0.000119668 | 24.10982982 | 8 mo | WILD |
| **1011080637** | 2 | MEPO | 452 | 0.041476858 | 0.000229512 | 2 | 0.005533493 | 48.21965964 | 8 mo | HOM |
| **1013609713** | 2 | MEPO | 452 | 0.041476858 | 0 | 0 | 0 | 0 | 2 mo | WILD |
| **1010418842** | 1 | MEPO | 452 | 0.041476858 | 0.000685178 | 2 | 0.016519515 | 48.21965964 | 8 mo | HOM |

| | | | | | | | | | | |
|---|---|---|---|---|---|---|---|---|---|---|
| **1013288213** | 2 | MEPO | 452 | 0.041476858 | 8.25E-06 | 1 | 0.000198837 | 24.10982982 | 4 mo | HOM |
| **1008976060** | 2 | MEPO | 452 | 0.041476858 | 0.00013255 | 1 | 0.003195754 | 24.10982982 | 8 mo | WILD |
| **1013446243** | 1 | MEPO | 452 | 0.041476858 | 0.000238468 | 0 | 0.005749414 | 0 | 4 mo | HOM |
| **1013288951** | 2 | MEPO | 452 | 0.041476858 | 5.74E-05 | 4 | 0.001384406 | 96.43931929 | 4 mo | HOM |
| **1013978665** | 1 | SS | 453 | 34.32736505 | 8.07E-05 | 5 | 2.35E-06 | 0.145656388 | 2 mo | HOM |
| **1013445609** | 1 | SS | 453 | 34.32736505 | 0.00410054 | 226 | 0.000119454 | 6.583668734 | 4 mo | HOM |
| **1013609015** | 1 | SS | 453 | 34.32736505 | 0.010802211 | 375 | 0.000314682 | 10.92422909 | 4 mo | HOM |
| **1013608106** | 2 | SS | 453 | 34.32736505 | 0 | 0 | 0 | 0 | 4 mo | HOM |
| **1011937543** | 1 | SS | 453 | 34.32736505 | 0.062772338 | 1074 | 0.001828638 | 31.28699213 | 8 mo | HOM |
| **1010419538** | 2 | SS | 453 | 34.32736505 | 5.95E-05 | 5 | 1.73E-06 | 0.145656388 | 4 mo | WILD |
| **1010419187** | 1 | SS | 453 | 34.32736505 | 0.056659495 | 1817 | 0.001650563 | 52.93153137 | 8 mo | HOM |
| **1013978366** | 1 | SS | 453 | 34.32736505 | 0.000269429 | 14 | 7.85E-06 | 0.407837886 | 2 mo | HOM |
| **1014416155** | 2 | SS | 453 | 34.32736505 | 4.49E-06 | 1 | 1.31E-07 | 0.029131278 | 2 mo | HOM |
| **1013978065** | 2 | SS | 453 | 34.32736505 | 6.3E-05 | 3 | 1.83E-06 | 0.087393833 | 2 mo | HOM |
| **1009934102** | 2 | SS | 453 | 34.32736505 | 0.000121777 | 10 | 3.55E-06 | 0.291312776 | 4 mo | WILD |
| **1009322937** | 2 | SS | 453 | 34.32736505 | 5.95E-05 | 9 | 1.73E-06 | 0.262181498 | 8 mo | WILD |
| **1011080637** | 2 | SS | 453 | 34.32736505 | 0.034369529 | 1261 | 0.001001228 | 36.73454103 | 8 mo | HOM |
| **1013609713** | 2 | SS | 453 | 34.32736505 | 6.76E-05 | 5 | 1.97E-06 | 0.145656388 | 2 mo | WILD |
| **1010418842** | 1 | SS | 453 | 34.32736505 | 0.051930093 | 1579 | 0.00151279 | 45.99828731 | 8 mo | HOM |
| **1013288213** | 2 | SS | 453 | 34.32736505 | 0.002523832 | 198 | 7.35E-05 | 5.767992962 | 4 mo | HOM |
| **1008976060** | 2 | SS | 453 | 34.32736505 | 0.001760562 | 77 | 5.13E-05 | 2.243108374 | 8 mo | WILD |
| **1013446243** | 1 | SS | 453 | 34.32736505 | 0.008443174 | 410 | 0.00024596 | 11.94382381 | 4 mo | HOM |
| **1013288951** | 2 | SS | 453 | 34.32736505 | 0.008812758 | 380 | 0.000256727 | 11.06988548 | 4 mo | HOM |
| **1013978665** | 1 | AUDpo6b | 456 | 0.018171109 | 0 | 0 | 0 | 0 | 2 mo | HOM |
| **1013445609** | 1 | AUDpo6b | 456 | 0.018171109 | 0 | 0 | 0 | 0 | 4 mo | HOM |
| **1013609015** | 1 | AUDpo6b | 456 | 0.018171109 | 9.24E-07 | 1 | 5.08E-05 | 55.03241573 | 4 mo | HOM |
| **1013608106** | 2 | AUDpo6b | 456 | 0.018171109 | 0 | 0 | 0 | 0 | 4 mo | HOM |
| **1011937543** | 1 | AUDpo6b | 456 | 0.018171109 | 1.31E-06 | 0 | 7.23E-05 | 0 | 8 mo | HOM |
| **1010419538** | 2 | AUDpo6b | 456 | 0.018171109 | 0 | 0 | 0 | 0 | 4 mo | WILD |
| **1010419187** | 1 | AUDpo6b | 456 | 0.018171109 | 1.39E-05 | 1 | 0.000762446 | 55.03241573 | 8 mo | HOM |
| **1013978366** | 1 | AUDpo6b | 456 | 0.018171109 | 0 | 0 | 0 | 0 | 2 mo | HOM |
| **1014416155** | 2 | AUDpo6b | 456 | 0.018171109 | 0 | 0 | 0 | 0 | 2 mo | HOM |
| **1013978065** | 2 | AUDpo6b | 456 | 0.018171109 | 0 | 0 | 0 | 0 | 2 mo | HOM |
| **1009934102** | 2 | AUDpo6b | 456 | 0.018171109 | 0 | 0 | 0 | 0 | 4 mo | WILD |
| **1009322937** | 2 | AUDpo6b | 456 | 0.018171109 | 0 | 0 | 0 | 0 | 8 mo | WILD |
| **1011080637** | 2 | AUDpo6b | 456 | 0.018171109 | 2.84E-06 | 0 | 0.000156023 | 0 | 8 mo | HOM |
| **1013609713** | 2 | AUDpo6b | 456 | 0.018171109 | 0 | 0 | 0 | 0 | 2 mo | WILD |
| **1010418842** | 1 | AUDpo6b | 456 | 0.018171109 | 4.41E-05 | 2 | 0.002425331 | 110.0648315 | 8 mo | HOM |
| **1013288213** | 2 | AUDpo6b | 456 | 0.018171109 | 0 | 0 | 0 | 0 | 4 mo | HOM |
| **1008976060** | 2 | AUDpo6b | 456 | 0.018171109 | 0 | 0 | 0 | 0 | 8 mo | WILD |
| **1013446243** | 1 | AUDpo6b | 456 | 0.018171109 | 2.5E-06 | 0 | 0.000137586 | 0 | 4 mo | HOM |
| **1013288951** | 2 | AUDpo6b | 456 | 0.018171109 | 0 | 0 | 0 | 0 | 4 mo | HOM |
| **1013978665** | 1 | MEV | 460 | 0.009770654 | 1.1E-07 | 0 | 1.13E-05 | 0 | 2 mo | HOM |
| **1013445609** | 1 | MEV | 460 | 0.009770654 | 0 | 0 | 0 | 0 | 4 mo | HOM |
| **1013609015** | 1 | MEV | 460 | 0.009770654 | 3.94E-08 | 0 | 4.04E-06 | 0 | 4 mo | HOM |
| **1013608106** | 2 | MEV | 460 | 0.009770654 | 0 | 0 | 0 | 0 | 4 mo | HOM |
| **1011937543** | 1 | MEV | 460 | 0.009770654 | 2.16E-06 | 0 | 0.0002208 | 0 | 8 mo | HOM |
| **1010419538** | 2 | MEV | 460 | 0.009770654 | 0 | 0 | 0 | 0 | 4 mo | WILD |
| **1010419187** | 1 | MEV | 460 | 0.009770654 | 6.29E-08 | 0 | 6.44E-06 | 0 | 8 mo | HOM |
| **1013978366** | 1 | MEV | 460 | 0.009770654 | 2.48E-06 | 0 | 0.000254229 | 0 | 2 mo | HOM |
| **1014416155** | 2 | MEV | 460 | 0.009770654 | 0 | 0 | 0 | 0 | 2 mo | HOM |
| **1013978065** | 2 | MEV | 460 | 0.009770654 | 0 | 0 | 0 | 0 | 2 mo | HOM |
| **1009934102** | 2 | MEV | 460 | 0.009770654 | 0 | 0 | 0 | 0 | 4 mo | WILD |
| **1009322937** | 2 | MEV | 460 | 0.009770654 | 2.01E-09 | 0 | 2.06E-07 | 0 | 8 mo | WILD |
| **1011080637** | 2 | MEV | 460 | 0.009770654 | 0 | 0 | 0 | 0 | 8 mo | HOM |
| **1013609713** | 2 | MEV | 460 | 0.009770654 | 0 | 0 | 0 | 0 | 2 mo | WILD |
| **1010418842** | 1 | MEV | 460 | 0.009770654 | 4.66E-06 | 0 | 0.000476924 | 0 | 8 mo | HOM |
| **1013288213** | 2 | MEV | 460 | 0.009770654 | 2.95E-08 | 0 | 3.02E-06 | 0 | 4 mo | HOM |
| **1008976060** | 2 | MEV | 460 | 0.009770654 | 2.65E-08 | 0 | 2.72E-06 | 0 | 8 mo | WILD |

| | | | | | | | | | | |
|---|---|---|---|---|---|---|---|---|---|---|
| 1013446243 | 1 | MEV | 460 | 0.009770654 | 1.71E-06 | 0 | 0.000174682 | 0 | 4 mo | HOM |
| 1013288951 | 2 | MEV | 460 | 0.009770654 | 0 | 0 | 0 | 0 | 4 mo | HOM |
| 1013978665 | 1 | SSp-tr6b | 461 | 0.034683555 | 0 | 0 | 0 | 0 | 2 mo | HOM |
| 1013445609 | 1 | SSp-tr6b | 461 | 0.034683555 | 0 | 0 | 0 | 0 | 4 mo | HOM |
| 1013609015 | 1 | SSp-tr6b | 461 | 0.034683555 | 2.24E-06 | 0 | 6.47E-05 | 0 | 4 mo | HOM |
| 1013608106 | 2 | SSp-tr6b | 461 | 0.034683555 | 0 | 0 | 0 | 0 | 4 mo | HOM |
| 1011937543 | 1 | SSp-tr6b | 461 | 0.034683555 | 2.64E-05 | 0 | 0.000760752 | 0 | 8 mo | HOM |
| 1010419538 | 2 | SSp-tr6b | 461 | 0.034683555 | 0 | 0 | 0 | 0 | 4 mo | WILD |
| 1010419187 | 1 | SSp-tr6b | 461 | 0.034683555 | 2.7E-05 | 0 | 0.000777354 | 0 | 8 mo | HOM |
| 1013978366 | 1 | SSp-tr6b | 461 | 0.034683555 | 0 | 0 | 0 | 0 | 2 mo | HOM |
| 1014416155 | 2 | SSp-tr6b | 461 | 0.034683555 | 0 | 0 | 0 | 0 | 2 mo | HOM |
| 1013978065 | 2 | SSp-tr6b | 461 | 0.034683555 | 0 | 0 | 0 | 0 | 2 mo | HOM |
| 1009934102 | 2 | SSp-tr6b | 461 | 0.034683555 | 0 | 0 | 0 | 0 | 4 mo | WILD |
| 1009322937 | 2 | SSp-tr6b | 461 | 0.034683555 | 0 | 0 | 0 | 0 | 8 mo | WILD |
| 1011080637 | 2 | SSp-tr6b | 461 | 0.034683555 | 2.89E-06 | 0 | 8.33E-05 | 0 | 8 mo | HOM |
| 1013609713 | 2 | SSp-tr6b | 461 | 0.034683555 | 0 | 0 | 0 | 0 | 2 mo | WILD |
| 1010418842 | 1 | SSp-tr6b | 461 | 0.034683555 | 0.000105291 | 4 | 0.003035756 | 115.3284321 | 8 mo | HOM |
| 1013288213 | 2 | SSp-tr6b | 461 | 0.034683555 | 0 | 0 | 0 | 0 | 4 mo | HOM |
| 1008976060 | 2 | SSp-tr6b | 461 | 0.034683555 | 0 | 0 | 0 | 0 | 8 mo | WILD |
| 1013446243 | 1 | SSp-tr6b | 461 | 0.034683555 | 1.39E-06 | 0 | 4E-05 | 0 | 4 mo | HOM |
| 1013288951 | 2 | SSp-tr6b | 461 | 0.034683555 | 2.08E-05 | 1 | 0.000598603 | 28.83210802 | 4 mo | HOM |
| 1013978665 | 1 | CA3 | 463 | 6.452738342 | 0 | 0 | 0 | 0 | 2 mo | HOM |
| 1013445609 | 1 | CA3 | 463 | 6.452738342 | 0.000114955 | 9 | 1.78E-05 | 1.394756694 | 4 mo | HOM |
| 1013609015 | 1 | CA3 | 463 | 6.452738342 | 0.001081482 | 33 | 0.0001676 | 5.114107879 | 4 mo | HOM |
| 1013608106 | 2 | CA3 | 463 | 6.452738342 | 0 | 0 | 0 | 0 | 4 mo | HOM |
| 1011937543 | 1 | CA3 | 463 | 6.452738342 | 0.005524618 | 97 | 0.000856166 | 15.03237771 | 8 mo | HOM |
| 1010419538 | 2 | CA3 | 463 | 6.452738342 | 2.97E-07 | 0 | 4.6E-08 | 0 | 4 mo | WILD |
| 1010419187 | 1 | CA3 | 463 | 6.452738342 | 0.008426605 | 240 | 0.001305896 | 37.19351185 | 8 mo | HOM |
| 1013978366 | 1 | CA3 | 463 | 6.452738342 | 0 | 0 | 0 | 0 | 2 mo | HOM |
| 1014416155 | 2 | CA3 | 463 | 6.452738342 | 0 | 0 | 0 | 0 | 2 mo | HOM |
| 1013978065 | 2 | CA3 | 463 | 6.452738342 | 0 | 0 | 0 | 0 | 2 mo | HOM |
| 1009934102 | 2 | CA3 | 463 | 6.452738342 | 6.35E-07 | 0 | 9.84E-08 | 0 | 4 mo | WILD |
| 1009322937 | 2 | CA3 | 463 | 6.452738342 | 1.08E-06 | 0 | 1.67E-07 | 0 | 8 mo | WILD |
| 1011080637 | 2 | CA3 | 463 | 6.452738342 | 0.005744931 | 216 | 0.000890309 | 33.47416066 | 8 mo | HOM |
| 1013609713 | 2 | CA3 | 463 | 6.452738342 | 0 | 0 | 0 | 0 | 2 mo | WILD |
| 1010418842 | 1 | CA3 | 463 | 6.452738342 | 0.009338383 | 261 | 0.001447197 | 40.44794414 | 8 mo | HOM |
| 1013288213 | 2 | CA3 | 463 | 6.452738342 | 0.000290194 | 21 | 4.5E-05 | 3.254432287 | 4 mo | HOM |
| 1008976060 | 2 | CA3 | 463 | 6.452738342 | 0.000152113 | 20 | 2.36E-05 | 3.099459321 | 8 mo | WILD |
| 1013446243 | 1 | CA3 | 463 | 6.452738342 | 0.000406671 | 25 | 6.3E-05 | 3.874324151 | 4 mo | HOM |
| 1013288951 | 2 | CA3 | 463 | 6.452738342 | 0.000483212 | 19 | 7.49E-05 | 2.944486355 | 4 mo | HOM |
| 1013978665 | 1 | alv | 466 | 1.434738659 | 0 | 0 | 0 | 0 | 2 mo | HOM |
| 1013445609 | 1 | alv | 466 | 1.434738659 | 6.24E-06 | 0 | 4.35E-06 | 0 | 4 mo | HOM |
| 1013609015 | 1 | alv | 466 | 1.434738659 | 0.000533813 | 6 | 0.000372063 | 4.181946281 | 4 mo | HOM |
| 1013608106 | 2 | alv | 466 | 1.434738659 | 0 | 0 | 0 | 0 | 4 mo | HOM |
| 1011937543 | 1 | alv | 466 | 1.434738659 | 0.001398717 | 25 | 0.000974894 | 17.42477617 | 8 mo | HOM |
| 1010419538 | 2 | alv | 466 | 1.434738659 | 2.33E-06 | 0 | 1.62E-06 | 0 | 4 mo | WILD |
| 1010419187 | 1 | alv | 466 | 1.434738659 | 0.001234495 | 29 | 0.000860432 | 20.21274036 | 8 mo | HOM |
| 1013978366 | 1 | alv | 466 | 1.434738659 | 6.53E-07 | 0 | 4.55E-07 | 0 | 2 mo | HOM |
| 1014416155 | 2 | alv | 466 | 1.434738659 | 0 | 0 | 0 | 0 | 2 mo | HOM |
| 1013978065 | 2 | alv | 466 | 1.434738659 | 0 | 0 | 0 | 0 | 2 mo | HOM |
| 1009934102 | 2 | alv | 466 | 1.434738659 | 4.07E-08 | 0 | 2.84E-08 | 0 | 4 mo | WILD |
| 1009322937 | 2 | alv | 466 | 1.434738659 | 3.59E-07 | 0 | 2.5E-07 | 0 | 8 mo | WILD |
| 1011080637 | 2 | alv | 466 | 1.434738659 | 0.000935119 | 24 | 0.00065177 | 16.72778512 | 8 mo | HOM |
| 1013609713 | 2 | alv | 466 | 1.434738659 | 0 | 0 | 0 | 0 | 2 mo | WILD |
| 1010418842 | 1 | alv | 466 | 1.434738659 | 0.001739265 | 51 | 0.001212252 | 35.54654339 | 8 mo | HOM |
| 1013288213 | 2 | alv | 466 | 1.434738659 | 2.53E-06 | 0 | 1.76E-06 | 0 | 4 mo | HOM |
| 1008976060 | 2 | alv | 466 | 1.434738659 | 4.66E-05 | 5 | 3.24E-05 | 3.484955234 | 8 mo | WILD |
| 1013446243 | 1 | alv | 466 | 1.434738659 | 6.83E-05 | 2 | 4.76E-05 | 1.393982094 | 4 mo | HOM |
| 1013288951 | 2 | alv | 466 | 1.434738659 | 0.000262668 | 5 | 0.000183077 | 3.484955234 | 4 mo | HOM |

| | | | | | | | | | | |
|---|---|---|---|---|---|---|---|---|---|---|
| **1013978665** | 1 | MEZ | 467 | 3.963549607 | 1.52E-06 | 0 | 3.84E-07 | 0 | 2 mo | HOM |
| **1013445609** | 1 | MEZ | 467 | 3.963549607 | 2.09E-07 | 0 | 5.27E-08 | 0 | 4 mo | HOM |
| **1013609015** | 1 | MEZ | 467 | 3.963549607 | 2.91E-05 | 3 | 7.35E-06 | 0.756897301 | 4 mo | HOM |
| **1013608106** | 2 | MEZ | 467 | 3.963549607 | 0 | 0 | 0 | 0 | 4 mo | HOM |
| **1011937543** | 1 | MEZ | 467 | 3.963549607 | 0.007300764 | 50 | 0.001841976 | 12.61495502 | 8 mo | HOM |
| **1010419538** | 2 | MEZ | 467 | 3.963549607 | 3.06E-06 | 0 | 7.72E-07 | 0 | 4 mo | WILD |
| **1010419187** | 1 | MEZ | 467 | 3.963549607 | 0.003220191 | 31 | 0.000812451 | 7.82127211 | 8 mo | HOM |
| **1013978366** | 1 | MEZ | 467 | 3.963549607 | 1.37E-05 | 1 | 3.45E-06 | 0.2522991 | 2 mo | HOM |
| **1014416155** | 2 | MEZ | 467 | 3.963549607 | 0 | 0 | 0 | 0 | 2 mo | HOM |
| **1013978065** | 2 | MEZ | 467 | 3.963549607 | 4.08E-06 | 1 | 1.03E-06 | 0.2522991 | 2 mo | HOM |
| **1009934102** | 2 | MEZ | 467 | 3.963549607 | 7.92E-06 | 1 | 2E-06 | 0.2522991 | 4 mo | WILD |
| **1009322937** | 2 | MEZ | 467 | 3.963549607 | 1.34E-06 | 0 | 3.38E-07 | 0 | 8 mo | WILD |
| **1011080637** | 2 | MEZ | 467 | 3.963549607 | 0.000618533 | 27 | 0.000156055 | 6.812075708 | 8 mo | HOM |
| **1013609713** | 2 | MEZ | 467 | 3.963549607 | 0 | 0 | 0 | 0 | 2 mo | WILD |
| **1010418842** | 1 | MEZ | 467 | 3.963549607 | 0.000919969 | 46 | 0.000232107 | 11.60575861 | 8 mo | HOM |
| **1013288213** | 2 | MEZ | 467 | 3.963549607 | 1.16E-05 | 2 | 2.92E-06 | 0.504598201 | 4 mo | HOM |
| **1008976060** | 2 | MEZ | 467 | 3.963549607 | 0.000317557 | 11 | 8.01E-05 | 2.775290103 | 8 mo | WILD |
| **1013446243** | 1 | MEZ | 467 | 3.963549607 | 0 | 0 | 0 | 0 | 4 mo | HOM |
| **1013288951** | 2 | MEZ | 467 | 3.963549607 | 5.22E-06 | 1 | 1.32E-06 | 0.2522991 | 4 mo | HOM |
| **1013978665** | 1 | VISpm6b | 469 | 0.028842179 | 0 | 0 | 0 | 0 | 2 mo | HOM |
| **1013445609** | 1 | VISpm6b | 469 | 0.028842179 | 4.28E-06 | 1 | 0.000148554 | 34.67144409 | 4 mo | HOM |
| **1013609015** | 1 | VISpm6b | 469 | 0.028842179 | 0 | 0 | 0 | 0 | 4 mo | HOM |
| **1013608106** | 2 | VISpm6b | 469 | 0.028842179 | 0 | 0 | 0 | 0 | 4 mo | HOM |
| **1011937543** | 1 | VISpm6b | 469 | 0.028842179 | 2.54E-05 | 0 | 0.000881839 | 0 | 8 mo | HOM |
| **1010419538** | 2 | VISpm6b | 469 | 0.028842179 | 0 | 0 | 0 | 0 | 4 mo | WILD |
| **1010419187** | 1 | VISpm6b | 469 | 0.028842179 | 2.93E-05 | 0 | 0.001014293 | 0 | 8 mo | HOM |
| **1013978366** | 1 | VISpm6b | 469 | 0.028842179 | 0 | 0 | 0 | 0 | 2 mo | HOM |
| **1014416155** | 2 | VISpm6b | 469 | 0.028842179 | 0 | 0 | 0 | 0 | 2 mo | HOM |
| **1013978065** | 2 | VISpm6b | 469 | 0.028842179 | 0 | 0 | 0 | 0 | 2 mo | HOM |
| **1009934102** | 2 | VISpm6b | 469 | 0.028842179 | 0 | 0 | 0 | 0 | 4 mo | WILD |
| **1009322937** | 2 | VISpm6b | 469 | 0.028842179 | 0 | 0 | 0 | 0 | 8 mo | WILD |
| **1011080637** | 2 | VISpm6b | 469 | 0.028842179 | 4.49E-07 | 0 | 1.56E-05 | 0 | 8 mo | HOM |
| **1013609713** | 2 | VISpm6b | 469 | 0.028842179 | 0 | 0 | 0 | 0 | 2 mo | WILD |
| **1010418842** | 1 | VISpm6b | 469 | 0.028842179 | 1.81E-05 | 1 | 0.000626075 | 34.67144409 | 8 mo | HOM |
| **1013288213** | 2 | VISpm6b | 469 | 0.028842179 | 0 | 0 | 0 | 0 | 4 mo | HOM |
| **1008976060** | 2 | VISpm6b | 469 | 0.028842179 | 0 | 0 | 0 | 0 | 8 mo | WILD |
| **1013446243** | 1 | VISpm6b | 469 | 0.028842179 | 1.55E-06 | 0 | 5.38E-05 | 0 | 4 mo | HOM |
| **1013288951** | 2 | VISpm6b | 469 | 0.028842179 | 0 | 0 | 0 | 0 | 4 mo | HOM |
| **1013978665** | 1 | STN | 470 | 0.200481775 | 0 | 0 | 0 | 0 | 2 mo | HOM |
| **1013445609** | 1 | STN | 470 | 0.200481775 | 0 | 0 | 0 | 0 | 4 mo | HOM |
| **1013609015** | 1 | STN | 470 | 0.200481775 | 0 | 0 | 0 | 0 | 4 mo | HOM |
| **1013608106** | 2 | STN | 470 | 0.200481775 | 0 | 0 | 0 | 0 | 4 mo | HOM |
| **1011937543** | 1 | STN | 470 | 0.200481775 | 0.00058696 | 3 | 0.002927748 | 14.96395369 | 8 mo | HOM |
| **1010419538** | 2 | STN | 470 | 0.200481775 | 0 | 0 | 0 | 0 | 4 mo | WILD |
| **1010419187** | 1 | STN | 470 | 0.200481775 | 0 | 0 | 0 | 0 | 8 mo | HOM |
| **1013978366** | 1 | STN | 470 | 0.200481775 | 0 | 0 | 0 | 0 | 2 mo | HOM |
| **1014416155** | 2 | STN | 470 | 0.200481775 | 0 | 0 | 0 | 0 | 2 mo | HOM |
| **1013978065** | 2 | STN | 470 | 0.200481775 | 0 | 0 | 0 | 0 | 2 mo | HOM |
| **1009934102** | 2 | STN | 470 | 0.200481775 | 0 | 0 | 0 | 0 | 4 mo | WILD |
| **1009322937** | 2 | STN | 470 | 0.200481775 | 0 | 0 | 0 | 0 | 8 mo | WILD |
| **1011080637** | 2 | STN | 470 | 0.200481775 | 0 | 0 | 0 | 0 | 8 mo | HOM |
| **1013609713** | 2 | STN | 470 | 0.200481775 | 0 | 0 | 0 | 0 | 2 mo | WILD |
| **1010418842** | 1 | STN | 470 | 0.200481775 | 9.1E-05 | 4 | 0.000453943 | 19.95193825 | 8 mo | HOM |
| **1013288213** | 2 | STN | 470 | 0.200481775 | 0 | 0 | 0 | 0 | 4 mo | HOM |
| **1008976060** | 2 | STN | 470 | 0.200481775 | 3.53E-05 | 1 | 0.000175873 | 4.987984564 | 8 mo | WILD |
| **1013446243** | 1 | STN | 470 | 0.200481775 | 0 | 0 | 0 | 0 | 4 mo | HOM |
| **1013288951** | 2 | STN | 470 | 0.200481775 | 0 | 0 | 0 | 0 | 4 mo | HOM |
| **1013978665** | 1 | MG | 475 | 0.72099678 | 0 | 0 | 0 | 0 | 2 mo | HOM |
| **1013445609** | 1 | MG | 475 | 0.72099678 | 9.65E-06 | 1 | 1.34E-05 | 1.386968746 | 4 mo | HOM |

| | | | | | | | | | | |
|---|---|---|---|---|---|---|---|---|---|---|
| **1013609015** | 1 | MG | 475 | 0.72099678 | 0 | 0 | 0 | 0 | 4 mo | HOM |
| **1013608106** | 2 | MG | 475 | 0.72099678 | 0 | 0 | 0 | 0 | 4 mo | HOM |
| **1011937543** | 1 | MG | 475 | 0.72099678 | 0.00090577 | 19 | 0.001256275 | 26.35240618 | 8 mo | HOM |
| **1010419538** | 2 | MG | 475 | 0.72099678 | 0 | 0 | 0 | 0 | 4 mo | WILD |
| **1010419187** | 1 | MG | 475 | 0.72099678 | 0.000162239 | 7 | 0.00022502 | 9.708781224 | 8 mo | HOM |
| **1013978366** | 1 | MG | 475 | 0.72099678 | 0 | 0 | 0 | 0 | 2 mo | HOM |
| **1014416155** | 2 | MG | 475 | 0.72099678 | 0 | 0 | 0 | 0 | 2 mo | HOM |
| **1013978065** | 2 | MG | 475 | 0.72099678 | 0 | 0 | 0 | 0 | 2 mo | HOM |
| **1009934102** | 2 | MG | 475 | 0.72099678 | 0 | 0 | 0 | 0 | 4 mo | WILD |
| **1009322937** | 2 | MG | 475 | 0.72099678 | 0 | 0 | 0 | 0 | 8 mo | WILD |
| **1011080637** | 2 | MG | 475 | 0.72099678 | 0.00014873 | 6 | 0.000206284 | 8.321812477 | 8 mo | HOM |
| **1013609713** | 2 | MG | 475 | 0.72099678 | 0 | 0 | 0 | 0 | 2 mo | WILD |
| **1010418842** | 1 | MG | 475 | 0.72099678 | 0.000391064 | 14 | 0.000542394 | 19.41756245 | 8 mo | HOM |
| **1013288213** | 2 | MG | 475 | 0.72099678 | 0 | 0 | 0 | 0 | 4 mo | HOM |
| **1008976060** | 2 | MG | 475 | 0.72099678 | 4.79E-05 | 3 | 6.64E-05 | 4.160906239 | 8 mo | WILD |
| **1013446243** | 1 | MG | 475 | 0.72099678 | 0 | 0 | 0 | 0 | 4 mo | HOM |
| **1013288951** | 2 | MG | 475 | 0.72099678 | 0 | 0 | 0 | 0 | 4 mo | HOM |
| **1013978665** | 1 | STR | 477 | 46.27580447 | 2.91E-06 | 1 | 6.29E-08 | 0.021609565 | 2 mo | HOM |
| **1013445609** | 1 | STR | 477 | 46.32076718 | 0.000477646 | 38 | 1.03E-05 | 0.820366378 | 4 mo | HOM |
| **1013609015** | 1 | STR | 477 | 46.32076718 | 0.004823585 | 125 | 0.000104134 | 2.698573612 | 4 mo | HOM |
| **1013608106** | 2 | STR | 477 | 46.32076718 | 0 | 0 | 0 | 0 | 4 mo | HOM |
| **1011937543** | 1 | STR | 477 | 46.32076718 | 0.030258674 | 646 | 0.000653242 | 13.94622843 | 8 mo | HOM |
| **1010419538** | 2 | STR | 477 | 46.32076718 | 5.34E-06 | 1 | 1.15E-07 | 0.021588589 | 4 mo | WILD |
| **1010419187** | 1 | STR | 477 | 46.32076718 | 0.023653025 | 924 | 0.000510635 | 19.94785614 | 8 mo | HOM |
| **1013978366** | 1 | STR | 477 | 46.32076718 | 2.09E-06 | 0 | 4.52E-08 | 0 | 2 mo | HOM |
| **1014416155** | 2 | STR | 477 | 46.32076718 | 0 | 0 | 0 | 0 | 2 mo | HOM |
| **1013978065** | 2 | STR | 477 | 46.32076718 | 1.07E-05 | 2 | 2.3E-07 | 0.043177178 | 2 mo | HOM |
| **1009934102** | 2 | STR | 477 | 46.32076718 | 3.9E-05 | 3 | 8.42E-07 | 0.064765767 | 4 mo | WILD |
| **1009322937** | 2 | STR | 477 | 46.32076718 | 8.03E-05 | 3 | 1.73E-06 | 0.064765767 | 8 mo | WILD |
| **1011080637** | 2 | STR | 477 | 46.32076718 | 0.015701679 | 731 | 0.000338977 | 15.78125848 | 8 mo | HOM |
| **1013609713** | 2 | STR | 477 | 46.32076718 | 0 | 0 | 0 | 0 | 2 mo | WILD |
| **1010418842** | 1 | STR | 477 | 46.32076718 | 0.036071649 | 1145 | 0.000778736 | 24.71893429 | 8 mo | HOM |
| **1013288213** | 2 | STR | 477 | 46.32076718 | 0.001721253 | 44 | 3.72E-05 | 0.949897911 | 4 mo | HOM |
| **1008976060** | 2 | STR | 477 | 46.32076718 | 0.001426259 | 106 | 3.08E-05 | 2.288390423 | 8 mo | WILD |
| **1013446243** | 1 | STR | 477 | 46.32076718 | 0.001502952 | 94 | 3.24E-05 | 2.029327356 | 4 mo | HOM |
| **1013288951** | 2 | STR | 477 | 46.32076718 | 0.002642581 | 105 | 5.7E-05 | 2.266801834 | 4 mo | HOM |
| **1013978665** | 1 | SSp-ll6a | 478 | 0.544035044 | 0 | 0 | 0 | 0 | 2 mo | HOM |
| **1013445609** | 1 | SSp-ll6a | 478 | 0.544035044 | 6.04E-05 | 3 | 0.000110992 | 5.514350653 | 4 mo | HOM |
| **1013609015** | 1 | SSp-ll6a | 478 | 0.544035044 | 0.000180465 | 4 | 0.000331715 | 7.352467537 | 4 mo | HOM |
| **1013608106** | 2 | SSp-ll6a | 478 | 0.544035044 | 0 | 0 | 0 | 0 | 4 mo | HOM |
| **1011937543** | 1 | SSp-ll6a | 478 | 0.544035044 | 0.000811637 | 14 | 0.001491883 | 25.73363638 | 8 mo | HOM |
| **1010419538** | 2 | SSp-ll6a | 478 | 0.544035044 | 0 | 0 | 0 | 0 | 4 mo | WILD |
| **1010419187** | 1 | SSp-ll6a | 478 | 0.544035044 | 0.00062764 | 36 | 0.001153676 | 66.17220783 | 8 mo | HOM |
| **1013978366** | 1 | SSp-ll6a | 478 | 0.544035044 | 0 | 0 | 0 | 0 | 2 mo | HOM |
| **1014416155** | 2 | SSp-ll6a | 478 | 0.544035044 | 0 | 0 | 0 | 0 | 2 mo | HOM |
| **1013978065** | 2 | SSp-ll6a | 478 | 0.544035044 | 0 | 0 | 0 | 0 | 2 mo | HOM |
| **1009934102** | 2 | SSp-ll6a | 478 | 0.544035044 | 0 | 0 | 0 | 0 | 4 mo | WILD |
| **1009322937** | 2 | SSp-ll6a | 478 | 0.544035044 | 0 | 0 | 0 | 0 | 8 mo | WILD |
| **1011080637** | 2 | SSp-ll6a | 478 | 0.544035044 | 0.00039462 | 21 | 0.000725358 | 38.60045457 | 8 mo | HOM |
| **1013609713** | 2 | SSp-ll6a | 478 | 0.544035044 | 0 | 0 | 0 | 0 | 2 mo | WILD |
| **1010418842** | 1 | SSp-ll6a | 478 | 0.544035044 | 0.000786682 | 36 | 0.001446013 | 66.17220783 | 8 mo | HOM |
| **1013288213** | 2 | SSp-ll6a | 478 | 0.544035044 | 4E-07 | 0 | 7.36E-07 | 0 | 4 mo | HOM |
| **1008976060** | 2 | SSp-ll6a | 478 | 0.544035044 | 9.73E-06 | 3 | 1.79E-05 | 5.514350653 | 8 mo | WILD |
| **1013446243** | 1 | SSp-ll6a | 478 | 0.544035044 | 0.000170083 | 8 | 0.000312632 | 14.70493507 | 4 mo | HOM |
| **1013288951** | 2 | SSp-ll6a | 478 | 0.544035044 | 0.000155454 | 5 | 0.000285742 | 9.190584421 | 4 mo | HOM |
| **1013978665** | 1 | bic | 482 | 0.2539484 | 0 | 0 | 0 | 0 | 2 mo | HOM |
| **1013445609** | 1 | bic | 482 | 0.2539484 | 4.46E-06 | 1 | 1.76E-05 | 3.937807844 | 4 mo | HOM |
| **1013609015** | 1 | bic | 482 | 0.2539484 | 2.59E-07 | 0 | 1.02E-06 | 0 | 4 mo | HOM |
| **1013608106** | 2 | bic | 482 | 0.2539484 | 0 | 0 | 0 | 0 | 4 mo | HOM |

| | | | | | | | | | | |
|---|---|---|---|---|---|---|---|---|---|---|
| 1011937543 | 1 | bic | 482 | 0.2539484 | 9.53E-05 | 2 | 0.000375137 | 7.875615689 | 8 mo | HOM |
| 1010419538 | 2 | bic | 482 | 0.2539484 | 0 | 0 | 0 | 0 | 4 mo | WILD |
| 1010419187 | 1 | bic | 482 | 0.2539484 | 0.001869665 | 0 | 0.00736238 | 0 | 8 mo | HOM |
| 1013978366 | 1 | bic | 482 | 0.2539484 | 3.89E-05 | 1 | 0.00015326 | 3.937807844 | 2 mo | HOM |
| 1014416155 | 2 | bic | 482 | 0.2539484 | 0 | 0 | 0 | 0 | 2 mo | HOM |
| 1013978065 | 2 | bic | 482 | 0.2539484 | 0 | 0 | 0 | 0 | 2 mo | HOM |
| 1009934102 | 2 | bic | 482 | 0.2539484 | 1.14E-06 | 0 | 4.51E-06 | 0 | 4 mo | WILD |
| 1009322937 | 2 | bic | 482 | 0.2539484 | 0 | 0 | 0 | 0 | 8 mo | WILD |
| 1011080637 | 2 | bic | 482 | 0.2539484 | 6.21E-06 | 0 | 2.44E-05 | 0 | 8 mo | HOM |
| 1013609713 | 2 | bic | 482 | 0.2539484 | 0 | 0 | 0 | 0 | 2 mo | WILD |
| 1010418842 | 1 | bic | 482 | 0.2539484 | 0.000302353 | 1 | 0.001190607 | 3.937807844 | 8 mo | HOM |
| 1013288213 | 2 | bic | 482 | 0.2539484 | 0 | 0 | 0 | 0 | 4 mo | HOM |
| 1008976060 | 2 | bic | 482 | 0.2539484 | 1.86E-05 | 3 | 7.31E-05 | 11.81342353 | 8 mo | WILD |
| 1013446243 | 1 | bic | 482 | 0.2539484 | 0 | 0 | 0 | 0 | 4 mo | HOM |
| 1013288951 | 2 | bic | 482 | 0.2539484 | 0 | 0 | 0 | 0 | 4 mo | HOM |
| 1013978665 | 1 | MH | 483 | 0.329700851 | 0 | 0 | 0 | 0 | 2 mo | HOM |
| 1013445609 | 1 | MH | 483 | 0.329700851 | 0 | 0 | 0 | 0 | 4 mo | HOM |
| 1013609015 | 1 | MH | 483 | 0.329700851 | 0.000127078 | 3 | 0.000385433 | 9.099157582 | 4 mo | HOM |
| 1013608106 | 2 | MH | 483 | 0.329700851 | 0 | 0 | 0 | 0 | 4 mo | HOM |
| 1011937543 | 1 | MH | 483 | 0.329700851 | 0.000909615 | 12 | 0.00275891 | 36.39663033 | 8 mo | HOM |
| 1010419538 | 2 | MH | 483 | 0.329700851 | 4.02E-07 | 0 | 1.22E-06 | 0 | 4 mo | WILD |
| 1010419187 | 1 | MH | 483 | 0.329700851 | 0.000111417 | 6 | 0.000337934 | 18.19831516 | 8 mo | HOM |
| 1013978366 | 1 | MH | 483 | 0.329700851 | 0 | 0 | 0 | 0 | 2 mo | HOM |
| 1014416155 | 2 | MH | 483 | 0.329700851 | 0 | 0 | 0 | 0 | 2 mo | HOM |
| 1013978065 | 2 | MH | 483 | 0.329700851 | 0 | 0 | 0 | 0 | 2 mo | HOM |
| 1009934102 | 2 | MH | 483 | 0.329700851 | 0 | 0 | 0 | 0 | 4 mo | WILD |
| 1009322937 | 2 | MH | 483 | 0.329700851 | 1.03E-05 | 2 | 3.14E-05 | 6.066105055 | 8 mo | WILD |
| 1011080637 | 2 | MH | 483 | 0.329700851 | 0.001865525 | 4 | 0.005658234 | 12.13221011 | 8 mo | HOM |
| 1013609713 | 2 | MH | 483 | 0.329700851 | 0 | 0 | 0 | 0 | 2 mo | WILD |
| 1010418842 | 1 | MH | 483 | 0.329700851 | 0.000104356 | 4 | 0.000316518 | 12.13221011 | 8 mo | HOM |
| 1013288213 | 2 | MH | 483 | 0.329700851 | 0 | 0 | 0 | 0 | 4 mo | HOM |
| 1008976060 | 2 | MH | 483 | 0.329700851 | 0.000389923 | 5 | 0.001182657 | 15.16526264 | 8 mo | WILD |
| 1013446243 | 1 | MH | 483 | 0.329700851 | 2.3E-05 | 1 | 6.96E-05 | 3.033052527 | 4 mo | HOM |
| 1013288951 | 2 | MH | 483 | 0.329700851 | 0.000574355 | 3 | 0.001742049 | 9.099157582 | 4 mo | HOM |
| 1013978665 | 1 | ORBm1 | 484 | 0.433561957 | 3.36E-08 | 0 | 7.75E-08 | 0 | 2 mo | HOM |
| 1013445609 | 1 | ORBm1 | 484 | 0.433561957 | 0 | 0 | 0 | 0 | 4 mo | HOM |
| 1013609015 | 1 | ORBm1 | 484 | 0.433561957 | 0.000399798 | 5 | 0.000922124 | 11.53237714 | 4 mo | HOM |
| 1013608106 | 2 | ORBm1 | 484 | 0.433561957 | 0 | 0 | 0 | 0 | 4 mo | HOM |
| 1011937543 | 1 | ORBm1 | 484 | 0.433561957 | 0.000760193 | 10 | 0.001753366 | 23.06475428 | 8 mo | HOM |
| 1010419538 | 2 | ORBm1 | 484 | 0.433561957 | 0 | 0 | 0 | 0 | 4 mo | WILD |
| 1010419187 | 1 | ORBm1 | 484 | 0.433561957 | 0.000149383 | 9 | 0.000344548 | 20.75827885 | 8 mo | HOM |
| 1013978366 | 1 | ORBm1 | 484 | 0.433561957 | 6.68E-06 | 0 | 1.54E-05 | 0 | 2 mo | HOM |
| 1014416155 | 2 | ORBm1 | 484 | 0.433561957 | 1.66E-05 | 1 | 3.84E-05 | 2.306475428 | 2 mo | HOM |
| 1013978065 | 2 | ORBm1 | 484 | 0.433561957 | 3.57E-08 | 0 | 8.24E-08 | 0 | 2 mo | HOM |
| 1009934102 | 2 | ORBm1 | 484 | 0.433561957 | 1.04E-05 | 1 | 2.41E-05 | 2.306475428 | 4 mo | WILD |
| 1009322937 | 2 | ORBm1 | 484 | 0.433561957 | 7.16E-05 | 3 | 0.0001651 | 6.919426285 | 8 mo | WILD |
| 1011080637 | 2 | ORBm1 | 484 | 0.433561957 | 0.001350704 | 12 | 0.003115365 | 27.67770514 | 8 mo | HOM |
| 1013609713 | 2 | ORBm1 | 484 | 0.433561957 | 0 | 0 | 0 | 0 | 2 mo | WILD |
| 1010418842 | 1 | ORBm1 | 484 | 0.433561957 | 0.001946067 | 6 | 0.004488556 | 13.83885257 | 8 mo | HOM |
| 1013288213 | 2 | ORBm1 | 484 | 0.433561957 | 0.002219774 | 12 | 0.005119854 | 27.67770514 | 4 mo | HOM |
| 1008976060 | 2 | ORBm1 | 484 | 0.433561957 | 6.95E-05 | 6 | 0.000160249 | 13.83885257 | 8 mo | WILD |
| 1013446243 | 1 | ORBm1 | 484 | 0.433561957 | 4.78E-05 | 1 | 0.000110208 | 2.306475428 | 4 mo | HOM |
| 1013288951 | 2 | ORBm1 | 484 | 0.433561957 | 0.000643671 | 19 | 0.001484611 | 43.82303314 | 4 mo | HOM |
| 1013978665 | 1 | STRd | 485 | 26.8051883 | 3.05E-07 | 0 | 1.14E-08 | 0 | 2 mo | HOM |
| 1013445609 | 1 | STRd | 485 | 26.8051883 | 0.000235332 | 18 | 8.78E-06 | 0.671511791 | 4 mo | HOM |
| 1013609015 | 1 | STRd | 485 | 26.8051883 | 0.002482428 | 74 | 9.26E-05 | 2.760659585 | 4 mo | HOM |
| 1013608106 | 2 | STRd | 485 | 26.8051883 | 0 | 0 | 0 | 0 | 4 mo | HOM |
| 1011937543 | 1 | STRd | 485 | 26.8051883 | 0.01161648 | 284 | 0.000433367 | 10.59496381 | 8 mo | HOM |
| 1010419538 | 2 | STRd | 485 | 26.8051883 | 0 | 0 | 0 | 0 | 4 mo | WILD |

| | | | | | | | | | | |
|---|---|---|---|---|---|---|---|---|---|---|
| 1010419187 | 1 | STRd | 485 | 26.8051883 | 0.009902259 | 413 | 0.000369416 | 15.40746498 | 8 mo | HOM |
| 1013978366 | 1 | STRd | 485 | 26.8051883 | 2.81E-07 | 0 | 1.05E-08 | 0 | 2 mo | HOM |
| 1014416155 | 2 | STRd | 485 | 26.8051883 | 0 | 0 | 0 | 0 | 2 mo | HOM |
| 1013978065 | 2 | STRd | 485 | 26.8051883 | 0 | 0 | 0 | 0 | 2 mo | HOM |
| 1009934102 | 2 | STRd | 485 | 26.8051883 | 0 | 0 | 0 | 0 | 4 mo | WILD |
| 1009322937 | 2 | STRd | 485 | 26.8051883 | 1.37E-05 | 3 | 5.11E-07 | 0.111918632 | 8 mo | WILD |
| 1011080637 | 2 | STRd | 485 | 26.8051883 | 0.005699317 | 333 | 0.00021262 | 12.42296813 | 8 mo | HOM |
| 1013609713 | 2 | STRd | 485 | 26.8051883 | 0 | 0 | 0 | 0 | 2 mo | WILD |
| 1010418842 | 1 | STRd | 485 | 26.8051883 | 0.015405613 | 562 | 0.000574725 | 20.96609036 | 8 mo | HOM |
| 1013288213 | 2 | STRd | 485 | 26.8051883 | 0.000157199 | 19 | 5.86E-06 | 0.708818001 | 4 mo | HOM |
| 1008976060 | 2 | STRd | 485 | 26.8051883 | 0.000599894 | 56 | 2.24E-05 | 2.089147794 | 8 mo | WILD |
| 1013446243 | 1 | STRd | 485 | 26.8051883 | 0.000745304 | 50 | 2.78E-05 | 1.86531053 | 4 mo | HOM |
| 1013288951 | 2 | STRd | 485 | 26.8051883 | 0.000918342 | 55 | 3.43E-05 | 2.051841583 | 4 mo | HOM |
| 1013978665 | 1 | ORBl6b | 488 | 0.045055863 | 0 | 0 | 0 | 0 | 2 mo | HOM |
| 1013445609 | 1 | ORBl6b | 488 | 0.045055863 | 0 | 0 | 0 | 0 | 4 mo | HOM |
| 1013609015 | 1 | ORBl6b | 488 | 0.045055863 | 0 | 0 | 0 | 0 | 4 mo | HOM |
| 1013608106 | 2 | ORBl6b | 488 | 0.045055863 | 0 | 0 | 0 | 0 | 4 mo | HOM |
| 1011937543 | 1 | ORBl6b | 488 | 0.045055863 | 1.5E-05 | 0 | 0.000333262 | 0 | 8 mo | HOM |
| 1010419538 | 2 | ORBl6b | 488 | 0.045055863 | 0 | 0 | 0 | 0 | 4 mo | WILD |
| 1010419187 | 1 | ORBl6b | 488 | 0.045055863 | 5.74E-06 | 0 | 0.000127337 | 0 | 8 mo | HOM |
| 1013978366 | 1 | ORBl6b | 488 | 0.045055863 | 0 | 0 | 0 | 0 | 2 mo | HOM |
| 1014416155 | 2 | ORBl6b | 488 | 0.045055863 | 0 | 0 | 0 | 0 | 2 mo | HOM |
| 1013978065 | 2 | ORBl6b | 488 | 0.045055863 | 0 | 0 | 0 | 0 | 2 mo | HOM |
| 1009934102 | 2 | ORBl6b | 488 | 0.045055863 | 0 | 0 | 0 | 0 | 4 mo | WILD |
| 1009322937 | 2 | ORBl6b | 488 | 0.045055863 | 0 | 0 | 0 | 0 | 8 mo | WILD |
| 1011080637 | 2 | ORBl6b | 488 | 0.045055863 | 1.48E-05 | 0 | 0.000328456 | 0 | 8 mo | HOM |
| 1013609713 | 2 | ORBl6b | 488 | 0.045055863 | 0 | 0 | 0 | 0 | 2 mo | WILD |
| 1010418842 | 1 | ORBl6b | 488 | 0.045055863 | 3.12E-05 | 1 | 0.000692484 | 22.19466996 | 8 mo | HOM |
| 1013288213 | 2 | ORBl6b | 488 | 0.045055863 | 0 | 0 | 0 | 0 | 4 mo | HOM |
| 1008976060 | 2 | ORBl6b | 488 | 0.045055863 | 0 | 0 | 0 | 0 | 8 mo | WILD |
| 1013446243 | 1 | ORBl6b | 488 | 0.045055863 | 3.39E-08 | 0 | 7.53E-07 | 0 | 4 mo | HOM |
| 1013288951 | 2 | ORBl6b | 488 | 0.045055863 | 0 | 0 | 0 | 0 | 4 mo | HOM |
| 1013978665 | 1 | MM | 491 | 0.556461608 | 1.41E-06 | 0 | 2.54E-06 | 0 | 2 mo | HOM |
| 1013445609 | 1 | MM | 491 | 0.556461608 | 0 | 0 | 0 | 0 | 4 mo | HOM |
| 1013609015 | 1 | MM | 491 | 0.556461608 | 1.44E-05 | 1 | 2.58E-05 | 1.797069169 | 4 mo | HOM |
| 1013608106 | 2 | MM | 491 | 0.556461608 | 0 | 0 | 0 | 0 | 4 mo | HOM |
| 1011937543 | 1 | MM | 491 | 0.556461608 | 0.001968689 | 10 | 0.00353787 | 17.97069169 | 8 mo | HOM |
| 1010419538 | 2 | MM | 491 | 0.556461608 | 1.79E-06 | 0 | 3.21E-06 | 0 | 4 mo | WILD |
| 1010419187 | 1 | MM | 491 | 0.556461608 | 0.001577333 | 4 | 0.002834576 | 7.188276678 | 8 mo | HOM |
| 1013978366 | 1 | MM | 491 | 0.556461608 | 1.21E-05 | 1 | 2.17E-05 | 1.797069169 | 2 mo | HOM |
| 1014416155 | 2 | MM | 491 | 0.556461608 | 0 | 0 | 0 | 0 | 2 mo | HOM |
| 1013978065 | 2 | MM | 491 | 0.556461608 | 4.08E-06 | 1 | 7.32E-06 | 1.797069169 | 2 mo | HOM |
| 1009934102 | 2 | MM | 491 | 0.556461608 | 7.82E-06 | 1 | 1.41E-05 | 1.797069169 | 4 mo | WILD |
| 1009322937 | 2 | MM | 491 | 0.556461608 | 0 | 0 | 0 | 0 | 8 mo | WILD |
| 1011080637 | 2 | MM | 491 | 0.556461608 | 0.000116763 | 5 | 0.000209831 | 8.985345847 | 8 mo | HOM |
| 1013609713 | 2 | MM | 491 | 0.556461608 | 0 | 0 | 0 | 0 | 2 mo | WILD |
| 1010418842 | 1 | MM | 491 | 0.556461608 | 6.48E-05 | 7 | 0.000116448 | 12.57948419 | 8 mo | HOM |
| 1013288213 | 2 | MM | 491 | 0.556461608 | 9.27E-06 | 2 | 1.67E-05 | 3.594138339 | 4 mo | HOM |
| 1008976060 | 2 | MM | 491 | 0.556461608 | 8.75E-05 | 6 | 0.000157328 | 10.78241502 | 8 mo | WILD |
| 1013446243 | 1 | MM | 491 | 0.556461608 | 0 | 0 | 0 | 0 | 4 mo | HOM |
| 1013288951 | 2 | MM | 491 | 0.556461608 | 3.97E-06 | 1 | 7.13E-06 | 1.797069169 | 4 mo | HOM |
| 1013978665 | 1 | STRv | 493 | 8.915072544 | 1.23E-10 | 0 | 1.38E-11 | 0 | 2 mo | HOM |
| 1013445609 | 1 | STRv | 493 | 8.915072662 | 2.1E-05 | 3 | 2.36E-06 | 0.336508755 | 4 mo | HOM |
| 1013609015 | 1 | STRv | 493 | 8.915072662 | 0.000186399 | 9 | 2.09E-05 | 1.009526264 | 4 mo | HOM |
| 1013608106 | 2 | STRv | 493 | 8.915072662 | 0 | 0 | 0 | 0 | 4 mo | HOM |
| 1011937543 | 1 | STRv | 493 | 8.915072662 | 0.005858414 | 127 | 0.000657136 | 14.24553729 | 8 mo | HOM |
| 1010419538 | 2 | STRv | 493 | 8.915072662 | 5.33E-06 | 1 | 5.98E-07 | 0.112169585 | 4 mo | WILD |
| 1010419187 | 1 | STRv | 493 | 8.915072662 | 0.003611572 | 176 | 0.000405109 | 19.74184695 | 8 mo | HOM |
| 1013978366 | 1 | STRv | 493 | 8.915072662 | 0 | 0 | 0 | 0 | 2 mo | HOM |

| | | | | | | | | | | |
|---|---|---|---|---|---|---|---|---|---|---|
| 1014416155 | 2 | STRv | 493 | 8.915072662 | 0 | 0 | 0 | 0 | 2 mo | HOM |
| 1013978065 | 2 | STRv | 493 | 8.915072662 | 0 | 0 | 0 | 0 | 2 mo | HOM |
| 1009934102 | 2 | STRv | 493 | 8.915072662 | 9.72E-07 | 0 | 1.09E-07 | 0 | 4 mo | WILD |
| 1009322937 | 2 | STRv | 493 | 8.915072662 | 0 | 0 | 0 | 0 | 8 mo | WILD |
| 1011080637 | 2 | STRv | 493 | 8.915072662 | 0.001900891 | 113 | 0.000213222 | 12.6751631 | 8 mo | HOM |
| 1013609713 | 2 | STRv | 493 | 8.915072662 | 0 | 0 | 0 | 0 | 2 mo | WILD |
| 1010418842 | 1 | STRv | 493 | 8.915072662 | 0.003485567 | 180 | 0.000390975 | 20.19052529 | 8 mo | HOM |
| 1013288213 | 2 | STRv | 493 | 8.915072662 | 8.83E-05 | 4 | 9.91E-06 | 0.44867834 | 4 mo | HOM |
| 1008976060 | 2 | STRv | 493 | 8.915072662 | 2.1E-05 | 4 | 2.35E-06 | 0.44867834 | 8 mo | WILD |
| 1013446243 | 1 | STRv | 493 | 8.915072662 | 8.85E-05 | 10 | 9.93E-06 | 1.121695849 | 4 mo | HOM |
| 1013288951 | 2 | STRv | 493 | 8.915072662 | 0.000111039 | 6 | 1.25E-05 | 0.67301751 | 4 mo | HOM |
| 1013978665 | 1 | MO | 500 | 25.18937428 | 0.000300012 | 19 | 1.19E-05 | 0.754286303 | 2 mo | HOM |
| 1013445609 | 1 | MO | 500 | 25.18937428 | 0.001825111 | 116 | 7.25E-05 | 4.605116377 | 4 mo | HOM |
| 1013609015 | 1 | MO | 500 | 25.18937428 | 0.005623036 | 204 | 0.00022323 | 8.098652938 | 4 mo | HOM |
| 1013608106 | 2 | MO | 500 | 25.18937428 | 0 | 0 | 0 | 0 | 4 mo | HOM |
| 1011937543 | 1 | MO | 500 | 25.18937428 | 0.032471223 | 633 | 0.001289084 | 25.12964368 | 8 mo | HOM |
| 1010419538 | 2 | MO | 500 | 25.18937428 | 7.09E-05 | 4 | 2.81E-06 | 0.158797116 | 4 mo | WILD |
| 1010419187 | 1 | MO | 500 | 25.18937428 | 0.034812061 | 1301 | 0.001382014 | 51.64876212 | 8 mo | HOM |
| 1013978366 | 1 | MO | 500 | 25.18937428 | 3.27E-05 | 4 | 1.3E-06 | 0.158797116 | 2 mo | HOM |
| 1014416155 | 2 | MO | 500 | 25.18937428 | 0 | 0 | 0 | 0 | 2 mo | HOM |
| 1013978065 | 2 | MO | 500 | 25.18937428 | 0 | 0 | 0 | 0 | 2 mo | HOM |
| 1009934102 | 2 | MO | 500 | 25.18937428 | 0.000148414 | 5 | 5.89E-06 | 0.198496396 | 4 mo | WILD |
| 1009322937 | 2 | MO | 500 | 25.18937428 | 4.98E-05 | 4 | 1.98E-06 | 0.158797116 | 8 mo | WILD |
| 1011080637 | 2 | MO | 500 | 25.18937428 | 0.014514678 | 685 | 0.000576222 | 27.19400619 | 8 mo | HOM |
| 1013609713 | 2 | MO | 500 | 25.18937428 | 2.17E-05 | 1 | 8.62E-07 | 0.039699279 | 2 mo | WILD |
| 1010418842 | 1 | MO | 500 | 25.18937428 | 0.022178921 | 1034 | 0.000880487 | 41.0490546 | 8 mo | HOM |
| 1013288213 | 2 | MO | 500 | 25.18937428 | 0.001268775 | 87 | 5.04E-05 | 3.453837282 | 4 mo | HOM |
| 1008976060 | 2 | MO | 500 | 25.18937428 | 0.001075086 | 44 | 4.27E-05 | 1.746768281 | 8 mo | WILD |
| 1013446243 | 1 | MO | 500 | 25.18937428 | 0.004320542 | 189 | 0.000171522 | 7.503163752 | 4 mo | HOM |
| 1013288951 | 2 | MO | 500 | 25.18937428 | 0.007101606 | 284 | 0.000281929 | 11.27459527 | 4 mo | HOM |
| 1013978665 | 1 | VISpm4 | 501 | 0.101804772 | 0 | 0 | 0 | 0 | 2 mo | HOM |
| 1013445609 | 1 | VISpm4 | 501 | 0.101804772 | 2.45E-05 | 3 | 0.000240327 | 29.46816689 | 4 mo | HOM |
| 1013609015 | 1 | VISpm4 | 501 | 0.101804772 | 3.43E-05 | 2 | 0.000337083 | 19.6454446 | 4 mo | HOM |
| 1013608106 | 2 | VISpm4 | 501 | 0.101804772 | 0 | 0 | 0 | 0 | 4 mo | HOM |
| 1011937543 | 1 | VISpm4 | 501 | 0.101804772 | 0.000162773 | 5 | 0.001598873 | 49.11361149 | 8 mo | HOM |
| 1010419538 | 2 | VISpm4 | 501 | 0.101804772 | 0 | 0 | 0 | 0 | 4 mo | WILD |
| 1010419187 | 1 | VISpm4 | 501 | 0.101804772 | 0.000416102 | 7 | 0.004087258 | 68.75905608 | 8 mo | HOM |
| 1013978366 | 1 | VISpm4 | 501 | 0.101804772 | 0 | 0 | 0 | 0 | 2 mo | HOM |
| 1014416155 | 2 | VISpm4 | 501 | 0.101804772 | 0 | 0 | 0 | 0 | 2 mo | HOM |
| 1013978065 | 2 | VISpm4 | 501 | 0.101804772 | 0 | 0 | 0 | 0 | 2 mo | HOM |
| 1009934102 | 2 | VISpm4 | 501 | 0.101804772 | 0 | 0 | 0 | 0 | 4 mo | WILD |
| 1009322937 | 2 | VISpm4 | 501 | 0.101804772 | 0 | 0 | 0 | 0 | 8 mo | WILD |
| 1011080637 | 2 | VISpm4 | 501 | 0.101804772 | 5.54E-05 | 5 | 0.00054377 | 49.11361149 | 8 mo | HOM |
| 1013609713 | 2 | VISpm4 | 501 | 0.101804772 | 0 | 0 | 0 | 0 | 2 mo | WILD |
| 1010418842 | 1 | VISpm4 | 501 | 0.101804772 | 0.000323931 | 4 | 0.003181881 | 39.29088919 | 8 mo | HOM |
| 1013288213 | 2 | VISpm4 | 501 | 0.101804772 | 6.77E-05 | 4 | 0.000664636 | 39.29088919 | 4 mo | HOM |
| 1008976060 | 2 | VISpm4 | 501 | 0.101804772 | 0 | 0 | 0 | 0 | 8 mo | WILD |
| 1013446243 | 1 | VISpm4 | 501 | 0.101804772 | 5.86E-06 | 1 | 5.75E-05 | 9.822722298 | 4 mo | HOM |
| 1013288951 | 2 | VISpm4 | 501 | 0.101804772 | 1.05E-05 | 1 | 0.000103063 | 9.822722298 | 4 mo | HOM |
| 1013978665 | 1 | SUB | 502 | 2.161199802 | 0 | 0 | 0 | 0 | 2 mo | HOM |
| 1013445609 | 1 | SUB | 502 | 2.161199802 | 6.31E-05 | 8 | 2.92E-05 | 3.701647573 | 4 mo | HOM |
| 1013609015 | 1 | SUB | 502 | 2.161199802 | 0.000155354 | 9 | 7.19E-05 | 4.164353519 | 4 mo | HOM |
| 1013608106 | 2 | SUB | 502 | 2.161199802 | 0 | 0 | 0 | 0 | 4 mo | HOM |
| 1011937543 | 1 | SUB | 502 | 2.161199802 | 0.002837616 | 36 | 0.001312982 | 16.65741408 | 8 mo | HOM |
| 1010419538 | 2 | SUB | 502 | 2.161199802 | 1.25E-06 | 0 | 5.77E-07 | 0 | 4 mo | WILD |
| 1010419187 | 1 | SUB | 502 | 2.161199802 | 0.004603152 | 113 | 0.002129906 | 52.28577197 | 8 mo | HOM |
| 1013978366 | 1 | SUB | 502 | 2.161199802 | 0.000174278 | 5 | 8.06E-05 | 2.313529733 | 2 mo | HOM |
| 1014416155 | 2 | SUB | 502 | 2.161199802 | 0 | 0 | 0 | 0 | 2 mo | HOM |
| 1013978065 | 2 | SUB | 502 | 2.161199802 | 0 | 0 | 0 | 0 | 2 mo | HOM |

| | | | | | | | | | | |
|---|---|---|---|---|---|---|---|---|---|---|
| **1009934102** | 2 | SUB | 502 | 2.161199802 | 0 | 0 | 0 | 0 | 4 mo | WILD |
| **1009322937** | 2 | SUB | 502 | 2.161199802 | 0 | 0 | 0 | 0 | 8 mo | WILD |
| **1011080637** | 2 | SUB | 502 | 2.161199802 | 0.002954242 | 94 | 0.001366945 | 43.49435898 | 8 mo | HOM |
| **1013609713** | 2 | SUB | 502 | 2.161199802 | 0 | 0 | 0 | 0 | 2 mo | WILD |
| **1010418842** | 1 | SUB | 502 | 2.161199802 | 0.004346443 | 102 | 0.002011125 | 47.19600655 | 8 mo | HOM |
| **1013288213** | 2 | SUB | 502 | 2.161199802 | 6.68E-05 | 7 | 3.09E-05 | 3.238941626 | 4 mo | HOM |
| **1008976060** | 2 | SUB | 502 | 2.161199802 | 4.42E-05 | 6 | 2.05E-05 | 2.77623568 | 8 mo | WILD |
| **1013446243** | 1 | SUB | 502 | 2.161199802 | 2.22E-05 | 2 | 1.03E-05 | 0.925411893 | 4 mo | HOM |
| **1013288951** | 2 | SUB | 502 | 2.161199802 | 8.96E-05 | 6 | 4.15E-05 | 2.77623568 | 4 mo | HOM |
| **1013978665** | 1 | das | 506 | 0.012204045 | 8.63E-09 | 0 | 7.08E-07 | 0 | 2 mo | HOM |
| **1013445609** | 1 | das | 506 | 0.012204045 | 0 | 0 | 0 | 0 | 4 mo | HOM |
| **1013609015** | 1 | das | 506 | 0.012204045 | 1.64E-07 | 0 | 1.34E-05 | 0 | 4 mo | HOM |
| **1013608106** | 2 | das | 506 | 0.012204045 | 0 | 0 | 0 | 0 | 4 mo | HOM |
| **1011937543** | 1 | das | 506 | 0.012204045 | 0 | 0 | 0 | 0 | 8 mo | HOM |
| **1010419538** | 2 | das | 506 | 0.012204045 | 0 | 0 | 0 | 0 | 4 mo | WILD |
| **1010419187** | 1 | das | 506 | 0.012204045 | 0 | 0 | 0 | 0 | 8 mo | HOM |
| **1013978366** | 1 | das | 506 | 0.012204045 | 0 | 0 | 0 | 0 | 2 mo | HOM |
| **1014416155** | 2 | das | 506 | 0.012204045 | 0 | 0 | 0 | 0 | 2 mo | HOM |
| **1013978065** | 2 | das | 506 | 0.012204045 | 0 | 0 | 0 | 0 | 2 mo | HOM |
| **1009934102** | 2 | das | 506 | 0.012204045 | 0 | 0 | 0 | 0 | 4 mo | WILD |
| **1009322937** | 2 | das | 506 | 0.012204045 | 4.9E-06 | 0 | 0.000401235 | 0 | 8 mo | WILD |
| **1011080637** | 2 | das | 506 | 0.012204045 | 0 | 0 | 0 | 0 | 8 mo | HOM |
| **1013609713** | 2 | das | 506 | 0.012204045 | 0 | 0 | 0 | 0 | 2 mo | WILD |
| **1010418842** | 1 | das | 506 | 0.012204045 | 4.25E-07 | 0 | 3.48E-05 | 0 | 8 mo | HOM |
| **1013288213** | 2 | das | 506 | 0.012204045 | 0 | 0 | 0 | 0 | 4 mo | HOM |
| **1008976060** | 2 | das | 506 | 0.012204045 | 2.62E-06 | 0 | 0.000214651 | 0 | 8 mo | WILD |
| **1013446243** | 1 | das | 506 | 0.012204045 | 0 | 0 | 0 | 0 | 4 mo | HOM |
| **1013288951** | 2 | das | 506 | 0.012204045 | 0 | 0 | 0 | 0 | 4 mo | HOM |
| **1013978665** | 1 | MOB | 507 | 15.46618627 | 0.004715497 | 209 | 0.000304891 | 13.51335076 | 2 mo | HOM |
| **1013445609** | 1 | MOB | 507 | 15.57542569 | 0.002067471 | 210 | 0.000132739 | 13.48277756 | 4 mo | HOM |
| **1013609015** | 1 | MOB | 507 | 13.4618796 | 0.015611067 | 260 | 0.00115965 | 19.31379626 | 4 mo | HOM |
| **1013608106** | 2 | MOB | 507 | 15.26649833 | 0 | 0 | 0 | 0 | 4 mo | HOM |
| **1011937543** | 1 | MOB | 507 | 15.95590653 | 0.016027014 | 132 | 0.001004457 | 8.272798522 | 8 mo | HOM |
| **1010419538** | 2 | MOB | 507 | 16.32539633 | 0.000690646 | 52 | 4.23E-05 | 3.185221293 | 4 mo | WILD |
| **1010419187** | 1 | MOB | 507 | 12.66055258 | 0.019071664 | 279 | 0.001506385 | 22.03695284 | 8 mo | HOM |
| **1013978366** | 1 | MOB | 507 | 16.96193408 | 0.003293204 | 200 | 0.000194153 | 11.7911082 | 2 mo | HOM |
| **1014416155** | 2 | MOB | 507 | 16.37244636 | 2.65E-05 | 2 | 1.62E-06 | 0.122156455 | 2 mo | HOM |
| **1013978065** | 2 | MOB | 507 | 16.96193408 | 0.001533435 | 118 | 9.04E-05 | 6.956753837 | 2 mo | HOM |
| **1009934102** | 2 | MOB | 507 | 14.66295757 | 0.001634896 | 120 | 0.000111498 | 8.183887831 | 4 mo | WILD |
| **1009322937** | 2 | MOB | 507 | 12.31511967 | 0.000296804 | 30 | 2.41E-05 | 2.436029922 | 8 mo | WILD |
| **1011080637** | 2 | MOB | 507 | 11.93800814 | 0.006603404 | 226 | 0.000553141 | 18.93113134 | 8 mo | HOM |
| **1013609713** | 2 | MOB | 507 | 16.96193408 | 0.000311877 | 32 | 1.84E-05 | 1.886577312 | 2 mo | WILD |
| **1010418842** | 1 | MOB | 507 | 14.40886129 | 0.013898157 | 385 | 0.000964556 | 26.71966871 | 8 mo | HOM |
| **1013288213** | 2 | MOB | 507 | 15.94949404 | 0.006393385 | 130 | 0.000400852 | 8.150728773 | 4 mo | HOM |
| **1008976060** | 2 | MOB | 507 | 16.74958548 | 0.00683009 | 228 | 0.000407777 | 13.61227716 | 8 mo | WILD |
| **1013446243** | 1 | MOB | 507 | 15.07992515 | 0.004228525 | 209 | 0.000280408 | 13.85948524 | 4 mo | HOM |
| **1013288951** | 2 | MOB | 507 | 14.81035333 | 0.007187834 | 224 | 0.000485325 | 15.12455476 | 4 mo | HOM |
| **1013978665** | 1 | SSp-ll6b | 510 | 0.05132993 | 0 | 0 | 0 | 0 | 2 mo | HOM |
| **1013445609** | 1 | SSp-ll6b | 510 | 0.05132993 | 9.32E-07 | 0 | 1.82E-05 | 0 | 4 mo | HOM |
| **1013609015** | 1 | SSp-ll6b | 510 | 0.05132993 | 0 | 0 | 0 | 0 | 4 mo | HOM |
| **1013608106** | 2 | SSp-ll6b | 510 | 0.05132993 | 0 | 0 | 0 | 0 | 4 mo | HOM |
| **1011937543** | 1 | SSp-ll6b | 510 | 0.05132993 | 9.88E-05 | 3 | 0.001925341 | 58.44543284 | 8 mo | HOM |
| **1010419538** | 2 | SSp-ll6b | 510 | 0.05132993 | 0 | 0 | 0 | 0 | 4 mo | WILD |
| **1010419187** | 1 | SSp-ll6b | 510 | 0.05132993 | 4.02E-05 | 3 | 0.000783811 | 58.44543284 | 8 mo | HOM |
| **1013978366** | 1 | SSp-ll6b | 510 | 0.05132993 | 0 | 0 | 0 | 0 | 2 mo | HOM |
| **1014416155** | 2 | SSp-ll6b | 510 | 0.05132993 | 0 | 0 | 0 | 0 | 2 mo | HOM |
| **1013978065** | 2 | SSp-ll6b | 510 | 0.05132993 | 0 | 0 | 0 | 0 | 2 mo | HOM |
| **1009934102** | 2 | SSp-ll6b | 510 | 0.05132993 | 0 | 0 | 0 | 0 | 4 mo | WILD |
| **1009322937** | 2 | SSp-ll6b | 510 | 0.05132993 | 0 | 0 | 0 | 0 | 8 mo | WILD |

| | | | | | | | | | | |
|---|---|---|---|---|---|---|---|---|---|---|
| **1011080637** | 2 | SSp-Il6b | 510 | 0.05132993 | 2.04E-08 | 0 | 3.98E-07 | 0 | 8 mo | HOM |
| **1013609713** | 2 | SSp-Il6b | 510 | 0.05132993 | 0 | 0 | 0 | 0 | 2 mo | WILD |
| **1010418842** | 1 | SSp-Il6b | 510 | 0.05132993 | 0.000111943 | 2 | 0.002180852 | 38.96362189 | 8 mo | HOM |
| **1013288213** | 2 | SSp-Il6b | 510 | 0.05132993 | 0 | 0 | 0 | 0 | 4 mo | HOM |
| **1008976060** | 2 | SSp-Il6b | 510 | 0.05132993 | 5.18E-07 | 0 | 1.01E-05 | 0 | 8 mo | WILD |
| **1013446243** | 1 | SSp-Il6b | 510 | 0.05132993 | 4.44E-06 | 0 | 8.64E-05 | 0 | 4 mo | HOM |
| **1013288951** | 2 | SSp-Il6b | 510 | 0.05132993 | 0 | 0 | 0 | 0 | 4 mo | HOM |
| **1013978665** | 1 | CB | 512 | 55.46977874 | 0.004468899 | 190 | 8.06E-05 | 3.425288586 | 2 mo | HOM |
| **1013445609** | 1 | CB | 512 | 55.46977874 | 0.001090898 | 29 | 1.97E-05 | 0.522807205 | 4 mo | HOM |
| **1013609015** | 1 | CB | 512 | 55.46977874 | 0.01682533 | 282 | 0.000303324 | 5.083849375 | 4 mo | HOM |
| **1013608106** | 2 | CB | 512 | 55.46977874 | 0 | 0 | 0 | 0 | 4 mo | HOM |
| **1011937543** | 1 | CB | 512 | 55.46977874 | 0.086741221 | 335 | 0.001563756 | 6.039324613 | 8 mo | HOM |
| **1010419538** | 2 | CB | 512 | 55.3935614 | 0.001720933 | 86 | 3.11E-05 | 1.552527006 | 4 mo | WILD |
| **1010419187** | 1 | CB | 512 | 55.46977874 | 0.055405946 | 658 | 0.000998849 | 11.86231521 | 8 mo | HOM |
| **1013978366** | 1 | CB | 512 | 55.46977874 | 0.017080933 | 333 | 0.000307932 | 6.003268943 | 2 mo | HOM |
| **1014416155** | 2 | CB | 512 | 55.46977874 | 0 | 0 | 0 | 0 | 2 mo | HOM |
| **1013978065** | 2 | CB | 512 | 55.46977874 | 0.001767801 | 61 | 3.19E-05 | 1.099697915 | 2 mo | HOM |
| **1009934102** | 2 | CB | 512 | 55.46977874 | 0.001987887 | 109 | 3.58E-05 | 1.965033978 | 4 mo | WILD |
| **1009322937** | 2 | CB | 512 | 55.46977874 | 0.003261443 | 158 | 5.88E-05 | 2.848397877 | 8 mo | WILD |
| **1011080637** | 2 | CB | 512 | 55.46977874 | 0.057254805 | 699 | 0.00103218 | 12.60145643 | 8 mo | HOM |
| **1013609713** | 2 | CB | 512 | 55.36351601 | 0.001414769 | 54 | 2.56E-05 | 0.975371579 | 2 mo | WILD |
| **1010418842** | 1 | CB | 512 | 55.46977874 | 0.086181964 | 924 | 0.001553674 | 16.65771923 | 8 mo | HOM |
| **1013288213** | 2 | CB | 512 | 55.46977874 | 0.001453405 | 67 | 2.62E-05 | 1.207864923 | 4 mo | HOM |
| **1008976060** | 2 | CB | 512 | 55.46977874 | 0.015495427 | 361 | 0.000279349 | 6.508048314 | 8 mo | WILD |
| **1013446243** | 1 | CB | 512 | 55.46977874 | 0.003380643 | 116 | 6.09E-05 | 2.091228821 | 4 mo | HOM |
| **1013288951** | 2 | CB | 512 | 55.46977874 | 0.022348082 | 229 | 0.000402888 | 4.128374138 | 4 mo | HOM |
| **1013978665** | 1 | dc | 514 | 0.02298638 | 0 | 0 | 0 | 0 | 2 mo | HOM |
| **1013445609** | 1 | dc | 514 | 0.02298638 | 0 | 0 | 0 | 0 | 4 mo | HOM |
| **1013609015** | 1 | dc | 514 | 0.02298638 | 0 | 0 | 0 | 0 | 4 mo | HOM |
| **1013608106** | 2 | dc | 514 | 0.02298638 | 0 | 0 | 0 | 0 | 4 mo | HOM |
| **1011937543** | 1 | dc | 514 | 0.02298638 | 0 | 0 | 0 | 0 | 8 mo | HOM |
| **1010419538** | 2 | dc | 514 | 0.02298638 | 0 | 0 | 0 | 0 | 4 mo | WILD |
| **1010419187** | 1 | dc | 514 | 0.02298638 | 0 | 0 | 0 | 0 | 8 mo | HOM |
| **1013978366** | 1 | dc | 514 | 0.02298638 | 0 | 0 | 0 | 0 | 2 mo | HOM |
| **1014416155** | 2 | dc | 514 | 0.02298638 | 0 | 0 | 0 | 0 | 2 mo | HOM |
| **1013978065** | 2 | dc | 514 | 0.02298638 | 0 | 0 | 0 | 0 | 2 mo | HOM |
| **1009934102** | 2 | dc | 514 | 0.02298638 | 0 | 0 | 0 | 0 | 4 mo | WILD |
| **1009322937** | 2 | dc | 514 | 0.02298638 | 0 | 0 | 0 | 0 | 8 mo | WILD |
| **1011080637** | 2 | dc | 514 | 0.02298638 | 0 | 0 | 0 | 0 | 8 mo | HOM |
| **1013609713** | 2 | dc | 514 | 0.02298638 | 0 | 0 | 0 | 0 | 2 mo | WILD |
| **1010418842** | 1 | dc | 514 | 0.02298638 | 0 | 0 | 0 | 0 | 8 mo | HOM |
| **1013288213** | 2 | dc | 514 | 0.02298638 | 0 | 0 | 0 | 0 | 4 mo | HOM |
| **1008976060** | 2 | dc | 514 | 0.02298638 | 0 | 0 | 0 | 0 | 8 mo | WILD |
| **1013446243** | 1 | dc | 514 | 0.02298638 | 0 | 0 | 0 | 0 | 4 mo | HOM |
| **1013288951** | 2 | dc | 514 | 0.02298638 | 0 | 0 | 0 | 0 | 4 mo | HOM |
| **1013978665** | 1 | MPN | 515 | 0.418704059 | 0 | 0 | 0 | 0 | 2 mo | HOM |
| **1013445609** | 1 | MPN | 515 | 0.418704059 | 0 | 0 | 0 | 0 | 4 mo | HOM |
| **1013609015** | 1 | MPN | 515 | 0.418704059 | 0 | 0 | 0 | 0 | 4 mo | HOM |
| **1013608106** | 2 | MPN | 515 | 0.418704059 | 0 | 0 | 0 | 0 | 4 mo | HOM |
| **1011937543** | 1 | MPN | 515 | 0.418704059 | 2.45E-05 | 1 | 5.85E-05 | 2.388321723 | 8 mo | HOM |
| **1010419538** | 2 | MPN | 515 | 0.418704059 | 0 | 0 | 0 | 0 | 4 mo | WILD |
| **1010419187** | 1 | MPN | 515 | 0.418704059 | 9.81E-05 | 6 | 0.000234326 | 14.32993034 | 8 mo | HOM |
| **1013978366** | 1 | MPN | 515 | 0.418704059 | 0 | 0 | 0 | 0 | 2 mo | HOM |
| **1014416155** | 2 | MPN | 515 | 0.418704059 | 0 | 0 | 0 | 0 | 2 mo | HOM |
| **1013978065** | 2 | MPN | 515 | 0.418704059 | 0 | 0 | 0 | 0 | 2 mo | HOM |
| **1009934102** | 2 | MPN | 515 | 0.418704059 | 0 | 0 | 0 | 0 | 4 mo | WILD |
| **1009322937** | 2 | MPN | 515 | 0.418704059 | 0 | 0 | 0 | 0 | 8 mo | WILD |
| **1011080637** | 2 | MPN | 515 | 0.418704059 | 3.43E-05 | 4 | 8.2E-05 | 9.553286891 | 8 mo | HOM |
| **1013609713** | 2 | MPN | 515 | 0.418704059 | 0 | 0 | 0 | 0 | 2 mo | WILD |

| | | | | | | | | | | |
|---|---|---|---|---|---|---|---|---|---|---|
| **1010418842** | 1 | MPN | 515 | 0.418704059 | 8.81E-05 | 5 | 0.000210471 | 11.94160861 | 8 mo | HOM |
| **1013288213** | 2 | MPN | 515 | 0.418704059 | 0 | 0 | 0 | 0 | 4 mo | HOM |
| **1008976060** | 2 | MPN | 515 | 0.418704059 | 0 | 0 | 0 | 0 | 8 mo | WILD |
| **1013446243** | 1 | MPN | 515 | 0.418704059 | 0 | 0 | 0 | 0 | 4 mo | HOM |
| **1013288951** | 2 | MPN | 515 | 0.418704059 | 0 | 0 | 0 | 0 | 4 mo | HOM |
| **1013978665** | 1 | CBN | 519 | 1.835902179 | 0 | 0 | 0 | 0 | 2 mo | HOM |
| **1013445609** | 1 | CBN | 519 | 1.835902179 | 1.21E-07 | 0 | 6.59E-08 | 0 | 4 mo | HOM |
| **1013609015** | 1 | CBN | 519 | 1.835902179 | 0 | 0 | 0 | 0 | 4 mo | HOM |
| **1013608106** | 2 | CBN | 519 | 1.835902179 | 0 | 0 | 0 | 0 | 4 mo | HOM |
| **1011937543** | 1 | CBN | 519 | 1.835902179 | 0.00093931 | 10 | 0.000511634 | 5.446913303 | 8 mo | HOM |
| **1010419538** | 2 | CBN | 519 | 1.79529874 | 1.16E-05 | 1 | 6.44E-06 | 0.557010361 | 4 mo | WILD |
| **1010419187** | 1 | CBN | 519 | 1.835902179 | 2.55E-05 | 1 | 1.39E-05 | 0.54469133 | 8 mo | HOM |
| **1013978366** | 1 | CBN | 519 | 1.835902179 | 1.99E-06 | 0 | 1.09E-06 | 0 | 2 mo | HOM |
| **1014416155** | 2 | CBN | 519 | 1.835902179 | 0 | 0 | 0 | 0 | 2 mo | HOM |
| **1013978065** | 2 | CBN | 519 | 1.835902179 | 0 | 0 | 0 | 0 | 2 mo | HOM |
| **1009934102** | 2 | CBN | 519 | 1.835902179 | 0 | 0 | 0 | 0 | 4 mo | WILD |
| **1009322937** | 2 | CBN | 519 | 1.835902179 | 0 | 0 | 0 | 0 | 8 mo | WILD |
| **1011080637** | 2 | CBN | 519 | 1.835902179 | 0.000113822 | 3 | 6.2E-05 | 1.634073991 | 8 mo | HOM |
| **1013609713** | 2 | CBN | 519 | 1.835902179 | 0 | 0 | 0 | 0 | 2 mo | WILD |
| **1010418842** | 1 | CBN | 519 | 1.835902179 | 4.05E-05 | 0 | 2.21E-05 | 0 | 8 mo | HOM |
| **1013288213** | 2 | CBN | 519 | 1.835902179 | 1.06E-05 | 1 | 5.75E-06 | 0.54469133 | 4 mo | HOM |
| **1008976060** | 2 | CBN | 519 | 1.835902179 | 0.000160791 | 14 | 8.76E-05 | 7.625678624 | 8 mo | WILD |
| **1013446243** | 1 | CBN | 519 | 1.835902179 | 0 | 0 | 0 | 0 | 4 mo | HOM |
| **1013288951** | 2 | CBN | 519 | 1.835902179 | 3.3E-05 | 0 | 1.8E-05 | 0 | 4 mo | HOM |
| **1013978665** | 1 | AUDv6a | 520 | 0.28959831 | 0 | 0 | 0 | 0 | 2 mo | HOM |
| **1013445609** | 1 | AUDv6a | 520 | 0.28959831 | 3.44E-05 | 2 | 0.000118829 | 6.90611764 | 4 mo | HOM |
| **1013609015** | 1 | AUDv6a | 520 | 0.28959831 | 5.36E-05 | 2 | 0.000185022 | 6.90611764 | 4 mo | HOM |
| **1013608106** | 2 | AUDv6a | 520 | 0.28959831 | 0 | 0 | 0 | 0 | 4 mo | HOM |
| **1011937543** | 1 | AUDv6a | 520 | 0.28959831 | 0.000346324 | 12 | 0.001195877 | 41.43670584 | 8 mo | HOM |
| **1010419538** | 2 | AUDv6a | 520 | 0.28959831 | 0 | 0 | 0 | 0 | 4 mo | WILD |
| **1010419187** | 1 | AUDv6a | 520 | 0.28959831 | 0.000284002 | 16 | 0.000980675 | 55.24894112 | 8 mo | HOM |
| **1013978366** | 1 | AUDv6a | 520 | 0.28959831 | 0 | 0 | 0 | 0 | 2 mo | HOM |
| **1014416155** | 2 | AUDv6a | 520 | 0.28959831 | 0 | 0 | 0 | 0 | 2 mo | HOM |
| **1013978065** | 2 | AUDv6a | 520 | 0.28959831 | 0 | 0 | 0 | 0 | 2 mo | HOM |
| **1009934102** | 2 | AUDv6a | 520 | 0.28959831 | 0 | 0 | 0 | 0 | 4 mo | WILD |
| **1009322937** | 2 | AUDv6a | 520 | 0.28959831 | 0 | 0 | 0 | 0 | 8 mo | WILD |
| **1011080637** | 2 | AUDv6a | 520 | 0.28959831 | 0.000279426 | 10 | 0.000964873 | 34.5305882 | 8 mo | HOM |
| **1013609713** | 2 | AUDv6a | 520 | 0.28959831 | 0 | 0 | 0 | 0 | 2 mo | WILD |
| **1010418842** | 1 | AUDv6a | 520 | 0.28959831 | 0.000621672 | 14 | 0.002146669 | 48.34282348 | 8 mo | HOM |
| **1013288213** | 2 | AUDv6a | 520 | 0.28959831 | 0 | 0 | 0 | 0 | 4 mo | HOM |
| **1008976060** | 2 | AUDv6a | 520 | 0.28959831 | 0 | 0 | 0 | 0 | 8 mo | WILD |
| **1013446243** | 1 | AUDv6a | 520 | 0.28959831 | 4.18E-05 | 4 | 0.000144403 | 13.81223528 | 4 mo | HOM |
| **1013288951** | 2 | AUDv6a | 520 | 0.28959831 | 3.12E-05 | 3 | 0.000107853 | 10.35917646 | 4 mo | HOM |
| **1013978665** | 1 | MPO | 523 | 0.572320902 | 0 | 0 | 0 | 0 | 2 mo | HOM |
| **1013445609** | 1 | MPO | 523 | 0.572320902 | 6.79E-06 | 1 | 1.19E-05 | 1.7472715 | 4 mo | HOM |
| **1013609015** | 1 | MPO | 523 | 0.572320902 | 0 | 0 | 0 | 0 | 4 mo | HOM |
| **1013608106** | 2 | MPO | 523 | 0.572320902 | 0 | 0 | 0 | 0 | 4 mo | HOM |
| **1011937543** | 1 | MPO | 523 | 0.572320902 | 0.000154396 | 4 | 0.000269772 | 6.989085998 | 8 mo | HOM |
| **1010419538** | 2 | MPO | 523 | 0.572320902 | 0 | 0 | 0 | 0 | 4 mo | WILD |
| **1010419187** | 1 | MPO | 523 | 0.572320902 | 0.000228305 | 10 | 0.000398911 | 17.472715 | 8 mo | HOM |
| **1013978366** | 1 | MPO | 523 | 0.572320902 | 0 | 0 | 0 | 0 | 2 mo | HOM |
| **1014416155** | 2 | MPO | 523 | 0.572320902 | 0 | 0 | 0 | 0 | 2 mo | HOM |
| **1013978065** | 2 | MPO | 523 | 0.572320902 | 0 | 0 | 0 | 0 | 2 mo | HOM |
| **1009934102** | 2 | MPO | 523 | 0.572320902 | 0 | 0 | 0 | 0 | 4 mo | WILD |
| **1009322937** | 2 | MPO | 523 | 0.572320902 | 0 | 0 | 0 | 0 | 8 mo | WILD |
| **1011080637** | 2 | MPO | 523 | 0.572320902 | 0.00022972 | 9 | 0.000401383 | 15.7254435 | 8 mo | HOM |
| **1013609713** | 2 | MPO | 523 | 0.572320902 | 0 | 0 | 0 | 0 | 2 mo | WILD |
| **1010418842** | 1 | MPO | 523 | 0.572320902 | 0.00047168 | 11 | 0.000824153 | 19.2199865 | 8 mo | HOM |
| **1013288213** | 2 | MPO | 523 | 0.572320902 | 0 | 0 | 0 | 0 | 4 mo | HOM |

| | | | | | | | | | | |
|---|---|---|---|---|---|---|---|---|---|---|
| **1008976060** | 2 | MPO | 523 | 0.572320902 | 0 | 0 | 0 | 0 | 8 mo | WILD |
| **1013446243** | 1 | MPO | 523 | 0.572320902 | 0 | 0 | 0 | 0 | 4 mo | HOM |
| **1013288951** | 2 | MPO | 523 | 0.572320902 | 0 | 0 | 0 | 0 | 4 mo | HOM |
| **1013978665** | 1 | SUM | 525 | 0.26242097 | 1.1E-07 | 0 | 4.19E-07 | 0 | 2 mo | HOM |
| **1013445609** | 1 | SUM | 525 | 0.26242097 | 0 | 0 | 0 | 0 | 4 mo | HOM |
| **1013609015** | 1 | SUM | 525 | 0.26242097 | 0 | 0 | 0 | 0 | 4 mo | HOM |
| **1013608106** | 2 | SUM | 525 | 0.26242097 | 0 | 0 | 0 | 0 | 4 mo | HOM |
| **1011937543** | 1 | SUM | 525 | 0.26242097 | 0.000511309 | 3 | 0.001948429 | 11.43201323 | 8 mo | HOM |
| **1010419538** | 2 | SUM | 525 | 0.26242097 | 7.91E-07 | 0 | 3.01E-06 | 0 | 4 mo | WILD |
| **1010419187** | 1 | SUM | 525 | 0.26242097 | 0.000513608 | 1 | 0.00195719 | 3.810671076 | 8 mo | HOM |
| **1013978366** | 1 | SUM | 525 | 0.26242097 | 0 | 0 | 0 | 0 | 2 mo | HOM |
| **1014416155** | 2 | SUM | 525 | 0.26242097 | 0 | 0 | 0 | 0 | 2 mo | HOM |
| **1013978065** | 2 | SUM | 525 | 0.26242097 | 0 | 0 | 0 | 0 | 2 mo | HOM |
| **1009934102** | 2 | SUM | 525 | 0.26242097 | 0 | 0 | 0 | 0 | 4 mo | WILD |
| **1009322937** | 2 | SUM | 525 | 0.26242097 | 5.57E-11 | 0 | 2.12E-10 | 0 | 8 mo | WILD |
| **1011080637** | 2 | SUM | 525 | 0.26242097 | 0 | 0 | 0 | 0 | 8 mo | HOM |
| **1013609713** | 2 | SUM | 525 | 0.26242097 | 0 | 0 | 0 | 0 | 2 mo | WILD |
| **1010418842** | 1 | SUM | 525 | 0.26242097 | 5.79E-05 | 0 | 0.000220462 | 0 | 8 mo | HOM |
| **1013288213** | 2 | SUM | 525 | 0.26242097 | 0 | 0 | 0 | 0 | 4 mo | HOM |
| **1008976060** | 2 | SUM | 525 | 0.26242097 | 3.06E-05 | 3 | 0.000116734 | 11.43201323 | 8 mo | WILD |
| **1013446243** | 1 | SUM | 525 | 0.26242097 | 0 | 0 | 0 | 0 | 4 mo | HOM |
| **1013288951** | 2 | SUM | 525 | 0.26242097 | 0 | 0 | 0 | 0 | 4 mo | HOM |
| **1013978665** | 1 | ENTm1 | 526 | 1.283763387 | 1.39E-05 | 2 | 1.08E-05 | 1.557919489 | 2 mo | HOM |
| **1013445609** | 1 | ENTm1 | 526 | 1.283763387 | 0.000198293 | 9 | 0.000154462 | 7.0106377 | 4 mo | HOM |
| **1013609015** | 1 | ENTm1 | 526 | 1.283763387 | 0.002434886 | 29 | 0.001896678 | 22.58983259 | 4 mo | HOM |
| **1013608106** | 2 | ENTm1 | 526 | 1.283763387 | 0 | 0 | 0 | 0 | 4 mo | HOM |
| **1011937543** | 1 | ENTm1 | 526 | 1.283763387 | 0.008822149 | 44 | 0.006872098 | 34.27422875 | 8 mo | HOM |
| **1010419538** | 2 | ENTm1 | 526 | 1.283763387 | 0.000141363 | 14 | 0.000110116 | 10.90543642 | 4 mo | WILD |
| **1010419187** | 1 | ENTm1 | 526 | 1.283763387 | 0.008178651 | 101 | 0.00637084 | 78.67493419 | 8 mo | HOM |
| **1013978366** | 1 | ENTm1 | 526 | 1.283763387 | 0.000407792 | 21 | 0.000317654 | 16.35815463 | 2 mo | HOM |
| **1014416155** | 2 | ENTm1 | 526 | 1.283763387 | 0 | 0 | 0 | 0 | 2 mo | HOM |
| **1013978065** | 2 | ENTm1 | 526 | 1.283763387 | 2.29E-05 | 2 | 1.79E-05 | 1.557919489 | 2 mo | HOM |
| **1009934102** | 2 | ENTm1 | 526 | 1.283763387 | 0.000485125 | 32 | 0.000377893 | 24.92671182 | 4 mo | WILD |
| **1009322937** | 2 | ENTm1 | 526 | 1.283763387 | 0.000259139 | 11 | 0.000201859 | 8.568557189 | 8 mo | WILD |
| **1011080637** | 2 | ENTm1 | 526 | 1.283763387 | 0.003064251 | 92 | 0.002386928 | 71.66429649 | 8 mo | HOM |
| **1013609713** | 2 | ENTm1 | 526 | 1.283763387 | 0.000129652 | 12 | 0.000100994 | 9.347516933 | 2 mo | WILD |
| **1010418842** | 1 | ENTm1 | 526 | 1.283763387 | 0.00173917 | 55 | 0.001354743 | 42.84278594 | 8 mo | HOM |
| **1013288213** | 2 | ENTm1 | 526 | 1.283763387 | 0.000245867 | 16 | 0.00019152 | 12.46335591 | 4 mo | HOM |
| **1008976060** | 2 | ENTm1 | 526 | 1.283763387 | 8.13E-05 | 9 | 6.34E-05 | 7.0106377 | 8 mo | WILD |
| **1013446243** | 1 | ENTm1 | 526 | 1.283763387 | 0.000221577 | 12 | 0.0001726 | 9.347516933 | 4 mo | HOM |
| **1013288951** | 2 | ENTm1 | 526 | 1.283763387 | 0.004946503 | 34 | 0.003853127 | 26.48463131 | 4 mo | HOM |
| **1013978665** | 1 | AUDd1 | 527 | 0.20284924 | 4.82E-06 | 1 | 2.38E-05 | 4.929769527 | 2 mo | HOM |
| **1013445609** | 1 | AUDd1 | 527 | 0.20284924 | 1.63E-05 | 1 | 8.02E-05 | 4.929769527 | 4 mo | HOM |
| **1013609015** | 1 | AUDd1 | 527 | 0.20284924 | 1.58E-05 | 2 | 7.79E-05 | 9.859539054 | 4 mo | HOM |
| **1013608106** | 2 | AUDd1 | 527 | 0.20284924 | 0 | 0 | 0 | 0 | 4 mo | HOM |
| **1011937543** | 1 | AUDd1 | 527 | 0.20284924 | 0.000914371 | 8 | 0.004507637 | 39.43815622 | 8 mo | HOM |
| **1010419538** | 2 | AUDd1 | 527 | 0.20284924 | 2.02E-06 | 0 | 9.96E-06 | 0 | 4 mo | WILD |
| **1010419187** | 1 | AUDd1 | 527 | 0.20284924 | 0.000430522 | 10 | 0.002122372 | 49.29769527 | 8 mo | HOM |
| **1013978366** | 1 | AUDd1 | 527 | 0.20284924 | 8.56E-06 | 1 | 4.22E-05 | 4.929769527 | 2 mo | HOM |
| **1014416155** | 2 | AUDd1 | 527 | 0.20284924 | 0 | 0 | 0 | 0 | 2 mo | HOM |
| **1013978065** | 2 | AUDd1 | 527 | 0.20284924 | 0 | 0 | 0 | 0 | 2 mo | HOM |
| **1009934102** | 2 | AUDd1 | 527 | 0.20284924 | 1.24E-06 | 0 | 6.12E-06 | 0 | 4 mo | WILD |
| **1009322937** | 2 | AUDd1 | 527 | 0.20284924 | 1.33E-06 | 1 | 6.54E-06 | 4.929769527 | 8 mo | WILD |
| **1011080637** | 2 | AUDd1 | 527 | 0.20284924 | 0.000309809 | 11 | 0.001527289 | 54.2274648 | 8 mo | HOM |
| **1013609713** | 2 | AUDd1 | 527 | 0.20284924 | 2.6E-06 | 0 | 1.28E-05 | 0 | 2 mo | WILD |
| **1010418842** | 1 | AUDd1 | 527 | 0.20284924 | 0.000759281 | 12 | 0.003743079 | 59.15723433 | 8 mo | HOM |
| **1013288213** | 2 | AUDd1 | 527 | 0.20284924 | 6.15E-05 | 4 | 0.000303023 | 19.71907811 | 4 mo | HOM |
| **1008976060** | 2 | AUDd1 | 527 | 0.20284924 | 6.8E-06 | 1 | 3.35E-05 | 4.929769527 | 8 mo | WILD |
| **1013446243** | 1 | AUDd1 | 527 | 0.20284924 | 5.69E-05 | 4 | 0.000280564 | 19.71907811 | 4 mo | HOM |

| | | | | | | | | | | |
|---|---|---|---|---|---|---|---|---|---|---|
| **1013288951** | 2 | AUDd1 | 527 | 0.20284924 | 0.000478652 | 10 | 0.002359642 | 49.29769527 | 4 mo | HOM |
| **1013978665** | 1 | CBX | 528 | 52.77954742 | 0.004385574 | 190 | 8.31E-05 | 3.599879296 | 2 mo | HOM |
| **1013445609** | 1 | CBX | 528 | 52.77954742 | 0.001057416 | 26 | 2E-05 | 0.492615062 | 4 mo | HOM |
| **1013609015** | 1 | CBX | 528 | 52.77954742 | 0.016527039 | 281 | 0.000313133 | 5.324032011 | 4 mo | HOM |
| **1013608106** | 2 | CBX | 528 | 52.77954742 | 0 | 0 | 0 | 0 | 4 mo | HOM |
| **1011937543** | 1 | CBX | 528 | 52.77954742 | 0.084953474 | 320 | 0.001609591 | 6.062954604 | 8 mo | HOM |
| **1010419538** | 2 | CBX | 528 | 52.7439323 | 0.001673697 | 83 | 3.17E-05 | 1.573640728 | 4 mo | WILD |
| **1010419187** | 1 | CBX | 528 | 52.77954742 | 0.054099587 | 645 | 0.00102501 | 12.22064287 | 8 mo | HOM |
| **1013978366** | 1 | CBX | 528 | 52.77954742 | 0.016647815 | 325 | 0.000315422 | 6.157688269 | 2 mo | HOM |
| **1014416155** | 2 | CBX | 528 | 52.77954742 | 0 | 0 | 0 | 0 | 2 mo | HOM |
| **1013978065** | 2 | CBX | 528 | 52.77954742 | 0.001695301 | 58 | 3.21E-05 | 1.098910522 | 2 mo | HOM |
| **1009934102** | 2 | CBX | 528 | 52.77954742 | 0.001945901 | 108 | 3.69E-05 | 2.046247179 | 4 mo | WILD |
| **1009322937** | 2 | CBX | 528 | 52.77954742 | 0.003196878 | 155 | 6.06E-05 | 2.936743636 | 8 mo | WILD |
| **1011080637** | 2 | CBX | 528 | 52.77954742 | 0.05593195 | 681 | 0.001059728 | 12.90272527 | 8 mo | HOM |
| **1013609713** | 2 | CBX | 528 | 52.67328884 | 0.001386011 | 54 | 2.63E-05 | 1.025187551 | 2 mo | WILD |
| **1010418842** | 1 | CBX | 528 | 52.77954742 | 0.08428647 | 914 | 0.001596953 | 17.31731409 | 8 mo | HOM |
| **1013288213** | 2 | CBX | 528 | 52.77954742 | 0.001228001 | 59 | 2.33E-05 | 1.117857255 | 4 mo | HOM |
| **1008976060** | 2 | CBX | 528 | 52.77954742 | 0.014749972 | 332 | 0.000279464 | 6.290315401 | 8 mo | WILD |
| **1013446243** | 1 | CBX | 528 | 52.77954742 | 0.003342914 | 110 | 6.33E-05 | 2.084140645 | 4 mo | HOM |
| **1013288951** | 2 | CBX | 528 | 52.77954742 | 0.021697939 | 227 | 0.000411105 | 4.300908422 | 4 mo | HOM |
| **1013978665** | 1 | df | 530 | 0.038664344 | 0 | 0 | 0 | 0 | 2 mo | HOM |
| **1013445609** | 1 | df | 530 | 0.038664344 | 0 | 0 | 0 | 0 | 4 mo | HOM |
| **1013609015** | 1 | df | 530 | 0.038664344 | 0 | 0 | 0 | 0 | 4 mo | HOM |
| **1013608106** | 2 | df | 530 | 0.038664344 | 0 | 0 | 0 | 0 | 4 mo | HOM |
| **1011937543** | 1 | df | 530 | 0.038664344 | 7.7E-06 | 1 | 0.000199095 | 25.8636224 | 8 mo | HOM |
| **1010419538** | 2 | df | 530 | 0.038664344 | 0 | 0 | 0 | 0 | 4 mo | WILD |
| **1010419187** | 1 | df | 530 | 0.038664344 | 1.94E-05 | 1 | 0.000500525 | 25.8636224 | 8 mo | HOM |
| **1013978366** | 1 | df | 530 | 0.038664344 | 0 | 0 | 0 | 0 | 2 mo | HOM |
| **1014416155** | 2 | df | 530 | 0.038664344 | 0 | 0 | 0 | 0 | 2 mo | HOM |
| **1013978065** | 2 | df | 530 | 0.038664344 | 0 | 0 | 0 | 0 | 2 mo | HOM |
| **1009934102** | 2 | df | 530 | 0.038664344 | 0 | 0 | 0 | 0 | 4 mo | WILD |
| **1009322937** | 2 | df | 530 | 0.038664344 | 0 | 0 | 0 | 0 | 8 mo | WILD |
| **1011080637** | 2 | df | 530 | 0.038664344 | 2.28E-07 | 0 | 5.89E-06 | 0 | 8 mo | HOM |
| **1013609713** | 2 | df | 530 | 0.038664344 | 0 | 0 | 0 | 0 | 2 mo | WILD |
| **1010418842** | 1 | df | 530 | 0.038664344 | 6.89E-06 | 0 | 0.000178274 | 0 | 8 mo | HOM |
| **1013288213** | 2 | df | 530 | 0.038664344 | 4.64E-05 | 1 | 0.001199472 | 25.8636224 | 4 mo | HOM |
| **1008976060** | 2 | df | 530 | 0.038664344 | 0 | 0 | 0 | 0 | 8 mo | WILD |
| **1013446243** | 1 | df | 530 | 0.038664344 | 0 | 0 | 0 | 0 | 4 mo | HOM |
| **1013288951** | 2 | df | 530 | 0.038664344 | 0 | 0 | 0 | 0 | 4 mo | HOM |
| **1013978665** | 1 | MPT | 531 | 0.045111492 | 0 | 0 | 0 | 0 | 2 mo | HOM |
| **1013445609** | 1 | MPT | 531 | 0.045111492 | 0 | 0 | 0 | 0 | 4 mo | HOM |
| **1013609015** | 1 | MPT | 531 | 0.045111492 | 0 | 0 | 0 | 0 | 4 mo | HOM |
| **1013608106** | 2 | MPT | 531 | 0.045111492 | 0 | 0 | 0 | 0 | 4 mo | HOM |
| **1011937543** | 1 | MPT | 531 | 0.045111492 | 3.19E-05 | 1 | 0.000707179 | 22.16730037 | 8 mo | HOM |
| **1010419538** | 2 | MPT | 531 | 0.045111492 | 2.45E-07 | 0 | 5.43E-06 | 0 | 4 mo | WILD |
| **1010419187** | 1 | MPT | 531 | 0.045111492 | 1.65E-10 | 0 | 3.66E-09 | 0 | 8 mo | HOM |
| **1013978366** | 1 | MPT | 531 | 0.045111492 | 0 | 0 | 0 | 0 | 2 mo | HOM |
| **1014416155** | 2 | MPT | 531 | 0.045111492 | 0 | 0 | 0 | 0 | 2 mo | HOM |
| **1013978065** | 2 | MPT | 531 | 0.045111492 | 0 | 0 | 0 | 0 | 2 mo | HOM |
| **1009934102** | 2 | MPT | 531 | 0.045111492 | 0 | 0 | 0 | 0 | 4 mo | WILD |
| **1009322937** | 2 | MPT | 531 | 0.045111492 | 3.1E-07 | 0 | 6.86E-06 | 0 | 8 mo | WILD |
| **1011080637** | 2 | MPT | 531 | 0.045111492 | 1.51E-10 | 0 | 3.34E-09 | 0 | 8 mo | HOM |
| **1013609713** | 2 | MPT | 531 | 0.045111492 | 0 | 0 | 0 | 0 | 2 mo | WILD |
| **1010418842** | 1 | MPT | 531 | 0.045111492 | 0 | 0 | 0 | 0 | 8 mo | HOM |
| **1013288213** | 2 | MPT | 531 | 0.045111492 | 0 | 0 | 0 | 0 | 4 mo | HOM |
| **1008976060** | 2 | MPT | 531 | 0.045111492 | 1.09E-07 | 0 | 2.42E-06 | 0 | 8 mo | WILD |
| **1013446243** | 1 | MPT | 531 | 0.045111492 | 0 | 0 | 0 | 0 | 4 mo | HOM |
| **1013288951** | 2 | MPT | 531 | 0.045111492 | 2.01E-06 | 0 | 4.45E-05 | 0 | 4 mo | HOM |
| **1013978665** | 1 | VISpm | 533 | 1.080462902 | 5.31E-06 | 1 | 4.92E-06 | 0.925529232 | 2 mo | HOM |

| | | | | | | | | | | |
|---|---|---|---|---|---|---|---|---|---|---|
| 1013445609 | 1 | VISpm | 533 | 1.080462902 | 0.000156932 | 12 | 0.000145245 | 11.10635079 | 4 mo | HOM |
| 1013609015 | 1 | VISpm | 533 | 1.080462902 | 0.000385701 | 17 | 0.000356977 | 15.73399695 | 4 mo | HOM |
| 1013608106 | 2 | VISpm | 533 | 1.080462902 | 0 | 0 | 0 | 0 | 4 mo | HOM |
| 1011937543 | 1 | VISpm | 533 | 1.080462902 | 0.001866636 | 20 | 0.001727626 | 18.51058465 | 8 mo | HOM |
| 1010419538 | 2 | VISpm | 533 | 1.080462902 | 0 | 0 | 0 | 0 | 4 mo | WILD |
| 1010419187 | 1 | VISpm | 533 | 1.080462902 | 0.002421673 | 61 | 0.002241329 | 56.45728317 | 8 mo | HOM |
| 1013978366 | 1 | VISpm | 533 | 1.080462902 | 0 | 0 | 0 | 0 | 2 mo | HOM |
| 1014416155 | 2 | VISpm | 533 | 1.080462902 | 0 | 0 | 0 | 0 | 2 mo | HOM |
| 1013978065 | 2 | VISpm | 533 | 1.080462902 | 0 | 0 | 0 | 0 | 2 mo | HOM |
| 1009934102 | 2 | VISpm | 533 | 1.080462902 | 0 | 0 | 0 | 0 | 4 mo | WILD |
| 1009322937 | 2 | VISpm | 533 | 1.080462902 | 5.52E-06 | 1 | 5.11E-06 | 0.925529232 | 8 mo | WILD |
| 1011080637 | 2 | VISpm | 533 | 1.080462902 | 0.000857562 | 38 | 0.000793699 | 35.17011083 | 8 mo | HOM |
| 1013609713 | 2 | VISpm | 533 | 1.080462902 | 0 | 0 | 0 | 0 | 2 mo | WILD |
| 1010418842 | 1 | VISpm | 533 | 1.080462902 | 0.002108046 | 47 | 0.001951058 | 43.49987392 | 8 mo | HOM |
| 1013288213 | 2 | VISpm | 533 | 1.080462902 | 0.000111912 | 9 | 0.000103577 | 8.329763091 | 4 mo | HOM |
| 1008976060 | 2 | VISpm | 533 | 1.080462902 | 0.000720046 | 5 | 0.000666423 | 4.627646162 | 8 mo | WILD |
| 1013446243 | 1 | VISpm | 533 | 1.080462902 | 0.000531346 | 14 | 0.000491776 | 12.95740925 | 4 mo | HOM |
| 1013288951 | 2 | VISpm | 533 | 1.080462902 | 9.75E-05 | 6 | 9.02E-05 | 5.553175394 | 4 mo | HOM |
| 1013978665 | 1 | SUT | 534 | 0.26744641 | 0 | 0 | 0 | 0 | 2 mo | HOM |
| 1013445609 | 1 | SUT | 534 | 0.26744641 | 0 | 0 | 0 | 0 | 4 mo | HOM |
| 1013609015 | 1 | SUT | 534 | 0.26744641 | 0 | 0 | 0 | 0 | 4 mo | HOM |
| 1013608106 | 2 | SUT | 534 | 0.26744641 | 0 | 0 | 0 | 0 | 4 mo | HOM |
| 1011937543 | 1 | SUT | 534 | 0.26744641 | 6.7E-05 | 4 | 0.000250686 | 14.95626731 | 8 mo | HOM |
| 1010419538 | 2 | SUT | 534 | 0.26744641 | 0 | 0 | 0 | 0 | 4 mo | WILD |
| 1010419187 | 1 | SUT | 534 | 0.26744641 | 0 | 0 | 0 | 0 | 8 mo | HOM |
| 1013978366 | 1 | SUT | 534 | 0.26744641 | 0 | 0 | 0 | 0 | 2 mo | HOM |
| 1014416155 | 2 | SUT | 534 | 0.26744641 | 0 | 0 | 0 | 0 | 2 mo | HOM |
| 1013978065 | 2 | SUT | 534 | 0.26744641 | 0 | 0 | 0 | 0 | 2 mo | HOM |
| 1009934102 | 2 | SUT | 534 | 0.26744641 | 0 | 0 | 0 | 0 | 4 mo | WILD |
| 1009322937 | 2 | SUT | 534 | 0.26744641 | 0 | 0 | 0 | 0 | 8 mo | WILD |
| 1011080637 | 2 | SUT | 534 | 0.26744641 | 0 | 0 | 0 | 0 | 8 mo | HOM |
| 1013609713 | 2 | SUT | 534 | 0.26744641 | 0 | 0 | 0 | 0 | 2 mo | WILD |
| 1010418842 | 1 | SUT | 534 | 0.26744641 | 1.44E-05 | 1 | 5.39E-05 | 3.739066828 | 8 mo | HOM |
| 1013288213 | 2 | SUT | 534 | 0.26744641 | 0 | 0 | 0 | 0 | 4 mo | HOM |
| 1008976060 | 2 | SUT | 534 | 0.26744641 | 0 | 0 | 0 | 0 | 8 mo | WILD |
| 1013446243 | 1 | SUT | 534 | 0.26744641 | 0 | 0 | 0 | 0 | 4 mo | HOM |
| 1013288951 | 2 | SUT | 534 | 0.26744641 | 0 | 0 | 0 | 0 | 4 mo | HOM |
| 1013978665 | 1 | CEA | 536 | 1.356806296 | 0 | 0 | 0 | 0 | 2 mo | HOM |
| 1013445609 | 1 | CEA | 536 | 1.356806296 | 0.000111456 | 4 | 8.21E-05 | 2.948099528 | 4 mo | HOM |
| 1013609015 | 1 | CEA | 536 | 1.356806296 | 0.000182955 | 9 | 0.000134842 | 6.633223939 | 4 mo | HOM |
| 1013608106 | 2 | CEA | 536 | 1.356806296 | 0 | 0 | 0 | 0 | 4 mo | HOM |
| 1011937543 | 1 | CEA | 536 | 1.356806296 | 0.002986773 | 45 | 0.002201326 | 33.16611969 | 8 mo | HOM |
| 1010419538 | 2 | CEA | 536 | 1.356806296 | 0 | 0 | 0 | 0 | 4 mo | WILD |
| 1010419187 | 1 | CEA | 536 | 1.356806296 | 0.001915016 | 67 | 0.001411414 | 49.3806671 | 8 mo | HOM |
| 1013978366 | 1 | CEA | 536 | 1.356806296 | 0 | 0 | 0 | 0 | 2 mo | HOM |
| 1014416155 | 2 | CEA | 536 | 1.356806296 | 0 | 0 | 0 | 0 | 2 mo | HOM |
| 1013978065 | 2 | CEA | 536 | 1.356806296 | 4.81E-06 | 1 | 3.55E-06 | 0.737024882 | 2 mo | HOM |
| 1009934102 | 2 | CEA | 536 | 1.356806296 | 0 | 0 | 0 | 0 | 4 mo | WILD |
| 1009322937 | 2 | CEA | 536 | 1.356806296 | 0 | 0 | 0 | 0 | 8 mo | WILD |
| 1011080637 | 2 | CEA | 536 | 1.356806296 | 0.001717854 | 57 | 0.001266101 | 42.01041828 | 8 mo | HOM |
| 1013609713 | 2 | CEA | 536 | 1.356806296 | 0 | 0 | 0 | 0 | 2 mo | WILD |
| 1010418842 | 1 | CEA | 536 | 1.356806296 | 0.002658065 | 67 | 0.00195906 | 49.3806671 | 8 mo | HOM |
| 1013288213 | 2 | CEA | 536 | 1.356806296 | 4.19E-05 | 3 | 3.09E-05 | 2.211074646 | 4 mo | HOM |
| 1008976060 | 2 | CEA | 536 | 1.356806296 | 4.53E-06 | 1 | 3.34E-06 | 0.737024882 | 8 mo | WILD |
| 1013446243 | 1 | CEA | 536 | 1.356806296 | 0.000227012 | 12 | 0.000167314 | 8.844298585 | 4 mo | HOM |
| 1013288951 | 2 | CEA | 536 | 1.356806296 | 0.000160754 | 10 | 0.00011848 | 7.370248821 | 4 mo | HOM |
| 1013978665 | 1 | lotd | 538 | 0.162495329 | 8.51E-07 | 0 | 5.24E-06 | 0 | 2 mo | HOM |
| 1013445609 | 1 | lotd | 538 | 0.162495329 | 0 | 0 | 0 | 0 | 4 mo | HOM |
| 1013609015 | 1 | lotd | 538 | 0.162495329 | 0 | 0 | 0 | 0 | 4 mo | HOM |

| | | | | | | | | | | |
|---|---|---|---|---|---|---|---|---|---|---|
| **1013608106** | 2 | lotd | 538 | 0.162495329 | 0 | 0 | 0 | 0 | 4 mo | HOM |
| **1011937543** | 1 | lotd | 538 | 0.162495329 | 0.000188592 | 1 | 0.001160597 | 6.154023056 | 8 mo | HOM |
| **1010419538** | 2 | lotd | 538 | 0.162495329 | 0 | 0 | 0 | 0 | 4 mo | WILD |
| **1010419187** | 1 | lotd | 538 | 0.162495329 | 8.76E-05 | 1 | 0.000539131 | 6.154023056 | 8 mo | HOM |
| **1013978366** | 1 | lotd | 538 | 0.162495329 | 0 | 0 | 0 | 0 | 2 mo | HOM |
| **1014416155** | 2 | lotd | 538 | 0.162495329 | 0 | 0 | 0 | 0 | 2 mo | HOM |
| **1013978065** | 2 | lotd | 538 | 0.162495329 | 0 | 0 | 0 | 0 | 2 mo | HOM |
| **1009934102** | 2 | lotd | 538 | 0.162495329 | 2.16E-06 | 0 | 1.33E-05 | 0 | 4 mo | WILD |
| **1009322937** | 2 | lotd | 538 | 0.162495329 | 1.9E-05 | 3 | 0.000117034 | 18.46206917 | 8 mo | WILD |
| **1011080637** | 2 | lotd | 538 | 0.162495329 | 7.84E-06 | 1 | 4.82E-05 | 6.154023056 | 8 mo | HOM |
| **1013609713** | 2 | lotd | 538 | 0.162495329 | 0 | 0 | 0 | 0 | 2 mo | WILD |
| **1010418842** | 1 | lotd | 538 | 0.162495329 | 0.000108445 | 2 | 0.000667374 | 12.30804611 | 8 mo | HOM |
| **1013288213** | 2 | lotd | 538 | 0.162495329 | 0.000457686 | 2 | 0.00281661 | 12.30804611 | 4 mo | HOM |
| **1008976060** | 2 | lotd | 538 | 0.162495329 | 3.42E-09 | 0 | 2.1E-08 | 0 | 8 mo | WILD |
| **1013446243** | 1 | lotd | 538 | 0.162495329 | 0 | 0 | 0 | 0 | 4 mo | HOM |
| **1013288951** | 2 | lotd | 538 | 0.162495329 | 2.59E-05 | 1 | 0.000159541 | 6.154023056 | 4 mo | HOM |
| **1013978665** | 1 | PERI1 | 540 | 0.227953762 | 0 | 0 | 0 | 0 | 2 mo | HOM |
| **1013445609** | 1 | PERI1 | 540 | 0.227953762 | 1.97E-05 | 2 | 8.66E-05 | 8.773709102 | 4 mo | HOM |
| **1013609015** | 1 | PERI1 | 540 | 0.227953762 | 6.98E-06 | 0 | 3.06E-05 | 0 | 4 mo | HOM |
| **1013608106** | 2 | PERI1 | 540 | 0.227953762 | 0 | 0 | 0 | 0 | 4 mo | HOM |
| **1011937543** | 1 | PERI1 | 540 | 0.227953762 | 0.000933124 | 5 | 0.004093478 | 21.93427275 | 8 mo | HOM |
| **1010419538** | 2 | PERI1 | 540 | 0.227953762 | 0 | 0 | 0 | 0 | 4 mo | WILD |
| **1010419187** | 1 | PERI1 | 540 | 0.227953762 | 0.000313334 | 14 | 0.001374552 | 61.41596371 | 8 mo | HOM |
| **1013978366** | 1 | PERI1 | 540 | 0.227953762 | 1.55E-06 | 0 | 6.81E-06 | 0 | 2 mo | HOM |
| **1014416155** | 2 | PERI1 | 540 | 0.227953762 | 0 | 0 | 0 | 0 | 2 mo | HOM |
| **1013978065** | 2 | PERI1 | 540 | 0.227953762 | 0 | 0 | 0 | 0 | 2 mo | HOM |
| **1009934102** | 2 | PERI1 | 540 | 0.227953762 | 1.28E-06 | 0 | 5.6E-06 | 0 | 4 mo | WILD |
| **1009322937** | 2 | PERI1 | 540 | 0.227953762 | 0 | 0 | 0 | 0 | 8 mo | WILD |
| **1011080637** | 2 | PERI1 | 540 | 0.227953762 | 6.77E-05 | 3 | 0.000296851 | 13.16056365 | 8 mo | HOM |
| **1013609713** | 2 | PERI1 | 540 | 0.227953762 | 0 | 0 | 0 | 0 | 2 mo | WILD |
| **1010418842** | 1 | PERI1 | 540 | 0.227953762 | 9.58E-05 | 6 | 0.000420074 | 26.32112731 | 8 mo | HOM |
| **1013288213** | 2 | PERI1 | 540 | 0.227953762 | 2.28E-07 | 0 | 1E-06 | 0 | 4 mo | HOM |
| **1008976060** | 2 | PERI1 | 540 | 0.227953762 | 7.11E-06 | 0 | 3.12E-05 | 0 | 8 mo | WILD |
| **1013446243** | 1 | PERI1 | 540 | 0.227953762 | 6.54E-06 | 0 | 2.87E-05 | 0 | 4 mo | HOM |
| **1013288951** | 2 | PERI1 | 540 | 0.227953762 | 1.21E-05 | 1 | 5.31E-05 | 4.386854551 | 4 mo | HOM |
| **1013978665** | 1 | TEa | 541 | 3.196048461 | 2.41E-07 | 0 | 7.55E-08 | 0 | 2 mo | HOM |
| **1013445609** | 1 | TEa | 541 | 3.196048461 | 0.000325122 | 20 | 0.000101726 | 6.257727391 | 4 mo | HOM |
| **1013609015** | 1 | TEa | 541 | 3.196048461 | 0.001164858 | 34 | 0.000364468 | 10.63813657 | 4 mo | HOM |
| **1013608106** | 2 | TEa | 541 | 3.196048461 | 0 | 0 | 0 | 0 | 4 mo | HOM |
| **1011937543** | 1 | TEa | 541 | 3.196048461 | 0.008472178 | 110 | 0.002650829 | 34.41750065 | 8 mo | HOM |
| **1010419538** | 2 | TEa | 541 | 3.196048461 | 8.25E-06 | 1 | 2.58E-06 | 0.31288637 | 4 mo | WILD |
| **1010419187** | 1 | TEa | 541 | 3.196048461 | 0.006582588 | 194 | 0.002059602 | 60.6999557 | 8 mo | HOM |
| **1013978366** | 1 | TEa | 541 | 3.196048461 | 0 | 0 | 0 | 0 | 2 mo | HOM |
| **1014416155** | 2 | TEa | 541 | 3.196048461 | 0 | 0 | 0 | 0 | 2 mo | HOM |
| **1013978065** | 2 | TEa | 541 | 3.196048461 | 0 | 0 | 0 | 0 | 2 mo | HOM |
| **1009934102** | 2 | TEa | 541 | 3.196048461 | 0 | 0 | 0 | 0 | 4 mo | WILD |
| **1009322937** | 2 | TEa | 541 | 3.196048461 | 0 | 0 | 0 | 0 | 8 mo | WILD |
| **1011080637** | 2 | TEa | 541 | 3.196048461 | 0.002655055 | 109 | 0.000830731 | 34.10461428 | 8 mo | HOM |
| **1013609713** | 2 | TEa | 541 | 3.196048461 | 9.31E-06 | 2 | 2.91E-06 | 0.625772739 | 2 mo | WILD |
| **1010418842** | 1 | TEa | 541 | 3.196048461 | 0.005072537 | 160 | 0.001587128 | 50.06181913 | 8 mo | HOM |
| **1013288213** | 2 | TEa | 541 | 3.196048461 | 0.000276697 | 18 | 8.66E-05 | 5.631954652 | 4 mo | HOM |
| **1008976060** | 2 | TEa | 541 | 3.196048461 | 4.91E-05 | 7 | 1.54E-05 | 2.190204587 | 8 mo | WILD |
| **1013446243** | 1 | TEa | 541 | 3.196048461 | 0.000651921 | 35 | 0.000203977 | 10.95102294 | 4 mo | HOM |
| **1013288951** | 2 | TEa | 541 | 3.196048461 | 0.000620412 | 30 | 0.000194119 | 9.386591087 | 4 mo | HOM |
| **1013978665** | 1 | RSPv1 | 542 | 0.976051619 | 3.82E-06 | 0 | 3.92E-06 | 0 | 2 mo | HOM |
| **1013445609** | 1 | RSPv1 | 542 | 0.976051619 | 5.34E-06 | 1 | 5.47E-06 | 1.024535978 | 4 mo | HOM |
| **1013609015** | 1 | RSPv1 | 542 | 0.976051619 | 0.000136357 | 5 | 0.000139703 | 5.122679889 | 4 mo | HOM |
| **1013608106** | 2 | RSPv1 | 542 | 0.976051619 | 0 | 0 | 0 | 0 | 4 mo | HOM |
| **1011937543** | 1 | RSPv1 | 542 | 0.976051619 | 0.001300404 | 20 | 0.001332311 | 20.49071955 | 8 mo | HOM |

| | | | | | | | | | | |
|---|---|---|---|---|---|---|---|---|---|---|
| **1010419538** | 2 | RSPv1 | 542 | 0.974991808 | 7.33E-05 | 3 | 7.52E-05 | 3.07694893 | 4 mo | WILD |
| **1010419187** | 1 | RSPv1 | 542 | 0.976051619 | 0.001965331 | 33 | 0.002013552 | 33.80968727 | 8 mo | HOM |
| **1013978366** | 1 | RSPv1 | 542 | 0.976051619 | 0.001795306 | 25 | 0.001839355 | 25.61339944 | 2 mo | HOM |
| **1014416155** | 2 | RSPv1 | 542 | 0.976051619 | 0 | 0 | 0 | 0 | 2 mo | HOM |
| **1013978065** | 2 | RSPv1 | 542 | 0.976051619 | 0 | 0 | 0 | 0 | 2 mo | HOM |
| **1009934102** | 2 | RSPv1 | 542 | 0.976051619 | 0.000228485 | 8 | 0.000234091 | 8.196287822 | 4 mo | WILD |
| **1009322937** | 2 | RSPv1 | 542 | 0.976051619 | 1.84E-05 | 1 | 1.89E-05 | 1.024535978 | 8 mo | WILD |
| **1011080637** | 2 | RSPv1 | 542 | 0.976051619 | 0.002948214 | 37 | 0.003020552 | 37.90783118 | 8 mo | HOM |
| **1013609713** | 2 | RSPv1 | 542 | 0.976051619 | 6.66E-10 | 0 | 6.82E-10 | 0 | 2 mo | WILD |
| **1010418842** | 1 | RSPv1 | 542 | 0.976051619 | 0.001132605 | 43 | 0.001160395 | 44.05504704 | 8 mo | HOM |
| **1013288213** | 2 | RSPv1 | 542 | 0.976051619 | 0.000256627 | 4 | 0.000262924 | 4.098143911 | 4 mo | HOM |
| **1008976060** | 2 | RSPv1 | 542 | 0.976051619 | 0.000254956 | 9 | 0.000261212 | 9.2208238 | 8 mo | WILD |
| **1013446243** | 1 | RSPv1 | 542 | 0.976051619 | 0.000173587 | 3 | 0.000177846 | 3.073607933 | 4 mo | HOM |
| **1013288951** | 2 | RSPv1 | 542 | 0.976051619 | 0.001056851 | 4 | 0.001082782 | 4.098143911 | 4 mo | HOM |
| **1013978665** | 1 | ENTm2 | 543 | 1.293379461 | 0 | 0 | 0 | 0 | 2 mo | HOM |
| **1013445609** | 1 | ENTm2 | 543 | 1.293379461 | 4.4E-05 | 4 | 3.4E-05 | 3.092673203 | 4 mo | HOM |
| **1013609015** | 1 | ENTm2 | 543 | 1.293379461 | 0.000101465 | 2 | 7.84E-05 | 1.546336602 | 4 mo | HOM |
| **1013608106** | 2 | ENTm2 | 543 | 1.293379461 | 0 | 0 | 0 | 0 | 4 mo | HOM |
| **1011937543** | 1 | ENTm2 | 543 | 1.293379461 | 0.001269155 | 24 | 0.00098127 | 18.55603922 | 8 mo | HOM |
| **1010419538** | 2 | ENTm2 | 543 | 1.293379461 | 6.12E-07 | 0 | 4.73E-07 | 0 | 4 mo | WILD |
| **1010419187** | 1 | ENTm2 | 543 | 1.293379461 | 0.001431436 | 35 | 0.001106741 | 27.06089053 | 8 mo | HOM |
| **1013978366** | 1 | ENTm2 | 543 | 1.293379461 | 2.23E-06 | 0 | 1.72E-06 | 0 | 2 mo | HOM |
| **1014416155** | 2 | ENTm2 | 543 | 1.293379461 | 0 | 0 | 0 | 0 | 2 mo | HOM |
| **1013978065** | 2 | ENTm2 | 543 | 1.293379461 | 1.12E-08 | 0 | 8.62E-09 | 0 | 2 mo | HOM |
| **1009934102** | 2 | ENTm2 | 543 | 1.293379461 | 5.23E-06 | 0 | 4.04E-06 | 0 | 4 mo | WILD |
| **1009322937** | 2 | ENTm2 | 543 | 1.293379461 | 3.4E-06 | 0 | 2.63E-06 | 0 | 8 mo | WILD |
| **1011080637** | 2 | ENTm2 | 543 | 1.293379461 | 0.000448083 | 21 | 0.000346444 | 16.23653432 | 8 mo | HOM |
| **1013609713** | 2 | ENTm2 | 543 | 1.293379461 | 1.48E-07 | 0 | 1.15E-07 | 0 | 2 mo | WILD |
| **1010418842** | 1 | ENTm2 | 543 | 1.293379461 | 0.001237095 | 34 | 0.000956483 | 26.28772223 | 8 mo | HOM |
| **1013288213** | 2 | ENTm2 | 543 | 1.293379461 | 2.62E-05 | 2 | 2.02E-05 | 1.546336602 | 4 mo | HOM |
| **1008976060** | 2 | ENTm2 | 543 | 1.293379461 | 6.68E-08 | 0 | 5.16E-08 | 0 | 8 mo | WILD |
| **1013446243** | 1 | ENTm2 | 543 | 1.293379461 | 0.000140598 | 8 | 0.000108706 | 6.185346406 | 4 mo | HOM |
| **1013288951** | 2 | ENTm2 | 543 | 1.293379461 | 8.47E-05 | 6 | 6.55E-05 | 4.639009805 | 4 mo | HOM |
| **1013978665** | 1 | CEAc | 544 | 0.318669187 | 0 | 0 | 0 | 0 | 2 mo | HOM |
| **1013445609** | 1 | CEAc | 544 | 0.318669187 | 1.13E-05 | 0 | 3.55E-05 | 0 | 4 mo | HOM |
| **1013609015** | 1 | CEAc | 544 | 0.318669187 | 4.32E-05 | 4 | 0.000135493 | 12.55220197 | 4 mo | HOM |
| **1013608106** | 2 | CEAc | 544 | 0.318669187 | 0 | 0 | 0 | 0 | 4 mo | HOM |
| **1011937543** | 1 | CEAc | 544 | 0.318669187 | 0.000835708 | 14 | 0.002622494 | 43.9327069 | 8 mo | HOM |
| **1010419538** | 2 | CEAc | 544 | 0.318669187 | 0 | 0 | 0 | 0 | 4 mo | WILD |
| **1010419187** | 1 | CEAc | 544 | 0.318669187 | 0.000616439 | 21 | 0.001934415 | 65.89906035 | 8 mo | HOM |
| **1013978366** | 1 | CEAc | 544 | 0.318669187 | 0 | 0 | 0 | 0 | 2 mo | HOM |
| **1014416155** | 2 | CEAc | 544 | 0.318669187 | 0 | 0 | 0 | 0 | 2 mo | HOM |
| **1013978065** | 2 | CEAc | 544 | 0.318669187 | 0 | 0 | 0 | 0 | 2 mo | HOM |
| **1009934102** | 2 | CEAc | 544 | 0.318669187 | 0 | 0 | 0 | 0 | 4 mo | WILD |
| **1009322937** | 2 | CEAc | 544 | 0.318669187 | 0 | 0 | 0 | 0 | 8 mo | WILD |
| **1011080637** | 2 | CEAc | 544 | 0.318669187 | 0.000428902 | 19 | 0.001345917 | 59.62295937 | 8 mo | HOM |
| **1013609713** | 2 | CEAc | 544 | 0.318669187 | 0 | 0 | 0 | 0 | 2 mo | WILD |
| **1010418842** | 1 | CEAc | 544 | 0.318669187 | 0.000884939 | 20 | 0.002776982 | 62.76100986 | 8 mo | HOM |
| **1013288213** | 2 | CEAc | 544 | 0.318669187 | 3.48E-05 | 2 | 0.00010915 | 6.276100986 | 4 mo | HOM |
| **1008976060** | 2 | CEAc | 544 | 0.318669187 | 4.53E-06 | 1 | 1.42E-05 | 3.138050493 | 8 mo | WILD |
| **1013446243** | 1 | CEAc | 544 | 0.318669187 | 0.000111123 | 5 | 0.00034871 | 15.69025247 | 4 mo | HOM |
| **1013288951** | 2 | CEAc | 544 | 0.318669187 | 8.18E-05 | 5 | 0.000256565 | 15.69025247 | 4 mo | HOM |
| **1013978665** | 1 | RSPd4 | 545 | 2.06E-06 | 0 | 0 | 0 | 0 | 2 mo | HOM |
| **1013445609** | 1 | RSPd4 | 545 | 2.06E-06 | 0 | 0 | 0 | 0 | 4 mo | HOM |
| **1013609015** | 1 | RSPd4 | 545 | 2.06E-06 | 0 | 0 | 0 | 0 | 4 mo | HOM |
| **1013608106** | 2 | RSPd4 | 545 | 2.06E-06 | 0 | 0 | 0 | 0 | 4 mo | HOM |
| **1011937543** | 1 | RSPd4 | 545 | 2.06E-06 | 0 | 0 | 0 | 0 | 8 mo | HOM |
| **1010419538** | 2 | RSPd4 | 545 | 2.06E-06 | 0 | 0 | 0 | 0 | 4 mo | WILD |
| **1010419187** | 1 | RSPd4 | 545 | 2.06E-06 | 0 | 0 | 0 | 0 | 8 mo | HOM |

| | | | | | | | | | | |
|---|---|---|---|---|---|---|---|---|---|---|
| 1013978366 | 1 | RSPd4 | 545 | 2.06E-06 | 0 | 0 | 0 | 0 | 2 mo | HOM |
| 1014416155 | 2 | RSPd4 | 545 | 2.06E-06 | 0 | 0 | 0 | 0 | 2 mo | HOM |
| 1013978065 | 2 | RSPd4 | 545 | 2.06E-06 | 0 | 0 | 0 | 0 | 2 mo | HOM |
| 1009934102 | 2 | RSPd4 | 545 | 2.06E-06 | 0 | 0 | 0 | 0 | 4 mo | WILD |
| 1009322937 | 2 | RSPd4 | 545 | 2.06E-06 | 0 | 0 | 0 | 0 | 8 mo | WILD |
| 1011080637 | 2 | RSPd4 | 545 | 2.06E-06 | 0 | 0 | 0 | 0 | 8 mo | HOM |
| 1013609713 | 2 | RSPd4 | 545 | 2.06E-06 | 0 | 0 | 0 | 0 | 2 mo | WILD |
| 1010418842 | 1 | RSPd4 | 545 | 2.06E-06 | 0 | 0 | 0 | 0 | 8 mo | HOM |
| 1013288213 | 2 | RSPd4 | 545 | 2.06E-06 | 0 | 0 | 0 | 0 | 4 mo | HOM |
| 1008976060 | 2 | RSPd4 | 545 | 2.06E-06 | 0 | 0 | 0 | 0 | 8 mo | WILD |
| 1013446243 | 1 | RSPd4 | 545 | 2.06E-06 | 0 | 0 | 0 | 0 | 4 mo | HOM |
| 1013288951 | 2 | RSPd4 | 545 | 2.06E-06 | 0 | 0 | 0 | 0 | 4 mo | HOM |
| 1013978665 | 1 | TH | 549 | 20.88755881 | 2.18E-08 | 0 | 1.04E-09 | 0 | 2 mo | HOM |
| 1013445609 | 1 | TH | 549 | 20.88755881 | 0.000134328 | 10 | 6.43E-06 | 0.478753888 | 4 mo | HOM |
| 1013609015 | 1 | TH | 549 | 20.88755881 | 0.001399364 | 46 | 6.7E-05 | 2.202267887 | 4 mo | HOM |
| 1013608106 | 2 | TH | 549 | 20.88755756 | 0 | 0 | 0 | 0 | 4 mo | HOM |
| 1011937543 | 1 | TH | 549 | 20.88755881 | 0.019024706 | 268 | 0.000910815 | 12.83060421 | 8 mo | HOM |
| 1010419538 | 2 | TH | 549 | 20.88755756 | 3.97E-05 | 1 | 1.9E-06 | 0.047875392 | 4 mo | WILD |
| 1010419187 | 1 | TH | 549 | 20.88755881 | 0.012034558 | 291 | 0.000576159 | 13.93173815 | 8 mo | HOM |
| 1013978366 | 1 | TH | 549 | 20.88755881 | 1.62E-05 | 3 | 7.77E-07 | 0.143626167 | 2 mo | HOM |
| 1014416155 | 2 | TH | 549 | 20.88755756 | 0 | 0 | 0 | 0 | 2 mo | HOM |
| 1013978065 | 2 | TH | 549 | 20.88755756 | 9.78E-07 | 0 | 4.68E-08 | 0 | 2 mo | HOM |
| 1009934102 | 2 | TH | 549 | 20.88755756 | 5.3E-06 | 1 | 2.54E-07 | 0.047875392 | 4 mo | WILD |
| 1009322937 | 2 | TH | 549 | 20.88755756 | 0.000115559 | 16 | 5.53E-06 | 0.766006267 | 8 mo | WILD |
| 1011080637 | 2 | TH | 549 | 20.88755756 | 0.010825418 | 278 | 0.000518271 | 13.3093589 | 8 mo | HOM |
| 1013609713 | 2 | TH | 549 | 20.88755756 | 0 | 0 | 0 | 0 | 2 mo | WILD |
| 1010418842 | 1 | TH | 549 | 20.88755881 | 0.017287831 | 397 | 0.000827662 | 19.00652937 | 8 mo | HOM |
| 1013288213 | 2 | TH | 549 | 20.88755756 | 0.000107379 | 12 | 5.14E-06 | 0.574504701 | 4 mo | HOM |
| 1008976060 | 2 | TH | 549 | 20.88755756 | 0.001058264 | 70 | 5.07E-05 | 3.35127742 | 8 mo | WILD |
| 1013446243 | 1 | TH | 549 | 20.88755881 | 0.00044382 | 24 | 2.12E-05 | 1.149009332 | 4 mo | HOM |
| 1013288951 | 2 | TH | 549 | 20.88755756 | 0.000787939 | 17 | 3.77E-05 | 0.813881659 | 4 mo | HOM |
| 1013978665 | 1 | CEAI | 551 | 0.267522654 | 0 | 0 | 0 | 0 | 2 mo | HOM |
| 1013445609 | 1 | CEAI | 551 | 0.267522654 | 6.19E-06 | 1 | 2.31E-05 | 3.738001194 | 4 mo | HOM |
| 1013609015 | 1 | CEAI | 551 | 0.267522654 | 5.57E-05 | 1 | 0.000208364 | 3.738001194 | 4 mo | HOM |
| 1013608106 | 2 | CEAI | 551 | 0.267522654 | 0 | 0 | 0 | 0 | 4 mo | HOM |
| 1011937543 | 1 | CEAI | 551 | 0.267522654 | 0.000602592 | 14 | 0.00225249 | 52.33201671 | 8 mo | HOM |
| 1010419538 | 2 | CEAI | 551 | 0.267522654 | 0 | 0 | 0 | 0 | 4 mo | WILD |
| 1010419187 | 1 | CEAI | 551 | 0.267522654 | 0.000202001 | 9 | 0.000755081 | 33.64201074 | 8 mo | HOM |
| 1013978366 | 1 | CEAI | 551 | 0.267522654 | 0 | 0 | 0 | 0 | 2 mo | HOM |
| 1014416155 | 2 | CEAI | 551 | 0.267522654 | 0 | 0 | 0 | 0 | 2 mo | HOM |
| 1013978065 | 2 | CEAI | 551 | 0.267522654 | 0 | 0 | 0 | 0 | 2 mo | HOM |
| 1009934102 | 2 | CEAI | 551 | 0.267522654 | 0 | 0 | 0 | 0 | 4 mo | WILD |
| 1009322937 | 2 | CEAI | 551 | 0.267522654 | 0 | 0 | 0 | 0 | 8 mo | WILD |
| 1011080637 | 2 | CEAI | 551 | 0.267522654 | 0.000463124 | 11 | 0.001731158 | 41.11801313 | 8 mo | HOM |
| 1013609713 | 2 | CEAI | 551 | 0.267522654 | 0 | 0 | 0 | 0 | 2 mo | WILD |
| 1010418842 | 1 | CEAI | 551 | 0.267522654 | 0.000390621 | 9 | 0.001460141 | 33.64201074 | 8 mo | HOM |
| 1013288213 | 2 | CEAI | 551 | 0.267522654 | 6.88E-06 | 1 | 2.57E-05 | 3.738001194 | 4 mo | HOM |
| 1008976060 | 2 | CEAI | 551 | 0.267522654 | 0 | 0 | 0 | 0 | 8 mo | WILD |
| 1013446243 | 1 | CEAI | 551 | 0.267522654 | 3.49E-05 | 3 | 0.000130545 | 11.21400358 | 4 mo | HOM |
| 1013288951 | 2 | CEAI | 551 | 0.267522654 | 3.41E-05 | 2 | 0.000127441 | 7.476002387 | 4 mo | HOM |
| 1013978665 | 1 | sctd | 553 | 0.15059417 | 0 | 0 | 0 | 0 | 2 mo | HOM |
| 1013445609 | 1 | sctd | 553 | 0.15059417 | 0 | 0 | 0 | 0 | 4 mo | HOM |
| 1013609015 | 1 | sctd | 553 | 0.15059417 | 0 | 0 | 0 | 0 | 4 mo | HOM |
| 1013608106 | 2 | sctd | 553 | 0.15059417 | 0 | 0 | 0 | 0 | 4 mo | HOM |
| 1011937543 | 1 | sctd | 553 | 0.15059417 | 0 | 0 | 0 | 0 | 8 mo | HOM |
| 1010419538 | 2 | sctd | 553 | 0.15059417 | 0 | 0 | 0 | 0 | 4 mo | WILD |
| 1010419187 | 1 | sctd | 553 | 0.15059417 | 0 | 0 | 0 | 0 | 8 mo | HOM |
| 1013978366 | 1 | sctd | 553 | 0.15059417 | 0 | 0 | 0 | 0 | 2 mo | HOM |
| 1014416155 | 2 | sctd | 553 | 0.15059417 | 0 | 0 | 0 | 0 | 2 mo | HOM |

| | | | | | | | | | | |
|---|---|---|---|---|---|---|---|---|---|---|
| 1013978065 | 2 | sctd | 553 | 0.15059417 | 0 | 0 | 0 | 0 | 2 mo | HOM |
| 1009934102 | 2 | sctd | 553 | 0.15059417 | 0 | 0 | 0 | 0 | 4 mo | WILD |
| 1009322937 | 2 | sctd | 553 | 0.15059417 | 0 | 0 | 0 | 0 | 8 mo | WILD |
| 1011080637 | 2 | sctd | 553 | 0.15059417 | 0 | 0 | 0 | 0 | 8 mo | HOM |
| 1013609713 | 2 | sctd | 553 | 0.15059417 | 0 | 0 | 0 | 0 | 2 mo | WILD |
| 1010418842 | 1 | sctd | 553 | 0.15059417 | 1.1E-08 | 0 | 7.3E-08 | 0 | 8 mo | HOM |
| 1013288213 | 2 | sctd | 553 | 0.15059417 | 0 | 0 | 0 | 0 | 4 mo | HOM |
| 1008976060 | 2 | sctd | 553 | 0.15059417 | 0 | 0 | 0 | 0 | 8 mo | WILD |
| 1013446243 | 1 | sctd | 553 | 0.15059417 | 0 | 0 | 0 | 0 | 4 mo | HOM |
| 1013288951 | 2 | sctd | 553 | 0.15059417 | 0 | 0 | 0 | 0 | 4 mo | HOM |
| 1013978665 | 1 | ILA2/3 | 556 | 0.1452164 | 0 | 0 | 0 | 0 | 2 mo | HOM |
| 1013445609 | 1 | ILA2/3 | 556 | 0.1452164 | 2.22E-07 | 0 | 1.53E-06 | 0 | 4 mo | HOM |
| 1013609015 | 1 | ILA2/3 | 556 | 0.1452164 | 1.81E-05 | 1 | 0.000124312 | 6.886274553 | 4 mo | HOM |
| 1013608106 | 2 | ILA2/3 | 556 | 0.1452164 | 0 | 0 | 0 | 0 | 4 mo | HOM |
| 1011937543 | 1 | ILA2/3 | 556 | 0.1452164 | 0.000297542 | 4 | 0.002048957 | 27.54509821 | 8 mo | HOM |
| 1010419538 | 2 | ILA2/3 | 556 | 0.1452164 | 0 | 0 | 0 | 0 | 4 mo | WILD |
| 1010419187 | 1 | ILA2/3 | 556 | 0.1452164 | 0.000115147 | 4 | 0.000792934 | 27.54509821 | 8 mo | HOM |
| 1013978366 | 1 | ILA2/3 | 556 | 0.1452164 | 0 | 0 | 0 | 0 | 2 mo | HOM |
| 1014416155 | 2 | ILA2/3 | 556 | 0.1452164 | 0 | 0 | 0 | 0 | 2 mo | HOM |
| 1013978065 | 2 | ILA2/3 | 556 | 0.1452164 | 0 | 0 | 0 | 0 | 2 mo | HOM |
| 1009934102 | 2 | ILA2/3 | 556 | 0.1452164 | 0 | 0 | 0 | 0 | 4 mo | WILD |
| 1009322937 | 2 | ILA2/3 | 556 | 0.1452164 | 5.46E-06 | 1 | 3.76E-05 | 6.886274553 | 8 mo | WILD |
| 1011080637 | 2 | ILA2/3 | 556 | 0.1452164 | 0.000142522 | 4 | 0.000981444 | 27.54509821 | 8 mo | HOM |
| 1013609713 | 2 | ILA2/3 | 556 | 0.1452164 | 0 | 0 | 0 | 0 | 2 mo | WILD |
| 1010418842 | 1 | ILA2/3 | 556 | 0.1452164 | 0.000295978 | 6 | 0.002038184 | 41.31764732 | 8 mo | HOM |
| 1013288213 | 2 | ILA2/3 | 556 | 0.1452164 | 5.04E-05 | 0 | 0.00034681 | 0 | 4 mo | HOM |
| 1008976060 | 2 | ILA2/3 | 556 | 0.1452164 | 0 | 0 | 0 | 0 | 8 mo | WILD |
| 1013446243 | 1 | ILA2/3 | 556 | 0.1452164 | 4.14E-05 | 1 | 0.000284975 | 6.886274553 | 4 mo | HOM |
| 1013288951 | 2 | ILA2/3 | 556 | 0.1452164 | 7.61E-05 | 1 | 0.000524096 | 6.886274553 | 4 mo | HOM |
| 1013978665 | 1 | TM | 557 | 0.147845527 | 0 | 0 | 0 | 0 | 2 mo | HOM |
| 1013445609 | 1 | TM | 557 | 0.147845527 | 0 | 0 | 0 | 0 | 4 mo | HOM |
| 1013609015 | 1 | TM | 557 | 0.147845527 | 5.81E-06 | 1 | 3.93E-05 | 6.763816406 | 4 mo | HOM |
| 1013608106 | 2 | TM | 557 | 0.147845527 | 0 | 0 | 0 | 0 | 4 mo | HOM |
| 1011937543 | 1 | TM | 557 | 0.147845527 | 0.000517241 | 7 | 0.00349852 | 47.34671484 | 8 mo | HOM |
| 1010419538 | 2 | TM | 557 | 0.147845527 | 6.59E-22 | 0 | 4.46E-21 | 0 | 4 mo | WILD |
| 1010419187 | 1 | TM | 557 | 0.147845527 | 9.7E-05 | 2 | 0.000655938 | 13.52763281 | 8 mo | HOM |
| 1013978366 | 1 | TM | 557 | 0.147845527 | 8.84E-07 | 0 | 5.98E-06 | 0 | 2 mo | HOM |
| 1014416155 | 2 | TM | 557 | 0.147845527 | 0 | 0 | 0 | 0 | 2 mo | HOM |
| 1013978065 | 2 | TM | 557 | 0.147845527 | 0 | 0 | 0 | 0 | 2 mo | HOM |
| 1009934102 | 2 | TM | 557 | 0.147845527 | 1.03E-07 | 0 | 6.96E-07 | 0 | 4 mo | WILD |
| 1009322937 | 2 | TM | 557 | 0.147845527 | 0 | 0 | 0 | 0 | 8 mo | WILD |
| 1011080637 | 2 | TM | 557 | 0.147845527 | 2.3E-06 | 0 | 1.55E-05 | 0 | 8 mo | HOM |
| 1013609713 | 2 | TM | 557 | 0.147845527 | 0 | 0 | 0 | 0 | 2 mo | WILD |
| 1010418842 | 1 | TM | 557 | 0.147845527 | 1.77E-05 | 2 | 0.0001199 | 13.52763281 | 8 mo | HOM |
| 1013288213 | 2 | TM | 557 | 0.147845527 | 0 | 0 | 0 | 0 | 4 mo | HOM |
| 1008976060 | 2 | TM | 557 | 0.147845527 | 1.43E-06 | 0 | 9.65E-06 | 0 | 8 mo | WILD |
| 1013446243 | 1 | TM | 557 | 0.147845527 | 0 | 0 | 0 | 0 | 4 mo | HOM |
| 1013288951 | 2 | TM | 557 | 0.147845527 | 1.25E-06 | 0 | 8.48E-06 | 0 | 4 mo | HOM |
| 1013978665 | 1 | SSp-n1 | 558 | 0.416546766 | 5.48E-06 | 1 | 1.32E-05 | 2.400690829 | 2 mo | HOM |
| 1013445609 | 1 | SSp-n1 | 558 | 0.416546766 | 0.000207432 | 6 | 0.00049798 | 14.40414497 | 4 mo | HOM |
| 1013609015 | 1 | SSp-n1 | 558 | 0.416546766 | 0.000261663 | 9 | 0.000628172 | 21.60621746 | 4 mo | HOM |
| 1013608106 | 2 | SSp-n1 | 558 | 0.416546766 | 0 | 0 | 0 | 0 | 4 mo | HOM |
| 1011937543 | 1 | SSp-n1 | 558 | 0.416546766 | 0.0017275 | 19 | 0.004147193 | 45.61312575 | 8 mo | HOM |
| 1010419538 | 2 | SSp-n1 | 558 | 0.416546766 | 0 | 0 | 0 | 0 | 4 mo | WILD |
| 1010419187 | 1 | SSp-n1 | 558 | 0.416546766 | 0.000886005 | 21 | 0.002127024 | 50.41450741 | 8 mo | HOM |
| 1013978366 | 1 | SSp-n1 | 558 | 0.416546766 | 0 | 0 | 0 | 0 | 2 mo | HOM |
| 1014416155 | 2 | SSp-n1 | 558 | 0.416546766 | 0 | 0 | 0 | 0 | 2 mo | HOM |
| 1013978065 | 2 | SSp-n1 | 558 | 0.416546766 | 0 | 0 | 0 | 0 | 2 mo | HOM |
| 1009934102 | 2 | SSp-n1 | 558 | 0.416546766 | 0 | 0 | 0 | 0 | 4 mo | WILD |

| | | | | | | | | | | |
|---|---|---|---|---|---|---|---|---|---|---|
| **1009322937** | 2 | SSp-n1 | 558 | 0.416546766 | 0 | 0 | 0 | 0 | 8 mo | WILD |
| **1011080637** | 2 | SSp-n1 | 558 | 0.416546766 | 0.000545134 | 18 | 0.001308697 | 43.21243492 | 8 mo | HOM |
| **1013609713** | 2 | SSp-n1 | 558 | 0.416546766 | 4.56E-06 | 0 | 1.09E-05 | 0 | 2 mo | WILD |
| **1010418842** | 1 | SSp-n1 | 558 | 0.416546766 | 0.002584854 | 28 | 0.006205435 | 67.21934321 | 8 mo | HOM |
| **1013288213** | 2 | SSp-n1 | 558 | 0.416546766 | 2.52E-05 | 1 | 6.04E-05 | 2.400690829 | 4 mo | HOM |
| **1008976060** | 2 | SSp-n1 | 558 | 0.416546766 | 1.89E-05 | 1 | 4.54E-05 | 2.400690829 | 8 mo | WILD |
| **1013446243** | 1 | SSp-n1 | 558 | 0.416546766 | 0.000123848 | 6 | 0.000297321 | 14.40414497 | 4 mo | HOM |
| **1013288951** | 2 | SSp-n1 | 558 | 0.416546766 | 0.000234774 | 16 | 0.000563621 | 38.41105326 | 4 mo | HOM |
| **1013978665** | 1 | CEAm | 559 | 0.770614454 | 0 | 0 | 0 | 0 | 2 mo | HOM |
| **1013445609** | 1 | CEAm | 559 | 0.770614454 | 9.4E-05 | 3 | 0.000121935 | 3.892997313 | 4 mo | HOM |
| **1013609015** | 1 | CEAm | 559 | 0.770614454 | 8.4E-05 | 4 | 0.00010905 | 5.190663083 | 4 mo | HOM |
| **1013608106** | 2 | CEAm | 559 | 0.770614454 | 0 | 0 | 0 | 0 | 4 mo | HOM |
| **1011937543** | 1 | CEAm | 559 | 0.770614454 | 0.001548473 | 17 | 0.0020094 | 22.0603181 | 8 mo | HOM |
| **1010419538** | 2 | CEAm | 559 | 0.770614454 | 0 | 0 | 0 | 0 | 4 mo | WILD |
| **1010419187** | 1 | CEAm | 559 | 0.770614454 | 0.001096576 | 37 | 0.001422989 | 48.01363352 | 8 mo | HOM |
| **1013978366** | 1 | CEAm | 559 | 0.770614454 | 0 | 0 | 0 | 0 | 2 mo | HOM |
| **1014416155** | 2 | CEAm | 559 | 0.770614454 | 0 | 0 | 0 | 0 | 2 mo | HOM |
| **1013978065** | 2 | CEAm | 559 | 0.770614454 | 4.81E-06 | 1 | 6.24E-06 | 1.297665771 | 2 mo | HOM |
| **1009934102** | 2 | CEAm | 559 | 0.770614454 | 0 | 0 | 0 | 0 | 4 mo | WILD |
| **1009322937** | 2 | CEAm | 559 | 0.770614454 | 0 | 0 | 0 | 0 | 8 mo | WILD |
| **1011080637** | 2 | CEAm | 559 | 0.770614454 | 0.000825828 | 27 | 0.001071649 | 35.03697581 | 8 mo | HOM |
| **1013609713** | 2 | CEAm | 559 | 0.770614454 | 0 | 0 | 0 | 0 | 2 mo | WILD |
| **1010418842** | 1 | CEAm | 559 | 0.770614454 | 0.001382505 | 38 | 0.00179403 | 49.31129929 | 8 mo | HOM |
| **1013288213** | 2 | CEAm | 559 | 0.770614454 | 2.38E-07 | 0 | 3.08E-07 | 0 | 4 mo | HOM |
| **1008976060** | 2 | CEAm | 559 | 0.770614454 | 0 | 0 | 0 | 0 | 8 mo | WILD |
| **1013446243** | 1 | CEAm | 559 | 0.770614454 | 8.1E-05 | 4 | 0.000105066 | 5.190663083 | 4 mo | HOM |
| **1013288951** | 2 | CEAm | 559 | 0.770614454 | 4.49E-05 | 3 | 5.83E-05 | 3.892997313 | 4 mo | HOM |
| **1013978665** | 1 | MS | 564 | 0.427716453 | 0 | 0 | 0 | 0 | 2 mo | HOM |
| **1013445609** | 1 | MS | 564 | 0.427716453 | 0 | 0 | 0 | 0 | 4 mo | HOM |
| **1013609015** | 1 | MS | 564 | 0.427716453 | 0 | 0 | 0 | 0 | 4 mo | HOM |
| **1013608106** | 2 | MS | 564 | 0.427716453 | 0 | 0 | 0 | 0 | 4 mo | HOM |
| **1011937543** | 1 | MS | 564 | 0.427716453 | 0.000453899 | 13 | 0.001061214 | 30.39396758 | 8 mo | HOM |
| **1010419538** | 2 | MS | 564 | 0.427716453 | 0 | 0 | 0 | 0 | 4 mo | WILD |
| **1010419187** | 1 | MS | 564 | 0.427716453 | 3.99E-05 | 1 | 9.32E-05 | 2.337997506 | 8 mo | HOM |
| **1013978366** | 1 | MS | 564 | 0.427716453 | 0 | 0 | 0 | 0 | 2 mo | HOM |
| **1014416155** | 2 | MS | 564 | 0.427716453 | 0 | 0 | 0 | 0 | 2 mo | HOM |
| **1013978065** | 2 | MS | 564 | 0.427716453 | 0 | 0 | 0 | 0 | 2 mo | HOM |
| **1009934102** | 2 | MS | 564 | 0.427716453 | 0 | 0 | 0 | 0 | 4 mo | WILD |
| **1009322937** | 2 | MS | 564 | 0.427716453 | 2.5E-05 | 1 | 5.84E-05 | 2.337997506 | 8 mo | WILD |
| **1011080637** | 2 | MS | 564 | 0.427716453 | 4.72E-05 | 3 | 0.000110358 | 7.013992519 | 8 mo | HOM |
| **1013609713** | 2 | MS | 564 | 0.427716453 | 0 | 0 | 0 | 0 | 2 mo | WILD |
| **1010418842** | 1 | MS | 564 | 0.427716453 | 0.000137213 | 5 | 0.000320804 | 11.68998753 | 8 mo | HOM |
| **1013288213** | 2 | MS | 564 | 0.427716453 | 0 | 0 | 0 | 0 | 4 mo | HOM |
| **1008976060** | 2 | MS | 564 | 0.427716453 | 0 | 0 | 0 | 0 | 8 mo | WILD |
| **1013446243** | 1 | MS | 564 | 0.427716453 | 0 | 0 | 0 | 0 | 4 mo | HOM |
| **1013288951** | 2 | MS | 564 | 0.427716453 | 0 | 0 | 0 | 0 | 4 mo | HOM |
| **1013978665** | 1 | VISpm5 | 565 | 0.295798202 | 0 | 0 | 0 | 0 | 2 mo | HOM |
| **1013445609** | 1 | VISpm5 | 565 | 0.295798202 | 9.17E-05 | 6 | 0.000309892 | 20.28409895 | 4 mo | HOM |
| **1013609015** | 1 | VISpm5 | 565 | 0.295798202 | 0.000190676 | 9 | 0.000644615 | 30.42614843 | 4 mo | HOM |
| **1013608106** | 2 | VISpm5 | 565 | 0.295798202 | 0 | 0 | 0 | 0 | 4 mo | HOM |
| **1011937543** | 1 | VISpm5 | 565 | 0.295798202 | 0.000492625 | 6 | 0.00166541 | 20.28409895 | 8 mo | HOM |
| **1010419538** | 2 | VISpm5 | 565 | 0.295798202 | 0 | 0 | 0 | 0 | 4 mo | WILD |
| **1010419187** | 1 | VISpm5 | 565 | 0.295798202 | 0.00056369 | 19 | 0.001905658 | 64.23298002 | 8 mo | HOM |
| **1013978366** | 1 | VISpm5 | 565 | 0.295798202 | 0 | 0 | 0 | 0 | 2 mo | HOM |
| **1014416155** | 2 | VISpm5 | 565 | 0.295798202 | 0 | 0 | 0 | 0 | 2 mo | HOM |
| **1013978065** | 2 | VISpm5 | 565 | 0.295798202 | 0 | 0 | 0 | 0 | 2 mo | HOM |
| **1009934102** | 2 | VISpm5 | 565 | 0.295798202 | 0 | 0 | 0 | 0 | 4 mo | WILD |
| **1009322937** | 2 | VISpm5 | 565 | 0.295798202 | 0 | 0 | 0 | 0 | 8 mo | WILD |
| **1011080637** | 2 | VISpm5 | 565 | 0.295798202 | 0.000192258 | 7 | 0.000649963 | 23.66478211 | 8 mo | HOM |

| | | | | | | | | | | |
|---|---|---|---|---|---|---|---|---|---|---|
| **1013609713** | 2 | VISpm5 | 565 | 0.295798202 | 0 | 0 | 0 | 0 | 2 mo | WILD |
| **1010418842** | 1 | VISpm5 | 565 | 0.295798202 | 0.000370065 | 12 | 0.001251072 | 40.56819791 | 8 mo | HOM |
| **1013288213** | 2 | VISpm5 | 565 | 0.295798202 | 1.23E-05 | 1 | 4.16E-05 | 3.380683159 | 4 mo | HOM |
| **1008976060** | 2 | VISpm5 | 565 | 0.295798202 | 0 | 0 | 0 | 0 | 8 mo | WILD |
| **1013446243** | 1 | VISpm5 | 565 | 0.295798202 | 4.12E-05 | 4 | 0.000139347 | 13.52273264 | 4 mo | HOM |
| **1013288951** | 2 | VISpm5 | 565 | 0.295798202 | 4.94E-05 | 3 | 0.000167071 | 10.14204948 | 4 mo | HOM |
| **1013978665** | 1 | TR | 566 | 1.447179798 | 0 | 0 | 0 | 0 | 2 mo | HOM |
| **1013445609** | 1 | TR | 566 | 1.447179798 | 8.86E-05 | 6 | 6.12E-05 | 4.145994856 | 4 mo | HOM |
| **1013609015** | 1 | TR | 566 | 1.447179798 | 0.000198672 | 6 | 0.000137282 | 4.145994856 | 4 mo | HOM |
| **1013608106** | 2 | TR | 566 | 1.447179798 | 0 | 0 | 0 | 0 | 4 mo | HOM |
| **1011937543** | 1 | TR | 566 | 1.447179798 | 0.002729094 | 48 | 0.001885801 | 33.16795885 | 8 mo | HOM |
| **1010419538** | 2 | TR | 566 | 1.447179798 | 2.01E-06 | 0 | 1.39E-06 | 0 | 4 mo | WILD |
| **1010419187** | 1 | TR | 566 | 1.447179798 | 0.003020514 | 53 | 0.002087173 | 36.62295456 | 8 mo | HOM |
| **1013978366** | 1 | TR | 566 | 1.447179798 | 0 | 0 | 0 | 0 | 2 mo | HOM |
| **1014416155** | 2 | TR | 566 | 1.447179798 | 0 | 0 | 0 | 0 | 2 mo | HOM |
| **1013978065** | 2 | TR | 566 | 1.447179798 | 1.05E-05 | 0 | 7.26E-06 | 0 | 2 mo | HOM |
| **1009934102** | 2 | TR | 566 | 1.447179798 | 0 | 0 | 0 | 0 | 4 mo | WILD |
| **1009322937** | 2 | TR | 566 | 1.447179798 | 5.68E-07 | 0 | 3.92E-07 | 0 | 8 mo | WILD |
| **1011080637** | 2 | TR | 566 | 1.447179798 | 0.001129314 | 48 | 0.000780355 | 33.16795885 | 8 mo | HOM |
| **1013609713** | 2 | TR | 566 | 1.447179798 | 5.66E-07 | 0 | 3.91E-07 | 0 | 2 mo | WILD |
| **1010418842** | 1 | TR | 566 | 1.447179798 | 0.001920605 | 51 | 0.001327137 | 35.24095628 | 8 mo | HOM |
| **1013288213** | 2 | TR | 566 | 1.447179798 | 0.00012813 | 6 | 8.85E-05 | 4.145994856 | 4 mo | HOM |
| **1008976060** | 2 | TR | 566 | 1.447179798 | 2.49E-07 | 0 | 1.72E-07 | 0 | 8 mo | WILD |
| **1013446243** | 1 | TR | 566 | 1.447179798 | 0.00019966 | 11 | 0.000137965 | 7.600990569 | 4 mo | HOM |
| **1013288951** | 2 | TR | 566 | 1.447179798 | 0.000206477 | 8 | 0.000142675 | 5.527993141 | 4 mo | HOM |
| **1013978665** | 1 | CH | 567 | 280.5856527 | 0.006424195 | 300 | 2.29E-05 | 1.069192231 | 2 mo | HOM |
| **1013445609** | 1 | CH | 567 | 280.8190016 | 0.016393829 | 1150 | 5.84E-05 | 4.095164478 | 4 mo | HOM |
| **1013609015** | 1 | CH | 567 | 278.5863944 | 0.070728179 | 2079 | 0.000253882 | 7.462676002 | 4 mo | HOM |
| **1013608106** | 2 | CH | 567 | 280.5528038 | 8.41E-06 | 1 | 3E-08 | 0.003564391 | 4 mo | HOM |
| **1011937543** | 1 | CH | 567 | 281.2276776 | 0.366533806 | 5906 | 0.001303335 | 21.00077791 | 8 mo | HOM |
| **1010419538** | 2 | CH | 567 | 281.5976542 | 0.002495091 | 147 | 8.86E-06 | 0.522021394 | 4 mo | WILD |
| **1010419187** | 1 | CH | 567 | 277.7601895 | 0.39261786 | 10769 | 0.001413514 | 38.77085488 | 8 mo | HOM |
| **1013978366** | 1 | CH | 567 | 282.9222644 | 0.009584127 | 351 | 3.39E-05 | 1.240623465 | 2 mo | HOM |
| **1014416155** | 2 | CH | 567 | 281.7472093 | 0.000107661 | 11 | 3.82E-07 | 0.03904209 | 2 mo | HOM |
| **1013978065** | 2 | CH | 567 | 283.3783388 | 0.001789817 | 131 | 6.32E-06 | 0.462279511 | 2 mo | HOM |
| **1009934102** | 2 | CH | 567 | 279.8406815 | 0.004787893 | 252 | 1.71E-05 | 0.900512387 | 4 mo | WILD |
| **1009322937** | 2 | CH | 567 | 277.4171839 | 0.003903907 | 219 | 1.41E-05 | 0.789424782 | 8 mo | WILD |
| **1011080637** | 2 | CH | 567 | 277.0297662 | 0.205627739 | 7859 | 0.000742259 | 28.36879267 | 8 mo | HOM |
| **1013609713** | 2 | CH | 567 | 284.1955585 | 0.001041412 | 75 | 3.66E-06 | 0.263902787 | 2 mo | WILD |
| **1010418842** | 1 | CH | 567 | 279.6201912 | 0.329437928 | 10093 | 0.001178162 | 36.09539052 | 8 mo | HOM |
| **1013288213** | 2 | CH | 567 | 281.2862828 | 0.060207529 | 1135 | 0.000214044 | 4.035035014 | 4 mo | HOM |
| **1008976060** | 2 | CH | 567 | 282.4684291 | 0.01928528 | 816 | 6.83E-05 | 2.888818417 | 8 mo | WILD |
| **1013446243** | 1 | CH | 567 | 280.2221969 | 0.040269488 | 1924 | 0.000143706 | 6.865980001 | 4 mo | HOM |
| **1013288951** | 2 | CH | 567 | 280.0301599 | 0.074738394 | 2297 | 0.000266894 | 8.202687885 | 4 mo | HOM |
| **1013978665** | 1 | MTN | 571 | 1.221904533 | 6.06E-09 | 0 | 4.96E-09 | 0 | 2 mo | HOM |
| **1013445609** | 1 | MTN | 571 | 1.221904533 | 2.25E-07 | 0 | 1.84E-07 | 0 | 4 mo | HOM |
| **1013609015** | 1 | MTN | 571 | 1.221904533 | 0.00036194 | 13 | 0.00029621 | 10.63912904 | 4 mo | HOM |
| **1013608106** | 2 | MTN | 571 | 1.221904533 | 0 | 0 | 0 | 0 | 4 mo | HOM |
| **1011937543** | 1 | MTN | 571 | 1.221904533 | 0.000542203 | 10 | 0.000443736 | 8.183945416 | 8 mo | HOM |
| **1010419538** | 2 | MTN | 571 | 1.221904533 | 0 | 0 | 0 | 0 | 4 mo | WILD |
| **1010419187** | 1 | MTN | 571 | 1.221904533 | 0.000622034 | 20 | 0.000509069 | 16.36789083 | 8 mo | HOM |
| **1013978366** | 1 | MTN | 571 | 1.221904533 | 1.26E-06 | 0 | 1.03E-06 | 0 | 2 mo | HOM |
| **1014416155** | 2 | MTN | 571 | 1.221904533 | 0 | 0 | 0 | 0 | 2 mo | HOM |
| **1013978065** | 2 | MTN | 571 | 1.221904533 | 9.78E-07 | 0 | 8E-07 | 0 | 2 mo | HOM |
| **1009934102** | 2 | MTN | 571 | 1.221904533 | 0 | 0 | 0 | 0 | 4 mo | WILD |
| **1009322937** | 2 | MTN | 571 | 1.221904533 | 1.19E-05 | 1 | 9.71E-06 | 0.818394542 | 8 mo | WILD |
| **1011080637** | 2 | MTN | 571 | 1.221904533 | 0.00024038 | 10 | 0.000196726 | 8.183945416 | 8 mo | HOM |
| **1013609713** | 2 | MTN | 571 | 1.221904533 | 0 | 0 | 0 | 0 | 2 mo | WILD |
| **1010418842** | 1 | MTN | 571 | 1.221904533 | 0.000302512 | 9 | 0.000247574 | 7.365550874 | 8 mo | HOM |

| | | | | | | | | | | |
|---|---|---|---|---|---|---|---|---|---|---|
| **1013288213** | 2 | MTN | 571 | 1.221904533 | 9.91E-08 | 0 | 8.11E-08 | 0 | 4 mo | HOM |
| **1008976060** | 2 | MTN | 571 | 1.221904533 | 3.76E-05 | 3 | 3.07E-05 | 2.455183625 | 8 mo | WILD |
| **1013446243** | 1 | MTN | 571 | 1.221904533 | 3.21E-05 | 2 | 2.63E-05 | 1.636789083 | 4 mo | HOM |
| **1013288951** | 2 | MTN | 571 | 1.221904533 | 0 | 0 | 0 | 0 | 4 mo | HOM |
| **1013978665** | 1 | VISl4 | 573 | 0.184496819 | 0 | 0 | 0 | 0 | 2 mo | HOM |
| **1013445609** | 1 | VISl4 | 573 | 0.184496819 | 1.28E-05 | 2 | 6.93E-05 | 10.84029531 | 4 mo | HOM |
| **1013609015** | 1 | VISl4 | 573 | 0.184496819 | 4.85E-05 | 3 | 0.000262926 | 16.26044297 | 4 mo | HOM |
| **1013608106** | 2 | VISl4 | 573 | 0.184496819 | 0 | 0 | 0 | 0 | 4 mo | HOM |
| **1011937543** | 1 | VISl4 | 573 | 0.184496819 | 0.000372606 | 9 | 0.002019577 | 48.78132891 | 8 mo | HOM |
| **1010419538** | 2 | VISl4 | 573 | 0.184496819 | 0 | 0 | 0 | 0 | 4 mo | WILD |
| **1010419187** | 1 | VISl4 | 573 | 0.184496819 | 0.000402371 | 15 | 0.00218091 | 81.30221484 | 8 mo | HOM |
| **1013978366** | 1 | VISl4 | 573 | 0.184496819 | 0 | 0 | 0 | 0 | 2 mo | HOM |
| **1014416155** | 2 | VISl4 | 573 | 0.184496819 | 0 | 0 | 0 | 0 | 2 mo | HOM |
| **1013978065** | 2 | VISl4 | 573 | 0.184496819 | 0 | 0 | 0 | 0 | 2 mo | HOM |
| **1009934102** | 2 | VISl4 | 573 | 0.184496819 | 0 | 0 | 0 | 0 | 4 mo | WILD |
| **1009322937** | 2 | VISl4 | 573 | 0.184496819 | 0 | 0 | 0 | 0 | 8 mo | WILD |
| **1011080637** | 2 | VISl4 | 573 | 0.184496819 | 0.00015421 | 5 | 0.00083584 | 27.10073828 | 8 mo | HOM |
| **1013609713** | 2 | VISl4 | 573 | 0.184496819 | 0 | 0 | 0 | 0 | 2 mo | WILD |
| **1010418842** | 1 | VISl4 | 573 | 0.184496819 | 0.000220867 | 11 | 0.001197131 | 59.62162422 | 8 mo | HOM |
| **1013288213** | 2 | VISl4 | 573 | 0.184496819 | 6.32E-05 | 1 | 0.000342471 | 5.420147656 | 4 mo | HOM |
| **1008976060** | 2 | VISl4 | 573 | 0.184496819 | 0 | 0 | 0 | 0 | 8 mo | WILD |
| **1013446243** | 1 | VISl4 | 573 | 0.184496819 | 5.56E-05 | 3 | 0.000301176 | 16.26044297 | 4 mo | HOM |
| **1013288951** | 2 | VISl4 | 573 | 0.184496819 | 8.62E-05 | 4 | 0.000467039 | 21.68059062 | 4 mo | HOM |
| **1013978665** | 1 | TRN | 574 | 0.710509094 | 0 | 0 | 0 | 0 | 2 mo | HOM |
| **1013445609** | 1 | TRN | 574 | 0.710509094 | 0 | 0 | 0 | 0 | 4 mo | HOM |
| **1013609015** | 1 | TRN | 574 | 0.710509094 | 0 | 0 | 0 | 0 | 4 mo | HOM |
| **1013608106** | 2 | TRN | 574 | 0.710509094 | 0 | 0 | 0 | 0 | 4 mo | HOM |
| **1011937543** | 1 | TRN | 574 | 0.710509094 | 6.91E-05 | 1 | 9.73E-05 | 1.40744152 | 8 mo | HOM |
| **1010419538** | 2 | TRN | 574 | 0.710509094 | 0 | 0 | 0 | 0 | 4 mo | WILD |
| **1010419187** | 1 | TRN | 574 | 0.710509094 | 0 | 0 | 0 | 0 | 8 mo | HOM |
| **1013978366** | 1 | TRN | 574 | 0.710509094 | 0 | 0 | 0 | 0 | 2 mo | HOM |
| **1014416155** | 2 | TRN | 574 | 0.710509094 | 0 | 0 | 0 | 0 | 2 mo | HOM |
| **1013978065** | 2 | TRN | 574 | 0.710509094 | 0 | 0 | 0 | 0 | 2 mo | HOM |
| **1009934102** | 2 | TRN | 574 | 0.710509094 | 0 | 0 | 0 | 0 | 4 mo | WILD |
| **1009322937** | 2 | TRN | 574 | 0.710509094 | 0 | 0 | 0 | 0 | 8 mo | WILD |
| **1011080637** | 2 | TRN | 574 | 0.710509094 | 1.11E-05 | 1 | 1.57E-05 | 1.40744152 | 8 mo | HOM |
| **1013609713** | 2 | TRN | 574 | 0.710509094 | 0 | 0 | 0 | 0 | 2 mo | WILD |
| **1010418842** | 1 | TRN | 574 | 0.710509094 | 0.000191001 | 3 | 0.000268823 | 4.222324561 | 8 mo | HOM |
| **1013288213** | 2 | TRN | 574 | 0.710509094 | 0 | 0 | 0 | 0 | 4 mo | HOM |
| **1008976060** | 2 | TRN | 574 | 0.710509094 | 5.63E-06 | 1 | 7.93E-06 | 1.40744152 | 8 mo | WILD |
| **1013446243** | 1 | TRN | 574 | 0.710509094 | 0 | 0 | 0 | 0 | 4 mo | HOM |
| **1013288951** | 2 | TRN | 574 | 0.710509094 | 0 | 0 | 0 | 0 | 4 mo | HOM |
| **1013978665** | 1 | CL | 575 | 0.371845281 | 0 | 0 | 0 | 0 | 2 mo | HOM |
| **1013445609** | 1 | CL | 575 | 0.371845281 | 0 | 0 | 0 | 0 | 4 mo | HOM |
| **1013609015** | 1 | CL | 575 | 0.371845281 | 0 | 0 | 0 | 0 | 4 mo | HOM |
| **1013608106** | 2 | CL | 575 | 0.371845281 | 0 | 0 | 0 | 0 | 4 mo | HOM |
| **1011937543** | 1 | CL | 575 | 0.371845281 | 0.000354809 | 6 | 0.000954185 | 16.1357433 | 8 mo | HOM |
| **1010419538** | 2 | CL | 575 | 0.371845281 | 0 | 0 | 0 | 0 | 4 mo | WILD |
| **1010419187** | 1 | CL | 575 | 0.371845281 | 0.000103883 | 3 | 0.000279371 | 8.067871652 | 8 mo | HOM |
| **1013978366** | 1 | CL | 575 | 0.371845281 | 0 | 0 | 0 | 0 | 2 mo | HOM |
| **1014416155** | 2 | CL | 575 | 0.371845281 | 0 | 0 | 0 | 0 | 2 mo | HOM |
| **1013978065** | 2 | CL | 575 | 0.371845281 | 0 | 0 | 0 | 0 | 2 mo | HOM |
| **1009934102** | 2 | CL | 575 | 0.371845281 | 0 | 0 | 0 | 0 | 4 mo | WILD |
| **1009322937** | 2 | CL | 575 | 0.371845281 | 0 | 0 | 0 | 0 | 8 mo | WILD |
| **1011080637** | 2 | CL | 575 | 0.371845281 | 6.49E-05 | 3 | 0.000174489 | 8.067871652 | 8 mo | HOM |
| **1013609713** | 2 | CL | 575 | 0.371845281 | 0 | 0 | 0 | 0 | 2 mo | WILD |
| **1010418842** | 1 | CL | 575 | 0.371845281 | 0.000120923 | 5 | 0.000325198 | 13.44645275 | 8 mo | HOM |
| **1013288213** | 2 | CL | 575 | 0.371845281 | 0 | 0 | 0 | 0 | 4 mo | HOM |
| **1008976060** | 2 | CL | 575 | 0.371845281 | 3.56E-06 | 1 | 9.58E-06 | 2.689290551 | 8 mo | WILD |

| | | | | | | | | | | |
|---|---|---|---|---|---|---|---|---|---|---|
| 1013446243 | 1 | CL | 575 | 0.371845281 | 1.61E-05 | 1 | 4.33E-05 | 2.689290551 | 4 mo | HOM |
| 1013288951 | 2 | CL | 575 | 0.371845281 | 0 | 0 | 0 | 0 | 4 mo | HOM |
| 1013978665 | 1 | ACVII | 576 | 0.005798106 | 0 | 0 | 0 | 0 | 2 mo | HOM |
| 1013445609 | 1 | ACVII | 576 | 0.005798106 | 0 | 0 | 0 | 0 | 4 mo | HOM |
| 1013609015 | 1 | ACVII | 576 | 0.005798106 | 0 | 0 | 0 | 0 | 4 mo | HOM |
| 1013608106 | 2 | ACVII | 576 | 0.005798106 | 0 | 0 | 0 | 0 | 4 mo | HOM |
| 1011937543 | 1 | ACVII | 576 | 0.005798106 | 0 | 0 | 0 | 0 | 8 mo | HOM |
| 1010419538 | 2 | ACVII | 576 | 0.005798106 | 0 | 0 | 0 | 0 | 4 mo | WILD |
| 1010419187 | 1 | ACVII | 576 | 0.005798106 | 0 | 0 | 0 | 0 | 8 mo | HOM |
| 1013978366 | 1 | ACVII | 576 | 0.005798106 | 0 | 0 | 0 | 0 | 2 mo | HOM |
| 1014416155 | 2 | ACVII | 576 | 0.005798106 | 0 | 0 | 0 | 0 | 2 mo | HOM |
| 1013978065 | 2 | ACVII | 576 | 0.005798106 | 0 | 0 | 0 | 0 | 2 mo | HOM |
| 1009934102 | 2 | ACVII | 576 | 0.005798106 | 0 | 0 | 0 | 0 | 4 mo | WILD |
| 1009322937 | 2 | ACVII | 576 | 0.005798106 | 0 | 0 | 0 | 0 | 8 mo | WILD |
| 1011080637 | 2 | ACVII | 576 | 0.005798106 | 0 | 0 | 0 | 0 | 8 mo | HOM |
| 1013609713 | 2 | ACVII | 576 | 0.005798106 | 0 | 0 | 0 | 0 | 2 mo | WILD |
| 1010418842 | 1 | ACVII | 576 | 0.005798106 | 0 | 0 | 0 | 0 | 8 mo | HOM |
| 1013288213 | 2 | ACVII | 576 | 0.005798106 | 0 | 0 | 0 | 0 | 4 mo | HOM |
| 1008976060 | 2 | ACVII | 576 | 0.005798106 | 0 | 0 | 0 | 0 | 8 mo | WILD |
| 1013446243 | 1 | ACVII | 576 | 0.005798106 | 0 | 0 | 0 | 0 | 4 mo | HOM |
| 1013288951 | 2 | ACVII | 576 | 0.005798106 | 0 | 0 | 0 | 0 | 4 mo | HOM |
| 1013978665 | 1 | SSp-ul4 | 577 | 0.538941592 | 0 | 0 | 0 | 0 | 2 mo | HOM |
| 1013445609 | 1 | SSp-ul4 | 577 | 0.538941592 | 3.9E-05 | 2 | 7.23E-05 | 3.710977274 | 4 mo | HOM |
| 1013609015 | 1 | SSp-ul4 | 577 | 0.538941592 | 0.000160124 | 8 | 0.000297108 | 14.8439091 | 4 mo | HOM |
| 1013608106 | 2 | SSp-ul4 | 577 | 0.538941592 | 0 | 0 | 0 | 0 | 4 mo | HOM |
| 1011937543 | 1 | SSp-ul4 | 577 | 0.538941592 | 0.000856294 | 16 | 0.001588844 | 29.68781819 | 8 mo | HOM |
| 1010419538 | 2 | SSp-ul4 | 577 | 0.538941592 | 0 | 0 | 0 | 0 | 4 mo | WILD |
| 1010419187 | 1 | SSp-ul4 | 577 | 0.538941592 | 0.00086222 | 23 | 0.00159984 | 42.67623865 | 8 mo | HOM |
| 1013978366 | 1 | SSp-ul4 | 577 | 0.538941592 | 0 | 0 | 0 | 0 | 2 mo | HOM |
| 1014416155 | 2 | SSp-ul4 | 577 | 0.538941592 | 0 | 0 | 0 | 0 | 2 mo | HOM |
| 1013978065 | 2 | SSp-ul4 | 577 | 0.538941592 | 0 | 0 | 0 | 0 | 2 mo | HOM |
| 1009934102 | 2 | SSp-ul4 | 577 | 0.538941592 | 0 | 0 | 0 | 0 | 4 mo | WILD |
| 1009322937 | 2 | SSp-ul4 | 577 | 0.538941592 | 0 | 0 | 0 | 0 | 8 mo | WILD |
| 1011080637 | 2 | SSp-ul4 | 577 | 0.538941592 | 0.000631621 | 18 | 0.001171965 | 33.39879547 | 8 mo | HOM |
| 1013609713 | 2 | SSp-ul4 | 577 | 0.538941592 | 0 | 0 | 0 | 0 | 2 mo | WILD |
| 1010418842 | 1 | SSp-ul4 | 577 | 0.538941592 | 0.000614835 | 21 | 0.00114082 | 38.96526138 | 8 mo | HOM |
| 1013288213 | 2 | SSp-ul4 | 577 | 0.538941592 | 2.77E-05 | 4 | 5.14E-05 | 7.421954548 | 4 mo | HOM |
| 1008976060 | 2 | SSp-ul4 | 577 | 0.538941592 | 0 | 0 | 0 | 0 | 8 mo | WILD |
| 1013446243 | 1 | SSp-ul4 | 577 | 0.538941592 | 8.26E-05 | 7 | 0.000153338 | 12.98842046 | 4 mo | HOM |
| 1013288951 | 2 | SSp-ul4 | 577 | 0.538941592 | 0.000129797 | 4 | 0.000240837 | 7.421954548 | 4 mo | HOM |
| 1013978665 | 1 | ec | 579 | 0.895764139 | 0 | 0 | 0 | 0 | 2 mo | HOM |
| 1013445609 | 1 | ec | 579 | 0.895764139 | 1.77E-05 | 2 | 1.97E-05 | 2.232730596 | 4 mo | HOM |
| 1013609015 | 1 | ec | 579 | 0.895764139 | 1.8E-05 | 1 | 2.01E-05 | 1.116365298 | 4 mo | HOM |
| 1013608106 | 2 | ec | 579 | 0.895764139 | 0 | 0 | 0 | 0 | 4 mo | HOM |
| 1011937543 | 1 | ec | 579 | 0.895764139 | 0.001237376 | 23 | 0.001381364 | 25.67640185 | 8 mo | HOM |
| 1010419538 | 2 | ec | 579 | 0.895764139 | 0 | 0 | 0 | 0 | 4 mo | WILD |
| 1010419187 | 1 | ec | 579 | 0.895764139 | 0.000296073 | 17 | 0.000330525 | 18.97821006 | 8 mo | HOM |
| 1013978366 | 1 | ec | 579 | 0.895764139 | 0 | 0 | 0 | 0 | 2 mo | HOM |
| 1014416155 | 2 | ec | 579 | 0.895764139 | 0 | 0 | 0 | 0 | 2 mo | HOM |
| 1013978065 | 2 | ec | 579 | 0.895764139 | 0 | 0 | 0 | 0 | 2 mo | HOM |
| 1009934102 | 2 | ec | 579 | 0.895764139 | 0 | 0 | 0 | 0 | 4 mo | WILD |
| 1009322937 | 2 | ec | 579 | 0.895764139 | 0 | 0 | 0 | 0 | 8 mo | WILD |
| 1011080637 | 2 | ec | 579 | 0.895764139 | 0.000160913 | 8 | 0.000179638 | 8.930922382 | 8 mo | HOM |
| 1013609713 | 2 | ec | 579 | 0.895764139 | 0 | 0 | 0 | 0 | 2 mo | WILD |
| 1010418842 | 1 | ec | 579 | 0.895764139 | 0.000536085 | 23 | 0.000598466 | 25.67640185 | 8 mo | HOM |
| 1013288213 | 2 | ec | 579 | 0.895764139 | 4E-07 | 0 | 4.47E-07 | 0 | 4 mo | HOM |
| 1008976060 | 2 | ec | 579 | 0.895764139 | 6.2E-06 | 2 | 6.93E-06 | 2.232730596 | 8 mo | WILD |
| 1013446243 | 1 | ec | 579 | 0.895764139 | 4.65E-05 | 4 | 5.19E-05 | 4.465461191 | 4 mo | HOM |
| 1013288951 | 2 | ec | 579 | 0.895764139 | 4.16E-05 | 0 | 4.64E-05 | 0 | 4 mo | HOM |

| | | | | | | | | | | |
|---|---|---|---|---|---|---|---|---|---|---|
| **1013978665** | 1 | NB | 580 | 0.090583566 | 0 | 0 | 0 | 0 | 2 mo | HOM |
| **1013445609** | 1 | NB | 580 | 0.090583566 | 0 | 0 | 0 | 0 | 4 mo | HOM |
| **1013609015** | 1 | NB | 580 | 0.090583566 | 0 | 0 | 0 | 0 | 4 mo | HOM |
| **1013608106** | 2 | NB | 580 | 0.090583566 | 0 | 0 | 0 | 0 | 4 mo | HOM |
| **1011937543** | 1 | NB | 580 | 0.090583566 | 0.000186589 | 2 | 0.002059856 | 22.07906017 | 8 mo | HOM |
| **1010419538** | 2 | NB | 580 | 0.090583566 | 0 | 0 | 0 | 0 | 4 mo | WILD |
| **1010419187** | 1 | NB | 580 | 0.090583566 | 0 | 0 | 0 | 0 | 8 mo | HOM |
| **1013978366** | 1 | NB | 580 | 0.090583566 | 0 | 0 | 0 | 0 | 2 mo | HOM |
| **1014416155** | 2 | NB | 580 | 0.090583566 | 0 | 0 | 0 | 0 | 2 mo | HOM |
| **1013978065** | 2 | NB | 580 | 0.090583566 | 0 | 0 | 0 | 0 | 2 mo | HOM |
| **1009934102** | 2 | NB | 580 | 0.090583566 | 0 | 0 | 0 | 0 | 4 mo | WILD |
| **1009322937** | 2 | NB | 580 | 0.090583566 | 0 | 0 | 0 | 0 | 8 mo | WILD |
| **1011080637** | 2 | NB | 580 | 0.090583566 | 0 | 0 | 0 | 0 | 8 mo | HOM |
| **1013609713** | 2 | NB | 580 | 0.090583566 | 0 | 0 | 0 | 0 | 2 mo | WILD |
| **1010418842** | 1 | NB | 580 | 0.090583566 | 0 | 0 | 0 | 0 | 8 mo | HOM |
| **1013288213** | 2 | NB | 580 | 0.090583566 | 0 | 0 | 0 | 0 | 4 mo | HOM |
| **1008976060** | 2 | NB | 580 | 0.090583566 | 0 | 0 | 0 | 0 | 8 mo | WILD |
| **1013446243** | 1 | NB | 580 | 0.090583566 | 0 | 0 | 0 | 0 | 4 mo | HOM |
| **1013288951** | 2 | NB | 580 | 0.090583566 | 0 | 0 | 0 | 0 | 4 mo | HOM |
| **1013978665** | 1 | TRS | 581 | 0.366605574 | 3.78E-05 | 0 | 0.000103076 | 0 | 2 mo | HOM |
| **1013445609** | 1 | TRS | 581 | 0.366605574 | 2.46E-05 | 1 | 6.71E-05 | 2.727727214 | 4 mo | HOM |
| **1013609015** | 1 | TRS | 581 | 0.366605574 | 0.000832678 | 7 | 0.002271318 | 19.0940905 | 4 mo | HOM |
| **1013608106** | 2 | TRS | 581 | 0.366605574 | 0 | 0 | 0 | 0 | 4 mo | HOM |
| **1011937543** | 1 | TRS | 581 | 0.366605574 | 0.000123908 | 7 | 0.000337988 | 19.0940905 | 8 mo | HOM |
| **1010419538** | 2 | TRS | 581 | 0.366605574 | 5.13E-06 | 0 | 1.4E-05 | 0 | 4 mo | WILD |
| **1010419187** | 1 | TRS | 581 | 0.366605574 | 0.000600044 | 14 | 0.001636756 | 38.188181 | 8 mo | HOM |
| **1013978366** | 1 | TRS | 581 | 0.366605574 | 3.75E-06 | 1 | 1.02E-05 | 2.727727214 | 2 mo | HOM |
| **1014416155** | 2 | TRS | 581 | 0.366605574 | 0 | 0 | 0 | 0 | 2 mo | HOM |
| **1013978065** | 2 | TRS | 581 | 0.366605574 | 0 | 0 | 0 | 0 | 2 mo | HOM |
| **1009934102** | 2 | TRS | 581 | 0.366605574 | 1.88E-05 | 2 | 5.13E-05 | 5.455454428 | 4 mo | WILD |
| **1009322937** | 2 | TRS | 581 | 0.366605574 | 0.000129964 | 7 | 0.000354506 | 19.0940905 | 8 mo | WILD |
| **1011080637** | 2 | TRS | 581 | 0.366605574 | 0.000846522 | 12 | 0.002309082 | 32.73272657 | 8 mo | HOM |
| **1013609713** | 2 | TRS | 581 | 0.366605574 | 1.86E-06 | 0 | 5.07E-06 | 0 | 2 mo | WILD |
| **1010418842** | 1 | TRS | 581 | 0.366605574 | 0.001363112 | 14 | 0.003718199 | 38.188181 | 8 mo | HOM |
| **1013288213** | 2 | TRS | 581 | 0.366605574 | 2.11E-05 | 0 | 5.75E-05 | 0 | 4 mo | HOM |
| **1008976060** | 2 | TRS | 581 | 0.366605574 | 0.00054828 | 3 | 0.001495558 | 8.183181643 | 8 mo | WILD |
| **1013446243** | 1 | TRS | 581 | 0.366605574 | 0.000497113 | 9 | 0.00135599 | 24.54954493 | 4 mo | HOM |
| **1013288951** | 2 | TRS | 581 | 0.366605574 | 0.000669224 | 7 | 0.00182546 | 19.0940905 | 4 mo | HOM |
| **1013978665** | 1 | ORBm2/3 | 582 | 0.326548348 | 0 | 0 | 0 | 0 | 2 mo | HOM |
| **1013445609** | 1 | ORBm2/3 | 582 | 0.326548348 | 5.99E-06 | 1 | 1.84E-05 | 3.062333667 | 4 mo | HOM |
| **1013609015** | 1 | ORBm2/3 | 582 | 0.326548348 | 4.25E-05 | 3 | 0.000130208 | 9.187001001 | 4 mo | HOM |
| **1013608106** | 2 | ORBm2/3 | 582 | 0.326548348 | 0 | 0 | 0 | 0 | 4 mo | HOM |
| **1011937543** | 1 | ORBm2/3 | 582 | 0.326548348 | 0.000523895 | 8 | 0.001604341 | 24.49866934 | 8 mo | HOM |
| **1010419538** | 2 | ORBm2/3 | 582 | 0.326548348 | 0 | 0 | 0 | 0 | 4 mo | WILD |
| **1010419187** | 1 | ORBm2/3 | 582 | 0.326548348 | 0.000323866 | 11 | 0.000991787 | 33.68567034 | 8 mo | HOM |
| **1013978366** | 1 | ORBm2/3 | 582 | 0.326548348 | 0 | 0 | 0 | 0 | 2 mo | HOM |
| **1014416155** | 2 | ORBm2/3 | 582 | 0.326548348 | 0 | 0 | 0 | 0 | 2 mo | HOM |
| **1013978065** | 2 | ORBm2/3 | 582 | 0.326548348 | 0 | 0 | 0 | 0 | 2 mo | HOM |
| **1009934102** | 2 | ORBm2/3 | 582 | 0.326548348 | 4.25E-08 | 0 | 1.3E-07 | 0 | 4 mo | WILD |
| **1009322937** | 2 | ORBm2/3 | 582 | 0.326548348 | 0 | 0 | 0 | 0 | 8 mo | WILD |
| **1011080637** | 2 | ORBm2/3 | 582 | 0.326548348 | 0.000150061 | 8 | 0.000459536 | 24.49866934 | 8 mo | HOM |
| **1013609713** | 2 | ORBm2/3 | 582 | 0.326548348 | 0 | 0 | 0 | 0 | 2 mo | WILD |
| **1010418842** | 1 | ORBm2/3 | 582 | 0.326548348 | 0.000359386 | 12 | 0.001100559 | 36.748004 | 8 mo | HOM |
| **1013288213** | 2 | ORBm2/3 | 582 | 0.326548348 | 0.000148248 | 5 | 0.000453984 | 15.31166834 | 4 mo | HOM |
| **1008976060** | 2 | ORBm2/3 | 582 | 0.326548348 | 0 | 0 | 0 | 0 | 8 mo | WILD |
| **1013446243** | 1 | ORBm2/3 | 582 | 0.326548348 | 2.06E-05 | 1 | 6.31E-05 | 3.062333667 | 4 mo | HOM |
| **1013288951** | 2 | ORBm2/3 | 582 | 0.326548348 | 4.39E-05 | 2 | 0.000134457 | 6.124667334 | 4 mo | HOM |
| **1013978665** | 1 | CLA | 583 | 0.567384054 | 0 | 0 | 0 | 0 | 2 mo | HOM |
| **1013445609** | 1 | CLA | 583 | 0.567384054 | 8.3E-06 | 1 | 1.46E-05 | 1.76247463 | 4 mo | HOM |

| | | | | | | | | | | |
|---|---|---|---|---|---|---|---|---|---|---|
| **1013609015** | 1 | CLA | 583 | 0.567384054 | 3.34E-05 | 2 | 5.89E-05 | 3.52494926 | 4 mo | HOM |
| **1013608106** | 2 | CLA | 583 | 0.567384054 | 0 | 0 | 0 | 0 | 4 mo | HOM |
| **1011937543** | 1 | CLA | 583 | 0.567384054 | 0.000601328 | 16 | 0.001059826 | 28.19959408 | 8 mo | HOM |
| **1010419538** | 2 | CLA | 583 | 0.567384054 | 0 | 0 | 0 | 0 | 4 mo | WILD |
| **1010419187** | 1 | CLA | 583 | 0.567384054 | 0.000660035 | 31 | 0.001163294 | 54.63671353 | 8 mo | HOM |
| **1013978366** | 1 | CLA | 583 | 0.567384054 | 0 | 0 | 0 | 0 | 2 mo | HOM |
| **1014416155** | 2 | CLA | 583 | 0.567384054 | 0 | 0 | 0 | 0 | 2 mo | HOM |
| **1013978065** | 2 | CLA | 583 | 0.567384054 | 0 | 0 | 0 | 0 | 2 mo | HOM |
| **1009934102** | 2 | CLA | 583 | 0.567384054 | 0 | 0 | 0 | 0 | 4 mo | WILD |
| **1009322937** | 2 | CLA | 583 | 0.567384054 | 0 | 0 | 0 | 0 | 8 mo | WILD |
| **1011080637** | 2 | CLA | 583 | 0.567384054 | 0.000577258 | 31 | 0.001017403 | 54.63671353 | 8 mo | HOM |
| **1013609713** | 2 | CLA | 583 | 0.567384054 | 0 | 0 | 0 | 0 | 2 mo | WILD |
| **1010418842** | 1 | CLA | 583 | 0.567384054 | 0.000848335 | 28 | 0.001495169 | 49.34928964 | 8 mo | HOM |
| **1013288213** | 2 | CLA | 583 | 0.567384054 | 7.71E-06 | 1 | 1.36E-05 | 1.76247463 | 4 mo | HOM |
| **1008976060** | 2 | CLA | 583 | 0.567384054 | 2.87E-05 | 4 | 5.05E-05 | 7.04989852 | 8 mo | WILD |
| **1013446243** | 1 | CLA | 583 | 0.567384054 | 6.34E-05 | 4 | 0.000111712 | 7.04989852 | 4 mo | HOM |
| **1013288951** | 2 | CLA | 583 | 0.567384054 | 0 | 0 | 0 | 0 | 4 mo | HOM |
| **1013978665** | 1 | ND | 587 | 0.095163939 | 0 | 0 | 0 | 0 | 2 mo | HOM |
| **1013445609** | 1 | ND | 587 | 0.095163939 | 0 | 0 | 0 | 0 | 4 mo | HOM |
| **1013609015** | 1 | ND | 587 | 0.095163939 | 0 | 0 | 0 | 0 | 4 mo | HOM |
| **1013608106** | 2 | ND | 587 | 0.095163939 | 0 | 0 | 0 | 0 | 4 mo | HOM |
| **1011937543** | 1 | ND | 587 | 0.095163939 | 4.11E-05 | 1 | 0.000431998 | 10.50818212 | 8 mo | HOM |
| **1010419538** | 2 | ND | 587 | 0.095163939 | 0 | 0 | 0 | 0 | 4 mo | WILD |
| **1010419187** | 1 | ND | 587 | 0.095163939 | 0 | 0 | 0 | 0 | 8 mo | HOM |
| **1013978366** | 1 | ND | 587 | 0.095163939 | 0 | 0 | 0 | 0 | 2 mo | HOM |
| **1014416155** | 2 | ND | 587 | 0.095163939 | 0 | 0 | 0 | 0 | 2 mo | HOM |
| **1013978065** | 2 | ND | 587 | 0.095163939 | 0 | 0 | 0 | 0 | 2 mo | HOM |
| **1009934102** | 2 | ND | 587 | 0.095163939 | 0 | 0 | 0 | 0 | 4 mo | WILD |
| **1009322937** | 2 | ND | 587 | 0.095163939 | 0 | 0 | 0 | 0 | 8 mo | WILD |
| **1011080637** | 2 | ND | 587 | 0.095163939 | 0 | 0 | 0 | 0 | 8 mo | HOM |
| **1013609713** | 2 | ND | 587 | 0.095163939 | 0 | 0 | 0 | 0 | 2 mo | WILD |
| **1010418842** | 1 | ND | 587 | 0.095163939 | 0 | 0 | 0 | 0 | 8 mo | HOM |
| **1013288213** | 2 | ND | 587 | 0.095163939 | 0 | 0 | 0 | 0 | 4 mo | HOM |
| **1008976060** | 2 | ND | 587 | 0.095163939 | 6.24E-08 | 0 | 6.56E-07 | 0 | 8 mo | WILD |
| **1013446243** | 1 | ND | 587 | 0.095163939 | 0 | 0 | 0 | 0 | 4 mo | HOM |
| **1013288951** | 2 | ND | 587 | 0.095163939 | 0 | 0 | 0 | 0 | 4 mo | HOM |
| **1013978665** | 1 | ACAv1 | 588 | 0.492694804 | 0 | 0 | 0 | 0 | 2 mo | HOM |
| **1013445609** | 1 | ACAv1 | 588 | 0.492694804 | 4.23E-05 | 5 | 8.58E-05 | 10.14827021 | 4 mo | HOM |
| **1013609015** | 1 | ACAv1 | 588 | 0.492694804 | 8.93E-05 | 3 | 0.000181234 | 6.088962123 | 4 mo | HOM |
| **1013608106** | 2 | ACAv1 | 588 | 0.492694804 | 0 | 0 | 0 | 0 | 4 mo | HOM |
| **1011937543** | 1 | ACAv1 | 588 | 0.492694804 | 0.000353839 | 5 | 0.000718172 | 10.14827021 | 8 mo | HOM |
| **1010419538** | 2 | ACAv1 | 588 | 0.492694804 | 0 | 0 | 0 | 0 | 4 mo | WILD |
| **1010419187** | 1 | ACAv1 | 588 | 0.492694804 | 0.000798029 | 15 | 0.001619724 | 30.44481062 | 8 mo | HOM |
| **1013978366** | 1 | ACAv1 | 588 | 0.492694804 | 0.000353081 | 4 | 0.000716632 | 8.118616165 | 2 mo | HOM |
| **1014416155** | 2 | ACAv1 | 588 | 0.492694804 | 0 | 0 | 0 | 0 | 2 mo | HOM |
| **1013978065** | 2 | ACAv1 | 588 | 0.492694804 | 0 | 0 | 0 | 0 | 2 mo | HOM |
| **1009934102** | 2 | ACAv1 | 588 | 0.492694804 | 9.37E-05 | 3 | 0.000190215 | 6.088962123 | 4 mo | WILD |
| **1009322937** | 2 | ACAv1 | 588 | 0.492694804 | 1.62E-05 | 1 | 3.29E-05 | 2.029654041 | 8 mo | WILD |
| **1011080637** | 2 | ACAv1 | 588 | 0.492694804 | 0.000936661 | 20 | 0.001901097 | 40.59308082 | 8 mo | HOM |
| **1013609713** | 2 | ACAv1 | 588 | 0.492694804 | 1.44E-06 | 0 | 2.92E-06 | 0 | 2 mo | WILD |
| **1010418842** | 1 | ACAv1 | 588 | 0.492694804 | 0.001120373 | 16 | 0.00227397 | 32.47446466 | 8 mo | HOM |
| **1013288213** | 2 | ACAv1 | 588 | 0.492694804 | 0.000319805 | 5 | 0.000649094 | 10.14827021 | 4 mo | HOM |
| **1008976060** | 2 | ACAv1 | 588 | 0.492694804 | 0 | 0 | 0 | 0 | 8 mo | WILD |
| **1013446243** | 1 | ACAv1 | 588 | 0.492694804 | 6.15E-06 | 0 | 1.25E-05 | 0 | 4 mo | HOM |
| **1013288951** | 2 | ACAv1 | 588 | 0.492694804 | 0.00019029 | 8 | 0.000386222 | 16.23723233 | 4 mo | HOM |
| **1013978665** | 1 | TT | 589 | 1.484552138 | 0 | 0 | 0 | 0 | 2 mo | HOM |
| **1013445609** | 1 | TT | 589 | 1.484552138 | 5.37E-06 | 1 | 3.62E-06 | 0.673603826 | 4 mo | HOM |
| **1013609015** | 1 | TT | 589 | 1.484552138 | 0.000136627 | 7 | 9.2E-05 | 4.715226783 | 4 mo | HOM |
| **1013608106** | 2 | TT | 589 | 1.484552138 | 0 | 0 | 0 | 0 | 4 mo | HOM |

| | | | | | | | | | | |
|---|---|---|---|---|---|---|---|---|---|---|
| **1011937543** | 1 | TT | 589 | 1.484552138 | 0.00046302 | 11 | 0.000311892 | 7.409642088 | 8 mo | HOM |
| **1010419538** | 2 | TT | 589 | 1.484552138 | 0 | 0 | 0 | 0 | 4 mo | WILD |
| **1010419187** | 1 | TT | 589 | 1.484552138 | 0.000890596 | 24 | 0.000599909 | 16.16649183 | 8 mo | HOM |
| **1013978366** | 1 | TT | 589 | 1.484552138 | 0 | 0 | 0 | 0 | 2 mo | HOM |
| **1014416155** | 2 | TT | 589 | 1.484552138 | 7.99E-06 | 2 | 5.38E-06 | 1.347207652 | 2 mo | HOM |
| **1013978065** | 2 | TT | 589 | 1.484552138 | 1.17E-06 | 0 | 7.85E-07 | 0 | 2 mo | HOM |
| **1009934102** | 2 | TT | 589 | 1.484552138 | 6.23E-06 | 1 | 4.2E-06 | 0.673603826 | 4 mo | WILD |
| **1009322937** | 2 | TT | 589 | 1.484552138 | 0.000620045 | 10 | 0.000417665 | 6.736038261 | 8 mo | WILD |
| **1011080637** | 2 | TT | 589 | 1.484552138 | 0.000586731 | 25 | 0.000395224 | 16.84009565 | 8 mo | HOM |
| **1013609713** | 2 | TT | 589 | 1.484552138 | 0 | 0 | 0 | 0 | 2 mo | WILD |
| **1010418842** | 1 | TT | 589 | 1.484552138 | 0.003466181 | 28 | 0.002334833 | 18.86090713 | 8 mo | HOM |
| **1013288213** | 2 | TT | 589 | 1.484552138 | 0.010638678 | 10 | 0.007166255 | 6.736038261 | 4 mo | HOM |
| **1008976060** | 2 | TT | 589 | 1.484552138 | 0.000295796 | 1 | 0.00019925 | 0.673603826 | 8 mo | WILD |
| **1013446243** | 1 | TT | 589 | 1.484552138 | 0.001020169 | 3 | 0.00068719 | 2.020811478 | 4 mo | HOM |
| **1013288951** | 2 | TT | 589 | 1.484552138 | 0.001775044 | 19 | 0.001195677 | 12.7984727 | 4 mo | HOM |
| **1013978665** | 1 | RSPv6a | 590 | 0.679670376 | 0 | 0 | 0 | 0 | 2 mo | HOM |
| **1013445609** | 1 | RSPv6a | 590 | 0.679670376 | 1.44E-05 | 1 | 2.12E-05 | 1.471301436 | 4 mo | HOM |
| **1013609015** | 1 | RSPv6a | 590 | 0.679670376 | 2.17E-05 | 2 | 3.2E-05 | 2.942602871 | 4 mo | HOM |
| **1013608106** | 2 | RSPv6a | 590 | 0.679670376 | 0 | 0 | 0 | 0 | 4 mo | HOM |
| **1011937543** | 1 | RSPv6a | 590 | 0.679670376 | 0.00128556 | 13 | 0.001891446 | 19.12691866 | 8 mo | HOM |
| **1010419538** | 2 | RSPv6a | 590 | 0.679668651 | 0 | 0 | 0 | 0 | 4 mo | WILD |
| **1010419187** | 1 | RSPv6a | 590 | 0.679670376 | 0.000762543 | 19 | 0.00112193 | 27.95472728 | 8 mo | HOM |
| **1013978366** | 1 | RSPv6a | 590 | 0.679670376 | 0 | 0 | 0 | 0 | 2 mo | HOM |
| **1014416155** | 2 | RSPv6a | 590 | 0.679670376 | 0 | 0 | 0 | 0 | 2 mo | HOM |
| **1013978065** | 2 | RSPv6a | 590 | 0.679670376 | 0 | 0 | 0 | 0 | 2 mo | HOM |
| **1009934102** | 2 | RSPv6a | 590 | 0.679670376 | 0 | 0 | 0 | 0 | 4 mo | WILD |
| **1009322937** | 2 | RSPv6a | 590 | 0.679670376 | 0 | 0 | 0 | 0 | 8 mo | WILD |
| **1011080637** | 2 | RSPv6a | 590 | 0.679670376 | 0.000686218 | 21 | 0.001009634 | 30.89733015 | 8 mo | HOM |
| **1013609713** | 2 | RSPv6a | 590 | 0.679670376 | 0 | 0 | 0 | 0 | 2 mo | WILD |
| **1010418842** | 1 | RSPv6a | 590 | 0.679670376 | 0.001217792 | 31 | 0.001791739 | 45.61034451 | 8 mo | HOM |
| **1013288213** | 2 | RSPv6a | 590 | 0.679670376 | 1.22E-05 | 1 | 1.79E-05 | 1.471301436 | 4 mo | HOM |
| **1008976060** | 2 | RSPv6a | 590 | 0.679670376 | 0 | 0 | 0 | 0 | 8 mo | WILD |
| **1013446243** | 1 | RSPv6a | 590 | 0.679670376 | 0.000122304 | 6 | 0.000179946 | 8.827808614 | 4 mo | HOM |
| **1013288951** | 2 | RSPv6a | 590 | 0.679670376 | 1.82E-05 | 2 | 2.68E-05 | 2.942602871 | 4 mo | HOM |
| **1013978665** | 1 | CLI | 591 | 0.085689995 | 0 | 0 | 0 | 0 | 2 mo | HOM |
| **1013445609** | 1 | CLI | 591 | 0.085689995 | 0 | 0 | 0 | 0 | 4 mo | HOM |
| **1013609015** | 1 | CLI | 591 | 0.085689995 | 0 | 0 | 0 | 0 | 4 mo | HOM |
| **1013608106** | 2 | CLI | 591 | 0.085689995 | 0 | 0 | 0 | 0 | 4 mo | HOM |
| **1011937543** | 1 | CLI | 591 | 0.085689995 | 0 | 0 | 0 | 0 | 8 mo | HOM |
| **1010419538** | 2 | CLI | 591 | 0.085689995 | 0 | 0 | 0 | 0 | 4 mo | WILD |
| **1010419187** | 1 | CLI | 591 | 0.085689995 | 0 | 0 | 0 | 0 | 8 mo | HOM |
| **1013978366** | 1 | CLI | 591 | 0.085689995 | 0 | 0 | 0 | 0 | 2 mo | HOM |
| **1014416155** | 2 | CLI | 591 | 0.085689995 | 0 | 0 | 0 | 0 | 2 mo | HOM |
| **1013978065** | 2 | CLI | 591 | 0.085689995 | 0 | 0 | 0 | 0 | 2 mo | HOM |
| **1009934102** | 2 | CLI | 591 | 0.085689995 | 0 | 0 | 0 | 0 | 4 mo | WILD |
| **1009322937** | 2 | CLI | 591 | 0.085689995 | 0 | 0 | 0 | 0 | 8 mo | WILD |
| **1011080637** | 2 | CLI | 591 | 0.085689995 | 0 | 0 | 0 | 0 | 8 mo | HOM |
| **1013609713** | 2 | CLI | 591 | 0.085689995 | 0 | 0 | 0 | 0 | 2 mo | WILD |
| **1010418842** | 1 | CLI | 591 | 0.085689995 | 0 | 0 | 0 | 0 | 8 mo | HOM |
| **1013288213** | 2 | CLI | 591 | 0.085689995 | 0 | 0 | 0 | 0 | 4 mo | HOM |
| **1008976060** | 2 | CLI | 591 | 0.085689995 | 0 | 0 | 0 | 0 | 8 mo | WILD |
| **1013446243** | 1 | CLI | 591 | 0.085689995 | 0 | 0 | 0 | 0 | 4 mo | HOM |
| **1013288951** | 2 | CLI | 591 | 0.085689995 | 0 | 0 | 0 | 0 | 4 mo | HOM |
| **1013978665** | 1 | VISp1 | 593 | 1.301617184 | 0.000668456 | 24 | 0.000513558 | 18.43860107 | 2 mo | HOM |
| **1013445609** | 1 | VISp1 | 593 | 1.301617184 | 0.000188622 | 14 | 0.000144913 | 10.75585062 | 4 mo | HOM |
| **1013609015** | 1 | VISp1 | 593 | 1.301617184 | 0.00068444 | 18 | 0.000525839 | 13.8289508 | 4 mo | HOM |
| **1013608106** | 2 | VISp1 | 593 | 1.301617184 | 0 | 0 | 0 | 0 | 4 mo | HOM |
| **1011937543** | 1 | VISp1 | 593 | 1.301617184 | 0.004928357 | 32 | 0.003786334 | 24.58480142 | 8 mo | HOM |
| **1010419538** | 2 | VISp1 | 593 | 1.301617184 | 0.000297713 | 10 | 0.000228726 | 7.682750445 | 4 mo | WILD |

| | | | | | | | | | | |
|---|---|---|---|---|---|---|---|---|---|---|
| **1010419187** | 1 | VISp1 | 593 | 1.301617184 | 0.013842115 | 68 | 0.010634552 | 52.24270303 | 8 mo | HOM |
| **1013978366** | 1 | VISp1 | 593 | 1.301617184 | 0.000618251 | 13 | 0.000474987 | 9.987575579 | 2 mo | HOM |
| **1014416155** | 2 | VISp1 | 593 | 1.301617184 | 0 | 0 | 0 | 0 | 2 mo | HOM |
| **1013978065** | 2 | VISp1 | 593 | 1.301617184 | 1.4E-05 | 2 | 1.08E-05 | 1.536550089 | 2 mo | HOM |
| **1009934102** | 2 | VISp1 | 593 | 1.301617184 | 0.000257224 | 8 | 0.000197618 | 6.146200356 | 4 mo | WILD |
| **1009322937** | 2 | VISp1 | 593 | 1.301617184 | 7.66E-05 | 7 | 5.89E-05 | 5.377925312 | 8 mo | WILD |
| **1011080637** | 2 | VISp1 | 593 | 1.301617184 | 0.003372859 | 68 | 0.002591283 | 52.24270303 | 8 mo | HOM |
| **1013609713** | 2 | VISp1 | 593 | 1.301617184 | 5.43E-05 | 4 | 4.17E-05 | 3.073100178 | 2 mo | WILD |
| **1010418842** | 1 | VISp1 | 593 | 1.301617184 | 0.004629387 | 55 | 0.003556643 | 42.25512745 | 8 mo | HOM |
| **1013288213** | 2 | VISp1 | 593 | 1.301617184 | 0.000163522 | 13 | 0.00012563 | 9.987575579 | 4 mo | HOM |
| **1008976060** | 2 | VISp1 | 593 | 1.301617184 | 0.00190459 | 21 | 0.001463249 | 16.13377594 | 8 mo | WILD |
| **1013446243** | 1 | VISp1 | 593 | 1.301617184 | 0.000473471 | 23 | 0.000363756 | 17.67032602 | 4 mo | HOM |
| **1013288951** | 2 | VISp1 | 593 | 1.301617184 | 0.003659686 | 25 | 0.002811646 | 19.20687611 | 4 mo | HOM |
| **1013978665** | 1 | fr | 595 | 0.172688368 | 0 | 0 | 0 | 0 | 2 mo | HOM |
| **1013445609** | 1 | fr | 595 | 0.172688368 | 0 | 0 | 0 | 0 | 4 mo | HOM |
| **1013609015** | 1 | fr | 595 | 0.172688368 | 0 | 0 | 0 | 0 | 4 mo | HOM |
| **1013608106** | 2 | fr | 595 | 0.172688368 | 0 | 0 | 0 | 0 | 4 mo | HOM |
| **1011937543** | 1 | fr | 595 | 0.172688368 | 2.68E-05 | 0 | 0.00015493 | 0 | 8 mo | HOM |
| **1010419538** | 2 | fr | 595 | 0.172688368 | 0 | 0 | 0 | 0 | 4 mo | WILD |
| **1010419187** | 1 | fr | 595 | 0.172688368 | 0 | 0 | 0 | 0 | 8 mo | HOM |
| **1013978366** | 1 | fr | 595 | 0.172688368 | 0 | 0 | 0 | 0 | 2 mo | HOM |
| **1014416155** | 2 | fr | 595 | 0.172688368 | 0 | 0 | 0 | 0 | 2 mo | HOM |
| **1013978065** | 2 | fr | 595 | 0.172688368 | 0 | 0 | 0 | 0 | 2 mo | HOM |
| **1009934102** | 2 | fr | 595 | 0.172688368 | 0 | 0 | 0 | 0 | 4 mo | WILD |
| **1009322937** | 2 | fr | 595 | 0.172688368 | 0 | 0 | 0 | 0 | 8 mo | WILD |
| **1011080637** | 2 | fr | 595 | 0.172688368 | 1.49E-06 | 0 | 8.63E-06 | 0 | 8 mo | HOM |
| **1013609713** | 2 | fr | 595 | 0.172688368 | 0 | 0 | 0 | 0 | 2 mo | WILD |
| **1010418842** | 1 | fr | 595 | 0.172688368 | 1.19E-08 | 0 | 6.9E-08 | 0 | 8 mo | HOM |
| **1013288213** | 2 | fr | 595 | 0.172688368 | 0 | 0 | 0 | 0 | 4 mo | HOM |
| **1008976060** | 2 | fr | 595 | 0.172688368 | 0 | 0 | 0 | 0 | 8 mo | WILD |
| **1013446243** | 1 | fr | 595 | 0.172688368 | 0 | 0 | 0 | 0 | 4 mo | HOM |
| **1013288951** | 2 | fr | 595 | 0.172688368 | 0 | 0 | 0 | 0 | 4 mo | HOM |
| **1013978665** | 1 | NDB | 596 | 0.76229645 | 0 | 0 | 0 | 0 | 2 mo | HOM |
| **1013445609** | 1 | NDB | 596 | 0.76229645 | 0 | 0 | 0 | 0 | 4 mo | HOM |
| **1013609015** | 1 | NDB | 596 | 0.76229645 | 0 | 0 | 0 | 0 | 4 mo | HOM |
| **1013608106** | 2 | NDB | 596 | 0.76229645 | 0 | 0 | 0 | 0 | 4 mo | HOM |
| **1011937543** | 1 | NDB | 596 | 0.76229645 | 0.00011959 | 3 | 0.000156881 | 3.93547681 | 8 mo | HOM |
| **1010419538** | 2 | NDB | 596 | 0.76229645 | 0 | 0 | 0 | 0 | 4 mo | WILD |
| **1010419187** | 1 | NDB | 596 | 0.76229645 | 8.95E-06 | 1 | 1.17E-05 | 1.311825603 | 8 mo | HOM |
| **1013978366** | 1 | NDB | 596 | 0.76229645 | 0 | 0 | 0 | 0 | 2 mo | HOM |
| **1014416155** | 2 | NDB | 596 | 0.76229645 | 0 | 0 | 0 | 0 | 2 mo | HOM |
| **1013978065** | 2 | NDB | 596 | 0.76229645 | 3.07E-09 | 0 | 4.03E-09 | 0 | 2 mo | HOM |
| **1009934102** | 2 | NDB | 596 | 0.76229645 | 7.99E-07 | 0 | 1.05E-06 | 0 | 4 mo | WILD |
| **1009322937** | 2 | NDB | 596 | 0.76229645 | 0 | 0 | 0 | 0 | 8 mo | WILD |
| **1011080637** | 2 | NDB | 596 | 0.76229645 | 5.56E-06 | 1 | 7.3E-06 | 1.311825603 | 8 mo | HOM |
| **1013609713** | 2 | NDB | 596 | 0.76229645 | 0 | 0 | 0 | 0 | 2 mo | WILD |
| **1010418842** | 1 | NDB | 596 | 0.76229645 | 0.000143161 | 5 | 0.000187802 | 6.559128017 | 8 mo | HOM |
| **1013288213** | 2 | NDB | 596 | 0.76229645 | 0 | 0 | 0 | 0 | 4 mo | HOM |
| **1008976060** | 2 | NDB | 596 | 0.76229645 | 2.09E-07 | 0 | 2.74E-07 | 0 | 8 mo | WILD |
| **1013446243** | 1 | NDB | 596 | 0.76229645 | 0 | 0 | 0 | 0 | 4 mo | HOM |
| **1013288951** | 2 | NDB | 596 | 0.76229645 | 0 | 0 | 0 | 0 | 4 mo | HOM |
| **1013978665** | 1 | TTd | 597 | 0.745233866 | 0 | 0 | 0 | 0 | 2 mo | HOM |
| **1013445609** | 1 | TTd | 597 | 0.745233866 | 7.19E-08 | 0 | 9.65E-08 | 0 | 4 mo | HOM |
| **1013609015** | 1 | TTd | 597 | 0.745233866 | 6.72E-05 | 3 | 9.01E-05 | 4.025581953 | 4 mo | HOM |
| **1013608106** | 2 | TTd | 597 | 0.745233866 | 0 | 0 | 0 | 0 | 4 mo | HOM |
| **1011937543** | 1 | TTd | 597 | 0.745233866 | 0.000306817 | 6 | 0.000411706 | 8.051163906 | 8 mo | HOM |
| **1010419538** | 2 | TTd | 597 | 0.745233866 | 0 | 0 | 0 | 0 | 4 mo | WILD |
| **1010419187** | 1 | TTd | 597 | 0.745233866 | 0.000365252 | 16 | 0.000490117 | 21.46977042 | 8 mo | HOM |
| **1013978366** | 1 | TTd | 597 | 0.745233866 | 0 | 0 | 0 | 0 | 2 mo | HOM |

| | | | | | | | | | | |
|---|---|---|---|---|---|---|---|---|---|---|
| 1014416155 | 2 | TTd | 597 | 0.745233866 | 3.19E-06 | 1 | 4.28E-06 | 1.341860651 | 2 mo | HOM |
| 1013978065 | 2 | TTd | 597 | 0.745233866 | 0 | 0 | 0 | 0 | 2 mo | HOM |
| 1009934102 | 2 | TTd | 597 | 0.745233866 | 2.98E-06 | 1 | 4E-06 | 1.341860651 | 4 mo | WILD |
| 1009322937 | 2 | TTd | 597 | 0.745233866 | 0.000535812 | 8 | 0.000718985 | 10.73488521 | 8 mo | WILD |
| 1011080637 | 2 | TTd | 597 | 0.745233866 | 0.000361489 | 17 | 0.000485068 | 22.81163107 | 8 mo | HOM |
| 1013609713 | 2 | TTd | 597 | 0.745233866 | 0 | 0 | 0 | 0 | 2 mo | WILD |
| 1010418842 | 1 | TTd | 597 | 0.745233866 | 0.003194371 | 20 | 0.004286401 | 26.83721302 | 8 mo | HOM |
| 1013288213 | 2 | TTd | 597 | 0.745233866 | 0.004782404 | 6 | 0.00641732 | 8.051163906 | 4 mo | HOM |
| 1008976060 | 2 | TTd | 597 | 0.745233866 | 9.05E-05 | 1 | 0.000121483 | 1.341860651 | 8 mo | WILD |
| 1013446243 | 1 | TTd | 597 | 0.745233866 | 0.000427728 | 1 | 0.000573951 | 1.341860651 | 4 mo | HOM |
| 1013288951 | 2 | TTd | 597 | 0.745233866 | 0.000227932 | 8 | 0.000305854 | 10.73488521 | 4 mo | HOM |
| 1013978665 | 1 | AUDv6b | 598 | 0.047441859 | 0 | 0 | 0 | 0 | 2 mo | HOM |
| 1013445609 | 1 | AUDv6b | 598 | 0.047441859 | 4.42E-06 | 0 | 9.32E-05 | 0 | 4 mo | HOM |
| 1013609015 | 1 | AUDv6b | 598 | 0.047441859 | 0 | 0 | 0 | 0 | 4 mo | HOM |
| 1013608106 | 2 | AUDv6b | 598 | 0.047441859 | 0 | 0 | 0 | 0 | 4 mo | HOM |
| 1011937543 | 1 | AUDv6b | 598 | 0.047441859 | 7.32E-05 | 3 | 0.001542475 | 63.23529634 | 8 mo | HOM |
| 1010419538 | 2 | AUDv6b | 598 | 0.047441859 | 0 | 0 | 0 | 0 | 4 mo | WILD |
| 1010419187 | 1 | AUDv6b | 598 | 0.047441859 | 8.09E-05 | 6 | 0.001705092 | 126.4705927 | 8 mo | HOM |
| 1013978366 | 1 | AUDv6b | 598 | 0.047441859 | 0 | 0 | 0 | 0 | 2 mo | HOM |
| 1014416155 | 2 | AUDv6b | 598 | 0.047441859 | 0 | 0 | 0 | 0 | 2 mo | HOM |
| 1013978065 | 2 | AUDv6b | 598 | 0.047441859 | 0 | 0 | 0 | 0 | 2 mo | HOM |
| 1009934102 | 2 | AUDv6b | 598 | 0.047441859 | 0 | 0 | 0 | 0 | 4 mo | WILD |
| 1009322937 | 2 | AUDv6b | 598 | 0.047441859 | 0 | 0 | 0 | 0 | 8 mo | WILD |
| 1011080637 | 2 | AUDv6b | 598 | 0.047441859 | 7.92E-07 | 0 | 1.67E-05 | 0 | 8 mo | HOM |
| 1013609713 | 2 | AUDv6b | 598 | 0.047441859 | 0 | 0 | 0 | 0 | 2 mo | WILD |
| 1010418842 | 1 | AUDv6b | 598 | 0.047441859 | 3.95E-05 | 3 | 0.000832997 | 63.23529634 | 8 mo | HOM |
| 1013288213 | 2 | AUDv6b | 598 | 0.047441859 | 0 | 0 | 0 | 0 | 4 mo | HOM |
| 1008976060 | 2 | AUDv6b | 598 | 0.047441859 | 0 | 0 | 0 | 0 | 8 mo | WILD |
| 1013446243 | 1 | AUDv6b | 598 | 0.047441859 | 1.47E-05 | 1 | 0.000310227 | 21.07843211 | 4 mo | HOM |
| 1013288951 | 2 | AUDv6b | 598 | 0.047441859 | 1.56E-05 | 1 | 0.000329752 | 21.07843211 | 4 mo | HOM |
| 1013978665 | 1 | CM | 599 | 0.257113251 | 0 | 0 | 0 | 0 | 2 mo | HOM |
| 1013445609 | 1 | CM | 599 | 0.257113251 | 0 | 0 | 0 | 0 | 4 mo | HOM |
| 1013609015 | 1 | CM | 599 | 0.257113251 | 3.84E-06 | 1 | 1.49E-05 | 3.889336691 | 4 mo | HOM |
| 1013608106 | 2 | CM | 599 | 0.257113251 | 0 | 0 | 0 | 0 | 4 mo | HOM |
| 1011937543 | 1 | CM | 599 | 0.257113251 | 1.62E-05 | 1 | 6.31E-05 | 3.889336691 | 8 mo | HOM |
| 1010419538 | 2 | CM | 599 | 0.257113251 | 0 | 0 | 0 | 0 | 4 mo | WILD |
| 1010419187 | 1 | CM | 599 | 0.257113251 | 1.48E-05 | 1 | 5.77E-05 | 3.889336691 | 8 mo | HOM |
| 1013978366 | 1 | CM | 599 | 0.257113251 | 0 | 0 | 0 | 0 | 2 mo | HOM |
| 1014416155 | 2 | CM | 599 | 0.257113251 | 0 | 0 | 0 | 0 | 2 mo | HOM |
| 1013978065 | 2 | CM | 599 | 0.257113251 | 0 | 0 | 0 | 0 | 2 mo | HOM |
| 1009934102 | 2 | CM | 599 | 0.257113251 | 0 | 0 | 0 | 0 | 4 mo | WILD |
| 1009322937 | 2 | CM | 599 | 0.257113251 | 0 | 0 | 0 | 0 | 8 mo | WILD |
| 1011080637 | 2 | CM | 599 | 0.257113251 | 4.86E-06 | 0 | 1.89E-05 | 0 | 8 mo | HOM |
| 1013609713 | 2 | CM | 599 | 0.257113251 | 0 | 0 | 0 | 0 | 2 mo | WILD |
| 1010418842 | 1 | CM | 599 | 0.257113251 | 9.46E-05 | 3 | 0.000367745 | 11.66801007 | 8 mo | HOM |
| 1013288213 | 2 | CM | 599 | 0.257113251 | 0 | 0 | 0 | 0 | 4 mo | HOM |
| 1008976060 | 2 | CM | 599 | 0.257113251 | 2.47E-05 | 2 | 9.62E-05 | 7.778673381 | 8 mo | WILD |
| 1013446243 | 1 | CM | 599 | 0.257113251 | 1.25E-05 | 1 | 4.85E-05 | 3.889336691 | 4 mo | HOM |
| 1013288951 | 2 | CM | 599 | 0.257113251 | 0 | 0 | 0 | 0 | 4 mo | HOM |
| 1013978665 | 1 | AUDd2/3 | 600 | 0.33725034 | 0 | 0 | 0 | 0 | 2 mo | HOM |
| 1013445609 | 1 | AUDd2/3 | 600 | 0.33725034 | 2.68E-05 | 2 | 7.96E-05 | 5.930312776 | 4 mo | HOM |
| 1013609015 | 1 | AUDd2/3 | 600 | 0.33725034 | 0.000103175 | 4 | 0.000305931 | 11.86062555 | 4 mo | HOM |
| 1013608106 | 2 | AUDd2/3 | 600 | 0.33725034 | 0 | 0 | 0 | 0 | 4 mo | HOM |
| 1011937543 | 1 | AUDd2/3 | 600 | 0.33725034 | 0.000219337 | 6 | 0.000650368 | 17.79093833 | 8 mo | HOM |
| 1010419538 | 2 | AUDd2/3 | 600 | 0.33725034 | 0 | 0 | 0 | 0 | 4 mo | WILD |
| 1010419187 | 1 | AUDd2/3 | 600 | 0.33725034 | 0.000567279 | 16 | 0.00168207 | 47.44250221 | 8 mo | HOM |
| 1013978366 | 1 | AUDd2/3 | 600 | 0.33725034 | 0 | 0 | 0 | 0 | 2 mo | HOM |
| 1014416155 | 2 | AUDd2/3 | 600 | 0.33725034 | 0 | 0 | 0 | 0 | 2 mo | HOM |
| 1013978065 | 2 | AUDd2/3 | 600 | 0.33725034 | 0 | 0 | 0 | 0 | 2 mo | HOM |

| | | | | | | | | | | |
|---|---|---|---|---|---|---|---|---|---|---|
| **1009934102** | 2 | AUDd2/3 | 600 | 0.33725034 | 0 | 0 | 0 | 0 | 4 mo | WILD |
| **1009322937** | 2 | AUDd2/3 | 600 | 0.33725034 | 0 | 0 | 0 | 0 | 8 mo | WILD |
| **1011080637** | 2 | AUDd2/3 | 600 | 0.33725034 | 8.36E-05 | 5 | 0.000248005 | 14.82578194 | 8 mo | HOM |
| **1013609713** | 2 | AUDd2/3 | 600 | 0.33725034 | 0 | 0 | 0 | 0 | 2 mo | WILD |
| **1010418842** | 1 | AUDd2/3 | 600 | 0.33725034 | 0.000161061 | 4 | 0.000477571 | 11.86062555 | 8 mo | HOM |
| **1013288213** | 2 | AUDd2/3 | 600 | 0.33725034 | 2.42E-05 | 2 | 7.16E-05 | 5.930312776 | 4 mo | HOM |
| **1008976060** | 2 | AUDd2/3 | 600 | 0.33725034 | 0 | 0 | 0 | 0 | 8 mo | WILD |
| **1013446243** | 1 | AUDd2/3 | 600 | 0.33725034 | 0.000129027 | 3 | 0.000382586 | 8.895469164 | 4 mo | HOM |
| **1013288951** | 2 | AUDd2/3 | 600 | 0.33725034 | 3.06E-05 | 3 | 9.08E-05 | 8.895469164 | 4 mo | HOM |
| **1013978665** | 1 | VISal6a | 601 | 0.115049344 | 0 | 0 | 0 | 0 | 2 mo | HOM |
| **1013445609** | 1 | VISal6a | 601 | 0.115049344 | 9.19E-06 | 1 | 7.99E-05 | 8.691922644 | 4 mo | HOM |
| **1013609015** | 1 | VISal6a | 601 | 0.115049344 | 1.96E-05 | 1 | 0.000170076 | 8.691922644 | 4 mo | HOM |
| **1013608106** | 2 | VISal6a | 601 | 0.115049344 | 0 | 0 | 0 | 0 | 4 mo | HOM |
| **1011937543** | 1 | VISal6a | 601 | 0.115049344 | 0.000105125 | 4 | 0.000913738 | 34.76769057 | 8 mo | HOM |
| **1010419538** | 2 | VISal6a | 601 | 0.115049344 | 0 | 0 | 0 | 0 | 4 mo | WILD |
| **1010419187** | 1 | VISal6a | 601 | 0.115049344 | 0.000120696 | 6 | 0.001049081 | 52.15153586 | 8 mo | HOM |
| **1013978366** | 1 | VISal6a | 601 | 0.115049344 | 0 | 0 | 0 | 0 | 2 mo | HOM |
| **1014416155** | 2 | VISal6a | 601 | 0.115049344 | 0 | 0 | 0 | 0 | 2 mo | HOM |
| **1013978065** | 2 | VISal6a | 601 | 0.115049344 | 0 | 0 | 0 | 0 | 2 mo | HOM |
| **1009934102** | 2 | VISal6a | 601 | 0.115049344 | 0 | 0 | 0 | 0 | 4 mo | WILD |
| **1009322937** | 2 | VISal6a | 601 | 0.115049344 | 0 | 0 | 0 | 0 | 8 mo | WILD |
| **1011080637** | 2 | VISal6a | 601 | 0.115049344 | 8E-05 | 3 | 0.000695289 | 26.07576793 | 8 mo | HOM |
| **1013609713** | 2 | VISal6a | 601 | 0.115049344 | 0 | 0 | 0 | 0 | 2 mo | WILD |
| **1010418842** | 1 | VISal6a | 601 | 0.115049344 | 3.2E-05 | 3 | 0.000277821 | 26.07576793 | 8 mo | HOM |
| **1013288213** | 2 | VISal6a | 601 | 0.115049344 | 6.6E-07 | 0 | 5.73E-06 | 0 | 4 mo | HOM |
| **1008976060** | 2 | VISal6a | 601 | 0.115049344 | 0 | 0 | 0 | 0 | 8 mo | WILD |
| **1013446243** | 1 | VISal6a | 601 | 0.115049344 | 1.14E-05 | 2 | 9.88E-05 | 17.38384529 | 4 mo | HOM |
| **1013288951** | 2 | VISal6a | 601 | 0.115049344 | 0 | 0 | 0 | 0 | 4 mo | HOM |
| **1013978665** | 1 | fi | 603 | 1.559507242 | 0 | 0 | 0 | 0 | 2 mo | HOM |
| **1013445609** | 1 | fi | 603 | 1.559507242 | 6.43E-06 | 1 | 4.12E-06 | 0.641228186 | 4 mo | HOM |
| **1013609015** | 1 | fi | 603 | 1.559507242 | 0.001077474 | 16 | 0.000690907 | 10.25965098 | 4 mo | HOM |
| **1013608106** | 2 | fi | 603 | 1.559507242 | 0 | 0 | 0 | 0 | 4 mo | HOM |
| **1011937543** | 1 | fi | 603 | 1.559507242 | 0.000690262 | 11 | 0.000442615 | 7.053510049 | 8 mo | HOM |
| **1010419538** | 2 | fi | 603 | 1.559507242 | 0 | 0 | 0 | 0 | 4 mo | WILD |
| **1010419187** | 1 | fi | 603 | 1.559507242 | 0.001860365 | 33 | 0.001192919 | 21.16053015 | 8 mo | HOM |
| **1013978366** | 1 | fi | 603 | 1.559507242 | 2.21E-05 | 1 | 1.41E-05 | 0.641228186 | 2 mo | HOM |
| **1014416155** | 2 | fi | 603 | 1.559507242 | 0 | 0 | 0 | 0 | 2 mo | HOM |
| **1013978065** | 2 | fi | 603 | 1.559507242 | 0 | 0 | 0 | 0 | 2 mo | HOM |
| **1009934102** | 2 | fi | 603 | 1.559507242 | 3.86E-09 | 0 | 2.47E-09 | 0 | 4 mo | WILD |
| **1009322937** | 2 | fi | 603 | 1.559507242 | 0 | 0 | 0 | 0 | 8 mo | WILD |
| **1011080637** | 2 | fi | 603 | 1.559507242 | 5.17E-05 | 1 | 3.32E-05 | 0.641228186 | 8 mo | HOM |
| **1013609713** | 2 | fi | 603 | 1.559507242 | 0 | 0 | 0 | 0 | 2 mo | WILD |
| **1010418842** | 1 | fi | 603 | 1.559507242 | 0.001762519 | 33 | 0.001130177 | 21.16053015 | 8 mo | HOM |
| **1013288213** | 2 | fi | 603 | 1.559507242 | 0 | 0 | 0 | 0 | 4 mo | HOM |
| **1008976060** | 2 | fi | 603 | 1.559507242 | 4.03E-05 | 0 | 2.58E-05 | 0 | 8 mo | WILD |
| **1013446243** | 1 | fi | 603 | 1.559507242 | 0.000358436 | 7 | 0.000229839 | 4.488597304 | 4 mo | HOM |
| **1013288951** | 2 | fi | 603 | 1.559507242 | 0.000449803 | 1 | 0.000288426 | 0.641228186 | 4 mo | HOM |
| **1013978665** | 1 | NI | 604 | 0.111501254 | 0 | 0 | 0 | 0 | 2 mo | HOM |
| **1013445609** | 1 | NI | 604 | 0.111501254 | 0 | 0 | 0 | 0 | 4 mo | HOM |
| **1013609015** | 1 | NI | 604 | 0.111501254 | 0 | 0 | 0 | 0 | 4 mo | HOM |
| **1013608106** | 2 | NI | 604 | 0.111501254 | 0 | 0 | 0 | 0 | 4 mo | HOM |
| **1011937543** | 1 | NI | 604 | 0.111501254 | 0 | 0 | 0 | 0 | 8 mo | HOM |
| **1010419538** | 2 | NI | 604 | 0.111501254 | 0 | 0 | 0 | 0 | 4 mo | WILD |
| **1010419187** | 1 | NI | 604 | 0.111501254 | 0 | 0 | 0 | 0 | 8 mo | HOM |
| **1013978366** | 1 | NI | 604 | 0.111501254 | 0 | 0 | 0 | 0 | 2 mo | HOM |
| **1014416155** | 2 | NI | 604 | 0.111501254 | 0 | 0 | 0 | 0 | 2 mo | HOM |
| **1013978065** | 2 | NI | 604 | 0.111501254 | 0 | 0 | 0 | 0 | 2 mo | HOM |
| **1009934102** | 2 | NI | 604 | 0.111501254 | 0 | 0 | 0 | 0 | 4 mo | WILD |
| **1009322937** | 2 | NI | 604 | 0.111501254 | 0 | 0 | 0 | 0 | 8 mo | WILD |

| | | | | | | | | | | |
|---|---|---|---|---|---|---|---|---|---|---|
| **1011080637** | 2 | NI | 604 | 0.111501254 | 3.26E-05 | 1 | 0.000292223 | 8.968508984 | 8 mo | HOM |
| **1013609713** | 2 | NI | 604 | 0.111501254 | 0 | 0 | 0 | 0 | 2 mo | WILD |
| **1010418842** | 1 | NI | 604 | 0.111501254 | 7.19E-06 | 1 | 6.45E-05 | 8.968508984 | 8 mo | HOM |
| **1013288213** | 2 | NI | 604 | 0.111501254 | 0 | 0 | 0 | 0 | 4 mo | HOM |
| **1008976060** | 2 | NI | 604 | 0.111501254 | 2.4E-06 | 0 | 2.15E-05 | 0 | 8 mo | WILD |
| **1013446243** | 1 | NI | 604 | 0.111501254 | 5.37E-08 | 0 | 4.82E-07 | 0 | 4 mo | HOM |
| **1013288951** | 2 | NI | 604 | 0.111501254 | 0 | 0 | 0 | 0 | 4 mo | HOM |
| **1013978665** | 1 | TTv | 605 | 0.739318272 | 0 | 0 | 0 | 0 | 2 mo | HOM |
| **1013445609** | 1 | TTv | 605 | 0.739318272 | 5.3E-06 | 1 | 7.17E-06 | 1.352597437 | 4 mo | HOM |
| **1013609015** | 1 | TTv | 605 | 0.739318272 | 6.95E-05 | 4 | 9.4E-05 | 5.410389749 | 4 mo | HOM |
| **1013608106** | 2 | TTv | 605 | 0.739318272 | 0 | 0 | 0 | 0 | 4 mo | HOM |
| **1011937543** | 1 | TTv | 605 | 0.739318272 | 0.000156203 | 5 | 0.00021128 | 6.762987186 | 8 mo | HOM |
| **1010419538** | 2 | TTv | 605 | 0.739318272 | 0 | 0 | 0 | 0 | 4 mo | WILD |
| **1010419187** | 1 | TTv | 605 | 0.739318272 | 0.000525343 | 8 | 0.000710578 | 10.8207795 | 8 mo | HOM |
| **1013978366** | 1 | TTv | 605 | 0.739318272 | 0 | 0 | 0 | 0 | 2 mo | HOM |
| **1014416155** | 2 | TTv | 605 | 0.739318272 | 4.8E-06 | 1 | 6.49E-06 | 1.352597437 | 2 mo | HOM |
| **1013978065** | 2 | TTv | 605 | 0.739318272 | 1.17E-06 | 0 | 1.58E-06 | 0 | 2 mo | HOM |
| **1009934102** | 2 | TTv | 605 | 0.739318272 | 3.25E-06 | 0 | 4.4E-06 | 0 | 4 mo | WILD |
| **1009322937** | 2 | TTv | 605 | 0.739318272 | 8.42E-05 | 2 | 0.000113933 | 2.705194874 | 8 mo | WILD |
| **1011080637** | 2 | TTv | 605 | 0.739318272 | 0.000225241 | 8 | 0.000304661 | 10.8207795 | 8 mo | HOM |
| **1013609713** | 2 | TTv | 605 | 0.739318272 | 0 | 0 | 0 | 0 | 2 mo | WILD |
| **1010418842** | 1 | TTv | 605 | 0.739318272 | 0.00027181 | 8 | 0.000367649 | 10.8207795 | 8 mo | HOM |
| **1013288213** | 2 | TTv | 605 | 0.739318272 | 0.005856274 | 4 | 0.007921182 | 5.410389749 | 4 mo | HOM |
| **1008976060** | 2 | TTv | 605 | 0.739318272 | 0.000205263 | 0 | 0.000277639 | 0 | 8 mo | WILD |
| **1013446243** | 1 | TTv | 605 | 0.739318272 | 0.000592441 | 2 | 0.000801334 | 2.705194874 | 4 mo | HOM |
| **1013288951** | 2 | TTv | 605 | 0.739318272 | 0.001547112 | 11 | 0.002092619 | 14.87857181 | 4 mo | HOM |
| **1013978665** | 1 | CN | 607 | 1.678527066 | 3.34E-05 | 4 | 1.99E-05 | 2.383041705 | 2 mo | HOM |
| **1013445609** | 1 | CN | 607 | 1.678527066 | 0.000160936 | 4 | 9.59E-05 | 2.383041705 | 4 mo | HOM |
| **1013609015** | 1 | CN | 607 | 1.678527066 | 0.00291303 | 8 | 0.001735468 | 4.76608341 | 4 mo | HOM |
| **1013608106** | 2 | CN | 607 | 1.678527066 | 0 | 0 | 0 | 0 | 4 mo | HOM |
| **1011937543** | 1 | CN | 607 | 1.678527066 | 0.004647481 | 6 | 0.002768785 | 3.574562557 | 8 mo | HOM |
| **1010419538** | 2 | CN | 607 | 1.678527066 | 2.57E-05 | 5 | 1.53E-05 | 2.978802131 | 4 mo | WILD |
| **1010419187** | 1 | CN | 607 | 1.678527066 | 0.000264002 | 8 | 0.000157282 | 4.76608341 | 8 mo | HOM |
| **1013978366** | 1 | CN | 607 | 1.678527066 | 0.001062282 | 5 | 0.000632866 | 2.978802131 | 2 mo | HOM |
| **1014416155** | 2 | CN | 607 | 1.678527066 | 0 | 0 | 0 | 0 | 2 mo | HOM |
| **1013978065** | 2 | CN | 607 | 1.678527066 | 7.26E-05 | 3 | 4.33E-05 | 1.787281279 | 2 mo | HOM |
| **1009934102** | 2 | CN | 607 | 1.678527066 | 1.91E-05 | 2 | 1.14E-05 | 1.191520852 | 4 mo | WILD |
| **1009322937** | 2 | CN | 607 | 1.678527066 | 0.000151711 | 6 | 9.04E-05 | 3.574562557 | 8 mo | WILD |
| **1011080637** | 2 | CN | 607 | 1.678527066 | 0.000617894 | 20 | 0.000368117 | 11.91520852 | 8 mo | HOM |
| **1013609713** | 2 | CN | 607 | 1.678527066 | 3.69E-07 | 0 | 2.2E-07 | 0 | 2 mo | WILD |
| **1010418842** | 1 | CN | 607 | 1.678527066 | 0.000168863 | 11 | 0.000100602 | 6.553364688 | 8 mo | HOM |
| **1013288213** | 2 | CN | 607 | 1.678527066 | 2.66E-06 | 0 | 1.58E-06 | 0 | 4 mo | HOM |
| **1008976060** | 2 | CN | 607 | 1.678527066 | 0.001560098 | 20 | 0.000929445 | 11.91520852 | 8 mo | WILD |
| **1013446243** | 1 | CN | 607 | 1.678527066 | 8.54E-05 | 3 | 5.09E-05 | 1.787281279 | 4 mo | HOM |
| **1013288951** | 2 | CN | 607 | 1.678527066 | 0.000249182 | 4 | 0.000148453 | 2.383041705 | 4 mo | HOM |
| **1013978665** | 1 | ORBvl6a | 608 | 0.245999188 | 0 | 0 | 0 | 0 | 2 mo | HOM |
| **1013445609** | 1 | ORBvl6a | 608 | 0.245999188 | 1.76E-06 | 0 | 7.14E-06 | 0 | 4 mo | HOM |
| **1013609015** | 1 | ORBvl6a | 608 | 0.245999188 | 2.47E-05 | 1 | 0.000100489 | 4.065054062 | 4 mo | HOM |
| **1013608106** | 2 | ORBvl6a | 608 | 0.245999188 | 0 | 0 | 0 | 0 | 4 mo | HOM |
| **1011937543** | 1 | ORBvl6a | 608 | 0.245999188 | 0.000100469 | 3 | 0.000408411 | 12.19516219 | 8 mo | HOM |
| **1010419538** | 2 | ORBvl6a | 608 | 0.245999188 | 0 | 0 | 0 | 0 | 4 mo | WILD |
| **1010419187** | 1 | ORBvl6a | 608 | 0.245999188 | 0.000111772 | 5 | 0.000454358 | 20.32527031 | 8 mo | HOM |
| **1013978366** | 1 | ORBvl6a | 608 | 0.245999188 | 0 | 0 | 0 | 0 | 2 mo | HOM |
| **1014416155** | 2 | ORBvl6a | 608 | 0.245999188 | 0 | 0 | 0 | 0 | 2 mo | HOM |
| **1013978065** | 2 | ORBvl6a | 608 | 0.245999188 | 0 | 0 | 0 | 0 | 2 mo | HOM |
| **1009934102** | 2 | ORBvl6a | 608 | 0.245999188 | 0 | 0 | 0 | 0 | 4 mo | WILD |
| **1009322937** | 2 | ORBvl6a | 608 | 0.245999188 | 4.2E-06 | 1 | 1.71E-05 | 4.065054062 | 8 mo | WILD |
| **1011080637** | 2 | ORBvl6a | 608 | 0.245999188 | 8.45E-05 | 7 | 0.000343662 | 28.45537843 | 8 mo | HOM |
| **1013609713** | 2 | ORBvl6a | 608 | 0.245999188 | 0 | 0 | 0 | 0 | 2 mo | WILD |

| | | | | | | | | | | |
|---|---|---|---|---|---|---|---|---|---|---|
| **1010418842** | 1 | ORBvl6a | 608 | 0.245999188 | 0.000135606 | 5 | 0.000551247 | 20.32527031 | 8 mo | HOM |
| **1013288213** | 2 | ORBvl6a | 608 | 0.245999188 | 0 | 0 | 0 | 0 | 4 mo | HOM |
| **1008976060** | 2 | ORBvl6a | 608 | 0.245999188 | 0 | 0 | 0 | 0 | 8 mo | WILD |
| **1013446243** | 1 | ORBvl6a | 608 | 0.245999188 | 8.36E-06 | 0 | 3.4E-05 | 0 | 4 mo | HOM |
| **1013288951** | 2 | ORBvl6a | 608 | 0.245999188 | 4.19E-05 | 1 | 0.000170435 | 4.065054062 | 4 mo | HOM |
| **1013978665** | 1 | SPA | 609 | 0.136636676 | 0 | 0 | 0 | 0 | 2 mo | HOM |
| **1013445609** | 1 | SPA | 609 | 0.136636676 | 0 | 0 | 0 | 0 | 4 mo | HOM |
| **1013609015** | 1 | SPA | 609 | 0.136636676 | 1.06E-05 | 1 | 7.72E-05 | 7.318679189 | 4 mo | HOM |
| **1013608106** | 2 | SPA | 609 | 0.136636676 | 0 | 0 | 0 | 0 | 4 mo | HOM |
| **1011937543** | 1 | SPA | 609 | 0.136636676 | 4.14E-06 | 0 | 3.03E-05 | 0 | 8 mo | HOM |
| **1010419538** | 2 | SPA | 609 | 0.136636676 | 0 | 0 | 0 | 0 | 4 mo | WILD |
| **1010419187** | 1 | SPA | 609 | 0.136636676 | 3.92E-05 | 2 | 0.000286645 | 14.63735838 | 8 mo | HOM |
| **1013978366** | 1 | SPA | 609 | 0.136636676 | 0 | 0 | 0 | 0 | 2 mo | HOM |
| **1014416155** | 2 | SPA | 609 | 0.136636676 | 0 | 0 | 0 | 0 | 2 mo | HOM |
| **1013978065** | 2 | SPA | 609 | 0.136636676 | 0 | 0 | 0 | 0 | 2 mo | HOM |
| **1009934102** | 2 | SPA | 609 | 0.136636676 | 0 | 0 | 0 | 0 | 4 mo | WILD |
| **1009322937** | 2 | SPA | 609 | 0.136636676 | 0 | 0 | 0 | 0 | 8 mo | WILD |
| **1011080637** | 2 | SPA | 609 | 0.136636676 | 5.49E-08 | 0 | 4.02E-07 | 0 | 8 mo | HOM |
| **1013609713** | 2 | SPA | 609 | 0.136636676 | 0 | 0 | 0 | 0 | 2 mo | WILD |
| **1010418842** | 1 | SPA | 609 | 0.136636676 | 9.79E-06 | 0 | 7.16E-05 | 0 | 8 mo | HOM |
| **1013288213** | 2 | SPA | 609 | 0.136636676 | 0 | 0 | 0 | 0 | 4 mo | HOM |
| **1008976060** | 2 | SPA | 609 | 0.136636676 | 3.29E-06 | 0 | 2.4E-05 | 0 | 8 mo | WILD |
| **1013446243** | 1 | SPA | 609 | 0.136636676 | 0 | 0 | 0 | 0 | 4 mo | HOM |
| **1013288951** | 2 | SPA | 609 | 0.136636676 | 0 | 0 | 0 | 0 | 4 mo | HOM |
| **1013978665** | 1 | RSPd5 | 610 | 1.242441021 | 0 | 0 | 0 | 0 | 2 mo | HOM |
| **1013445609** | 1 | RSPd5 | 610 | 1.242441021 | 8.42E-05 | 5 | 6.78E-05 | 4.024335897 | 4 mo | HOM |
| **1013609015** | 1 | RSPd5 | 610 | 1.242441021 | 0.000374172 | 14 | 0.000301159 | 11.26814051 | 4 mo | HOM |
| **1013608106** | 2 | RSPd5 | 610 | 1.242441021 | 0 | 0 | 0 | 0 | 4 mo | HOM |
| **1011937543** | 1 | RSPd5 | 610 | 1.242441021 | 0.002000244 | 21 | 0.001609931 | 16.90221077 | 8 mo | HOM |
| **1010419538** | 2 | RSPd5 | 610 | 1.212294653 | 0 | 0 | 0 | 0 | 4 mo | WILD |
| **1010419187** | 1 | RSPd5 | 610 | 1.242441021 | 0.001916412 | 52 | 0.001542457 | 41.85309333 | 8 mo | HOM |
| **1013978366** | 1 | RSPd5 | 610 | 1.242441021 | 0 | 0 | 0 | 0 | 2 mo | HOM |
| **1014416155** | 2 | RSPd5 | 610 | 1.242441021 | 0 | 0 | 0 | 0 | 2 mo | HOM |
| **1013978065** | 2 | RSPd5 | 610 | 1.242441021 | 0 | 0 | 0 | 0 | 2 mo | HOM |
| **1009934102** | 2 | RSPd5 | 610 | 1.242441021 | 0 | 0 | 0 | 0 | 4 mo | WILD |
| **1009322937** | 2 | RSPd5 | 610 | 1.242441021 | 0 | 0 | 0 | 0 | 8 mo | WILD |
| **1011080637** | 2 | RSPd5 | 610 | 1.242441021 | 0.00106484 | 44 | 0.000857055 | 35.41415589 | 8 mo | HOM |
| **1013609713** | 2 | RSPd5 | 610 | 1.242441021 | 0 | 0 | 0 | 0 | 2 mo | WILD |
| **1010418842** | 1 | RSPd5 | 610 | 1.242441021 | 0.001479281 | 36 | 0.001190625 | 28.97521846 | 8 mo | HOM |
| **1013288213** | 2 | RSPd5 | 610 | 1.242441021 | 2.35E-05 | 4 | 1.89E-05 | 3.219468718 | 4 mo | HOM |
| **1008976060** | 2 | RSPd5 | 610 | 1.242441021 | 0 | 0 | 0 | 0 | 8 mo | WILD |
| **1013446243** | 1 | RSPd5 | 610 | 1.242441021 | 0.000197152 | 13 | 0.000158681 | 10.46327333 | 4 mo | HOM |
| **1013288951** | 2 | RSPd5 | 610 | 1.242441021 | 0.000114039 | 5 | 9.18E-05 | 4.024335897 | 4 mo | HOM |
| **1013978665** | 1 | hbc | 611 | 0.049234455 | 0 | 0 | 0 | 0 | 2 mo | HOM |
| **1013445609** | 1 | hbc | 611 | 0.049234455 | 2.29E-06 | 1 | 4.64E-05 | 20.31097929 | 4 mo | HOM |
| **1013609015** | 1 | hbc | 611 | 0.049234455 | 6.07E-05 | 1 | 0.001231908 | 20.31097929 | 4 mo | HOM |
| **1013608106** | 2 | hbc | 611 | 0.049234455 | 0 | 0 | 0 | 0 | 4 mo | HOM |
| **1011937543** | 1 | hbc | 611 | 0.049234455 | 0.000125562 | 1 | 0.002550285 | 20.31097929 | 8 mo | HOM |
| **1010419538** | 2 | hbc | 611 | 0.049234455 | 0 | 0 | 0 | 0 | 4 mo | WILD |
| **1010419187** | 1 | hbc | 611 | 0.049234455 | 0.000112104 | 2 | 0.002276951 | 40.62195858 | 8 mo | HOM |
| **1013978366** | 1 | hbc | 611 | 0.049234455 | 0 | 0 | 0 | 0 | 2 mo | HOM |
| **1014416155** | 2 | hbc | 611 | 0.049234455 | 0 | 0 | 0 | 0 | 2 mo | HOM |
| **1013978065** | 2 | hbc | 611 | 0.049234455 | 0 | 0 | 0 | 0 | 2 mo | HOM |
| **1009934102** | 2 | hbc | 611 | 0.049234455 | 0 | 0 | 0 | 0 | 4 mo | WILD |
| **1009322937** | 2 | hbc | 611 | 0.049234455 | 0 | 0 | 0 | 0 | 8 mo | WILD |
| **1011080637** | 2 | hbc | 611 | 0.049234455 | 0.000140936 | 1 | 0.002862542 | 20.31097929 | 8 mo | HOM |
| **1013609713** | 2 | hbc | 611 | 0.049234455 | 0 | 0 | 0 | 0 | 2 mo | WILD |
| **1010418842** | 1 | hbc | 611 | 0.049234455 | 0.00010081 | 2 | 0.002047549 | 40.62195858 | 8 mo | HOM |
| **1013288213** | 2 | hbc | 611 | 0.049234455 | 1.75E-11 | 0 | 3.56E-10 | 0 | 4 mo | HOM |

| | | | | | | | | | | |
|---|---|---|---|---|---|---|---|---|---|---|
| **1008976060** | 2 | hbc | 611 | 0.049234455 | 5.95E-08 | 0 | 1.21E-06 | 0 | 8 mo | WILD |
| **1013446243** | 1 | hbc | 611 | 0.049234455 | 0 | 0 | 0 | 0 | 4 mo | HOM |
| **1013288951** | 2 | hbc | 611 | 0.049234455 | 0.000526298 | 2 | 0.01068962 | 40.62195858 | 4 mo | HOM |
| **1013978665** | 1 | NLL | 612 | 0.745660362 | 4.81E-06 | 1 | 6.44E-06 | 1.341093146 | 2 mo | HOM |
| **1013445609** | 1 | NLL | 612 | 0.745660362 | 2.34E-06 | 1 | 3.14E-06 | 1.341093146 | 4 mo | HOM |
| **1013609015** | 1 | NLL | 612 | 0.745660362 | 0.000599605 | 2 | 0.000804126 | 2.682186292 | 4 mo | HOM |
| **1013608106** | 2 | NLL | 612 | 0.745660362 | 0 | 0 | 0 | 0 | 4 mo | HOM |
| **1011937543** | 1 | NLL | 612 | 0.745660362 | 0.001628244 | 5 | 0.002183627 | 6.705465729 | 8 mo | HOM |
| **1010419538** | 2 | NLL | 612 | 0.745660362 | 0 | 0 | 0 | 0 | 4 mo | WILD |
| **1010419187** | 1 | NLL | 612 | 0.745660362 | 0.000289822 | 1 | 0.000388678 | 1.341093146 | 8 mo | HOM |
| **1013978366** | 1 | NLL | 612 | 0.745660362 | 0 | 0 | 0 | 0 | 2 mo | HOM |
| **1014416155** | 2 | NLL | 612 | 0.745660362 | 0 | 0 | 0 | 0 | 2 mo | HOM |
| **1013978065** | 2 | NLL | 612 | 0.745660362 | 0 | 0 | 0 | 0 | 2 mo | HOM |
| **1009934102** | 2 | NLL | 612 | 0.745660362 | 0 | 0 | 0 | 0 | 4 mo | WILD |
| **1009322937** | 2 | NLL | 612 | 0.745660362 | 0 | 0 | 0 | 0 | 8 mo | WILD |
| **1011080637** | 2 | NLL | 612 | 0.745660362 | 0 | 0 | 0 | 0 | 8 mo | HOM |
| **1013609713** | 2 | NLL | 612 | 0.745660362 | 0 | 0 | 0 | 0 | 2 mo | WILD |
| **1010418842** | 1 | NLL | 612 | 0.745660362 | 0.000106829 | 5 | 0.000143268 | 6.705465729 | 8 mo | HOM |
| **1013288213** | 2 | NLL | 612 | 0.745660362 | 0 | 0 | 0 | 0 | 4 mo | HOM |
| **1008976060** | 2 | NLL | 612 | 0.745660362 | 1.39E-05 | 2 | 1.86E-05 | 2.682186292 | 8 mo | WILD |
| **1013446243** | 1 | NLL | 612 | 0.745660362 | 0 | 0 | 0 | 0 | 4 mo | HOM |
| **1013288951** | 2 | NLL | 612 | 0.745660362 | 0 | 0 | 0 | 0 | 4 mo | HOM |
| **1013978665** | 1 | VISl5 | 613 | 0.324028435 | 0 | 0 | 0 | 0 | 2 mo | HOM |
| **1013445609** | 1 | VISl5 | 613 | 0.324028435 | 3.42E-05 | 3 | 0.000105535 | 9.258446722 | 4 mo | HOM |
| **1013609015** | 1 | VISl5 | 613 | 0.324028435 | 6.95E-05 | 3 | 0.000214374 | 9.258446722 | 4 mo | HOM |
| **1013608106** | 2 | VISl5 | 613 | 0.324028435 | 0 | 0 | 0 | 0 | 4 mo | HOM |
| **1011937543** | 1 | VISl5 | 613 | 0.324028435 | 0.000481705 | 10 | 0.001486613 | 30.86148907 | 8 mo | HOM |
| **1010419538** | 2 | VISl5 | 613 | 0.324028435 | 0 | 0 | 0 | 0 | 4 mo | WILD |
| **1010419187** | 1 | VISl5 | 613 | 0.324028435 | 0.000471875 | 23 | 0.001456277 | 70.98142487 | 8 mo | HOM |
| **1013978366** | 1 | VISl5 | 613 | 0.324028435 | 0 | 0 | 0 | 0 | 2 mo | HOM |
| **1014416155** | 2 | VISl5 | 613 | 0.324028435 | 0 | 0 | 0 | 0 | 2 mo | HOM |
| **1013978065** | 2 | VISl5 | 613 | 0.324028435 | 0 | 0 | 0 | 0 | 2 mo | HOM |
| **1009934102** | 2 | VISl5 | 613 | 0.324028435 | 0 | 0 | 0 | 0 | 4 mo | WILD |
| **1009322937** | 2 | VISl5 | 613 | 0.324028435 | 0 | 0 | 0 | 0 | 8 mo | WILD |
| **1011080637** | 2 | VISl5 | 613 | 0.324028435 | 0.000220407 | 14 | 0.000680208 | 43.2060847 | 8 mo | HOM |
| **1013609713** | 2 | VISl5 | 613 | 0.324028435 | 0 | 0 | 0 | 0 | 2 mo | WILD |
| **1010418842** | 1 | VISl5 | 613 | 0.324028435 | 0.000542984 | 19 | 0.00167573 | 58.63682924 | 8 mo | HOM |
| **1013288213** | 2 | VISl5 | 613 | 0.324028435 | 0.000124949 | 5 | 0.000385611 | 15.43074454 | 4 mo | HOM |
| **1008976060** | 2 | VISl5 | 613 | 0.324028435 | 0 | 0 | 0 | 0 | 8 mo | WILD |
| **1013446243** | 1 | VISl5 | 613 | 0.324028435 | 0.000136769 | 9 | 0.00042209 | 27.77534017 | 4 mo | HOM |
| **1013288951** | 2 | VISl5 | 613 | 0.324028435 | 4.04E-05 | 4 | 0.000124714 | 12.34459563 | 4 mo | HOM |
| **1013978665** | 1 | TU | 614 | 0.539030184 | 0 | 0 | 0 | 0 | 2 mo | HOM |
| **1013445609** | 1 | TU | 614 | 0.539030184 | 0 | 0 | 0 | 0 | 4 mo | HOM |
| **1013609015** | 1 | TU | 614 | 0.539030184 | 0 | 0 | 0 | 0 | 4 mo | HOM |
| **1013608106** | 2 | TU | 614 | 0.539030184 | 0 | 0 | 0 | 0 | 4 mo | HOM |
| **1011937543** | 1 | TU | 614 | 0.539030184 | 0.001483147 | 10 | 0.00275151 | 18.55183679 | 8 mo | HOM |
| **1010419538** | 2 | TU | 614 | 0.539030184 | 1.71E-07 | 0 | 3.18E-07 | 0 | 4 mo | WILD |
| **1010419187** | 1 | TU | 614 | 0.539030184 | 6.59E-05 | 1 | 0.000122319 | 1.855183679 | 8 mo | HOM |
| **1013978366** | 1 | TU | 614 | 0.539030184 | 0 | 0 | 0 | 0 | 2 mo | HOM |
| **1014416155** | 2 | TU | 614 | 0.539030184 | 0 | 0 | 0 | 0 | 2 mo | HOM |
| **1013978065** | 2 | TU | 614 | 0.539030184 | 4.75E-06 | 0 | 8.81E-06 | 0 | 2 mo | HOM |
| **1009934102** | 2 | TU | 614 | 0.539030184 | 0 | 0 | 0 | 0 | 4 mo | WILD |
| **1009322937** | 2 | TU | 614 | 0.539030184 | 0 | 0 | 0 | 0 | 8 mo | WILD |
| **1011080637** | 2 | TU | 614 | 0.539030184 | 4.95E-05 | 3 | 9.18E-05 | 5.565551038 | 8 mo | HOM |
| **1013609713** | 2 | TU | 614 | 0.539030184 | 0 | 0 | 0 | 0 | 2 mo | WILD |
| **1010418842** | 1 | TU | 614 | 0.539030184 | 0.000156445 | 9 | 0.000290235 | 16.69665311 | 8 mo | HOM |
| **1013288213** | 2 | TU | 614 | 0.539030184 | 0 | 0 | 0 | 0 | 4 mo | HOM |
| **1008976060** | 2 | TU | 614 | 0.539030184 | 0 | 0 | 0 | 0 | 8 mo | WILD |
| **1013446243** | 1 | TU | 614 | 0.539030184 | 9.1E-09 | 0 | 1.69E-08 | 0 | 4 mo | HOM |

| | | | | | | | | | | |
|---|---|---|---|---|---|---|---|---|---|---|
| 1013288951 | 2 | TU | 614 | 0.539030184 | 0 | 0 | 0 | 0 | 4 mo | HOM |
| 1013978665 | 1 | CUN | 616 | 0.568094906 | 0 | 0 | 0 | 0 | 2 mo | HOM |
| 1013445609 | 1 | CUN | 616 | 0.568094906 | 2.17E-06 | 1 | 3.82E-06 | 1.760269259 | 4 mo | HOM |
| 1013609015 | 1 | CUN | 616 | 0.568094906 | 0.000186068 | 1 | 0.000327529 | 1.760269259 | 4 mo | HOM |
| 1013608106 | 2 | CUN | 616 | 0.568094906 | 0 | 0 | 0 | 0 | 4 mo | HOM |
| 1011937543 | 1 | CUN | 616 | 0.568094906 | 0.00128123 | 12 | 0.002255309 | 21.12323111 | 8 mo | HOM |
| 1010419538 | 2 | CUN | 616 | 0.568094906 | 0 | 0 | 0 | 0 | 4 mo | WILD |
| 1010419187 | 1 | CUN | 616 | 0.568094906 | 5.1E-05 | 4 | 8.97E-05 | 7.041077036 | 8 mo | HOM |
| 1013978366 | 1 | CUN | 616 | 0.568094906 | 2.2E-06 | 0 | 3.87E-06 | 0 | 2 mo | HOM |
| 1014416155 | 2 | CUN | 616 | 0.568094906 | 0 | 0 | 0 | 0 | 2 mo | HOM |
| 1013978065 | 2 | CUN | 616 | 0.568094906 | 0 | 0 | 0 | 0 | 2 mo | HOM |
| 1009934102 | 2 | CUN | 616 | 0.568094906 | 1.46E-07 | 0 | 2.56E-07 | 0 | 4 mo | WILD |
| 1009322937 | 2 | CUN | 616 | 0.568094906 | 1E-05 | 0 | 1.76E-05 | 0 | 8 mo | WILD |
| 1011080637 | 2 | CUN | 616 | 0.568094906 | 0.000195205 | 5 | 0.000343613 | 8.801346296 | 8 mo | HOM |
| 1013609713 | 2 | CUN | 616 | 0.568094906 | 0 | 0 | 0 | 0 | 2 mo | WILD |
| 1010418842 | 1 | CUN | 616 | 0.568094906 | 5.24E-05 | 2 | 9.22E-05 | 3.520538518 | 8 mo | HOM |
| 1013288213 | 2 | CUN | 616 | 0.568094906 | 3.62E-06 | 0 | 6.37E-06 | 0 | 4 mo | HOM |
| 1008976060 | 2 | CUN | 616 | 0.568094906 | 0.000131382 | 5 | 0.000231267 | 8.801346296 | 8 mo | WILD |
| 1013446243 | 1 | CUN | 616 | 0.568094906 | 7.17E-07 | 0 | 1.26E-06 | 0 | 4 mo | HOM |
| 1013288951 | 2 | CUN | 616 | 0.568094906 | 3.35E-08 | 0 | 5.89E-08 | 0 | 4 mo | HOM |
| 1013978665 | 1 | hc | 618 | 1.04928208 | 0 | 0 | 0 | 0 | 2 mo | HOM |
| 1013445609 | 1 | hc | 618 | 1.04928208 | 8.5E-06 | 1 | 8.1E-06 | 0.953032573 | 4 mo | HOM |
| 1013609015 | 1 | hc | 618 | 1.04928208 | 1.83E-05 | 2 | 1.75E-05 | 1.906065145 | 4 mo | HOM |
| 1013608106 | 2 | hc | 618 | 1.04928208 | 0 | 0 | 0 | 0 | 4 mo | HOM |
| 1011937543 | 1 | hc | 618 | 1.04928208 | 0.001217608 | 12 | 0.00116042 | 11.43639087 | 8 mo | HOM |
| 1010419538 | 2 | hc | 618 | 1.04928208 | 0 | 0 | 0 | 0 | 4 mo | WILD |
| 1010419187 | 1 | hc | 618 | 1.04928208 | 0.000413521 | 21 | 0.000394099 | 20.01368402 | 8 mo | HOM |
| 1013978366 | 1 | hc | 618 | 1.04928208 | 7.61E-08 | 0 | 7.25E-08 | 0 | 2 mo | HOM |
| 1014416155 | 2 | hc | 618 | 1.04928208 | 0 | 0 | 0 | 0 | 2 mo | HOM |
| 1013978065 | 2 | hc | 618 | 1.04928208 | 0 | 0 | 0 | 0 | 2 mo | HOM |
| 1009934102 | 2 | hc | 618 | 1.04928208 | 4.93E-06 | 0 | 4.69E-06 | 0 | 4 mo | WILD |
| 1009322937 | 2 | hc | 618 | 1.04928208 | 3.28E-07 | 0 | 3.13E-07 | 0 | 8 mo | WILD |
| 1011080637 | 2 | hc | 618 | 1.04928208 | 4.39E-05 | 3 | 4.19E-05 | 2.859097718 | 8 mo | HOM |
| 1013609713 | 2 | hc | 618 | 1.04928208 | 0 | 0 | 0 | 0 | 2 mo | WILD |
| 1010418842 | 1 | hc | 618 | 1.04928208 | 0.001299886 | 29 | 0.001238834 | 27.6379446 | 8 mo | HOM |
| 1013288213 | 2 | hc | 618 | 1.04928208 | 0 | 0 | 0 | 0 | 4 mo | HOM |
| 1008976060 | 2 | hc | 618 | 1.04928208 | 3.97E-06 | 1 | 3.78E-06 | 0.953032573 | 8 mo | WILD |
| 1013446243 | 1 | hc | 618 | 1.04928208 | 1.28E-05 | 1 | 1.22E-05 | 0.953032573 | 4 mo | HOM |
| 1013288951 | 2 | hc | 618 | 1.04928208 | 2.3E-05 | 1 | 2.19E-05 | 0.953032573 | 4 mo | HOM |
| 1013978665 | 1 | NLOT | 619 | 0.329135141 | 0 | 0 | 0 | 0 | 2 mo | HOM |
| 1013445609 | 1 | NLOT | 619 | 0.329136288 | 0 | 0 | 0 | 0 | 4 mo | HOM |
| 1013609015 | 1 | NLOT | 619 | 0.329136288 | 0 | 0 | 0 | 0 | 4 mo | HOM |
| 1013608106 | 2 | NLOT | 619 | 0.329136288 | 0 | 0 | 0 | 0 | 4 mo | HOM |
| 1011937543 | 1 | NLOT | 619 | 0.329136288 | 0.000113934 | 1 | 0.00034616 | 3.038255084 | 8 mo | HOM |
| 1010419538 | 2 | NLOT | 619 | 0.329136288 | 0 | 0 | 0 | 0 | 4 mo | WILD |
| 1010419187 | 1 | NLOT | 619 | 0.329136288 | 9.79E-05 | 5 | 0.000297323 | 15.19127542 | 8 mo | HOM |
| 1013978366 | 1 | NLOT | 619 | 0.329136288 | 0 | 0 | 0 | 0 | 2 mo | HOM |
| 1014416155 | 2 | NLOT | 619 | 0.329136288 | 0 | 0 | 0 | 0 | 2 mo | HOM |
| 1013978065 | 2 | NLOT | 619 | 0.329136288 | 0 | 0 | 0 | 0 | 2 mo | HOM |
| 1009934102 | 2 | NLOT | 619 | 0.329136288 | 0 | 0 | 0 | 0 | 4 mo | WILD |
| 1009322937 | 2 | NLOT | 619 | 0.329136288 | 0 | 0 | 0 | 0 | 8 mo | WILD |
| 1011080637 | 2 | NLOT | 619 | 0.329136288 | 0.000164628 | 5 | 0.000500181 | 15.19127542 | 8 mo | HOM |
| 1013609713 | 2 | NLOT | 619 | 0.329136288 | 0 | 0 | 0 | 0 | 2 mo | WILD |
| 1010418842 | 1 | NLOT | 619 | 0.329136288 | 0.000262002 | 9 | 0.000796029 | 27.34429575 | 8 mo | HOM |
| 1013288213 | 2 | NLOT | 619 | 0.329136288 | 5.2E-07 | 0 | 1.58E-06 | 0 | 4 mo | HOM |
| 1008976060 | 2 | NLOT | 619 | 0.329136288 | 0 | 0 | 0 | 0 | 8 mo | WILD |
| 1013446243 | 1 | NLOT | 619 | 0.329136288 | 1.41E-05 | 1 | 4.29E-05 | 3.038255084 | 4 mo | HOM |
| 1013288951 | 2 | NLOT | 619 | 0.329136288 | 0 | 0 | 0 | 0 | 4 mo | HOM |
| 1013978665 | 1 | ORBm5 | 620 | 0.485380202 | 0 | 0 | 0 | 0 | 2 mo | HOM |

| | | | | | | | | | | |
|---|---|---|---|---|---|---|---|---|---|---|
| **1013445609** | 1 | ORBm5 | 620 | 0.485380202 | 0 | 0 | 0 | 0 | 4 mo | HOM |
| **1013609015** | 1 | ORBm5 | 620 | 0.485380202 | 7.78E-05 | 3 | 0.000160353 | 6.180721814 | 4 mo | HOM |
| **1013608106** | 2 | ORBm5 | 620 | 0.485380202 | 0 | 0 | 0 | 0 | 4 mo | HOM |
| **1011937543** | 1 | ORBm5 | 620 | 0.485380202 | 0.000653726 | 15 | 0.001346833 | 30.90360907 | 8 mo | HOM |
| **1010419538** | 2 | ORBm5 | 620 | 0.485380202 | 0 | 0 | 0 | 0 | 4 mo | WILD |
| **1010419187** | 1 | ORBm5 | 620 | 0.485380202 | 0.000346634 | 12 | 0.00071415 | 24.72288726 | 8 mo | HOM |
| **1013978366** | 1 | ORBm5 | 620 | 0.485380202 | 0 | 0 | 0 | 0 | 2 mo | HOM |
| **1014416155** | 2 | ORBm5 | 620 | 0.485380202 | 0 | 0 | 0 | 0 | 2 mo | HOM |
| **1013978065** | 2 | ORBm5 | 620 | 0.485380202 | 0 | 0 | 0 | 0 | 2 mo | HOM |
| **1009934102** | 2 | ORBm5 | 620 | 0.485380202 | 4.51E-06 | 1 | 9.28E-06 | 2.060240605 | 4 mo | WILD |
| **1009322937** | 2 | ORBm5 | 620 | 0.485380202 | 0 | 0 | 0 | 0 | 8 mo | WILD |
| **1011080637** | 2 | ORBm5 | 620 | 0.485380202 | 0.000404943 | 15 | 0.000834281 | 30.90360907 | 8 mo | HOM |
| **1013609713** | 2 | ORBm5 | 620 | 0.485380202 | 0 | 0 | 0 | 0 | 2 mo | WILD |
| **1010418842** | 1 | ORBm5 | 620 | 0.485380202 | 0.00052049 | 13 | 0.001072335 | 26.78312786 | 8 mo | HOM |
| **1013288213** | 2 | ORBm5 | 620 | 0.485380202 | 2.54E-05 | 1 | 5.24E-05 | 2.060240605 | 4 mo | HOM |
| **1008976060** | 2 | ORBm5 | 620 | 0.485380202 | 0 | 0 | 0 | 0 | 8 mo | WILD |
| **1013446243** | 1 | ORBm5 | 620 | 0.485380202 | 2.86E-05 | 2 | 5.89E-05 | 4.120481209 | 4 mo | HOM |
| **1013288951** | 2 | ORBm5 | 620 | 0.485380202 | 4.66E-05 | 4 | 9.6E-05 | 8.240962418 | 4 mo | HOM |
| **1013978665** | 1 | V | 621 | 0.365480573 | 0 | 0 | 0 | 0 | 2 mo | HOM |
| **1013445609** | 1 | V | 621 | 0.365480573 | 0 | 0 | 0 | 0 | 4 mo | HOM |
| **1013609015** | 1 | V | 621 | 0.365480573 | 0 | 0 | 0 | 0 | 4 mo | HOM |
| **1013608106** | 2 | V | 621 | 0.365480573 | 0 | 0 | 0 | 0 | 4 mo | HOM |
| **1011937543** | 1 | V | 621 | 0.365480573 | 0.00055553 | 2 | 0.001519999 | 5.472247087 | 8 mo | HOM |
| **1010419538** | 2 | V | 621 | 0.365480573 | 0 | 0 | 0 | 0 | 4 mo | WILD |
| **1010419187** | 1 | V | 621 | 0.365480573 | 0 | 0 | 0 | 0 | 8 mo | HOM |
| **1013978366** | 1 | V | 621 | 0.365480573 | 0 | 0 | 0 | 0 | 2 mo | HOM |
| **1014416155** | 2 | V | 621 | 0.365480573 | 0 | 0 | 0 | 0 | 2 mo | HOM |
| **1013978065** | 2 | V | 621 | 0.365480573 | 0 | 0 | 0 | 0 | 2 mo | HOM |
| **1009934102** | 2 | V | 621 | 0.365480573 | 0 | 0 | 0 | 0 | 4 mo | WILD |
| **1009322937** | 2 | V | 621 | 0.365480573 | 0 | 0 | 0 | 0 | 8 mo | WILD |
| **1011080637** | 2 | V | 621 | 0.365480573 | 8.51E-06 | 1 | 2.33E-05 | 2.736123544 | 8 mo | HOM |
| **1013609713** | 2 | V | 621 | 0.365480573 | 0 | 0 | 0 | 0 | 2 mo | WILD |
| **1010418842** | 1 | V | 621 | 0.365480573 | 2.1E-05 | 2 | 5.73E-05 | 5.472247087 | 8 mo | HOM |
| **1013288213** | 2 | V | 621 | 0.365480573 | 0 | 0 | 0 | 0 | 4 mo | HOM |
| **1008976060** | 2 | V | 621 | 0.365480573 | 0 | 0 | 0 | 0 | 8 mo | WILD |
| **1013446243** | 1 | V | 621 | 0.365480573 | 0 | 0 | 0 | 0 | 4 mo | HOM |
| **1013288951** | 2 | V | 621 | 0.365480573 | 0 | 0 | 0 | 0 | 4 mo | HOM |
| **1013978665** | 1 | RSPv6b | 622 | 0.049759867 | 0 | 0 | 0 | 0 | 2 mo | HOM |
| **1013445609** | 1 | RSPv6b | 622 | 0.049759867 | 0 | 0 | 0 | 0 | 4 mo | HOM |
| **1013609015** | 1 | RSPv6b | 622 | 0.049759867 | 0 | 0 | 0 | 0 | 4 mo | HOM |
| **1013608106** | 2 | RSPv6b | 622 | 0.049759867 | 0 | 0 | 0 | 0 | 4 mo | HOM |
| **1011937543** | 1 | RSPv6b | 622 | 0.049759867 | 3.25E-05 | 0 | 0.000652374 | 0 | 8 mo | HOM |
| **1010419538** | 2 | RSPv6b | 622 | 0.049759867 | 0 | 0 | 0 | 0 | 4 mo | WILD |
| **1010419187** | 1 | RSPv6b | 622 | 0.049759867 | 1.82E-05 | 1 | 0.000365898 | 20.09651654 | 8 mo | HOM |
| **1013978366** | 1 | RSPv6b | 622 | 0.049759867 | 0 | 0 | 0 | 0 | 2 mo | HOM |
| **1014416155** | 2 | RSPv6b | 622 | 0.049759867 | 0 | 0 | 0 | 0 | 2 mo | HOM |
| **1013978065** | 2 | RSPv6b | 622 | 0.049759867 | 0 | 0 | 0 | 0 | 2 mo | HOM |
| **1009934102** | 2 | RSPv6b | 622 | 0.049759867 | 0 | 0 | 0 | 0 | 4 mo | WILD |
| **1009322937** | 2 | RSPv6b | 622 | 0.049759867 | 0 | 0 | 0 | 0 | 8 mo | WILD |
| **1011080637** | 2 | RSPv6b | 622 | 0.049759867 | 9.1E-05 | 2 | 0.001828514 | 40.19303307 | 8 mo | HOM |
| **1013609713** | 2 | RSPv6b | 622 | 0.049759867 | 0 | 0 | 0 | 0 | 2 mo | WILD |
| **1010418842** | 1 | RSPv6b | 622 | 0.049759867 | 6.11E-05 | 4 | 0.001227391 | 80.38606614 | 8 mo | HOM |
| **1013288213** | 2 | RSPv6b | 622 | 0.049759867 | 0 | 0 | 0 | 0 | 4 mo | HOM |
| **1008976060** | 2 | RSPv6b | 622 | 0.049759867 | 0 | 0 | 0 | 0 | 8 mo | WILD |
| **1013446243** | 1 | RSPv6b | 622 | 0.049759867 | 0 | 0 | 0 | 0 | 4 mo | HOM |
| **1013288951** | 2 | RSPv6b | 622 | 0.049759867 | 1.6E-09 | 0 | 3.21E-08 | 0 | 4 mo | HOM |
| **1013978665** | 1 | CNU | 623 | 55.92184878 | 4.94E-05 | 2 | 8.84E-07 | 0.035764197 | 2 mo | HOM |
| **1013445609** | 1 | CNU | 623 | 55.96863084 | 0.000552829 | 45 | 9.88E-06 | 0.804021812 | 4 mo | HOM |
| **1013609015** | 1 | CNU | 623 | 55.96863084 | 0.00600812 | 140 | 0.000107348 | 2.501401194 | 4 mo | HOM |

| | | | | | | | | | | |
|---|---|---|---|---|---|---|---|---|---|---|
| **1013608106** | 2 | CNU | 623 | 55.96863084 | 0 | 0 | 0 | 0 | 4 mo | HOM |
| **1011937543** | 1 | CNU | 623 | 55.96863084 | 0.036689927 | 783 | 0.000655544 | 13.98997953 | 8 mo | HOM |
| **1010419538** | 2 | CNU | 623 | 55.96863084 | 6.39E-05 | 2 | 1.14E-06 | 0.035734303 | 4 mo | WILD |
| **1010419187** | 1 | CNU | 623 | 55.96863084 | 0.02714464 | 1088 | 0.000484997 | 19.43946071 | 8 mo | HOM |
| **1013978366** | 1 | CNU | 623 | 55.96863084 | 5.84E-06 | 1 | 1.04E-07 | 0.017867151 | 2 mo | HOM |
| **1014416155** | 2 | CNU | 623 | 55.96863084 | 0 | 0 | 0 | 0 | 2 mo | HOM |
| **1013978065** | 2 | CNU | 623 | 55.96863084 | 1.07E-05 | 2 | 1.92E-07 | 0.035734303 | 2 mo | HOM |
| **1009934102** | 2 | CNU | 623 | 55.96863084 | 6.95E-05 | 6 | 1.24E-06 | 0.107202908 | 4 mo | WILD |
| **1009322937** | 2 | CNU | 623 | 55.96863084 | 0.000236828 | 11 | 4.23E-06 | 0.196538665 | 8 mo | WILD |
| **1011080637** | 2 | CNU | 623 | 55.96863084 | 0.019503334 | 877 | 0.000348469 | 15.66949176 | 8 mo | HOM |
| **1013609713** | 2 | CNU | 623 | 55.96863084 | 1.86E-06 | 0 | 3.32E-08 | 0 | 2 mo | WILD |
| **1010418842** | 1 | CNU | 623 | 55.96863084 | 0.042464897 | 1366 | 0.000758727 | 24.40652879 | 8 mo | HOM |
| **1013288213** | 2 | CNU | 623 | 55.96863084 | 0.001815253 | 51 | 3.24E-05 | 0.911224721 | 4 mo | HOM |
| **1008976060** | 2 | CNU | 623 | 55.96863084 | 0.00206545 | 120 | 3.69E-05 | 2.144058166 | 8 mo | WILD |
| **1013446243** | 1 | CNU | 623 | 55.96863084 | 0.002141885 | 112 | 3.83E-05 | 2.001120955 | 4 mo | HOM |
| **1013288951** | 2 | CNU | 623 | 55.96863084 | 0.003359146 | 118 | 6E-05 | 2.108323863 | 4 mo | HOM |
| **1013978665** | 1 | SSp-ul5 | 625 | 0.816245214 | 0 | 0 | 0 | 0 | 2 mo | HOM |
| **1013445609** | 1 | SSp-ul5 | 625 | 0.816245214 | 8.54E-05 | 5 | 0.000104679 | 6.125610186 | 4 mo | HOM |
| **1013609015** | 1 | SSp-ul5 | 625 | 0.816245214 | 0.000161188 | 7 | 0.000197475 | 8.575854261 | 4 mo | HOM |
| **1013608106** | 2 | SSp-ul5 | 625 | 0.816245214 | 0 | 0 | 0 | 0 | 4 mo | HOM |
| **1011937543** | 1 | SSp-ul5 | 625 | 0.816245214 | 0.001271329 | 26 | 0.001557533 | 31.85317297 | 8 mo | HOM |
| **1010419538** | 2 | SSp-ul5 | 625 | 0.816245214 | 0 | 0 | 0 | 0 | 4 mo | WILD |
| **1010419187** | 1 | SSp-ul5 | 625 | 0.816245214 | 0.001642647 | 47 | 0.002012443 | 57.58073575 | 8 mo | HOM |
| **1013978366** | 1 | SSp-ul5 | 625 | 0.816245214 | 0 | 0 | 0 | 0 | 2 mo | HOM |
| **1014416155** | 2 | SSp-ul5 | 625 | 0.816245214 | 0 | 0 | 0 | 0 | 2 mo | HOM |
| **1013978065** | 2 | SSp-ul5 | 625 | 0.816245214 | 0 | 0 | 0 | 0 | 2 mo | HOM |
| **1009934102** | 2 | SSp-ul5 | 625 | 0.816245214 | 0 | 0 | 0 | 0 | 4 mo | WILD |
| **1009322937** | 2 | SSp-ul5 | 625 | 0.816245214 | 0 | 0 | 0 | 0 | 8 mo | WILD |
| **1011080637** | 2 | SSp-ul5 | 625 | 0.816245214 | 0.000643151 | 25 | 0.000787939 | 30.62805093 | 8 mo | HOM |
| **1013609713** | 2 | SSp-ul5 | 625 | 0.816245214 | 0 | 0 | 0 | 0 | 2 mo | WILD |
| **1010418842** | 1 | SSp-ul5 | 625 | 0.816245214 | 0.00077909 | 29 | 0.00095448 | 35.52853908 | 8 mo | HOM |
| **1013288213** | 2 | SSp-ul5 | 625 | 0.816245214 | 7.02E-05 | 5 | 8.6E-05 | 6.125610186 | 4 mo | HOM |
| **1008976060** | 2 | SSp-ul5 | 625 | 0.816245214 | 8.62E-08 | 0 | 1.06E-07 | 0 | 8 mo | WILD |
| **1013446243** | 1 | SSp-ul5 | 625 | 0.816245214 | 0.000186982 | 14 | 0.000229076 | 17.15170852 | 4 mo | HOM |
| **1013288951** | 2 | SSp-ul5 | 625 | 0.816245214 | 0.000196866 | 10 | 0.000241185 | 12.25122037 | 4 mo | HOM |
| **1013978665** | 1 | NOT | 628 | 0.211690623 | 0 | 0 | 0 | 0 | 2 mo | HOM |
| **1013445609** | 1 | NOT | 628 | 0.211690623 | 0 | 0 | 0 | 0 | 4 mo | HOM |
| **1013609015** | 1 | NOT | 628 | 0.211690623 | 0 | 0 | 0 | 0 | 4 mo | HOM |
| **1013608106** | 2 | NOT | 628 | 0.211690623 | 0 | 0 | 0 | 0 | 4 mo | HOM |
| **1011937543** | 1 | NOT | 628 | 0.211690623 | 0.000186077 | 7 | 0.000879006 | 33.06712359 | 8 mo | HOM |
| **1010419538** | 2 | NOT | 628 | 0.211690623 | 7.98E-08 | 0 | 3.77E-07 | 0 | 4 mo | WILD |
| **1010419187** | 1 | NOT | 628 | 0.211690623 | 1.84E-05 | 1 | 8.71E-05 | 4.723874798 | 8 mo | HOM |
| **1013978366** | 1 | NOT | 628 | 0.211690623 | 2.38E-07 | 0 | 1.13E-06 | 0 | 2 mo | HOM |
| **1014416155** | 2 | NOT | 628 | 0.211690623 | 0 | 0 | 0 | 0 | 2 mo | HOM |
| **1013978065** | 2 | NOT | 628 | 0.211690623 | 0 | 0 | 0 | 0 | 2 mo | HOM |
| **1009934102** | 2 | NOT | 628 | 0.211690623 | 0 | 0 | 0 | 0 | 4 mo | WILD |
| **1009322937** | 2 | NOT | 628 | 0.211690623 | 1.24E-05 | 1 | 5.88E-05 | 4.723874798 | 8 mo | WILD |
| **1011080637** | 2 | NOT | 628 | 0.211690623 | 9.75E-06 | 0 | 4.6E-05 | 0 | 8 mo | HOM |
| **1013609713** | 2 | NOT | 628 | 0.211690623 | 0 | 0 | 0 | 0 | 2 mo | WILD |
| **1010418842** | 1 | NOT | 628 | 0.211690623 | 9.02E-06 | 0 | 4.26E-05 | 0 | 8 mo | HOM |
| **1013288213** | 2 | NOT | 628 | 0.211690623 | 4.1E-05 | 1 | 0.000193739 | 4.723874798 | 4 mo | HOM |
| **1008976060** | 2 | NOT | 628 | 0.211690623 | 5.77E-06 | 0 | 2.73E-05 | 0 | 8 mo | WILD |
| **1013446243** | 1 | NOT | 628 | 0.211690623 | 8.19E-07 | 0 | 3.87E-06 | 0 | 4 mo | HOM |
| **1013288951** | 2 | NOT | 628 | 0.211690623 | 1.13E-07 | 0 | 5.32E-07 | 0 | 4 mo | HOM |
| **1013978665** | 1 | VAL | 629 | 0.84173917 | 0 | 0 | 0 | 0 | 2 mo | HOM |
| **1013445609** | 1 | VAL | 629 | 0.84173917 | 0 | 0 | 0 | 0 | 4 mo | HOM |
| **1013609015** | 1 | VAL | 629 | 0.84173917 | 0 | 0 | 0 | 0 | 4 mo | HOM |
| **1013608106** | 2 | VAL | 629 | 0.84173917 | 0 | 0 | 0 | 0 | 4 mo | HOM |
| **1011937543** | 1 | VAL | 629 | 0.84173917 | 0.000191149 | 3 | 0.000227088 | 3.564049421 | 8 mo | HOM |

| | | | | | | | | | | |
|---|---|---|---|---|---|---|---|---|---|---|
| **1010419538** | 2 | VAL | 629 | 0.84173917 | 0 | 0 | 0 | 0 | 4 mo | WILD |
| **1010419187** | 1 | VAL | 629 | 0.84173917 | 0.000255299 | 8 | 0.000303299 | 9.504131789 | 8 mo | HOM |
| **1013978366** | 1 | VAL | 629 | 0.84173917 | 0 | 0 | 0 | 0 | 2 mo | HOM |
| **1014416155** | 2 | VAL | 629 | 0.84173917 | 0 | 0 | 0 | 0 | 2 mo | HOM |
| **1013978065** | 2 | VAL | 629 | 0.84173917 | 0 | 0 | 0 | 0 | 2 mo | HOM |
| **1009934102** | 2 | VAL | 629 | 0.84173917 | 0 | 0 | 0 | 0 | 4 mo | WILD |
| **1009322937** | 2 | VAL | 629 | 0.84173917 | 0 | 0 | 0 | 0 | 8 mo | WILD |
| **1011080637** | 2 | VAL | 629 | 0.84173917 | 0.000313527 | 12 | 0.000372475 | 14.25619768 | 8 mo | HOM |
| **1013609713** | 2 | VAL | 629 | 0.84173917 | 0 | 0 | 0 | 0 | 2 mo | WILD |
| **1010418842** | 1 | VAL | 629 | 0.84173917 | 0.000401391 | 14 | 0.000476859 | 16.63223063 | 8 mo | HOM |
| **1013288213** | 2 | VAL | 629 | 0.84173917 | 1.53E-10 | 0 | 1.81E-10 | 0 | 4 mo | HOM |
| **1008976060** | 2 | VAL | 629 | 0.84173917 | 2.16E-05 | 2 | 2.56E-05 | 2.376032947 | 8 mo | WILD |
| **1013446243** | 1 | VAL | 629 | 0.84173917 | 6.16E-06 | 1 | 7.32E-06 | 1.188016474 | 4 mo | HOM |
| **1013288951** | 2 | VAL | 629 | 0.84173917 | 4.9E-07 | 0 | 5.83E-07 | 0 | 4 mo | HOM |
| **1013978665** | 1 | ORBI5 | 630 | 1.243382683 | 0 | 0 | 0 | 0 | 2 mo | HOM |
| **1013445609** | 1 | ORBI5 | 630 | 1.243382683 | 7.44E-05 | 7 | 5.98E-05 | 5.629803354 | 4 mo | HOM |
| **1013609015** | 1 | ORBI5 | 630 | 1.243382683 | 0.000221997 | 6 | 0.000178542 | 4.825545732 | 4 mo | HOM |
| **1013608106** | 2 | ORBI5 | 630 | 1.243382683 | 0 | 0 | 0 | 0 | 4 mo | HOM |
| **1011937543** | 1 | ORBI5 | 630 | 1.243382683 | 0.001155353 | 28 | 0.000929201 | 22.51921342 | 8 mo | HOM |
| **1010419538** | 2 | ORBI5 | 630 | 1.243382683 | 0 | 0 | 0 | 0 | 4 mo | WILD |
| **1010419187** | 1 | ORBI5 | 630 | 1.243382683 | 0.001595086 | 63 | 0.00128286 | 50.66823018 | 8 mo | HOM |
| **1013978366** | 1 | ORBI5 | 630 | 1.243382683 | 0 | 0 | 0 | 0 | 2 mo | HOM |
| **1014416155** | 2 | ORBI5 | 630 | 1.243382683 | 0 | 0 | 0 | 0 | 2 mo | HOM |
| **1013978065** | 2 | ORBI5 | 630 | 1.243382683 | 0 | 0 | 0 | 0 | 2 mo | HOM |
| **1009934102** | 2 | ORBI5 | 630 | 1.243382683 | 0 | 0 | 0 | 0 | 4 mo | WILD |
| **1009322937** | 2 | ORBI5 | 630 | 1.243382683 | 0 | 0 | 0 | 0 | 8 mo | WILD |
| **1011080637** | 2 | ORBI5 | 630 | 1.243382683 | 0.000875954 | 44 | 0.000704492 | 35.38733537 | 8 mo | HOM |
| **1013609713** | 2 | ORBI5 | 630 | 1.243382683 | 0 | 0 | 0 | 0 | 2 mo | WILD |
| **1010418842** | 1 | ORBI5 | 630 | 1.243382683 | 0.001243577 | 38 | 0.001000157 | 30.56178964 | 8 mo | HOM |
| **1013288213** | 2 | ORBI5 | 630 | 1.243382683 | 1.3E-05 | 2 | 1.05E-05 | 1.608515244 | 4 mo | HOM |
| **1008976060** | 2 | ORBI5 | 630 | 1.243382683 | 0 | 0 | 0 | 0 | 8 mo | WILD |
| **1013446243** | 1 | ORBI5 | 630 | 1.243382683 | 0.000113499 | 6 | 9.13E-05 | 4.825545732 | 4 mo | HOM |
| **1013288951** | 2 | ORBI5 | 630 | 1.243382683 | 0.000142647 | 8 | 0.000114725 | 6.434060976 | 4 mo | HOM |
| **1013978665** | 1 | COA | 631 | 3.341443515 | 1.88E-05 | 2 | 5.62E-06 | 0.59854371 | 2 mo | HOM |
| **1013445609** | 1 | COA | 631 | 3.363841219 | 0.000224904 | 14 | 6.69E-05 | 4.161908689 | 4 mo | HOM |
| **1013609015** | 1 | COA | 631 | 3.363841219 | 0.000651461 | 23 | 0.000193666 | 6.837421418 | 4 mo | HOM |
| **1013608106** | 2 | COA | 631 | 3.363841219 | 0 | 0 | 0 | 0 | 4 mo | HOM |
| **1011937543** | 1 | COA | 631 | 3.363841219 | 0.005839055 | 85 | 0.00173583 | 25.26873133 | 8 mo | HOM |
| **1010419538** | 2 | COA | 631 | 3.363841219 | 3.09E-05 | 5 | 9.2E-06 | 1.486395961 | 4 mo | WILD |
| **1010419187** | 1 | COA | 631 | 3.363841219 | 0.007024753 | 124 | 0.002088313 | 36.86261982 | 8 mo | HOM |
| **1013978366** | 1 | COA | 631 | 3.363841219 | 0 | 0 | 0 | 0 | 2 mo | HOM |
| **1014416155** | 2 | COA | 631 | 3.363841219 | 0 | 0 | 0 | 0 | 2 mo | HOM |
| **1013978065** | 2 | COA | 631 | 3.363841219 | 1.52E-06 | 0 | 4.52E-07 | 0 | 2 mo | HOM |
| **1009934102** | 2 | COA | 631 | 3.363841219 | 0 | 0 | 0 | 0 | 4 mo | WILD |
| **1009322937** | 2 | COA | 631 | 3.363841219 | 0 | 0 | 0 | 0 | 8 mo | WILD |
| **1011080637** | 2 | COA | 631 | 3.363841219 | 0.004357725 | 147 | 0.001295461 | 43.70004124 | 8 mo | HOM |
| **1013609713** | 2 | COA | 631 | 3.363841219 | 0 | 0 | 0 | 0 | 2 mo | WILD |
| **1010418842** | 1 | COA | 631 | 3.363841219 | 0.006085277 | 155 | 0.001809026 | 46.07827478 | 8 mo | HOM |
| **1013288213** | 2 | COA | 631 | 3.363841219 | 0.00038118 | 23 | 0.000113317 | 6.837421418 | 4 mo | HOM |
| **1008976060** | 2 | COA | 631 | 3.363841219 | 5.17E-05 | 10 | 1.54E-05 | 2.972791921 | 8 mo | WILD |
| **1013446243** | 1 | COA | 631 | 3.363841219 | 0.000513792 | 23 | 0.00015274 | 6.837421418 | 4 mo | HOM |
| **1013288951** | 2 | COA | 631 | 3.363841219 | 0.000945313 | 18 | 0.000281022 | 5.351025458 | 4 mo | HOM |
| **1013978665** | 1 | DG-sg | 632 | 1.653037187 | 0 | 0 | 0 | 0 | 2 mo | HOM |
| **1013445609** | 1 | DG-sg | 632 | 1.653037187 | 0 | 0 | 0 | 0 | 4 mo | HOM |
| **1013609015** | 1 | DG-sg | 632 | 1.653037187 | 7.16E-06 | 1 | 4.33E-06 | 0.604947068 | 4 mo | HOM |
| **1013608106** | 2 | DG-sg | 632 | 1.653037187 | 0 | 0 | 0 | 0 | 4 mo | HOM |
| **1011937543** | 1 | DG-sg | 632 | 1.653037187 | 0.001958644 | 19 | 0.001184876 | 11.4939943 | 8 mo | HOM |
| **1010419538** | 2 | DG-sg | 632 | 1.653037187 | 1.11E-05 | 1 | 6.69E-06 | 0.604947068 | 4 mo | WILD |
| **1010419187** | 1 | DG-sg | 632 | 1.653037187 | 0.000426803 | 21 | 0.000258193 | 12.70388843 | 8 mo | HOM |

| | | | | | | | | | | |
|---|---|---|---|---|---|---|---|---|---|---|
| **1013978366** | 1 | DG-sg | 632 | 1.653037187 | 0 | 0 | 0 | 0 | 2 mo | HOM |
| **1014416155** | 2 | DG-sg | 632 | 1.653037187 | 0 | 0 | 0 | 0 | 2 mo | HOM |
| **1013978065** | 2 | DG-sg | 632 | 1.653037187 | 0 | 0 | 0 | 0 | 2 mo | HOM |
| **1009934102** | 2 | DG-sg | 632 | 1.653037187 | 7.5E-06 | 0 | 4.53E-06 | 0 | 4 mo | WILD |
| **1009322937** | 2 | DG-sg | 632 | 1.653037187 | 0 | 0 | 0 | 0 | 8 mo | WILD |
| **1011080637** | 2 | DG-sg | 632 | 1.653037187 | 0.000210285 | 9 | 0.000127211 | 5.444523614 | 8 mo | HOM |
| **1013609713** | 2 | DG-sg | 632 | 1.653037187 | 0 | 0 | 0 | 0 | 2 mo | WILD |
| **1010418842** | 1 | DG-sg | 632 | 1.653037187 | 0.000477428 | 17 | 0.000288819 | 10.28410016 | 8 mo | HOM |
| **1013288213** | 2 | DG-sg | 632 | 1.653037187 | 2.26E-07 | 0 | 1.37E-07 | 0 | 4 mo | HOM |
| **1008976060** | 2 | DG-sg | 632 | 1.653037187 | 1.72E-05 | 2 | 1.04E-05 | 1.209894136 | 8 mo | WILD |
| **1013446243** | 1 | DG-sg | 632 | 1.653037187 | 1.04E-05 | 0 | 6.28E-06 | 0 | 4 mo | HOM |
| **1013288951** | 2 | DG-sg | 632 | 1.653037187 | 3.73E-05 | 2 | 2.25E-05 | 1.209894136 | 4 mo | HOM |
| **1013978665** | 1 | cic | 633 | 0.018702705 | 0 | 0 | 0 | 0 | 2 mo | HOM |
| **1013445609** | 1 | cic | 633 | 0.018702705 | 0 | 0 | 0 | 0 | 4 mo | HOM |
| **1013609015** | 1 | cic | 633 | 0.018702705 | 0 | 0 | 0 | 0 | 4 mo | HOM |
| **1013608106** | 2 | cic | 633 | 0.018702705 | 0 | 0 | 0 | 0 | 4 mo | HOM |
| **1011937543** | 1 | cic | 633 | 0.018702705 | 5.21E-05 | 0 | 0.002787949 | 0 | 8 mo | HOM |
| **1010419538** | 2 | cic | 633 | 0.018702705 | 0 | 0 | 0 | 0 | 4 mo | WILD |
| **1010419187** | 1 | cic | 633 | 0.018702705 | 0 | 0 | 0 | 0 | 8 mo | HOM |
| **1013978366** | 1 | cic | 633 | 0.018702705 | 3.09E-08 | 0 | 1.65E-06 | 0 | 2 mo | HOM |
| **1014416155** | 2 | cic | 633 | 0.018702705 | 0 | 0 | 0 | 0 | 2 mo | HOM |
| **1013978065** | 2 | cic | 633 | 0.018702705 | 0 | 0 | 0 | 0 | 2 mo | HOM |
| **1009934102** | 2 | cic | 633 | 0.018702705 | 0 | 0 | 0 | 0 | 4 mo | WILD |
| **1009322937** | 2 | cic | 633 | 0.018702705 | 0 | 0 | 0 | 0 | 8 mo | WILD |
| **1011080637** | 2 | cic | 633 | 0.018702705 | 0 | 0 | 0 | 0 | 8 mo | HOM |
| **1013609713** | 2 | cic | 633 | 0.018702705 | 0 | 0 | 0 | 0 | 2 mo | WILD |
| **1010418842** | 1 | cic | 633 | 0.018702705 | 1.09E-07 | 0 | 5.85E-06 | 0 | 8 mo | HOM |
| **1013288213** | 2 | cic | 633 | 0.018702705 | 0 | 0 | 0 | 0 | 4 mo | HOM |
| **1008976060** | 2 | cic | 633 | 0.018702705 | 0 | 0 | 0 | 0 | 8 mo | WILD |
| **1013446243** | 1 | cic | 633 | 0.018702705 | 0 | 0 | 0 | 0 | 4 mo | HOM |
| **1013288951** | 2 | cic | 633 | 0.018702705 | 0 | 0 | 0 | 0 | 4 mo | HOM |
| **1013978665** | 1 | NPC | 634 | 0.298464429 | 0 | 0 | 0 | 0 | 2 mo | HOM |
| **1013445609** | 1 | NPC | 634 | 0.298464429 | 0 | 0 | 0 | 0 | 4 mo | HOM |
| **1013609015** | 1 | NPC | 634 | 0.298464429 | 0 | 0 | 0 | 0 | 4 mo | HOM |
| **1013608106** | 2 | NPC | 634 | 0.298464429 | 0 | 0 | 0 | 0 | 4 mo | HOM |
| **1011937543** | 1 | NPC | 634 | 0.298464429 | 0.000188799 | 9 | 0.000632567 | 30.15434712 | 8 mo | HOM |
| **1010419538** | 2 | NPC | 634 | 0.298464429 | 0 | 0 | 0 | 0 | 4 mo | WILD |
| **1010419187** | 1 | NPC | 634 | 0.298464429 | 0 | 0 | 0 | 0 | 8 mo | HOM |
| **1013978366** | 1 | NPC | 634 | 0.298464429 | 0 | 0 | 0 | 0 | 2 mo | HOM |
| **1014416155** | 2 | NPC | 634 | 0.298464429 | 0 | 0 | 0 | 0 | 2 mo | HOM |
| **1013978065** | 2 | NPC | 634 | 0.298464429 | 0 | 0 | 0 | 0 | 2 mo | HOM |
| **1009934102** | 2 | NPC | 634 | 0.298464429 | 0 | 0 | 0 | 0 | 4 mo | WILD |
| **1009322937** | 2 | NPC | 634 | 0.298464429 | 0 | 0 | 0 | 0 | 8 mo | WILD |
| **1011080637** | 2 | NPC | 634 | 0.298464429 | 2.18E-05 | 1 | 7.31E-05 | 3.350483013 | 8 mo | HOM |
| **1013609713** | 2 | NPC | 634 | 0.298464429 | 0 | 0 | 0 | 0 | 2 mo | WILD |
| **1010418842** | 1 | NPC | 634 | 0.298464429 | 6.35E-06 | 1 | 2.13E-05 | 3.350483013 | 8 mo | HOM |
| **1013288213** | 2 | NPC | 634 | 0.298464429 | 1.16E-05 | 0 | 3.89E-05 | 0 | 4 mo | HOM |
| **1008976060** | 2 | NPC | 634 | 0.298464429 | 1.76E-05 | 0 | 5.9E-05 | 0 | 8 mo | WILD |
| **1013446243** | 1 | NPC | 634 | 0.298464429 | 0 | 0 | 0 | 0 | 4 mo | HOM |
| **1013288951** | 2 | NPC | 634 | 0.298464429 | 2.81E-09 | 0 | 9.41E-09 | 0 | 4 mo | HOM |
| **1013978665** | 1 | VENT | 637 | 5.006679262 | 0 | 0 | 0 | 0 | 2 mo | HOM |
| **1013445609** | 1 | VENT | 637 | 5.006679262 | 0 | 0 | 0 | 0 | 4 mo | HOM |
| **1013609015** | 1 | VENT | 637 | 5.006679262 | 5.19E-05 | 2 | 1.04E-05 | 0.399466372 | 4 mo | HOM |
| **1013608106** | 2 | VENT | 637 | 5.006679262 | 0 | 0 | 0 | 0 | 4 mo | HOM |
| **1011937543** | 1 | VENT | 637 | 5.006679262 | 0.002258549 | 34 | 0.000451107 | 6.790928322 | 8 mo | HOM |
| **1010419538** | 2 | VENT | 637 | 5.006679262 | 0 | 0 | 0 | 0 | 4 mo | WILD |
| **1010419187** | 1 | VENT | 637 | 5.006679262 | 0.001065739 | 30 | 0.000212863 | 5.991995578 | 8 mo | HOM |
| **1013978366** | 1 | VENT | 637 | 5.006679262 | 0 | 0 | 0 | 0 | 2 mo | HOM |
| **1014416155** | 2 | VENT | 637 | 5.006679262 | 0 | 0 | 0 | 0 | 2 mo | HOM |

| | | | | | | | | | | |
|---|---|---|---|---|---|---|---|---|---|---|
| **1013978065** | 2 | VENT | 637 | 5.006679262 | 0 | 0 | 0 | 0 | 2 mo | HOM |
| **1009934102** | 2 | VENT | 637 | 5.006679262 | 0 | 0 | 0 | 0 | 4 mo | WILD |
| **1009322937** | 2 | VENT | 637 | 5.006679262 | 0 | 0 | 0 | 0 | 8 mo | WILD |
| **1011080637** | 2 | VENT | 637 | 5.006679262 | 0.001729812 | 50 | 0.000345501 | 9.986659297 | 8 mo | HOM |
| **1013609713** | 2 | VENT | 637 | 5.006679262 | 0 | 0 | 0 | 0 | 2 mo | WILD |
| **1010418842** | 1 | VENT | 637 | 5.006679262 | 0.002224471 | 56 | 0.000444301 | 11.18505841 | 8 mo | HOM |
| **1013288213** | 2 | VENT | 637 | 5.006679262 | 2.25E-05 | 3 | 4.49E-06 | 0.599199558 | 4 mo | HOM |
| **1008976060** | 2 | VENT | 637 | 5.006679262 | 8.82E-05 | 10 | 1.76E-05 | 1.997331859 | 8 mo | WILD |
| **1013446243** | 1 | VENT | 637 | 5.006679262 | 6.16E-06 | 1 | 1.23E-06 | 0.199733186 | 4 mo | HOM |
| **1013288951** | 2 | VENT | 637 | 5.006679262 | 5.58E-06 | 1 | 1.11E-06 | 0.199733186 | 4 mo | HOM |
| **1013978665** | 1 | GU6a | 638 | 0.452091562 | 0 | 0 | 0 | 0 | 2 mo | HOM |
| **1013445609** | 1 | GU6a | 638 | 0.452091562 | 3.55E-05 | 2 | 7.86E-05 | 4.423882616 | 4 mo | HOM |
| **1013609015** | 1 | GU6a | 638 | 0.452091562 | 8.07E-05 | 4 | 0.000178445 | 8.847765231 | 4 mo | HOM |
| **1013608106** | 2 | GU6a | 638 | 0.452091562 | 0 | 0 | 0 | 0 | 4 mo | HOM |
| **1011937543** | 1 | GU6a | 638 | 0.452091562 | 0.000983504 | 30 | 0.002175454 | 66.35823923 | 8 mo | HOM |
| **1010419538** | 2 | GU6a | 638 | 0.452091562 | 0 | 0 | 0 | 0 | 4 mo | WILD |
| **1010419187** | 1 | GU6a | 638 | 0.452091562 | 0.000403743 | 17 | 0.000893056 | 37.60300223 | 8 mo | HOM |
| **1013978366** | 1 | GU6a | 638 | 0.452091562 | 0 | 0 | 0 | 0 | 2 mo | HOM |
| **1014416155** | 2 | GU6a | 638 | 0.452091562 | 0 | 0 | 0 | 0 | 2 mo | HOM |
| **1013978065** | 2 | GU6a | 638 | 0.452091562 | 0 | 0 | 0 | 0 | 2 mo | HOM |
| **1009934102** | 2 | GU6a | 638 | 0.452091562 | 0 | 0 | 0 | 0 | 4 mo | WILD |
| **1009322937** | 2 | GU6a | 638 | 0.452091562 | 2.24E-06 | 1 | 4.94E-06 | 2.211941308 | 8 mo | WILD |
| **1011080637** | 2 | GU6a | 638 | 0.452091562 | 0.000137311 | 10 | 0.000303724 | 22.11941308 | 8 mo | HOM |
| **1013609713** | 2 | GU6a | 638 | 0.452091562 | 0 | 0 | 0 | 0 | 2 mo | WILD |
| **1010418842** | 1 | GU6a | 638 | 0.452091562 | 0.000462198 | 16 | 0.001022355 | 35.39106092 | 8 mo | HOM |
| **1013288213** | 2 | GU6a | 638 | 0.452091562 | 1.58E-06 | 0 | 3.49E-06 | 0 | 4 mo | HOM |
| **1008976060** | 2 | GU6a | 638 | 0.452091562 | 4.89E-06 | 1 | 1.08E-05 | 2.211941308 | 8 mo | WILD |
| **1013446243** | 1 | GU6a | 638 | 0.452091562 | 0.000101438 | 5 | 0.000224375 | 11.05970654 | 4 mo | HOM |
| **1013288951** | 2 | GU6a | 638 | 0.452091562 | 4.99E-05 | 2 | 0.000110293 | 4.423882616 | 4 mo | HOM |
| **1013978665** | 1 | COAa | 639 | 0.768275234 | 1.71E-05 | 2 | 2.23E-05 | 2.603233727 | 2 mo | HOM |
| **1013445609** | 1 | COAa | 639 | 0.790672938 | 7.07E-06 | 1 | 8.95E-06 | 1.26474545 | 4 mo | HOM |
| **1013609015** | 1 | COAa | 639 | 0.790672938 | 3.01E-06 | 0 | 3.8E-06 | 0 | 4 mo | HOM |
| **1013608106** | 2 | COAa | 639 | 0.790672938 | 0 | 0 | 0 | 0 | 4 mo | HOM |
| **1011937543** | 1 | COAa | 639 | 0.790672938 | 0.000213014 | 6 | 0.000269408 | 7.588472698 | 8 mo | HOM |
| **1010419538** | 2 | COAa | 639 | 0.790672938 | 0 | 0 | 0 | 0 | 4 mo | WILD |
| **1010419187** | 1 | COAa | 639 | 0.790672938 | 0.000352628 | 12 | 0.000445984 | 15.1769454 | 8 mo | HOM |
| **1013978366** | 1 | COAa | 639 | 0.790672938 | 0 | 0 | 0 | 0 | 2 mo | HOM |
| **1014416155** | 2 | COAa | 639 | 0.790672938 | 0 | 0 | 0 | 0 | 2 mo | HOM |
| **1013978065** | 2 | COAa | 639 | 0.790672938 | 0 | 0 | 0 | 0 | 2 mo | HOM |
| **1009934102** | 2 | COAa | 639 | 0.790672938 | 0 | 0 | 0 | 0 | 4 mo | WILD |
| **1009322937** | 2 | COAa | 639 | 0.790672938 | 0 | 0 | 0 | 0 | 8 mo | WILD |
| **1011080637** | 2 | COAa | 639 | 0.790672938 | 0.000459322 | 19 | 0.000580925 | 24.03016354 | 8 mo | HOM |
| **1013609713** | 2 | COAa | 639 | 0.790672938 | 0 | 0 | 0 | 0 | 2 mo | WILD |
| **1010418842** | 1 | COAa | 639 | 0.790672938 | 0.000356427 | 14 | 0.000450789 | 17.7064363 | 8 mo | HOM |
| **1013288213** | 2 | COAa | 639 | 0.790672938 | 2.11E-05 | 2 | 2.67E-05 | 2.529490899 | 4 mo | HOM |
| **1008976060** | 2 | COAa | 639 | 0.790672938 | 2.39E-20 | 0 | 3.03E-20 | 0 | 8 mo | WILD |
| **1013446243** | 1 | COAa | 639 | 0.790672938 | 6.91E-05 | 3 | 8.74E-05 | 3.794236349 | 4 mo | HOM |
| **1013288951** | 2 | COAa | 639 | 0.790672938 | 2.62E-05 | 2 | 3.32E-05 | 2.529490899 | 4 mo | HOM |
| **1013978665** | 1 | NTB | 642 | 0.159124433 | 0 | 0 | 0 | 0 | 2 mo | HOM |
| **1013445609** | 1 | NTB | 642 | 0.159124433 | 0 | 0 | 0 | 0 | 4 mo | HOM |
| **1013609015** | 1 | NTB | 642 | 0.159124433 | 0 | 0 | 0 | 0 | 4 mo | HOM |
| **1013608106** | 2 | NTB | 642 | 0.159124433 | 0 | 0 | 0 | 0 | 4 mo | HOM |
| **1011937543** | 1 | NTB | 642 | 0.159124433 | 2.98E-05 | 0 | 0.000187111 | 0 | 8 mo | HOM |
| **1010419538** | 2 | NTB | 642 | 0.159124433 | 0 | 0 | 0 | 0 | 4 mo | WILD |
| **1010419187** | 1 | NTB | 642 | 0.159124433 | 0 | 0 | 0 | 0 | 8 mo | HOM |
| **1013978366** | 1 | NTB | 642 | 0.159124433 | 0 | 0 | 0 | 0 | 2 mo | HOM |
| **1014416155** | 2 | NTB | 642 | 0.159124433 | 0 | 0 | 0 | 0 | 2 mo | HOM |
| **1013978065** | 2 | NTB | 642 | 0.159124433 | 0 | 0 | 0 | 0 | 2 mo | HOM |
| **1009934102** | 2 | NTB | 642 | 0.159124433 | 0 | 0 | 0 | 0 | 4 mo | WILD |

| | | | | | | | | | | |
|---|---|---|---|---|---|---|---|---|---|---|
| **1009322937** | 2 | NTB | 642 | 0.159124433 | 0 | 0 | 0 | 0 | 8 mo | WILD |
| **1011080637** | 2 | NTB | 642 | 0.159124433 | 0 | 0 | 0 | 0 | 8 mo | HOM |
| **1013609713** | 2 | NTB | 642 | 0.159124433 | 0 | 0 | 0 | 0 | 2 mo | WILD |
| **1010418842** | 1 | NTB | 642 | 0.159124433 | 0 | 0 | 0 | 0 | 8 mo | HOM |
| **1013288213** | 2 | NTB | 642 | 0.159124433 | 0 | 0 | 0 | 0 | 4 mo | HOM |
| **1008976060** | 2 | NTB | 642 | 0.159124433 | 0 | 0 | 0 | 0 | 8 mo | WILD |
| **1013446243** | 1 | NTB | 642 | 0.159124433 | 0 | 0 | 0 | 0 | 4 mo | HOM |
| **1013288951** | 2 | NTB | 642 | 0.159124433 | 0 | 0 | 0 | 0 | 4 mo | HOM |
| **1013978665** | 1 | AUDpo2/3 | 643 | 0.15138744 | 0 | 0 | 0 | 0 | 2 mo | HOM |
| **1013445609** | 1 | AUDpo2/3 | 643 | 0.15138744 | 1.39E-06 | 0 | 9.17E-06 | 0 | 4 mo | HOM |
| **1013609015** | 1 | AUDpo2/3 | 643 | 0.15138744 | 8.89E-05 | 3 | 0.000587267 | 19.81670337 | 4 mo | HOM |
| **1013608106** | 2 | AUDpo2/3 | 643 | 0.15138744 | 0 | 0 | 0 | 0 | 4 mo | HOM |
| **1011937543** | 1 | AUDpo2/3 | 643 | 0.15138744 | 0.000120333 | 2 | 0.00079487 | 13.21113558 | 8 mo | HOM |
| **1010419538** | 2 | AUDpo2/3 | 643 | 0.15138744 | 0 | 0 | 0 | 0 | 4 mo | WILD |
| **1010419187** | 1 | AUDpo2/3 | 643 | 0.15138744 | 0.000171362 | 5 | 0.001131941 | 33.02783895 | 8 mo | HOM |
| **1013978366** | 1 | AUDpo2/3 | 643 | 0.15138744 | 0 | 0 | 0 | 0 | 2 mo | HOM |
| **1014416155** | 2 | AUDpo2/3 | 643 | 0.15138744 | 0 | 0 | 0 | 0 | 2 mo | HOM |
| **1013978065** | 2 | AUDpo2/3 | 643 | 0.15138744 | 0 | 0 | 0 | 0 | 2 mo | HOM |
| **1009934102** | 2 | AUDpo2/3 | 643 | 0.15138744 | 0 | 0 | 0 | 0 | 4 mo | WILD |
| **1009322937** | 2 | AUDpo2/3 | 643 | 0.15138744 | 0 | 0 | 0 | 0 | 8 mo | WILD |
| **1011080637** | 2 | AUDpo2/3 | 643 | 0.15138744 | 0.000103642 | 5 | 0.000684613 | 33.02783895 | 8 mo | HOM |
| **1013609713** | 2 | AUDpo2/3 | 643 | 0.15138744 | 0 | 0 | 0 | 0 | 2 mo | WILD |
| **1010418842** | 1 | AUDpo2/3 | 643 | 0.15138744 | 0.000155822 | 7 | 0.001029292 | 46.23897452 | 8 mo | HOM |
| **1013288213** | 2 | AUDpo2/3 | 643 | 0.15138744 | 6.25E-06 | 1 | 4.13E-05 | 6.605567789 | 4 mo | HOM |
| **1008976060** | 2 | AUDpo2/3 | 643 | 0.15138744 | 0 | 0 | 0 | 0 | 8 mo | WILD |
| **1013446243** | 1 | AUDpo2/3 | 643 | 0.15138744 | 1.42E-06 | 0 | 9.37E-06 | 0 | 4 mo | HOM |
| **1013288951** | 2 | AUDpo2/3 | 643 | 0.15138744 | 9.2E-06 | 1 | 6.07E-05 | 6.605567789 | 4 mo | HOM |
| **1013978665** | 1 | VERM | 645 | 20.88639041 | 0.002571292 | 91 | 0.000123108 | 4.356904099 | 2 mo | HOM |
| **1013445609** | 1 | VERM | 645 | 20.88639041 | 0.000958625 | 22 | 4.59E-05 | 1.053317474 | 4 mo | HOM |
| **1013609015** | 1 | VERM | 645 | 20.88639041 | 0.008375416 | 167 | 0.000400999 | 7.995637192 | 4 mo | HOM |
| **1013608106** | 2 | VERM | 645 | 20.88639041 | 0 | 0 | 0 | 0 | 4 mo | HOM |
| **1011937543** | 1 | VERM | 645 | 20.88639041 | 0.02315135 | 163 | 0.001108442 | 7.804124924 | 8 mo | HOM |
| **1010419538** | 2 | VERM | 645 | 20.85077529 | 0.001137785 | 50 | 5.46E-05 | 2.397992367 | 4 mo | WILD |
| **1010419187** | 1 | VERM | 645 | 20.88639041 | 0.017780312 | 298 | 0.000851287 | 14.26766397 | 8 mo | HOM |
| **1013978366** | 1 | VERM | 645 | 20.88639041 | 0.005247321 | 109 | 0.000251232 | 5.218709305 | 2 mo | HOM |
| **1014416155** | 2 | VERM | 645 | 20.88639041 | 0 | 0 | 0 | 0 | 2 mo | HOM |
| **1013978065** | 2 | VERM | 645 | 20.88639041 | 0.001396784 | 55 | 6.69E-05 | 2.633293686 | 2 mo | HOM |
| **1009934102** | 2 | VERM | 645 | 20.88639041 | 0.00127583 | 72 | 6.11E-05 | 3.447220825 | 4 mo | WILD |
| **1009322937** | 2 | VERM | 645 | 20.88639041 | 0.002249906 | 92 | 0.000107721 | 4.404782166 | 8 mo | WILD |
| **1011080637** | 2 | VERM | 645 | 20.88639041 | 0.036390847 | 340 | 0.001742323 | 16.27854279 | 8 mo | HOM |
| **1013609713** | 2 | VERM | 645 | 20.87510387 | 0.000774927 | 32 | 3.71E-05 | 1.532926504 | 2 mo | WILD |
| **1010418842** | 1 | VERM | 645 | 20.88639041 | 0.036642063 | 443 | 0.001754351 | 21.20998369 | 8 mo | HOM |
| **1013288213** | 2 | VERM | 645 | 20.88639041 | 0.000830771 | 38 | 3.98E-05 | 1.819366547 | 4 mo | HOM |
| **1008976060** | 2 | VERM | 645 | 20.88639041 | 0.013557314 | 240 | 0.000649098 | 11.49073608 | 8 mo | WILD |
| **1013446243** | 1 | VERM | 645 | 20.88639041 | 0.002144853 | 73 | 0.000102691 | 3.495098892 | 4 mo | HOM |
| **1013288951** | 2 | VERM | 645 | 20.88639041 | 0.011822091 | 109 | 0.000566019 | 5.218709305 | 4 mo | HOM |
| **1013978665** | 1 | COAp | 647 | 2.573168282 | 1.67E-06 | 0 | 6.5E-07 | 0 | 2 mo | HOM |
| **1013445609** | 1 | COAp | 647 | 2.573168282 | 0.000217829 | 13 | 8.47E-05 | 5.052137512 | 4 mo | HOM |
| **1013609015** | 1 | COAp | 647 | 2.573168282 | 0.000648456 | 23 | 0.000252007 | 8.938397137 | 4 mo | HOM |
| **1013608106** | 2 | COAp | 647 | 2.573168282 | 0 | 0 | 0 | 0 | 4 mo | HOM |
| **1011937543** | 1 | COAp | 647 | 2.573168282 | 0.005626042 | 79 | 0.002186426 | 30.70145103 | 8 mo | HOM |
| **1010419538** | 2 | COAp | 647 | 2.573168282 | 3.09E-05 | 5 | 1.2E-05 | 1.943129812 | 4 mo | WILD |
| **1010419187** | 1 | COAp | 647 | 2.573168282 | 0.006672125 | 112 | 0.002592961 | 43.5261078 | 8 mo | HOM |
| **1013978366** | 1 | COAp | 647 | 2.573168282 | 0 | 0 | 0 | 0 | 2 mo | HOM |
| **1014416155** | 2 | COAp | 647 | 2.573168282 | 0 | 0 | 0 | 0 | 2 mo | HOM |
| **1013978065** | 2 | COAp | 647 | 2.573168282 | 1.52E-06 | 0 | 5.91E-07 | 0 | 2 mo | HOM |
| **1009934102** | 2 | COAp | 647 | 2.573168282 | 0 | 0 | 0 | 0 | 4 mo | WILD |
| **1009322937** | 2 | COAp | 647 | 2.573168282 | 0 | 0 | 0 | 0 | 8 mo | WILD |
| **1011080637** | 2 | COAp | 647 | 2.573168282 | 0.003898403 | 128 | 0.001515021 | 49.74412319 | 8 mo | HOM |

| | | | | | | | | | | |
|---|---|---|---|---|---|---|---|---|---|---|
| **1013609713** | 2 | COAp | 647 | 2.573168282 | 0 | 0 | 0 | 0 | 2 mo | WILD |
| **1010418842** | 1 | COAp | 647 | 2.573168282 | 0.00572885 | 141 | 0.00222638 | 54.79626071 | 8 mo | HOM |
| **1013288213** | 2 | COAp | 647 | 2.573168282 | 0.000360063 | 21 | 0.00013993 | 8.161145212 | 4 mo | HOM |
| **1008976060** | 2 | COAp | 647 | 2.573168282 | 5.17E-05 | 10 | 2.01E-05 | 3.886259625 | 8 mo | WILD |
| **1013446243** | 1 | COAp | 647 | 2.573168282 | 0.000444677 | 20 | 0.000172813 | 7.772519249 | 4 mo | HOM |
| **1013288951** | 2 | COAp | 647 | 2.573168282 | 0.000919069 | 16 | 0.000357174 | 6.218015399 | 4 mo | HOM |
| **1013978665** | 1 | MOp5 | 648 | 3.371765734 | 0 | 0 | 0 | 0 | 2 mo | HOM |
| **1013445609** | 1 | MOp5 | 648 | 3.371765734 | 0.000315886 | 20 | 9.37E-05 | 5.931610193 | 4 mo | HOM |
| **1013609015** | 1 | MOp5 | 648 | 3.371765734 | 0.00080354 | 29 | 0.000238314 | 8.60083478 | 4 mo | HOM |
| **1013608106** | 2 | MOp5 | 648 | 3.371765734 | 0 | 0 | 0 | 0 | 4 mo | HOM |
| **1011937543** | 1 | MOp5 | 648 | 3.371765734 | 0.004881843 | 100 | 0.001447859 | 29.65805097 | 8 mo | HOM |
| **1010419538** | 2 | MOp5 | 648 | 3.371765734 | 0 | 0 | 0 | 0 | 4 mo | WILD |
| **1010419187** | 1 | MOp5 | 648 | 3.371765734 | 0.004513331 | 185 | 0.001338566 | 54.86739429 | 8 mo | HOM |
| **1013978366** | 1 | MOp5 | 648 | 3.371765734 | 1.03E-05 | 2 | 3.06E-06 | 0.593161019 | 2 mo | HOM |
| **1014416155** | 2 | MOp5 | 648 | 3.371765734 | 0 | 0 | 0 | 0 | 2 mo | HOM |
| **1013978065** | 2 | MOp5 | 648 | 3.371765734 | 0 | 0 | 0 | 0 | 2 mo | HOM |
| **1009934102** | 2 | MOp5 | 648 | 3.371765734 | 0 | 0 | 0 | 0 | 4 mo | WILD |
| **1009322937** | 2 | MOp5 | 648 | 3.371765734 | 0 | 0 | 0 | 0 | 8 mo | WILD |
| **1011080637** | 2 | MOp5 | 648 | 3.371765734 | 0.002226199 | 107 | 0.000660247 | 31.73411453 | 8 mo | HOM |
| **1013609713** | 2 | MOp5 | 648 | 3.371765734 | 0 | 0 | 0 | 0 | 2 mo | WILD |
| **1010418842** | 1 | MOp5 | 648 | 3.371765734 | 0.003215247 | 152 | 0.00095358 | 45.08023747 | 8 mo | HOM |
| **1013288213** | 2 | MOp5 | 648 | 3.371765734 | 0.00022648 | 12 | 6.72E-05 | 3.558966116 | 4 mo | HOM |
| **1008976060** | 2 | MOp5 | 648 | 3.371765734 | 1.71E-05 | 2 | 5.08E-06 | 0.593161019 | 8 mo | WILD |
| **1013446243** | 1 | MOp5 | 648 | 3.371765734 | 0.000847004 | 34 | 0.000251205 | 10.08373733 | 4 mo | HOM |
| **1013288951** | 2 | MOp5 | 648 | 3.371765734 | 0.000749102 | 32 | 0.000222169 | 9.490576309 | 4 mo | HOM |
| **1013978665** | 1 | VISal6b | 649 | 0.022922506 | 0 | 0 | 0 | 0 | 2 mo | HOM |
| **1013445609** | 1 | VISal6b | 649 | 0.022922506 | 0 | 0 | 0 | 0 | 4 mo | HOM |
| **1013609015** | 1 | VISal6b | 649 | 0.022922506 | 7.11E-06 | 0 | 0.000310338 | 0 | 4 mo | HOM |
| **1013608106** | 2 | VISal6b | 649 | 0.022922506 | 0 | 0 | 0 | 0 | 4 mo | HOM |
| **1011937543** | 1 | VISal6b | 649 | 0.022922506 | 2.2E-05 | 0 | 0.000961499 | 0 | 8 mo | HOM |
| **1010419538** | 2 | VISal6b | 649 | 0.022922506 | 0 | 0 | 0 | 0 | 4 mo | WILD |
| **1010419187** | 1 | VISal6b | 649 | 0.022922506 | 6.36E-05 | 1 | 0.002775397 | 43.62524713 | 8 mo | HOM |
| **1013978366** | 1 | VISal6b | 649 | 0.022922506 | 0 | 0 | 0 | 0 | 2 mo | HOM |
| **1014416155** | 2 | VISal6b | 649 | 0.022922506 | 0 | 0 | 0 | 0 | 2 mo | HOM |
| **1013978065** | 2 | VISal6b | 649 | 0.022922506 | 0 | 0 | 0 | 0 | 2 mo | HOM |
| **1009934102** | 2 | VISal6b | 649 | 0.022922506 | 0 | 0 | 0 | 0 | 4 mo | WILD |
| **1009322937** | 2 | VISal6b | 649 | 0.022922506 | 0 | 0 | 0 | 0 | 8 mo | WILD |
| **1011080637** | 2 | VISal6b | 649 | 0.022922506 | 1.14E-05 | 1 | 0.000498977 | 43.62524713 | 8 mo | HOM |
| **1013609713** | 2 | VISal6b | 649 | 0.022922506 | 0 | 0 | 0 | 0 | 2 mo | WILD |
| **1010418842** | 1 | VISal6b | 649 | 0.022922506 | 4.71E-05 | 0 | 0.002054467 | 0 | 8 mo | HOM |
| **1013288213** | 2 | VISal6b | 649 | 0.022922506 | 6.41E-06 | 1 | 0.000279836 | 43.62524713 | 4 mo | HOM |
| **1008976060** | 2 | VISal6b | 649 | 0.022922506 | 0 | 0 | 0 | 0 | 8 mo | WILD |
| **1013446243** | 1 | VISal6b | 649 | 0.022922506 | 4.81E-06 | 1 | 0.000209917 | 43.62524713 | 4 mo | HOM |
| **1013288951** | 2 | VISal6b | 649 | 0.022922506 | 0 | 0 | 0 | 0 | 4 mo | HOM |
| **1013978665** | 1 | NTS | 651 | 0.874483792 | 0 | 0 | 0 | 0 | 2 mo | HOM |
| **1013445609** | 1 | NTS | 651 | 0.874483792 | 0 | 0 | 0 | 0 | 4 mo | HOM |
| **1013609015** | 1 | NTS | 651 | 0.874483792 | 0 | 0 | 0 | 0 | 4 mo | HOM |
| **1013608106** | 2 | NTS | 651 | 0.874483792 | 0 | 0 | 0 | 0 | 4 mo | HOM |
| **1011937543** | 1 | NTS | 651 | 0.874483792 | 0.003045722 | 3 | 0.00348288 | 3.430595315 | 8 mo | HOM |
| **1010419538** | 2 | NTS | 651 | 0.874483792 | 0 | 0 | 0 | 0 | 4 mo | WILD |
| **1010419187** | 1 | NTS | 651 | 0.874483792 | 1.39E-05 | 1 | 1.59E-05 | 1.143531772 | 8 mo | HOM |
| **1013978366** | 1 | NTS | 651 | 0.874483792 | 0 | 0 | 0 | 0 | 2 mo | HOM |
| **1014416155** | 2 | NTS | 651 | 0.874483792 | 0 | 0 | 0 | 0 | 2 mo | HOM |
| **1013978065** | 2 | NTS | 651 | 0.874483792 | 4.22E-09 | 0 | 4.83E-09 | 0 | 2 mo | HOM |
| **1009934102** | 2 | NTS | 651 | 0.874483792 | 0 | 0 | 0 | 0 | 4 mo | WILD |
| **1009322937** | 2 | NTS | 651 | 0.874483792 | 0.000117932 | 2 | 0.000134859 | 2.287063543 | 8 mo | WILD |
| **1011080637** | 2 | NTS | 651 | 0.874483792 | 0.000167025 | 5 | 0.000190998 | 5.717658859 | 8 mo | HOM |
| **1013609713** | 2 | NTS | 651 | 0.874483792 | 0 | 0 | 0 | 0 | 2 mo | WILD |
| **1010418842** | 1 | NTS | 651 | 0.874483792 | 0.00028521 | 8 | 0.000326147 | 9.148254174 | 8 mo | HOM |

| | | | | | | | | | | |
|---|---|---|---|---|---|---|---|---|---|---|
| 1013288213 | 2 | NTS | 651 | 0.874483792 | 0 | 0 | 0 | 0 | 4 mo | HOM |
| 1008976060 | 2 | NTS | 651 | 0.874483792 | 7.66E-05 | 6 | 8.76E-05 | 6.86119063 | 8 mo | WILD |
| 1013446243 | 1 | NTS | 651 | 0.874483792 | 0.000329181 | 2 | 0.000376429 | 2.287063543 | 4 mo | HOM |
| 1013288951 | 2 | NTS | 651 | 0.874483792 | 2.54E-07 | 0 | 2.9E-07 | 0 | 4 mo | HOM |
| 1013978665 | 1 | VI | 653 | 0.031248785 | 0 | 0 | 0 | 0 | 2 mo | HOM |
| 1013445609 | 1 | VI | 653 | 0.031248785 | 0 | 0 | 0 | 0 | 4 mo | HOM |
| 1013609015 | 1 | VI | 653 | 0.031248785 | 0 | 0 | 0 | 0 | 4 mo | HOM |
| 1013608106 | 2 | VI | 653 | 0.031248785 | 0 | 0 | 0 | 0 | 4 mo | HOM |
| 1011937543 | 1 | VI | 653 | 0.031248785 | 0 | 0 | 0 | 0 | 8 mo | HOM |
| 1010419538 | 2 | VI | 653 | 0.031248785 | 0 | 0 | 0 | 0 | 4 mo | WILD |
| 1010419187 | 1 | VI | 653 | 0.031248785 | 0 | 0 | 0 | 0 | 8 mo | HOM |
| 1013978366 | 1 | VI | 653 | 0.031248785 | 0 | 0 | 0 | 0 | 2 mo | HOM |
| 1014416155 | 2 | VI | 653 | 0.031248785 | 0 | 0 | 0 | 0 | 2 mo | HOM |
| 1013978065 | 2 | VI | 653 | 0.031248785 | 0 | 0 | 0 | 0 | 2 mo | HOM |
| 1009934102 | 2 | VI | 653 | 0.031248785 | 0 | 0 | 0 | 0 | 4 mo | WILD |
| 1009322937 | 2 | VI | 653 | 0.031248785 | 0 | 0 | 0 | 0 | 8 mo | WILD |
| 1011080637 | 2 | VI | 653 | 0.031248785 | 0 | 0 | 0 | 0 | 8 mo | HOM |
| 1013609713 | 2 | VI | 653 | 0.031248785 | 0 | 0 | 0 | 0 | 2 mo | WILD |
| 1010418842 | 1 | VI | 653 | 0.031248785 | 0 | 0 | 0 | 0 | 8 mo | HOM |
| 1013288213 | 2 | VI | 653 | 0.031248785 | 0 | 0 | 0 | 0 | 4 mo | HOM |
| 1008976060 | 2 | VI | 653 | 0.031248785 | 0 | 0 | 0 | 0 | 8 mo | WILD |
| 1013446243 | 1 | VI | 653 | 0.031248785 | 0 | 0 | 0 | 0 | 4 mo | HOM |
| 1013288951 | 2 | VI | 653 | 0.031248785 | 0 | 0 | 0 | 0 | 4 mo | HOM |
| 1013978665 | 1 | SSp-n4 | 654 | 0.564695169 | 0 | 0 | 0 | 0 | 2 mo | HOM |
| 1013445609 | 1 | SSp-n4 | 654 | 0.564695169 | 7.08E-05 | 2 | 0.000125365 | 3.54173386 | 4 mo | HOM |
| 1013609015 | 1 | SSp-n4 | 654 | 0.564695169 | 0.000139402 | 6 | 0.000246863 | 10.62520158 | 4 mo | HOM |
| 1013608106 | 2 | SSp-n4 | 654 | 0.564695169 | 0 | 0 | 0 | 0 | 4 mo | HOM |
| 1011937543 | 1 | SSp-n4 | 654 | 0.564695169 | 0.000958825 | 21 | 0.001697951 | 37.18820553 | 8 mo | HOM |
| 1010419538 | 2 | SSp-n4 | 654 | 0.564695169 | 4.94E-06 | 1 | 8.74E-06 | 1.77086693 | 4 mo | WILD |
| 1010419187 | 1 | SSp-n4 | 654 | 0.564695169 | 0.000662918 | 25 | 0.00117394 | 44.27167325 | 8 mo | HOM |
| 1013978366 | 1 | SSp-n4 | 654 | 0.564695169 | 0 | 0 | 0 | 0 | 2 mo | HOM |
| 1014416155 | 2 | SSp-n4 | 654 | 0.564695169 | 0 | 0 | 0 | 0 | 2 mo | HOM |
| 1013978065 | 2 | SSp-n4 | 654 | 0.564695169 | 0 | 0 | 0 | 0 | 2 mo | HOM |
| 1009934102 | 2 | SSp-n4 | 654 | 0.564695169 | 3.77E-06 | 1 | 6.68E-06 | 1.77086693 | 4 mo | WILD |
| 1009322937 | 2 | SSp-n4 | 654 | 0.564695169 | 0 | 0 | 0 | 0 | 8 mo | WILD |
| 1011080637 | 2 | SSp-n4 | 654 | 0.564695169 | 0.0006862 | 30 | 0.00121517 | 53.1260079 | 8 mo | HOM |
| 1013609713 | 2 | SSp-n4 | 654 | 0.564695169 | 5.96E-06 | 1 | 1.06E-05 | 1.77086693 | 2 mo | WILD |
| 1010418842 | 1 | SSp-n4 | 654 | 0.564695169 | 0.000618848 | 28 | 0.001095898 | 49.58427404 | 8 mo | HOM |
| 1013288213 | 2 | SSp-n4 | 654 | 0.564695169 | 7.47E-05 | 5 | 0.000132217 | 8.85433465 | 4 mo | HOM |
| 1008976060 | 2 | SSp-n4 | 654 | 0.564695169 | 8.27E-06 | 1 | 1.47E-05 | 1.77086693 | 8 mo | WILD |
| 1013446243 | 1 | SSp-n4 | 654 | 0.564695169 | 0.000245126 | 10 | 0.000434086 | 17.7086693 | 4 mo | HOM |
| 1013288951 | 2 | SSp-n4 | 654 | 0.564695169 | 7.26E-05 | 3 | 0.000128568 | 5.31260079 | 4 mo | HOM |
| 1013978665 | 1 | COApl | 655 | 1.238676566 | 0 | 0 | 0 | 0 | 2 mo | HOM |
| 1013445609 | 1 | COApl | 655 | 1.238676566 | 7.85E-05 | 6 | 6.34E-05 | 4.843879478 | 4 mo | HOM |
| 1013609015 | 1 | COApl | 655 | 1.238676566 | 0.000116276 | 5 | 9.39E-05 | 4.036566232 | 4 mo | HOM |
| 1013608106 | 2 | COApl | 655 | 1.238676566 | 0 | 0 | 0 | 0 | 4 mo | HOM |
| 1011937543 | 1 | COApl | 655 | 1.238676566 | 0.002099799 | 29 | 0.001695195 | 23.41208414 | 8 mo | HOM |
| 1010419538 | 2 | COApl | 655 | 1.238676566 | 2.98E-05 | 4 | 2.41E-05 | 3.229252985 | 4 mo | WILD |
| 1010419187 | 1 | COApl | 655 | 1.238676566 | 0.002809282 | 57 | 0.002267971 | 46.01685504 | 8 mo | HOM |
| 1013978366 | 1 | COApl | 655 | 1.238676566 | 0 | 0 | 0 | 0 | 2 mo | HOM |
| 1014416155 | 2 | COApl | 655 | 1.238676566 | 0 | 0 | 0 | 0 | 2 mo | HOM |
| 1013978065 | 2 | COApl | 655 | 1.238676566 | 1.52E-06 | 0 | 1.23E-06 | 0 | 2 mo | HOM |
| 1009934102 | 2 | COApl | 655 | 1.238676566 | 0 | 0 | 0 | 0 | 4 mo | WILD |
| 1009322937 | 2 | COApl | 655 | 1.238676566 | 0 | 0 | 0 | 0 | 8 mo | WILD |
| 1011080637 | 2 | COApl | 655 | 1.238676566 | 0.001625516 | 45 | 0.0013123 | 36.32909609 | 8 mo | HOM |
| 1013609713 | 2 | COApl | 655 | 1.238676566 | 0 | 0 | 0 | 0 | 2 mo | WILD |
| 1010418842 | 1 | COApl | 655 | 1.238676566 | 0.002494585 | 60 | 0.002013911 | 48.43879478 | 8 mo | HOM |
| 1013288213 | 2 | COApl | 655 | 1.238676566 | 0.000149721 | 9 | 0.000120872 | 7.265819217 | 4 mo | HOM |
| 1008976060 | 2 | COApl | 655 | 1.238676566 | 4.93E-06 | 1 | 3.98E-06 | 0.807313246 | 8 mo | WILD |

| | | | | | | | | | | |
|---|---|---|---|---|---|---|---|---|---|---|
| **1013446243** | 1 | COApl | 655 | 1.238676566 | 0.000238813 | 9 | 0.000192797 | 7.265819217 | 4 mo | HOM |
| **1013288951** | 2 | COApl | 655 | 1.238676566 | 0.000501803 | 8 | 0.000405112 | 6.458505971 | 4 mo | HOM |
| **1013978665** | 1 | MOs1 | 656 | 2.390560883 | 0.000262127 | 13 | 0.000109651 | 5.438054346 | 2 mo | HOM |
| **1013445609** | 1 | MOs1 | 656 | 2.390560883 | 0.000213897 | 12 | 8.95E-05 | 5.019742473 | 4 mo | HOM |
| **1013609015** | 1 | MOs1 | 656 | 2.390560883 | 0.000517038 | 19 | 0.000216283 | 7.947925582 | 4 mo | HOM |
| **1013608106** | 2 | MOs1 | 656 | 2.390560883 | 0 | 0 | 0 | 0 | 4 mo | HOM |
| **1011937543** | 1 | MOs1 | 656 | 2.390560883 | 0.005475438 | 91 | 0.00229044 | 38.06638042 | 8 mo | HOM |
| **1010419538** | 2 | MOs1 | 656 | 2.390560883 | 6.15E-05 | 3 | 2.57E-05 | 1.254935618 | 4 mo | WILD |
| **1010419187** | 1 | MOs1 | 656 | 2.390560883 | 0.005809863 | 172 | 0.002430334 | 71.94964211 | 8 mo | HOM |
| **1013978366** | 1 | MOs1 | 656 | 2.390560883 | 2.1E-06 | 0 | 8.77E-07 | 0 | 2 mo | HOM |
| **1014416155** | 2 | MOs1 | 656 | 2.390560883 | 0 | 0 | 0 | 0 | 2 mo | HOM |
| **1013978065** | 2 | MOs1 | 656 | 2.390560883 | 0 | 0 | 0 | 0 | 2 mo | HOM |
| **1009934102** | 2 | MOs1 | 656 | 2.390560883 | 0.000145472 | 4 | 6.09E-05 | 1.673247491 | 4 mo | WILD |
| **1009322937** | 2 | MOs1 | 656 | 2.390560883 | 4.41E-05 | 3 | 1.85E-05 | 1.254935618 | 8 mo | WILD |
| **1011080637** | 2 | MOs1 | 656 | 2.390560883 | 0.00122453 | 40 | 0.000512236 | 16.73247491 | 8 mo | HOM |
| **1013609713** | 2 | MOs1 | 656 | 2.390560883 | 2.97E-06 | 0 | 1.24E-06 | 0 | 2 mo | WILD |
| **1010418842** | 1 | MOs1 | 656 | 2.390560883 | 0.002298959 | 92 | 0.000961682 | 38.48469229 | 8 mo | HOM |
| **1013288213** | 2 | MOs1 | 656 | 2.390560883 | 6.39E-05 | 4 | 2.67E-05 | 1.673247491 | 4 mo | HOM |
| **1008976060** | 2 | MOs1 | 656 | 2.390560883 | 0.00077877 | 24 | 0.000325769 | 10.03948495 | 8 mo | WILD |
| **1013446243** | 1 | MOs1 | 656 | 2.390560883 | 0.00101933 | 17 | 0.000426398 | 7.111301837 | 4 mo | HOM |
| **1013288951** | 2 | MOs1 | 656 | 2.390560883 | 0.001759902 | 68 | 0.000736188 | 28.44520735 | 4 mo | HOM |
| **1013978665** | 1 | SSp-m2/3 | 657 | 1.542498205 | 9.08E-09 | 0 | 5.89E-09 | 0 | 2 mo | HOM |
| **1013445609** | 1 | SSp-m2/3 | 657 | 1.542498205 | 0.0002117 | 9 | 0.000137245 | 5.834690745 | 4 mo | HOM |
| **1013609015** | 1 | SSp-m2/3 | 657 | 1.542498205 | 0.000544583 | 16 | 0.000353053 | 10.37278355 | 4 mo | HOM |
| **1013608106** | 2 | SSp-m2/3 | 657 | 1.542498205 | 0 | 0 | 0 | 0 | 4 mo | HOM |
| **1011937543** | 1 | SSp-m2/3 | 657 | 1.542498205 | 0.002499494 | 41 | 0.001620419 | 26.58025784 | 8 mo | HOM |
| **1010419538** | 2 | SSp-m2/3 | 657 | 1.542498205 | 0 | 0 | 0 | 0 | 4 mo | WILD |
| **1010419187** | 1 | SSp-m2/3 | 657 | 1.542498205 | 0.002491337 | 70 | 0.001615131 | 45.38092802 | 8 mo | HOM |
| **1013978366** | 1 | SSp-m2/3 | 657 | 1.542498205 | 0 | 0 | 0 | 0 | 2 mo | HOM |
| **1014416155** | 2 | SSp-m2/3 | 657 | 1.542498205 | 4E-07 | 0 | 2.6E-07 | 0 | 2 mo | HOM |
| **1013978065** | 2 | SSp-m2/3 | 657 | 1.542498205 | 0 | 0 | 0 | 0 | 2 mo | HOM |
| **1009934102** | 2 | SSp-m2/3 | 657 | 1.542498205 | 0 | 0 | 0 | 0 | 4 mo | WILD |
| **1009322937** | 2 | SSp-m2/3 | 657 | 1.542498205 | 0 | 0 | 0 | 0 | 8 mo | WILD |
| **1011080637** | 2 | SSp-m2/3 | 657 | 1.542498205 | 0.001155748 | 47 | 0.00074927 | 30.47005167 | 8 mo | HOM |
| **1013609713** | 2 | SSp-m2/3 | 657 | 1.542498205 | 0 | 0 | 0 | 0 | 2 mo | WILD |
| **1010418842** | 1 | SSp-m2/3 | 657 | 1.542498205 | 0.001316747 | 57 | 0.000853645 | 36.95304139 | 8 mo | HOM |
| **1013288213** | 2 | SSp-m2/3 | 657 | 1.542498205 | 4.41E-05 | 4 | 2.86E-05 | 2.593195887 | 4 mo | HOM |
| **1008976060** | 2 | SSp-m2/3 | 657 | 1.542498205 | 0 | 0 | 0 | 0 | 8 mo | WILD |
| **1013446243** | 1 | SSp-m2/3 | 657 | 1.542498205 | 0.00047122 | 21 | 0.000305491 | 13.61427841 | 4 mo | HOM |
| **1013288951** | 2 | SSp-m2/3 | 657 | 1.542498205 | 0.000395525 | 16 | 0.000256418 | 10.37278355 | 4 mo | HOM |
| **1013978665** | 1 | II | 658 | 0.860369754 | 3.84E-07 | 0 | 4.46E-07 | 0 | 2 mo | HOM |
| **1013445609** | 1 | II | 658 | 0.860369754 | 6.23E-07 | 0 | 7.24E-07 | 0 | 4 mo | HOM |
| **1013609015** | 1 | II | 658 | 0.860369754 | 2.03E-05 | 0 | 2.36E-05 | 0 | 4 mo | HOM |
| **1013608106** | 2 | II | 658 | 0.860369754 | 0 | 0 | 0 | 0 | 4 mo | HOM |
| **1011937543** | 1 | II | 658 | 0.860369754 | 0.001641673 | 15 | 0.001908102 | 17.43436463 | 8 mo | HOM |
| **1010419538** | 2 | II | 658 | 0.860369754 | 0 | 0 | 0 | 0 | 4 mo | WILD |
| **1010419187** | 1 | II | 658 | 0.860369754 | 0.000453689 | 2 | 0.000527319 | 2.324581951 | 8 mo | HOM |
| **1013978366** | 1 | II | 658 | 0.860369754 | 2.19E-07 | 0 | 2.55E-07 | 0 | 2 mo | HOM |
| **1014416155** | 2 | II | 658 | 0.860369754 | 0 | 0 | 0 | 0 | 2 mo | HOM |
| **1013978065** | 2 | II | 658 | 0.860369754 | 0 | 0 | 0 | 0 | 2 mo | HOM |
| **1009934102** | 2 | II | 658 | 0.860369754 | 0 | 0 | 0 | 0 | 4 mo | WILD |
| **1009322937** | 2 | II | 658 | 0.860369754 | 0 | 0 | 0 | 0 | 8 mo | WILD |
| **1011080637** | 2 | II | 658 | 0.860369754 | 2.88E-07 | 0 | 3.34E-07 | 0 | 8 mo | HOM |
| **1013609713** | 2 | II | 658 | 0.860369754 | 0 | 0 | 0 | 0 | 2 mo | WILD |
| **1010418842** | 1 | II | 658 | 0.860369754 | 0.000101209 | 3 | 0.000117634 | 3.486872926 | 8 mo | HOM |
| **1013288213** | 2 | II | 658 | 0.860369754 | 8.77E-08 | 0 | 1.02E-07 | 0 | 4 mo | HOM |
| **1008976060** | 2 | II | 658 | 0.860369754 | 2.54E-05 | 0 | 2.95E-05 | 0 | 8 mo | WILD |
| **1013446243** | 1 | II | 658 | 0.860369754 | 1.36E-06 | 0 | 1.58E-06 | 0 | 4 mo | HOM |
| **1013288951** | 2 | II | 658 | 0.860369754 | 0 | 0 | 0 | 0 | 4 mo | HOM |

| | | | | | | | | | | |
|---|---|---|---|---|---|---|---|---|---|---|
| 1013978665 | 1 | VII | 661 | 0.944611259 | 0 | 0 | 0 | 0 | 2 mo | HOM |
| 1013445609 | 1 | VII | 661 | 0.944611259 | 0 | 0 | 0 | 0 | 4 mo | HOM |
| 1013609015 | 1 | VII | 661 | 0.944611259 | 0 | 0 | 0 | 0 | 4 mo | HOM |
| 1013608106 | 2 | VII | 661 | 0.944611259 | 0 | 0 | 0 | 0 | 4 mo | HOM |
| 1011937543 | 1 | VII | 661 | 0.944611259 | 0.000659867 | 2 | 0.000698559 | 2.11727309 | 8 mo | HOM |
| 1010419538 | 2 | VII | 661 | 0.944611259 | 0 | 0 | 0 | 0 | 4 mo | WILD |
| 1010419187 | 1 | VII | 661 | 0.944611259 | 0 | 0 | 0 | 0 | 8 mo | HOM |
| 1013978366 | 1 | VII | 661 | 0.944611259 | 0 | 0 | 0 | 0 | 2 mo | HOM |
| 1014416155 | 2 | VII | 661 | 0.944611259 | 0 | 0 | 0 | 0 | 2 mo | HOM |
| 1013978065 | 2 | VII | 661 | 0.944611259 | 0 | 0 | 0 | 0 | 2 mo | HOM |
| 1009934102 | 2 | VII | 661 | 0.944611259 | 0 | 0 | 0 | 0 | 4 mo | WILD |
| 1009322937 | 2 | VII | 661 | 0.944611259 | 0 | 0 | 0 | 0 | 8 mo | WILD |
| 1011080637 | 2 | VII | 661 | 0.944611259 | 3.64E-05 | 1 | 3.86E-05 | 1.058636545 | 8 mo | HOM |
| 1013609713 | 2 | VII | 661 | 0.944611259 | 0 | 0 | 0 | 0 | 2 mo | WILD |
| 1010418842 | 1 | VII | 661 | 0.944611259 | 3.33E-05 | 1 | 3.53E-05 | 1.058636545 | 8 mo | HOM |
| 1013288213 | 2 | VII | 661 | 0.944611259 | 0 | 0 | 0 | 0 | 4 mo | HOM |
| 1008976060 | 2 | VII | 661 | 0.944611259 | 6.12E-06 | 1 | 6.48E-06 | 1.058636545 | 8 mo | WILD |
| 1013446243 | 1 | VII | 661 | 0.944611259 | 0 | 0 | 0 | 0 | 4 mo | HOM |
| 1013288951 | 2 | VII | 661 | 0.944611259 | 0 | 0 | 0 | 0 | 4 mo | HOM |
| 1013978665 | 1 | GU6b | 662 | 0.040533172 | 0 | 0 | 0 | 0 | 2 mo | HOM |
| 1013445609 | 1 | GU6b | 662 | 0.040533172 | 0 | 0 | 0 | 0 | 4 mo | HOM |
| 1013609015 | 1 | GU6b | 662 | 0.040533172 | 5.09E-06 | 0 | 0.000125542 | 0 | 4 mo | HOM |
| 1013608106 | 2 | GU6b | 662 | 0.040533172 | 0 | 0 | 0 | 0 | 4 mo | HOM |
| 1011937543 | 1 | GU6b | 662 | 0.040533172 | 7.58E-05 | 3 | 0.001869557 | 74.01345176 | 8 mo | HOM |
| 1010419538 | 2 | GU6b | 662 | 0.040533172 | 0 | 0 | 0 | 0 | 4 mo | WILD |
| 1010419187 | 1 | GU6b | 662 | 0.040533172 | 1.66E-05 | 1 | 0.000408611 | 24.67115059 | 8 mo | HOM |
| 1013978366 | 1 | GU6b | 662 | 0.040533172 | 0 | 0 | 0 | 0 | 2 mo | HOM |
| 1014416155 | 2 | GU6b | 662 | 0.040533172 | 0 | 0 | 0 | 0 | 2 mo | HOM |
| 1013978065 | 2 | GU6b | 662 | 0.040533172 | 0 | 0 | 0 | 0 | 2 mo | HOM |
| 1009934102 | 2 | GU6b | 662 | 0.040533172 | 0 | 0 | 0 | 0 | 4 mo | WILD |
| 1009322937 | 2 | GU6b | 662 | 0.040533172 | 0 | 0 | 0 | 0 | 8 mo | WILD |
| 1011080637 | 2 | GU6b | 662 | 0.040533172 | 2.52E-06 | 0 | 6.21E-05 | 0 | 8 mo | HOM |
| 1013609713 | 2 | GU6b | 662 | 0.040533172 | 0 | 0 | 0 | 0 | 2 mo | WILD |
| 1010418842 | 1 | GU6b | 662 | 0.040533172 | 1.03E-05 | 2 | 0.000253252 | 49.34230117 | 8 mo | HOM |
| 1013288213 | 2 | GU6b | 662 | 0.040533172 | 0 | 0 | 0 | 0 | 4 mo | HOM |
| 1008976060 | 2 | GU6b | 662 | 0.040533172 | 1.4E-07 | 0 | 3.45E-06 | 0 | 8 mo | WILD |
| 1013446243 | 1 | GU6b | 662 | 0.040533172 | 8.97E-06 | 0 | 0.000221398 | 0 | 4 mo | HOM |
| 1013288951 | 2 | GU6b | 662 | 0.040533172 | 0 | 0 | 0 | 0 | 4 mo | HOM |
| 1013978665 | 1 | COApm | 663 | 1.334491715 | 1.67E-06 | 0 | 1.25E-06 | 0 | 2 mo | HOM |
| 1013445609 | 1 | COApm | 663 | 1.334491715 | 0.000139347 | 7 | 0.000104419 | 5.245442831 | 4 mo | HOM |
| 1013609015 | 1 | COApm | 663 | 1.334491715 | 0.00053218 | 18 | 0.000398789 | 13.48828156 | 4 mo | HOM |
| 1013608106 | 2 | COApm | 663 | 1.334491715 | 0 | 0 | 0 | 0 | 4 mo | HOM |
| 1011937543 | 1 | COApm | 663 | 1.334491715 | 0.003526243 | 50 | 0.002642386 | 37.46744879 | 8 mo | HOM |
| 1010419538 | 2 | COApm | 663 | 1.334491715 | 1.13E-06 | 1 | 8.46E-07 | 0.749348976 | 4 mo | WILD |
| 1010419187 | 1 | COApm | 663 | 1.334491715 | 0.003862842 | 55 | 0.002894617 | 41.21419367 | 8 mo | HOM |
| 1013978366 | 1 | COApm | 663 | 1.334491715 | 0 | 0 | 0 | 0 | 2 mo | HOM |
| 1014416155 | 2 | COApm | 663 | 1.334491715 | 0 | 0 | 0 | 0 | 2 mo | HOM |
| 1013978065 | 2 | COApm | 663 | 1.334491715 | 0 | 0 | 0 | 0 | 2 mo | HOM |
| 1009934102 | 2 | COApm | 663 | 1.334491715 | 0 | 0 | 0 | 0 | 4 mo | WILD |
| 1009322937 | 2 | COApm | 663 | 1.334491715 | 0 | 0 | 0 | 0 | 8 mo | WILD |
| 1011080637 | 2 | COApm | 663 | 1.334491715 | 0.002272888 | 83 | 0.001703186 | 62.19596499 | 8 mo | HOM |
| 1013609713 | 2 | COApm | 663 | 1.334491715 | 0 | 0 | 0 | 0 | 2 mo | WILD |
| 1010418842 | 1 | COApm | 663 | 1.334491715 | 0.003234265 | 81 | 0.002423593 | 60.69726704 | 8 mo | HOM |
| 1013288213 | 2 | COApm | 663 | 1.334491715 | 0.000210342 | 12 | 0.000157619 | 8.99218771 | 4 mo | HOM |
| 1008976060 | 2 | COApm | 663 | 1.334491715 | 4.68E-05 | 9 | 3.51E-05 | 6.744140782 | 8 mo | WILD |
| 1013446243 | 1 | COApm | 663 | 1.334491715 | 0.000205865 | 11 | 0.000154264 | 8.242838734 | 4 mo | HOM |
| 1013288951 | 2 | COApm | 663 | 1.334491715 | 0.000417267 | 8 | 0.000312678 | 5.994791806 | 4 mo | HOM |
| 1013978665 | 1 | ENTm3 | 664 | 1.012863634 | 0 | 0 | 0 | 0 | 2 mo | HOM |
| 1013445609 | 1 | ENTm3 | 664 | 1.012863634 | 1.32E-05 | 2 | 1.31E-05 | 1.974599476 | 4 mo | HOM |

| | | | | | | | | | | |
|---|---|---|---|---|---|---|---|---|---|---|
| **1013609015** | 1 | ENTm3 | 664 | 1.012863634 | 0.000106287 | 3 | 0.000104937 | 2.961899214 | 4 mo | HOM |
| **1013608106** | 2 | ENTm3 | 664 | 1.012863634 | 0 | 0 | 0 | 0 | 4 mo | HOM |
| **1011937543** | 1 | ENTm3 | 664 | 1.012863634 | 0.000705663 | 16 | 0.0006967 | 15.79679581 | 8 mo | HOM |
| **1010419538** | 2 | ENTm3 | 664 | 1.012863634 | 0 | 0 | 0 | 0 | 4 mo | WILD |
| **1010419187** | 1 | ENTm3 | 664 | 1.012863634 | 0.000813166 | 33 | 0.000802839 | 32.58089135 | 8 mo | HOM |
| **1013978366** | 1 | ENTm3 | 664 | 1.012863634 | 0 | 0 | 0 | 0 | 2 mo | HOM |
| **1014416155** | 2 | ENTm3 | 664 | 1.012863634 | 0 | 0 | 0 | 0 | 2 mo | HOM |
| **1013978065** | 2 | ENTm3 | 664 | 1.012863634 | 0 | 0 | 0 | 0 | 2 mo | HOM |
| **1009934102** | 2 | ENTm3 | 664 | 1.012863634 | 0 | 0 | 0 | 0 | 4 mo | WILD |
| **1009322937** | 2 | ENTm3 | 664 | 1.012863634 | 0 | 0 | 0 | 0 | 8 mo | WILD |
| **1011080637** | 2 | ENTm3 | 664 | 1.012863634 | 0.000496759 | 33 | 0.00049045 | 32.58089135 | 8 mo | HOM |
| **1013609713** | 2 | ENTm3 | 664 | 1.012863634 | 0 | 0 | 0 | 0 | 2 mo | WILD |
| **1010418842** | 1 | ENTm3 | 664 | 1.012863634 | 0.001396046 | 38 | 0.001378316 | 37.51739004 | 8 mo | HOM |
| **1013288213** | 2 | ENTm3 | 664 | 1.012863634 | 2.05E-05 | 2 | 2.02E-05 | 1.974599476 | 4 mo | HOM |
| **1008976060** | 2 | ENTm3 | 664 | 1.012863634 | 0 | 0 | 0 | 0 | 8 mo | WILD |
| **1013446243** | 1 | ENTm3 | 664 | 1.012863634 | 6.05E-05 | 4 | 5.98E-05 | 3.949198952 | 4 mo | HOM |
| **1013288951** | 2 | ENTm3 | 664 | 1.012863634 | 0.000208487 | 8 | 0.00020584 | 7.898397903 | 4 mo | HOM |
| **1013978665** | 1 | lot | 665 | 0.931556248 | 0 | 0 | 0 | 0 | 2 mo | HOM |
| **1013445609** | 1 | lot | 665 | 0.931556248 | 3.89E-06 | 0 | 4.18E-06 | 0 | 4 mo | HOM |
| **1013609015** | 1 | lot | 665 | 0.931556248 | 3.14E-06 | 0 | 3.37E-06 | 0 | 4 mo | HOM |
| **1013608106** | 2 | lot | 665 | 0.931556248 | 0 | 0 | 0 | 0 | 4 mo | HOM |
| **1011937543** | 1 | lot | 665 | 0.931556248 | 5.36E-05 | 2 | 5.75E-05 | 2.146944969 | 8 mo | HOM |
| **1010419538** | 2 | lot | 665 | 0.931556248 | 0 | 0 | 0 | 0 | 4 mo | WILD |
| **1010419187** | 1 | lot | 665 | 0.931556248 | 0.000110219 | 3 | 0.000118318 | 3.220417454 | 8 mo | HOM |
| **1013978366** | 1 | lot | 665 | 0.931556248 | 0 | 0 | 0 | 0 | 2 mo | HOM |
| **1014416155** | 2 | lot | 665 | 0.931556248 | 0 | 0 | 0 | 0 | 2 mo | HOM |
| **1013978065** | 2 | lot | 665 | 0.931556248 | 0 | 0 | 0 | 0 | 2 mo | HOM |
| **1009934102** | 2 | lot | 665 | 0.931556248 | 0 | 0 | 0 | 0 | 4 mo | WILD |
| **1009322937** | 2 | lot | 665 | 0.931556248 | 3.73E-06 | 0 | 4E-06 | 0 | 8 mo | WILD |
| **1011080637** | 2 | lot | 665 | 0.931556248 | 0.00023901 | 3 | 0.000256571 | 3.220417454 | 8 mo | HOM |
| **1013609713** | 2 | lot | 665 | 0.931556248 | 0 | 0 | 0 | 0 | 2 mo | WILD |
| **1010418842** | 1 | lot | 665 | 0.931556248 | 1E-04 | 1 | 0.000107298 | 1.073472485 | 8 mo | HOM |
| **1013288213** | 2 | lot | 665 | 0.931556248 | 7.9E-05 | 0 | 8.48E-05 | 0 | 4 mo | HOM |
| **1008976060** | 2 | lot | 665 | 0.931556248 | 5.61E-06 | 1 | 6.02E-06 | 1.073472485 | 8 mo | WILD |
| **1013446243** | 1 | lot | 665 | 0.931556248 | 2.62E-06 | 0 | 2.81E-06 | 0 | 4 mo | HOM |
| **1013288951** | 2 | lot | 665 | 0.931556248 | 6.18E-06 | 1 | 6.63E-06 | 1.073472485 | 4 mo | HOM |
| **1013978665** | 1 | FRP2/3 | 667 | 0.239638577 | 0 | 0 | 0 | 0 | 2 mo | HOM |
| **1013445609** | 1 | FRP2/3 | 667 | 0.239638577 | 0 | 0 | 0 | 0 | 4 mo | HOM |
| **1013609015** | 1 | FRP2/3 | 667 | 0.239638577 | 6.63E-07 | 0 | 2.77E-06 | 0 | 4 mo | HOM |
| **1013608106** | 2 | FRP2/3 | 667 | 0.239638577 | 0 | 0 | 0 | 0 | 4 mo | HOM |
| **1011937543** | 1 | FRP2/3 | 667 | 0.239638577 | 6.91E-05 | 4 | 0.000288444 | 16.69180335 | 8 mo | HOM |
| **1010419538** | 2 | FRP2/3 | 667 | 0.239638577 | 0 | 0 | 0 | 0 | 4 mo | WILD |
| **1010419187** | 1 | FRP2/3 | 667 | 0.239638577 | 5.91E-05 | 3 | 0.000246466 | 12.51885252 | 8 mo | HOM |
| **1013978366** | 1 | FRP2/3 | 667 | 0.239638577 | 0 | 0 | 0 | 0 | 2 mo | HOM |
| **1014416155** | 2 | FRP2/3 | 667 | 0.239638577 | 0 | 0 | 0 | 0 | 2 mo | HOM |
| **1013978065** | 2 | FRP2/3 | 667 | 0.239638577 | 0 | 0 | 0 | 0 | 2 mo | HOM |
| **1009934102** | 2 | FRP2/3 | 667 | 0.239638577 | 0 | 0 | 0 | 0 | 4 mo | WILD |
| **1009322937** | 2 | FRP2/3 | 667 | 0.239638577 | 5.74E-06 | 1 | 2.39E-05 | 4.172950839 | 8 mo | WILD |
| **1011080637** | 2 | FRP2/3 | 667 | 0.239638577 | 2.48E-05 | 1 | 0.000103594 | 4.172950839 | 8 mo | HOM |
| **1013609713** | 2 | FRP2/3 | 667 | 0.239638577 | 0 | 0 | 0 | 0 | 2 mo | WILD |
| **1010418842** | 1 | FRP2/3 | 667 | 0.239638577 | 0.000100569 | 6 | 0.00041967 | 25.03770503 | 8 mo | HOM |
| **1013288213** | 2 | FRP2/3 | 667 | 0.239638577 | 2.09E-05 | 1 | 8.71E-05 | 4.172950839 | 4 mo | HOM |
| **1008976060** | 2 | FRP2/3 | 667 | 0.239638577 | 0 | 0 | 0 | 0 | 8 mo | WILD |
| **1013446243** | 1 | FRP2/3 | 667 | 0.239638577 | 8.65E-06 | 1 | 3.61E-05 | 4.172950839 | 4 mo | HOM |
| **1013288951** | 2 | FRP2/3 | 667 | 0.239638577 | 6.64E-05 | 6 | 0.000277211 | 25.03770503 | 4 mo | HOM |
| **1013978665** | 1 | VIS | 669 | 13.87533814 | 0.000738573 | 32 | 5.32E-05 | 2.306250102 | 2 mo | HOM |
| **1013445609** | 1 | VIS | 669 | 13.87533814 | 0.001603725 | 113 | 0.000115581 | 8.143945672 | 4 mo | HOM |
| **1013609015** | 1 | VIS | 669 | 13.87533814 | 0.005536905 | 193 | 0.000399047 | 13.90957093 | 4 mo | HOM |
| **1013608106** | 2 | VIS | 669 | 13.87533814 | 0 | 0 | 0 | 0 | 4 mo | HOM |

| | | | | | | | | | | |
|---|---|---|---|---|---|---|---|---|---|---|
| **1011937543** | 1 | VIS | 669 | 13.87533814 | 0.034555447 | 349 | 0.002490422 | 25.15254017 | 8 mo | HOM |
| **1010419538** | 2 | VIS | 669 | 13.87533814 | 0.000410976 | 13 | 2.96E-05 | 0.936914104 | 4 mo | WILD |
| **1010419187** | 1 | VIS | 669 | 13.87533814 | 0.046012802 | 764 | 0.003316157 | 55.06172118 | 8 mo | HOM |
| **1013978366** | 1 | VIS | 669 | 13.87533814 | 0.000893129 | 22 | 6.44E-05 | 1.585546945 | 2 mo | HOM |
| **1014416155** | 2 | VIS | 669 | 13.87533814 | 0 | 0 | 0 | 0 | 2 mo | HOM |
| **1013978065** | 2 | VIS | 669 | 13.87533814 | 2.54E-05 | 2 | 1.83E-06 | 0.144140631 | 2 mo | HOM |
| **1009934102** | 2 | VIS | 669 | 13.87533814 | 0.000297374 | 13 | 2.14E-05 | 0.936914104 | 4 mo | WILD |
| **1009322937** | 2 | VIS | 669 | 13.87533814 | 9.55E-05 | 11 | 6.88E-06 | 0.792773472 | 8 mo | WILD |
| **1011080637** | 2 | VIS | 669 | 13.87533814 | 0.015992585 | 585 | 0.001152591 | 42.16113467 | 8 mo | HOM |
| **1013609713** | 2 | VIS | 669 | 13.87533814 | 7.69E-05 | 6 | 5.54E-06 | 0.432421894 | 2 mo | WILD |
| **1010418842** | 1 | VIS | 669 | 13.87533814 | 0.023265335 | 623 | 0.00167674 | 44.89980667 | 8 mo | HOM |
| **1013288213** | 2 | VIS | 669 | 13.87533814 | 0.001580011 | 117 | 0.000113872 | 8.432226934 | 4 mo | HOM |
| **1008976060** | 2 | VIS | 669 | 13.87533814 | 0.00300787 | 55 | 0.000216778 | 3.963867362 | 8 mo | WILD |
| **1013446243** | 1 | VIS | 669 | 13.87533814 | 0.004368252 | 199 | 0.000314821 | 14.34199282 | 4 mo | HOM |
| **1013288951** | 2 | VIS | 669 | 13.87533814 | 0.009945147 | 206 | 0.00071675 | 14.84648503 | 4 mo | HOM |
| **1013978665** | 1 | SSp-tr2/3 | 670 | 0.436541353 | 0 | 0 | 0 | 0 | 2 mo | HOM |
| **1013445609** | 1 | SSp-tr2/3 | 670 | 0.436541353 | 5.82E-05 | 4 | 0.000133299 | 9.162934907 | 4 mo | HOM |
| **1013609015** | 1 | SSp-tr2/3 | 670 | 0.436541353 | 0.000165601 | 6 | 0.000379348 | 13.74440236 | 4 mo | HOM |
| **1013608106** | 2 | SSp-tr2/3 | 670 | 0.436541353 | 0 | 0 | 0 | 0 | 4 mo | HOM |
| **1011937543** | 1 | SSp-tr2/3 | 670 | 0.436541353 | 0.000803549 | 6 | 0.001840718 | 13.74440236 | 8 mo | HOM |
| **1010419538** | 2 | SSp-tr2/3 | 670 | 0.436541353 | 0 | 0 | 0 | 0 | 4 mo | WILD |
| **1010419187** | 1 | SSp-tr2/3 | 670 | 0.436541353 | 0.000849464 | 21 | 0.001945895 | 48.10540826 | 8 mo | HOM |
| **1013978366** | 1 | SSp-tr2/3 | 670 | 0.436541353 | 0 | 0 | 0 | 0 | 2 mo | HOM |
| **1014416155** | 2 | SSp-tr2/3 | 670 | 0.436541353 | 0 | 0 | 0 | 0 | 2 mo | HOM |
| **1013978065** | 2 | SSp-tr2/3 | 670 | 0.436541353 | 0 | 0 | 0 | 0 | 2 mo | HOM |
| **1009934102** | 2 | SSp-tr2/3 | 670 | 0.436541353 | 0 | 0 | 0 | 0 | 4 mo | WILD |
| **1009322937** | 2 | SSp-tr2/3 | 670 | 0.436541353 | 0 | 0 | 0 | 0 | 8 mo | WILD |
| **1011080637** | 2 | SSp-tr2/3 | 670 | 0.436541353 | 0.000260201 | 11 | 0.000596052 | 25.19807099 | 8 mo | HOM |
| **1013609713** | 2 | SSp-tr2/3 | 670 | 0.436541353 | 0 | 0 | 0 | 0 | 2 mo | WILD |
| **1010418842** | 1 | SSp-tr2/3 | 670 | 0.436541353 | 0.000391798 | 19 | 0.000897505 | 43.52394081 | 8 mo | HOM |
| **1013288213** | 2 | SSp-tr2/3 | 670 | 0.436541353 | 2.04E-05 | 2 | 4.68E-05 | 4.581467454 | 4 mo | HOM |
| **1008976060** | 2 | SSp-tr2/3 | 670 | 0.436541353 | 0 | 0 | 0 | 0 | 8 mo | WILD |
| **1013446243** | 1 | SSp-tr2/3 | 670 | 0.436541353 | 0.000192873 | 7 | 0.00044182 | 16.03513609 | 4 mo | HOM |
| **1013288951** | 2 | SSp-tr2/3 | 670 | 0.436541353 | 0.000131759 | 3 | 0.000301826 | 6.87220118 | 4 mo | HOM |
| **1013978665** | 1 | RSPagl1 | 671 | 0.522629022 | 3.43E-05 | 3 | 6.57E-05 | 5.740209357 | 2 mo | HOM |
| **1013445609** | 1 | RSPagl1 | 671 | 0.522629022 | 6.74E-05 | 4 | 0.000128904 | 7.653612476 | 4 mo | HOM |
| **1013609015** | 1 | RSPagl1 | 671 | 0.522629022 | 0.000541256 | 11 | 0.00103564 | 21.04743431 | 4 mo | HOM |
| **1013608106** | 2 | RSPagl1 | 671 | 0.522629022 | 0 | 0 | 0 | 0 | 4 mo | HOM |
| **1011937543** | 1 | RSPagl1 | 671 | 0.522629022 | 0.001520767 | 11 | 0.00290984 | 21.04743431 | 8 mo | HOM |
| **1010419538** | 2 | RSPagl1 | 671 | 0.522629022 | 9.61E-05 | 6 | 0.000183855 | 11.48041871 | 4 mo | WILD |
| **1010419187** | 1 | RSPagl1 | 671 | 0.522629022 | 0.006523251 | 33 | 0.012481608 | 63.14230293 | 8 mo | HOM |
| **1013978366** | 1 | RSPagl1 | 671 | 0.522629022 | 2.74E-06 | 0 | 5.24E-06 | 0 | 2 mo | HOM |
| **1014416155** | 2 | RSPagl1 | 671 | 0.522629022 | 0 | 0 | 0 | 0 | 2 mo | HOM |
| **1013978065** | 2 | RSPagl1 | 671 | 0.522629022 | 8.63E-09 | 0 | 1.65E-08 | 0 | 2 mo | HOM |
| **1009934102** | 2 | RSPagl1 | 671 | 0.522629022 | 7.45E-05 | 3 | 0.000142619 | 5.740209357 | 4 mo | WILD |
| **1009322937** | 2 | RSPagl1 | 671 | 0.522629022 | 1.56E-05 | 1 | 2.98E-05 | 1.913403119 | 8 mo | WILD |
| **1011080637** | 2 | RSPagl1 | 671 | 0.522629022 | 0.000928609 | 23 | 0.001776803 | 44.00827174 | 8 mo | HOM |
| **1013609713** | 2 | RSPagl1 | 671 | 0.522629022 | 3.78E-05 | 2 | 7.23E-05 | 3.826806238 | 2 mo | WILD |
| **1010418842** | 1 | RSPagl1 | 671 | 0.522629022 | 0.00068167 | 25 | 0.00130431 | 47.83507797 | 8 mo | HOM |
| **1013288213** | 2 | RSPagl1 | 671 | 0.522629022 | 0.000107816 | 5 | 0.000206296 | 9.567015595 | 4 mo | HOM |
| **1008976060** | 2 | RSPagl1 | 671 | 0.522629022 | 0.000292816 | 11 | 0.000560275 | 21.04743431 | 8 mo | WILD |
| **1013446243** | 1 | RSPagl1 | 671 | 0.522629022 | 0.000335883 | 6 | 0.000642679 | 11.48041871 | 4 mo | HOM |
| **1013288951** | 2 | RSPagl1 | 671 | 0.522629022 | 0.001267123 | 14 | 0.002424516 | 26.78764367 | 4 mo | HOM |
| **1013978665** | 1 | CP | 672 | 26.8051883 | 3.05E-07 | 0 | 1.14E-08 | 0 | 2 mo | HOM |
| **1013445609** | 1 | CP | 672 | 26.8051883 | 0.000235332 | 18 | 8.78E-06 | 0.671511791 | 4 mo | HOM |
| **1013609015** | 1 | CP | 672 | 26.8051883 | 0.002482428 | 74 | 9.26E-05 | 2.760659585 | 4 mo | HOM |
| **1013608106** | 2 | CP | 672 | 26.8051883 | 0 | 0 | 0 | 0 | 4 mo | HOM |
| **1011937543** | 1 | CP | 672 | 26.8051883 | 0.01161648 | 284 | 0.000433367 | 10.59496381 | 8 mo | HOM |
| **1010419538** | 2 | CP | 672 | 26.8051883 | 0 | 0 | 0 | 0 | 4 mo | WILD |

| | | | | | | | | | | |
|---|---|---|---|---|---|---|---|---|---|---|
| **1010419187** | 1 | CP | 672 | 26.8051883 | 0.009902259 | 413 | 0.000369416 | 15.40746498 | 8 mo | HOM |
| **1013978366** | 1 | CP | 672 | 26.8051883 | 2.81E-07 | 0 | 1.05E-08 | 0 | 2 mo | HOM |
| **1014416155** | 2 | CP | 672 | 26.8051883 | 0 | 0 | 0 | 0 | 2 mo | HOM |
| **1013978065** | 2 | CP | 672 | 26.8051883 | 0 | 0 | 0 | 0 | 2 mo | HOM |
| **1009934102** | 2 | CP | 672 | 26.8051883 | 0 | 0 | 0 | 0 | 4 mo | WILD |
| **1009322937** | 2 | CP | 672 | 26.8051883 | 1.37E-05 | 3 | 5.11E-07 | 0.111918632 | 8 mo | WILD |
| **1011080637** | 2 | CP | 672 | 26.8051883 | 0.005699317 | 333 | 0.00021262 | 12.42296813 | 8 mo | HOM |
| **1013609713** | 2 | CP | 672 | 26.8051883 | 0 | 0 | 0 | 0 | 2 mo | WILD |
| **1010418842** | 1 | CP | 672 | 26.8051883 | 0.015405613 | 562 | 0.000574725 | 20.96609036 | 8 mo | HOM |
| **1013288213** | 2 | CP | 672 | 26.8051883 | 0.000157199 | 19 | 5.86E-06 | 0.708818001 | 4 mo | HOM |
| **1008976060** | 2 | CP | 672 | 26.8051883 | 0.000599894 | 56 | 2.24E-05 | 2.089147794 | 8 mo | WILD |
| **1013446243** | 1 | CP | 672 | 26.8051883 | 0.000745304 | 50 | 2.78E-05 | 1.86531053 | 4 mo | HOM |
| **1013288951** | 2 | CP | 672 | 26.8051883 | 0.000918342 | 55 | 3.43E-05 | 2.051841583 | 4 mo | HOM |
| **1013978665** | 1 | mp | 673 | 0.032701402 | 0 | 0 | 0 | 0 | 2 mo | HOM |
| **1013445609** | 1 | mp | 673 | 0.032701402 | 0 | 0 | 0 | 0 | 4 mo | HOM |
| **1013609015** | 1 | mp | 673 | 0.032701402 | 0 | 0 | 0 | 0 | 4 mo | HOM |
| **1013608106** | 2 | mp | 673 | 0.032701402 | 0 | 0 | 0 | 0 | 4 mo | HOM |
| **1011937543** | 1 | mp | 673 | 0.032701402 | 2.86E-06 | 0 | 8.76E-05 | 0 | 8 mo | HOM |
| **1010419538** | 2 | mp | 673 | 0.032701402 | 0 | 0 | 0 | 0 | 4 mo | WILD |
| **1010419187** | 1 | mp | 673 | 0.032701402 | 0 | 0 | 0 | 0 | 8 mo | HOM |
| **1013978366** | 1 | mp | 673 | 0.032701402 | 0 | 0 | 0 | 0 | 2 mo | HOM |
| **1014416155** | 2 | mp | 673 | 0.032701402 | 0 | 0 | 0 | 0 | 2 mo | HOM |
| **1013978065** | 2 | mp | 673 | 0.032701402 | 0 | 0 | 0 | 0 | 2 mo | HOM |
| **1009934102** | 2 | mp | 673 | 0.032701402 | 0 | 0 | 0 | 0 | 4 mo | WILD |
| **1009322937** | 2 | mp | 673 | 0.032701402 | 0 | 0 | 0 | 0 | 8 mo | WILD |
| **1011080637** | 2 | mp | 673 | 0.032701402 | 0 | 0 | 0 | 0 | 8 mo | HOM |
| **1013609713** | 2 | mp | 673 | 0.032701402 | 0 | 0 | 0 | 0 | 2 mo | WILD |
| **1010418842** | 1 | mp | 673 | 0.032701402 | 0 | 0 | 0 | 0 | 8 mo | HOM |
| **1013288213** | 2 | mp | 673 | 0.032701402 | 0 | 0 | 0 | 0 | 4 mo | HOM |
| **1008976060** | 2 | mp | 673 | 0.032701402 | 0 | 0 | 0 | 0 | 8 mo | WILD |
| **1013446243** | 1 | mp | 673 | 0.032701402 | 0 | 0 | 0 | 0 | 4 mo | HOM |
| **1013288951** | 2 | mp | 673 | 0.032701402 | 0 | 0 | 0 | 0 | 4 mo | HOM |
| **1013978665** | 1 | Alv6a | 675 | 0.173003634 | 0 | 0 | 0 | 0 | 2 mo | HOM |
| **1013445609** | 1 | Alv6a | 675 | 0.173003634 | 7.84E-07 | 0 | 4.53E-06 | 0 | 4 mo | HOM |
| **1013609015** | 1 | Alv6a | 675 | 0.173003634 | 0 | 0 | 0 | 0 | 4 mo | HOM |
| **1013608106** | 2 | Alv6a | 675 | 0.173003634 | 0 | 0 | 0 | 0 | 4 mo | HOM |
| **1011937543** | 1 | Alv6a | 675 | 0.173003634 | 0.000196392 | 4 | 0.001135189 | 23.12090166 | 8 mo | HOM |
| **1010419538** | 2 | Alv6a | 675 | 0.173003634 | 0 | 0 | 0 | 0 | 4 mo | WILD |
| **1010419187** | 1 | Alv6a | 675 | 0.173003634 | 0.000285641 | 10 | 0.001651072 | 57.80225416 | 8 mo | HOM |
| **1013978366** | 1 | Alv6a | 675 | 0.173003634 | 0 | 0 | 0 | 0 | 2 mo | HOM |
| **1014416155** | 2 | Alv6a | 675 | 0.173003634 | 0 | 0 | 0 | 0 | 2 mo | HOM |
| **1013978065** | 2 | Alv6a | 675 | 0.173003634 | 0 | 0 | 0 | 0 | 2 mo | HOM |
| **1009934102** | 2 | Alv6a | 675 | 0.173003634 | 0 | 0 | 0 | 0 | 4 mo | WILD |
| **1009322937** | 2 | Alv6a | 675 | 0.173003634 | 0 | 0 | 0 | 0 | 8 mo | WILD |
| **1011080637** | 2 | Alv6a | 675 | 0.173003634 | 9.12E-05 | 4 | 0.000526888 | 23.12090166 | 8 mo | HOM |
| **1013609713** | 2 | Alv6a | 675 | 0.173003634 | 0 | 0 | 0 | 0 | 2 mo | WILD |
| **1010418842** | 1 | Alv6a | 675 | 0.173003634 | 0.000177968 | 9 | 0.001028697 | 52.02202874 | 8 mo | HOM |
| **1013288213** | 2 | Alv6a | 675 | 0.173003634 | 2.87E-06 | 0 | 1.66E-05 | 0 | 4 mo | HOM |
| **1008976060** | 2 | Alv6a | 675 | 0.173003634 | 1.18E-05 | 1 | 6.83E-05 | 5.780225416 | 8 mo | WILD |
| **1013446243** | 1 | Alv6a | 675 | 0.173003634 | 4.77E-06 | 0 | 2.76E-05 | 0 | 4 mo | HOM |
| **1013288951** | 2 | Alv6a | 675 | 0.173003634 | 0 | 0 | 0 | 0 | 4 mo | HOM |
| **1013978665** | 1 | VISC | 677 | 2.420575402 | 4.87E-06 | 1 | 2.01E-06 | 0.413124912 | 2 mo | HOM |
| **1013445609** | 1 | VISC | 677 | 2.420575402 | 0.000270617 | 16 | 0.000111798 | 6.609998592 | 4 mo | HOM |
| **1013609015** | 1 | VISC | 677 | 2.420575402 | 0.000494156 | 22 | 0.000204148 | 9.088748065 | 4 mo | HOM |
| **1013608106** | 2 | VISC | 677 | 2.420575402 | 0 | 0 | 0 | 0 | 4 mo | HOM |
| **1011937543** | 1 | VISC | 677 | 2.420575402 | 0.003676125 | 112 | 0.001518699 | 46.26999015 | 8 mo | HOM |
| **1010419538** | 2 | VISC | 677 | 2.420575402 | 0 | 0 | 0 | 0 | 4 mo | WILD |
| **1010419187** | 1 | VISC | 677 | 2.420575402 | 0.002423508 | 111 | 0.001001212 | 45.85686523 | 8 mo | HOM |
| **1013978366** | 1 | VISC | 677 | 2.420575402 | 6.77E-06 | 1 | 2.8E-06 | 0.413124912 | 2 mo | HOM |

| | | | | | | | | | | |
|---|---|---|---|---|---|---|---|---|---|---|
| **1014416155** | 2 | VISC | 677 | 2.420575402 | 0 | 0 | 0 | 0 | 2 mo | HOM |
| **1013978065** | 2 | VISC | 677 | 2.420575402 | 0 | 0 | 0 | 0 | 2 mo | HOM |
| **1009934102** | 2 | VISC | 677 | 2.420575402 | 0 | 0 | 0 | 0 | 4 mo | WILD |
| **1009322937** | 2 | VISC | 677 | 2.420575402 | 1.7E-06 | 0 | 7.01E-07 | 0 | 8 mo | WILD |
| **1011080637** | 2 | VISC | 677 | 2.420575402 | 0.001481819 | 79 | 0.000612176 | 32.63686805 | 8 mo | HOM |
| **1013609713** | 2 | VISC | 677 | 2.420575402 | 6.41E-07 | 0 | 2.65E-07 | 0 | 2 mo | WILD |
| **1010418842** | 1 | VISC | 677 | 2.420575402 | 0.002302385 | 98 | 0.000951172 | 40.48624138 | 8 mo | HOM |
| **1013288213** | 2 | VISC | 677 | 2.420575402 | 8.85E-05 | 9 | 3.65E-05 | 3.718124208 | 4 mo | HOM |
| **1008976060** | 2 | VISC | 677 | 2.420575402 | 1.29E-05 | 2 | 5.35E-06 | 0.826249824 | 8 mo | WILD |
| **1013446243** | 1 | VISC | 677 | 2.420575402 | 0.000584877 | 30 | 0.000241627 | 12.39374736 | 4 mo | HOM |
| **1013288951** | 2 | VISC | 677 | 2.420575402 | 0.000593581 | 35 | 0.000245223 | 14.45937192 | 4 mo | HOM |
| **1013978665** | 1 | AUDd4 | 678 | 0.136513059 | 0 | 0 | 0 | 0 | 2 mo | HOM |
| **1013445609** | 1 | AUDd4 | 678 | 0.136513059 | 2.13E-05 | 1 | 0.00015601 | 7.325306501 | 4 mo | HOM |
| **1013609015** | 1 | AUDd4 | 678 | 0.136513059 | 1.42E-05 | 1 | 0.000104315 | 7.325306501 | 4 mo | HOM |
| **1013608106** | 2 | AUDd4 | 678 | 0.136513059 | 0 | 0 | 0 | 0 | 4 mo | HOM |
| **1011937543** | 1 | AUDd4 | 678 | 0.136513059 | 7.98E-05 | 1 | 0.00058488 | 7.325306501 | 8 mo | HOM |
| **1010419538** | 2 | AUDd4 | 678 | 0.136513059 | 0 | 0 | 0 | 0 | 4 mo | WILD |
| **1010419187** | 1 | AUDd4 | 678 | 0.136513059 | 0.000135042 | 5 | 0.000989224 | 36.6265325 | 8 mo | HOM |
| **1013978366** | 1 | AUDd4 | 678 | 0.136513059 | 0 | 0 | 0 | 0 | 2 mo | HOM |
| **1014416155** | 2 | AUDd4 | 678 | 0.136513059 | 0 | 0 | 0 | 0 | 2 mo | HOM |
| **1013978065** | 2 | AUDd4 | 678 | 0.136513059 | 0 | 0 | 0 | 0 | 2 mo | HOM |
| **1009934102** | 2 | AUDd4 | 678 | 0.136513059 | 0 | 0 | 0 | 0 | 4 mo | WILD |
| **1009322937** | 2 | AUDd4 | 678 | 0.136513059 | 0 | 0 | 0 | 0 | 8 mo | WILD |
| **1011080637** | 2 | AUDd4 | 678 | 0.136513059 | 0.000124836 | 3 | 0.000914462 | 21.9759195 | 8 mo | HOM |
| **1013609713** | 2 | AUDd4 | 678 | 0.136513059 | 0 | 0 | 0 | 0 | 2 mo | WILD |
| **1010418842** | 1 | AUDd4 | 678 | 0.136513059 | 0.000138616 | 5 | 0.001015407 | 36.6265325 | 8 mo | HOM |
| **1013288213** | 2 | AUDd4 | 678 | 0.136513059 | 4.34E-05 | 3 | 0.000317588 | 21.9759195 | 4 mo | HOM |
| **1008976060** | 2 | AUDd4 | 678 | 0.136513059 | 0 | 0 | 0 | 0 | 8 mo | WILD |
| **1013446243** | 1 | AUDd4 | 678 | 0.136513059 | 3.94E-06 | 0 | 2.89E-05 | 0 | 4 mo | HOM |
| **1013288951** | 2 | AUDd4 | 678 | 0.136513059 | 4.9E-07 | 0 | 3.59E-06 | 0 | 4 mo | HOM |
| **1013978665** | 1 | CS | 679 | 0.619836946 | 0 | 0 | 0 | 0 | 2 mo | HOM |
| **1013445609** | 1 | CS | 679 | 0.619836946 | 0 | 0 | 0 | 0 | 4 mo | HOM |
| **1013609015** | 1 | CS | 679 | 0.619836946 | 0 | 0 | 0 | 0 | 4 mo | HOM |
| **1013608106** | 2 | CS | 679 | 0.619836946 | 0 | 0 | 0 | 0 | 4 mo | HOM |
| **1011937543** | 1 | CS | 679 | 0.619836946 | 0.000149571 | 0 | 0.000241306 | 0 | 8 mo | HOM |
| **1010419538** | 2 | CS | 679 | 0.619836946 | 0 | 0 | 0 | 0 | 4 mo | WILD |
| **1010419187** | 1 | CS | 679 | 0.619836946 | 0 | 0 | 0 | 0 | 8 mo | HOM |
| **1013978366** | 1 | CS | 679 | 0.619836946 | 0 | 0 | 0 | 0 | 2 mo | HOM |
| **1014416155** | 2 | CS | 679 | 0.619836946 | 0 | 0 | 0 | 0 | 2 mo | HOM |
| **1013978065** | 2 | CS | 679 | 0.619836946 | 0 | 0 | 0 | 0 | 2 mo | HOM |
| **1009934102** | 2 | CS | 679 | 0.619836946 | 0 | 0 | 0 | 0 | 4 mo | WILD |
| **1009322937** | 2 | CS | 679 | 0.619836946 | 0 | 0 | 0 | 0 | 8 mo | WILD |
| **1011080637** | 2 | CS | 679 | 0.619836946 | 0 | 0 | 0 | 0 | 8 mo | HOM |
| **1013609713** | 2 | CS | 679 | 0.619836946 | 0 | 0 | 0 | 0 | 2 mo | WILD |
| **1010418842** | 1 | CS | 679 | 0.619836946 | 5.98E-06 | 1 | 9.64E-06 | 1.613327514 | 8 mo | HOM |
| **1013288213** | 2 | CS | 679 | 0.619836946 | 0 | 0 | 0 | 0 | 4 mo | HOM |
| **1008976060** | 2 | CS | 679 | 0.619836946 | 0 | 0 | 0 | 0 | 8 mo | WILD |
| **1013446243** | 1 | CS | 679 | 0.619836946 | 0 | 0 | 0 | 0 | 4 mo | HOM |
| **1013288951** | 2 | CS | 679 | 0.619836946 | 0 | 0 | 0 | 0 | 4 mo | HOM |
| **1013978665** | 1 | ORBvl6b | 680 | 0.009921067 | 0 | 0 | 0 | 0 | 2 mo | HOM |
| **1013445609** | 1 | ORBvl6b | 680 | 0.009921067 | 0 | 0 | 0 | 0 | 4 mo | HOM |
| **1013609015** | 1 | ORBvl6b | 680 | 0.009921067 | 0 | 0 | 0 | 0 | 4 mo | HOM |
| **1013608106** | 2 | ORBvl6b | 680 | 0.009921067 | 0 | 0 | 0 | 0 | 4 mo | HOM |
| **1011937543** | 1 | ORBvl6b | 680 | 0.009921067 | 4.88E-06 | 0 | 0.00049213 | 0 | 8 mo | HOM |
| **1010419538** | 2 | ORBvl6b | 680 | 0.009921067 | 0 | 0 | 0 | 0 | 4 mo | WILD |
| **1010419187** | 1 | ORBvl6b | 680 | 0.009921067 | 2.37E-06 | 0 | 0.000238958 | 0 | 8 mo | HOM |
| **1013978366** | 1 | ORBvl6b | 680 | 0.009921067 | 0 | 0 | 0 | 0 | 2 mo | HOM |
| **1014416155** | 2 | ORBvl6b | 680 | 0.009921067 | 0 | 0 | 0 | 0 | 2 mo | HOM |
| **1013978065** | 2 | ORBvl6b | 680 | 0.009921067 | 0 | 0 | 0 | 0 | 2 mo | HOM |

| | | | | | | | | | | |
|---|---|---|---|---|---|---|---|---|---|---|
| **1009934102** | 2 | ORBvl6b | 680 | 0.009921067 | 0 | 0 | 0 | 0 | 4 mo | WILD |
| **1009322937** | 2 | ORBvl6b | 680 | 0.009921067 | 0 | 0 | 0 | 0 | 8 mo | WILD |
| **1011080637** | 2 | ORBvl6b | 680 | 0.009921067 | 1.61E-06 | 0 | 0.000162703 | 0 | 8 mo | HOM |
| **1013609713** | 2 | ORBvl6b | 680 | 0.009921067 | 0 | 0 | 0 | 0 | 2 mo | WILD |
| **1010418842** | 1 | ORBvl6b | 680 | 0.009921067 | 0 | 0 | 0 | 0 | 8 mo | HOM |
| **1013288213** | 2 | ORBvl6b | 680 | 0.009921067 | 0 | 0 | 0 | 0 | 4 mo | HOM |
| **1008976060** | 2 | ORBvl6b | 680 | 0.009921067 | 0 | 0 | 0 | 0 | 8 mo | WILD |
| **1013446243** | 1 | ORBvl6b | 680 | 0.009921067 | 0 | 0 | 0 | 0 | 4 mo | HOM |
| **1013288951** | 2 | ORBvl6b | 680 | 0.009921067 | 0 | 0 | 0 | 0 | 4 mo | HOM |
| **1013978665** | 1 | mtg | 681 | 0.067790863 | 0 | 0 | 0 | 0 | 2 mo | HOM |
| **1013445609** | 1 | mtg | 681 | 0.067790863 | 0 | 0 | 0 | 0 | 4 mo | HOM |
| **1013609015** | 1 | mtg | 681 | 0.067790863 | 0 | 0 | 0 | 0 | 4 mo | HOM |
| **1013608106** | 2 | mtg | 681 | 0.067790863 | 0 | 0 | 0 | 0 | 4 mo | HOM |
| **1011937543** | 1 | mtg | 681 | 0.067790863 | 0 | 0 | 0 | 0 | 8 mo | HOM |
| **1010419538** | 2 | mtg | 681 | 0.067790863 | 0 | 0 | 0 | 0 | 4 mo | WILD |
| **1010419187** | 1 | mtg | 681 | 0.067790863 | 0 | 0 | 0 | 0 | 8 mo | HOM |
| **1013978366** | 1 | mtg | 681 | 0.067790863 | 0 | 0 | 0 | 0 | 2 mo | HOM |
| **1014416155** | 2 | mtg | 681 | 0.067790863 | 0 | 0 | 0 | 0 | 2 mo | HOM |
| **1013978065** | 2 | mtg | 681 | 0.067790863 | 0 | 0 | 0 | 0 | 2 mo | HOM |
| **1009934102** | 2 | mtg | 681 | 0.067790863 | 0 | 0 | 0 | 0 | 4 mo | WILD |
| **1009322937** | 2 | mtg | 681 | 0.067790863 | 0 | 0 | 0 | 0 | 8 mo | WILD |
| **1011080637** | 2 | mtg | 681 | 0.067790863 | 0 | 0 | 0 | 0 | 8 mo | HOM |
| **1013609713** | 2 | mtg | 681 | 0.067790863 | 0 | 0 | 0 | 0 | 2 mo | WILD |
| **1010418842** | 1 | mtg | 681 | 0.067790863 | 1.22E-05 | 0 | 0.000180507 | 0 | 8 mo | HOM |
| **1013288213** | 2 | mtg | 681 | 0.067790863 | 0 | 0 | 0 | 0 | 4 mo | HOM |
| **1008976060** | 2 | mtg | 681 | 0.067790863 | 0 | 0 | 0 | 0 | 8 mo | WILD |
| **1013446243** | 1 | mtg | 681 | 0.067790863 | 0 | 0 | 0 | 0 | 4 mo | HOM |
| **1013288951** | 2 | mtg | 681 | 0.067790863 | 0 | 0 | 0 | 0 | 4 mo | HOM |
| **1013978665** | 1 | VM | 685 | 0.956800891 | 0 | 0 | 0 | 0 | 2 mo | HOM |
| **1013445609** | 1 | VM | 685 | 0.956800891 | 0 | 0 | 0 | 0 | 4 mo | HOM |
| **1013609015** | 1 | VM | 685 | 0.956800891 | 0 | 0 | 0 | 0 | 4 mo | HOM |
| **1013608106** | 2 | VM | 685 | 0.956800891 | 0 | 0 | 0 | 0 | 4 mo | HOM |
| **1011937543** | 1 | VM | 685 | 0.956800891 | 2.14E-07 | 0 | 2.23E-07 | 0 | 8 mo | HOM |
| **1010419538** | 2 | VM | 685 | 0.956800891 | 0 | 0 | 0 | 0 | 4 mo | WILD |
| **1010419187** | 1 | VM | 685 | 0.956800891 | 0.000130288 | 4 | 0.00013617 | 4.180598113 | 8 mo | HOM |
| **1013978366** | 1 | VM | 685 | 0.956800891 | 0 | 0 | 0 | 0 | 2 mo | HOM |
| **1014416155** | 2 | VM | 685 | 0.956800891 | 0 | 0 | 0 | 0 | 2 mo | HOM |
| **1013978065** | 2 | VM | 685 | 0.956800891 | 0 | 0 | 0 | 0 | 2 mo | HOM |
| **1009934102** | 2 | VM | 685 | 0.956800891 | 0 | 0 | 0 | 0 | 4 mo | WILD |
| **1009322937** | 2 | VM | 685 | 0.956800891 | 0 | 0 | 0 | 0 | 8 mo | WILD |
| **1011080637** | 2 | VM | 685 | 0.956800891 | 0.000104898 | 4 | 0.000109634 | 4.180598113 | 8 mo | HOM |
| **1013609713** | 2 | VM | 685 | 0.956800891 | 0 | 0 | 0 | 0 | 2 mo | WILD |
| **1010418842** | 1 | VM | 685 | 0.956800891 | 0.000192179 | 6 | 0.000200856 | 6.270897169 | 8 mo | HOM |
| **1013288213** | 2 | VM | 685 | 0.956800891 | 0 | 0 | 0 | 0 | 4 mo | HOM |
| **1008976060** | 2 | VM | 685 | 0.956800891 | 1.4E-05 | 2 | 1.47E-05 | 2.090299056 | 8 mo | WILD |
| **1013446243** | 1 | VM | 685 | 0.956800891 | 0 | 0 | 0 | 0 | 4 mo | HOM |
| **1013288951** | 2 | VM | 685 | 0.956800891 | 0 | 0 | 0 | 0 | 4 mo | HOM |
| **1013978665** | 1 | RSPv5 | 687 | 1.729490202 | 0 | 0 | 0 | 0 | 2 mo | HOM |
| **1013445609** | 1 | RSPv5 | 687 | 1.729490202 | 0.000127247 | 7 | 7.36E-05 | 4.047435478 | 4 mo | HOM |
| **1013609015** | 1 | RSPv5 | 687 | 1.729490202 | 0.000787339 | 19 | 0.000455243 | 10.9858963 | 4 mo | HOM |
| **1013608106** | 2 | RSPv5 | 687 | 1.729490202 | 0 | 0 | 0 | 0 | 4 mo | HOM |
| **1011937543** | 1 | RSPv5 | 687 | 1.729490202 | 0.001916212 | 34 | 0.001107964 | 19.65897232 | 8 mo | HOM |
| **1010419538** | 2 | RSPv5 | 687 | 1.722305155 | 0 | 0 | 0 | 0 | 4 mo | WILD |
| **1010419187** | 1 | RSPv5 | 687 | 1.729490202 | 0.003167121 | 75 | 0.001831246 | 43.36538012 | 8 mo | HOM |
| **1013978366** | 1 | RSPv5 | 687 | 1.729490202 | 0 | 0 | 0 | 0 | 2 mo | HOM |
| **1014416155** | 2 | RSPv5 | 687 | 1.729490202 | 0 | 0 | 0 | 0 | 2 mo | HOM |
| **1013978065** | 2 | RSPv5 | 687 | 1.729490202 | 0 | 0 | 0 | 0 | 2 mo | HOM |
| **1009934102** | 2 | RSPv5 | 687 | 1.729490202 | 0 | 0 | 0 | 0 | 4 mo | WILD |
| **1009322937** | 2 | RSPv5 | 687 | 1.729490202 | 0 | 0 | 0 | 0 | 8 mo | WILD |

| | | | | | | | | | | |
|---|---|---|---|---|---|---|---|---|---|---|
| **1011080637** | 2 | RSPv5 | 687 | 1.729490202 | 0.002138934 | 53 | 0.001236743 | 30.64486862 | 8 mo | HOM |
| **1013609713** | 2 | RSPv5 | 687 | 1.729490202 | 0 | 0 | 0 | 0 | 2 mo | WILD |
| **1010418842** | 1 | RSPv5 | 687 | 1.729490202 | 0.002600405 | 76 | 0.001503567 | 43.94358518 | 8 mo | HOM |
| **1013288213** | 2 | RSPv5 | 687 | 1.729490202 | 4.68E-05 | 6 | 2.71E-05 | 3.469230409 | 4 mo | HOM |
| **1008976060** | 2 | RSPv5 | 687 | 1.729490202 | 8.69E-06 | 1 | 5.02E-06 | 0.578205068 | 8 mo | WILD |
| **1013446243** | 1 | RSPv5 | 687 | 1.729490202 | 0.0001235 | 9 | 7.14E-05 | 5.203845614 | 4 mo | HOM |
| **1013288951** | 2 | RSPv5 | 687 | 1.729490202 | 0.000160339 | 10 | 9.27E-05 | 5.782050682 | 4 mo | HOM |
| **1013978665** | 1 | CTX | 688 | 224.6638039 | 0.006374763 | 298 | 2.84E-05 | 1.326426397 | 2 mo | HOM |
| **1013445609** | 1 | CTX | 688 | 224.8503707 | 0.015841 | 1105 | 7.05E-05 | 4.914379267 | 4 mo | HOM |
| **1013609015** | 1 | CTX | 688 | 222.6177636 | 0.06472006 | 1939 | 0.000290723 | 8.709996762 | 4 mo | HOM |
| **1013608106** | 2 | CTX | 688 | 224.5841729 | 8.41E-06 | 1 | 3.74E-08 | 0.004452674 | 4 mo | HOM |
| **1011937543** | 1 | CTX | 688 | 225.2590467 | 0.32984388 | 5123 | 0.001464287 | 22.74270479 | 8 mo | HOM |
| **1010419538** | 2 | CTX | 688 | 225.6290234 | 0.002431227 | 145 | 1.08E-05 | 0.64264782 | 4 mo | WILD |
| **1010419187** | 1 | CTX | 688 | 221.7915587 | 0.36547322 | 9681 | 0.001647823 | 43.64909133 | 8 mo | HOM |
| **1013978366** | 1 | CTX | 688 | 226.9536336 | 0.009578283 | 350 | 4.22E-05 | 1.542165219 | 2 mo | HOM |
| **1014416155** | 2 | CTX | 688 | 225.7785784 | 0.000107661 | 11 | 4.77E-07 | 0.0487203 | 2 mo | HOM |
| **1013978065** | 2 | CTX | 688 | 227.409708 | 0.001779068 | 129 | 7.82E-06 | 0.567258105 | 2 mo | HOM |
| **1009934102** | 2 | CTX | 688 | 223.8720507 | 0.004718414 | 246 | 2.11E-05 | 1.098841947 | 4 mo | WILD |
| **1009322937** | 2 | CTX | 688 | 221.448553 | 0.003667079 | 208 | 1.66E-05 | 0.93927008 | 8 mo | WILD |
| **1011080637** | 2 | CTX | 688 | 221.0611353 | 0.186124404 | 6982 | 0.000841959 | 31.58402308 | 8 mo | HOM |
| **1013609713** | 2 | CTX | 688 | 228.2269276 | 0.001039553 | 75 | 4.55E-06 | 0.328620294 | 2 mo | WILD |
| **1010418842** | 1 | CTX | 688 | 223.6515604 | 0.286973031 | 8727 | 0.001283126 | 39.02051917 | 8 mo | HOM |
| **1013288213** | 2 | CTX | 688 | 225.317652 | 0.058392276 | 1084 | 0.000259155 | 4.810985693 | 4 mo | HOM |
| **1008976060** | 2 | CTX | 688 | 226.4997982 | 0.017219831 | 696 | 7.6E-05 | 3.07285042 | 8 mo | WILD |
| **1013446243** | 1 | CTX | 688 | 224.2535661 | 0.038127603 | 1812 | 0.00017002 | 8.080139066 | 4 mo | HOM |
| **1013288951** | 2 | CTX | 688 | 224.0615291 | 0.071379247 | 2179 | 0.00031857 | 9.725007274 | 4 mo | HOM |
| **1013978665** | 1 | VLPO | 689 | 0.067403498 | 0 | 0 | 0 | 0 | 2 mo | HOM |
| **1013445609** | 1 | VLPO | 689 | 0.067403498 | 0 | 0 | 0 | 0 | 4 mo | HOM |
| **1013609015** | 1 | VLPO | 689 | 0.067403498 | 0 | 0 | 0 | 0 | 4 mo | HOM |
| **1013608106** | 2 | VLPO | 689 | 0.067403498 | 0 | 0 | 0 | 0 | 4 mo | HOM |
| **1011937543** | 1 | VLPO | 689 | 0.067403498 | 4.69E-06 | 0 | 6.96E-05 | 0 | 8 mo | HOM |
| **1010419538** | 2 | VLPO | 689 | 0.067403498 | 0 | 0 | 0 | 0 | 4 mo | WILD |
| **1010419187** | 1 | VLPO | 689 | 0.067403498 | 0 | 0 | 0 | 0 | 8 mo | HOM |
| **1013978366** | 1 | VLPO | 689 | 0.067403498 | 0 | 0 | 0 | 0 | 2 mo | HOM |
| **1014416155** | 2 | VLPO | 689 | 0.067403498 | 0 | 0 | 0 | 0 | 2 mo | HOM |
| **1013978065** | 2 | VLPO | 689 | 0.067403498 | 0 | 0 | 0 | 0 | 2 mo | HOM |
| **1009934102** | 2 | VLPO | 689 | 0.067403498 | 0 | 0 | 0 | 0 | 4 mo | WILD |
| **1009322937** | 2 | VLPO | 689 | 0.067403498 | 0 | 0 | 0 | 0 | 8 mo | WILD |
| **1011080637** | 2 | VLPO | 689 | 0.067403498 | 0 | 0 | 0 | 0 | 8 mo | HOM |
| **1013609713** | 2 | VLPO | 689 | 0.067403498 | 0 | 0 | 0 | 0 | 2 mo | WILD |
| **1010418842** | 1 | VLPO | 689 | 0.067403498 | 2E-06 | 0 | 2.96E-05 | 0 | 8 mo | HOM |
| **1013288213** | 2 | VLPO | 689 | 0.067403498 | 0 | 0 | 0 | 0 | 4 mo | HOM |
| **1008976060** | 2 | VLPO | 689 | 0.067403498 | 0 | 0 | 0 | 0 | 8 mo | WILD |
| **1013446243** | 1 | VLPO | 689 | 0.067403498 | 0 | 0 | 0 | 0 | 4 mo | HOM |
| **1013288951** | 2 | VLPO | 689 | 0.067403498 | 0 | 0 | 0 | 0 | 4 mo | HOM |
| **1013978665** | 1 | mtt | 690 | 0.133471825 | 0 | 0 | 0 | 0 | 2 mo | HOM |
| **1013445609** | 1 | mtt | 690 | 0.133471825 | 0 | 0 | 0 | 0 | 4 mo | HOM |
| **1013609015** | 1 | mtt | 690 | 0.133471825 | 0 | 0 | 0 | 0 | 4 mo | HOM |
| **1013608106** | 2 | mtt | 690 | 0.133471825 | 0 | 0 | 0 | 0 | 4 mo | HOM |
| **1011937543** | 1 | mtt | 690 | 0.133471825 | 0 | 0 | 0 | 0 | 8 mo | HOM |
| **1010419538** | 2 | mtt | 690 | 0.133471825 | 0 | 0 | 0 | 0 | 4 mo | WILD |
| **1010419187** | 1 | mtt | 690 | 0.133471825 | 1.29E-10 | 0 | 9.64E-10 | 0 | 8 mo | HOM |
| **1013978366** | 1 | mtt | 690 | 0.133471825 | 0 | 0 | 0 | 0 | 2 mo | HOM |
| **1014416155** | 2 | mtt | 690 | 0.133471825 | 0 | 0 | 0 | 0 | 2 mo | HOM |
| **1013978065** | 2 | mtt | 690 | 0.133471825 | 0 | 0 | 0 | 0 | 2 mo | HOM |
| **1009934102** | 2 | mtt | 690 | 0.133471825 | 0 | 0 | 0 | 0 | 4 mo | WILD |
| **1009322937** | 2 | mtt | 690 | 0.133471825 | 0 | 0 | 0 | 0 | 8 mo | WILD |
| **1011080637** | 2 | mtt | 690 | 0.133471825 | 1.32E-07 | 0 | 9.9E-07 | 0 | 8 mo | HOM |
| **1013609713** | 2 | mtt | 690 | 0.133471825 | 0 | 0 | 0 | 0 | 2 mo | WILD |

| | | | | | | | | | | |
|---|---|---|---|---|---|---|---|---|---|---|
| **1010418842** | 1 | mtt | 690 | 0.133471825 | 0 | 0 | 0 | 0 | 8 mo | HOM |
| **1013288213** | 2 | mtt | 690 | 0.133471825 | 0 | 0 | 0 | 0 | 4 mo | HOM |
| **1008976060** | 2 | mtt | 690 | 0.133471825 | 0 | 0 | 0 | 0 | 8 mo | WILD |
| **1013446243** | 1 | mtt | 690 | 0.133471825 | 0 | 0 | 0 | 0 | 4 mo | HOM |
| **1013288951** | 2 | mtt | 690 | 0.133471825 | 0 | 0 | 0 | 0 | 4 mo | HOM |
| **1013978665** | 1 | PERI5 | 692 | 0.173463097 | 0 | 0 | 0 | 0 | 2 mo | HOM |
| **1013445609** | 1 | PERI5 | 692 | 0.173463097 | 1.24E-05 | 1 | 7.16E-05 | 5.76491495 | 4 mo | HOM |
| **1013609015** | 1 | PERI5 | 692 | 0.173463097 | 3.64E-05 | 4 | 0.000209911 | 23.0596598 | 4 mo | HOM |
| **1013608106** | 2 | PERI5 | 692 | 0.173463097 | 0 | 0 | 0 | 0 | 4 mo | HOM |
| **1011937543** | 1 | PERI5 | 692 | 0.173463097 | 0.000193297 | 8 | 0.001114343 | 46.1193196 | 8 mo | HOM |
| **1010419538** | 2 | PERI5 | 692 | 0.173463097 | 0 | 0 | 0 | 0 | 4 mo | WILD |
| **1010419187** | 1 | PERI5 | 692 | 0.173463097 | 0.000292885 | 10 | 0.001688455 | 57.6491495 | 8 mo | HOM |
| **1013978366** | 1 | PERI5 | 692 | 0.173463097 | 0 | 0 | 0 | 0 | 2 mo | HOM |
| **1014416155** | 2 | PERI5 | 692 | 0.173463097 | 0 | 0 | 0 | 0 | 2 mo | HOM |
| **1013978065** | 2 | PERI5 | 692 | 0.173463097 | 0 | 0 | 0 | 0 | 2 mo | HOM |
| **1009934102** | 2 | PERI5 | 692 | 0.173463097 | 0 | 0 | 0 | 0 | 4 mo | WILD |
| **1009322937** | 2 | PERI5 | 692 | 0.173463097 | 0 | 0 | 0 | 0 | 8 mo | WILD |
| **1011080637** | 2 | PERI5 | 692 | 0.173463097 | 0.000130183 | 6 | 0.000750495 | 34.5894897 | 8 mo | HOM |
| **1013609713** | 2 | PERI5 | 692 | 0.173463097 | 0 | 0 | 0 | 0 | 2 mo | WILD |
| **1010418842** | 1 | PERI5 | 692 | 0.173463097 | 0.000211149 | 11 | 0.001217255 | 63.41406445 | 8 mo | HOM |
| **1013288213** | 2 | PERI5 | 692 | 0.173463097 | 0 | 0 | 0 | 0 | 4 mo | HOM |
| **1008976060** | 2 | PERI5 | 692 | 0.173463097 | 0 | 0 | 0 | 0 | 8 mo | WILD |
| **1013446243** | 1 | PERI5 | 692 | 0.173463097 | 3.98E-05 | 5 | 0.000229471 | 28.82457475 | 4 mo | HOM |
| **1013288951** | 2 | PERI5 | 692 | 0.173463097 | 5.38E-05 | 5 | 0.000310148 | 28.82457475 | 4 mo | HOM |
| **1013978665** | 1 | VMH | 693 | 0.56606125 | 0 | 0 | 0 | 0 | 2 mo | HOM |
| **1013445609** | 1 | VMH | 693 | 0.56606125 | 0 | 0 | 0 | 0 | 4 mo | HOM |
| **1013609015** | 1 | VMH | 693 | 0.56606125 | 0 | 0 | 0 | 0 | 4 mo | HOM |
| **1013608106** | 2 | VMH | 693 | 0.56606125 | 0 | 0 | 0 | 0 | 4 mo | HOM |
| **1011937543** | 1 | VMH | 693 | 0.56606125 | 0.001405773 | 7 | 0.002483429 | 12.36615296 | 8 mo | HOM |
| **1010419538** | 2 | VMH | 693 | 0.56606125 | 0 | 0 | 0 | 0 | 4 mo | WILD |
| **1010419187** | 1 | VMH | 693 | 0.56606125 | 0.000120577 | 5 | 0.00021301 | 8.832966403 | 8 mo | HOM |
| **1013978366** | 1 | VMH | 693 | 0.56606125 | 0 | 0 | 0 | 0 | 2 mo | HOM |
| **1014416155** | 2 | VMH | 693 | 0.56606125 | 0 | 0 | 0 | 0 | 2 mo | HOM |
| **1013978065** | 2 | VMH | 693 | 0.56606125 | 0 | 0 | 0 | 0 | 2 mo | HOM |
| **1009934102** | 2 | VMH | 693 | 0.56606125 | 0 | 0 | 0 | 0 | 4 mo | WILD |
| **1009322937** | 2 | VMH | 693 | 0.56606125 | 0 | 0 | 0 | 0 | 8 mo | WILD |
| **1011080637** | 2 | VMH | 693 | 0.56606125 | 6.57E-05 | 2 | 0.000116085 | 3.533186561 | 8 mo | HOM |
| **1013609713** | 2 | VMH | 693 | 0.56606125 | 0 | 0 | 0 | 0 | 2 mo | WILD |
| **1010418842** | 1 | VMH | 693 | 0.56606125 | 0.00011243 | 6 | 0.000198618 | 10.59955968 | 8 mo | HOM |
| **1013288213** | 2 | VMH | 693 | 0.56606125 | 0 | 0 | 0 | 0 | 4 mo | HOM |
| **1008976060** | 2 | VMH | 693 | 0.56606125 | 0 | 0 | 0 | 0 | 8 mo | WILD |
| **1013446243** | 1 | VMH | 693 | 0.56606125 | 0 | 0 | 0 | 0 | 4 mo | HOM |
| **1013288951** | 2 | VMH | 693 | 0.56606125 | 0 | 0 | 0 | 0 | 4 mo | HOM |
| **1013978665** | 1 | AIv2/3 | 694 | 0.612909718 | 0 | 0 | 0 | 0 | 2 mo | HOM |
| **1013445609** | 1 | AIv2/3 | 694 | 0.612909718 | 1.09E-05 | 0 | 1.77E-05 | 0 | 4 mo | HOM |
| **1013609015** | 1 | AIv2/3 | 694 | 0.612909718 | 3.67E-05 | 3 | 5.99E-05 | 4.894684992 | 4 mo | HOM |
| **1013608106** | 2 | AIv2/3 | 694 | 0.612909718 | 0 | 0 | 0 | 0 | 4 mo | HOM |
| **1011937543** | 1 | AIv2/3 | 694 | 0.612909718 | 0.000469776 | 7 | 0.000766468 | 11.42093165 | 8 mo | HOM |
| **1010419538** | 2 | AIv2/3 | 694 | 0.612909718 | 0 | 0 | 0 | 0 | 4 mo | WILD |
| **1010419187** | 1 | AIv2/3 | 694 | 0.612909718 | 0.000723139 | 30 | 0.001179846 | 48.94684992 | 8 mo | HOM |
| **1013978366** | 1 | AIv2/3 | 694 | 0.612909718 | 5.79E-08 | 0 | 9.44E-08 | 0 | 2 mo | HOM |
| **1014416155** | 2 | AIv2/3 | 694 | 0.612909718 | 0 | 0 | 0 | 0 | 2 mo | HOM |
| **1013978065** | 2 | AIv2/3 | 694 | 0.612909718 | 0 | 0 | 0 | 0 | 2 mo | HOM |
| **1009934102** | 2 | AIv2/3 | 694 | 0.612909718 | 0 | 0 | 0 | 0 | 4 mo | WILD |
| **1009322937** | 2 | AIv2/3 | 694 | 0.612909718 | 1.9E-05 | 1 | 3.1E-05 | 1.631561664 | 8 mo | WILD |
| **1011080637** | 2 | AIv2/3 | 694 | 0.612909718 | 0.000291487 | 14 | 0.000475579 | 22.8418633 | 8 mo | HOM |
| **1013609713** | 2 | AIv2/3 | 694 | 0.612909718 | 0 | 0 | 0 | 0 | 2 mo | WILD |
| **1010418842** | 1 | AIv2/3 | 694 | 0.612909718 | 0.000606 | 27 | 0.000988727 | 44.05216493 | 8 mo | HOM |
| **1013288213** | 2 | AIv2/3 | 694 | 0.612909718 | 1.98E-06 | 0 | 3.24E-06 | 0 | 4 mo | HOM |

| | | | | | | | | | | |
|---|---|---|---|---|---|---|---|---|---|---|
| 1008976060 | 2 | AIv2/3 | 694 | 0.612909718 | 4.82E-07 | 0 | 7.86E-07 | 0 | 8 mo | WILD |
| 1013446243 | 1 | AIv2/3 | 694 | 0.612909718 | 2.35E-05 | 1 | 3.84E-05 | 1.631561664 | 4 mo | HOM |
| 1013288951 | 2 | AIv2/3 | 694 | 0.612909718 | 4.29E-05 | 2 | 6.99E-05 | 3.263123328 | 4 mo | HOM |
| 1013978665 | 1 | CTXpl | 695 | 215.5023612 | 0.006374763 | 298 | 2.96E-05 | 1.382815475 | 2 mo | HOM |
| 1013445609 | 1 | CTXpl | 695 | 215.6732274 | 0.015122576 | 1055 | 7.01E-05 | 4.891659538 | 4 mo | HOM |
| 1013609015 | 1 | CTXpl | 695 | 213.4406203 | 0.062995928 | 1874 | 0.000295145 | 8.779959493 | 4 mo | HOM |
| 1013608106 | 2 | CTXpl | 695 | 215.4070296 | 8.41E-06 | 1 | 3.9E-08 | 0.004642374 | 4 mo | HOM |
| 1011937543 | 1 | CTXpl | 695 | 216.0819034 | 0.314024979 | 4815 | 0.001453268 | 22.28321726 | 8 mo | HOM |
| 1010419538 | 2 | CTXpl | 695 | 216.4518801 | 0.00242732 | 144 | 1.12E-05 | 0.665274887 | 4 mo | WILD |
| 1010419187 | 1 | CTXpl | 695 | 212.6144153 | 0.35211378 | 9165 | 0.001656114 | 43.1062023 | 8 mo | HOM |
| 1013978366 | 1 | CTXpl | 695 | 217.7764902 | 0.009578283 | 350 | 4.4E-05 | 1.607152359 | 2 mo | HOM |
| 1014416155 | 2 | CTXpl | 695 | 216.6014351 | 0.000107661 | 11 | 4.97E-07 | 0.05078452 | 2 mo | HOM |
| 1013978065 | 2 | CTXpl | 695 | 218.2325647 | 0.001779068 | 129 | 8.15E-06 | 0.591112514 | 2 mo | HOM |
| 1009934102 | 2 | CTXpl | 695 | 214.6949074 | 0.004718414 | 246 | 2.2E-05 | 1.145811994 | 4 mo | WILD |
| 1009322937 | 2 | CTXpl | 695 | 212.2714097 | 0.003658867 | 206 | 1.72E-05 | 0.970455702 | 8 mo | WILD |
| 1011080637 | 2 | CTXpl | 695 | 211.883992 | 0.17676234 | 6536 | 0.000834241 | 30.84706842 | 8 mo | HOM |
| 1013609713 | 2 | CTXpl | 695 | 219.0497843 | 0.001039553 | 75 | 4.75E-06 | 0.34238792 | 2 mo | WILD |
| 1010418842 | 1 | CTXpl | 695 | 214.4744171 | 0.270994978 | 8247 | 0.001263531 | 38.45213854 | 8 mo | HOM |
| 1013288213 | 2 | CTXpl | 695 | 216.1405087 | 0.057982886 | 1049 | 0.000268265 | 4.853324379 | 4 mo | HOM |
| 1008976060 | 2 | CTXpl | 695 | 217.3226549 | 0.017128304 | 683 | 7.88E-05 | 3.142792454 | 8 mo | WILD |
| 1013446243 | 1 | CTXpl | 695 | 215.0764228 | 0.036926422 | 1741 | 0.00017169 | 8.094797085 | 4 mo | HOM |
| 1013288951 | 2 | CTXpl | 695 | 214.8843857 | 0.069861677 | 2103 | 0.000325113 | 9.786658034 | 4 mo | HOM |
| 1013978665 | 1 | AUDpo1 | 696 | 0.116093995 | 0 | 0 | 0 | 0 | 2 mo | HOM |
| 1013445609 | 1 | AUDpo1 | 696 | 0.116093995 | 1.95E-05 | 2 | 0.000168084 | 17.22741993 | 4 mo | HOM |
| 1013609015 | 1 | AUDpo1 | 696 | 0.116093995 | 0.000118709 | 3 | 0.001022524 | 25.8411299 | 4 mo | HOM |
| 1013608106 | 2 | AUDpo1 | 696 | 0.116093995 | 0 | 0 | 0 | 0 | 4 mo | HOM |
| 1011937543 | 1 | AUDpo1 | 696 | 0.116093995 | 0.000439421 | 6 | 0.003785043 | 51.6822598 | 8 mo | HOM |
| 1010419538 | 2 | AUDpo1 | 696 | 0.116093995 | 0 | 0 | 0 | 0 | 4 mo | WILD |
| 1010419187 | 1 | AUDpo1 | 696 | 0.116093995 | 0.000285858 | 5 | 0.0024623 | 43.06854983 | 8 mo | HOM |
| 1013978366 | 1 | AUDpo1 | 696 | 0.116093995 | 0 | 0 | 0 | 0 | 2 mo | HOM |
| 1014416155 | 2 | AUDpo1 | 696 | 0.116093995 | 0 | 0 | 0 | 0 | 2 mo | HOM |
| 1013978065 | 2 | AUDpo1 | 696 | 0.116093995 | 0 | 0 | 0 | 0 | 2 mo | HOM |
| 1009934102 | 2 | AUDpo1 | 696 | 0.116093995 | 0 | 0 | 0 | 0 | 4 mo | WILD |
| 1009322937 | 2 | AUDpo1 | 696 | 0.116093995 | 0 | 0 | 0 | 0 | 8 mo | WILD |
| 1011080637 | 2 | AUDpo1 | 696 | 0.116093995 | 0.00011195 | 3 | 0.000964306 | 25.8411299 | 8 mo | HOM |
| 1013609713 | 2 | AUDpo1 | 696 | 0.116093995 | 0 | 0 | 0 | 0 | 2 mo | WILD |
| 1010418842 | 1 | AUDpo1 | 696 | 0.116093995 | 0.00040528 | 8 | 0.003490961 | 68.90967973 | 8 mo | HOM |
| 1013288213 | 2 | AUDpo1 | 696 | 0.116093995 | 1.23E-05 | 2 | 0.000105566 | 17.22741993 | 4 mo | HOM |
| 1008976060 | 2 | AUDpo1 | 696 | 0.116093995 | 3.92E-05 | 3 | 0.00033786 | 25.8411299 | 8 mo | WILD |
| 1013446243 | 1 | AUDpo1 | 696 | 0.116093995 | 3.5E-05 | 1 | 0.000301897 | 8.613709966 | 4 mo | HOM |
| 1013288951 | 2 | AUDpo1 | 696 | 0.116093995 | 3.41E-05 | 3 | 0.000293695 | 25.8411299 | 4 mo | HOM |
| 1013978665 | 1 | ml | 697 | 0.685793965 | 0 | 0 | 0 | 0 | 2 mo | HOM |
| 1013445609 | 1 | ml | 697 | 0.685793965 | 0 | 0 | 0 | 0 | 4 mo | HOM |
| 1013609015 | 1 | ml | 697 | 0.685793965 | 0 | 0 | 0 | 0 | 4 mo | HOM |
| 1013608106 | 2 | ml | 697 | 0.684687349 | 0 | 0 | 0 | 0 | 4 mo | HOM |
| 1011937543 | 1 | ml | 697 | 0.685793965 | 0.001106121 | 8 | 0.001612906 | 11.66531117 | 8 mo | HOM |
| 1010419538 | 2 | ml | 697 | 0.685793965 | 0 | 0 | 0 | 0 | 4 mo | WILD |
| 1010419187 | 1 | ml | 697 | 0.685793965 | 0 | 0 | 0 | 0 | 8 mo | HOM |
| 1013978366 | 1 | ml | 697 | 0.685793965 | 0 | 0 | 0 | 0 | 2 mo | HOM |
| 1014416155 | 2 | ml | 697 | 0.685793965 | 0 | 0 | 0 | 0 | 2 mo | HOM |
| 1013978065 | 2 | ml | 697 | 0.685793965 | 0 | 0 | 0 | 0 | 2 mo | HOM |
| 1009934102 | 2 | ml | 697 | 0.685793965 | 0 | 0 | 0 | 0 | 4 mo | WILD |
| 1009322937 | 2 | ml | 697 | 0.685793965 | 0 | 0 | 0 | 0 | 8 mo | WILD |
| 1011080637 | 2 | ml | 697 | 0.685793965 | 1.08E-08 | 0 | 1.58E-08 | 0 | 8 mo | HOM |
| 1013609713 | 2 | ml | 697 | 0.685793965 | 0 | 0 | 0 | 0 | 2 mo | WILD |
| 1010418842 | 1 | ml | 697 | 0.685793965 | 2.85E-05 | 2 | 4.16E-05 | 2.916327793 | 8 mo | HOM |
| 1013288213 | 2 | ml | 697 | 0.685793965 | 0 | 0 | 0 | 0 | 4 mo | HOM |
| 1008976060 | 2 | ml | 697 | 0.685793965 | 6.86E-06 | 1 | 1E-05 | 1.458163897 | 8 mo | WILD |
| 1013446243 | 1 | ml | 697 | 0.685793965 | 0 | 0 | 0 | 0 | 4 mo | HOM |

| | | | | | | | | | | |
|---|---|---|---|---|---|---|---|---|---|---|
| **1013288951** | 2 | ml | 697 | 0.685793965 | 0 | 0 | 0 | 0 | 4 mo | HOM |
| **1013978665** | 1 | OLF | 698 | 44.56360366 | 0.004824468 | 216 | 0.00010826 | 4.847004781 | 2 mo | HOM |
| **1013445609** | 1 | OLF | 698 | 44.73446799 | 0.002757013 | 261 | 6.16E-05 | 5.834427271 | 4 mo | HOM |
| **1013609015** | 1 | OLF | 698 | 42.50186179 | 0.019171325 | 367 | 0.00045107 | 8.63491585 | 4 mo | HOM |
| **1013608106** | 2 | OLF | 698 | 44.46826926 | 8.18E-06 | 1 | 1.84E-07 | 0.022487945 | 4 mo | HOM |
| **1011937543** | 1 | OLF | 698 | 45.14314148 | 0.04356026 | 647 | 0.000964936 | 14.33218821 | 8 mo | HOM |
| **1010419538** | 2 | OLF | 698 | 45.60813752 | 0.000750715 | 59 | 1.65E-05 | 1.293628795 | 4 mo | WILD |
| **1010419187** | 1 | OLF | 698 | 41.67565624 | 0.047780522 | 1059 | 0.001146485 | 25.41051768 | 8 mo | HOM |
| **1013978366** | 1 | OLF | 698 | 46.83773051 | 0.003968663 | 221 | 8.47E-05 | 4.718418198 | 2 mo | HOM |
| **1014416155** | 2 | OLF | 698 | 45.66267884 | 8.65E-05 | 9 | 1.9E-06 | 0.197097503 | 2 mo | HOM |
| **1013978065** | 2 | OLF | 698 | 47.29380024 | 0.00164801 | 119 | 3.48E-05 | 2.516186041 | 2 mo | HOM |
| **1009934102** | 2 | OLF | 698 | 43.7561514 | 0.001757269 | 127 | 4.02E-05 | 2.902449049 | 4 mo | WILD |
| **1009322937** | 2 | OLF | 698 | 41.3326506 | 0.001783415 | 95 | 4.31E-05 | 2.298425062 | 8 mo | WILD |
| **1011080637** | 2 | OLF | 698 | 40.94523668 | 0.023902643 | 950 | 0.000583771 | 23.20172203 | 8 mo | HOM |
| **1013609713** | 2 | OLF | 698 | 48.11102679 | 0.000329395 | 35 | 6.85E-06 | 0.727483954 | 2 mo | WILD |
| **1010418842** | 1 | OLF | 698 | 43.53565798 | 0.045083758 | 1206 | 0.001035559 | 27.70143041 | 8 mo | HOM |
| **1013288213** | 2 | OLF | 698 | 45.20175206 | 0.036456098 | 257 | 0.00080652 | 5.685620319 | 4 mo | HOM |
| **1008976060** | 2 | OLF | 698 | 46.38389862 | 0.008535597 | 295 | 0.000184021 | 6.359965608 | 8 mo | WILD |
| **1013446243** | 1 | OLF | 698 | 44.13766461 | 0.007395806 | 298 | 0.000167562 | 6.751603253 | 4 mo | HOM |
| **1013288951** | 2 | OLF | 698 | 43.94562601 | 0.015120969 | 424 | 0.000344084 | 9.648286724 | 4 mo | HOM |
| **1013978665** | 1 | Alv6b | 699 | 0.003152488 | 0 | 0 | 0 | 0 | 2 mo | HOM |
| **1013445609** | 1 | Alv6b | 699 | 0.003152488 | 0 | 0 | 0 | 0 | 4 mo | HOM |
| **1013609015** | 1 | Alv6b | 699 | 0.003152488 | 0 | 0 | 0 | 0 | 4 mo | HOM |
| **1013608106** | 2 | Alv6b | 699 | 0.003152488 | 0 | 0 | 0 | 0 | 4 mo | HOM |
| **1011937543** | 1 | Alv6b | 699 | 0.003152488 | 0 | 0 | 0 | 0 | 8 mo | HOM |
| **1010419538** | 2 | Alv6b | 699 | 0.003152488 | 0 | 0 | 0 | 0 | 4 mo | WILD |
| **1010419187** | 1 | Alv6b | 699 | 0.003152488 | 0 | 0 | 0 | 0 | 8 mo | HOM |
| **1013978366** | 1 | Alv6b | 699 | 0.003152488 | 0 | 0 | 0 | 0 | 2 mo | HOM |
| **1014416155** | 2 | Alv6b | 699 | 0.003152488 | 0 | 0 | 0 | 0 | 2 mo | HOM |
| **1013978065** | 2 | Alv6b | 699 | 0.003152488 | 0 | 0 | 0 | 0 | 2 mo | HOM |
| **1009934102** | 2 | Alv6b | 699 | 0.003152488 | 0 | 0 | 0 | 0 | 4 mo | WILD |
| **1009322937** | 2 | Alv6b | 699 | 0.003152488 | 0 | 0 | 0 | 0 | 8 mo | WILD |
| **1011080637** | 2 | Alv6b | 699 | 0.003152488 | 0 | 0 | 0 | 0 | 8 mo | HOM |
| **1013609713** | 2 | Alv6b | 699 | 0.003152488 | 0 | 0 | 0 | 0 | 2 mo | WILD |
| **1010418842** | 1 | Alv6b | 699 | 0.003152488 | 2.66E-07 | 0 | 8.44E-05 | 0 | 8 mo | HOM |
| **1013288213** | 2 | Alv6b | 699 | 0.003152488 | 0 | 0 | 0 | 0 | 4 mo | HOM |
| **1008976060** | 2 | Alv6b | 699 | 0.003152488 | 0 | 0 | 0 | 0 | 8 mo | WILD |
| **1013446243** | 1 | Alv6b | 699 | 0.003152488 | 0 | 0 | 0 | 0 | 4 mo | HOM |
| **1013288951** | 2 | Alv6b | 699 | 0.003152488 | 0 | 0 | 0 | 0 | 4 mo | HOM |
| **1013978665** | 1 | VNC | 701 | 3.304411667 | 0.001003127 | 8 | 0.000303572 | 2.421005857 | 2 mo | HOM |
| **1013445609** | 1 | VNC | 701 | 3.304411667 | 3.39E-05 | 2 | 1.02E-05 | 0.605251464 | 4 mo | HOM |
| **1013609015** | 1 | VNC | 701 | 3.304411667 | 0.001936001 | 10 | 0.000585884 | 3.026257321 | 4 mo | HOM |
| **1013608106** | 2 | VNC | 701 | 3.304411667 | 0 | 0 | 0 | 0 | 4 mo | HOM |
| **1011937543** | 1 | VNC | 701 | 3.304411667 | 0.009089097 | 18 | 0.002750595 | 5.447263178 | 8 mo | HOM |
| **1010419538** | 2 | VNC | 701 | 3.304411667 | 4.62E-05 | 3 | 1.4E-05 | 0.907877196 | 4 mo | WILD |
| **1010419187** | 1 | VNC | 701 | 3.304411667 | 0.002301735 | 11 | 0.000696564 | 3.328883053 | 8 mo | HOM |
| **1013978366** | 1 | VNC | 701 | 3.304411667 | 0.000322949 | 4 | 9.77E-05 | 1.210502928 | 2 mo | HOM |
| **1014416155** | 2 | VNC | 701 | 3.304411667 | 0 | 0 | 0 | 0 | 2 mo | HOM |
| **1013978065** | 2 | VNC | 701 | 3.304411667 | 0.000183994 | 2 | 5.57E-05 | 0.605251464 | 2 mo | HOM |
| **1009934102** | 2 | VNC | 701 | 3.304411667 | 2.63E-10 | 0 | 7.97E-11 | 0 | 4 mo | WILD |
| **1009322937** | 2 | VNC | 701 | 3.304411667 | 6.38E-05 | 5 | 1.93E-05 | 1.513128661 | 8 mo | WILD |
| **1011080637** | 2 | VNC | 701 | 3.304411667 | 0.001803667 | 12 | 0.000545836 | 3.631508785 | 8 mo | HOM |
| **1013609713** | 2 | VNC | 701 | 3.304411667 | 0 | 0 | 0 | 0 | 2 mo | WILD |
| **1010418842** | 1 | VNC | 701 | 3.304411667 | 0.002429478 | 56 | 0.000735222 | 16.947041 | 8 mo | HOM |
| **1013288213** | 2 | VNC | 701 | 3.304411667 | 0.000525373 | 7 | 0.000158991 | 2.118380125 | 4 mo | HOM |
| **1008976060** | 2 | VNC | 701 | 3.304411667 | 0.004467676 | 71 | 0.001352034 | 21.48642698 | 8 mo | WILD |
| **1013446243** | 1 | VNC | 701 | 3.304411667 | 4.61E-05 | 3 | 1.39E-05 | 0.907877196 | 4 mo | HOM |
| **1013288951** | 2 | VNC | 701 | 3.304411667 | 0.00116136 | 5 | 0.000351457 | 1.513128661 | 4 mo | HOM |
| **1013978665** | 1 | SSp-n5 | 702 | 0.58451465 | 0 | 0 | 0 | 0 | 2 mo | HOM |

| | | | | | | | | | | |
|---|---|---|---|---|---|---|---|---|---|---|
| 1013445609 | 1 | SSp-n5 | 702 | 0.58451465 | 5.29E-05 | 7 | 9.06E-05 | 11.97574775 | 4 mo | HOM |
| 1013609015 | 1 | SSp-n5 | 702 | 0.58451465 | 9.16E-05 | 3 | 0.000156728 | 5.132463322 | 4 mo | HOM |
| 1013608106 | 2 | SSp-n5 | 702 | 0.58451465 | 0 | 0 | 0 | 0 | 4 mo | HOM |
| 1011937543 | 1 | SSp-n5 | 702 | 0.58451465 | 0.000708377 | 20 | 0.001211906 | 34.21642214 | 8 mo | HOM |
| 1010419538 | 2 | SSp-n5 | 702 | 0.58451465 | 0 | 0 | 0 | 0 | 4 mo | WILD |
| 1010419187 | 1 | SSp-n5 | 702 | 0.58451465 | 0.000888923 | 39 | 0.001520787 | 66.72202318 | 8 mo | HOM |
| 1013978366 | 1 | SSp-n5 | 702 | 0.58451465 | 0 | 0 | 0 | 0 | 2 mo | HOM |
| 1014416155 | 2 | SSp-n5 | 702 | 0.58451465 | 0 | 0 | 0 | 0 | 2 mo | HOM |
| 1013978065 | 2 | SSp-n5 | 702 | 0.58451465 | 0 | 0 | 0 | 0 | 2 mo | HOM |
| 1009934102 | 2 | SSp-n5 | 702 | 0.58451465 | 1.15E-09 | 0 | 1.97E-09 | 0 | 4 mo | WILD |
| 1009322937 | 2 | SSp-n5 | 702 | 0.58451465 | 0 | 0 | 0 | 0 | 8 mo | WILD |
| 1011080637 | 2 | SSp-n5 | 702 | 0.58451465 | 0.000607614 | 21 | 0.001039518 | 35.92724325 | 8 mo | HOM |
| 1013609713 | 2 | SSp-n5 | 702 | 0.58451465 | 0 | 0 | 0 | 0 | 2 mo | WILD |
| 1010418842 | 1 | SSp-n5 | 702 | 0.58451465 | 0.000844225 | 19 | 0.001444318 | 32.50560104 | 8 mo | HOM |
| 1013288213 | 2 | SSp-n5 | 702 | 0.58451465 | 3.92E-05 | 4 | 6.71E-05 | 6.843284429 | 4 mo | HOM |
| 1008976060 | 2 | SSp-n5 | 702 | 0.58451465 | 1.15E-05 | 2 | 1.97E-05 | 3.421642214 | 8 mo | WILD |
| 1013446243 | 1 | SSp-n5 | 702 | 0.58451465 | 8.46E-05 | 5 | 0.000144744 | 8.554105536 | 4 mo | HOM |
| 1013288951 | 2 | SSp-n5 | 702 | 0.58451465 | 9.52E-05 | 5 | 0.00016284 | 8.554105536 | 4 mo | HOM |
| 1013978665 | 1 | CTXsp | 703 | 9.161442662 | 0 | 0 | 0 | 0 | 2 mo | HOM |
| 1013445609 | 1 | CTXsp | 703 | 9.177143987 | 0.000718424 | 50 | 7.83E-05 | 5.448318134 | 4 mo | HOM |
| 1013609015 | 1 | CTXsp | 703 | 9.177143987 | 0.001724131 | 65 | 0.000187872 | 7.082813574 | 4 mo | HOM |
| 1013608106 | 2 | CTXsp | 703 | 9.17714336 | 0 | 0 | 0 | 0 | 4 mo | HOM |
| 1011937543 | 1 | CTXsp | 703 | 9.177143987 | 0.015818902 | 308 | 0.001723728 | 33.5616397 | 8 mo | HOM |
| 1010419538 | 2 | CTXsp | 703 | 9.17714336 | 3.91E-06 | 1 | 4.26E-07 | 0.10896637 | 4 mo | WILD |
| 1010419187 | 1 | CTXsp | 703 | 9.177143987 | 0.013359438 | 516 | 0.001455729 | 56.22664314 | 8 mo | HOM |
| 1013978366 | 1 | CTXsp | 703 | 9.177143987 | 0 | 0 | 0 | 0 | 2 mo | HOM |
| 1014416155 | 2 | CTXsp | 703 | 9.17714336 | 0 | 0 | 0 | 0 | 2 mo | HOM |
| 1013978065 | 2 | CTXsp | 703 | 9.17714336 | 0 | 0 | 0 | 0 | 2 mo | HOM |
| 1009934102 | 2 | CTXsp | 703 | 9.17714336 | 0 | 0 | 0 | 0 | 4 mo | WILD |
| 1009322937 | 2 | CTXsp | 703 | 9.17714336 | 8.21E-06 | 2 | 8.95E-07 | 0.21793274 | 8 mo | WILD |
| 1011080637 | 2 | CTXsp | 703 | 9.17714336 | 0.009362065 | 446 | 0.00102015 | 48.59900107 | 8 mo | HOM |
| 1013609713 | 2 | CTXsp | 703 | 9.17714336 | 0 | 0 | 0 | 0 | 2 mo | WILD |
| 1010418842 | 1 | CTXsp | 703 | 9.177143987 | 0.015978054 | 480 | 0.00174107 | 52.30385408 | 8 mo | HOM |
| 1013288213 | 2 | CTXsp | 703 | 9.17714336 | 0.00040939 | 35 | 4.46E-05 | 3.813822954 | 4 mo | HOM |
| 1008976060 | 2 | CTXsp | 703 | 9.17714336 | 9.15E-05 | 13 | 9.97E-06 | 1.416562812 | 8 mo | WILD |
| 1013446243 | 1 | CTXsp | 703 | 9.177143987 | 0.001201181 | 71 | 0.000130888 | 7.73661175 | 4 mo | HOM |
| 1013288951 | 2 | CTXsp | 703 | 9.17714336 | 0.00151757 | 76 | 0.000165364 | 8.281444129 | 4 mo | HOM |
| 1013978665 | 1 | AIv1 | 704 | 0.255992366 | 0 | 0 | 0 | 0 | 2 mo | HOM |
| 1013445609 | 1 | AIv1 | 704 | 0.255992366 | 2.93E-06 | 0 | 1.14E-05 | 0 | 4 mo | HOM |
| 1013609015 | 1 | AIv1 | 704 | 0.255992366 | 3.31E-05 | 1 | 0.000129379 | 3.906366489 | 4 mo | HOM |
| 1013608106 | 2 | AIv1 | 704 | 0.255992366 | 2.26E-07 | 0 | 8.82E-07 | 0 | 4 mo | HOM |
| 1011937543 | 1 | AIv1 | 704 | 0.255992366 | 8.1E-05 | 5 | 0.000316581 | 19.53183245 | 8 mo | HOM |
| 1010419538 | 2 | AIv1 | 704 | 0.255992366 | 0 | 0 | 0 | 0 | 4 mo | WILD |
| 1010419187 | 1 | AIv1 | 704 | 0.255992366 | 0.000294825 | 10 | 0.001151694 | 39.06366489 | 8 mo | HOM |
| 1013978366 | 1 | AIv1 | 704 | 0.255992366 | 5.65E-06 | 1 | 2.21E-05 | 3.906366489 | 2 mo | HOM |
| 1014416155 | 2 | AIv1 | 704 | 0.255992366 | 0 | 0 | 0 | 0 | 2 mo | HOM |
| 1013978065 | 2 | AIv1 | 704 | 0.255992366 | 0 | 0 | 0 | 0 | 2 mo | HOM |
| 1009934102 | 2 | AIv1 | 704 | 0.255992366 | 0 | 0 | 0 | 0 | 4 mo | WILD |
| 1009322937 | 2 | AIv1 | 704 | 0.255992366 | 4.92E-05 | 4 | 0.000192244 | 15.62546596 | 8 mo | WILD |
| 1011080637 | 2 | AIv1 | 704 | 0.255992366 | 0.00019149 | 15 | 0.000748029 | 58.59549734 | 8 mo | HOM |
| 1013609713 | 2 | AIv1 | 704 | 0.255992366 | 0 | 0 | 0 | 0 | 2 mo | WILD |
| 1010418842 | 1 | AIv1 | 704 | 0.255992366 | 0.000198332 | 12 | 0.000774759 | 46.87639787 | 8 mo | HOM |
| 1013288213 | 2 | AIv1 | 704 | 0.255992366 | 5.99E-05 | 1 | 0.000233884 | 3.906366489 | 4 mo | HOM |
| 1008976060 | 2 | AIv1 | 704 | 0.255992366 | 2.14E-05 | 1 | 8.35E-05 | 3.906366489 | 8 mo | WILD |
| 1013446243 | 1 | AIv1 | 704 | 0.255992366 | 3.36E-05 | 4 | 0.000131111 | 15.62546596 | 4 mo | HOM |
| 1013288951 | 2 | AIv1 | 704 | 0.255992366 | 0.000154477 | 2 | 0.000603445 | 7.812732978 | 4 mo | HOM |
| 1013978665 | 1 | OP | 706 | 0.059882861 | 0 | 0 | 0 | 0 | 2 mo | HOM |
| 1013445609 | 1 | OP | 706 | 0.059882861 | 0 | 0 | 0 | 0 | 4 mo | HOM |
| 1013609015 | 1 | OP | 706 | 0.059882861 | 0 | 0 | 0 | 0 | 4 mo | HOM |

| | | | | | | | | | | |
|---|---|---|---|---|---|---|---|---|---|---|
| 1013608106 | 2 | OP | 706 | 0.059882861 | 0 | 0 | 0 | 0 | 4 mo | HOM |
| 1011937543 | 1 | OP | 706 | 0.059882861 | 5.92E-05 | 1 | 0.000987884 | 16.69926898 | 8 mo | HOM |
| 1010419538 | 2 | OP | 706 | 0.059882861 | 0 | 0 | 0 | 0 | 4 mo | WILD |
| 1010419187 | 1 | OP | 706 | 0.059882861 | 2.25E-05 | 1 | 0.000376209 | 16.69926898 | 8 mo | HOM |
| 1013978366 | 1 | OP | 706 | 0.059882861 | 5.26E-07 | 0 | 8.78E-06 | 0 | 2 mo | HOM |
| 1014416155 | 2 | OP | 706 | 0.059882861 | 0 | 0 | 0 | 0 | 2 mo | HOM |
| 1013978065 | 2 | OP | 706 | 0.059882861 | 0 | 0 | 0 | 0 | 2 mo | HOM |
| 1009934102 | 2 | OP | 706 | 0.059882861 | 0 | 0 | 0 | 0 | 4 mo | WILD |
| 1009322937 | 2 | OP | 706 | 0.059882861 | 4.11E-06 | 1 | 6.87E-05 | 16.69926898 | 8 mo | WILD |
| 1011080637 | 2 | OP | 706 | 0.059882861 | 4.26E-06 | 1 | 7.12E-05 | 16.69926898 | 8 mo | HOM |
| 1013609713 | 2 | OP | 706 | 0.059882861 | 0 | 0 | 0 | 0 | 2 mo | WILD |
| 1010418842 | 1 | OP | 706 | 0.059882861 | 9.62E-07 | 0 | 1.61E-05 | 0 | 8 mo | HOM |
| 1013288213 | 2 | OP | 706 | 0.059882861 | 0 | 0 | 0 | 0 | 4 mo | HOM |
| 1008976060 | 2 | OP | 706 | 0.059882861 | 2.03E-07 | 0 | 3.39E-06 | 0 | 8 mo | WILD |
| 1013446243 | 1 | OP | 706 | 0.059882861 | 0 | 0 | 0 | 0 | 4 mo | HOM |
| 1013288951 | 2 | OP | 706 | 0.059882861 | 0 | 0 | 0 | 0 | 4 mo | HOM |
| 1013978665 | 1 | ILA1 | 707 | 0.155823587 | 0 | 0 | 0 | 0 | 2 mo | HOM |
| 1013445609 | 1 | ILA1 | 707 | 0.155823587 | 0 | 0 | 0 | 0 | 4 mo | HOM |
| 1013609015 | 1 | ILA1 | 707 | 0.155823587 | 0.000317496 | 3 | 0.002037534 | 19.25254104 | 4 mo | HOM |
| 1013608106 | 2 | ILA1 | 707 | 0.155823587 | 0 | 0 | 0 | 0 | 4 mo | HOM |
| 1011937543 | 1 | ILA1 | 707 | 0.155823587 | 0.000143462 | 3 | 0.000920666 | 19.25254104 | 8 mo | HOM |
| 1010419538 | 2 | ILA1 | 707 | 0.155823587 | 0 | 0 | 0 | 0 | 4 mo | WILD |
| 1010419187 | 1 | ILA1 | 707 | 0.155823587 | 0.000240649 | 5 | 0.001544368 | 32.0875684 | 8 mo | HOM |
| 1013978366 | 1 | ILA1 | 707 | 0.155823587 | 0 | 0 | 0 | 0 | 2 mo | HOM |
| 1014416155 | 2 | ILA1 | 707 | 0.155823587 | 0 | 0 | 0 | 0 | 2 mo | HOM |
| 1013978065 | 2 | ILA1 | 707 | 0.155823587 | 0 | 0 | 0 | 0 | 2 mo | HOM |
| 1009934102 | 2 | ILA1 | 707 | 0.155823587 | 0 | 0 | 0 | 0 | 4 mo | WILD |
| 1009322937 | 2 | ILA1 | 707 | 0.155823587 | 0.000218103 | 3 | 0.001399682 | 19.25254104 | 8 mo | WILD |
| 1011080637 | 2 | ILA1 | 707 | 0.155823587 | 9.35E-05 | 5 | 0.000600167 | 32.0875684 | 8 mo | HOM |
| 1013609713 | 2 | ILA1 | 707 | 0.155823587 | 0 | 0 | 0 | 0 | 2 mo | WILD |
| 1010418842 | 1 | ILA1 | 707 | 0.155823587 | 0.001649121 | 2 | 0.010583256 | 12.83502736 | 8 mo | HOM |
| 1013288213 | 2 | ILA1 | 707 | 0.155823587 | 0.001289292 | 2 | 0.00827405 | 12.83502736 | 4 mo | HOM |
| 1008976060 | 2 | ILA1 | 707 | 0.155823587 | 7.9E-05 | 2 | 0.000506988 | 12.83502736 | 8 mo | WILD |
| 1013446243 | 1 | ILA1 | 707 | 0.155823587 | 6.09E-08 | 0 | 3.91E-07 | 0 | 4 mo | HOM |
| 1013288951 | 2 | ILA1 | 707 | 0.155823587 | 0.000304165 | 10 | 0.001951983 | 64.1751368 | 4 mo | HOM |
| 1013978665 | 1 | VP | 709 | 2.920086233 | 0 | 0 | 0 | 0 | 2 mo | HOM |
| 1013445609 | 1 | VP | 709 | 2.920086233 | 0 | 0 | 0 | 0 | 4 mo | HOM |
| 1013609015 | 1 | VP | 709 | 2.920086233 | 5.19E-05 | 2 | 1.78E-05 | 0.68491128 | 4 mo | HOM |
| 1013608106 | 2 | VP | 709 | 2.920086233 | 0 | 0 | 0 | 0 | 4 mo | HOM |
| 1011937543 | 1 | VP | 709 | 2.920086233 | 0.00162038 | 21 | 0.000554908 | 7.191568441 | 8 mo | HOM |
| 1010419538 | 2 | VP | 709 | 2.920086233 | 0 | 0 | 0 | 0 | 4 mo | WILD |
| 1010419187 | 1 | VP | 709 | 2.920086233 | 0.000677406 | 18 | 0.000231982 | 6.164201521 | 8 mo | HOM |
| 1013978366 | 1 | VP | 709 | 2.920086233 | 0 | 0 | 0 | 0 | 2 mo | HOM |
| 1014416155 | 2 | VP | 709 | 2.920086233 | 0 | 0 | 0 | 0 | 2 mo | HOM |
| 1013978065 | 2 | VP | 709 | 2.920086233 | 0 | 0 | 0 | 0 | 2 mo | HOM |
| 1009934102 | 2 | VP | 709 | 2.920086233 | 0 | 0 | 0 | 0 | 4 mo | WILD |
| 1009322937 | 2 | VP | 709 | 2.920086233 | 0 | 0 | 0 | 0 | 8 mo | WILD |
| 1011080637 | 2 | VP | 709 | 2.920086233 | 0.001285559 | 33 | 0.000440247 | 11.30103612 | 8 mo | HOM |
| 1013609713 | 2 | VP | 709 | 2.920086233 | 0 | 0 | 0 | 0 | 2 mo | WILD |
| 1010418842 | 1 | VP | 709 | 2.920086233 | 0.001406614 | 33 | 0.000481703 | 11.30103612 | 8 mo | HOM |
| 1013288213 | 2 | VP | 709 | 2.920086233 | 2.25E-05 | 3 | 7.7E-06 | 1.02736692 | 4 mo | HOM |
| 1008976060 | 2 | VP | 709 | 2.920086233 | 3.41E-05 | 5 | 1.17E-05 | 1.7122782 | 8 mo | WILD |
| 1013446243 | 1 | VP | 709 | 2.920086233 | 0 | 0 | 0 | 0 | 4 mo | HOM |
| 1013288951 | 2 | VP | 709 | 2.920086233 | 5.09E-06 | 1 | 1.74E-06 | 0.34245564 | 4 mo | HOM |
| 1013978665 | 1 | CU | 711 | 0.343192668 | 0 | 0 | 0 | 0 | 2 mo | HOM |
| 1013445609 | 1 | CU | 711 | 0.343192668 | 0 | 0 | 0 | 0 | 4 mo | HOM |
| 1013609015 | 1 | CU | 711 | 0.343192668 | 0 | 0 | 0 | 0 | 4 mo | HOM |
| 1013608106 | 2 | CU | 711 | 0.343192668 | 0 | 0 | 0 | 0 | 4 mo | HOM |
| 1011937543 | 1 | CU | 711 | 0.343192668 | 0.000853494 | 1 | 0.002486924 | 2.913815163 | 8 mo | HOM |

| | | | | | | | | | | |
|---|---|---|---|---|---|---|---|---|---|---|
| **1010419538** | 2 | CU | 711 | 0.343192668 | 0 | 0 | 0 | 0 | 4 mo | WILD |
| **1010419187** | 1 | CU | 711 | 0.343192668 | 6.63E-05 | 0 | 0.000193184 | 0 | 8 mo | HOM |
| **1013978366** | 1 | CU | 711 | 0.343192668 | 0 | 0 | 0 | 0 | 2 mo | HOM |
| **1014416155** | 2 | CU | 711 | 0.343192668 | 0 | 0 | 0 | 0 | 2 mo | HOM |
| **1013978065** | 2 | CU | 711 | 0.343192668 | 0 | 0 | 0 | 0 | 2 mo | HOM |
| **1009934102** | 2 | CU | 711 | 0.343192668 | 0 | 0 | 0 | 0 | 4 mo | WILD |
| **1009322937** | 2 | CU | 711 | 0.343192668 | 0 | 0 | 0 | 0 | 8 mo | WILD |
| **1011080637** | 2 | CU | 711 | 0.343192668 | 0 | 0 | 0 | 0 | 8 mo | HOM |
| **1013609713** | 2 | CU | 711 | 0.343192668 | 0 | 0 | 0 | 0 | 2 mo | WILD |
| **1010418842** | 1 | CU | 711 | 0.343192668 | 0 | 0 | 0 | 0 | 8 mo | HOM |
| **1013288213** | 2 | CU | 711 | 0.343192668 | 0 | 0 | 0 | 0 | 4 mo | HOM |
| **1008976060** | 2 | CU | 711 | 0.343192668 | 3.9E-06 | 1 | 1.14E-05 | 2.913815163 | 8 mo | WILD |
| **1013446243** | 1 | CU | 711 | 0.343192668 | 2E-06 | 0 | 5.83E-06 | 0 | 4 mo | HOM |
| **1013288951** | 2 | CU | 711 | 0.343192668 | 4.18E-05 | 0 | 0.000121773 | 0 | 4 mo | HOM |
| **1013978665** | 1 | ORB | 714 | 6.082736875 | 4.25E-05 | 4 | 6.98E-06 | 0.657598723 | 2 mo | HOM |
| **1013445609** | 1 | ORB | 714 | 6.082736875 | 0.000213952 | 14 | 3.52E-05 | 2.30159553 | 4 mo | HOM |
| **1013609015** | 1 | ORB | 714 | 6.082736875 | 0.00107359 | 30 | 0.000176498 | 4.931990421 | 4 mo | HOM |
| **1013608106** | 2 | ORB | 714 | 6.082736875 | 0 | 0 | 0 | 0 | 4 mo | HOM |
| **1011937543** | 1 | ORB | 714 | 6.082736875 | 0.005788106 | 130 | 0.000951563 | 21.37195849 | 8 mo | HOM |
| **1010419538** | 2 | ORB | 714 | 6.082736875 | 0 | 0 | 0 | 0 | 4 mo | WILD |
| **1010419187** | 1 | ORB | 714 | 6.082736875 | 0.005605472 | 214 | 0.000921538 | 35.18153167 | 8 mo | HOM |
| **1013978366** | 1 | ORB | 714 | 6.082736875 | 2.79E-05 | 0 | 4.58E-06 | 0 | 2 mo | HOM |
| **1014416155** | 2 | ORB | 714 | 6.082736875 | 1.66E-05 | 1 | 2.73E-06 | 0.164399681 | 2 mo | HOM |
| **1013978065** | 2 | ORB | 714 | 6.082736875 | 3.21E-06 | 0 | 5.27E-07 | 0 | 2 mo | HOM |
| **1009934102** | 2 | ORB | 714 | 6.082736875 | 2.7E-05 | 2 | 4.44E-06 | 0.328799361 | 4 mo | WILD |
| **1009322937** | 2 | ORB | 714 | 6.082736875 | 0.000284717 | 17 | 4.68E-05 | 2.794794572 | 8 mo | WILD |
| **1011080637** | 2 | ORB | 714 | 6.082736875 | 0.004902868 | 159 | 0.00080603 | 26.13954923 | 8 mo | HOM |
| **1013609713** | 2 | ORB | 714 | 6.082736875 | 0 | 0 | 0 | 0 | 2 mo | WILD |
| **1010418842** | 1 | ORB | 714 | 6.082736875 | 0.007292953 | 177 | 0.001198959 | 29.09874348 | 8 mo | HOM |
| **1013288213** | 2 | ORB | 714 | 6.082736875 | 0.004254495 | 33 | 0.000699438 | 5.425189463 | 4 mo | HOM |
| **1008976060** | 2 | ORB | 714 | 6.082736875 | 9.52E-05 | 9 | 1.57E-05 | 1.479597126 | 8 mo | WILD |
| **1013446243** | 1 | ORB | 714 | 6.082736875 | 0.000464601 | 21 | 7.64E-05 | 3.452393294 | 4 mo | HOM |
| **1013288951** | 2 | ORB | 714 | 6.082736875 | 0.001913051 | 61 | 0.000314505 | 10.02838052 | 4 mo | HOM |
| **1013978665** | 1 | VPL | 718 | 0.902141274 | 0 | 0 | 0 | 0 | 2 mo | HOM |
| **1013445609** | 1 | VPL | 718 | 0.902141274 | 0 | 0 | 0 | 0 | 4 mo | HOM |
| **1013609015** | 1 | VPL | 718 | 0.902141274 | 0 | 0 | 0 | 0 | 4 mo | HOM |
| **1013608106** | 2 | VPL | 718 | 0.902141274 | 0 | 0 | 0 | 0 | 4 mo | HOM |
| **1011937543** | 1 | VPL | 718 | 0.902141274 | 0.000650383 | 14 | 0.000720932 | 15.51863373 | 8 mo | HOM |
| **1010419538** | 2 | VPL | 718 | 0.902141274 | 0 | 0 | 0 | 0 | 4 mo | WILD |
| **1010419187** | 1 | VPL | 718 | 0.902141274 | 7.64E-05 | 4 | 8.47E-05 | 4.433895352 | 8 mo | HOM |
| **1013978366** | 1 | VPL | 718 | 0.902141274 | 0 | 0 | 0 | 0 | 2 mo | HOM |
| **1014416155** | 2 | VPL | 718 | 0.902141274 | 0 | 0 | 0 | 0 | 2 mo | HOM |
| **1013978065** | 2 | VPL | 718 | 0.902141274 | 0 | 0 | 0 | 0 | 2 mo | HOM |
| **1009934102** | 2 | VPL | 718 | 0.902141274 | 0 | 0 | 0 | 0 | 4 mo | WILD |
| **1009322937** | 2 | VPL | 718 | 0.902141274 | 0 | 0 | 0 | 0 | 8 mo | WILD |
| **1011080637** | 2 | VPL | 718 | 0.902141274 | 0.000371451 | 14 | 0.000411743 | 15.51863373 | 8 mo | HOM |
| **1013609713** | 2 | VPL | 718 | 0.902141274 | 0 | 0 | 0 | 0 | 2 mo | WILD |
| **1010418842** | 1 | VPL | 718 | 0.902141274 | 0.000362067 | 10 | 0.000401341 | 11.08473838 | 8 mo | HOM |
| **1013288213** | 2 | VPL | 718 | 0.902141274 | 4.58E-06 | 1 | 5.07E-06 | 1.108473838 | 4 mo | HOM |
| **1008976060** | 2 | VPL | 718 | 0.902141274 | 6.03E-06 | 1 | 6.69E-06 | 1.108473838 | 8 mo | WILD |
| **1013446243** | 1 | VPL | 718 | 0.902141274 | 0 | 0 | 0 | 0 | 4 mo | HOM |
| **1013288951** | 2 | VPL | 718 | 0.902141274 | 0 | 0 | 0 | 0 | 4 mo | HOM |
| **1013978665** | 1 | DCN | 720 | 0.427283753 | 0 | 0 | 0 | 0 | 2 mo | HOM |
| **1013445609** | 1 | DCN | 720 | 0.427283753 | 0 | 0 | 0 | 0 | 4 mo | HOM |
| **1013609015** | 1 | DCN | 720 | 0.427283753 | 0 | 0 | 0 | 0 | 4 mo | HOM |
| **1013608106** | 2 | DCN | 720 | 0.427283753 | 0 | 0 | 0 | 0 | 4 mo | HOM |
| **1011937543** | 1 | DCN | 720 | 0.427283753 | 0.000853494 | 1 | 0.001997488 | 2.340365137 | 8 mo | HOM |
| **1010419538** | 2 | DCN | 720 | 0.427283753 | 1.06E-06 | 0 | 2.49E-06 | 0 | 4 mo | WILD |
| **1010419187** | 1 | DCN | 720 | 0.427283753 | 0.00229802 | 1 | 0.005378206 | 2.340365137 | 8 mo | HOM |

| | | | | | | | | | | |
|---|---|---|---|---|---|---|---|---|---|---|
| **1013978366** | 1 | DCN | 720 | 0.427283753 | 3.66E-06 | 1 | 8.55E-06 | 2.340365137 | 2 mo | HOM |
| **1014416155** | 2 | DCN | 720 | 0.427283753 | 0 | 0 | 0 | 0 | 2 mo | HOM |
| **1013978065** | 2 | DCN | 720 | 0.427283753 | 2.33E-06 | 0 | 5.44E-06 | 0 | 2 mo | HOM |
| **1009934102** | 2 | DCN | 720 | 0.427283753 | 0 | 0 | 0 | 0 | 4 mo | WILD |
| **1009322937** | 2 | DCN | 720 | 0.427283753 | 1.96E-06 | 0 | 4.58E-06 | 0 | 8 mo | WILD |
| **1011080637** | 2 | DCN | 720 | 0.427283753 | 0 | 0 | 0 | 0 | 8 mo | HOM |
| **1013609713** | 2 | DCN | 720 | 0.427283753 | 0 | 0 | 0 | 0 | 2 mo | WILD |
| **1010418842** | 1 | DCN | 720 | 0.427283753 | 8.39E-06 | 1 | 1.96E-05 | 2.340365137 | 8 mo | HOM |
| **1013288213** | 2 | DCN | 720 | 0.427283753 | 0 | 0 | 0 | 0 | 4 mo | HOM |
| **1008976060** | 2 | DCN | 720 | 0.427283753 | 3.9E-06 | 1 | 9.14E-06 | 2.340365137 | 8 mo | WILD |
| **1013446243** | 1 | DCN | 720 | 0.427283753 | 0.000225533 | 0 | 0.000527829 | 0 | 4 mo | HOM |
| **1013288951** | 2 | DCN | 720 | 0.427283753 | 0.001217275 | 3 | 0.002848868 | 7.021095411 | 4 mo | HOM |
| **1013978665** | 1 | VISp4 | 721 | 1.05457949 | 0 | 0 | 0 | 0 | 2 mo | HOM |
| **1013445609** | 1 | VISp4 | 721 | 1.05457949 | 0.000133826 | 11 | 0.0001269 | 10.43069784 | 4 mo | HOM |
| **1013609015** | 1 | VISp4 | 721 | 1.05457949 | 0.000663844 | 22 | 0.000629487 | 20.86139567 | 4 mo | HOM |
| **1013608106** | 2 | VISp4 | 721 | 1.05457949 | 0 | 0 | 0 | 0 | 4 mo | HOM |
| **1011937543** | 1 | VISp4 | 721 | 1.05457949 | 0.001876495 | 29 | 0.001779377 | 27.49911248 | 8 mo | HOM |
| **1010419538** | 2 | VISp4 | 721 | 1.05457949 | 0 | 0 | 0 | 0 | 4 mo | WILD |
| **1010419187** | 1 | VISp4 | 721 | 1.05457949 | 0.002607378 | 69 | 0.002472434 | 65.42892279 | 8 mo | HOM |
| **1013978366** | 1 | VISp4 | 721 | 1.05457949 | 1.81E-06 | 1 | 1.71E-06 | 0.948245258 | 2 mo | HOM |
| **1014416155** | 2 | VISp4 | 721 | 1.05457949 | 0 | 0 | 0 | 0 | 2 mo | HOM |
| **1013978065** | 2 | VISp4 | 721 | 1.05457949 | 0 | 0 | 0 | 0 | 2 mo | HOM |
| **1009934102** | 2 | VISp4 | 721 | 1.05457949 | 3.15E-06 | 0 | 2.99E-06 | 0 | 4 mo | WILD |
| **1009322937** | 2 | VISp4 | 721 | 1.05457949 | 0 | 0 | 0 | 0 | 8 mo | WILD |
| **1011080637** | 2 | VISp4 | 721 | 1.05457949 | 0.001365382 | 50 | 0.001294717 | 47.41226289 | 8 mo | HOM |
| **1013609713** | 2 | VISp4 | 721 | 1.05457949 | 0 | 0 | 0 | 0 | 2 mo | WILD |
| **1010418842** | 1 | VISp4 | 721 | 1.05457949 | 0.001905782 | 50 | 0.001807149 | 47.41226289 | 8 mo | HOM |
| **1013288213** | 2 | VISp4 | 721 | 1.05457949 | 6.95E-05 | 4 | 6.59E-05 | 3.792981031 | 4 mo | HOM |
| **1008976060** | 2 | VISp4 | 721 | 1.05457949 | 0 | 0 | 0 | 0 | 8 mo | WILD |
| **1013446243** | 1 | VISp4 | 721 | 1.05457949 | 0.00035444 | 14 | 0.000336096 | 13.27543361 | 4 mo | HOM |
| **1013288951** | 2 | VISp4 | 721 | 1.05457949 | 0.000344978 | 20 | 0.000327124 | 18.96490516 | 4 mo | HOM |
| **1013978665** | 1 | ORBl | 723 | 2.810581563 | 2.91E-05 | 4 | 1.04E-05 | 1.423192998 | 2 mo | HOM |
| **1013445609** | 1 | ORBl | 723 | 2.810581563 | 0.000117946 | 9 | 4.2E-05 | 3.202184245 | 4 mo | HOM |
| **1013609015** | 1 | ORBl | 723 | 2.810581563 | 0.000283838 | 10 | 0.000100989 | 3.557982495 | 4 mo | HOM |
| **1013608106** | 2 | ORBl | 723 | 2.810581563 | 0 | 0 | 0 | 0 | 4 mo | HOM |
| **1011937543** | 1 | ORBl | 723 | 2.810581563 | 0.002378193 | 54 | 0.000846157 | 19.21310547 | 8 mo | HOM |
| **1010419538** | 2 | ORBl | 723 | 2.810581563 | 0 | 0 | 0 | 0 | 4 mo | WILD |
| **1010419187** | 1 | ORBl | 723 | 2.810581563 | 0.003010892 | 118 | 0.00107127 | 41.98419344 | 8 mo | HOM |
| **1013978366** | 1 | ORBl | 723 | 2.810581563 | 2.12E-05 | 0 | 7.54E-06 | 0 | 2 mo | HOM |
| **1014416155** | 2 | ORBl | 723 | 2.810581563 | 0 | 0 | 0 | 0 | 2 mo | HOM |
| **1013978065** | 2 | ORBl | 723 | 2.810581563 | 0 | 0 | 0 | 0 | 2 mo | HOM |
| **1009934102** | 2 | ORBl | 723 | 2.810581563 | 1.21E-05 | 0 | 4.29E-06 | 0 | 4 mo | WILD |
| **1009322937** | 2 | ORBl | 723 | 2.810581563 | 0.000126654 | 6 | 4.51E-05 | 2.134789497 | 8 mo | WILD |
| **1011080637** | 2 | ORBl | 723 | 2.810581563 | 0.001547568 | 76 | 0.000550622 | 27.04066696 | 8 mo | HOM |
| **1013609713** | 2 | ORBl | 723 | 2.810581563 | 0 | 0 | 0 | 0 | 2 mo | WILD |
| **1010418842** | 1 | ORBl | 723 | 2.810581563 | 0.002340983 | 85 | 0.000832918 | 30.24285121 | 8 mo | HOM |
| **1013288213** | 2 | ORBl | 723 | 2.810581563 | 0.000795634 | 9 | 0.000283085 | 3.202184245 | 4 mo | HOM |
| **1008976060** | 2 | ORBl | 723 | 2.810581563 | 2.07E-05 | 2 | 7.36E-06 | 0.711596499 | 8 mo | WILD |
| **1013446243** | 1 | ORBl | 723 | 2.810581563 | 0.000214787 | 12 | 7.64E-05 | 4.269578994 | 4 mo | HOM |
| **1013288951** | 2 | ORBl | 723 | 2.810581563 | 0.000595499 | 19 | 0.000211878 | 6.76016674 | 4 mo | HOM |
| **1013978665** | 1 | VPLpc | 725 | 0.101810955 | 0 | 0 | 0 | 0 | 2 mo | HOM |
| **1013445609** | 1 | VPLpc | 725 | 0.101810955 | 0 | 0 | 0 | 0 | 4 mo | HOM |
| **1013609015** | 1 | VPLpc | 725 | 0.101810955 | 0 | 0 | 0 | 0 | 4 mo | HOM |
| **1013608106** | 2 | VPLpc | 725 | 0.101810955 | 0 | 0 | 0 | 0 | 4 mo | HOM |
| **1011937543** | 1 | VPLpc | 725 | 0.101810955 | 1.45E-05 | 1 | 0.000142886 | 9.822125685 | 8 mo | HOM |
| **1010419538** | 2 | VPLpc | 725 | 0.101810955 | 0 | 0 | 0 | 0 | 4 mo | WILD |
| **1010419187** | 1 | VPLpc | 725 | 0.101810955 | 8.51E-05 | 1 | 0.000835374 | 9.822125685 | 8 mo | HOM |
| **1013978366** | 1 | VPLpc | 725 | 0.101810955 | 0 | 0 | 0 | 0 | 2 mo | HOM |
| **1014416155** | 2 | VPLpc | 725 | 0.101810955 | 0 | 0 | 0 | 0 | 2 mo | HOM |

| | | | | | | | | | | |
|---|---|---|---|---|---|---|---|---|---|---|
| **1013978065** | 2 | VPLpc | 725 | 0.101810955 | 0 | 0 | 0 | 0 | 2 mo | HOM |
| **1009934102** | 2 | VPLpc | 725 | 0.101810955 | 0 | 0 | 0 | 0 | 4 mo | WILD |
| **1009322937** | 2 | VPLpc | 725 | 0.101810955 | 0 | 0 | 0 | 0 | 8 mo | WILD |
| **1011080637** | 2 | VPLpc | 725 | 0.101810955 | 5.17E-05 | 2 | 0.000507768 | 19.64425137 | 8 mo | HOM |
| **1013609713** | 2 | VPLpc | 725 | 0.101810955 | 0 | 0 | 0 | 0 | 2 mo | WILD |
| **1010418842** | 1 | VPLpc | 725 | 0.101810955 | 7.16E-06 | 1 | 7.03E-05 | 9.822125685 | 8 mo | HOM |
| **1013288213** | 2 | VPLpc | 725 | 0.101810955 | 0 | 0 | 0 | 0 | 4 mo | HOM |
| **1008976060** | 2 | VPLpc | 725 | 0.101810955 | 0 | 0 | 0 | 0 | 8 mo | WILD |
| **1013446243** | 1 | VPLpc | 725 | 0.101810955 | 0 | 0 | 0 | 0 | 4 mo | HOM |
| **1013288951** | 2 | VPLpc | 725 | 0.101810955 | 0 | 0 | 0 | 0 | 4 mo | HOM |
| **1013978665** | 1 | DG | 726 | 6.796575654 | 7.13E-06 | 1 | 1.05E-06 | 0.147132917 | 2 mo | HOM |
| **1013445609** | 1 | DG | 726 | 6.796575654 | 5.08E-05 | 4 | 7.48E-06 | 0.588531667 | 4 mo | HOM |
| **1013609015** | 1 | DG | 726 | 6.796575654 | 0.000253898 | 14 | 3.74E-05 | 2.059860835 | 4 mo | HOM |
| **1013608106** | 2 | DG | 726 | 6.796575654 | 0 | 0 | 0 | 0 | 4 mo | HOM |
| **1011937543** | 1 | DG | 726 | 6.796575654 | 0.009318522 | 97 | 0.001371061 | 14.27189293 | 8 mo | HOM |
| **1010419538** | 2 | DG | 726 | 6.796575654 | 0.000100986 | 7 | 1.49E-05 | 1.029930417 | 4 mo | WILD |
| **1010419187** | 1 | DG | 726 | 6.796575654 | 0.010154579 | 164 | 0.001494073 | 24.12979835 | 8 mo | HOM |
| **1013978366** | 1 | DG | 726 | 6.796575654 | 0 | 0 | 0 | 0 | 2 mo | HOM |
| **1014416155** | 2 | DG | 726 | 6.796575654 | 0 | 0 | 0 | 0 | 2 mo | HOM |
| **1013978065** | 2 | DG | 726 | 6.796575654 | 3.91E-06 | 1 | 5.75E-07 | 0.147132917 | 2 mo | HOM |
| **1009934102** | 2 | DG | 726 | 6.796575654 | 0.000129631 | 10 | 1.91E-05 | 1.471329168 | 4 mo | WILD |
| **1009322937** | 2 | DG | 726 | 6.796575654 | 5.04E-05 | 4 | 7.42E-06 | 0.588531667 | 8 mo | WILD |
| **1011080637** | 2 | DG | 726 | 6.796575654 | 0.002363662 | 103 | 0.000347772 | 15.15469043 | 8 mo | HOM |
| **1013609713** | 2 | DG | 726 | 6.796575654 | 6.14E-06 | 0 | 9.03E-07 | 0 | 2 mo | WILD |
| **1010418842** | 1 | DG | 726 | 6.796575654 | 0.005589801 | 193 | 0.000822444 | 28.39665293 | 8 mo | HOM |
| **1013288213** | 2 | DG | 726 | 6.796575654 | 7.98E-05 | 10 | 1.17E-05 | 1.471329168 | 4 mo | HOM |
| **1008976060** | 2 | DG | 726 | 6.796575654 | 0.000195765 | 18 | 2.88E-05 | 2.648392502 | 8 mo | WILD |
| **1013446243** | 1 | DG | 726 | 6.796575654 | 0.00030857 | 22 | 4.54E-05 | 3.236924169 | 4 mo | HOM |
| **1013288951** | 2 | DG | 726 | 6.796575654 | 0.000340901 | 18 | 5.02E-05 | 2.648392502 | 4 mo | HOM |
| **1013978665** | 1 | ENTm5 | 727 | 0.939078963 | 0 | 0 | 0 | 0 | 2 mo | HOM |
| **1013445609** | 1 | ENTm5 | 727 | 0.939078963 | 4.29E-06 | 0 | 4.57E-06 | 0 | 4 mo | HOM |
| **1013609015** | 1 | ENTm5 | 727 | 0.939078963 | 0.000105964 | 5 | 0.000112838 | 5.32436589 | 4 mo | HOM |
| **1013608106** | 2 | ENTm5 | 727 | 0.939078963 | 0 | 0 | 0 | 0 | 4 mo | HOM |
| **1011937543** | 1 | ENTm5 | 727 | 0.939078963 | 0.000758133 | 13 | 0.000807315 | 13.84335131 | 8 mo | HOM |
| **1010419538** | 2 | ENTm5 | 727 | 0.939078963 | 0 | 0 | 0 | 0 | 4 mo | WILD |
| **1010419187** | 1 | ENTm5 | 727 | 0.939078963 | 0.000703689 | 34 | 0.000749339 | 36.20568805 | 8 mo | HOM |
| **1013978366** | 1 | ENTm5 | 727 | 0.939078963 | 0 | 0 | 0 | 0 | 2 mo | HOM |
| **1014416155** | 2 | ENTm5 | 727 | 0.939078963 | 0 | 0 | 0 | 0 | 2 mo | HOM |
| **1013978065** | 2 | ENTm5 | 727 | 0.939078963 | 0 | 0 | 0 | 0 | 2 mo | HOM |
| **1009934102** | 2 | ENTm5 | 727 | 0.939078963 | 0 | 0 | 0 | 0 | 4 mo | WILD |
| **1009322937** | 2 | ENTm5 | 727 | 0.939078963 | 0 | 0 | 0 | 0 | 8 mo | WILD |
| **1011080637** | 2 | ENTm5 | 727 | 0.939078963 | 0.00026101 | 19 | 0.000277943 | 20.23259038 | 8 mo | HOM |
| **1013609713** | 2 | ENTm5 | 727 | 0.939078963 | 0 | 0 | 0 | 0 | 2 mo | WILD |
| **1010418842** | 1 | ENTm5 | 727 | 0.939078963 | 0.001123628 | 32 | 0.001196521 | 34.0759417 | 8 mo | HOM |
| **1013288213** | 2 | ENTm5 | 727 | 0.939078963 | 1.1E-06 | 0 | 1.17E-06 | 0 | 4 mo | HOM |
| **1008976060** | 2 | ENTm5 | 727 | 0.939078963 | 0 | 0 | 0 | 0 | 8 mo | WILD |
| **1013446243** | 1 | ENTm5 | 727 | 0.939078963 | 6.84E-05 | 3 | 7.28E-05 | 3.194619534 | 4 mo | HOM |
| **1013288951** | 2 | ENTm5 | 727 | 0.939078963 | 5.55E-05 | 1 | 5.91E-05 | 1.064873178 | 4 mo | HOM |
| **1013978665** | 1 | arb | 728 | 7.129526029 | 0.000142499 | 9 | 2E-05 | 1.262356006 | 2 mo | HOM |
| **1013445609** | 1 | arb | 728 | 7.129526029 | 6.38E-06 | 1 | 8.95E-07 | 0.140261778 | 4 mo | HOM |
| **1013609015** | 1 | arb | 728 | 7.129526029 | 0.000172419 | 8 | 2.42E-05 | 1.122094227 | 4 mo | HOM |
| **1013608106** | 2 | arb | 728 | 7.129526029 | 0 | 0 | 0 | 0 | 4 mo | HOM |
| **1011937543** | 1 | arb | 728 | 7.129526029 | 0.004277225 | 23 | 0.000599931 | 3.226020903 | 8 mo | HOM |
| **1010419538** | 2 | arb | 728 | 7.095511091 | 9.31E-05 | 5 | 1.31E-05 | 0.704670874 | 4 mo | WILD |
| **1010419187** | 1 | arb | 728 | 7.129526029 | 6.31E-05 | 3 | 8.85E-06 | 0.420785335 | 8 mo | HOM |
| **1013978366** | 1 | arb | 728 | 7.129526029 | 0.000469515 | 6 | 6.59E-05 | 0.84157067 | 2 mo | HOM |
| **1014416155** | 2 | arb | 728 | 7.129526029 | 0 | 0 | 0 | 0 | 2 mo | HOM |
| **1013978065** | 2 | arb | 728 | 7.129526029 | 0.000405163 | 7 | 5.68E-05 | 0.981832449 | 2 mo | HOM |
| **1009934102** | 2 | arb | 728 | 7.129526029 | 2.17E-05 | 2 | 3.04E-06 | 0.280523557 | 4 mo | WILD |

| | | | | | | | | | | |
|---|---|---|---|---|---|---|---|---|---|---|
| **1009322937** | 2 | arb | 728 | 7.129526029 | 2E-05 | 1 | 2.81E-06 | 0.140261778 | 8 mo | WILD |
| **1011080637** | 2 | arb | 728 | 7.129526029 | 0.001556998 | 8 | 0.000218387 | 1.122094227 | 8 mo | HOM |
| **1013609713** | 2 | arb | 728 | 7.124121446 | 5.72E-06 | 0 | 8.04E-07 | 0 | 2 mo | WILD |
| **1010418842** | 1 | arb | 728 | 7.129526029 | 0.000757908 | 15 | 0.000106306 | 2.103926676 | 8 mo | HOM |
| **1013288213** | 2 | arb | 728 | 7.129526029 | 6.5E-06 | 0 | 9.12E-07 | 0 | 4 mo | HOM |
| **1008976060** | 2 | arb | 728 | 7.129526029 | 0.000907514 | 16 | 0.00012729 | 2.244188455 | 8 mo | WILD |
| **1013446243** | 1 | arb | 728 | 7.129526029 | 1.27E-05 | 2 | 1.79E-06 | 0.280523557 | 4 mo | HOM |
| **1013288951** | 2 | arb | 728 | 7.129526029 | 0.001384218 | 5 | 0.000194153 | 0.701308892 | 4 mo | HOM |
| **1013978665** | 1 | TEa6a | 729 | 0.539293922 | 0 | 0 | 0 | 0 | 2 mo | HOM |
| **1013445609** | 1 | TEa6a | 729 | 0.539293922 | 2.41E-05 | 2 | 4.46E-05 | 3.708552832 | 4 mo | HOM |
| **1013609015** | 1 | TEa6a | 729 | 0.539293922 | 0.000179534 | 8 | 0.000332905 | 14.83421133 | 4 mo | HOM |
| **1013608106** | 2 | TEa6a | 729 | 0.539293922 | 0 | 0 | 0 | 0 | 4 mo | HOM |
| **1011937543** | 1 | TEa6a | 729 | 0.539293922 | 0.000914114 | 15 | 0.001695021 | 27.81414624 | 8 mo | HOM |
| **1010419538** | 2 | TEa6a | 729 | 0.539293922 | 0 | 0 | 0 | 0 | 4 mo | WILD |
| **1010419187** | 1 | TEa6a | 729 | 0.539293922 | 0.000764514 | 34 | 0.001417619 | 63.04539814 | 8 mo | HOM |
| **1013978366** | 1 | TEa6a | 729 | 0.539293922 | 0 | 0 | 0 | 0 | 2 mo | HOM |
| **1014416155** | 2 | TEa6a | 729 | 0.539293922 | 0 | 0 | 0 | 0 | 2 mo | HOM |
| **1013978065** | 2 | TEa6a | 729 | 0.539293922 | 0 | 0 | 0 | 0 | 2 mo | HOM |
| **1009934102** | 2 | TEa6a | 729 | 0.539293922 | 0 | 0 | 0 | 0 | 4 mo | WILD |
| **1009322937** | 2 | TEa6a | 729 | 0.539293922 | 0 | 0 | 0 | 0 | 8 mo | WILD |
| **1011080637** | 2 | TEa6a | 729 | 0.539293922 | 0.000436784 | 14 | 0.000809918 | 25.95986982 | 8 mo | HOM |
| **1013609713** | 2 | TEa6a | 729 | 0.539293922 | 0 | 0 | 0 | 0 | 2 mo | WILD |
| **1010418842** | 1 | TEa6a | 729 | 0.539293922 | 0.000824494 | 30 | 0.001528839 | 55.62829247 | 8 mo | HOM |
| **1013288213** | 2 | TEa6a | 729 | 0.539293922 | 1.32E-05 | 1 | 2.44E-05 | 1.854276416 | 4 mo | HOM |
| **1008976060** | 2 | TEa6a | 729 | 0.539293922 | 0 | 0 | 0 | 0 | 8 mo | WILD |
| **1013446243** | 1 | TEa6a | 729 | 0.539293922 | 7.26E-05 | 2 | 0.000134532 | 3.708552832 | 4 mo | HOM |
| **1013288951** | 2 | TEa6a | 729 | 0.539293922 | 6.73E-05 | 5 | 0.000124874 | 9.271382079 | 4 mo | HOM |
| **1013978665** | 1 | ORBm | 731 | 1.441665961 | 3.36E-08 | 0 | 2.33E-08 | 0 | 2 mo | HOM |
| **1013445609** | 1 | ORBm | 731 | 1.441665961 | 5.99E-06 | 1 | 4.16E-06 | 0.693641958 | 4 mo | HOM |
| **1013609015** | 1 | ORBm | 731 | 1.441665961 | 0.000520921 | 11 | 0.000361333 | 7.63006154 | 4 mo | HOM |
| **1013608106** | 2 | ORBm | 731 | 1.441665961 | 0 | 0 | 0 | 0 | 4 mo | HOM |
| **1011937543** | 1 | ORBm | 731 | 1.441665961 | 0.002022333 | 35 | 0.001402775 | 24.27746854 | 8 mo | HOM |
| **1010419538** | 2 | ORBm | 731 | 1.441665961 | 0 | 0 | 0 | 0 | 4 mo | WILD |
| **1010419187** | 1 | ORBm | 731 | 1.441665961 | 0.001017328 | 39 | 0.000705662 | 27.05203637 | 8 mo | HOM |
| **1013978366** | 1 | ORBm | 731 | 1.441665961 | 6.68E-06 | 0 | 4.64E-06 | 0 | 2 mo | HOM |
| **1014416155** | 2 | ORBm | 731 | 1.441665961 | 1.66E-05 | 1 | 1.15E-05 | 0.693641958 | 2 mo | HOM |
| **1013978065** | 2 | ORBm | 731 | 1.441665961 | 3.57E-08 | 0 | 2.48E-08 | 0 | 2 mo | HOM |
| **1009934102** | 2 | ORBm | 731 | 1.441665961 | 1.5E-05 | 2 | 1.04E-05 | 1.387283916 | 4 mo | WILD |
| **1009322937** | 2 | ORBm | 731 | 1.441665961 | 7.41E-05 | 3 | 5.14E-05 | 2.080925874 | 8 mo | WILD |
| **1011080637** | 2 | ORBm | 731 | 1.441665961 | 0.002029742 | 42 | 0.001407914 | 29.13296224 | 8 mo | HOM |
| **1013609713** | 2 | ORBm | 731 | 1.441665961 | 0 | 0 | 0 | 0 | 2 mo | WILD |
| **1010418842** | 1 | ORBm | 731 | 1.441665961 | 0.002958763 | 32 | 0.002052322 | 22.19654266 | 8 mo | HOM |
| **1013288213** | 2 | ORBm | 731 | 1.441665961 | 0.002393457 | 18 | 0.001660203 | 12.48555525 | 4 mo | HOM |
| **1008976060** | 2 | ORBm | 731 | 1.441665961 | 6.95E-05 | 6 | 4.82E-05 | 4.161851749 | 8 mo | WILD |
| **1013446243** | 1 | ORBm | 731 | 1.441665961 | 9.7E-05 | 4 | 6.73E-05 | 2.774567833 | 4 mo | HOM |
| **1013288951** | 2 | ORBm | 731 | 1.441665961 | 0.000750865 | 26 | 0.000520831 | 18.03469091 | 4 mo | HOM |
| **1013978665** | 1 | MMme | 732 | 0.079284058 | 0 | 0 | 0 | 0 | 2 mo | HOM |
| **1013445609** | 1 | MMme | 732 | 0.079284058 | 0 | 0 | 0 | 0 | 4 mo | HOM |
| **1013609015** | 1 | MMme | 732 | 0.079284058 | 0 | 0 | 0 | 0 | 4 mo | HOM |
| **1013608106** | 2 | MMme | 732 | 0.079284058 | 0 | 0 | 0 | 0 | 4 mo | HOM |
| **1011937543** | 1 | MMme | 732 | 0.079284058 | 8.99E-05 | 1 | 0.001133556 | 12.6128761 | 8 mo | HOM |
| **1010419538** | 2 | MMme | 732 | 0.079284058 | 0 | 0 | 0 | 0 | 4 mo | WILD |
| **1010419187** | 1 | MMme | 732 | 0.079284058 | 5.36E-06 | 1 | 6.77E-05 | 12.6128761 | 8 mo | HOM |
| **1013978366** | 1 | MMme | 732 | 0.079284058 | 0 | 0 | 0 | 0 | 2 mo | HOM |
| **1014416155** | 2 | MMme | 732 | 0.079284058 | 0 | 0 | 0 | 0 | 2 mo | HOM |
| **1013978065** | 2 | MMme | 732 | 0.079284058 | 0 | 0 | 0 | 0 | 2 mo | HOM |
| **1009934102** | 2 | MMme | 732 | 0.079284058 | 0 | 0 | 0 | 0 | 4 mo | WILD |
| **1009322937** | 2 | MMme | 732 | 0.079284058 | 0 | 0 | 0 | 0 | 8 mo | WILD |
| **1011080637** | 2 | MMme | 732 | 0.079284058 | 0 | 0 | 0 | 0 | 8 mo | HOM |

| | | | | | | | | | | |
|---|---|---|---|---|---|---|---|---|---|---|
| 1013609713 | 2 | MMme | 732 | 0.079284058 | 0 | 0 | 0 | 0 | 2 mo | WILD |
| 1010418842 | 1 | MMme | 732 | 0.079284058 | 3.47E-06 | 0 | 4.37E-05 | 0 | 8 mo | HOM |
| 1013288213 | 2 | MMme | 732 | 0.079284058 | 0 | 0 | 0 | 0 | 4 mo | HOM |
| 1008976060 | 2 | MMme | 732 | 0.079284058 | 0 | 0 | 0 | 0 | 8 mo | WILD |
| 1013446243 | 1 | MMme | 732 | 0.079284058 | 0 | 0 | 0 | 0 | 4 mo | HOM |
| 1013288951 | 2 | MMme | 732 | 0.079284058 | 0 | 0 | 0 | 0 | 4 mo | HOM |
| 1013978665 | 1 | VPM | 733 | 1.694369824 | 0 | 0 | 0 | 0 | 2 mo | HOM |
| 1013445609 | 1 | VPM | 733 | 1.694369824 | 0 | 0 | 0 | 0 | 4 mo | HOM |
| 1013609015 | 1 | VPM | 733 | 1.694369824 | 5.19E-05 | 2 | 3.07E-05 | 1.180379851 | 4 mo | HOM |
| 1013608106 | 2 | VPM | 733 | 1.694369824 | 0 | 0 | 0 | 0 | 4 mo | HOM |
| 1011937543 | 1 | VPM | 733 | 1.694369824 | 0.000926708 | 6 | 0.000546934 | 3.541139552 | 8 mo | HOM |
| 1010419538 | 2 | VPM | 733 | 1.694369824 | 0 | 0 | 0 | 0 | 4 mo | WILD |
| 1010419187 | 1 | VPM | 733 | 1.694369824 | 0.000397186 | 9 | 0.000234415 | 5.311709328 | 8 mo | HOM |
| 1013978366 | 1 | VPM | 733 | 1.694369824 | 0 | 0 | 0 | 0 | 2 mo | HOM |
| 1014416155 | 2 | VPM | 733 | 1.694369824 | 0 | 0 | 0 | 0 | 2 mo | HOM |
| 1013978065 | 2 | VPM | 733 | 1.694369824 | 0 | 0 | 0 | 0 | 2 mo | HOM |
| 1009934102 | 2 | VPM | 733 | 1.694369824 | 0 | 0 | 0 | 0 | 4 mo | WILD |
| 1009322937 | 2 | VPM | 733 | 1.694369824 | 0 | 0 | 0 | 0 | 8 mo | WILD |
| 1011080637 | 2 | VPM | 733 | 1.694369824 | 0.00085725 | 17 | 0.00050594 | 10.03322873 | 8 mo | HOM |
| 1013609713 | 2 | VPM | 733 | 1.694369824 | 0 | 0 | 0 | 0 | 2 mo | WILD |
| 1010418842 | 1 | VPM | 733 | 1.694369824 | 0.000852068 | 15 | 0.000502882 | 8.852848881 | 8 mo | HOM |
| 1013288213 | 2 | VPM | 733 | 1.694369824 | 1.79E-05 | 2 | 1.06E-05 | 1.180379851 | 4 mo | HOM |
| 1008976060 | 2 | VPM | 733 | 1.694369824 | 2.81E-05 | 4 | 1.66E-05 | 2.360759702 | 8 mo | WILD |
| 1013446243 | 1 | VPM | 733 | 1.694369824 | 0 | 0 | 0 | 0 | 4 mo | HOM |
| 1013288951 | 2 | VPM | 733 | 1.694369824 | 5.09E-06 | 1 | 3E-06 | 0.590189925 | 4 mo | HOM |
| 1013978665 | 1 | AUDp1 | 735 | 0.386321998 | 5.9E-06 | 1 | 1.53E-05 | 2.588514261 | 2 mo | HOM |
| 1013445609 | 1 | AUDp1 | 735 | 0.386321998 | 4.56E-05 | 4 | 0.000118035 | 10.35405704 | 4 mo | HOM |
| 1013609015 | 1 | AUDp1 | 735 | 0.386321998 | 7.78E-05 | 3 | 0.000201304 | 7.765542782 | 4 mo | HOM |
| 1013608106 | 2 | AUDp1 | 735 | 0.386321998 | 0 | 0 | 0 | 0 | 4 mo | HOM |
| 1011937543 | 1 | AUDp1 | 735 | 0.386321998 | 0.00152837 | 19 | 0.003956207 | 49.18177095 | 8 mo | HOM |
| 1010419538 | 2 | AUDp1 | 735 | 0.386321998 | 0 | 0 | 0 | 0 | 4 mo | WILD |
| 1010419187 | 1 | AUDp1 | 735 | 0.386321998 | 0.001040627 | 22 | 0.002693677 | 56.94731373 | 8 mo | HOM |
| 1013978366 | 1 | AUDp1 | 735 | 0.386321998 | 0 | 0 | 0 | 0 | 2 mo | HOM |
| 1014416155 | 2 | AUDp1 | 735 | 0.386321998 | 0 | 0 | 0 | 0 | 2 mo | HOM |
| 1013978065 | 2 | AUDp1 | 735 | 0.386321998 | 0 | 0 | 0 | 0 | 2 mo | HOM |
| 1009934102 | 2 | AUDp1 | 735 | 0.386321998 | 1.07E-06 | 1 | 2.78E-06 | 2.588514261 | 4 mo | WILD |
| 1009322937 | 2 | AUDp1 | 735 | 0.386321998 | 1.6E-06 | 0 | 4.13E-06 | 0 | 8 mo | WILD |
| 1011080637 | 2 | AUDp1 | 735 | 0.386321998 | 0.000499871 | 20 | 0.001293923 | 51.77028521 | 8 mo | HOM |
| 1013609713 | 2 | AUDp1 | 735 | 0.386321998 | 1.28E-05 | 1 | 3.32E-05 | 2.588514261 | 2 mo | WILD |
| 1010418842 | 1 | AUDp1 | 735 | 0.386321998 | 0.001987221 | 18 | 0.00514395 | 46.59325669 | 8 mo | HOM |
| 1013288213 | 2 | AUDp1 | 735 | 0.386321998 | 9.59E-05 | 6 | 0.000248329 | 15.53108556 | 4 mo | HOM |
| 1008976060 | 2 | AUDp1 | 735 | 0.386321998 | 9.54E-06 | 2 | 2.47E-05 | 5.177028521 | 8 mo | WILD |
| 1013446243 | 1 | AUDp1 | 735 | 0.386321998 | 5.3E-05 | 3 | 0.000137232 | 7.765542782 | 4 mo | HOM |
| 1013288951 | 2 | AUDp1 | 735 | 0.386321998 | 0.000857344 | 13 | 0.002219247 | 33.65068539 | 4 mo | HOM |
| 1013978665 | 1 | fxpo | 737 | 0.286348976 | 4.87E-09 | 0 | 1.7E-08 | 0 | 2 mo | HOM |
| 1013445609 | 1 | fxpo | 737 | 0.286348976 | 5.59E-08 | 0 | 1.95E-07 | 0 | 4 mo | HOM |
| 1013609015 | 1 | fxpo | 737 | 0.286348976 | 4.18E-07 | 0 | 1.46E-06 | 0 | 4 mo | HOM |
| 1013608106 | 2 | fxpo | 737 | 0.286348976 | 0 | 0 | 0 | 0 | 4 mo | HOM |
| 1011937543 | 1 | fxpo | 737 | 0.286348976 | 4.42E-05 | 1 | 0.000154397 | 3.492242279 | 8 mo | HOM |
| 1010419538 | 2 | fxpo | 737 | 0.286348976 | 0 | 0 | 0 | 0 | 4 mo | WILD |
| 1010419187 | 1 | fxpo | 737 | 0.286348976 | 4.29E-06 | 0 | 1.5E-05 | 0 | 8 mo | HOM |
| 1013978366 | 1 | fxpo | 737 | 0.286348976 | 4.25E-08 | 0 | 1.48E-07 | 0 | 2 mo | HOM |
| 1014416155 | 2 | fxpo | 737 | 0.286348976 | 0 | 0 | 0 | 0 | 2 mo | HOM |
| 1013978065 | 2 | fxpo | 737 | 0.286348976 | 0 | 0 | 0 | 0 | 2 mo | HOM |
| 1009934102 | 2 | fxpo | 737 | 0.286348976 | 3.23E-06 | 0 | 1.13E-05 | 0 | 4 mo | WILD |
| 1009322937 | 2 | fxpo | 737 | 0.286348976 | 0 | 0 | 0 | 0 | 8 mo | WILD |
| 1011080637 | 2 | fxpo | 737 | 0.286348976 | 6.07E-05 | 0 | 0.000212095 | 0 | 8 mo | HOM |
| 1013609713 | 2 | fxpo | 737 | 0.286348976 | 1.56E-06 | 0 | 5.43E-06 | 0 | 2 mo | WILD |
| 1010418842 | 1 | fxpo | 737 | 0.286348976 | 0.0001054 | 4 | 0.000368083 | 13.96896912 | 8 mo | HOM |

| | | | | | | | | | | |
|---|---|---|---|---|---|---|---|---|---|---|
| 1013288213 | 2 | fxpo | 737 | 0.286348976 | 1.45E-06 | 0 | 5.08E-06 | 0 | 4 mo | HOM |
| 1008976060 | 2 | fxpo | 737 | 0.286348976 | 2.22E-05 | 0 | 7.75E-05 | 0 | 8 mo | WILD |
| 1013446243 | 1 | fxpo | 737 | 0.286348976 | 2.38E-06 | 0 | 8.3E-06 | 0 | 4 mo | HOM |
| 1013288951 | 2 | fxpo | 737 | 0.286348976 | 4.93E-05 | 0 | 0.000172296 | 0 | 4 mo | HOM |
| 1013978665 | 1 | VPMpc | 741 | 0.22176418 | 0 | 0 | 0 | 0 | 2 mo | HOM |
| 1013445609 | 1 | VPMpc | 741 | 0.22176418 | 0 | 0 | 0 | 0 | 4 mo | HOM |
| 1013609015 | 1 | VPMpc | 741 | 0.22176418 | 0 | 0 | 0 | 0 | 4 mo | HOM |
| 1013608106 | 2 | VPMpc | 741 | 0.22176418 | 0 | 0 | 0 | 0 | 4 mo | HOM |
| 1011937543 | 1 | VPMpc | 741 | 0.22176418 | 2.87E-05 | 0 | 0.000129608 | 0 | 8 mo | HOM |
| 1010419538 | 2 | VPMpc | 741 | 0.22176418 | 0 | 0 | 0 | 0 | 4 mo | WILD |
| 1010419187 | 1 | VPMpc | 741 | 0.22176418 | 0.000118778 | 4 | 0.000535603 | 18.03717802 | 8 mo | HOM |
| 1013978366 | 1 | VPMpc | 741 | 0.22176418 | 0 | 0 | 0 | 0 | 2 mo | HOM |
| 1014416155 | 2 | VPMpc | 741 | 0.22176418 | 0 | 0 | 0 | 0 | 2 mo | HOM |
| 1013978065 | 2 | VPMpc | 741 | 0.22176418 | 0 | 0 | 0 | 0 | 2 mo | HOM |
| 1009934102 | 2 | VPMpc | 741 | 0.22176418 | 0 | 0 | 0 | 0 | 4 mo | WILD |
| 1009322937 | 2 | VPMpc | 741 | 0.22176418 | 0 | 0 | 0 | 0 | 8 mo | WILD |
| 1011080637 | 2 | VPMpc | 741 | 0.22176418 | 5.16E-06 | 0 | 2.33E-05 | 0 | 8 mo | HOM |
| 1013609713 | 2 | VPMpc | 741 | 0.22176418 | 0 | 0 | 0 | 0 | 2 mo | WILD |
| 1010418842 | 1 | VPMpc | 741 | 0.22176418 | 0.000185321 | 7 | 0.000835669 | 31.56506153 | 8 mo | HOM |
| 1013288213 | 2 | VPMpc | 741 | 0.22176418 | 0 | 0 | 0 | 0 | 4 mo | HOM |
| 1008976060 | 2 | VPMpc | 741 | 0.22176418 | 0 | 0 | 0 | 0 | 8 mo | WILD |
| 1013446243 | 1 | VPMpc | 741 | 0.22176418 | 0 | 0 | 0 | 0 | 4 mo | HOM |
| 1013288951 | 2 | VPMpc | 741 | 0.22176418 | 0 | 0 | 0 | 0 | 4 mo | HOM |
| 1013978665 | 1 | ENTm6 | 743 | 0.641407766 | 0 | 0 | 0 | 0 | 2 mo | HOM |
| 1013445609 | 1 | ENTm6 | 743 | 0.641407766 | 9.4E-12 | 0 | 1.47E-11 | 0 | 4 mo | HOM |
| 1013609015 | 1 | ENTm6 | 743 | 0.641407766 | 5.47E-05 | 1 | 8.54E-05 | 1.559070616 | 4 mo | HOM |
| 1013608106 | 2 | ENTm6 | 743 | 0.641407766 | 0 | 0 | 0 | 0 | 4 mo | HOM |
| 1011937543 | 1 | ENTm6 | 743 | 0.641407766 | 0.000866863 | 10 | 0.001351501 | 15.59070616 | 8 mo | HOM |
| 1010419538 | 2 | ENTm6 | 743 | 0.641407766 | 0 | 0 | 0 | 0 | 4 mo | WILD |
| 1010419187 | 1 | ENTm6 | 743 | 0.641407766 | 0.000438121 | 10 | 0.000683062 | 15.59070616 | 8 mo | HOM |
| 1013978366 | 1 | ENTm6 | 743 | 0.641407766 | 0 | 0 | 0 | 0 | 2 mo | HOM |
| 1014416155 | 2 | ENTm6 | 743 | 0.641407766 | 0 | 0 | 0 | 0 | 2 mo | HOM |
| 1013978065 | 2 | ENTm6 | 743 | 0.641407766 | 0 | 0 | 0 | 0 | 2 mo | HOM |
| 1009934102 | 2 | ENTm6 | 743 | 0.641407766 | 0 | 0 | 0 | 0 | 4 mo | WILD |
| 1009322937 | 2 | ENTm6 | 743 | 0.641407766 | 0 | 0 | 0 | 0 | 8 mo | WILD |
| 1011080637 | 2 | ENTm6 | 743 | 0.641407766 | 0.000184378 | 6 | 0.000287459 | 9.354423694 | 8 mo | HOM |
| 1013609713 | 2 | ENTm6 | 743 | 0.641407766 | 0 | 0 | 0 | 0 | 2 mo | WILD |
| 1010418842 | 1 | ENTm6 | 743 | 0.641407766 | 0.000355862 | 16 | 0.000554814 | 24.94512985 | 8 mo | HOM |
| 1013288213 | 2 | ENTm6 | 743 | 0.641407766 | 4.45E-07 | 0 | 6.93E-07 | 0 | 4 mo | HOM |
| 1008976060 | 2 | ENTm6 | 743 | 0.641407766 | 0 | 0 | 0 | 0 | 8 mo | WILD |
| 1013446243 | 1 | ENTm6 | 743 | 0.641407766 | 2.66E-05 | 2 | 4.15E-05 | 3.118141231 | 4 mo | HOM |
| 1013288951 | 2 | ENTm6 | 743 | 0.641407766 | 6.18E-05 | 3 | 9.64E-05 | 4.677211847 | 4 mo | HOM |
| 1013978665 | 1 | cbc | 744 | 0.068959141 | 0 | 0 | 0 | 0 | 2 mo | HOM |
| 1013445609 | 1 | cbc | 744 | 0.068959141 | 0 | 0 | 0 | 0 | 4 mo | HOM |
| 1013609015 | 1 | cbc | 744 | 0.068959141 | 0 | 0 | 0 | 0 | 4 mo | HOM |
| 1013608106 | 2 | cbc | 744 | 0.068959141 | 0 | 0 | 0 | 0 | 4 mo | HOM |
| 1011937543 | 1 | cbc | 744 | 0.068959141 | 0 | 0 | 0 | 0 | 8 mo | HOM |
| 1010419538 | 2 | cbc | 744 | 0.061372828 | 0 | 0 | 0 | 0 | 4 mo | WILD |
| 1010419187 | 1 | cbc | 744 | 0.068959141 | 0 | 0 | 0 | 0 | 8 mo | HOM |
| 1013978366 | 1 | cbc | 744 | 0.068959141 | 0 | 0 | 0 | 0 | 2 mo | HOM |
| 1014416155 | 2 | cbc | 744 | 0.068959141 | 0 | 0 | 0 | 0 | 2 mo | HOM |
| 1013978065 | 2 | cbc | 744 | 0.068959141 | 0.00020781 | 1 | 0.003013523 | 14.50134081 | 2 mo | HOM |
| 1009934102 | 2 | cbc | 744 | 0.068959141 | 0 | 0 | 0 | 0 | 4 mo | WILD |
| 1009322937 | 2 | cbc | 744 | 0.068959141 | 0 | 0 | 0 | 0 | 8 mo | WILD |
| 1011080637 | 2 | cbc | 744 | 0.068959141 | 1.07E-05 | 0 | 0.000155072 | 0 | 8 mo | HOM |
| 1013609713 | 2 | cbc | 744 | 0.068959141 | 9.92E-07 | 0 | 1.44E-05 | 0 | 2 mo | WILD |
| 1010418842 | 1 | cbc | 744 | 0.068959141 | 0 | 0 | 0 | 0 | 8 mo | HOM |
| 1013288213 | 2 | cbc | 744 | 0.068959141 | 0 | 0 | 0 | 0 | 4 mo | HOM |
| 1008976060 | 2 | cbc | 744 | 0.068959141 | 6.84E-05 | 1 | 0.000991801 | 14.50134081 | 8 mo | WILD |

| | | | | | | | | | | |
|---|---|---|---|---|---|---|---|---|---|---|
| **1013446243** | 1 | cbc | 744 | 0.068959141 | 0 | 0 | 0 | 0 | 4 mo | HOM |
| **1013288951** | 2 | cbc | 744 | 0.068959141 | 2.5E-09 | 0 | 3.63E-08 | 0 | 4 mo | HOM |
| **1013978665** | 1 | ORBvl | 746 | 1.830489352 | 1.33E-05 | 0 | 7.27E-06 | 0 | 2 mo | HOM |
| **1013445609** | 1 | ORBvl | 746 | 1.830489352 | 9E-05 | 4 | 4.92E-05 | 2.185208013 | 4 mo | HOM |
| **1013609015** | 1 | ORBvl | 746 | 1.830489352 | 0.000268831 | 9 | 0.000146863 | 4.91671803 | 4 mo | HOM |
| **1013608106** | 2 | ORBvl | 746 | 1.830489352 | 0 | 0 | 0 | 0 | 4 mo | HOM |
| **1011937543** | 1 | ORBvl | 746 | 1.830489352 | 0.001387579 | 41 | 0.000758037 | 22.39838214 | 8 mo | HOM |
| **1010419538** | 2 | ORBvl | 746 | 1.830489352 | 0 | 0 | 0 | 0 | 4 mo | WILD |
| **1010419187** | 1 | ORBvl | 746 | 1.830489352 | 0.001577252 | 57 | 0.000861656 | 31.13921419 | 8 mo | HOM |
| **1013978366** | 1 | ORBvl | 746 | 1.830489352 | 0 | 0 | 0 | 0 | 2 mo | HOM |
| **1014416155** | 2 | ORBvl | 746 | 1.830489352 | 0 | 0 | 0 | 0 | 2 mo | HOM |
| **1013978065** | 2 | ORBvl | 746 | 1.830489352 | 3.17E-06 | 0 | 1.73E-06 | 0 | 2 mo | HOM |
| **1009934102** | 2 | ORBvl | 746 | 1.830489352 | 0 | 0 | 0 | 0 | 4 mo | WILD |
| **1009322937** | 2 | ORBvl | 746 | 1.830489352 | 8.4E-05 | 8 | 4.59E-05 | 4.370416026 | 8 mo | WILD |
| **1011080637** | 2 | ORBvl | 746 | 1.830489352 | 0.001325558 | 41 | 0.000724155 | 22.39838214 | 8 mo | HOM |
| **1013609713** | 2 | ORBvl | 746 | 1.830489352 | 0 | 0 | 0 | 0 | 2 mo | WILD |
| **1010418842** | 1 | ORBvl | 746 | 1.830489352 | 0.001993208 | 60 | 0.001088894 | 32.7781202 | 8 mo | HOM |
| **1013288213** | 2 | ORBvl | 746 | 1.830489352 | 0.001065404 | 6 | 0.000582032 | 3.27781202 | 4 mo | HOM |
| **1008976060** | 2 | ORBvl | 746 | 1.830489352 | 5.05E-06 | 1 | 2.76E-06 | 0.546302003 | 8 mo | WILD |
| **1013446243** | 1 | ORBvl | 746 | 1.830489352 | 0.000152846 | 5 | 8.35E-05 | 2.731510017 | 4 mo | HOM |
| **1013288951** | 2 | ORBvl | 746 | 1.830489352 | 0.000566687 | 16 | 0.000309582 | 8.740832053 | 4 mo | HOM |
| **1013978665** | 1 | VTA | 749 | 0.441572986 | 0 | 0 | 0 | 0 | 2 mo | HOM |
| **1013445609** | 1 | VTA | 749 | 0.441572986 | 0 | 0 | 0 | 0 | 4 mo | HOM |
| **1013609015** | 1 | VTA | 749 | 0.441572986 | 2.34E-07 | 0 | 5.3E-07 | 0 | 4 mo | HOM |
| **1013608106** | 2 | VTA | 749 | 0.441572986 | 0 | 0 | 0 | 0 | 4 mo | HOM |
| **1011937543** | 1 | VTA | 749 | 0.441572986 | 0.000145298 | 4 | 0.000329047 | 9.058525144 | 8 mo | HOM |
| **1010419538** | 2 | VTA | 749 | 0.441572986 | 0 | 0 | 0 | 0 | 4 mo | WILD |
| **1010419187** | 1 | VTA | 749 | 0.441572986 | 4.05E-06 | 0 | 9.17E-06 | 0 | 8 mo | HOM |
| **1013978366** | 1 | VTA | 749 | 0.441572986 | 0 | 0 | 0 | 0 | 2 mo | HOM |
| **1014416155** | 2 | VTA | 749 | 0.441572986 | 0 | 0 | 0 | 0 | 2 mo | HOM |
| **1013978065** | 2 | VTA | 749 | 0.441572986 | 0 | 0 | 0 | 0 | 2 mo | HOM |
| **1009934102** | 2 | VTA | 749 | 0.441572986 | 0 | 0 | 0 | 0 | 4 mo | WILD |
| **1009322937** | 2 | VTA | 749 | 0.441572986 | 0 | 0 | 0 | 0 | 8 mo | WILD |
| **1011080637** | 2 | VTA | 749 | 0.441572986 | 1.17E-05 | 1 | 2.65E-05 | 2.264631286 | 8 mo | HOM |
| **1013609713** | 2 | VTA | 749 | 0.441572986 | 0 | 0 | 0 | 0 | 2 mo | WILD |
| **1010418842** | 1 | VTA | 749 | 0.441572986 | 2.29E-05 | 1 | 5.19E-05 | 2.264631286 | 8 mo | HOM |
| **1013288213** | 2 | VTA | 749 | 0.441572986 | 0 | 0 | 0 | 0 | 4 mo | HOM |
| **1008976060** | 2 | VTA | 749 | 0.441572986 | 1.31E-05 | 1 | 2.97E-05 | 2.264631286 | 8 mo | WILD |
| **1013446243** | 1 | VTA | 749 | 0.441572986 | 0 | 0 | 0 | 0 | 4 mo | HOM |
| **1013288951** | 2 | VTA | 749 | 0.441572986 | 0 | 0 | 0 | 0 | 4 mo | HOM |
| **1013978665** | 1 | VISpl1 | 750 | 0.199647305 | 9.44E-06 | 1 | 4.73E-05 | 5.008832947 | 2 mo | HOM |
| **1013445609** | 1 | VISpl1 | 750 | 0.199647305 | 8.27E-06 | 0 | 4.14E-05 | 0 | 4 mo | HOM |
| **1013609015** | 1 | VISpl1 | 750 | 0.199647305 | 4.46E-05 | 3 | 0.000223172 | 15.02649884 | 4 mo | HOM |
| **1013608106** | 2 | VISpl1 | 750 | 0.199647305 | 0 | 0 | 0 | 0 | 4 mo | HOM |
| **1011937543** | 1 | VISpl1 | 750 | 0.199647305 | 0.004282471 | 6 | 0.021450181 | 30.05299768 | 8 mo | HOM |
| **1010419538** | 2 | VISpl1 | 750 | 0.199647305 | 5.04E-05 | 2 | 0.000252615 | 10.01766589 | 4 mo | WILD |
| **1010419187** | 1 | VISpl1 | 750 | 0.199647305 | 0.003199944 | 8 | 0.016027987 | 40.07066357 | 8 mo | HOM |
| **1013978366** | 1 | VISpl1 | 750 | 0.199647305 | 0.000231724 | 6 | 0.001160668 | 30.05299768 | 2 mo | HOM |
| **1014416155** | 2 | VISpl1 | 750 | 0.199647305 | 0 | 0 | 0 | 0 | 2 mo | HOM |
| **1013978065** | 2 | VISpl1 | 750 | 0.199647305 | 5.36E-06 | 0 | 2.68E-05 | 0 | 2 mo | HOM |
| **1009934102** | 2 | VISpl1 | 750 | 0.199647305 | 8.97E-06 | 2 | 4.49E-05 | 10.01766589 | 4 mo | WILD |
| **1009322937** | 2 | VISpl1 | 750 | 0.199647305 | 6.58E-06 | 1 | 3.29E-05 | 5.008832947 | 8 mo | WILD |
| **1011080637** | 2 | VISpl1 | 750 | 0.199647305 | 0.000244336 | 11 | 0.001223838 | 55.09716241 | 8 mo | HOM |
| **1013609713** | 2 | VISpl1 | 750 | 0.199647305 | 1.88E-05 | 1 | 9.43E-05 | 5.008832947 | 2 mo | WILD |
| **1010418842** | 1 | VISpl1 | 750 | 0.199647305 | 0.000335467 | 6 | 0.001680301 | 30.05299768 | 8 mo | HOM |
| **1013288213** | 2 | VISpl1 | 750 | 0.199647305 | 7.16E-06 | 1 | 3.58E-05 | 5.008832947 | 4 mo | HOM |
| **1008976060** | 2 | VISpl1 | 750 | 0.199647305 | 0 | 0 | 0 | 0 | 8 mo | WILD |
| **1013446243** | 1 | VISpl1 | 750 | 0.199647305 | 0.000182823 | 5 | 0.000915728 | 25.04416473 | 4 mo | HOM |
| **1013288951** | 2 | VISpl1 | 750 | 0.199647305 | 0.002282956 | 3 | 0.011434947 | 15.02649884 | 4 mo | HOM |

| | | | | | | | | | | |
|---|---|---|---|---|---|---|---|---|---|---|
| 1013978665 | 1 | cbp | 752 | 2.904360889 | 0 | 0 | 0 | 0 | 2 mo | HOM |
| 1013445609 | 1 | cbp | 752 | 2.904360889 | 2.64E-06 | 0 | 9.08E-07 | 0 | 4 mo | HOM |
| 1013609015 | 1 | cbp | 752 | 2.904360889 | 3.6E-06 | 0 | 1.24E-06 | 0 | 4 mo | HOM |
| 1013608106 | 2 | cbp | 752 | 2.904360889 | 0 | 0 | 0 | 0 | 4 mo | HOM |
| 1011937543 | 1 | cbp | 752 | 2.904360889 | 0.004499185 | 20 | 0.001549113 | 6.886196573 | 8 mo | HOM |
| 1010419538 | 2 | cbp | 752 | 2.904360889 | 0 | 0 | 0 | 0 | 4 mo | WILD |
| 1010419187 | 1 | cbp | 752 | 2.904360889 | 3.18E-05 | 1 | 1.1E-05 | 0.344309829 | 8 mo | HOM |
| 1013978366 | 1 | cbp | 752 | 2.904360889 | 1.71E-06 | 0 | 5.88E-07 | 0 | 2 mo | HOM |
| 1014416155 | 2 | cbp | 752 | 2.904360889 | 0 | 0 | 0 | 0 | 2 mo | HOM |
| 1013978065 | 2 | cbp | 752 | 2.904360889 | 0 | 0 | 0 | 0 | 2 mo | HOM |
| 1009934102 | 2 | cbp | 752 | 2.904360889 | 6.01E-06 | 0 | 2.07E-06 | 0 | 4 mo | WILD |
| 1009322937 | 2 | cbp | 752 | 2.904360889 | 8.62E-06 | 0 | 2.97E-06 | 0 | 8 mo | WILD |
| 1011080637 | 2 | cbp | 752 | 2.904360889 | 1.26E-05 | 1 | 4.32E-06 | 0.344309829 | 8 mo | HOM |
| 1013609713 | 2 | cbp | 752 | 2.904360889 | 3.65E-10 | 0 | 1.26E-10 | 0 | 2 mo | WILD |
| 1010418842 | 1 | cbp | 752 | 2.904360889 | 0.00024457 | 3 | 8.42E-05 | 1.032929486 | 8 mo | HOM |
| 1013288213 | 2 | cbp | 752 | 2.904360889 | 0 | 0 | 0 | 0 | 4 mo | HOM |
| 1008976060 | 2 | cbp | 752 | 2.904360889 | 4.14E-05 | 2 | 1.43E-05 | 0.688619657 | 8 mo | WILD |
| 1013446243 | 1 | cbp | 752 | 2.904360889 | 7.42E-06 | 0 | 2.55E-06 | 0 | 4 mo | HOM |
| 1013288951 | 2 | cbp | 752 | 2.904360889 | 1.15E-06 | 0 | 3.95E-07 | 0 | 4 mo | HOM |
| 1013978665 | 1 | pm | 753 | 0.034304432 | 0 | 0 | 0 | 0 | 2 mo | HOM |
| 1013445609 | 1 | pm | 753 | 0.034304432 | 0 | 0 | 0 | 0 | 4 mo | HOM |
| 1013609015 | 1 | pm | 753 | 0.034304432 | 0 | 0 | 0 | 0 | 4 mo | HOM |
| 1013608106 | 2 | pm | 753 | 0.034304432 | 0 | 0 | 0 | 0 | 4 mo | HOM |
| 1011937543 | 1 | pm | 753 | 0.034304432 | 0 | 0 | 0 | 0 | 8 mo | HOM |
| 1010419538 | 2 | pm | 753 | 0.034304432 | 0 | 0 | 0 | 0 | 4 mo | WILD |
| 1010419187 | 1 | pm | 753 | 0.034304432 | 0 | 0 | 0 | 0 | 8 mo | HOM |
| 1013978366 | 1 | pm | 753 | 0.034304432 | 0 | 0 | 0 | 0 | 2 mo | HOM |
| 1014416155 | 2 | pm | 753 | 0.034304432 | 0 | 0 | 0 | 0 | 2 mo | HOM |
| 1013978065 | 2 | pm | 753 | 0.034304432 | 0 | 0 | 0 | 0 | 2 mo | HOM |
| 1009934102 | 2 | pm | 753 | 0.034304432 | 0 | 0 | 0 | 0 | 4 mo | WILD |
| 1009322937 | 2 | pm | 753 | 0.034304432 | 0 | 0 | 0 | 0 | 8 mo | WILD |
| 1011080637 | 2 | pm | 753 | 0.034304432 | 0 | 0 | 0 | 0 | 8 mo | HOM |
| 1013609713 | 2 | pm | 753 | 0.034304432 | 0 | 0 | 0 | 0 | 2 mo | WILD |
| 1010418842 | 1 | pm | 753 | 0.034304432 | 0 | 0 | 0 | 0 | 8 mo | HOM |
| 1013288213 | 2 | pm | 753 | 0.034304432 | 0 | 0 | 0 | 0 | 4 mo | HOM |
| 1008976060 | 2 | pm | 753 | 0.034304432 | 0 | 0 | 0 | 0 | 8 mo | WILD |
| 1013446243 | 1 | pm | 753 | 0.034304432 | 0 | 0 | 0 | 0 | 4 mo | HOM |
| 1013288951 | 2 | pm | 753 | 0.034304432 | 0 | 0 | 0 | 0 | 4 mo | HOM |
| 1013978665 | 1 | OT | 754 | 3.939256922 | 0 | 0 | 0 | 0 | 2 mo | HOM |
| 1013445609 | 1 | OT | 754 | 3.939256922 | 0 | 0 | 0 | 0 | 4 mo | HOM |
| 1013609015 | 1 | OT | 754 | 3.939256922 | 8.39E-05 | 2 | 2.13E-05 | 0.507709967 | 4 mo | HOM |
| 1013608106 | 2 | OT | 754 | 3.939256922 | 0 | 0 | 0 | 0 | 4 mo | HOM |
| 1011937543 | 1 | OT | 754 | 3.939256922 | 0.001529601 | 35 | 0.000388297 | 8.884924415 | 8 mo | HOM |
| 1010419538 | 2 | OT | 754 | 3.939256922 | 5.33E-06 | 1 | 1.35E-06 | 0.253854983 | 4 mo | WILD |
| 1010419187 | 1 | OT | 754 | 3.939256922 | 0.000513368 | 24 | 0.000130321 | 6.092519599 | 8 mo | HOM |
| 1013978366 | 1 | OT | 754 | 3.939256922 | 0 | 0 | 0 | 0 | 2 mo | HOM |
| 1014416155 | 2 | OT | 754 | 3.939256922 | 0 | 0 | 0 | 0 | 2 mo | HOM |
| 1013978065 | 2 | OT | 754 | 3.939256922 | 0 | 0 | 0 | 0 | 2 mo | HOM |
| 1009934102 | 2 | OT | 754 | 3.939256922 | 9.72E-07 | 0 | 2.47E-07 | 0 | 4 mo | WILD |
| 1009322937 | 2 | OT | 754 | 3.939256922 | 0 | 0 | 0 | 0 | 8 mo | WILD |
| 1011080637 | 2 | OT | 754 | 3.939256922 | 0.000313292 | 15 | 7.95E-05 | 3.807824749 | 8 mo | HOM |
| 1013609713 | 2 | OT | 754 | 3.939256922 | 0 | 0 | 0 | 0 | 2 mo | WILD |
| 1010418842 | 1 | OT | 754 | 3.939256922 | 0.000574161 | 42 | 0.000145754 | 10.6619093 | 8 mo | HOM |
| 1013288213 | 2 | OT | 754 | 3.939256922 | 4.51E-05 | 0 | 1.14E-05 | 0 | 4 mo | HOM |
| 1008976060 | 2 | OT | 754 | 3.939256922 | 8.46E-06 | 1 | 2.15E-06 | 0.253854983 | 8 mo | WILD |
| 1013446243 | 1 | OT | 754 | 3.939256922 | 2.44E-10 | 0 | 6.2E-11 | 0 | 4 mo | HOM |
| 1013288951 | 2 | OT | 754 | 3.939256922 | 8.58E-05 | 3 | 2.18E-05 | 0.76156495 | 4 mo | HOM |
| 1013978665 | 1 | AUDv2/3 | 755 | 0.399187477 | 0 | 0 | 0 | 0 | 2 mo | HOM |
| 1013445609 | 1 | AUDv2/3 | 755 | 0.399187477 | 1.55E-07 | 0 | 3.89E-07 | 0 | 4 mo | HOM |

| | | | | | | | | | | |
|---|---|---|---|---|---|---|---|---|---|---|
| **1013609015** | 1 | AUDv2/3 | 755 | 0.399187477 | 2.95E-05 | 2 | 7.39E-05 | 5.010177213 | 4 mo | HOM |
| **1013608106** | 2 | AUDv2/3 | 755 | 0.399187477 | 0 | 0 | 0 | 0 | 4 mo | HOM |
| **1011937543** | 1 | AUDv2/3 | 755 | 0.399187477 | 0.000243775 | 8 | 0.000610678 | 20.04070885 | 8 mo | HOM |
| **1010419538** | 2 | AUDv2/3 | 755 | 0.399187477 | 0 | 0 | 0 | 0 | 4 mo | WILD |
| **1010419187** | 1 | AUDv2/3 | 755 | 0.399187477 | 0.000498785 | 13 | 0.0012495 | 32.56615188 | 8 mo | HOM |
| **1013978366** | 1 | AUDv2/3 | 755 | 0.399187477 | 0 | 0 | 0 | 0 | 2 mo | HOM |
| **1014416155** | 2 | AUDv2/3 | 755 | 0.399187477 | 0 | 0 | 0 | 0 | 2 mo | HOM |
| **1013978065** | 2 | AUDv2/3 | 755 | 0.399187477 | 0 | 0 | 0 | 0 | 2 mo | HOM |
| **1009934102** | 2 | AUDv2/3 | 755 | 0.399187477 | 0 | 0 | 0 | 0 | 4 mo | WILD |
| **1009322937** | 2 | AUDv2/3 | 755 | 0.399187477 | 0 | 0 | 0 | 0 | 8 mo | WILD |
| **1011080637** | 2 | AUDv2/3 | 755 | 0.399187477 | 0.000254485 | 7 | 0.000637509 | 17.53562025 | 8 mo | HOM |
| **1013609713** | 2 | AUDv2/3 | 755 | 0.399187477 | 0 | 0 | 0 | 0 | 2 mo | WILD |
| **1010418842** | 1 | AUDv2/3 | 755 | 0.399187477 | 0.000350784 | 13 | 0.000878745 | 32.56615188 | 8 mo | HOM |
| **1013288213** | 2 | AUDv2/3 | 755 | 0.399187477 | 5.81E-05 | 3 | 0.000145482 | 7.51526582 | 4 mo | HOM |
| **1008976060** | 2 | AUDv2/3 | 755 | 0.399187477 | 0 | 0 | 0 | 0 | 8 mo | WILD |
| **1013446243** | 1 | AUDv2/3 | 755 | 0.399187477 | 3.94E-05 | 2 | 9.88E-05 | 5.010177213 | 4 mo | HOM |
| **1013288951** | 2 | AUDv2/3 | 755 | 0.399187477 | 4.63E-06 | 0 | 1.16E-05 | 0 | 4 mo | HOM |
| **1013978665** | 1 | VTN | 757 | 0.038664344 | 0 | 0 | 0 | 0 | 2 mo | HOM |
| **1013445609** | 1 | VTN | 757 | 0.038664344 | 0 | 0 | 0 | 0 | 4 mo | HOM |
| **1013609015** | 1 | VTN | 757 | 0.038664344 | 0 | 0 | 0 | 0 | 4 mo | HOM |
| **1013608106** | 2 | VTN | 757 | 0.038664344 | 0 | 0 | 0 | 0 | 4 mo | HOM |
| **1011937543** | 1 | VTN | 757 | 0.038664344 | 0 | 0 | 0 | 0 | 8 mo | HOM |
| **1010419538** | 2 | VTN | 757 | 0.038664344 | 0 | 0 | 0 | 0 | 4 mo | WILD |
| **1010419187** | 1 | VTN | 757 | 0.038664344 | 0 | 0 | 0 | 0 | 8 mo | HOM |
| **1013978366** | 1 | VTN | 757 | 0.038664344 | 0 | 0 | 0 | 0 | 2 mo | HOM |
| **1014416155** | 2 | VTN | 757 | 0.038664344 | 0 | 0 | 0 | 0 | 2 mo | HOM |
| **1013978065** | 2 | VTN | 757 | 0.038664344 | 0 | 0 | 0 | 0 | 2 mo | HOM |
| **1009934102** | 2 | VTN | 757 | 0.038664344 | 0 | 0 | 0 | 0 | 4 mo | WILD |
| **1009322937** | 2 | VTN | 757 | 0.038664344 | 0 | 0 | 0 | 0 | 8 mo | WILD |
| **1011080637** | 2 | VTN | 757 | 0.038664344 | 0 | 0 | 0 | 0 | 8 mo | HOM |
| **1013609713** | 2 | VTN | 757 | 0.038664344 | 0 | 0 | 0 | 0 | 2 mo | WILD |
| **1010418842** | 1 | VTN | 757 | 0.038664344 | 0 | 0 | 0 | 0 | 8 mo | HOM |
| **1013288213** | 2 | VTN | 757 | 0.038664344 | 0 | 0 | 0 | 0 | 4 mo | HOM |
| **1008976060** | 2 | VTN | 757 | 0.038664344 | 0 | 0 | 0 | 0 | 8 mo | WILD |
| **1013446243** | 1 | VTN | 757 | 0.038664344 | 0 | 0 | 0 | 0 | 4 mo | HOM |
| **1013288951** | 2 | VTN | 757 | 0.038664344 | 0 | 0 | 0 | 0 | 4 mo | HOM |
| **1013978665** | 1 | AUDpo4 | 759 | 0.076440637 | 0 | 0 | 0 | 0 | 2 mo | HOM |
| **1013445609** | 1 | AUDpo4 | 759 | 0.076440637 | 1.41E-06 | 0 | 1.85E-05 | 0 | 4 mo | HOM |
| **1013609015** | 1 | AUDpo4 | 759 | 0.076440637 | 3.43E-05 | 1 | 0.000448576 | 13.08204692 | 4 mo | HOM |
| **1013608106** | 2 | AUDpo4 | 759 | 0.076440637 | 0 | 0 | 0 | 0 | 4 mo | HOM |
| **1011937543** | 1 | AUDpo4 | 759 | 0.076440637 | 0.000154458 | 5 | 0.002020622 | 65.41023461 | 8 mo | HOM |
| **1010419538** | 2 | AUDpo4 | 759 | 0.076440637 | 0 | 0 | 0 | 0 | 4 mo | WILD |
| **1010419187** | 1 | AUDpo4 | 759 | 0.076440637 | 0.000161929 | 4 | 0.002118356 | 52.32818769 | 8 mo | HOM |
| **1013978366** | 1 | AUDpo4 | 759 | 0.076440637 | 0 | 0 | 0 | 0 | 2 mo | HOM |
| **1014416155** | 2 | AUDpo4 | 759 | 0.076440637 | 0 | 0 | 0 | 0 | 2 mo | HOM |
| **1013978065** | 2 | AUDpo4 | 759 | 0.076440637 | 0 | 0 | 0 | 0 | 2 mo | HOM |
| **1009934102** | 2 | AUDpo4 | 759 | 0.076440637 | 0 | 0 | 0 | 0 | 4 mo | WILD |
| **1009322937** | 2 | AUDpo4 | 759 | 0.076440637 | 0 | 0 | 0 | 0 | 8 mo | WILD |
| **1011080637** | 2 | AUDpo4 | 759 | 0.076440637 | 6.08E-05 | 2 | 0.000794862 | 26.16409384 | 8 mo | HOM |
| **1013609713** | 2 | AUDpo4 | 759 | 0.076440637 | 0 | 0 | 0 | 0 | 2 mo | WILD |
| **1010418842** | 1 | AUDpo4 | 759 | 0.076440637 | 0.000132781 | 4 | 0.001737048 | 52.32818769 | 8 mo | HOM |
| **1013288213** | 2 | AUDpo4 | 759 | 0.076440637 | 1.4E-06 | 0 | 1.83E-05 | 0 | 4 mo | HOM |
| **1008976060** | 2 | AUDpo4 | 759 | 0.076440637 | 0 | 0 | 0 | 0 | 8 mo | WILD |
| **1013446243** | 1 | AUDpo4 | 759 | 0.076440637 | 5.28E-06 | 0 | 6.9E-05 | 0 | 4 mo | HOM |
| **1013288951** | 2 | AUDpo4 | 759 | 0.076440637 | 3.43E-07 | 0 | 4.48E-06 | 0 | 4 mo | HOM |
| **1013978665** | 1 | epsc | 760 | 0.115760207 | 0 | 0 | 0 | 0 | 2 mo | HOM |
| **1013445609** | 1 | epsc | 760 | 0.115760207 | 0 | 0 | 0 | 0 | 4 mo | HOM |
| **1013609015** | 1 | epsc | 760 | 0.115760207 | 0 | 0 | 0 | 0 | 4 mo | HOM |
| **1013608106** | 2 | epsc | 760 | 0.115760207 | 0 | 0 | 0 | 0 | 4 mo | HOM |

| | | | | | | | | | | |
|---|---|---|---|---|---|---|---|---|---|---|
| **1011937543** | 1 | epsc | 760 | 0.115760207 | 0.000165372 | 3 | 0.001428578 | 25.91564133 | 8 mo | HOM |
| **1010419538** | 2 | epsc | 760 | 0.115760207 | 0 | 0 | 0 | 0 | 4 mo | WILD |
| **1010419187** | 1 | epsc | 760 | 0.115760207 | 0 | 0 | 0 | 0 | 8 mo | HOM |
| **1013978366** | 1 | epsc | 760 | 0.115760207 | 0 | 0 | 0 | 0 | 2 mo | HOM |
| **1014416155** | 2 | epsc | 760 | 0.115760207 | 0 | 0 | 0 | 0 | 2 mo | HOM |
| **1013978065** | 2 | epsc | 760 | 0.115760207 | 0 | 0 | 0 | 0 | 2 mo | HOM |
| **1009934102** | 2 | epsc | 760 | 0.115760207 | 0 | 0 | 0 | 0 | 4 mo | WILD |
| **1009322937** | 2 | epsc | 760 | 0.115760207 | 0 | 0 | 0 | 0 | 8 mo | WILD |
| **1011080637** | 2 | epsc | 760 | 0.115760207 | 0 | 0 | 0 | 0 | 8 mo | HOM |
| **1013609713** | 2 | epsc | 760 | 0.115760207 | 0 | 0 | 0 | 0 | 2 mo | WILD |
| **1010418842** | 1 | epsc | 760 | 0.115760207 | 2.23E-05 | 2 | 0.000192795 | 17.27709422 | 8 mo | HOM |
| **1013288213** | 2 | epsc | 760 | 0.115760207 | 0 | 0 | 0 | 0 | 4 mo | HOM |
| **1008976060** | 2 | epsc | 760 | 0.115760207 | 0 | 0 | 0 | 0 | 8 mo | WILD |
| **1013446243** | 1 | epsc | 760 | 0.115760207 | 0 | 0 | 0 | 0 | 4 mo | HOM |
| **1013288951** | 2 | epsc | 760 | 0.115760207 | 0 | 0 | 0 | 0 | 4 mo | HOM |
| **1013978665** | 1 | OV | 763 | 0.008925869 | 0 | 0 | 0 | 0 | 2 mo | HOM |
| **1013445609** | 1 | OV | 763 | 0.008925869 | 0 | 0 | 0 | 0 | 4 mo | HOM |
| **1013609015** | 1 | OV | 763 | 0.008925869 | 1.45E-05 | 0 | 0.001624765 | 0 | 4 mo | HOM |
| **1013608106** | 2 | OV | 763 | 0.008925869 | 0 | 0 | 0 | 0 | 4 mo | HOM |
| **1011937543** | 1 | OV | 763 | 0.008925869 | 0 | 0 | 0 | 0 | 8 mo | HOM |
| **1010419538** | 2 | OV | 763 | 0.008925869 | 0 | 0 | 0 | 0 | 4 mo | WILD |
| **1010419187** | 1 | OV | 763 | 0.008925869 | 0 | 0 | 0 | 0 | 8 mo | HOM |
| **1013978366** | 1 | OV | 763 | 0.008925869 | 0 | 0 | 0 | 0 | 2 mo | HOM |
| **1014416155** | 2 | OV | 763 | 0.008925869 | 0 | 0 | 0 | 0 | 2 mo | HOM |
| **1013978065** | 2 | OV | 763 | 0.008925869 | 4.84E-06 | 1 | 0.000542567 | 112.0339045 | 2 mo | HOM |
| **1009934102** | 2 | OV | 763 | 0.008925869 | 7.46E-06 | 2 | 0.000836076 | 224.067809 | 4 mo | WILD |
| **1009322937** | 2 | OV | 763 | 0.008925869 | 0 | 0 | 0 | 0 | 8 mo | WILD |
| **1011080637** | 2 | OV | 763 | 0.008925869 | 2.23E-06 | 0 | 0.00024969 | 0 | 8 mo | HOM |
| **1013609713** | 2 | OV | 763 | 0.008925869 | 0 | 0 | 0 | 0 | 2 mo | WILD |
| **1010418842** | 1 | OV | 763 | 0.008925869 | 8.91E-06 | 1 | 0.000997793 | 112.0339045 | 8 mo | HOM |
| **1013288213** | 2 | OV | 763 | 0.008925869 | 2.37E-05 | 2 | 0.002650444 | 224.067809 | 4 mo | HOM |
| **1008976060** | 2 | OV | 763 | 0.008925869 | 2.52E-06 | 1 | 0.000282134 | 112.0339045 | 8 mo | WILD |
| **1013446243** | 1 | OV | 763 | 0.008925869 | 0 | 0 | 0 | 0 | 4 mo | HOM |
| **1013288951** | 2 | OV | 763 | 0.008925869 | 9.1E-07 | 0 | 0.000102001 | 0 | 4 mo | HOM |
| **1013978665** | 1 | x | 765 | 0.058776397 | 0 | 0 | 0 | 0 | 2 mo | HOM |
| **1013445609** | 1 | x | 765 | 0.058776397 | 0 | 0 | 0 | 0 | 4 mo | HOM |
| **1013609015** | 1 | x | 765 | 0.058776397 | 0 | 0 | 0 | 0 | 4 mo | HOM |
| **1013608106** | 2 | x | 765 | 0.058776397 | 0 | 0 | 0 | 0 | 4 mo | HOM |
| **1011937543** | 1 | x | 765 | 0.058776397 | 4.74E-05 | 1 | 0.000806486 | 17.01363228 | 8 mo | HOM |
| **1010419538** | 2 | x | 765 | 0.058776397 | 0 | 0 | 0 | 0 | 4 mo | WILD |
| **1010419187** | 1 | x | 765 | 0.058776397 | 0 | 0 | 0 | 0 | 8 mo | HOM |
| **1013978366** | 1 | x | 765 | 0.058776397 | 0 | 0 | 0 | 0 | 2 mo | HOM |
| **1014416155** | 2 | x | 765 | 0.058776397 | 0 | 0 | 0 | 0 | 2 mo | HOM |
| **1013978065** | 2 | x | 765 | 0.058776397 | 0 | 0 | 0 | 0 | 2 mo | HOM |
| **1009934102** | 2 | x | 765 | 0.058776397 | 0 | 0 | 0 | 0 | 4 mo | WILD |
| **1009322937** | 2 | x | 765 | 0.058776397 | 0 | 0 | 0 | 0 | 8 mo | WILD |
| **1011080637** | 2 | x | 765 | 0.058776397 | 0 | 0 | 0 | 0 | 8 mo | HOM |
| **1013609713** | 2 | x | 765 | 0.058776397 | 0 | 0 | 0 | 0 | 2 mo | WILD |
| **1010418842** | 1 | x | 765 | 0.058776397 | 0 | 0 | 0 | 0 | 8 mo | HOM |
| **1013288213** | 2 | x | 765 | 0.058776397 | 0 | 0 | 0 | 0 | 4 mo | HOM |
| **1008976060** | 2 | x | 765 | 0.058776397 | 2.72E-07 | 0 | 4.62E-06 | 0 | 8 mo | WILD |
| **1013446243** | 1 | x | 765 | 0.058776397 | 0 | 0 | 0 | 0 | 4 mo | HOM |
| **1013288951** | 2 | x | 765 | 0.058776397 | 0 | 0 | 0 | 0 | 4 mo | HOM |
| **1013978665** | 1 | MOs5 | 767 | 4.65976327 | 0 | 0 | 0 | 0 | 2 mo | HOM |
| **1013445609** | 1 | MOs5 | 767 | 4.65976327 | 0.00034277 | 20 | 7.36E-05 | 4.292063532 | 4 mo | HOM |
| **1013609015** | 1 | MOs5 | 767 | 4.65976327 | 0.00157926 | 48 | 0.000338914 | 10.30095248 | 4 mo | HOM |
| **1013608106** | 2 | MOs5 | 767 | 4.65976327 | 0 | 0 | 0 | 0 | 4 mo | HOM |
| **1011937543** | 1 | MOs5 | 767 | 4.65976327 | 0.004438294 | 109 | 0.000952472 | 23.39174625 | 8 mo | HOM |
| **1010419538** | 2 | MOs5 | 767 | 4.65976327 | 0 | 0 | 0 | 0 | 4 mo | WILD |

| | | | | | | | | | | |
|---|---|---|---|---|---|---|---|---|---|---|
| **1010419187** | 1 | MOs5 | 767 | 4.65976327 | 0.006543156 | 265 | 0.001404182 | 56.8698418 | 8 mo | HOM |
| **1013978366** | 1 | MOs5 | 767 | 4.65976327 | 1.42E-05 | 1 | 3.05E-06 | 0.214603177 | 2 mo | HOM |
| **1014416155** | 2 | MOs5 | 767 | 4.65976327 | 0 | 0 | 0 | 0 | 2 mo | HOM |
| **1013978065** | 2 | MOs5 | 767 | 4.65976327 | 0 | 0 | 0 | 0 | 2 mo | HOM |
| **1009934102** | 2 | MOs5 | 767 | 4.65976327 | 0 | 0 | 0 | 0 | 4 mo | WILD |
| **1009322937** | 2 | MOs5 | 767 | 4.65976327 | 0 | 0 | 0 | 0 | 8 mo | WILD |
| **1011080637** | 2 | MOs5 | 767 | 4.65976327 | 0.003150314 | 164 | 0.000676067 | 35.19492096 | 8 mo | HOM |
| **1013609713** | 2 | MOs5 | 767 | 4.65976327 | 0 | 0 | 0 | 0 | 2 mo | WILD |
| **1010418842** | 1 | MOs5 | 767 | 4.65976327 | 0.004298602 | 219 | 0.000922494 | 46.99809567 | 8 mo | HOM |
| **1013288213** | 2 | MOs5 | 767 | 4.65976327 | 0.000256515 | 21 | 5.5E-05 | 4.506666708 | 4 mo | HOM |
| **1008976060** | 2 | MOs5 | 767 | 4.65976327 | 7.39E-06 | 2 | 1.59E-06 | 0.429206353 | 8 mo | WILD |
| **1013446243** | 1 | MOs5 | 767 | 4.65976327 | 0.000681711 | 34 | 0.000146297 | 7.296508004 | 4 mo | HOM |
| **1013288951** | 2 | MOs5 | 767 | 4.65976327 | 0.001019721 | 46 | 0.000218835 | 9.871746123 | 4 mo | HOM |
| **1013978665** | 1 | mfbc | 768 | 6.410816054 | 6.29E-07 | 0 | 9.8E-08 | 0 | 2 mo | HOM |
| **1013445609** | 1 | mfbc | 768 | 6.410816054 | 2.64E-05 | 2 | 4.12E-06 | 0.311972763 | 4 mo | HOM |
| **1013609015** | 1 | mfbc | 768 | 6.410816054 | 0.001658681 | 26 | 0.000258732 | 4.055645924 | 4 mo | HOM |
| **1013608106** | 2 | mfbc | 768 | 6.410816054 | 0 | 0 | 0 | 0 | 4 mo | HOM |
| **1011937543** | 1 | mfbc | 768 | 6.410816054 | 0.005218226 | 75 | 0.000813972 | 11.69897863 | 8 mo | HOM |
| **1010419538** | 2 | mfbc | 768 | 6.410816054 | 2.33E-06 | 0 | 3.64E-07 | 0 | 4 mo | WILD |
| **1010419187** | 1 | mfbc | 768 | 6.410816054 | 0.003955005 | 116 | 0.000616927 | 18.09442028 | 8 mo | HOM |
| **1013978366** | 1 | mfbc | 768 | 6.410816054 | 2.28E-05 | 1 | 3.56E-06 | 0.155986382 | 2 mo | HOM |
| **1014416155** | 2 | mfbc | 768 | 6.410816054 | 0 | 0 | 0 | 0 | 2 mo | HOM |
| **1013978065** | 2 | mfbc | 768 | 6.410816054 | 0 | 0 | 0 | 0 | 2 mo | HOM |
| **1009934102** | 2 | mfbc | 768 | 6.410816054 | 8.22E-06 | 0 | 1.28E-06 | 0 | 4 mo | WILD |
| **1009322937** | 2 | mfbc | 768 | 6.410816054 | 6.87E-07 | 0 | 1.07E-07 | 0 | 8 mo | WILD |
| **1011080637** | 2 | mfbc | 768 | 6.410816054 | 0.00148402 | 50 | 0.000231487 | 7.799319085 | 8 mo | HOM |
| **1013609713** | 2 | mfbc | 768 | 6.410816054 | 1.56E-06 | 0 | 2.43E-07 | 0 | 2 mo | WILD |
| **1010418842** | 1 | mfbc | 768 | 6.410816054 | 0.005958018 | 151 | 0.00092937 | 23.55394364 | 8 mo | HOM |
| **1013288213** | 2 | mfbc | 768 | 6.410816054 | 5.55E-05 | 1 | 8.66E-06 | 0.155986382 | 4 mo | HOM |
| **1008976060** | 2 | mfbc | 768 | 6.410816054 | 0.000136947 | 8 | 2.14E-05 | 1.247891054 | 8 mo | WILD |
| **1013446243** | 1 | mfbc | 768 | 6.410816054 | 0.000474409 | 13 | 7.4E-05 | 2.027822962 | 4 mo | HOM |
| **1013288951** | 2 | mfbc | 768 | 6.410816054 | 0.000981113 | 10 | 0.00015304 | 1.559863817 | 4 mo | HOM |
| **1013978665** | 1 | P | 771 | 17.21408228 | 0.00011209 | 11 | 6.51E-06 | 0.639011701 | 2 mo | HOM |
| **1013445609** | 1 | P | 771 | 17.21408228 | 1.03E-05 | 2 | 6.01E-07 | 0.116183946 | 4 mo | HOM |
| **1013609015** | 1 | P | 771 | 17.21408228 | 0.00262509 | 17 | 0.000152497 | 0.987563538 | 4 mo | HOM |
| **1013608106** | 2 | P | 771 | 17.21408353 | 0 | 0 | 0 | 0 | 4 mo | HOM |
| **1011937543** | 1 | P | 771 | 17.21408228 | 0.021778765 | 78 | 0.001265171 | 4.531173881 | 8 mo | HOM |
| **1010419538** | 2 | P | 771 | 17.21408353 | 0 | 0 | 0 | 0 | 4 mo | WILD |
| **1010419187** | 1 | P | 771 | 17.21408228 | 0.00226629 | 40 | 0.000131653 | 2.323678913 | 8 mo | HOM |
| **1013978366** | 1 | P | 771 | 17.21408228 | 0.000111585 | 7 | 6.48E-06 | 0.40664381 | 2 mo | HOM |
| **1014416155** | 2 | P | 771 | 17.21408353 | 0 | 0 | 0 | 0 | 2 mo | HOM |
| **1013978065** | 2 | P | 771 | 17.21408353 | 0 | 0 | 0 | 0 | 2 mo | HOM |
| **1009934102** | 2 | P | 771 | 17.21408353 | 1.11E-05 | 0 | 6.43E-07 | 0 | 4 mo | WILD |
| **1009322937** | 2 | P | 771 | 17.21408353 | 0.000141546 | 11 | 8.22E-06 | 0.639011655 | 8 mo | WILD |
| **1011080637** | 2 | P | 771 | 17.21408353 | 0.003261149 | 34 | 0.000189447 | 1.975126932 | 8 mo | HOM |
| **1013609713** | 2 | P | 771 | 17.21408353 | 1.32E-05 | 0 | 7.7E-07 | 0 | 2 mo | WILD |
| **1010418842** | 1 | P | 771 | 17.21408228 | 0.003675552 | 112 | 0.00021352 | 6.506300957 | 8 mo | HOM |
| **1013288213** | 2 | P | 771 | 17.21408353 | 3.05E-05 | 3 | 1.77E-06 | 0.174275906 | 4 mo | HOM |
| **1008976060** | 2 | P | 771 | 17.21408353 | 0.003412469 | 116 | 0.000198237 | 6.738668357 | 8 mo | WILD |
| **1013446243** | 1 | P | 771 | 17.21408228 | 0.000203388 | 16 | 1.18E-05 | 0.929471565 | 4 mo | HOM |
| **1013288951** | 2 | P | 771 | 17.21408353 | 0.000268073 | 18 | 1.56E-05 | 1.045655435 | 4 mo | HOM |
| **1013978665** | 1 | ACAv5 | 772 | 1.032862376 | 0 | 0 | 0 | 0 | 2 mo | HOM |
| **1013445609** | 1 | ACAv5 | 772 | 1.032862376 | 4.94E-05 | 3 | 4.78E-05 | 2.904549599 | 4 mo | HOM |
| **1013609015** | 1 | ACAv5 | 772 | 1.032862376 | 0.000143293 | 6 | 0.000138734 | 5.809099198 | 4 mo | HOM |
| **1013608106** | 2 | ACAv5 | 772 | 1.032862376 | 0 | 0 | 0 | 0 | 4 mo | HOM |
| **1011937543** | 1 | ACAv5 | 772 | 1.032862376 | 0.001003936 | 20 | 0.000971994 | 19.36366399 | 8 mo | HOM |
| **1010419538** | 2 | ACAv5 | 772 | 1.032862376 | 0 | 0 | 0 | 0 | 4 mo | WILD |
| **1010419187** | 1 | ACAv5 | 772 | 1.032862376 | 0.001669542 | 60 | 0.001616422 | 58.09099198 | 8 mo | HOM |
| **1013978366** | 1 | ACAv5 | 772 | 1.032862376 | 0 | 0 | 0 | 0 | 2 mo | HOM |

| | | | | | | | | | | |
|---|---|---|---|---|---|---|---|---|---|---|
| **1014416155** | 2 | ACAv5 | 772 | 1.032862376 | 0 | 0 | 0 | 0 | 2 mo | HOM |
| **1013978065** | 2 | ACAv5 | 772 | 1.032862376 | 0 | 0 | 0 | 0 | 2 mo | HOM |
| **1009934102** | 2 | ACAv5 | 772 | 1.032862376 | 0 | 0 | 0 | 0 | 4 mo | WILD |
| **1009322937** | 2 | ACAv5 | 772 | 1.032862376 | 0 | 0 | 0 | 0 | 8 mo | WILD |
| **1011080637** | 2 | ACAv5 | 772 | 1.032862376 | 0.000876931 | 40 | 0.00084903 | 38.72732799 | 8 mo | HOM |
| **1013609713** | 2 | ACAv5 | 772 | 1.032862376 | 0 | 0 | 0 | 0 | 2 mo | WILD |
| **1010418842** | 1 | ACAv5 | 772 | 1.032862376 | 0.001354332 | 42 | 0.001311242 | 40.66369439 | 8 mo | HOM |
| **1013288213** | 2 | ACAv5 | 772 | 1.032862376 | 5.5E-05 | 4 | 5.32E-05 | 3.872732799 | 4 mo | HOM |
| **1008976060** | 2 | ACAv5 | 772 | 1.032862376 | 1.12E-05 | 1 | 1.08E-05 | 0.9681832 | 8 mo | WILD |
| **1013446243** | 1 | ACAv5 | 772 | 1.032862376 | 9.61E-05 | 3 | 9.3E-05 | 2.904549599 | 4 mo | HOM |
| **1013288951** | 2 | ACAv5 | 772 | 1.032862376 | 0.000163487 | 7 | 0.000158286 | 6.777282398 | 4 mo | HOM |
| **1013978665** | 1 | XII | 773 | 0.291184852 | 0 | 0 | 0 | 0 | 2 mo | HOM |
| **1013445609** | 1 | XII | 773 | 0.291184852 | 0 | 0 | 0 | 0 | 4 mo | HOM |
| **1013609015** | 1 | XII | 773 | 0.291184852 | 0 | 0 | 0 | 0 | 4 mo | HOM |
| **1013608106** | 2 | XII | 773 | 0.291184852 | 0 | 0 | 0 | 0 | 4 mo | HOM |
| **1011937543** | 1 | XII | 773 | 0.291184852 | 0 | 0 | 0 | 0 | 8 mo | HOM |
| **1010419538** | 2 | XII | 773 | 0.291184852 | 0 | 0 | 0 | 0 | 4 mo | WILD |
| **1010419187** | 1 | XII | 773 | 0.291184852 | 3.76E-06 | 1 | 1.29E-05 | 3.434244581 | 8 mo | HOM |
| **1013978366** | 1 | XII | 773 | 0.291184852 | 0 | 0 | 0 | 0 | 2 mo | HOM |
| **1014416155** | 2 | XII | 773 | 0.291184852 | 0 | 0 | 0 | 0 | 2 mo | HOM |
| **1013978065** | 2 | XII | 773 | 0.291184852 | 0 | 0 | 0 | 0 | 2 mo | HOM |
| **1009934102** | 2 | XII | 773 | 0.291184852 | 0 | 0 | 0 | 0 | 4 mo | WILD |
| **1009322937** | 2 | XII | 773 | 0.291184852 | 0 | 0 | 0 | 0 | 8 mo | WILD |
| **1011080637** | 2 | XII | 773 | 0.291184852 | 0 | 0 | 0 | 0 | 8 mo | HOM |
| **1013609713** | 2 | XII | 773 | 0.291184852 | 0 | 0 | 0 | 0 | 2 mo | WILD |
| **1010418842** | 1 | XII | 773 | 0.291184852 | 1.7E-06 | 0 | 5.84E-06 | 0 | 8 mo | HOM |
| **1013288213** | 2 | XII | 773 | 0.291184852 | 0 | 0 | 0 | 0 | 4 mo | HOM |
| **1008976060** | 2 | XII | 773 | 0.291184852 | 4.52E-06 | 1 | 1.55E-05 | 3.434244581 | 8 mo | WILD |
| **1013446243** | 1 | XII | 773 | 0.291184852 | 0 | 0 | 0 | 0 | 4 mo | HOM |
| **1013288951** | 2 | XII | 773 | 0.291184852 | 0 | 0 | 0 | 0 | 4 mo | HOM |
| **1013978665** | 1 | RSPagl5 | 774 | 0.755404227 | 0 | 0 | 0 | 0 | 2 mo | HOM |
| **1013445609** | 1 | RSPagl5 | 774 | 0.755404227 | 8.87E-05 | 6 | 0.000117404 | 7.942767308 | 4 mo | HOM |
| **1013609015** | 1 | RSPagl5 | 774 | 0.755404227 | 0.00032226 | 9 | 0.000426605 | 11.91415096 | 4 mo | HOM |
| **1013608106** | 2 | RSPagl5 | 774 | 0.755404227 | 0 | 0 | 0 | 0 | 4 mo | HOM |
| **1011937543** | 1 | RSPagl5 | 774 | 0.755404227 | 0.001153019 | 11 | 0.00152636 | 14.56174006 | 8 mo | HOM |
| **1010419538** | 2 | RSPagl5 | 774 | 0.755404227 | 0 | 0 | 0 | 0 | 4 mo | WILD |
| **1010419187** | 1 | RSPagl5 | 774 | 0.755404227 | 0.00162428 | 48 | 0.002150213 | 63.54213846 | 8 mo | HOM |
| **1013978366** | 1 | RSPagl5 | 774 | 0.755404227 | 6.07E-08 | 0 | 8.04E-08 | 0 | 2 mo | HOM |
| **1014416155** | 2 | RSPagl5 | 774 | 0.755404227 | 0 | 0 | 0 | 0 | 2 mo | HOM |
| **1013978065** | 2 | RSPagl5 | 774 | 0.755404227 | 0 | 0 | 0 | 0 | 2 mo | HOM |
| **1009934102** | 2 | RSPagl5 | 774 | 0.755404227 | 0 | 0 | 0 | 0 | 4 mo | WILD |
| **1009322937** | 2 | RSPagl5 | 774 | 0.755404227 | 0 | 0 | 0 | 0 | 8 mo | WILD |
| **1011080637** | 2 | RSPagl5 | 774 | 0.755404227 | 0.000434509 | 23 | 0.000575201 | 30.44727468 | 8 mo | HOM |
| **1013609713** | 2 | RSPagl5 | 774 | 0.755404227 | 0 | 0 | 0 | 0 | 2 mo | WILD |
| **1010418842** | 1 | RSPagl5 | 774 | 0.755404227 | 0.000963722 | 28 | 0.00127577 | 37.06624744 | 8 mo | HOM |
| **1013288213** | 2 | RSPagl5 | 774 | 0.755404227 | 5.9E-05 | 6 | 7.81E-05 | 7.942767308 | 4 mo | HOM |
| **1008976060** | 2 | RSPagl5 | 774 | 0.755404227 | 0 | 0 | 0 | 0 | 8 mo | WILD |
| **1013446243** | 1 | RSPagl5 | 774 | 0.755404227 | 0.000235667 | 14 | 0.000311974 | 18.53312372 | 4 mo | HOM |
| **1013288951** | 2 | RSPagl5 | 774 | 0.755404227 | 0.000197209 | 9 | 0.000261064 | 11.91415096 | 4 mo | HOM |
| **1013978665** | 1 | cc | 776 | 8.036162964 | 0 | 0 | 0 | 0 | 2 mo | HOM |
| **1013445609** | 1 | cc | 776 | 8.036162964 | 5.79E-05 | 4 | 7.21E-06 | 0.497749986 | 4 mo | HOM |
| **1013609015** | 1 | cc | 776 | 8.036162964 | 0.000119604 | 5 | 1.49E-05 | 0.622187482 | 4 mo | HOM |
| **1013608106** | 2 | cc | 776 | 8.036162964 | 0 | 0 | 0 | 0 | 4 mo | HOM |
| **1011937543** | 1 | cc | 776 | 8.036162964 | 0.00535188 | 86 | 0.000665974 | 10.70162469 | 8 mo | HOM |
| **1010419538** | 2 | cc | 776 | 8.036162964 | 0 | 0 | 0 | 0 | 4 mo | WILD |
| **1010419187** | 1 | cc | 776 | 8.036162964 | 0.002198524 | 95 | 0.000273579 | 11.82156216 | 8 mo | HOM |
| **1013978366** | 1 | cc | 776 | 8.036162964 | 3.05E-06 | 0 | 3.8E-07 | 0 | 2 mo | HOM |
| **1014416155** | 2 | cc | 776 | 8.036162964 | 0 | 0 | 0 | 0 | 2 mo | HOM |
| **1013978065** | 2 | cc | 776 | 8.036162964 | 7.35E-08 | 0 | 9.14E-09 | 0 | 2 mo | HOM |

| | | | | | | | | | | |
|---|---|---|---|---|---|---|---|---|---|---|
| **1009934102** | 2 | cc | 776 | 8.036162964 | 7.03E-06 | 0 | 8.74E-07 | 0 | 4 mo | WILD |
| **1009322937** | 2 | cc | 776 | 8.036162964 | 6.19E-06 | 0 | 7.7E-07 | 0 | 8 mo | WILD |
| **1011080637** | 2 | cc | 776 | 8.036162964 | 0.001731495 | 66 | 0.000215463 | 8.212874763 | 8 mo | HOM |
| **1013609713** | 2 | cc | 776 | 8.036162964 | 0 | 0 | 0 | 0 | 2 mo | WILD |
| **1010418842** | 1 | cc | 776 | 8.036162964 | 0.004260644 | 125 | 0.000530184 | 15.55468705 | 8 mo | HOM |
| **1013288213** | 2 | cc | 776 | 8.036162964 | 0.000311305 | 3 | 3.87E-05 | 0.373312489 | 4 mo | HOM |
| **1008976060** | 2 | cc | 776 | 8.036162964 | 8.23E-05 | 8 | 1.02E-05 | 0.995499971 | 8 mo | WILD |
| **1013446243** | 1 | cc | 776 | 8.036162964 | 0.000101038 | 6 | 1.26E-05 | 0.746624978 | 4 mo | HOM |
| **1013288951** | 2 | cc | 776 | 8.036162964 | 0.000121303 | 4 | 1.51E-05 | 0.497749986 | 4 mo | HOM |
| **1013978665** | 1 | VISp5 | 778 | 1.599617936 | 0 | 0 | 0 | 0 | 2 mo | HOM |
| **1013445609** | 1 | VISp5 | 778 | 1.599617936 | 0.00030915 | 17 | 0.000193265 | 10.62753775 | 4 mo | HOM |
| **1013609015** | 1 | VISp5 | 778 | 1.599617936 | 0.000736148 | 27 | 0.000460202 | 16.87903054 | 4 mo | HOM |
| **1013608106** | 2 | VISp5 | 778 | 1.599617936 | 0 | 0 | 0 | 0 | 4 mo | HOM |
| **1011937543** | 1 | VISp5 | 778 | 1.599617936 | 0.002240113 | 34 | 0.001400405 | 21.2550755 | 8 mo | HOM |
| **1010419538** | 2 | VISp5 | 778 | 1.599617936 | 0 | 0 | 0 | 0 | 4 mo | WILD |
| **1010419187** | 1 | VISp5 | 778 | 1.599617936 | 0.003310776 | 94 | 0.002069729 | 58.76403226 | 8 mo | HOM |
| **1013978366** | 1 | VISp5 | 778 | 1.599617936 | 9.57E-07 | 0 | 5.98E-07 | 0 | 2 mo | HOM |
| **1014416155** | 2 | VISp5 | 778 | 1.599617936 | 0 | 0 | 0 | 0 | 2 mo | HOM |
| **1013978065** | 2 | VISp5 | 778 | 1.599617936 | 0 | 0 | 0 | 0 | 2 mo | HOM |
| **1009934102** | 2 | VISp5 | 778 | 1.599617936 | 3.89E-06 | 1 | 2.43E-06 | 0.625149279 | 4 mo | WILD |
| **1009322937** | 2 | VISp5 | 778 | 1.599617936 | 0 | 0 | 0 | 0 | 8 mo | WILD |
| **1011080637** | 2 | VISp5 | 778 | 1.599617936 | 0.001724989 | 80 | 0.001078376 | 50.01194235 | 8 mo | HOM |
| **1013609713** | 2 | VISp5 | 778 | 1.599617936 | 0 | 0 | 0 | 0 | 2 mo | WILD |
| **1010418842** | 1 | VISp5 | 778 | 1.599617936 | 0.00200128 | 74 | 0.001251099 | 46.26104668 | 8 mo | HOM |
| **1013288213** | 2 | VISp5 | 778 | 1.599617936 | 0.000172287 | 16 | 0.000107705 | 10.00238847 | 4 mo | HOM |
| **1008976060** | 2 | VISp5 | 778 | 1.599617936 | 0 | 0 | 0 | 0 | 8 mo | WILD |
| **1013446243** | 1 | VISp5 | 778 | 1.599617936 | 0.000469153 | 24 | 0.000293291 | 15.00358271 | 4 mo | HOM |
| **1013288951** | 2 | VISp5 | 778 | 1.599617936 | 0.000666251 | 31 | 0.000416507 | 19.37962766 | 4 mo | HOM |
| **1013978665** | 1 | PA | 780 | 0.995875216 | 0 | 0 | 0 | 0 | 2 mo | HOM |
| **1013445609** | 1 | PA | 780 | 0.995875216 | 4.36E-05 | 4 | 4.37E-05 | 4.016567473 | 4 mo | HOM |
| **1013609015** | 1 | PA | 780 | 0.995875216 | 0.000103806 | 2 | 0.000104236 | 2.008283737 | 4 mo | HOM |
| **1013608106** | 2 | PA | 780 | 0.995875216 | 0 | 0 | 0 | 0 | 4 mo | HOM |
| **1011937543** | 1 | PA | 780 | 0.995875216 | 0.002103232 | 40 | 0.002111943 | 40.16567473 | 8 mo | HOM |
| **1010419538** | 2 | PA | 780 | 0.995875216 | 3.91E-06 | 1 | 3.92E-06 | 1.004141868 | 4 mo | WILD |
| **1010419187** | 1 | PA | 780 | 0.995875216 | 0.002245906 | 68 | 0.002255208 | 68.28164705 | 8 mo | HOM |
| **1013978366** | 1 | PA | 780 | 0.995875216 | 0 | 0 | 0 | 0 | 2 mo | HOM |
| **1014416155** | 2 | PA | 780 | 0.995875216 | 0 | 0 | 0 | 0 | 2 mo | HOM |
| **1013978065** | 2 | PA | 780 | 0.995875216 | 0 | 0 | 0 | 0 | 2 mo | HOM |
| **1009934102** | 2 | PA | 780 | 0.995875216 | 0 | 0 | 0 | 0 | 4 mo | WILD |
| **1009322937** | 2 | PA | 780 | 0.995875216 | 0 | 0 | 0 | 0 | 8 mo | WILD |
| **1011080637** | 2 | PA | 780 | 0.995875216 | 0.000914034 | 50 | 0.00091782 | 50.20709342 | 8 mo | HOM |
| **1013609713** | 2 | PA | 780 | 0.995875216 | 0 | 0 | 0 | 0 | 2 mo | WILD |
| **1010418842** | 1 | PA | 780 | 0.995875216 | 0.00166026 | 56 | 0.001667136 | 56.23194463 | 8 mo | HOM |
| **1013288213** | 2 | PA | 780 | 0.995875216 | 9.57E-05 | 8 | 9.61E-05 | 8.033134946 | 4 mo | HOM |
| **1008976060** | 2 | PA | 780 | 0.995875216 | 2.21E-05 | 2 | 2.22E-05 | 2.008283737 | 8 mo | WILD |
| **1013446243** | 1 | PA | 780 | 0.995875216 | 0.000109544 | 5 | 0.000109998 | 5.020709342 | 4 mo | HOM |
| **1013288951** | 2 | PA | 780 | 0.995875216 | 0.000121221 | 6 | 0.000121723 | 6.02485121 | 4 mo | HOM |
| **1013978665** | 1 | y | 781 | 0.024185562 | 0 | 0 | 0 | 0 | 2 mo | HOM |
| **1013445609** | 1 | y | 781 | 0.024185562 | 0 | 0 | 0 | 0 | 4 mo | HOM |
| **1013609015** | 1 | y | 781 | 0.024185562 | 9.71E-06 | 0 | 0.000401364 | 0 | 4 mo | HOM |
| **1013608106** | 2 | y | 781 | 0.024185562 | 0 | 0 | 0 | 0 | 4 mo | HOM |
| **1011937543** | 1 | y | 781 | 0.024185562 | 0 | 0 | 0 | 0 | 8 mo | HOM |
| **1010419538** | 2 | y | 781 | 0.024185562 | 2.8E-06 | 0 | 0.00011596 | 0 | 4 mo | WILD |
| **1010419187** | 1 | y | 781 | 0.024185562 | 1.72E-05 | 1 | 0.000709143 | 41.34698197 | 8 mo | HOM |
| **1013978366** | 1 | y | 781 | 0.024185562 | 0 | 0 | 0 | 0 | 2 mo | HOM |
| **1014416155** | 2 | y | 781 | 0.024185562 | 0 | 0 | 0 | 0 | 2 mo | HOM |
| **1013978065** | 2 | y | 781 | 0.024185562 | 0 | 0 | 0 | 0 | 2 mo | HOM |
| **1009934102** | 2 | y | 781 | 0.024185562 | 0 | 0 | 0 | 0 | 4 mo | WILD |
| **1009322937** | 2 | y | 781 | 0.024185562 | 0 | 0 | 0 | 0 | 8 mo | WILD |

| | | | | | | | | | | |
|---|---|---|---|---|---|---|---|---|---|---|
| **1011080637** | 2 | y | 781 | 0.024185562 | 0 | 0 | 0 | 0 | 8 mo | HOM |
| **1013609713** | 2 | y | 781 | 0.024185562 | 0 | 0 | 0 | 0 | 2 mo | WILD |
| **1010418842** | 1 | y | 781 | 0.024185562 | 6.44E-08 | 0 | 2.66E-06 | 0 | 8 mo | HOM |
| **1013288213** | 2 | y | 781 | 0.024185562 | 0 | 0 | 0 | 0 | 4 mo | HOM |
| **1008976060** | 2 | y | 781 | 0.024185562 | 8.31E-07 | 0 | 3.44E-05 | 0 | 8 mo | WILD |
| **1013446243** | 1 | y | 781 | 0.024185562 | 0 | 0 | 0 | 0 | 4 mo | HOM |
| **1013288951** | 2 | y | 781 | 0.024185562 | 0 | 0 | 0 | 0 | 4 mo | HOM |
| **1013978665** | 1 | Ald6a | 783 | 0.805351613 | 0 | 0 | 0 | 0 | 2 mo | HOM |
| **1013445609** | 1 | Ald6a | 783 | 0.805351613 | 1.71E-05 | 1 | 2.12E-05 | 1.24169367 | 4 mo | HOM |
| **1013609015** | 1 | Ald6a | 783 | 0.805351613 | 0.000217459 | 7 | 0.000270017 | 8.691855692 | 4 mo | HOM |
| **1013608106** | 2 | Ald6a | 783 | 0.805351613 | 0 | 0 | 0 | 0 | 4 mo | HOM |
| **1011937543** | 1 | Ald6a | 783 | 0.805351613 | 0.000871182 | 22 | 0.001081742 | 27.31726075 | 8 mo | HOM |
| **1010419538** | 2 | Ald6a | 783 | 0.805351613 | 0 | 0 | 0 | 0 | 4 mo | WILD |
| **1010419187** | 1 | Ald6a | 783 | 0.805351613 | 0.000863879 | 41 | 0.001072673 | 50.90944048 | 8 mo | HOM |
| **1013978366** | 1 | Ald6a | 783 | 0.805351613 | 0 | 0 | 0 | 0 | 2 mo | HOM |
| **1014416155** | 2 | Ald6a | 783 | 0.805351613 | 0 | 0 | 0 | 0 | 2 mo | HOM |
| **1013978065** | 2 | Ald6a | 783 | 0.805351613 | 0 | 0 | 0 | 0 | 2 mo | HOM |
| **1009934102** | 2 | Ald6a | 783 | 0.805351613 | 0 | 0 | 0 | 0 | 4 mo | WILD |
| **1009322937** | 2 | Ald6a | 783 | 0.805351613 | 3.44E-06 | 0 | 4.27E-06 | 0 | 8 mo | WILD |
| **1011080637** | 2 | Ald6a | 783 | 0.805351613 | 0.000541147 | 20 | 0.000671939 | 24.83387341 | 8 mo | HOM |
| **1013609713** | 2 | Ald6a | 783 | 0.805351613 | 0 | 0 | 0 | 0 | 2 mo | WILD |
| **1010418842** | 1 | Ald6a | 783 | 0.805351613 | 0.000801279 | 28 | 0.000994943 | 34.76742277 | 8 mo | HOM |
| **1013288213** | 2 | Ald6a | 783 | 0.805351613 | 1.48E-05 | 2 | 1.84E-05 | 2.483387341 | 4 mo | HOM |
| **1008976060** | 2 | Ald6a | 783 | 0.805351613 | 1.18E-06 | 0 | 1.47E-06 | 0 | 8 mo | WILD |
| **1013446243** | 1 | Ald6a | 783 | 0.805351613 | 0.000143482 | 9 | 0.000178161 | 11.17524303 | 4 mo | HOM |
| **1013288951** | 2 | Ald6a | 783 | 0.805351613 | 5.62E-05 | 4 | 6.97E-05 | 4.966774681 | 4 mo | HOM |
| **1013978665** | 1 | cst | 784 | 3.806992358 | 0 | 0 | 0 | 0 | 2 mo | HOM |
| **1013445609** | 1 | cst | 784 | 3.806992358 | 0 | 0 | 0 | 0 | 4 mo | HOM |
| **1013609015** | 1 | cst | 784 | 3.806992358 | 0 | 0 | 0 | 0 | 4 mo | HOM |
| **1013608106** | 2 | cst | 784 | 3.806976678 | 0 | 0 | 0 | 0 | 4 mo | HOM |
| **1011937543** | 1 | cst | 784 | 3.806992358 | 0.003683117 | 50 | 0.000967461 | 13.13372744 | 8 mo | HOM |
| **1010419538** | 2 | cst | 784 | 3.806992358 | 1.13E-06 | 0 | 2.97E-07 | 0 | 4 mo | WILD |
| **1010419187** | 1 | cst | 784 | 3.806992358 | 0.000216413 | 4 | 5.68E-05 | 1.050698195 | 8 mo | HOM |
| **1013978366** | 1 | cst | 784 | 3.806992358 | 0 | 0 | 0 | 0 | 2 mo | HOM |
| **1014416155** | 2 | cst | 784 | 3.806992358 | 0 | 0 | 0 | 0 | 2 mo | HOM |
| **1013978065** | 2 | cst | 784 | 3.806992358 | 0 | 0 | 0 | 0 | 2 mo | HOM |
| **1009934102** | 2 | cst | 784 | 3.806992358 | 2.16E-06 | 0 | 5.66E-07 | 0 | 4 mo | WILD |
| **1009322937** | 2 | cst | 784 | 3.806992358 | 6.58E-11 | 0 | 1.73E-11 | 0 | 8 mo | WILD |
| **1011080637** | 2 | cst | 784 | 3.806992358 | 7.7E-05 | 6 | 2.02E-05 | 1.576047293 | 8 mo | HOM |
| **1013609713** | 2 | cst | 784 | 3.806992358 | 0 | 0 | 0 | 0 | 2 mo | WILD |
| **1010418842** | 1 | cst | 784 | 3.806992358 | 0.00021294 | 13 | 5.59E-05 | 3.414769134 | 8 mo | HOM |
| **1013288213** | 2 | cst | 784 | 3.806992358 | 0 | 0 | 0 | 0 | 4 mo | HOM |
| **1008976060** | 2 | cst | 784 | 3.806992358 | 2.04E-05 | 3 | 5.35E-06 | 0.788023646 | 8 mo | WILD |
| **1013446243** | 1 | cst | 784 | 3.806992358 | 1.79E-05 | 3 | 4.7E-06 | 0.788023646 | 4 mo | HOM |
| **1013288951** | 2 | cst | 784 | 3.806992358 | 3.18E-05 | 1 | 8.34E-06 | 0.262674549 | 4 mo | HOM |
| **1013978665** | 1 | TEa6b | 786 | 0.092470938 | 0 | 0 | 0 | 0 | 2 mo | HOM |
| **1013445609** | 1 | TEa6b | 786 | 0.092470938 | 3.63E-06 | 0 | 3.92E-05 | 0 | 4 mo | HOM |
| **1013609015** | 1 | TEa6b | 786 | 0.092470938 | 4.75E-06 | 0 | 5.14E-05 | 0 | 4 mo | HOM |
| **1013608106** | 2 | TEa6b | 786 | 0.092470938 | 0 | 0 | 0 | 0 | 4 mo | HOM |
| **1011937543** | 1 | TEa6b | 786 | 0.092470938 | 7.29E-05 | 2 | 0.000788421 | 21.62841692 | 8 mo | HOM |
| **1010419538** | 2 | TEa6b | 786 | 0.092470938 | 0 | 0 | 0 | 0 | 4 mo | WILD |
| **1010419187** | 1 | TEa6b | 786 | 0.092470938 | 0.000144321 | 6 | 0.001560719 | 64.88525076 | 8 mo | HOM |
| **1013978366** | 1 | TEa6b | 786 | 0.092470938 | 0 | 0 | 0 | 0 | 2 mo | HOM |
| **1014416155** | 2 | TEa6b | 786 | 0.092470938 | 0 | 0 | 0 | 0 | 2 mo | HOM |
| **1013978065** | 2 | TEa6b | 786 | 0.092470938 | 0 | 0 | 0 | 0 | 2 mo | HOM |
| **1009934102** | 2 | TEa6b | 786 | 0.092470938 | 0 | 0 | 0 | 0 | 4 mo | WILD |
| **1009322937** | 2 | TEa6b | 786 | 0.092470938 | 0 | 0 | 0 | 0 | 8 mo | WILD |
| **1011080637** | 2 | TEa6b | 786 | 0.092470938 | 4.15E-05 | 2 | 0.000448905 | 21.62841692 | 8 mo | HOM |
| **1013609713** | 2 | TEa6b | 786 | 0.092470938 | 0 | 0 | 0 | 0 | 2 mo | WILD |

| | | | | | | | | | | |
|---|---|---|---|---|---|---|---|---|---|---|
| **1010418842** | 1 | TEa6b | 786 | 0.092470938 | 0.000188319 | 6 | 0.002036519 | 64.88525076 | 8 mo | HOM |
| **1013288213** | 2 | TEa6b | 786 | 0.092470938 | 0 | 0 | 0 | 0 | 4 mo | HOM |
| **1008976060** | 2 | TEa6b | 786 | 0.092470938 | 0 | 0 | 0 | 0 | 8 mo | WILD |
| **1013446243** | 1 | TEa6b | 786 | 0.092470938 | 4.48E-09 | 0 | 4.85E-08 | 0 | 4 mo | HOM |
| **1013288951** | 2 | TEa6b | 786 | 0.092470938 | 9.1E-06 | 0 | 9.84E-05 | 0 | 4 mo | HOM |
| **1013978665** | 1 | PAA | 788 | 1.212198064 | 1.63E-09 | 0 | 1.34E-09 | 0 | 2 mo | HOM |
| **1013445609** | 1 | PAA | 788 | 1.227628438 | 2.47E-05 | 2 | 2.01E-05 | 1.629157437 | 4 mo | HOM |
| **1013609015** | 1 | PAA | 788 | 1.227628438 | 6.17E-05 | 3 | 5.03E-05 | 2.443736155 | 4 mo | HOM |
| **1013608106** | 2 | PAA | 788 | 1.227628438 | 0 | 0 | 0 | 0 | 4 mo | HOM |
| **1011937543** | 1 | PAA | 788 | 1.227628438 | 0.001435364 | 18 | 0.001169217 | 14.66241693 | 8 mo | HOM |
| **1010419538** | 2 | PAA | 788 | 1.227628438 | 3.13E-25 | 0 | 2.55E-25 | 0 | 4 mo | WILD |
| **1010419187** | 1 | PAA | 788 | 1.227628438 | 0.001658284 | 17 | 0.001350803 | 13.84783821 | 8 mo | HOM |
| **1013978366** | 1 | PAA | 788 | 1.227628438 | 0 | 0 | 0 | 0 | 2 mo | HOM |
| **1014416155** | 2 | PAA | 788 | 1.227628438 | 0 | 0 | 0 | 0 | 2 mo | HOM |
| **1013978065** | 2 | PAA | 788 | 1.227628438 | 3.04E-06 | 0 | 2.47E-06 | 0 | 2 mo | HOM |
| **1009934102** | 2 | PAA | 788 | 1.227628438 | 0 | 0 | 0 | 0 | 4 mo | WILD |
| **1009322937** | 2 | PAA | 788 | 1.227628438 | 6.58E-07 | 0 | 5.36E-07 | 0 | 8 mo | WILD |
| **1011080637** | 2 | PAA | 788 | 1.227628438 | 0.000722833 | 36 | 0.000588804 | 29.32483386 | 8 mo | HOM |
| **1013609713** | 2 | PAA | 788 | 1.227628438 | 0 | 0 | 0 | 0 | 2 mo | WILD |
| **1010418842** | 1 | PAA | 788 | 1.227628438 | 0.000886173 | 27 | 0.000721858 | 21.9936254 | 8 mo | HOM |
| **1013288213** | 2 | PAA | 788 | 1.227628438 | 2.7E-05 | 2 | 2.2E-05 | 1.629157437 | 4 mo | HOM |
| **1008976060** | 2 | PAA | 788 | 1.227628438 | 2.19E-06 | 0 | 1.79E-06 | 0 | 8 mo | WILD |
| **1013446243** | 1 | PAA | 788 | 1.227628438 | 0.000106377 | 4 | 8.67E-05 | 3.258314874 | 4 mo | HOM |
| **1013288951** | 2 | PAA | 788 | 1.227628438 | 4.73E-05 | 3 | 3.85E-05 | 2.443736155 | 4 mo | HOM |
| **1013978665** | 1 | AUDpo5 | 791 | 0.186831296 | 0 | 0 | 0 | 0 | 2 mo | HOM |
| **1013445609** | 1 | AUDpo5 | 791 | 0.186831296 | 6.55E-05 | 5 | 0.000350554 | 26.76211158 | 4 mo | HOM |
| **1013609015** | 1 | AUDpo5 | 791 | 0.186831296 | 8.32E-05 | 4 | 0.000445457 | 21.40968926 | 4 mo | HOM |
| **1013608106** | 2 | AUDpo5 | 791 | 0.186831296 | 0 | 0 | 0 | 0 | 4 mo | HOM |
| **1011937543** | 1 | AUDpo5 | 791 | 0.186831296 | 0.000239787 | 7 | 0.001283442 | 37.46695621 | 8 mo | HOM |
| **1010419538** | 2 | AUDpo5 | 791 | 0.186831296 | 0 | 0 | 0 | 0 | 4 mo | WILD |
| **1010419187** | 1 | AUDpo5 | 791 | 0.186831296 | 0.000230859 | 16 | 0.001235657 | 85.63875704 | 8 mo | HOM |
| **1013978366** | 1 | AUDpo5 | 791 | 0.186831296 | 0 | 0 | 0 | 0 | 2 mo | HOM |
| **1014416155** | 2 | AUDpo5 | 791 | 0.186831296 | 0 | 0 | 0 | 0 | 2 mo | HOM |
| **1013978065** | 2 | AUDpo5 | 791 | 0.186831296 | 0 | 0 | 0 | 0 | 2 mo | HOM |
| **1009934102** | 2 | AUDpo5 | 791 | 0.186831296 | 0 | 0 | 0 | 0 | 4 mo | WILD |
| **1009322937** | 2 | AUDpo5 | 791 | 0.186831296 | 0 | 0 | 0 | 0 | 8 mo | WILD |
| **1011080637** | 2 | AUDpo5 | 791 | 0.186831296 | 0.000164169 | 7 | 0.0008787 | 37.46695621 | 8 mo | HOM |
| **1013609713** | 2 | AUDpo5 | 791 | 0.186831296 | 0 | 0 | 0 | 0 | 2 mo | WILD |
| **1010418842** | 1 | AUDpo5 | 791 | 0.186831296 | 0.000121519 | 4 | 0.000650421 | 21.40968926 | 8 mo | HOM |
| **1013288213** | 2 | AUDpo5 | 791 | 0.186831296 | 6.59E-10 | 0 | 3.53E-09 | 0 | 4 mo | HOM |
| **1008976060** | 2 | AUDpo5 | 791 | 0.186831296 | 0 | 0 | 0 | 0 | 8 mo | WILD |
| **1013446243** | 1 | AUDpo5 | 791 | 0.186831296 | 5.07E-05 | 3 | 0.000271413 | 16.05726695 | 4 mo | HOM |
| **1013288951** | 2 | AUDpo5 | 791 | 0.186831296 | 4.09E-05 | 5 | 0.000218972 | 26.76211158 | 4 mo | HOM |
| **1013978665** | 1 | drt | 792 | 0.708780345 | 0 | 0 | 0 | 0 | 2 mo | HOM |
| **1013445609** | 1 | drt | 792 | 0.708780345 | 0 | 0 | 0 | 0 | 4 mo | HOM |
| **1013609015** | 1 | drt | 792 | 0.708780345 | 0 | 0 | 0 | 0 | 4 mo | HOM |
| **1013608106** | 2 | drt | 792 | 0.707673729 | 0 | 0 | 0 | 0 | 4 mo | HOM |
| **1011937543** | 1 | drt | 792 | 0.708780345 | 0.001106121 | 8 | 0.001560598 | 11.2869947 | 8 mo | HOM |
| **1010419538** | 2 | drt | 792 | 0.708780345 | 0 | 0 | 0 | 0 | 4 mo | WILD |
| **1010419187** | 1 | drt | 792 | 0.708780345 | 0 | 0 | 0 | 0 | 8 mo | HOM |
| **1013978366** | 1 | drt | 792 | 0.708780345 | 0 | 0 | 0 | 0 | 2 mo | HOM |
| **1014416155** | 2 | drt | 792 | 0.708780345 | 0 | 0 | 0 | 0 | 2 mo | HOM |
| **1013978065** | 2 | drt | 792 | 0.708780345 | 0 | 0 | 0 | 0 | 2 mo | HOM |
| **1009934102** | 2 | drt | 792 | 0.708780345 | 0 | 0 | 0 | 0 | 4 mo | WILD |
| **1009322937** | 2 | drt | 792 | 0.708780345 | 0 | 0 | 0 | 0 | 8 mo | WILD |
| **1011080637** | 2 | drt | 792 | 0.708780345 | 1.08E-08 | 0 | 1.53E-08 | 0 | 8 mo | HOM |
| **1013609713** | 2 | drt | 792 | 0.708780345 | 0 | 0 | 0 | 0 | 2 mo | WILD |
| **1010418842** | 1 | drt | 792 | 0.708780345 | 2.85E-05 | 2 | 4.02E-05 | 2.821748676 | 8 mo | HOM |
| **1013288213** | 2 | drt | 792 | 0.708780345 | 0 | 0 | 0 | 0 | 4 mo | HOM |

| | | | | | | | | | | |
|---|---|---|---|---|---|---|---|---|---|---|
| **1008976060** | 2 | drt | 792 | 0.708780345 | 6.86E-06 | 1 | 9.68E-06 | 1.410874338 | 8 mo | WILD |
| **1013446243** | 1 | drt | 792 | 0.708780345 | 0 | 0 | 0 | 0 | 4 mo | HOM |
| **1013288951** | 2 | drt | 792 | 0.708780345 | 0 | 0 | 0 | 0 | 4 mo | HOM |
| **1013978665** | 1 | sptV | 794 | 1.670783811 | 0 | 0 | 0 | 0 | 2 mo | HOM |
| **1013445609** | 1 | sptV | 794 | 1.670783811 | 0 | 0 | 0 | 0 | 4 mo | HOM |
| **1013609015** | 1 | sptV | 794 | 1.670783811 | 0 | 0 | 0 | 0 | 4 mo | HOM |
| **1013608106** | 2 | sptV | 794 | 1.670783811 | 0 | 0 | 0 | 0 | 4 mo | HOM |
| **1011937543** | 1 | sptV | 794 | 1.670783811 | 0.00426879 | 3 | 0.002554962 | 1.795564441 | 8 mo | HOM |
| **1010419538** | 2 | sptV | 794 | 1.670783811 | 0 | 0 | 0 | 0 | 4 mo | WILD |
| **1010419187** | 1 | sptV | 794 | 1.670783811 | 0.000807922 | 1 | 0.000483558 | 0.59852148 | 8 mo | HOM |
| **1013978366** | 1 | sptV | 794 | 1.670783811 | 1.01E-05 | 0 | 6.06E-06 | 0 | 2 mo | HOM |
| **1014416155** | 2 | sptV | 794 | 1.670783811 | 0 | 0 | 0 | 0 | 2 mo | HOM |
| **1013978065** | 2 | sptV | 794 | 1.670783811 | 0 | 0 | 0 | 0 | 2 mo | HOM |
| **1009934102** | 2 | sptV | 794 | 1.670783811 | 1.95E-09 | 0 | 1.17E-09 | 0 | 4 mo | WILD |
| **1009322937** | 2 | sptV | 794 | 1.670783811 | 0 | 0 | 0 | 0 | 8 mo | WILD |
| **1011080637** | 2 | sptV | 794 | 1.670783811 | 0 | 0 | 0 | 0 | 8 mo | HOM |
| **1013609713** | 2 | sptV | 794 | 1.670783811 | 0 | 0 | 0 | 0 | 2 mo | WILD |
| **1010418842** | 1 | sptV | 794 | 1.670783811 | 4.1E-07 | 0 | 2.46E-07 | 0 | 8 mo | HOM |
| **1013288213** | 2 | sptV | 794 | 1.670783811 | 0 | 0 | 0 | 0 | 4 mo | HOM |
| **1008976060** | 2 | sptV | 794 | 1.670783811 | 2.88E-05 | 3 | 1.72E-05 | 1.795564441 | 8 mo | WILD |
| **1013446243** | 1 | sptV | 794 | 1.670783811 | 2.71E-07 | 0 | 1.62E-07 | 0 | 4 mo | HOM |
| **1013288951** | 2 | sptV | 794 | 1.670783811 | 3.91E-08 | 0 | 2.34E-08 | 0 | 4 mo | HOM |
| **1013978665** | 1 | PAG | 795 | 4.83566151 | 4.92E-05 | 6 | 1.02E-05 | 1.240781636 | 2 mo | HOM |
| **1013445609** | 1 | PAG | 795 | 4.83566151 | 0.000128844 | 8 | 2.66E-05 | 1.654375515 | 4 mo | HOM |
| **1013609015** | 1 | PAG | 795 | 4.83566151 | 0.006078147 | 9 | 0.001256942 | 1.861172454 | 4 mo | HOM |
| **1013608106** | 2 | PAG | 795 | 4.835661824 | 0 | 0 | 0 | 0 | 4 mo | HOM |
| **1011937543** | 1 | PAG | 795 | 4.83566151 | 0.025577314 | 65 | 0.00528931 | 13.44180106 | 8 mo | HOM |
| **1010419538** | 2 | PAG | 795 | 4.835661824 | 3.76E-05 | 4 | 7.78E-06 | 0.827187704 | 4 mo | WILD |
| **1010419187** | 1 | PAG | 795 | 4.83566151 | 0.003297857 | 26 | 0.000681987 | 5.376720423 | 8 mo | HOM |
| **1013978366** | 1 | PAG | 795 | 4.83566151 | 0.00029331 | 14 | 6.07E-05 | 2.895157151 | 2 mo | HOM |
| **1014416155** | 2 | PAG | 795 | 4.835661824 | 0 | 0 | 0 | 0 | 2 mo | HOM |
| **1013978065** | 2 | PAG | 795 | 4.835661824 | 1.28E-05 | 1 | 2.65E-06 | 0.206796926 | 2 mo | HOM |
| **1009934102** | 2 | PAG | 795 | 4.835661824 | 4.72E-05 | 1 | 9.77E-06 | 0.206796926 | 4 mo | WILD |
| **1009322937** | 2 | PAG | 795 | 4.835661824 | 0.000652705 | 20 | 0.000134977 | 4.135938518 | 8 mo | WILD |
| **1011080637** | 2 | PAG | 795 | 4.835661824 | 0.01100711 | 20 | 0.002276236 | 4.135938518 | 8 mo | HOM |
| **1013609713** | 2 | PAG | 795 | 4.835661824 | 3.27E-06 | 0 | 6.77E-07 | 0 | 2 mo | WILD |
| **1010418842** | 1 | PAG | 795 | 4.83566151 | 0.004931362 | 36 | 0.00101979 | 7.444689816 | 8 mo | HOM |
| **1013288213** | 2 | PAG | 795 | 4.835661824 | 6.41E-05 | 4 | 1.32E-05 | 0.827187704 | 4 mo | HOM |
| **1008976060** | 2 | PAG | 795 | 4.835661824 | 0.003098789 | 58 | 0.00064082 | 11.9942217 | 8 mo | WILD |
| **1013446243** | 1 | PAG | 795 | 4.83566151 | 0.001572573 | 8 | 0.000325203 | 1.654375515 | 4 mo | HOM |
| **1013288951** | 2 | PAG | 795 | 4.835661824 | 0.002188244 | 17 | 0.000452522 | 3.515547741 | 4 mo | HOM |
| **1013978665** | 1 | Zl | 797 | 1.854139146 | 0 | 0 | 0 | 0 | 2 mo | HOM |
| **1013445609** | 1 | Zl | 797 | 1.854139146 | 0 | 0 | 0 | 0 | 4 mo | HOM |
| **1013609015** | 1 | Zl | 797 | 1.854139146 | 0 | 0 | 0 | 0 | 4 mo | HOM |
| **1013608106** | 2 | Zl | 797 | 1.854139146 | 0 | 0 | 0 | 0 | 4 mo | HOM |
| **1011937543** | 1 | Zl | 797 | 1.854139146 | 0.001330345 | 29 | 0.0007175 | 15.64068159 | 8 mo | HOM |
| **1010419538** | 2 | Zl | 797 | 1.854139146 | 0 | 0 | 0 | 0 | 4 mo | WILD |
| **1010419187** | 1 | Zl | 797 | 1.854139146 | 0.0001077 | 5 | 5.81E-05 | 2.69666924 | 8 mo | HOM |
| **1013978366** | 1 | Zl | 797 | 1.854139146 | 0 | 0 | 0 | 0 | 2 mo | HOM |
| **1014416155** | 2 | Zl | 797 | 1.854139146 | 0 | 0 | 0 | 0 | 2 mo | HOM |
| **1013978065** | 2 | Zl | 797 | 1.854139146 | 2.55E-05 | 1 | 1.38E-05 | 0.539333848 | 2 mo | HOM |
| **1009934102** | 2 | Zl | 797 | 1.854139146 | 0 | 0 | 0 | 0 | 4 mo | WILD |
| **1009322937** | 2 | Zl | 797 | 1.854139146 | 0 | 0 | 0 | 0 | 8 mo | WILD |
| **1011080637** | 2 | Zl | 797 | 1.854139146 | 0.000126198 | 9 | 6.81E-05 | 4.854004631 | 8 mo | HOM |
| **1013609713** | 2 | Zl | 797 | 1.854139146 | 0 | 0 | 0 | 0 | 2 mo | WILD |
| **1010418842** | 1 | Zl | 797 | 1.854139146 | 0.000716641 | 26 | 0.000386509 | 14.02268005 | 8 mo | HOM |
| **1013288213** | 2 | Zl | 797 | 1.854139146 | 0 | 0 | 0 | 0 | 4 mo | HOM |
| **1008976060** | 2 | Zl | 797 | 1.854139146 | 8.82E-06 | 2 | 4.76E-06 | 1.078667696 | 8 mo | WILD |
| **1013446243** | 1 | Zl | 797 | 1.854139146 | 0 | 0 | 0 | 0 | 4 mo | HOM |

| | | | | | | | | | | |
|---|---|---|---|---|---|---|---|---|---|---|
| **1013288951** | 2 | ZI | 797 | 1.854139146 | 0 | 0 | 0 | 0 | 4 mo | HOM |
| **1013978665** | 1 | VIIn | 798 | 0.111328176 | 0 | 0 | 0 | 0 | 2 mo | HOM |
| **1013445609** | 1 | VIIn | 798 | 0.111328176 | 0 | 0 | 0 | 0 | 4 mo | HOM |
| **1013609015** | 1 | VIIn | 798 | 0.111328176 | 0 | 0 | 0 | 0 | 4 mo | HOM |
| **1013608106** | 2 | VIIn | 798 | 0.111328176 | 0 | 0 | 0 | 0 | 4 mo | HOM |
| **1011937543** | 1 | VIIn | 798 | 0.111328176 | 0 | 0 | 0 | 0 | 8 mo | HOM |
| **1010419538** | 2 | VIIn | 798 | 0.111328176 | 0 | 0 | 0 | 0 | 4 mo | WILD |
| **1010419187** | 1 | VIIn | 798 | 0.111328176 | 0 | 0 | 0 | 0 | 8 mo | HOM |
| **1013978366** | 1 | VIIn | 798 | 0.111328176 | 0 | 0 | 0 | 0 | 2 mo | HOM |
| **1014416155** | 2 | VIIn | 798 | 0.111328176 | 0 | 0 | 0 | 0 | 2 mo | HOM |
| **1013978065** | 2 | VIIn | 798 | 0.111328176 | 0 | 0 | 0 | 0 | 2 mo | HOM |
| **1009934102** | 2 | VIIn | 798 | 0.111328176 | 0 | 0 | 0 | 0 | 4 mo | WILD |
| **1009322937** | 2 | VIIn | 798 | 0.111328176 | 0 | 0 | 0 | 0 | 8 mo | WILD |
| **1011080637** | 2 | VIIn | 798 | 0.111328176 | 0 | 0 | 0 | 0 | 8 mo | HOM |
| **1013609713** | 2 | VIIn | 798 | 0.111328176 | 0 | 0 | 0 | 0 | 2 mo | WILD |
| **1010418842** | 1 | VIIn | 798 | 0.111328176 | 7.6E-07 | 0 | 6.83E-06 | 0 | 8 mo | HOM |
| **1013288213** | 2 | VIIn | 798 | 0.111328176 | 0 | 0 | 0 | 0 | 4 mo | HOM |
| **1008976060** | 2 | VIIn | 798 | 0.111328176 | 0 | 0 | 0 | 0 | 8 mo | WILD |
| **1013446243** | 1 | VIIn | 798 | 0.111328176 | 0 | 0 | 0 | 0 | 4 mo | HOM |
| **1013288951** | 2 | VIIn | 798 | 0.111328176 | 0 | 0 | 0 | 0 | 4 mo | HOM |
| **1013978665** | 1 | AIv5 | 800 | 0.742207077 | 0 | 0 | 0 | 0 | 2 mo | HOM |
| **1013445609** | 1 | AIv5 | 800 | 0.742207077 | 4.36E-05 | 3 | 5.88E-05 | 4.041998647 | 4 mo | HOM |
| **1013609015** | 1 | AIv5 | 800 | 0.742207077 | 2.98E-05 | 3 | 4.01E-05 | 4.041998647 | 4 mo | HOM |
| **1013608106** | 2 | AIv5 | 800 | 0.742207077 | 0 | 0 | 0 | 0 | 4 mo | HOM |
| **1011937543** | 1 | AIv5 | 800 | 0.742207077 | 0.001024468 | 22 | 0.0013803 | 29.64132341 | 8 mo | HOM |
| **1010419538** | 2 | AIv5 | 800 | 0.742207077 | 0 | 0 | 0 | 0 | 4 mo | WILD |
| **1010419187** | 1 | AIv5 | 800 | 0.742207077 | 0.001169722 | 36 | 0.001576004 | 48.50398376 | 8 mo | HOM |
| **1013978366** | 1 | AIv5 | 800 | 0.742207077 | 0 | 0 | 0 | 0 | 2 mo | HOM |
| **1014416155** | 2 | AIv5 | 800 | 0.742207077 | 0 | 0 | 0 | 0 | 2 mo | HOM |
| **1013978065** | 2 | AIv5 | 800 | 0.742207077 | 0 | 0 | 0 | 0 | 2 mo | HOM |
| **1009934102** | 2 | AIv5 | 800 | 0.742207077 | 0 | 0 | 0 | 0 | 4 mo | WILD |
| **1009322937** | 2 | AIv5 | 800 | 0.742207077 | 3E-07 | 0 | 4.04E-07 | 0 | 8 mo | WILD |
| **1011080637** | 2 | AIv5 | 800 | 0.742207077 | 0.000428904 | 20 | 0.000577876 | 26.94665764 | 8 mo | HOM |
| **1013609713** | 2 | AIv5 | 800 | 0.742207077 | 0 | 0 | 0 | 0 | 2 mo | WILD |
| **1010418842** | 1 | AIv5 | 800 | 0.742207077 | 0.000430985 | 14 | 0.00058068 | 18.86266035 | 8 mo | HOM |
| **1013288213** | 2 | AIv5 | 800 | 0.742207077 | 2.9E-05 | 2 | 3.91E-05 | 2.694665764 | 4 mo | HOM |
| **1008976060** | 2 | AIv5 | 800 | 0.742207077 | 0.000111803 | 4 | 0.000150636 | 5.389331529 | 8 mo | WILD |
| **1013446243** | 1 | AIv5 | 800 | 0.742207077 | 3.73E-05 | 6 | 5.02E-05 | 8.083997293 | 4 mo | HOM |
| **1013288951** | 2 | AIv5 | 800 | 0.742207077 | 2.59E-06 | 0 | 3.49E-06 | 0 | 4 mo | HOM |
| **1013978665** | 1 | sm | 802 | 0.278881912 | 4.27E-05 | 0 | 0.000153179 | 0 | 2 mo | HOM |
| **1013445609** | 1 | sm | 802 | 0.278881912 | 1.26E-05 | 1 | 4.53E-05 | 3.585747069 | 4 mo | HOM |
| **1013609015** | 1 | sm | 802 | 0.278881912 | 1.79E-05 | 0 | 6.41E-05 | 0 | 4 mo | HOM |
| **1013608106** | 2 | sm | 802 | 0.278881912 | 0 | 0 | 0 | 0 | 4 mo | HOM |
| **1011937543** | 1 | sm | 802 | 0.278881912 | 0.000104289 | 3 | 0.000373955 | 10.75724121 | 8 mo | HOM |
| **1010419538** | 2 | sm | 802 | 0.278881912 | 3.14E-05 | 2 | 0.000112736 | 7.171494138 | 4 mo | WILD |
| **1010419187** | 1 | sm | 802 | 0.278881912 | 0.000415191 | 8 | 0.001488768 | 28.68597655 | 8 mo | HOM |
| **1013978366** | 1 | sm | 802 | 0.278881912 | 4.12E-05 | 2 | 0.000147735 | 7.171494138 | 2 mo | HOM |
| **1014416155** | 2 | sm | 802 | 0.278881912 | 0 | 0 | 0 | 0 | 2 mo | HOM |
| **1013978065** | 2 | sm | 802 | 0.278881912 | 0 | 0 | 0 | 0 | 2 mo | HOM |
| **1009934102** | 2 | sm | 802 | 0.278881912 | 0 | 0 | 0 | 0 | 4 mo | WILD |
| **1009322937** | 2 | sm | 802 | 0.278881912 | 3.15E-08 | 0 | 1.13E-07 | 0 | 8 mo | WILD |
| **1011080637** | 2 | sm | 802 | 0.278881912 | 0.000196645 | 4 | 0.00070512 | 14.34298828 | 8 mo | HOM |
| **1013609713** | 2 | sm | 802 | 0.278881912 | 0 | 0 | 0 | 0 | 2 mo | WILD |
| **1010418842** | 1 | sm | 802 | 0.278881912 | 0.000232981 | 4 | 0.00083541 | 14.34298828 | 8 mo | HOM |
| **1013288213** | 2 | sm | 802 | 0.278881912 | 0 | 0 | 0 | 0 | 4 mo | HOM |
| **1008976060** | 2 | sm | 802 | 0.278881912 | 9.82E-05 | 1 | 0.00035206 | 3.585747069 | 8 mo | WILD |
| **1013446243** | 1 | sm | 802 | 0.278881912 | 4.11E-05 | 2 | 0.00014754 | 7.171494138 | 4 mo | HOM |
| **1013288951** | 2 | sm | 802 | 0.278881912 | 1.59E-05 | 1 | 5.69E-05 | 3.585747069 | 4 mo | HOM |
| **1013978665** | 1 | PAL | 803 | 9.646043098 | 4.65E-05 | 1 | 4.82E-06 | 0.103669452 | 2 mo | HOM |

| | | | | | | | | | | |
|---|---|---|---|---|---|---|---|---|---|---|
| 1013445609 | 1 | PAL | 803 | 9.64786135 | 7.52E-05 | 7 | 7.79E-06 | 0.725549399 | 4 mo | HOM |
| 1013609015 | 1 | PAL | 803 | 9.64786135 | 0.001184534 | 15 | 0.000122777 | 1.554748711 | 4 mo | HOM |
| 1013608106 | 2 | PAL | 803 | 9.64786135 | 0 | 0 | 0 | 0 | 4 mo | HOM |
| 1011937543 | 1 | PAL | 803 | 9.64786135 | 0.006431254 | 137 | 0.000666599 | 14.20003823 | 8 mo | HOM |
| 1010419538 | 2 | PAL | 803 | 9.64786135 | 5.85E-05 | 1 | 6.07E-06 | 0.103649914 | 4 mo | WILD |
| 1010419187 | 1 | PAL | 803 | 9.64786135 | 0.003491611 | 164 | 0.000361905 | 16.99858591 | 8 mo | HOM |
| 1013978366 | 1 | PAL | 803 | 9.64786135 | 3.75E-06 | 1 | 3.89E-07 | 0.103649914 | 2 mo | HOM |
| 1014416155 | 2 | PAL | 803 | 9.64786135 | 0 | 0 | 0 | 0 | 2 mo | HOM |
| 1013978065 | 2 | PAL | 803 | 9.64786135 | 8.72E-08 | 0 | 9.04E-09 | 0 | 2 mo | HOM |
| 1009934102 | 2 | PAL | 803 | 9.64786135 | 3.05E-05 | 3 | 3.16E-06 | 0.310949742 | 4 mo | WILD |
| 1009322937 | 2 | PAL | 803 | 9.64786135 | 0.000156521 | 8 | 1.62E-05 | 0.829199313 | 8 mo | WILD |
| 1011080637 | 2 | PAL | 803 | 9.64786135 | 0.003801657 | 146 | 0.000394041 | 15.13288746 | 8 mo | HOM |
| 1013609713 | 2 | PAL | 803 | 9.64786135 | 1.86E-06 | 0 | 1.93E-07 | 0 | 2 mo | WILD |
| 1010418842 | 1 | PAL | 803 | 9.64786135 | 0.006393251 | 221 | 0.00066266 | 22.90663101 | 8 mo | HOM |
| 1013288213 | 2 | PAL | 803 | 9.64786135 | 9.4E-05 | 7 | 9.74E-06 | 0.725549399 | 4 mo | HOM |
| 1008976060 | 2 | PAL | 803 | 9.64786135 | 0.000639191 | 14 | 6.63E-05 | 1.451098797 | 8 mo | WILD |
| 1013446243 | 1 | PAL | 803 | 9.64786135 | 0.000638933 | 18 | 6.62E-05 | 1.865698453 | 4 mo | HOM |
| 1013288951 | 2 | PAL | 803 | 9.64786135 | 0.000716566 | 13 | 7.43E-05 | 1.347448883 | 4 mo | HOM |
| 1013978665 | 1 | FF | 804 | 0.248092606 | 0 | 0 | 0 | 0 | 2 mo | HOM |
| 1013445609 | 1 | FF | 804 | 0.248092606 | 0 | 0 | 0 | 0 | 4 mo | HOM |
| 1013609015 | 1 | FF | 804 | 0.248092606 | 0 | 0 | 0 | 0 | 4 mo | HOM |
| 1013608106 | 2 | FF | 804 | 0.248092606 | 0 | 0 | 0 | 0 | 4 mo | HOM |
| 1011937543 | 1 | FF | 804 | 0.248092606 | 8.84E-05 | 2 | 0.000356331 | 8.061505885 | 8 mo | HOM |
| 1010419538 | 2 | FF | 804 | 0.248092606 | 0 | 0 | 0 | 0 | 4 mo | WILD |
| 1010419187 | 1 | FF | 804 | 0.248092606 | 0 | 0 | 0 | 0 | 8 mo | HOM |
| 1013978366 | 1 | FF | 804 | 0.248092606 | 0 | 0 | 0 | 0 | 2 mo | HOM |
| 1014416155 | 2 | FF | 804 | 0.248092606 | 0 | 0 | 0 | 0 | 2 mo | HOM |
| 1013978065 | 2 | FF | 804 | 0.248092606 | 0 | 0 | 0 | 0 | 2 mo | HOM |
| 1009934102 | 2 | FF | 804 | 0.248092606 | 0 | 0 | 0 | 0 | 4 mo | WILD |
| 1009322937 | 2 | FF | 804 | 0.248092606 | 0 | 0 | 0 | 0 | 8 mo | WILD |
| 1011080637 | 2 | FF | 804 | 0.248092606 | 6.31E-07 | 0 | 2.54E-06 | 0 | 8 mo | HOM |
| 1013609713 | 2 | FF | 804 | 0.248092606 | 0 | 0 | 0 | 0 | 2 mo | WILD |
| 1010418842 | 1 | FF | 804 | 0.248092606 | 1.49E-08 | 0 | 6.03E-08 | 0 | 8 mo | HOM |
| 1013288213 | 2 | FF | 804 | 0.248092606 | 0 | 0 | 0 | 0 | 4 mo | HOM |
| 1008976060 | 2 | FF | 804 | 0.248092606 | 0 | 0 | 0 | 0 | 8 mo | WILD |
| 1013446243 | 1 | FF | 804 | 0.248092606 | 0 | 0 | 0 | 0 | 4 mo | HOM |
| 1013288951 | 2 | FF | 804 | 0.248092606 | 0 | 0 | 0 | 0 | 4 mo | HOM |
| 1013978665 | 1 | VISpm1 | 805 | 0.196412404 | 5.31E-06 | 1 | 2.71E-05 | 5.091328153 | 2 mo | HOM |
| 1013445609 | 1 | VISpm1 | 805 | 0.196412404 | 2.07E-05 | 1 | 0.000105337 | 5.091328153 | 4 mo | HOM |
| 1013609015 | 1 | VISpm1 | 805 | 0.196412404 | 8.24E-05 | 3 | 0.000419659 | 15.27398446 | 4 mo | HOM |
| 1013608106 | 2 | VISpm1 | 805 | 0.196412404 | 0 | 0 | 0 | 0 | 4 mo | HOM |
| 1011937543 | 1 | VISpm1 | 805 | 0.196412404 | 0.000303887 | 1 | 0.001547187 | 5.091328153 | 8 mo | HOM |
| 1010419538 | 2 | VISpm1 | 805 | 0.196412404 | 0 | 0 | 0 | 0 | 4 mo | WILD |
| 1010419187 | 1 | VISpm1 | 805 | 0.196412404 | 0.000743357 | 11 | 0.003784677 | 56.00460968 | 8 mo | HOM |
| 1013978366 | 1 | VISpm1 | 805 | 0.196412404 | 0 | 0 | 0 | 0 | 2 mo | HOM |
| 1014416155 | 2 | VISpm1 | 805 | 0.196412404 | 0 | 0 | 0 | 0 | 2 mo | HOM |
| 1013978065 | 2 | VISpm1 | 805 | 0.196412404 | 0 | 0 | 0 | 0 | 2 mo | HOM |
| 1009934102 | 2 | VISpm1 | 805 | 0.196412404 | 0 | 0 | 0 | 0 | 4 mo | WILD |
| 1009322937 | 2 | VISpm1 | 805 | 0.196412404 | 5.52E-06 | 1 | 2.81E-05 | 5.091328153 | 8 mo | WILD |
| 1011080637 | 2 | VISpm1 | 805 | 0.196412404 | 0.000413327 | 17 | 0.002104385 | 86.5525786 | 8 mo | HOM |
| 1013609713 | 2 | VISpm1 | 805 | 0.196412404 | 0 | 0 | 0 | 0 | 2 mo | WILD |
| 1010418842 | 1 | VISpm1 | 805 | 0.196412404 | 0.000738129 | 13 | 0.003758057 | 66.18726599 | 8 mo | HOM |
| 1013288213 | 2 | VISpm1 | 805 | 0.196412404 | 1.08E-05 | 1 | 5.51E-05 | 5.091328153 | 4 mo | HOM |
| 1008976060 | 2 | VISpm1 | 805 | 0.196412404 | 0.000715995 | 5 | 0.003645365 | 25.45664076 | 8 mo | WILD |
| 1013446243 | 1 | VISpm1 | 805 | 0.196412404 | 0.000286264 | 4 | 0.001457465 | 20.36531261 | 4 mo | HOM |
| 1013288951 | 2 | VISpm1 | 805 | 0.196412404 | 2.23E-05 | 2 | 0.000113788 | 10.18265631 | 4 mo | HOM |
| 1013978665 | 1 | SSs2/3 | 806 | 2.21110391 | 0 | 0 | 0 | 0 | 2 mo | HOM |
| 1013445609 | 1 | SSs2/3 | 806 | 2.21110391 | 0.000305425 | 17 | 0.000138133 | 7.688467249 | 4 mo | HOM |
| 1013609015 | 1 | SSs2/3 | 806 | 2.21110391 | 0.000617562 | 19 | 0.0002793 | 8.592992808 | 4 mo | HOM |

| | | | | | | | | | | |
|---|---|---|---|---|---|---|---|---|---|---|
| **1013608106** | 2 | SSs2/3 | 806 | 2.21110391 | 0 | 0 | 0 | 0 | 4 mo | HOM |
| **1011937543** | 1 | SSs2/3 | 806 | 2.21110391 | 0.002822172 | 60 | 0.001276363 | 27.13576676 | 8 mo | HOM |
| **1010419538** | 2 | SSs2/3 | 806 | 2.21110391 | 0 | 0 | 0 | 0 | 4 mo | WILD |
| **1010419187** | 1 | SSs2/3 | 806 | 2.21110391 | 0.003498782 | 104 | 0.001582369 | 47.03532905 | 8 mo | HOM |
| **1013978366** | 1 | SSs2/3 | 806 | 2.21110391 | 1.77E-05 | 1 | 8E-06 | 0.452262779 | 2 mo | HOM |
| **1014416155** | 2 | SSs2/3 | 806 | 2.21110391 | 0 | 0 | 0 | 0 | 2 mo | HOM |
| **1013978065** | 2 | SSs2/3 | 806 | 2.21110391 | 1.41E-05 | 1 | 6.39E-06 | 0.452262779 | 2 mo | HOM |
| **1009934102** | 2 | SSs2/3 | 806 | 2.21110391 | 0 | 0 | 0 | 0 | 4 mo | WILD |
| **1009322937** | 2 | SSs2/3 | 806 | 2.21110391 | 0 | 0 | 0 | 0 | 8 mo | WILD |
| **1011080637** | 2 | SSs2/3 | 806 | 2.21110391 | 0.002198401 | 74 | 0.000994255 | 33.46744567 | 8 mo | HOM |
| **1013609713** | 2 | SSs2/3 | 806 | 2.21110391 | 0 | 0 | 0 | 0 | 2 mo | WILD |
| **1010418842** | 1 | SSs2/3 | 806 | 2.21110391 | 0.002029015 | 78 | 0.000917648 | 35.27649679 | 8 mo | HOM |
| **1013288213** | 2 | SSs2/3 | 806 | 2.21110391 | 8.49E-05 | 8 | 3.84E-05 | 3.618102235 | 4 mo | HOM |
| **1008976060** | 2 | SSs2/3 | 806 | 2.21110391 | 1.29E-05 | 0 | 5.83E-06 | 0 | 8 mo | WILD |
| **1013446243** | 1 | SSs2/3 | 806 | 2.21110391 | 0.000558507 | 18 | 0.000252592 | 8.140730029 | 4 mo | HOM |
| **1013288951** | 2 | SSs2/3 | 806 | 2.21110391 | 0.000479532 | 17 | 0.000216874 | 7.688467249 | 4 mo | HOM |
| **1013978665** | 1 | PALc | 809 | 1.389215126 | 0 | 0 | 0 | 0 | 2 mo | HOM |
| **1013445609** | 1 | PALc | 809 | 1.389215126 | 3.17E-05 | 3 | 2.28E-05 | 2.159492755 | 4 mo | HOM |
| **1013609015** | 1 | PALc | 809 | 1.389215126 | 1.21E-05 | 2 | 8.72E-06 | 1.439661837 | 4 mo | HOM |
| **1013608106** | 2 | PALc | 809 | 1.389215126 | 0 | 0 | 0 | 0 | 4 mo | HOM |
| **1011937543** | 1 | PALc | 809 | 1.389215126 | 0.000494286 | 21 | 0.000355802 | 15.11644929 | 8 mo | HOM |
| **1010419538** | 2 | PALc | 809 | 1.389215126 | 0 | 0 | 0 | 0 | 4 mo | WILD |
| **1010419187** | 1 | PALc | 809 | 1.389215126 | 0.001028202 | 39 | 0.000740131 | 28.07340582 | 8 mo | HOM |
| **1013978366** | 1 | PALc | 809 | 1.389215126 | 0 | 0 | 0 | 0 | 2 mo | HOM |
| **1014416155** | 2 | PALc | 809 | 1.389215126 | 0 | 0 | 0 | 0 | 2 mo | HOM |
| **1013978065** | 2 | PALc | 809 | 1.389215126 | 0 | 0 | 0 | 0 | 2 mo | HOM |
| **1009934102** | 2 | PALc | 809 | 1.389215126 | 0 | 0 | 0 | 0 | 4 mo | WILD |
| **1009322937** | 2 | PALc | 809 | 1.389215126 | 1.56E-06 | 0 | 1.12E-06 | 0 | 8 mo | WILD |
| **1011080637** | 2 | PALc | 809 | 1.389215126 | 0.001052575 | 40 | 0.000757676 | 28.79323674 | 8 mo | HOM |
| **1013609713** | 2 | PALc | 809 | 1.389215126 | 0 | 0 | 0 | 0 | 2 mo | WILD |
| **1010418842** | 1 | PALc | 809 | 1.389215126 | 0.001455435 | 42 | 0.001047667 | 30.23289857 | 8 mo | HOM |
| **1013288213** | 2 | PALc | 809 | 1.389215126 | 4.92E-05 | 4 | 3.54E-05 | 2.879323674 | 4 mo | HOM |
| **1008976060** | 2 | PALc | 809 | 1.389215126 | 5E-05 | 3 | 3.6E-05 | 2.159492755 | 8 mo | WILD |
| **1013446243** | 1 | PALc | 809 | 1.389215126 | 5.41E-05 | 4 | 3.9E-05 | 2.879323674 | 4 mo | HOM |
| **1013288951** | 2 | PALc | 809 | 1.389215126 | 3.56E-05 | 4 | 2.56E-05 | 2.879323674 | 4 mo | HOM |
| **1013978665** | 1 | ACAv6a | 810 | 0.374375523 | 0 | 0 | 0 | 0 | 2 mo | HOM |
| **1013445609** | 1 | ACAv6a | 810 | 0.374375523 | 1.43E-05 | 1 | 3.81E-05 | 2.671114798 | 4 mo | HOM |
| **1013609015** | 1 | ACAv6a | 810 | 0.374375523 | 2.26E-05 | 2 | 6.03E-05 | 5.342229596 | 4 mo | HOM |
| **1013608106** | 2 | ACAv6a | 810 | 0.374375523 | 0 | 0 | 0 | 0 | 4 mo | HOM |
| **1011937543** | 1 | ACAv6a | 810 | 0.374375523 | 7.13E-05 | 3 | 0.000190415 | 8.013344394 | 8 mo | HOM |
| **1010419538** | 2 | ACAv6a | 810 | 0.374375523 | 0 | 0 | 0 | 0 | 4 mo | WILD |
| **1010419187** | 1 | ACAv6a | 810 | 0.374375523 | 0.000554628 | 16 | 0.001481474 | 42.73783677 | 8 mo | HOM |
| **1013978366** | 1 | ACAv6a | 810 | 0.374375523 | 0 | 0 | 0 | 0 | 2 mo | HOM |
| **1014416155** | 2 | ACAv6a | 810 | 0.374375523 | 0 | 0 | 0 | 0 | 2 mo | HOM |
| **1013978065** | 2 | ACAv6a | 810 | 0.374375523 | 0 | 0 | 0 | 0 | 2 mo | HOM |
| **1009934102** | 2 | ACAv6a | 810 | 0.374375523 | 0 | 0 | 0 | 0 | 4 mo | WILD |
| **1009322937** | 2 | ACAv6a | 810 | 0.374375523 | 0 | 0 | 0 | 0 | 8 mo | WILD |
| **1011080637** | 2 | ACAv6a | 810 | 0.374375523 | 0.00022301 | 14 | 0.000595685 | 37.39560717 | 8 mo | HOM |
| **1013609713** | 2 | ACAv6a | 810 | 0.374375523 | 0 | 0 | 0 | 0 | 2 mo | WILD |
| **1010418842** | 1 | ACAv6a | 810 | 0.374375523 | 0.000210803 | 11 | 0.000563078 | 29.38226278 | 8 mo | HOM |
| **1013288213** | 2 | ACAv6a | 810 | 0.374375523 | 5.94E-06 | 1 | 1.59E-05 | 2.671114798 | 4 mo | HOM |
| **1008976060** | 2 | ACAv6a | 810 | 0.374375523 | 2.39E-06 | 1 | 6.4E-06 | 2.671114798 | 8 mo | WILD |
| **1013446243** | 1 | ACAv6a | 810 | 0.374375523 | 2.17E-05 | 1 | 5.81E-05 | 2.671114798 | 4 mo | HOM |
| **1013288951** | 2 | ACAv6a | 810 | 0.374375523 | 5.71E-05 | 3 | 0.00015251 | 8.013344394 | 4 mo | HOM |
| **1013978665** | 1 | ICc | 811 | 1.160624662 | 0 | 0 | 0 | 0 | 2 mo | HOM |
| **1013445609** | 1 | ICc | 811 | 1.160624662 | 0 | 0 | 0 | 0 | 4 mo | HOM |
| **1013609015** | 1 | ICc | 811 | 1.160624662 | 1.34E-06 | 0 | 1.15E-06 | 0 | 4 mo | HOM |
| **1013608106** | 2 | ICc | 811 | 1.160624662 | 0 | 0 | 0 | 0 | 4 mo | HOM |
| **1011937543** | 1 | ICc | 811 | 1.160624662 | 0.002929465 | 18 | 0.002524042 | 15.50888981 | 8 mo | HOM |

| | | | | | | | | | | |
|---|---|---|---|---|---|---|---|---|---|---|
| **1010419538** | 2 | ICc | 811 | 1.160624662 | 0 | 0 | 0 | 0 | 4 mo | WILD |
| **1010419187** | 1 | ICc | 811 | 1.160624662 | 0.000133756 | 5 | 0.000115244 | 4.308024947 | 8 mo | HOM |
| **1013978366** | 1 | ICc | 811 | 1.160624662 | 1.71E-05 | 1 | 1.47E-05 | 0.861604989 | 2 mo | HOM |
| **1014416155** | 2 | ICc | 811 | 1.160624662 | 0 | 0 | 0 | 0 | 2 mo | HOM |
| **1013978065** | 2 | ICc | 811 | 1.160624662 | 0 | 0 | 0 | 0 | 2 mo | HOM |
| **1009934102** | 2 | ICc | 811 | 1.160624662 | 0 | 0 | 0 | 0 | 4 mo | WILD |
| **1009322937** | 2 | ICc | 811 | 1.160624662 | 0 | 0 | 0 | 0 | 8 mo | WILD |
| **1011080637** | 2 | ICc | 811 | 1.160624662 | 0.000110835 | 8 | 9.55E-05 | 6.892839916 | 8 mo | HOM |
| **1013609713** | 2 | ICc | 811 | 1.160624662 | 0 | 0 | 0 | 0 | 2 mo | WILD |
| **1010418842** | 1 | ICc | 811 | 1.160624662 | 0.0001948 | 7 | 0.00016784 | 6.031234926 | 8 mo | HOM |
| **1013288213** | 2 | ICc | 811 | 1.160624662 | 0 | 0 | 0 | 0 | 4 mo | HOM |
| **1008976060** | 2 | ICc | 811 | 1.160624662 | 2.75E-05 | 4 | 2.37E-05 | 3.446419958 | 8 mo | WILD |
| **1013446243** | 1 | ICc | 811 | 1.160624662 | 0 | 0 | 0 | 0 | 4 mo | HOM |
| **1013288951** | 2 | ICc | 811 | 1.160624662 | 6.79E-06 | 1 | 5.85E-06 | 0.861604989 | 4 mo | HOM |
| **1013978665** | 1 | dscp | 812 | 0.047720022 | 0 | 0 | 0 | 0 | 2 mo | HOM |
| **1013445609** | 1 | dscp | 812 | 0.047720022 | 0 | 0 | 0 | 0 | 4 mo | HOM |
| **1013609015** | 1 | dscp | 812 | 0.047720022 | 0 | 0 | 0 | 0 | 4 mo | HOM |
| **1013608106** | 2 | dscp | 812 | 0.047720022 | 0 | 0 | 0 | 0 | 4 mo | HOM |
| **1011937543** | 1 | dscp | 812 | 0.047720022 | 0 | 0 | 0 | 0 | 8 mo | HOM |
| **1010419538** | 2 | dscp | 812 | 0.047720022 | 0 | 0 | 0 | 0 | 4 mo | WILD |
| **1010419187** | 1 | dscp | 812 | 0.047720022 | 0 | 0 | 0 | 0 | 8 mo | HOM |
| **1013978366** | 1 | dscp | 812 | 0.047720022 | 0 | 0 | 0 | 0 | 2 mo | HOM |
| **1014416155** | 2 | dscp | 812 | 0.047720022 | 0 | 0 | 0 | 0 | 2 mo | HOM |
| **1013978065** | 2 | dscp | 812 | 0.047720022 | 0 | 0 | 0 | 0 | 2 mo | HOM |
| **1009934102** | 2 | dscp | 812 | 0.047720022 | 0 | 0 | 0 | 0 | 4 mo | WILD |
| **1009322937** | 2 | dscp | 812 | 0.047720022 | 0 | 0 | 0 | 0 | 8 mo | WILD |
| **1011080637** | 2 | dscp | 812 | 0.047720022 | 0 | 0 | 0 | 0 | 8 mo | HOM |
| **1013609713** | 2 | dscp | 812 | 0.047720022 | 0 | 0 | 0 | 0 | 2 mo | WILD |
| **1010418842** | 1 | dscp | 812 | 0.047720022 | 0 | 0 | 0 | 0 | 8 mo | HOM |
| **1013288213** | 2 | dscp | 812 | 0.047720022 | 0 | 0 | 0 | 0 | 4 mo | HOM |
| **1008976060** | 2 | dscp | 812 | 0.047720022 | 0 | 0 | 0 | 0 | 8 mo | WILD |
| **1013446243** | 1 | dscp | 812 | 0.047720022 | 0 | 0 | 0 | 0 | 4 mo | HOM |
| **1013288951** | 2 | dscp | 812 | 0.047720022 | 0 | 0 | 0 | 0 | 4 mo | HOM |
| **1013978665** | 1 | DP | 814 | 0.519975142 | 0 | 0 | 0 | 0 | 2 mo | HOM |
| **1013445609** | 1 | DP | 814 | 0.519975142 | 1.32E-05 | 1 | 2.55E-05 | 1.923168856 | 4 mo | HOM |
| **1013609015** | 1 | DP | 814 | 0.519975142 | 2.75E-06 | 0 | 5.29E-06 | 0 | 4 mo | HOM |
| **1013608106** | 2 | DP | 814 | 0.519975142 | 0 | 0 | 0 | 0 | 4 mo | HOM |
| **1011937543** | 1 | DP | 814 | 0.519975142 | 0.000190734 | 4 | 0.000366813 | 7.692675426 | 8 mo | HOM |
| **1010419538** | 2 | DP | 814 | 0.519975142 | 0 | 0 | 0 | 0 | 4 mo | WILD |
| **1010419187** | 1 | DP | 814 | 0.519975142 | 0.000607631 | 14 | 0.001168577 | 26.92436399 | 8 mo | HOM |
| **1013978366** | 1 | DP | 814 | 0.519975142 | 0 | 0 | 0 | 0 | 2 mo | HOM |
| **1014416155** | 2 | DP | 814 | 0.519975142 | 8.48E-06 | 1 | 1.63E-05 | 1.923168856 | 2 mo | HOM |
| **1013978065** | 2 | DP | 814 | 0.519975142 | 0 | 0 | 0 | 0 | 2 mo | HOM |
| **1009934102** | 2 | DP | 814 | 0.519975142 | 4.23E-07 | 0 | 8.14E-07 | 0 | 4 mo | WILD |
| **1009322937** | 2 | DP | 814 | 0.519975142 | 4.8E-05 | 2 | 9.23E-05 | 3.846337713 | 8 mo | WILD |
| **1011080637** | 2 | DP | 814 | 0.519975142 | 0.000407574 | 20 | 0.000783834 | 38.46337713 | 8 mo | HOM |
| **1013609713** | 2 | DP | 814 | 0.519975142 | 0 | 0 | 0 | 0 | 2 mo | WILD |
| **1010418842** | 1 | DP | 814 | 0.519975142 | 0.001672591 | 24 | 0.003216675 | 46.15605256 | 8 mo | HOM |
| **1013288213** | 2 | DP | 814 | 0.519975142 | 0.00214836 | 8 | 0.004131658 | 15.38535085 | 4 mo | HOM |
| **1008976060** | 2 | DP | 814 | 0.519975142 | 5.28E-05 | 1 | 0.000101635 | 1.923168856 | 8 mo | WILD |
| **1013446243** | 1 | DP | 814 | 0.519975142 | 0.000108602 | 0 | 0.00020886 | 0 | 4 mo | HOM |
| **1013288951** | 2 | DP | 814 | 0.519975142 | 0.000199419 | 12 | 0.000383516 | 23.07802628 | 4 mo | HOM |
| **1013978665** | 1 | AUDp4 | 816 | 0.242026641 | 0 | 0 | 0 | 0 | 2 mo | HOM |
| **1013445609** | 1 | AUDp4 | 816 | 0.242026641 | 2.44E-05 | 1 | 0.000100845 | 4.131776554 | 4 mo | HOM |
| **1013609015** | 1 | AUDp4 | 816 | 0.242026641 | 6.92E-05 | 2 | 0.000286109 | 8.263553109 | 4 mo | HOM |
| **1013608106** | 2 | AUDp4 | 816 | 0.242026641 | 0 | 0 | 0 | 0 | 4 mo | HOM |
| **1011937543** | 1 | AUDp4 | 816 | 0.242026641 | 0.000475642 | 9 | 0.001965246 | 37.18598899 | 8 mo | HOM |
| **1010419538** | 2 | AUDp4 | 816 | 0.242026641 | 0 | 0 | 0 | 0 | 4 mo | WILD |
| **1010419187** | 1 | AUDp4 | 816 | 0.242026641 | 0.000334312 | 11 | 0.001381303 | 45.4495421 | 8 mo | HOM |

| | | | | | | | | | | |
|---|---|---|---|---|---|---|---|---|---|---|
| 1013978366 | 1 | AUDp4 | 816 | 0.242026641 | 0 | 0 | 0 | 0 | 2 mo | HOM |
| 1014416155 | 2 | AUDp4 | 816 | 0.242026641 | 0 | 0 | 0 | 0 | 2 mo | HOM |
| 1013978065 | 2 | AUDp4 | 816 | 0.242026641 | 0 | 0 | 0 | 0 | 2 mo | HOM |
| 1009934102 | 2 | AUDp4 | 816 | 0.242026641 | 0 | 0 | 0 | 0 | 4 mo | WILD |
| 1009322937 | 2 | AUDp4 | 816 | 0.242026641 | 0 | 0 | 0 | 0 | 8 mo | WILD |
| 1011080637 | 2 | AUDp4 | 816 | 0.242026641 | 0.000290502 | 7 | 0.001200288 | 28.92243588 | 8 mo | HOM |
| 1013609713 | 2 | AUDp4 | 816 | 0.242026641 | 0 | 0 | 0 | 0 | 2 mo | WILD |
| 1010418842 | 1 | AUDp4 | 816 | 0.242026641 | 0.000558652 | 12 | 0.002308226 | 49.58131865 | 8 mo | HOM |
| 1013288213 | 2 | AUDp4 | 816 | 0.242026641 | 1.38E-05 | 1 | 5.71E-05 | 4.131776554 | 4 mo | HOM |
| 1008976060 | 2 | AUDp4 | 816 | 0.242026641 | 0 | 0 | 0 | 0 | 8 mo | WILD |
| 1013446243 | 1 | AUDp4 | 816 | 0.242026641 | 0.000129349 | 5 | 0.000534441 | 20.65888277 | 4 mo | HOM |
| 1013288951 | 2 | AUDp4 | 816 | 0.242026641 | 7.85E-06 | 0 | 3.24E-05 | 0 | 4 mo | HOM |
| 1013978665 | 1 | PALd | 818 | 2.048493177 | 0 | 0 | 0 | 0 | 2 mo | HOM |
| 1013445609 | 1 | PALd | 818 | 2.048493177 | 1.3E-05 | 1 | 6.33E-06 | 0.488163696 | 4 mo | HOM |
| 1013609015 | 1 | PALd | 818 | 2.048493177 | 1.6E-05 | 1 | 7.8E-06 | 0.488163696 | 4 mo | HOM |
| 1013608106 | 2 | PALd | 818 | 2.048493177 | 0 | 0 | 0 | 0 | 4 mo | HOM |
| 1011937543 | 1 | PALd | 818 | 2.048493177 | 0.002451889 | 42 | 0.001196923 | 20.50287522 | 8 mo | HOM |
| 1010419538 | 2 | PALd | 818 | 2.048493177 | 0 | 0 | 0 | 0 | 4 mo | WILD |
| 1010419187 | 1 | PALd | 818 | 2.048493177 | 0.000381174 | 28 | 0.000186075 | 13.66858348 | 8 mo | HOM |
| 1013978366 | 1 | PALd | 818 | 2.048493177 | 0 | 0 | 0 | 0 | 2 mo | HOM |
| 1014416155 | 2 | PALd | 818 | 2.048493177 | 0 | 0 | 0 | 0 | 2 mo | HOM |
| 1013978065 | 2 | PALd | 818 | 2.048493177 | 0 | 0 | 0 | 0 | 2 mo | HOM |
| 1009934102 | 2 | PALd | 818 | 2.048493177 | 0 | 0 | 0 | 0 | 4 mo | WILD |
| 1009322937 | 2 | PALd | 818 | 2.048493177 | 0 | 0 | 0 | 0 | 8 mo | WILD |
| 1011080637 | 2 | PALd | 818 | 2.048493177 | 0.000458844 | 19 | 0.000223991 | 9.27511022 | 8 mo | HOM |
| 1013609713 | 2 | PALd | 818 | 2.048493177 | 0 | 0 | 0 | 0 | 2 mo | WILD |
| 1010418842 | 1 | PALd | 818 | 2.048493177 | 0.000857067 | 45 | 0.000418389 | 21.96736631 | 8 mo | HOM |
| 1013288213 | 2 | PALd | 818 | 2.048493177 | 7.94E-06 | 1 | 3.87E-06 | 0.488163696 | 4 mo | HOM |
| 1008976060 | 2 | PALd | 818 | 2.048493177 | 5.02E-06 | 1 | 2.45E-06 | 0.488163696 | 8 mo | WILD |
| 1013446243 | 1 | PALd | 818 | 2.048493177 | 9.66E-06 | 1 | 4.72E-06 | 0.488163696 | 4 mo | HOM |
| 1013288951 | 2 | PALd | 818 | 2.048493177 | 7.23E-06 | 1 | 3.53E-06 | 0.488163696 | 4 mo | HOM |
| 1013978665 | 1 | ACAv6b | 819 | 0.063937826 | 0 | 0 | 0 | 0 | 2 mo | HOM |
| 1013445609 | 1 | ACAv6b | 819 | 0.063937826 | 0 | 0 | 0 | 0 | 4 mo | HOM |
| 1013609015 | 1 | ACAv6b | 819 | 0.063937826 | 0 | 0 | 0 | 0 | 4 mo | HOM |
| 1013608106 | 2 | ACAv6b | 819 | 0.063937826 | 0 | 0 | 0 | 0 | 4 mo | HOM |
| 1011937543 | 1 | ACAv6b | 819 | 0.063937826 | 2.65E-05 | 1 | 0.000414863 | 15.64019386 | 8 mo | HOM |
| 1010419538 | 2 | ACAv6b | 819 | 0.063937826 | 0 | 0 | 0 | 0 | 4 mo | WILD |
| 1010419187 | 1 | ACAv6b | 819 | 0.063937826 | 1.16E-05 | 0 | 0.000180798 | 0 | 8 mo | HOM |
| 1013978366 | 1 | ACAv6b | 819 | 0.063937826 | 0 | 0 | 0 | 0 | 2 mo | HOM |
| 1014416155 | 2 | ACAv6b | 819 | 0.063937826 | 0 | 0 | 0 | 0 | 2 mo | HOM |
| 1013978065 | 2 | ACAv6b | 819 | 0.063937826 | 0 | 0 | 0 | 0 | 2 mo | HOM |
| 1009934102 | 2 | ACAv6b | 819 | 0.063937826 | 0 | 0 | 0 | 0 | 4 mo | WILD |
| 1009322937 | 2 | ACAv6b | 819 | 0.063937826 | 0 | 0 | 0 | 0 | 8 mo | WILD |
| 1011080637 | 2 | ACAv6b | 819 | 0.063937826 | 1.97E-06 | 0 | 3.09E-05 | 0 | 8 mo | HOM |
| 1013609713 | 2 | ACAv6b | 819 | 0.063937826 | 0 | 0 | 0 | 0 | 2 mo | WILD |
| 1010418842 | 1 | ACAv6b | 819 | 0.063937826 | 2.38E-05 | 1 | 0.000371484 | 15.64019386 | 8 mo | HOM |
| 1013288213 | 2 | ACAv6b | 819 | 0.063937826 | 0 | 0 | 0 | 0 | 4 mo | HOM |
| 1008976060 | 2 | ACAv6b | 819 | 0.063937826 | 1.75E-06 | 0 | 2.74E-05 | 0 | 8 mo | WILD |
| 1013446243 | 1 | ACAv6b | 819 | 0.063937826 | 0 | 0 | 0 | 0 | 4 mo | HOM |
| 1013288951 | 2 | ACAv6b | 819 | 0.063937826 | 1.45E-07 | 0 | 2.27E-06 | 0 | 4 mo | HOM |
| 1013978665 | 1 | ICd | 820 | 1.363813942 | 1.86E-05 | 1 | 1.36E-05 | 0.733237848 | 2 mo | HOM |
| 1013445609 | 1 | ICd | 820 | 1.363813942 | 0 | 0 | 0 | 0 | 4 mo | HOM |
| 1013609015 | 1 | ICd | 820 | 1.363813942 | 1.63E-05 | 2 | 1.19E-05 | 1.466475696 | 4 mo | HOM |
| 1013608106 | 2 | ICd | 820 | 1.363813942 | 0 | 0 | 0 | 0 | 4 mo | HOM |
| 1011937543 | 1 | ICd | 820 | 1.363813942 | 0.005271324 | 17 | 0.003865134 | 12.46504341 | 8 mo | HOM |
| 1010419538 | 2 | ICd | 820 | 1.363813942 | 0 | 0 | 0 | 0 | 4 mo | WILD |
| 1010419187 | 1 | ICd | 820 | 1.363813942 | 0.001028094 | 22 | 0.000753838 | 16.13123265 | 8 mo | HOM |
| 1013978366 | 1 | ICd | 820 | 1.363813942 | 2.24E-05 | 1 | 1.64E-05 | 0.733237848 | 2 mo | HOM |
| 1014416155 | 2 | ICd | 820 | 1.363813942 | 0 | 0 | 0 | 0 | 2 mo | HOM |

| | | | | | | | | | | |
|---|---|---|---|---|---|---|---|---|---|---|
| **1013978065** | 2 | ICd | 820 | 1.363813942 | 0 | 0 | 0 | 0 | 2 mo | HOM |
| **1009934102** | 2 | ICd | 820 | 1.363813942 | 0 | 0 | 0 | 0 | 4 mo | WILD |
| **1009322937** | 2 | ICd | 820 | 1.363813942 | 2.55E-07 | 0 | 1.87E-07 | 0 | 8 mo | WILD |
| **1011080637** | 2 | ICd | 820 | 1.363813942 | 0.000391898 | 13 | 0.000287355 | 9.532092022 | 8 mo | HOM |
| **1013609713** | 2 | ICd | 820 | 1.363813942 | 0 | 0 | 0 | 0 | 2 mo | WILD |
| **1010418842** | 1 | ICd | 820 | 1.363813942 | 0.001176781 | 34 | 0.00086286 | 24.93008683 | 8 mo | HOM |
| **1013288213** | 2 | ICd | 820 | 1.363813942 | 0 | 0 | 0 | 0 | 4 mo | HOM |
| **1008976060** | 2 | ICd | 820 | 1.363813942 | 6.2E-05 | 4 | 4.55E-05 | 2.932951391 | 8 mo | WILD |
| **1013446243** | 1 | ICd | 820 | 1.363813942 | 0 | 0 | 0 | 0 | 4 mo | HOM |
| **1013288951** | 2 | ICd | 820 | 1.363813942 | 2.2E-05 | 1 | 1.61E-05 | 0.733237848 | 4 mo | HOM |
| **1013978665** | 1 | VISp2/3 | 821 | 2.059461062 | 5.32E-08 | 0 | 2.58E-08 | 0 | 2 mo | HOM |
| **1013445609** | 1 | VISp2/3 | 821 | 2.059461062 | 0.000221021 | 13 | 0.00010732 | 6.312331045 | 4 mo | HOM |
| **1013609015** | 1 | VISp2/3 | 821 | 2.059461062 | 0.000811359 | 16 | 0.000393967 | 7.769022825 | 4 mo | HOM |
| **1013608106** | 2 | VISp2/3 | 821 | 2.059461062 | 0 | 0 | 0 | 0 | 4 mo | HOM |
| **1011937543** | 1 | VISp2/3 | 821 | 2.059461062 | 0.003482683 | 54 | 0.001691065 | 26.22045203 | 8 mo | HOM |
| **1010419538** | 2 | VISp2/3 | 821 | 2.059461062 | 7.53E-06 | 0 | 3.66E-06 | 0 | 4 mo | WILD |
| **1010419187** | 1 | VISp2/3 | 821 | 2.059461062 | 0.003861092 | 72 | 0.001874807 | 34.96060271 | 8 mo | HOM |
| **1013978366** | 1 | VISp2/3 | 821 | 2.059461062 | 7.21E-06 | 0 | 3.5E-06 | 0 | 2 mo | HOM |
| **1014416155** | 2 | VISp2/3 | 821 | 2.059461062 | 0 | 0 | 0 | 0 | 2 mo | HOM |
| **1013978065** | 2 | VISp2/3 | 821 | 2.059461062 | 0 | 0 | 0 | 0 | 2 mo | HOM |
| **1009934102** | 2 | VISp2/3 | 821 | 2.059461062 | 2.09E-05 | 1 | 1.02E-05 | 0.485563927 | 4 mo | WILD |
| **1009322937** | 2 | VISp2/3 | 821 | 2.059461062 | 8.64E-08 | 0 | 4.19E-08 | 0 | 8 mo | WILD |
| **1011080637** | 2 | VISp2/3 | 821 | 2.059461062 | 0.002032048 | 63 | 0.000986689 | 30.59052737 | 8 mo | HOM |
| **1013609713** | 2 | VISp2/3 | 821 | 2.059461062 | 4.6E-07 | 0 | 2.23E-07 | 0 | 2 mo | WILD |
| **1010418842** | 1 | VISp2/3 | 821 | 2.059461062 | 0.002794174 | 80 | 0.00135675 | 38.84511412 | 8 mo | HOM |
| **1013288213** | 2 | VISp2/3 | 821 | 2.059461062 | 0.000305458 | 23 | 0.000148319 | 11.16797031 | 4 mo | HOM |
| **1008976060** | 2 | VISp2/3 | 821 | 2.059461062 | 4.77E-06 | 0 | 2.32E-06 | 0 | 8 mo | WILD |
| **1013446243** | 1 | VISp2/3 | 821 | 2.059461062 | 0.000403834 | 19 | 0.000196087 | 9.225714604 | 4 mo | HOM |
| **1013288951** | 2 | VISp2/3 | 821 | 2.059461062 | 0.000763453 | 29 | 0.000370705 | 14.08135387 | 4 mo | HOM |
| **1013978665** | 1 | RHP | 822 | 19.06389527 | 0.000107785 | 5 | 5.65E-06 | 0.262275885 | 2 mo | HOM |
| **1013445609** | 1 | RHP | 822 | 19.06389527 | 0.000671692 | 54 | 3.52E-05 | 2.832579556 | 4 mo | HOM |
| **1013609015** | 1 | RHP | 822 | 19.06389527 | 0.004682583 | 116 | 0.000245626 | 6.084800527 | 4 mo | HOM |
| **1013608106** | 2 | RHP | 822 | 19.06389527 | 0 | 0 | 0 | 0 | 4 mo | HOM |
| **1011937543** | 1 | RHP | 822 | 19.06389527 | 0.036605843 | 389 | 0.001920166 | 20.40506384 | 8 mo | HOM |
| **1010419538** | 2 | RHP | 822 | 19.06389527 | 0.000437013 | 33 | 2.29E-05 | 1.73102084 | 4 mo | WILD |
| **1010419187** | 1 | RHP | 822 | 19.06389527 | 0.039343597 | 792 | 0.002063775 | 41.54450015 | 8 mo | HOM |
| **1013978366** | 1 | RHP | 822 | 19.06389527 | 0.001432968 | 40 | 7.52E-05 | 2.098207078 | 2 mo | HOM |
| **1014416155** | 2 | RHP | 822 | 19.06389527 | 0 | 0 | 0 | 0 | 2 mo | HOM |
| **1013978065** | 2 | RHP | 822 | 19.06389527 | 2.65E-05 | 2 | 1.39E-06 | 0.104910354 | 2 mo | HOM |
| **1009934102** | 2 | RHP | 822 | 19.06389527 | 0.001547466 | 54 | 8.12E-05 | 2.832579556 | 4 mo | WILD |
| **1009322937** | 2 | RHP | 822 | 19.06389527 | 0.000369104 | 19 | 1.94E-05 | 0.996648362 | 8 mo | WILD |
| **1011080637** | 2 | RHP | 822 | 19.06389527 | 0.020148605 | 589 | 0.001056899 | 30.89609923 | 8 mo | HOM |
| **1013609713** | 2 | RHP | 822 | 19.06389527 | 0.000410924 | 20 | 2.16E-05 | 1.049103539 | 2 mo | WILD |
| **1010418842** | 1 | RHP | 822 | 19.06389527 | 0.025423269 | 691 | 0.001333582 | 36.24652728 | 8 mo | HOM |
| **1013288213** | 2 | RHP | 822 | 19.06389527 | 0.000969291 | 66 | 5.08E-05 | 3.462041679 | 4 mo | HOM |
| **1008976060** | 2 | RHP | 822 | 19.06389527 | 0.000661842 | 49 | 3.47E-05 | 2.570303671 | 8 mo | WILD |
| **1013446243** | 1 | RHP | 822 | 19.06389527 | 0.001695117 | 92 | 8.89E-05 | 4.82587628 | 4 mo | HOM |
| **1013288951** | 2 | RHP | 822 | 19.06389527 | 0.008645536 | 135 | 0.000453503 | 7.081448889 | 4 mo | HOM |
| **1013978665** | 1 | mfsbshy | 824 | 0.845693151 | 4.27E-05 | 0 | 5.05E-05 | 0 | 2 mo | HOM |
| **1013445609** | 1 | mfsbshy | 824 | 0.845693151 | 1.49E-05 | 2 | 1.76E-05 | 2.364923965 | 4 mo | HOM |
| **1013609015** | 1 | mfsbshy | 824 | 0.845693151 | 7.85E-05 | 1 | 9.29E-05 | 1.182461982 | 4 mo | HOM |
| **1013608106** | 2 | mfsbshy | 824 | 0.845693151 | 0 | 0 | 0 | 0 | 4 mo | HOM |
| **1011937543** | 1 | mfsbshy | 824 | 0.845693151 | 0.000364273 | 7 | 0.000430738 | 8.277233877 | 8 mo | HOM |
| **1010419538** | 2 | mfsbshy | 824 | 0.845693151 | 3.14E-05 | 2 | 3.72E-05 | 2.364923965 | 4 mo | WILD |
| **1010419187** | 1 | mfsbshy | 824 | 0.845693151 | 0.000527367 | 10 | 0.000623592 | 11.82461982 | 8 mo | HOM |
| **1013978366** | 1 | mfsbshy | 824 | 0.845693151 | 4.12E-05 | 2 | 4.87E-05 | 2.364923965 | 2 mo | HOM |
| **1014416155** | 2 | mfsbshy | 824 | 0.845693151 | 0 | 0 | 0 | 0 | 2 mo | HOM |
| **1013978065** | 2 | mfsbshy | 824 | 0.845693151 | 0 | 0 | 0 | 0 | 2 mo | HOM |
| **1009934102** | 2 | mfsbshy | 824 | 0.845693151 | 0 | 0 | 0 | 0 | 4 mo | WILD |

| | | | | | | | | | | |
|---|---|---|---|---|---|---|---|---|---|---|
| 1009322937 | 2 | mfsbshy | 824 | 0.845693151 | 3.15E-08 | 0 | 3.72E-08 | 0 | 8 mo | WILD |
| 1011080637 | 2 | mfsbshy | 824 | 0.845693151 | 0.000339203 | 5 | 0.000401094 | 5.912309912 | 8 mo | HOM |
| 1013609713 | 2 | mfsbshy | 824 | 0.845693151 | 0 | 0 | 0 | 0 | 2 mo | WILD |
| 1010418842 | 1 | mfsbshy | 824 | 0.845693151 | 0.000347789 | 7 | 0.000411247 | 8.277233877 | 8 mo | HOM |
| 1013288213 | 2 | mfsbshy | 824 | 0.845693151 | 1.75E-11 | 0 | 2.07E-11 | 0 | 4 mo | HOM |
| 1008976060 | 2 | mfsbshy | 824 | 0.845693151 | 9.82E-05 | 1 | 0.000116168 | 1.182461982 | 8 mo | WILD |
| 1013446243 | 1 | mfsbshy | 824 | 0.845693151 | 4.11E-05 | 2 | 4.87E-05 | 2.364923965 | 4 mo | HOM |
| 1013288951 | 2 | mfsbshy | 824 | 0.845693151 | 0.000542161 | 3 | 0.000641085 | 3.547385947 | 4 mo | HOM |
| 1013978665 | 1 | PALm | 826 | 1.556618476 | 3.78E-05 | 0 | 2.43E-05 | 0 | 2 mo | HOM |
| 1013445609 | 1 | PALm | 826 | 1.556618476 | 2.46E-05 | 1 | 1.58E-05 | 0.642418175 | 4 mo | HOM |
| 1013609015 | 1 | PALm | 826 | 1.556618476 | 0.000832678 | 7 | 0.000534927 | 4.496927223 | 4 mo | HOM |
| 1013608106 | 2 | PALm | 826 | 1.556618476 | 0 | 0 | 0 | 0 | 4 mo | HOM |
| 1011937543 | 1 | PALm | 826 | 1.556618476 | 0.000697397 | 23 | 0.00044802 | 14.77561802 | 8 mo | HOM |
| 1010419538 | 2 | PALm | 826 | 1.556618476 | 5.13E-06 | 0 | 3.29E-06 | 0 | 4 mo | WILD |
| 1010419187 | 1 | PALm | 826 | 1.556618476 | 0.000648873 | 16 | 0.000416848 | 10.27869079 | 8 mo | HOM |
| 1013978366 | 1 | PALm | 826 | 1.556618476 | 3.75E-06 | 1 | 2.41E-06 | 0.642418175 | 2 mo | HOM |
| 1014416155 | 2 | PALm | 826 | 1.556618476 | 0 | 0 | 0 | 0 | 2 mo | HOM |
| 1013978065 | 2 | PALm | 826 | 1.556618476 | 3.07E-09 | 0 | 1.97E-09 | 0 | 2 mo | HOM |
| 1009934102 | 2 | PALm | 826 | 1.556618476 | 1.96E-05 | 2 | 1.26E-05 | 1.284836349 | 4 mo | WILD |
| 1009322937 | 2 | PALm | 826 | 1.556618476 | 0.00015496 | 8 | 9.95E-05 | 5.139345397 | 8 mo | WILD |
| 1011080637 | 2 | PALm | 826 | 1.556618476 | 0.000899287 | 16 | 0.000577718 | 10.27869079 | 8 mo | HOM |
| 1013609713 | 2 | PALm | 826 | 1.556618476 | 1.86E-06 | 0 | 1.19E-06 | 0 | 2 mo | WILD |
| 1010418842 | 1 | PALm | 826 | 1.556618476 | 0.001643486 | 24 | 0.001055806 | 15.41803619 | 8 mo | HOM |
| 1013288213 | 2 | PALm | 826 | 1.556618476 | 2.11E-05 | 0 | 1.35E-05 | 0 | 4 mo | HOM |
| 1008976060 | 2 | PALm | 826 | 1.556618476 | 0.000548489 | 3 | 0.000352359 | 1.927254524 | 8 mo | WILD |
| 1013446243 | 1 | PALm | 826 | 1.556618476 | 0.000497113 | 9 | 0.000319355 | 5.781763572 | 4 mo | HOM |
| 1013288951 | 2 | PALm | 826 | 1.556618476 | 0.000669224 | 7 | 0.000429921 | 4.496927223 | 4 mo | HOM |
| 1013978665 | 1 | ILA5 | 827 | 0.346371945 | 0 | 0 | 0 | 0 | 2 mo | HOM |
| 1013445609 | 1 | ILA5 | 827 | 0.346371945 | 3.93E-05 | 1 | 0.000113452 | 2.887069854 | 4 mo | HOM |
| 1013609015 | 1 | ILA5 | 827 | 0.346371945 | 0.000131887 | 5 | 0.000380767 | 14.43534927 | 4 mo | HOM |
| 1013608106 | 2 | ILA5 | 827 | 0.346371945 | 0 | 0 | 0 | 0 | 4 mo | HOM |
| 1011937543 | 1 | ILA5 | 827 | 0.346371945 | 0.000224949 | 6 | 0.000649443 | 17.32241912 | 8 mo | HOM |
| 1010419538 | 2 | ILA5 | 827 | 0.346371945 | 0 | 0 | 0 | 0 | 4 mo | WILD |
| 1010419187 | 1 | ILA5 | 827 | 0.346371945 | 0.000808444 | 14 | 0.002334035 | 40.41897795 | 8 mo | HOM |
| 1013978366 | 1 | ILA5 | 827 | 0.346371945 | 0 | 0 | 0 | 0 | 2 mo | HOM |
| 1014416155 | 2 | ILA5 | 827 | 0.346371945 | 0 | 0 | 0 | 0 | 2 mo | HOM |
| 1013978065 | 2 | ILA5 | 827 | 0.346371945 | 0 | 0 | 0 | 0 | 2 mo | HOM |
| 1009934102 | 2 | ILA5 | 827 | 0.346371945 | 0 | 0 | 0 | 0 | 4 mo | WILD |
| 1009322937 | 2 | ILA5 | 827 | 0.346371945 | 0 | 0 | 0 | 0 | 8 mo | WILD |
| 1011080637 | 2 | ILA5 | 827 | 0.346371945 | 0.000585838 | 17 | 0.001691354 | 49.08018751 | 8 mo | HOM |
| 1013609713 | 2 | ILA5 | 827 | 0.346371945 | 0 | 0 | 0 | 0 | 2 mo | WILD |
| 1010418842 | 1 | ILA5 | 827 | 0.346371945 | 0.000313895 | 8 | 0.000906237 | 23.09655883 | 8 mo | HOM |
| 1013288213 | 2 | ILA5 | 827 | 0.346371945 | 3.42E-05 | 3 | 9.87E-05 | 8.661209561 | 4 mo | HOM |
| 1008976060 | 2 | ILA5 | 827 | 0.346371945 | 0 | 0 | 0 | 0 | 8 mo | WILD |
| 1013446243 | 1 | ILA5 | 827 | 0.346371945 | 4.49E-05 | 1 | 0.000129753 | 2.887069854 | 4 mo | HOM |
| 1013288951 | 2 | ILA5 | 827 | 0.346371945 | 8.03E-05 | 2 | 0.000231885 | 5.774139707 | 4 mo | HOM |
| 1013978665 | 1 | ICe | 828 | 2.05549465 | 4.95E-05 | 5 | 2.41E-05 | 2.432504507 | 2 mo | HOM |
| 1013445609 | 1 | ICe | 828 | 2.05549465 | 0 | 0 | 0 | 0 | 4 mo | HOM |
| 1013609015 | 1 | ICe | 828 | 2.05549465 | 0.000104086 | 3 | 5.06E-05 | 1.459502704 | 4 mo | HOM |
| 1013608106 | 2 | ICe | 828 | 2.05549465 | 0 | 0 | 0 | 0 | 4 mo | HOM |
| 1011937543 | 1 | ICe | 828 | 2.05549465 | 0.003344883 | 24 | 0.001627288 | 11.67602164 | 8 mo | HOM |
| 1010419538 | 2 | ICe | 828 | 2.05549465 | 0 | 0 | 0 | 0 | 4 mo | WILD |
| 1010419187 | 1 | ICe | 828 | 2.05549465 | 0.000512363 | 14 | 0.000249265 | 6.811012621 | 8 mo | HOM |
| 1013978366 | 1 | ICe | 828 | 2.05549465 | 9.72E-09 | 0 | 4.73E-09 | 0 | 2 mo | HOM |
| 1014416155 | 2 | ICe | 828 | 2.05549465 | 0 | 0 | 0 | 0 | 2 mo | HOM |
| 1013978065 | 2 | ICe | 828 | 2.05549465 | 0 | 0 | 0 | 0 | 2 mo | HOM |
| 1009934102 | 2 | ICe | 828 | 2.05549465 | 0 | 0 | 0 | 0 | 4 mo | WILD |
| 1009322937 | 2 | ICe | 828 | 2.05549465 | 1.22E-10 | 0 | 5.93E-11 | 0 | 8 mo | WILD |
| 1011080637 | 2 | ICe | 828 | 2.05549465 | 0.000109936 | 6 | 5.35E-05 | 2.919005409 | 8 mo | HOM |

| | | | | | | | | | | |
|---|---|---|---|---|---|---|---|---|---|---|
| 1013609713 | 2 | ICe | 828 | 2.05549465 | 6.94E-07 | 0 | 3.38E-07 | 0 | 2 mo | WILD |
| 1010418842 | 1 | ICe | 828 | 2.05549465 | 0.000672558 | 21 | 0.0003272 | 10.21651893 | 8 mo | HOM |
| 1013288213 | 2 | ICe | 828 | 2.05549465 | 0 | 0 | 0 | 0 | 4 mo | HOM |
| 1008976060 | 2 | ICe | 828 | 2.05549465 | 0.000136103 | 10 | 6.62E-05 | 4.865009015 | 8 mo | WILD |
| 1013446243 | 1 | ICe | 828 | 2.05549465 | 9.31E-06 | 1 | 4.53E-06 | 0.486500901 | 4 mo | HOM |
| 1013288951 | 2 | ICe | 828 | 2.05549465 | 4.73E-06 | 0 | 2.3E-06 | 0 | 4 mo | HOM |
| 1013978665 | 1 | DMH | 830 | 0.387428457 | 0 | 0 | 0 | 0 | 2 mo | HOM |
| 1013445609 | 1 | DMH | 830 | 0.387428457 | 0 | 0 | 0 | 0 | 4 mo | HOM |
| 1013609015 | 1 | DMH | 830 | 0.387428457 | 0 | 0 | 0 | 0 | 4 mo | HOM |
| 1013608106 | 2 | DMH | 830 | 0.387428457 | 0 | 0 | 0 | 0 | 4 mo | HOM |
| 1011937543 | 1 | DMH | 830 | 0.387428457 | 0.000342314 | 2 | 0.000883553 | 5.162243415 | 8 mo | HOM |
| 1010419538 | 2 | DMH | 830 | 0.387428457 | 0 | 0 | 0 | 0 | 4 mo | WILD |
| 1010419187 | 1 | DMH | 830 | 0.387428457 | 3.12E-05 | 2 | 8.05E-05 | 5.162243415 | 8 mo | HOM |
| 1013978366 | 1 | DMH | 830 | 0.387428457 | 0 | 0 | 0 | 0 | 2 mo | HOM |
| 1014416155 | 2 | DMH | 830 | 0.387428457 | 0 | 0 | 0 | 0 | 2 mo | HOM |
| 1013978065 | 2 | DMH | 830 | 0.387428457 | 0 | 0 | 0 | 0 | 2 mo | HOM |
| 1009934102 | 2 | DMH | 830 | 0.387428457 | 0 | 0 | 0 | 0 | 4 mo | WILD |
| 1009322937 | 2 | DMH | 830 | 0.387428457 | 0 | 0 | 0 | 0 | 8 mo | WILD |
| 1011080637 | 2 | DMH | 830 | 0.387428457 | 5.52E-06 | 1 | 1.42E-05 | 2.581121708 | 8 mo | HOM |
| 1013609713 | 2 | DMH | 830 | 0.387428457 | 0 | 0 | 0 | 0 | 2 mo | WILD |
| 1010418842 | 1 | DMH | 830 | 0.387428457 | 0.000128247 | 4 | 0.00033102 | 10.32448683 | 8 mo | HOM |
| 1013288213 | 2 | DMH | 830 | 0.387428457 | 0 | 0 | 0 | 0 | 4 mo | HOM |
| 1008976060 | 2 | DMH | 830 | 0.387428457 | 1.99E-08 | 0 | 5.13E-08 | 0 | 8 mo | WILD |
| 1013446243 | 1 | DMH | 830 | 0.387428457 | 0 | 0 | 0 | 0 | 4 mo | HOM |
| 1013288951 | 2 | DMH | 830 | 0.387428457 | 0 | 0 | 0 | 0 | 4 mo | HOM |
| 1013978665 | 1 | Ald6b | 831 | 0.049741321 | 0 | 0 | 0 | 0 | 2 mo | HOM |
| 1013445609 | 1 | Ald6b | 831 | 0.049741321 | 5.59E-06 | 0 | 0.00011233 | 0 | 4 mo | HOM |
| 1013609015 | 1 | Ald6b | 831 | 0.049741321 | 5.42E-09 | 0 | 1.09E-07 | 0 | 4 mo | HOM |
| 1013608106 | 2 | Ald6b | 831 | 0.049741321 | 0 | 0 | 0 | 0 | 4 mo | HOM |
| 1011937543 | 1 | Ald6b | 831 | 0.049741321 | 9.23E-05 | 3 | 0.001855945 | 60.31202911 | 8 mo | HOM |
| 1010419538 | 2 | Ald6b | 831 | 0.049741321 | 0 | 0 | 0 | 0 | 4 mo | WILD |
| 1010419187 | 1 | Ald6b | 831 | 0.049741321 | 3.4E-05 | 2 | 0.000683406 | 40.20801941 | 8 mo | HOM |
| 1013978366 | 1 | Ald6b | 831 | 0.049741321 | 0 | 0 | 0 | 0 | 2 mo | HOM |
| 1014416155 | 2 | Ald6b | 831 | 0.049741321 | 0 | 0 | 0 | 0 | 2 mo | HOM |
| 1013978065 | 2 | Ald6b | 831 | 0.049741321 | 0 | 0 | 0 | 0 | 2 mo | HOM |
| 1009934102 | 2 | Ald6b | 831 | 0.049741321 | 0 | 0 | 0 | 0 | 4 mo | WILD |
| 1009322937 | 2 | Ald6b | 831 | 0.049741321 | 3.6E-06 | 1 | 7.25E-05 | 20.1040097 | 8 mo | WILD |
| 1011080637 | 2 | Ald6b | 831 | 0.049741321 | 6.96E-06 | 0 | 0.000139906 | 0 | 8 mo | HOM |
| 1013609713 | 2 | Ald6b | 831 | 0.049741321 | 0 | 0 | 0 | 0 | 2 mo | WILD |
| 1010418842 | 1 | Ald6b | 831 | 0.049741321 | 2.08E-05 | 0 | 0.000418842 | 0 | 8 mo | HOM |
| 1013288213 | 2 | Ald6b | 831 | 0.049741321 | 0 | 0 | 0 | 0 | 4 mo | HOM |
| 1008976060 | 2 | Ald6b | 831 | 0.049741321 | 3.38E-06 | 0 | 6.8E-05 | 0 | 8 mo | WILD |
| 1013446243 | 1 | Ald6b | 831 | 0.049741321 | 7.9E-06 | 0 | 0.000158858 | 0 | 4 mo | HOM |
| 1013288951 | 2 | Ald6b | 831 | 0.049741321 | 0 | 0 | 0 | 0 | 4 mo | HOM |
| 1013978665 | 1 | IIIn | 832 | 0.215710564 | 0 | 0 | 0 | 0 | 2 mo | HOM |
| 1013445609 | 1 | IIIn | 832 | 0.215710564 | 0 | 0 | 0 | 0 | 4 mo | HOM |
| 1013609015 | 1 | IIIn | 832 | 0.215710564 | 0 | 0 | 0 | 0 | 4 mo | HOM |
| 1013608106 | 2 | IIIn | 832 | 0.215710564 | 0 | 0 | 0 | 0 | 4 mo | HOM |
| 1011937543 | 1 | IIIn | 832 | 0.215710564 | 1.55E-05 | 0 | 7.2E-05 | 0 | 8 mo | HOM |
| 1010419538 | 2 | IIIn | 832 | 0.215710564 | 0 | 0 | 0 | 0 | 4 mo | WILD |
| 1010419187 | 1 | IIIn | 832 | 0.215710564 | 0 | 0 | 0 | 0 | 8 mo | HOM |
| 1013978366 | 1 | IIIn | 832 | 0.215710564 | 0 | 0 | 0 | 0 | 2 mo | HOM |
| 1014416155 | 2 | IIIn | 832 | 0.215710564 | 0 | 0 | 0 | 0 | 2 mo | HOM |
| 1013978065 | 2 | IIIn | 832 | 0.215710564 | 0 | 0 | 0 | 0 | 2 mo | HOM |
| 1009934102 | 2 | IIIn | 832 | 0.215710564 | 0 | 0 | 0 | 0 | 4 mo | WILD |
| 1009322937 | 2 | IIIn | 832 | 0.215710564 | 0 | 0 | 0 | 0 | 8 mo | WILD |
| 1011080637 | 2 | IIIn | 832 | 0.215710564 | 0 | 0 | 0 | 0 | 8 mo | HOM |
| 1013609713 | 2 | IIIn | 832 | 0.215710564 | 0 | 0 | 0 | 0 | 2 mo | WILD |
| 1010418842 | 1 | IIIn | 832 | 0.215710564 | 1.02E-05 | 0 | 4.71E-05 | 0 | 8 mo | HOM |

| | | | | | | | | | | |
|---|---|---|---|---|---|---|---|---|---|---|
| **1013288213** | 2 | IIIn | 832 | 0.215710564 | 3.01E-09 | 0 | 1.4E-08 | 0 | 4 mo | HOM |
| **1008976060** | 2 | IIIn | 832 | 0.215710564 | 4.85E-08 | 0 | 2.25E-07 | 0 | 8 mo | WILD |
| **1013446243** | 1 | IIIn | 832 | 0.215710564 | 0 | 0 | 0 | 0 | 4 mo | HOM |
| **1013288951** | 2 | IIIn | 832 | 0.215710564 | 0 | 0 | 0 | 0 | 4 mo | HOM |
| **1013978665** | 1 | SCzo | 834 | 0.372160527 | 3.65E-06 | 0 | 9.81E-06 | 0 | 2 mo | HOM |
| **1013445609** | 1 | SCzo | 834 | 0.372160527 | 0 | 0 | 0 | 0 | 4 mo | HOM |
| **1013609015** | 1 | SCzo | 834 | 0.372160527 | 2.1E-05 | 1 | 5.65E-05 | 2.687012531 | 4 mo | HOM |
| **1013608106** | 2 | SCzo | 834 | 0.372160527 | 0 | 0 | 0 | 0 | 4 mo | HOM |
| **1011937543** | 1 | SCzo | 834 | 0.372160527 | 0.000755673 | 10 | 0.002030502 | 26.87012531 | 8 mo | HOM |
| **1010419538** | 2 | SCzo | 834 | 0.372160527 | 4.19E-06 | 1 | 1.13E-05 | 2.687012531 | 4 mo | WILD |
| **1010419187** | 1 | SCzo | 834 | 0.372160527 | 0.000310615 | 12 | 0.000834627 | 32.24415037 | 8 mo | HOM |
| **1013978366** | 1 | SCzo | 834 | 0.372160527 | 5.05E-05 | 0 | 0.000135767 | 0 | 2 mo | HOM |
| **1014416155** | 2 | SCzo | 834 | 0.372160527 | 0 | 0 | 0 | 0 | 2 mo | HOM |
| **1013978065** | 2 | SCzo | 834 | 0.372160527 | 0 | 0 | 0 | 0 | 2 mo | HOM |
| **1009934102** | 2 | SCzo | 834 | 0.372160527 | 8.54E-06 | 2 | 2.29E-05 | 5.374025061 | 4 mo | WILD |
| **1009322937** | 2 | SCzo | 834 | 0.372160527 | 2.31E-05 | 1 | 6.21E-05 | 2.687012531 | 8 mo | WILD |
| **1011080637** | 2 | SCzo | 834 | 0.372160527 | 0.000332935 | 17 | 0.000894601 | 45.67921302 | 8 mo | HOM |
| **1013609713** | 2 | SCzo | 834 | 0.372160527 | 1.11E-05 | 2 | 2.99E-05 | 5.374025061 | 2 mo | WILD |
| **1010418842** | 1 | SCzo | 834 | 0.372160527 | 0.000232143 | 18 | 0.00062377 | 48.36622555 | 8 mo | HOM |
| **1013288213** | 2 | SCzo | 834 | 0.372160527 | 5.94E-06 | 0 | 1.6E-05 | 0 | 4 mo | HOM |
| **1008976060** | 2 | SCzo | 834 | 0.372160527 | 9.92E-05 | 3 | 0.00026652 | 8.061037592 | 8 mo | WILD |
| **1013446243** | 1 | SCzo | 834 | 0.372160527 | 2.87E-05 | 1 | 7.72E-05 | 2.687012531 | 4 mo | HOM |
| **1013288951** | 2 | SCzo | 834 | 0.372160527 | 0.000157369 | 12 | 0.000422853 | 32.24415037 | 4 mo | HOM |
| **1013978665** | 1 | PALv | 835 | 3.492731799 | 4.64E-06 | 1 | 1.33E-06 | 0.286308843 | 2 mo | HOM |
| **1013445609** | 1 | PALv | 835 | 3.493892746 | 2.17E-06 | 1 | 6.22E-07 | 0.286213708 | 4 mo | HOM |
| **1013609015** | 1 | PALv | 835 | 3.493892746 | 0.000318601 | 4 | 9.12E-05 | 1.144854834 | 4 mo | HOM |
| **1013608106** | 2 | PALv | 835 | 3.493892746 | 0 | 0 | 0 | 0 | 4 mo | HOM |
| **1011937543** | 1 | PALv | 835 | 3.493892746 | 0.001784014 | 37 | 0.000510609 | 10.58990721 | 8 mo | HOM |
| **1010419538** | 2 | PALv | 835 | 3.493892746 | 0 | 0 | 0 | 0 | 4 mo | WILD |
| **1010419187** | 1 | PALv | 835 | 3.493892746 | 0.001241072 | 67 | 0.000355212 | 19.17631847 | 8 mo | HOM |
| **1013978366** | 1 | PALv | 835 | 3.493892746 | 0 | 0 | 0 | 0 | 2 mo | HOM |
| **1014416155** | 2 | PALv | 835 | 3.493892746 | 0 | 0 | 0 | 0 | 2 mo | HOM |
| **1013978065** | 2 | PALv | 835 | 3.493892746 | 0 | 0 | 0 | 0 | 2 mo | HOM |
| **1009934102** | 2 | PALv | 835 | 3.493892746 | 0 | 0 | 0 | 0 | 4 mo | WILD |
| **1009322937** | 2 | PALv | 835 | 3.493892746 | 0 | 0 | 0 | 0 | 8 mo | WILD |
| **1011080637** | 2 | PALv | 835 | 3.493892746 | 0.001182636 | 57 | 0.000338486 | 16.31418138 | 8 mo | HOM |
| **1013609713** | 2 | PALv | 835 | 3.493892746 | 0 | 0 | 0 | 0 | 2 mo | WILD |
| **1010418842** | 1 | PALv | 835 | 3.493892746 | 0.002066768 | 89 | 0.000591537 | 25.47302006 | 8 mo | HOM |
| **1013288213** | 2 | PALv | 835 | 3.493892746 | 1.24E-05 | 1 | 3.54E-06 | 0.286213708 | 4 mo | HOM |
| **1008976060** | 2 | PALv | 835 | 3.493892746 | 2.06E-05 | 4 | 5.9E-06 | 1.144854834 | 8 mo | WILD |
| **1013446243** | 1 | PALv | 835 | 3.493892746 | 7.75E-05 | 4 | 2.22E-05 | 1.144854834 | 4 mo | HOM |
| **1013288951** | 2 | PALv | 835 | 3.493892746 | 4.51E-06 | 1 | 1.29E-06 | 0.286213708 | 4 mo | HOM |
| **1013978665** | 1 | ECT1 | 836 | 0.266333777 | 0 | 0 | 0 | 0 | 2 mo | HOM |
| **1013445609** | 1 | ECT1 | 836 | 0.266333777 | 9.2E-06 | 1 | 3.45E-05 | 3.754687115 | 4 mo | HOM |
| **1013609015** | 1 | ECT1 | 836 | 0.266333777 | 5.97E-05 | 1 | 0.000224071 | 3.754687115 | 4 mo | HOM |
| **1013608106** | 2 | ECT1 | 836 | 0.266333777 | 0 | 0 | 0 | 0 | 4 mo | HOM |
| **1011937543** | 1 | ECT1 | 836 | 0.266333777 | 0.002550126 | 3 | 0.009574926 | 11.26406134 | 8 mo | HOM |
| **1010419538** | 2 | ECT1 | 836 | 0.266333777 | 1.91E-06 | 1 | 7.18E-06 | 3.754687115 | 4 mo | WILD |
| **1010419187** | 1 | ECT1 | 836 | 0.266333777 | 0.000754491 | 10 | 0.002832879 | 37.54687115 | 8 mo | HOM |
| **1013978366** | 1 | ECT1 | 836 | 0.266333777 | 3.73E-06 | 0 | 1.4E-05 | 0 | 2 mo | HOM |
| **1014416155** | 2 | ECT1 | 836 | 0.266333777 | 0 | 0 | 0 | 0 | 2 mo | HOM |
| **1013978065** | 2 | ECT1 | 836 | 0.266333777 | 0 | 0 | 0 | 0 | 2 mo | HOM |
| **1009934102** | 2 | ECT1 | 836 | 0.266333777 | 7.23E-06 | 0 | 2.72E-05 | 0 | 4 mo | WILD |
| **1009322937** | 2 | ECT1 | 836 | 0.266333777 | 6.22E-07 | 0 | 2.33E-06 | 0 | 8 mo | WILD |
| **1011080637** | 2 | ECT1 | 836 | 0.266333777 | 0.000100634 | 4 | 0.000377851 | 15.01874846 | 8 mo | HOM |
| **1013609713** | 2 | ECT1 | 836 | 0.266333777 | 0 | 0 | 0 | 0 | 2 mo | WILD |
| **1010418842** | 1 | ECT1 | 836 | 0.266333777 | 0.000112942 | 7 | 0.000424061 | 26.2828098 | 8 mo | HOM |
| **1013288213** | 2 | ECT1 | 836 | 0.266333777 | 7.93E-06 | 1 | 2.98E-05 | 3.754687115 | 4 mo | HOM |
| **1008976060** | 2 | ECT1 | 836 | 0.266333777 | 3.62E-05 | 1 | 0.000135774 | 3.754687115 | 8 mo | WILD |

| | | | | | | | | | | |
|---|---|---|---|---|---|---|---|---|---|---|
| **1013446243** | 1 | ECT1 | 836 | 0.266333777 | 9.39E-06 | 1 | 3.53E-05 | 3.754687115 | 4 mo | HOM |
| **1013288951** | 2 | ECT1 | 836 | 0.266333777 | 1.29E-05 | 0 | 4.84E-05 | 0 | 4 mo | HOM |
| **1013978665** | 1 | SSp-n2/3 | 838 | 0.66030832 | 0 | 0 | 0 | 0 | 2 mo | HOM |
| **1013445609** | 1 | SSp-n2/3 | 838 | 0.66030832 | 6.79E-05 | 5 | 0.000102832 | 7.572220202 | 4 mo | HOM |
| **1013609015** | 1 | SSp-n2/3 | 838 | 0.66030832 | 0.000222864 | 6 | 0.000337514 | 9.086664242 | 4 mo | HOM |
| **1013608106** | 2 | SSp-n2/3 | 838 | 0.66030832 | 0 | 0 | 0 | 0 | 4 mo | HOM |
| **1011937543** | 1 | SSp-n2/3 | 838 | 0.66030832 | 0.001569314 | 21 | 0.002376638 | 31.80332485 | 8 mo | HOM |
| **1010419538** | 2 | SSp-n2/3 | 838 | 0.66030832 | 4.54E-09 | 0 | 6.87E-09 | 0 | 4 mo | WILD |
| **1010419187** | 1 | SSp-n2/3 | 838 | 0.66030832 | 0.001090601 | 46 | 0.001651654 | 69.66442585 | 8 mo | HOM |
| **1013978366** | 1 | SSp-n2/3 | 838 | 0.66030832 | 0 | 0 | 0 | 0 | 2 mo | HOM |
| **1014416155** | 2 | SSp-n2/3 | 838 | 0.66030832 | 0 | 0 | 0 | 0 | 2 mo | HOM |
| **1013978065** | 2 | SSp-n2/3 | 838 | 0.66030832 | 0 | 0 | 0 | 0 | 2 mo | HOM |
| **1009934102** | 2 | SSp-n2/3 | 838 | 0.66030832 | 0 | 0 | 0 | 0 | 4 mo | WILD |
| **1009322937** | 2 | SSp-n2/3 | 838 | 0.66030832 | 0 | 0 | 0 | 0 | 8 mo | WILD |
| **1011080637** | 2 | SSp-n2/3 | 838 | 0.66030832 | 0.000967322 | 26 | 0.001464955 | 39.37554505 | 8 mo | HOM |
| **1013609713** | 2 | SSp-n2/3 | 838 | 0.66030832 | 0 | 0 | 0 | 0 | 2 mo | WILD |
| **1010418842** | 1 | SSp-n2/3 | 838 | 0.66030832 | 0.000863632 | 33 | 0.001307922 | 49.97665333 | 8 mo | HOM |
| **1013288213** | 2 | SSp-n2/3 | 838 | 0.66030832 | 5.04E-05 | 5 | 7.63E-05 | 7.572220202 | 4 mo | HOM |
| **1008976060** | 2 | SSp-n2/3 | 838 | 0.66030832 | 7.89E-06 | 1 | 1.2E-05 | 1.51444404 | 8 mo | WILD |
| **1013446243** | 1 | SSp-n2/3 | 838 | 0.66030832 | 0.000140132 | 10 | 0.000212222 | 15.1444404 | 4 mo | HOM |
| **1013288951** | 2 | SSp-n2/3 | 838 | 0.66030832 | 0.000207659 | 9 | 0.000314488 | 13.62999636 | 4 mo | HOM |
| **1013978665** | 1 | DMX | 839 | 0.179459011 | 0 | 0 | 0 | 0 | 2 mo | HOM |
| **1013445609** | 1 | DMX | 839 | 0.179459011 | 0 | 0 | 0 | 0 | 4 mo | HOM |
| **1013609015** | 1 | DMX | 839 | 0.179459011 | 0 | 0 | 0 | 0 | 4 mo | HOM |
| **1013608106** | 2 | DMX | 839 | 0.179459011 | 0 | 0 | 0 | 0 | 4 mo | HOM |
| **1011937543** | 1 | DMX | 839 | 0.179459011 | 0 | 0 | 0 | 0 | 8 mo | HOM |
| **1010419538** | 2 | DMX | 839 | 0.179459011 | 0 | 0 | 0 | 0 | 4 mo | WILD |
| **1010419187** | 1 | DMX | 839 | 0.179459011 | 5.18E-05 | 2 | 0.000288882 | 11.14460615 | 8 mo | HOM |
| **1013978366** | 1 | DMX | 839 | 0.179459011 | 0 | 0 | 0 | 0 | 2 mo | HOM |
| **1014416155** | 2 | DMX | 839 | 0.179459011 | 0 | 0 | 0 | 0 | 2 mo | HOM |
| **1013978065** | 2 | DMX | 839 | 0.179459011 | 0 | 0 | 0 | 0 | 2 mo | HOM |
| **1009934102** | 2 | DMX | 839 | 0.179459011 | 0 | 0 | 0 | 0 | 4 mo | WILD |
| **1009322937** | 2 | DMX | 839 | 0.179459011 | 4.25E-06 | 0 | 2.37E-05 | 0 | 8 mo | WILD |
| **1011080637** | 2 | DMX | 839 | 0.179459011 | 9.95E-06 | 1 | 5.54E-05 | 5.572303075 | 8 mo | HOM |
| **1013609713** | 2 | DMX | 839 | 0.179459011 | 0 | 0 | 0 | 0 | 2 mo | WILD |
| **1010418842** | 1 | DMX | 839 | 0.179459011 | 3.12E-05 | 5 | 0.000173743 | 27.86151538 | 8 mo | HOM |
| **1013288213** | 2 | DMX | 839 | 0.179459011 | 0 | 0 | 0 | 0 | 4 mo | HOM |
| **1008976060** | 2 | DMX | 839 | 0.179459011 | 0 | 0 | 0 | 0 | 8 mo | WILD |
| **1013446243** | 1 | DMX | 839 | 0.179459011 | 0 | 0 | 0 | 0 | 4 mo | HOM |
| **1013288951** | 2 | DMX | 839 | 0.179459011 | 0 | 0 | 0 | 0 | 4 mo | HOM |
| **1013978665** | 1 | In | 840 | 5.254376114 | 0.001774669 | 54 | 0.000337751 | 10.27714781 | 2 mo | HOM |
| **1013445609** | 1 | In | 840 | 5.289803506 | 0.00104648 | 56 | 0.00019783 | 10.58640457 | 4 mo | HOM |
| **1013609015** | 1 | In | 840 | 4.41480055 | 0.002972628 | 108 | 0.000673332 | 24.46316629 | 4 mo | HOM |
| **1013608106** | 2 | In | 840 | 5.388062226 | 0 | 0 | 0 | 0 | 4 mo | HOM |
| **1011937543** | 1 | In | 840 | 5.425300972 | 0.004673506 | 71 | 0.000861428 | 13.08683156 | 8 mo | HOM |
| **1010419538** | 2 | In | 840 | 5.705188345 | 0.000235049 | 20 | 4.12E-05 | 3.505581024 | 4 mo | WILD |
| **1010419187** | 1 | In | 840 | 4.379706985 | 0.00700789 | 65 | 0.001600082 | 14.8411755 | 8 mo | HOM |
| **1013978366** | 1 | In | 840 | 6.034505977 | 0.001373778 | 73 | 0.000227654 | 12.09709631 | 2 mo | HOM |
| **1014416155** | 2 | In | 840 | 5.777268991 | 1.89E-06 | 0 | 3.28E-07 | 0 | 2 mo | HOM |
| **1013978065** | 2 | In | 840 | 6.089171474 | 0.001847231 | 80 | 0.000303363 | 13.13807639 | 2 mo | HOM |
| **1009934102** | 2 | In | 840 | 4.970281484 | 0.000667805 | 50 | 0.00013436 | 10.05979242 | 4 mo | WILD |
| **1009322937** | 2 | In | 840 | 4.194886668 | 9.38E-05 | 13 | 2.24E-05 | 3.099011017 | 8 mo | WILD |
| **1011080637** | 2 | In | 840 | 4.123790726 | 0.008225411 | 70 | 0.001994624 | 16.97467322 | 8 mo | HOM |
| **1013609713** | 2 | In | 840 | 6.153725407 | 9.71E-05 | 15 | 1.58E-05 | 2.437547828 | 2 mo | WILD |
| **1010418842** | 1 | In | 840 | 5.070489228 | 0.00551337 | 128 | 0.001087345 | 25.2441124 | 8 mo | HOM |
| **1013288213** | 2 | In | 840 | 5.662738822 | 0.002179484 | 92 | 0.000384882 | 16.2465554 | 4 mo | HOM |
| **1008976060** | 2 | In | 840 | 5.437702598 | 0.001548377 | 72 | 0.000284748 | 13.24088596 | 8 mo | WILD |
| **1013446243** | 1 | In | 840 | 4.790911065 | 0.001270402 | 84 | 0.000265169 | 17.53319961 | 4 mo | HOM |
| **1013288951** | 2 | In | 840 | 5.180035039 | 0.001732271 | 71 | 0.000334413 | 13.70647099 | 4 mo | HOM |

| | | | | | | | | | | |
|---|---|---|---|---|---|---|---|---|---|---|
| 1013978665 | 1 | tb | 841 | 0.30849469 | 0 | 0 | 0 | 0 | 2 mo | HOM |
| 1013445609 | 1 | tb | 841 | 0.30849469 | 0 | 0 | 0 | 0 | 4 mo | HOM |
| 1013609015 | 1 | tb | 841 | 0.30849469 | 0 | 0 | 0 | 0 | 4 mo | HOM |
| 1013608106 | 2 | tb | 841 | 0.30849469 | 0 | 0 | 0 | 0 | 4 mo | HOM |
| 1011937543 | 1 | tb | 841 | 0.30849469 | 0 | 0 | 0 | 0 | 8 mo | HOM |
| 1010419538 | 2 | tb | 841 | 0.30849469 | 0 | 0 | 0 | 0 | 4 mo | WILD |
| 1010419187 | 1 | tb | 841 | 0.30849469 | 0 | 0 | 0 | 0 | 8 mo | HOM |
| 1013978366 | 1 | tb | 841 | 0.30849469 | 0 | 0 | 0 | 0 | 2 mo | HOM |
| 1014416155 | 2 | tb | 841 | 0.30849469 | 0 | 0 | 0 | 0 | 2 mo | HOM |
| 1013978065 | 2 | tb | 841 | 0.30849469 | 0 | 0 | 0 | 0 | 2 mo | HOM |
| 1009934102 | 2 | tb | 841 | 0.30849469 | 0 | 0 | 0 | 0 | 4 mo | WILD |
| 1009322937 | 2 | tb | 841 | 0.30849469 | 0 | 0 | 0 | 0 | 8 mo | WILD |
| 1011080637 | 2 | tb | 841 | 0.30849469 | 0 | 0 | 0 | 0 | 8 mo | HOM |
| 1013609713 | 2 | tb | 841 | 0.30849469 | 0 | 0 | 0 | 0 | 2 mo | WILD |
| 1010418842 | 1 | tb | 841 | 0.30849469 | 0 | 0 | 0 | 0 | 8 mo | HOM |
| 1013288213 | 2 | tb | 841 | 0.30849469 | 0 | 0 | 0 | 0 | 4 mo | HOM |
| 1008976060 | 2 | tb | 841 | 0.30849469 | 1.04E-05 | 1 | 3.38E-05 | 3.24154688 | 8 mo | WILD |
| 1013446243 | 1 | tb | 841 | 0.30849469 | 0 | 0 | 0 | 0 | 4 mo | HOM |
| 1013288951 | 2 | tb | 841 | 0.30849469 | 0 | 0 | 0 | 0 | 4 mo | HOM |
| 1013978665 | 1 | SCsg | 842 | 1.187668115 | 1.41E-05 | 2 | 1.19E-05 | 1.683972125 | 2 mo | HOM |
| 1013445609 | 1 | SCsg | 842 | 1.187668115 | 0 | 0 | 0 | 0 | 4 mo | HOM |
| 1013609015 | 1 | SCsg | 842 | 1.187668115 | 6.06E-06 | 0 | 5.1E-06 | 0 | 4 mo | HOM |
| 1013608106 | 2 | SCsg | 842 | 1.187668115 | 0 | 0 | 0 | 0 | 4 mo | HOM |
| 1011937543 | 1 | SCsg | 842 | 1.187668115 | 0.002012335 | 38 | 0.001694358 | 31.99547038 | 8 mo | HOM |
| 1010419538 | 2 | SCsg | 842 | 1.187668115 | 2.04E-06 | 0 | 1.72E-06 | 0 | 4 mo | WILD |
| 1010419187 | 1 | SCsg | 842 | 1.187668115 | 0.000243513 | 6 | 0.000205034 | 5.051916376 | 8 mo | HOM |
| 1013978366 | 1 | SCsg | 842 | 1.187668115 | 8.1E-06 | 1 | 6.82E-06 | 0.841986063 | 2 mo | HOM |
| 1014416155 | 2 | SCsg | 842 | 1.187668115 | 0 | 0 | 0 | 0 | 2 mo | HOM |
| 1013978065 | 2 | SCsg | 842 | 1.187668115 | 0 | 0 | 0 | 0 | 2 mo | HOM |
| 1009934102 | 2 | SCsg | 842 | 1.187668115 | 1.18E-05 | 0 | 9.92E-06 | 0 | 4 mo | WILD |
| 1009322937 | 2 | SCsg | 842 | 1.187668115 | 2.12E-05 | 1 | 1.78E-05 | 0.841986063 | 8 mo | WILD |
| 1011080637 | 2 | SCsg | 842 | 1.187668115 | 0.00029443 | 6 | 0.000247906 | 5.051916376 | 8 mo | HOM |
| 1013609713 | 2 | SCsg | 842 | 1.187668115 | 2.25E-06 | 0 | 1.89E-06 | 0 | 2 mo | WILD |
| 1010418842 | 1 | SCsg | 842 | 1.187668115 | 0.00022533 | 5 | 0.000189724 | 4.209930313 | 8 mo | HOM |
| 1013288213 | 2 | SCsg | 842 | 1.187668115 | 1.83E-06 | 0 | 1.54E-06 | 0 | 4 mo | HOM |
| 1008976060 | 2 | SCsg | 842 | 1.187668115 | 3.83E-05 | 3 | 3.22E-05 | 2.525958188 | 8 mo | WILD |
| 1013446243 | 1 | SCsg | 842 | 1.187668115 | 1.06E-05 | 0 | 8.89E-06 | 0 | 4 mo | HOM |
| 1013288951 | 2 | SCsg | 842 | 1.187668115 | 0.000184081 | 6 | 0.000154993 | 5.051916376 | 4 mo | HOM |
| 1013978665 | 1 | PAR | 843 | 0.961072202 | 0 | 0 | 0 | 0 | 2 mo | HOM |
| 1013445609 | 1 | PAR | 843 | 0.961072202 | 1.73E-05 | 1 | 1.8E-05 | 1.040504551 | 4 mo | HOM |
| 1013609015 | 1 | PAR | 843 | 0.961072202 | 0.000638253 | 16 | 0.000664105 | 16.64807282 | 4 mo | HOM |
| 1013608106 | 2 | PAR | 843 | 0.961072202 | 0 | 0 | 0 | 0 | 4 mo | HOM |
| 1011937543 | 1 | PAR | 843 | 0.961072202 | 0.00164344 | 11 | 0.001710007 | 11.44555007 | 8 mo | HOM |
| 1010419538 | 2 | PAR | 843 | 0.961072202 | 0.000132401 | 10 | 0.000137764 | 10.40504551 | 4 mo | WILD |
| 1010419187 | 1 | PAR | 843 | 0.961072202 | 0.004923529 | 25 | 0.005122954 | 26.01261379 | 8 mo | HOM |
| 1013978366 | 1 | PAR | 843 | 0.961072202 | 0.000743504 | 9 | 0.000773619 | 9.364540963 | 2 mo | HOM |
| 1014416155 | 2 | PAR | 843 | 0.961072202 | 0 | 0 | 0 | 0 | 2 mo | HOM |
| 1013978065 | 2 | PAR | 843 | 0.961072202 | 3.51E-06 | 0 | 3.65E-06 | 0 | 2 mo | HOM |
| 1009934102 | 2 | PAR | 843 | 0.961072202 | 0.000269915 | 8 | 0.000280848 | 8.324036411 | 4 mo | WILD |
| 1009322937 | 2 | PAR | 843 | 0.961072202 | 3.11E-05 | 3 | 3.23E-05 | 3.121513654 | 8 mo | WILD |
| 1011080637 | 2 | PAR | 843 | 0.961072202 | 0.00119108 | 42 | 0.001239324 | 43.70119116 | 8 mo | HOM |
| 1013609713 | 2 | PAR | 843 | 0.961072202 | 5.12E-05 | 3 | 5.33E-05 | 3.121513654 | 2 mo | WILD |
| 1010418842 | 1 | PAR | 843 | 0.961072202 | 0.001093854 | 35 | 0.00113816 | 36.4176593 | 8 mo | HOM |
| 1013288213 | 2 | PAR | 843 | 0.961072202 | 8.09E-05 | 8 | 8.42E-05 | 8.324036411 | 4 mo | HOM |
| 1008976060 | 2 | PAR | 843 | 0.961072202 | 2.58E-05 | 5 | 2.68E-05 | 5.202522757 | 8 mo | WILD |
| 1013446243 | 1 | PAR | 843 | 0.961072202 | 0.000186501 | 10 | 0.000194055 | 10.40504551 | 4 mo | HOM |
| 1013288951 | 2 | PAR | 843 | 0.961072202 | 0.002164117 | 16 | 0.002251774 | 16.64807282 | 4 mo | HOM |
| 1013978665 | 1 | MOp6a | 844 | 2.963367904 | 0 | 0 | 0 | 0 | 2 mo | HOM |
| 1013445609 | 1 | MOp6a | 844 | 2.963367904 | 0.00014395 | 15 | 4.86E-05 | 5.061808215 | 4 mo | HOM |

| | | | | | | | | | | |
|---|---|---|---|---|---|---|---|---|---|---|
| **1013609015** | 1 | MOp6a | 844 | 2.963367904 | 0.000363618 | 16 | 0.000122704 | 5.399262096 | 4 mo | HOM |
| **1013608106** | 2 | MOp6a | 844 | 2.963367904 | 0 | 0 | 0 | 0 | 4 mo | HOM |
| **1011937543** | 1 | MOp6a | 844 | 2.963367904 | 0.00383725 | 83 | 0.001294895 | 28.00867212 | 8 mo | HOM |
| **1010419538** | 2 | MOp6a | 844 | 2.963367904 | 0 | 0 | 0 | 0 | 4 mo | WILD |
| **1010419187** | 1 | MOp6a | 844 | 2.963367904 | 0.002777767 | 166 | 0.000937368 | 56.01734424 | 8 mo | HOM |
| **1013978366** | 1 | MOp6a | 844 | 2.963367904 | 0 | 0 | 0 | 0 | 2 mo | HOM |
| **1014416155** | 2 | MOp6a | 844 | 2.963367904 | 0 | 0 | 0 | 0 | 2 mo | HOM |
| **1013978065** | 2 | MOp6a | 844 | 2.963367904 | 0 | 0 | 0 | 0 | 2 mo | HOM |
| **1009934102** | 2 | MOp6a | 844 | 2.963367904 | 0 | 0 | 0 | 0 | 4 mo | WILD |
| **1009322937** | 2 | MOp6a | 844 | 2.963367904 | 0 | 0 | 0 | 0 | 8 mo | WILD |
| **1011080637** | 2 | MOp6a | 844 | 2.963367904 | 0.001893003 | 85 | 0.000638801 | 28.68357988 | 8 mo | HOM |
| **1013609713** | 2 | MOp6a | 844 | 2.963367904 | 0 | 0 | 0 | 0 | 2 mo | WILD |
| **1010418842** | 1 | MOp6a | 844 | 2.963367904 | 0.00286428 | 127 | 0.000966562 | 42.85664289 | 8 mo | HOM |
| **1013288213** | 2 | MOp6a | 844 | 2.963367904 | 5.05E-05 | 4 | 1.7E-05 | 1.349815524 | 4 mo | HOM |
| **1008976060** | 2 | MOp6a | 844 | 2.963367904 | 5.71E-06 | 1 | 1.93E-06 | 0.337453881 | 8 mo | WILD |
| **1013446243** | 1 | MOp6a | 844 | 2.963367904 | 0.000291759 | 21 | 9.85E-05 | 7.086531501 | 4 mo | HOM |
| **1013288951** | 2 | MOp6a | 844 | 2.963367904 | 0.000546605 | 23 | 0.000184454 | 7.761439263 | 4 mo | HOM |
| **1013978665** | 1 | DN | 846 | 0.345108882 | 0 | 0 | 0 | 0 | 2 mo | HOM |
| **1013445609** | 1 | DN | 846 | 0.345108882 | 0 | 0 | 0 | 0 | 4 mo | HOM |
| **1013609015** | 1 | DN | 846 | 0.345108882 | 0 | 0 | 0 | 0 | 4 mo | HOM |
| **1013608106** | 2 | DN | 846 | 0.345108882 | 0 | 0 | 0 | 0 | 4 mo | HOM |
| **1011937543** | 1 | DN | 846 | 0.345108882 | 0.000246743 | 2 | 0.000714972 | 5.795272468 | 8 mo | HOM |
| **1010419538** | 2 | DN | 846 | 0.345108882 | 0 | 0 | 0 | 0 | 4 mo | WILD |
| **1010419187** | 1 | DN | 846 | 0.345108882 | 9.59E-06 | 1 | 2.78E-05 | 2.897636234 | 8 mo | HOM |
| **1013978366** | 1 | DN | 846 | 0.345108882 | 0 | 0 | 0 | 0 | 2 mo | HOM |
| **1014416155** | 2 | DN | 846 | 0.345108882 | 0 | 0 | 0 | 0 | 2 mo | HOM |
| **1013978065** | 2 | DN | 846 | 0.345108882 | 0 | 0 | 0 | 0 | 2 mo | HOM |
| **1009934102** | 2 | DN | 846 | 0.345108882 | 0 | 0 | 0 | 0 | 4 mo | WILD |
| **1009322937** | 2 | DN | 846 | 0.345108882 | 0 | 0 | 0 | 0 | 8 mo | WILD |
| **1011080637** | 2 | DN | 846 | 0.345108882 | 0 | 0 | 0 | 0 | 8 mo | HOM |
| **1013609713** | 2 | DN | 846 | 0.345108882 | 0 | 0 | 0 | 0 | 2 mo | WILD |
| **1010418842** | 1 | DN | 846 | 0.345108882 | 6.38E-08 | 0 | 1.85E-07 | 0 | 8 mo | HOM |
| **1013288213** | 2 | DN | 846 | 0.345108882 | 0 | 0 | 0 | 0 | 4 mo | HOM |
| **1008976060** | 2 | DN | 846 | 0.345108882 | 2.26E-05 | 3 | 6.55E-05 | 8.692908702 | 8 mo | WILD |
| **1013446243** | 1 | DN | 846 | 0.345108882 | 0 | 0 | 0 | 0 | 4 mo | HOM |
| **1013288951** | 2 | DN | 846 | 0.345108882 | 0 | 0 | 0 | 0 | 4 mo | HOM |
| **1013978665** | 1 | AUDp5 | 847 | 0.735469381 | 0 | 0 | 0 | 0 | 2 mo | HOM |
| **1013445609** | 1 | AUDp5 | 847 | 0.735469381 | 4.91E-05 | 5 | 6.67E-05 | 6.798379553 | 4 mo | HOM |
| **1013609015** | 1 | AUDp5 | 847 | 0.735469381 | 0.000227627 | 6 | 0.000309499 | 8.158055463 | 4 mo | HOM |
| **1013608106** | 2 | AUDp5 | 847 | 0.735469381 | 0 | 0 | 0 | 0 | 4 mo | HOM |
| **1011937543** | 1 | AUDp5 | 847 | 0.735469381 | 0.001100927 | 24 | 0.001496904 | 32.63222185 | 8 mo | HOM |
| **1010419538** | 2 | AUDp5 | 847 | 0.735469381 | 0 | 0 | 0 | 0 | 4 mo | WILD |
| **1010419187** | 1 | AUDp5 | 847 | 0.735469381 | 0.00090658 | 32 | 0.001232655 | 43.50962914 | 8 mo | HOM |
| **1013978366** | 1 | AUDp5 | 847 | 0.735469381 | 1.01E-05 | 1 | 1.37E-05 | 1.359675911 | 2 mo | HOM |
| **1014416155** | 2 | AUDp5 | 847 | 0.735469381 | 0 | 0 | 0 | 0 | 2 mo | HOM |
| **1013978065** | 2 | AUDp5 | 847 | 0.735469381 | 0 | 0 | 0 | 0 | 2 mo | HOM |
| **1009934102** | 2 | AUDp5 | 847 | 0.735469381 | 0 | 0 | 0 | 0 | 4 mo | WILD |
| **1009322937** | 2 | AUDp5 | 847 | 0.735469381 | 0 | 0 | 0 | 0 | 8 mo | WILD |
| **1011080637** | 2 | AUDp5 | 847 | 0.735469381 | 0.00061401 | 27 | 0.000834854 | 36.71124958 | 8 mo | HOM |
| **1013609713** | 2 | AUDp5 | 847 | 0.735469381 | 0 | 0 | 0 | 0 | 2 mo | WILD |
| **1010418842** | 1 | AUDp5 | 847 | 0.735469381 | 0.000594442 | 23 | 0.000808248 | 31.27254594 | 8 mo | HOM |
| **1013288213** | 2 | AUDp5 | 847 | 0.735469381 | 2.15E-05 | 2 | 2.93E-05 | 2.719351821 | 4 mo | HOM |
| **1008976060** | 2 | AUDp5 | 847 | 0.735469381 | 0 | 0 | 0 | 0 | 8 mo | WILD |
| **1013446243** | 1 | AUDp5 | 847 | 0.735469381 | 0.000112177 | 6 | 0.000152525 | 8.158055463 | 4 mo | HOM |
| **1013288951** | 2 | AUDp5 | 847 | 0.735469381 | 0.00013301 | 10 | 0.000180851 | 13.59675911 | 4 mo | HOM |
| **1013978665** | 1 | IIn | 848 | 1.346854298 | 7.15E-11 | 0 | 5.31E-11 | 0 | 2 mo | HOM |
| **1013445609** | 1 | IIn | 848 | 1.346854298 | 0 | 0 | 0 | 0 | 4 mo | HOM |
| **1013609015** | 1 | IIn | 848 | 1.346854298 | 2.46E-07 | 0 | 1.83E-07 | 0 | 4 mo | HOM |
| **1013608106** | 2 | IIn | 848 | 1.346854298 | 0 | 0 | 0 | 0 | 4 mo | HOM |

| | | | | | | | | | | |
|---|---|---|---|---|---|---|---|---|---|---|
| **1011937543** | 1 | IIn | 848 | 1.346854298 | 0.004264109 | 17 | 0.003165977 | 12.62200375 | 8 mo | HOM |
| **1010419538** | 2 | IIn | 848 | 1.346854298 | 4.23E-08 | 0 | 3.14E-08 | 0 | 4 mo | WILD |
| **1010419187** | 1 | IIn | 848 | 1.346854298 | 0.000383544 | 7 | 0.00028477 | 5.197295663 | 8 mo | HOM |
| **1013978366** | 1 | IIn | 848 | 1.346854298 | 3.75E-05 | 1 | 2.78E-05 | 0.742470809 | 2 mo | HOM |
| **1014416155** | 2 | IIn | 848 | 1.346854298 | 0 | 0 | 0 | 0 | 2 mo | HOM |
| **1013978065** | 2 | IIn | 848 | 1.346854298 | 0 | 0 | 0 | 0 | 2 mo | HOM |
| **1009934102** | 2 | IIn | 848 | 1.346854298 | 5.69E-06 | 1 | 4.22E-06 | 0.742470809 | 4 mo | WILD |
| **1009322937** | 2 | IIn | 848 | 1.346854298 | 5.22E-07 | 0 | 3.88E-07 | 0 | 8 mo | WILD |
| **1011080637** | 2 | IIn | 848 | 1.346854298 | 2.52E-05 | 1 | 1.87E-05 | 0.742470809 | 8 mo | HOM |
| **1013609713** | 2 | IIn | 848 | 1.346854298 | 0 | 0 | 0 | 0 | 2 mo | WILD |
| **1010418842** | 1 | IIn | 848 | 1.346854298 | 0.000426594 | 7 | 0.000316734 | 5.197295663 | 8 mo | HOM |
| **1013288213** | 2 | IIn | 848 | 1.346854298 | 0 | 0 | 0 | 0 | 4 mo | HOM |
| **1008976060** | 2 | IIn | 848 | 1.346854298 | 6.1E-05 | 2 | 4.53E-05 | 1.484941618 | 8 mo | WILD |
| **1013446243** | 1 | IIn | 848 | 1.346854298 | 7.67E-07 | 0 | 5.7E-07 | 0 | 4 mo | HOM |
| **1013288951** | 2 | IIn | 848 | 1.346854298 | 1.32E-07 | 0 | 9.78E-08 | 0 | 4 mo | HOM |
| **1013978665** | 1 | VISC6b | 849 | 0.053734478 | 0 | 0 | 0 | 0 | 2 mo | HOM |
| **1013445609** | 1 | VISC6b | 849 | 0.053734478 | 0 | 0 | 0 | 0 | 4 mo | HOM |
| **1013609015** | 1 | VISC6b | 849 | 0.053734478 | 1.16E-05 | 0 | 0.000216672 | 0 | 4 mo | HOM |
| **1013608106** | 2 | VISC6b | 849 | 0.053734478 | 0 | 0 | 0 | 0 | 4 mo | HOM |
| **1011937543** | 1 | VISC6b | 849 | 0.053734478 | 6.53E-05 | 2 | 0.001215953 | 37.22005078 | 8 mo | HOM |
| **1010419538** | 2 | VISC6b | 849 | 0.053734478 | 0 | 0 | 0 | 0 | 4 mo | WILD |
| **1010419187** | 1 | VISC6b | 849 | 0.053734478 | 2.98E-05 | 3 | 0.000554153 | 55.83007618 | 8 mo | HOM |
| **1013978366** | 1 | VISC6b | 849 | 0.053734478 | 0 | 0 | 0 | 0 | 2 mo | HOM |
| **1014416155** | 2 | VISC6b | 849 | 0.053734478 | 0 | 0 | 0 | 0 | 2 mo | HOM |
| **1013978065** | 2 | VISC6b | 849 | 0.053734478 | 0 | 0 | 0 | 0 | 2 mo | HOM |
| **1009934102** | 2 | VISC6b | 849 | 0.053734478 | 0 | 0 | 0 | 0 | 4 mo | WILD |
| **1009322937** | 2 | VISC6b | 849 | 0.053734478 | 0 | 0 | 0 | 0 | 8 mo | WILD |
| **1011080637** | 2 | VISC6b | 849 | 0.053734478 | 2.23E-05 | 2 | 0.000415727 | 37.22005078 | 8 mo | HOM |
| **1013609713** | 2 | VISC6b | 849 | 0.053734478 | 0 | 0 | 0 | 0 | 2 mo | WILD |
| **1010418842** | 1 | VISC6b | 849 | 0.053734478 | 2.86E-05 | 3 | 0.000531439 | 55.83007618 | 8 mo | HOM |
| **1013288213** | 2 | VISC6b | 849 | 0.053734478 | 1.42E-06 | 0 | 2.64E-05 | 0 | 4 mo | HOM |
| **1008976060** | 2 | VISC6b | 849 | 0.053734478 | 2.87E-06 | 1 | 5.35E-05 | 18.61002539 | 8 mo | WILD |
| **1013446243** | 1 | VISC6b | 849 | 0.053734478 | 3.55E-06 | 1 | 6.6E-05 | 18.61002539 | 4 mo | HOM |
| **1013288951** | 2 | VISC6b | 849 | 0.053734478 | 1.79E-07 | 0 | 3.33E-06 | 0 | 4 mo | HOM |
| **1013978665** | 1 | uf | 850 | 0.087639185 | 0 | 0 | 0 | 0 | 2 mo | HOM |
| **1013445609** | 1 | uf | 850 | 0.087639185 | 0 | 0 | 0 | 0 | 4 mo | HOM |
| **1013609015** | 1 | uf | 850 | 0.087639185 | 0 | 0 | 0 | 0 | 4 mo | HOM |
| **1013608106** | 2 | uf | 850 | 0.087639185 | 0 | 0 | 0 | 0 | 4 mo | HOM |
| **1011937543** | 1 | uf | 850 | 0.087639185 | 4.66E-08 | 0 | 5.32E-07 | 0 | 8 mo | HOM |
| **1010419538** | 2 | uf | 850 | 0.087639185 | 0 | 0 | 0 | 0 | 4 mo | WILD |
| **1010419187** | 1 | uf | 850 | 0.087639185 | 0 | 0 | 0 | 0 | 8 mo | HOM |
| **1013978366** | 1 | uf | 850 | 0.087639185 | 0 | 0 | 0 | 0 | 2 mo | HOM |
| **1014416155** | 2 | uf | 850 | 0.087639185 | 0 | 0 | 0 | 0 | 2 mo | HOM |
| **1013978065** | 2 | uf | 850 | 0.087639185 | 0 | 0 | 0 | 0 | 2 mo | HOM |
| **1009934102** | 2 | uf | 850 | 0.087639185 | 0 | 0 | 0 | 0 | 4 mo | WILD |
| **1009322937** | 2 | uf | 850 | 0.087639185 | 0 | 0 | 0 | 0 | 8 mo | WILD |
| **1011080637** | 2 | uf | 850 | 0.087639185 | 0 | 0 | 0 | 0 | 8 mo | HOM |
| **1013609713** | 2 | uf | 850 | 0.087639185 | 0 | 0 | 0 | 0 | 2 mo | WILD |
| **1010418842** | 1 | uf | 850 | 0.087639185 | 0 | 0 | 0 | 0 | 8 mo | HOM |
| **1013288213** | 2 | uf | 850 | 0.087639185 | 0 | 0 | 0 | 0 | 4 mo | HOM |
| **1008976060** | 2 | uf | 850 | 0.087639185 | 5.99E-06 | 1 | 6.83E-05 | 11.41042101 | 8 mo | WILD |
| **1013446243** | 1 | uf | 850 | 0.087639185 | 0 | 0 | 0 | 0 | 4 mo | HOM |
| **1013288951** | 2 | uf | 850 | 0.087639185 | 0 | 0 | 0 | 0 | 4 mo | HOM |
| **1013978665** | 1 | SCop | 851 | 0.661804192 | 9.04E-07 | 0 | 1.37E-06 | 0 | 2 mo | HOM |
| **1013445609** | 1 | SCop | 851 | 0.661804192 | 0 | 0 | 0 | 0 | 4 mo | HOM |
| **1013609015** | 1 | SCop | 851 | 0.661804192 | 3.23E-07 | 0 | 4.88E-07 | 0 | 4 mo | HOM |
| **1013608106** | 2 | SCop | 851 | 0.661804192 | 0 | 0 | 0 | 0 | 4 mo | HOM |
| **1011937543** | 1 | SCop | 851 | 0.661804192 | 0.001400022 | 27 | 0.002115463 | 40.79756569 | 8 mo | HOM |
| **1010419538** | 2 | SCop | 851 | 0.661804192 | 0 | 0 | 0 | 0 | 4 mo | WILD |

| | | | | | | | | | | |
|---|---|---|---|---|---|---|---|---|---|---|
| **1010419187** | 1 | SCop | 851 | 0.661804192 | 4.4E-05 | 3 | 6.65E-05 | 4.533062855 | 8 mo | HOM |
| **1013978366** | 1 | SCop | 851 | 0.661804192 | 0 | 0 | 0 | 0 | 2 mo | HOM |
| **1014416155** | 2 | SCop | 851 | 0.661804192 | 0 | 0 | 0 | 0 | 2 mo | HOM |
| **1013978065** | 2 | SCop | 851 | 0.661804192 | 0 | 0 | 0 | 0 | 2 mo | HOM |
| **1009934102** | 2 | SCop | 851 | 0.661804192 | 0 | 0 | 0 | 0 | 4 mo | WILD |
| **1009322937** | 2 | SCop | 851 | 0.661804192 | 6.17E-06 | 0 | 9.33E-06 | 0 | 8 mo | WILD |
| **1011080637** | 2 | SCop | 851 | 0.661804192 | 8.64E-05 | 5 | 0.00013057 | 7.555104758 | 8 mo | HOM |
| **1013609713** | 2 | SCop | 851 | 0.661804192 | 0 | 0 | 0 | 0 | 2 mo | WILD |
| **1010418842** | 1 | SCop | 851 | 0.661804192 | 1.56E-05 | 1 | 2.36E-05 | 1.511020952 | 8 mo | HOM |
| **1013288213** | 2 | SCop | 851 | 0.661804192 | 1.59E-06 | 0 | 2.41E-06 | 0 | 4 mo | HOM |
| **1008976060** | 2 | SCop | 851 | 0.661804192 | 1.22E-05 | 1 | 1.85E-05 | 1.511020952 | 8 mo | WILD |
| **1013446243** | 1 | SCop | 851 | 0.661804192 | 0 | 0 | 0 | 0 | 4 mo | HOM |
| **1013288951** | 2 | SCop | 851 | 0.661804192 | 0 | 0 | 0 | 0 | 4 mo | HOM |
| **1013978665** | 1 | PARN | 852 | 2.298034301 | 0 | 0 | 0 | 0 | 2 mo | HOM |
| **1013445609** | 1 | PARN | 852 | 2.298034301 | 0 | 0 | 0 | 0 | 4 mo | HOM |
| **1013609015** | 1 | PARN | 852 | 2.298034301 | 0 | 0 | 0 | 0 | 4 mo | HOM |
| **1013608106** | 2 | PARN | 852 | 2.298034301 | 0 | 0 | 0 | 0 | 4 mo | HOM |
| **1011937543** | 1 | PARN | 852 | 2.298034301 | 0.002213894 | 4 | 0.000963386 | 1.740618057 | 8 mo | HOM |
| **1010419538** | 2 | PARN | 852 | 2.298034301 | 0 | 0 | 0 | 0 | 4 mo | WILD |
| **1010419187** | 1 | PARN | 852 | 2.298034301 | 0 | 0 | 0 | 0 | 8 mo | HOM |
| **1013978366** | 1 | PARN | 852 | 2.298034301 | 0 | 0 | 0 | 0 | 2 mo | HOM |
| **1014416155** | 2 | PARN | 852 | 2.298034301 | 0 | 0 | 0 | 0 | 2 mo | HOM |
| **1013978065** | 2 | PARN | 852 | 2.298034301 | 0 | 0 | 0 | 0 | 2 mo | HOM |
| **1009934102** | 2 | PARN | 852 | 2.298034301 | 0 | 0 | 0 | 0 | 4 mo | WILD |
| **1009322937** | 2 | PARN | 852 | 2.298034301 | 0 | 0 | 0 | 0 | 8 mo | WILD |
| **1011080637** | 2 | PARN | 852 | 2.298034301 | 1.54E-06 | 0 | 6.69E-07 | 0 | 8 mo | HOM |
| **1013609713** | 2 | PARN | 852 | 2.298034301 | 0 | 0 | 0 | 0 | 2 mo | WILD |
| **1010418842** | 1 | PARN | 852 | 2.298034301 | 0.000292001 | 13 | 0.000127066 | 5.657008686 | 8 mo | HOM |
| **1013288213** | 2 | PARN | 852 | 2.298034301 | 0 | 0 | 0 | 0 | 4 mo | HOM |
| **1008976060** | 2 | PARN | 852 | 2.298034301 | 2.64E-05 | 2 | 1.15E-05 | 0.870309029 | 8 mo | WILD |
| **1013446243** | 1 | PARN | 852 | 2.298034301 | 0 | 0 | 0 | 0 | 4 mo | HOM |
| **1013288951** | 2 | PARN | 852 | 2.298034301 | 0 | 0 | 0 | 0 | 4 mo | HOM |
| **1013978665** | 1 | SSp-ul2/3 | 854 | 1.00854497 | 0 | 0 | 0 | 0 | 2 mo | HOM |
| **1013445609** | 1 | SSp-ul2/3 | 854 | 1.00854497 | 0.000109037 | 8 | 0.000108113 | 7.932219426 | 4 mo | HOM |
| **1013609015** | 1 | SSp-ul2/3 | 854 | 1.00854497 | 0.000334265 | 12 | 0.000331433 | 11.89832914 | 4 mo | HOM |
| **1013608106** | 2 | SSp-ul2/3 | 854 | 1.00854497 | 0 | 0 | 0 | 0 | 4 mo | HOM |
| **1011937543** | 1 | SSp-ul2/3 | 854 | 1.00854497 | 0.001453913 | 36 | 0.001441595 | 35.69498742 | 8 mo | HOM |
| **1010419538** | 2 | SSp-ul2/3 | 854 | 1.00854497 | 0 | 0 | 0 | 0 | 4 mo | WILD |
| **1010419187** | 1 | SSp-ul2/3 | 854 | 1.00854497 | 0.001747732 | 57 | 0.001732924 | 56.51706341 | 8 mo | HOM |
| **1013978366** | 1 | SSp-ul2/3 | 854 | 1.00854497 | 0 | 0 | 0 | 0 | 2 mo | HOM |
| **1014416155** | 2 | SSp-ul2/3 | 854 | 1.00854497 | 0 | 0 | 0 | 0 | 2 mo | HOM |
| **1013978065** | 2 | SSp-ul2/3 | 854 | 1.00854497 | 0 | 0 | 0 | 0 | 2 mo | HOM |
| **1009934102** | 2 | SSp-ul2/3 | 854 | 1.00854497 | 0 | 0 | 0 | 0 | 4 mo | WILD |
| **1009322937** | 2 | SSp-ul2/3 | 854 | 1.00854497 | 0 | 0 | 0 | 0 | 8 mo | WILD |
| **1011080637** | 2 | SSp-ul2/3 | 854 | 1.00854497 | 0.001026142 | 31 | 0.001017448 | 30.73735028 | 8 mo | HOM |
| **1013609713** | 2 | SSp-ul2/3 | 854 | 1.00854497 | 0 | 0 | 0 | 0 | 2 mo | WILD |
| **1010418842** | 1 | SSp-ul2/3 | 854 | 1.00854497 | 0.000762453 | 33 | 0.000755993 | 32.72040513 | 8 mo | HOM |
| **1013288213** | 2 | SSp-ul2/3 | 854 | 1.00854497 | 9.3E-05 | 6 | 9.22E-05 | 5.94916457 | 4 mo | HOM |
| **1008976060** | 2 | SSp-ul2/3 | 854 | 1.00854497 | 0 | 0 | 0 | 0 | 8 mo | WILD |
| **1013446243** | 1 | SSp-ul2/3 | 854 | 1.00854497 | 0.00014353 | 8 | 0.000142314 | 7.932219426 | 4 mo | HOM |
| **1013288951** | 2 | SSp-ul2/3 | 854 | 1.00854497 | 0.000235059 | 8 | 0.000233067 | 7.932219426 | 4 mo | HOM |
| **1013978665** | 1 | DORpm | 856 | 13.0573252 | 6.06E-09 | 0 | 4.64E-10 | 0 | 2 mo | HOM |
| **1013445609** | 1 | DORpm | 856 | 13.0573252 | 9.42E-05 | 7 | 7.22E-06 | 0.536097546 | 4 mo | HOM |
| **1013609015** | 1 | DORpm | 856 | 13.0573252 | 0.001115528 | 38 | 8.54E-05 | 2.910243823 | 4 mo | HOM |
| **1013608106** | 2 | DORpm | 856 | 13.0573252 | 0 | 0 | 0 | 0 | 4 mo | HOM |
| **1011937543** | 1 | DORpm | 856 | 13.0573252 | 0.013822406 | 183 | 0.001058594 | 14.01512157 | 8 mo | HOM |
| **1010419538** | 2 | DORpm | 856 | 13.0573252 | 2.57E-05 | 1 | 1.97E-06 | 0.076585364 | 4 mo | WILD |
| **1010419187** | 1 | DORpm | 856 | 13.0573252 | 0.010139793 | 230 | 0.00077656 | 17.61463367 | 8 mo | HOM |
| **1013978366** | 1 | DORpm | 856 | 13.0573252 | 1.51E-05 | 2 | 1.15E-06 | 0.153170728 | 2 mo | HOM |

| | | | | | | | | | | |
|---|---|---|---|---|---|---|---|---|---|---|
| **1014416155** | 2 | DORpm | 856 | 13.0573252 | 0 | 0 | 0 | 0 | 2 mo | HOM |
| **1013978065** | 2 | DORpm | 856 | 13.0573252 | 9.78E-07 | 0 | 7.49E-08 | 0 | 2 mo | HOM |
| **1009934102** | 2 | DORpm | 856 | 13.0573252 | 5.12E-06 | 1 | 3.92E-07 | 0.076585364 | 4 mo | WILD |
| **1009322937** | 2 | DORpm | 856 | 13.0573252 | 9.37E-05 | 13 | 7.18E-06 | 0.995609729 | 8 mo | WILD |
| **1011080637** | 2 | DORpm | 856 | 13.0573252 | 0.008389266 | 208 | 0.000642495 | 15.92975566 | 8 mo | HOM |
| **1013609713** | 2 | DORpm | 856 | 13.0573252 | 0 | 0 | 0 | 0 | 2 mo | WILD |
| **1010418842** | 1 | DORpm | 856 | 13.0573252 | 0.013023297 | 289 | 0.000997394 | 22.13317013 | 8 mo | HOM |
| **1013288213** | 2 | DORpm | 856 | 13.0573252 | 7.99E-05 | 8 | 6.12E-06 | 0.61268291 | 4 mo | HOM |
| **1008976060** | 2 | DORpm | 856 | 13.0573252 | 0.000813586 | 50 | 6.23E-05 | 3.829268189 | 8 mo | WILD |
| **1013446243** | 1 | DORpm | 856 | 13.0573252 | 0.000390855 | 20 | 2.99E-05 | 1.531707275 | 4 mo | HOM |
| **1013288951** | 2 | DORpm | 856 | 13.0573252 | 0.000762067 | 13 | 5.84E-05 | 0.995609729 | 4 mo | HOM |
| **1013978665** | 1 | VISC6a | 857 | 0.550918995 | 0 | 0 | 0 | 0 | 2 mo | HOM |
| **1013445609** | 1 | VISC6a | 857 | 0.550918995 | 4.21E-05 | 4 | 7.64E-05 | 7.260595541 | 4 mo | HOM |
| **1013609015** | 1 | VISC6a | 857 | 0.550918995 | 0.000196581 | 10 | 0.000356824 | 18.15148885 | 4 mo | HOM |
| **1013608106** | 2 | VISC6a | 857 | 0.550918995 | 0 | 0 | 0 | 0 | 4 mo | HOM |
| **1011937543** | 1 | VISC6a | 857 | 0.550918995 | 0.001393388 | 27 | 0.002529206 | 49.0090199 | 8 mo | HOM |
| **1010419538** | 2 | VISC6a | 857 | 0.550918995 | 0 | 0 | 0 | 0 | 4 mo | WILD |
| **1010419187** | 1 | VISC6a | 857 | 0.550918995 | 0.000462368 | 25 | 0.000839267 | 45.37872213 | 8 mo | HOM |
| **1013978366** | 1 | VISC6a | 857 | 0.550918995 | 0 | 0 | 0 | 0 | 2 mo | HOM |
| **1014416155** | 2 | VISC6a | 857 | 0.550918995 | 0 | 0 | 0 | 0 | 2 mo | HOM |
| **1013978065** | 2 | VISC6a | 857 | 0.550918995 | 0 | 0 | 0 | 0 | 2 mo | HOM |
| **1009934102** | 2 | VISC6a | 857 | 0.550918995 | 0 | 0 | 0 | 0 | 4 mo | WILD |
| **1009322937** | 2 | VISC6a | 857 | 0.550918995 | 0 | 0 | 0 | 0 | 8 mo | WILD |
| **1011080637** | 2 | VISC6a | 857 | 0.550918995 | 0.000422375 | 28 | 0.000766673 | 50.82416879 | 8 mo | HOM |
| **1013609713** | 2 | VISC6a | 857 | 0.550918995 | 0 | 0 | 0 | 0 | 2 mo | WILD |
| **1010418842** | 1 | VISC6a | 857 | 0.550918995 | 0.000417917 | 29 | 0.000758581 | 52.63931767 | 8 mo | HOM |
| **1013288213** | 2 | VISC6a | 857 | 0.550918995 | 5.89E-06 | 1 | 1.07E-05 | 1.815148885 | 4 mo | HOM |
| **1008976060** | 2 | VISC6a | 857 | 0.550918995 | 2.37E-07 | 0 | 4.3E-07 | 0 | 8 mo | WILD |
| **1013446243** | 1 | VISC6a | 857 | 0.550918995 | 0.000174825 | 11 | 0.000317333 | 19.96663774 | 4 mo | HOM |
| **1013288951** | 2 | VISC6a | 857 | 0.550918995 | 6.14E-05 | 7 | 0.000111424 | 12.7060422 | 4 mo | HOM |
| **1013978665** | 1 | PAS | 859 | 0.028751519 | 0 | 0 | 0 | 0 | 2 mo | HOM |
| **1013445609** | 1 | PAS | 859 | 0.028751519 | 0 | 0 | 0 | 0 | 4 mo | HOM |
| **1013609015** | 1 | PAS | 859 | 0.028751519 | 0 | 0 | 0 | 0 | 4 mo | HOM |
| **1013608106** | 2 | PAS | 859 | 0.028751519 | 0 | 0 | 0 | 0 | 4 mo | HOM |
| **1011937543** | 1 | PAS | 859 | 0.028751519 | 9.66E-06 | 1 | 0.000336139 | 34.7807707 | 8 mo | HOM |
| **1010419538** | 2 | PAS | 859 | 0.028751519 | 0 | 0 | 0 | 0 | 4 mo | WILD |
| **1010419187** | 1 | PAS | 859 | 0.028751519 | 0 | 0 | 0 | 0 | 8 mo | HOM |
| **1013978366** | 1 | PAS | 859 | 0.028751519 | 0 | 0 | 0 | 0 | 2 mo | HOM |
| **1014416155** | 2 | PAS | 859 | 0.028751519 | 0 | 0 | 0 | 0 | 2 mo | HOM |
| **1013978065** | 2 | PAS | 859 | 0.028751519 | 0 | 0 | 0 | 0 | 2 mo | HOM |
| **1009934102** | 2 | PAS | 859 | 0.028751519 | 0 | 0 | 0 | 0 | 4 mo | WILD |
| **1009322937** | 2 | PAS | 859 | 0.028751519 | 0 | 0 | 0 | 0 | 8 mo | WILD |
| **1011080637** | 2 | PAS | 859 | 0.028751519 | 0 | 0 | 0 | 0 | 8 mo | HOM |
| **1013609713** | 2 | PAS | 859 | 0.028751519 | 0 | 0 | 0 | 0 | 2 mo | WILD |
| **1010418842** | 1 | PAS | 859 | 0.028751519 | 2.94E-06 | 1 | 0.000102422 | 34.7807707 | 8 mo | HOM |
| **1013288213** | 2 | PAS | 859 | 0.028751519 | 0 | 0 | 0 | 0 | 4 mo | HOM |
| **1008976060** | 2 | PAS | 859 | 0.028751519 | 8.56E-07 | 0 | 2.98E-05 | 0 | 8 mo | WILD |
| **1013446243** | 1 | PAS | 859 | 0.028751519 | 0 | 0 | 0 | 0 | 4 mo | HOM |
| **1013288951** | 2 | PAS | 859 | 0.028751519 | 0 | 0 | 0 | 0 | 4 mo | HOM |
| **1013978665** | 1 | SSs6a | 862 | 2.135808864 | 0 | 0 | 0 | 0 | 2 mo | HOM |
| **1013445609** | 1 | SSs6a | 862 | 2.135808864 | 0.000135536 | 11 | 6.35E-05 | 5.150273597 | 4 mo | HOM |
| **1013609015** | 1 | SSs6a | 862 | 2.135808864 | 0.000361313 | 21 | 0.000169169 | 9.832340503 | 4 mo | HOM |
| **1013608106** | 2 | SSs6a | 862 | 2.135808864 | 0 | 0 | 0 | 0 | 4 mo | HOM |
| **1011937543** | 1 | SSs6a | 862 | 2.135808864 | 0.002442912 | 56 | 0.001143788 | 26.21957467 | 8 mo | HOM |
| **1010419538** | 2 | SSs6a | 862 | 2.135808864 | 0 | 0 | 0 | 0 | 4 mo | WILD |
| **1010419187** | 1 | SSs6a | 862 | 2.135808864 | 0.001938118 | 102 | 0.00090744 | 47.75708244 | 8 mo | HOM |
| **1013978366** | 1 | SSs6a | 862 | 2.135808864 | 0 | 0 | 0 | 0 | 2 mo | HOM |
| **1014416155** | 2 | SSs6a | 862 | 2.135808864 | 0 | 0 | 0 | 0 | 2 mo | HOM |
| **1013978065** | 2 | SSs6a | 862 | 2.135808864 | 0 | 0 | 0 | 0 | 2 mo | HOM |

| | | | | | | | | | | |
|---|---|---|---|---|---|---|---|---|---|---|
| **1009934102** | 2 | SSs6a | 862 | 2.135808864 | 0 | 0 | 0 | 0 | 4 mo | WILD |
| **1009322937** | 2 | SSs6a | 862 | 2.135808864 | 0 | 0 | 0 | 0 | 8 mo | WILD |
| **1011080637** | 2 | SSs6a | 862 | 2.135808864 | 0.001356748 | 68 | 0.000635238 | 31.83805496 | 8 mo | HOM |
| **1013609713** | 2 | SSs6a | 862 | 2.135808864 | 0 | 0 | 0 | 0 | 2 mo | WILD |
| **1010418842** | 1 | SSs6a | 862 | 2.135808864 | 0.001982077 | 84 | 0.000928022 | 39.32936201 | 8 mo | HOM |
| **1013288213** | 2 | SSs6a | 862 | 2.135808864 | 2.29E-05 | 3 | 1.07E-05 | 1.404620072 | 4 mo | HOM |
| **1008976060** | 2 | SSs6a | 862 | 2.135808864 | 2.32E-05 | 3 | 1.09E-05 | 1.404620072 | 8 mo | WILD |
| **1013446243** | 1 | SSs6a | 862 | 2.135808864 | 0.000231263 | 20 | 0.000108279 | 9.364133812 | 4 mo | HOM |
| **1013288951** | 2 | SSs6a | 862 | 2.135808864 | 0.000248694 | 13 | 0.00011644 | 6.086686978 | 4 mo | HOM |
| **1013978665** | 1 | rust | 863 | 0.624161843 | 0 | 0 | 0 | 0 | 2 mo | HOM |
| **1013445609** | 1 | rust | 863 | 0.624161843 | 0 | 0 | 0 | 0 | 4 mo | HOM |
| **1013609015** | 1 | rust | 863 | 0.624161843 | 0 | 0 | 0 | 0 | 4 mo | HOM |
| **1013608106** | 2 | rust | 863 | 0.624161843 | 0 | 0 | 0 | 0 | 4 mo | HOM |
| **1011937543** | 1 | rust | 863 | 0.624161843 | 0.00458189 | 3 | 0.007340869 | 4.806445688 | 8 mo | HOM |
| **1010419538** | 2 | rust | 863 | 0.624161843 | 0 | 0 | 0 | 0 | 4 mo | WILD |
| **1010419187** | 1 | rust | 863 | 0.624161843 | 0 | 0 | 0 | 0 | 8 mo | HOM |
| **1013978366** | 1 | rust | 863 | 0.624161843 | 0 | 0 | 0 | 0 | 2 mo | HOM |
| **1014416155** | 2 | rust | 863 | 0.624161843 | 0 | 0 | 0 | 0 | 2 mo | HOM |
| **1013978065** | 2 | rust | 863 | 0.624161843 | 0 | 0 | 0 | 0 | 2 mo | HOM |
| **1009934102** | 2 | rust | 863 | 0.624161843 | 0 | 0 | 0 | 0 | 4 mo | WILD |
| **1009322937** | 2 | rust | 863 | 0.624161843 | 0 | 0 | 0 | 0 | 8 mo | WILD |
| **1011080637** | 2 | rust | 863 | 0.624161843 | 0 | 0 | 0 | 0 | 8 mo | HOM |
| **1013609713** | 2 | rust | 863 | 0.624161843 | 0 | 0 | 0 | 0 | 2 mo | WILD |
| **1010418842** | 1 | rust | 863 | 0.624161843 | 5.02E-06 | 0 | 8.04E-06 | 0 | 8 mo | HOM |
| **1013288213** | 2 | rust | 863 | 0.624161451 | 0 | 0 | 0 | 0 | 4 mo | HOM |
| **1008976060** | 2 | rust | 863 | 0.624161843 | 0 | 0 | 0 | 0 | 8 mo | WILD |
| **1013446243** | 1 | rust | 863 | 0.624161843 | 0 | 0 | 0 | 0 | 4 mo | HOM |
| **1013288951** | 2 | rust | 863 | 0.624161843 | 0 | 0 | 0 | 0 | 4 mo | HOM |
| **1013978665** | 1 | DORsm | 864 | 6.882650965 | 0 | 0 | 0 | 0 | 2 mo | HOM |
| **1013445609** | 1 | DORsm | 864 | 6.882650965 | 3.3E-05 | 3 | 4.79E-06 | 0.435878561 | 4 mo | HOM |
| **1013609015** | 1 | DORsm | 864 | 6.882650965 | 0.000249106 | 7 | 3.62E-05 | 1.017049976 | 4 mo | HOM |
| **1013608106** | 2 | DORsm | 864 | 6.882650965 | 0 | 0 | 0 | 0 | 4 mo | HOM |
| **1011937543** | 1 | DORsm | 864 | 6.882650965 | 0.004463551 | 72 | 0.000648522 | 10.46108547 | 8 mo | HOM |
| **1010419538** | 2 | DORsm | 864 | 6.882650965 | 0 | 0 | 0 | 0 | 4 mo | WILD |
| **1010419187** | 1 | DORsm | 864 | 6.882650965 | 0.00160048 | 50 | 0.000232538 | 7.264642687 | 8 mo | HOM |
| **1013978366** | 1 | DORsm | 864 | 6.882650965 | 0 | 0 | 0 | 0 | 2 mo | HOM |
| **1014416155** | 2 | DORsm | 864 | 6.882650965 | 0 | 0 | 0 | 0 | 2 mo | HOM |
| **1013978065** | 2 | DORsm | 864 | 6.882650965 | 0 | 0 | 0 | 0 | 2 mo | HOM |
| **1009934102** | 2 | DORsm | 864 | 6.882650965 | 0 | 0 | 0 | 0 | 4 mo | WILD |
| **1009322937** | 2 | DORsm | 864 | 6.882650965 | 1.48E-05 | 3 | 2.15E-06 | 0.435878561 | 8 mo | WILD |
| **1011080637** | 2 | DORsm | 864 | 6.882650965 | 0.002269386 | 66 | 0.000329726 | 9.589328346 | 8 mo | HOM |
| **1013609713** | 2 | DORsm | 864 | 6.882650965 | 0 | 0 | 0 | 0 | 2 mo | WILD |
| **1010418842** | 1 | DORsm | 864 | 6.882650965 | 0.003785918 | 94 | 0.000550067 | 13.65752825 | 8 mo | HOM |
| **1013288213** | 2 | DORsm | 864 | 6.882650965 | 2.25E-05 | 3 | 3.27E-06 | 0.435878561 | 4 mo | HOM |
| **1008976060** | 2 | DORsm | 864 | 6.882650965 | 0.000179988 | 15 | 2.62E-05 | 2.179392806 | 8 mo | WILD |
| **1013446243** | 1 | DORsm | 864 | 6.882650965 | 4.48E-05 | 3 | 6.51E-06 | 0.435878561 | 4 mo | HOM |
| **1013288951** | 2 | DORsm | 864 | 6.882650965 | 2.58E-05 | 4 | 3.75E-06 | 0.581171415 | 4 mo | HOM |
| **1013978665** | 1 | sctv | 866 | 0.292763162 | 0 | 0 | 0 | 0 | 2 mo | HOM |
| **1013445609** | 1 | sctv | 866 | 0.292763162 | 2.64E-06 | 0 | 9.01E-06 | 0 | 4 mo | HOM |
| **1013609015** | 1 | sctv | 866 | 0.292763162 | 3.08E-06 | 0 | 1.05E-05 | 0 | 4 mo | HOM |
| **1013608106** | 2 | sctv | 866 | 0.292763162 | 0 | 0 | 0 | 0 | 4 mo | HOM |
| **1011937543** | 1 | sctv | 866 | 0.292763162 | 0.000936486 | 3 | 0.003198784 | 10.24719088 | 8 mo | HOM |
| **1010419538** | 2 | sctv | 866 | 0.292763162 | 0 | 0 | 0 | 0 | 4 mo | WILD |
| **1010419187** | 1 | sctv | 866 | 0.292763162 | 5.86E-06 | 0 | 2E-05 | 0 | 8 mo | HOM |
| **1013978366** | 1 | sctv | 866 | 0.292763162 | 0 | 0 | 0 | 0 | 2 mo | HOM |
| **1014416155** | 2 | sctv | 866 | 0.292763162 | 0 | 0 | 0 | 0 | 2 mo | HOM |
| **1013978065** | 2 | sctv | 866 | 0.292763162 | 0 | 0 | 0 | 0 | 2 mo | HOM |
| **1009934102** | 2 | sctv | 866 | 0.292763162 | 0 | 0 | 0 | 0 | 4 mo | WILD |
| **1009322937** | 2 | sctv | 866 | 0.292763162 | 8.18E-06 | 0 | 2.79E-05 | 0 | 8 mo | WILD |

| | | | | | | | | | | |
|---|---|---|---|---|---|---|---|---|---|---|
| **1011080637** | 2 | sctv | 866 | 0.292763162 | 3.32E-06 | 0 | 1.13E-05 | 0 | 8 mo | HOM |
| **1013609713** | 2 | sctv | 866 | 0.292763162 | 3.65E-10 | 0 | 1.25E-09 | 0 | 2 mo | WILD |
| **1010418842** | 1 | sctv | 866 | 0.292763162 | 0.000206168 | 1 | 0.000704215 | 3.415730294 | 8 mo | HOM |
| **1013288213** | 2 | sctv | 866 | 0.292763162 | 0 | 0 | 0 | 0 | 4 mo | HOM |
| **1008976060** | 2 | sctv | 866 | 0.292763162 | 1.25E-05 | 0 | 4.27E-05 | 0 | 8 mo | WILD |
| **1013446243** | 1 | sctv | 866 | 0.292763162 | 7.42E-06 | 0 | 2.53E-05 | 0 | 4 mo | HOM |
| **1013288951** | 2 | sctv | 866 | 0.292763162 | 1.15E-06 | 0 | 3.92E-06 | 0 | 4 mo | HOM |
| **1013978665** | 1 | PB | 867 | 1.155566294 | 4.19E-05 | 5 | 3.63E-05 | 4.326882866 | 2 mo | HOM |
| **1013445609** | 1 | PB | 867 | 1.155566294 | 7.72E-06 | 1 | 6.68E-06 | 0.865376573 | 4 mo | HOM |
| **1013609015** | 1 | PB | 867 | 1.155566294 | 7.36E-05 | 4 | 6.37E-05 | 3.461506293 | 4 mo | HOM |
| **1013608106** | 2 | PB | 867 | 1.155566294 | 0 | 0 | 0 | 0 | 4 mo | HOM |
| **1011937543** | 1 | PB | 867 | 1.155566294 | 0.001166096 | 15 | 0.001009112 | 12.9806486 | 8 mo | HOM |
| **1010419538** | 2 | PB | 867 | 1.155566294 | 0 | 0 | 0 | 0 | 4 mo | WILD |
| **1010419187** | 1 | PB | 867 | 1.155566294 | 0.000124614 | 8 | 0.000107838 | 6.923012586 | 8 mo | HOM |
| **1013978366** | 1 | PB | 867 | 1.155566294 | 1.1E-05 | 2 | 9.54E-06 | 1.730753146 | 2 mo | HOM |
| **1014416155** | 2 | PB | 867 | 1.155566294 | 0 | 0 | 0 | 0 | 2 mo | HOM |
| **1013978065** | 2 | PB | 867 | 1.155566294 | 0 | 0 | 0 | 0 | 2 mo | HOM |
| **1009934102** | 2 | PB | 867 | 1.155566294 | 3.15E-08 | 0 | 2.73E-08 | 0 | 4 mo | WILD |
| **1009322937** | 2 | PB | 867 | 1.155566294 | 6.45E-05 | 3 | 5.58E-05 | 2.59612972 | 8 mo | WILD |
| **1011080637** | 2 | PB | 867 | 1.155566294 | 0.000172151 | 9 | 0.000148975 | 7.788389159 | 8 mo | HOM |
| **1013609713** | 2 | PB | 867 | 1.155566294 | 0 | 0 | 0 | 0 | 2 mo | WILD |
| **1010418842** | 1 | PB | 867 | 1.155566294 | 0.00018225 | 7 | 0.000157715 | 6.057636013 | 8 mo | HOM |
| **1013288213** | 2 | PB | 867 | 1.155566294 | 7.08E-06 | 1 | 6.13E-06 | 0.865376573 | 4 mo | HOM |
| **1008976060** | 2 | PB | 867 | 1.155566294 | 0.000255307 | 10 | 0.000220937 | 8.653765732 | 8 mo | WILD |
| **1013446243** | 1 | PB | 867 | 1.155566294 | 6.82E-05 | 5 | 5.9E-05 | 4.326882866 | 4 mo | HOM |
| **1013288951** | 2 | PB | 867 | 1.155566294 | 1.91E-05 | 2 | 1.65E-05 | 1.730753146 | 4 mo | HOM |
| **1013978665** | 1 | VISpl4 | 869 | 0.037745384 | 0 | 0 | 0 | 0 | 2 mo | HOM |
| **1013445609** | 1 | VISpl4 | 869 | 0.037745384 | 8.82E-06 | 1 | 0.000233622 | 26.49330609 | 4 mo | HOM |
| **1013609015** | 1 | VISpl4 | 869 | 0.037745384 | 4.02E-06 | 0 | 0.000106524 | 0 | 4 mo | HOM |
| **1013608106** | 2 | VISpl4 | 869 | 0.037745384 | 0 | 0 | 0 | 0 | 4 mo | HOM |
| **1011937543** | 1 | VISpl4 | 869 | 0.037745384 | 2.6E-05 | 2 | 0.00068873 | 52.98661219 | 8 mo | HOM |
| **1010419538** | 2 | VISpl4 | 869 | 0.037745384 | 0 | 0 | 0 | 0 | 4 mo | WILD |
| **1010419187** | 1 | VISpl4 | 869 | 0.037745384 | 3.31E-05 | 0 | 0.000875703 | 0 | 8 mo | HOM |
| **1013978366** | 1 | VISpl4 | 869 | 0.037745384 | 6.05E-10 | 0 | 1.6E-08 | 0 | 2 mo | HOM |
| **1014416155** | 2 | VISpl4 | 869 | 0.037745384 | 0 | 0 | 0 | 0 | 2 mo | HOM |
| **1013978065** | 2 | VISpl4 | 869 | 0.037745384 | 0 | 0 | 0 | 0 | 2 mo | HOM |
| **1009934102** | 2 | VISpl4 | 869 | 0.037745384 | 0 | 0 | 0 | 0 | 4 mo | WILD |
| **1009322937** | 2 | VISpl4 | 869 | 0.037745384 | 0 | 0 | 0 | 0 | 8 mo | WILD |
| **1011080637** | 2 | VISpl4 | 869 | 0.037745384 | 3.58E-05 | 0 | 0.000947681 | 0 | 8 mo | HOM |
| **1013609713** | 2 | VISpl4 | 869 | 0.037745384 | 0 | 0 | 0 | 0 | 2 mo | WILD |
| **1010418842** | 1 | VISpl4 | 869 | 0.037745384 | 0.000114305 | 3 | 0.003028313 | 79.47991828 | 8 mo | HOM |
| **1013288213** | 2 | VISpl4 | 869 | 0.037745384 | 1.79E-06 | 0 | 4.75E-05 | 0 | 4 mo | HOM |
| **1008976060** | 2 | VISpl4 | 869 | 0.037745384 | 0 | 0 | 0 | 0 | 8 mo | WILD |
| **1013446243** | 1 | VISpl4 | 869 | 0.037745384 | 3.7E-05 | 1 | 0.000979037 | 26.49330609 | 4 mo | HOM |
| **1013288951** | 2 | VISpl4 | 869 | 0.037745384 | 1.52E-06 | 0 | 4.03E-05 | 0 | 4 mo | HOM |
| **1013978665** | 1 | DR | 872 | 0.157513166 | 0 | 0 | 0 | 0 | 2 mo | HOM |
| **1013445609** | 1 | DR | 872 | 0.157513166 | 0 | 0 | 0 | 0 | 4 mo | HOM |
| **1013609015** | 1 | DR | 872 | 0.157513166 | 6.49E-06 | 1 | 4.12E-05 | 6.348675656 | 4 mo | HOM |
| **1013608106** | 2 | DR | 872 | 0.157513166 | 0 | 0 | 0 | 0 | 4 mo | HOM |
| **1011937543** | 1 | DR | 872 | 0.157513166 | 0.000326428 | 1 | 0.002072386 | 6.348675656 | 8 mo | HOM |
| **1010419538** | 2 | DR | 872 | 0.157513166 | 0 | 0 | 0 | 0 | 4 mo | WILD |
| **1010419187** | 1 | DR | 872 | 0.157513166 | 3.89E-05 | 1 | 0.000247203 | 6.348675656 | 8 mo | HOM |
| **1013978366** | 1 | DR | 872 | 0.157513166 | 0 | 0 | 0 | 0 | 2 mo | HOM |
| **1014416155** | 2 | DR | 872 | 0.157513166 | 0 | 0 | 0 | 0 | 2 mo | HOM |
| **1013978065** | 2 | DR | 872 | 0.157513166 | 0 | 0 | 0 | 0 | 2 mo | HOM |
| **1009934102** | 2 | DR | 872 | 0.157513166 | 0 | 0 | 0 | 0 | 4 mo | WILD |
| **1009322937** | 2 | DR | 872 | 0.157513166 | 5.38E-06 | 1 | 3.42E-05 | 6.348675656 | 8 mo | WILD |
| **1011080637** | 2 | DR | 872 | 0.157513166 | 1.48E-06 | 0 | 9.38E-06 | 0 | 8 mo | HOM |
| **1013609713** | 2 | DR | 872 | 0.157513166 | 0 | 0 | 0 | 0 | 2 mo | WILD |

| | | | | | | | | | | |
|---|---|---|---|---|---|---|---|---|---|---|
| **1010418842** | 1 | DR | 872 | 0.157513166 | 1E-05 | 1 | 6.38E-05 | 6.348675656 | 8 mo | HOM |
| **1013288213** | 2 | DR | 872 | 0.157513166 | 0 | 0 | 0 | 0 | 4 mo | HOM |
| **1008976060** | 2 | DR | 872 | 0.157513166 | 2.84E-09 | 0 | 1.8E-08 | 0 | 8 mo | WILD |
| **1013446243** | 1 | DR | 872 | 0.157513166 | 4.97E-09 | 0 | 3.16E-08 | 0 | 4 mo | HOM |
| **1013288951** | 2 | DR | 872 | 0.157513166 | 1.32E-05 | 0 | 8.36E-05 | 0 | 4 mo | HOM |
| **1013978665** | 1 | SSs1 | 873 | 1.275484504 | 3.02E-05 | 2 | 2.37E-05 | 1.568031594 | 2 mo | HOM |
| **1013445609** | 1 | SSs1 | 873 | 1.275484504 | 0.000342086 | 14 | 0.000268201 | 10.97622116 | 4 mo | HOM |
| **1013609015** | 1 | SSs1 | 873 | 1.275484504 | 0.001038967 | 23 | 0.000814566 | 18.03236333 | 4 mo | HOM |
| **1013608106** | 2 | SSs1 | 873 | 1.275484504 | 0 | 0 | 0 | 0 | 4 mo | HOM |
| **1011937543** | 1 | SSs1 | 873 | 1.275484504 | 0.0054948 | 69 | 0.00430801 | 54.09708999 | 8 mo | HOM |
| **1010419538** | 2 | SSs1 | 873 | 1.275484504 | 1.93E-06 | 0 | 1.51E-06 | 0 | 4 mo | WILD |
| **1010419187** | 1 | SSs1 | 873 | 1.275484504 | 0.002395866 | 57 | 0.001878397 | 44.68890043 | 8 mo | HOM |
| **1013978366** | 1 | SSs1 | 873 | 1.275484504 | 0.000128377 | 1 | 0.000100649 | 0.784015797 | 2 mo | HOM |
| **1014416155** | 2 | SSs1 | 873 | 1.275484504 | 0 | 0 | 0 | 0 | 2 mo | HOM |
| **1013978065** | 2 | SSs1 | 873 | 1.275484504 | 0 | 0 | 0 | 0 | 2 mo | HOM |
| **1009934102** | 2 | SSs1 | 873 | 1.275484504 | 1.53E-05 | 2 | 1.2E-05 | 1.568031594 | 4 mo | WILD |
| **1009322937** | 2 | SSs1 | 873 | 1.275484504 | 2.76E-05 | 4 | 2.16E-05 | 3.136063188 | 8 mo | WILD |
| **1011080637** | 2 | SSs1 | 873 | 1.275484504 | 0.001624068 | 56 | 0.001273295 | 43.90488463 | 8 mo | HOM |
| **1013609713** | 2 | SSs1 | 873 | 1.275484504 | 1.74E-05 | 0 | 1.36E-05 | 0 | 2 mo | WILD |
| **1010418842** | 1 | SSs1 | 873 | 1.275484504 | 0.005866343 | 85 | 0.004599306 | 66.64134275 | 8 mo | HOM |
| **1013288213** | 2 | SSs1 | 873 | 1.275484504 | 0.000205771 | 13 | 0.000161328 | 10.19220536 | 4 mo | HOM |
| **1008976060** | 2 | SSs1 | 873 | 1.275484504 | 0.000205924 | 8 | 0.000161448 | 6.272126376 | 8 mo | WILD |
| **1013446243** | 1 | SSs1 | 873 | 1.275484504 | 0.000385898 | 17 | 0.00030255 | 13.32826855 | 4 mo | HOM |
| **1013288951** | 2 | SSs1 | 873 | 1.275484504 | 0.000912183 | 31 | 0.000715166 | 24.30448971 | 4 mo | HOM |
| **1013978665** | 1 | PBG | 874 | 0.046116992 | 0 | 0 | 0 | 0 | 2 mo | HOM |
| **1013445609** | 1 | PBG | 874 | 0.046116992 | 0 | 0 | 0 | 0 | 4 mo | HOM |
| **1013609015** | 1 | PBG | 874 | 0.046116992 | 0 | 0 | 0 | 0 | 4 mo | HOM |
| **1013608106** | 2 | PBG | 874 | 0.046116992 | 0 | 0 | 0 | 0 | 4 mo | HOM |
| **1011937543** | 1 | PBG | 874 | 0.046116992 | 7.13E-05 | 2 | 0.001545558 | 43.36796303 | 8 mo | HOM |
| **1010419538** | 2 | PBG | 874 | 0.046116992 | 0 | 0 | 0 | 0 | 4 mo | WILD |
| **1010419187** | 1 | PBG | 874 | 0.046116992 | 0 | 0 | 0 | 0 | 8 mo | HOM |
| **1013978366** | 1 | PBG | 874 | 0.046116992 | 0 | 0 | 0 | 0 | 2 mo | HOM |
| **1014416155** | 2 | PBG | 874 | 0.046116992 | 0 | 0 | 0 | 0 | 2 mo | HOM |
| **1013978065** | 2 | PBG | 874 | 0.046116992 | 0 | 0 | 0 | 0 | 2 mo | HOM |
| **1009934102** | 2 | PBG | 874 | 0.046116992 | 0 | 0 | 0 | 0 | 4 mo | WILD |
| **1009322937** | 2 | PBG | 874 | 0.046116992 | 0 | 0 | 0 | 0 | 8 mo | WILD |
| **1011080637** | 2 | PBG | 874 | 0.046116992 | 0 | 0 | 0 | 0 | 8 mo | HOM |
| **1013609713** | 2 | PBG | 874 | 0.046116992 | 0 | 0 | 0 | 0 | 2 mo | WILD |
| **1010418842** | 1 | PBG | 874 | 0.046116992 | 4.07E-11 | 0 | 8.83E-10 | 0 | 8 mo | HOM |
| **1013288213** | 2 | PBG | 874 | 0.046116992 | 0 | 0 | 0 | 0 | 4 mo | HOM |
| **1008976060** | 2 | PBG | 874 | 0.046116992 | 0 | 0 | 0 | 0 | 8 mo | WILD |
| **1013446243** | 1 | PBG | 874 | 0.046116992 | 0 | 0 | 0 | 0 | 4 mo | HOM |
| **1013288951** | 2 | PBG | 874 | 0.046116992 | 0 | 0 | 0 | 0 | 4 mo | HOM |
| **1013978665** | 1 | tsp | 877 | 0.525940175 | 0 | 0 | 0 | 0 | 2 mo | HOM |
| **1013445609** | 1 | tsp | 877 | 0.525940175 | 0 | 0 | 0 | 0 | 4 mo | HOM |
| **1013609015** | 1 | tsp | 877 | 0.525940175 | 0 | 0 | 0 | 0 | 4 mo | HOM |
| **1013608106** | 2 | tsp | 877 | 0.520452763 | 0 | 0 | 0 | 0 | 4 mo | HOM |
| **1011937543** | 1 | tsp | 877 | 0.525940175 | 9.78E-05 | 3 | 0.00018596 | 5.704070815 | 8 mo | HOM |
| **1010419538** | 2 | tsp | 877 | 0.525940175 | 0 | 0 | 0 | 0 | 4 mo | WILD |
| **1010419187** | 1 | tsp | 877 | 0.525940175 | 0 | 0 | 0 | 0 | 8 mo | HOM |
| **1013978366** | 1 | tsp | 877 | 0.525940175 | 0 | 0 | 0 | 0 | 2 mo | HOM |
| **1014416155** | 2 | tsp | 877 | 0.525940175 | 0 | 0 | 0 | 0 | 2 mo | HOM |
| **1013978065** | 2 | tsp | 877 | 0.525940175 | 0 | 0 | 0 | 0 | 2 mo | HOM |
| **1009934102** | 2 | tsp | 877 | 0.525940175 | 0 | 0 | 0 | 0 | 4 mo | WILD |
| **1009322937** | 2 | tsp | 877 | 0.525940175 | 0 | 0 | 0 | 0 | 8 mo | WILD |
| **1011080637** | 2 | tsp | 877 | 0.525940175 | 0 | 0 | 0 | 0 | 8 mo | HOM |
| **1013609713** | 2 | tsp | 877 | 0.525940175 | 0 | 0 | 0 | 0 | 2 mo | WILD |
| **1010418842** | 1 | tsp | 877 | 0.525940175 | 2.36E-05 | 2 | 4.49E-05 | 3.802713877 | 8 mo | HOM |
| **1013288213** | 2 | tsp | 877 | 0.525940175 | 0 | 0 | 0 | 0 | 4 mo | HOM |

| | | | | | | | | | | |
|---|---|---|---|---|---|---|---|---|---|---|
| **1008976060** | 2 | tsp | 877 | 0.525940175 | 1.47E-05 | 1 | 2.79E-05 | 1.901356938 | 8 mo | WILD |
| **1013446243** | 1 | tsp | 877 | 0.525940175 | 0 | 0 | 0 | 0 | 4 mo | HOM |
| **1013288951** | 2 | tsp | 877 | 0.525940175 | 0 | 0 | 0 | 0 | 4 mo | HOM |
| **1013978665** | 1 | SSp-m1 | 878 | 0.804206032 | 1.86E-05 | 1 | 2.31E-05 | 1.243462446 | 2 mo | HOM |
| **1013445609** | 1 | SSp-m1 | 878 | 0.804206032 | 6.06E-05 | 4 | 7.53E-05 | 4.973849786 | 4 mo | HOM |
| **1013609015** | 1 | SSp-m1 | 878 | 0.804206032 | 0.000251042 | 11 | 0.000312161 | 13.67808691 | 4 mo | HOM |
| **1013608106** | 2 | SSp-m1 | 878 | 0.804206032 | 0 | 0 | 0 | 0 | 4 mo | HOM |
| **1011937543** | 1 | SSp-m1 | 878 | 0.804206032 | 0.001599904 | 30 | 0.00198942 | 37.30387339 | 8 mo | HOM |
| **1010419538** | 2 | SSp-m1 | 878 | 0.804206032 | 0 | 0 | 0 | 0 | 4 mo | WILD |
| **1010419187** | 1 | SSp-m1 | 878 | 0.804206032 | 0.001747176 | 67 | 0.002172548 | 83.31198391 | 8 mo | HOM |
| **1013978366** | 1 | SSp-m1 | 878 | 0.804206032 | 0 | 0 | 0 | 0 | 2 mo | HOM |
| **1014416155** | 2 | SSp-m1 | 878 | 0.804206032 | 4.09E-06 | 1 | 5.09E-06 | 1.243462446 | 2 mo | HOM |
| **1013978065** | 2 | SSp-m1 | 878 | 0.804206032 | 0 | 0 | 0 | 0 | 2 mo | HOM |
| **1009934102** | 2 | SSp-m1 | 878 | 0.804206032 | 1.13E-05 | 2 | 1.41E-05 | 2.486924893 | 4 mo | WILD |
| **1009322937** | 2 | SSp-m1 | 878 | 0.804206032 | 0 | 0 | 0 | 0 | 8 mo | WILD |
| **1011080637** | 2 | SSp-m1 | 878 | 0.804206032 | 0.000900751 | 28 | 0.00112005 | 34.8169485 | 8 mo | HOM |
| **1013609713** | 2 | SSp-m1 | 878 | 0.804206032 | 4.37E-06 | 1 | 5.43E-06 | 1.243462446 | 2 mo | WILD |
| **1010418842** | 1 | SSp-m1 | 878 | 0.804206032 | 0.002012416 | 58 | 0.002502364 | 72.12082189 | 8 mo | HOM |
| **1013288213** | 2 | SSp-m1 | 878 | 0.804206032 | 5.9E-05 | 5 | 7.34E-05 | 6.217312232 | 4 mo | HOM |
| **1008976060** | 2 | SSp-m1 | 878 | 0.804206032 | 0.000345374 | 6 | 0.00042946 | 7.460774679 | 8 mo | WILD |
| **1013446243** | 1 | SSp-m1 | 878 | 0.804206032 | 0.000189488 | 11 | 0.000235621 | 13.67808691 | 4 mo | HOM |
| **1013288951** | 2 | SSp-m1 | 878 | 0.804206032 | 0.000206396 | 11 | 0.000256646 | 13.67808691 | 4 mo | HOM |
| **1013978665** | 1 | RSPd | 879 | 3.914222395 | 0.000111698 | 2 | 2.85E-05 | 0.510957171 | 2 mo | HOM |
| **1013445609** | 1 | RSPd | 879 | 3.914222395 | 0.000186141 | 13 | 4.76E-05 | 3.321221609 | 4 mo | HOM |
| **1013609015** | 1 | RSPd | 879 | 3.914222395 | 0.001032663 | 43 | 0.000263823 | 10.98557917 | 4 mo | HOM |
| **1013608106** | 2 | RSPd | 879 | 3.914222395 | 0 | 0 | 0 | 0 | 4 mo | HOM |
| **1011937543** | 1 | RSPd | 879 | 3.914222395 | 0.006833419 | 64 | 0.001745792 | 16.35062946 | 8 mo | HOM |
| **1010419538** | 2 | RSPd | 879 | 3.833028611 | 0.000372022 | 9 | 9.71E-05 | 2.348012737 | 4 mo | WILD |
| **1010419187** | 1 | RSPd | 879 | 3.914222395 | 0.010282395 | 150 | 0.002626932 | 38.32178779 | 8 mo | HOM |
| **1013978366** | 1 | RSPd | 879 | 3.914222395 | 0.000170251 | 2 | 4.35E-05 | 0.510957171 | 2 mo | HOM |
| **1014416155** | 2 | RSPd | 879 | 3.914222395 | 0 | 0 | 0 | 0 | 2 mo | HOM |
| **1013978065** | 2 | RSPd | 879 | 3.914222395 | 0 | 0 | 0 | 0 | 2 mo | HOM |
| **1009934102** | 2 | RSPd | 879 | 3.914222395 | 6.41E-05 | 4 | 1.64E-05 | 1.021914341 | 4 mo | WILD |
| **1009322937** | 2 | RSPd | 879 | 3.914222395 | 5.09E-05 | 4 | 1.3E-05 | 1.021914341 | 8 mo | WILD |
| **1011080637** | 2 | RSPd | 879 | 3.914222395 | 0.004261212 | 127 | 0.001088648 | 32.44578033 | 8 mo | HOM |
| **1013609713** | 2 | RSPd | 879 | 3.914222395 | 0 | 0 | 0 | 0 | 2 mo | WILD |
| **1010418842** | 1 | RSPd | 879 | 3.914222395 | 0.004119819 | 139 | 0.001052526 | 35.51152335 | 8 mo | HOM |
| **1013288213** | 2 | RSPd | 879 | 3.914222395 | 7.1E-05 | 8 | 1.81E-05 | 2.043828682 | 4 mo | HOM |
| **1008976060** | 2 | RSPd | 879 | 3.914222395 | 0.000131213 | 9 | 3.35E-05 | 2.299307267 | 8 mo | WILD |
| **1013446243** | 1 | RSPd | 879 | 3.914222395 | 0.000507241 | 29 | 0.000129589 | 7.408878973 | 4 mo | HOM |
| **1013288951** | 2 | RSPd | 879 | 3.914222395 | 0.002595883 | 29 | 0.000663192 | 7.408878973 | 4 mo | HOM |
| **1013978665** | 1 | DTN | 880 | 0.106869363 | 0 | 0 | 0 | 0 | 2 mo | HOM |
| **1013445609** | 1 | DTN | 880 | 0.106869363 | 0 | 0 | 0 | 0 | 4 mo | HOM |
| **1013609015** | 1 | DTN | 880 | 0.106869363 | 1.51E-06 | 0 | 1.41E-05 | 0 | 4 mo | HOM |
| **1013608106** | 2 | DTN | 880 | 0.106869363 | 0 | 0 | 0 | 0 | 4 mo | HOM |
| **1011937543** | 1 | DTN | 880 | 0.106869363 | 0.000582034 | 1 | 0.00544622 | 9.357218717 | 8 mo | HOM |
| **1010419538** | 2 | DTN | 880 | 0.106869363 | 0 | 0 | 0 | 0 | 4 mo | WILD |
| **1010419187** | 1 | DTN | 880 | 0.106869363 | 1.27E-06 | 0 | 1.18E-05 | 0 | 8 mo | HOM |
| **1013978366** | 1 | DTN | 880 | 0.106869363 | 0 | 0 | 0 | 0 | 2 mo | HOM |
| **1014416155** | 2 | DTN | 880 | 0.106869363 | 0 | 0 | 0 | 0 | 2 mo | HOM |
| **1013978065** | 2 | DTN | 880 | 0.106869363 | 0 | 0 | 0 | 0 | 2 mo | HOM |
| **1009934102** | 2 | DTN | 880 | 0.106869363 | 0 | 0 | 0 | 0 | 4 mo | WILD |
| **1009322937** | 2 | DTN | 880 | 0.106869363 | 5.05E-11 | 0 | 4.72E-10 | 0 | 8 mo | WILD |
| **1011080637** | 2 | DTN | 880 | 0.106869363 | 4.13E-06 | 0 | 3.86E-05 | 0 | 8 mo | HOM |
| **1013609713** | 2 | DTN | 880 | 0.106869363 | 0 | 0 | 0 | 0 | 2 mo | WILD |
| **1010418842** | 1 | DTN | 880 | 0.106869363 | 3.15E-08 | 0 | 2.94E-07 | 0 | 8 mo | HOM |
| **1013288213** | 2 | DTN | 880 | 0.106869363 | 0 | 0 | 0 | 0 | 4 mo | HOM |
| **1008976060** | 2 | DTN | 880 | 0.106869363 | 6.84E-05 | 0 | 0.000640189 | 0 | 8 mo | WILD |
| **1013446243** | 1 | DTN | 880 | 0.106869363 | 0 | 0 | 0 | 0 | 4 mo | HOM |

| | | | | | | | | | | |
|---|---|---|---|---|---|---|---|---|---|---|
| **1013288951** | 2 | DTN | 880 | 0.106869363 | 8.74E-06 | 0 | 8.17E-05 | 0 | 4 mo | HOM |
| **1013978665** | 1 | MOp6b | 882 | 0.189342997 | 0 | 0 | 0 | 0 | 2 mo | HOM |
| **1013445609** | 1 | MOp6b | 882 | 0.189342997 | 0 | 0 | 0 | 0 | 4 mo | HOM |
| **1013609015** | 1 | MOp6b | 882 | 0.189342997 | 2.05E-05 | 1 | 0.000108339 | 5.281420569 | 4 mo | HOM |
| **1013608106** | 2 | MOp6b | 882 | 0.189342997 | 0 | 0 | 0 | 0 | 4 mo | HOM |
| **1011937543** | 1 | MOp6b | 882 | 0.189342997 | 0.000133718 | 2 | 0.000706222 | 10.56284114 | 8 mo | HOM |
| **1010419538** | 2 | MOp6b | 882 | 0.189342997 | 0 | 0 | 0 | 0 | 4 mo | WILD |
| **1010419187** | 1 | MOp6b | 882 | 0.189342997 | 3.39E-05 | 2 | 0.000179051 | 10.56284114 | 8 mo | HOM |
| **1013978366** | 1 | MOp6b | 882 | 0.189342997 | 0 | 0 | 0 | 0 | 2 mo | HOM |
| **1014416155** | 2 | MOp6b | 882 | 0.189342997 | 0 | 0 | 0 | 0 | 2 mo | HOM |
| **1013978065** | 2 | MOp6b | 882 | 0.189342997 | 0 | 0 | 0 | 0 | 2 mo | HOM |
| **1009934102** | 2 | MOp6b | 882 | 0.189342997 | 0 | 0 | 0 | 0 | 4 mo | WILD |
| **1009322937** | 2 | MOp6b | 882 | 0.189342997 | 2.71E-06 | 0 | 1.43E-05 | 0 | 8 mo | WILD |
| **1011080637** | 2 | MOp6b | 882 | 0.189342997 | 2.06E-05 | 2 | 0.000109015 | 10.56284114 | 8 mo | HOM |
| **1013609713** | 2 | MOp6b | 882 | 0.189342997 | 0 | 0 | 0 | 0 | 2 mo | WILD |
| **1010418842** | 1 | MOp6b | 882 | 0.189342997 | 0.000189305 | 5 | 0.000999797 | 26.40710284 | 8 mo | HOM |
| **1013288213** | 2 | MOp6b | 882 | 0.189342997 | 0 | 0 | 0 | 0 | 4 mo | HOM |
| **1008976060** | 2 | MOp6b | 882 | 0.189342997 | 3.03E-06 | 1 | 1.6E-05 | 5.281420569 | 8 mo | WILD |
| **1013446243** | 1 | MOp6b | 882 | 0.189342997 | 4.39E-06 | 1 | 2.32E-05 | 5.281420569 | 4 mo | HOM |
| **1013288951** | 2 | MOp6b | 882 | 0.189342997 | 4.3E-06 | 0 | 2.27E-05 | 0 | 4 mo | HOM |
| **1013978665** | 1 | amc | 884 | 0.168435599 | 0 | 0 | 0 | 0 | 2 mo | HOM |
| **1013445609** | 1 | amc | 884 | 0.168435599 | 4.64E-06 | 0 | 2.75E-05 | 0 | 4 mo | HOM |
| **1013609015** | 1 | amc | 884 | 0.168435599 | 2.4E-05 | 2 | 0.000142636 | 11.87397447 | 4 mo | HOM |
| **1013608106** | 2 | amc | 884 | 0.168435599 | 0 | 0 | 0 | 0 | 4 mo | HOM |
| **1011937543** | 1 | amc | 884 | 0.168435599 | 0.000157027 | 8 | 0.000932269 | 47.49589788 | 8 mo | HOM |
| **1010419538** | 2 | amc | 884 | 0.168435599 | 0 | 0 | 0 | 0 | 4 mo | WILD |
| **1010419187** | 1 | amc | 884 | 0.168435599 | 0.000145303 | 10 | 0.000862664 | 59.36987235 | 8 mo | HOM |
| **1013978366** | 1 | amc | 884 | 0.168435599 | 0 | 0 | 0 | 0 | 2 mo | HOM |
| **1014416155** | 2 | amc | 884 | 0.168435599 | 0 | 0 | 0 | 0 | 2 mo | HOM |
| **1013978065** | 2 | amc | 884 | 0.168435599 | 0 | 0 | 0 | 0 | 2 mo | HOM |
| **1009934102** | 2 | amc | 884 | 0.168435599 | 0 | 0 | 0 | 0 | 4 mo | WILD |
| **1009322937** | 2 | amc | 884 | 0.168435599 | 0 | 0 | 0 | 0 | 8 mo | WILD |
| **1011080637** | 2 | amc | 884 | 0.168435599 | 0.000140623 | 6 | 0.000834875 | 35.62192341 | 8 mo | HOM |
| **1013609713** | 2 | amc | 884 | 0.168435599 | 0 | 0 | 0 | 0 | 2 mo | WILD |
| **1010418842** | 1 | amc | 884 | 0.168435599 | 0.000184598 | 6 | 0.001095953 | 35.62192341 | 8 mo | HOM |
| **1013288213** | 2 | amc | 884 | 0.168435599 | 1.97E-06 | 0 | 1.17E-05 | 0 | 4 mo | HOM |
| **1008976060** | 2 | amc | 884 | 0.168435599 | 0 | 0 | 0 | 0 | 8 mo | WILD |
| **1013446243** | 1 | amc | 884 | 0.168435599 | 6.03E-06 | 1 | 3.58E-05 | 5.936987235 | 4 mo | HOM |
| **1013288951** | 2 | amc | 884 | 0.168435599 | 2.89E-05 | 2 | 0.0001713 | 11.87397447 | 4 mo | HOM |
| **1013978665** | 1 | RSPv | 886 | 4.47855909 | 3.82E-06 | 0 | 8.54E-07 | 0 | 2 mo | HOM |
| **1013445609** | 1 | RSPv | 886 | 4.47855909 | 0.000180606 | 11 | 4.03E-05 | 2.456147118 | 4 mo | HOM |
| **1013609015** | 1 | RSPv | 886 | 4.47855909 | 0.001040745 | 29 | 0.000232384 | 6.475296947 | 4 mo | HOM |
| **1013608106** | 2 | RSPv | 886 | 4.47855909 | 0 | 0 | 0 | 0 | 4 mo | HOM |
| **1011937543** | 1 | RSPv | 886 | 4.47855909 | 0.006045146 | 88 | 0.001349797 | 19.64917694 | 8 mo | HOM |
| **1010419538** | 2 | RSPv | 886 | 4.46473568 | 7.37E-05 | 3 | 1.65E-05 | 0.671932274 | 4 mo | WILD |
| **1010419187** | 1 | RSPv | 886 | 4.47855909 | 0.00727281 | 162 | 0.001623917 | 36.17234846 | 8 mo | HOM |
| **1013978366** | 1 | RSPv | 886 | 4.47855909 | 0.002108033 | 28 | 0.000470695 | 6.252010845 | 2 mo | HOM |
| **1014416155** | 2 | RSPv | 886 | 4.47855909 | 0 | 0 | 0 | 0 | 2 mo | HOM |
| **1013978065** | 2 | RSPv | 886 | 4.47855909 | 0 | 0 | 0 | 0 | 2 mo | HOM |
| **1009934102** | 2 | RSPv | 886 | 4.47855909 | 0.000228485 | 8 | 5.1E-05 | 1.786288813 | 4 mo | WILD |
| **1009322937** | 2 | RSPv | 886 | 4.47855909 | 1.84E-05 | 1 | 4.12E-06 | 0.223286102 | 8 mo | WILD |
| **1011080637** | 2 | RSPv | 886 | 4.47855909 | 0.006719622 | 145 | 0.001500398 | 32.37648473 | 8 mo | HOM |
| **1013609713** | 2 | RSPv | 886 | 4.47855909 | 6.66E-10 | 0 | 1.49E-10 | 0 | 2 mo | WILD |
| **1010418842** | 1 | RSPv | 886 | 4.47855909 | 0.006214479 | 190 | 0.001387607 | 42.42435931 | 8 mo | HOM |
| **1013288213** | 2 | RSPv | 886 | 4.47855909 | 0.000317708 | 11 | 7.09E-05 | 2.456147118 | 4 mo | HOM |
| **1008976060** | 2 | RSPv | 886 | 4.47855909 | 0.000269699 | 11 | 6.02E-05 | 2.456147118 | 8 mo | WILD |
| **1013446243** | 1 | RSPv | 886 | 4.47855909 | 0.000434772 | 19 | 9.71E-05 | 4.242435931 | 4 mo | HOM |
| **1013288951** | 2 | RSPv | 886 | 4.47855909 | 0.00127024 | 18 | 0.000283627 | 4.019149829 | 4 mo | HOM |
| **1013978665** | 1 | PERI2/3 | 888 | 0.343178242 | 0 | 0 | 0 | 0 | 2 mo | HOM |

| | | | | | | | | | | |
|---|---|---|---|---|---|---|---|---|---|---|
| **1013445609** | 1 | PERI2/3 | 888 | 0.343178242 | 0 | 0 | 0 | 0 | 4 mo | HOM |
| **1013609015** | 1 | PERI2/3 | 888 | 0.343178242 | 3.3E-05 | 3 | 9.6E-05 | 8.74181294 | 4 mo | HOM |
| **1013608106** | 2 | PERI2/3 | 888 | 0.343178242 | 0 | 0 | 0 | 0 | 4 mo | HOM |
| **1011937543** | 1 | PERI2/3 | 888 | 0.343178242 | 0.000300724 | 12 | 0.00087629 | 34.96725176 | 8 mo | HOM |
| **1010419538** | 2 | PERI2/3 | 888 | 0.343178242 | 0 | 0 | 0 | 0 | 4 mo | WILD |
| **1010419187** | 1 | PERI2/3 | 888 | 0.343178242 | 0.00026055 | 9 | 0.000759227 | 26.22543882 | 8 mo | HOM |
| **1013978366** | 1 | PERI2/3 | 888 | 0.343178242 | 0 | 0 | 0 | 0 | 2 mo | HOM |
| **1014416155** | 2 | PERI2/3 | 888 | 0.343178242 | 0 | 0 | 0 | 0 | 2 mo | HOM |
| **1013978065** | 2 | PERI2/3 | 888 | 0.343178242 | 0 | 0 | 0 | 0 | 2 mo | HOM |
| **1009934102** | 2 | PERI2/3 | 888 | 0.343178242 | 0 | 0 | 0 | 0 | 4 mo | WILD |
| **1009322937** | 2 | PERI2/3 | 888 | 0.343178242 | 0 | 0 | 0 | 0 | 8 mo | WILD |
| **1011080637** | 2 | PERI2/3 | 888 | 0.343178242 | 0.000145593 | 9 | 0.000424248 | 26.22543882 | 8 mo | HOM |
| **1013609713** | 2 | PERI2/3 | 888 | 0.343178242 | 0 | 0 | 0 | 0 | 2 mo | WILD |
| **1010418842** | 1 | PERI2/3 | 888 | 0.343178242 | 0.000306406 | 13 | 0.000892848 | 37.8811894 | 8 mo | HOM |
| **1013288213** | 2 | PERI2/3 | 888 | 0.343178242 | 6.88E-07 | 0 | 2E-06 | 0 | 4 mo | HOM |
| **1008976060** | 2 | PERI2/3 | 888 | 0.343178242 | 0 | 0 | 0 | 0 | 8 mo | WILD |
| **1013446243** | 1 | PERI2/3 | 888 | 0.343178242 | 0 | 0 | 0 | 0 | 4 mo | HOM |
| **1013288951** | 2 | PERI2/3 | 888 | 0.343178242 | 2.84E-05 | 2 | 8.27E-05 | 5.827875293 | 4 mo | HOM |
| **1013978665** | 1 | SSp-n6a | 889 | 0.829133312 | 0 | 0 | 0 | 0 | 2 mo | HOM |
| **1013445609** | 1 | SSp-n6a | 889 | 0.829133312 | 8.32E-05 | 6 | 0.000100358 | 7.236472004 | 4 mo | HOM |
| **1013609015** | 1 | SSp-n6a | 889 | 0.829133312 | 0.00020708 | 6 | 0.000249755 | 7.236472004 | 4 mo | HOM |
| **1013608106** | 2 | SSp-n6a | 889 | 0.829133312 | 0 | 0 | 0 | 0 | 4 mo | HOM |
| **1011937543** | 1 | SSp-n6a | 889 | 0.829133312 | 0.001049749 | 27 | 0.00126608 | 32.56412402 | 8 mo | HOM |
| **1010419538** | 2 | SSp-n6a | 889 | 0.829133312 | 0 | 0 | 0 | 0 | 4 mo | WILD |
| **1010419187** | 1 | SSp-n6a | 889 | 0.829133312 | 0.001107046 | 57 | 0.001335185 | 68.74648404 | 8 mo | HOM |
| **1013978366** | 1 | SSp-n6a | 889 | 0.829133312 | 4.2E-05 | 1 | 5.07E-05 | 1.206078667 | 2 mo | HOM |
| **1014416155** | 2 | SSp-n6a | 889 | 0.829133312 | 0 | 0 | 0 | 0 | 2 mo | HOM |
| **1013978065** | 2 | SSp-n6a | 889 | 0.829133312 | 0 | 0 | 0 | 0 | 2 mo | HOM |
| **1009934102** | 2 | SSp-n6a | 889 | 0.829133312 | 0 | 0 | 0 | 0 | 4 mo | WILD |
| **1009322937** | 2 | SSp-n6a | 889 | 0.829133312 | 0 | 0 | 0 | 0 | 8 mo | WILD |
| **1011080637** | 2 | SSp-n6a | 889 | 0.829133312 | 0.000924523 | 33 | 0.001115047 | 39.80059602 | 8 mo | HOM |
| **1013609713** | 2 | SSp-n6a | 889 | 0.829133312 | 0 | 0 | 0 | 0 | 2 mo | WILD |
| **1010418842** | 1 | SSp-n6a | 889 | 0.829133312 | 0.001246253 | 52 | 0.001503079 | 62.7160907 | 8 mo | HOM |
| **1013288213** | 2 | SSp-n6a | 889 | 0.829133312 | 4.61E-05 | 3 | 5.56E-05 | 3.618236002 | 4 mo | HOM |
| **1008976060** | 2 | SSp-n6a | 889 | 0.829133312 | 6.76E-06 | 1 | 8.16E-06 | 1.206078667 | 8 mo | WILD |
| **1013446243** | 1 | SSp-n6a | 889 | 0.829133312 | 0.000249908 | 15 | 0.000301409 | 18.09118001 | 4 mo | HOM |
| **1013288951** | 2 | SSp-n6a | 889 | 0.829133312 | 0.000196081 | 8 | 0.000236489 | 9.648629339 | 4 mo | HOM |
| **1013978665** | 1 | SSs6b | 893 | 0.214323883 | 0 | 0 | 0 | 0 | 2 mo | HOM |
| **1013445609** | 1 | SSs6b | 893 | 0.214323883 | 1.48E-05 | 2 | 6.9E-05 | 9.331671161 | 4 mo | HOM |
| **1013609015** | 1 | SSs6b | 893 | 0.214323883 | 2.18E-05 | 2 | 0.000101648 | 9.331671161 | 4 mo | HOM |
| **1013608106** | 2 | SSs6b | 893 | 0.214323883 | 0 | 0 | 0 | 0 | 4 mo | HOM |
| **1011937543** | 1 | SSs6b | 893 | 0.214323883 | 0.00024232 | 7 | 0.001130627 | 32.66084906 | 8 mo | HOM |
| **1010419538** | 2 | SSs6b | 893 | 0.214323883 | 0 | 0 | 0 | 0 | 4 mo | WILD |
| **1010419187** | 1 | SSs6b | 893 | 0.214323883 | 0.000152848 | 11 | 0.000713163 | 51.32419139 | 8 mo | HOM |
| **1013978366** | 1 | SSs6b | 893 | 0.214323883 | 0 | 0 | 0 | 0 | 2 mo | HOM |
| **1014416155** | 2 | SSs6b | 893 | 0.214323883 | 0 | 0 | 0 | 0 | 2 mo | HOM |
| **1013978065** | 2 | SSs6b | 893 | 0.214323883 | 0 | 0 | 0 | 0 | 2 mo | HOM |
| **1009934102** | 2 | SSs6b | 893 | 0.214323883 | 0 | 0 | 0 | 0 | 4 mo | WILD |
| **1009322937** | 2 | SSs6b | 893 | 0.214323883 | 0 | 0 | 0 | 0 | 8 mo | WILD |
| **1011080637** | 2 | SSs6b | 893 | 0.214323883 | 0.000105508 | 8 | 0.000492285 | 37.32668464 | 8 mo | HOM |
| **1013609713** | 2 | SSs6b | 893 | 0.214323883 | 0 | 0 | 0 | 0 | 2 mo | WILD |
| **1010418842** | 1 | SSs6b | 893 | 0.214323883 | 0.000204905 | 10 | 0.000956051 | 46.65835581 | 8 mo | HOM |
| **1013288213** | 2 | SSs6b | 893 | 0.214323883 | 5.91E-06 | 1 | 2.76E-05 | 4.665835581 | 4 mo | HOM |
| **1008976060** | 2 | SSs6b | 893 | 0.214323883 | 3.38E-06 | 1 | 1.58E-05 | 4.665835581 | 8 mo | WILD |
| **1013446243** | 1 | SSs6b | 893 | 0.214323883 | 2.16E-05 | 0 | 0.000100686 | 0 | 4 mo | HOM |
| **1013288951** | 2 | SSs6b | 893 | 0.214323883 | 0 | 0 | 0 | 0 | 4 mo | HOM |
| **1013978665** | 1 | RSPagl | 894 | 2.428326842 | 3.45E-05 | 3 | 1.42E-05 | 1.235418539 | 2 mo | HOM |
| **1013445609** | 1 | RSPagl | 894 | 2.428326842 | 0.000212268 | 12 | 8.74E-05 | 4.941674157 | 4 mo | HOM |
| **1013609015** | 1 | RSPagl | 894 | 2.428326842 | 0.001036478 | 29 | 0.000426828 | 11.94237921 | 4 mo | HOM |

| | | | | | | | | | | |
|---|---|---|---|---|---|---|---|---|---|---|
| **1013608106** | 2 | RSPagl | 894 | 2.428326842 | 0 | 0 | 0 | 0 | 4 mo | HOM |
| **1011937543** | 1 | RSPagl | 894 | 2.428326842 | 0.004578766 | 42 | 0.001885564 | 17.29585955 | 8 mo | HOM |
| **1010419538** | 2 | RSPagl | 894 | 2.428326842 | 0.000108815 | 7 | 4.48E-05 | 2.882643258 | 4 mo | WILD |
| **1010419187** | 1 | RSPagl | 894 | 2.428326842 | 0.010381666 | 143 | 0.004275234 | 58.88828371 | 8 mo | HOM |
| **1013978366** | 1 | RSPagl | 894 | 2.428326842 | 2.8E-06 | 0 | 1.15E-06 | 0 | 2 mo | HOM |
| **1014416155** | 2 | RSPagl | 894 | 2.428326842 | 0 | 0 | 0 | 0 | 2 mo | HOM |
| **1013978065** | 2 | RSPagl | 894 | 2.428326842 | 8.63E-09 | 0 | 3.56E-09 | 0 | 2 mo | HOM |
| **1009934102** | 2 | RSPagl | 894 | 2.428326842 | 7.74E-05 | 3 | 3.19E-05 | 1.235418539 | 4 mo | WILD |
| **1009322937** | 2 | RSPagl | 894 | 2.428326842 | 1.56E-05 | 1 | 6.41E-06 | 0.41180618 | 8 mo | WILD |
| **1011080637** | 2 | RSPagl | 894 | 2.428326842 | 0.002141518 | 84 | 0.00088189 | 34.5917191 | 8 mo | HOM |
| **1013609713** | 2 | RSPagl | 894 | 2.428326842 | 3.83E-05 | 2 | 1.58E-05 | 0.82361236 | 2 mo | WILD |
| **1010418842** | 1 | RSPagl | 894 | 2.428326842 | 0.002957364 | 96 | 0.001217861 | 39.53339326 | 8 mo | HOM |
| **1013288213** | 2 | RSPagl | 894 | 2.428326842 | 0.000214148 | 15 | 8.82E-05 | 6.177092697 | 4 mo | HOM |
| **1008976060** | 2 | RSPagl | 894 | 2.428326842 | 0.000319419 | 12 | 0.000131539 | 4.941674157 | 8 mo | WILD |
| **1013446243** | 1 | RSPagl | 894 | 2.428326842 | 0.000723593 | 33 | 0.00029798 | 13.58960393 | 4 mo | HOM |
| **1013288951** | 2 | RSPagl | 894 | 2.428326842 | 0.001708654 | 35 | 0.000703634 | 14.41321629 | 4 mo | HOM |
| **1013978665** | 1 | ECT | 895 | 1.763990386 | 1.85E-06 | 0 | 1.05E-06 | 0 | 2 mo | HOM |
| **1013445609** | 1 | ECT | 895 | 1.763990386 | 9.35E-05 | 10 | 5.3E-05 | 5.668965136 | 4 mo | HOM |
| **1013609015** | 1 | ECT | 895 | 1.763990386 | 0.000450069 | 11 | 0.000255143 | 6.23586165 | 4 mo | HOM |
| **1013608106** | 2 | ECT | 895 | 1.763990386 | 0 | 0 | 0 | 0 | 4 mo | HOM |
| **1011937543** | 1 | ECT | 895 | 1.763990386 | 0.004755285 | 58 | 0.002695754 | 32.87999779 | 8 mo | HOM |
| **1010419538** | 2 | ECT | 895 | 1.763990386 | 1.91E-06 | 1 | 1.08E-06 | 0.566896514 | 4 mo | WILD |
| **1010419187** | 1 | ECT | 895 | 1.763990386 | 0.003039432 | 102 | 0.001723043 | 57.82344439 | 8 mo | HOM |
| **1013978366** | 1 | ECT | 895 | 1.763990386 | 3.73E-06 | 0 | 2.12E-06 | 0 | 2 mo | HOM |
| **1014416155** | 2 | ECT | 895 | 1.763990386 | 0 | 0 | 0 | 0 | 2 mo | HOM |
| **1013978065** | 2 | ECT | 895 | 1.763990386 | 0 | 0 | 0 | 0 | 2 mo | HOM |
| **1009934102** | 2 | ECT | 895 | 1.763990386 | 7.23E-06 | 0 | 4.1E-06 | 0 | 4 mo | WILD |
| **1009322937** | 2 | ECT | 895 | 1.763990386 | 6.22E-07 | 0 | 3.53E-07 | 0 | 8 mo | WILD |
| **1011080637** | 2 | ECT | 895 | 1.763990386 | 0.001269174 | 55 | 0.00071949 | 31.17930825 | 8 mo | HOM |
| **1013609713** | 2 | ECT | 895 | 1.763990386 | 0 | 0 | 0 | 0 | 2 mo | WILD |
| **1010418842** | 1 | ECT | 895 | 1.763990386 | 0.002104425 | 94 | 0.001192991 | 53.28827228 | 8 mo | HOM |
| **1013288213** | 2 | ECT | 895 | 1.763990386 | 0.000140464 | 9 | 7.96E-05 | 5.102068623 | 4 mo | HOM |
| **1008976060** | 2 | ECT | 895 | 1.763990386 | 4.81E-05 | 3 | 2.73E-05 | 1.700689541 | 8 mo | WILD |
| **1013446243** | 1 | ECT | 895 | 1.763990386 | 0.000328324 | 18 | 0.000186126 | 10.20413725 | 4 mo | HOM |
| **1013288951** | 2 | ECT | 895 | 1.763990386 | 0.000222059 | 13 | 0.000125885 | 7.369654677 | 4 mo | HOM |
| **1013978665** | 1 | lfbst | 896 | 2.440419667 | 0 | 0 | 0 | 0 | 2 mo | HOM |
| **1013445609** | 1 | lfbst | 896 | 2.440419667 | 2.28E-05 | 2 | 9.34E-06 | 0.819531176 | 4 mo | HOM |
| **1013609015** | 1 | lfbst | 896 | 2.440419667 | 2.93E-05 | 1 | 1.2E-05 | 0.409765588 | 4 mo | HOM |
| **1013608106** | 2 | lfbst | 896 | 2.440419667 | 0 | 0 | 0 | 0 | 4 mo | HOM |
| **1011937543** | 1 | lfbst | 896 | 2.440419667 | 0.002197648 | 32 | 0.00090052 | 13.11249882 | 8 mo | HOM |
| **1010419538** | 2 | lfbst | 896 | 2.440419667 | 0 | 0 | 0 | 0 | 4 mo | WILD |
| **1010419187** | 1 | lfbst | 896 | 2.440419667 | 0.000519294 | 28 | 0.000212789 | 11.47343647 | 8 mo | HOM |
| **1013978366** | 1 | lfbst | 896 | 2.440419667 | 0 | 0 | 0 | 0 | 2 mo | HOM |
| **1014416155** | 2 | lfbst | 896 | 2.440419667 | 0 | 0 | 0 | 0 | 2 mo | HOM |
| **1013978065** | 2 | lfbst | 896 | 2.440419667 | 0 | 0 | 0 | 0 | 2 mo | HOM |
| **1009934102** | 2 | lfbst | 896 | 2.440419667 | 0 | 0 | 0 | 0 | 4 mo | WILD |
| **1009322937** | 2 | lfbst | 896 | 2.440419667 | 1.3E-06 | 0 | 5.34E-07 | 0 | 8 mo | WILD |
| **1011080637** | 2 | lfbst | 896 | 2.440419667 | 0.000402117 | 24 | 0.000164774 | 9.834374114 | 8 mo | HOM |
| **1013609713** | 2 | lfbst | 896 | 2.440419667 | 0 | 0 | 0 | 0 | 2 mo | WILD |
| **1010418842** | 1 | lfbst | 896 | 2.440419667 | 0.001075291 | 42 | 0.000440617 | 17.2101547 | 8 mo | HOM |
| **1013288213** | 2 | lfbst | 896 | 2.440419667 | 6.76E-06 | 1 | 2.77E-06 | 0.409765588 | 4 mo | HOM |
| **1008976060** | 2 | lfbst | 896 | 2.440419667 | 3.28E-05 | 5 | 1.34E-05 | 2.04882794 | 8 mo | WILD |
| **1013446243** | 1 | lfbst | 896 | 2.440419667 | 5.9E-05 | 2 | 2.42E-05 | 0.819531176 | 4 mo | HOM |
| **1013288951** | 2 | lfbst | 896 | 2.440419667 | 0.000147732 | 1 | 6.05E-05 | 0.409765588 | 4 mo | HOM |
| **1013978665** | 1 | VISC1 | 897 | 0.345197474 | 0 | 0 | 0 | 0 | 2 mo | HOM |
| **1013445609** | 1 | VISC1 | 897 | 0.345197474 | 3.39E-05 | 2 | 9.83E-05 | 5.793785161 | 4 mo | HOM |
| **1013609015** | 1 | VISC1 | 897 | 0.345197474 | 2.95E-05 | 2 | 8.54E-05 | 5.793785161 | 4 mo | HOM |
| **1013608106** | 2 | VISC1 | 897 | 0.345197474 | 0 | 0 | 0 | 0 | 4 mo | HOM |
| **1011937543** | 1 | VISC1 | 897 | 0.345197474 | 0.000626004 | 14 | 0.001813467 | 40.55649612 | 8 mo | HOM |

| | | | | | | | | | | |
|---|---|---|---|---|---|---|---|---|---|---|
| **1010419538** | 2 | VISC1 | 897 | 0.345197474 | 0 | 0 | 0 | 0 | 4 mo | WILD |
| **1010419187** | 1 | VISC1 | 897 | 0.345197474 | 0.000428485 | 17 | 0.001241274 | 49.24717386 | 8 mo | HOM |
| **1013978366** | 1 | VISC1 | 897 | 0.345197474 | 6.77E-06 | 1 | 1.96E-05 | 2.89689258 | 2 mo | HOM |
| **1014416155** | 2 | VISC1 | 897 | 0.345197474 | 0 | 0 | 0 | 0 | 2 mo | HOM |
| **1013978065** | 2 | VISC1 | 897 | 0.345197474 | 0 | 0 | 0 | 0 | 2 mo | HOM |
| **1009934102** | 2 | VISC1 | 897 | 0.345197474 | 0 | 0 | 0 | 0 | 4 mo | WILD |
| **1009322937** | 2 | VISC1 | 897 | 0.345197474 | 1.7E-06 | 0 | 4.91E-06 | 0 | 8 mo | WILD |
| **1011080637** | 2 | VISC1 | 897 | 0.345197474 | 0.000278988 | 11 | 0.000808199 | 31.86581838 | 8 mo | HOM |
| **1013609713** | 2 | VISC1 | 897 | 0.345197474 | 6.41E-07 | 0 | 1.86E-06 | 0 | 2 mo | WILD |
| **1010418842** | 1 | VISC1 | 897 | 0.345197474 | 0.000821226 | 19 | 0.002379002 | 55.04095903 | 8 mo | HOM |
| **1013288213** | 2 | VISC1 | 897 | 0.345197474 | 7.98E-06 | 1 | 2.31E-05 | 2.89689258 | 4 mo | HOM |
| **1008976060** | 2 | VISC1 | 897 | 0.345197474 | 4.27E-06 | 0 | 1.24E-05 | 0 | 8 mo | WILD |
| **1013446243** | 1 | VISC1 | 897 | 0.345197474 | 0.00012665 | 5 | 0.000366892 | 14.4844629 | 4 mo | HOM |
| **1013288951** | 2 | VISC1 | 897 | 0.345197474 | 0.000206069 | 11 | 0.000596961 | 31.86581838 | 4 mo | HOM |
| **1013978665** | 1 | PCG | 898 | 0.557786482 | 2.88E-05 | 2 | 5.16E-05 | 3.585601414 | 2 mo | HOM |
| **1013445609** | 1 | PCG | 898 | 0.557786482 | 0 | 0 | 0 | 0 | 4 mo | HOM |
| **1013609015** | 1 | PCG | 898 | 0.557786482 | 0.00171064 | 10 | 0.003066837 | 17.92800707 | 4 mo | HOM |
| **1013608106** | 2 | PCG | 898 | 0.557786482 | 0 | 0 | 0 | 0 | 4 mo | HOM |
| **1011937543** | 1 | PCG | 898 | 0.557786482 | 0.000287613 | 1 | 0.000515633 | 1.792800707 | 8 mo | HOM |
| **1010419538** | 2 | PCG | 898 | 0.557786482 | 0 | 0 | 0 | 0 | 4 mo | WILD |
| **1010419187** | 1 | PCG | 898 | 0.557786482 | 0.000918595 | 20 | 0.001646858 | 35.85601414 | 8 mo | HOM |
| **1013978366** | 1 | PCG | 898 | 0.557786482 | 2.47E-05 | 2 | 4.43E-05 | 3.585601414 | 2 mo | HOM |
| **1014416155** | 2 | PCG | 898 | 0.557786482 | 0 | 0 | 0 | 0 | 2 mo | HOM |
| **1013978065** | 2 | PCG | 898 | 0.557786482 | 0 | 0 | 0 | 0 | 2 mo | HOM |
| **1009934102** | 2 | PCG | 898 | 0.557786482 | 9.29E-06 | 0 | 1.66E-05 | 0 | 4 mo | WILD |
| **1009322937** | 2 | PCG | 898 | 0.557786482 | 5.77E-05 | 6 | 0.000103489 | 10.75680424 | 8 mo | WILD |
| **1011080637** | 2 | PCG | 898 | 0.557786482 | 0.002180286 | 13 | 0.003908819 | 23.30640919 | 8 mo | HOM |
| **1013609713** | 2 | PCG | 898 | 0.557786482 | 1.32E-05 | 0 | 2.38E-05 | 0 | 2 mo | WILD |
| **1010418842** | 1 | PCG | 898 | 0.557786482 | 0.001009692 | 5 | 0.001810177 | 8.964003535 | 8 mo | HOM |
| **1013288213** | 2 | PCG | 898 | 0.557786482 | 4.77E-06 | 1 | 8.56E-06 | 1.792800707 | 4 mo | HOM |
| **1008976060** | 2 | PCG | 898 | 0.557786482 | 0.001763443 | 26 | 0.003161501 | 46.61281838 | 8 mo | WILD |
| **1013446243** | 1 | PCG | 898 | 0.557786482 | 9.68E-05 | 8 | 0.000173582 | 14.34240566 | 4 mo | HOM |
| **1013288951** | 2 | PCG | 898 | 0.557786482 | 0.000210979 | 13 | 0.000378244 | 23.30640919 | 4 mo | HOM |
| **1013978665** | 1 | aco | 900 | 0.783098714 | 8.88E-08 | 0 | 1.13E-07 | 0 | 2 mo | HOM |
| **1013445609** | 1 | aco | 900 | 0.783098714 | 3.13E-07 | 0 | 4E-07 | 0 | 4 mo | HOM |
| **1013609015** | 1 | aco | 900 | 0.783098714 | 7.39E-06 | 1 | 9.43E-06 | 1.276978218 | 4 mo | HOM |
| **1013608106** | 2 | aco | 900 | 0.783098714 | 0 | 0 | 0 | 0 | 4 mo | HOM |
| **1011937543** | 1 | aco | 900 | 0.783098714 | 0.000347747 | 10 | 0.000444065 | 12.76978218 | 8 mo | HOM |
| **1010419538** | 2 | aco | 900 | 0.783098714 | 0 | 0 | 0 | 0 | 4 mo | WILD |
| **1010419187** | 1 | aco | 900 | 0.783098714 | 2.91E-05 | 2 | 3.72E-05 | 2.553956436 | 8 mo | HOM |
| **1013978366** | 1 | aco | 900 | 0.783098714 | 0 | 0 | 0 | 0 | 2 mo | HOM |
| **1014416155** | 2 | aco | 900 | 0.783098714 | 0 | 0 | 0 | 0 | 2 mo | HOM |
| **1013978065** | 2 | aco | 900 | 0.783098714 | 0 | 0 | 0 | 0 | 2 mo | HOM |
| **1009934102** | 2 | aco | 900 | 0.783098714 | 0 | 0 | 0 | 0 | 4 mo | WILD |
| **1009322937** | 2 | aco | 900 | 0.783098714 | 0 | 0 | 0 | 0 | 8 mo | WILD |
| **1011080637** | 2 | aco | 900 | 0.783098714 | 1.71E-05 | 1 | 2.18E-05 | 1.276978218 | 8 mo | HOM |
| **1013609713** | 2 | aco | 900 | 0.783098714 | 0 | 0 | 0 | 0 | 2 mo | WILD |
| **1010418842** | 1 | aco | 900 | 0.783098714 | 4.74E-05 | 1 | 6.05E-05 | 1.276978218 | 8 mo | HOM |
| **1013288213** | 2 | aco | 900 | 0.783098714 | 0 | 0 | 0 | 0 | 4 mo | HOM |
| **1008976060** | 2 | aco | 900 | 0.783098714 | 1.22E-05 | 1 | 1.56E-05 | 1.276978218 | 8 mo | WILD |
| **1013446243** | 1 | aco | 900 | 0.783098714 | 1.59E-06 | 0 | 2.03E-06 | 0 | 4 mo | HOM |
| **1013288951** | 2 | aco | 900 | 0.783098714 | 3.3E-06 | 0 | 4.21E-06 | 0 | 4 mo | HOM |
| **1013978665** | 1 | Vn | 901 | 2.449456708 | 0 | 0 | 0 | 0 | 2 mo | HOM |
| **1013445609** | 1 | Vn | 901 | 2.449456708 | 0 | 0 | 0 | 0 | 4 mo | HOM |
| **1013609015** | 1 | Vn | 901 | 2.449456708 | 0 | 0 | 0 | 0 | 4 mo | HOM |
| **1013608106** | 2 | Vn | 901 | 2.449456708 | 0 | 0 | 0 | 0 | 4 mo | HOM |
| **1011937543** | 1 | Vn | 901 | 2.449456708 | 0.004559896 | 6 | 0.001861595 | 2.449522778 | 8 mo | HOM |
| **1010419538** | 2 | Vn | 901 | 2.449456708 | 0 | 0 | 0 | 0 | 4 mo | WILD |
| **1010419187** | 1 | Vn | 901 | 2.449456708 | 0.000807922 | 1 | 0.000329837 | 0.408253796 | 8 mo | HOM |

| | | | | | | | | | | |
|---|---|---|---|---|---|---|---|---|---|---|
| **1013978366** | 1 | Vn | 901 | 2.449456708 | 1.01E-05 | 0 | 4.13E-06 | 0 | 2 mo | HOM |
| **1014416155** | 2 | Vn | 901 | 2.449456708 | 0 | 0 | 0 | 0 | 2 mo | HOM |
| **1013978065** | 2 | Vn | 901 | 2.449456708 | 0 | 0 | 0 | 0 | 2 mo | HOM |
| **1009934102** | 2 | Vn | 901 | 2.449456708 | 1.95E-09 | 0 | 7.95E-10 | 0 | 4 mo | WILD |
| **1009322937** | 2 | Vn | 901 | 2.449456708 | 0 | 0 | 0 | 0 | 8 mo | WILD |
| **1011080637** | 2 | Vn | 901 | 2.449456708 | 0 | 0 | 0 | 0 | 8 mo | HOM |
| **1013609713** | 2 | Vn | 901 | 2.449456708 | 0 | 0 | 0 | 0 | 2 mo | WILD |
| **1010418842** | 1 | Vn | 901 | 2.449456708 | 4.1E-07 | 0 | 1.68E-07 | 0 | 8 mo | HOM |
| **1013288213** | 2 | Vn | 901 | 2.449456708 | 0 | 0 | 0 | 0 | 4 mo | HOM |
| **1008976060** | 2 | Vn | 901 | 2.449456708 | 3.49E-05 | 4 | 1.42E-05 | 1.633015186 | 8 mo | WILD |
| **1013446243** | 1 | Vn | 901 | 2.449456708 | 2.71E-07 | 0 | 1.11E-07 | 0 | 4 mo | HOM |
| **1013288951** | 2 | Vn | 901 | 2.449456708 | 3.91E-08 | 0 | 1.6E-08 | 0 | 4 mo | HOM |
| **1013978665** | 1 | VISpl5 | 902 | 0.250150998 | 0 | 0 | 0 | 0 | 2 mo | HOM |
| **1013445609** | 1 | VISpl5 | 902 | 0.250150998 | 4.3E-05 | 4 | 0.000171816 | 15.99034199 | 4 mo | HOM |
| **1013609015** | 1 | VISpl5 | 902 | 0.250150998 | 8.66E-05 | 5 | 0.000346119 | 19.98792748 | 4 mo | HOM |
| **1013608106** | 2 | VISpl5 | 902 | 0.250150998 | 0 | 0 | 0 | 0 | 4 mo | HOM |
| **1011937543** | 1 | VISpl5 | 902 | 0.250150998 | 0.000114775 | 3 | 0.000458824 | 11.99275649 | 8 mo | HOM |
| **1010419538** | 2 | VISpl5 | 902 | 0.250150998 | 0 | 0 | 0 | 0 | 4 mo | WILD |
| **1010419187** | 1 | VISpl5 | 902 | 0.250150998 | 0.00043669 | 12 | 0.001745704 | 47.97102596 | 8 mo | HOM |
| **1013978366** | 1 | VISpl5 | 902 | 0.250150998 | 0 | 0 | 0 | 0 | 2 mo | HOM |
| **1014416155** | 2 | VISpl5 | 902 | 0.250150998 | 0 | 0 | 0 | 0 | 2 mo | HOM |
| **1013978065** | 2 | VISpl5 | 902 | 0.250150998 | 0 | 0 | 0 | 0 | 2 mo | HOM |
| **1009934102** | 2 | VISpl5 | 902 | 0.250150998 | 0 | 0 | 0 | 0 | 4 mo | WILD |
| **1009322937** | 2 | VISpl5 | 902 | 0.250150998 | 0 | 0 | 0 | 0 | 8 mo | WILD |
| **1011080637** | 2 | VISpl5 | 902 | 0.250150998 | 0.000196749 | 11 | 0.00078652 | 43.97344046 | 8 mo | HOM |
| **1013609713** | 2 | VISpl5 | 902 | 0.250150998 | 0 | 0 | 0 | 0 | 2 mo | WILD |
| **1010418842** | 1 | VISpl5 | 902 | 0.250150998 | 0.000285821 | 7 | 0.001142595 | 27.98309848 | 8 mo | HOM |
| **1013288213** | 2 | VISpl5 | 902 | 0.250150998 | 8.78E-06 | 0 | 3.51E-05 | 0 | 4 mo | HOM |
| **1008976060** | 2 | VISpl5 | 902 | 0.250150998 | 0 | 0 | 0 | 0 | 8 mo | WILD |
| **1013446243** | 1 | VISpl5 | 902 | 0.250150998 | 5.16E-05 | 3 | 0.000206423 | 11.99275649 | 4 mo | HOM |
| **1013288951** | 2 | VISpl5 | 902 | 0.250150998 | 5.32E-06 | 1 | 2.13E-05 | 3.997585497 | 4 mo | HOM |
| **1013978665** | 1 | ECU | 903 | 0.210911778 | 0 | 0 | 0 | 0 | 2 mo | HOM |
| **1013445609** | 1 | ECU | 903 | 0.210911778 | 0 | 0 | 0 | 0 | 4 mo | HOM |
| **1013609015** | 1 | ECU | 903 | 0.210911778 | 0 | 0 | 0 | 0 | 4 mo | HOM |
| **1013608106** | 2 | ECU | 903 | 0.210911778 | 0 | 0 | 0 | 0 | 4 mo | HOM |
| **1011937543** | 1 | ECU | 903 | 0.210911778 | 0.000200013 | 0 | 0.000948324 | 0 | 8 mo | HOM |
| **1010419538** | 2 | ECU | 903 | 0.210911778 | 0 | 0 | 0 | 0 | 4 mo | WILD |
| **1010419187** | 1 | ECU | 903 | 0.210911778 | 0 | 0 | 0 | 0 | 8 mo | HOM |
| **1013978366** | 1 | ECU | 903 | 0.210911778 | 0 | 0 | 0 | 0 | 2 mo | HOM |
| **1014416155** | 2 | ECU | 903 | 0.210911778 | 0 | 0 | 0 | 0 | 2 mo | HOM |
| **1013978065** | 2 | ECU | 903 | 0.210911778 | 0 | 0 | 0 | 0 | 2 mo | HOM |
| **1009934102** | 2 | ECU | 903 | 0.210911778 | 0 | 0 | 0 | 0 | 4 mo | WILD |
| **1009322937** | 2 | ECU | 903 | 0.210911778 | 0 | 0 | 0 | 0 | 8 mo | WILD |
| **1011080637** | 2 | ECU | 903 | 0.210911778 | 0 | 0 | 0 | 0 | 8 mo | HOM |
| **1013609713** | 2 | ECU | 903 | 0.210911778 | 0 | 0 | 0 | 0 | 2 mo | WILD |
| **1010418842** | 1 | ECU | 903 | 0.210911778 | 0 | 0 | 0 | 0 | 8 mo | HOM |
| **1013288213** | 2 | ECU | 903 | 0.210911778 | 0 | 0 | 0 | 0 | 4 mo | HOM |
| **1008976060** | 2 | ECU | 903 | 0.210911778 | 5.55E-06 | 1 | 2.63E-05 | 4.741318903 | 8 mo | WILD |
| **1013446243** | 1 | ECU | 903 | 0.210911778 | 9.26E-06 | 1 | 4.39E-05 | 4.741318903 | 4 mo | HOM |
| **1013288951** | 2 | ECU | 903 | 0.210911778 | 0 | 0 | 0 | 0 | 4 mo | HOM |
| **1013978665** | 1 | MSC | 904 | 1.190012902 | 0 | 0 | 0 | 0 | 2 mo | HOM |
| **1013445609** | 1 | MSC | 904 | 1.190012902 | 0 | 0 | 0 | 0 | 4 mo | HOM |
| **1013609015** | 1 | MSC | 904 | 1.190012902 | 0 | 0 | 0 | 0 | 4 mo | HOM |
| **1013608106** | 2 | MSC | 904 | 1.190012902 | 0 | 0 | 0 | 0 | 4 mo | HOM |
| **1011937543** | 1 | MSC | 904 | 1.190012902 | 0.000573489 | 16 | 0.000481918 | 13.44523237 | 8 mo | HOM |
| **1010419538** | 2 | MSC | 904 | 1.190012902 | 0 | 0 | 0 | 0 | 4 mo | WILD |
| **1010419187** | 1 | MSC | 904 | 1.190012902 | 4.88E-05 | 2 | 4.1E-05 | 1.680654047 | 8 mo | HOM |
| **1013978366** | 1 | MSC | 904 | 1.190012902 | 0 | 0 | 0 | 0 | 2 mo | HOM |
| **1014416155** | 2 | MSC | 904 | 1.190012902 | 0 | 0 | 0 | 0 | 2 mo | HOM |

| | | | | | | | | | | |
|---|---|---|---|---|---|---|---|---|---|---|
| 1013978065 | 2 | MSC | 904 | 1.190012902 | 3.07E-09 | 0 | 2.58E-09 | 0 | 2 mo | HOM |
| 1009934102 | 2 | MSC | 904 | 1.190012902 | 7.99E-07 | 0 | 6.72E-07 | 0 | 4 mo | WILD |
| 1009322937 | 2 | MSC | 904 | 1.190012902 | 2.5E-05 | 1 | 2.1E-05 | 0.840327023 | 8 mo | WILD |
| 1011080637 | 2 | MSC | 904 | 1.190012902 | 5.28E-05 | 4 | 4.43E-05 | 3.361308093 | 8 mo | HOM |
| 1013609713 | 2 | MSC | 904 | 1.190012902 | 0 | 0 | 0 | 0 | 2 mo | WILD |
| 1010418842 | 1 | MSC | 904 | 1.190012902 | 0.000280374 | 10 | 0.000235606 | 8.403270233 | 8 mo | HOM |
| 1013288213 | 2 | MSC | 904 | 1.190012902 | 0 | 0 | 0 | 0 | 4 mo | HOM |
| 1008976060 | 2 | MSC | 904 | 1.190012902 | 2.09E-07 | 0 | 1.75E-07 | 0 | 8 mo | WILD |
| 1013446243 | 1 | MSC | 904 | 1.190012902 | 0 | 0 | 0 | 0 | 4 mo | HOM |
| 1013288951 | 2 | MSC | 904 | 1.190012902 | 0 | 0 | 0 | 0 | 4 mo | HOM |
| 1013978665 | 1 | VISal2/3 | 905 | 0.205338263 | 0 | 0 | 0 | 0 | 2 mo | HOM |
| 1013445609 | 1 | VISal2/3 | 905 | 0.205338263 | 3.72E-06 | 0 | 1.81E-05 | 0 | 4 mo | HOM |
| 1013609015 | 1 | VISal2/3 | 905 | 0.205338263 | 0.000120615 | 3 | 0.000587398 | 14.61003884 | 4 mo | HOM |
| 1013608106 | 2 | VISal2/3 | 905 | 0.205338263 | 0 | 0 | 0 | 0 | 4 mo | HOM |
| 1011937543 | 1 | VISal2/3 | 905 | 0.205338263 | 0.000298381 | 5 | 0.001453118 | 24.35006473 | 8 mo | HOM |
| 1010419538 | 2 | VISal2/3 | 905 | 0.205338263 | 0 | 0 | 0 | 0 | 4 mo | WILD |
| 1010419187 | 1 | VISal2/3 | 905 | 0.205338263 | 0.000306974 | 8 | 0.001494968 | 38.96010356 | 8 mo | HOM |
| 1013978366 | 1 | VISal2/3 | 905 | 0.205338263 | 8.2E-07 | 0 | 4E-06 | 0 | 2 mo | HOM |
| 1014416155 | 2 | VISal2/3 | 905 | 0.205338263 | 0 | 0 | 0 | 0 | 2 mo | HOM |
| 1013978065 | 2 | VISal2/3 | 905 | 0.205338263 | 0 | 0 | 0 | 0 | 2 mo | HOM |
| 1009934102 | 2 | VISal2/3 | 905 | 0.205338263 | 0 | 0 | 0 | 0 | 4 mo | WILD |
| 1009322937 | 2 | VISal2/3 | 905 | 0.205338263 | 0 | 0 | 0 | 0 | 8 mo | WILD |
| 1011080637 | 2 | VISal2/3 | 905 | 0.205338263 | 7.75E-05 | 4 | 0.000377222 | 19.48005178 | 8 mo | HOM |
| 1013609713 | 2 | VISal2/3 | 905 | 0.205338263 | 0 | 0 | 0 | 0 | 2 mo | WILD |
| 1010418842 | 1 | VISal2/3 | 905 | 0.205338263 | 0.00017338 | 8 | 0.000844361 | 38.96010356 | 8 mo | HOM |
| 1013288213 | 2 | VISal2/3 | 905 | 0.205338263 | 1.99E-05 | 2 | 9.71E-05 | 9.740025891 | 4 mo | HOM |
| 1008976060 | 2 | VISal2/3 | 905 | 0.205338263 | 0 | 0 | 0 | 0 | 8 mo | WILD |
| 1013446243 | 1 | VISal2/3 | 905 | 0.205338263 | 6.6E-05 | 2 | 0.000321416 | 9.740025891 | 4 mo | HOM |
| 1013288951 | 2 | VISal2/3 | 905 | 0.205338263 | 0.000110345 | 3 | 0.000537382 | 14.61003884 | 4 mo | HOM |
| 1013978665 | 1 | RSPagl6a | 906 | 0.424697896 | 0 | 0 | 0 | 0 | 2 mo | HOM |
| 1013445609 | 1 | RSPagl6a | 906 | 0.424697896 | 2.2E-05 | 1 | 5.19E-05 | 2.354614914 | 4 mo | HOM |
| 1013609015 | 1 | RSPagl6a | 906 | 0.424697896 | 2.27E-05 | 3 | 5.35E-05 | 7.063844743 | 4 mo | HOM |
| 1013608106 | 2 | RSPagl6a | 906 | 0.424697896 | 0 | 0 | 0 | 0 | 4 mo | HOM |
| 1011937543 | 1 | RSPagl6a | 906 | 0.424697896 | 0.000890632 | 6 | 0.002097096 | 14.12768949 | 8 mo | HOM |
| 1010419538 | 2 | RSPagl6a | 906 | 0.424697896 | 0 | 0 | 0 | 0 | 4 mo | WILD |
| 1010419187 | 1 | RSPagl6a | 906 | 0.424697896 | 0.000881532 | 28 | 0.002075668 | 65.9292176 | 8 mo | HOM |
| 1013978366 | 1 | RSPagl6a | 906 | 0.424697896 | 0 | 0 | 0 | 0 | 2 mo | HOM |
| 1014416155 | 2 | RSPagl6a | 906 | 0.424697896 | 0 | 0 | 0 | 0 | 2 mo | HOM |
| 1013978065 | 2 | RSPagl6a | 906 | 0.424697896 | 0 | 0 | 0 | 0 | 2 mo | HOM |
| 1009934102 | 2 | RSPagl6a | 906 | 0.424697896 | 0 | 0 | 0 | 0 | 4 mo | WILD |
| 1009322937 | 2 | RSPagl6a | 906 | 0.424697896 | 0 | 0 | 0 | 0 | 8 mo | WILD |
| 1011080637 | 2 | RSPagl6a | 906 | 0.424697896 | 0.000301131 | 18 | 0.000709047 | 42.38306846 | 8 mo | HOM |
| 1013609713 | 2 | RSPagl6a | 906 | 0.424697896 | 0 | 0 | 0 | 0 | 2 mo | WILD |
| 1010418842 | 1 | RSPagl6a | 906 | 0.424697896 | 0.000407739 | 15 | 0.000960068 | 35.31922371 | 8 mo | HOM |
| 1013288213 | 2 | RSPagl6a | 906 | 0.424697896 | 1.53E-05 | 2 | 3.6E-05 | 4.709229829 | 4 mo | HOM |
| 1008976060 | 2 | RSPagl6a | 906 | 0.424697896 | 0 | 0 | 0 | 0 | 8 mo | WILD |
| 1013446243 | 1 | RSPagl6a | 906 | 0.424697896 | 0.000121996 | 11 | 0.000287253 | 25.90076406 | 4 mo | HOM |
| 1013288951 | 2 | RSPagl6a | 906 | 0.424697896 | 6.25E-05 | 5 | 0.00014717 | 11.77307457 | 4 mo | HOM |
| 1013978665 | 1 | PCN | 907 | 0.245308935 | 0 | 0 | 0 | 0 | 2 mo | HOM |
| 1013445609 | 1 | PCN | 907 | 0.245308935 | 0 | 0 | 0 | 0 | 4 mo | HOM |
| 1013609015 | 1 | PCN | 907 | 0.245308935 | 0 | 0 | 0 | 0 | 4 mo | HOM |
| 1013608106 | 2 | PCN | 907 | 0.245308935 | 0 | 0 | 0 | 0 | 4 mo | HOM |
| 1011937543 | 1 | PCN | 907 | 0.245308935 | 0.000108466 | 1 | 0.000442161 | 4.076492359 | 8 mo | HOM |
| 1010419538 | 2 | PCN | 907 | 0.245308935 | 0 | 0 | 0 | 0 | 4 mo | WILD |
| 1010419187 | 1 | PCN | 907 | 0.245308935 | 3.64E-05 | 2 | 0.000148386 | 8.152984719 | 8 mo | HOM |
| 1013978366 | 1 | PCN | 907 | 0.245308935 | 0 | 0 | 0 | 0 | 2 mo | HOM |
| 1014416155 | 2 | PCN | 907 | 0.245308935 | 0 | 0 | 0 | 0 | 2 mo | HOM |
| 1013978065 | 2 | PCN | 907 | 0.245308935 | 0 | 0 | 0 | 0 | 2 mo | HOM |
| 1009934102 | 2 | PCN | 907 | 0.245308935 | 0 | 0 | 0 | 0 | 4 mo | WILD |

| | | | | | | | | | | |
|---|---|---|---|---|---|---|---|---|---|---|
| **1009322937** | 2 | PCN | 907 | 0.245308935 | 7.39E-06 | 2 | 3.01E-05 | 8.152984719 | 8 mo | WILD |
| **1011080637** | 2 | PCN | 907 | 0.245308935 | 0.00013465 | 4 | 0.0005489 | 16.30596944 | 8 mo | HOM |
| **1013609713** | 2 | PCN | 907 | 0.245308935 | 0 | 0 | 0 | 0 | 2 mo | WILD |
| **1010418842** | 1 | PCN | 907 | 0.245308935 | 9.62E-05 | 4 | 0.0003921 | 16.30596944 | 8 mo | HOM |
| **1013288213** | 2 | PCN | 907 | 0.245308935 | 0 | 0 | 0 | 0 | 4 mo | HOM |
| **1008976060** | 2 | PCN | 907 | 0.245308935 | 0 | 0 | 0 | 0 | 8 mo | WILD |
| **1013446243** | 1 | PCN | 907 | 0.245308935 | 0 | 0 | 0 | 0 | 4 mo | HOM |
| **1013288951** | 2 | PCN | 907 | 0.245308935 | 0 | 0 | 0 | 0 | 4 mo | HOM |
| **1013978665** | 1 | act | 908 | 0.218465402 | 6.24E-07 | 0 | 2.85E-06 | 0 | 2 mo | HOM |
| **1013445609** | 1 | act | 908 | 0.218465402 | 5.62E-07 | 0 | 2.57E-06 | 0 | 4 mo | HOM |
| **1013609015** | 1 | act | 908 | 0.218465402 | 6.4E-07 | 0 | 2.93E-06 | 0 | 4 mo | HOM |
| **1013608106** | 2 | act | 908 | 0.218465402 | 0 | 0 | 0 | 0 | 4 mo | HOM |
| **1011937543** | 1 | act | 908 | 0.218465402 | 6.15E-05 | 1 | 0.000281374 | 4.577383828 | 8 mo | HOM |
| **1010419538** | 2 | act | 908 | 0.218465402 | 0 | 0 | 0 | 0 | 4 mo | WILD |
| **1010419187** | 1 | act | 908 | 0.218465402 | 3.22E-05 | 1 | 0.000147348 | 4.577383828 | 8 mo | HOM |
| **1013978366** | 1 | act | 908 | 0.218465402 | 0 | 0 | 0 | 0 | 2 mo | HOM |
| **1014416155** | 2 | act | 908 | 0.218465402 | 0 | 0 | 0 | 0 | 2 mo | HOM |
| **1013978065** | 2 | act | 908 | 0.218465402 | 0 | 0 | 0 | 0 | 2 mo | HOM |
| **1009934102** | 2 | act | 908 | 0.218465402 | 1.9E-08 | 0 | 8.71E-08 | 0 | 4 mo | WILD |
| **1009322937** | 2 | act | 908 | 0.218465402 | 0 | 0 | 0 | 0 | 8 mo | WILD |
| **1011080637** | 2 | act | 908 | 0.218465402 | 5.53E-05 | 3 | 0.00025295 | 13.73215149 | 8 mo | HOM |
| **1013609713** | 2 | act | 908 | 0.218465402 | 0 | 0 | 0 | 0 | 2 mo | WILD |
| **1010418842** | 1 | act | 908 | 0.218465402 | 0.000245939 | 4 | 0.001125756 | 18.30953531 | 8 mo | HOM |
| **1013288213** | 2 | act | 908 | 0.218465402 | 3.08E-06 | 0 | 1.41E-05 | 0 | 4 mo | HOM |
| **1008976060** | 2 | act | 908 | 0.218465402 | 3.17E-07 | 0 | 1.45E-06 | 0 | 8 mo | WILD |
| **1013446243** | 1 | act | 908 | 0.218465402 | 1.77E-05 | 1 | 8.09E-05 | 4.577383828 | 4 mo | HOM |
| **1013288951** | 2 | act | 908 | 0.218465402 | 1.57E-05 | 0 | 7.19E-05 | 0 | 4 mo | HOM |
| **1013978665** | 1 | ENT | 909 | 11.74135887 | 0.000107785 | 5 | 9.18E-06 | 0.425845088 | 2 mo | HOM |
| **1013445609** | 1 | ENT | 909 | 11.74135887 | 0.00046339 | 34 | 3.95E-05 | 2.895746597 | 4 mo | HOM |
| **1013609015** | 1 | ENT | 909 | 11.74135887 | 0.003492523 | 72 | 0.000297455 | 6.132169265 | 4 mo | HOM |
| **1013608106** | 2 | ENT | 909 | 11.74135887 | 0 | 0 | 0 | 0 | 4 mo | HOM |
| **1011937543** | 1 | ENT | 909 | 11.74135887 | 0.026771627 | 277 | 0.002280113 | 23.59181787 | 8 mo | HOM |
| **1010419538** | 2 | ENT | 909 | 11.74135887 | 0.000151402 | 14 | 1.29E-05 | 1.192366246 | 4 mo | WILD |
| **1010419187** | 1 | ENT | 909 | 11.74135887 | 0.021517431 | 480 | 0.001832618 | 40.88112843 | 8 mo | HOM |
| **1013978366** | 1 | ENT | 909 | 11.74135887 | 0.000478819 | 23 | 4.08E-05 | 1.958887404 | 2 mo | HOM |
| **1014416155** | 2 | ENT | 909 | 11.74135887 | 0 | 0 | 0 | 0 | 2 mo | HOM |
| **1013978065** | 2 | ENT | 909 | 11.74135887 | 2.29E-05 | 2 | 1.95E-06 | 0.170338035 | 2 mo | HOM |
| **1009934102** | 2 | ENT | 909 | 11.74135887 | 0.00052722 | 34 | 4.49E-05 | 2.895746597 | 4 mo | WILD |
| **1009322937** | 2 | ENT | 909 | 11.74135887 | 0.000327283 | 15 | 2.79E-05 | 1.277535264 | 8 mo | WILD |
| **1011080637** | 2 | ENT | 909 | 11.74135887 | 0.00881462 | 340 | 0.000750733 | 28.95746597 | 8 mo | HOM |
| **1013609713** | 2 | ENT | 909 | 11.74135887 | 0.000135455 | 13 | 1.15E-05 | 1.107197228 | 2 mo | WILD |
| **1010418842** | 1 | ENT | 909 | 11.74135887 | 0.01361838 | 390 | 0.001159864 | 33.21591685 | 8 mo | HOM |
| **1013288213** | 2 | ENT | 909 | 11.74135887 | 0.000607777 | 42 | 5.18E-05 | 3.577098738 | 4 mo | HOM |
| **1008976060** | 2 | ENT | 909 | 11.74135887 | 0.000170594 | 17 | 1.45E-05 | 1.447873299 | 8 mo | WILD |
| **1013446243** | 1 | ENT | 909 | 11.74135887 | 0.001082139 | 55 | 9.22E-05 | 4.684295966 | 4 mo | HOM |
| **1013288951** | 2 | ENT | 909 | 11.74135887 | 0.005932767 | 89 | 0.000505288 | 7.580042564 | 4 mo | HOM |
| **1013978665** | 1 | ORBm6a | 910 | 0.19235538 | 0 | 0 | 0 | 0 | 2 mo | HOM |
| **1013445609** | 1 | ORBm6a | 910 | 0.19235538 | 0 | 0 | 0 | 0 | 4 mo | HOM |
| **1013609015** | 1 | ORBm6a | 910 | 0.19235538 | 7.72E-07 | 0 | 4.01E-06 | 0 | 4 mo | HOM |
| **1013608106** | 2 | ORBm6a | 910 | 0.19235538 | 0 | 0 | 0 | 0 | 4 mo | HOM |
| **1011937543** | 1 | ORBm6a | 910 | 0.19235538 | 8.45E-05 | 2 | 0.00043939 | 10.39742169 | 8 mo | HOM |
| **1010419538** | 2 | ORBm6a | 910 | 0.19235538 | 0 | 0 | 0 | 0 | 4 mo | WILD |
| **1010419187** | 1 | ORBm6a | 910 | 0.19235538 | 0.000197445 | 7 | 0.001026458 | 36.39097591 | 8 mo | HOM |
| **1013978366** | 1 | ORBm6a | 910 | 0.19235538 | 0 | 0 | 0 | 0 | 2 mo | HOM |
| **1014416155** | 2 | ORBm6a | 910 | 0.19235538 | 0 | 0 | 0 | 0 | 2 mo | HOM |
| **1013978065** | 2 | ORBm6a | 910 | 0.19235538 | 0 | 0 | 0 | 0 | 2 mo | HOM |
| **1009934102** | 2 | ORBm6a | 910 | 0.19235538 | 0 | 0 | 0 | 0 | 4 mo | WILD |
| **1009322937** | 2 | ORBm6a | 910 | 0.19235538 | 2.51E-06 | 0 | 1.3E-05 | 0 | 8 mo | WILD |
| **1011080637** | 2 | ORBm6a | 910 | 0.19235538 | 0.000124034 | 7 | 0.000644817 | 36.39097591 | 8 mo | HOM |

| | | | | | | | | | | |
|---|---|---|---|---|---|---|---|---|---|---|
| 1013609713 | 2 | ORBm6a | 910 | 0.19235538 | 0 | 0 | 0 | 0 | 2 mo | WILD |
| 1010418842 | 1 | ORBm6a | 910 | 0.19235538 | 0.000132819 | 1 | 0.00069049 | 5.198710844 | 8 mo | HOM |
| 1013288213 | 2 | ORBm6a | 910 | 0.19235538 | 0 | 0 | 0 | 0 | 4 mo | HOM |
| 1008976060 | 2 | ORBm6a | 910 | 0.19235538 | 0 | 0 | 0 | 0 | 8 mo | WILD |
| 1013446243 | 1 | ORBm6a | 910 | 0.19235538 | 0 | 0 | 0 | 0 | 4 mo | HOM |
| 1013288951 | 2 | ORBm6a | 910 | 0.19235538 | 1.67E-05 | 1 | 8.67E-05 | 5.198710844 | 4 mo | HOM |
| 1013978665 | 1 | IVn | 911 | 0.007009651 | 0 | 0 | 0 | 0 | 2 mo | HOM |
| 1013445609 | 1 | IVn | 911 | 0.007009651 | 0 | 0 | 0 | 0 | 4 mo | HOM |
| 1013609015 | 1 | IVn | 911 | 0.007009651 | 9.82E-07 | 0 | 0.000140121 | 0 | 4 mo | HOM |
| 1013608106 | 2 | IVn | 911 | 0.007009651 | 0 | 0 | 0 | 0 | 4 mo | HOM |
| 1011937543 | 1 | IVn | 911 | 0.007009651 | 7E-05 | 1 | 0.009984082 | 142.6604569 | 8 mo | HOM |
| 1010419538 | 2 | IVn | 911 | 0.007009651 | 0 | 0 | 0 | 0 | 4 mo | WILD |
| 1010419187 | 1 | IVn | 911 | 0.007009651 | 0 | 0 | 0 | 0 | 8 mo | HOM |
| 1013978366 | 1 | IVn | 911 | 0.007009651 | 0 | 0 | 0 | 0 | 2 mo | HOM |
| 1014416155 | 2 | IVn | 911 | 0.007009651 | 0 | 0 | 0 | 0 | 2 mo | HOM |
| 1013978065 | 2 | IVn | 911 | 0.007009651 | 0 | 0 | 0 | 0 | 2 mo | HOM |
| 1009934102 | 2 | IVn | 911 | 0.007009651 | 0 | 0 | 0 | 0 | 4 mo | WILD |
| 1009322937 | 2 | IVn | 911 | 0.007009651 | 6.28E-07 | 0 | 8.96E-05 | 0 | 8 mo | WILD |
| 1011080637 | 2 | IVn | 911 | 0.007009651 | 0 | 0 | 0 | 0 | 8 mo | HOM |
| 1013609713 | 2 | IVn | 911 | 0.007009651 | 0 | 0 | 0 | 0 | 2 mo | WILD |
| 1010418842 | 1 | IVn | 911 | 0.007009651 | 1.68E-05 | 0 | 0.002390542 | 0 | 8 mo | HOM |
| 1013288213 | 2 | IVn | 911 | 0.007009651 | 1.94E-06 | 0 | 0.000277163 | 0 | 4 mo | HOM |
| 1008976060 | 2 | IVn | 911 | 0.007009651 | 4.91E-06 | 0 | 0.000700742 | 0 | 8 mo | WILD |
| 1013446243 | 1 | IVn | 911 | 0.007009651 | 1.84E-05 | 1 | 0.002632034 | 142.6604569 | 4 mo | HOM |
| 1013288951 | 2 | IVn | 911 | 0.007009651 | 0 | 0 | 0 | 0 | 4 mo | HOM |
| 1013978665 | 1 | LING | 912 | 0.125304211 | 0.000124815 | 1 | 0.000996098 | 7.980577751 | 2 mo | HOM |
| 1013445609 | 1 | LING | 912 | 0.125304211 | 4.21E-06 | 1 | 3.36E-05 | 7.980577751 | 4 mo | HOM |
| 1013609015 | 1 | LING | 912 | 0.125304211 | 8.12E-05 | 1 | 0.000648401 | 7.980577751 | 4 mo | HOM |
| 1013608106 | 2 | LING | 912 | 0.125304211 | 0 | 0 | 0 | 0 | 4 mo | HOM |
| 1011937543 | 1 | LING | 912 | 0.125304211 | 1.65E-06 | 0 | 1.32E-05 | 0 | 8 mo | HOM |
| 1010419538 | 2 | LING | 912 | 0.11938309 | 9.53E-06 | 1 | 7.98E-05 | 8.376395657 | 4 mo | WILD |
| 1010419187 | 1 | LING | 912 | 0.125304211 | 0.000460995 | 8 | 0.003679008 | 63.84462201 | 8 mo | HOM |
| 1013978366 | 1 | LING | 912 | 0.125304211 | 1.99E-06 | 0 | 1.59E-05 | 0 | 2 mo | HOM |
| 1014416155 | 2 | LING | 912 | 0.125304211 | 0 | 0 | 0 | 0 | 2 mo | HOM |
| 1013978065 | 2 | LING | 912 | 0.125304211 | 2.54E-06 | 0 | 2.02E-05 | 0 | 2 mo | HOM |
| 1009934102 | 2 | LING | 912 | 0.125304211 | 2.88E-05 | 1 | 0.000229846 | 7.980577751 | 4 mo | WILD |
| 1009322937 | 2 | LING | 912 | 0.125304211 | 1.11E-05 | 0 | 8.88E-05 | 0 | 8 mo | WILD |
| 1011080637 | 2 | LING | 912 | 0.125304211 | 0.001966878 | 4 | 0.01569682 | 31.922311 | 8 mo | HOM |
| 1013609713 | 2 | LING | 912 | 0.125304211 | 4.37E-05 | 2 | 0.000348468 | 15.9611555 | 2 mo | WILD |
| 1010418842 | 1 | LING | 912 | 0.125304211 | 0.000517302 | 3 | 0.004128368 | 23.94173325 | 8 mo | HOM |
| 1013288213 | 2 | LING | 912 | 0.125304211 | 5.21E-07 | 0 | 4.16E-06 | 0 | 4 mo | HOM |
| 1008976060 | 2 | LING | 912 | 0.125304211 | 0.001034187 | 5 | 0.008253407 | 39.90288875 | 8 mo | WILD |
| 1013446243 | 1 | LING | 912 | 0.125304211 | 7.7E-05 | 0 | 0.000614383 | 0 | 4 mo | HOM |
| 1013288951 | 2 | LING | 912 | 0.125304211 | 0.001409166 | 5 | 0.011245959 | 39.90288875 | 4 mo | HOM |
| 1013978665 | 1 | PD | 914 | 0.012018605 | 0 | 0 | 0 | 0 | 2 mo | HOM |
| 1013445609 | 1 | PD | 914 | 0.012018605 | 0 | 0 | 0 | 0 | 4 mo | HOM |
| 1013609015 | 1 | PD | 914 | 0.012018605 | 0 | 0 | 0 | 0 | 4 mo | HOM |
| 1013608106 | 2 | PD | 914 | 0.012018605 | 0 | 0 | 0 | 0 | 4 mo | HOM |
| 1011937543 | 1 | PD | 914 | 0.012018605 | 3.49E-07 | 0 | 2.9E-05 | 0 | 8 mo | HOM |
| 1010419538 | 2 | PD | 914 | 0.012018605 | 0 | 0 | 0 | 0 | 4 mo | WILD |
| 1010419187 | 1 | PD | 914 | 0.012018605 | 0 | 0 | 0 | 0 | 8 mo | HOM |
| 1013978366 | 1 | PD | 914 | 0.012018605 | 0 | 0 | 0 | 0 | 2 mo | HOM |
| 1014416155 | 2 | PD | 914 | 0.012018605 | 0 | 0 | 0 | 0 | 2 mo | HOM |
| 1013978065 | 2 | PD | 914 | 0.012018605 | 0 | 0 | 0 | 0 | 2 mo | HOM |
| 1009934102 | 2 | PD | 914 | 0.012018605 | 0 | 0 | 0 | 0 | 4 mo | WILD |
| 1009322937 | 2 | PD | 914 | 0.012018605 | 0 | 0 | 0 | 0 | 8 mo | WILD |
| 1011080637 | 2 | PD | 914 | 0.012018605 | 0 | 0 | 0 | 0 | 8 mo | HOM |
| 1013609713 | 2 | PD | 914 | 0.012018605 | 0 | 0 | 0 | 0 | 2 mo | WILD |
| 1010418842 | 1 | PD | 914 | 0.012018605 | 0 | 0 | 0 | 0 | 8 mo | HOM |

| | | | | | | | | | | |
|---|---|---|---|---|---|---|---|---|---|---|
| 1013288213 | 2 | PD | 914 | 0.012018605 | 0 | 0 | 0 | 0 | 4 mo | HOM |
| 1008976060 | 2 | PD | 914 | 0.012018605 | 0 | 0 | 0 | 0 | 8 mo | WILD |
| 1013446243 | 1 | PD | 914 | 0.012018605 | 0 | 0 | 0 | 0 | 4 mo | HOM |
| 1013288951 | 2 | PD | 914 | 0.012018605 | 0 | 0 | 0 | 0 | 4 mo | HOM |
| 1013978665 | 1 | bsc | 916 | 0.206844445 | 7.15E-11 | 0 | 3.46E-10 | 0 | 2 mo | HOM |
| 1013445609 | 1 | bsc | 916 | 0.206844445 | 0 | 0 | 0 | 0 | 4 mo | HOM |
| 1013609015 | 1 | bsc | 916 | 0.206844445 | 2.46E-07 | 0 | 1.19E-06 | 0 | 4 mo | HOM |
| 1013608106 | 2 | bsc | 916 | 0.206844445 | 0 | 0 | 0 | 0 | 4 mo | HOM |
| 1011937543 | 1 | bsc | 916 | 0.206844445 | 0.000327406 | 3 | 0.001582863 | 14.50365275 | 8 mo | HOM |
| 1010419538 | 2 | bsc | 916 | 0.206844445 | 4.23E-08 | 0 | 2.04E-07 | 0 | 4 mo | WILD |
| 1010419187 | 1 | bsc | 916 | 0.206844445 | 0.000260153 | 2 | 0.001257723 | 9.669101831 | 8 mo | HOM |
| 1013978366 | 1 | bsc | 916 | 0.206844445 | 0 | 0 | 0 | 0 | 2 mo | HOM |
| 1014416155 | 2 | bsc | 916 | 0.206844445 | 0 | 0 | 0 | 0 | 2 mo | HOM |
| 1013978065 | 2 | bsc | 916 | 0.206844445 | 0 | 0 | 0 | 0 | 2 mo | HOM |
| 1009934102 | 2 | bsc | 916 | 0.206844445 | 2.07E-08 | 0 | 9.99E-08 | 0 | 4 mo | WILD |
| 1009322937 | 2 | bsc | 916 | 0.206844445 | 5.22E-07 | 0 | 2.52E-06 | 0 | 8 mo | WILD |
| 1011080637 | 2 | bsc | 916 | 0.206844445 | 1.56E-06 | 0 | 7.53E-06 | 0 | 8 mo | HOM |
| 1013609713 | 2 | bsc | 916 | 0.206844445 | 0 | 0 | 0 | 0 | 2 mo | WILD |
| 1010418842 | 1 | bsc | 916 | 0.206844445 | 0.00026609 | 3 | 0.001286426 | 14.50365275 | 8 mo | HOM |
| 1013288213 | 2 | bsc | 916 | 0.206844445 | 0 | 0 | 0 | 0 | 4 mo | HOM |
| 1008976060 | 2 | bsc | 916 | 0.206844445 | 3.95E-05 | 1 | 0.000190824 | 4.834550916 | 8 mo | WILD |
| 1013446243 | 1 | bsc | 916 | 0.206844445 | 7.67E-07 | 0 | 3.71E-06 | 0 | 4 mo | HOM |
| 1013288951 | 2 | bsc | 916 | 0.206844445 | 1.32E-07 | 0 | 6.37E-07 | 0 | 4 mo | HOM |
| 1013978665 | 1 | Xn | 917 | 0.009593455 | 0 | 0 | 0 | 0 | 2 mo | HOM |
| 1013445609 | 1 | Xn | 917 | 0.009593455 | 0 | 0 | 0 | 0 | 4 mo | HOM |
| 1013609015 | 1 | Xn | 917 | 0.009593455 | 0 | 0 | 0 | 0 | 4 mo | HOM |
| 1013608106 | 2 | Xn | 917 | 0.009593455 | 0 | 0 | 0 | 0 | 4 mo | HOM |
| 1011937543 | 1 | Xn | 917 | 0.009593455 | 0 | 0 | 0 | 0 | 8 mo | HOM |
| 1010419538 | 2 | Xn | 917 | 0.009593455 | 0 | 0 | 0 | 0 | 4 mo | WILD |
| 1010419187 | 1 | Xn | 917 | 0.009593455 | 0 | 0 | 0 | 0 | 8 mo | HOM |
| 1013978366 | 1 | Xn | 917 | 0.009593455 | 0 | 0 | 0 | 0 | 2 mo | HOM |
| 1014416155 | 2 | Xn | 917 | 0.009593455 | 0 | 0 | 0 | 0 | 2 mo | HOM |
| 1013978065 | 2 | Xn | 917 | 0.009593455 | 0 | 0 | 0 | 0 | 2 mo | HOM |
| 1009934102 | 2 | Xn | 917 | 0.009593455 | 0 | 0 | 0 | 0 | 4 mo | WILD |
| 1009322937 | 2 | Xn | 917 | 0.009593455 | 0 | 0 | 0 | 0 | 8 mo | WILD |
| 1011080637 | 2 | Xn | 917 | 0.009593455 | 0 | 0 | 0 | 0 | 8 mo | HOM |
| 1013609713 | 2 | Xn | 917 | 0.009593455 | 0 | 0 | 0 | 0 | 2 mo | WILD |
| 1010418842 | 1 | Xn | 917 | 0.009593455 | 0 | 0 | 0 | 0 | 8 mo | HOM |
| 1013288213 | 2 | Xn | 917 | 0.009593455 | 0 | 0 | 0 | 0 | 4 mo | HOM |
| 1008976060 | 2 | Xn | 917 | 0.009593455 | 0 | 0 | 0 | 0 | 8 mo | WILD |
| 1013446243 | 1 | Xn | 917 | 0.009593455 | 0 | 0 | 0 | 0 | 4 mo | HOM |
| 1013288951 | 2 | Xn | 917 | 0.009593455 | 0 | 0 | 0 | 0 | 4 mo | HOM |
| 1013978665 | 1 | ENTI | 918 | 6.570865661 | 9.39E-05 | 3 | 1.43E-05 | 0.456560848 | 2 mo | HOM |
| 1013445609 | 1 | ENTI | 918 | 6.570865661 | 0.000203535 | 19 | 3.1E-05 | 2.891552039 | 4 mo | HOM |
| 1013609015 | 1 | ENTI | 918 | 6.570865661 | 0.000689175 | 32 | 0.000104884 | 4.869982382 | 4 mo | HOM |
| 1013608106 | 2 | ENTI | 918 | 6.570865661 | 0 | 0 | 0 | 0 | 4 mo | HOM |
| 1011937543 | 1 | ENTI | 918 | 6.570865661 | 0.014349665 | 170 | 0.002183832 | 25.8717814 | 8 mo | HOM |
| 1010419538 | 2 | ENTI | 918 | 6.570865661 | 9.43E-06 | 0 | 1.43E-06 | 0 | 4 mo | WILD |
| 1010419187 | 1 | ENTI | 918 | 6.570865661 | 0.009952369 | 267 | 0.001514621 | 40.6339155 | 8 mo | HOM |
| 1013978366 | 1 | ENTI | 918 | 6.570865661 | 6.88E-05 | 2 | 1.05E-05 | 0.304373899 | 2 mo | HOM |
| 1014416155 | 2 | ENTI | 918 | 6.570865661 | 0 | 0 | 0 | 0 | 2 mo | HOM |
| 1013978065 | 2 | ENTI | 918 | 6.570865661 | 0 | 0 | 0 | 0 | 2 mo | HOM |
| 1009934102 | 2 | ENTI | 918 | 6.570865661 | 3.69E-05 | 2 | 5.61E-06 | 0.304373899 | 4 mo | WILD |
| 1009322937 | 2 | ENTI | 918 | 6.570865661 | 6.47E-05 | 4 | 9.85E-06 | 0.608747798 | 8 mo | WILD |
| 1011080637 | 2 | ENTI | 918 | 6.570865661 | 0.004360138 | 169 | 0.000663556 | 25.71959445 | 8 mo | HOM |
| 1013609713 | 2 | ENTI | 918 | 6.570865661 | 5.66E-06 | 1 | 8.61E-07 | 0.152186949 | 2 mo | WILD |
| 1010418842 | 1 | ENTI | 918 | 6.570865661 | 0.00776658 | 215 | 0.001181972 | 32.72019413 | 8 mo | HOM |
| 1013288213 | 2 | ENTI | 918 | 6.570865661 | 0.000313712 | 22 | 4.77E-05 | 3.348112887 | 4 mo | HOM |
| 1008976060 | 2 | ENTI | 918 | 6.570865661 | 8.92E-05 | 8 | 1.36E-05 | 1.217495595 | 8 mo | WILD |

| | | | | | | | | | | |
|---|---|---|---|---|---|---|---|---|---|---|
| **1013446243** | 1 | ENTI | 918 | 6.570865661 | 0.000564417 | 26 | 8.59E-05 | 3.956860685 | 4 mo | HOM |
| **1013288951** | 2 | ENTI | 918 | 6.570865661 | 0.000575774 | 37 | 8.76E-05 | 5.630917129 | 4 mo | HOM |
| **1013978665** | 1 | ACAd6a | 919 | 0.720053118 | 0 | 0 | 0 | 0 | 2 mo | HOM |
| **1013445609** | 1 | ACAd6a | 919 | 0.720053118 | 2.93E-05 | 2 | 4.07E-05 | 2.777572861 | 4 mo | HOM |
| **1013609015** | 1 | ACAd6a | 919 | 0.720053118 | 0.000241259 | 8 | 0.000335057 | 11.11029144 | 4 mo | HOM |
| **1013608106** | 2 | ACAd6a | 919 | 0.720053118 | 0 | 0 | 0 | 0 | 4 mo | HOM |
| **1011937543** | 1 | ACAd6a | 919 | 0.720053118 | 0.000468635 | 7 | 0.000650834 | 9.721505013 | 8 mo | HOM |
| **1010419538** | 2 | ACAd6a | 919 | 0.720053118 | 0 | 0 | 0 | 0 | 4 mo | WILD |
| **1010419187** | 1 | ACAd6a | 919 | 0.720053118 | 0.000760945 | 29 | 0.00105679 | 40.27480648 | 8 mo | HOM |
| **1013978366** | 1 | ACAd6a | 919 | 0.720053118 | 0 | 0 | 0 | 0 | 2 mo | HOM |
| **1014416155** | 2 | ACAd6a | 919 | 0.720053118 | 0 | 0 | 0 | 0 | 2 mo | HOM |
| **1013978065** | 2 | ACAd6a | 919 | 0.720053118 | 0 | 0 | 0 | 0 | 2 mo | HOM |
| **1009934102** | 2 | ACAd6a | 919 | 0.720053118 | 0 | 0 | 0 | 0 | 4 mo | WILD |
| **1009322937** | 2 | ACAd6a | 919 | 0.720053118 | 0 | 0 | 0 | 0 | 8 mo | WILD |
| **1011080637** | 2 | ACAd6a | 919 | 0.720053118 | 0.000469545 | 41 | 0.000652097 | 56.94024365 | 8 mo | HOM |
| **1013609713** | 2 | ACAd6a | 919 | 0.720053118 | 0 | 0 | 0 | 0 | 2 mo | WILD |
| **1010418842** | 1 | ACAd6a | 919 | 0.720053118 | 0.000604401 | 26 | 0.000839383 | 36.10844719 | 8 mo | HOM |
| **1013288213** | 2 | ACAd6a | 919 | 0.720053118 | 1.34E-23 | 0 | 1.86E-23 | 0 | 4 mo | HOM |
| **1008976060** | 2 | ACAd6a | 919 | 0.720053118 | 0 | 0 | 0 | 0 | 8 mo | WILD |
| **1013446243** | 1 | ACAd6a | 919 | 0.720053118 | 7.9E-05 | 3 | 0.00010978 | 4.166359291 | 4 mo | HOM |
| **1013288951** | 2 | ACAd6a | 919 | 0.720053118 | 4.61E-05 | 4 | 6.41E-05 | 5.555145722 | 4 mo | HOM |
| **1013978665** | 1 | CENT | 920 | 4.202720435 | 0.000906002 | 32 | 0.000215575 | 7.614115784 | 2 mo | HOM |
| **1013445609** | 1 | CENT | 920 | 4.202720435 | 0.000189131 | 3 | 4.5E-05 | 0.713823355 | 4 mo | HOM |
| **1013609015** | 1 | CENT | 920 | 4.202720435 | 0.003883627 | 43 | 0.000924074 | 10.23146808 | 4 mo | HOM |
| **1013608106** | 2 | CENT | 920 | 4.202720435 | 0 | 0 | 0 | 0 | 4 mo | HOM |
| **1011937543** | 1 | CENT | 920 | 4.202720435 | 0.004302474 | 45 | 0.001023736 | 10.70735032 | 8 mo | HOM |
| **1010419538** | 2 | CENT | 920 | 4.186029389 | 0.00017053 | 8 | 4.07E-05 | 1.911118928 | 4 mo | WILD |
| **1010419187** | 1 | CENT | 920 | 4.202720435 | 0.002367177 | 60 | 0.000563249 | 14.27646709 | 8 mo | HOM |
| **1013978366** | 1 | CENT | 920 | 4.202720435 | 0.001151692 | 16 | 0.000274035 | 3.807057892 | 2 mo | HOM |
| **1014416155** | 2 | CENT | 920 | 4.202720435 | 0 | 0 | 0 | 0 | 2 mo | HOM |
| **1013978065** | 2 | CENT | 920 | 4.202720435 | 0.000267551 | 12 | 6.37E-05 | 2.855293419 | 2 mo | HOM |
| **1009934102** | 2 | CENT | 920 | 4.202720435 | 0.000366838 | 13 | 8.73E-05 | 3.093234537 | 4 mo | WILD |
| **1009322937** | 2 | CENT | 920 | 4.202720435 | 0.000526027 | 24 | 0.000125164 | 5.710586838 | 8 mo | WILD |
| **1011080637** | 2 | CENT | 920 | 4.202720435 | 0.010481892 | 92 | 0.002494073 | 21.89058288 | 8 mo | HOM |
| **1013609713** | 2 | CENT | 920 | 4.202720435 | 0.000216332 | 9 | 5.15E-05 | 2.141470064 | 2 mo | WILD |
| **1010418842** | 1 | CENT | 920 | 4.202720435 | 0.01261278 | 141 | 0.003001099 | 33.54969767 | 8 mo | HOM |
| **1013288213** | 2 | CENT | 920 | 4.202720435 | 0.000341914 | 15 | 8.14E-05 | 3.569116774 | 4 mo | HOM |
| **1008976060** | 2 | CENT | 920 | 4.202720435 | 0.002719435 | 62 | 0.000647065 | 14.75234933 | 8 mo | WILD |
| **1013446243** | 1 | CENT | 920 | 4.202720435 | 0.000516849 | 15 | 0.00012298 | 3.569116774 | 4 mo | HOM |
| **1013288951** | 2 | CENT | 920 | 4.202720435 | 0.000425475 | 17 | 0.000101238 | 4.04499901 | 4 mo | HOM |
| **1013978665** | 1 | PERI | 922 | 0.813663448 | 0 | 0 | 0 | 0 | 2 mo | HOM |
| **1013445609** | 1 | PERI | 922 | 0.813663448 | 3.22E-05 | 3 | 3.95E-05 | 3.687028102 | 4 mo | HOM |
| **1013609015** | 1 | PERI | 922 | 0.813663448 | 7.76E-05 | 7 | 9.53E-05 | 8.603065571 | 4 mo | HOM |
| **1013608106** | 2 | PERI | 922 | 0.813663448 | 0 | 0 | 0 | 0 | 4 mo | HOM |
| **1011937543** | 1 | PERI | 922 | 0.813663448 | 0.001579091 | 30 | 0.001940718 | 36.87028102 | 8 mo | HOM |
| **1010419538** | 2 | PERI | 922 | 0.813663448 | 0 | 0 | 0 | 0 | 4 mo | WILD |
| **1010419187** | 1 | PERI | 922 | 0.813663448 | 0.000921193 | 35 | 0.001132155 | 43.01532785 | 8 mo | HOM |
| **1013978366** | 1 | PERI | 922 | 0.813663448 | 1.55E-06 | 0 | 1.91E-06 | 0 | 2 mo | HOM |
| **1014416155** | 2 | PERI | 922 | 0.813663448 | 0 | 0 | 0 | 0 | 2 mo | HOM |
| **1013978065** | 2 | PERI | 922 | 0.813663448 | 0 | 0 | 0 | 0 | 2 mo | HOM |
| **1009934102** | 2 | PERI | 922 | 0.813663448 | 1.28E-06 | 0 | 1.57E-06 | 0 | 4 mo | WILD |
| **1009322937** | 2 | PERI | 922 | 0.813663448 | 0 | 0 | 0 | 0 | 8 mo | WILD |
| **1011080637** | 2 | PERI | 922 | 0.813663448 | 0.000403449 | 21 | 0.000495843 | 25.80919671 | 8 mo | HOM |
| **1013609713** | 2 | PERI | 922 | 0.813663448 | 0 | 0 | 0 | 0 | 2 mo | WILD |
| **1010418842** | 1 | PERI | 922 | 0.813663448 | 0.000703057 | 34 | 0.000864064 | 41.78631849 | 8 mo | HOM |
| **1013288213** | 2 | PERI | 922 | 0.813663448 | 1.06E-06 | 0 | 1.3E-06 | 0 | 4 mo | HOM |
| **1008976060** | 2 | PERI | 922 | 0.813663448 | 7.11E-06 | 0 | 8.74E-06 | 0 | 8 mo | WILD |
| **1013446243** | 1 | PERI | 922 | 0.813663448 | 6.52E-05 | 5 | 8.01E-05 | 6.145046836 | 4 mo | HOM |
| **1013288951** | 2 | PERI | 922 | 0.813663448 | 9.43E-05 | 8 | 0.000115863 | 9.832074938 | 4 mo | HOM |

| | | | | | | | | | | |
|---|---|---|---|---|---|---|---|---|---|---|
| **1013978665** | 1 | cpd | 924 | 0.912495013 | 0 | 0 | 0 | 0 | 2 mo | HOM |
| **1013445609** | 1 | cpd | 924 | 0.912495013 | 0 | 0 | 0 | 0 | 4 mo | HOM |
| **1013609015** | 1 | cpd | 924 | 0.912495013 | 0 | 0 | 0 | 0 | 4 mo | HOM |
| **1013608106** | 2 | cpd | 924 | 0.912495013 | 0 | 0 | 0 | 0 | 4 mo | HOM |
| **1011937543** | 1 | cpd | 924 | 0.912495013 | 0.001869215 | 18 | 0.002048466 | 19.72613521 | 8 mo | HOM |
| **1010419538** | 2 | cpd | 924 | 0.912495013 | 1.13E-06 | 0 | 1.24E-06 | 0 | 4 mo | WILD |
| **1010419187** | 1 | cpd | 924 | 0.912495013 | 0.000149685 | 0 | 0.000164039 | 0 | 8 mo | HOM |
| **1013978366** | 1 | cpd | 924 | 0.912495013 | 0 | 0 | 0 | 0 | 2 mo | HOM |
| **1014416155** | 2 | cpd | 924 | 0.912495013 | 0 | 0 | 0 | 0 | 2 mo | HOM |
| **1013978065** | 2 | cpd | 924 | 0.912495013 | 0 | 0 | 0 | 0 | 2 mo | HOM |
| **1009934102** | 2 | cpd | 924 | 0.912495013 | 2.16E-06 | 0 | 2.36E-06 | 0 | 4 mo | WILD |
| **1009322937** | 2 | cpd | 924 | 0.912495013 | 6.58E-11 | 0 | 7.21E-11 | 0 | 8 mo | WILD |
| **1011080637** | 2 | cpd | 924 | 0.912495013 | 3.53E-05 | 1 | 3.87E-05 | 1.0958964 | 8 mo | HOM |
| **1013609713** | 2 | cpd | 924 | 0.912495013 | 0 | 0 | 0 | 0 | 2 mo | WILD |
| **1010418842** | 1 | cpd | 924 | 0.912495013 | 6.69E-05 | 3 | 7.34E-05 | 3.287689201 | 8 mo | HOM |
| **1013288213** | 2 | cpd | 924 | 0.912495013 | 0 | 0 | 0 | 0 | 4 mo | HOM |
| **1008976060** | 2 | cpd | 924 | 0.912495013 | 3.62E-07 | 0 | 3.97E-07 | 0 | 8 mo | WILD |
| **1013446243** | 1 | cpd | 924 | 0.912495013 | 0 | 0 | 0 | 0 | 4 mo | HOM |
| **1013288951** | 2 | cpd | 924 | 0.912495013 | 8.8E-07 | 0 | 9.65E-07 | 0 | 4 mo | HOM |
| **1013978665** | 1 | ENTm | 926 | 5.17049321 | 1.39E-05 | 2 | 2.69E-06 | 0.386810294 | 2 mo | HOM |
| **1013445609** | 1 | ENTm | 926 | 5.17049321 | 0.000259856 | 15 | 5.03E-05 | 2.901077207 | 4 mo | HOM |
| **1013609015** | 1 | ENTm | 926 | 5.17049321 | 0.002803348 | 40 | 0.000542182 | 7.736205884 | 4 mo | HOM |
| **1013608106** | 2 | ENTm | 926 | 5.17049321 | 0 | 0 | 0 | 0 | 4 mo | HOM |
| **1011937543** | 1 | ENTm | 926 | 5.17049321 | 0.012421962 | 107 | 0.002402471 | 20.69435074 | 8 mo | HOM |
| **1010419538** | 2 | ENTm | 926 | 5.17049321 | 0.000141975 | 14 | 2.75E-05 | 2.70767206 | 4 mo | WILD |
| **1010419187** | 1 | ENTm | 926 | 5.17049321 | 0.011565063 | 213 | 0.002236743 | 41.19529633 | 8 mo | HOM |
| **1013978366** | 1 | ENTm | 926 | 5.17049321 | 0.000410018 | 21 | 7.93E-05 | 4.061508089 | 2 mo | HOM |
| **1014416155** | 2 | ENTm | 926 | 5.17049321 | 0 | 0 | 0 | 0 | 2 mo | HOM |
| **1013978065** | 2 | ENTm | 926 | 5.17049321 | 2.29E-05 | 2 | 4.44E-06 | 0.386810294 | 2 mo | HOM |
| **1009934102** | 2 | ENTm | 926 | 5.17049321 | 0.000490354 | 32 | 9.48E-05 | 6.188964708 | 4 mo | WILD |
| **1009322937** | 2 | ENTm | 926 | 5.17049321 | 0.00026254 | 11 | 5.08E-05 | 2.127456618 | 8 mo | WILD |
| **1011080637** | 2 | ENTm | 926 | 5.17049321 | 0.004454482 | 171 | 0.00086152 | 33.07228016 | 8 mo | HOM |
| **1013609713** | 2 | ENTm | 926 | 5.17049321 | 0.0001298 | 12 | 2.51E-05 | 2.320861765 | 2 mo | WILD |
| **1010418842** | 1 | ENTm | 926 | 5.17049321 | 0.0058518 | 175 | 0.001131768 | 33.84590074 | 8 mo | HOM |
| **1013288213** | 2 | ENTm | 926 | 5.17049321 | 0.000294066 | 20 | 5.69E-05 | 3.868102942 | 4 mo | HOM |
| **1008976060** | 2 | ENTm | 926 | 5.17049321 | 8.14E-05 | 9 | 1.57E-05 | 1.740646324 | 8 mo | WILD |
| **1013446243** | 1 | ENTm | 926 | 5.17049321 | 0.000517723 | 29 | 0.00010013 | 5.608749266 | 4 mo | HOM |
| **1013288951** | 2 | ENTm | 926 | 5.17049321 | 0.005356994 | 52 | 0.00103607 | 10.05706765 | 4 mo | HOM |
| **1013978665** | 1 | ACAd6b | 927 | 0.025463041 | 0 | 0 | 0 | 0 | 2 mo | HOM |
| **1013445609** | 1 | ACAd6b | 927 | 0.025463041 | 0 | 0 | 0 | 0 | 4 mo | HOM |
| **1013609015** | 1 | ACAd6b | 927 | 0.025463041 | 0 | 0 | 0 | 0 | 4 mo | HOM |
| **1013608106** | 2 | ACAd6b | 927 | 0.025463041 | 0 | 0 | 0 | 0 | 4 mo | HOM |
| **1011937543** | 1 | ACAd6b | 927 | 0.025463041 | 2.6E-05 | 0 | 0.001022534 | 0 | 8 mo | HOM |
| **1010419538** | 2 | ACAd6b | 927 | 0.025463041 | 0 | 0 | 0 | 0 | 4 mo | WILD |
| **1010419187** | 1 | ACAd6b | 927 | 0.025463041 | 7.7E-06 | 1 | 0.000302446 | 39.27260676 | 8 mo | HOM |
| **1013978366** | 1 | ACAd6b | 927 | 0.025463041 | 0 | 0 | 0 | 0 | 2 mo | HOM |
| **1014416155** | 2 | ACAd6b | 927 | 0.025463041 | 0 | 0 | 0 | 0 | 2 mo | HOM |
| **1013978065** | 2 | ACAd6b | 927 | 0.025463041 | 0 | 0 | 0 | 0 | 2 mo | HOM |
| **1009934102** | 2 | ACAd6b | 927 | 0.025463041 | 0 | 0 | 0 | 0 | 4 mo | WILD |
| **1009322937** | 2 | ACAd6b | 927 | 0.025463041 | 0 | 0 | 0 | 0 | 8 mo | WILD |
| **1011080637** | 2 | ACAd6b | 927 | 0.025463041 | 1.57E-05 | 1 | 0.000614893 | 39.27260676 | 8 mo | HOM |
| **1013609713** | 2 | ACAd6b | 927 | 0.025463041 | 0 | 0 | 0 | 0 | 2 mo | WILD |
| **1010418842** | 1 | ACAd6b | 927 | 0.025463041 | 6.08E-06 | 0 | 0.000238669 | 0 | 8 mo | HOM |
| **1013288213** | 2 | ACAd6b | 927 | 0.025463041 | 0 | 0 | 0 | 0 | 4 mo | HOM |
| **1008976060** | 2 | ACAd6b | 927 | 0.025463041 | 0 | 0 | 0 | 0 | 8 mo | WILD |
| **1013446243** | 1 | ACAd6b | 927 | 0.025463041 | 0 | 0 | 0 | 0 | 4 mo | HOM |
| **1013288951** | 2 | ACAd6b | 927 | 0.025463041 | 0 | 0 | 0 | 0 | 4 mo | HOM |
| **1013978665** | 1 | CUL | 928 | 6.929062874 | 0.000729491 | 26 | 0.00010528 | 3.752311167 | 2 mo | HOM |
| **1013445609** | 1 | CUL | 928 | 6.929062874 | 0.000132449 | 3 | 1.91E-05 | 0.432958981 | 4 mo | HOM |

| | | | | | | | | | | |
|---|---|---|---|---|---|---|---|---|---|---|
| **1013609015** | 1 | CUL | 928 | 6.929062874 | 0.002160155 | 51 | 0.000311753 | 7.360302674 | 4 mo | HOM |
| **1013608106** | 2 | CUL | 928 | 6.929062874 | 0 | 0 | 0 | 0 | 4 mo | HOM |
| **1011937543** | 1 | CUL | 928 | 6.929062874 | 0.00736756 | 47 | 0.001063284 | 6.783024033 | 8 mo | HOM |
| **1010419538** | 2 | CUL | 928 | 6.919467341 | 0.000641261 | 18 | 9.27E-05 | 2.601356306 | 4 mo | WILD |
| **1010419187** | 1 | CUL | 928 | 6.929062874 | 0.005305846 | 122 | 0.000765738 | 17.60699855 | 8 mo | HOM |
| **1013978366** | 1 | CUL | 928 | 6.929062874 | 0.000597636 | 25 | 8.63E-05 | 3.607991507 | 2 mo | HOM |
| **1014416155** | 2 | CUL | 928 | 6.929062874 | 0 | 0 | 0 | 0 | 2 mo | HOM |
| **1013978065** | 2 | CUL | 928 | 6.929062874 | 0.000243308 | 6 | 3.51E-05 | 0.865917962 | 2 mo | HOM |
| **1009934102** | 2 | CUL | 928 | 6.929062874 | 0.00038076 | 12 | 5.5E-05 | 1.731835923 | 4 mo | WILD |
| **1009322937** | 2 | CUL | 928 | 6.929062874 | 0.001061446 | 41 | 0.000153187 | 5.917106072 | 8 mo | WILD |
| **1011080637** | 2 | CUL | 928 | 6.929062874 | 0.009674886 | 136 | 0.001396276 | 19.6274738 | 8 mo | HOM |
| **1013609713** | 2 | CUL | 928 | 6.929062874 | 0.000118733 | 3 | 1.71E-05 | 0.432958981 | 2 mo | WILD |
| **1010418842** | 1 | CUL | 928 | 6.929062874 | 0.015365877 | 182 | 0.002217598 | 26.26617817 | 8 mo | HOM |
| **1013288213** | 2 | CUL | 928 | 6.929062874 | 0.000307292 | 11 | 4.43E-05 | 1.587516263 | 4 mo | HOM |
| **1008976060** | 2 | CUL | 928 | 6.929062874 | 0.000940445 | 71 | 0.000135725 | 10.24669588 | 8 mo | WILD |
| **1013446243** | 1 | CUL | 928 | 6.929062874 | 0.000145307 | 16 | 2.1E-05 | 2.309114565 | 4 mo | HOM |
| **1013288951** | 2 | CUL | 928 | 6.929062874 | 0.002316034 | 34 | 0.000334249 | 4.90686845 | 4 mo | HOM |
| **1013978665** | 1 | SSp-n6b | 929 | 0.055133521 | 0 | 0 | 0 | 0 | 2 mo | HOM |
| **1013445609** | 1 | SSp-n6b | 929 | 0.055133521 | 0 | 0 | 0 | 0 | 4 mo | HOM |
| **1013609015** | 1 | SSp-n6b | 929 | 0.055133521 | 0 | 0 | 0 | 0 | 4 mo | HOM |
| **1013608106** | 2 | SSp-n6b | 929 | 0.055133521 | 0 | 0 | 0 | 0 | 4 mo | HOM |
| **1011937543** | 1 | SSp-n6b | 929 | 0.055133521 | 4.55E-05 | 0 | 0.000824459 | 0 | 8 mo | HOM |
| **1010419538** | 2 | SSp-n6b | 929 | 0.055133521 | 0 | 0 | 0 | 0 | 4 mo | WILD |
| **1010419187** | 1 | SSp-n6b | 929 | 0.055133521 | 2.33E-05 | 2 | 0.000423338 | 36.27557174 | 8 mo | HOM |
| **1013978366** | 1 | SSp-n6b | 929 | 0.055133521 | 0 | 0 | 0 | 0 | 2 mo | HOM |
| **1014416155** | 2 | SSp-n6b | 929 | 0.055133521 | 0 | 0 | 0 | 0 | 2 mo | HOM |
| **1013978065** | 2 | SSp-n6b | 929 | 0.055133521 | 0 | 0 | 0 | 0 | 2 mo | HOM |
| **1009934102** | 2 | SSp-n6b | 929 | 0.055133521 | 0 | 0 | 0 | 0 | 4 mo | WILD |
| **1009322937** | 2 | SSp-n6b | 929 | 0.055133521 | 0 | 0 | 0 | 0 | 8 mo | WILD |
| **1011080637** | 2 | SSp-n6b | 929 | 0.055133521 | 2.78E-05 | 4 | 0.000504078 | 72.55114348 | 8 mo | HOM |
| **1013609713** | 2 | SSp-n6b | 929 | 0.055133521 | 0 | 0 | 0 | 0 | 2 mo | WILD |
| **1010418842** | 1 | SSp-n6b | 929 | 0.055133521 | 1.08E-05 | 2 | 0.000196314 | 36.27557174 | 8 mo | HOM |
| **1013288213** | 2 | SSp-n6b | 929 | 0.055133521 | 0 | 0 | 0 | 0 | 4 mo | HOM |
| **1008976060** | 2 | SSp-n6b | 929 | 0.055133521 | 0 | 0 | 0 | 0 | 8 mo | WILD |
| **1013446243** | 1 | SSp-n6b | 929 | 0.055133521 | 8.36E-06 | 0 | 0.000151705 | 0 | 4 mo | HOM |
| **1013288951** | 2 | SSp-n6b | 929 | 0.055133521 | 0 | 0 | 0 | 0 | 4 mo | HOM |
| **1013978665** | 1 | PF | 930 | 0.500683176 | 0 | 0 | 0 | 0 | 2 mo | HOM |
| **1013445609** | 1 | PF | 930 | 0.500683176 | 0 | 0 | 0 | 0 | 4 mo | HOM |
| **1013609015** | 1 | PF | 930 | 0.500683176 | 4.77E-07 | 0 | 9.53E-07 | 0 | 4 mo | HOM |
| **1013608106** | 2 | PF | 930 | 0.500683176 | 0 | 0 | 0 | 0 | 4 mo | HOM |
| **1011937543** | 1 | PF | 930 | 0.500683176 | 0.000234133 | 7 | 0.000467627 | 13.98089717 | 8 mo | HOM |
| **1010419538** | 2 | PF | 930 | 0.500683176 | 0 | 0 | 0 | 0 | 4 mo | WILD |
| **1010419187** | 1 | PF | 930 | 0.500683176 | 4.91E-05 | 2 | 9.81E-05 | 3.994542049 | 8 mo | HOM |
| **1013978366** | 1 | PF | 930 | 0.500683176 | 0 | 0 | 0 | 0 | 2 mo | HOM |
| **1014416155** | 2 | PF | 930 | 0.500683176 | 0 | 0 | 0 | 0 | 2 mo | HOM |
| **1013978065** | 2 | PF | 930 | 0.500683176 | 0 | 0 | 0 | 0 | 2 mo | HOM |
| **1009934102** | 2 | PF | 930 | 0.500683176 | 0 | 0 | 0 | 0 | 4 mo | WILD |
| **1009322937** | 2 | PF | 930 | 0.500683176 | 0 | 0 | 0 | 0 | 8 mo | WILD |
| **1011080637** | 2 | PF | 930 | 0.500683176 | 6.99E-05 | 2 | 0.000139702 | 3.994542049 | 8 mo | HOM |
| **1013609713** | 2 | PF | 930 | 0.500683176 | 0 | 0 | 0 | 0 | 2 mo | WILD |
| **1010418842** | 1 | PF | 930 | 0.500683176 | 0.000277525 | 4 | 0.000554293 | 7.989084099 | 8 mo | HOM |
| **1013288213** | 2 | PF | 930 | 0.500683176 | 0 | 0 | 0 | 0 | 4 mo | HOM |
| **1008976060** | 2 | PF | 930 | 0.500683176 | 9.12E-06 | 1 | 1.82E-05 | 1.997271025 | 8 mo | WILD |
| **1013446243** | 1 | PF | 930 | 0.500683176 | 0 | 0 | 0 | 0 | 4 mo | HOM |
| **1013288951** | 2 | PF | 930 | 0.500683176 | 0 | 0 | 0 | 0 | 4 mo | HOM |
| **1013978665** | 1 | PG | 931 | 0.994214547 | 0 | 0 | 0 | 0 | 2 mo | HOM |
| **1013445609** | 1 | PG | 931 | 0.994214547 | 0 | 0 | 0 | 0 | 4 mo | HOM |
| **1013609015** | 1 | PG | 931 | 0.994214547 | 0 | 0 | 0 | 0 | 4 mo | HOM |
| **1013608106** | 2 | PG | 931 | 0.994214547 | 0 | 0 | 0 | 0 | 4 mo | HOM |

| | | | | | | | | | | |
|---|---|---|---|---|---|---|---|---|---|---|
| **1011937543** | 1 | PG | 931 | 0.994214547 | 0.000703585 | 4 | 0.00070768 | 4.023276476 | 8 mo | HOM |
| **1010419538** | 2 | PG | 931 | 0.994214547 | 0 | 0 | 0 | 0 | 4 mo | WILD |
| **1010419187** | 1 | PG | 931 | 0.994214547 | 0.000249841 | 0 | 0.000251295 | 0 | 8 mo | HOM |
| **1013978366** | 1 | PG | 931 | 0.994214547 | 0 | 0 | 0 | 0 | 2 mo | HOM |
| **1014416155** | 2 | PG | 931 | 0.994214547 | 0 | 0 | 0 | 0 | 2 mo | HOM |
| **1013978065** | 2 | PG | 931 | 0.994214547 | 0 | 0 | 0 | 0 | 2 mo | HOM |
| **1009934102** | 2 | PG | 931 | 0.994214547 | 0 | 0 | 0 | 0 | 4 mo | WILD |
| **1009322937** | 2 | PG | 931 | 0.994214547 | 0 | 0 | 0 | 0 | 8 mo | WILD |
| **1011080637** | 2 | PG | 931 | 0.994214547 | 0 | 0 | 0 | 0 | 8 mo | HOM |
| **1013609713** | 2 | PG | 931 | 0.994214547 | 0 | 0 | 0 | 0 | 2 mo | WILD |
| **1010418842** | 1 | PG | 931 | 0.994214547 | 0.00029465 | 2 | 0.000296365 | 2.011638238 | 8 mo | HOM |
| **1013288213** | 2 | PG | 931 | 0.994214547 | 0 | 0 | 0 | 0 | 4 mo | HOM |
| **1008976060** | 2 | PG | 931 | 0.994214547 | 4.78E-05 | 3 | 4.8E-05 | 3.017457357 | 8 mo | WILD |
| **1013446243** | 1 | PG | 931 | 0.994214547 | 0 | 0 | 0 | 0 | 4 mo | HOM |
| **1013288951** | 2 | PG | 931 | 0.994214547 | 0 | 0 | 0 | 0 | 4 mo | HOM |
| **1013978665** | 1 | cett | 932 | 0.708780345 | 0 | 0 | 0 | 0 | 2 mo | HOM |
| **1013445609** | 1 | cett | 932 | 0.708780345 | 0 | 0 | 0 | 0 | 4 mo | HOM |
| **1013609015** | 1 | cett | 932 | 0.708780345 | 0 | 0 | 0 | 0 | 4 mo | HOM |
| **1013608106** | 2 | cett | 932 | 0.707673729 | 0 | 0 | 0 | 0 | 4 mo | HOM |
| **1011937543** | 1 | cett | 932 | 0.708780345 | 0.001106121 | 8 | 0.001560598 | 11.2869947 | 8 mo | HOM |
| **1010419538** | 2 | cett | 932 | 0.708780345 | 0 | 0 | 0 | 0 | 4 mo | WILD |
| **1010419187** | 1 | cett | 932 | 0.708780345 | 0 | 0 | 0 | 0 | 8 mo | HOM |
| **1013978366** | 1 | cett | 932 | 0.708780345 | 0 | 0 | 0 | 0 | 2 mo | HOM |
| **1014416155** | 2 | cett | 932 | 0.708780345 | 0 | 0 | 0 | 0 | 2 mo | HOM |
| **1013978065** | 2 | cett | 932 | 0.708780345 | 0 | 0 | 0 | 0 | 2 mo | HOM |
| **1009934102** | 2 | cett | 932 | 0.708780345 | 0 | 0 | 0 | 0 | 4 mo | WILD |
| **1009322937** | 2 | cett | 932 | 0.708780345 | 0 | 0 | 0 | 0 | 8 mo | WILD |
| **1011080637** | 2 | cett | 932 | 0.708780345 | 1.08E-08 | 0 | 1.53E-08 | 0 | 8 mo | HOM |
| **1013609713** | 2 | cett | 932 | 0.708780345 | 0 | 0 | 0 | 0 | 2 mo | WILD |
| **1010418842** | 1 | cett | 932 | 0.708780345 | 2.85E-05 | 2 | 4.02E-05 | 2.821748676 | 8 mo | HOM |
| **1013288213** | 2 | cett | 932 | 0.708780345 | 0 | 0 | 0 | 0 | 4 mo | HOM |
| **1008976060** | 2 | cett | 932 | 0.708780345 | 6.86E-06 | 1 | 9.68E-06 | 1.410874338 | 8 mo | WILD |
| **1013446243** | 1 | cett | 932 | 0.708780345 | 0 | 0 | 0 | 0 | 4 mo | HOM |
| **1013288951** | 2 | cett | 932 | 0.708780345 | 0 | 0 | 0 | 0 | 4 mo | HOM |
| **1013978665** | 1 | VIIIn | 933 | 1.711436563 | 3.92E-07 | 0 | 2.29E-07 | 0 | 2 mo | HOM |
| **1013445609** | 1 | VIIIn | 933 | 1.711436563 | 5.08E-06 | 1 | 2.97E-06 | 0.58430445 | 4 mo | HOM |
| **1013609015** | 1 | VIIIn | 933 | 1.711436563 | 2.07E-05 | 0 | 1.21E-05 | 0 | 4 mo | HOM |
| **1013608106** | 2 | VIIIn | 933 | 1.711436563 | 0 | 0 | 0 | 0 | 4 mo | HOM |
| **1011937543** | 1 | VIIIn | 933 | 1.711436563 | 0.002009968 | 17 | 0.001174433 | 9.933175652 | 8 mo | HOM |
| **1010419538** | 2 | VIIIn | 933 | 1.711436563 | 0 | 0 | 0 | 0 | 4 mo | WILD |
| **1010419187** | 1 | VIIIn | 933 | 1.711436563 | 0.002334375 | 3 | 0.001363985 | 1.75291335 | 8 mo | HOM |
| **1013978366** | 1 | VIIIn | 933 | 1.711436563 | 3.92E-05 | 1 | 2.29E-05 | 0.58430445 | 2 mo | HOM |
| **1014416155** | 2 | VIIIn | 933 | 1.711436563 | 0 | 0 | 0 | 0 | 2 mo | HOM |
| **1013978065** | 2 | VIIIn | 933 | 1.711436563 | 0 | 0 | 0 | 0 | 2 mo | HOM |
| **1009934102** | 2 | VIIIn | 933 | 1.711436563 | 1.14E-06 | 0 | 6.69E-07 | 0 | 4 mo | WILD |
| **1009322937** | 2 | VIIIn | 933 | 1.711436563 | 4.9E-06 | 0 | 2.86E-06 | 0 | 8 mo | WILD |
| **1011080637** | 2 | VIIIn | 933 | 1.711436563 | 6.49E-06 | 0 | 3.79E-06 | 0 | 8 mo | HOM |
| **1013609713** | 2 | VIIIn | 933 | 1.711436563 | 0 | 0 | 0 | 0 | 2 mo | WILD |
| **1010418842** | 1 | VIIIn | 933 | 1.711436563 | 0.000404274 | 4 | 0.000236219 | 2.337217801 | 8 mo | HOM |
| **1013288213** | 2 | VIIIn | 933 | 1.711436563 | 8.77E-08 | 0 | 5.12E-08 | 0 | 4 mo | HOM |
| **1008976060** | 2 | VIIIn | 933 | 1.711436563 | 5.7E-05 | 4 | 3.33E-05 | 2.337217801 | 8 mo | WILD |
| **1013446243** | 1 | VIIIn | 933 | 1.711436563 | 1.36E-06 | 0 | 7.93E-07 | 0 | 4 mo | HOM |
| **1013288951** | 2 | VIIIn | 933 | 1.711436563 | 0 | 0 | 0 | 0 | 4 mo | HOM |
| **1013978665** | 1 | ACAd1 | 935 | 0.62416592 | 4.77E-05 | 1 | 7.64E-05 | 1.602138098 | 2 mo | HOM |
| **1013445609** | 1 | ACAd1 | 935 | 0.62416592 | 2.26E-05 | 4 | 3.62E-05 | 6.408552393 | 4 mo | HOM |
| **1013609015** | 1 | ACAd1 | 935 | 0.62416592 | 0.000300114 | 7 | 0.000480823 | 11.21496669 | 4 mo | HOM |
| **1013608106** | 2 | ACAd1 | 935 | 0.62416592 | 0 | 0 | 0 | 0 | 4 mo | HOM |
| **1011937543** | 1 | ACAd1 | 935 | 0.62416592 | 0.00077826 | 12 | 0.00124688 | 19.22565718 | 8 mo | HOM |
| **1010419538** | 2 | ACAd1 | 935 | 0.62416592 | 4.21E-06 | 0 | 6.75E-06 | 0 | 4 mo | WILD |

| | | | | | | | | | | |
|---|---|---|---|---|---|---|---|---|---|---|
| 1010419187 | 1 | ACAd1 | 935 | 0.62416592 | 0.000958783 | 22 | 0.001536103 | 35.24703816 | 8 mo | HOM |
| 1013978366 | 1 | ACAd1 | 935 | 0.62416592 | 0.000132034 | 3 | 0.000211536 | 4.806414294 | 2 mo | HOM |
| 1014416155 | 2 | ACAd1 | 935 | 0.62416592 | 0 | 0 | 0 | 0 | 2 mo | HOM |
| 1013978065 | 2 | ACAd1 | 935 | 0.62416592 | 0 | 0 | 0 | 0 | 2 mo | HOM |
| 1009934102 | 2 | ACAd1 | 935 | 0.62416592 | 7.15E-05 | 2 | 0.000114499 | 3.204276196 | 4 mo | WILD |
| 1009322937 | 2 | ACAd1 | 935 | 0.62416592 | 1.9E-05 | 2 | 3.04E-05 | 3.204276196 | 8 mo | WILD |
| 1011080637 | 2 | ACAd1 | 935 | 0.62416592 | 0.000439386 | 14 | 0.000703957 | 22.42993337 | 8 mo | HOM |
| 1013609713 | 2 | ACAd1 | 935 | 0.62416592 | 3.98E-05 | 1 | 6.37E-05 | 1.602138098 | 2 mo | WILD |
| 1010418842 | 1 | ACAd1 | 935 | 0.62416592 | 0.000501316 | 15 | 0.000803177 | 24.03207147 | 8 mo | HOM |
| 1013288213 | 2 | ACAd1 | 935 | 0.62416592 | 0.00018613 | 4 | 0.000298205 | 6.408552393 | 4 mo | HOM |
| 1008976060 | 2 | ACAd1 | 935 | 0.62416592 | 4.5E-05 | 2 | 7.22E-05 | 3.204276196 | 8 mo | WILD |
| 1013446243 | 1 | ACAd1 | 935 | 0.62416592 | 0.000228781 | 10 | 0.000366539 | 16.02138098 | 4 mo | HOM |
| 1013288951 | 2 | ACAd1 | 935 | 0.62416592 | 0.000581449 | 14 | 0.000931562 | 22.42993337 | 4 mo | HOM |
| 1013978665 | 1 | DEC | 936 | 3.435540685 | 4.49E-05 | 5 | 1.31E-05 | 1.455374993 | 2 mo | HOM |
| 1013445609 | 1 | DEC | 936 | 3.435540685 | 8.68E-05 | 6 | 2.53E-05 | 1.746449992 | 4 mo | HOM |
| 1013609015 | 1 | DEC | 936 | 3.435540685 | 0.00119138 | 17 | 0.000346781 | 4.948274976 | 4 mo | HOM |
| 1013608106 | 2 | DEC | 936 | 3.435540685 | 0 | 0 | 0 | 0 | 4 mo | HOM |
| 1011937543 | 1 | DEC | 936 | 3.435540685 | 0.006058513 | 26 | 0.001763482 | 7.567949963 | 8 mo | HOM |
| 1010419538 | 2 | DEC | 936 | 3.435540685 | 0.000129478 | 12 | 3.77E-05 | 3.492899983 | 4 mo | WILD |
| 1010419187 | 1 | DEC | 936 | 3.435540685 | 0.004525053 | 32 | 0.00131713 | 9.314399955 | 8 mo | HOM |
| 1013978366 | 1 | DEC | 936 | 3.435540685 | 0.001470272 | 20 | 0.00042796 | 5.821499972 | 2 mo | HOM |
| 1014416155 | 2 | DEC | 936 | 3.435540685 | 0 | 0 | 0 | 0 | 2 mo | HOM |
| 1013978065 | 2 | DEC | 936 | 3.435540685 | 2.72E-05 | 2 | 7.92E-06 | 0.582149997 | 2 mo | HOM |
| 1009934102 | 2 | DEC | 936 | 3.435540685 | 0.000296739 | 24 | 8.64E-05 | 6.985799966 | 4 mo | WILD |
| 1009322937 | 2 | DEC | 936 | 3.435540685 | 4.55E-05 | 3 | 1.32E-05 | 0.873224996 | 8 mo | WILD |
| 1011080637 | 2 | DEC | 936 | 3.435540685 | 0.006961785 | 35 | 0.002026401 | 10.18762495 | 8 mo | HOM |
| 1013609713 | 2 | DEC | 936 | 3.435540685 | 4.27E-05 | 3 | 1.24E-05 | 0.873224996 | 2 mo | WILD |
| 1010418842 | 1 | DEC | 936 | 3.435540685 | 0.001183656 | 21 | 0.000344533 | 6.11257497 | 8 mo | HOM |
| 1013288213 | 2 | DEC | 936 | 3.435540685 | 9.39E-05 | 5 | 2.73E-05 | 1.455374993 | 4 mo | HOM |
| 1008976060 | 2 | DEC | 936 | 3.435540685 | 0.001690353 | 21 | 0.000492019 | 6.11257497 | 8 mo | WILD |
| 1013446243 | 1 | DEC | 936 | 3.435540685 | 0.000677369 | 17 | 0.000197165 | 4.948274976 | 4 mo | HOM |
| 1013288951 | 2 | DEC | 936 | 3.435540685 | 0.001240045 | 12 | 0.000360946 | 3.492899983 | 4 mo | HOM |
| 1013978665 | 1 | PGRN | 938 | 0.975625104 | 0 | 0 | 0 | 0 | 2 mo | HOM |
| 1013445609 | 1 | PGRN | 938 | 0.975625104 | 0 | 0 | 0 | 0 | 4 mo | HOM |
| 1013609015 | 1 | PGRN | 938 | 0.975625104 | 0 | 0 | 0 | 0 | 4 mo | HOM |
| 1013608106 | 2 | PGRN | 938 | 0.975625104 | 0 | 0 | 0 | 0 | 4 mo | HOM |
| 1011937543 | 1 | PGRN | 938 | 0.975625104 | 2.45E-05 | 1 | 2.52E-05 | 1.024983876 | 8 mo | HOM |
| 1010419538 | 2 | PGRN | 938 | 0.975625104 | 0 | 0 | 0 | 0 | 4 mo | WILD |
| 1010419187 | 1 | PGRN | 938 | 0.975625104 | 0 | 0 | 0 | 0 | 8 mo | HOM |
| 1013978366 | 1 | PGRN | 938 | 0.975625104 | 0 | 0 | 0 | 0 | 2 mo | HOM |
| 1014416155 | 2 | PGRN | 938 | 0.975625104 | 0 | 0 | 0 | 0 | 2 mo | HOM |
| 1013978065 | 2 | PGRN | 938 | 0.975625104 | 0 | 0 | 0 | 0 | 2 mo | HOM |
| 1009934102 | 2 | PGRN | 938 | 0.975625104 | 0 | 0 | 0 | 0 | 4 mo | WILD |
| 1009322937 | 2 | PGRN | 938 | 0.975625104 | 0 | 0 | 0 | 0 | 8 mo | WILD |
| 1011080637 | 2 | PGRN | 938 | 0.975625104 | 0 | 0 | 0 | 0 | 8 mo | HOM |
| 1013609713 | 2 | PGRN | 938 | 0.975625104 | 0 | 0 | 0 | 0 | 2 mo | WILD |
| 1010418842 | 1 | PGRN | 938 | 0.975625104 | 1.16E-06 | 0 | 1.19E-06 | 0 | 8 mo | HOM |
| 1013288213 | 2 | PGRN | 938 | 0.975625104 | 0 | 0 | 0 | 0 | 4 mo | HOM |
| 1008976060 | 2 | PGRN | 938 | 0.975625104 | 2.92E-06 | 0 | 3E-06 | 0 | 8 mo | WILD |
| 1013446243 | 1 | PGRN | 938 | 0.975625104 | 0 | 0 | 0 | 0 | 4 mo | HOM |
| 1013288951 | 2 | PGRN | 938 | 0.975625104 | 0 | 0 | 0 | 0 | 4 mo | HOM |
| 1013978665 | 1 | AMBd | 939 | 0.021849012 | 0 | 0 | 0 | 0 | 2 mo | HOM |
| 1013445609 | 1 | AMBd | 939 | 0.021849012 | 0 | 0 | 0 | 0 | 4 mo | HOM |
| 1013609015 | 1 | AMBd | 939 | 0.021849012 | 0 | 0 | 0 | 0 | 4 mo | HOM |
| 1013608106 | 2 | AMBd | 939 | 0.021849012 | 0 | 0 | 0 | 0 | 4 mo | HOM |
| 1011937543 | 1 | AMBd | 939 | 0.021849012 | 0 | 0 | 0 | 0 | 8 mo | HOM |
| 1010419538 | 2 | AMBd | 939 | 0.021849012 | 0 | 0 | 0 | 0 | 4 mo | WILD |
| 1010419187 | 1 | AMBd | 939 | 0.021849012 | 0 | 0 | 0 | 0 | 8 mo | HOM |
| 1013978366 | 1 | AMBd | 939 | 0.021849012 | 0 | 0 | 0 | 0 | 2 mo | HOM |

| | | | | | | | | | | |
|---|---|---|---|---|---|---|---|---|---|---|
| **1014416155** | 2 | AMBd | 939 | 0.021849012 | 0 | 0 | 0 | 0 | 2 mo | HOM |
| **1013978065** | 2 | AMBd | 939 | 0.021849012 | 0 | 0 | 0 | 0 | 2 mo | HOM |
| **1009934102** | 2 | AMBd | 939 | 0.021849012 | 0 | 0 | 0 | 0 | 4 mo | WILD |
| **1009322937** | 2 | AMBd | 939 | 0.021849012 | 0 | 0 | 0 | 0 | 8 mo | WILD |
| **1011080637** | 2 | AMBd | 939 | 0.021849012 | 0 | 0 | 0 | 0 | 8 mo | HOM |
| **1013609713** | 2 | AMBd | 939 | 0.021849012 | 0 | 0 | 0 | 0 | 2 mo | WILD |
| **1010418842** | 1 | AMBd | 939 | 0.021849012 | 0 | 0 | 0 | 0 | 8 mo | HOM |
| **1013288213** | 2 | AMBd | 939 | 0.021849012 | 0 | 0 | 0 | 0 | 4 mo | HOM |
| **1008976060** | 2 | AMBd | 939 | 0.021849012 | 0 | 0 | 0 | 0 | 8 mo | WILD |
| **1013446243** | 1 | AMBd | 939 | 0.021849012 | 0 | 0 | 0 | 0 | 4 mo | HOM |
| **1013288951** | 2 | AMBd | 939 | 0.021849012 | 0 | 0 | 0 | 0 | 4 mo | HOM |
| **1013978665** | 1 | cing | 940 | 1.304001093 | 0 | 0 | 0 | 0 | 2 mo | HOM |
| **1013445609** | 1 | cing | 940 | 1.304001093 | 0 | 0 | 0 | 0 | 4 mo | HOM |
| **1013609015** | 1 | cing | 940 | 1.304001093 | 3.99E-06 | 0 | 3.06E-06 | 0 | 4 mo | HOM |
| **1013608106** | 2 | cing | 940 | 1.304001093 | 0 | 0 | 0 | 0 | 4 mo | HOM |
| **1011937543** | 1 | cing | 940 | 1.304001093 | 0.001493935 | 15 | 0.001145655 | 11.50305784 | 8 mo | HOM |
| **1010419538** | 2 | cing | 940 | 1.304001093 | 0 | 0 | 0 | 0 | 4 mo | WILD |
| **1010419187** | 1 | cing | 940 | 1.304001093 | 0.000173377 | 16 | 0.000132958 | 12.26992837 | 8 mo | HOM |
| **1013978366** | 1 | cing | 940 | 1.304001093 | 0 | 0 | 0 | 0 | 2 mo | HOM |
| **1014416155** | 2 | cing | 940 | 1.304001093 | 0 | 0 | 0 | 0 | 2 mo | HOM |
| **1013978065** | 2 | cing | 940 | 1.304001093 | 0 | 0 | 0 | 0 | 2 mo | HOM |
| **1009934102** | 2 | cing | 940 | 1.304001093 | 0 | 0 | 0 | 0 | 4 mo | WILD |
| **1009322937** | 2 | cing | 940 | 1.304001093 | 0 | 0 | 0 | 0 | 8 mo | WILD |
| **1011080637** | 2 | cing | 940 | 1.304001093 | 0.000141765 | 10 | 0.000108715 | 7.66870523 | 8 mo | HOM |
| **1013609713** | 2 | cing | 940 | 1.304001093 | 0 | 0 | 0 | 0 | 2 mo | WILD |
| **1010418842** | 1 | cing | 940 | 1.304001093 | 0.00040502 | 16 | 0.000310598 | 12.26992837 | 8 mo | HOM |
| **1013288213** | 2 | cing | 940 | 1.304001093 | 0 | 0 | 0 | 0 | 4 mo | HOM |
| **1008976060** | 2 | cing | 940 | 1.304001093 | 1.39E-05 | 1 | 1.06E-05 | 0.766870523 | 8 mo | WILD |
| **1013446243** | 1 | cing | 940 | 1.304001093 | 8.44E-06 | 1 | 6.48E-06 | 0.766870523 | 4 mo | HOM |
| **1013288951** | 2 | cing | 940 | 1.304001093 | 0.000143943 | 1 | 0.000110386 | 0.766870523 | 4 mo | HOM |
| **1013978665** | 1 | EP | 942 | 2.875703219 | 0 | 0 | 0 | 0 | 2 mo | HOM |
| **1013445609** | 1 | EP | 942 | 2.876907365 | 0.000129049 | 12 | 4.49E-05 | 4.171145775 | 4 mo | HOM |
| **1013609015** | 1 | EP | 942 | 2.876907365 | 0.000300514 | 15 | 0.000104457 | 5.213932219 | 4 mo | HOM |
| **1013608106** | 2 | EP | 942 | 2.876907365 | 0 | 0 | 0 | 0 | 4 mo | HOM |
| **1011937543** | 1 | EP | 942 | 2.876907365 | 0.003697777 | 99 | 0.001285331 | 34.41195264 | 8 mo | HOM |
| **1010419538** | 2 | EP | 942 | 2.876907365 | 0 | 0 | 0 | 0 | 4 mo | WILD |
| **1010419187** | 1 | EP | 942 | 2.876907365 | 0.002898229 | 139 | 0.001007411 | 48.31577189 | 8 mo | HOM |
| **1013978366** | 1 | EP | 942 | 2.876907365 | 0 | 0 | 0 | 0 | 2 mo | HOM |
| **1014416155** | 2 | EP | 942 | 2.876907365 | 0 | 0 | 0 | 0 | 2 mo | HOM |
| **1013978065** | 2 | EP | 942 | 2.876907365 | 0 | 0 | 0 | 0 | 2 mo | HOM |
| **1009934102** | 2 | EP | 942 | 2.876907365 | 0 | 0 | 0 | 0 | 4 mo | WILD |
| **1009322937** | 2 | EP | 942 | 2.876907365 | 0 | 0 | 0 | 0 | 8 mo | WILD |
| **1011080637** | 2 | EP | 942 | 2.876907365 | 0.002555623 | 122 | 0.000888323 | 42.40664871 | 8 mo | HOM |
| **1013609713** | 2 | EP | 942 | 2.876907365 | 0 | 0 | 0 | 0 | 2 mo | WILD |
| **1010418842** | 1 | EP | 942 | 2.876907365 | 0.004401437 | 145 | 0.00152992 | 50.40134478 | 8 mo | HOM |
| **1013288213** | 2 | EP | 942 | 2.876907365 | 7.45E-05 | 7 | 2.59E-05 | 2.433168369 | 4 mo | HOM |
| **1008976060** | 2 | EP | 942 | 2.876907365 | 2.34E-05 | 5 | 8.13E-06 | 1.737977406 | 8 mo | WILD |
| **1013446243** | 1 | EP | 942 | 2.876907365 | 0.000284521 | 20 | 9.89E-05 | 6.951909625 | 4 mo | HOM |
| **1013288951** | 2 | EP | 942 | 2.876907365 | 0.000311361 | 25 | 0.000108228 | 8.689887031 | 4 mo | HOM |
| **1013978665** | 1 | MOp2/3 | 943 | 3.800800954 | 0 | 0 | 0 | 0 | 2 mo | HOM |
| **1013445609** | 1 | MOp2/3 | 943 | 3.800800954 | 0.000346778 | 19 | 9.12E-05 | 4.998946336 | 4 mo | HOM |
| **1013609015** | 1 | MOp2/3 | 943 | 3.800800954 | 0.000784376 | 32 | 0.000206371 | 8.419278039 | 4 mo | HOM |
| **1013608106** | 2 | MOp2/3 | 943 | 3.800800954 | 0 | 0 | 0 | 0 | 4 mo | HOM |
| **1011937543** | 1 | MOp2/3 | 943 | 3.800800954 | 0.003540391 | 63 | 0.000931486 | 16.57545364 | 8 mo | HOM |
| **1010419538** | 2 | MOp2/3 | 943 | 3.800800954 | 0 | 0 | 0 | 0 | 4 mo | WILD |
| **1010419187** | 1 | MOp2/3 | 943 | 3.800800954 | 0.005231799 | 162 | 0.001376499 | 42.62259507 | 8 mo | HOM |
| **1013978366** | 1 | MOp2/3 | 943 | 3.800800954 | 5.64E-06 | 1 | 1.48E-06 | 0.263102439 | 2 mo | HOM |
| **1014416155** | 2 | MOp2/3 | 943 | 3.800800954 | 0 | 0 | 0 | 0 | 2 mo | HOM |
| **1013978065** | 2 | MOp2/3 | 943 | 3.800800954 | 0 | 0 | 0 | 0 | 2 mo | HOM |

| | | | | | | | | | | |
|---|---|---|---|---|---|---|---|---|---|---|
| **1009934102** | 2 | MOp2/3 | 943 | 3.800800954 | 0 | 0 | 0 | 0 | 4 mo | WILD |
| **1009322937** | 2 | MOp2/3 | 943 | 3.800800954 | 0 | 0 | 0 | 0 | 8 mo | WILD |
| **1011080637** | 2 | MOp2/3 | 943 | 3.800800954 | 0.001958491 | 84 | 0.000515284 | 22.10060485 | 8 mo | HOM |
| **1013609713** | 2 | MOp2/3 | 943 | 3.800800954 | 0 | 0 | 0 | 0 | 2 mo | WILD |
| **1010418842** | 1 | MOp2/3 | 943 | 3.800800954 | 0.002437855 | 125 | 0.000641405 | 32.88780484 | 8 mo | HOM |
| **1013288213** | 2 | MOp2/3 | 943 | 3.800800954 | 0.00036443 | 22 | 9.59E-05 | 5.788253652 | 4 mo | HOM |
| **1008976060** | 2 | MOp2/3 | 943 | 3.800800954 | 1.78E-05 | 1 | 4.7E-06 | 0.263102439 | 8 mo | WILD |
| **1013446243** | 1 | MOp2/3 | 943 | 3.800800954 | 0.00035939 | 19 | 9.46E-05 | 4.998946336 | 4 mo | HOM |
| **1013288951** | 2 | MOp2/3 | 943 | 3.800800954 | 0.000665035 | 28 | 0.000174972 | 7.366868284 | 4 mo | HOM |
| **1013978665** | 1 | FOTU | 944 | 1.085236947 | 5.09E-05 | 2 | 4.69E-05 | 1.842915508 | 2 mo | HOM |
| **1013445609** | 1 | FOTU | 944 | 1.085236947 | 1.66E-05 | 1 | 1.53E-05 | 0.921457754 | 4 mo | HOM |
| **1013609015** | 1 | FOTU | 944 | 1.085236947 | 0.000105434 | 11 | 9.72E-05 | 10.13603529 | 4 mo | HOM |
| **1013608106** | 2 | FOTU | 944 | 1.085236947 | 0 | 0 | 0 | 0 | 4 mo | HOM |
| **1011937543** | 1 | FOTU | 944 | 1.085236947 | 0.001682788 | 15 | 0.001550618 | 13.82186631 | 8 mo | HOM |
| **1010419538** | 2 | FOTU | 944 | 1.085236947 | 9.55E-06 | 3 | 8.8E-06 | 2.764373262 | 4 mo | WILD |
| **1010419187** | 1 | FOTU | 944 | 1.085236947 | 0.002581675 | 9 | 0.002378904 | 8.293119787 | 8 mo | HOM |
| **1013978366** | 1 | FOTU | 944 | 1.085236947 | 0.000733729 | 16 | 0.0006761 | 14.74332407 | 2 mo | HOM |
| **1014416155** | 2 | FOTU | 944 | 1.085236947 | 0 | 0 | 0 | 0 | 2 mo | HOM |
| **1013978065** | 2 | FOTU | 944 | 1.085236947 | 2.49E-09 | 0 | 2.3E-09 | 0 | 2 mo | HOM |
| **1009934102** | 2 | FOTU | 944 | 1.085236947 | 6.55E-05 | 8 | 6.04E-05 | 7.371662033 | 4 mo | WILD |
| **1009322937** | 2 | FOTU | 944 | 1.085236947 | 2.09E-05 | 2 | 1.93E-05 | 1.842915508 | 8 mo | WILD |
| **1011080637** | 2 | FOTU | 944 | 1.085236947 | 0.002055879 | 14 | 0.001894406 | 12.90040856 | 8 mo | HOM |
| **1013609713** | 2 | FOTU | 944 | 1.085236947 | 3.03E-05 | 2 | 2.79E-05 | 1.842915508 | 2 mo | WILD |
| **1010418842** | 1 | FOTU | 944 | 1.085236947 | 0.00196627 | 21 | 0.001811835 | 19.35061284 | 8 mo | HOM |
| **1013288213** | 2 | FOTU | 944 | 1.085236947 | 2.94E-05 | 2 | 2.71E-05 | 1.842915508 | 4 mo | HOM |
| **1008976060** | 2 | FOTU | 944 | 1.085236947 | 0.000143216 | 9 | 0.000131968 | 8.293119787 | 8 mo | WILD |
| **1013446243** | 1 | FOTU | 944 | 1.085236947 | 0.000298862 | 6 | 0.000275389 | 5.528746524 | 4 mo | HOM |
| **1013288951** | 2 | FOTU | 944 | 1.085236947 | 0.003678447 | 8 | 0.003389534 | 7.371662033 | 4 mo | HOM |
| **1013978665** | 1 | SSp-ul6a | 945 | 0.970486395 | 0 | 0 | 0 | 0 | 2 mo | HOM |
| **1013445609** | 1 | SSp-ul6a | 945 | 0.970486395 | 0.000106915 | 5 | 0.000110166 | 5.152055737 | 4 mo | HOM |
| **1013609015** | 1 | SSp-ul6a | 945 | 0.970486395 | 0.00020743 | 11 | 0.000213738 | 11.33452262 | 4 mo | HOM |
| **1013608106** | 2 | SSp-ul6a | 945 | 0.970486395 | 0 | 0 | 0 | 0 | 4 mo | HOM |
| **1011937543** | 1 | SSp-ul6a | 945 | 0.970486395 | 0.001403584 | 28 | 0.001446268 | 28.85151213 | 8 mo | HOM |
| **1010419538** | 2 | SSp-ul6a | 945 | 0.970486395 | 0 | 0 | 0 | 0 | 4 mo | WILD |
| **1010419187** | 1 | SSp-ul6a | 945 | 0.970486395 | 0.001326446 | 54 | 0.001366785 | 55.64220196 | 8 mo | HOM |
| **1013978366** | 1 | SSp-ul6a | 945 | 0.970486395 | 0 | 0 | 0 | 0 | 2 mo | HOM |
| **1014416155** | 2 | SSp-ul6a | 945 | 0.970486395 | 0 | 0 | 0 | 0 | 2 mo | HOM |
| **1013978065** | 2 | SSp-ul6a | 945 | 0.970486395 | 0 | 0 | 0 | 0 | 2 mo | HOM |
| **1009934102** | 2 | SSp-ul6a | 945 | 0.970486395 | 0 | 0 | 0 | 0 | 4 mo | WILD |
| **1009322937** | 2 | SSp-ul6a | 945 | 0.970486395 | 0 | 0 | 0 | 0 | 8 mo | WILD |
| **1011080637** | 2 | SSp-ul6a | 945 | 0.970486395 | 0.000785363 | 31 | 0.000809247 | 31.94274557 | 8 mo | HOM |
| **1013609713** | 2 | SSp-ul6a | 945 | 0.970486395 | 0 | 0 | 0 | 0 | 2 mo | WILD |
| **1010418842** | 1 | SSp-ul6a | 945 | 0.970486395 | 0.000952704 | 45 | 0.000981677 | 46.36850163 | 8 mo | HOM |
| **1013288213** | 2 | SSp-ul6a | 945 | 0.970486395 | 4.21E-05 | 5 | 4.33E-05 | 5.152055737 | 4 mo | HOM |
| **1008976060** | 2 | SSp-ul6a | 945 | 0.970486395 | 1.65E-05 | 2 | 1.7E-05 | 2.060822295 | 8 mo | WILD |
| **1013446243** | 1 | SSp-ul6a | 945 | 0.970486395 | 0.000114084 | 12 | 0.000117553 | 12.36493377 | 4 mo | HOM |
| **1013288951** | 2 | SSp-ul6a | 945 | 0.970486395 | 0.000170285 | 11 | 0.000175463 | 11.33452262 | 4 mo | HOM |
| **1013978665** | 1 | PH | 946 | 0.728486528 | 0 | 0 | 0 | 0 | 2 mo | HOM |
| **1013445609** | 1 | PH | 946 | 0.728486528 | 0 | 0 | 0 | 0 | 4 mo | HOM |
| **1013609015** | 1 | PH | 946 | 0.728486528 | 0 | 0 | 0 | 0 | 4 mo | HOM |
| **1013608106** | 2 | PH | 946 | 0.728486528 | 0 | 0 | 0 | 0 | 4 mo | HOM |
| **1011937543** | 1 | PH | 946 | 0.728486528 | 0.000257291 | 9 | 0.000353186 | 12.35438083 | 8 mo | HOM |
| **1010419538** | 2 | PH | 946 | 0.728486528 | 0 | 0 | 0 | 0 | 4 mo | WILD |
| **1010419187** | 1 | PH | 946 | 0.728486528 | 5.11E-05 | 4 | 7.02E-05 | 5.490835927 | 8 mo | HOM |
| **1013978366** | 1 | PH | 946 | 0.728486528 | 0 | 0 | 0 | 0 | 2 mo | HOM |
| **1014416155** | 2 | PH | 946 | 0.728486528 | 0 | 0 | 0 | 0 | 2 mo | HOM |
| **1013978065** | 2 | PH | 946 | 0.728486528 | 0 | 0 | 0 | 0 | 2 mo | HOM |
| **1009934102** | 2 | PH | 946 | 0.728486528 | 0 | 0 | 0 | 0 | 4 mo | WILD |
| **1009322937** | 2 | PH | 946 | 0.728486528 | 0 | 0 | 0 | 0 | 8 mo | WILD |

| | | | | | | | | | | |
|---|---|---|---|---|---|---|---|---|---|---|
| **1011080637** | 2 | PH | 946 | 0.728486528 | 6.06E-05 | 4 | 8.33E-05 | 5.490835927 | 8 mo | HOM |
| **1013609713** | 2 | PH | 946 | 0.728486528 | 0 | 0 | 0 | 0 | 2 mo | WILD |
| **1010418842** | 1 | PH | 946 | 0.728486528 | 0.000118297 | 7 | 0.000162387 | 9.608962872 | 8 mo | HOM |
| **1013288213** | 2 | PH | 946 | 0.728486528 | 0 | 0 | 0 | 0 | 4 mo | HOM |
| **1008976060** | 2 | PH | 946 | 0.728486528 | 4.24E-05 | 1 | 5.82E-05 | 1.372708982 | 8 mo | WILD |
| **1013446243** | 1 | PH | 946 | 0.728486528 | 0 | 0 | 0 | 0 | 4 mo | HOM |
| **1013288951** | 2 | PH | 946 | 0.728486528 | 0 | 0 | 0 | 0 | 4 mo | HOM |
| **1013978665** | 1 | cVIIIn | 948 | 1.453719593 | 3.92E-07 | 0 | 2.7E-07 | 0 | 2 mo | HOM |
| **1013445609** | 1 | cVIIIn | 948 | 1.453719593 | 5.08E-06 | 1 | 3.5E-06 | 0.68789057 | 4 mo | HOM |
| **1013609015** | 1 | cVIIIn | 948 | 1.453719593 | 2.07E-05 | 0 | 1.42E-05 | 0 | 4 mo | HOM |
| **1013608106** | 2 | cVIIIn | 948 | 1.453719593 | 0 | 0 | 0 | 0 | 4 mo | HOM |
| **1011937543** | 1 | cVIIIn | 948 | 1.453719593 | 0.001789081 | 17 | 0.001230692 | 11.69413969 | 8 mo | HOM |
| **1010419538** | 2 | cVIIIn | 948 | 1.453719593 | 0 | 0 | 0 | 0 | 4 mo | WILD |
| **1010419187** | 1 | cVIIIn | 948 | 1.453719593 | 0.002323354 | 2 | 0.001598213 | 1.375781141 | 8 mo | HOM |
| **1013978366** | 1 | cVIIIn | 948 | 1.453719593 | 3.92E-05 | 1 | 2.69E-05 | 0.68789057 | 2 mo | HOM |
| **1014416155** | 2 | cVIIIn | 948 | 1.453719593 | 0 | 0 | 0 | 0 | 2 mo | HOM |
| **1013978065** | 2 | cVIIIn | 948 | 1.453719593 | 0 | 0 | 0 | 0 | 2 mo | HOM |
| **1009934102** | 2 | cVIIIn | 948 | 1.453719593 | 1.14E-06 | 0 | 7.87E-07 | 0 | 4 mo | WILD |
| **1009322937** | 2 | cVIIIn | 948 | 1.453719593 | 4.9E-06 | 0 | 3.37E-06 | 0 | 8 mo | WILD |
| **1011080637** | 2 | cVIIIn | 948 | 1.453719593 | 6.49E-06 | 0 | 4.47E-06 | 0 | 8 mo | HOM |
| **1013609713** | 2 | cVIIIn | 948 | 1.453719593 | 0 | 0 | 0 | 0 | 2 mo | WILD |
| **1010418842** | 1 | cVIIIn | 948 | 1.453719593 | 0.000404096 | 4 | 0.000277974 | 2.751562281 | 8 mo | HOM |
| **1013288213** | 2 | cVIIIn | 948 | 1.453719593 | 8.77E-08 | 0 | 6.03E-08 | 0 | 4 mo | HOM |
| **1008976060** | 2 | cVIIIn | 948 | 1.453719593 | 5.7E-05 | 4 | 3.92E-05 | 2.751562281 | 8 mo | WILD |
| **1013446243** | 1 | cVIIIn | 948 | 1.453719593 | 1.36E-06 | 0 | 9.34E-07 | 0 | 4 mo | HOM |
| **1013288951** | 2 | cVIIIn | 948 | 1.453719593 | 0 | 0 | 0 | 0 | 4 mo | HOM |
| **1013978665** | 1 | von | 949 | 0.013833861 | 1.86E-08 | 0 | 1.35E-06 | 0 | 2 mo | HOM |
| **1013445609** | 1 | von | 949 | 0.013833861 | 0 | 0 | 0 | 0 | 4 mo | HOM |
| **1013609015** | 1 | von | 949 | 0.013833861 | 8.47E-07 | 0 | 6.12E-05 | 0 | 4 mo | HOM |
| **1013608106** | 2 | von | 949 | 0.013833861 | 0 | 0 | 0 | 0 | 4 mo | HOM |
| **1011937543** | 1 | von | 949 | 0.013833861 | 3.61E-05 | 1 | 0.002606127 | 72.28639772 | 8 mo | HOM |
| **1010419538** | 2 | von | 949 | 0.013833861 | 0 | 0 | 0 | 0 | 4 mo | WILD |
| **1010419187** | 1 | von | 949 | 0.013833861 | 4.15E-05 | 2 | 0.00299686 | 144.5727954 | 8 mo | HOM |
| **1013978366** | 1 | von | 949 | 0.013833861 | 0 | 0 | 0 | 0 | 2 mo | HOM |
| **1014416155** | 2 | von | 949 | 0.013833861 | 0 | 0 | 0 | 0 | 2 mo | HOM |
| **1013978065** | 2 | von | 949 | 0.013833861 | 4.28E-06 | 1 | 0.000309069 | 72.28639772 | 2 mo | HOM |
| **1009934102** | 2 | von | 949 | 0.013833861 | 0 | 0 | 0 | 0 | 4 mo | WILD |
| **1009322937** | 2 | von | 949 | 0.013833861 | 0 | 0 | 0 | 0 | 8 mo | WILD |
| **1011080637** | 2 | von | 949 | 0.013833861 | 1.06E-05 | 1 | 0.000768856 | 72.28639772 | 8 mo | HOM |
| **1013609713** | 2 | von | 949 | 0.013833861 | 0 | 0 | 0 | 0 | 2 mo | WILD |
| **1010418842** | 1 | von | 949 | 0.013833861 | 2.65E-05 | 3 | 0.00191642 | 216.8591932 | 8 mo | HOM |
| **1013288213** | 2 | von | 949 | 0.013833861 | 0.000134765 | 3 | 0.009741706 | 216.8591932 | 4 mo | HOM |
| **1008976060** | 2 | von | 949 | 0.013833861 | 0 | 0 | 0 | 0 | 8 mo | WILD |
| **1013446243** | 1 | von | 949 | 0.013833861 | 2.74E-06 | 1 | 0.000198423 | 72.28639772 | 4 mo | HOM |
| **1013288951** | 2 | von | 949 | 0.013833861 | 2.86E-05 | 1 | 0.002066057 | 72.28639772 | 4 mo | HOM |
| **1013978665** | 1 | SSp-m4 | 950 | 0.977425971 | 0 | 0 | 0 | 0 | 2 mo | HOM |
| **1013445609** | 1 | SSp-m4 | 950 | 0.977425971 | 0.000121748 | 7 | 0.000124559 | 7.161667693 | 4 mo | HOM |
| **1013609015** | 1 | SSp-m4 | 950 | 0.977425971 | 0.000133954 | 10 | 0.000137047 | 10.23095385 | 4 mo | HOM |
| **1013608106** | 2 | SSp-m4 | 950 | 0.977425971 | 0 | 0 | 0 | 0 | 4 mo | HOM |
| **1011937543** | 1 | SSp-m4 | 950 | 0.977425971 | 0.001034716 | 26 | 0.001058613 | 26.60048 | 8 mo | HOM |
| **1010419538** | 2 | SSp-m4 | 950 | 0.977425971 | 0 | 0 | 0 | 0 | 4 mo | WILD |
| **1010419187** | 1 | SSp-m4 | 950 | 0.977425971 | 0.001451903 | 45 | 0.001485435 | 46.03929231 | 8 mo | HOM |
| **1013978366** | 1 | SSp-m4 | 950 | 0.977425971 | 7.55E-07 | 0 | 7.73E-07 | 0 | 2 mo | HOM |
| **1014416155** | 2 | SSp-m4 | 950 | 0.977425971 | 0 | 0 | 0 | 0 | 2 mo | HOM |
| **1013978065** | 2 | SSp-m4 | 950 | 0.977425971 | 0 | 0 | 0 | 0 | 2 mo | HOM |
| **1009934102** | 2 | SSp-m4 | 950 | 0.977425971 | 9.92E-06 | 1 | 1.01E-05 | 1.023095385 | 4 mo | WILD |
| **1009322937** | 2 | SSp-m4 | 950 | 0.977425971 | 8.25E-06 | 1 | 8.45E-06 | 1.023095385 | 8 mo | WILD |
| **1011080637** | 2 | SSp-m4 | 950 | 0.977425971 | 0.000726847 | 28 | 0.000743633 | 28.64667077 | 8 mo | HOM |
| **1013609713** | 2 | SSp-m4 | 950 | 0.977425971 | 8.35E-06 | 1 | 8.54E-06 | 1.023095385 | 2 mo | WILD |

| | | | | | | | | | | |
|---|---|---|---|---|---|---|---|---|---|---|
| **1010418842** | 1 | SSp-m4 | 950 | 0.977425971 | 0.001096288 | 46 | 0.001121607 | 47.0623877 | 8 mo | HOM |
| **1013288213** | 2 | SSp-m4 | 950 | 0.977425971 | 0.000106659 | 9 | 0.000109122 | 9.207858462 | 4 mo | HOM |
| **1008976060** | 2 | SSp-m4 | 950 | 0.977425971 | 3.26E-05 | 1 | 3.34E-05 | 1.023095385 | 8 mo | WILD |
| **1013446243** | 1 | SSp-m4 | 950 | 0.977425971 | 0.000265528 | 13 | 0.00027166 | 13.30024 | 4 mo | HOM |
| **1013288951** | 2 | SSp-m4 | 950 | 0.977425971 | 0.000110234 | 7 | 0.00011278 | 7.161667693 | 4 mo | HOM |
| **1013978665** | 1 | PYR | 951 | 1.285739381 | 5.11E-05 | 5 | 3.97E-05 | 3.888812985 | 2 mo | HOM |
| **1013445609** | 1 | PYR | 951 | 1.285739381 | 2.12E-10 | 0 | 1.65E-10 | 0 | 4 mo | HOM |
| **1013609015** | 1 | PYR | 951 | 1.285739381 | 8.85E-05 | 10 | 6.88E-05 | 7.777625971 | 4 mo | HOM |
| **1013608106** | 2 | PYR | 951 | 1.285739381 | 0 | 0 | 0 | 0 | 4 mo | HOM |
| **1011937543** | 1 | PYR | 951 | 1.285739381 | 0.00174749 | 5 | 0.001359132 | 3.888812985 | 8 mo | HOM |
| **1010419538** | 2 | PYR | 951 | 1.285739381 | 9.27E-06 | 1 | 7.21E-06 | 0.777762597 | 4 mo | WILD |
| **1010419187** | 1 | PYR | 951 | 1.285739381 | 0.000518887 | 17 | 0.000403571 | 13.22196415 | 8 mo | HOM |
| **1013978366** | 1 | PYR | 951 | 1.285739381 | 0.000691724 | 7 | 0.000537997 | 5.44433818 | 2 mo | HOM |
| **1014416155** | 2 | PYR | 951 | 1.285739381 | 0 | 0 | 0 | 0 | 2 mo | HOM |
| **1013978065** | 2 | PYR | 951 | 1.285739381 | 0.00012389 | 7 | 9.64E-05 | 5.44433818 | 2 mo | HOM |
| **1009934102** | 2 | PYR | 951 | 1.285739381 | 3.33E-05 | 5 | 2.59E-05 | 3.888812985 | 4 mo | WILD |
| **1009322937** | 2 | PYR | 951 | 1.285739381 | 1.58E-05 | 2 | 1.23E-05 | 1.555525194 | 8 mo | WILD |
| **1011080637** | 2 | PYR | 951 | 1.285739381 | 0.000885025 | 8 | 0.000688339 | 6.222100777 | 8 mo | HOM |
| **1013609713** | 2 | PYR | 951 | 1.276385398 | 0.000207695 | 2 | 0.000162721 | 1.566924851 | 2 mo | WILD |
| **1010418842** | 1 | PYR | 951 | 1.285739381 | 0.001404952 | 24 | 0.001092719 | 18.66630233 | 8 mo | HOM |
| **1013288213** | 2 | PYR | 951 | 1.285739381 | 1.45E-05 | 2 | 1.13E-05 | 1.555525194 | 4 mo | HOM |
| **1008976060** | 2 | PYR | 951 | 1.285739381 | 7.14E-05 | 6 | 5.55E-05 | 4.666575583 | 8 mo | WILD |
| **1013446243** | 1 | PYR | 951 | 1.285739381 | 3.16E-05 | 5 | 2.46E-05 | 3.888812985 | 4 mo | HOM |
| **1013288951** | 2 | PYR | 951 | 1.285739381 | 0.001061708 | 10 | 0.000825757 | 7.777625971 | 4 mo | HOM |
| **1013978665** | 1 | EPd | 952 | 1.874784845 | 0 | 0 | 0 | 0 | 2 mo | HOM |
| **1013445609** | 1 | EPd | 952 | 1.874784845 | 9.3E-05 | 9 | 4.96E-05 | 4.800550861 | 4 mo | HOM |
| **1013609015** | 1 | EPd | 952 | 1.874784845 | 0.000159438 | 7 | 8.5E-05 | 3.73376178 | 4 mo | HOM |
| **1013608106** | 2 | EPd | 952 | 1.874784845 | 0 | 0 | 0 | 0 | 4 mo | HOM |
| **1011937543** | 1 | EPd | 952 | 1.874784845 | 0.002234347 | 60 | 0.001191788 | 32.0036724 | 8 mo | HOM |
| **1010419538** | 2 | EPd | 952 | 1.874784845 | 0 | 0 | 0 | 0 | 4 mo | WILD |
| **1010419187** | 1 | EPd | 952 | 1.874784845 | 0.002102414 | 93 | 0.001121416 | 49.60569223 | 8 mo | HOM |
| **1013978366** | 1 | EPd | 952 | 1.874784845 | 0 | 0 | 0 | 0 | 2 mo | HOM |
| **1014416155** | 2 | EPd | 952 | 1.874784845 | 0 | 0 | 0 | 0 | 2 mo | HOM |
| **1013978065** | 2 | EPd | 952 | 1.874784845 | 0 | 0 | 0 | 0 | 2 mo | HOM |
| **1009934102** | 2 | EPd | 952 | 1.874784845 | 0 | 0 | 0 | 0 | 4 mo | WILD |
| **1009322937** | 2 | EPd | 952 | 1.874784845 | 0 | 0 | 0 | 0 | 8 mo | WILD |
| **1011080637** | 2 | EPd | 952 | 1.874784845 | 0.001510728 | 75 | 0.000805814 | 40.0045905 | 8 mo | HOM |
| **1013609713** | 2 | EPd | 952 | 1.874784845 | 0 | 0 | 0 | 0 | 2 mo | WILD |
| **1010418842** | 1 | EPd | 952 | 1.874784845 | 0.002931466 | 81 | 0.001563628 | 43.20495774 | 8 mo | HOM |
| **1013288213** | 2 | EPd | 952 | 1.874784845 | 5.1E-05 | 4 | 2.72E-05 | 2.13357816 | 4 mo | HOM |
| **1008976060** | 2 | EPd | 952 | 1.874784845 | 5.45E-06 | 1 | 2.91E-06 | 0.53339454 | 8 mo | WILD |
| **1013446243** | 1 | EPd | 952 | 1.874784845 | 0.000149565 | 8 | 7.98E-05 | 4.26715632 | 4 mo | HOM |
| **1013288951** | 2 | EPd | 952 | 1.874784845 | 0.000174496 | 14 | 9.31E-05 | 7.467523561 | 4 mo | HOM |
| **1013978665** | 1 | AUDp6a | 954 | 0.306735061 | 0 | 0 | 0 | 0 | 2 mo | HOM |
| **1013445609** | 1 | AUDp6a | 954 | 0.306735061 | 1.04E-05 | 1 | 3.39E-05 | 3.260142474 | 4 mo | HOM |
| **1013609015** | 1 | AUDp6a | 954 | 0.306735061 | 3.2E-05 | 3 | 0.000104364 | 9.780427422 | 4 mo | HOM |
| **1013608106** | 2 | AUDp6a | 954 | 0.306735061 | 0 | 0 | 0 | 0 | 4 mo | HOM |
| **1011937543** | 1 | AUDp6a | 954 | 0.306735061 | 0.000565023 | 10 | 0.001842057 | 32.60142474 | 8 mo | HOM |
| **1010419538** | 2 | AUDp6a | 954 | 0.306735061 | 0 | 0 | 0 | 0 | 4 mo | WILD |
| **1010419187** | 1 | AUDp6a | 954 | 0.306735061 | 0.000371663 | 13 | 0.001211675 | 42.38185216 | 8 mo | HOM |
| **1013978366** | 1 | AUDp6a | 954 | 0.306735061 | 0 | 0 | 0 | 0 | 2 mo | HOM |
| **1014416155** | 2 | AUDp6a | 954 | 0.306735061 | 0 | 0 | 0 | 0 | 2 mo | HOM |
| **1013978065** | 2 | AUDp6a | 954 | 0.306735061 | 0 | 0 | 0 | 0 | 2 mo | HOM |
| **1009934102** | 2 | AUDp6a | 954 | 0.306735061 | 0 | 0 | 0 | 0 | 4 mo | WILD |
| **1009322937** | 2 | AUDp6a | 954 | 0.306735061 | 0 | 0 | 0 | 0 | 8 mo | WILD |
| **1011080637** | 2 | AUDp6a | 954 | 0.306735061 | 0.000281043 | 15 | 0.00091624 | 48.90213711 | 8 mo | HOM |
| **1013609713** | 2 | AUDp6a | 954 | 0.306735061 | 0 | 0 | 0 | 0 | 2 mo | WILD |
| **1010418842** | 1 | AUDp6a | 954 | 0.306735061 | 0.000331378 | 13 | 0.001080339 | 42.38185216 | 8 mo | HOM |
| **1013288213** | 2 | AUDp6a | 954 | 0.306735061 | 1.02E-05 | 1 | 3.34E-05 | 3.260142474 | 4 mo | HOM |

| | | | | | | | | | | |
|---|---|---|---|---|---|---|---|---|---|---|
| 1008976060 | 2 | AUDp6a | 954 | 0.306735061 | 4.18E-06 | 1 | 1.36E-05 | 3.260142474 | 8 mo | WILD |
| 1013446243 | 1 | AUDp6a | 954 | 0.306735061 | 5.15E-05 | 4 | 0.000167973 | 13.0405699 | 4 mo | HOM |
| 1013288951 | 2 | AUDp6a | 954 | 0.306735061 | 4.62E-05 | 1 | 0.000150631 | 3.260142474 | 4 mo | HOM |
| 1013978665 | 1 | LRNm | 955 | 0.528359121 | 0 | 0 | 0 | 0 | 2 mo | HOM |
| 1013445609 | 1 | LRNm | 955 | 0.528359121 | 0 | 0 | 0 | 0 | 4 mo | HOM |
| 1013609015 | 1 | LRNm | 955 | 0.528359121 | 0 | 0 | 0 | 0 | 4 mo | HOM |
| 1013608106 | 2 | LRNm | 955 | 0.528359121 | 0 | 0 | 0 | 0 | 4 mo | HOM |
| 1011937543 | 1 | LRNm | 955 | 0.528359121 | 0 | 0 | 0 | 0 | 8 mo | HOM |
| 1010419538 | 2 | LRNm | 955 | 0.528359121 | 0 | 0 | 0 | 0 | 4 mo | WILD |
| 1010419187 | 1 | LRNm | 955 | 0.528359121 | 9.03E-09 | 0 | 1.71E-08 | 0 | 8 mo | HOM |
| 1013978366 | 1 | LRNm | 955 | 0.528359121 | 0 | 0 | 0 | 0 | 2 mo | HOM |
| 1014416155 | 2 | LRNm | 955 | 0.528359121 | 0 | 0 | 0 | 0 | 2 mo | HOM |
| 1013978065 | 2 | LRNm | 955 | 0.528359121 | 0 | 0 | 0 | 0 | 2 mo | HOM |
| 1009934102 | 2 | LRNm | 955 | 0.528359121 | 0 | 0 | 0 | 0 | 4 mo | WILD |
| 1009322937 | 2 | LRNm | 955 | 0.528359121 | 0 | 0 | 0 | 0 | 8 mo | WILD |
| 1011080637 | 2 | LRNm | 955 | 0.528359121 | 0 | 0 | 0 | 0 | 8 mo | HOM |
| 1013609713 | 2 | LRNm | 955 | 0.528359121 | 0 | 0 | 0 | 0 | 2 mo | WILD |
| 1010418842 | 1 | LRNm | 955 | 0.528359121 | 2.26E-05 | 4 | 4.27E-05 | 7.570608404 | 8 mo | HOM |
| 1013288213 | 2 | LRNm | 955 | 0.528359121 | 0 | 0 | 0 | 0 | 4 mo | HOM |
| 1008976060 | 2 | LRNm | 955 | 0.528359121 | 6.95E-05 | 10 | 0.000131519 | 18.92652101 | 8 mo | WILD |
| 1013446243 | 1 | LRNm | 955 | 0.528359121 | 0 | 0 | 0 | 0 | 4 mo | HOM |
| 1013288951 | 2 | LRNm | 955 | 0.528359121 | 0 | 0 | 0 | 0 | 4 mo | HOM |
| 1013978665 | 1 | fa | 956 | 1.778477344 | 0 | 0 | 0 | 0 | 2 mo | HOM |
| 1013445609 | 1 | fa | 956 | 1.778477344 | 1.9E-05 | 2 | 1.07E-05 | 1.12455748 | 4 mo | HOM |
| 1013609015 | 1 | fa | 956 | 1.778477344 | 1.8E-05 | 1 | 1.01E-05 | 0.56227874 | 4 mo | HOM |
| 1013608106 | 2 | fa | 956 | 1.778477344 | 0 | 0 | 0 | 0 | 4 mo | HOM |
| 1011937543 | 1 | fa | 956 | 1.778477344 | 0.001392496 | 26 | 0.000782971 | 14.61924724 | 8 mo | HOM |
| 1010419538 | 2 | fa | 956 | 1.778477344 | 0 | 0 | 0 | 0 | 4 mo | WILD |
| 1010419187 | 1 | fa | 956 | 1.778477344 | 0.000377653 | 22 | 0.000212346 | 12.37013228 | 8 mo | HOM |
| 1013978366 | 1 | fa | 956 | 1.778477344 | 1.35E-06 | 0 | 7.57E-07 | 0 | 2 mo | HOM |
| 1014416155 | 2 | fa | 956 | 1.778477344 | 0 | 0 | 0 | 0 | 2 mo | HOM |
| 1013978065 | 2 | fa | 956 | 1.778477344 | 0 | 0 | 0 | 0 | 2 mo | HOM |
| 1009934102 | 2 | fa | 956 | 1.778477344 | 0 | 0 | 0 | 0 | 4 mo | WILD |
| 1009322937 | 2 | fa | 956 | 1.778477344 | 3.12E-06 | 0 | 1.75E-06 | 0 | 8 mo | WILD |
| 1011080637 | 2 | fa | 956 | 1.778477344 | 0.000391404 | 20 | 0.000220078 | 11.2455748 | 8 mo | HOM |
| 1013609713 | 2 | fa | 956 | 1.778477344 | 0 | 0 | 0 | 0 | 2 mo | WILD |
| 1010418842 | 1 | fa | 956 | 1.778477344 | 0.000635967 | 26 | 0.000357591 | 14.61924724 | 8 mo | HOM |
| 1013288213 | 2 | fa | 956 | 1.778477344 | 4.83E-05 | 1 | 2.71E-05 | 0.56227874 | 4 mo | HOM |
| 1008976060 | 2 | fa | 956 | 1.778477344 | 6.45E-06 | 2 | 3.63E-06 | 1.12455748 | 8 mo | WILD |
| 1013446243 | 1 | fa | 956 | 1.778477344 | 4.94E-05 | 4 | 2.78E-05 | 2.24911496 | 4 mo | HOM |
| 1013288951 | 2 | fa | 956 | 1.778477344 | 4.33E-05 | 0 | 2.44E-05 | 0 | 4 mo | HOM |
| 1013978665 | 1 | UVU | 957 | 2.257773078 | 0.000536483 | 17 | 0.000237616 | 7.529543231 | 2 mo | HOM |
| 1013445609 | 1 | UVU | 957 | 2.257773078 | 2.69E-05 | 2 | 1.19E-05 | 0.885828615 | 4 mo | HOM |
| 1013609015 | 1 | UVU | 957 | 2.257773078 | 0.000498509 | 20 | 0.000220797 | 8.858286154 | 4 mo | HOM |
| 1013608106 | 2 | UVU | 957 | 2.257773078 | 0 | 0 | 0 | 0 | 4 mo | HOM |
| 1011937543 | 1 | UVU | 957 | 2.257773078 | 0.001080529 | 10 | 0.000478582 | 4.429143077 | 8 mo | HOM |
| 1010419538 | 2 | UVU | 957 | 2.257773078 | 4.71E-05 | 4 | 2.09E-05 | 1.771657231 | 4 mo | WILD |
| 1010419187 | 1 | UVU | 957 | 2.257773078 | 0.000548467 | 25 | 0.000242924 | 11.07285769 | 8 mo | HOM |
| 1013978366 | 1 | UVU | 957 | 2.257773078 | 0.000317821 | 16 | 0.000140767 | 7.086628923 | 2 mo | HOM |
| 1014416155 | 2 | UVU | 957 | 2.257773078 | 0 | 0 | 0 | 0 | 2 mo | HOM |
| 1013978065 | 2 | UVU | 957 | 2.257773078 | 0.000463515 | 17 | 0.000205298 | 7.529543231 | 2 mo | HOM |
| 1009934102 | 2 | UVU | 957 | 2.257773078 | 9.33E-05 | 8 | 4.13E-05 | 3.543314462 | 4 mo | WILD |
| 1009322937 | 2 | UVU | 957 | 2.257773078 | 0.00019153 | 9 | 8.48E-05 | 3.986228769 | 8 mo | WILD |
| 1011080637 | 2 | UVU | 957 | 2.257773078 | 0.000890005 | 26 | 0.000394196 | 11.515772 | 8 mo | HOM |
| 1013609713 | 2 | UVU | 957 | 2.255840518 | 2.72E-05 | 0 | 1.21E-05 | 0 | 2 mo | WILD |
| 1010418842 | 1 | UVU | 957 | 2.257773078 | 0.001431748 | 21 | 0.000634142 | 9.301200462 | 8 mo | HOM |
| 1013288213 | 2 | UVU | 957 | 2.257773078 | 3.45E-05 | 3 | 1.53E-05 | 1.328742923 | 4 mo | HOM |
| 1008976060 | 2 | UVU | 957 | 2.257773078 | 0.002051649 | 35 | 0.000908705 | 15.50200077 | 8 mo | WILD |
| 1013446243 | 1 | UVU | 957 | 2.257773078 | 0.000143392 | 7 | 6.35E-05 | 3.100400154 | 4 mo | HOM |

| | | | | | | | | | | |
|---|---|---|---|---|---|---|---|---|---|---|
| 1013288951 | 2 | UVU | 957 | 2.257773078 | 0.001045313 | 15 | 0.000462984 | 6.643714616 | 4 mo | HOM |
| 1013978665 | 1 | EPI | 958 | 0.684837955 | 0 | 0 | 0 | 0 | 2 mo | HOM |
| 1013445609 | 1 | EPI | 958 | 0.684837955 | 0 | 0 | 0 | 0 | 4 mo | HOM |
| 1013609015 | 1 | EPI | 958 | 0.684837955 | 0.000127586 | 3 | 0.000186301 | 4.380598326 | 4 mo | HOM |
| 1013608106 | 2 | EPI | 958 | 0.684837955 | 0 | 0 | 0 | 0 | 4 mo | HOM |
| 1011937543 | 1 | EPI | 958 | 0.684837955 | 0.001917206 | 19 | 0.002799503 | 27.7437894 | 8 mo | HOM |
| 1010419538 | 2 | EPI | 958 | 0.684837955 | 2.54E-06 | 0 | 3.71E-06 | 0 | 4 mo | WILD |
| 1010419187 | 1 | EPI | 958 | 0.684837955 | 0.000111417 | 6 | 0.000162691 | 8.761196652 | 8 mo | HOM |
| 1013978366 | 1 | EPI | 958 | 0.684837955 | 0 | 0 | 0 | 0 | 2 mo | HOM |
| 1014416155 | 2 | EPI | 958 | 0.684837955 | 0 | 0 | 0 | 0 | 2 mo | HOM |
| 1013978065 | 2 | EPI | 958 | 0.684837955 | 0 | 0 | 0 | 0 | 2 mo | HOM |
| 1009934102 | 2 | EPI | 958 | 0.684837955 | 0 | 0 | 0 | 0 | 4 mo | WILD |
| 1009322937 | 2 | EPI | 958 | 0.684837955 | 1.05E-05 | 2 | 1.53E-05 | 2.920398884 | 8 mo | WILD |
| 1011080637 | 2 | EPI | 958 | 0.684837955 | 0.001957544 | 7 | 0.002858405 | 10.22139609 | 8 mo | HOM |
| 1013609713 | 2 | EPI | 958 | 0.684837955 | 0 | 0 | 0 | 0 | 2 mo | WILD |
| 1010418842 | 1 | EPI | 958 | 0.684837955 | 0.000104356 | 4 | 0.000152381 | 5.840797768 | 8 mo | HOM |
| 1013288213 | 2 | EPI | 958 | 0.684837955 | 0 | 0 | 0 | 0 | 4 mo | HOM |
| 1008976060 | 2 | EPI | 958 | 0.684837955 | 0.000389923 | 5 | 0.000569365 | 7.30099721 | 8 mo | WILD |
| 1013446243 | 1 | EPI | 958 | 0.684837955 | 3.04E-05 | 2 | 4.43E-05 | 2.920398884 | 4 mo | HOM |
| 1013288951 | 2 | EPI | 958 | 0.684837955 | 0.000630202 | 5 | 0.000920221 | 7.30099721 | 4 mo | HOM |
| 1013978665 | 1 | AUDv1 | 959 | 0.293933497 | 0 | 0 | 0 | 0 | 2 mo | HOM |
| 1013445609 | 1 | AUDv1 | 959 | 0.293933497 | 6.59E-05 | 3 | 0.000224204 | 10.20639032 | 4 mo | HOM |
| 1013609015 | 1 | AUDv1 | 959 | 0.293933497 | 5.41E-05 | 4 | 0.000184154 | 13.60852042 | 4 mo | HOM |
| 1013608106 | 2 | AUDv1 | 959 | 0.293933497 | 0 | 0 | 0 | 0 | 4 mo | HOM |
| 1011937543 | 1 | AUDv1 | 959 | 0.293933497 | 0.001063449 | 20 | 0.003617991 | 68.04260212 | 8 mo | HOM |
| 1010419538 | 2 | AUDv1 | 959 | 0.293933497 | 0 | 0 | 0 | 0 | 4 mo | WILD |
| 1010419187 | 1 | AUDv1 | 959 | 0.293933497 | 0.000898989 | 19 | 0.003058478 | 64.64047201 | 8 mo | HOM |
| 1013978366 | 1 | AUDv1 | 959 | 0.293933497 | 0 | 0 | 0 | 0 | 2 mo | HOM |
| 1014416155 | 2 | AUDv1 | 959 | 0.293933497 | 0 | 0 | 0 | 0 | 2 mo | HOM |
| 1013978065 | 2 | AUDv1 | 959 | 0.293933497 | 0 | 0 | 0 | 0 | 2 mo | HOM |
| 1009934102 | 2 | AUDv1 | 959 | 0.293933497 | 0 | 0 | 0 | 0 | 4 mo | WILD |
| 1009322937 | 2 | AUDv1 | 959 | 0.293933497 | 8.27E-06 | 0 | 2.81E-05 | 0 | 8 mo | WILD |
| 1011080637 | 2 | AUDv1 | 959 | 0.293933497 | 0.000222859 | 9 | 0.000758197 | 30.61917095 | 8 mo | HOM |
| 1013609713 | 2 | AUDv1 | 959 | 0.293933497 | 0 | 0 | 0 | 0 | 2 mo | WILD |
| 1010418842 | 1 | AUDv1 | 959 | 0.293933497 | 0.000922099 | 22 | 0.003137101 | 74.84686233 | 8 mo | HOM |
| 1013288213 | 2 | AUDv1 | 959 | 0.293933497 | 2.95E-05 | 3 | 0.000100447 | 10.20639032 | 4 mo | HOM |
| 1008976060 | 2 | AUDv1 | 959 | 0.293933497 | 2.18E-05 | 2 | 7.43E-05 | 6.804260212 | 8 mo | WILD |
| 1013446243 | 1 | AUDv1 | 959 | 0.293933497 | 3.71E-05 | 3 | 0.000126338 | 10.20639032 | 4 mo | HOM |
| 1013288951 | 2 | AUDv1 | 959 | 0.293933497 | 0.000621889 | 18 | 0.002115749 | 61.23834191 | 4 mo | HOM |
| 1013978665 | 1 | cbf | 960 | 10.10284606 | 0.000142499 | 9 | 1.41E-05 | 0.890838081 | 2 mo | HOM |
| 1013445609 | 1 | cbf | 960 | 10.10284606 | 9.02E-06 | 1 | 8.93E-07 | 0.098982009 | 4 mo | HOM |
| 1013609015 | 1 | cbf | 960 | 10.10284606 | 0.000176018 | 8 | 1.74E-05 | 0.791856072 | 4 mo | HOM |
| 1013608106 | 2 | cbf | 960 | 10.10284606 | 0 | 0 | 0 | 0 | 4 mo | HOM |
| 1011937543 | 1 | cbf | 960 | 10.10284606 | 0.00877641 | 43 | 0.000868707 | 4.256226389 | 8 mo | HOM |
| 1010419538 | 2 | cbf | 960 | 10.06124481 | 9.31E-05 | 5 | 9.25E-06 | 0.4969564 | 4 mo | WILD |
| 1010419187 | 1 | cbf | 960 | 10.10284606 | 9.49E-05 | 4 | 9.39E-06 | 0.395928036 | 8 mo | HOM |
| 1013978366 | 1 | cbf | 960 | 10.10284606 | 0.000471223 | 6 | 4.66E-05 | 0.593892054 | 2 mo | HOM |
| 1014416155 | 2 | cbf | 960 | 10.10284606 | 0 | 0 | 0 | 0 | 2 mo | HOM |
| 1013978065 | 2 | cbf | 960 | 10.10284606 | 0.000612973 | 8 | 6.07E-05 | 0.791856072 | 2 mo | HOM |
| 1009934102 | 2 | cbf | 960 | 10.10284606 | 2.77E-05 | 2 | 2.74E-06 | 0.197964018 | 4 mo | WILD |
| 1009322937 | 2 | cbf | 960 | 10.10284606 | 2.86E-05 | 1 | 2.83E-06 | 0.098982009 | 8 mo | WILD |
| 1011080637 | 2 | cbf | 960 | 10.10284606 | 0.001580248 | 9 | 0.000156416 | 0.890838081 | 8 mo | HOM |
| 1013609713 | 2 | cbf | 960 | 10.09744148 | 6.72E-06 | 0 | 6.65E-07 | 0 | 2 mo | WILD |
| 1010418842 | 1 | cbf | 960 | 10.10284606 | 0.001002478 | 18 | 9.92E-05 | 1.781676163 | 8 mo | HOM |
| 1013288213 | 2 | cbf | 960 | 10.10284606 | 6.5E-06 | 0 | 6.44E-07 | 0 | 4 mo | HOM |
| 1008976060 | 2 | cbf | 960 | 10.10284606 | 0.001017323 | 19 | 0.000100697 | 1.880658172 | 8 mo | WILD |
| 1013446243 | 1 | cbf | 960 | 10.10284606 | 2.01E-05 | 2 | 1.99E-06 | 0.197964018 | 4 mo | HOM |
| 1013288951 | 2 | cbf | 960 | 10.10284606 | 0.001385368 | 5 | 0.000137126 | 0.494910045 | 4 mo | HOM |
| 1013978665 | 1 | PIR | 961 | 11.90382574 | 2.48E-05 | 1 | 2.09E-06 | 0.084006606 | 2 mo | HOM |

| | | | | | | | | | | |
|---|---|---|---|---|---|---|---|---|---|---|
| **1013445609** | 1 | PIR | 961 | 11.91642117 | 0.000235465 | 21 | 1.98E-05 | 1.762274067 | 4 mo | HOM |
| **1013609015** | 1 | PIR | 961 | 11.91642117 | 0.00065152 | 39 | 5.47E-05 | 3.272794696 | 4 mo | HOM |
| **1013608106** | 2 | PIR | 961 | 11.91642117 | 6.86E-07 | 0 | 5.76E-08 | 0 | 4 mo | HOM |
| **1011937543** | 1 | PIR | 961 | 11.91642117 | 0.009711077 | 250 | 0.000814932 | 20.97945318 | 8 mo | HOM |
| **1010419538** | 2 | PIR | 961 | 11.91642117 | 0 | 0 | 0 | 0 | 4 mo | WILD |
| **1010419187** | 1 | PIR | 961 | 11.91642117 | 0.008552153 | 363 | 0.000717678 | 30.46216601 | 8 mo | HOM |
| **1013978366** | 1 | PIR | 961 | 11.91642117 | 0 | 0 | 0 | 0 | 2 mo | HOM |
| **1014416155** | 2 | PIR | 961 | 11.91642117 | 0 | 0 | 0 | 0 | 2 mo | HOM |
| **1013978065** | 2 | PIR | 961 | 11.91642117 | 2E-09 | 0 | 1.68E-10 | 0 | 2 mo | HOM |
| **1009934102** | 2 | PIR | 961 | 11.91642117 | 1.19E-05 | 1 | 1E-06 | 0.083917813 | 4 mo | WILD |
| **1009322937** | 2 | PIR | 961 | 11.91642117 | 2.71E-05 | 2 | 2.27E-06 | 0.167835625 | 8 mo | WILD |
| **1011080637** | 2 | PIR | 961 | 11.91642117 | 0.00589693 | 286 | 0.000494857 | 24.00049443 | 8 mo | HOM |
| **1013609713** | 2 | PIR | 961 | 11.91642117 | 0 | 0 | 0 | 0 | 2 mo | WILD |
| **1010418842** | 1 | PIR | 961 | 11.91642117 | 0.008748244 | 339 | 0.000734133 | 28.44813851 | 8 mo | HOM |
| **1013288213** | 2 | PIR | 961 | 11.91642117 | 0.000258564 | 21 | 2.17E-05 | 1.762274067 | 4 mo | HOM |
| **1008976060** | 2 | PIR | 961 | 11.91642117 | 6.15E-05 | 5 | 5.16E-06 | 0.419589064 | 8 mo | WILD |
| **1013446243** | 1 | PIR | 961 | 11.91642117 | 0.000452793 | 34 | 3.8E-05 | 2.853205632 | 4 mo | HOM |
| **1013288951** | 2 | PIR | 961 | 11.91642117 | 0.000699657 | 51 | 5.87E-05 | 4.279808448 | 4 mo | HOM |
| **1013978665** | 1 | MOs2/3 | 962 | 3.919215059 | 0 | 0 | 0 | 0 | 2 mo | HOM |
| **1013445609** | 1 | MOs2/3 | 962 | 3.919215059 | 0.000220006 | 12 | 5.61E-05 | 3.061837592 | 4 mo | HOM |
| **1013609015** | 1 | MOs2/3 | 962 | 3.919215059 | 0.000738799 | 22 | 0.000188507 | 5.613368919 | 4 mo | HOM |
| **1013608106** | 2 | MOs2/3 | 962 | 3.919215059 | 0 | 0 | 0 | 0 | 4 mo | HOM |
| **1011937543** | 1 | MOs2/3 | 962 | 3.919215059 | 0.004206834 | 70 | 0.001073387 | 17.86071929 | 8 mo | HOM |
| **1010419538** | 2 | MOs2/3 | 962 | 3.919215059 | 0 | 0 | 0 | 0 | 4 mo | WILD |
| **1010419187** | 1 | MOs2/3 | 962 | 3.919215059 | 0.004281871 | 125 | 0.001092533 | 31.89414158 | 8 mo | HOM |
| **1013978366** | 1 | MOs2/3 | 962 | 3.919215059 | 4.07E-07 | 0 | 1.04E-07 | 0 | 2 mo | HOM |
| **1014416155** | 2 | MOs2/3 | 962 | 3.919215059 | 0 | 0 | 0 | 0 | 2 mo | HOM |
| **1013978065** | 2 | MOs2/3 | 962 | 3.919215059 | 0 | 0 | 0 | 0 | 2 mo | HOM |
| **1009934102** | 2 | MOs2/3 | 962 | 3.919215059 | 2.49E-07 | 0 | 6.35E-08 | 0 | 4 mo | WILD |
| **1009322937** | 2 | MOs2/3 | 962 | 3.919215059 | 0 | 0 | 0 | 0 | 8 mo | WILD |
| **1011080637** | 2 | MOs2/3 | 962 | 3.919215059 | 0.001403082 | 72 | 0.000358001 | 18.37102555 | 8 mo | HOM |
| **1013609713** | 2 | MOs2/3 | 962 | 3.919215059 | 0 | 0 | 0 | 0 | 2 mo | WILD |
| **1010418842** | 1 | MOs2/3 | 962 | 3.919215059 | 0.002361685 | 126 | 0.000602591 | 32.14929472 | 8 mo | HOM |
| **1013288213** | 2 | MOs2/3 | 962 | 3.919215059 | 0.000165271 | 10 | 4.22E-05 | 2.551531327 | 4 mo | HOM |
| **1008976060** | 2 | MOs2/3 | 962 | 3.919215059 | 0 | 0 | 0 | 0 | 8 mo | WILD |
| **1013446243** | 1 | MOs2/3 | 962 | 3.919215059 | 0.000380279 | 22 | 9.7E-05 | 5.613368919 | 4 mo | HOM |
| **1013288951** | 2 | MOs2/3 | 962 | 3.919215059 | 0.000508262 | 18 | 0.000129685 | 4.592756388 | 4 mo | HOM |
| **1013978665** | 1 | LRNp | 963 | 0.058595077 | 0 | 0 | 0 | 0 | 2 mo | HOM |
| **1013445609** | 1 | LRNp | 963 | 0.058595077 | 0 | 0 | 0 | 0 | 4 mo | HOM |
| **1013609015** | 1 | LRNp | 963 | 0.058595077 | 0 | 0 | 0 | 0 | 4 mo | HOM |
| **1013608106** | 2 | LRNp | 963 | 0.058595077 | 0 | 0 | 0 | 0 | 4 mo | HOM |
| **1011937543** | 1 | LRNp | 963 | 0.058595077 | 0.000410433 | 0 | 0.007004569 | 0 | 8 mo | HOM |
| **1010419538** | 2 | LRNp | 963 | 0.058595077 | 0 | 0 | 0 | 0 | 4 mo | WILD |
| **1010419187** | 1 | LRNp | 963 | 0.058595077 | 0 | 0 | 0 | 0 | 8 mo | HOM |
| **1013978366** | 1 | LRNp | 963 | 0.058595077 | 0 | 0 | 0 | 0 | 2 mo | HOM |
| **1014416155** | 2 | LRNp | 963 | 0.058595077 | 0 | 0 | 0 | 0 | 2 mo | HOM |
| **1013978065** | 2 | LRNp | 963 | 0.058595077 | 0 | 0 | 0 | 0 | 2 mo | HOM |
| **1009934102** | 2 | LRNp | 963 | 0.058595077 | 0 | 0 | 0 | 0 | 4 mo | WILD |
| **1009322937** | 2 | LRNp | 963 | 0.058595077 | 0 | 0 | 0 | 0 | 8 mo | WILD |
| **1011080637** | 2 | LRNp | 963 | 0.058595077 | 0 | 0 | 0 | 0 | 8 mo | HOM |
| **1013609713** | 2 | LRNp | 963 | 0.058595077 | 0 | 0 | 0 | 0 | 2 mo | WILD |
| **1010418842** | 1 | LRNp | 963 | 0.058595077 | 0 | 0 | 0 | 0 | 8 mo | HOM |
| **1013288213** | 2 | LRNp | 963 | 0.058595077 | 0 | 0 | 0 | 0 | 4 mo | HOM |
| **1008976060** | 2 | LRNp | 963 | 0.058595077 | 0 | 0 | 0 | 0 | 8 mo | WILD |
| **1013446243** | 1 | LRNp | 963 | 0.058595077 | 0 | 0 | 0 | 0 | 4 mo | HOM |
| **1013288951** | 2 | LRNp | 963 | 0.058595077 | 0 | 0 | 0 | 0 | 4 mo | HOM |
| **1013978665** | 1 | ee | 964 | 0.096159139 | 0 | 0 | 0 | 0 | 2 mo | HOM |
| **1013445609** | 1 | ee | 964 | 0.096159139 | 1.22E-06 | 0 | 1.27E-05 | 0 | 4 mo | HOM |
| **1013609015** | 1 | ee | 964 | 0.096159139 | 2.66E-11 | 0 | 2.77E-10 | 0 | 4 mo | HOM |

| | | | | | | | | | | |
|---|---|---|---|---|---|---|---|---|---|---|
| **1013608106** | 2 | ee | 964 | 0.096159139 | 0 | 0 | 0 | 0 | 4 mo | HOM |
| **1011937543** | 1 | ee | 964 | 0.096159139 | 0.000195572 | 5 | 0.002033839 | 51.997138 | 8 mo | HOM |
| **1010419538** | 2 | ee | 964 | 0.096159139 | 0 | 0 | 0 | 0 | 4 mo | WILD |
| **1010419187** | 1 | ee | 964 | 0.096159139 | 0.00014064 | 5 | 0.001462574 | 51.997138 | 8 mo | HOM |
| **1013978366** | 1 | ee | 964 | 0.096159139 | 0 | 0 | 0 | 0 | 2 mo | HOM |
| **1014416155** | 2 | ee | 964 | 0.096159139 | 0 | 0 | 0 | 0 | 2 mo | HOM |
| **1013978065** | 2 | ee | 964 | 0.096159139 | 0 | 0 | 0 | 0 | 2 mo | HOM |
| **1009934102** | 2 | ee | 964 | 0.096159139 | 0 | 0 | 0 | 0 | 4 mo | WILD |
| **1009322937** | 2 | ee | 964 | 0.096159139 | 0 | 0 | 0 | 0 | 8 mo | WILD |
| **1011080637** | 2 | ee | 964 | 0.096159139 | 6.46E-05 | 6 | 0.000671977 | 62.3965656 | 8 mo | HOM |
| **1013609713** | 2 | ee | 964 | 0.096159139 | 0 | 0 | 0 | 0 | 2 mo | WILD |
| **1010418842** | 1 | ee | 964 | 0.096159139 | 0.000273348 | 5 | 0.002842662 | 51.997138 | 8 mo | HOM |
| **1013288213** | 2 | ee | 964 | 0.096159139 | 1.62E-06 | 0 | 1.68E-05 | 0 | 4 mo | HOM |
| **1008976060** | 2 | ee | 964 | 0.096159139 | 0 | 0 | 0 | 0 | 8 mo | WILD |
| **1013446243** | 1 | ee | 964 | 0.096159139 | 2.62E-05 | 1 | 0.000272921 | 10.3994276 | 4 mo | HOM |
| **1013288951** | 2 | ee | 964 | 0.096159139 | 1E-05 | 1 | 0.000104006 | 10.3994276 | 4 mo | HOM |
| **1013978665** | 1 | RSPagl2/3 | 965 | 0.690195106 | 1.68E-07 | 0 | 2.43E-07 | 0 | 2 mo | HOM |
| **1013445609** | 1 | RSPagl2/3 | 965 | 0.690195106 | 3.42E-05 | 1 | 4.95E-05 | 1.448865679 | 4 mo | HOM |
| **1013609015** | 1 | RSPagl2/3 | 965 | 0.690195106 | 0.000149693 | 6 | 0.000216885 | 8.693194071 | 4 mo | HOM |
| **1013608106** | 2 | RSPagl2/3 | 965 | 0.690195106 | 0 | 0 | 0 | 0 | 4 mo | HOM |
| **1011937543** | 1 | RSPagl2/3 | 965 | 0.690195106 | 0.000938256 | 13 | 0.001359406 | 18.83525382 | 8 mo | HOM |
| **1010419538** | 2 | RSPagl2/3 | 965 | 0.690195106 | 1.27E-05 | 1 | 1.84E-05 | 1.448865679 | 4 mo | WILD |
| **1010419187** | 1 | RSPagl2/3 | 965 | 0.690195106 | 0.001293112 | 31 | 0.001873545 | 44.91483603 | 8 mo | HOM |
| **1013978366** | 1 | RSPagl2/3 | 965 | 0.690195106 | 0 | 0 | 0 | 0 | 2 mo | HOM |
| **1014416155** | 2 | RSPagl2/3 | 965 | 0.690195106 | 0 | 0 | 0 | 0 | 2 mo | HOM |
| **1013978065** | 2 | RSPagl2/3 | 965 | 0.690195106 | 0 | 0 | 0 | 0 | 2 mo | HOM |
| **1009934102** | 2 | RSPagl2/3 | 965 | 0.690195106 | 2.9E-06 | 0 | 4.2E-06 | 0 | 4 mo | WILD |
| **1009322937** | 2 | RSPagl2/3 | 965 | 0.690195106 | 0 | 0 | 0 | 0 | 8 mo | WILD |
| **1011080637** | 2 | RSPagl2/3 | 965 | 0.690195106 | 0.000471699 | 19 | 0.000683428 | 27.52844789 | 8 mo | HOM |
| **1013609713** | 2 | RSPagl2/3 | 965 | 0.690195106 | 5.03E-07 | 0 | 7.29E-07 | 0 | 2 mo | WILD |
| **1010418842** | 1 | RSPagl2/3 | 965 | 0.690195106 | 0.00080152 | 24 | 0.001161294 | 34.77277628 | 8 mo | HOM |
| **1013288213** | 2 | RSPagl2/3 | 965 | 0.690195106 | 3.2E-05 | 2 | 4.64E-05 | 2.897731357 | 4 mo | HOM |
| **1008976060** | 2 | RSPagl2/3 | 965 | 0.690195106 | 2.66E-05 | 1 | 3.85E-05 | 1.448865679 | 8 mo | WILD |
| **1013446243** | 1 | RSPagl2/3 | 965 | 0.690195106 | 1.19E-05 | 1 | 1.72E-05 | 1.448865679 | 4 mo | HOM |
| **1013288951** | 2 | RSPagl2/3 | 965 | 0.690195106 | 0.000176447 | 6 | 0.000255648 | 8.693194071 | 4 mo | HOM |
| **1013978665** | 1 | EPv | 966 | 1.000918374 | 0 | 0 | 0 | 0 | 2 mo | HOM |
| **1013445609** | 1 | EPv | 966 | 1.00212252 | 3.61E-05 | 3 | 3.6E-05 | 2.993645927 | 4 mo | HOM |
| **1013609015** | 1 | EPv | 966 | 1.00212252 | 0.000141077 | 8 | 0.000140778 | 7.983055804 | 4 mo | HOM |
| **1013608106** | 2 | EPv | 966 | 1.00212252 | 0 | 0 | 0 | 0 | 4 mo | HOM |
| **1011937543** | 1 | EPv | 966 | 1.00212252 | 0.00146343 | 39 | 0.00146033 | 38.91739705 | 8 mo | HOM |
| **1010419538** | 2 | EPv | 966 | 1.00212252 | 0 | 0 | 0 | 0 | 4 mo | WILD |
| **1010419187** | 1 | EPv | 966 | 1.00212252 | 0.000795815 | 46 | 0.000794129 | 45.90257088 | 8 mo | HOM |
| **1013978366** | 1 | EPv | 966 | 1.00212252 | 0 | 0 | 0 | 0 | 2 mo | HOM |
| **1014416155** | 2 | EPv | 966 | 1.00212252 | 0 | 0 | 0 | 0 | 2 mo | HOM |
| **1013978065** | 2 | EPv | 966 | 1.00212252 | 0 | 0 | 0 | 0 | 2 mo | HOM |
| **1009934102** | 2 | EPv | 966 | 1.00212252 | 0 | 0 | 0 | 0 | 4 mo | WILD |
| **1009322937** | 2 | EPv | 966 | 1.00212252 | 0 | 0 | 0 | 0 | 8 mo | WILD |
| **1011080637** | 2 | EPv | 966 | 1.00212252 | 0.001044895 | 47 | 0.001042682 | 46.90045285 | 8 mo | HOM |
| **1013609713** | 2 | EPv | 966 | 1.00212252 | 0 | 0 | 0 | 0 | 2 mo | WILD |
| **1010418842** | 1 | EPv | 966 | 1.00212252 | 0.001469971 | 64 | 0.001466858 | 63.86444644 | 8 mo | HOM |
| **1013288213** | 2 | EPv | 966 | 1.00212252 | 2.35E-05 | 3 | 2.35E-05 | 2.993645927 | 4 mo | HOM |
| **1008976060** | 2 | EPv | 966 | 1.00212252 | 1.79E-05 | 4 | 1.79E-05 | 3.991527902 | 8 mo | WILD |
| **1013446243** | 1 | EPv | 966 | 1.00212252 | 0.000134956 | 12 | 0.00013467 | 11.97458371 | 4 mo | HOM |
| **1013288951** | 2 | EPv | 966 | 1.00212252 | 0.000136864 | 11 | 0.000136574 | 10.97670173 | 4 mo | HOM |
| **1013978665** | 1 | cm | 967 | 11.8283798 | 0.00177508 | 54 | 0.00015007 | 4.565291351 | 2 mo | HOM |
| **1013445609** | 1 | cm | 967 | 11.86380719 | 0.001051565 | 57 | 8.86E-05 | 4.804528519 | 4 mo | HOM |
| **1013609015** | 1 | cm | 967 | 10.98880424 | 0.002995398 | 108 | 0.000272586 | 9.828184912 | 4 mo | HOM |
| **1013608106** | 2 | cm | 967 | 11.9609593 | 0 | 0 | 0 | 0 | 4 mo | HOM |
| **1011937543** | 1 | cm | 967 | 11.99930466 | 0.016735169 | 121 | 0.001394678 | 10.08391765 | 8 mo | HOM |

| | | | | | | | | | | |
|---|---|---|---|---|---|---|---|---|---|---|
| **1010419538** | 2 | cm | 967 | 12.27919203 | 0.000235091 | 20 | 1.91E-05 | 1.628771661 | 4 mo | WILD |
| **1010419187** | 1 | cm | 967 | 10.95371067 | 0.010575188 | 78 | 0.000965443 | 7.120874591 | 8 mo | HOM |
| **1013978366** | 1 | cm | 967 | 12.60850966 | 0.001460541 | 75 | 0.000115838 | 5.948363605 | 2 mo | HOM |
| **1014416155** | 2 | cm | 967 | 12.35127268 | 1.89E-06 | 0 | 1.53E-07 | 0 | 2 mo | HOM |
| **1013978065** | 2 | cm | 967 | 12.66317516 | 0.001851507 | 81 | 0.000146212 | 6.396500006 | 2 mo | HOM |
| **1009934102** | 2 | cm | 967 | 11.54428517 | 0.000674639 | 51 | 5.84E-05 | 4.417770286 | 4 mo | WILD |
| **1009322937** | 2 | cm | 967 | 10.76889035 | 9.98E-05 | 13 | 9.27E-06 | 1.207181016 | 8 mo | WILD |
| **1011080637** | 2 | cm | 967 | 10.69779441 | 0.008267728 | 72 | 0.000772844 | 6.730359289 | 8 mo | HOM |
| **1013609713** | 2 | cm | 967 | 12.72772909 | 9.71E-05 | 15 | 7.63E-06 | 1.17852917 | 2 mo | WILD |
| **1010418842** | 1 | cm | 967 | 11.64449291 | 0.006427356 | 144 | 0.000551965 | 12.36636074 | 8 mo | HOM |
| **1013288213** | 2 | cm | 967 | 12.23674251 | 0.002316283 | 95 | 0.000189289 | 7.763504048 | 4 mo | HOM |
| **1008976060** | 2 | cm | 967 | 12.01170628 | 0.001713134 | 83 | 0.000142622 | 6.90992587 | 8 mo | WILD |
| **1013446243** | 1 | cm | 967 | 11.36491475 | 0.001293993 | 86 | 0.000113859 | 7.56714871 | 4 mo | HOM |
| **1013288951** | 2 | cm | 967 | 11.75403873 | 0.001761023 | 72 | 0.000149823 | 6.125554091 | 4 mo | HOM |
| **1013978665** | 1 | NOD | 968 | 1.565012803 | 0.000127529 | 3 | 8.15E-05 | 1.916917225 | 2 mo | HOM |
| **1013445609** | 1 | NOD | 968 | 1.565012803 | 0.000502511 | 6 | 0.000321091 | 3.83383445 | 4 mo | HOM |
| **1013609015** | 1 | NOD | 968 | 1.565012803 | 0.000366545 | 14 | 0.000234212 | 8.945613717 | 4 mo | HOM |
| **1013608106** | 2 | NOD | 968 | 1.565012803 | 0 | 0 | 0 | 0 | 4 mo | HOM |
| **1011937543** | 1 | NOD | 968 | 1.565012803 | 0.000910345 | 15 | 0.000581685 | 9.584586126 | 8 mo | HOM |
| **1010419538** | 2 | NOD | 968 | 1.561605382 | 0.000121093 | 3 | 7.75E-05 | 1.921099936 | 4 mo | WILD |
| **1010419187** | 1 | NOD | 968 | 1.565012803 | 0.001472212 | 25 | 0.000940703 | 15.97431021 | 8 mo | HOM |
| **1013978366** | 1 | NOD | 968 | 1.565012803 | 0.000282461 | 9 | 0.000180485 | 5.750751675 | 2 mo | HOM |
| **1014416155** | 2 | NOD | 968 | 1.565012803 | 0 | 0 | 0 | 0 | 2 mo | HOM |
| **1013978065** | 2 | NOD | 968 | 1.565012803 | 0.000268759 | 11 | 0.00017173 | 7.028696492 | 2 mo | HOM |
| **1009934102** | 2 | NOD | 968 | 1.565012803 | 1.05E-05 | 1 | 6.71E-06 | 0.638972408 | 4 mo | WILD |
| **1009322937** | 2 | NOD | 968 | 1.565012803 | 0.000377497 | 11 | 0.00024121 | 7.028696492 | 8 mo | WILD |
| **1011080637** | 2 | NOD | 968 | 1.565012803 | 0.003474497 | 25 | 0.002220108 | 15.97431021 | 8 mo | HOM |
| **1013609713** | 2 | NOD | 968 | 1.565012803 | 8.83E-05 | 11 | 5.64E-05 | 7.028696492 | 2 mo | WILD |
| **1010418842** | 1 | NOD | 968 | 1.565012803 | 0.002159477 | 30 | 0.001379846 | 19.16917225 | 8 mo | HOM |
| **1013288213** | 2 | NOD | 968 | 1.565012803 | 8.74E-06 | 0 | 5.58E-06 | 0 | 4 mo | HOM |
| **1008976060** | 2 | NOD | 968 | 1.565012803 | 0.004906614 | 31 | 0.003135191 | 19.80814466 | 8 mo | WILD |
| **1013446243** | 1 | NOD | 968 | 1.565012803 | 0.000254467 | 7 | 0.000162597 | 4.472806859 | 4 mo | HOM |
| **1013288951** | 2 | NOD | 968 | 1.565012803 | 0.000645901 | 8 | 0.000412713 | 5.111779267 | 4 mo | HOM |
| **1013978665** | 1 | ORBvl1 | 969 | 0.396754058 | 1.33E-05 | 0 | 3.36E-05 | 0 | 2 mo | HOM |
| **1013445609** | 1 | ORBvl1 | 969 | 0.396754058 | 1.52E-07 | 0 | 3.82E-07 | 0 | 4 mo | HOM |
| **1013609015** | 1 | ORBvl1 | 969 | 0.396754058 | 4.27E-05 | 1 | 0.000107725 | 2.520453109 | 4 mo | HOM |
| **1013608106** | 2 | ORBvl1 | 969 | 0.396754058 | 0 | 0 | 0 | 0 | 4 mo | HOM |
| **1011937543** | 1 | ORBvl1 | 969 | 0.396754058 | 0.000323955 | 6 | 0.000816513 | 15.12271865 | 8 mo | HOM |
| **1010419538** | 2 | ORBvl1 | 969 | 0.396754058 | 0 | 0 | 0 | 0 | 4 mo | WILD |
| **1010419187** | 1 | ORBvl1 | 969 | 0.396754058 | 0.000351196 | 13 | 0.000885173 | 32.76589042 | 8 mo | HOM |
| **1013978366** | 1 | ORBvl1 | 969 | 0.396754058 | 0 | 0 | 0 | 0 | 2 mo | HOM |
| **1014416155** | 2 | ORBvl1 | 969 | 0.396754058 | 0 | 0 | 0 | 0 | 2 mo | HOM |
| **1013978065** | 2 | ORBvl1 | 969 | 0.396754058 | 3.17E-06 | 0 | 7.99E-06 | 0 | 2 mo | HOM |
| **1009934102** | 2 | ORBvl1 | 969 | 0.396754058 | 0 | 0 | 0 | 0 | 4 mo | WILD |
| **1009322937** | 2 | ORBvl1 | 969 | 0.396754058 | 3.35E-05 | 4 | 8.44E-05 | 10.08181244 | 8 mo | WILD |
| **1011080637** | 2 | ORBvl1 | 969 | 0.396754058 | 0.00076707 | 7 | 0.001933365 | 17.64317176 | 8 mo | HOM |
| **1013609713** | 2 | ORBvl1 | 969 | 0.396754058 | 0 | 0 | 0 | 0 | 2 mo | WILD |
| **1010418842** | 1 | ORBvl1 | 969 | 0.396754058 | 0.000208812 | 8 | 0.000526302 | 20.16362487 | 8 mo | HOM |
| **1013288213** | 2 | ORBvl1 | 969 | 0.396754058 | 0.000827373 | 4 | 0.002085355 | 10.08181244 | 4 mo | HOM |
| **1008976060** | 2 | ORBvl1 | 969 | 0.396754058 | 4.51E-06 | 1 | 1.14E-05 | 2.520453109 | 8 mo | WILD |
| **1013446243** | 1 | ORBvl1 | 969 | 0.396754058 | 0 | 0 | 0 | 0 | 4 mo | HOM |
| **1013288951** | 2 | ORBvl1 | 969 | 0.396754058 | 0.000375851 | 6 | 0.000947314 | 15.12271865 | 4 mo | HOM |
| **1013978665** | 1 | PGRNd | 970 | 0.246541089 | 0 | 0 | 0 | 0 | 2 mo | HOM |
| **1013445609** | 1 | PGRNd | 970 | 0.246541089 | 0 | 0 | 0 | 0 | 4 mo | HOM |
| **1013609015** | 1 | PGRNd | 970 | 0.246541089 | 0 | 0 | 0 | 0 | 4 mo | HOM |
| **1013608106** | 2 | PGRNd | 970 | 0.246541089 | 0 | 0 | 0 | 0 | 4 mo | HOM |
| **1011937543** | 1 | PGRNd | 970 | 0.246541089 | 2.45E-05 | 1 | 9.95E-05 | 4.056119015 | 8 mo | HOM |
| **1010419538** | 2 | PGRNd | 970 | 0.246541089 | 0 | 0 | 0 | 0 | 4 mo | WILD |
| **1010419187** | 1 | PGRNd | 970 | 0.246541089 | 0 | 0 | 0 | 0 | 8 mo | HOM |

| | | | | | | | | | | |
|---|---|---|---|---|---|---|---|---|---|---|
| 1013978366 | 1 | PGRNd | 970 | 0.246541089 | 0 | 0 | 0 | 0 | 2 mo | HOM |
| 1014416155 | 2 | PGRNd | 970 | 0.246541089 | 0 | 0 | 0 | 0 | 2 mo | HOM |
| 1013978065 | 2 | PGRNd | 970 | 0.246541089 | 0 | 0 | 0 | 0 | 2 mo | HOM |
| 1009934102 | 2 | PGRNd | 970 | 0.246541089 | 0 | 0 | 0 | 0 | 4 mo | WILD |
| 1009322937 | 2 | PGRNd | 970 | 0.246541089 | 0 | 0 | 0 | 0 | 8 mo | WILD |
| 1011080637 | 2 | PGRNd | 970 | 0.246541089 | 0 | 0 | 0 | 0 | 8 mo | HOM |
| 1013609713 | 2 | PGRNd | 970 | 0.246541089 | 0 | 0 | 0 | 0 | 2 mo | WILD |
| 1010418842 | 1 | PGRNd | 970 | 0.246541089 | 1.16E-06 | 0 | 4.72E-06 | 0 | 8 mo | HOM |
| 1013288213 | 2 | PGRNd | 970 | 0.246541089 | 0 | 0 | 0 | 0 | 4 mo | HOM |
| 1008976060 | 2 | PGRNd | 970 | 0.246541089 | 0 | 0 | 0 | 0 | 8 mo | WILD |
| 1013446243 | 1 | PGRNd | 970 | 0.246541089 | 0 | 0 | 0 | 0 | 4 mo | HOM |
| 1013288951 | 2 | PGRNd | 970 | 0.246541089 | 0 | 0 | 0 | 0 | 4 mo | HOM |
| 1013978665 | 1 | fp | 971 | 1.406656877 | 0 | 0 | 0 | 0 | 2 mo | HOM |
| 1013445609 | 1 | fp | 971 | 1.406656877 | 1.46E-05 | 1 | 1.04E-05 | 0.710905422 | 4 mo | HOM |
| 1013609015 | 1 | fp | 971 | 1.406656877 | 2E-05 | 2 | 1.42E-05 | 1.421810843 | 4 mo | HOM |
| 1013608106 | 2 | fp | 971 | 1.406656877 | 0 | 0 | 0 | 0 | 4 mo | HOM |
| 1011937543 | 1 | fp | 971 | 1.406656877 | 0.001063547 | 17 | 0.000756082 | 12.08539217 | 8 mo | HOM |
| 1010419538 | 2 | fp | 971 | 1.406656877 | 0 | 0 | 0 | 0 | 4 mo | WILD |
| 1010419187 | 1 | fp | 971 | 1.406656877 | 0.0004795 | 23 | 0.000340879 | 16.3508247 | 8 mo | HOM |
| 1013978366 | 1 | fp | 971 | 1.406656877 | 0 | 0 | 0 | 0 | 2 mo | HOM |
| 1014416155 | 2 | fp | 971 | 1.406656877 | 0 | 0 | 0 | 0 | 2 mo | HOM |
| 1013978065 | 2 | fp | 971 | 1.406656877 | 0 | 0 | 0 | 0 | 2 mo | HOM |
| 1009934102 | 2 | fp | 971 | 1.406656877 | 0 | 0 | 0 | 0 | 4 mo | WILD |
| 1009322937 | 2 | fp | 971 | 1.406656877 | 0 | 0 | 0 | 0 | 8 mo | WILD |
| 1011080637 | 2 | fp | 971 | 1.406656877 | 0.000476665 | 8 | 0.000338864 | 5.687243373 | 8 mo | HOM |
| 1013609713 | 2 | fp | 971 | 1.406656877 | 0 | 0 | 0 | 0 | 2 mo | WILD |
| 1010418842 | 1 | fp | 971 | 1.406656877 | 0.001199314 | 28 | 0.000852599 | 19.9053518 | 8 mo | HOM |
| 1013288213 | 2 | fp | 971 | 1.406656877 | 0 | 0 | 0 | 0 | 4 mo | HOM |
| 1008976060 | 2 | fp | 971 | 1.406656877 | 3.83E-06 | 1 | 2.72E-06 | 0.710905422 | 8 mo | WILD |
| 1013446243 | 1 | fp | 971 | 1.406656877 | 7.12E-08 | 0 | 5.06E-08 | 0 | 4 mo | HOM |
| 1013288951 | 2 | fp | 971 | 1.406656877 | 0 | 0 | 0 | 0 | 4 mo | HOM |
| 1013978665 | 1 | PL | 972 | 2.483252253 | 5.29E-06 | 0 | 2.13E-06 | 0 | 2 mo | HOM |
| 1013445609 | 1 | PL | 972 | 2.483252253 | 8.2E-05 | 4 | 3.3E-05 | 1.610790847 | 4 mo | HOM |
| 1013609015 | 1 | PL | 972 | 2.483252253 | 0.000742273 | 23 | 0.000298912 | 9.262047372 | 4 mo | HOM |
| 1013608106 | 2 | PL | 972 | 2.483252253 | 0 | 0 | 0 | 0 | 4 mo | HOM |
| 1011937543 | 1 | PL | 972 | 2.483252253 | 0.001381687 | 34 | 0.000556402 | 13.6917222 | 8 mo | HOM |
| 1010419538 | 2 | PL | 972 | 2.483252253 | 0 | 0 | 0 | 0 | 4 mo | WILD |
| 1010419187 | 1 | PL | 972 | 2.483252253 | 0.004248665 | 113 | 0.001710928 | 45.50484144 | 8 mo | HOM |
| 1013978366 | 1 | PL | 972 | 2.483252253 | 7.84E-05 | 2 | 3.16E-05 | 0.805395424 | 2 mo | HOM |
| 1014416155 | 2 | PL | 972 | 2.483252253 | 0 | 0 | 0 | 0 | 2 mo | HOM |
| 1013978065 | 2 | PL | 972 | 2.483252253 | 0 | 0 | 0 | 0 | 2 mo | HOM |
| 1009934102 | 2 | PL | 972 | 2.483252253 | 4.9E-09 | 0 | 1.97E-09 | 0 | 4 mo | WILD |
| 1009322937 | 2 | PL | 972 | 2.483252253 | 1.99E-05 | 3 | 8.02E-06 | 1.208093135 | 8 mo | WILD |
| 1011080637 | 2 | PL | 972 | 2.483252253 | 0.001894463 | 79 | 0.000762896 | 31.81311923 | 8 mo | HOM |
| 1013609713 | 2 | PL | 972 | 2.483252253 | 1.09E-05 | 1 | 4.37E-06 | 0.402697712 | 2 mo | WILD |
| 1010418842 | 1 | PL | 972 | 2.483252253 | 0.003865724 | 101 | 0.001556718 | 40.67246889 | 8 mo | HOM |
| 1013288213 | 2 | PL | 972 | 2.483252253 | 0.001805053 | 24 | 0.000726891 | 9.664745084 | 4 mo | HOM |
| 1008976060 | 2 | PL | 972 | 2.483252253 | 1.9E-05 | 3 | 7.65E-06 | 1.208093135 | 8 mo | WILD |
| 1013446243 | 1 | PL | 972 | 2.483252253 | 0.000208911 | 11 | 8.41E-05 | 4.42967483 | 4 mo | HOM |
| 1013288951 | 2 | PL | 972 | 2.483252253 | 0.000825807 | 21 | 0.00033255 | 8.456651948 | 4 mo | HOM |
| 1013978665 | 1 | VISl2/3 | 973 | 0.310703492 | 0 | 0 | 0 | 0 | 2 mo | HOM |
| 1013445609 | 1 | VISl2/3 | 973 | 0.310703492 | 3E-05 | 3 | 9.64E-05 | 9.655507818 | 4 mo | HOM |
| 1013609015 | 1 | VISl2/3 | 973 | 0.310703492 | 6.67E-05 | 2 | 0.000214817 | 6.437005212 | 4 mo | HOM |
| 1013608106 | 2 | VISl2/3 | 973 | 0.310703492 | 0 | 0 | 0 | 0 | 4 mo | HOM |
| 1011937543 | 1 | VISl2/3 | 973 | 0.310703492 | 0.000335961 | 5 | 0.001081291 | 16.09251303 | 8 mo | HOM |
| 1010419538 | 2 | VISl2/3 | 973 | 0.310703492 | 0 | 0 | 0 | 0 | 4 mo | WILD |
| 1010419187 | 1 | VISl2/3 | 973 | 0.310703492 | 0.000612434 | 16 | 0.001971122 | 51.4960417 | 8 mo | HOM |
| 1013978366 | 1 | VISl2/3 | 973 | 0.310703492 | 1.5E-05 | 1 | 4.82E-05 | 3.218502606 | 2 mo | HOM |
| 1014416155 | 2 | VISl2/3 | 973 | 0.310703492 | 0 | 0 | 0 | 0 | 2 mo | HOM |

| | | | | | | | | | | |
|---|---|---|---|---|---|---|---|---|---|---|
| 1013978065 | 2 | VISl2/3 | 973 | 0.310703492 | 0 | 0 | 0 | 0 | 2 mo | HOM |
| 1009934102 | 2 | VISl2/3 | 973 | 0.310703492 | 0 | 0 | 0 | 0 | 4 mo | WILD |
| 1009322937 | 2 | VISl2/3 | 973 | 0.310703492 | 0 | 0 | 0 | 0 | 8 mo | WILD |
| 1011080637 | 2 | VISl2/3 | 973 | 0.310703492 | 0.000493286 | 14 | 0.001587642 | 45.05903648 | 8 mo | HOM |
| 1013609713 | 2 | VISl2/3 | 973 | 0.310703492 | 0 | 0 | 0 | 0 | 2 mo | WILD |
| 1010418842 | 1 | VISl2/3 | 973 | 0.310703492 | 0.00030405 | 11 | 0.000978587 | 35.40352867 | 8 mo | HOM |
| 1013288213 | 2 | VISl2/3 | 973 | 0.310703492 | 2.79E-05 | 2 | 8.99E-05 | 6.437005212 | 4 mo | HOM |
| 1008976060 | 2 | VISl2/3 | 973 | 0.310703492 | 0 | 0 | 0 | 0 | 8 mo | WILD |
| 1013446243 | 1 | VISl2/3 | 973 | 0.310703492 | 8.66E-05 | 7 | 0.000278709 | 22.52951824 | 4 mo | HOM |
| 1013288951 | 2 | VISl2/3 | 973 | 0.310703492 | 2.29E-05 | 1 | 7.36E-05 | 3.218502606 | 4 mo | HOM |
| 1013978665 | 1 | SSp-m5 | 974 | 1.253866253 | 0 | 0 | 0 | 0 | 2 mo | HOM |
| 1013445609 | 1 | SSp-m5 | 974 | 1.253866253 | 0.000109099 | 4 | 8.7E-05 | 3.190132912 | 4 mo | HOM |
| 1013609015 | 1 | SSp-m5 | 974 | 1.253866253 | 0.000364977 | 10 | 0.000291081 | 7.975332279 | 4 mo | HOM |
| 1013608106 | 2 | SSp-m5 | 974 | 1.253866253 | 0 | 0 | 0 | 0 | 4 mo | HOM |
| 1011937543 | 1 | SSp-m5 | 974 | 1.253866253 | 0.001935534 | 47 | 0.001543652 | 37.48406171 | 8 mo | HOM |
| 1010419538 | 2 | SSp-m5 | 974 | 1.253866253 | 0 | 0 | 0 | 0 | 4 mo | WILD |
| 1010419187 | 1 | SSp-m5 | 974 | 1.253866253 | 0.001285957 | 47 | 0.001025593 | 37.48406171 | 8 mo | HOM |
| 1013978366 | 1 | SSp-m5 | 974 | 1.253866253 | 1.65E-05 | 2 | 1.31E-05 | 1.595066456 | 2 mo | HOM |
| 1014416155 | 2 | SSp-m5 | 974 | 1.253866253 | 0 | 0 | 0 | 0 | 2 mo | HOM |
| 1013978065 | 2 | SSp-m5 | 974 | 1.253866253 | 0 | 0 | 0 | 0 | 2 mo | HOM |
| 1009934102 | 2 | SSp-m5 | 974 | 1.253866253 | 7.46E-07 | 0 | 5.95E-07 | 0 | 4 mo | WILD |
| 1009322937 | 2 | SSp-m5 | 974 | 1.253866253 | 2.7E-10 | 0 | 2.16E-10 | 0 | 8 mo | WILD |
| 1011080637 | 2 | SSp-m5 | 974 | 1.253866253 | 0.00117919 | 47 | 0.000940443 | 37.48406171 | 8 mo | HOM |
| 1013609713 | 2 | SSp-m5 | 974 | 1.253866253 | 0 | 0 | 0 | 0 | 2 mo | WILD |
| 1010418842 | 1 | SSp-m5 | 974 | 1.253866253 | 0.001438165 | 65 | 0.001146984 | 51.83965982 | 8 mo | HOM |
| 1013288213 | 2 | SSp-m5 | 974 | 1.253866253 | 0.000141441 | 11 | 0.000112804 | 8.772865507 | 4 mo | HOM |
| 1008976060 | 2 | SSp-m5 | 974 | 1.253866253 | 9.27E-06 | 1 | 7.39E-06 | 0.797533228 | 8 mo | WILD |
| 1013446243 | 1 | SSp-m5 | 974 | 1.253866253 | 0.000232097 | 10 | 0.000185105 | 7.975332279 | 4 mo | HOM |
| 1013288951 | 2 | SSp-m5 | 974 | 1.253866253 | 0.000431765 | 16 | 0.000344347 | 12.76053165 | 4 mo | HOM |
| 1013978665 | 1 | EW | 975 | 0.026227468 | 0 | 0 | 0 | 0 | 2 mo | HOM |
| 1013445609 | 1 | EW | 975 | 0.026227468 | 0 | 0 | 0 | 0 | 4 mo | HOM |
| 1013609015 | 1 | EW | 975 | 0.026227468 | 0 | 0 | 0 | 0 | 4 mo | HOM |
| 1013608106 | 2 | EW | 975 | 0.026227468 | 0 | 0 | 0 | 0 | 4 mo | HOM |
| 1011937543 | 1 | EW | 975 | 0.026227468 | 2.1E-05 | 1 | 0.000802364 | 38.12796562 | 8 mo | HOM |
| 1010419538 | 2 | EW | 975 | 0.026227468 | 0 | 0 | 0 | 0 | 4 mo | WILD |
| 1010419187 | 1 | EW | 975 | 0.026227468 | 0 | 0 | 0 | 0 | 8 mo | HOM |
| 1013978366 | 1 | EW | 975 | 0.026227468 | 0 | 0 | 0 | 0 | 2 mo | HOM |
| 1014416155 | 2 | EW | 975 | 0.026227468 | 0 | 0 | 0 | 0 | 2 mo | HOM |
| 1013978065 | 2 | EW | 975 | 0.026227468 | 0 | 0 | 0 | 0 | 2 mo | HOM |
| 1009934102 | 2 | EW | 975 | 0.026227468 | 0 | 0 | 0 | 0 | 4 mo | WILD |
| 1009322937 | 2 | EW | 975 | 0.026227468 | 0 | 0 | 0 | 0 | 8 mo | WILD |
| 1011080637 | 2 | EW | 975 | 0.026227468 | 0 | 0 | 0 | 0 | 8 mo | HOM |
| 1013609713 | 2 | EW | 975 | 0.026227468 | 0 | 0 | 0 | 0 | 2 mo | WILD |
| 1010418842 | 1 | EW | 975 | 0.026227468 | 1.2E-05 | 0 | 0.000458002 | 0 | 8 mo | HOM |
| 1013288213 | 2 | EW | 975 | 0.026227468 | 0 | 0 | 0 | 0 | 4 mo | HOM |
| 1008976060 | 2 | EW | 975 | 0.026227468 | 0 | 0 | 0 | 0 | 8 mo | WILD |
| 1013446243 | 1 | EW | 975 | 0.026227468 | 0 | 0 | 0 | 0 | 4 mo | HOM |
| 1013288951 | 2 | EW | 975 | 0.026227468 | 0 | 0 | 0 | 0 | 4 mo | HOM |
| 1013978665 | 1 | CENT2 | 976 | 1.376030387 | 0.000260626 | 9 | 0.000189404 | 6.540553235 | 2 mo | HOM |
| 1013445609 | 1 | CENT2 | 976 | 1.376030387 | 2.64E-05 | 1 | 1.92E-05 | 0.726728137 | 4 mo | HOM |
| 1013609015 | 1 | CENT2 | 976 | 1.376030387 | 0.002674409 | 13 | 0.001943568 | 9.447465783 | 4 mo | HOM |
| 1013608106 | 2 | CENT2 | 976 | 1.376030387 | 0 | 0 | 0 | 0 | 4 mo | HOM |
| 1011937543 | 1 | CENT2 | 976 | 1.376030387 | 0.000812344 | 14 | 0.000590353 | 10.17419392 | 8 mo | HOM |
| 1010419538 | 2 | CENT2 | 976 | 1.376030387 | 3.07E-05 | 3 | 2.23E-05 | 2.180184412 | 4 mo | WILD |
| 1010419187 | 1 | CENT2 | 976 | 1.376030387 | 0.000890254 | 16 | 0.000646972 | 11.62765019 | 8 mo | HOM |
| 1013978366 | 1 | CENT2 | 976 | 1.376030387 | 0.001065244 | 8 | 0.000774143 | 5.813825097 | 2 mo | HOM |
| 1014416155 | 2 | CENT2 | 976 | 1.376030387 | 0 | 0 | 0 | 0 | 2 mo | HOM |
| 1013978065 | 2 | CENT2 | 976 | 1.376030387 | 2.3E-05 | 1 | 1.67E-05 | 0.726728137 | 2 mo | HOM |
| 1009934102 | 2 | CENT2 | 976 | 1.376030387 | 0.000343948 | 10 | 0.000249957 | 7.267281372 | 4 mo | WILD |

| | | | | | | | | | | |
|---|---|---|---|---|---|---|---|---|---|---|
| **1009322937** | 2 | CENT2 | 976 | 1.376030387 | 0.000176787 | 4 | 0.000128476 | 2.906912549 | 8 mo | WILD |
| **1011080637** | 2 | CENT2 | 976 | 1.376030387 | 0.003822763 | 22 | 0.002778109 | 15.98801902 | 8 mo | HOM |
| **1013609713** | 2 | CENT2 | 976 | 1.376030387 | 0.000174935 | 7 | 0.00012713 | 5.08709696 | 2 mo | WILD |
| **1010418842** | 1 | CENT2 | 976 | 1.376030387 | 0.004582917 | 37 | 0.003330535 | 26.88894108 | 8 mo | HOM |
| **1013288213** | 2 | CENT2 | 976 | 1.376030387 | 8.99E-05 | 6 | 6.53E-05 | 4.360368823 | 4 mo | HOM |
| **1008976060** | 2 | CENT2 | 976 | 1.376030387 | 0.001239426 | 20 | 0.000900726 | 14.53456274 | 8 mo | WILD |
| **1013446243** | 1 | CENT2 | 976 | 1.376030387 | 0.00036688 | 7 | 0.000266622 | 5.08709696 | 4 mo | HOM |
| **1013288951** | 2 | CENT2 | 976 | 1.376030387 | 0.000350347 | 11 | 0.000254607 | 7.994009509 | 4 mo | HOM |
| **1013978665** | 1 | ECT6a | 977 | 0.400153796 | 0 | 0 | 0 | 0 | 2 mo | HOM |
| **1013445609** | 1 | ECT6a | 977 | 0.400153796 | 2.43E-05 | 3 | 6.07E-05 | 7.497117433 | 4 mo | HOM |
| **1013609015** | 1 | ECT6a | 977 | 0.400153796 | 4.89E-05 | 2 | 0.000122087 | 4.998078289 | 4 mo | HOM |
| **1013608106** | 2 | ECT6a | 977 | 0.400153796 | 0 | 0 | 0 | 0 | 4 mo | HOM |
| **1011937543** | 1 | ECT6a | 977 | 0.400153796 | 0.000852706 | 21 | 0.002130945 | 52.47982203 | 8 mo | HOM |
| **1010419538** | 2 | ECT6a | 977 | 0.400153796 | 0 | 0 | 0 | 0 | 4 mo | WILD |
| **1010419187** | 1 | ECT6a | 977 | 0.400153796 | 0.000469182 | 23 | 0.001172504 | 57.47790032 | 8 mo | HOM |
| **1013978366** | 1 | ECT6a | 977 | 0.400153796 | 0 | 0 | 0 | 0 | 2 mo | HOM |
| **1014416155** | 2 | ECT6a | 977 | 0.400153796 | 0 | 0 | 0 | 0 | 2 mo | HOM |
| **1013978065** | 2 | ECT6a | 977 | 0.400153796 | 0 | 0 | 0 | 0 | 2 mo | HOM |
| **1009934102** | 2 | ECT6a | 977 | 0.400153796 | 0 | 0 | 0 | 0 | 4 mo | WILD |
| **1009322937** | 2 | ECT6a | 977 | 0.400153796 | 0 | 0 | 0 | 0 | 8 mo | WILD |
| **1011080637** | 2 | ECT6a | 977 | 0.400153796 | 0.000545696 | 19 | 0.001363715 | 47.48174374 | 8 mo | HOM |
| **1013609713** | 2 | ECT6a | 977 | 0.400153796 | 0 | 0 | 0 | 0 | 2 mo | WILD |
| **1010418842** | 1 | ECT6a | 977 | 0.400153796 | 0.000768632 | 24 | 0.001920841 | 59.97693947 | 8 mo | HOM |
| **1013288213** | 2 | ECT6a | 977 | 0.400153796 | 1.85E-05 | 2 | 4.63E-05 | 4.998078289 | 4 mo | HOM |
| **1008976060** | 2 | ECT6a | 977 | 0.400153796 | 1.04E-05 | 2 | 2.59E-05 | 4.998078289 | 8 mo | WILD |
| **1013446243** | 1 | ECT6a | 977 | 0.400153796 | 0.00015624 | 7 | 0.000390449 | 17.49327401 | 4 mo | HOM |
| **1013288951** | 2 | ECT6a | 977 | 0.400153796 | 3.04E-05 | 3 | 7.59E-05 | 7.497117433 | 4 mo | HOM |
| **1013978665** | 1 | PGRNI | 978 | 0.729084014 | 0 | 0 | 0 | 0 | 2 mo | HOM |
| **1013445609** | 1 | PGRNI | 978 | 0.729084014 | 0 | 0 | 0 | 0 | 4 mo | HOM |
| **1013609015** | 1 | PGRNI | 978 | 0.729084014 | 0 | 0 | 0 | 0 | 4 mo | HOM |
| **1013608106** | 2 | PGRNI | 978 | 0.729084014 | 0 | 0 | 0 | 0 | 4 mo | HOM |
| **1011937543** | 1 | PGRNI | 978 | 0.729084014 | 0 | 0 | 0 | 0 | 8 mo | HOM |
| **1010419538** | 2 | PGRNI | 978 | 0.729084014 | 0 | 0 | 0 | 0 | 4 mo | WILD |
| **1010419187** | 1 | PGRNI | 978 | 0.729084014 | 0 | 0 | 0 | 0 | 8 mo | HOM |
| **1013978366** | 1 | PGRNI | 978 | 0.729084014 | 0 | 0 | 0 | 0 | 2 mo | HOM |
| **1014416155** | 2 | PGRNI | 978 | 0.729084014 | 0 | 0 | 0 | 0 | 2 mo | HOM |
| **1013978065** | 2 | PGRNI | 978 | 0.729084014 | 0 | 0 | 0 | 0 | 2 mo | HOM |
| **1009934102** | 2 | PGRNI | 978 | 0.729084014 | 0 | 0 | 0 | 0 | 4 mo | WILD |
| **1009322937** | 2 | PGRNI | 978 | 0.729084014 | 0 | 0 | 0 | 0 | 8 mo | WILD |
| **1011080637** | 2 | PGRNI | 978 | 0.729084014 | 0 | 0 | 0 | 0 | 8 mo | HOM |
| **1013609713** | 2 | PGRNI | 978 | 0.729084014 | 0 | 0 | 0 | 0 | 2 mo | WILD |
| **1010418842** | 1 | PGRNI | 978 | 0.729084014 | 0 | 0 | 0 | 0 | 8 mo | HOM |
| **1013288213** | 2 | PGRNI | 978 | 0.729084014 | 0 | 0 | 0 | 0 | 4 mo | HOM |
| **1008976060** | 2 | PGRNI | 978 | 0.729084014 | 2.92E-06 | 0 | 4.01E-06 | 0 | 8 mo | WILD |
| **1013446243** | 1 | PGRNI | 978 | 0.729084014 | 0 | 0 | 0 | 0 | 4 mo | HOM |
| **1013288951** | 2 | PGRNI | 978 | 0.729084014 | 0 | 0 | 0 | 0 | 4 mo | HOM |
| **1013978665** | 1 | PMd | 980 | 0.136206045 | 0 | 0 | 0 | 0 | 2 mo | HOM |
| **1013445609** | 1 | PMd | 980 | 0.136206045 | 0 | 0 | 0 | 0 | 4 mo | HOM |
| **1013609015** | 1 | PMd | 980 | 0.136206045 | 0 | 0 | 0 | 0 | 4 mo | HOM |
| **1013608106** | 2 | PMd | 980 | 0.136206045 | 0 | 0 | 0 | 0 | 4 mo | HOM |
| **1011937543** | 1 | PMd | 980 | 0.136206045 | 4.62E-05 | 1 | 0.000339058 | 7.341818063 | 8 mo | HOM |
| **1010419538** | 2 | PMd | 980 | 0.136206045 | 0 | 0 | 0 | 0 | 4 mo | WILD |
| **1010419187** | 1 | PMd | 980 | 0.136206045 | 0 | 0 | 0 | 0 | 8 mo | HOM |
| **1013978366** | 1 | PMd | 980 | 0.136206045 | 0 | 0 | 0 | 0 | 2 mo | HOM |
| **1014416155** | 2 | PMd | 980 | 0.136206045 | 0 | 0 | 0 | 0 | 2 mo | HOM |
| **1013978065** | 2 | PMd | 980 | 0.136206045 | 0 | 0 | 0 | 0 | 2 mo | HOM |
| **1009934102** | 2 | PMd | 980 | 0.136206045 | 0 | 0 | 0 | 0 | 4 mo | WILD |
| **1009322937** | 2 | PMd | 980 | 0.136206045 | 0 | 0 | 0 | 0 | 8 mo | WILD |
| **1011080637** | 2 | PMd | 980 | 0.136206045 | 4.73E-10 | 0 | 3.47E-09 | 0 | 8 mo | HOM |

| | | | | | | | | | | |
|---|---|---|---|---|---|---|---|---|---|---|
| **1013609713** | 2 | PMd | 980 | 0.136206045 | 0 | 0 | 0 | 0 | 2 mo | WILD |
| **1010418842** | 1 | PMd | 980 | 0.136206045 | 0 | 0 | 0 | 0 | 8 mo | HOM |
| **1013288213** | 2 | PMd | 980 | 0.136206045 | 0 | 0 | 0 | 0 | 4 mo | HOM |
| **1008976060** | 2 | PMd | 980 | 0.136206045 | 0 | 0 | 0 | 0 | 8 mo | WILD |
| **1013446243** | 1 | PMd | 980 | 0.136206045 | 0 | 0 | 0 | 0 | 4 mo | HOM |
| **1013288951** | 2 | PMd | 980 | 0.136206045 | 0 | 0 | 0 | 0 | 4 mo | HOM |
| **1013978665** | 1 | SSp-bfd1 | 981 | 0.864873912 | 0 | 0 | 0 | 0 | 2 mo | HOM |
| **1013445609** | 1 | SSp-bfd1 | 981 | 0.864873912 | 0.000207008 | 10 | 0.000239351 | 11.56237905 | 4 mo | HOM |
| **1013609015** | 1 | SSp-bfd1 | 981 | 0.864873912 | 0.000414782 | 19 | 0.000479587 | 21.96852019 | 4 mo | HOM |
| **1013608106** | 2 | SSp-bfd1 | 981 | 0.864873912 | 0 | 0 | 0 | 0 | 4 mo | HOM |
| **1011937543** | 1 | SSp-bfd1 | 981 | 0.864873912 | 0.005100579 | 33 | 0.005897483 | 38.15585086 | 8 mo | HOM |
| **1010419538** | 2 | SSp-bfd1 | 981 | 0.864873912 | 0 | 0 | 0 | 0 | 4 mo | WILD |
| **1010419187** | 1 | SSp-bfd1 | 981 | 0.864873912 | 0.003738576 | 73 | 0.004322683 | 84.40536706 | 8 mo | HOM |
| **1013978366** | 1 | SSp-bfd1 | 981 | 0.864873912 | 1.66E-05 | 3 | 1.92E-05 | 3.468713715 | 2 mo | HOM |
| **1014416155** | 2 | SSp-bfd1 | 981 | 0.864873912 | 0 | 0 | 0 | 0 | 2 mo | HOM |
| **1013978065** | 2 | SSp-bfd1 | 981 | 0.864873912 | 0 | 0 | 0 | 0 | 2 mo | HOM |
| **1009934102** | 2 | SSp-bfd1 | 981 | 0.864873912 | 0 | 0 | 0 | 0 | 4 mo | WILD |
| **1009322937** | 2 | SSp-bfd1 | 981 | 0.864873912 | 3.93E-06 | 1 | 4.55E-06 | 1.156237905 | 8 mo | WILD |
| **1011080637** | 2 | SSp-bfd1 | 981 | 0.864873912 | 0.001292871 | 42 | 0.001494867 | 48.56199201 | 8 mo | HOM |
| **1013609713** | 2 | SSp-bfd1 | 981 | 0.864873912 | 1.88E-05 | 1 | 2.17E-05 | 1.156237905 | 2 mo | WILD |
| **1010418842** | 1 | SSp-bfd1 | 981 | 0.864873912 | 0.005077845 | 41 | 0.005871197 | 47.4057541 | 8 mo | HOM |
| **1013288213** | 2 | SSp-bfd1 | 981 | 0.864873912 | 0.000121268 | 10 | 0.000140214 | 11.56237905 | 4 mo | HOM |
| **1008976060** | 2 | SSp-bfd1 | 981 | 0.864873912 | 0.000262602 | 13 | 0.000303631 | 15.03109276 | 8 mo | WILD |
| **1013446243** | 1 | SSp-bfd1 | 981 | 0.864873912 | 0.000522941 | 18 | 0.000604644 | 20.81228229 | 4 mo | HOM |
| **1013288951** | 2 | SSp-bfd1 | 981 | 0.864873912 | 0.0004572 | 19 | 0.000528632 | 21.96852019 | 4 mo | HOM |
| **1013978665** | 1 | FC | 982 | 0.063544278 | 0 | 0 | 0 | 0 | 2 mo | HOM |
| **1013445609** | 1 | FC | 982 | 0.063544278 | 0 | 0 | 0 | 0 | 4 mo | HOM |
| **1013609015** | 1 | FC | 982 | 0.063544278 | 0 | 0 | 0 | 0 | 4 mo | HOM |
| **1013608106** | 2 | FC | 982 | 0.063544278 | 0 | 0 | 0 | 0 | 4 mo | HOM |
| **1011937543** | 1 | FC | 982 | 0.063544278 | 0.000241227 | 2 | 0.003796204 | 31.47411636 | 8 mo | HOM |
| **1010419538** | 2 | FC | 982 | 0.063544278 | 0 | 0 | 0 | 0 | 4 mo | WILD |
| **1010419187** | 1 | FC | 982 | 0.063544278 | 4.43E-05 | 1 | 0.000696791 | 15.73705818 | 8 mo | HOM |
| **1013978366** | 1 | FC | 982 | 0.063544278 | 0 | 0 | 0 | 0 | 2 mo | HOM |
| **1014416155** | 2 | FC | 982 | 0.063544278 | 0 | 0 | 0 | 0 | 2 mo | HOM |
| **1013978065** | 2 | FC | 982 | 0.063544278 | 6.17E-06 | 2 | 9.71E-05 | 31.47411636 | 2 mo | HOM |
| **1009934102** | 2 | FC | 982 | 0.063544278 | 0 | 0 | 0 | 0 | 4 mo | WILD |
| **1009322937** | 2 | FC | 982 | 0.063544278 | 0 | 0 | 0 | 0 | 8 mo | WILD |
| **1011080637** | 2 | FC | 982 | 0.063544278 | 8.61E-05 | 1 | 0.001355027 | 15.73705818 | 8 mo | HOM |
| **1013609713** | 2 | FC | 982 | 0.063544278 | 0 | 0 | 0 | 0 | 2 mo | WILD |
| **1010418842** | 1 | FC | 982 | 0.063544278 | 0.000188527 | 2 | 0.00296686 | 31.47411636 | 8 mo | HOM |
| **1013288213** | 2 | FC | 982 | 0.063544278 | 0 | 0 | 0 | 0 | 4 mo | HOM |
| **1008976060** | 2 | FC | 982 | 0.063544278 | 4.54E-06 | 0 | 7.15E-05 | 0 | 8 mo | WILD |
| **1013446243** | 1 | FC | 982 | 0.063544278 | 0 | 0 | 0 | 0 | 4 mo | HOM |
| **1013288951** | 2 | FC | 982 | 0.063544278 | 1.22E-05 | 1 | 0.00019179 | 15.73705818 | 4 mo | HOM |
| **1013978665** | 1 | lfbs | 983 | 14.28357511 | 0 | 0 | 0 | 0 | 2 mo | HOM |
| **1013445609** | 1 | lfbs | 983 | 14.28357511 | 8.07E-05 | 6 | 5.65E-06 | 0.420062901 | 4 mo | HOM |
| **1013609015** | 1 | lfbs | 983 | 14.28357511 | 0.000148894 | 6 | 1.04E-05 | 0.420062901 | 4 mo | HOM |
| **1013608106** | 2 | lfbs | 983 | 14.28355923 | 0 | 0 | 0 | 0 | 4 mo | HOM |
| **1011937543** | 1 | lfbs | 983 | 14.28357511 | 0.011232644 | 168 | 0.000786403 | 11.76176124 | 8 mo | HOM |
| **1010419538** | 2 | lfbs | 983 | 14.28357511 | 1.13E-06 | 0 | 7.92E-08 | 0 | 4 mo | WILD |
| **1010419187** | 1 | lfbs | 983 | 14.28357511 | 0.002934231 | 127 | 0.000205427 | 8.89133141 | 8 mo | HOM |
| **1013978366** | 1 | lfbs | 983 | 14.28357511 | 3.05E-06 | 0 | 2.14E-07 | 0 | 2 mo | HOM |
| **1014416155** | 2 | lfbs | 983 | 14.28357511 | 0 | 0 | 0 | 0 | 2 mo | HOM |
| **1013978065** | 2 | lfbs | 983 | 14.28357511 | 7.35E-08 | 0 | 5.14E-09 | 0 | 2 mo | HOM |
| **1009934102** | 2 | lfbs | 983 | 14.28357511 | 9.18E-06 | 0 | 6.43E-07 | 0 | 4 mo | WILD |
| **1009322937** | 2 | lfbs | 983 | 14.28357511 | 7.49E-06 | 0 | 5.24E-07 | 0 | 8 mo | WILD |
| **1011080637** | 2 | lfbs | 983 | 14.28357511 | 0.002210581 | 96 | 0.000154764 | 6.72100642 | 8 mo | HOM |
| **1013609713** | 2 | lfbs | 983 | 14.28357511 | 0 | 0 | 0 | 0 | 2 mo | WILD |
| **1010418842** | 1 | lfbs | 983 | 14.28357511 | 0.005548875 | 180 | 0.000388479 | 12.60188704 | 8 mo | HOM |

| | | | | | | | | | | |
|---|---|---|---|---|---|---|---|---|---|---|
| 1013288213 | 2 | lfbs | 983 | 14.28357511 | 0.000318066 | 4 | 2.23E-05 | 0.280041934 | 4 mo | HOM |
| 1008976060 | 2 | lfbs | 983 | 14.28357511 | 0.000135393 | 16 | 9.48E-06 | 1.120167737 | 8 mo | WILD |
| 1013446243 | 1 | lfbs | 983 | 14.28357511 | 0.00017797 | 11 | 1.25E-05 | 0.770115319 | 4 mo | HOM |
| 1013288951 | 2 | lfbs | 983 | 14.28357511 | 0.000300791 | 6 | 2.11E-05 | 0.420062901 | 4 mo | HOM |
| 1013978665 | 1 | CENT3 | 984 | 2.826690048 | 0.000645377 | 23 | 0.000228315 | 8.136725148 | 2 mo | HOM |
| 1013445609 | 1 | CENT3 | 984 | 2.826690048 | 0.00016275 | 2 | 5.76E-05 | 0.707541317 | 4 mo | HOM |
| 1013609015 | 1 | CENT3 | 984 | 2.826690048 | 0.001209218 | 30 | 0.000427786 | 10.61311976 | 4 mo | HOM |
| 1013608106 | 2 | CENT3 | 984 | 2.826690048 | 0 | 0 | 0 | 0 | 4 mo | HOM |
| 1011937543 | 1 | CENT3 | 984 | 2.826690048 | 0.00349013 | 31 | 0.001234706 | 10.96689042 | 8 mo | HOM |
| 1010419538 | 2 | CENT3 | 984 | 2.809999002 | 0.000139858 | 5 | 4.98E-05 | 1.779360063 | 4 mo | WILD |
| 1010419187 | 1 | CENT3 | 984 | 2.826690048 | 0.001476923 | 44 | 0.000522492 | 15.56590898 | 8 mo | HOM |
| 1013978366 | 1 | CENT3 | 984 | 2.826690048 | 8.64E-05 | 8 | 3.06E-05 | 2.830165269 | 2 mo | HOM |
| 1014416155 | 2 | CENT3 | 984 | 2.826690048 | 0 | 0 | 0 | 0 | 2 mo | HOM |
| 1013978065 | 2 | CENT3 | 984 | 2.826690048 | 0.00024454 | 11 | 8.65E-05 | 3.891477245 | 2 mo | HOM |
| 1009934102 | 2 | CENT3 | 984 | 2.826690048 | 2.29E-05 | 3 | 8.1E-06 | 1.061311976 | 4 mo | WILD |
| 1009322937 | 2 | CENT3 | 984 | 2.826690048 | 0.00034924 | 20 | 0.000123551 | 7.075413172 | 8 mo | WILD |
| 1011080637 | 2 | CENT3 | 984 | 2.826690048 | 0.006659129 | 70 | 0.002355804 | 24.7639461 | 8 mo | HOM |
| 1013609713 | 2 | CENT3 | 984 | 2.826690048 | 4.14E-05 | 2 | 1.46E-05 | 0.707541317 | 2 mo | WILD |
| 1010418842 | 1 | CENT3 | 984 | 2.826690048 | 0.008029863 | 104 | 0.00284073 | 36.7921485 | 8 mo | HOM |
| 1013288213 | 2 | CENT3 | 984 | 2.826690048 | 0.000252055 | 9 | 8.92E-05 | 3.183935928 | 4 mo | HOM |
| 1008976060 | 2 | CENT3 | 984 | 2.826690048 | 0.001480008 | 42 | 0.000523584 | 14.85836766 | 8 mo | WILD |
| 1013446243 | 1 | CENT3 | 984 | 2.826690048 | 0.000149969 | 8 | 5.31E-05 | 2.830165269 | 4 mo | HOM |
| 1013288951 | 2 | CENT3 | 984 | 2.826690048 | 7.51E-05 | 6 | 2.66E-05 | 2.122623952 | 4 mo | HOM |
| 1013978665 | 1 | MOp | 985 | 11.69442806 | 3.79E-05 | 6 | 3.24E-06 | 0.513064852 | 2 mo | HOM |
| 1013445609 | 1 | MOp | 985 | 11.69442806 | 0.000959184 | 65 | 8.2E-05 | 5.55820256 | 4 mo | HOM |
| 1013609015 | 1 | MOp | 985 | 11.69442806 | 0.00232825 | 98 | 0.000199091 | 8.380059245 | 4 mo | HOM |
| 1013608106 | 2 | MOp | 985 | 11.69442806 | 0 | 0 | 0 | 0 | 4 mo | HOM |
| 1011937543 | 1 | MOp | 985 | 11.69442806 | 0.015667328 | 313 | 0.001339726 | 26.7648831 | 8 mo | HOM |
| 1010419538 | 2 | MOp | 985 | 11.69442806 | 9.44E-06 | 1 | 8.07E-07 | 0.085510809 | 4 mo | WILD |
| 1010419187 | 1 | MOp | 985 | 11.69442806 | 0.01597985 | 627 | 0.00136645 | 53.615277 | 8 mo | HOM |
| 1013978366 | 1 | MOp | 985 | 11.69442806 | 1.6E-05 | 3 | 1.37E-06 | 0.256532426 | 2 mo | HOM |
| 1014416155 | 2 | MOp | 985 | 11.69442806 | 0 | 0 | 0 | 0 | 2 mo | HOM |
| 1013978065 | 2 | MOp | 985 | 11.69442806 | 0 | 0 | 0 | 0 | 2 mo | HOM |
| 1009934102 | 2 | MOp | 985 | 11.69442806 | 2.69E-06 | 1 | 2.3E-07 | 0.085510809 | 4 mo | WILD |
| 1009322937 | 2 | MOp | 985 | 11.69442806 | 5.71E-06 | 1 | 4.88E-07 | 0.085510809 | 8 mo | WILD |
| 1011080637 | 2 | MOp | 985 | 11.69442806 | 0.007085767 | 316 | 0.00060591 | 27.02141552 | 8 mo | HOM |
| 1013609713 | 2 | MOp | 985 | 11.69442806 | 1.87E-05 | 1 | 1.6E-06 | 0.085510809 | 2 mo | WILD |
| 1010418842 | 1 | MOp | 985 | 11.69442806 | 0.011561433 | 505 | 0.000988627 | 43.18295835 | 8 mo | HOM |
| 1013288213 | 2 | MOp | 985 | 11.69442806 | 0.000739804 | 46 | 6.33E-05 | 3.933497197 | 4 mo | HOM |
| 1008976060 | 2 | MOp | 985 | 11.69442806 | 0.000288922 | 18 | 2.47E-05 | 1.539194555 | 8 mo | WILD |
| 1013446243 | 1 | MOp | 985 | 11.69442806 | 0.001991505 | 100 | 0.000170295 | 8.551080862 | 4 mo | HOM |
| 1013288951 | 2 | MOp | 985 | 11.69442806 | 0.002973375 | 124 | 0.000254256 | 10.60334027 | 4 mo | HOM |
| 1013978665 | 1 | ccs | 986 | 0.767474378 | 0 | 0 | 0 | 0 | 2 mo | HOM |
| 1013445609 | 1 | ccs | 986 | 0.767474378 | 2.29E-06 | 1 | 2.98E-06 | 1.302975095 | 4 mo | HOM |
| 1013609015 | 1 | ccs | 986 | 0.767474378 | 1.34E-05 | 1 | 1.74E-05 | 1.302975095 | 4 mo | HOM |
| 1013608106 | 2 | ccs | 986 | 0.767474378 | 0 | 0 | 0 | 0 | 4 mo | HOM |
| 1011937543 | 1 | ccs | 986 | 0.767474378 | 0.001121564 | 10 | 0.00146137 | 13.02975095 | 8 mo | HOM |
| 1010419538 | 2 | ccs | 986 | 0.767474378 | 0 | 0 | 0 | 0 | 4 mo | WILD |
| 1010419187 | 1 | ccs | 986 | 0.767474378 | 0.00034215 | 12 | 0.000445813 | 15.63570114 | 8 mo | HOM |
| 1013978366 | 1 | ccs | 986 | 0.767474378 | 1.7E-06 | 0 | 2.22E-06 | 0 | 2 mo | HOM |
| 1014416155 | 2 | ccs | 986 | 0.767474378 | 0 | 0 | 0 | 0 | 2 mo | HOM |
| 1013978065 | 2 | ccs | 986 | 0.767474378 | 0 | 0 | 0 | 0 | 2 mo | HOM |
| 1009934102 | 2 | ccs | 986 | 0.767474378 | 0 | 0 | 0 | 0 | 4 mo | WILD |
| 1009322937 | 2 | ccs | 986 | 0.767474378 | 0 | 0 | 0 | 0 | 8 mo | WILD |
| 1011080637 | 2 | ccs | 986 | 0.767474378 | 0.000133713 | 7 | 0.000174225 | 9.120825664 | 8 mo | HOM |
| 1013609713 | 2 | ccs | 986 | 0.767474378 | 0 | 0 | 0 | 0 | 2 mo | WILD |
| 1010418842 | 1 | ccs | 986 | 0.767474378 | 0.001042926 | 22 | 0.001358907 | 28.66545209 | 8 mo | HOM |
| 1013288213 | 2 | ccs | 986 | 0.767474378 | 0 | 0 | 0 | 0 | 4 mo | HOM |
| 1008976060 | 2 | ccs | 986 | 0.767474378 | 1.02E-06 | 0 | 1.33E-06 | 0 | 8 mo | WILD |

| | | | | | | | | | | |
|---|---|---|---|---|---|---|---|---|---|---|
| **1013446243** | 1 | ccs | 986 | 0.767474378 | 0 | 0 | 0 | 0 | 4 mo | HOM |
| **1013288951** | 2 | ccs | 986 | 0.767474378 | 1.95E-05 | 2 | 2.54E-05 | 2.60595019 | 4 mo | HOM |
| **1013978665** | 1 | P-mot | 987 | 5.969910735 | 3.79E-05 | 3 | 6.35E-06 | 0.502520077 | 2 mo | HOM |
| **1013445609** | 1 | P-mot | 987 | 5.969910735 | 0 | 0 | 0 | 0 | 4 mo | HOM |
| **1013609015** | 1 | P-mot | 987 | 5.969910735 | 0.001723956 | 10 | 0.000288774 | 1.675066922 | 4 mo | HOM |
| **1013608106** | 2 | P-mot | 987 | 5.969910735 | 0 | 0 | 0 | 0 | 4 mo | HOM |
| **1011937543** | 1 | P-mot | 987 | 5.969910735 | 0.003635352 | 17 | 0.000608946 | 2.847613768 | 8 mo | HOM |
| **1010419538** | 2 | P-mot | 987 | 5.969910735 | 0 | 0 | 0 | 0 | 4 mo | WILD |
| **1010419187** | 1 | P-mot | 987 | 5.969910735 | 0.001178642 | 20 | 0.00019743 | 3.350133844 | 8 mo | HOM |
| **1013978366** | 1 | P-mot | 987 | 5.969910735 | 2.48E-05 | 2 | 4.15E-06 | 0.335013384 | 2 mo | HOM |
| **1014416155** | 2 | P-mot | 987 | 5.969910735 | 0 | 0 | 0 | 0 | 2 mo | HOM |
| **1013978065** | 2 | P-mot | 987 | 5.969910735 | 0 | 0 | 0 | 0 | 2 mo | HOM |
| **1009934102** | 2 | P-mot | 987 | 5.969910735 | 9.29E-06 | 0 | 1.56E-06 | 0 | 4 mo | WILD |
| **1009322937** | 2 | P-mot | 987 | 5.969910735 | 5.77E-05 | 6 | 9.67E-06 | 1.005040153 | 8 mo | WILD |
| **1011080637** | 2 | P-mot | 987 | 5.969910735 | 0.002278003 | 16 | 0.000381581 | 2.680107076 | 8 mo | HOM |
| **1013609713** | 2 | P-mot | 987 | 5.969910735 | 1.32E-05 | 0 | 2.22E-06 | 0 | 2 mo | WILD |
| **1010418842** | 1 | P-mot | 987 | 5.969910735 | 0.002072352 | 50 | 0.000347133 | 8.375334611 | 8 mo | HOM |
| **1013288213** | 2 | P-mot | 987 | 5.969910735 | 4.77E-06 | 1 | 7.99E-07 | 0.167506692 | 4 mo | HOM |
| **1008976060** | 2 | P-mot | 987 | 5.969910735 | 0.002361183 | 57 | 0.000395514 | 9.547881457 | 8 mo | WILD |
| **1013446243** | 1 | P-mot | 987 | 5.969910735 | 9.74E-05 | 8 | 1.63E-05 | 1.340053538 | 4 mo | HOM |
| **1013288951** | 2 | P-mot | 987 | 5.969910735 | 0.000222004 | 13 | 3.72E-05 | 2.177586999 | 4 mo | HOM |
| **1013978665** | 1 | ECT5 | 988 | 0.545815273 | 1.85E-06 | 0 | 3.39E-06 | 0 | 2 mo | HOM |
| **1013445609** | 1 | ECT5 | 988 | 0.545815273 | 4.52E-05 | 4 | 8.28E-05 | 7.32848676 | 4 mo | HOM |
| **1013609015** | 1 | ECT5 | 988 | 0.545815273 | 0.000186203 | 6 | 0.000341147 | 10.99273014 | 4 mo | HOM |
| **1013608106** | 2 | ECT5 | 988 | 0.545815273 | 0 | 0 | 0 | 0 | 4 mo | HOM |
| **1011937543** | 1 | ECT5 | 988 | 0.545815273 | 0.000595321 | 21 | 0.001090701 | 38.47455549 | 8 mo | HOM |
| **1010419538** | 2 | ECT5 | 988 | 0.545815273 | 0 | 0 | 0 | 0 | 4 mo | WILD |
| **1010419187** | 1 | ECT5 | 988 | 0.545815273 | 0.001059834 | 38 | 0.001941745 | 69.62062422 | 8 mo | HOM |
| **1013978366** | 1 | ECT5 | 988 | 0.545815273 | 0 | 0 | 0 | 0 | 2 mo | HOM |
| **1014416155** | 2 | ECT5 | 988 | 0.545815273 | 0 | 0 | 0 | 0 | 2 mo | HOM |
| **1013978065** | 2 | ECT5 | 988 | 0.545815273 | 0 | 0 | 0 | 0 | 2 mo | HOM |
| **1009934102** | 2 | ECT5 | 988 | 0.545815273 | 0 | 0 | 0 | 0 | 4 mo | WILD |
| **1009322937** | 2 | ECT5 | 988 | 0.545815273 | 0 | 0 | 0 | 0 | 8 mo | WILD |
| **1011080637** | 2 | ECT5 | 988 | 0.545815273 | 0.000345233 | 18 | 0.000632508 | 32.97819042 | 8 mo | HOM |
| **1013609713** | 2 | ECT5 | 988 | 0.545815273 | 0 | 0 | 0 | 0 | 2 mo | WILD |
| **1010418842** | 1 | ECT5 | 988 | 0.545815273 | 0.000583176 | 30 | 0.001068449 | 54.9636507 | 8 mo | HOM |
| **1013288213** | 2 | ECT5 | 988 | 0.545815273 | 8.6E-05 | 4 | 0.000157578 | 7.32848676 | 4 mo | HOM |
| **1008976060** | 2 | ECT5 | 988 | 0.545815273 | 1.56E-06 | 0 | 2.85E-06 | 0 | 8 mo | WILD |
| **1013446243** | 1 | ECT5 | 988 | 0.545815273 | 6.18E-05 | 5 | 0.000113289 | 9.16060845 | 4 mo | HOM |
| **1013288951** | 2 | ECT5 | 988 | 0.545815273 | 8.51E-05 | 6 | 0.000155897 | 10.99273014 | 4 mo | HOM |
| **1013978665** | 1 | FN | 989 | 0.517210053 | 0 | 0 | 0 | 0 | 2 mo | HOM |
| **1013445609** | 1 | FN | 989 | 0.517210053 | 0 | 0 | 0 | 0 | 4 mo | HOM |
| **1013609015** | 1 | FN | 989 | 0.517210053 | 0 | 0 | 0 | 0 | 4 mo | HOM |
| **1013608106** | 2 | FN | 989 | 0.517210053 | 0 | 0 | 0 | 0 | 4 mo | HOM |
| **1011937543** | 1 | FN | 989 | 0.517210053 | 0 | 0 | 0 | 0 | 8 mo | HOM |
| **1010419538** | 2 | FN | 989 | 0.476606614 | 1.16E-05 | 1 | 2.43E-05 | 2.098166433 | 4 mo | WILD |
| **1010419187** | 1 | FN | 989 | 0.517210053 | 4.11E-12 | 0 | 7.94E-12 | 0 | 8 mo | HOM |
| **1013978366** | 1 | FN | 989 | 0.517210053 | 0 | 0 | 0 | 0 | 2 mo | HOM |
| **1014416155** | 2 | FN | 989 | 0.517210053 | 0 | 0 | 0 | 0 | 2 mo | HOM |
| **1013978065** | 2 | FN | 989 | 0.517210053 | 0 | 0 | 0 | 0 | 2 mo | HOM |
| **1009934102** | 2 | FN | 989 | 0.517210053 | 0 | 0 | 0 | 0 | 4 mo | WILD |
| **1009322937** | 2 | FN | 989 | 0.517210053 | 0 | 0 | 0 | 0 | 8 mo | WILD |
| **1011080637** | 2 | FN | 989 | 0.517210053 | 0 | 0 | 0 | 0 | 8 mo | HOM |
| **1013609713** | 2 | FN | 989 | 0.517210053 | 0 | 0 | 0 | 0 | 2 mo | WILD |
| **1010418842** | 1 | FN | 989 | 0.517210053 | 0 | 0 | 0 | 0 | 8 mo | HOM |
| **1013288213** | 2 | FN | 989 | 0.517210053 | 0 | 0 | 0 | 0 | 4 mo | HOM |
| **1008976060** | 2 | FN | 989 | 0.517210053 | 5.84E-05 | 9 | 0.000112984 | 17.40105389 | 8 mo | WILD |
| **1013446243** | 1 | FN | 989 | 0.517210053 | 0 | 0 | 0 | 0 | 4 mo | HOM |
| **1013288951** | 2 | FN | 989 | 0.517210053 | 0 | 0 | 0 | 0 | 4 mo | HOM |

| | | | | | | | | | | |
|---|---|---|---|---|---|---|---|---|---|---|
| **1013978665** | 1 | AUDv4 | 990 | 0.220072544 | 0 | 0 | 0 | 0 | 2 mo | HOM |
| **1013445609** | 1 | AUDv4 | 990 | 0.220072544 | 1.27E-05 | 2 | 5.78E-05 | 9.08791241 | 4 mo | HOM |
| **1013609015** | 1 | AUDv4 | 990 | 0.220072544 | 0.00017121 | 5 | 0.000777972 | 22.71978102 | 4 mo | HOM |
| **1013608106** | 2 | AUDv4 | 990 | 0.220072544 | 0 | 0 | 0 | 0 | 4 mo | HOM |
| **1011937543** | 1 | AUDv4 | 990 | 0.220072544 | 0.000333324 | 6 | 0.001514609 | 27.26373723 | 8 mo | HOM |
| **1010419538** | 2 | AUDv4 | 990 | 0.220072544 | 0 | 0 | 0 | 0 | 4 mo | WILD |
| **1010419187** | 1 | AUDv4 | 990 | 0.220072544 | 0.000185967 | 10 | 0.000845027 | 45.43956205 | 8 mo | HOM |
| **1013978366** | 1 | AUDv4 | 990 | 0.220072544 | 0 | 0 | 0 | 0 | 2 mo | HOM |
| **1014416155** | 2 | AUDv4 | 990 | 0.220072544 | 0 | 0 | 0 | 0 | 2 mo | HOM |
| **1013978065** | 2 | AUDv4 | 990 | 0.220072544 | 0 | 0 | 0 | 0 | 2 mo | HOM |
| **1009934102** | 2 | AUDv4 | 990 | 0.220072544 | 0 | 0 | 0 | 0 | 4 mo | WILD |
| **1009322937** | 2 | AUDv4 | 990 | 0.220072544 | 0 | 0 | 0 | 0 | 8 mo | WILD |
| **1011080637** | 2 | AUDv4 | 990 | 0.220072544 | 0.000123951 | 7 | 0.000563226 | 31.80769343 | 8 mo | HOM |
| **1013609713** | 2 | AUDv4 | 990 | 0.220072544 | 0 | 0 | 0 | 0 | 2 mo | WILD |
| **1010418842** | 1 | AUDv4 | 990 | 0.220072544 | 0.000265116 | 11 | 0.001204675 | 49.98351825 | 8 mo | HOM |
| **1013288213** | 2 | AUDv4 | 990 | 0.220072544 | 1.93E-05 | 1 | 8.76E-05 | 4.543956205 | 4 mo | HOM |
| **1008976060** | 2 | AUDv4 | 990 | 0.220072544 | 0 | 0 | 0 | 0 | 8 mo | WILD |
| **1013446243** | 1 | AUDv4 | 990 | 0.220072544 | 4.55E-05 | 3 | 0.000206679 | 13.63186861 | 4 mo | HOM |
| **1013288951** | 2 | AUDv4 | 990 | 0.220072544 | 0.000101993 | 3 | 0.00046345 | 13.63186861 | 4 mo | HOM |
| **1013978665** | 1 | mfbs | 991 | 7.256509205 | 4.33E-05 | 0 | 5.97E-06 | 0 | 2 mo | HOM |
| **1013445609** | 1 | mfbs | 991 | 7.256509205 | 4.13E-05 | 4 | 5.7E-06 | 0.551229233 | 4 mo | HOM |
| **1013609015** | 1 | mfbs | 991 | 7.256509205 | 0.001737222 | 27 | 0.000239402 | 3.720797319 | 4 mo | HOM |
| **1013608106** | 2 | mfbs | 991 | 7.256509205 | 0 | 0 | 0 | 0 | 4 mo | HOM |
| **1011937543** | 1 | mfbs | 991 | 7.256509205 | 0.005582499 | 82 | 0.000769309 | 11.30019927 | 8 mo | HOM |
| **1010419538** | 2 | mfbs | 991 | 7.256509205 | 3.38E-05 | 2 | 4.65E-06 | 0.275614616 | 4 mo | WILD |
| **1010419187** | 1 | mfbs | 991 | 7.256509205 | 0.004482372 | 126 | 0.000617704 | 17.36372082 | 8 mo | HOM |
| **1013978366** | 1 | mfbs | 991 | 7.256509205 | 6.4E-05 | 3 | 8.82E-06 | 0.413421924 | 2 mo | HOM |
| **1014416155** | 2 | mfbs | 991 | 7.256509205 | 0 | 0 | 0 | 0 | 2 mo | HOM |
| **1013978065** | 2 | mfbs | 991 | 7.256509205 | 0 | 0 | 0 | 0 | 2 mo | HOM |
| **1009934102** | 2 | mfbs | 991 | 7.256509205 | 8.22E-06 | 0 | 1.13E-06 | 0 | 4 mo | WILD |
| **1009322937** | 2 | mfbs | 991 | 7.256509205 | 7.19E-07 | 0 | 9.9E-08 | 0 | 8 mo | WILD |
| **1011080637** | 2 | mfbs | 991 | 7.256509205 | 0.001823223 | 55 | 0.000251253 | 7.579401947 | 8 mo | HOM |
| **1013609713** | 2 | mfbs | 991 | 7.256509205 | 1.56E-06 | 0 | 2.14E-07 | 0 | 2 mo | WILD |
| **1010418842** | 1 | mfbs | 991 | 7.256509205 | 0.006305807 | 158 | 0.000868986 | 21.77355468 | 8 mo | HOM |
| **1013288213** | 2 | mfbs | 991 | 7.256509205 | 5.55E-05 | 1 | 7.65E-06 | 0.137807308 | 4 mo | HOM |
| **1008976060** | 2 | mfbs | 991 | 7.256509205 | 0.000235189 | 9 | 3.24E-05 | 1.240265773 | 8 mo | WILD |
| **1013446243** | 1 | mfbs | 991 | 7.256509205 | 0.000515555 | 15 | 7.1E-05 | 2.067109622 | 4 mo | HOM |
| **1013288951** | 2 | mfbs | 991 | 7.256509205 | 0.001523274 | 13 | 0.000209918 | 1.791495006 | 4 mo | HOM |
| **1013978665** | 1 | MOs | 993 | 13.49494621 | 0.000262127 | 13 | 1.94E-05 | 0.963323587 | 2 mo | HOM |
| **1013445609** | 1 | MOs | 993 | 13.49494621 | 0.000865926 | 51 | 6.42E-05 | 3.779192535 | 4 mo | HOM |
| **1013609015** | 1 | MOs | 993 | 13.49494621 | 0.003294786 | 106 | 0.00024415 | 7.854792329 | 4 mo | HOM |
| **1013608106** | 2 | MOs | 993 | 13.49494621 | 0 | 0 | 0 | 0 | 4 mo | HOM |
| **1011937543** | 1 | MOs | 993 | 13.49494621 | 0.016803895 | 320 | 0.001245199 | 23.71258061 | 8 mo | HOM |
| **1010419538** | 2 | MOs | 993 | 13.49494621 | 6.15E-05 | 3 | 4.55E-06 | 0.222305443 | 4 mo | WILD |
| **1010419187** | 1 | MOs | 993 | 13.49494621 | 0.018832212 | 674 | 0.001395501 | 49.94462292 | 8 mo | HOM |
| **1013978366** | 1 | MOs | 993 | 13.49494621 | 1.67E-05 | 1 | 1.24E-06 | 0.074101814 | 2 mo | HOM |
| **1014416155** | 2 | MOs | 993 | 13.49494621 | 0 | 0 | 0 | 0 | 2 mo | HOM |
| **1013978065** | 2 | MOs | 993 | 13.49494621 | 0 | 0 | 0 | 0 | 2 mo | HOM |
| **1009934102** | 2 | MOs | 993 | 13.49494621 | 0.000145721 | 4 | 1.08E-05 | 0.296407258 | 4 mo | WILD |
| **1009322937** | 2 | MOs | 993 | 13.49494621 | 4.41E-05 | 3 | 3.27E-06 | 0.222305443 | 8 mo | WILD |
| **1011080637** | 2 | MOs | 993 | 13.49494621 | 0.007428911 | 369 | 0.000550496 | 27.34356952 | 8 mo | HOM |
| **1013609713** | 2 | MOs | 993 | 13.49494621 | 2.97E-06 | 0 | 2.2E-07 | 0 | 2 mo | WILD |
| **1010418842** | 1 | MOs | 993 | 13.49494621 | 0.010617488 | 529 | 0.000786775 | 39.19985983 | 8 mo | HOM |
| **1013288213** | 2 | MOs | 993 | 13.49494621 | 0.000528972 | 41 | 3.92E-05 | 3.038174391 | 4 mo | HOM |
| **1008976060** | 2 | MOs | 993 | 13.49494621 | 0.000786164 | 26 | 5.83E-05 | 1.926647175 | 8 mo | WILD |
| **1013446243** | 1 | MOs | 993 | 13.49494621 | 0.002329037 | 89 | 0.000172586 | 6.595061483 | 4 mo | HOM |
| **1013288951** | 2 | MOs | 993 | 13.49494621 | 0.004128231 | 160 | 0.000305909 | 11.85629031 | 4 mo | HOM |
| **1013978665** | 1 | Ald1 | 996 | 0.473435766 | 0 | 0 | 0 | 0 | 2 mo | HOM |
| **1013445609** | 1 | Ald1 | 996 | 0.473435766 | 0.000100874 | 5 | 0.000213068 | 10.5610948 | 4 mo | HOM |

| | | | | | | | | | | |
|---|---|---|---|---|---|---|---|---|---|---|
| **1013609015** | 1 | Ald1 | 996 | 0.473435766 | 7.94E-05 | 4 | 0.000167608 | 8.448875836 | 4 mo | HOM |
| **1013608106** | 2 | Ald1 | 996 | 0.473435766 | 0 | 0 | 0 | 0 | 4 mo | HOM |
| **1011937543** | 1 | Ald1 | 996 | 0.473435766 | 0.000529633 | 21 | 0.0011187 | 44.35659814 | 8 mo | HOM |
| **1010419538** | 2 | Ald1 | 996 | 0.473435766 | 1.68E-05 | 0 | 3.54E-05 | 0 | 4 mo | WILD |
| **1010419187** | 1 | Ald1 | 996 | 0.473435766 | 0.000492869 | 27 | 0.001041048 | 57.02991189 | 8 mo | HOM |
| **1013978366** | 1 | Ald1 | 996 | 0.473435766 | 2.55E-08 | 0 | 5.38E-08 | 0 | 2 mo | HOM |
| **1014416155** | 2 | Ald1 | 996 | 0.473435766 | 0 | 0 | 0 | 0 | 2 mo | HOM |
| **1013978065** | 2 | Ald1 | 996 | 0.473435766 | 0 | 0 | 0 | 0 | 2 mo | HOM |
| **1009934102** | 2 | Ald1 | 996 | 0.473435766 | 0 | 0 | 0 | 0 | 4 mo | WILD |
| **1009322937** | 2 | Ald1 | 996 | 0.473435766 | 1.66E-05 | 3 | 3.5E-05 | 6.336656877 | 8 mo | WILD |
| **1011080637** | 2 | Ald1 | 996 | 0.473435766 | 0.00015559 | 7 | 0.000328641 | 14.78553271 | 8 mo | HOM |
| **1013609713** | 2 | Ald1 | 996 | 0.473435766 | 0 | 0 | 0 | 0 | 2 mo | WILD |
| **1010418842** | 1 | Ald1 | 996 | 0.473435766 | 0.000385381 | 19 | 0.000814009 | 40.13216022 | 8 mo | HOM |
| **1013288213** | 2 | Ald1 | 996 | 0.473435766 | 2.99E-05 | 2 | 6.32E-05 | 4.224437918 | 4 mo | HOM |
| **1008976060** | 2 | Ald1 | 996 | 0.473435766 | 1.96E-05 | 4 | 4.14E-05 | 8.448875836 | 8 mo | WILD |
| **1013446243** | 1 | Ald1 | 996 | 0.473435766 | 7.53E-05 | 3 | 0.000159084 | 6.336656877 | 4 mo | HOM |
| **1013288951** | 2 | Ald1 | 996 | 0.473435766 | 0.000257845 | 8 | 0.000544625 | 16.89775167 | 4 mo | HOM |
| **1013978665** | 1 | root | 997 | 516.8152486 | 0.015815462 | 635 | 3.06E-05 | 1.228678917 | 2 mo | HOM |
| **1013445609** | 1 | root | 997 | 517.0840715 | 0.021612974 | 1372 | 4.18E-05 | 2.653340289 | 4 mo | HOM |
| **1013609015** | 1 | root | 997 | 513.9764511 | 0.143233576 | 2765 | 0.000278677 | 5.3796239 | 4 mo | HOM |
| **1013608106** | 2 | root | 997 | 516.8397345 | 8.41E-06 | 1 | 1.63E-08 | 0.001934836 | 4 mo | HOM |
| **1011937543** | 1 | root | 997 | 517.6282202 | 0.70336709 | 8157 | 0.001358827 | 15.7584144 | 8 mo | HOM |
| **1010419538** | 2 | root | 997 | 518.1603668 | 0.00536203 | 307 | 1.03E-05 | 0.592480668 | 4 mo | WILD |
| **1010419187** | 1 | root | 997 | 513.1151499 | 0.576364342 | 12722 | 0.001123265 | 24.793655 | 8 mo | HOM |
| **1013978366** | 1 | root | 997 | 519.9320211 | 0.034868758 | 889 | 6.71E-05 | 1.709838909 | 2 mo | HOM |
| **1014416155** | 2 | root | 997 | 518.4997974 | 0.000111494 | 12 | 2.15E-07 | 0.023143693 | 2 mo | HOM |
| **1013978065** | 2 | root | 997 | 520.4427727 | 0.007251773 | 323 | 1.39E-05 | 0.620625392 | 2 mo | HOM |
| **1009934102** | 2 | root | 997 | 515.7862793 | 0.008069532 | 446 | 1.56E-05 | 0.864699233 | 4 mo | WILD |
| **1009322937** | 2 | root | 997 | 512.5874188 | 0.010253082 | 521 | 2E-05 | 1.016411993 | 8 mo | WILD |
| **1011080637** | 2 | root | 997 | 512.1288905 | 0.339840911 | 9676 | 0.000663585 | 18.89368122 | 8 mo | HOM |
| **1013609713** | 2 | root | 997 | 521.2129141 | 0.002841349 | 155 | 5.45E-06 | 0.297383269 | 2 mo | WILD |
| **1010418842** | 1 | root | 997 | 515.6659372 | 0.535161262 | 12999 | 0.001037806 | 25.20818046 | 8 mo | HOM |
| **1013288213** | 2 | root | 997 | 517.9227734 | 0.067296551 | 1411 | 0.000129935 | 2.724344386 | 4 mo | HOM |
| **1008976060** | 2 | root | 997 | 518.8814561 | 0.071586295 | 2096 | 0.000137963 | 4.039458291 | 8 mo | WILD |
| **1013446243** | 1 | root | 997 | 515.9883481 | 0.061969874 | 2393 | 0.000120099 | 4.637701624 | 4 mo | HOM |
| **1013288951** | 2 | root | 997 | 516.1854796 | 0.136090601 | 2898 | 0.000263647 | 5.614260987 | 4 mo | HOM |
| **1013978665** | 1 | FS | 998 | 0.441418342 | 1.23E-10 | 0 | 2.78E-10 | 0 | 2 mo | HOM |
| **1013445609** | 1 | FS | 998 | 0.44141846 | 6.8E-06 | 1 | 1.54E-05 | 2.26542406 | 4 mo | HOM |
| **1013609015** | 1 | FS | 998 | 0.44141846 | 2.26E-05 | 3 | 5.12E-05 | 6.796272181 | 4 mo | HOM |
| **1013608106** | 2 | FS | 998 | 0.44141846 | 0 | 0 | 0 | 0 | 4 mo | HOM |
| **1011937543** | 1 | FS | 998 | 0.44141846 | 0.001124271 | 14 | 0.002546951 | 31.71593685 | 8 mo | HOM |
| **1010419538** | 2 | FS | 998 | 0.44141846 | 0 | 0 | 0 | 0 | 4 mo | WILD |
| **1010419187** | 1 | FS | 998 | 0.44141846 | 0.00049887 | 25 | 0.001130153 | 56.63560151 | 8 mo | HOM |
| **1013978366** | 1 | FS | 998 | 0.44141846 | 0 | 0 | 0 | 0 | 2 mo | HOM |
| **1014416155** | 2 | FS | 998 | 0.44141846 | 0 | 0 | 0 | 0 | 2 mo | HOM |
| **1013978065** | 2 | FS | 998 | 0.44141846 | 0 | 0 | 0 | 0 | 2 mo | HOM |
| **1009934102** | 2 | FS | 998 | 0.44141846 | 0 | 0 | 0 | 0 | 4 mo | WILD |
| **1009322937** | 2 | FS | 998 | 0.44141846 | 0 | 0 | 0 | 0 | 8 mo | WILD |
| **1011080637** | 2 | FS | 998 | 0.44141846 | 0.000248443 | 21 | 0.000562829 | 47.57390527 | 8 mo | HOM |
| **1013609713** | 2 | FS | 998 | 0.44141846 | 0 | 0 | 0 | 0 | 2 mo | WILD |
| **1010418842** | 1 | FS | 998 | 0.44141846 | 0.000643087 | 34 | 0.001456865 | 77.02441805 | 8 mo | HOM |
| **1013288213** | 2 | FS | 998 | 0.44141846 | 2.48E-05 | 3 | 5.62E-05 | 6.796272181 | 4 mo | HOM |
| **1008976060** | 2 | FS | 998 | 0.44141846 | 5.44E-06 | 1 | 1.23E-05 | 2.26542406 | 8 mo | WILD |
| **1013446243** | 1 | FS | 998 | 0.44141846 | 1.96E-05 | 2 | 4.43E-05 | 4.530848121 | 4 mo | HOM |
| **1013288951** | 2 | FS | 998 | 0.44141846 | 7.75E-06 | 1 | 1.75E-05 | 2.26542406 | 4 mo | HOM |
| **1013978665** | 1 | eps | 1000 | 1.265862205 | 0 | 0 | 0 | 0 | 2 mo | HOM |
| **1013445609** | 1 | eps | 1000 | 1.265862205 | 0 | 0 | 0 | 0 | 4 mo | HOM |
| **1013609015** | 1 | eps | 1000 | 1.265862205 | 0 | 0 | 0 | 0 | 4 mo | HOM |
| **1013608106** | 2 | eps | 1000 | 1.260374793 | 0 | 0 | 0 | 0 | 4 mo | HOM |

| | | | | | | | | | | |
|---|---|---|---|---|---|---|---|---|---|---|
| **1011937543** | 1 | eps | 1000 | 1.265862205 | 0.004845067 | 9 | 0.003827484 | 7.109778588 | 8 mo | HOM |
| **1010419538** | 2 | eps | 1000 | 1.265862205 | 0 | 0 | 0 | 0 | 4 mo | WILD |
| **1010419187** | 1 | eps | 1000 | 1.265862205 | 0 | 0 | 0 | 0 | 8 mo | HOM |
| **1013978366** | 1 | eps | 1000 | 1.265862205 | 0 | 0 | 0 | 0 | 2 mo | HOM |
| **1014416155** | 2 | eps | 1000 | 1.265862205 | 0 | 0 | 0 | 0 | 2 mo | HOM |
| **1013978065** | 2 | eps | 1000 | 1.265862205 | 0 | 0 | 0 | 0 | 2 mo | HOM |
| **1009934102** | 2 | eps | 1000 | 1.265862205 | 0 | 0 | 0 | 0 | 4 mo | WILD |
| **1009322937** | 2 | eps | 1000 | 1.265862205 | 0 | 0 | 0 | 0 | 8 mo | WILD |
| **1011080637** | 2 | eps | 1000 | 1.265862205 | 0 | 0 | 0 | 0 | 8 mo | HOM |
| **1013609713** | 2 | eps | 1000 | 1.265862205 | 0 | 0 | 0 | 0 | 2 mo | WILD |
| **1010418842** | 1 | eps | 1000 | 1.265862205 | 5.1E-05 | 4 | 4.02E-05 | 3.159901595 | 8 mo | HOM |
| **1013288213** | 2 | eps | 1000 | 1.265861852 | 0 | 0 | 0 | 0 | 4 mo | HOM |
| **1008976060** | 2 | eps | 1000 | 1.265862205 | 1.47E-05 | 1 | 1.16E-05 | 0.789975399 | 8 mo | WILD |
| **1013446243** | 1 | eps | 1000 | 1.265862205 | 0 | 0 | 0 | 0 | 4 mo | HOM |
| **1013288951** | 2 | eps | 1000 | 1.265862205 | 0 | 0 | 0 | 0 | 4 mo | HOM |
| **1013978665** | 1 | AUDp | 1002 | 2.21394936 | 5.9E-06 | 1 | 2.66E-06 | 0.451681515 | 2 mo | HOM |
| **1013445609** | 1 | AUDp | 1002 | 2.21394936 | 0.000193938 | 16 | 8.76E-05 | 7.226904232 | 4 mo | HOM |
| **1013609015** | 1 | AUDp | 1002 | 2.21394936 | 0.000705602 | 21 | 0.000318708 | 9.485311805 | 4 mo | HOM |
| **1013608106** | 2 | AUDp | 1002 | 2.21394936 | 0 | 0 | 0 | 0 | 4 mo | HOM |
| **1011937543** | 1 | AUDp | 1002 | 2.21394936 | 0.004229717 | 77 | 0.001910485 | 34.77947662 | 8 mo | HOM |
| **1010419538** | 2 | AUDp | 1002 | 2.21394936 | 0 | 0 | 0 | 0 | 4 mo | WILD |
| **1010419187** | 1 | AUDp | 1002 | 2.21394936 | 0.003526024 | 100 | 0.00159264 | 45.16815145 | 8 mo | HOM |
| **1013978366** | 1 | AUDp | 1002 | 2.21394936 | 1.01E-05 | 1 | 4.56E-06 | 0.451681515 | 2 mo | HOM |
| **1014416155** | 2 | AUDp | 1002 | 2.21394936 | 0 | 0 | 0 | 0 | 2 mo | HOM |
| **1013978065** | 2 | AUDp | 1002 | 2.21394936 | 0 | 0 | 0 | 0 | 2 mo | HOM |
| **1009934102** | 2 | AUDp | 1002 | 2.21394936 | 1.07E-06 | 1 | 4.86E-07 | 0.451681515 | 4 mo | WILD |
| **1009322937** | 2 | AUDp | 1002 | 2.21394936 | 1.6E-06 | 0 | 7.21E-07 | 0 | 8 mo | WILD |
| **1011080637** | 2 | AUDp | 1002 | 2.21394936 | 0.002022674 | 83 | 0.000913605 | 37.48956571 | 8 mo | HOM |
| **1013609713** | 2 | AUDp | 1002 | 2.21394936 | 1.28E-05 | 1 | 5.78E-06 | 0.451681515 | 2 mo | WILD |
| **1010418842** | 1 | AUDp | 1002 | 2.21394936 | 0.003816825 | 82 | 0.001723989 | 37.03788419 | 8 mo | HOM |
| **1013288213** | 2 | AUDp | 1002 | 2.21394936 | 0.000183055 | 12 | 8.27E-05 | 5.420178174 | 4 mo | HOM |
| **1008976060** | 2 | AUDp | 1002 | 2.21394936 | 1.53E-05 | 3 | 6.9E-06 | 1.355044544 | 8 mo | WILD |
| **1013446243** | 1 | AUDp | 1002 | 2.21394936 | 0.000381041 | 21 | 0.000172109 | 9.485311805 | 4 mo | HOM |
| **1013288951** | 2 | AUDp | 1002 | 2.21394936 | 0.001093301 | 27 | 0.000493824 | 12.19540089 | 4 mo | HOM |
| **1013978665** | 1 | PMv | 1004 | 0.200916542 | 0 | 0 | 0 | 0 | 2 mo | HOM |
| **1013445609** | 1 | PMv | 1004 | 0.200916542 | 0 | 0 | 0 | 0 | 4 mo | HOM |
| **1013609015** | 1 | PMv | 1004 | 0.200916542 | 1.28E-06 | 0 | 6.38E-06 | 0 | 4 mo | HOM |
| **1013608106** | 2 | PMv | 1004 | 0.200916542 | 0 | 0 | 0 | 0 | 4 mo | HOM |
| **1011937543** | 1 | PMv | 1004 | 0.200916542 | 0.002199863 | 3 | 0.010949139 | 14.9315729 | 8 mo | HOM |
| **1010419538** | 2 | PMv | 1004 | 0.200916542 | 0 | 0 | 0 | 0 | 4 mo | WILD |
| **1010419187** | 1 | PMv | 1004 | 0.200916542 | 0.000188676 | 2 | 0.000939077 | 9.954381934 | 8 mo | HOM |
| **1013978366** | 1 | PMv | 1004 | 0.200916542 | 0 | 0 | 0 | 0 | 2 mo | HOM |
| **1014416155** | 2 | PMv | 1004 | 0.200916542 | 0 | 0 | 0 | 0 | 2 mo | HOM |
| **1013978065** | 2 | PMv | 1004 | 0.200916542 | 0 | 0 | 0 | 0 | 2 mo | HOM |
| **1009934102** | 2 | PMv | 1004 | 0.200916542 | 0 | 0 | 0 | 0 | 4 mo | WILD |
| **1009322937** | 2 | PMv | 1004 | 0.200916542 | 1.34E-06 | 0 | 6.66E-06 | 0 | 8 mo | WILD |
| **1011080637** | 2 | PMv | 1004 | 0.200916542 | 1.33E-05 | 1 | 6.61E-05 | 4.977190967 | 8 mo | HOM |
| **1013609713** | 2 | PMv | 1004 | 0.200916542 | 0 | 0 | 0 | 0 | 2 mo | WILD |
| **1010418842** | 1 | PMv | 1004 | 0.200916542 | 4.88E-05 | 4 | 0.000242904 | 19.90876387 | 8 mo | HOM |
| **1013288213** | 2 | PMv | 1004 | 0.200916542 | 0 | 0 | 0 | 0 | 4 mo | HOM |
| **1008976060** | 2 | PMv | 1004 | 0.200916542 | 0 | 0 | 0 | 0 | 8 mo | WILD |
| **1013446243** | 1 | PMv | 1004 | 0.200916542 | 0 | 0 | 0 | 0 | 4 mo | HOM |
| **1013288951** | 2 | PMv | 1004 | 0.200916542 | 0 | 0 | 0 | 0 | 4 mo | HOM |
| **1013978665** | 1 | AUDp6b | 1005 | 0.056915813 | 0 | 0 | 0 | 0 | 2 mo | HOM |
| **1013445609** | 1 | AUDp6b | 1005 | 0.056915813 | 3.27E-06 | 1 | 5.75E-05 | 17.5698097 | 4 mo | HOM |
| **1013609015** | 1 | AUDp6b | 1005 | 0.056915813 | 1.34E-05 | 0 | 0.000235221 | 0 | 4 mo | HOM |
| **1013608106** | 2 | AUDp6b | 1005 | 0.056915813 | 0 | 0 | 0 | 0 | 4 mo | HOM |
| **1011937543** | 1 | AUDp6b | 1005 | 0.056915813 | 3.54E-05 | 1 | 0.000622799 | 17.5698097 | 8 mo | HOM |
| **1010419538** | 2 | AUDp6b | 1005 | 0.056915813 | 0 | 0 | 0 | 0 | 4 mo | WILD |

| | | | | | | | | | | |
|---|---|---|---|---|---|---|---|---|---|---|
| 1010419187 | 1 | AUDp6b | 1005 | 0.056915813 | 8.2E-05 | 4 | 0.001440678 | 70.27923881 | 8 mo | HOM |
| 1013978366 | 1 | AUDp6b | 1005 | 0.056915813 | 0 | 0 | 0 | 0 | 2 mo | HOM |
| 1014416155 | 2 | AUDp6b | 1005 | 0.056915813 | 0 | 0 | 0 | 0 | 2 mo | HOM |
| 1013978065 | 2 | AUDp6b | 1005 | 0.056915813 | 0 | 0 | 0 | 0 | 2 mo | HOM |
| 1009934102 | 2 | AUDp6b | 1005 | 0.056915813 | 0 | 0 | 0 | 0 | 4 mo | WILD |
| 1009322937 | 2 | AUDp6b | 1005 | 0.056915813 | 0 | 0 | 0 | 0 | 8 mo | WILD |
| 1011080637 | 2 | AUDp6b | 1005 | 0.056915813 | 4.05E-05 | 4 | 0.000711636 | 70.27923881 | 8 mo | HOM |
| 1013609713 | 2 | AUDp6b | 1005 | 0.056915813 | 0 | 0 | 0 | 0 | 2 mo | WILD |
| 1010418842 | 1 | AUDp6b | 1005 | 0.056915813 | 7.66E-05 | 4 | 0.001346487 | 70.27923881 | 8 mo | HOM |
| 1013288213 | 2 | AUDp6b | 1005 | 0.056915813 | 0 | 0 | 0 | 0 | 4 mo | HOM |
| 1008976060 | 2 | AUDp6b | 1005 | 0.056915813 | 1.54E-06 | 0 | 2.71E-05 | 0 | 8 mo | WILD |
| 1013446243 | 1 | AUDp6b | 1005 | 0.056915813 | 1.29E-05 | 1 | 0.000225871 | 17.5698097 | 4 mo | HOM |
| 1013288951 | 2 | AUDp6b | 1005 | 0.056915813 | 1.04E-05 | 1 | 0.000181955 | 17.5698097 | 4 mo | HOM |
| 1013978665 | 1 | SSp-tr1 | 1006 | 0.211855459 | 0 | 0 | 0 | 0 | 2 mo | HOM |
| 1013445609 | 1 | SSp-tr1 | 1006 | 0.211855459 | 2.9E-06 | 0 | 1.37E-05 | 0 | 4 mo | HOM |
| 1013609015 | 1 | SSp-tr1 | 1006 | 0.211855459 | 0.000164983 | 6 | 0.000778752 | 28.32119608 | 4 mo | HOM |
| 1013608106 | 2 | SSp-tr1 | 1006 | 0.211855459 | 0 | 0 | 0 | 0 | 4 mo | HOM |
| 1011937543 | 1 | SSp-tr1 | 1006 | 0.211855459 | 0.001191833 | 13 | 0.005625689 | 61.3625915 | 8 mo | HOM |
| 1010419538 | 2 | SSp-tr1 | 1006 | 0.211855459 | 0 | 0 | 0 | 0 | 4 mo | WILD |
| 1010419187 | 1 | SSp-tr1 | 1006 | 0.211855459 | 0.000667065 | 20 | 0.003148681 | 94.40398692 | 8 mo | HOM |
| 1013978366 | 1 | SSp-tr1 | 1006 | 0.211855459 | 0 | 0 | 0 | 0 | 2 mo | HOM |
| 1014416155 | 2 | SSp-tr1 | 1006 | 0.211855459 | 0 | 0 | 0 | 0 | 2 mo | HOM |
| 1013978065 | 2 | SSp-tr1 | 1006 | 0.211855459 | 0 | 0 | 0 | 0 | 2 mo | HOM |
| 1009934102 | 2 | SSp-tr1 | 1006 | 0.211855459 | 0 | 0 | 0 | 0 | 4 mo | WILD |
| 1009322937 | 2 | SSp-tr1 | 1006 | 0.211855459 | 0 | 0 | 0 | 0 | 8 mo | WILD |
| 1011080637 | 2 | SSp-tr1 | 1006 | 0.211855459 | 0.000842124 | 9 | 0.003974991 | 42.48179412 | 8 mo | HOM |
| 1013609713 | 2 | SSp-tr1 | 1006 | 0.211855459 | 0 | 0 | 0 | 0 | 2 mo | WILD |
| 1010418842 | 1 | SSp-tr1 | 1006 | 0.211855459 | 0.000603108 | 18 | 0.002846789 | 84.96358823 | 8 mo | HOM |
| 1013288213 | 2 | SSp-tr1 | 1006 | 0.211855459 | 8.91E-05 | 5 | 0.000420453 | 23.60099673 | 4 mo | HOM |
| 1008976060 | 2 | SSp-tr1 | 1006 | 0.211855459 | 2.11E-05 | 3 | 9.98E-05 | 14.16059804 | 8 mo | WILD |
| 1013446243 | 1 | SSp-tr1 | 1006 | 0.211855459 | 7.03E-05 | 5 | 0.000332025 | 23.60099673 | 4 mo | HOM |
| 1013288951 | 2 | SSp-tr1 | 1006 | 0.211855459 | 0.000106422 | 5 | 0.000502332 | 23.60099673 | 4 mo | HOM |
| 1013978665 | 1 | SIM | 1007 | 5.868211149 | 0.000755102 | 22 | 0.000128677 | 3.74901302 | 2 mo | HOM |
| 1013445609 | 1 | SIM | 1007 | 5.868211149 | 0 | 0 | 0 | 0 | 4 mo | HOM |
| 1013609015 | 1 | SIM | 1007 | 5.868211149 | 0.002155594 | 25 | 0.000367334 | 4.260242068 | 4 mo | HOM |
| 1013608106 | 2 | SIM | 1007 | 5.868211149 | 0 | 0 | 0 | 0 | 4 mo | HOM |
| 1011937543 | 1 | SIM | 1007 | 5.868211149 | 0.009979805 | 37 | 0.001700655 | 6.305158261 | 8 mo | HOM |
| 1010419538 | 2 | SIM | 1007 | 5.868211149 | 0.000153768 | 6 | 2.62E-05 | 1.022458096 | 4 mo | WILD |
| 1010419187 | 1 | SIM | 1007 | 5.868211149 | 0.007915242 | 82 | 0.001348834 | 13.97359398 | 8 mo | HOM |
| 1013978366 | 1 | SIM | 1007 | 5.868211149 | 0.001182501 | 27 | 0.00020151 | 4.601061433 | 2 mo | HOM |
| 1014416155 | 2 | SIM | 1007 | 5.868211149 | 0 | 0 | 0 | 0 | 2 mo | HOM |
| 1013978065 | 2 | SIM | 1007 | 5.868211149 | 0.000117903 | 2 | 2.01E-05 | 0.340819365 | 2 mo | HOM |
| 1009934102 | 2 | SIM | 1007 | 5.868211149 | 0.000219692 | 9 | 3.74E-05 | 1.533687144 | 4 mo | WILD |
| 1009322937 | 2 | SIM | 1007 | 5.868211149 | 0.000367411 | 19 | 6.26E-05 | 3.237783972 | 8 mo | WILD |
| 1011080637 | 2 | SIM | 1007 | 5.868211149 | 0.00338886 | 63 | 0.000577495 | 10.73581001 | 8 mo | HOM |
| 1013609713 | 2 | SIM | 1007 | 5.868211149 | 5.14E-05 | 3 | 8.76E-06 | 0.511229048 | 2 mo | WILD |
| 1010418842 | 1 | SIM | 1007 | 5.868211149 | 0.009764501 | 88 | 0.001663966 | 14.99605208 | 8 mo | HOM |
| 1013288213 | 2 | SIM | 1007 | 5.868211149 | 5.62E-05 | 5 | 9.57E-06 | 0.852048414 | 4 mo | HOM |
| 1008976060 | 2 | SIM | 1007 | 5.868211149 | 0.000333486 | 25 | 5.68E-05 | 4.260242068 | 8 mo | WILD |
| 1013446243 | 1 | SIM | 1007 | 5.868211149 | 3.17E-05 | 2 | 5.4E-06 | 0.340819365 | 4 mo | HOM |
| 1013288951 | 2 | SIM | 1007 | 5.868211149 | 0.002398275 | 27 | 0.000408689 | 4.601061433 | 4 mo | HOM |
| 1013978665 | 1 | GENd | 1008 | 1.465155066 | 0 | 0 | 0 | 0 | 2 mo | HOM |
| 1013445609 | 1 | GENd | 1008 | 1.465155066 | 3.3E-05 | 3 | 2.25E-05 | 2.047564841 | 4 mo | HOM |
| 1013609015 | 1 | GENd | 1008 | 1.465155066 | 0.00018662 | 4 | 0.000127372 | 2.730086455 | 4 mo | HOM |
| 1013608106 | 2 | GENd | 1008 | 1.465155066 | 0 | 0 | 0 | 0 | 4 mo | HOM |
| 1011937543 | 1 | GENd | 1008 | 1.465155066 | 0.002077826 | 35 | 0.001418161 | 23.88825648 | 8 mo | HOM |
| 1010419538 | 2 | GENd | 1008 | 1.465155066 | 0 | 0 | 0 | 0 | 4 mo | WILD |
| 1010419187 | 1 | GENd | 1008 | 1.465155066 | 0.000490796 | 18 | 0.000334979 | 12.28538905 | 8 mo | HOM |
| 1013978366 | 1 | GENd | 1008 | 1.465155066 | 0 | 0 | 0 | 0 | 2 mo | HOM |

| | | | | | | | | | | |
|---|---|---|---|---|---|---|---|---|---|---|
| **1014416155** | 2 | GENd | 1008 | 1.465155066 | 0 | 0 | 0 | 0 | 2 mo | HOM |
| **1013978065** | 2 | GENd | 1008 | 1.465155066 | 0 | 0 | 0 | 0 | 2 mo | HOM |
| **1009934102** | 2 | GENd | 1008 | 1.465155066 | 0 | 0 | 0 | 0 | 4 mo | WILD |
| **1009322937** | 2 | GENd | 1008 | 1.465155066 | 1.48E-05 | 3 | 1.01E-05 | 2.047564841 | 8 mo | WILD |
| **1011080637** | 2 | GENd | 1008 | 1.465155066 | 0.000539519 | 16 | 0.000368233 | 10.92034582 | 8 mo | HOM |
| **1013609713** | 2 | GENd | 1008 | 1.465155066 | 0 | 0 | 0 | 0 | 2 mo | WILD |
| **1010418842** | 1 | GENd | 1008 | 1.465155066 | 0.001496309 | 36 | 0.001021263 | 24.57077809 | 8 mo | HOM |
| **1013288213** | 2 | GENd | 1008 | 1.465155066 | 0 | 0 | 0 | 0 | 4 mo | HOM |
| **1008976060** | 2 | GENd | 1008 | 1.465155066 | 8.57E-05 | 5 | 5.85E-05 | 3.412608068 | 8 mo | WILD |
| **1013446243** | 1 | GENd | 1008 | 1.465155066 | 3.87E-05 | 2 | 2.64E-05 | 1.365043227 | 4 mo | HOM |
| **1013288951** | 2 | GENd | 1008 | 1.465155066 | 2.02E-05 | 3 | 1.38E-05 | 2.047564841 | 4 mo | HOM |
| **1013978665** | 1 | fiber tracts | 1009 | 47.44023388 | 0.001966185 | 64 | 4.14E-05 | 1.349065862 | 2 mo | HOM |
| **1013445609** | 1 | fiber tracts | 1009 | 47.47566316 | 0.001191327 | 69 | 2.51E-05 | 1.453376223 | 4 mo | HOM |
| **1013609015** | 1 | fiber tracts | 1009 | 46.60066396 | 0.005102135 | 154 | 0.000109486 | 3.304673945 | 4 mo | HOM |
| **1013608106** | 2 | fiber tracts | 1009 | 47.56730962 | 0 | 0 | 0 | 0 | 4 mo | HOM |
| **1011937543** | 1 | fiber tracts | 1009 | 47.61115843 | 0.050181263 | 466 | 0.001053981 | 9.787621545 | 8 mo | HOM |
| **1010419538** | 2 | fiber tracts | 1009 | 47.84944425 | 0.000363437 | 27 | 7.6E-06 | 0.564269877 | 4 mo | WILD |
| **1010419187** | 1 | fiber tracts | 1009 | 46.56557087 | 0.0193381 | 365 | 0.000415288 | 7.83840922 | 8 mo | HOM |
| **1013978366** | 1 | fiber tracts | 1009 | 48.22036029 | 0.002000468 | 84 | 4.15E-05 | 1.742002745 | 2 mo | HOM |
| **1014416155** | 2 | fiber tracts | 1009 | 47.96312801 | 1.89E-06 | 0 | 3.95E-08 | 0 | 2 mo | HOM |
| **1013978065** | 2 | fiber tracts | 1009 | 48.27503206 | 0.002464554 | 89 | 5.11E-05 | 1.843603126 | 2 mo | HOM |
| **1009934102** | 2 | fiber tracts | 1009 | 47.15614239 | 0.000743275 | 54 | 1.58E-05 | 1.145131838 | 4 mo | WILD |
| **1009322937** | 2 | fiber tracts | 1009 | 46.38074757 | 0.000160296 | 15 | 3.46E-06 | 0.323410052 | 8 mo | WILD |
| **1011080637** | 2 | fiber tracts | 1009 | 46.3096532 | 0.014729502 | 260 | 0.000318065 | 5.6143802 | 8 mo | HOM |
| **1013609713** | 2 | fiber tracts | 1009 | 48.33417953 | 0.000108433 | 15 | 2.24E-06 | 0.310339394 | 2 mo | WILD |
| **1010418842** | 1 | fiber tracts | 1009 | 47.25634888 | 0.021269506 | 556 | 0.000450088 | 11.76561485 | 8 mo | HOM |
| **1013288213** | 2 | fiber tracts | 1009 | 47.84859628 | 0.002712787 | 102 | 5.67E-05 | 2.131723978 | 4 mo | HOM |
| **1008976060** | 2 | fiber tracts | 1009 | 47.62356193 | 0.003462917 | 144 | 7.27E-05 | 3.02371335 | 8 mo | WILD |
| **1013446243** | 1 | fiber tracts | 1009 | 46.9767682 | 0.002125052 | 120 | 4.52E-05 | 2.554454139 | 4 mo | HOM |
| **1013288951** | 2 | fiber tracts | 1009 | 47.36589312 | 0.005237328 | 98 | 0.000110572 | 2.068999306 | 4 mo | HOM |
| **1013978665** | 1 | VISC4 | 1010 | 0.186810697 | 0 | 0 | 0 | 0 | 2 mo | HOM |
| **1013445609** | 1 | VISC4 | 1010 | 0.186810697 | 4.81E-05 | 4 | 0.000257718 | 21.4120501 | 4 mo | HOM |
| **1013609015** | 1 | VISC4 | 1010 | 0.186810697 | 6.62E-05 | 1 | 0.000354288 | 5.353012526 | 4 mo | HOM |
| **1013608106** | 2 | VISC4 | 1010 | 0.186810697 | 0 | 0 | 0 | 0 | 4 mo | HOM |
| **1011937543** | 1 | VISC4 | 1010 | 0.186810697 | 0.000189697 | 8 | 0.001015451 | 42.82410021 | 8 mo | HOM |
| **1010419538** | 2 | VISC4 | 1010 | 0.186810697 | 0 | 0 | 0 | 0 | 4 mo | WILD |
| **1010419187** | 1 | VISC4 | 1010 | 0.186810697 | 0.000135456 | 5 | 0.000725095 | 26.76506263 | 8 mo | HOM |
| **1013978366** | 1 | VISC4 | 1010 | 0.186810697 | 0 | 0 | 0 | 0 | 2 mo | HOM |
| **1014416155** | 2 | VISC4 | 1010 | 0.186810697 | 0 | 0 | 0 | 0 | 2 mo | HOM |
| **1013978065** | 2 | VISC4 | 1010 | 0.186810697 | 0 | 0 | 0 | 0 | 2 mo | HOM |
| **1009934102** | 2 | VISC4 | 1010 | 0.186810697 | 0 | 0 | 0 | 0 | 4 mo | WILD |
| **1009322937** | 2 | VISC4 | 1010 | 0.186810697 | 0 | 0 | 0 | 0 | 8 mo | WILD |
| **1011080637** | 2 | VISC4 | 1010 | 0.186810697 | 0.000112949 | 4 | 0.000604616 | 21.4120501 | 8 mo | HOM |
| **1013609713** | 2 | VISC4 | 1010 | 0.186810697 | 0 | 0 | 0 | 0 | 2 mo | WILD |
| **1010418842** | 1 | VISC4 | 1010 | 0.186810697 | 5.1E-05 | 3 | 0.000273144 | 16.05903758 | 8 mo | HOM |
| **1013288213** | 2 | VISC4 | 1010 | 0.186810697 | 0 | 0 | 0 | 0 | 4 mo | HOM |
| **1008976060** | 2 | VISC4 | 1010 | 0.186810697 | 0 | 0 | 0 | 0 | 8 mo | WILD |
| **1013446243** | 1 | VISC4 | 1010 | 0.186810697 | 3.38E-05 | 2 | 0.000181118 | 10.70602505 | 4 mo | HOM |
| **1013288951** | 2 | VISC4 | 1010 | 0.186810697 | 8.92E-05 | 4 | 0.000477446 | 21.4120501 | 4 mo | HOM |
| **1013978665** | 1 | AUDd | 1011 | 1.251544118 | 4.82E-06 | 1 | 3.85E-06 | 0.799012984 | 2 mo | HOM |
| **1013445609** | 1 | AUDd | 1011 | 1.251544118 | 0.000112258 | 9 | 8.97E-05 | 7.191116852 | 4 mo | HOM |
| **1013609015** | 1 | AUDd | 1011 | 1.251544118 | 0.000379176 | 17 | 0.000302967 | 13.58322072 | 4 mo | HOM |
| **1013608106** | 2 | AUDd | 1011 | 1.251544118 | 0 | 0 | 0 | 0 | 4 mo | HOM |
| **1011937543** | 1 | AUDd | 1011 | 1.251544118 | 0.001960868 | 35 | 0.001566759 | 27.96545442 | 8 mo | HOM |
| **1010419538** | 2 | AUDd | 1011 | 1.251544118 | 2.02E-06 | 0 | 1.61E-06 | 0 | 4 mo | WILD |
| **1010419187** | 1 | AUDd | 1011 | 1.251544118 | 0.001950307 | 67 | 0.001558321 | 53.5338699 | 8 mo | HOM |
| **1013978366** | 1 | AUDd | 1011 | 1.251544118 | 8.58E-06 | 1 | 6.86E-06 | 0.799012984 | 2 mo | HOM |
| **1014416155** | 2 | AUDd | 1011 | 1.251544118 | 0 | 0 | 0 | 0 | 2 mo | HOM |
| **1013978065** | 2 | AUDd | 1011 | 1.251544118 | 0 | 0 | 0 | 0 | 2 mo | HOM |

| | | | | | | | | | | |
|---|---|---|---|---|---|---|---|---|---|---|
| **1009934102** | 2 | AUDd | 1011 | 1.251544118 | 1.24E-06 | 0 | 9.92E-07 | 0 | 4 mo | WILD |
| **1009322937** | 2 | AUDd | 1011 | 1.251544118 | 1.33E-06 | 1 | 1.06E-06 | 0.799012984 | 8 mo | WILD |
| **1011080637** | 2 | AUDd | 1011 | 1.251544118 | 0.001000303 | 36 | 0.000799255 | 28.76446741 | 8 mo | HOM |
| **1013609713** | 2 | AUDd | 1011 | 1.251544118 | 2.6E-06 | 0 | 2.08E-06 | 0 | 2 mo | WILD |
| **1010418842** | 1 | AUDd | 1011 | 1.251544118 | 0.001573254 | 42 | 0.00125705 | 33.55854531 | 8 mo | HOM |
| **1013288213** | 2 | AUDd | 1011 | 1.251544118 | 0.000187838 | 12 | 0.000150085 | 9.588155803 | 4 mo | HOM |
| **1008976060** | 2 | AUDd | 1011 | 1.251544118 | 8.34E-06 | 1 | 6.66E-06 | 0.799012984 | 8 mo | WILD |
| **1013446243** | 1 | AUDd | 1011 | 1.251544118 | 0.000348879 | 14 | 0.000278759 | 11.18618177 | 4 mo | HOM |
| **1013288951** | 2 | AUDd | 1011 | 1.251544118 | 0.000618119 | 19 | 0.000493885 | 15.18124669 | 4 mo | HOM |
| **1013978665** | 1 | GENv | 1014 | 0.539345455 | 0 | 0 | 0 | 0 | 2 mo | HOM |
| **1013445609** | 1 | GENv | 1014 | 0.539345455 | 2.82E-06 | 1 | 5.23E-06 | 1.854099244 | 4 mo | HOM |
| **1013609015** | 1 | GENv | 1014 | 0.539345455 | 2.48E-05 | 1 | 4.6E-05 | 1.854099244 | 4 mo | HOM |
| **1013608106** | 2 | GENv | 1014 | 0.539345455 | 0 | 0 | 0 | 0 | 4 mo | HOM |
| **1011937543** | 1 | GENv | 1014 | 0.539345455 | 0.001341844 | 7 | 0.002487913 | 12.97869471 | 8 mo | HOM |
| **1010419538** | 2 | GENv | 1014 | 0.539345455 | 0 | 0 | 0 | 0 | 4 mo | WILD |
| **1010419187** | 1 | GENv | 1014 | 0.539345455 | 0.000205534 | 12 | 0.00038108 | 22.24919092 | 8 mo | HOM |
| **1013978366** | 1 | GENv | 1014 | 0.539345455 | 0 | 0 | 0 | 0 | 2 mo | HOM |
| **1014416155** | 2 | GENv | 1014 | 0.539345455 | 0 | 0 | 0 | 0 | 2 mo | HOM |
| **1013978065** | 2 | GENv | 1014 | 0.539345455 | 0 | 0 | 0 | 0 | 2 mo | HOM |
| **1009934102** | 2 | GENv | 1014 | 0.539345455 | 0 | 0 | 0 | 0 | 4 mo | WILD |
| **1009322937** | 2 | GENv | 1014 | 0.539345455 | 0 | 0 | 0 | 0 | 8 mo | WILD |
| **1011080637** | 2 | GENv | 1014 | 0.539345455 | 8.81E-05 | 6 | 0.000163335 | 11.12459546 | 8 mo | HOM |
| **1013609713** | 2 | GENv | 1014 | 0.539345455 | 0 | 0 | 0 | 0 | 2 mo | WILD |
| **1010418842** | 1 | GENv | 1014 | 0.539345455 | 0.000901158 | 17 | 0.001670837 | 31.51968714 | 8 mo | HOM |
| **1013288213** | 2 | GENv | 1014 | 0.539345455 | 0 | 0 | 0 | 0 | 4 mo | HOM |
| **1008976060** | 2 | GENv | 1014 | 0.539345455 | 3.94E-05 | 2 | 7.31E-05 | 3.708198487 | 8 mo | WILD |
| **1013446243** | 1 | GENv | 1014 | 0.539345455 | 0 | 0 | 0 | 0 | 4 mo | HOM |
| **1013288951** | 2 | GENv | 1014 | 0.539345455 | 0 | 0 | 0 | 0 | 4 mo | HOM |
| **1013978665** | 1 | ACAd5 | 1015 | 1.233717139 | 0 | 0 | 0 | 0 | 2 mo | HOM |
| **1013445609** | 1 | ACAd5 | 1015 | 1.233717139 | 0.000131206 | 5 | 0.00010635 | 4.052792849 | 4 mo | HOM |
| **1013609015** | 1 | ACAd5 | 1015 | 1.233717139 | 0.000448465 | 16 | 0.000363507 | 12.96893712 | 4 mo | HOM |
| **1013608106** | 2 | ACAd5 | 1015 | 1.233717139 | 0 | 0 | 0 | 0 | 4 mo | HOM |
| **1011937543** | 1 | ACAd5 | 1015 | 1.233717139 | 0.001089699 | 19 | 0.000883265 | 15.40061283 | 8 mo | HOM |
| **1010419538** | 2 | ACAd5 | 1015 | 1.233717139 | 0 | 0 | 0 | 0 | 4 mo | WILD |
| **1010419187** | 1 | ACAd5 | 1015 | 1.233717139 | 0.002816799 | 73 | 0.00228318 | 59.1707756 | 8 mo | HOM |
| **1013978366** | 1 | ACAd5 | 1015 | 1.233717139 | 0 | 0 | 0 | 0 | 2 mo | HOM |
| **1014416155** | 2 | ACAd5 | 1015 | 1.233717139 | 0 | 0 | 0 | 0 | 2 mo | HOM |
| **1013978065** | 2 | ACAd5 | 1015 | 1.233717139 | 0 | 0 | 0 | 0 | 2 mo | HOM |
| **1009934102** | 2 | ACAd5 | 1015 | 1.233717139 | 0 | 0 | 0 | 0 | 4 mo | WILD |
| **1009322937** | 2 | ACAd5 | 1015 | 1.233717139 | 0 | 0 | 0 | 0 | 8 mo | WILD |
| **1011080637** | 2 | ACAd5 | 1015 | 1.233717139 | 0.000813414 | 41 | 0.00065932 | 33.23290137 | 8 mo | HOM |
| **1013609713** | 2 | ACAd5 | 1015 | 1.233717139 | 0 | 0 | 0 | 0 | 2 mo | WILD |
| **1010418842** | 1 | ACAd5 | 1015 | 1.233717139 | 0.001211064 | 59 | 0.000981639 | 47.82295562 | 8 mo | HOM |
| **1013288213** | 2 | ACAd5 | 1015 | 1.233717139 | 6.07E-05 | 4 | 4.92E-05 | 3.24223428 | 4 mo | HOM |
| **1008976060** | 2 | ACAd5 | 1015 | 1.233717139 | 0 | 0 | 0 | 0 | 8 mo | WILD |
| **1013446243** | 1 | ACAd5 | 1015 | 1.233717139 | 0.000288545 | 9 | 0.000233883 | 7.295027129 | 4 mo | HOM |
| **1013288951** | 2 | ACAd5 | 1015 | 1.233717139 | 0.000404786 | 13 | 0.000328103 | 10.53726141 | 4 mo | HOM |
| **1013978665** | 1 | onl | 1016 | 3.377225824 | 0.001773729 | 54 | 0.000525203 | 15.9894549 | 2 mo | HOM |
| **1013445609** | 1 | onl | 1016 | 3.412653216 | 0.001042276 | 56 | 0.000305415 | 16.40951965 | 4 mo | HOM |
| **1013609015** | 1 | onl | 1016 | 2.537650259 | 0.002962098 | 107 | 0.00116726 | 42.16499087 | 4 mo | HOM |
| **1013608106** | 2 | onl | 1016 | 3.510911936 | 0 | 0 | 0 | 0 | 4 mo | HOM |
| **1011937543** | 1 | onl | 1016 | 3.548150682 | 0.004083566 | 58 | 0.0011509 | 16.34654365 | 8 mo | HOM |
| **1010419538** | 2 | onl | 1016 | 3.828038054 | 0.000235049 | 20 | 6.14E-05 | 5.224608459 | 4 mo | WILD |
| **1010419187** | 1 | onl | 1016 | 2.502556694 | 0.006780969 | 59 | 0.002709616 | 23.57588946 | 8 mo | HOM |
| **1013978366** | 1 | onl | 1016 | 4.157355686 | 0.001373778 | 73 | 0.000330445 | 17.55923849 | 2 mo | HOM |
| **1014416155** | 2 | onl | 1016 | 3.900118701 | 1.89E-06 | 0 | 4.86E-07 | 0 | 2 mo | HOM |
| **1013978065** | 2 | onl | 1016 | 4.212021184 | 0.001847231 | 80 | 0.000438562 | 18.9932568 | 2 mo | HOM |
| **1009934102** | 2 | onl | 1016 | 3.093131194 | 0.00066565 | 50 | 0.000215202 | 16.16484943 | 4 mo | WILD |
| **1009322937** | 2 | onl | 1016 | 2.317736378 | 7.1E-05 | 10 | 3.06E-05 | 4.314554535 | 8 mo | WILD |

| | | | | | | | | | | |
|---|---|---|---|---|---|---|---|---|---|---|
| **1011080637** | 2 | onl | 1016 | 2.246640435 | 0.007961491 | 65 | 0.003543732 | 28.93208855 | 8 mo | HOM |
| **1013609713** | 2 | onl | 1016 | 4.276575117 | 9.71E-05 | 15 | 2.27E-05 | 3.507479605 | 2 mo | WILD |
| **1010418842** | 1 | onl | 1016 | 3.193338938 | 0.005257556 | 124 | 0.001646413 | 38.83082956 | 8 mo | HOM |
| **1013288213** | 2 | onl | 1016 | 3.785588531 | 0.001642768 | 90 | 0.000433953 | 23.77437465 | 4 mo | HOM |
| **1008976060** | 2 | onl | 1016 | 3.560552307 | 0.001530567 | 70 | 0.000429868 | 19.65987127 | 8 mo | WILD |
| **1013446243** | 1 | onl | 1016 | 2.913760774 | 0.001266195 | 84 | 0.000434557 | 28.82872223 | 4 mo | HOM |
| **1013288951** | 2 | onl | 1016 | 3.302884749 | 0.001696866 | 69 | 0.000513753 | 20.89082885 | 4 mo | HOM |
| **1013978665** | 1 | AN | 1017 | 11.13223176 | 0.000283254 | 27 | 2.54E-05 | 2.425389678 | 2 mo | HOM |
| **1013445609** | 1 | AN | 1017 | 11.13223176 | 6.13E-06 | 1 | 5.51E-07 | 0.089829247 | 4 mo | HOM |
| **1013609015** | 1 | AN | 1017 | 11.13223176 | 0.002097645 | 22 | 0.00018843 | 1.976243442 | 4 mo | HOM |
| **1013608106** | 2 | AN | 1017 | 11.13223176 | 0 | 0 | 0 | 0 | 4 mo | HOM |
| **1011937543** | 1 | AN | 1017 | 11.13223176 | 0.02005045 | 53 | 0.001801117 | 4.760950109 | 8 mo | HOM |
| **1010419538** | 2 | AN | 1017 | 11.13223176 | 0.000253633 | 15 | 2.28E-05 | 1.34743871 | 4 mo | WILD |
| **1010419187** | 1 | AN | 1017 | 11.13223176 | 0.014707905 | 156 | 0.0013212 | 14.01336259 | 8 mo | HOM |
| **1013978366** | 1 | AN | 1017 | 11.13223176 | 0.005725202 | 94 | 0.000514291 | 8.443949251 | 2 mo | HOM |
| **1014416155** | 2 | AN | 1017 | 11.13223176 | 0 | 0 | 0 | 0 | 2 mo | HOM |
| **1013978065** | 2 | AN | 1017 | 11.13223176 | 1.21E-05 | 0 | 1.09E-06 | 0 | 2 mo | HOM |
| **1009934102** | 2 | AN | 1017 | 11.13223176 | 0.000213116 | 16 | 1.91E-05 | 1.437267958 | 4 mo | WILD |
| **1009322937** | 2 | AN | 1017 | 11.13223176 | 0.00010991 | 13 | 9.87E-06 | 1.167780216 | 8 mo | WILD |
| **1011080637** | 2 | AN | 1017 | 11.13223176 | 0.011739083 | 185 | 0.001054513 | 16.61841076 | 8 mo | HOM |
| **1013609713** | 2 | AN | 1017 | 11.13223176 | 9.79E-05 | 10 | 8.8E-06 | 0.898292473 | 2 mo | WILD |
| **1010418842** | 1 | AN | 1017 | 11.13223176 | 0.020687202 | 175 | 0.001858316 | 15.72011829 | 8 mo | HOM |
| **1013288213** | 2 | AN | 1017 | 11.13223176 | 0.000230659 | 7 | 2.07E-05 | 0.628804731 | 4 mo | HOM |
| **1008976060** | 2 | AN | 1017 | 11.13223176 | 0.000236747 | 27 | 2.13E-05 | 2.425389678 | 8 mo | WILD |
| **1013446243** | 1 | AN | 1017 | 11.13223176 | 0.000189039 | 6 | 1.7E-05 | 0.538975484 | 4 mo | HOM |
| **1013288951** | 2 | AN | 1017 | 11.13223176 | 0.000853509 | 38 | 7.67E-05 | 3.413511399 | 4 mo | HOM |
| **1013978665** | 1 | AUDv | 1018 | 1.873177782 | 0 | 0 | 0 | 0 | 2 mo | HOM |
| **1013445609** | 1 | AUDv | 1018 | 1.873177782 | 0.000125618 | 8 | 6.71E-05 | 4.270817259 | 4 mo | HOM |
| **1013609015** | 1 | AUDv | 1018 | 1.873177782 | 0.000434421 | 17 | 0.000231917 | 9.075486675 | 4 mo | HOM |
| **1013608106** | 2 | AUDv | 1018 | 1.873177782 | 0 | 0 | 0 | 0 | 4 mo | HOM |
| **1011937543** | 1 | AUDv | 1018 | 1.873177782 | 0.002542488 | 68 | 0.001357313 | 36.3019467 | 8 mo | HOM |
| **1010419538** | 2 | AUDv | 1018 | 1.873177782 | 0 | 0 | 0 | 0 | 4 mo | WILD |
| **1010419187** | 1 | AUDv | 1018 | 1.873177782 | 0.002729424 | 99 | 0.001457109 | 52.85136358 | 8 mo | HOM |
| **1013978366** | 1 | AUDv | 1018 | 1.873177782 | 6.07E-07 | 0 | 3.24E-07 | 0 | 2 mo | HOM |
| **1014416155** | 2 | AUDv | 1018 | 1.873177782 | 0 | 0 | 0 | 0 | 2 mo | HOM |
| **1013978065** | 2 | AUDv | 1018 | 1.873177782 | 0 | 0 | 0 | 0 | 2 mo | HOM |
| **1009934102** | 2 | AUDv | 1018 | 1.873177782 | 0 | 0 | 0 | 0 | 4 mo | WILD |
| **1009322937** | 2 | AUDv | 1018 | 1.873177782 | 8.27E-06 | 0 | 4.42E-06 | 0 | 8 mo | WILD |
| **1011080637** | 2 | AUDv | 1018 | 1.873177782 | 0.001292614 | 56 | 0.000690065 | 29.89572081 | 8 mo | HOM |
| **1013609713** | 2 | AUDv | 1018 | 1.873177782 | 0 | 0 | 0 | 0 | 2 mo | WILD |
| **1010418842** | 1 | AUDv | 1018 | 1.873177782 | 0.003135488 | 92 | 0.001673887 | 49.11439848 | 8 mo | HOM |
| **1013288213** | 2 | AUDv | 1018 | 1.873177782 | 0.000144461 | 10 | 7.71E-05 | 5.338521573 | 4 mo | HOM |
| **1008976060** | 2 | AUDv | 1018 | 1.873177782 | 2.59E-05 | 3 | 1.38E-05 | 1.601556472 | 8 mo | WILD |
| **1013446243** | 1 | AUDv | 1018 | 1.873177782 | 0.000228876 | 18 | 0.000122186 | 9.609338832 | 4 mo | HOM |
| **1013288951** | 2 | AUDv | 1018 | 1.873177782 | 0.000948966 | 32 | 0.000506608 | 17.08326903 | 4 mo | HOM |
| **1013978665** | 1 | PO | 1020 | 1.290478347 | 0 | 0 | 0 | 0 | 2 mo | HOM |
| **1013445609** | 1 | PO | 1020 | 1.290478347 | 0 | 0 | 0 | 0 | 4 mo | HOM |
| **1013609015** | 1 | PO | 1020 | 1.290478347 | 2.58E-05 | 2 | 2E-05 | 1.549812908 | 4 mo | HOM |
| **1013608106** | 2 | PO | 1020 | 1.290478347 | 0 | 0 | 0 | 0 | 4 mo | HOM |
| **1011937543** | 1 | PO | 1020 | 1.290478347 | 0.001176445 | 19 | 0.000911635 | 14.72322263 | 8 mo | HOM |
| **1010419538** | 2 | PO | 1020 | 1.290478347 | 0 | 0 | 0 | 0 | 4 mo | WILD |
| **1010419187** | 1 | PO | 1020 | 1.290478347 | 0.00112051 | 26 | 0.00086829 | 20.1475678 | 8 mo | HOM |
| **1013978366** | 1 | PO | 1020 | 1.290478347 | 0 | 0 | 0 | 0 | 2 mo | HOM |
| **1014416155** | 2 | PO | 1020 | 1.290478347 | 0 | 0 | 0 | 0 | 2 mo | HOM |
| **1013978065** | 2 | PO | 1020 | 1.290478347 | 0 | 0 | 0 | 0 | 2 mo | HOM |
| **1009934102** | 2 | PO | 1020 | 1.290478347 | 0 | 0 | 0 | 0 | 4 mo | WILD |
| **1009322937** | 2 | PO | 1020 | 1.290478347 | 7.92E-06 | 1 | 6.14E-06 | 0.774906454 | 8 mo | WILD |
| **1011080637** | 2 | PO | 1020 | 1.290478347 | 0.001374495 | 39 | 0.001065105 | 30.22135171 | 8 mo | HOM |
| **1013609713** | 2 | PO | 1020 | 1.290478347 | 0 | 0 | 0 | 0 | 2 mo | WILD |

| | | | | | | | | | | |
|---|---|---|---|---|---|---|---|---|---|---|
| **1010418842** | 1 | PO | 1020 | 1.290478347 | 0.003061786 | 48 | 0.002372597 | 37.19550979 | 8 mo | HOM |
| **1013288213** | 2 | PO | 1020 | 1.290478347 | 1.97E-05 | 1 | 1.53E-05 | 0.774906454 | 4 mo | HOM |
| **1008976060** | 2 | PO | 1020 | 1.290478347 | 2.13E-05 | 4 | 1.65E-05 | 3.099625816 | 8 mo | WILD |
| **1013446243** | 1 | PO | 1020 | 1.290478347 | 7.82E-06 | 1 | 6.06E-06 | 0.774906454 | 4 mo | HOM |
| **1013288951** | 2 | PO | 1020 | 1.290478347 | 1.42E-05 | 1 | 1.1E-05 | 0.774906454 | 4 mo | HOM |
| **1013978665** | 1 | MOs6a | 1021 | 2.449596867 | 0 | 0 | 0 | 0 | 2 mo | HOM |
| **1013445609** | 1 | MOs6a | 1021 | 2.449596867 | 8.93E-05 | 7 | 3.64E-05 | 2.85761306 | 4 mo | HOM |
| **1013609015** | 1 | MOs6a | 1021 | 2.449596867 | 0.000459689 | 17 | 0.000187659 | 6.939917432 | 4 mo | HOM |
| **1013608106** | 2 | MOs6a | 1021 | 2.449596867 | 0 | 0 | 0 | 0 | 4 mo | HOM |
| **1011937543** | 1 | MOs6a | 1021 | 2.449596867 | 0.002631692 | 48 | 0.001074337 | 19.59506098 | 8 mo | HOM |
| **1010419538** | 2 | MOs6a | 1021 | 2.449596867 | 0 | 0 | 0 | 0 | 4 mo | WILD |
| **1010419187** | 1 | MOs6a | 1021 | 2.449596867 | 0.002162368 | 110 | 0.000882744 | 44.90534809 | 8 mo | HOM |
| **1013978366** | 1 | MOs6a | 1021 | 2.449596867 | 0 | 0 | 0 | 0 | 2 mo | HOM |
| **1014416155** | 2 | MOs6a | 1021 | 2.449596867 | 0 | 0 | 0 | 0 | 2 mo | HOM |
| **1013978065** | 2 | MOs6a | 1021 | 2.449596867 | 0 | 0 | 0 | 0 | 2 mo | HOM |
| **1009934102** | 2 | MOs6a | 1021 | 2.449596867 | 0 | 0 | 0 | 0 | 4 mo | WILD |
| **1009322937** | 2 | MOs6a | 1021 | 2.449596867 | 0 | 0 | 0 | 0 | 8 mo | WILD |
| **1011080637** | 2 | MOs6a | 1021 | 2.449596867 | 0.001643253 | 92 | 0.000670826 | 37.55720022 | 8 mo | HOM |
| **1013609713** | 2 | MOs6a | 1021 | 2.449596867 | 0 | 0 | 0 | 0 | 2 mo | WILD |
| **1010418842** | 1 | MOs6a | 1021 | 2.449596867 | 0.001638948 | 90 | 0.000669069 | 36.74073935 | 8 mo | HOM |
| **1013288213** | 2 | MOs6a | 1021 | 2.449596867 | 4.33E-05 | 6 | 1.77E-05 | 2.449382623 | 4 mo | HOM |
| **1008976060** | 2 | MOs6a | 1021 | 2.449596867 | 0 | 0 | 0 | 0 | 8 mo | WILD |
| **1013446243** | 1 | MOs6a | 1021 | 2.449596867 | 0.000247716 | 16 | 0.000101125 | 6.531686995 | 4 mo | HOM |
| **1013288951** | 2 | MOs6a | 1021 | 2.449596867 | 0.000827529 | 28 | 0.000337822 | 11.43045224 | 4 mo | HOM |
| **1013978665** | 1 | GPe | 1022 | 1.612112544 | 0 | 0 | 0 | 0 | 2 mo | HOM |
| **1013445609** | 1 | GPe | 1022 | 1.612112544 | 1.3E-05 | 1 | 8.04E-06 | 0.620304087 | 4 mo | HOM |
| **1013609015** | 1 | GPe | 1022 | 1.612112544 | 1.6E-05 | 1 | 9.91E-06 | 0.620304087 | 4 mo | HOM |
| **1013608106** | 2 | GPe | 1022 | 1.612112544 | 0 | 0 | 0 | 0 | 4 mo | HOM |
| **1011937543** | 1 | GPe | 1022 | 1.612112544 | 0.001366765 | 30 | 0.00084781 | 18.60912261 | 8 mo | HOM |
| **1010419538** | 2 | GPe | 1022 | 1.612112544 | 0 | 0 | 0 | 0 | 4 mo | WILD |
| **1010419187** | 1 | GPe | 1022 | 1.612112544 | 0.000376436 | 27 | 0.000233505 | 16.74821035 | 8 mo | HOM |
| **1013978366** | 1 | GPe | 1022 | 1.612112544 | 0 | 0 | 0 | 0 | 2 mo | HOM |
| **1014416155** | 2 | GPe | 1022 | 1.612112544 | 0 | 0 | 0 | 0 | 2 mo | HOM |
| **1013978065** | 2 | GPe | 1022 | 1.612112544 | 0 | 0 | 0 | 0 | 2 mo | HOM |
| **1009934102** | 2 | GPe | 1022 | 1.612112544 | 0 | 0 | 0 | 0 | 4 mo | WILD |
| **1009322937** | 2 | GPe | 1022 | 1.612112544 | 0 | 0 | 0 | 0 | 8 mo | WILD |
| **1011080637** | 2 | GPe | 1022 | 1.612112544 | 0.000458844 | 19 | 0.000284623 | 11.78577766 | 8 mo | HOM |
| **1013609713** | 2 | GPe | 1022 | 1.612112544 | 0 | 0 | 0 | 0 | 2 mo | WILD |
| **1010418842** | 1 | GPe | 1022 | 1.612112544 | 0.000838822 | 42 | 0.000520325 | 26.05277166 | 8 mo | HOM |
| **1013288213** | 2 | GPe | 1022 | 1.612112544 | 7.94E-06 | 1 | 4.92E-06 | 0.620304087 | 4 mo | HOM |
| **1008976060** | 2 | GPe | 1022 | 1.612112544 | 5.02E-06 | 1 | 3.11E-06 | 0.620304087 | 8 mo | WILD |
| **1013446243** | 1 | GPe | 1022 | 1.612112544 | 9.66E-06 | 1 | 5.99E-06 | 0.620304087 | 4 mo | HOM |
| **1013288951** | 2 | GPe | 1022 | 1.612112544 | 7.23E-06 | 1 | 4.48E-06 | 0.620304087 | 4 mo | HOM |
| **1013978665** | 1 | AUDv5 | 1023 | 0.622944095 | 0 | 0 | 0 | 0 | 2 mo | HOM |
| **1013445609** | 1 | AUDv5 | 1023 | 0.622944095 | 8E-06 | 1 | 1.28E-05 | 1.605280486 | 4 mo | HOM |
| **1013609015** | 1 | AUDv5 | 1023 | 0.622944095 | 0.000126015 | 4 | 0.000202289 | 6.421121945 | 4 mo | HOM |
| **1013608106** | 2 | AUDv5 | 1023 | 0.622944095 | 0 | 0 | 0 | 0 | 4 mo | HOM |
| **1011937543** | 1 | AUDv5 | 1023 | 0.622944095 | 0.000482438 | 19 | 0.000774449 | 30.50032924 | 8 mo | HOM |
| **1010419538** | 2 | AUDv5 | 1023 | 0.622944095 | 0 | 0 | 0 | 0 | 4 mo | WILD |
| **1010419187** | 1 | AUDv5 | 1023 | 0.622944095 | 0.000780788 | 35 | 0.001253384 | 56.18481702 | 8 mo | HOM |
| **1013978366** | 1 | AUDv5 | 1023 | 0.622944095 | 6.07E-07 | 0 | 9.74E-07 | 0 | 2 mo | HOM |
| **1014416155** | 2 | AUDv5 | 1023 | 0.622944095 | 0 | 0 | 0 | 0 | 2 mo | HOM |
| **1013978065** | 2 | AUDv5 | 1023 | 0.622944095 | 0 | 0 | 0 | 0 | 2 mo | HOM |
| **1009934102** | 2 | AUDv5 | 1023 | 0.622944095 | 0 | 0 | 0 | 0 | 4 mo | WILD |
| **1009322937** | 2 | AUDv5 | 1023 | 0.622944095 | 0 | 0 | 0 | 0 | 8 mo | WILD |
| **1011080637** | 2 | AUDv5 | 1023 | 0.622944095 | 0.000411101 | 23 | 0.000659933 | 36.92145118 | 8 mo | HOM |
| **1013609713** | 2 | AUDv5 | 1023 | 0.622944095 | 0 | 0 | 0 | 0 | 2 mo | WILD |
| **1010418842** | 1 | AUDv5 | 1023 | 0.622944095 | 0.000936299 | 29 | 0.001503022 | 46.5531341 | 8 mo | HOM |
| **1013288213** | 2 | AUDv5 | 1023 | 0.622944095 | 3.76E-05 | 3 | 6.03E-05 | 4.815841459 | 4 mo | HOM |

| | | | | | | | | | | |
|---|---|---|---|---|---|---|---|---|---|---|
| **1008976060** | 2 | AUDv5 | 1023 | 0.622944095 | 4.05E-06 | 1 | 6.51E-06 | 1.605280486 | 8 mo | WILD |
| **1013446243** | 1 | AUDv5 | 1023 | 0.622944095 | 5.03E-05 | 5 | 8.07E-05 | 8.026402431 | 4 mo | HOM |
| **1013288951** | 2 | AUDv5 | 1023 | 0.622944095 | 0.00017358 | 7 | 0.000278645 | 11.2369634 | 4 mo | HOM |
| **1013978665** | 1 | PRM | 1025 | 5.030817299 | 0.000366461 | 14 | 7.28E-05 | 2.782848028 | 2 mo | HOM |
| **1013445609** | 1 | PRM | 1025 | 5.030817299 | 6.75E-05 | 2 | 1.34E-05 | 0.397549718 | 4 mo | HOM |
| **1013609015** | 1 | PRM | 1025 | 5.030817299 | 0.000987361 | 14 | 0.000196262 | 2.782848028 | 4 mo | HOM |
| **1013608106** | 2 | PRM | 1025 | 5.030817299 | 0 | 0 | 0 | 0 | 4 mo | HOM |
| **1011937543** | 1 | PRM | 1025 | 5.030817299 | 0.009746629 | 29 | 0.001937385 | 5.764470915 | 8 mo | HOM |
| **1010419538** | 2 | PRM | 1025 | 5.030817299 | 2.9E-05 | 4 | 5.77E-06 | 0.795099437 | 4 mo | WILD |
| **1010419187** | 1 | PRM | 1025 | 5.030817299 | 0.006054246 | 58 | 0.001203432 | 11.52894183 | 8 mo | HOM |
| **1013978366** | 1 | PRM | 1025 | 5.030817299 | 0.001634326 | 35 | 0.000324863 | 6.95712007 | 2 mo | HOM |
| **1014416155** | 2 | PRM | 1025 | 5.030817299 | 0 | 0 | 0 | 0 | 2 mo | HOM |
| **1013978065** | 2 | PRM | 1025 | 5.030817299 | 0 | 0 | 0 | 0 | 2 mo | HOM |
| **1009934102** | 2 | PRM | 1025 | 5.030817299 | 0.000151346 | 7 | 3.01E-05 | 1.391424014 | 4 mo | WILD |
| **1009322937** | 2 | PRM | 1025 | 5.030817299 | 0.000250482 | 18 | 4.98E-05 | 3.577947464 | 8 mo | WILD |
| **1011080637** | 2 | PRM | 1025 | 5.030817299 | 0.002083373 | 52 | 0.000414122 | 10.33629268 | 8 mo | HOM |
| **1013609713** | 2 | PRM | 1025 | 5.00709785 | 0.000424207 | 5 | 8.47E-05 | 0.998582442 | 2 mo | WILD |
| **1010418842** | 1 | PRM | 1025 | 5.030817299 | 0.011254062 | 79 | 0.002237024 | 15.70321387 | 8 mo | HOM |
| **1013288213** | 2 | PRM | 1025 | 5.030817299 | 4.61E-05 | 3 | 9.16E-06 | 0.596324577 | 4 mo | HOM |
| **1008976060** | 2 | PRM | 1025 | 5.030817299 | 0.000165492 | 11 | 3.29E-05 | 2.186523451 | 8 mo | WILD |
| **1013446243** | 1 | PRM | 1025 | 5.030817299 | 0.000534068 | 10 | 0.000106159 | 1.987748591 | 4 mo | HOM |
| **1013288951** | 2 | PRM | 1025 | 5.030817299 | 0.002609229 | 14 | 0.000518649 | 2.782848028 | 4 mo | HOM |
| **1013978665** | 1 | SSp-ul6b | 1026 | 0.0754578 | 0 | 0 | 0 | 0 | 2 mo | HOM |
| **1013445609** | 1 | SSp-ul6b | 1026 | 0.0754578 | 0 | 0 | 0 | 0 | 4 mo | HOM |
| **1013609015** | 1 | SSp-ul6b | 1026 | 0.0754578 | 3.16E-07 | 0 | 4.19E-06 | 0 | 4 mo | HOM |
| **1013608106** | 2 | SSp-ul6b | 1026 | 0.0754578 | 0 | 0 | 0 | 0 | 4 mo | HOM |
| **1011937543** | 1 | SSp-ul6b | 1026 | 0.0754578 | 1.79E-05 | 0 | 0.000236987 | 0 | 8 mo | HOM |
| **1010419538** | 2 | SSp-ul6b | 1026 | 0.0754578 | 0 | 0 | 0 | 0 | 4 mo | WILD |
| **1010419187** | 1 | SSp-ul6b | 1026 | 0.0754578 | 3.87E-05 | 1 | 0.000512913 | 13.25244045 | 8 mo | HOM |
| **1013978366** | 1 | SSp-ul6b | 1026 | 0.0754578 | 0 | 0 | 0 | 0 | 2 mo | HOM |
| **1014416155** | 2 | SSp-ul6b | 1026 | 0.0754578 | 0 | 0 | 0 | 0 | 2 mo | HOM |
| **1013978065** | 2 | SSp-ul6b | 1026 | 0.0754578 | 0 | 0 | 0 | 0 | 2 mo | HOM |
| **1009934102** | 2 | SSp-ul6b | 1026 | 0.0754578 | 0 | 0 | 0 | 0 | 4 mo | WILD |
| **1009322937** | 2 | SSp-ul6b | 1026 | 0.0754578 | 0 | 0 | 0 | 0 | 8 mo | WILD |
| **1011080637** | 2 | SSp-ul6b | 1026 | 0.0754578 | 4.7E-05 | 2 | 0.000622533 | 26.50488091 | 8 mo | HOM |
| **1013609713** | 2 | SSp-ul6b | 1026 | 0.0754578 | 0 | 0 | 0 | 0 | 2 mo | WILD |
| **1010418842** | 1 | SSp-ul6b | 1026 | 0.0754578 | 8.8E-06 | 0 | 0.000116563 | 0 | 8 mo | HOM |
| **1013288213** | 2 | SSp-ul6b | 1026 | 0.0754578 | 1.22E-07 | 0 | 1.62E-06 | 0 | 4 mo | HOM |
| **1008976060** | 2 | SSp-ul6b | 1026 | 0.0754578 | 0 | 0 | 0 | 0 | 8 mo | WILD |
| **1013446243** | 1 | SSp-ul6b | 1026 | 0.0754578 | 4.24E-09 | 0 | 5.62E-08 | 0 | 4 mo | HOM |
| **1013288951** | 2 | SSp-ul6b | 1026 | 0.0754578 | 4.15E-06 | 1 | 5.49E-05 | 13.25244045 | 4 mo | HOM |
| **1013978665** | 1 | AUDpo | 1027 | 0.630707859 | 0 | 0 | 0 | 0 | 2 mo | HOM |
| **1013445609** | 1 | AUDpo | 1027 | 0.630707859 | 9.74E-05 | 9 | 0.000154422 | 14.26968109 | 4 mo | HOM |
| **1013609015** | 1 | AUDpo | 1027 | 0.630707859 | 0.000350836 | 14 | 0.000556258 | 22.19728169 | 4 mo | HOM |
| **1013608106** | 2 | AUDpo | 1027 | 0.630707859 | 0 | 0 | 0 | 0 | 4 mo | HOM |
| **1011937543** | 1 | AUDpo | 1027 | 0.630707859 | 0.001262692 | 22 | 0.002002024 | 34.88144265 | 8 mo | HOM |
| **1010419538** | 2 | AUDpo | 1027 | 0.630707859 | 0 | 0 | 0 | 0 | 4 mo | WILD |
| **1010419187** | 1 | AUDpo | 1027 | 0.630707859 | 0.000906571 | 33 | 0.001437386 | 52.32216398 | 8 mo | HOM |
| **1013978366** | 1 | AUDpo | 1027 | 0.630707859 | 0 | 0 | 0 | 0 | 2 mo | HOM |
| **1014416155** | 2 | AUDpo | 1027 | 0.630707859 | 0 | 0 | 0 | 0 | 2 mo | HOM |
| **1013978065** | 2 | AUDpo | 1027 | 0.630707859 | 0 | 0 | 0 | 0 | 2 mo | HOM |
| **1009934102** | 2 | AUDpo | 1027 | 0.630707859 | 0 | 0 | 0 | 0 | 4 mo | WILD |
| **1009322937** | 2 | AUDpo | 1027 | 0.630707859 | 0 | 0 | 0 | 0 | 8 mo | WILD |
| **1011080637** | 2 | AUDpo | 1027 | 0.630707859 | 0.000483457 | 18 | 0.000766531 | 28.53936217 | 8 mo | HOM |
| **1013609713** | 2 | AUDpo | 1027 | 0.630707859 | 0 | 0 | 0 | 0 | 2 mo | WILD |
| **1010418842** | 1 | AUDpo | 1027 | 0.630707859 | 0.000907379 | 27 | 0.001438668 | 42.80904326 | 8 mo | HOM |
| **1013288213** | 2 | AUDpo | 1027 | 0.630707859 | 1.99E-05 | 3 | 3.16E-05 | 4.756560362 | 4 mo | HOM |
| **1008976060** | 2 | AUDpo | 1027 | 0.630707859 | 3.92E-05 | 3 | 6.22E-05 | 4.756560362 | 8 mo | WILD |
| **1013446243** | 1 | AUDpo | 1027 | 0.630707859 | 0.000133731 | 7 | 0.000212034 | 11.09864084 | 4 mo | HOM |

| | | | | | | | | | | |
|---|---|---|---|---|---|---|---|---|---|---|
| 1013288951 | 2 | AUDpo | 1027 | 0.630707859 | 9.07E-05 | 9 | 0.000143774 | 14.26968109 | 4 mo | HOM |
| 1013978665 | 1 | POL | 1029 | 0.211700933 | 0 | 0 | 0 | 0 | 2 mo | HOM |
| 1013445609 | 1 | POL | 1029 | 0.211700933 | 0 | 0 | 0 | 0 | 4 mo | HOM |
| 1013609015 | 1 | POL | 1029 | 0.211700933 | 0 | 0 | 0 | 0 | 4 mo | HOM |
| 1013608106 | 2 | POL | 1029 | 0.211700933 | 0 | 0 | 0 | 0 | 4 mo | HOM |
| 1011937543 | 1 | POL | 1029 | 0.211700933 | 0.000106117 | 4 | 0.000501261 | 18.894579 | 8 mo | HOM |
| 1010419538 | 2 | POL | 1029 | 0.211700933 | 0 | 0 | 0 | 0 | 4 mo | WILD |
| 1010419187 | 1 | POL | 1029 | 0.211700933 | 1.59E-05 | 3 | 7.52E-05 | 14.17093425 | 8 mo | HOM |
| 1013978366 | 1 | POL | 1029 | 0.211700933 | 0 | 0 | 0 | 0 | 2 mo | HOM |
| 1014416155 | 2 | POL | 1029 | 0.211700933 | 0 | 0 | 0 | 0 | 2 mo | HOM |
| 1013978065 | 2 | POL | 1029 | 0.211700933 | 0 | 0 | 0 | 0 | 2 mo | HOM |
| 1009934102 | 2 | POL | 1029 | 0.211700933 | 0 | 0 | 0 | 0 | 4 mo | WILD |
| 1009322937 | 2 | POL | 1029 | 0.211700933 | 0 | 0 | 0 | 0 | 8 mo | WILD |
| 1011080637 | 2 | POL | 1029 | 0.211700933 | 5.82E-06 | 0 | 2.75E-05 | 0 | 8 mo | HOM |
| 1013609713 | 2 | POL | 1029 | 0.211700933 | 0 | 0 | 0 | 0 | 2 mo | WILD |
| 1010418842 | 1 | POL | 1029 | 0.211700933 | 0.000129908 | 3 | 0.00061364 | 14.17093425 | 8 mo | HOM |
| 1013288213 | 2 | POL | 1029 | 0.211700933 | 0 | 0 | 0 | 0 | 4 mo | HOM |
| 1008976060 | 2 | POL | 1029 | 0.211700933 | 9.1E-06 | 1 | 4.3E-05 | 4.723644751 | 8 mo | WILD |
| 1013446243 | 1 | POL | 1029 | 0.211700933 | 0 | 0 | 0 | 0 | 4 mo | HOM |
| 1013288951 | 2 | POL | 1029 | 0.211700933 | 0 | 0 | 0 | 0 | 4 mo | HOM |
| 1013978665 | 1 | SSp-ll1 | 1030 | 0.308626559 | 2.64E-05 | 1 | 8.57E-05 | 3.240161844 | 2 mo | HOM |
| 1013445609 | 1 | SSp-ll1 | 1030 | 0.308626559 | 1.56E-06 | 0 | 5.07E-06 | 0 | 4 mo | HOM |
| 1013609015 | 1 | SSp-ll1 | 1030 | 0.308626559 | 0.000229008 | 11 | 0.000742023 | 35.64178029 | 4 mo | HOM |
| 1013608106 | 2 | SSp-ll1 | 1030 | 0.308626559 | 0 | 0 | 0 | 0 | 4 mo | HOM |
| 1011937543 | 1 | SSp-ll1 | 1030 | 0.308626559 | 0.0020839 | 13 | 0.006752173 | 42.12210398 | 8 mo | HOM |
| 1010419538 | 2 | SSp-ll1 | 1030 | 0.308626559 | 0 | 0 | 0 | 0 | 4 mo | WILD |
| 1010419187 | 1 | SSp-ll1 | 1030 | 0.308626559 | 0.001069508 | 23 | 0.003465379 | 74.52372242 | 8 mo | HOM |
| 1013978366 | 1 | SSp-ll1 | 1030 | 0.308626559 | 0 | 0 | 0 | 0 | 2 mo | HOM |
| 1014416155 | 2 | SSp-ll1 | 1030 | 0.308626559 | 0 | 0 | 0 | 0 | 2 mo | HOM |
| 1013978065 | 2 | SSp-ll1 | 1030 | 0.308626559 | 4.42E-06 | 1 | 1.43E-05 | 3.240161844 | 2 mo | HOM |
| 1009934102 | 2 | SSp-ll1 | 1030 | 0.308626559 | 7.31E-05 | 3 | 0.000236954 | 9.720485533 | 4 mo | WILD |
| 1009322937 | 2 | SSp-ll1 | 1030 | 0.308626559 | 0 | 0 | 0 | 0 | 8 mo | WILD |
| 1011080637 | 2 | SSp-ll1 | 1030 | 0.308626559 | 0.000322725 | 11 | 0.001045681 | 35.64178029 | 8 mo | HOM |
| 1013609713 | 2 | SSp-ll1 | 1030 | 0.308626559 | 3.73E-06 | 1 | 1.21E-05 | 3.240161844 | 2 mo | WILD |
| 1010418842 | 1 | SSp-ll1 | 1030 | 0.308626559 | 0.000952759 | 22 | 0.003087092 | 71.28356058 | 8 mo | HOM |
| 1013288213 | 2 | SSp-ll1 | 1030 | 0.308626559 | 7.2E-05 | 4 | 0.000233237 | 12.96064738 | 4 mo | HOM |
| 1008976060 | 2 | SSp-ll1 | 1030 | 0.308626559 | 0.000134644 | 5 | 0.000436267 | 16.20080922 | 8 mo | WILD |
| 1013446243 | 1 | SSp-ll1 | 1030 | 0.308626559 | 0.000386808 | 5 | 0.001253321 | 16.20080922 | 4 mo | HOM |
| 1013288951 | 2 | SSp-ll1 | 1030 | 0.308626559 | 0.000128755 | 6 | 0.000417188 | 19.44097107 | 4 mo | HOM |
| 1013978665 | 1 | GPi | 1031 | 0.436380633 | 0 | 0 | 0 | 0 | 2 mo | HOM |
| 1013445609 | 1 | GPi | 1031 | 0.436380633 | 0 | 0 | 0 | 0 | 4 mo | HOM |
| 1013609015 | 1 | GPi | 1031 | 0.436380633 | 0 | 0 | 0 | 0 | 4 mo | HOM |
| 1013608106 | 2 | GPi | 1031 | 0.436380633 | 0 | 0 | 0 | 0 | 4 mo | HOM |
| 1011937543 | 1 | GPi | 1031 | 0.436380633 | 0.001085124 | 12 | 0.002486646 | 27.49892891 | 8 mo | HOM |
| 1010419538 | 2 | GPi | 1031 | 0.436380633 | 0 | 0 | 0 | 0 | 4 mo | WILD |
| 1010419187 | 1 | GPi | 1031 | 0.436380633 | 4.74E-06 | 1 | 1.09E-05 | 2.291577409 | 8 mo | HOM |
| 1013978366 | 1 | GPi | 1031 | 0.436380633 | 0 | 0 | 0 | 0 | 2 mo | HOM |
| 1014416155 | 2 | GPi | 1031 | 0.436380633 | 0 | 0 | 0 | 0 | 2 mo | HOM |
| 1013978065 | 2 | GPi | 1031 | 0.436380633 | 0 | 0 | 0 | 0 | 2 mo | HOM |
| 1009934102 | 2 | GPi | 1031 | 0.436380633 | 0 | 0 | 0 | 0 | 4 mo | WILD |
| 1009322937 | 2 | GPi | 1031 | 0.436380633 | 0 | 0 | 0 | 0 | 8 mo | WILD |
| 1011080637 | 2 | GPi | 1031 | 0.436380633 | 0 | 0 | 0 | 0 | 8 mo | HOM |
| 1013609713 | 2 | GPi | 1031 | 0.436380633 | 0 | 0 | 0 | 0 | 2 mo | WILD |
| 1010418842 | 1 | GPi | 1031 | 0.436380633 | 1.82E-05 | 3 | 4.18E-05 | 6.874732228 | 8 mo | HOM |
| 1013288213 | 2 | GPi | 1031 | 0.436380633 | 0 | 0 | 0 | 0 | 4 mo | HOM |
| 1008976060 | 2 | GPi | 1031 | 0.436380633 | 0 | 0 | 0 | 0 | 8 mo | WILD |
| 1013446243 | 1 | GPi | 1031 | 0.436380633 | 0 | 0 | 0 | 0 | 4 mo | HOM |
| 1013288951 | 2 | GPi | 1031 | 0.436380633 | 0 | 0 | 0 | 0 | 4 mo | HOM |
| 1013978665 | 1 | COPY | 1033 | 2.581908627 | 8.04E-05 | 9 | 3.11E-05 | 3.48579338 | 2 mo | HOM |

| | | | | | | | | | | |
|---|---|---|---|---|---|---|---|---|---|---|
| **1013445609** | 1 | COPY | 1033 | 2.581908627 | 4.81E-06 | 0 | 1.86E-06 | 0 | 4 mo | HOM |
| **1013609015** | 1 | COPY | 1033 | 2.581908627 | 0.000162728 | 16 | 6.3E-05 | 6.196966009 | 4 mo | HOM |
| **1013608106** | 2 | COPY | 1033 | 2.581908627 | 0 | 0 | 0 | 0 | 4 mo | HOM |
| **1011937543** | 1 | COPY | 1033 | 2.581908627 | 0.002965086 | 7 | 0.001148409 | 2.711172629 | 8 mo | HOM |
| **1010419538** | 2 | COPY | 1033 | 2.581908627 | 2.35E-05 | 4 | 9.12E-06 | 1.549241502 | 4 mo | WILD |
| **1010419187** | 1 | COPY | 1033 | 2.581908627 | 0.006680368 | 35 | 0.002587376 | 13.55586314 | 8 mo | HOM |
| **1013978366** | 1 | COPY | 1033 | 2.581908627 | 0.00051747 | 9 | 0.000200421 | 3.48579338 | 2 mo | HOM |
| **1014416155** | 2 | COPY | 1033 | 2.581908627 | 0 | 0 | 0 | 0 | 2 mo | HOM |
| **1013978065** | 2 | COPY | 1033 | 2.581908627 | 0 | 0 | 0 | 0 | 2 mo | HOM |
| **1009934102** | 2 | COPY | 1033 | 2.581908627 | 5.89E-05 | 3 | 2.28E-05 | 1.161931127 | 4 mo | WILD |
| **1009322937** | 2 | COPY | 1033 | 2.581908627 | 3.75E-05 | 4 | 1.45E-05 | 1.549241502 | 8 mo | WILD |
| **1011080637** | 2 | COPY | 1033 | 2.581908627 | 0.000253914 | 10 | 9.83E-05 | 3.873103755 | 8 mo | HOM |
| **1013609713** | 2 | COPY | 1033 | 2.510656042 | 2.56E-05 | 2 | 1.02E-05 | 0.79660454 | 2 mo | WILD |
| **1010418842** | 1 | COPY | 1033 | 2.581908627 | 0.003522291 | 52 | 0.00136422 | 20.14013953 | 8 mo | HOM |
| **1013288213** | 2 | COPY | 1033 | 2.581908627 | 5.51E-06 | 1 | 2.13E-06 | 0.387310376 | 4 mo | HOM |
| **1008976060** | 2 | COPY | 1033 | 2.581908627 | 0.000334762 | 18 | 0.000129657 | 6.97158676 | 8 mo | WILD |
| **1013446243** | 1 | COPY | 1033 | 2.581908627 | 5.39E-06 | 2 | 2.09E-06 | 0.774620751 | 4 mo | HOM |
| **1013288951** | 2 | COPY | 1033 | 2.581908627 | 0.001059472 | 9 | 0.000410344 | 3.48579338 | 4 mo | HOM |
| **1013978665** | 1 | SSs4 | 1035 | 1.202441498 | 0 | 0 | 0 | 0 | 2 mo | HOM |
| **1013445609** | 1 | SSs4 | 1035 | 1.202441498 | 7.86E-05 | 5 | 6.53E-05 | 4.158206457 | 4 mo | HOM |
| **1013609015** | 1 | SSs4 | 1035 | 1.202441498 | 0.000304083 | 9 | 0.000252888 | 7.484771623 | 4 mo | HOM |
| **1013608106** | 2 | SSs4 | 1035 | 1.202441498 | 0 | 0 | 0 | 0 | 4 mo | HOM |
| **1011937543** | 1 | SSs4 | 1035 | 1.202441498 | 0.001070686 | 35 | 0.000890426 | 29.1074452 | 8 mo | HOM |
| **1010419538** | 2 | SSs4 | 1035 | 1.202441498 | 1.28E-05 | 1 | 1.06E-05 | 0.831641291 | 4 mo | WILD |
| **1010419187** | 1 | SSs4 | 1035 | 1.202441498 | 0.001377079 | 51 | 0.001145235 | 42.41370587 | 8 mo | HOM |
| **1013978366** | 1 | SSs4 | 1035 | 1.202441498 | 1.88E-05 | 2 | 1.56E-05 | 1.663282583 | 2 mo | HOM |
| **1014416155** | 2 | SSs4 | 1035 | 1.202441498 | 0 | 0 | 0 | 0 | 2 mo | HOM |
| **1013978065** | 2 | SSs4 | 1035 | 1.202441498 | 4.44E-05 | 1 | 3.69E-05 | 0.831641291 | 2 mo | HOM |
| **1009934102** | 2 | SSs4 | 1035 | 1.202441498 | 0 | 0 | 0 | 0 | 4 mo | WILD |
| **1009322937** | 2 | SSs4 | 1035 | 1.202441498 | 0 | 0 | 0 | 0 | 8 mo | WILD |
| **1011080637** | 2 | SSs4 | 1035 | 1.202441498 | 0.001032842 | 50 | 0.000858954 | 41.58206457 | 8 mo | HOM |
| **1013609713** | 2 | SSs4 | 1035 | 1.202441498 | 0 | 0 | 0 | 0 | 2 mo | WILD |
| **1010418842** | 1 | SSs4 | 1035 | 1.202441498 | 0.001002251 | 40 | 0.000833513 | 33.26565166 | 8 mo | HOM |
| **1013288213** | 2 | SSs4 | 1035 | 1.202441498 | 5.54E-05 | 5 | 4.61E-05 | 4.158206457 | 4 mo | HOM |
| **1008976060** | 2 | SSs4 | 1035 | 1.202441498 | 1.64E-05 | 1 | 1.36E-05 | 0.831641291 | 8 mo | WILD |
| **1013446243** | 1 | SSs4 | 1035 | 1.202441498 | 0.000100033 | 5 | 8.32E-05 | 4.158206457 | 4 mo | HOM |
| **1013288951** | 2 | SSs4 | 1035 | 1.202441498 | 0.000219635 | 9 | 0.000182658 | 7.484771623 | 4 mo | HOM |
| **1013978665** | 1 | POST | 1037 | 1.117091494 | 0 | 0 | 0 | 0 | 2 mo | HOM |
| **1013445609** | 1 | POST | 1037 | 1.117091494 | 2.78E-05 | 2 | 2.49E-05 | 1.790363645 | 4 mo | HOM |
| **1013609015** | 1 | POST | 1037 | 1.117091494 | 8.39E-05 | 4 | 7.51E-05 | 3.580727291 | 4 mo | HOM |
| **1013608106** | 2 | POST | 1037 | 1.117091494 | 0 | 0 | 0 | 0 | 4 mo | HOM |
| **1011937543** | 1 | POST | 1037 | 1.117091494 | 0.000885269 | 10 | 0.000792477 | 8.951818226 | 8 mo | HOM |
| **1010419538** | 2 | POST | 1037 | 1.117091494 | 7.13E-06 | 0 | 6.38E-06 | 0 | 4 mo | WILD |
| **1010419187** | 1 | POST | 1037 | 1.117091494 | 0.002177234 | 42 | 0.001949021 | 37.59763655 | 8 mo | HOM |
| **1013978366** | 1 | POST | 1037 | 1.117091494 | 3.45E-05 | 3 | 3.08E-05 | 2.685545468 | 2 mo | HOM |
| **1014416155** | 2 | POST | 1037 | 1.117091494 | 0 | 0 | 0 | 0 | 2 mo | HOM |
| **1013978065** | 2 | POST | 1037 | 1.117091494 | 0 | 0 | 0 | 0 | 2 mo | HOM |
| **1009934102** | 2 | POST | 1037 | 1.117091494 | 0.000472054 | 9 | 0.000422575 | 8.056636404 | 4 mo | WILD |
| **1009322937** | 2 | POST | 1037 | 1.117091494 | 0 | 0 | 0 | 0 | 8 mo | WILD |
| **1011080637** | 2 | POST | 1037 | 1.117091494 | 0.001228546 | 28 | 0.001099772 | 25.06509103 | 8 mo | HOM |
| **1013609713** | 2 | POST | 1037 | 1.117091494 | 0.000109241 | 1 | 9.78E-05 | 0.895181823 | 2 mo | WILD |
| **1010418842** | 1 | POST | 1037 | 1.117091494 | 0.001464145 | 28 | 0.001310676 | 25.06509103 | 8 mo | HOM |
| **1013288213** | 2 | POST | 1037 | 1.117091494 | 3.45E-05 | 0 | 3.09E-05 | 0 | 4 mo | HOM |
| **1008976060** | 2 | POST | 1037 | 1.117091494 | 0.000252312 | 12 | 0.000225865 | 10.74218187 | 8 mo | WILD |
| **1013446243** | 1 | POST | 1037 | 1.117091494 | 4.51E-05 | 3 | 4.04E-05 | 2.685545468 | 4 mo | HOM |
| **1013288951** | 2 | POST | 1037 | 1.117091494 | 4.17E-05 | 6 | 3.73E-05 | 5.371090936 | 4 mo | HOM |
| **1013978665** | 1 | SSp-bfd6a | 1038 | 1.335216915 | 0 | 0 | 0 | 0 | 2 mo | HOM |
| **1013445609** | 1 | SSp-bfd6a | 1038 | 1.335216915 | 0.000105507 | 8 | 7.9E-05 | 5.991535839 | 4 mo | HOM |
| **1013609015** | 1 | SSp-bfd6a | 1038 | 1.335216915 | 0.000475447 | 14 | 0.000356082 | 10.48518772 | 4 mo | HOM |

| | | | | | | | | | | |
|---|---|---|---|---|---|---|---|---|---|---|
| **1013608106** | 2 | SSp-bfd6a | 1038 | 1.335216915 | 0 | 0 | 0 | 0 | 4 mo | HOM |
| **1011937543** | 1 | SSp-bfd6a | 1038 | 1.335216915 | 0.00204429 | 33 | 0.001531055 | 24.71508534 | 8 mo | HOM |
| **1010419538** | 2 | SSp-bfd6a | 1038 | 1.335216915 | 0 | 0 | 0 | 0 | 4 mo | WILD |
| **1010419187** | 1 | SSp-bfd6a | 1038 | 1.335216915 | 0.00229384 | 80 | 0.001717953 | 59.91535839 | 8 mo | HOM |
| **1013978366** | 1 | SSp-bfd6a | 1038 | 1.335216915 | 0 | 0 | 0 | 0 | 2 mo | HOM |
| **1014416155** | 2 | SSp-bfd6a | 1038 | 1.335216915 | 0 | 0 | 0 | 0 | 2 mo | HOM |
| **1013978065** | 2 | SSp-bfd6a | 1038 | 1.335216915 | 0 | 0 | 0 | 0 | 2 mo | HOM |
| **1009934102** | 2 | SSp-bfd6a | 1038 | 1.335216915 | 0 | 0 | 0 | 0 | 4 mo | WILD |
| **1009322937** | 2 | SSp-bfd6a | 1038 | 1.335216915 | 5.11E-06 | 1 | 3.83E-06 | 0.74894198 | 8 mo | WILD |
| **1011080637** | 2 | SSp-bfd6a | 1038 | 1.335216915 | 0.001722981 | 68 | 0.001290413 | 50.92805463 | 8 mo | HOM |
| **1013609713** | 2 | SSp-bfd6a | 1038 | 1.335216915 | 0 | 0 | 0 | 0 | 2 mo | WILD |
| **1010418842** | 1 | SSp-bfd6a | 1038 | 1.335216915 | 0.00186511 | 73 | 0.001396859 | 54.67276453 | 8 mo | HOM |
| **1013288213** | 2 | SSp-bfd6a | 1038 | 1.335216915 | 6.61E-05 | 8 | 4.95E-05 | 5.991535839 | 4 mo | HOM |
| **1008976060** | 2 | SSp-bfd6a | 1038 | 1.335216915 | 1.01E-05 | 2 | 7.58E-06 | 1.49788396 | 8 mo | WILD |
| **1013446243** | 1 | SSp-bfd6a | 1038 | 1.335216915 | 0.000253649 | 17 | 0.000189969 | 12.73201366 | 4 mo | HOM |
| **1013288951** | 2 | SSp-bfd6a | 1038 | 1.335216915 | 0.000357103 | 20 | 0.00026745 | 14.9788396 | 4 mo | HOM |
| **1013978665** | 1 | GR | 1039 | 0.084091085 | 0 | 0 | 0 | 0 | 2 mo | HOM |
| **1013445609** | 1 | GR | 1039 | 0.084091085 | 0 | 0 | 0 | 0 | 4 mo | HOM |
| **1013609015** | 1 | GR | 1039 | 0.084091085 | 0 | 0 | 0 | 0 | 4 mo | HOM |
| **1013608106** | 2 | GR | 1039 | 0.084091085 | 0 | 0 | 0 | 0 | 4 mo | HOM |
| **1011937543** | 1 | GR | 1039 | 0.084091085 | 0 | 0 | 0 | 0 | 8 mo | HOM |
| **1010419538** | 2 | GR | 1039 | 0.084091085 | 1.06E-06 | 0 | 1.27E-05 | 0 | 4 mo | WILD |
| **1010419187** | 1 | GR | 1039 | 0.084091085 | 0.002231721 | 1 | 0.026539326 | 11.89186696 | 8 mo | HOM |
| **1013978366** | 1 | GR | 1039 | 0.084091085 | 3.66E-06 | 1 | 4.35E-05 | 11.89186696 | 2 mo | HOM |
| **1014416155** | 2 | GR | 1039 | 0.084091085 | 0 | 0 | 0 | 0 | 2 mo | HOM |
| **1013978065** | 2 | GR | 1039 | 0.084091085 | 2.33E-06 | 0 | 2.77E-05 | 0 | 2 mo | HOM |
| **1009934102** | 2 | GR | 1039 | 0.084091085 | 0 | 0 | 0 | 0 | 4 mo | WILD |
| **1009322937** | 2 | GR | 1039 | 0.084091085 | 1.96E-06 | 0 | 2.33E-05 | 0 | 8 mo | WILD |
| **1011080637** | 2 | GR | 1039 | 0.084091085 | 0 | 0 | 0 | 0 | 8 mo | HOM |
| **1013609713** | 2 | GR | 1039 | 0.084091085 | 0 | 0 | 0 | 0 | 2 mo | WILD |
| **1010418842** | 1 | GR | 1039 | 0.084091085 | 8.39E-06 | 1 | 9.98E-05 | 11.89186696 | 8 mo | HOM |
| **1013288213** | 2 | GR | 1039 | 0.084091085 | 0 | 0 | 0 | 0 | 4 mo | HOM |
| **1008976060** | 2 | GR | 1039 | 0.084091085 | 0 | 0 | 0 | 0 | 8 mo | WILD |
| **1013446243** | 1 | GR | 1039 | 0.084091085 | 0.000223532 | 0 | 0.002658217 | 0 | 4 mo | HOM |
| **1013288951** | 2 | GR | 1039 | 0.084091085 | 0.001175484 | 3 | 0.013978694 | 35.67560088 | 4 mo | HOM |
| **1013978665** | 1 | PFL | 1041 | 5.902303232 | 0.000165352 | 13 | 2.8E-05 | 2.202530011 | 2 mo | HOM |
| **1013445609** | 1 | PFL | 1041 | 5.902303232 | 2.04E-05 | 1 | 3.45E-06 | 0.169425385 | 4 mo | HOM |
| **1013609015** | 1 | PFL | 1041 | 5.902303232 | 0.002706559 | 33 | 0.00045856 | 5.591037719 | 4 mo | HOM |
| **1013608106** | 2 | PFL | 1041 | 5.902303232 | 0 | 0 | 0 | 0 | 4 mo | HOM |
| **1011937543** | 1 | PFL | 1041 | 5.902303232 | 0.01873516 | 27 | 0.003174212 | 4.574485407 | 8 mo | HOM |
| **1010419538** | 2 | PFL | 1041 | 5.902303232 | 4.79E-05 | 3 | 8.12E-06 | 0.508276156 | 4 mo | WILD |
| **1010419187** | 1 | PFL | 1041 | 5.902303232 | 0.000619784 | 5 | 0.000105007 | 0.847126927 | 8 mo | HOM |
| **1013978366** | 1 | PFL | 1041 | 5.902303232 | 0.002317241 | 49 | 0.000392599 | 8.301843886 | 2 mo | HOM |
| **1014416155** | 2 | PFL | 1041 | 5.902303232 | 0 | 0 | 0 | 0 | 2 mo | HOM |
| **1013978065** | 2 | PFL | 1041 | 5.902303232 | 1.53E-05 | 0 | 2.6E-06 | 0 | 2 mo | HOM |
| **1009934102** | 2 | PFL | 1041 | 5.902303232 | 2.42E-05 | 1 | 4.1E-06 | 0.169425385 | 4 mo | WILD |
| **1009322937** | 2 | PFL | 1041 | 5.902303232 | 0.000148745 | 7 | 2.52E-05 | 1.185977698 | 8 mo | WILD |
| **1011080637** | 2 | PFL | 1041 | 5.902303232 | 0.001973294 | 25 | 0.000334326 | 4.235634636 | 8 mo | HOM |
| **1013609713** | 2 | PFL | 1041 | 5.902303232 | 8.84E-06 | 1 | 1.5E-06 | 0.169425385 | 2 mo | WILD |
| **1010418842** | 1 | PFL | 1041 | 5.902303232 | 0.00201344 | 58 | 0.000341128 | 9.826672355 | 8 mo | HOM |
| **1013288213** | 2 | PFL | 1041 | 5.902303232 | 5.29E-05 | 5 | 8.96E-06 | 0.847126927 | 4 mo | HOM |
| **1008976060** | 2 | PFL | 1041 | 5.902303232 | 9.53E-05 | 9 | 1.61E-05 | 1.524828469 | 8 mo | WILD |
| **1013446243** | 1 | PFL | 1041 | 5.902303232 | 0.000381648 | 15 | 6.47E-05 | 2.541380781 | 4 mo | HOM |
| **1013288951** | 2 | PFL | 1041 | 5.902303232 | 0.002786111 | 28 | 0.000472038 | 4.743910792 | 4 mo | HOM |
| **1013978665** | 1 | tspc | 1043 | 0.518796594 | 0 | 0 | 0 | 0 | 2 mo | HOM |
| **1013445609** | 1 | tspc | 1043 | 0.518796594 | 0 | 0 | 0 | 0 | 4 mo | HOM |
| **1013609015** | 1 | tspc | 1043 | 0.518796594 | 0 | 0 | 0 | 0 | 4 mo | HOM |
| **1013608106** | 2 | tspc | 1043 | 0.513309182 | 0 | 0 | 0 | 0 | 4 mo | HOM |
| **1011937543** | 1 | tspc | 1043 | 0.518796594 | 9.78E-05 | 3 | 0.000188521 | 5.782613133 | 8 mo | HOM |

| | | | | | | | | | | |
|---|---|---|---|---|---|---|---|---|---|---|
| **1010419538** | 2 | tspc | 1043 | 0.518796594 | 0 | 0 | 0 | 0 | 4 mo | WILD |
| **1010419187** | 1 | tspc | 1043 | 0.518796594 | 0 | 0 | 0 | 0 | 8 mo | HOM |
| **1013978366** | 1 | tspc | 1043 | 0.518796594 | 0 | 0 | 0 | 0 | 2 mo | HOM |
| **1014416155** | 2 | tspc | 1043 | 0.518796594 | 0 | 0 | 0 | 0 | 2 mo | HOM |
| **1013978065** | 2 | tspc | 1043 | 0.518796594 | 0 | 0 | 0 | 0 | 2 mo | HOM |
| **1009934102** | 2 | tspc | 1043 | 0.518796594 | 0 | 0 | 0 | 0 | 4 mo | WILD |
| **1009322937** | 2 | tspc | 1043 | 0.518796594 | 0 | 0 | 0 | 0 | 8 mo | WILD |
| **1011080637** | 2 | tspc | 1043 | 0.518796594 | 0 | 0 | 0 | 0 | 8 mo | HOM |
| **1013609713** | 2 | tspc | 1043 | 0.518796594 | 0 | 0 | 0 | 0 | 2 mo | WILD |
| **1010418842** | 1 | tspc | 1043 | 0.518796594 | 1.43E-05 | 1 | 2.76E-05 | 1.927537711 | 8 mo | HOM |
| **1013288213** | 2 | tspc | 1043 | 0.518796594 | 0 | 0 | 0 | 0 | 4 mo | HOM |
| **1008976060** | 2 | tspc | 1043 | 0.518796594 | 1.47E-05 | 1 | 2.83E-05 | 1.927537711 | 8 mo | WILD |
| **1013446243** | 1 | tspc | 1043 | 0.518796594 | 0 | 0 | 0 | 0 | 4 mo | HOM |
| **1013288951** | 2 | tspc | 1043 | 0.518796594 | 0 | 0 | 0 | 0 | 4 mo | HOM |
| **1013978665** | 1 | PP | 1044 | 0.062992077 | 0 | 0 | 0 | 0 | 2 mo | HOM |
| **1013445609** | 1 | PP | 1044 | 0.062992077 | 0 | 0 | 0 | 0 | 4 mo | HOM |
| **1013609015** | 1 | PP | 1044 | 0.062992077 | 0 | 0 | 0 | 0 | 4 mo | HOM |
| **1013608106** | 2 | PP | 1044 | 0.062992077 | 0 | 0 | 0 | 0 | 4 mo | HOM |
| **1011937543** | 1 | PP | 1044 | 0.062992077 | 0.00010781 | 2 | 0.00171148 | 31.75002449 | 8 mo | HOM |
| **1010419538** | 2 | PP | 1044 | 0.062992077 | 0 | 0 | 0 | 0 | 4 mo | WILD |
| **1010419187** | 1 | PP | 1044 | 0.062992077 | 0 | 0 | 0 | 0 | 8 mo | HOM |
| **1013978366** | 1 | PP | 1044 | 0.062992077 | 0 | 0 | 0 | 0 | 2 mo | HOM |
| **1014416155** | 2 | PP | 1044 | 0.062992077 | 0 | 0 | 0 | 0 | 2 mo | HOM |
| **1013978065** | 2 | PP | 1044 | 0.062992077 | 0 | 0 | 0 | 0 | 2 mo | HOM |
| **1009934102** | 2 | PP | 1044 | 0.062992077 | 0 | 0 | 0 | 0 | 4 mo | WILD |
| **1009322937** | 2 | PP | 1044 | 0.062992077 | 0 | 0 | 0 | 0 | 8 mo | WILD |
| **1011080637** | 2 | PP | 1044 | 0.062992077 | 0 | 0 | 0 | 0 | 8 mo | HOM |
| **1013609713** | 2 | PP | 1044 | 0.062992077 | 0 | 0 | 0 | 0 | 2 mo | WILD |
| **1010418842** | 1 | PP | 1044 | 0.062992077 | 2.91E-07 | 0 | 4.62E-06 | 0 | 8 mo | HOM |
| **1013288213** | 2 | PP | 1044 | 0.062992077 | 0 | 0 | 0 | 0 | 4 mo | HOM |
| **1008976060** | 2 | PP | 1044 | 0.062992077 | 2.82E-06 | 0 | 4.48E-05 | 0 | 8 mo | WILD |
| **1013446243** | 1 | PP | 1044 | 0.062992077 | 0 | 0 | 0 | 0 | 4 mo | HOM |
| **1013288951** | 2 | PP | 1044 | 0.062992077 | 0 | 0 | 0 | 0 | 4 mo | HOM |
| **1013978665** | 1 | ECT6b | 1045 | 0.061145914 | 0 | 0 | 0 | 0 | 2 mo | HOM |
| **1013445609** | 1 | ECT6b | 1045 | 0.061145914 | 3.85E-09 | 0 | 6.29E-08 | 0 | 4 mo | HOM |
| **1013609015** | 1 | ECT6b | 1045 | 0.061145914 | 0 | 0 | 0 | 0 | 4 mo | HOM |
| **1013608106** | 2 | ECT6b | 1045 | 0.061145914 | 0 | 0 | 0 | 0 | 4 mo | HOM |
| **1011937543** | 1 | ECT6b | 1045 | 0.061145914 | 5.97E-05 | 1 | 0.000976863 | 16.35432249 | 8 mo | HOM |
| **1010419538** | 2 | ECT6b | 1045 | 0.061145914 | 0 | 0 | 0 | 0 | 4 mo | WILD |
| **1010419187** | 1 | ECT6b | 1045 | 0.061145914 | 0.000104229 | 4 | 0.001704589 | 65.41728997 | 8 mo | HOM |
| **1013978366** | 1 | ECT6b | 1045 | 0.061145914 | 0 | 0 | 0 | 0 | 2 mo | HOM |
| **1014416155** | 2 | ECT6b | 1045 | 0.061145914 | 0 | 0 | 0 | 0 | 2 mo | HOM |
| **1013978065** | 2 | ECT6b | 1045 | 0.061145914 | 0 | 0 | 0 | 0 | 2 mo | HOM |
| **1009934102** | 2 | ECT6b | 1045 | 0.061145914 | 0 | 0 | 0 | 0 | 4 mo | WILD |
| **1009322937** | 2 | ECT6b | 1045 | 0.061145914 | 0 | 0 | 0 | 0 | 8 mo | WILD |
| **1011080637** | 2 | ECT6b | 1045 | 0.061145914 | 3.65E-05 | 3 | 0.000597144 | 49.06296748 | 8 mo | HOM |
| **1013609713** | 2 | ECT6b | 1045 | 0.061145914 | 0 | 0 | 0 | 0 | 2 mo | WILD |
| **1010418842** | 1 | ECT6b | 1045 | 0.061145914 | 0.000189043 | 8 | 0.003091666 | 130.8345799 | 8 mo | HOM |
| **1013288213** | 2 | ECT6b | 1045 | 0.061145914 | 0 | 0 | 0 | 0 | 4 mo | HOM |
| **1008976060** | 2 | ECT6b | 1045 | 0.061145914 | 0 | 0 | 0 | 0 | 8 mo | WILD |
| **1013446243** | 1 | ECT6b | 1045 | 0.061145914 | 1.27E-05 | 1 | 0.00020712 | 16.35432249 | 4 mo | HOM |
| **1013288951** | 2 | ECT6b | 1045 | 0.061145914 | 0 | 0 | 0 | 0 | 4 mo | HOM |
| **1013978665** | 1 | VISam6a | 1046 | 0.142809794 | 0 | 0 | 0 | 0 | 2 mo | HOM |
| **1013445609** | 1 | VISam6a | 1046 | 0.142809794 | 1.94E-05 | 2 | 0.000135866 | 14.00464169 | 4 mo | HOM |
| **1013609015** | 1 | VISam6a | 1046 | 0.142809794 | 3.4E-05 | 1 | 0.000238163 | 7.002320844 | 4 mo | HOM |
| **1013608106** | 2 | VISam6a | 1046 | 0.142809794 | 0 | 0 | 0 | 0 | 4 mo | HOM |
| **1011937543** | 1 | VISam6a | 1046 | 0.142809794 | 0.000200723 | 3 | 0.001405529 | 21.00696253 | 8 mo | HOM |
| **1010419538** | 2 | VISam6a | 1046 | 0.142809794 | 0 | 0 | 0 | 0 | 4 mo | WILD |
| **1010419187** | 1 | VISam6a | 1046 | 0.142809794 | 0.000218749 | 9 | 0.001531751 | 63.02088759 | 8 mo | HOM |

| | | | | | | | | | | |
|---|---|---|---|---|---|---|---|---|---|---|
| **1013978366** | 1 | VISam6a | 1046 | 0.142809794 | 0 | 0 | 0 | 0 | 2 mo | HOM |
| **1014416155** | 2 | VISam6a | 1046 | 0.142809794 | 0 | 0 | 0 | 0 | 2 mo | HOM |
| **1013978065** | 2 | VISam6a | 1046 | 0.142809794 | 0 | 0 | 0 | 0 | 2 mo | HOM |
| **1009934102** | 2 | VISam6a | 1046 | 0.142809794 | 0 | 0 | 0 | 0 | 4 mo | WILD |
| **1009322937** | 2 | VISam6a | 1046 | 0.142809794 | 0 | 0 | 0 | 0 | 8 mo | WILD |
| **1011080637** | 2 | VISam6a | 1046 | 0.142809794 | 8.91E-05 | 7 | 0.000624024 | 49.01624591 | 8 mo | HOM |
| **1013609713** | 2 | VISam6a | 1046 | 0.142809794 | 0 | 0 | 0 | 0 | 2 mo | WILD |
| **1010418842** | 1 | VISam6a | 1046 | 0.142809794 | 8.2E-05 | 5 | 0.000573871 | 35.01160422 | 8 mo | HOM |
| **1013288213** | 2 | VISam6a | 1046 | 0.142809794 | 2.55E-05 | 3 | 0.000178504 | 21.00696253 | 4 mo | HOM |
| **1008976060** | 2 | VISam6a | 1046 | 0.142809794 | 0 | 0 | 0 | 0 | 8 mo | WILD |
| **1013446243** | 1 | VISam6a | 1046 | 0.142809794 | 1.78E-05 | 2 | 0.000124619 | 14.00464169 | 4 mo | HOM |
| **1013288951** | 2 | VISam6a | 1046 | 0.142809794 | 2.68E-05 | 2 | 0.000187325 | 14.00464169 | 4 mo | HOM |
| **1013978665** | 1 | SSp-bfd4 | 1047 | 1.320340515 | 0 | 0 | 0 | 0 | 2 mo | HOM |
| **1013445609** | 1 | SSp-bfd4 | 1047 | 1.320340515 | 7.15E-05 | 7 | 5.42E-05 | 5.301662654 | 4 mo | HOM |
| **1013609015** | 1 | SSp-bfd4 | 1047 | 1.320340515 | 0.000347227 | 20 | 0.000262983 | 15.14760758 | 4 mo | HOM |
| **1013608106** | 2 | SSp-bfd4 | 1047 | 1.320340515 | 0 | 0 | 0 | 0 | 4 mo | HOM |
| **1011937543** | 1 | SSp-bfd4 | 1047 | 1.320340515 | 0.003055443 | 47 | 0.002314133 | 35.59687782 | 8 mo | HOM |
| **1010419538** | 2 | SSp-bfd4 | 1047 | 1.320340515 | 0 | 0 | 0 | 0 | 4 mo | WILD |
| **1010419187** | 1 | SSp-bfd4 | 1047 | 1.320340515 | 0.002702284 | 71 | 0.002046657 | 53.77400692 | 8 mo | HOM |
| **1013978366** | 1 | SSp-bfd4 | 1047 | 1.320340515 | 8.74E-06 | 1 | 6.62E-06 | 0.757380379 | 2 mo | HOM |
| **1014416155** | 2 | SSp-bfd4 | 1047 | 1.320340515 | 0 | 0 | 0 | 0 | 2 mo | HOM |
| **1013978065** | 2 | SSp-bfd4 | 1047 | 1.320340515 | 0 | 0 | 0 | 0 | 2 mo | HOM |
| **1009934102** | 2 | SSp-bfd4 | 1047 | 1.320340515 | 0 | 0 | 0 | 0 | 4 mo | WILD |
| **1009322937** | 2 | SSp-bfd4 | 1047 | 1.320340515 | 6.36E-06 | 1 | 4.81E-06 | 0.757380379 | 8 mo | WILD |
| **1011080637** | 2 | SSp-bfd4 | 1047 | 1.320340515 | 0.002104551 | 63 | 0.001593945 | 47.71496389 | 8 mo | HOM |
| **1013609713** | 2 | SSp-bfd4 | 1047 | 1.320340515 | 0 | 0 | 0 | 0 | 2 mo | WILD |
| **1010418842** | 1 | SSp-bfd4 | 1047 | 1.320340515 | 0.002224849 | 67 | 0.001685057 | 50.7444854 | 8 mo | HOM |
| **1013288213** | 2 | SSp-bfd4 | 1047 | 1.320340515 | 0.000122174 | 13 | 9.25E-05 | 9.845944929 | 4 mo | HOM |
| **1008976060** | 2 | SSp-bfd4 | 1047 | 1.320340515 | 2.11E-05 | 3 | 1.59E-05 | 2.272141137 | 8 mo | WILD |
| **1013446243** | 1 | SSp-bfd4 | 1047 | 1.320340515 | 0.000201391 | 9 | 0.000152529 | 6.816423412 | 4 mo | HOM |
| **1013288951** | 2 | SSp-bfd4 | 1047 | 1.320340515 | 0.000311357 | 12 | 0.000235816 | 9.08856455 | 4 mo | HOM |
| **1013978665** | 1 | GRN | 1048 | 2.703057011 | 0 | 0 | 0 | 0 | 2 mo | HOM |
| **1013445609** | 1 | GRN | 1048 | 2.703057011 | 0 | 0 | 0 | 0 | 4 mo | HOM |
| **1013609015** | 1 | GRN | 1048 | 2.703057011 | 0 | 0 | 0 | 0 | 4 mo | HOM |
| **1013608106** | 2 | GRN | 1048 | 2.688563674 | 0 | 0 | 0 | 0 | 4 mo | HOM |
| **1011937543** | 1 | GRN | 1048 | 2.703057011 | 6.17E-05 | 1 | 2.28E-05 | 0.369951502 | 8 mo | HOM |
| **1010419538** | 2 | GRN | 1048 | 2.703057011 | 0 | 0 | 0 | 0 | 4 mo | WILD |
| **1010419187** | 1 | GRN | 1048 | 2.703057011 | 0 | 0 | 0 | 0 | 8 mo | HOM |
| **1013978366** | 1 | GRN | 1048 | 2.703057011 | 0 | 0 | 0 | 0 | 2 mo | HOM |
| **1014416155** | 2 | GRN | 1048 | 2.703057011 | 0 | 0 | 0 | 0 | 2 mo | HOM |
| **1013978065** | 2 | GRN | 1048 | 2.703057011 | 0 | 0 | 0 | 0 | 2 mo | HOM |
| **1009934102** | 2 | GRN | 1048 | 2.703057011 | 0 | 0 | 0 | 0 | 4 mo | WILD |
| **1009322937** | 2 | GRN | 1048 | 2.703057011 | 8.85E-06 | 1 | 3.27E-06 | 0.369951502 | 8 mo | WILD |
| **1011080637** | 2 | GRN | 1048 | 2.703057011 | 1.34E-05 | 2 | 4.96E-06 | 0.739903003 | 8 mo | HOM |
| **1013609713** | 2 | GRN | 1048 | 2.703057011 | 0 | 0 | 0 | 0 | 2 mo | WILD |
| **1010418842** | 1 | GRN | 1048 | 2.703057011 | 0.000426313 | 34 | 0.000157715 | 12.57835105 | 8 mo | HOM |
| **1013288213** | 2 | GRN | 1048 | 2.703057011 | 5.38E-06 | 1 | 1.99E-06 | 0.369951502 | 4 mo | HOM |
| **1008976060** | 2 | GRN | 1048 | 2.703057011 | 0.000488343 | 34 | 0.000180663 | 12.57835105 | 8 mo | WILD |
| **1013446243** | 1 | GRN | 1048 | 2.703057011 | 0 | 0 | 0 | 0 | 4 mo | HOM |
| **1013288951** | 2 | GRN | 1048 | 2.703057011 | 0 | 0 | 0 | 0 | 4 mo | HOM |
| **1013978665** | 1 | FL | 1049 | 1.377684941 | 0.000163758 | 14 | 0.000118865 | 10.16197505 | 2 mo | HOM |
| **1013445609** | 1 | FL | 1049 | 1.377684941 | 0 | 0 | 0 | 0 | 4 mo | HOM |
| **1013609015** | 1 | FL | 1049 | 1.377684941 | 4.17E-05 | 4 | 3.03E-05 | 2.903421444 | 4 mo | HOM |
| **1013608106** | 2 | FL | 1049 | 1.377684941 | 0 | 0 | 0 | 0 | 4 mo | HOM |
| **1011937543** | 1 | FL | 1049 | 1.377684941 | 0.000324994 | 4 | 0.000235899 | 2.903421444 | 8 mo | HOM |
| **1010419538** | 2 | FL | 1049 | 1.377684941 | 2.8E-05 | 1 | 2.04E-05 | 0.725855361 | 4 mo | WILD |
| **1010419187** | 1 | FL | 1049 | 1.377684941 | 0.00034173 | 11 | 0.000248046 | 7.984408971 | 8 mo | HOM |
| **1013978366** | 1 | FL | 1049 | 1.377684941 | 2.38E-05 | 2 | 1.72E-05 | 1.451710722 | 2 mo | HOM |
| **1014416155** | 2 | FL | 1049 | 1.377684941 | 0 | 0 | 0 | 0 | 2 mo | HOM |

| | | | | | | | | | | |
|---|---|---|---|---|---|---|---|---|---|---|
| **1013978065** | 2 | FL | 1049 | 1.377684941 | 0.000153156 | 1 | 0.000111169 | 0.725855361 | 2 mo | HOM |
| **1009934102** | 2 | FL | 1049 | 1.377684941 | 2.81E-06 | 0 | 2.04E-06 | 0 | 4 mo | WILD |
| **1009322937** | 2 | FL | 1049 | 1.377684941 | 3.29E-05 | 2 | 2.39E-05 | 1.451710722 | 8 mo | WILD |
| **1011080637** | 2 | FL | 1049 | 1.377684941 | 0.00010258 | 6 | 7.45E-05 | 4.355132166 | 8 mo | HOM |
| **1013609713** | 2 | FL | 1049 | 1.377684941 | 3.05E-06 | 1 | 2.21E-06 | 0.725855361 | 2 mo | WILD |
| **1010418842** | 1 | FL | 1049 | 1.377684941 | 0.000402912 | 19 | 0.000292456 | 13.79125186 | 8 mo | HOM |
| **1013288213** | 2 | FL | 1049 | 1.377684941 | 5.92E-06 | 0 | 4.3E-06 | 0 | 4 mo | HOM |
| **1008976060** | 2 | FL | 1049 | 1.377684941 | 2.69E-05 | 2 | 1.95E-05 | 1.451710722 | 8 mo | WILD |
| **1013446243** | 1 | FL | 1049 | 1.377684941 | 5.62E-05 | 2 | 4.08E-05 | 1.451710722 | 4 mo | HOM |
| **1013288951** | 2 | FL | 1049 | 1.377684941 | 0.000169253 | 2 | 0.000122853 | 1.451710722 | 4 mo | HOM |
| **1013978665** | 1 | tspd | 1051 | 4.12E-06 | 0 | 0 | 0 | 0 | 2 mo | HOM |
| **1013445609** | 1 | tspd | 1051 | 4.12E-06 | 0 | 0 | 0 | 0 | 4 mo | HOM |
| **1013609015** | 1 | tspd | 1051 | 4.12E-06 | 0 | 0 | 0 | 0 | 4 mo | HOM |
| **1013608106** | 2 | tspd | 1051 | 4.12E-06 | 0 | 0 | 0 | 0 | 4 mo | HOM |
| **1011937543** | 1 | tspd | 1051 | 4.12E-06 | 0 | 0 | 0 | 0 | 8 mo | HOM |
| **1010419538** | 2 | tspd | 1051 | 4.12E-06 | 0 | 0 | 0 | 0 | 4 mo | WILD |
| **1010419187** | 1 | tspd | 1051 | 4.12E-06 | 0 | 0 | 0 | 0 | 8 mo | HOM |
| **1013978366** | 1 | tspd | 1051 | 4.12E-06 | 0 | 0 | 0 | 0 | 2 mo | HOM |
| **1014416155** | 2 | tspd | 1051 | 4.12E-06 | 0 | 0 | 0 | 0 | 2 mo | HOM |
| **1013978065** | 2 | tspd | 1051 | 4.12E-06 | 0 | 0 | 0 | 0 | 2 mo | HOM |
| **1009934102** | 2 | tspd | 1051 | 4.12E-06 | 0 | 0 | 0 | 0 | 4 mo | WILD |
| **1009322937** | 2 | tspd | 1051 | 4.12E-06 | 0 | 0 | 0 | 0 | 8 mo | WILD |
| **1011080637** | 2 | tspd | 1051 | 4.12E-06 | 0 | 0 | 0 | 0 | 8 mo | HOM |
| **1013609713** | 2 | tspd | 1051 | 4.12E-06 | 0 | 0 | 0 | 0 | 2 mo | WILD |
| **1010418842** | 1 | tspd | 1051 | 4.12E-06 | 0 | 0 | 0 | 0 | 8 mo | HOM |
| **1013288213** | 2 | tspd | 1051 | 4.12E-06 | 0 | 0 | 0 | 0 | 4 mo | HOM |
| **1008976060** | 2 | tspd | 1051 | 4.12E-06 | 0 | 0 | 0 | 0 | 8 mo | WILD |
| **1013446243** | 1 | tspd | 1051 | 4.12E-06 | 0 | 0 | 0 | 0 | 4 mo | HOM |
| **1013288951** | 2 | tspd | 1051 | 4.12E-06 | 0 | 0 | 0 | 0 | 4 mo | HOM |
| **1013978665** | 1 | PPN | 1052 | 0.919461402 | 0 | 0 | 0 | 0 | 2 mo | HOM |
| **1013445609** | 1 | PPN | 1052 | 0.919461402 | 0 | 0 | 0 | 0 | 4 mo | HOM |
| **1013609015** | 1 | PPN | 1052 | 0.919461402 | 0 | 0 | 0 | 0 | 4 mo | HOM |
| **1013608106** | 2 | PPN | 1052 | 0.919461402 | 0 | 0 | 0 | 0 | 4 mo | HOM |
| **1011937543** | 1 | PPN | 1052 | 0.919461402 | 0.005205339 | 28 | 0.005661292 | 30.45261057 | 8 mo | HOM |
| **1010419538** | 2 | PPN | 1052 | 0.919461402 | 0 | 0 | 0 | 0 | 4 mo | WILD |
| **1010419187** | 1 | PPN | 1052 | 0.919461402 | 0 | 0 | 0 | 0 | 8 mo | HOM |
| **1013978366** | 1 | PPN | 1052 | 0.919461402 | 0 | 0 | 0 | 0 | 2 mo | HOM |
| **1014416155** | 2 | PPN | 1052 | 0.919461402 | 0 | 0 | 0 | 0 | 2 mo | HOM |
| **1013978065** | 2 | PPN | 1052 | 0.919461402 | 0 | 0 | 0 | 0 | 2 mo | HOM |
| **1009934102** | 2 | PPN | 1052 | 0.919461402 | 0 | 0 | 0 | 0 | 4 mo | WILD |
| **1009322937** | 2 | PPN | 1052 | 0.919461402 | 0 | 0 | 0 | 0 | 8 mo | WILD |
| **1011080637** | 2 | PPN | 1052 | 0.919461402 | 0 | 0 | 0 | 0 | 8 mo | HOM |
| **1013609713** | 2 | PPN | 1052 | 0.919461402 | 0 | 0 | 0 | 0 | 2 mo | WILD |
| **1010418842** | 1 | PPN | 1052 | 0.919461402 | 3.43E-05 | 2 | 3.73E-05 | 2.17518647 | 8 mo | HOM |
| **1013288213** | 2 | PPN | 1052 | 0.919461402 | 0 | 0 | 0 | 0 | 4 mo | HOM |
| **1008976060** | 2 | PPN | 1052 | 0.919461402 | 2.22E-06 | 0 | 2.42E-06 | 0 | 8 mo | WILD |
| **1013446243** | 1 | PPN | 1052 | 0.919461402 | 0 | 0 | 0 | 0 | 4 mo | HOM |
| **1013288951** | 2 | PPN | 1052 | 0.919461402 | 0 | 0 | 0 | 0 | 4 mo | HOM |
| **1013978665** | 1 | ILA6a | 1054 | 0.211274417 | 0 | 0 | 0 | 0 | 2 mo | HOM |
| **1013445609** | 1 | ILA6a | 1054 | 0.211274417 | 3.69E-06 | 0 | 1.75E-05 | 0 | 4 mo | HOM |
| **1013609015** | 1 | ILA6a | 1054 | 0.211274417 | 2.93E-05 | 2 | 0.000138755 | 9.466361458 | 4 mo | HOM |
| **1013608106** | 2 | ILA6a | 1054 | 0.211274417 | 0 | 0 | 0 | 0 | 4 mo | HOM |
| **1011937543** | 1 | ILA6a | 1054 | 0.211274417 | 5.02E-05 | 1 | 0.000237532 | 4.733180729 | 8 mo | HOM |
| **1010419538** | 2 | ILA6a | 1054 | 0.211274417 | 0 | 0 | 0 | 0 | 4 mo | WILD |
| **1010419187** | 1 | ILA6a | 1054 | 0.211274417 | 0.000296717 | 5 | 0.001404414 | 23.66590364 | 8 mo | HOM |
| **1013978366** | 1 | ILA6a | 1054 | 0.211274417 | 0 | 0 | 0 | 0 | 2 mo | HOM |
| **1014416155** | 2 | ILA6a | 1054 | 0.211274417 | 0 | 0 | 0 | 0 | 2 mo | HOM |
| **1013978065** | 2 | ILA6a | 1054 | 0.211274417 | 0 | 0 | 0 | 0 | 2 mo | HOM |
| **1009934102** | 2 | ILA6a | 1054 | 0.211274417 | 0 | 0 | 0 | 0 | 4 mo | WILD |

| | | | | | | | | | | |
|---|---|---|---|---|---|---|---|---|---|---|
| **1009322937** | 2 | ILA6a | 1054 | 0.211274417 | 0 | 0 | 0 | 0 | 8 mo | WILD |
| **1011080637** | 2 | ILA6a | 1054 | 0.211274417 | 0.000116449 | 6 | 0.000551175 | 28.39908437 | 8 mo | HOM |
| **1013609713** | 2 | ILA6a | 1054 | 0.211274417 | 0 | 0 | 0 | 0 | 2 mo | WILD |
| **1010418842** | 1 | ILA6a | 1054 | 0.211274417 | 0.000171502 | 5 | 0.000811751 | 23.66590364 | 8 mo | HOM |
| **1013288213** | 2 | ILA6a | 1054 | 0.211274417 | 0 | 0 | 0 | 0 | 4 mo | HOM |
| **1008976060** | 2 | ILA6a | 1054 | 0.211274417 | 0 | 0 | 0 | 0 | 8 mo | WILD |
| **1013446243** | 1 | ILA6a | 1054 | 0.211274417 | 2.49E-05 | 2 | 0.000117938 | 9.466361458 | 4 mo | HOM |
| **1013288951** | 2 | ILA6a | 1054 | 0.211274417 | 4.09E-06 | 1 | 1.94E-05 | 4.733180729 | 4 mo | HOM |
| **1013978665** | 1 | ANcr1 | 1056 | 5.865874202 | 0.00021473 | 18 | 3.66E-05 | 3.068596322 | 2 mo | HOM |
| **1013445609** | 1 | ANcr1 | 1056 | 5.865874202 | 6.13E-06 | 1 | 1.05E-06 | 0.170477573 | 4 mo | HOM |
| **1013609015** | 1 | ANcr1 | 1056 | 5.865874202 | 0.001840997 | 11 | 0.000313849 | 1.875253308 | 4 mo | HOM |
| **1013608106** | 2 | ANcr1 | 1056 | 5.865874202 | 0 | 0 | 0 | 0 | 4 mo | HOM |
| **1011937543** | 1 | ANcr1 | 1056 | 5.865874202 | 0.01116556 | 20 | 0.001903478 | 3.409551469 | 8 mo | HOM |
| **1010419538** | 2 | ANcr1 | 1056 | 5.865874202 | 5.63E-05 | 6 | 9.6E-06 | 1.022865441 | 4 mo | WILD |
| **1010419187** | 1 | ANcr1 | 1056 | 5.865874202 | 0.004909616 | 68 | 0.000836979 | 11.59247499 | 8 mo | HOM |
| **1013978366** | 1 | ANcr1 | 1056 | 5.865874202 | 0.002295971 | 23 | 0.000391412 | 3.920984189 | 2 mo | HOM |
| **1014416155** | 2 | ANcr1 | 1056 | 5.865874202 | 0 | 0 | 0 | 0 | 2 mo | HOM |
| **1013978065** | 2 | ANcr1 | 1056 | 5.865874202 | 1.21E-05 | 0 | 2.07E-06 | 0 | 2 mo | HOM |
| **1009934102** | 2 | ANcr1 | 1056 | 5.865874202 | 6.83E-05 | 4 | 1.16E-05 | 0.681910294 | 4 mo | WILD |
| **1009322937** | 2 | ANcr1 | 1056 | 5.865874202 | 5.23E-05 | 6 | 8.91E-06 | 1.022865441 | 8 mo | WILD |
| **1011080637** | 2 | ANcr1 | 1056 | 5.865874202 | 0.005457363 | 85 | 0.000930358 | 14.49059374 | 8 mo | HOM |
| **1013609713** | 2 | ANcr1 | 1056 | 5.865874202 | 3.65E-05 | 3 | 6.23E-06 | 0.51143272 | 2 mo | WILD |
| **1010418842** | 1 | ANcr1 | 1056 | 5.865874202 | 0.010693796 | 95 | 0.001823052 | 16.19536948 | 8 mo | HOM |
| **1013288213** | 2 | ANcr1 | 1056 | 5.865874202 | 3.08E-05 | 3 | 5.26E-06 | 0.51143272 | 4 mo | HOM |
| **1008976060** | 2 | ANcr1 | 1056 | 5.865874202 | 0.000116987 | 16 | 1.99E-05 | 2.727641175 | 8 mo | WILD |
| **1013446243** | 1 | ANcr1 | 1056 | 5.865874202 | 5.98E-05 | 4 | 1.02E-05 | 0.681910294 | 4 mo | HOM |
| **1013288951** | 2 | ANcr1 | 1056 | 5.865874202 | 0.000574727 | 16 | 9.8E-05 | 2.727641175 | 4 mo | HOM |
| **1013978665** | 1 | GU | 1057 | 1.821761288 | 5.08E-06 | 1 | 2.79E-06 | 0.548919338 | 2 mo | HOM |
| **1013445609** | 1 | GU | 1057 | 1.821761288 | 0.000112555 | 6 | 6.18E-05 | 3.293516027 | 4 mo | HOM |
| **1013609015** | 1 | GU | 1057 | 1.821761288 | 0.000424649 | 14 | 0.000233098 | 7.68487073 | 4 mo | HOM |
| **1013608106** | 2 | GU | 1057 | 1.821761288 | 0 | 0 | 0 | 0 | 4 mo | HOM |
| **1011937543** | 1 | GU | 1057 | 1.821761288 | 0.002432725 | 68 | 0.00133537 | 37.32651497 | 8 mo | HOM |
| **1010419538** | 2 | GU | 1057 | 1.821761288 | 2.47E-06 | 0 | 1.36E-06 | 0 | 4 mo | WILD |
| **1010419187** | 1 | GU | 1057 | 1.821761288 | 0.001839812 | 83 | 0.001009909 | 45.56030504 | 8 mo | HOM |
| **1013978366** | 1 | GU | 1057 | 1.821761288 | 8.24E-21 | 0 | 4.52E-21 | 0 | 2 mo | HOM |
| **1014416155** | 2 | GU | 1057 | 1.821761288 | 0 | 0 | 0 | 0 | 2 mo | HOM |
| **1013978065** | 2 | GU | 1057 | 1.821761288 | 0 | 0 | 0 | 0 | 2 mo | HOM |
| **1009934102** | 2 | GU | 1057 | 1.821761288 | 0 | 0 | 0 | 0 | 4 mo | WILD |
| **1009322937** | 2 | GU | 1057 | 1.821761288 | 3.36E-06 | 1 | 1.84E-06 | 0.548919338 | 8 mo | WILD |
| **1011080637** | 2 | GU | 1057 | 1.821761288 | 0.001009824 | 68 | 0.000554312 | 37.32651497 | 8 mo | HOM |
| **1013609713** | 2 | GU | 1057 | 1.821761288 | 0 | 0 | 0 | 0 | 2 mo | WILD |
| **1010418842** | 1 | GU | 1057 | 1.821761288 | 0.001484542 | 72 | 0.000814894 | 39.52219232 | 8 mo | HOM |
| **1013288213** | 2 | GU | 1057 | 1.821761288 | 3.36E-05 | 3 | 1.85E-05 | 1.646758013 | 4 mo | HOM |
| **1008976060** | 2 | GU | 1057 | 1.821761288 | 1.49E-05 | 2 | 8.16E-06 | 1.097838676 | 8 mo | WILD |
| **1013446243** | 1 | GU | 1057 | 1.821761288 | 0.000247373 | 11 | 0.000135788 | 6.038112716 | 4 mo | HOM |
| **1013288951** | 2 | GU | 1057 | 1.821761288 | 0.000253307 | 15 | 0.000139045 | 8.233790067 | 4 mo | HOM |
| **1013978665** | 1 | VISC5 | 1058 | 0.727785946 | 4.87E-06 | 1 | 6.68E-06 | 1.374030381 | 2 mo | HOM |
| **1013445609** | 1 | VISC5 | 1058 | 0.727785946 | 6.11E-05 | 2 | 8.4E-05 | 2.748060762 | 4 mo | HOM |
| **1013609015** | 1 | VISC5 | 1058 | 0.727785946 | 0.00011139 | 4 | 0.000153053 | 5.496121523 | 4 mo | HOM |
| **1013608106** | 2 | VISC5 | 1058 | 0.727785946 | 0 | 0 | 0 | 0 | 4 mo | HOM |
| **1011937543** | 1 | VISC5 | 1058 | 0.727785946 | 0.000994317 | 45 | 0.001366222 | 61.83136714 | 8 mo | HOM |
| **1010419538** | 2 | VISC5 | 1058 | 0.727785946 | 0 | 0 | 0 | 0 | 4 mo | WILD |
| **1010419187** | 1 | VISC5 | 1058 | 0.727785946 | 0.000996953 | 40 | 0.001369844 | 54.96121523 | 8 mo | HOM |
| **1013978366** | 1 | VISC5 | 1058 | 0.727785946 | 0 | 0 | 0 | 0 | 2 mo | HOM |
| **1014416155** | 2 | VISC5 | 1058 | 0.727785946 | 0 | 0 | 0 | 0 | 2 mo | HOM |
| **1013978065** | 2 | VISC5 | 1058 | 0.727785946 | 0 | 0 | 0 | 0 | 2 mo | HOM |
| **1009934102** | 2 | VISC5 | 1058 | 0.727785946 | 0 | 0 | 0 | 0 | 4 mo | WILD |
| **1009322937** | 2 | VISC5 | 1058 | 0.727785946 | 0 | 0 | 0 | 0 | 8 mo | WILD |
| **1011080637** | 2 | VISC5 | 1058 | 0.727785946 | 0.00026228 | 16 | 0.00036038 | 21.98448609 | 8 mo | HOM |

| | | | | | | | | | | |
|---|---|---|---|---|---|---|---|---|---|---|
| **1013609713** | 2 | VISC5 | 1058 | 0.727785946 | 0 | 0 | 0 | 0 | 2 mo | WILD |
| **1010418842** | 1 | VISC5 | 1058 | 0.727785946 | 0.000446602 | 28 | 0.000613645 | 38.47285066 | 8 mo | HOM |
| **1013288213** | 2 | VISC5 | 1058 | 0.727785946 | 5.12E-05 | 5 | 7.04E-05 | 6.870151904 | 4 mo | HOM |
| **1008976060** | 2 | VISC5 | 1058 | 0.727785946 | 5.56E-06 | 1 | 7.64E-06 | 1.374030381 | 8 mo | WILD |
| **1013446243** | 1 | VISC5 | 1058 | 0.727785946 | 0.000137046 | 8 | 0.000188305 | 10.99224305 | 4 mo | HOM |
| **1013288951** | 2 | VISC5 | 1058 | 0.727785946 | 0.00016235 | 10 | 0.000223073 | 13.74030381 | 4 mo | HOM |
| **1013978665** | 1 | dtd | 1060 | 0.007139459 | 0 | 0 | 0 | 0 | 2 mo | HOM |
| **1013445609** | 1 | dtd | 1060 | 0.007139459 | 0 | 0 | 0 | 0 | 4 mo | HOM |
| **1013609015** | 1 | dtd | 1060 | 0.007139459 | 0 | 0 | 0 | 0 | 4 mo | HOM |
| **1013608106** | 2 | dtd | 1060 | 0.007139459 | 0 | 0 | 0 | 0 | 4 mo | HOM |
| **1011937543** | 1 | dtd | 1060 | 0.007139459 | 0 | 0 | 0 | 0 | 8 mo | HOM |
| **1010419538** | 2 | dtd | 1060 | 0.007139459 | 0 | 0 | 0 | 0 | 4 mo | WILD |
| **1010419187** | 1 | dtd | 1060 | 0.007139459 | 0 | 0 | 0 | 0 | 8 mo | HOM |
| **1013978366** | 1 | dtd | 1060 | 0.007139459 | 0 | 0 | 0 | 0 | 2 mo | HOM |
| **1014416155** | 2 | dtd | 1060 | 0.007139459 | 0 | 0 | 0 | 0 | 2 mo | HOM |
| **1013978065** | 2 | dtd | 1060 | 0.007139459 | 0 | 0 | 0 | 0 | 2 mo | HOM |
| **1009934102** | 2 | dtd | 1060 | 0.007139459 | 0 | 0 | 0 | 0 | 4 mo | WILD |
| **1009322937** | 2 | dtd | 1060 | 0.007139459 | 0 | 0 | 0 | 0 | 8 mo | WILD |
| **1011080637** | 2 | dtd | 1060 | 0.007139459 | 0 | 0 | 0 | 0 | 8 mo | HOM |
| **1013609713** | 2 | dtd | 1060 | 0.007139459 | 0 | 0 | 0 | 0 | 2 mo | WILD |
| **1010418842** | 1 | dtd | 1060 | 0.007139459 | 9.3E-06 | 1 | 0.001303115 | 140.0666304 | 8 mo | HOM |
| **1013288213** | 2 | dtd | 1060 | 0.007139459 | 0 | 0 | 0 | 0 | 4 mo | HOM |
| **1008976060** | 2 | dtd | 1060 | 0.007139459 | 0 | 0 | 0 | 0 | 8 mo | WILD |
| **1013446243** | 1 | dtd | 1060 | 0.007139459 | 0 | 0 | 0 | 0 | 4 mo | HOM |
| **1013288951** | 2 | dtd | 1060 | 0.007139459 | 0 | 0 | 0 | 0 | 4 mo | HOM |
| **1013978665** | 1 | PPT | 1061 | 0.149207489 | 0 | 0 | 0 | 0 | 2 mo | HOM |
| **1013445609** | 1 | PPT | 1061 | 0.149207489 | 0 | 0 | 0 | 0 | 4 mo | HOM |
| **1013609015** | 1 | PPT | 1061 | 0.149207489 | 0 | 0 | 0 | 0 | 4 mo | HOM |
| **1013608106** | 2 | PPT | 1061 | 0.149207489 | 0 | 0 | 0 | 0 | 4 mo | HOM |
| **1011937543** | 1 | PPT | 1061 | 0.149207489 | 2.97E-05 | 1 | 0.000199119 | 6.702076453 | 8 mo | HOM |
| **1010419538** | 2 | PPT | 1061 | 0.149207489 | 0 | 0 | 0 | 0 | 4 mo | WILD |
| **1010419187** | 1 | PPT | 1061 | 0.149207489 | 7.21E-07 | 0 | 4.84E-06 | 0 | 8 mo | HOM |
| **1013978366** | 1 | PPT | 1061 | 0.149207489 | 0 | 0 | 0 | 0 | 2 mo | HOM |
| **1014416155** | 2 | PPT | 1061 | 0.149207489 | 0 | 0 | 0 | 0 | 2 mo | HOM |
| **1013978065** | 2 | PPT | 1061 | 0.149207489 | 0 | 0 | 0 | 0 | 2 mo | HOM |
| **1009934102** | 2 | PPT | 1061 | 0.149207489 | 0 | 0 | 0 | 0 | 4 mo | WILD |
| **1009322937** | 2 | PPT | 1061 | 0.149207489 | 1.33E-05 | 1 | 8.94E-05 | 6.702076453 | 8 mo | WILD |
| **1011080637** | 2 | PPT | 1061 | 0.149207489 | 9.82E-06 | 1 | 6.58E-05 | 6.702076453 | 8 mo | HOM |
| **1013609713** | 2 | PPT | 1061 | 0.149207489 | 0 | 0 | 0 | 0 | 2 mo | WILD |
| **1010418842** | 1 | PPT | 1061 | 0.149207489 | 5.17E-05 | 2 | 0.00034634 | 13.40415291 | 8 mo | HOM |
| **1013288213** | 2 | PPT | 1061 | 0.149207489 | 0 | 0 | 0 | 0 | 4 mo | HOM |
| **1008976060** | 2 | PPT | 1061 | 0.149207489 | 0 | 0 | 0 | 0 | 8 mo | WILD |
| **1013446243** | 1 | PPT | 1061 | 0.149207489 | 0 | 0 | 0 | 0 | 4 mo | HOM |
| **1013288951** | 2 | PPT | 1061 | 0.149207489 | 0 | 0 | 0 | 0 | 4 mo | HOM |
| **1013978665** | 1 | SSp-bfd6b | 1062 | 0.150983592 | 0 | 0 | 0 | 0 | 2 mo | HOM |
| **1013445609** | 1 | SSp-bfd6b | 1062 | 0.150983592 | 3.65E-06 | 0 | 2.42E-05 | 0 | 4 mo | HOM |
| **1013609015** | 1 | SSp-bfd6b | 1062 | 0.150983592 | 2.15E-06 | 0 | 1.42E-05 | 0 | 4 mo | HOM |
| **1013608106** | 2 | SSp-bfd6b | 1062 | 0.150983592 | 0 | 0 | 0 | 0 | 4 mo | HOM |
| **1011937543** | 1 | SSp-bfd6b | 1062 | 0.150983592 | 0.000218979 | 3 | 0.001450352 | 19.86970873 | 8 mo | HOM |
| **1010419538** | 2 | SSp-bfd6b | 1062 | 0.150983592 | 0 | 0 | 0 | 0 | 4 mo | WILD |
| **1010419187** | 1 | SSp-bfd6b | 1062 | 0.150983592 | 9.93E-05 | 7 | 0.000657518 | 46.3626537 | 8 mo | HOM |
| **1013978366** | 1 | SSp-bfd6b | 1062 | 0.150983592 | 0 | 0 | 0 | 0 | 2 mo | HOM |
| **1014416155** | 2 | SSp-bfd6b | 1062 | 0.150983592 | 0 | 0 | 0 | 0 | 2 mo | HOM |
| **1013978065** | 2 | SSp-bfd6b | 1062 | 0.150983592 | 0 | 0 | 0 | 0 | 2 mo | HOM |
| **1009934102** | 2 | SSp-bfd6b | 1062 | 0.150983592 | 0 | 0 | 0 | 0 | 4 mo | WILD |
| **1009322937** | 2 | SSp-bfd6b | 1062 | 0.150983592 | 0 | 0 | 0 | 0 | 8 mo | WILD |
| **1011080637** | 2 | SSp-bfd6b | 1062 | 0.150983592 | 6.36E-05 | 4 | 0.000421388 | 26.49294497 | 8 mo | HOM |
| **1013609713** | 2 | SSp-bfd6b | 1062 | 0.150983592 | 0 | 0 | 0 | 0 | 2 mo | WILD |
| **1010418842** | 1 | SSp-bfd6b | 1062 | 0.150983592 | 0.000152892 | 3 | 0.001012642 | 19.86970873 | 8 mo | HOM |

| | | | | | | | | | | |
|---|---|---|---|---|---|---|---|---|---|---|
| 1013288213 | 2 | SSp-bfd6b | 1062 | 0.150983592 | 0 | 0 | 0 | 0 | 4 mo | HOM |
| 1008976060 | 2 | SSp-bfd6b | 1062 | 0.150983592 | 0 | 0 | 0 | 0 | 8 mo | WILD |
| 1013446243 | 1 | SSp-bfd6b | 1062 | 0.150983592 | 2.92E-05 | 3 | 0.000193499 | 19.86970873 | 4 mo | HOM |
| 1013288951 | 2 | SSp-bfd6b | 1062 | 0.150983592 | 6.04E-06 | 0 | 4E-05 | 0 | 4 mo | HOM |
| 1013978665 | 1 | ANcr2 | 1064 | 5.266357555 | 6.85E-05 | 9 | 1.3E-05 | 1.708961062 | 2 mo | HOM |
| 1013445609 | 1 | ANcr2 | 1064 | 5.266357555 | 0 | 0 | 0 | 0 | 4 mo | HOM |
| 1013609015 | 1 | ANcr2 | 1064 | 5.266357555 | 0.000256648 | 11 | 4.87E-05 | 2.088730187 | 4 mo | HOM |
| 1013608106 | 2 | ANcr2 | 1064 | 5.266357555 | 0 | 0 | 0 | 0 | 4 mo | HOM |
| 1011937543 | 1 | ANcr2 | 1064 | 5.266357555 | 0.008884889 | 33 | 0.001687103 | 6.26619056 | 8 mo | HOM |
| 1010419538 | 2 | ANcr2 | 1064 | 5.266357555 | 0.000197327 | 9 | 3.75E-05 | 1.708961062 | 4 mo | WILD |
| 1010419187 | 1 | ANcr2 | 1064 | 5.266357555 | 0.009798289 | 88 | 0.001860544 | 16.70984149 | 8 mo | HOM |
| 1013978366 | 1 | ANcr2 | 1064 | 5.266357555 | 0.003429231 | 71 | 0.000651158 | 13.48180393 | 2 mo | HOM |
| 1014416155 | 2 | ANcr2 | 1064 | 5.266357555 | 0 | 0 | 0 | 0 | 2 mo | HOM |
| 1013978065 | 2 | ANcr2 | 1064 | 5.266357555 | 0 | 0 | 0 | 0 | 2 mo | HOM |
| 1009934102 | 2 | ANcr2 | 1064 | 5.266357555 | 0.00014485 | 12 | 2.75E-05 | 2.278614749 | 4 mo | WILD |
| 1009322937 | 2 | ANcr2 | 1064 | 5.266357555 | 5.76E-05 | 7 | 1.09E-05 | 1.329191937 | 8 mo | WILD |
| 1011080637 | 2 | ANcr2 | 1064 | 5.266357555 | 0.006281719 | 100 | 0.001192802 | 18.98845624 | 8 mo | HOM |
| 1013609713 | 2 | ANcr2 | 1064 | 5.266357555 | 6.14E-05 | 7 | 1.17E-05 | 1.329191937 | 2 mo | WILD |
| 1010418842 | 1 | ANcr2 | 1064 | 5.266357555 | 0.009993406 | 80 | 0.001897593 | 15.19076499 | 8 mo | HOM |
| 1013288213 | 2 | ANcr2 | 1064 | 5.266357555 | 0.000199815 | 4 | 3.79E-05 | 0.75953825 | 4 mo | HOM |
| 1008976060 | 2 | ANcr2 | 1064 | 5.266357555 | 0.00011976 | 11 | 2.27E-05 | 2.088730187 | 8 mo | WILD |
| 1013446243 | 1 | ANcr2 | 1064 | 5.266357555 | 0.000129283 | 2 | 2.45E-05 | 0.379769125 | 4 mo | HOM |
| 1013288951 | 2 | ANcr2 | 1064 | 5.266357555 | 0.000278782 | 22 | 5.29E-05 | 4.177460374 | 4 mo | HOM |
| 1013978665 | 1 | HB | 1065 | 49.11529921 | 0.001382706 | 27 | 2.82E-05 | 0.549726876 | 2 mo | HOM |
| 1013445609 | 1 | HB | 1065 | 49.11529921 | 0.000212076 | 9 | 4.32E-06 | 0.183242292 | 4 mo | HOM |
| 1013609015 | 1 | HB | 1065 | 49.11529921 | 0.009811969 | 40 | 0.000199774 | 0.814410187 | 4 mo | HOM |
| 1013608106 | 2 | HB | 1065 | 49.04544284 | 0 | 0 | 0 | 0 | 4 mo | HOM |
| 1011937543 | 1 | HB | 1065 | 49.11529921 | 0.053562397 | 146 | 0.001090544 | 2.972597182 | 8 mo | HOM |
| 1010419538 | 2 | HB | 1065 | 49.11529921 | 0.000103332 | 10 | 2.1E-06 | 0.203602547 | 4 mo | WILD |
| 1010419187 | 1 | HB | 1065 | 49.11529921 | 0.016233813 | 86 | 0.000330525 | 1.750981902 | 8 mo | HOM |
| 1013978366 | 1 | HB | 1065 | 49.11529921 | 0.001967742 | 20 | 4.01E-05 | 0.407205093 | 2 mo | HOM |
| 1014416155 | 2 | HB | 1065 | 49.11529921 | 0 | 0 | 0 | 0 | 2 mo | HOM |
| 1013978065 | 2 | HB | 1065 | 49.11529921 | 0.000433514 | 8 | 8.83E-06 | 0.162882037 | 2 mo | HOM |
| 1009934102 | 2 | HB | 1065 | 49.11529921 | 3.84E-05 | 3 | 7.82E-07 | 0.061080764 | 4 mo | WILD |
| 1009322937 | 2 | HB | 1065 | 49.11529921 | 0.000531539 | 28 | 1.08E-05 | 0.570087131 | 8 mo | WILD |
| 1011080637 | 2 | HB | 1065 | 49.11529921 | 0.007157835 | 90 | 0.000145735 | 1.83242292 | 8 mo | HOM |
| 1013609713 | 2 | HB | 1065 | 49.11529921 | 1.54E-05 | 1 | 3.14E-07 | 0.020360255 | 2 mo | WILD |
| 1010418842 | 1 | HB | 1065 | 49.11529921 | 0.008758692 | 317 | 0.000178329 | 6.45420073 | 8 mo | HOM |
| 1013288213 | 2 | HB | 1065 | 49.11372643 | 0.000624627 | 12 | 1.27E-05 | 0.24433088 | 4 mo | HOM |
| 1008976060 | 2 | HB | 1065 | 49.11529921 | 0.011930204 | 303 | 0.000242902 | 6.169157165 | 8 mo | WILD |
| 1013446243 | 1 | HB | 1065 | 49.11529921 | 0.001193986 | 27 | 2.43E-05 | 0.549726876 | 4 mo | HOM |
| 1013288951 | 2 | HB | 1065 | 49.11529921 | 0.004157843 | 31 | 8.47E-05 | 0.631167895 | 4 mo | HOM |
| 1013978665 | 1 | VISam2/3 | 1066 | 0.213207114 | 0 | 0 | 0 | 0 | 2 mo | HOM |
| 1013445609 | 1 | VISam2/3 | 1066 | 0.213207114 | 4.25E-07 | 0 | 1.99E-06 | 0 | 4 mo | HOM |
| 1013609015 | 1 | VISam2/3 | 1066 | 0.213207114 | 3.19E-05 | 1 | 0.000149786 | 4.690275007 | 4 mo | HOM |
| 1013608106 | 2 | VISam2/3 | 1066 | 0.213207114 | 0 | 0 | 0 | 0 | 4 mo | HOM |
| 1011937543 | 1 | VISam2/3 | 1066 | 0.213207114 | 0.000702668 | 5 | 0.003295704 | 23.45137504 | 8 mo | HOM |
| 1010419538 | 2 | VISam2/3 | 1066 | 0.213207114 | 0 | 0 | 0 | 0 | 4 mo | WILD |
| 1010419187 | 1 | VISam2/3 | 1066 | 0.213207114 | 0.000350098 | 10 | 0.001642057 | 46.90275007 | 8 mo | HOM |
| 1013978366 | 1 | VISam2/3 | 1066 | 0.213207114 | 0 | 0 | 0 | 0 | 2 mo | HOM |
| 1014416155 | 2 | VISam2/3 | 1066 | 0.213207114 | 0 | 0 | 0 | 0 | 2 mo | HOM |
| 1013978065 | 2 | VISam2/3 | 1066 | 0.213207114 | 0 | 0 | 0 | 0 | 2 mo | HOM |
| 1009934102 | 2 | VISam2/3 | 1066 | 0.213207114 | 0 | 0 | 0 | 0 | 4 mo | WILD |
| 1009322937 | 2 | VISam2/3 | 1066 | 0.213207114 | 0 | 0 | 0 | 0 | 8 mo | WILD |
| 1011080637 | 2 | VISam2/3 | 1066 | 0.213207114 | 0.000134849 | 8 | 0.000632479 | 37.52220006 | 8 mo | HOM |
| 1013609713 | 2 | VISam2/3 | 1066 | 0.213207114 | 0 | 0 | 0 | 0 | 2 mo | WILD |
| 1010418842 | 1 | VISam2/3 | 1066 | 0.213207114 | 0.000205023 | 7 | 0.000961612 | 32.83192505 | 8 mo | HOM |
| 1013288213 | 2 | VISam2/3 | 1066 | 0.213207114 | 9.94E-06 | 1 | 4.66E-05 | 4.690275007 | 4 mo | HOM |
| 1008976060 | 2 | VISam2/3 | 1066 | 0.213207114 | 0 | 0 | 0 | 0 | 8 mo | WILD |

| | | | | | | | | | | |
|---|---|---|---|---|---|---|---|---|---|---|
| 1013446243 | 1 | VISam2/3 | 1066 | 0.213207114 | 6.33E-05 | 4 | 0.000296835 | 18.76110003 | 4 mo | HOM |
| 1013288951 | 2 | VISam2/3 | 1066 | 0.213207114 | 3.62E-05 | 3 | 0.000169982 | 14.07082502 | 4 mo | HOM |
| 1013978665 | 1 | PPY | 1069 | 0.097331542 | 0 | 0 | 0 | 0 | 2 mo | HOM |
| 1013445609 | 1 | PPY | 1069 | 0.097331542 | 0 | 0 | 0 | 0 | 4 mo | HOM |
| 1013609015 | 1 | PPY | 1069 | 0.097331542 | 0 | 0 | 0 | 0 | 4 mo | HOM |
| 1013608106 | 2 | PPY | 1069 | 0.097331542 | 0 | 0 | 0 | 0 | 4 mo | HOM |
| 1011937543 | 1 | PPY | 1069 | 0.097331542 | 0 | 0 | 0 | 0 | 8 mo | HOM |
| 1010419538 | 2 | PPY | 1069 | 0.097331542 | 0 | 0 | 0 | 0 | 4 mo | WILD |
| 1010419187 | 1 | PPY | 1069 | 0.097331542 | 0 | 0 | 0 | 0 | 8 mo | HOM |
| 1013978366 | 1 | PPY | 1069 | 0.097331542 | 0 | 0 | 0 | 0 | 2 mo | HOM |
| 1014416155 | 2 | PPY | 1069 | 0.097331542 | 0 | 0 | 0 | 0 | 2 mo | HOM |
| 1013978065 | 2 | PPY | 1069 | 0.097331542 | 0 | 0 | 0 | 0 | 2 mo | HOM |
| 1009934102 | 2 | PPY | 1069 | 0.097331542 | 0 | 0 | 0 | 0 | 4 mo | WILD |
| 1009322937 | 2 | PPY | 1069 | 0.097331542 | 0 | 0 | 0 | 0 | 8 mo | WILD |
| 1011080637 | 2 | PPY | 1069 | 0.097331542 | 0 | 0 | 0 | 0 | 8 mo | HOM |
| 1013609713 | 2 | PPY | 1069 | 0.097331542 | 0 | 0 | 0 | 0 | 2 mo | WILD |
| 1010418842 | 1 | PPY | 1069 | 0.097331542 | 0 | 0 | 0 | 0 | 8 mo | HOM |
| 1013288213 | 2 | PPY | 1069 | 0.097331542 | 0 | 0 | 0 | 0 | 4 mo | HOM |
| 1008976060 | 2 | PPY | 1069 | 0.097331542 | 0 | 0 | 0 | 0 | 8 mo | WILD |
| 1013446243 | 1 | PPY | 1069 | 0.097331542 | 0 | 0 | 0 | 0 | 4 mo | HOM |
| 1013288951 | 2 | PPY | 1069 | 0.097331542 | 0 | 0 | 0 | 0 | 4 mo | HOM |
| 1013978665 | 1 | SSp-bfd5 | 1070 | 1.244550922 | 0 | 0 | 0 | 0 | 2 mo | HOM |
| 1013445609 | 1 | SSp-bfd5 | 1070 | 1.244550922 | 0.000141203 | 8 | 0.000113457 | 6.428021434 | 4 mo | HOM |
| 1013609015 | 1 | SSp-bfd5 | 1070 | 1.244550922 | 0.000516465 | 21 | 0.000414981 | 16.87355626 | 4 mo | HOM |
| 1013608106 | 2 | SSp-bfd5 | 1070 | 1.244550922 | 0 | 0 | 0 | 0 | 4 mo | HOM |
| 1011937543 | 1 | SSp-bfd5 | 1070 | 1.244550922 | 0.001885521 | 36 | 0.001515021 | 28.92609645 | 8 mo | HOM |
| 1010419538 | 2 | SSp-bfd5 | 1070 | 1.244550922 | 0 | 0 | 0 | 0 | 4 mo | WILD |
| 1010419187 | 1 | SSp-bfd5 | 1070 | 1.244550922 | 0.002333284 | 78 | 0.0018748 | 62.67320898 | 8 mo | HOM |
| 1013978366 | 1 | SSp-bfd5 | 1070 | 1.244550922 | 0 | 0 | 0 | 0 | 2 mo | HOM |
| 1014416155 | 2 | SSp-bfd5 | 1070 | 1.244550922 | 0 | 0 | 0 | 0 | 2 mo | HOM |
| 1013978065 | 2 | SSp-bfd5 | 1070 | 1.244550922 | 0 | 0 | 0 | 0 | 2 mo | HOM |
| 1009934102 | 2 | SSp-bfd5 | 1070 | 1.244550922 | 0 | 0 | 0 | 0 | 4 mo | WILD |
| 1009322937 | 2 | SSp-bfd5 | 1070 | 1.244550922 | 0 | 0 | 0 | 0 | 8 mo | WILD |
| 1011080637 | 2 | SSp-bfd5 | 1070 | 1.244550922 | 0.001397039 | 56 | 0.001122525 | 44.99615004 | 8 mo | HOM |
| 1013609713 | 2 | SSp-bfd5 | 1070 | 1.244550922 | 0 | 0 | 0 | 0 | 2 mo | WILD |
| 1010418842 | 1 | SSp-bfd5 | 1070 | 1.244550922 | 0.001407406 | 57 | 0.001130854 | 45.79965272 | 8 mo | HOM |
| 1013288213 | 2 | SSp-bfd5 | 1070 | 1.244550922 | 0.000153197 | 9 | 0.000123094 | 7.231524113 | 4 mo | HOM |
| 1008976060 | 2 | SSp-bfd5 | 1070 | 1.244550922 | 0 | 0 | 0 | 0 | 8 mo | WILD |
| 1013446243 | 1 | SSp-bfd5 | 1070 | 1.244550922 | 0.00030437 | 11 | 0.000244562 | 8.838529472 | 4 mo | HOM |
| 1013288951 | 2 | SSp-bfd5 | 1070 | 1.244550922 | 0.00027238 | 9 | 0.000218858 | 7.231524113 | 4 mo | HOM |
| 1013978665 | 1 | MGd | 1072 | 0.178321643 | 0 | 0 | 0 | 0 | 2 mo | HOM |
| 1013445609 | 1 | MGd | 1072 | 0.178321643 | 9.61E-22 | 0 | 5.39E-21 | 0 | 4 mo | HOM |
| 1013609015 | 1 | MGd | 1072 | 0.178321643 | 0 | 0 | 0 | 0 | 4 mo | HOM |
| 1013608106 | 2 | MGd | 1072 | 0.178321643 | 0 | 0 | 0 | 0 | 4 mo | HOM |
| 1011937543 | 1 | MGd | 1072 | 0.178321643 | 0.000352911 | 6 | 0.001979067 | 33.64706553 | 8 mo | HOM |
| 1010419538 | 2 | MGd | 1072 | 0.178321643 | 0 | 0 | 0 | 0 | 4 mo | WILD |
| 1010419187 | 1 | MGd | 1072 | 0.178321643 | 8.36E-05 | 4 | 0.000468887 | 22.43137702 | 8 mo | HOM |
| 1013978366 | 1 | MGd | 1072 | 0.178321643 | 0 | 0 | 0 | 0 | 2 mo | HOM |
| 1014416155 | 2 | MGd | 1072 | 0.178321643 | 0 | 0 | 0 | 0 | 2 mo | HOM |
| 1013978065 | 2 | MGd | 1072 | 0.178321643 | 0 | 0 | 0 | 0 | 2 mo | HOM |
| 1009934102 | 2 | MGd | 1072 | 0.178321643 | 0 | 0 | 0 | 0 | 4 mo | WILD |
| 1009322937 | 2 | MGd | 1072 | 0.178321643 | 0 | 0 | 0 | 0 | 8 mo | WILD |
| 1011080637 | 2 | MGd | 1072 | 0.178321643 | 6.5E-05 | 2 | 0.000364727 | 11.21568851 | 8 mo | HOM |
| 1013609713 | 2 | MGd | 1072 | 0.178321643 | 0 | 0 | 0 | 0 | 2 mo | WILD |
| 1010418842 | 1 | MGd | 1072 | 0.178321643 | 5.78E-05 | 4 | 0.000324207 | 22.43137702 | 8 mo | HOM |
| 1013288213 | 2 | MGd | 1072 | 0.178321643 | 0 | 0 | 0 | 0 | 4 mo | HOM |
| 1008976060 | 2 | MGd | 1072 | 0.178321643 | 2.67E-06 | 0 | 1.5E-05 | 0 | 8 mo | WILD |
| 1013446243 | 1 | MGd | 1072 | 0.178321643 | 0 | 0 | 0 | 0 | 4 mo | HOM |
| 1013288951 | 2 | MGd | 1072 | 0.178321643 | 0 | 0 | 0 | 0 | 4 mo | HOM |

| | | | | | | | | | | |
|---|---|---|---|---|---|---|---|---|---|---|
| **1013978665** | 1 | HEM | 1073 | 31.893157 | 0.001814282 | 99 | 5.69E-05 | 3.104114152 | 2 mo | HOM |
| **1013445609** | 1 | HEM | 1073 | 31.893157 | 9.88E-05 | 4 | 3.1E-06 | 0.125418754 | 4 mo | HOM |
| **1013609015** | 1 | HEM | 1073 | 31.893157 | 0.008151623 | 114 | 0.000255592 | 3.574434478 | 4 mo | HOM |
| **1013608106** | 2 | HEM | 1073 | 31.893157 | 0 | 0 | 0 | 0 | 4 mo | HOM |
| **1011937543** | 1 | HEM | 1073 | 31.893157 | 0.061802124 | 157 | 0.001937786 | 4.922686079 | 8 mo | HOM |
| **1010419538** | 2 | HEM | 1073 | 31.893157 | 0.000535912 | 33 | 1.68E-05 | 1.034704717 | 4 mo | WILD |
| **1010419187** | 1 | HEM | 1073 | 31.893157 | 0.036319274 | 347 | 0.00113878 | 10.88007688 | 8 mo | HOM |
| **1013978366** | 1 | HEM | 1073 | 31.893157 | 0.011400494 | 216 | 0.000357459 | 6.772612695 | 2 mo | HOM |
| **1014416155** | 2 | HEM | 1073 | 31.893157 | 0 | 0 | 0 | 0 | 2 mo | HOM |
| **1013978065** | 2 | HEM | 1073 | 31.893157 | 0.000298517 | 3 | 9.36E-06 | 0.094064065 | 2 mo | HOM |
| **1009934102** | 2 | HEM | 1073 | 31.893157 | 0.000670071 | 36 | 2.1E-05 | 1.128768782 | 4 mo | WILD |
| **1009322937** | 2 | HEM | 1073 | 31.893157 | 0.000946972 | 63 | 2.97E-05 | 1.975345369 | 8 mo | WILD |
| **1011080637** | 2 | HEM | 1073 | 31.893157 | 0.019541104 | 341 | 0.000612705 | 10.69194874 | 8 mo | HOM |
| **1013609713** | 2 | HEM | 1073 | 31.79818497 | 0.000611084 | 22 | 1.92E-05 | 0.691863388 | 2 mo | WILD |
| **1010418842** | 1 | HEM | 1073 | 31.893157 | 0.047644407 | 471 | 0.001493876 | 14.76805824 | 8 mo | HOM |
| **1013288213** | 2 | HEM | 1073 | 31.893157 | 0.000397229 | 21 | 1.25E-05 | 0.658448456 | 4 mo | HOM |
| **1008976060** | 2 | HEM | 1073 | 31.893157 | 0.001192658 | 92 | 3.74E-05 | 2.884631333 | 8 mo | WILD |
| **1013446243** | 1 | HEM | 1073 | 31.893157 | 0.001198061 | 37 | 3.76E-05 | 1.160123471 | 4 mo | HOM |
| **1013288951** | 2 | HEM | 1073 | 31.893157 | 0.009875848 | 118 | 0.000309654 | 3.699853231 | 4 mo | HOM |
| **1013978665** | 1 | VISal1 | 1074 | 0.121455281 | 3.43E-10 | 0 | 2.83E-09 | 0 | 2 mo | HOM |
| **1013445609** | 1 | VISal1 | 1074 | 0.121455281 | 2.07E-05 | 2 | 0.000170055 | 16.46696617 | 4 mo | HOM |
| **1013609015** | 1 | VISal1 | 1074 | 0.121455281 | 5.79E-05 | 3 | 0.00047699 | 24.70044925 | 4 mo | HOM |
| **1013608106** | 2 | VISal1 | 1074 | 0.121455281 | 0 | 0 | 0 | 0 | 4 mo | HOM |
| **1011937543** | 1 | VISal1 | 1074 | 0.121455281 | 0.000646332 | 4 | 0.00532156 | 32.93393234 | 8 mo | HOM |
| **1010419538** | 2 | VISal1 | 1074 | 0.121455281 | 0 | 0 | 0 | 0 | 4 mo | WILD |
| **1010419187** | 1 | VISal1 | 1074 | 0.121455281 | 0.000190929 | 7 | 0.001572012 | 57.63438159 | 8 mo | HOM |
| **1013978366** | 1 | VISal1 | 1074 | 0.121455281 | 0 | 0 | 0 | 0 | 2 mo | HOM |
| **1014416155** | 2 | VISal1 | 1074 | 0.121455281 | 0 | 0 | 0 | 0 | 2 mo | HOM |
| **1013978065** | 2 | VISal1 | 1074 | 0.121455281 | 0 | 0 | 0 | 0 | 2 mo | HOM |
| **1009934102** | 2 | VISal1 | 1074 | 0.121455281 | 2.38E-06 | 1 | 1.96E-05 | 8.233483085 | 4 mo | WILD |
| **1009322937** | 2 | VISal1 | 1074 | 0.121455281 | 0 | 0 | 0 | 0 | 8 mo | WILD |
| **1011080637** | 2 | VISal1 | 1074 | 0.121455281 | 0.000272183 | 5 | 0.002241016 | 41.16741542 | 8 mo | HOM |
| **1013609713** | 2 | VISal1 | 1074 | 0.121455281 | 0 | 0 | 0 | 0 | 2 mo | WILD |
| **1010418842** | 1 | VISal1 | 1074 | 0.121455281 | 0.00054141 | 12 | 0.00445769 | 98.80179701 | 8 mo | HOM |
| **1013288213** | 2 | VISal1 | 1074 | 0.121455281 | 1.81E-05 | 2 | 0.000148969 | 16.46696617 | 4 mo | HOM |
| **1008976060** | 2 | VISal1 | 1074 | 0.121455281 | 5.19E-05 | 5 | 0.000427255 | 41.16741542 | 8 mo | WILD |
| **1013446243** | 1 | VISal1 | 1074 | 0.121455281 | 8.38E-05 | 5 | 0.000689818 | 41.16741542 | 4 mo | HOM |
| **1013288951** | 2 | VISal1 | 1074 | 0.121455281 | 3.01E-05 | 4 | 0.000247975 | 32.93393234 | 4 mo | HOM |
| **1013978665** | 1 | PR | 1077 | 0.155897773 | 0 | 0 | 0 | 0 | 2 mo | HOM |
| **1013445609** | 1 | PR | 1077 | 0.155897773 | 0 | 0 | 0 | 0 | 4 mo | HOM |
| **1013609015** | 1 | PR | 1077 | 0.155897773 | 0 | 0 | 0 | 0 | 4 mo | HOM |
| **1013608106** | 2 | PR | 1077 | 0.155897773 | 0 | 0 | 0 | 0 | 4 mo | HOM |
| **1011937543** | 1 | PR | 1077 | 0.155897773 | 0 | 0 | 0 | 0 | 8 mo | HOM |
| **1010419538** | 2 | PR | 1077 | 0.155897773 | 0 | 0 | 0 | 0 | 4 mo | WILD |
| **1010419187** | 1 | PR | 1077 | 0.155897773 | 4.3E-05 | 1 | 0.000275623 | 6.414459822 | 8 mo | HOM |
| **1013978366** | 1 | PR | 1077 | 0.155897773 | 0 | 0 | 0 | 0 | 2 mo | HOM |
| **1014416155** | 2 | PR | 1077 | 0.155897773 | 0 | 0 | 0 | 0 | 2 mo | HOM |
| **1013978065** | 2 | PR | 1077 | 0.155897773 | 0 | 0 | 0 | 0 | 2 mo | HOM |
| **1009934102** | 2 | PR | 1077 | 0.155897773 | 0 | 0 | 0 | 0 | 4 mo | WILD |
| **1009322937** | 2 | PR | 1077 | 0.155897773 | 0 | 0 | 0 | 0 | 8 mo | WILD |
| **1011080637** | 2 | PR | 1077 | 0.155897773 | 6.49E-05 | 1 | 0.000416406 | 6.414459822 | 8 mo | HOM |
| **1013609713** | 2 | PR | 1077 | 0.155897773 | 0 | 0 | 0 | 0 | 2 mo | WILD |
| **1010418842** | 1 | PR | 1077 | 0.155897773 | 4.35E-05 | 4 | 0.000279085 | 25.65783929 | 8 mo | HOM |
| **1013288213** | 2 | PR | 1077 | 0.155897773 | 0 | 0 | 0 | 0 | 4 mo | HOM |
| **1008976060** | 2 | PR | 1077 | 0.155897773 | 0 | 0 | 0 | 0 | 8 mo | WILD |
| **1013446243** | 1 | PR | 1077 | 0.155897773 | 0 | 0 | 0 | 0 | 4 mo | HOM |
| **1013288951** | 2 | PR | 1077 | 0.155897773 | 0 | 0 | 0 | 0 | 4 mo | HOM |
| **1013978665** | 1 | MGv | 1079 | 0.284799525 | 0 | 0 | 0 | 0 | 2 mo | HOM |
| **1013445609** | 1 | MGv | 1079 | 0.284799525 | 9.65E-06 | 1 | 3.39E-05 | 3.511241814 | 4 mo | HOM |

| | | | | | | | | | | |
|---|---|---|---|---|---|---|---|---|---|---|
| **1013609015** | 1 | MGv | 1079 | 0.284799525 | 0 | 0 | 0 | 0 | 4 mo | HOM |
| **1013608106** | 2 | MGv | 1079 | 0.284799525 | 0 | 0 | 0 | 0 | 4 mo | HOM |
| **1011937543** | 1 | MGv | 1079 | 0.284799525 | 0.000363937 | 8 | 0.001277871 | 28.08993451 | 8 mo | HOM |
| **1010419538** | 2 | MGv | 1079 | 0.284799525 | 0 | 0 | 0 | 0 | 4 mo | WILD |
| **1010419187** | 1 | MGv | 1079 | 0.284799525 | 7.16E-05 | 2 | 0.000251455 | 7.022483627 | 8 mo | HOM |
| **1013978366** | 1 | MGv | 1079 | 0.284799525 | 0 | 0 | 0 | 0 | 2 mo | HOM |
| **1014416155** | 2 | MGv | 1079 | 0.284799525 | 0 | 0 | 0 | 0 | 2 mo | HOM |
| **1013978065** | 2 | MGv | 1079 | 0.284799525 | 0 | 0 | 0 | 0 | 2 mo | HOM |
| **1009934102** | 2 | MGv | 1079 | 0.284799525 | 0 | 0 | 0 | 0 | 4 mo | WILD |
| **1009322937** | 2 | MGv | 1079 | 0.284799525 | 0 | 0 | 0 | 0 | 8 mo | WILD |
| **1011080637** | 2 | MGv | 1079 | 0.284799525 | 6.22E-05 | 3 | 0.000218297 | 10.53372544 | 8 mo | HOM |
| **1013609713** | 2 | MGv | 1079 | 0.284799525 | 0 | 0 | 0 | 0 | 2 mo | WILD |
| **1010418842** | 1 | MGv | 1079 | 0.284799525 | 0.000193106 | 5 | 0.000678041 | 17.55620907 | 8 mo | HOM |
| **1013288213** | 2 | MGv | 1079 | 0.284799525 | 0 | 0 | 0 | 0 | 4 mo | HOM |
| **1008976060** | 2 | MGv | 1079 | 0.284799525 | 4.35E-05 | 3 | 0.000152753 | 10.53372544 | 8 mo | WILD |
| **1013446243** | 1 | MGv | 1079 | 0.284799525 | 0 | 0 | 0 | 0 | 4 mo | HOM |
| **1013288951** | 2 | MGv | 1079 | 0.284799525 | 0 | 0 | 0 | 0 | 4 mo | HOM |
| **1013978665** | 1 | HIP | 1080 | 24.43237721 | 4.28E-05 | 6 | 1.75E-06 | 0.245575776 | 2 mo | HOM |
| **1013445609** | 1 | HIP | 1080 | 24.43237721 | 0.000584658 | 45 | 2.39E-05 | 1.841818322 | 4 mo | HOM |
| **1013609015** | 1 | HIP | 1080 | 24.43237721 | 0.00310662 | 139 | 0.000127152 | 5.689172151 | 4 mo | HOM |
| **1013608106** | 2 | HIP | 1080 | 24.43237721 | 0 | 0 | 0 | 0 | 4 mo | HOM |
| **1011937543** | 1 | HIP | 1080 | 24.43237721 | 0.027509607 | 363 | 0.001125949 | 14.85733447 | 8 mo | HOM |
| **1010419538** | 2 | HIP | 1080 | 24.43237721 | 0.000101283 | 7 | 4.15E-06 | 0.286505072 | 4 mo | WILD |
| **1010419187** | 1 | HIP | 1080 | 24.43237721 | 0.039555592 | 961 | 0.001618983 | 39.33305351 | 8 mo | HOM |
| **1013978366** | 1 | HIP | 1080 | 24.43237721 | 2.62E-05 | 2 | 1.07E-06 | 0.081858592 | 2 mo | HOM |
| **1014416155** | 2 | HIP | 1080 | 24.43237721 | 0 | 0 | 0 | 0 | 2 mo | HOM |
| **1013978065** | 2 | HIP | 1080 | 24.43237721 | 1.01E-05 | 3 | 4.12E-07 | 0.122787888 | 2 mo | HOM |
| **1009934102** | 2 | HIP | 1080 | 24.43237721 | 0.000265742 | 14 | 1.09E-05 | 0.573010145 | 4 mo | WILD |
| **1009322937** | 2 | HIP | 1080 | 24.43237721 | 0.000448692 | 14 | 1.84E-05 | 0.573010145 | 8 mo | WILD |
| **1011080637** | 2 | HIP | 1080 | 24.43237721 | 0.02386229 | 767 | 0.000976667 | 31.39277007 | 8 mo | HOM |
| **1013609713** | 2 | HIP | 1080 | 24.43237721 | 6.14E-06 | 0 | 2.51E-07 | 0 | 2 mo | WILD |
| **1010418842** | 1 | HIP | 1080 | 24.43237721 | 0.035910562 | 987 | 0.001469794 | 40.3972152 | 8 mo | HOM |
| **1013288213** | 2 | HIP | 1080 | 24.43237721 | 0.003869239 | 76 | 0.000158365 | 3.1106265 | 4 mo | HOM |
| **1008976060** | 2 | HIP | 1080 | 24.43237721 | 0.000589679 | 71 | 2.41E-05 | 2.90598002 | 8 mo | WILD |
| **1013446243** | 1 | HIP | 1080 | 24.43237721 | 0.00228646 | 132 | 9.36E-05 | 5.402667079 | 4 mo | HOM |
| **1013288951** | 2 | HIP | 1080 | 24.43237721 | 0.002490964 | 145 | 0.000101953 | 5.934747928 | 4 mo | HOM |
| **1013978665** | 1 | ILA6b | 1081 | 0.011155276 | 0 | 0 | 0 | 0 | 2 mo | HOM |
| **1013445609** | 1 | ILA6b | 1081 | 0.011155276 | 0 | 0 | 0 | 0 | 4 mo | HOM |
| **1013609015** | 1 | ILA6b | 1081 | 0.011155276 | 0 | 0 | 0 | 0 | 4 mo | HOM |
| **1013608106** | 2 | ILA6b | 1081 | 0.011155276 | 0 | 0 | 0 | 0 | 4 mo | HOM |
| **1011937543** | 1 | ILA6b | 1081 | 0.011155276 | 0 | 0 | 0 | 0 | 8 mo | HOM |
| **1010419538** | 2 | ILA6b | 1081 | 0.011155276 | 0 | 0 | 0 | 0 | 4 mo | WILD |
| **1010419187** | 1 | ILA6b | 1081 | 0.011155276 | 7.01E-06 | 1 | 0.000628673 | 89.6436783 | 8 mo | HOM |
| **1013978366** | 1 | ILA6b | 1081 | 0.011155276 | 0 | 0 | 0 | 0 | 2 mo | HOM |
| **1014416155** | 2 | ILA6b | 1081 | 0.011155276 | 0 | 0 | 0 | 0 | 2 mo | HOM |
| **1013978065** | 2 | ILA6b | 1081 | 0.011155276 | 0 | 0 | 0 | 0 | 2 mo | HOM |
| **1009934102** | 2 | ILA6b | 1081 | 0.011155276 | 0 | 0 | 0 | 0 | 4 mo | WILD |
| **1009322937** | 2 | ILA6b | 1081 | 0.011155276 | 0 | 0 | 0 | 0 | 8 mo | WILD |
| **1011080637** | 2 | ILA6b | 1081 | 0.011155276 | 0 | 0 | 0 | 0 | 8 mo | HOM |
| **1013609713** | 2 | ILA6b | 1081 | 0.011155276 | 0 | 0 | 0 | 0 | 2 mo | WILD |
| **1010418842** | 1 | ILA6b | 1081 | 0.011155276 | 0 | 0 | 0 | 0 | 8 mo | HOM |
| **1013288213** | 2 | ILA6b | 1081 | 0.011155276 | 0 | 0 | 0 | 0 | 4 mo | HOM |
| **1008976060** | 2 | ILA6b | 1081 | 0.011155276 | 0 | 0 | 0 | 0 | 8 mo | WILD |
| **1013446243** | 1 | ILA6b | 1081 | 0.011155276 | 0 | 0 | 0 | 0 | 4 mo | HOM |
| **1013288951** | 2 | ILA6b | 1081 | 0.011155276 | 2.91E-12 | 0 | 2.61E-10 | 0 | 4 mo | HOM |
| **1013978665** | 1 | mfbse | 1083 | 0.500804735 | 4.27E-05 | 0 | 8.53E-05 | 0 | 2 mo | HOM |
| **1013445609** | 1 | mfbse | 1083 | 0.500804735 | 1.49E-05 | 2 | 2.98E-05 | 3.993572463 | 4 mo | HOM |
| **1013609015** | 1 | mfbse | 1083 | 0.500804735 | 7.85E-05 | 1 | 0.00015683 | 1.996786232 | 4 mo | HOM |
| **1013608106** | 2 | mfbse | 1083 | 0.500804735 | 0 | 0 | 0 | 0 | 4 mo | HOM |

| | | | | | | | | | | |
|---|---|---|---|---|---|---|---|---|---|---|
| **1011937543** | 1 | mfbse | 1083 | 0.500804735 | 0.000256606 | 4 | 0.000512387 | 7.987144927 | 8 mo | HOM |
| **1010419538** | 2 | mfbse | 1083 | 0.500804735 | 3.14E-05 | 2 | 6.28E-05 | 3.993572463 | 4 mo | WILD |
| **1010419187** | 1 | mfbse | 1083 | 0.500804735 | 0.000527295 | 10 | 0.001052895 | 19.96786232 | 8 mo | HOM |
| **1013978366** | 1 | mfbse | 1083 | 0.500804735 | 4.12E-05 | 2 | 8.23E-05 | 3.993572463 | 2 mo | HOM |
| **1014416155** | 2 | mfbse | 1083 | 0.500804735 | 0 | 0 | 0 | 0 | 2 mo | HOM |
| **1013978065** | 2 | mfbse | 1083 | 0.500804735 | 0 | 0 | 0 | 0 | 2 mo | HOM |
| **1009934102** | 2 | mfbse | 1083 | 0.500804735 | 0 | 0 | 0 | 0 | 4 mo | WILD |
| **1009322937** | 2 | mfbse | 1083 | 0.500804735 | 3.15E-08 | 0 | 6.28E-08 | 0 | 8 mo | WILD |
| **1011080637** | 2 | mfbse | 1083 | 0.500804735 | 0.000339071 | 5 | 0.000677052 | 9.983931158 | 8 mo | HOM |
| **1013609713** | 2 | mfbse | 1083 | 0.500804735 | 0 | 0 | 0 | 0 | 2 mo | WILD |
| **1010418842** | 1 | mfbse | 1083 | 0.500804735 | 0.000333803 | 6 | 0.000666533 | 11.98071739 | 8 mo | HOM |
| **1013288213** | 2 | mfbse | 1083 | 0.500804735 | 1.75E-11 | 0 | 3.49E-11 | 0 | 4 mo | HOM |
| **1008976060** | 2 | mfbse | 1083 | 0.500804735 | 9.82E-05 | 1 | 0.000196169 | 1.996786232 | 8 mo | WILD |
| **1013446243** | 1 | mfbse | 1083 | 0.500804735 | 4.11E-05 | 2 | 8.22E-05 | 3.993572463 | 4 mo | HOM |
| **1013288951** | 2 | mfbse | 1083 | 0.500804735 | 0.000542161 | 3 | 0.00108258 | 5.990358695 | 4 mo | HOM |
| **1013978665** | 1 | PRE | 1084 | 0.944396992 | 0 | 0 | 0 | 0 | 2 mo | HOM |
| **1013445609** | 1 | PRE | 1084 | 0.944396992 | 5.95E-06 | 1 | 6.3E-06 | 1.058876731 | 4 mo | HOM |
| **1013609015** | 1 | PRE | 1084 | 0.944396992 | 3.35E-05 | 2 | 3.55E-05 | 2.117753463 | 4 mo | HOM |
| **1013608106** | 2 | PRE | 1084 | 0.944396992 | 0 | 0 | 0 | 0 | 4 mo | HOM |
| **1011937543** | 1 | PRE | 1084 | 0.944396992 | 0.001359066 | 13 | 0.001439083 | 13.76539751 | 8 mo | HOM |
| **1010419538** | 2 | PRE | 1084 | 0.944396992 | 4.29E-07 | 0 | 4.54E-07 | 0 | 4 mo | WILD |
| **1010419187** | 1 | PRE | 1084 | 0.944396992 | 0.001741194 | 37 | 0.00184371 | 39.17843906 | 8 mo | HOM |
| **1013978366** | 1 | PRE | 1084 | 0.944396992 | 0 | 0 | 0 | 0 | 2 mo | HOM |
| **1014416155** | 2 | PRE | 1084 | 0.944396992 | 0 | 0 | 0 | 0 | 2 mo | HOM |
| **1013978065** | 2 | PRE | 1084 | 0.944396992 | 0 | 0 | 0 | 0 | 2 mo | HOM |
| **1009934102** | 2 | PRE | 1084 | 0.944396992 | 0.000114896 | 1 | 0.000121661 | 1.058876731 | 4 mo | WILD |
| **1009322937** | 2 | PRE | 1084 | 0.944396992 | 4.28E-06 | 0 | 4.53E-06 | 0 | 8 mo | WILD |
| **1011080637** | 2 | PRE | 1084 | 0.944396992 | 0.002113853 | 16 | 0.002238309 | 16.9420277 | 8 mo | HOM |
| **1013609713** | 2 | PRE | 1084 | 0.944396992 | 4.02E-06 | 1 | 4.25E-06 | 1.058876731 | 2 mo | WILD |
| **1010418842** | 1 | PRE | 1084 | 0.944396992 | 0.001133777 | 31 | 0.00120053 | 32.82517867 | 8 mo | HOM |
| **1013288213** | 2 | PRE | 1084 | 0.944396992 | 0 | 0 | 0 | 0 | 4 mo | HOM |
| **1008976060** | 2 | PRE | 1084 | 0.944396992 | 0.00015624 | 6 | 0.000165439 | 6.353260388 | 8 mo | WILD |
| **1013446243** | 1 | PRE | 1084 | 0.944396992 | 8.94E-05 | 3 | 9.46E-05 | 3.176630194 | 4 mo | HOM |
| **1013288951** | 2 | PRE | 1084 | 0.944396992 | 7.83E-05 | 4 | 8.29E-05 | 4.235506925 | 4 mo | HOM |
| **1013978665** | 1 | MOs6b | 1085 | 0.075810134 | 0 | 0 | 0 | 0 | 2 mo | HOM |
| **1013445609** | 1 | MOs6b | 1085 | 0.075810134 | 0 | 0 | 0 | 0 | 4 mo | HOM |
| **1013609015** | 1 | MOs6b | 1085 | 0.075810134 | 0 | 0 | 0 | 0 | 4 mo | HOM |
| **1013608106** | 2 | MOs6b | 1085 | 0.075810134 | 0 | 0 | 0 | 0 | 4 mo | HOM |
| **1011937543** | 1 | MOs6b | 1085 | 0.075810134 | 5.16E-05 | 2 | 0.000681151 | 26.38169706 | 8 mo | HOM |
| **1010419538** | 2 | MOs6b | 1085 | 0.075810134 | 0 | 0 | 0 | 0 | 4 mo | WILD |
| **1010419187** | 1 | MOs6b | 1085 | 0.075810134 | 3.5E-05 | 2 | 0.000461076 | 26.38169706 | 8 mo | HOM |
| **1013978366** | 1 | MOs6b | 1085 | 0.075810134 | 0 | 0 | 0 | 0 | 2 mo | HOM |
| **1014416155** | 2 | MOs6b | 1085 | 0.075810134 | 0 | 0 | 0 | 0 | 2 mo | HOM |
| **1013978065** | 2 | MOs6b | 1085 | 0.075810134 | 0 | 0 | 0 | 0 | 2 mo | HOM |
| **1009934102** | 2 | MOs6b | 1085 | 0.075810134 | 0 | 0 | 0 | 0 | 4 mo | WILD |
| **1009322937** | 2 | MOs6b | 1085 | 0.075810134 | 0 | 0 | 0 | 0 | 8 mo | WILD |
| **1011080637** | 2 | MOs6b | 1085 | 0.075810134 | 7.73E-06 | 1 | 0.000101991 | 13.19084853 | 8 mo | HOM |
| **1013609713** | 2 | MOs6b | 1085 | 0.075810134 | 0 | 0 | 0 | 0 | 2 mo | WILD |
| **1010418842** | 1 | MOs6b | 1085 | 0.075810134 | 1.93E-05 | 2 | 0.000254517 | 26.38169706 | 8 mo | HOM |
| **1013288213** | 2 | MOs6b | 1085 | 0.075810134 | 0 | 0 | 0 | 0 | 4 mo | HOM |
| **1008976060** | 2 | MOs6b | 1085 | 0.075810134 | 0 | 0 | 0 | 0 | 8 mo | WILD |
| **1013446243** | 1 | MOs6b | 1085 | 0.075810134 | 0 | 0 | 0 | 0 | 4 mo | HOM |
| **1013288951** | 2 | MOs6b | 1085 | 0.075810134 | 1.28E-05 | 0 | 0.000169083 | 0 | 4 mo | HOM |
| **1013978665** | 1 | SSp-tr4 | 1086 | 0.157362745 | 0 | 0 | 0 | 0 | 2 mo | HOM |
| **1013445609** | 1 | SSp-tr4 | 1086 | 0.157362745 | 2.98E-05 | 2 | 0.000189107 | 12.70948848 | 4 mo | HOM |
| **1013609015** | 1 | SSp-tr4 | 1086 | 0.157362745 | 2.96E-05 | 0 | 0.000188239 | 0 | 4 mo | HOM |
| **1013608106** | 2 | SSp-tr4 | 1086 | 0.157362745 | 0 | 0 | 0 | 0 | 4 mo | HOM |
| **1011937543** | 1 | SSp-tr4 | 1086 | 0.157362745 | 0.000215418 | 3 | 0.001368927 | 19.06423272 | 8 mo | HOM |
| **1010419538** | 2 | SSp-tr4 | 1086 | 0.157362745 | 0 | 0 | 0 | 0 | 4 mo | WILD |

| | | | | | | | | | | |
|---|---|---|---|---|---|---|---|---|---|---|
| **1010419187** | 1 | SSp-tr4 | 1086 | 0.157362745 | 0.000303901 | 7 | 0.001931213 | 44.48320968 | 8 mo | HOM |
| **1013978366** | 1 | SSp-tr4 | 1086 | 0.157362745 | 0 | 0 | 0 | 0 | 2 mo | HOM |
| **1014416155** | 2 | SSp-tr4 | 1086 | 0.157362745 | 0 | 0 | 0 | 0 | 2 mo | HOM |
| **1013978065** | 2 | SSp-tr4 | 1086 | 0.157362745 | 0 | 0 | 0 | 0 | 2 mo | HOM |
| **1009934102** | 2 | SSp-tr4 | 1086 | 0.157362745 | 0 | 0 | 0 | 0 | 4 mo | WILD |
| **1009322937** | 2 | SSp-tr4 | 1086 | 0.157362745 | 0 | 0 | 0 | 0 | 8 mo | WILD |
| **1011080637** | 2 | SSp-tr4 | 1086 | 0.157362745 | 0.000174319 | 8 | 0.001107752 | 50.83795392 | 8 mo | HOM |
| **1013609713** | 2 | SSp-tr4 | 1086 | 0.157362745 | 0 | 0 | 0 | 0 | 2 mo | WILD |
| **1010418842** | 1 | SSp-tr4 | 1086 | 0.157362745 | 0.000155055 | 5 | 0.000985333 | 31.7737212 | 8 mo | HOM |
| **1013288213** | 2 | SSp-tr4 | 1086 | 0.157362745 | 0 | 0 | 0 | 0 | 4 mo | HOM |
| **1008976060** | 2 | SSp-tr4 | 1086 | 0.157362745 | 0 | 0 | 0 | 0 | 8 mo | WILD |
| **1013446243** | 1 | SSp-tr4 | 1086 | 0.157362745 | 0.00012238 | 6 | 0.000777697 | 38.12846544 | 4 mo | HOM |
| **1013288951** | 2 | SSp-tr4 | 1086 | 0.157362745 | 1.11E-05 | 2 | 7.04E-05 | 12.70948848 | 4 mo | HOM |
| **1013978665** | 1 | MGm | 1088 | 0.257875612 | 0 | 0 | 0 | 0 | 2 mo | HOM |
| **1013445609** | 1 | MGm | 1088 | 0.257875612 | 0 | 0 | 0 | 0 | 4 mo | HOM |
| **1013609015** | 1 | MGm | 1088 | 0.257875612 | 0 | 0 | 0 | 0 | 4 mo | HOM |
| **1013608106** | 2 | MGm | 1088 | 0.257875612 | 0 | 0 | 0 | 0 | 4 mo | HOM |
| **1011937543** | 1 | MGm | 1088 | 0.257875612 | 0.000188923 | 5 | 0.000732612 | 19.38919293 | 8 mo | HOM |
| **1010419538** | 2 | MGm | 1088 | 0.257875612 | 0 | 0 | 0 | 0 | 4 mo | WILD |
| **1010419187** | 1 | MGm | 1088 | 0.257875612 | 7.01E-06 | 1 | 2.72E-05 | 3.877838585 | 8 mo | HOM |
| **1013978366** | 1 | MGm | 1088 | 0.257875612 | 0 | 0 | 0 | 0 | 2 mo | HOM |
| **1014416155** | 2 | MGm | 1088 | 0.257875612 | 0 | 0 | 0 | 0 | 2 mo | HOM |
| **1013978065** | 2 | MGm | 1088 | 0.257875612 | 0 | 0 | 0 | 0 | 2 mo | HOM |
| **1009934102** | 2 | MGm | 1088 | 0.257875612 | 0 | 0 | 0 | 0 | 4 mo | WILD |
| **1009322937** | 2 | MGm | 1088 | 0.257875612 | 0 | 0 | 0 | 0 | 8 mo | WILD |
| **1011080637** | 2 | MGm | 1088 | 0.257875612 | 2.15E-05 | 1 | 8.35E-05 | 3.877838585 | 8 mo | HOM |
| **1013609713** | 2 | MGm | 1088 | 0.257875612 | 0 | 0 | 0 | 0 | 2 mo | WILD |
| **1010418842** | 1 | MGm | 1088 | 0.257875612 | 0.000140145 | 5 | 0.000543461 | 19.38919293 | 8 mo | HOM |
| **1013288213** | 2 | MGm | 1088 | 0.257875612 | 0 | 0 | 0 | 0 | 4 mo | HOM |
| **1008976060** | 2 | MGm | 1088 | 0.257875612 | 1.7E-06 | 0 | 6.6E-06 | 0 | 8 mo | WILD |
| **1013446243** | 1 | MGm | 1088 | 0.257875612 | 0 | 0 | 0 | 0 | 4 mo | HOM |
| **1013288951** | 2 | MGm | 1088 | 0.257875612 | 0 | 0 | 0 | 0 | 4 mo | HOM |
| **1013978665** | 1 | HPF | 1089 | 43.9299159 | 0.000150594 | 11 | 3.43E-06 | 0.25039884 | 2 mo | HOM |
| **1013445609** | 1 | HPF | 1089 | 43.9299159 | 0.001261256 | 99 | 2.87E-05 | 2.253589563 | 4 mo | HOM |
| **1013609015** | 1 | HPF | 1089 | 43.9299159 | 0.007800154 | 255 | 0.000177559 | 5.80470039 | 4 mo | HOM |
| **1013608106** | 2 | HPF | 1089 | 43.9299159 | 0 | 0 | 0 | 0 | 4 mo | HOM |
| **1011937543** | 1 | HPF | 1089 | 43.9299159 | 0.064732004 | 764 | 0.001473529 | 17.39133764 | 8 mo | HOM |
| **1010419538** | 2 | HPF | 1089 | 43.9299159 | 0.000545028 | 42 | 1.24E-05 | 0.9560683 | 4 mo | WILD |
| **1010419187** | 1 | HPF | 1089 | 43.9299159 | 0.079173543 | 1764 | 0.001802269 | 40.15486858 | 8 mo | HOM |
| **1013978366** | 1 | HPF | 1089 | 43.9299159 | 0.001468337 | 43 | 3.34E-05 | 0.978831831 | 2 mo | HOM |
| **1014416155** | 2 | HPF | 1089 | 43.9299159 | 0 | 0 | 0 | 0 | 2 mo | HOM |
| **1013978065** | 2 | HPF | 1089 | 43.9299159 | 3.8E-05 | 5 | 8.65E-07 | 0.113817655 | 2 mo | HOM |
| **1009934102** | 2 | HPF | 1089 | 43.9299159 | 0.001820403 | 68 | 4.14E-05 | 1.547920104 | 4 mo | WILD |
| **1009322937** | 2 | HPF | 1089 | 43.9299159 | 0.000831202 | 37 | 1.89E-05 | 0.842250645 | 8 mo | WILD |
| **1011080637** | 2 | HPF | 1089 | 43.9299159 | 0.044226351 | 1364 | 0.001006748 | 31.04945621 | 8 mo | HOM |
| **1013609713** | 2 | HPF | 1089 | 43.9299159 | 0.000417063 | 20 | 9.49E-06 | 0.455270619 | 2 mo | WILD |
| **1010418842** | 1 | HPF | 1089 | 43.9299159 | 0.061753656 | 1689 | 0.001405731 | 38.44760376 | 8 mo | HOM |
| **1013288213** | 2 | HPF | 1089 | 43.9299159 | 0.004838653 | 142 | 0.000110145 | 3.232421394 | 4 mo | HOM |
| **1008976060** | 2 | HPF | 1089 | 43.9299159 | 0.001291057 | 123 | 2.94E-05 | 2.799914306 | 8 mo | WILD |
| **1013446243** | 1 | HPF | 1089 | 43.9299159 | 0.003987468 | 224 | 9.08E-05 | 5.099030931 | 4 mo | HOM |
| **1013288951** | 2 | HPF | 1089 | 43.9299159 | 0.011148972 | 281 | 0.00025379 | 6.396552195 | 4 mo | HOM |
| **1013978665** | 1 | SSs5 | 1090 | 2.251573206 | 0 | 0 | 0 | 0 | 2 mo | HOM |
| **1013445609** | 1 | SSs5 | 1090 | 2.251573206 | 0.000228792 | 15 | 0.000101614 | 6.662008571 | 4 mo | HOM |
| **1013609015** | 1 | SSs5 | 1090 | 2.251573206 | 0.000396499 | 20 | 0.000176099 | 8.882678095 | 4 mo | HOM |
| **1013608106** | 2 | SSs5 | 1090 | 2.251573206 | 0 | 0 | 0 | 0 | 4 mo | HOM |
| **1011937543** | 1 | SSs5 | 1090 | 2.251573206 | 0.002653721 | 74 | 0.001178608 | 32.86590895 | 8 mo | HOM |
| **1010419538** | 2 | SSs5 | 1090 | 2.251573206 | 2.75E-05 | 1 | 1.22E-05 | 0.444133905 | 4 mo | WILD |
| **1010419187** | 1 | SSs5 | 1090 | 2.251573206 | 0.002103698 | 105 | 0.000934324 | 46.63406 | 8 mo | HOM |
| **1013978366** | 1 | SSs5 | 1090 | 2.251573206 | 1.43E-05 | 2 | 6.34E-06 | 0.88826781 | 2 mo | HOM |

| | | | | | | | | | | |
|---|---|---|---|---|---|---|---|---|---|---|
| 1014416155 | 2 | SSs5 | 1090 | 2.251573206 | 0 | 0 | 0 | 0 | 2 mo | HOM |
| 1013978065 | 2 | SSs5 | 1090 | 2.251573206 | 0 | 0 | 0 | 0 | 2 mo | HOM |
| 1009934102 | 2 | SSs5 | 1090 | 2.251573206 | 0 | 0 | 0 | 0 | 4 mo | WILD |
| 1009322937 | 2 | SSs5 | 1090 | 2.251573206 | 0 | 0 | 0 | 0 | 8 mo | WILD |
| 1011080637 | 2 | SSs5 | 1090 | 2.251573206 | 0.002548253 | 90 | 0.001131766 | 39.97205143 | 8 mo | HOM |
| 1013609713 | 2 | SSs5 | 1090 | 2.251573206 | 0 | 0 | 0 | 0 | 2 mo | WILD |
| 1010418842 | 1 | SSs5 | 1090 | 2.251573206 | 0.002102173 | 83 | 0.000933646 | 36.86311409 | 8 mo | HOM |
| 1013288213 | 2 | SSs5 | 1090 | 2.251573206 | 7.9E-05 | 7 | 3.51E-05 | 3.108937333 | 4 mo | HOM |
| 1008976060 | 2 | SSs5 | 1090 | 2.251573206 | 3.77E-06 | 1 | 1.68E-06 | 0.444133905 | 8 mo | WILD |
| 1013446243 | 1 | SSs5 | 1090 | 2.251573206 | 0.000479334 | 25 | 0.000212888 | 11.10334762 | 4 mo | HOM |
| 1013288951 | 2 | SSs5 | 1090 | 2.251573206 | 0.000477117 | 21 | 0.000211904 | 9.326812 | 4 mo | HOM |
| 1013978665 | 1 | CUL4, 5 | 1091 | 6.929062874 | 0.000729491 | 26 | 0.00010528 | 3.752311167 | 2 mo | HOM |
| 1013445609 | 1 | CUL4, 5 | 1091 | 6.929062874 | 0.000132449 | 3 | 1.91E-05 | 0.432958981 | 4 mo | HOM |
| 1013609015 | 1 | CUL4, 5 | 1091 | 6.929062874 | 0.002160155 | 51 | 0.000311753 | 7.360302674 | 4 mo | HOM |
| 1013608106 | 2 | CUL4, 5 | 1091 | 6.929062874 | 0 | 0 | 0 | 0 | 4 mo | HOM |
| 1011937543 | 1 | CUL4, 5 | 1091 | 6.929062874 | 0.00736756 | 47 | 0.001063284 | 6.783024033 | 8 mo | HOM |
| 1010419538 | 2 | CUL4, 5 | 1091 | 6.919467341 | 0.000641261 | 18 | 9.27E-05 | 2.601356306 | 4 mo | WILD |
| 1010419187 | 1 | CUL4, 5 | 1091 | 6.929062874 | 0.005305846 | 122 | 0.000765738 | 17.60699855 | 8 mo | HOM |
| 1013978366 | 1 | CUL4, 5 | 1091 | 6.929062874 | 0.000597636 | 25 | 8.63E-05 | 3.607991507 | 2 mo | HOM |
| 1014416155 | 2 | CUL4, 5 | 1091 | 6.929062874 | 0 | 0 | 0 | 0 | 2 mo | HOM |
| 1013978065 | 2 | CUL4, 5 | 1091 | 6.929062874 | 0.000243308 | 6 | 3.51E-05 | 0.865917962 | 2 mo | HOM |
| 1009934102 | 2 | CUL4, 5 | 1091 | 6.929062874 | 0.00038076 | 12 | 5.5E-05 | 1.731835923 | 4 mo | WILD |
| 1009322937 | 2 | CUL4, 5 | 1091 | 6.929062874 | 0.001061446 | 41 | 0.000153187 | 5.917106072 | 8 mo | WILD |
| 1011080637 | 2 | CUL4, 5 | 1091 | 6.929062874 | 0.009674886 | 136 | 0.001396276 | 19.6274738 | 8 mo | HOM |
| 1013609713 | 2 | CUL4, 5 | 1091 | 6.929062874 | 0.000118733 | 3 | 1.71E-05 | 0.432958981 | 2 mo | WILD |
| 1010418842 | 1 | CUL4, 5 | 1091 | 6.929062874 | 0.015365877 | 182 | 0.002217598 | 26.26617817 | 8 mo | HOM |
| 1013288213 | 2 | CUL4, 5 | 1091 | 6.929062874 | 0.000307292 | 11 | 4.43E-05 | 1.587516263 | 4 mo | HOM |
| 1008976060 | 2 | CUL4, 5 | 1091 | 6.929062874 | 0.000940445 | 71 | 0.000135725 | 10.24669588 | 8 mo | WILD |
| 1013446243 | 1 | CUL4, 5 | 1091 | 6.929062874 | 0.000145307 | 16 | 2.1E-05 | 2.309114565 | 4 mo | HOM |
| 1013288951 | 2 | CUL4, 5 | 1091 | 6.929062874 | 0.002316034 | 34 | 0.000334249 | 4.90686845 | 4 mo | HOM |
| 1013978665 | 1 | em | 1092 | 0.11397791 | 0 | 0 | 0 | 0 | 2 mo | HOM |
| 1013445609 | 1 | em | 1092 | 0.11397791 | 0 | 0 | 0 | 0 | 4 mo | HOM |
| 1013609015 | 1 | em | 1092 | 0.11397791 | 0 | 0 | 0 | 0 | 4 mo | HOM |
| 1013608106 | 2 | em | 1092 | 0.11397791 | 0 | 0 | 0 | 0 | 4 mo | HOM |
| 1011937543 | 1 | em | 1092 | 0.11397791 | 2.82E-05 | 0 | 0.000247105 | 0 | 8 mo | HOM |
| 1010419538 | 2 | em | 1092 | 0.11397791 | 0 | 0 | 0 | 0 | 4 mo | WILD |
| 1010419187 | 1 | em | 1092 | 0.11397791 | 2.28E-05 | 1 | 0.000200104 | 8.773629892 | 8 mo | HOM |
| 1013978366 | 1 | em | 1092 | 0.11397791 | 0 | 0 | 0 | 0 | 2 mo | HOM |
| 1014416155 | 2 | em | 1092 | 0.11397791 | 0 | 0 | 0 | 0 | 2 mo | HOM |
| 1013978065 | 2 | em | 1092 | 0.11397791 | 0 | 0 | 0 | 0 | 2 mo | HOM |
| 1009934102 | 2 | em | 1092 | 0.11397791 | 0 | 0 | 0 | 0 | 4 mo | WILD |
| 1009322937 | 2 | em | 1092 | 0.11397791 | 0 | 0 | 0 | 0 | 8 mo | WILD |
| 1011080637 | 2 | em | 1092 | 0.11397791 | 1.76E-05 | 2 | 0.000154403 | 17.54725978 | 8 mo | HOM |
| 1013609713 | 2 | em | 1092 | 0.11397791 | 0 | 0 | 0 | 0 | 2 mo | WILD |
| 1010418842 | 1 | em | 1092 | 0.11397791 | 3.8E-05 | 1 | 0.000333825 | 8.773629892 | 8 mo | HOM |
| 1013288213 | 2 | em | 1092 | 0.11397791 | 9.92E-07 | 0 | 8.71E-06 | 0 | 4 mo | HOM |
| 1008976060 | 2 | em | 1092 | 0.11397791 | 2.81E-06 | 1 | 2.46E-05 | 8.773629892 | 8 mo | WILD |
| 1013446243 | 1 | em | 1092 | 0.11397791 | 0 | 0 | 0 | 0 | 4 mo | HOM |
| 1013288951 | 2 | em | 1092 | 0.11397791 | 0 | 0 | 0 | 0 | 4 mo | HOM |
| 1013978665 | 1 | PRNc | 1093 | 2.420047907 | 9.14E-06 | 1 | 3.78E-06 | 0.41321496 | 2 mo | HOM |
| 1013445609 | 1 | PRNc | 1093 | 2.420047907 | 0 | 0 | 0 | 0 | 4 mo | HOM |
| 1013609015 | 1 | PRNc | 1093 | 2.420047907 | 0 | 0 | 0 | 0 | 4 mo | HOM |
| 1013608106 | 2 | PRNc | 1093 | 2.420047907 | 0 | 0 | 0 | 0 | 4 mo | HOM |
| 1011937543 | 1 | PRNc | 1093 | 2.420047907 | 1.9E-05 | 1 | 7.84E-06 | 0.41321496 | 8 mo | HOM |
| 1010419538 | 2 | PRNc | 1093 | 2.420047907 | 0 | 0 | 0 | 0 | 4 mo | WILD |
| 1010419187 | 1 | PRNc | 1093 | 2.420047907 | 0 | 0 | 0 | 0 | 8 mo | HOM |
| 1013978366 | 1 | PRNc | 1093 | 2.420047907 | 0 | 0 | 0 | 0 | 2 mo | HOM |
| 1014416155 | 2 | PRNc | 1093 | 2.420047907 | 0 | 0 | 0 | 0 | 2 mo | HOM |
| 1013978065 | 2 | PRNc | 1093 | 2.420047907 | 0 | 0 | 0 | 0 | 2 mo | HOM |

| | | | | | | | | | | |
|---|---|---|---|---|---|---|---|---|---|---|
| **1009934102** | 2 | PRNc | 1093 | 2.420047907 | 0 | 0 | 0 | 0 | 4 mo | WILD |
| **1009322937** | 2 | PRNc | 1093 | 2.420047907 | 0 | 0 | 0 | 0 | 8 mo | WILD |
| **1011080637** | 2 | PRNc | 1093 | 2.420047907 | 1.21E-05 | 1 | 5.01E-06 | 0.41321496 | 8 mo | HOM |
| **1013609713** | 2 | PRNc | 1093 | 2.420047907 | 0 | 0 | 0 | 0 | 2 mo | WILD |
| **1010418842** | 1 | PRNc | 1093 | 2.420047907 | 0.000398073 | 30 | 0.00016449 | 12.39644881 | 8 mo | HOM |
| **1013288213** | 2 | PRNc | 1093 | 2.420047907 | 0 | 0 | 0 | 0 | 4 mo | HOM |
| **1008976060** | 2 | PRNc | 1093 | 2.420047907 | 0.000430735 | 27 | 0.000177986 | 11.15680393 | 8 mo | WILD |
| **1013446243** | 1 | PRNc | 1093 | 2.420047907 | 0 | 0 | 0 | 0 | 4 mo | HOM |
| **1013288951** | 2 | PRNc | 1093 | 2.420047907 | 0 | 0 | 0 | 0 | 4 mo | HOM |
| **1013978665** | 1 | SSp-II4 | 1094 | 0.281070115 | 0 | 0 | 0 | 0 | 2 mo | HOM |
| **1013445609** | 1 | SSp-II4 | 1094 | 0.281070115 | 4.43E-05 | 3 | 0.000157442 | 10.67349333 | 4 mo | HOM |
| **1013609015** | 1 | SSp-II4 | 1094 | 0.281070115 | 0.000157486 | 5 | 0.000560307 | 17.78915555 | 4 mo | HOM |
| **1013608106** | 2 | SSp-II4 | 1094 | 0.281070115 | 0 | 0 | 0 | 0 | 4 mo | HOM |
| **1011937543** | 1 | SSp-II4 | 1094 | 0.281070115 | 0.000593083 | 7 | 0.002110088 | 24.90481777 | 8 mo | HOM |
| **1010419538** | 2 | SSp-II4 | 1094 | 0.281070115 | 0 | 0 | 0 | 0 | 4 mo | WILD |
| **1010419187** | 1 | SSp-II4 | 1094 | 0.281070115 | 0.000540455 | 13 | 0.001922848 | 46.25180443 | 8 mo | HOM |
| **1013978366** | 1 | SSp-II4 | 1094 | 0.281070115 | 0 | 0 | 0 | 0 | 2 mo | HOM |
| **1014416155** | 2 | SSp-II4 | 1094 | 0.281070115 | 0 | 0 | 0 | 0 | 2 mo | HOM |
| **1013978065** | 2 | SSp-II4 | 1094 | 0.281070115 | 0 | 0 | 0 | 0 | 2 mo | HOM |
| **1009934102** | 2 | SSp-II4 | 1094 | 0.281070115 | 0 | 0 | 0 | 0 | 4 mo | WILD |
| **1009322937** | 2 | SSp-II4 | 1094 | 0.281070115 | 0 | 0 | 0 | 0 | 8 mo | WILD |
| **1011080637** | 2 | SSp-II4 | 1094 | 0.281070115 | 0.000229371 | 9 | 0.000816065 | 32.02047999 | 8 mo | HOM |
| **1013609713** | 2 | SSp-II4 | 1094 | 0.281070115 | 0 | 0 | 0 | 0 | 2 mo | WILD |
| **1010418842** | 1 | SSp-II4 | 1094 | 0.281070115 | 0.000440243 | 20 | 0.001566309 | 71.1566222 | 8 mo | HOM |
| **1013288213** | 2 | SSp-II4 | 1094 | 0.281070115 | 9.17E-06 | 1 | 3.26E-05 | 3.55783111 | 4 mo | HOM |
| **1008976060** | 2 | SSp-II4 | 1094 | 0.281070115 | 0 | 0 | 0 | 0 | 8 mo | WILD |
| **1013446243** | 1 | SSp-II4 | 1094 | 0.281070115 | 0.000134414 | 6 | 0.000478224 | 21.34698666 | 4 mo | HOM |
| **1013288951** | 2 | SSp-II4 | 1094 | 0.281070115 | 8.28E-05 | 5 | 0.00029448 | 17.78915555 | 4 mo | HOM |
| **1013978665** | 1 | AMd | 1096 | 0.251653063 | 0 | 0 | 0 | 0 | 2 mo | HOM |
| **1013445609** | 1 | AMd | 1096 | 0.251653063 | 0 | 0 | 0 | 0 | 4 mo | HOM |
| **1013609015** | 1 | AMd | 1096 | 0.251653063 | 0 | 0 | 0 | 0 | 4 mo | HOM |
| **1013608106** | 2 | AMd | 1096 | 0.251653063 | 0 | 0 | 0 | 0 | 4 mo | HOM |
| **1011937543** | 1 | AMd | 1096 | 0.251653063 | 0.000139089 | 4 | 0.0005527 | 15.89489891 | 8 mo | HOM |
| **1010419538** | 2 | AMd | 1096 | 0.251653063 | 0 | 0 | 0 | 0 | 4 mo | WILD |
| **1010419187** | 1 | AMd | 1096 | 0.251653063 | 0.000112802 | 3 | 0.000448245 | 11.92117418 | 8 mo | HOM |
| **1013978366** | 1 | AMd | 1096 | 0.251653063 | 0 | 0 | 0 | 0 | 2 mo | HOM |
| **1014416155** | 2 | AMd | 1096 | 0.251653063 | 0 | 0 | 0 | 0 | 2 mo | HOM |
| **1013978065** | 2 | AMd | 1096 | 0.251653063 | 0 | 0 | 0 | 0 | 2 mo | HOM |
| **1009934102** | 2 | AMd | 1096 | 0.251653063 | 0 | 0 | 0 | 0 | 4 mo | WILD |
| **1009322937** | 2 | AMd | 1096 | 0.251653063 | 2.58E-06 | 0 | 1.02E-05 | 0 | 8 mo | WILD |
| **1011080637** | 2 | AMd | 1096 | 0.251653063 | 0.00026347 | 4 | 0.001046958 | 15.89489891 | 8 mo | HOM |
| **1013609713** | 2 | AMd | 1096 | 0.251653063 | 0 | 0 | 0 | 0 | 2 mo | WILD |
| **1010418842** | 1 | AMd | 1096 | 0.251653063 | 9.07E-05 | 5 | 0.000360413 | 19.86862364 | 8 mo | HOM |
| **1013288213** | 2 | AMd | 1096 | 0.251653063 | 1.08E-06 | 0 | 4.3E-06 | 0 | 4 mo | HOM |
| **1008976060** | 2 | AMd | 1096 | 0.251653063 | 0 | 0 | 0 | 0 | 8 mo | WILD |
| **1013446243** | 1 | AMd | 1096 | 0.251653063 | 7.46E-05 | 1 | 0.000296278 | 3.973724728 | 4 mo | HOM |
| **1013288951** | 2 | AMd | 1096 | 0.251653063 | 0 | 0 | 0 | 0 | 4 mo | HOM |
| **1013978665** | 1 | HY | 1097 | 15.66168952 | 4.06E-05 | 2 | 2.59E-06 | 0.127700144 | 2 mo | HOM |
| **1013445609** | 1 | HY | 1097 | 15.66168952 | 0.000208083 | 7 | 1.33E-05 | 0.446950502 | 4 mo | HOM |
| **1013609015** | 1 | HY | 1097 | 15.66168952 | 0.000813267 | 11 | 5.19E-05 | 0.702350789 | 4 mo | HOM |
| **1013608106** | 2 | HY | 1097 | 15.66169078 | 0 | 0 | 0 | 0 | 4 mo | HOM |
| **1011937543** | 1 | HY | 1097 | 15.66168952 | 0.020584971 | 209 | 0.001314352 | 13.344665 | 8 mo | HOM |
| **1010419538** | 2 | HY | 1097 | 15.66169078 | 1.53E-05 | 3 | 9.78E-07 | 0.1915502 | 4 mo | WILD |
| **1010419187** | 1 | HY | 1097 | 15.66168952 | 0.006764785 | 102 | 0.000431932 | 6.51270732 | 8 mo | HOM |
| **1013978366** | 1 | HY | 1097 | 15.66168952 | 0.000282313 | 4 | 1.8E-05 | 0.255400287 | 2 mo | HOM |
| **1014416155** | 2 | HY | 1097 | 15.66169078 | 0 | 0 | 0 | 0 | 2 mo | HOM |
| **1013978065** | 2 | HY | 1097 | 15.66169078 | 4.04E-05 | 3 | 2.58E-06 | 0.1915502 | 2 mo | HOM |
| **1009934102** | 2 | HY | 1097 | 15.66169078 | 2.85E-05 | 4 | 1.82E-06 | 0.255400267 | 4 mo | WILD |
| **1009322937** | 2 | HY | 1097 | 15.66169078 | 5.01E-05 | 4 | 3.2E-06 | 0.255400267 | 8 mo | WILD |

| | | | | | | | | | | |
|---|---|---|---|---|---|---|---|---|---|---|
| **1011080637** | 2 | HY | 1097 | 15.66169078 | 0.002926923 | 109 | 0.000186884 | 6.959657265 | 8 mo | HOM |
| **1013609713** | 2 | HY | 1097 | 15.66169078 | 0 | 0 | 0 | 0 | 2 mo | WILD |
| **1010418842** | 1 | HY | 1097 | 15.66168952 | 0.007090203 | 209 | 0.00045271 | 13.344665 | 8 mo | HOM |
| **1013288213** | 2 | HY | 1097 | 15.66169078 | 5.38E-05 | 6 | 3.44E-06 | 0.3831004 | 4 mo | HOM |
| **1008976060** | 2 | HY | 1097 | 15.66169078 | 0.00088339 | 44 | 5.64E-05 | 2.809402933 | 8 mo | WILD |
| **1013446243** | 1 | HY | 1097 | 15.66168952 | 0.000938843 | 4 | 5.99E-05 | 0.255400287 | 4 mo | HOM |
| **1013288951** | 2 | HY | 1097 | 15.66169078 | 0.000475979 | 5 | 3.04E-05 | 0.319250333 | 4 mo | HOM |
| **1013978665** | 1 | MDRNd | 1098 | 1.138567606 | 0 | 0 | 0 | 0 | 2 mo | HOM |
| **1013445609** | 1 | MDRNd | 1098 | 1.138567606 | 0 | 0 | 0 | 0 | 4 mo | HOM |
| **1013609015** | 1 | MDRNd | 1098 | 1.138567606 | 0 | 0 | 0 | 0 | 4 mo | HOM |
| **1013608106** | 2 | MDRNd | 1098 | 1.138567606 | 0 | 0 | 0 | 0 | 4 mo | HOM |
| **1011937543** | 1 | MDRNd | 1098 | 1.138567606 | 0 | 0 | 0 | 0 | 8 mo | HOM |
| **1010419538** | 2 | MDRNd | 1098 | 1.138567606 | 0 | 0 | 0 | 0 | 4 mo | WILD |
| **1010419187** | 1 | MDRNd | 1098 | 1.138567606 | 5.34E-05 | 5 | 4.69E-05 | 4.391482747 | 8 mo | HOM |
| **1013978366** | 1 | MDRNd | 1098 | 1.138567606 | 0 | 0 | 0 | 0 | 2 mo | HOM |
| **1014416155** | 2 | MDRNd | 1098 | 1.138567606 | 0 | 0 | 0 | 0 | 2 mo | HOM |
| **1013978065** | 2 | MDRNd | 1098 | 1.138567606 | 0 | 0 | 0 | 0 | 2 mo | HOM |
| **1009934102** | 2 | MDRNd | 1098 | 1.138567606 | 0 | 0 | 0 | 0 | 4 mo | WILD |
| **1009322937** | 2 | MDRNd | 1098 | 1.138567606 | 0 | 0 | 0 | 0 | 8 mo | WILD |
| **1011080637** | 2 | MDRNd | 1098 | 1.138567606 | 2.04E-06 | 1 | 1.79E-06 | 0.878296549 | 8 mo | HOM |
| **1013609713** | 2 | MDRNd | 1098 | 1.138567606 | 0 | 0 | 0 | 0 | 2 mo | WILD |
| **1010418842** | 1 | MDRNd | 1098 | 1.138567606 | 0.000266233 | 7 | 0.000233831 | 6.148075846 | 8 mo | HOM |
| **1013288213** | 2 | MDRNd | 1098 | 1.138566274 | 0 | 0 | 0 | 0 | 4 mo | HOM |
| **1008976060** | 2 | MDRNd | 1098 | 1.138567606 | 4.33E-06 | 1 | 3.8E-06 | 0.878296549 | 8 mo | WILD |
| **1013446243** | 1 | MDRNd | 1098 | 1.138567606 | 0 | 0 | 0 | 0 | 4 mo | HOM |
| **1013288951** | 2 | MDRNd | 1098 | 1.138567606 | 0 | 0 | 0 | 0 | 4 mo | HOM |
| **1013978665** | 1 | fxs | 1099 | 4.368541301 | 4.87E-09 | 0 | 1.11E-09 | 0 | 2 mo | HOM |
| **1013445609** | 1 | fxs | 1099 | 4.368541301 | 2.12E-05 | 2 | 4.86E-06 | 0.457818723 | 4 mo | HOM |
| **1013609015** | 1 | fxs | 1099 | 4.368541301 | 0.001630021 | 24 | 0.000373127 | 5.493824677 | 4 mo | HOM |
| **1013608106** | 2 | fxs | 1099 | 4.368541301 | 0 | 0 | 0 | 0 | 4 mo | HOM |
| **1011937543** | 1 | fxs | 1099 | 4.368541301 | 0.003358496 | 50 | 0.000768791 | 11.44546808 | 8 mo | HOM |
| **1010419538** | 2 | fxs | 1099 | 4.368541301 | 2.33E-06 | 0 | 5.33E-07 | 0 | 4 mo | WILD |
| **1010419187** | 1 | fxs | 1099 | 4.368541301 | 0.003532021 | 84 | 0.000808513 | 19.22838637 | 8 mo | HOM |
| **1013978366** | 1 | fxs | 1099 | 4.368541301 | 2.28E-05 | 1 | 5.23E-06 | 0.228909362 | 2 mo | HOM |
| **1014416155** | 2 | fxs | 1099 | 4.368541301 | 0 | 0 | 0 | 0 | 2 mo | HOM |
| **1013978065** | 2 | fxs | 1099 | 4.368541301 | 0 | 0 | 0 | 0 | 2 mo | HOM |
| **1009934102** | 2 | fxs | 1099 | 4.368541301 | 8.2E-06 | 0 | 1.88E-06 | 0 | 4 mo | WILD |
| **1009322937** | 2 | fxs | 1099 | 4.368541301 | 6.87E-07 | 0 | 1.57E-07 | 0 | 8 mo | WILD |
| **1011080637** | 2 | fxs | 1099 | 4.368541301 | 0.00109174 | 28 | 0.000249909 | 6.409462123 | 8 mo | HOM |
| **1013609713** | 2 | fxs | 1099 | 4.368541301 | 1.56E-06 | 0 | 3.56E-07 | 0 | 2 mo | WILD |
| **1010418842** | 1 | fxs | 1099 | 4.368541301 | 0.004913963 | 117 | 0.001124852 | 26.7823953 | 8 mo | HOM |
| **1013288213** | 2 | fxs | 1099 | 4.368541301 | 5.04E-05 | 1 | 1.15E-05 | 0.228909362 | 4 mo | HOM |
| **1008976060** | 2 | fxs | 1099 | 4.368541301 | 0.000113038 | 6 | 2.59E-05 | 1.373456169 | 8 mo | WILD |
| **1013446243** | 1 | fxs | 1099 | 4.368541301 | 0.000441864 | 10 | 0.000101147 | 2.289093615 | 4 mo | HOM |
| **1013288951** | 2 | fxs | 1099 | 4.368541301 | 0.000784798 | 7 | 0.000179648 | 1.602365531 | 4 mo | HOM |
| **1013978665** | 1 | PRT | 1100 | 2.142995758 | 0 | 0 | 0 | 0 | 2 mo | HOM |
| **1013445609** | 1 | PRT | 1100 | 2.142995758 | 0 | 0 | 0 | 0 | 4 mo | HOM |
| **1013609015** | 1 | PRT | 1100 | 2.142995758 | 0 | 0 | 0 | 0 | 4 mo | HOM |
| **1013608106** | 2 | PRT | 1100 | 2.142995758 | 0 | 0 | 0 | 0 | 4 mo | HOM |
| **1011937543** | 1 | PRT | 1100 | 2.142995758 | 0.002799401 | 69 | 0.001306303 | 32.19791722 | 8 mo | HOM |
| **1010419538** | 2 | PRT | 1100 | 2.142995758 | 3.25E-07 | 0 | 1.52E-07 | 0 | 4 mo | WILD |
| **1010419187** | 1 | PRT | 1100 | 2.142995758 | 0.000226276 | 10 | 0.000105589 | 4.666364814 | 8 mo | HOM |
| **1013978366** | 1 | PRT | 1100 | 2.142995758 | 6.06E-06 | 1 | 2.83E-06 | 0.466636481 | 2 mo | HOM |
| **1014416155** | 2 | PRT | 1100 | 2.142995758 | 0 | 0 | 0 | 0 | 2 mo | HOM |
| **1013978065** | 2 | PRT | 1100 | 2.142995758 | 0 | 0 | 0 | 0 | 2 mo | HOM |
| **1009934102** | 2 | PRT | 1100 | 2.142995758 | 0 | 0 | 0 | 0 | 4 mo | WILD |
| **1009322937** | 2 | PRT | 1100 | 2.142995758 | 3.02E-05 | 3 | 1.41E-05 | 1.399909444 | 8 mo | WILD |
| **1011080637** | 2 | PRT | 1100 | 2.142995758 | 0.000145756 | 10 | 6.8E-05 | 4.666364814 | 8 mo | HOM |
| **1013609713** | 2 | PRT | 1100 | 2.142995758 | 1.38E-06 | 0 | 6.43E-07 | 0 | 2 mo | WILD |

| | | | | | | | | | | |
|---|---|---|---|---|---|---|---|---|---|---|
| **1010418842** | 1 | PRT | 1100 | 2.142995758 | 0.000678907 | 26 | 0.000316803 | 12.13254852 | 8 mo | HOM |
| **1013288213** | 2 | PRT | 1100 | 2.142995758 | 5.3E-05 | 1 | 2.47E-05 | 0.466636481 | 4 mo | HOM |
| **1008976060** | 2 | PRT | 1100 | 2.142995758 | 5.37E-05 | 4 | 2.51E-05 | 1.866545926 | 8 mo | WILD |
| **1013446243** | 1 | PRT | 1100 | 2.142995758 | 8.19E-07 | 0 | 3.82E-07 | 0 | 4 mo | HOM |
| **1013288951** | 2 | PRT | 1100 | 2.142995758 | 1.67E-05 | 1 | 7.81E-06 | 0.466636481 | 4 mo | HOM |
| **1013978665** | 1 | Ald5 | 1101 | 1.555213274 | 0 | 0 | 0 | 0 | 2 mo | HOM |
| **1013445609** | 1 | Ald5 | 1101 | 1.555213274 | 9.1E-05 | 6 | 5.85E-05 | 3.857991763 | 4 mo | HOM |
| **1013609015** | 1 | Ald5 | 1101 | 1.555213274 | 0.000191004 | 10 | 0.000122815 | 6.429986272 | 4 mo | HOM |
| **1013608106** | 2 | Ald5 | 1101 | 1.555213274 | 0 | 0 | 0 | 0 | 4 mo | HOM |
| **1011937543** | 1 | Ald5 | 1101 | 1.555213274 | 0.001678465 | 49 | 0.001079251 | 31.50693274 | 8 mo | HOM |
| **1010419538** | 2 | Ald5 | 1101 | 1.555213274 | 0 | 0 | 0 | 0 | 4 mo | WILD |
| **1010419187** | 1 | Ald5 | 1101 | 1.555213274 | 0.001677303 | 69 | 0.001078504 | 44.36690528 | 8 mo | HOM |
| **1013978366** | 1 | Ald5 | 1101 | 1.555213274 | 1.24E-05 | 1 | 7.99E-06 | 0.642998627 | 2 mo | HOM |
| **1014416155** | 2 | Ald5 | 1101 | 1.555213274 | 0 | 0 | 0 | 0 | 2 mo | HOM |
| **1013978065** | 2 | Ald5 | 1101 | 1.555213274 | 0 | 0 | 0 | 0 | 2 mo | HOM |
| **1009934102** | 2 | Ald5 | 1101 | 1.555213274 | 0 | 0 | 0 | 0 | 4 mo | WILD |
| **1009322937** | 2 | Ald5 | 1101 | 1.555213274 | 5.22E-08 | 0 | 3.36E-08 | 0 | 8 mo | WILD |
| **1011080637** | 2 | Ald5 | 1101 | 1.555213274 | 0.000857826 | 48 | 0.000551581 | 30.86393411 | 8 mo | HOM |
| **1013609713** | 2 | Ald5 | 1101 | 1.555213274 | 0 | 0 | 0 | 0 | 2 mo | WILD |
| **1010418842** | 1 | Ald5 | 1101 | 1.555213274 | 0.001225662 | 58 | 0.000788099 | 37.29392038 | 8 mo | HOM |
| **1013288213** | 2 | Ald5 | 1101 | 1.555213274 | 5.96E-05 | 6 | 3.83E-05 | 3.857991763 | 4 mo | HOM |
| **1008976060** | 2 | Ald5 | 1101 | 1.555213274 | 0 | 0 | 0 | 0 | 8 mo | WILD |
| **1013446243** | 1 | Ald5 | 1101 | 1.555213274 | 0.0003459 | 18 | 0.000222413 | 11.57397529 | 4 mo | HOM |
| **1013288951** | 2 | Ald5 | 1101 | 1.555213274 | 0.000310981 | 11 | 0.000199961 | 7.0729849 | 4 mo | HOM |
| **1013978665** | 1 | SSp-m6a | 1102 | 1.701602067 | 0 | 0 | 0 | 0 | 2 mo | HOM |
| **1013445609** | 1 | SSp-m6a | 1102 | 1.701602067 | 0.000164821 | 11 | 9.69E-05 | 6.464496143 | 4 mo | HOM |
| **1013609015** | 1 | SSp-m6a | 1102 | 1.701602067 | 0.000313775 | 13 | 0.0001844 | 7.639859078 | 4 mo | HOM |
| **1013608106** | 2 | SSp-m6a | 1102 | 1.701602067 | 0 | 0 | 0 | 0 | 4 mo | HOM |
| **1011937543** | 1 | SSp-m6a | 1102 | 1.701602067 | 0.002423688 | 51 | 0.001424356 | 29.97175484 | 8 mo | HOM |
| **1010419538** | 2 | SSp-m6a | 1102 | 1.701602067 | 0 | 0 | 0 | 0 | 4 mo | WILD |
| **1010419187** | 1 | SSp-m6a | 1102 | 1.701602067 | 0.001345144 | 72 | 0.000790516 | 42.31306566 | 8 mo | HOM |
| **1013978366** | 1 | SSp-m6a | 1102 | 1.701602067 | 0 | 0 | 0 | 0 | 2 mo | HOM |
| **1014416155** | 2 | SSp-m6a | 1102 | 1.701602067 | 0 | 0 | 0 | 0 | 2 mo | HOM |
| **1013978065** | 2 | SSp-m6a | 1102 | 1.701602067 | 0 | 0 | 0 | 0 | 2 mo | HOM |
| **1009934102** | 2 | SSp-m6a | 1102 | 1.701602067 | 0 | 0 | 0 | 0 | 4 mo | WILD |
| **1009322937** | 2 | SSp-m6a | 1102 | 1.701602067 | 0 | 0 | 0 | 0 | 8 mo | WILD |
| **1011080637** | 2 | SSp-m6a | 1102 | 1.701602067 | 0.00128499 | 55 | 0.000755165 | 32.32248071 | 8 mo | HOM |
| **1013609713** | 2 | SSp-m6a | 1102 | 1.701602067 | 0 | 0 | 0 | 0 | 2 mo | WILD |
| **1010418842** | 1 | SSp-m6a | 1102 | 1.701602067 | 0.001470168 | 79 | 0.00086399 | 46.42683593 | 8 mo | HOM |
| **1013288213** | 2 | SSp-m6a | 1102 | 1.701602067 | 5.33E-05 | 5 | 3.14E-05 | 2.938407338 | 4 mo | HOM |
| **1008976060** | 2 | SSp-m6a | 1102 | 1.701602067 | 1.44E-05 | 2 | 8.45E-06 | 1.175362935 | 8 mo | WILD |
| **1013446243** | 1 | SSp-m6a | 1102 | 1.701602067 | 0.000210303 | 15 | 0.000123591 | 8.815222013 | 4 mo | HOM |
| **1013288951** | 2 | SSp-m6a | 1102 | 1.701602067 | 0.000299313 | 14 | 0.000175901 | 8.227540545 | 4 mo | HOM |
| **1013978665** | 1 | AMv | 1104 | 0.171379989 | 0 | 0 | 0 | 0 | 2 mo | HOM |
| **1013445609** | 1 | AMv | 1104 | 0.171379989 | 0 | 0 | 0 | 0 | 4 mo | HOM |
| **1013609015** | 1 | AMv | 1104 | 0.171379989 | 0 | 0 | 0 | 0 | 4 mo | HOM |
| **1013608106** | 2 | AMv | 1104 | 0.171379989 | 0 | 0 | 0 | 0 | 4 mo | HOM |
| **1011937543** | 1 | AMv | 1104 | 0.171379989 | 2.62E-05 | 1 | 0.00015286 | 5.834986947 | 8 mo | HOM |
| **1010419538** | 2 | AMv | 1104 | 0.171379989 | 0 | 0 | 0 | 0 | 4 mo | WILD |
| **1010419187** | 1 | AMv | 1104 | 0.171379989 | 0.000132399 | 6 | 0.000772544 | 35.00992168 | 8 mo | HOM |
| **1013978366** | 1 | AMv | 1104 | 0.171379989 | 0 | 0 | 0 | 0 | 2 mo | HOM |
| **1014416155** | 2 | AMv | 1104 | 0.171379989 | 0 | 0 | 0 | 0 | 2 mo | HOM |
| **1013978065** | 2 | AMv | 1104 | 0.171379989 | 0 | 0 | 0 | 0 | 2 mo | HOM |
| **1009934102** | 2 | AMv | 1104 | 0.171379989 | 0 | 0 | 0 | 0 | 4 mo | WILD |
| **1009322937** | 2 | AMv | 1104 | 0.171379989 | 8.09E-06 | 1 | 4.72E-05 | 5.834986947 | 8 mo | WILD |
| **1011080637** | 2 | AMv | 1104 | 0.171379989 | 0.000114584 | 4 | 0.000668596 | 23.33994779 | 8 mo | HOM |
| **1013609713** | 2 | AMv | 1104 | 0.171379989 | 0 | 0 | 0 | 0 | 2 mo | WILD |
| **1010418842** | 1 | AMv | 1104 | 0.171379989 | 0.000105251 | 5 | 0.000614137 | 29.17493474 | 8 mo | HOM |
| **1013288213** | 2 | AMv | 1104 | 0.171379989 | 6.41E-06 | 1 | 3.74E-05 | 5.834986947 | 4 mo | HOM |

| | | | | | | | | | | |
|---|---|---|---|---|---|---|---|---|---|---|
| **1008976060** | 2 | AMv | 1104 | 0.171379989 | 4.72E-06 | 1 | 2.75E-05 | 5.834986947 | 8 mo | WILD |
| **1013446243** | 1 | AMv | 1104 | 0.171379989 | 0 | 0 | 0 | 0 | 4 mo | HOM |
| **1013288951** | 2 | AMv | 1104 | 0.171379989 | 0 | 0 | 0 | 0 | 4 mo | HOM |
| **1013978665** | 1 | IA | 1105 | 0.188236538 | 0 | 0 | 0 | 0 | 2 mo | HOM |
| **1013445609** | 1 | IA | 1105 | 0.188236538 | 4.41E-06 | 1 | 2.34E-05 | 5.312464895 | 4 mo | HOM |
| **1013609015** | 1 | IA | 1105 | 0.188236538 | 0.000100936 | 2 | 0.000536217 | 10.62492979 | 4 mo | HOM |
| **1013608106** | 2 | IA | 1105 | 0.188236538 | 0 | 0 | 0 | 0 | 4 mo | HOM |
| **1011937543** | 1 | IA | 1105 | 0.188236538 | 0.000654626 | 5 | 0.003477677 | 26.56232447 | 8 mo | HOM |
| **1010419538** | 2 | IA | 1105 | 0.188236538 | 0 | 0 | 0 | 0 | 4 mo | WILD |
| **1010419187** | 1 | IA | 1105 | 0.188236538 | 0.00028029 | 8 | 0.001489033 | 42.49971916 | 8 mo | HOM |
| **1013978366** | 1 | IA | 1105 | 0.188236538 | 0 | 0 | 0 | 0 | 2 mo | HOM |
| **1014416155** | 2 | IA | 1105 | 0.188236538 | 0 | 0 | 0 | 0 | 2 mo | HOM |
| **1013978065** | 2 | IA | 1105 | 0.188236538 | 0 | 0 | 0 | 0 | 2 mo | HOM |
| **1009934102** | 2 | IA | 1105 | 0.188236538 | 0 | 0 | 0 | 0 | 4 mo | WILD |
| **1009322937** | 2 | IA | 1105 | 0.188236538 | 0 | 0 | 0 | 0 | 8 mo | WILD |
| **1011080637** | 2 | IA | 1105 | 0.188236538 | 0.000223295 | 12 | 0.001186249 | 63.74957873 | 8 mo | HOM |
| **1013609713** | 2 | IA | 1105 | 0.188236538 | 0 | 0 | 0 | 0 | 2 mo | WILD |
| **1010418842** | 1 | IA | 1105 | 0.188236538 | 0.000276648 | 12 | 0.001469684 | 63.74957873 | 8 mo | HOM |
| **1013288213** | 2 | IA | 1105 | 0.188236538 | 2.91E-05 | 2 | 0.0001548 | 10.62492979 | 4 mo | HOM |
| **1008976060** | 2 | IA | 1105 | 0.188236538 | 0 | 0 | 0 | 0 | 8 mo | WILD |
| **1013446243** | 1 | IA | 1105 | 0.188236538 | 5.42E-05 | 4 | 0.000288154 | 21.24985958 | 4 mo | HOM |
| **1013288951** | 2 | IA | 1105 | 0.188236538 | 2.12E-05 | 1 | 0.000112739 | 5.312464895 | 4 mo | HOM |
| **1013978665** | 1 | VISC2/3 | 1106 | 0.556127813 | 0 | 0 | 0 | 0 | 2 mo | HOM |
| **1013445609** | 1 | VISC2/3 | 1106 | 0.556127813 | 8.53E-05 | 4 | 0.000153395 | 7.192591178 | 4 mo | HOM |
| **1013609015** | 1 | VISC2/3 | 1106 | 0.556127813 | 7.89E-05 | 5 | 0.000141826 | 8.990738972 | 4 mo | HOM |
| **1013608106** | 2 | VISC2/3 | 1106 | 0.556127813 | 0 | 0 | 0 | 0 | 4 mo | HOM |
| **1011937543** | 1 | VISC2/3 | 1106 | 0.556127813 | 0.00040738 | 16 | 0.00073253 | 28.77036471 | 8 mo | HOM |
| **1010419538** | 2 | VISC2/3 | 1106 | 0.556127813 | 0 | 0 | 0 | 0 | 4 mo | WILD |
| **1010419187** | 1 | VISC2/3 | 1106 | 0.556127813 | 0.00037047 | 21 | 0.000666159 | 37.76110368 | 8 mo | HOM |
| **1013978366** | 1 | VISC2/3 | 1106 | 0.556127813 | 0 | 0 | 0 | 0 | 2 mo | HOM |
| **1014416155** | 2 | VISC2/3 | 1106 | 0.556127813 | 0 | 0 | 0 | 0 | 2 mo | HOM |
| **1013978065** | 2 | VISC2/3 | 1106 | 0.556127813 | 0 | 0 | 0 | 0 | 2 mo | HOM |
| **1009934102** | 2 | VISC2/3 | 1106 | 0.556127813 | 0 | 0 | 0 | 0 | 4 mo | WILD |
| **1009322937** | 2 | VISC2/3 | 1106 | 0.556127813 | 0 | 0 | 0 | 0 | 8 mo | WILD |
| **1011080637** | 2 | VISC2/3 | 1106 | 0.556127813 | 0.000382889 | 18 | 0.00068849 | 32.3666603 | 8 mo | HOM |
| **1013609713** | 2 | VISC2/3 | 1106 | 0.556127813 | 0 | 0 | 0 | 0 | 2 mo | WILD |
| **1010418842** | 1 | VISC2/3 | 1106 | 0.556127813 | 0.000537058 | 16 | 0.000965709 | 28.77036471 | 8 mo | HOM |
| **1013288213** | 2 | VISC2/3 | 1106 | 0.556127813 | 2.2E-05 | 2 | 3.95E-05 | 3.596295589 | 4 mo | HOM |
| **1008976060** | 2 | VISC2/3 | 1106 | 0.556127813 | 0 | 0 | 0 | 0 | 8 mo | WILD |
| **1013446243** | 1 | VISC2/3 | 1106 | 0.556127813 | 0.000108975 | 3 | 0.000195952 | 5.394443383 | 4 mo | HOM |
| **1013288951** | 2 | VISC2/3 | 1106 | 0.556127813 | 7.44E-05 | 3 | 0.000133793 | 5.394443383 | 4 mo | HOM |
| **1013978665** | 1 | MDRNv | 1107 | 0.959273392 | 0 | 0 | 0 | 0 | 2 mo | HOM |
| **1013445609** | 1 | MDRNv | 1107 | 0.959273392 | 0 | 0 | 0 | 0 | 4 mo | HOM |
| **1013609015** | 1 | MDRNv | 1107 | 0.959273392 | 0 | 0 | 0 | 0 | 4 mo | HOM |
| **1013608106** | 2 | MDRNv | 1107 | 0.958651288 | 0 | 0 | 0 | 0 | 4 mo | HOM |
| **1011937543** | 1 | MDRNv | 1107 | 0.959273392 | 4.48E-05 | 1 | 4.67E-05 | 1.042455684 | 8 mo | HOM |
| **1010419538** | 2 | MDRNv | 1107 | 0.959273392 | 0 | 0 | 0 | 0 | 4 mo | WILD |
| **1010419187** | 1 | MDRNv | 1107 | 0.959273392 | 9.96E-06 | 1 | 1.04E-05 | 1.042455684 | 8 mo | HOM |
| **1013978366** | 1 | MDRNv | 1107 | 0.959273392 | 0 | 0 | 0 | 0 | 2 mo | HOM |
| **1014416155** | 2 | MDRNv | 1107 | 0.959273392 | 0 | 0 | 0 | 0 | 2 mo | HOM |
| **1013978065** | 2 | MDRNv | 1107 | 0.959273392 | 0 | 0 | 0 | 0 | 2 mo | HOM |
| **1009934102** | 2 | MDRNv | 1107 | 0.959273392 | 0 | 0 | 0 | 0 | 4 mo | WILD |
| **1009322937** | 2 | MDRNv | 1107 | 0.959273392 | 0 | 0 | 0 | 0 | 8 mo | WILD |
| **1011080637** | 2 | MDRNv | 1107 | 0.959273392 | 4.28E-05 | 2 | 4.47E-05 | 2.084911368 | 8 mo | HOM |
| **1013609713** | 2 | MDRNv | 1107 | 0.959273392 | 0 | 0 | 0 | 0 | 2 mo | WILD |
| **1010418842** | 1 | MDRNv | 1107 | 0.959273392 | 5.45E-05 | 3 | 5.68E-05 | 3.127367052 | 8 mo | HOM |
| **1013288213** | 2 | MDRNv | 1107 | 0.959273392 | 0 | 0 | 0 | 0 | 4 mo | HOM |
| **1008976060** | 2 | MDRNv | 1107 | 0.959273392 | 1.61E-05 | 3 | 1.68E-05 | 3.127367052 | 8 mo | WILD |
| **1013446243** | 1 | MDRNv | 1107 | 0.959273392 | 0 | 0 | 0 | 0 | 4 mo | HOM |

| | | | | | | | | | | |
|---|---|---|---|---|---|---|---|---|---|---|
| **1013288951** | 2 | MDRNv | 1107 | 0.959273392 | 0 | 0 | 0 | 0 | 4 mo | HOM |
| **1013978665** | 1 | ccg | 1108 | 0.854433619 | 0 | 0 | 0 | 0 | 2 mo | HOM |
| **1013445609** | 1 | ccg | 1108 | 0.854433619 | 3.46E-06 | 0 | 4.05E-06 | 0 | 4 mo | HOM |
| **1013609015** | 1 | ccg | 1108 | 0.854433619 | 1.02E-05 | 1 | 1.2E-05 | 1.170365933 | 4 mo | HOM |
| **1013608106** | 2 | ccg | 1108 | 0.854433619 | 0 | 0 | 0 | 0 | 4 mo | HOM |
| **1011937543** | 1 | ccg | 1108 | 0.854433619 | 0.000194596 | 3 | 0.000227749 | 3.511097799 | 8 mo | HOM |
| **1010419538** | 2 | ccg | 1108 | 0.854433619 | 0 | 0 | 0 | 0 | 4 mo | WILD |
| **1010419187** | 1 | ccg | 1108 | 0.854433619 | 0.000162046 | 4 | 0.000189653 | 4.681463732 | 8 mo | HOM |
| **1013978366** | 1 | ccg | 1108 | 0.854433619 | 0 | 0 | 0 | 0 | 2 mo | HOM |
| **1014416155** | 2 | ccg | 1108 | 0.854433619 | 0 | 0 | 0 | 0 | 2 mo | HOM |
| **1013978065** | 2 | ccg | 1108 | 0.854433619 | 7.35E-08 | 0 | 8.6E-08 | 0 | 2 mo | HOM |
| **1009934102** | 2 | ccg | 1108 | 0.854433619 | 5.81E-06 | 0 | 6.8E-06 | 0 | 4 mo | WILD |
| **1009322937** | 2 | ccg | 1108 | 0.854433619 | 3.07E-06 | 0 | 3.59E-06 | 0 | 8 mo | WILD |
| **1011080637** | 2 | ccg | 1108 | 0.854433619 | 0.000108324 | 3 | 0.000126779 | 3.511097799 | 8 mo | HOM |
| **1013609713** | 2 | ccg | 1108 | 0.854433619 | 0 | 0 | 0 | 0 | 2 mo | WILD |
| **1010418842** | 1 | ccg | 1108 | 0.854433619 | 0.000203406 | 6 | 0.000238059 | 7.022195599 | 8 mo | HOM |
| **1013288213** | 2 | ccg | 1108 | 0.854433619 | 6.94E-05 | 1 | 8.12E-05 | 1.170365933 | 4 mo | HOM |
| **1008976060** | 2 | ccg | 1108 | 0.854433619 | 2.04E-06 | 0 | 2.39E-06 | 0 | 8 mo | WILD |
| **1013446243** | 1 | ccg | 1108 | 0.854433619 | 4.88E-06 | 0 | 5.71E-06 | 0 | 4 mo | HOM |
| **1013288951** | 2 | ccg | 1108 | 0.854433619 | 2.67E-06 | 0 | 3.13E-06 | 0 | 4 mo | HOM |
| **1013978665** | 1 | PS | 1109 | 0.099591853 | 0 | 0 | 0 | 0 | 2 mo | HOM |
| **1013445609** | 1 | PS | 1109 | 0.099591853 | 0 | 0 | 0 | 0 | 4 mo | HOM |
| **1013609015** | 1 | PS | 1109 | 0.099591853 | 3.57E-06 | 0 | 3.58E-05 | 0 | 4 mo | HOM |
| **1013608106** | 2 | PS | 1109 | 0.099591853 | 0 | 0 | 0 | 0 | 4 mo | HOM |
| **1011937543** | 1 | PS | 1109 | 0.099591853 | 4.48E-06 | 0 | 4.5E-05 | 0 | 8 mo | HOM |
| **1010419538** | 2 | PS | 1109 | 0.099591853 | 0 | 0 | 0 | 0 | 4 mo | WILD |
| **1010419187** | 1 | PS | 1109 | 0.099591853 | 4.88E-05 | 3 | 0.000490329 | 30.12294584 | 8 mo | HOM |
| **1013978366** | 1 | PS | 1109 | 0.099591853 | 0 | 0 | 0 | 0 | 2 mo | HOM |
| **1014416155** | 2 | PS | 1109 | 0.099591853 | 0 | 0 | 0 | 0 | 2 mo | HOM |
| **1013978065** | 2 | PS | 1109 | 0.099591853 | 0 | 0 | 0 | 0 | 2 mo | HOM |
| **1009934102** | 2 | PS | 1109 | 0.099591853 | 0 | 0 | 0 | 0 | 4 mo | WILD |
| **1009322937** | 2 | PS | 1109 | 0.099591853 | 0 | 0 | 0 | 0 | 8 mo | WILD |
| **1011080637** | 2 | PS | 1109 | 0.099591853 | 3.35E-05 | 2 | 0.000336791 | 20.08196389 | 8 mo | HOM |
| **1013609713** | 2 | PS | 1109 | 0.099591853 | 0 | 0 | 0 | 0 | 2 mo | WILD |
| **1010418842** | 1 | PS | 1109 | 0.099591853 | 2.21E-05 | 0 | 0.000221845 | 0 | 8 mo | HOM |
| **1013288213** | 2 | PS | 1109 | 0.099591853 | 0 | 0 | 0 | 0 | 4 mo | HOM |
| **1008976060** | 2 | PS | 1109 | 0.099591853 | 5.01E-06 | 1 | 5.03E-05 | 10.04098195 | 8 mo | WILD |
| **1013446243** | 1 | PS | 1109 | 0.099591853 | 0 | 0 | 0 | 0 | 4 mo | HOM |
| **1013288951** | 2 | PS | 1109 | 0.099591853 | 0 | 0 | 0 | 0 | 4 mo | HOM |
| **1013978665** | 1 | SSp-tr5 | 1111 | 0.363675609 | 0 | 0 | 0 | 0 | 2 mo | HOM |
| **1013445609** | 1 | SSp-tr5 | 1111 | 0.363675609 | 4.87E-05 | 4 | 0.00013402 | 10.99881296 | 4 mo | HOM |
| **1013609015** | 1 | SSp-tr5 | 1111 | 0.363675609 | 3.45E-05 | 2 | 9.48E-05 | 5.499406481 | 4 mo | HOM |
| **1013608106** | 2 | SSp-tr5 | 1111 | 0.363675609 | 0 | 0 | 0 | 0 | 4 mo | HOM |
| **1011937543** | 1 | SSp-tr5 | 1111 | 0.363675609 | 0.000926827 | 9 | 0.002548499 | 24.74732916 | 8 mo | HOM |
| **1010419538** | 2 | SSp-tr5 | 1111 | 0.363675609 | 0 | 0 | 0 | 0 | 4 mo | WILD |
| **1010419187** | 1 | SSp-tr5 | 1111 | 0.363675609 | 0.000514483 | 15 | 0.001414676 | 41.24554861 | 8 mo | HOM |
| **1013978366** | 1 | SSp-tr5 | 1111 | 0.363675609 | 0 | 0 | 0 | 0 | 2 mo | HOM |
| **1014416155** | 2 | SSp-tr5 | 1111 | 0.363675609 | 0 | 0 | 0 | 0 | 2 mo | HOM |
| **1013978065** | 2 | SSp-tr5 | 1111 | 0.363675609 | 0 | 0 | 0 | 0 | 2 mo | HOM |
| **1009934102** | 2 | SSp-tr5 | 1111 | 0.363675609 | 0 | 0 | 0 | 0 | 4 mo | WILD |
| **1009322937** | 2 | SSp-tr5 | 1111 | 0.363675609 | 0 | 0 | 0 | 0 | 8 mo | WILD |
| **1011080637** | 2 | SSp-tr5 | 1111 | 0.363675609 | 0.000267347 | 15 | 0.000735124 | 41.24554861 | 8 mo | HOM |
| **1013609713** | 2 | SSp-tr5 | 1111 | 0.363675609 | 0 | 0 | 0 | 0 | 2 mo | WILD |
| **1010418842** | 1 | SSp-tr5 | 1111 | 0.363675609 | 0.000348108 | 16 | 0.000957193 | 43.99525185 | 8 mo | HOM |
| **1013288213** | 2 | SSp-tr5 | 1111 | 0.363675609 | 5.94E-05 | 5 | 0.000163438 | 13.7485162 | 4 mo | HOM |
| **1008976060** | 2 | SSp-tr5 | 1111 | 0.363675609 | 8.17E-06 | 1 | 2.25E-05 | 2.74970324 | 8 mo | WILD |
| **1013446243** | 1 | SSp-tr5 | 1111 | 0.363675609 | 0.000163983 | 7 | 0.000450905 | 19.24792268 | 4 mo | HOM |
| **1013288951** | 2 | SSp-tr5 | 1111 | 0.363675609 | 8.8E-05 | 3 | 0.000242081 | 8.249109721 | 4 mo | HOM |
| **1013978665** | 1 | IAD | 1113 | 0.114929843 | 0 | 0 | 0 | 0 | 2 mo | HOM |

| | | | | | | | | | | |
|---|---|---|---|---|---|---|---|---|---|---|
| 1013445609 | 1 | IAD | 1113 | 0.114929843 | 0 | 0 | 0 | 0 | 4 mo | HOM |
| 1013609015 | 1 | IAD | 1113 | 0.114929843 | 0 | 0 | 0 | 0 | 4 mo | HOM |
| 1013608106 | 2 | IAD | 1113 | 0.114929843 | 0 | 0 | 0 | 0 | 4 mo | HOM |
| 1011937543 | 1 | IAD | 1113 | 0.114929843 | 1.54E-05 | 1 | 0.000133919 | 8.700960289 | 8 mo | HOM |
| 1010419538 | 2 | IAD | 1113 | 0.114929843 | 0 | 0 | 0 | 0 | 4 mo | WILD |
| 1010419187 | 1 | IAD | 1113 | 0.114929843 | 7.71E-05 | 2 | 0.000671081 | 17.40192058 | 8 mo | HOM |
| 1013978366 | 1 | IAD | 1113 | 0.114929843 | 0 | 0 | 0 | 0 | 2 mo | HOM |
| 1014416155 | 2 | IAD | 1113 | 0.114929843 | 0 | 0 | 0 | 0 | 2 mo | HOM |
| 1013978065 | 2 | IAD | 1113 | 0.114929843 | 0 | 0 | 0 | 0 | 2 mo | HOM |
| 1009934102 | 2 | IAD | 1113 | 0.114929843 | 0 | 0 | 0 | 0 | 4 mo | WILD |
| 1009322937 | 2 | IAD | 1113 | 0.114929843 | 1.66E-06 | 0 | 1.45E-05 | 0 | 8 mo | WILD |
| 1011080637 | 2 | IAD | 1113 | 0.114929843 | 2.25E-05 | 1 | 0.000195725 | 8.700960289 | 8 mo | HOM |
| 1013609713 | 2 | IAD | 1113 | 0.114929843 | 0 | 0 | 0 | 0 | 2 mo | WILD |
| 1010418842 | 1 | IAD | 1113 | 0.114929843 | 3.56E-05 | 1 | 0.000309925 | 8.700960289 | 8 mo | HOM |
| 1013288213 | 2 | IAD | 1113 | 0.114929843 | 0 | 0 | 0 | 0 | 4 mo | HOM |
| 1008976060 | 2 | IAD | 1113 | 0.114929843 | 0 | 0 | 0 | 0 | 8 mo | WILD |
| 1013446243 | 1 | IAD | 1113 | 0.114929843 | 0 | 0 | 0 | 0 | 4 mo | HOM |
| 1013288951 | 2 | IAD | 1113 | 0.114929843 | 0 | 0 | 0 | 0 | 4 mo | HOM |
| 1013978665 | 1 | VISal4 | 1114 | 0.107299994 | 0 | 0 | 0 | 0 | 2 mo | HOM |
| 1013445609 | 1 | VISal4 | 1114 | 0.107299994 | 2.89E-05 | 2 | 0.000269483 | 18.63932999 | 4 mo | HOM |
| 1013609015 | 1 | VISal4 | 1114 | 0.107299994 | 7.59E-05 | 4 | 0.000707623 | 37.27865998 | 4 mo | HOM |
| 1013608106 | 2 | VISal4 | 1114 | 0.107299994 | 0 | 0 | 0 | 0 | 4 mo | HOM |
| 1011937543 | 1 | VISal4 | 1114 | 0.107299994 | 0.000227091 | 8 | 0.002116414 | 74.55731997 | 8 mo | HOM |
| 1010419538 | 2 | VISal4 | 1114 | 0.107299994 | 0 | 0 | 0 | 0 | 4 mo | WILD |
| 1010419187 | 1 | VISal4 | 1114 | 0.107299994 | 0.000232273 | 10 | 0.002164705 | 93.19664996 | 8 mo | HOM |
| 1013978366 | 1 | VISal4 | 1114 | 0.107299994 | 0 | 0 | 0 | 0 | 2 mo | HOM |
| 1014416155 | 2 | VISal4 | 1114 | 0.107299994 | 0 | 0 | 0 | 0 | 2 mo | HOM |
| 1013978065 | 2 | VISal4 | 1114 | 0.107299994 | 0 | 0 | 0 | 0 | 2 mo | HOM |
| 1009934102 | 2 | VISal4 | 1114 | 0.107299994 | 0 | 0 | 0 | 0 | 4 mo | WILD |
| 1009322937 | 2 | VISal4 | 1114 | 0.107299994 | 0 | 0 | 0 | 0 | 8 mo | WILD |
| 1011080637 | 2 | VISal4 | 1114 | 0.107299994 | 4.75E-05 | 3 | 0.000442922 | 27.95899499 | 8 mo | HOM |
| 1013609713 | 2 | VISal4 | 1114 | 0.107299994 | 0 | 0 | 0 | 0 | 2 mo | WILD |
| 1010418842 | 1 | VISal4 | 1114 | 0.107299994 | 0.000137403 | 5 | 0.001280547 | 46.59832498 | 8 mo | HOM |
| 1013288213 | 2 | VISal4 | 1114 | 0.107299994 | 1.73E-05 | 1 | 0.000161659 | 9.319664996 | 4 mo | HOM |
| 1008976060 | 2 | VISal4 | 1114 | 0.107299994 | 0 | 0 | 0 | 0 | 8 mo | WILD |
| 1013446243 | 1 | VISal4 | 1114 | 0.107299994 | 8.14E-06 | 1 | 7.59E-05 | 9.319664996 | 4 mo | HOM |
| 1013288951 | 2 | VISal4 | 1114 | 0.107299994 | 0.000131672 | 3 | 0.001227142 | 27.95899499 | 4 mo | HOM |
| 1013978665 | 1 | gVIIn | 1116 | 0.029905371 | 0 | 0 | 0 | 0 | 2 mo | HOM |
| 1013445609 | 1 | gVIIn | 1116 | 0.029905371 | 0 | 0 | 0 | 0 | 4 mo | HOM |
| 1013609015 | 1 | gVIIn | 1116 | 0.029905371 | 0 | 0 | 0 | 0 | 4 mo | HOM |
| 1013608106 | 2 | gVIIn | 1116 | 0.029905371 | 0 | 0 | 0 | 0 | 4 mo | HOM |
| 1011937543 | 1 | gVIIn | 1116 | 0.029905371 | 0 | 0 | 0 | 0 | 8 mo | HOM |
| 1010419538 | 2 | gVIIn | 1116 | 0.029905371 | 0 | 0 | 0 | 0 | 4 mo | WILD |
| 1010419187 | 1 | gVIIn | 1116 | 0.029905371 | 0 | 0 | 0 | 0 | 8 mo | HOM |
| 1013978366 | 1 | gVIIn | 1116 | 0.029905371 | 0 | 0 | 0 | 0 | 2 mo | HOM |
| 1014416155 | 2 | gVIIn | 1116 | 0.029905371 | 0 | 0 | 0 | 0 | 2 mo | HOM |
| 1013978065 | 2 | gVIIn | 1116 | 0.029905371 | 0 | 0 | 0 | 0 | 2 mo | HOM |
| 1009934102 | 2 | gVIIn | 1116 | 0.029905371 | 0 | 0 | 0 | 0 | 4 mo | WILD |
| 1009322937 | 2 | gVIIn | 1116 | 0.029905371 | 0 | 0 | 0 | 0 | 8 mo | WILD |
| 1011080637 | 2 | gVIIn | 1116 | 0.029905371 | 0 | 0 | 0 | 0 | 8 mo | HOM |
| 1013609713 | 2 | gVIIn | 1116 | 0.029905371 | 0 | 0 | 0 | 0 | 2 mo | WILD |
| 1010418842 | 1 | gVIIn | 1116 | 0.029905371 | 1.94E-07 | 0 | 6.47E-06 | 0 | 8 mo | HOM |
| 1013288213 | 2 | gVIIn | 1116 | 0.029905371 | 0 | 0 | 0 | 0 | 4 mo | HOM |
| 1008976060 | 2 | gVIIn | 1116 | 0.029905371 | 0 | 0 | 0 | 0 | 8 mo | WILD |
| 1013446243 | 1 | gVIIn | 1116 | 0.029905371 | 0 | 0 | 0 | 0 | 4 mo | HOM |
| 1013288951 | 2 | gVIIn | 1116 | 0.029905371 | 0 | 0 | 0 | 0 | 4 mo | HOM |
| 1013978665 | 1 | P-sat | 1117 | 3.566653346 | 0 | 0 | 0 | 0 | 2 mo | HOM |
| 1013445609 | 1 | P-sat | 1117 | 3.566653346 | 0 | 0 | 0 | 0 | 4 mo | HOM |
| 1013609015 | 1 | P-sat | 1117 | 3.566653346 | 1.7E-05 | 0 | 4.78E-06 | 0 | 4 mo | HOM |

| | | | | | | | | | | |
|---|---|---|---|---|---|---|---|---|---|---|
| 1013608106 | 2 | P-sat | 1117 | 3.566653346 | 0 | 0 | 0 | 0 | 4 mo | HOM |
| 1011937543 | 1 | P-sat | 1117 | 3.566653346 | 0.007828948 | 16 | 0.00219504 | 4.485998062 | 8 mo | HOM |
| 1010419538 | 2 | P-sat | 1117 | 3.566653346 | 0 | 0 | 0 | 0 | 4 mo | WILD |
| 1010419187 | 1 | P-sat | 1117 | 3.566653346 | 2.98E-05 | 4 | 8.36E-06 | 1.121499516 | 8 mo | HOM |
| 1013978366 | 1 | P-sat | 1117 | 3.566653346 | 1.92E-06 | 0 | 5.38E-07 | 0 | 2 mo | HOM |
| 1014416155 | 2 | P-sat | 1117 | 3.566653346 | 0 | 0 | 0 | 0 | 2 mo | HOM |
| 1013978065 | 2 | P-sat | 1117 | 3.566653346 | 0 | 0 | 0 | 0 | 2 mo | HOM |
| 1009934102 | 2 | P-sat | 1117 | 3.566653346 | 0 | 0 | 0 | 0 | 4 mo | WILD |
| 1009322937 | 2 | P-sat | 1117 | 3.566653346 | 1.01E-05 | 2 | 2.84E-06 | 0.560749758 | 8 mo | WILD |
| 1011080637 | 2 | P-sat | 1117 | 3.566653346 | 4.84E-05 | 2 | 1.36E-05 | 0.560749758 | 8 mo | HOM |
| 1013609713 | 2 | P-sat | 1117 | 3.566653346 | 0 | 0 | 0 | 0 | 2 mo | WILD |
| 1010418842 | 1 | P-sat | 1117 | 3.566653346 | 0.000310715 | 24 | 8.71E-05 | 6.728997093 | 8 mo | HOM |
| 1013288213 | 2 | P-sat | 1117 | 3.566653346 | 0 | 0 | 0 | 0 | 4 mo | HOM |
| 1008976060 | 2 | P-sat | 1117 | 3.566653346 | 0.000257369 | 20 | 7.22E-05 | 5.607497578 | 8 mo | WILD |
| 1013446243 | 1 | P-sat | 1117 | 3.566653346 | 6.39E-08 | 0 | 1.79E-08 | 0 | 4 mo | HOM |
| 1013288951 | 2 | P-sat | 1117 | 3.566653346 | 5.47E-06 | 1 | 1.53E-06 | 0.280374879 | 4 mo | HOM |
| 1013978665 | 1 | IAM | 1120 | 0.044682919 | 0 | 0 | 0 | 0 | 2 mo | HOM |
| 1013445609 | 1 | IAM | 1120 | 0.044682919 | 0 | 0 | 0 | 0 | 4 mo | HOM |
| 1013609015 | 1 | IAM | 1120 | 0.044682919 | 0 | 0 | 0 | 0 | 4 mo | HOM |
| 1013608106 | 2 | IAM | 1120 | 0.044682919 | 0 | 0 | 0 | 0 | 4 mo | HOM |
| 1011937543 | 1 | IAM | 1120 | 0.044682919 | 2.27E-05 | 0 | 0.000508313 | 0 | 8 mo | HOM |
| 1010419538 | 2 | IAM | 1120 | 0.044682919 | 0 | 0 | 0 | 0 | 4 mo | WILD |
| 1010419187 | 1 | IAM | 1120 | 0.044682919 | 0 | 0 | 0 | 0 | 8 mo | HOM |
| 1013978366 | 1 | IAM | 1120 | 0.044682919 | 0 | 0 | 0 | 0 | 2 mo | HOM |
| 1014416155 | 2 | IAM | 1120 | 0.044682919 | 0 | 0 | 0 | 0 | 2 mo | HOM |
| 1013978065 | 2 | IAM | 1120 | 0.044682919 | 0 | 0 | 0 | 0 | 2 mo | HOM |
| 1009934102 | 2 | IAM | 1120 | 0.044682919 | 0 | 0 | 0 | 0 | 4 mo | WILD |
| 1009322937 | 2 | IAM | 1120 | 0.044682919 | 0 | 0 | 0 | 0 | 8 mo | WILD |
| 1011080637 | 2 | IAM | 1120 | 0.044682919 | 0 | 0 | 0 | 0 | 8 mo | HOM |
| 1013609713 | 2 | IAM | 1120 | 0.044682919 | 0 | 0 | 0 | 0 | 2 mo | WILD |
| 1010418842 | 1 | IAM | 1120 | 0.044682919 | 8.1E-06 | 1 | 0.00018117 | 22.37991674 | 8 mo | HOM |
| 1013288213 | 2 | IAM | 1120 | 0.044682919 | 0 | 0 | 0 | 0 | 4 mo | HOM |
| 1008976060 | 2 | IAM | 1120 | 0.044682919 | 0 | 0 | 0 | 0 | 8 mo | WILD |
| 1013446243 | 1 | IAM | 1120 | 0.044682919 | 0 | 0 | 0 | 0 | 4 mo | HOM |
| 1013288951 | 2 | IAM | 1120 | 0.044682919 | 0 | 0 | 0 | 0 | 4 mo | HOM |
| 1013978665 | 1 | ENTI1 | 1121 | 1.008736579 | 9.39E-05 | 3 | 9.31E-05 | 2.974017263 | 2 mo | HOM |
| 1013445609 | 1 | ENTI1 | 1121 | 1.008736579 | 2.34E-05 | 3 | 2.32E-05 | 2.974017263 | 4 mo | HOM |
| 1013609015 | 1 | ENTI1 | 1121 | 1.008736579 | 0.00018632 | 8 | 0.000184707 | 7.9307127 | 4 mo | HOM |
| 1013608106 | 2 | ENTI1 | 1121 | 1.008736579 | 0 | 0 | 0 | 0 | 4 mo | HOM |
| 1011937543 | 1 | ENTI1 | 1121 | 1.008736579 | 0.008182397 | 21 | 0.00811153 | 20.81812084 | 8 mo | HOM |
| 1010419538 | 2 | ENTI1 | 1121 | 1.008736579 | 9.43E-06 | 0 | 9.35E-06 | 0 | 4 mo | WILD |
| 1010419187 | 1 | ENTI1 | 1121 | 1.008736579 | 0.004848691 | 57 | 0.004806697 | 56.50632799 | 8 mo | HOM |
| 1013978366 | 1 | ENTI1 | 1121 | 1.008736579 | 6.88E-05 | 2 | 6.82E-05 | 1.982678175 | 2 mo | HOM |
| 1014416155 | 2 | ENTI1 | 1121 | 1.008736579 | 0 | 0 | 0 | 0 | 2 mo | HOM |
| 1013978065 | 2 | ENTI1 | 1121 | 1.008736579 | 0 | 0 | 0 | 0 | 2 mo | HOM |
| 1009934102 | 2 | ENTI1 | 1121 | 1.008736579 | 3.69E-05 | 2 | 3.65E-05 | 1.982678175 | 4 mo | WILD |
| 1009322937 | 2 | ENTI1 | 1121 | 1.008736579 | 6.47E-05 | 4 | 6.42E-05 | 3.96535635 | 8 mo | WILD |
| 1011080637 | 2 | ENTI1 | 1121 | 1.008736579 | 0.00097913 | 18 | 0.00097065 | 17.84410358 | 8 mo | HOM |
| 1013609713 | 2 | ENTI1 | 1121 | 1.008736579 | 5.66E-06 | 1 | 5.61E-06 | 0.991339088 | 2 mo | WILD |
| 1010418842 | 1 | ENTI1 | 1121 | 1.008736579 | 0.002648776 | 32 | 0.002625835 | 31.7228508 | 8 mo | HOM |
| 1013288213 | 2 | ENTI1 | 1121 | 1.008736579 | 5.88E-06 | 1 | 5.83E-06 | 0.991339088 | 4 mo | HOM |
| 1008976060 | 2 | ENTI1 | 1121 | 1.008736579 | 7.54E-05 | 5 | 7.48E-05 | 4.956695438 | 8 mo | WILD |
| 1013446243 | 1 | ENTI1 | 1121 | 1.008736579 | 8.76E-05 | 3 | 8.68E-05 | 2.974017263 | 4 mo | HOM |
| 1013288951 | 2 | ENTI1 | 1121 | 1.008736579 | 0.000198963 | 9 | 0.00019724 | 8.922051788 | 4 mo | HOM |
| 1013978665 | 1 | icp | 1123 | 0.894954424 | 0 | 0 | 0 | 0 | 2 mo | HOM |
| 1013445609 | 1 | icp | 1123 | 0.894954424 | 0 | 0 | 0 | 0 | 4 mo | HOM |
| 1013609015 | 1 | icp | 1123 | 0.894954424 | 5.21E-07 | 0 | 5.82E-07 | 0 | 4 mo | HOM |
| 1013608106 | 2 | icp | 1123 | 0.894954424 | 0 | 0 | 0 | 0 | 4 mo | HOM |
| 1011937543 | 1 | icp | 1123 | 0.894954424 | 0.0006714 | 3 | 0.000750205 | 3.352126007 | 8 mo | HOM |

| | | | | | | | | | | |
|---|---|---|---|---|---|---|---|---|---|---|
| **1010419538** | 2 | icp | 1123 | 0.894954424 | 0 | 0 | 0 | 0 | 4 mo | WILD |
| **1010419187** | 1 | icp | 1123 | 0.894954424 | 0 | 0 | 0 | 0 | 8 mo | HOM |
| **1013978366** | 1 | icp | 1123 | 0.894954424 | 0 | 0 | 0 | 0 | 2 mo | HOM |
| **1014416155** | 2 | icp | 1123 | 0.894954424 | 0 | 0 | 0 | 0 | 2 mo | HOM |
| **1013978065** | 2 | icp | 1123 | 0.894954424 | 0 | 0 | 0 | 0 | 2 mo | HOM |
| **1009934102** | 2 | icp | 1123 | 0.894954424 | 0 | 0 | 0 | 0 | 4 mo | WILD |
| **1009322937** | 2 | icp | 1123 | 0.894954424 | 0 | 0 | 0 | 0 | 8 mo | WILD |
| **1011080637** | 2 | icp | 1123 | 0.894954424 | 0 | 0 | 0 | 0 | 8 mo | HOM |
| **1013609713** | 2 | icp | 1123 | 0.894954424 | 0 | 0 | 0 | 0 | 2 mo | WILD |
| **1010418842** | 1 | icp | 1123 | 0.894954424 | 1.78E-08 | 0 | 1.99E-08 | 0 | 8 mo | HOM |
| **1013288213** | 2 | icp | 1123 | 0.894954424 | 0 | 0 | 0 | 0 | 4 mo | HOM |
| **1008976060** | 2 | icp | 1123 | 0.894954424 | 8.91E-07 | 0 | 9.96E-07 | 0 | 8 mo | WILD |
| **1013446243** | 1 | icp | 1123 | 0.894954424 | 0 | 0 | 0 | 0 | 4 mo | HOM |
| **1013288951** | 2 | icp | 1123 | 0.894954424 | 0 | 0 | 0 | 0 | 4 mo | HOM |
| **1013978665** | 1 | ORBvl5 | 1125 | 0.638681798 | 0 | 0 | 0 | 0 | 2 mo | HOM |
| **1013445609** | 1 | ORBvl5 | 1125 | 0.638681798 | 3.64E-05 | 3 | 5.7E-05 | 4.69717473 | 4 mo | HOM |
| **1013609015** | 1 | ORBvl5 | 1125 | 0.638681798 | 0.000188817 | 6 | 0.000295636 | 9.394349459 | 4 mo | HOM |
| **1013608106** | 2 | ORBvl5 | 1125 | 0.638681798 | 0 | 0 | 0 | 0 | 4 mo | HOM |
| **1011937543** | 1 | ORBvl5 | 1125 | 0.638681798 | 0.000548348 | 20 | 0.000858562 | 31.3144982 | 8 mo | HOM |
| **1010419538** | 2 | ORBvl5 | 1125 | 0.638681798 | 0 | 0 | 0 | 0 | 4 mo | WILD |
| **1010419187** | 1 | ORBvl5 | 1125 | 0.638681798 | 0.000617163 | 22 | 0.000966307 | 34.44594802 | 8 mo | HOM |
| **1013978366** | 1 | ORBvl5 | 1125 | 0.638681798 | 0 | 0 | 0 | 0 | 2 mo | HOM |
| **1014416155** | 2 | ORBvl5 | 1125 | 0.638681798 | 0 | 0 | 0 | 0 | 2 mo | HOM |
| **1013978065** | 2 | ORBvl5 | 1125 | 0.638681798 | 0 | 0 | 0 | 0 | 2 mo | HOM |
| **1009934102** | 2 | ORBvl5 | 1125 | 0.638681798 | 0 | 0 | 0 | 0 | 4 mo | WILD |
| **1009322937** | 2 | ORBvl5 | 1125 | 0.638681798 | 6.78E-06 | 1 | 1.06E-05 | 1.56572491 | 8 mo | WILD |
| **1011080637** | 2 | ORBvl5 | 1125 | 0.638681798 | 0.000296279 | 15 | 0.000463891 | 23.48587365 | 8 mo | HOM |
| **1013609713** | 2 | ORBvl5 | 1125 | 0.638681798 | 0 | 0 | 0 | 0 | 2 mo | WILD |
| **1010418842** | 1 | ORBvl5 | 1125 | 0.638681798 | 0.000915951 | 27 | 0.001434127 | 42.27457257 | 8 mo | HOM |
| **1013288213** | 2 | ORBvl5 | 1125 | 0.638681798 | 5.9E-06 | 0 | 9.24E-06 | 0 | 4 mo | HOM |
| **1008976060** | 2 | ORBvl5 | 1125 | 0.638681798 | 0 | 0 | 0 | 0 | 8 mo | WILD |
| **1013446243** | 1 | ORBvl5 | 1125 | 0.638681798 | 0.000102174 | 4 | 0.000159976 | 6.26289964 | 4 mo | HOM |
| **1013288951** | 2 | ORBvl5 | 1125 | 0.638681798 | 9.02E-05 | 5 | 0.000141187 | 7.828624549 | 4 mo | HOM |
| **1013978665** | 1 | TMd | 1126 | 0.036785214 | 0 | 0 | 0 | 0 | 2 mo | HOM |
| **1013445609** | 1 | TMd | 1126 | 0.036785214 | 0 | 0 | 0 | 0 | 4 mo | HOM |
| **1013609015** | 1 | TMd | 1126 | 0.036785214 | 0 | 0 | 0 | 0 | 4 mo | HOM |
| **1013608106** | 2 | TMd | 1126 | 0.036785214 | 0 | 0 | 0 | 0 | 4 mo | HOM |
| **1011937543** | 1 | TMd | 1126 | 0.036785214 | 0.000308325 | 2 | 0.008381752 | 54.36967169 | 8 mo | HOM |
| **1010419538** | 2 | TMd | 1126 | 0.036785214 | 0 | 0 | 0 | 0 | 4 mo | WILD |
| **1010419187** | 1 | TMd | 1126 | 0.036785214 | 0 | 0 | 0 | 0 | 8 mo | HOM |
| **1013978366** | 1 | TMd | 1126 | 0.036785214 | 0 | 0 | 0 | 0 | 2 mo | HOM |
| **1014416155** | 2 | TMd | 1126 | 0.036785214 | 0 | 0 | 0 | 0 | 2 mo | HOM |
| **1013978065** | 2 | TMd | 1126 | 0.036785214 | 0 | 0 | 0 | 0 | 2 mo | HOM |
| **1009934102** | 2 | TMd | 1126 | 0.036785214 | 0 | 0 | 0 | 0 | 4 mo | WILD |
| **1009322937** | 2 | TMd | 1126 | 0.036785214 | 0 | 0 | 0 | 0 | 8 mo | WILD |
| **1011080637** | 2 | TMd | 1126 | 0.036785214 | 0 | 0 | 0 | 0 | 8 mo | HOM |
| **1013609713** | 2 | TMd | 1126 | 0.036785214 | 0 | 0 | 0 | 0 | 2 mo | WILD |
| **1010418842** | 1 | TMd | 1126 | 0.036785214 | 0 | 0 | 0 | 0 | 8 mo | HOM |
| **1013288213** | 2 | TMd | 1126 | 0.036785214 | 0 | 0 | 0 | 0 | 4 mo | HOM |
| **1008976060** | 2 | TMd | 1126 | 0.036785214 | 1.43E-06 | 0 | 3.88E-05 | 0 | 8 mo | WILD |
| **1013446243** | 1 | TMd | 1126 | 0.036785214 | 0 | 0 | 0 | 0 | 4 mo | HOM |
| **1013288951** | 2 | TMd | 1126 | 0.036785214 | 0 | 0 | 0 | 0 | 4 mo | HOM |
| **1013978665** | 1 | TEa2/3 | 1127 | 0.722801744 | 0 | 0 | 0 | 0 | 2 mo | HOM |
| **1013445609** | 1 | TEa2/3 | 1127 | 0.722801744 | 8.95E-05 | 7 | 0.000123832 | 9.684536677 | 4 mo | HOM |
| **1013609015** | 1 | TEa2/3 | 1127 | 0.722801744 | 0.000183427 | 6 | 0.000253773 | 8.301031437 | 4 mo | HOM |
| **1013608106** | 2 | TEa2/3 | 1127 | 0.722801744 | 0 | 0 | 0 | 0 | 4 mo | HOM |
| **1011937543** | 1 | TEa2/3 | 1127 | 0.722801744 | 0.000674812 | 20 | 0.000933606 | 27.67010479 | 8 mo | HOM |
| **1010419538** | 2 | TEa2/3 | 1127 | 0.722801744 | 0 | 0 | 0 | 0 | 4 mo | WILD |
| **1010419187** | 1 | TEa2/3 | 1127 | 0.722801744 | 0.001302328 | 40 | 0.001801778 | 55.34020958 | 8 mo | HOM |

| | | | | | | | | | | |
|---|---|---|---|---|---|---|---|---|---|---|
| **1013978366** | 1 | TEa2/3 | 1127 | 0.722801744 | 0 | 0 | 0 | 0 | 2 mo | HOM |
| **1014416155** | 2 | TEa2/3 | 1127 | 0.722801744 | 0 | 0 | 0 | 0 | 2 mo | HOM |
| **1013978065** | 2 | TEa2/3 | 1127 | 0.722801744 | 0 | 0 | 0 | 0 | 2 mo | HOM |
| **1009934102** | 2 | TEa2/3 | 1127 | 0.722801744 | 0 | 0 | 0 | 0 | 4 mo | WILD |
| **1009322937** | 2 | TEa2/3 | 1127 | 0.722801744 | 0 | 0 | 0 | 0 | 8 mo | WILD |
| **1011080637** | 2 | TEa2/3 | 1127 | 0.722801744 | 0.000452126 | 14 | 0.000625519 | 19.36907335 | 8 mo | HOM |
| **1013609713** | 2 | TEa2/3 | 1127 | 0.722801744 | 0 | 0 | 0 | 0 | 2 mo | WILD |
| **1010418842** | 1 | TEa2/3 | 1127 | 0.722801744 | 0.000662456 | 25 | 0.000916512 | 34.58763099 | 8 mo | HOM |
| **1013288213** | 2 | TEa2/3 | 1127 | 0.722801744 | 8.34E-05 | 3 | 0.000115379 | 4.150515719 | 4 mo | HOM |
| **1008976060** | 2 | TEa2/3 | 1127 | 0.722801744 | 0 | 0 | 0 | 0 | 8 mo | WILD |
| **1013446243** | 1 | TEa2/3 | 1127 | 0.722801744 | 0.000108844 | 9 | 0.000150587 | 12.45154716 | 4 mo | HOM |
| **1013288951** | 2 | TEa2/3 | 1127 | 0.722801744 | 0.000115452 | 5 | 0.000159729 | 6.917526198 | 4 mo | HOM |
| **1013978665** | 1 | SSp-II5 | 1128 | 0.572042738 | 0 | 0 | 0 | 0 | 2 mo | HOM |
| **1013445609** | 1 | SSp-II5 | 1128 | 0.572042738 | 3.92E-05 | 2 | 6.86E-05 | 3.496242266 | 4 mo | HOM |
| **1013609015** | 1 | SSp-II5 | 1128 | 0.572042738 | 0.00019274 | 3 | 0.000336933 | 5.244363399 | 4 mo | HOM |
| **1013608106** | 2 | SSp-II5 | 1128 | 0.572042738 | 0 | 0 | 0 | 0 | 4 mo | HOM |
| **1011937543** | 1 | SSp-II5 | 1128 | 0.572042738 | 0.000839589 | 18 | 0.001467703 | 31.46618039 | 8 mo | HOM |
| **1010419538** | 2 | SSp-II5 | 1128 | 0.572042738 | 0 | 0 | 0 | 0 | 4 mo | WILD |
| **1010419187** | 1 | SSp-II5 | 1128 | 0.572042738 | 0.001171913 | 30 | 0.002048646 | 52.44363399 | 8 mo | HOM |
| **1013978366** | 1 | SSp-II5 | 1128 | 0.572042738 | 0 | 0 | 0 | 0 | 2 mo | HOM |
| **1014416155** | 2 | SSp-II5 | 1128 | 0.572042738 | 0 | 0 | 0 | 0 | 2 mo | HOM |
| **1013978065** | 2 | SSp-II5 | 1128 | 0.572042738 | 0 | 0 | 0 | 0 | 2 mo | HOM |
| **1009934102** | 2 | SSp-II5 | 1128 | 0.572042738 | 0 | 0 | 0 | 0 | 4 mo | WILD |
| **1009322937** | 2 | SSp-II5 | 1128 | 0.572042738 | 8.24E-06 | 1 | 1.44E-05 | 1.748121133 | 8 mo | WILD |
| **1011080637** | 2 | SSp-II5 | 1128 | 0.572042738 | 0.000545819 | 18 | 0.000954158 | 31.46618039 | 8 mo | HOM |
| **1013609713** | 2 | SSp-II5 | 1128 | 0.572042738 | 0 | 0 | 0 | 0 | 2 mo | WILD |
| **1010418842** | 1 | SSp-II5 | 1128 | 0.572042738 | 0.000727294 | 17 | 0.001271398 | 29.71805926 | 8 mo | HOM |
| **1013288213** | 2 | SSp-II5 | 1128 | 0.572042738 | 7.52E-07 | 0 | 1.31E-06 | 0 | 4 mo | HOM |
| **1008976060** | 2 | SSp-II5 | 1128 | 0.572042738 | 5.72E-06 | 1 | 9.99E-06 | 1.748121133 | 8 mo | WILD |
| **1013446243** | 1 | SSp-II5 | 1128 | 0.572042738 | 8.73E-05 | 4 | 0.000152546 | 6.992484532 | 4 mo | HOM |
| **1013288951** | 2 | SSp-II5 | 1128 | 0.572042738 | 0.000142105 | 7 | 0.000248417 | 12.23684793 | 4 mo | HOM |
| **1013978665** | 1 | IB | 1129 | 36.54924512 | 4.06E-05 | 2 | 1.11E-06 | 0.054720692 | 2 mo | HOM |
| **1013445609** | 1 | IB | 1129 | 36.54924512 | 0.000342411 | 17 | 9.37E-06 | 0.465125885 | 4 mo | HOM |
| **1013609015** | 1 | IB | 1129 | 36.54924512 | 0.002212631 | 57 | 6.05E-05 | 1.559539734 | 4 mo | HOM |
| **1013608106** | 2 | IB | 1129 | 36.54924512 | 0 | 0 | 0 | 0 | 4 mo | HOM |
| **1011937543** | 1 | IB | 1129 | 36.54924512 | 0.039609675 | 477 | 0.001083734 | 13.05088514 | 8 mo | HOM |
| **1010419538** | 2 | IB | 1129 | 36.54924512 | 5.5E-05 | 4 | 1.5E-06 | 0.109441385 | 4 mo | WILD |
| **1010419187** | 1 | IB | 1129 | 36.54924512 | 0.018799342 | 393 | 0.000514357 | 10.75261606 | 8 mo | HOM |
| **1013978366** | 1 | IB | 1129 | 36.54924512 | 0.000298546 | 7 | 8.17E-06 | 0.191522423 | 2 mo | HOM |
| **1014416155** | 2 | IB | 1129 | 36.54924512 | 0 | 0 | 0 | 0 | 2 mo | HOM |
| **1013978065** | 2 | IB | 1129 | 36.54924512 | 4.14E-05 | 3 | 1.13E-06 | 0.082081039 | 2 mo | HOM |
| **1009934102** | 2 | IB | 1129 | 36.54924512 | 3.38E-05 | 5 | 9.24E-07 | 0.136801731 | 4 mo | WILD |
| **1009322937** | 2 | IB | 1129 | 36.54924512 | 0.000165686 | 20 | 4.53E-06 | 0.547206924 | 8 mo | WILD |
| **1011080637** | 2 | IB | 1129 | 36.54924512 | 0.013752338 | 387 | 0.000376269 | 10.58845398 | 8 mo | HOM |
| **1013609713** | 2 | IB | 1129 | 36.54924512 | 0 | 0 | 0 | 0 | 2 mo | WILD |
| **1010418842** | 1 | IB | 1129 | 36.54924512 | 0.024378031 | 606 | 0.000666991 | 16.5803698 | 8 mo | HOM |
| **1013288213** | 2 | IB | 1129 | 36.54924512 | 0.000161187 | 18 | 4.41E-06 | 0.492486232 | 4 mo | HOM |
| **1008976060** | 2 | IB | 1129 | 36.54924512 | 0.001941654 | 114 | 5.31E-05 | 3.119079467 | 8 mo | WILD |
| **1013446243** | 1 | IB | 1129 | 36.54924512 | 0.001382663 | 28 | 3.78E-05 | 0.766089694 | 4 mo | HOM |
| **1013288951** | 2 | IB | 1129 | 36.54924512 | 0.001263918 | 22 | 3.46E-05 | 0.601927616 | 4 mo | HOM |
| **1013978665** | 1 | P-sen | 1132 | 3.96653728 | 5.17E-05 | 7 | 1.3E-05 | 1.764763446 | 2 mo | HOM |
| **1013445609** | 1 | P-sen | 1132 | 3.96653728 | 1.01E-05 | 2 | 2.54E-06 | 0.504218128 | 4 mo | HOM |
| **1013609015** | 1 | P-sen | 1132 | 3.96653728 | 0.000673194 | 6 | 0.000169718 | 1.512654383 | 4 mo | HOM |
| **1013608106** | 2 | P-sen | 1132 | 3.96653728 | 0 | 0 | 0 | 0 | 4 mo | HOM |
| **1011937543** | 1 | P-sen | 1132 | 3.96653728 | 0.004432185 | 26 | 0.001117394 | 6.554835658 | 8 mo | HOM |
| **1010419538** | 2 | P-sen | 1132 | 3.96653728 | 0 | 0 | 0 | 0 | 4 mo | WILD |
| **1010419187** | 1 | P-sen | 1132 | 3.96653728 | 0.000423632 | 10 | 0.000106801 | 2.521090638 | 8 mo | HOM |
| **1013978366** | 1 | P-sen | 1132 | 3.96653728 | 1.1E-05 | 2 | 2.78E-06 | 0.504218128 | 2 mo | HOM |
| **1014416155** | 2 | P-sen | 1132 | 3.96653728 | 0 | 0 | 0 | 0 | 2 mo | HOM |

| | | | | | | | | | | |
|---|---|---|---|---|---|---|---|---|---|---|
| 1013978065 | 2 | P-sen | 1132 | 3.96653728 | 0 | 0 | 0 | 0 | 2 mo | HOM |
| 1009934102 | 2 | P-sen | 1132 | 3.96653728 | 3.15E-08 | 0 | 7.95E-09 | 0 | 4 mo | WILD |
| 1009322937 | 2 | P-sen | 1132 | 3.96653728 | 6.45E-05 | 3 | 1.63E-05 | 0.756327191 | 8 mo | WILD |
| 1011080637 | 2 | P-sen | 1132 | 3.96653728 | 0.000172151 | 9 | 4.34E-05 | 2.268981574 | 8 mo | HOM |
| 1013609713 | 2 | P-sen | 1132 | 3.96653728 | 0 | 0 | 0 | 0 | 2 mo | WILD |
| 1010418842 | 1 | P-sen | 1132 | 3.96653728 | 0.000311287 | 13 | 7.85E-05 | 3.277417829 | 8 mo | HOM |
| 1013288213 | 2 | P-sen | 1132 | 3.96653728 | 7.08E-06 | 1 | 1.79E-06 | 0.252109064 | 4 mo | HOM |
| 1008976060 | 2 | P-sen | 1132 | 3.96653728 | 0.000294117 | 16 | 7.41E-05 | 4.03374502 | 8 mo | WILD |
| 1013446243 | 1 | P-sen | 1132 | 3.96653728 | 6.82E-05 | 5 | 1.72E-05 | 1.260545319 | 4 mo | HOM |
| 1013288951 | 2 | P-sen | 1132 | 3.96653728 | 1.91E-05 | 2 | 4.82E-06 | 0.504218128 | 4 mo | HOM |
| 1013978665 | 1 | NLOT3 | 1139 | 0.060161019 | 0 | 0 | 0 | 0 | 2 mo | HOM |
| 1013445609 | 1 | NLOT3 | 1139 | 0.060161019 | 0 | 0 | 0 | 0 | 4 mo | HOM |
| 1013609015 | 1 | NLOT3 | 1139 | 0.060161019 | 0 | 0 | 0 | 0 | 4 mo | HOM |
| 1013608106 | 2 | NLOT3 | 1139 | 0.060161019 | 0 | 0 | 0 | 0 | 4 mo | HOM |
| 1011937543 | 1 | NLOT3 | 1139 | 0.060161019 | 2.12E-06 | 0 | 3.52E-05 | 0 | 8 mo | HOM |
| 1010419538 | 2 | NLOT3 | 1139 | 0.060161019 | 0 | 0 | 0 | 0 | 4 mo | WILD |
| 1010419187 | 1 | NLOT3 | 1139 | 0.060161019 | 2.66E-05 | 2 | 0.000441897 | 33.2441177 | 8 mo | HOM |
| 1013978366 | 1 | NLOT3 | 1139 | 0.060161019 | 0 | 0 | 0 | 0 | 2 mo | HOM |
| 1014416155 | 2 | NLOT3 | 1139 | 0.060161019 | 0 | 0 | 0 | 0 | 2 mo | HOM |
| 1013978065 | 2 | NLOT3 | 1139 | 0.060161019 | 0 | 0 | 0 | 0 | 2 mo | HOM |
| 1009934102 | 2 | NLOT3 | 1139 | 0.060161019 | 0 | 0 | 0 | 0 | 4 mo | WILD |
| 1009322937 | 2 | NLOT3 | 1139 | 0.060161019 | 0 | 0 | 0 | 0 | 8 mo | WILD |
| 1011080637 | 2 | NLOT3 | 1139 | 0.060161019 | 1.13E-05 | 0 | 0.000187934 | 0 | 8 mo | HOM |
| 1013609713 | 2 | NLOT3 | 1139 | 0.060161019 | 0 | 0 | 0 | 0 | 2 mo | WILD |
| 1010418842 | 1 | NLOT3 | 1139 | 0.060161019 | 5.08E-05 | 2 | 0.00084484 | 33.2441177 | 8 mo | HOM |
| 1013288213 | 2 | NLOT3 | 1139 | 0.060161019 | 0 | 0 | 0 | 0 | 4 mo | HOM |
| 1008976060 | 2 | NLOT3 | 1139 | 0.060161019 | 0 | 0 | 0 | 0 | 8 mo | WILD |
| 1013446243 | 1 | NLOT3 | 1139 | 0.060161019 | 1.41E-05 | 1 | 0.000234858 | 16.62205885 | 4 mo | HOM |
| 1013288951 | 2 | NLOT3 | 1139 | 0.060161019 | 0 | 0 | 0 | 0 | 4 mo | HOM |
| 1013978665 | 1 | ME | 10671 | 0.084014851 | 0 | 0 | 0 | 0 | 2 mo | HOM |
| 1013445609 | 1 | ME | 10671 | 0.084014851 | 0 | 0 | 0 | 0 | 4 mo | HOM |
| 1013609015 | 1 | ME | 10671 | 0.084014851 | 0 | 0 | 0 | 0 | 4 mo | HOM |
| 1013608106 | 2 | ME | 10671 | 0.084014851 | 0 | 0 | 0 | 0 | 4 mo | HOM |
| 1011937543 | 1 | ME | 10671 | 0.084014851 | 0 | 0 | 0 | 0 | 8 mo | HOM |
| 1010419538 | 2 | ME | 10671 | 0.084014851 | 0 | 0 | 0 | 0 | 4 mo | WILD |
| 1010419187 | 1 | ME | 10671 | 0.084014851 | 0.000130283 | 3 | 0.00155071 | 35.70797254 | 8 mo | HOM |
| 1013978366 | 1 | ME | 10671 | 0.084014851 | 0 | 0 | 0 | 0 | 2 mo | HOM |
| 1014416155 | 2 | ME | 10671 | 0.084014851 | 0 | 0 | 0 | 0 | 2 mo | HOM |
| 1013978065 | 2 | ME | 10671 | 0.084014851 | 0 | 0 | 0 | 0 | 2 mo | HOM |
| 1009934102 | 2 | ME | 10671 | 0.084014851 | 0 | 0 | 0 | 0 | 4 mo | WILD |
| 1009322937 | 2 | ME | 10671 | 0.084014851 | 0 | 0 | 0 | 0 | 8 mo | WILD |
| 1011080637 | 2 | ME | 10671 | 0.084014851 | 5.01E-06 | 1 | 5.96E-05 | 11.90265751 | 8 mo | HOM |
| 1013609713 | 2 | ME | 10671 | 0.084014851 | 0 | 0 | 0 | 0 | 2 mo | WILD |
| 1010418842 | 1 | ME | 10671 | 0.084014851 | 6.87E-06 | 2 | 8.17E-05 | 23.80531503 | 8 mo | HOM |
| 1013288213 | 2 | ME | 10671 | 0.084014851 | 0 | 0 | 0 | 0 | 4 mo | HOM |
| 1008976060 | 2 | ME | 10671 | 0.084014851 | 0 | 0 | 0 | 0 | 8 mo | WILD |
| 1013446243 | 1 | ME | 10671 | 0.084014851 | 0 | 0 | 0 | 0 | 4 mo | HOM |
| 1013288951 | 2 | ME | 10671 | 0.084014851 | 2.63E-21 | 0 | 3.12E-20 | 0 | 4 mo | HOM |
| 1013978665 | 1 | DG-mo | 10703 | 4.436810291 | 7.13E-06 | 1 | 1.61E-06 | 0.225387144 | 2 mo | HOM |
| 1013445609 | 1 | DG-mo | 10703 | 4.436810291 | 5.08E-05 | 4 | 1.15E-05 | 0.901548576 | 4 mo | HOM |
| 1013609015 | 1 | DG-mo | 10703 | 4.436810291 | 0.000211763 | 12 | 4.77E-05 | 2.704645728 | 4 mo | HOM |
| 1013608106 | 2 | DG-mo | 10703 | 4.436810291 | 0 | 0 | 0 | 0 | 4 mo | HOM |
| 1011937543 | 1 | DG-mo | 10703 | 4.436810291 | 0.005947134 | 68 | 0.001340408 | 15.32632579 | 8 mo | HOM |
| 1010419538 | 2 | DG-mo | 10703 | 4.436810291 | 8.91E-05 | 6 | 2.01E-05 | 1.352322864 | 4 mo | WILD |
| 1010419187 | 1 | DG-mo | 10703 | 4.436810291 | 0.00938863 | 130 | 0.002116076 | 29.30032872 | 8 mo | HOM |
| 1013978366 | 1 | DG-mo | 10703 | 4.436810291 | 0 | 0 | 0 | 0 | 2 mo | HOM |
| 1014416155 | 2 | DG-mo | 10703 | 4.436810291 | 0 | 0 | 0 | 0 | 2 mo | HOM |
| 1013978065 | 2 | DG-mo | 10703 | 4.436810291 | 3.91E-06 | 1 | 8.81E-07 | 0.225387144 | 2 mo | HOM |
| 1009934102 | 2 | DG-mo | 10703 | 4.436810291 | 0.000122136 | 10 | 2.75E-05 | 2.25387144 | 4 mo | WILD |

| | | | | | | | | | | |
|---|---|---|---|---|---|---|---|---|---|---|
| **1009322937** | 2 | DG-mo | 10703 | 4.436810291 | 5.04E-05 | 4 | 1.14E-05 | 0.901548576 | 8 mo | WILD |
| **1011080637** | 2 | DG-mo | 10703 | 4.436810291 | 0.002028734 | 86 | 0.000457251 | 19.38329438 | 8 mo | HOM |
| **1013609713** | 2 | DG-mo | 10703 | 4.436810291 | 6.14E-06 | 0 | 1.38E-06 | 0 | 2 mo | WILD |
| **1010418842** | 1 | DG-mo | 10703 | 4.436810291 | 0.00487672 | 162 | 0.00109915 | 36.51271733 | 8 mo | HOM |
| **1013288213** | 2 | DG-mo | 10703 | 4.436810291 | 7.96E-05 | 10 | 1.79E-05 | 2.25387144 | 4 mo | HOM |
| **1008976060** | 2 | DG-mo | 10703 | 4.436810291 | 0.000178152 | 16 | 4.02E-05 | 3.606194304 | 8 mo | WILD |
| **1013446243** | 1 | DG-mo | 10703 | 4.436810291 | 0.000274039 | 21 | 6.18E-05 | 4.733130024 | 4 mo | HOM |
| **1013288951** | 2 | DG-mo | 10703 | 4.436810291 | 0.000280251 | 14 | 6.32E-05 | 3.155420016 | 4 mo | HOM |
| **1013978665** | 1 | DG-po | 10704 | 0.706728176 | 0 | 0 | 0 | 0 | 2 mo | HOM |
| **1013445609** | 1 | DG-po | 10704 | 0.706728176 | 0 | 0 | 0 | 0 | 4 mo | HOM |
| **1013609015** | 1 | DG-po | 10704 | 0.706728176 | 3.5E-05 | 1 | 4.95E-05 | 1.414971178 | 4 mo | HOM |
| **1013608106** | 2 | DG-po | 10704 | 0.706728176 | 0 | 0 | 0 | 0 | 4 mo | HOM |
| **1011937543** | 1 | DG-po | 10704 | 0.706728176 | 0.001412744 | 10 | 0.001998992 | 14.14971178 | 8 mo | HOM |
| **1010419538** | 2 | DG-po | 10704 | 0.706728176 | 7.8E-07 | 0 | 1.1E-06 | 0 | 4 mo | WILD |
| **1010419187** | 1 | DG-po | 10704 | 0.706728176 | 0.000339146 | 13 | 0.000479882 | 18.39462532 | 8 mo | HOM |
| **1013978366** | 1 | DG-po | 10704 | 0.706728176 | 0 | 0 | 0 | 0 | 2 mo | HOM |
| **1014416155** | 2 | DG-po | 10704 | 0.706728176 | 0 | 0 | 0 | 0 | 2 mo | HOM |
| **1013978065** | 2 | DG-po | 10704 | 0.706728176 | 0 | 0 | 0 | 0 | 2 mo | HOM |
| **1009934102** | 2 | DG-po | 10704 | 0.706728176 | 0 | 0 | 0 | 0 | 4 mo | WILD |
| **1009322937** | 2 | DG-po | 10704 | 0.706728176 | 0 | 0 | 0 | 0 | 8 mo | WILD |
| **1011080637** | 2 | DG-po | 10704 | 0.706728176 | 0.000124643 | 8 | 0.000176366 | 11.31976943 | 8 mo | HOM |
| **1013609713** | 2 | DG-po | 10704 | 0.706728176 | 0 | 0 | 0 | 0 | 2 mo | WILD |
| **1010418842** | 1 | DG-po | 10704 | 0.706728176 | 0.000235654 | 14 | 0.000333443 | 19.8095965 | 8 mo | HOM |
| **1013288213** | 2 | DG-po | 10704 | 0.706728176 | 0 | 0 | 0 | 0 | 4 mo | HOM |
| **1008976060** | 2 | DG-po | 10704 | 0.706728176 | 4.49E-07 | 0 | 6.36E-07 | 0 | 8 mo | WILD |
| **1013446243** | 1 | DG-po | 10704 | 0.706728176 | 2.41E-05 | 1 | 3.42E-05 | 1.414971178 | 4 mo | HOM |
| **1013288951** | 2 | DG-po | 10704 | 0.706728176 | 2.34E-05 | 2 | 3.31E-05 | 2.829942357 | 4 mo | HOM |
| **1013978665** | 1 | SSp-un | 182305689 | 1.300486017 | 0 | 0 | 0 | 0 | 2 mo | HOM |
| **1013445609** | 1 | SSp-un | 182305689 | 1.300486017 | 0.000163476 | 8 | 0.000125704 | 6.151546344 | 4 mo | HOM |
| **1013609015** | 1 | SSp-un | 182305689 | 1.300486017 | 0.000426794 | 10 | 0.00032818 | 7.689432929 | 4 mo | HOM |
| **1013608106** | 2 | SSp-un | 182305689 | 1.300486017 | 0 | 0 | 0 | 0 | 4 mo | HOM |
| **1011937543** | 1 | SSp-un | 182305689 | 1.300486017 | 0.001751845 | 36 | 0.001347069 | 27.68195855 | 8 mo | HOM |
| **1010419538** | 2 | SSp-un | 182305689 | 1.300486017 | 5.94E-06 | 1 | 4.57E-06 | 0.768943293 | 4 mo | WILD |
| **1010419187** | 1 | SSp-un | 182305689 | 1.300486017 | 0.002940483 | 78 | 0.002261064 | 59.97757685 | 8 mo | HOM |
| **1013978366** | 1 | SSp-un | 182305689 | 1.300486017 | 0 | 0 | 0 | 0 | 2 mo | HOM |
| **1014416155** | 2 | SSp-un | 182305689 | 1.300486017 | 0 | 0 | 0 | 0 | 2 mo | HOM |
| **1013978065** | 2 | SSp-un | 182305689 | 1.300486017 | 0 | 0 | 0 | 0 | 2 mo | HOM |
| **1009934102** | 2 | SSp-un | 182305689 | 1.300486017 | 0 | 0 | 0 | 0 | 4 mo | WILD |
| **1009322937** | 2 | SSp-un | 182305689 | 1.300486017 | 0 | 0 | 0 | 0 | 8 mo | WILD |
| **1011080637** | 2 | SSp-un | 182305689 | 1.300486017 | 0.001314864 | 44 | 0.001011056 | 33.83350489 | 8 mo | HOM |
| **1013609713** | 2 | SSp-un | 182305689 | 1.300486017 | 0 | 0 | 0 | 0 | 2 mo | WILD |
| **1010418842** | 1 | SSp-un | 182305689 | 1.300486017 | 0.001977343 | 70 | 0.001520464 | 53.82603051 | 8 mo | HOM |
| **1013288213** | 2 | SSp-un | 182305689 | 1.300486017 | 0.000216589 | 13 | 0.000166544 | 9.996262808 | 4 mo | HOM |
| **1008976060** | 2 | SSp-un | 182305689 | 1.300486017 | 1.79E-05 | 2 | 1.38E-05 | 1.537886586 | 8 mo | WILD |
| **1013446243** | 1 | SSp-un | 182305689 | 1.300486017 | 0.00027264 | 20 | 0.000209645 | 15.37886586 | 4 mo | HOM |
| **1013288951** | 2 | SSp-un | 182305689 | 1.300486017 | 0.000238455 | 12 | 0.000183359 | 9.227319515 | 4 mo | HOM |
| **1013978665** | 1 | SSp-un1 | 182305693 | 0.171264604 | 0 | 0 | 0 | 0 | 2 mo | HOM |
| **1013445609** | 1 | SSp-un1 | 182305693 | 0.171264604 | 4.67E-05 | 2 | 0.000272824 | 11.67783624 | 4 mo | HOM |
| **1013609015** | 1 | SSp-un1 | 182305693 | 0.171264604 | 0.000101465 | 1 | 0.000592446 | 5.838918122 | 4 mo | HOM |
| **1013608106** | 2 | SSp-un1 | 182305693 | 0.171264604 | 0 | 0 | 0 | 0 | 4 mo | HOM |
| **1011937543** | 1 | SSp-un1 | 182305693 | 0.171264604 | 0.000399778 | 8 | 0.002334269 | 46.71134498 | 8 mo | HOM |
| **1010419538** | 2 | SSp-un1 | 182305693 | 0.171264604 | 5.94E-06 | 1 | 3.47E-05 | 5.838918122 | 4 mo | WILD |
| **1010419187** | 1 | SSp-un1 | 182305693 | 0.171264604 | 0.000330113 | 11 | 0.0019275 | 64.22809935 | 8 mo | HOM |
| **1013978366** | 1 | SSp-un1 | 182305693 | 0.171264604 | 0 | 0 | 0 | 0 | 2 mo | HOM |
| **1014416155** | 2 | SSp-un1 | 182305693 | 0.171264604 | 0 | 0 | 0 | 0 | 2 mo | HOM |
| **1013978065** | 2 | SSp-un1 | 182305693 | 0.171264604 | 0 | 0 | 0 | 0 | 2 mo | HOM |
| **1009934102** | 2 | SSp-un1 | 182305693 | 0.171264604 | 0 | 0 | 0 | 0 | 4 mo | WILD |
| **1009322937** | 2 | SSp-un1 | 182305693 | 0.171264604 | 0 | 0 | 0 | 0 | 8 mo | WILD |
| **1011080637** | 2 | SSp-un1 | 182305693 | 0.171264604 | 0.000268678 | 6 | 0.00156879 | 35.03350873 | 8 mo | HOM |

| | | | | | | | | | | |
|---|---|---|---|---|---|---|---|---|---|---|
| **1013609713** | 2 | SSp-un1 | 182305693 | 0.171264604 | 0 | 0 | 0 | 0 | 2 mo | WILD |
| **1010418842** | 1 | SSp-un1 | 182305693 | 0.171264604 | 0.000572712 | 18 | 0.003344019 | 105.1005262 | 8 mo | HOM |
| **1013288213** | 2 | SSp-un1 | 182305693 | 0.171264604 | 5.44E-05 | 2 | 0.000317611 | 11.67783624 | 4 mo | HOM |
| **1008976060** | 2 | SSp-un1 | 182305693 | 0.171264604 | 7.83E-06 | 1 | 4.57E-05 | 5.838918122 | 8 mo | WILD |
| **1013446243** | 1 | SSp-un1 | 182305693 | 0.171264604 | 9.2E-05 | 7 | 0.000537389 | 40.87242686 | 4 mo | HOM |
| **1013288951** | 2 | SSp-un1 | 182305693 | 0.171264604 | 7.55E-05 | 4 | 0.000440694 | 23.35567249 | 4 mo | HOM |
| **1013978665** | 1 | SSp-un2/3 | 182305697 | 0.341494877 | 0 | 0 | 0 | 0 | 2 mo | HOM |
| **1013445609** | 1 | SSp-un2/3 | 182305697 | 0.341494877 | 2.11E-05 | 0 | 6.19E-05 | 0 | 4 mo | HOM |
| **1013609015** | 1 | SSp-un2/3 | 182305697 | 0.341494877 | 6.27E-05 | 3 | 0.000183723 | 8.784904852 | 4 mo | HOM |
| **1013608106** | 2 | SSp-un2/3 | 182305697 | 0.341494877 | 0 | 0 | 0 | 0 | 4 mo | HOM |
| **1011937543** | 1 | SSp-un2/3 | 182305697 | 0.341494877 | 0.00025936 | 7 | 0.000759483 | 20.49811132 | 8 mo | HOM |
| **1010419538** | 2 | SSp-un2/3 | 182305697 | 0.341494877 | 0 | 0 | 0 | 0 | 4 mo | WILD |
| **1010419187** | 1 | SSp-un2/3 | 182305697 | 0.341494877 | 0.001132703 | 21 | 0.003316895 | 61.49433396 | 8 mo | HOM |
| **1013978366** | 1 | SSp-un2/3 | 182305697 | 0.341494877 | 0 | 0 | 0 | 0 | 2 mo | HOM |
| **1014416155** | 2 | SSp-un2/3 | 182305697 | 0.341494877 | 0 | 0 | 0 | 0 | 2 mo | HOM |
| **1013978065** | 2 | SSp-un2/3 | 182305697 | 0.341494877 | 0 | 0 | 0 | 0 | 2 mo | HOM |
| **1009934102** | 2 | SSp-un2/3 | 182305697 | 0.341494877 | 0 | 0 | 0 | 0 | 4 mo | WILD |
| **1009322937** | 2 | SSp-un2/3 | 182305697 | 0.341494877 | 0 | 0 | 0 | 0 | 8 mo | WILD |
| **1011080637** | 2 | SSp-un2/3 | 182305697 | 0.341494877 | 0.000357565 | 13 | 0.001047057 | 38.06792102 | 8 mo | HOM |
| **1013609713** | 2 | SSp-un2/3 | 182305697 | 0.341494877 | 0 | 0 | 0 | 0 | 2 mo | WILD |
| **1010418842** | 1 | SSp-un2/3 | 182305697 | 0.341494877 | 0.000313256 | 13 | 0.000917308 | 38.06792102 | 8 mo | HOM |
| **1013288213** | 2 | SSp-un2/3 | 182305697 | 0.341494877 | 2.73E-05 | 2 | 7.98E-05 | 5.856603234 | 4 mo | HOM |
| **1008976060** | 2 | SSp-un2/3 | 182305697 | 0.341494877 | 0 | 0 | 0 | 0 | 8 mo | WILD |
| **1013446243** | 1 | SSp-un2/3 | 182305697 | 0.341494877 | 3.97E-05 | 1 | 0.000116272 | 2.928301617 | 4 mo | HOM |
| **1013288951** | 2 | SSp-un2/3 | 182305697 | 0.341494877 | 0.000113651 | 4 | 0.000332805 | 11.71320647 | 4 mo | HOM |
| **1013978665** | 1 | SSp-un4 | 182305701 | 0.193610192 | 0 | 0 | 0 | 0 | 2 mo | HOM |
| **1013445609** | 1 | SSp-un4 | 182305701 | 0.193610192 | 3.94E-05 | 1 | 0.000203588 | 5.165017356 | 4 mo | HOM |
| **1013609015** | 1 | SSp-un4 | 182305701 | 0.193610192 | 5.49E-05 | 1 | 0.000283577 | 5.165017356 | 4 mo | HOM |
| **1013608106** | 2 | SSp-un4 | 182305701 | 0.193610192 | 0 | 0 | 0 | 0 | 4 mo | HOM |
| **1011937543** | 1 | SSp-un4 | 182305701 | 0.193610192 | 0.000249138 | 4 | 0.001286802 | 20.66006943 | 8 mo | HOM |
| **1010419538** | 2 | SSp-un4 | 182305701 | 0.193610192 | 0 | 0 | 0 | 0 | 4 mo | WILD |
| **1010419187** | 1 | SSp-un4 | 182305701 | 0.193610192 | 0.000365717 | 10 | 0.001888935 | 51.65017356 | 8 mo | HOM |
| **1013978366** | 1 | SSp-un4 | 182305701 | 0.193610192 | 0 | 0 | 0 | 0 | 2 mo | HOM |
| **1014416155** | 2 | SSp-un4 | 182305701 | 0.193610192 | 0 | 0 | 0 | 0 | 2 mo | HOM |
| **1013978065** | 2 | SSp-un4 | 182305701 | 0.193610192 | 0 | 0 | 0 | 0 | 2 mo | HOM |
| **1009934102** | 2 | SSp-un4 | 182305701 | 0.193610192 | 0 | 0 | 0 | 0 | 4 mo | WILD |
| **1009322937** | 2 | SSp-un4 | 182305701 | 0.193610192 | 0 | 0 | 0 | 0 | 8 mo | WILD |
| **1011080637** | 2 | SSp-un4 | 182305701 | 0.193610192 | 0.000138004 | 7 | 0.000712792 | 36.1551215 | 8 mo | HOM |
| **1013609713** | 2 | SSp-un4 | 182305701 | 0.193610192 | 0 | 0 | 0 | 0 | 2 mo | WILD |
| **1010418842** | 1 | SSp-un4 | 182305701 | 0.193610192 | 0.000435926 | 8 | 0.002251567 | 41.32013885 | 8 mo | HOM |
| **1013288213** | 2 | SSp-un4 | 182305701 | 0.193610192 | 7.69E-05 | 6 | 0.000396968 | 30.99010414 | 4 mo | HOM |
| **1008976060** | 2 | SSp-un4 | 182305701 | 0.193610192 | 0 | 0 | 0 | 0 | 8 mo | WILD |
| **1013446243** | 1 | SSp-un4 | 182305701 | 0.193610192 | 3.09E-05 | 3 | 0.000159844 | 15.49505207 | 4 mo | HOM |
| **1013288951** | 2 | SSp-un4 | 182305701 | 0.193610192 | 1.56E-08 | 0 | 8.04E-08 | 0 | 4 mo | HOM |
| **1013978665** | 1 | SSp-un5 | 182305705 | 0.265775392 | 0 | 0 | 0 | 0 | 2 mo | HOM |
| **1013445609** | 1 | SSp-un5 | 182305705 | 0.265775392 | 2.59E-05 | 2 | 9.76E-05 | 7.525151151 | 4 mo | HOM |
| **1013609015** | 1 | SSp-un5 | 182305705 | 0.265775392 | 8.78E-05 | 4 | 0.000330422 | 15.0503023 | 4 mo | HOM |
| **1013608106** | 2 | SSp-un5 | 182305705 | 0.265775392 | 0 | 0 | 0 | 0 | 4 mo | HOM |
| **1011937543** | 1 | SSp-un5 | 182305705 | 0.265775392 | 0.000310382 | 7 | 0.001167836 | 26.33802903 | 8 mo | HOM |
| **1010419538** | 2 | SSp-un5 | 182305705 | 0.265775392 | 0 | 0 | 0 | 0 | 4 mo | WILD |
| **1010419187** | 1 | SSp-un5 | 182305705 | 0.265775392 | 0.000640832 | 20 | 0.002411178 | 75.25151151 | 8 mo | HOM |
| **1013978366** | 1 | SSp-un5 | 182305705 | 0.265775392 | 0 | 0 | 0 | 0 | 2 mo | HOM |
| **1014416155** | 2 | SSp-un5 | 182305705 | 0.265775392 | 0 | 0 | 0 | 0 | 2 mo | HOM |
| **1013978065** | 2 | SSp-un5 | 182305705 | 0.265775392 | 0 | 0 | 0 | 0 | 2 mo | HOM |
| **1009934102** | 2 | SSp-un5 | 182305705 | 0.265775392 | 0 | 0 | 0 | 0 | 4 mo | WILD |
| **1009322937** | 2 | SSp-un5 | 182305705 | 0.265775392 | 0 | 0 | 0 | 0 | 8 mo | WILD |
| **1011080637** | 2 | SSp-un5 | 182305705 | 0.265775392 | 0.000304027 | 9 | 0.001143923 | 33.86318018 | 8 mo | HOM |
| **1013609713** | 2 | SSp-un5 | 182305705 | 0.265775392 | 0 | 0 | 0 | 0 | 2 mo | WILD |
| **1010418842** | 1 | SSp-un5 | 182305705 | 0.265775392 | 0.000385792 | 20 | 0.001451572 | 75.25151151 | 8 mo | HOM |

| | | | | | | | | | | |
|---|---|---|---|---|---|---|---|---|---|---|
| 1013288213 | 2 | SSp-un5 | 182305705 | 0.265775392 | 5.42E-05 | 3 | 0.000203797 | 11.28772673 | 4 mo | HOM |
| 1008976060 | 2 | SSp-un5 | 182305705 | 0.265775392 | 0 | 0 | 0 | 0 | 8 mo | WILD |
| 1013446243 | 1 | SSp-un5 | 182305705 | 0.265775392 | 4.87E-05 | 3 | 0.000183286 | 11.28772673 | 4 mo | HOM |
| 1013288951 | 2 | SSp-un5 | 182305705 | 0.265775392 | 9.56E-06 | 1 | 3.6E-05 | 3.762575575 | 4 mo | HOM |
| 1013978665 | 1 | SSp-un6a | 182305709 | 0.303032444 | 0 | 0 | 0 | 0 | 2 mo | HOM |
| 1013445609 | 1 | SSp-un6a | 182305709 | 0.303032444 | 3.03E-05 | 3 | 9.98E-05 | 9.899930042 | 4 mo | HOM |
| 1013609015 | 1 | SSp-un6a | 182305709 | 0.303032444 | 0.000119867 | 1 | 0.000395558 | 3.299976681 | 4 mo | HOM |
| 1013608106 | 2 | SSp-un6a | 182305709 | 0.303032444 | 0 | 0 | 0 | 0 | 4 mo | HOM |
| 1011937543 | 1 | SSp-un6a | 182305709 | 0.303032444 | 0.000526627 | 10 | 0.001737858 | 32.99976681 | 8 mo | HOM |
| 1010419538 | 2 | SSp-un6a | 182305709 | 0.303032444 | 0 | 0 | 0 | 0 | 4 mo | WILD |
| 1010419187 | 1 | SSp-un6a | 182305709 | 0.303032444 | 0.000463186 | 15 | 0.001528503 | 49.49965021 | 8 mo | HOM |
| 1013978366 | 1 | SSp-un6a | 182305709 | 0.303032444 | 0 | 0 | 0 | 0 | 2 mo | HOM |
| 1014416155 | 2 | SSp-un6a | 182305709 | 0.303032444 | 0 | 0 | 0 | 0 | 2 mo | HOM |
| 1013978065 | 2 | SSp-un6a | 182305709 | 0.303032444 | 0 | 0 | 0 | 0 | 2 mo | HOM |
| 1009934102 | 2 | SSp-un6a | 182305709 | 0.303032444 | 0 | 0 | 0 | 0 | 4 mo | WILD |
| 1009322937 | 2 | SSp-un6a | 182305709 | 0.303032444 | 0 | 0 | 0 | 0 | 8 mo | WILD |
| 1011080637 | 2 | SSp-un6a | 182305709 | 0.303032444 | 0.000239495 | 8 | 0.000790326 | 26.39981345 | 8 mo | HOM |
| 1013609713 | 2 | SSp-un6a | 182305709 | 0.303032444 | 0 | 0 | 0 | 0 | 2 mo | WILD |
| 1010418842 | 1 | SSp-un6a | 182305709 | 0.303032444 | 0.000261501 | 10 | 0.000862947 | 32.99976681 | 8 mo | HOM |
| 1013288213 | 2 | SSp-un6a | 182305709 | 0.303032444 | 2.43E-06 | 0 | 8E-06 | 0 | 4 mo | HOM |
| 1008976060 | 2 | SSp-un6a | 182305709 | 0.303032444 | 1.01E-05 | 1 | 3.34E-05 | 3.299976681 | 8 mo | WILD |
| 1013446243 | 1 | SSp-un6a | 182305709 | 0.303032444 | 5.88E-05 | 6 | 0.000194114 | 19.79986008 | 4 mo | HOM |
| 1013288951 | 2 | SSp-un6a | 182305709 | 0.303032444 | 3.97E-05 | 3 | 0.000131134 | 9.899930042 | 4 mo | HOM |
| 1013978665 | 1 | SSp-un6b | 182305713 | 0.025308507 | 0 | 0 | 0 | 0 | 2 mo | HOM |
| 1013445609 | 1 | SSp-un6b | 182305713 | 0.025308507 | 0 | 0 | 0 | 0 | 4 mo | HOM |
| 1013609015 | 1 | SSp-un6b | 182305713 | 0.025308507 | 0 | 0 | 0 | 0 | 4 mo | HOM |
| 1013608106 | 2 | SSp-un6b | 182305713 | 0.025308507 | 0 | 0 | 0 | 0 | 4 mo | HOM |
| 1011937543 | 1 | SSp-un6b | 182305713 | 0.025308507 | 6.56E-06 | 0 | 0.000259212 | 0 | 8 mo | HOM |
| 1010419538 | 2 | SSp-un6b | 182305713 | 0.025308507 | 0 | 0 | 0 | 0 | 4 mo | WILD |
| 1010419187 | 1 | SSp-un6b | 182305713 | 0.025308507 | 7.93E-06 | 1 | 0.000313438 | 39.5124053 | 8 mo | HOM |
| 1013978366 | 1 | SSp-un6b | 182305713 | 0.025308507 | 0 | 0 | 0 | 0 | 2 mo | HOM |
| 1014416155 | 2 | SSp-un6b | 182305713 | 0.025308507 | 0 | 0 | 0 | 0 | 2 mo | HOM |
| 1013978065 | 2 | SSp-un6b | 182305713 | 0.025308507 | 0 | 0 | 0 | 0 | 2 mo | HOM |
| 1009934102 | 2 | SSp-un6b | 182305713 | 0.025308507 | 0 | 0 | 0 | 0 | 4 mo | WILD |
| 1009322937 | 2 | SSp-un6b | 182305713 | 0.025308507 | 0 | 0 | 0 | 0 | 8 mo | WILD |
| 1011080637 | 2 | SSp-un6b | 182305713 | 0.025308507 | 7.1E-06 | 1 | 0.000280411 | 39.5124053 | 8 mo | HOM |
| 1013609713 | 2 | SSp-un6b | 182305713 | 0.025308507 | 0 | 0 | 0 | 0 | 2 mo | WILD |
| 1010418842 | 1 | SSp-un6b | 182305713 | 0.025308507 | 8.16E-06 | 1 | 0.000322238 | 39.5124053 | 8 mo | HOM |
| 1013288213 | 2 | SSp-un6b | 182305713 | 0.025308507 | 1.48E-06 | 0 | 5.86E-05 | 0 | 4 mo | HOM |
| 1008976060 | 2 | SSp-un6b | 182305713 | 0.025308507 | 0 | 0 | 0 | 0 | 8 mo | WILD |
| 1013446243 | 1 | SSp-un6b | 182305713 | 0.025308507 | 2.41E-06 | 0 | 9.54E-05 | 0 | 4 mo | HOM |
| 1013288951 | 2 | SSp-un6b | 182305713 | 0.025308507 | 1.32E-08 | 0 | 5.23E-07 | 0 | 4 mo | HOM |
| 1013978665 | 1 | VISa | 312782546 | 1.482662737 | 0 | 0 | 0 | 0 | 2 mo | HOM |
| 1013445609 | 1 | VISa | 312782546 | 1.482662737 | 0.000163668 | 9 | 0.000110388 | 6.070159974 | 4 mo | HOM |
| 1013609015 | 1 | VISa | 312782546 | 1.482662737 | 0.000607369 | 19 | 0.000409647 | 12.81478217 | 4 mo | HOM |
| 1013608106 | 2 | VISa | 312782546 | 1.482662737 | 0 | 0 | 0 | 0 | 4 mo | HOM |
| 1011937543 | 1 | VISa | 312782546 | 1.482662737 | 0.002766704 | 26 | 0.001866037 | 17.5360177 | 8 mo | HOM |
| 1010419538 | 2 | VISa | 312782546 | 1.482662737 | 0 | 0 | 0 | 0 | 4 mo | WILD |
| 1010419187 | 1 | VISa | 312782546 | 1.482662737 | 0.002307834 | 82 | 0.001556547 | 55.30590199 | 8 mo | HOM |
| 1013978366 | 1 | VISa | 312782546 | 1.482662737 | 0 | 0 | 0 | 0 | 2 mo | HOM |
| 1014416155 | 2 | VISa | 312782546 | 1.482662737 | 0 | 0 | 0 | 0 | 2 mo | HOM |
| 1013978065 | 2 | VISa | 312782546 | 1.482662737 | 7.87E-10 | 0 | 5.31E-10 | 0 | 2 mo | HOM |
| 1009934102 | 2 | VISa | 312782546 | 1.482662737 | 0 | 0 | 0 | 0 | 4 mo | WILD |
| 1009322937 | 2 | VISa | 312782546 | 1.482662737 | 5.18E-05 | 2 | 3.49E-05 | 1.348924439 | 8 mo | WILD |
| 1011080637 | 2 | VISa | 312782546 | 1.482662737 | 0.000974555 | 48 | 0.0006573 | 32.37418653 | 8 mo | HOM |
| 1013609713 | 2 | VISa | 312782546 | 1.482662737 | 0 | 0 | 0 | 0 | 2 mo | WILD |
| 1010418842 | 1 | VISa | 312782546 | 1.482662737 | 0.002310361 | 74 | 0.001558251 | 49.91020423 | 8 mo | HOM |
| 1013288213 | 2 | VISa | 312782546 | 1.482662737 | 9.05E-05 | 9 | 6.11E-05 | 6.070159974 | 4 mo | HOM |
| 1008976060 | 2 | VISa | 312782546 | 1.482662737 | 1.35E-05 | 0 | 9.09E-06 | 0 | 8 mo | WILD |

| | | | | | | | | | | |
|---|---|---|---|---|---|---|---|---|---|---|
| **1013446243** | 1 | VISa | 312782546 | 1.482662737 | 0.000357566 | 18 | 0.000241165 | 12.14031995 | 4 mo | HOM |
| **1013288951** | 2 | VISa | 312782546 | 1.482662737 | 0.000678506 | 22 | 0.000457627 | 14.83816883 | 4 mo | HOM |
| **1013978665** | 1 | VISa1 | 312782550 | 0.246918154 | 0 | 0 | 0 | 0 | 2 mo | HOM |
| **1013445609** | 1 | VISa1 | 312782550 | 0.246918154 | 2.27E-05 | 2 | 9.18E-05 | 8.099849961 | 4 mo | HOM |
| **1013609015** | 1 | VISa1 | 312782550 | 0.246918154 | 6.01E-05 | 2 | 0.000243234 | 8.099849961 | 4 mo | HOM |
| **1013608106** | 2 | VISa1 | 312782550 | 0.246918154 | 0 | 0 | 0 | 0 | 4 mo | HOM |
| **1011937543** | 1 | VISa1 | 312782550 | 0.246918154 | 0.001436656 | 6 | 0.00581835 | 24.29954988 | 8 mo | HOM |
| **1010419538** | 2 | VISa1 | 312782550 | 0.246918154 | 0 | 0 | 0 | 0 | 4 mo | WILD |
| **1010419187** | 1 | VISa1 | 312782550 | 0.246918154 | 0.000395993 | 14 | 0.001603743 | 56.69894973 | 8 mo | HOM |
| **1013978366** | 1 | VISa1 | 312782550 | 0.246918154 | 0 | 0 | 0 | 0 | 2 mo | HOM |
| **1014416155** | 2 | VISa1 | 312782550 | 0.246918154 | 0 | 0 | 0 | 0 | 2 mo | HOM |
| **1013978065** | 2 | VISa1 | 312782550 | 0.246918154 | 7.87E-10 | 0 | 3.19E-09 | 0 | 2 mo | HOM |
| **1009934102** | 2 | VISa1 | 312782550 | 0.246918154 | 0 | 0 | 0 | 0 | 4 mo | WILD |
| **1009322937** | 2 | VISa1 | 312782550 | 0.246918154 | 4.33E-05 | 2 | 0.000175437 | 8.099849961 | 8 mo | WILD |
| **1011080637** | 2 | VISa1 | 312782550 | 0.246918154 | 0.000239415 | 12 | 0.000969611 | 48.59909976 | 8 mo | HOM |
| **1013609713** | 2 | VISa1 | 312782550 | 0.246918154 | 0 | 0 | 0 | 0 | 2 mo | WILD |
| **1010418842** | 1 | VISa1 | 312782550 | 0.246918154 | 0.000929253 | 18 | 0.003763405 | 72.89864965 | 8 mo | HOM |
| **1013288213** | 2 | VISa1 | 312782550 | 0.246918154 | 1.92E-05 | 3 | 7.76E-05 | 12.14977494 | 4 mo | HOM |
| **1008976060** | 2 | VISa1 | 312782550 | 0.246918154 | 1.35E-05 | 0 | 5.46E-05 | 0 | 8 mo | WILD |
| **1013446243** | 1 | VISa1 | 312782550 | 0.246918154 | 6.22E-05 | 5 | 0.000251944 | 20.2496249 | 4 mo | HOM |
| **1013288951** | 2 | VISa1 | 312782550 | 0.246918154 | 0.000140174 | 7 | 0.000567696 | 28.34947486 | 4 mo | HOM |
| **1013978665** | 1 | VISa2/3 | 312782554 | 0.439615534 | 0 | 0 | 0 | 0 | 2 mo | HOM |
| **1013445609** | 1 | VISa2/3 | 312782554 | 0.439615534 | 5.97E-05 | 2 | 0.000135707 | 4.549429771 | 4 mo | HOM |
| **1013609015** | 1 | VISa2/3 | 312782554 | 0.439615534 | 0.000246182 | 7 | 0.000559994 | 15.9230042 | 4 mo | HOM |
| **1013608106** | 2 | VISa2/3 | 312782554 | 0.439615534 | 0 | 0 | 0 | 0 | 4 mo | HOM |
| **1011937543** | 1 | VISa2/3 | 312782554 | 0.439615534 | 0.000639965 | 7 | 0.001455739 | 15.9230042 | 8 mo | HOM |
| **1010419538** | 2 | VISa2/3 | 312782554 | 0.439615534 | 0 | 0 | 0 | 0 | 4 mo | WILD |
| **1010419187** | 1 | VISa2/3 | 312782554 | 0.439615534 | 0.000821102 | 23 | 0.001867772 | 52.31844237 | 8 mo | HOM |
| **1013978366** | 1 | VISa2/3 | 312782554 | 0.439615534 | 0 | 0 | 0 | 0 | 2 mo | HOM |
| **1014416155** | 2 | VISa2/3 | 312782554 | 0.439615534 | 0 | 0 | 0 | 0 | 2 mo | HOM |
| **1013978065** | 2 | VISa2/3 | 312782554 | 0.439615534 | 0 | 0 | 0 | 0 | 2 mo | HOM |
| **1009934102** | 2 | VISa2/3 | 312782554 | 0.439615534 | 0 | 0 | 0 | 0 | 4 mo | WILD |
| **1009322937** | 2 | VISa2/3 | 312782554 | 0.439615534 | 8.45E-06 | 0 | 1.92E-05 | 0 | 8 mo | WILD |
| **1011080637** | 2 | VISa2/3 | 312782554 | 0.439615534 | 0.000184205 | 10 | 0.000419014 | 22.74714886 | 8 mo | HOM |
| **1013609713** | 2 | VISa2/3 | 312782554 | 0.439615534 | 0 | 0 | 0 | 0 | 2 mo | WILD |
| **1010418842** | 1 | VISa2/3 | 312782554 | 0.439615534 | 0.000442611 | 15 | 0.001006814 | 34.12072328 | 8 mo | HOM |
| **1013288213** | 2 | VISa2/3 | 312782554 | 0.439615534 | 3.33E-05 | 3 | 7.57E-05 | 6.824144657 | 4 mo | HOM |
| **1008976060** | 2 | VISa2/3 | 312782554 | 0.439615534 | 0 | 0 | 0 | 0 | 8 mo | WILD |
| **1013446243** | 1 | VISa2/3 | 312782554 | 0.439615534 | 6.52E-05 | 2 | 0.000148272 | 4.549429771 | 4 mo | HOM |
| **1013288951** | 2 | VISa2/3 | 312782554 | 0.439615534 | 0.000134989 | 3 | 0.000307061 | 6.824144657 | 4 mo | HOM |
| **1013978665** | 1 | VISa4 | 312782558 | 0.15947883 | 0 | 0 | 0 | 0 | 2 mo | HOM |
| **1013445609** | 1 | VISa4 | 312782558 | 0.15947883 | 1.12E-05 | 1 | 7.03E-05 | 6.270424733 | 4 mo | HOM |
| **1013609015** | 1 | VISa4 | 312782558 | 0.15947883 | 5.65E-05 | 1 | 0.00035401 | 6.270424733 | 4 mo | HOM |
| **1013608106** | 2 | VISa4 | 312782558 | 0.15947883 | 0 | 0 | 0 | 0 | 4 mo | HOM |
| **1011937543** | 1 | VISa4 | 312782558 | 0.15947883 | 0.000116751 | 1 | 0.000732078 | 6.270424733 | 8 mo | HOM |
| **1010419538** | 2 | VISa4 | 312782558 | 0.15947883 | 0 | 0 | 0 | 0 | 4 mo | WILD |
| **1010419187** | 1 | VISa4 | 312782558 | 0.15947883 | 0.000213241 | 8 | 0.001337114 | 50.16339786 | 8 mo | HOM |
| **1013978366** | 1 | VISa4 | 312782558 | 0.15947883 | 0 | 0 | 0 | 0 | 2 mo | HOM |
| **1014416155** | 2 | VISa4 | 312782558 | 0.15947883 | 0 | 0 | 0 | 0 | 2 mo | HOM |
| **1013978065** | 2 | VISa4 | 312782558 | 0.15947883 | 0 | 0 | 0 | 0 | 2 mo | HOM |
| **1009934102** | 2 | VISa4 | 312782558 | 0.15947883 | 0 | 0 | 0 | 0 | 4 mo | WILD |
| **1009322937** | 2 | VISa4 | 312782558 | 0.15947883 | 0 | 0 | 0 | 0 | 8 mo | WILD |
| **1011080637** | 2 | VISa4 | 312782558 | 0.15947883 | 0.000140024 | 5 | 0.000878009 | 31.35212366 | 8 mo | HOM |
| **1013609713** | 2 | VISa4 | 312782558 | 0.15947883 | 0 | 0 | 0 | 0 | 2 mo | WILD |
| **1010418842** | 1 | VISa4 | 312782558 | 0.15947883 | 0.000211695 | 9 | 0.00132742 | 56.4338226 | 8 mo | HOM |
| **1013288213** | 2 | VISa4 | 312782558 | 0.15947883 | 0 | 0 | 0 | 0 | 4 mo | HOM |
| **1008976060** | 2 | VISa4 | 312782558 | 0.15947883 | 0 | 0 | 0 | 0 | 8 mo | WILD |
| **1013446243** | 1 | VISa4 | 312782558 | 0.15947883 | 8.6E-05 | 4 | 0.000539435 | 25.08169893 | 4 mo | HOM |
| **1013288951** | 2 | VISa4 | 312782558 | 0.15947883 | 6.75E-05 | 2 | 0.00042345 | 12.54084947 | 4 mo | HOM |

| | | | | | | | | | | |
|---|---|---|---|---|---|---|---|---|---|---|
| **1013978665** | 1 | VISa5 | 312782562 | 0.365975081 | 0 | 0 | 0 | 0 | 2 mo | HOM |
| **1013445609** | 1 | VISa5 | 312782562 | 0.365975081 | 5.96E-05 | 3 | 0.00016281 | 8.197279425 | 4 mo | HOM |
| **1013609015** | 1 | VISa5 | 312782562 | 0.365975081 | 0.000112312 | 4 | 0.000306885 | 10.9297059 | 4 mo | HOM |
| **1013608106** | 2 | VISa5 | 312782562 | 0.365975081 | 0 | 0 | 0 | 0 | 4 mo | HOM |
| **1011937543** | 1 | VISa5 | 312782562 | 0.365975081 | 0.000253523 | 4 | 0.000692733 | 10.9297059 | 8 mo | HOM |
| **1010419538** | 2 | VISa5 | 312782562 | 0.365975081 | 0 | 0 | 0 | 0 | 4 mo | WILD |
| **1010419187** | 1 | VISa5 | 312782562 | 0.365975081 | 0.000552861 | 23 | 0.001510653 | 62.84580893 | 8 mo | HOM |
| **1013978366** | 1 | VISa5 | 312782562 | 0.365975081 | 0 | 0 | 0 | 0 | 2 mo | HOM |
| **1014416155** | 2 | VISa5 | 312782562 | 0.365975081 | 0 | 0 | 0 | 0 | 2 mo | HOM |
| **1013978065** | 2 | VISa5 | 312782562 | 0.365975081 | 0 | 0 | 0 | 0 | 2 mo | HOM |
| **1009934102** | 2 | VISa5 | 312782562 | 0.365975081 | 0 | 0 | 0 | 0 | 4 mo | WILD |
| **1009322937** | 2 | VISa5 | 312782562 | 0.365975081 | 0 | 0 | 0 | 0 | 8 mo | WILD |
| **1011080637** | 2 | VISa5 | 312782562 | 0.365975081 | 0.00022144 | 13 | 0.000605068 | 35.52154418 | 8 mo | HOM |
| **1013609713** | 2 | VISa5 | 312782562 | 0.365975081 | 0 | 0 | 0 | 0 | 2 mo | WILD |
| **1010418842** | 1 | VISa5 | 312782562 | 0.365975081 | 0.000446909 | 15 | 0.001221147 | 40.98639713 | 8 mo | HOM |
| **1013288213** | 2 | VISa5 | 312782562 | 0.365975081 | 2.56E-05 | 2 | 7E-05 | 5.46485295 | 4 mo | HOM |
| **1008976060** | 2 | VISa5 | 312782562 | 0.365975081 | 0 | 0 | 0 | 0 | 8 mo | WILD |
| **1013446243** | 1 | VISa5 | 312782562 | 0.365975081 | 0.000127427 | 4 | 0.000348185 | 10.9297059 | 4 mo | HOM |
| **1013288951** | 2 | VISa5 | 312782562 | 0.365975081 | 0.000285291 | 7 | 0.000779536 | 19.12698533 | 4 mo | HOM |
| **1013978665** | 1 | VISa6a | 312782566 | 0.230125482 | 0 | 0 | 0 | 0 | 2 mo | HOM |
| **1013445609** | 1 | VISa6a | 312782566 | 0.230125482 | 1.05E-05 | 1 | 4.58E-05 | 4.345455327 | 4 mo | HOM |
| **1013609015** | 1 | VISa6a | 312782566 | 0.230125482 | 0.000117044 | 4 | 0.000508608 | 17.38182131 | 4 mo | HOM |
| **1013608106** | 2 | VISa6a | 312782566 | 0.230125482 | 0 | 0 | 0 | 0 | 4 mo | HOM |
| **1011937543** | 1 | VISa6a | 312782566 | 0.230125482 | 0.000275536 | 7 | 0.001197329 | 30.41818729 | 8 mo | HOM |
| **1010419538** | 2 | VISa6a | 312782566 | 0.230125482 | 0 | 0 | 0 | 0 | 4 mo | WILD |
| **1010419187** | 1 | VISa6a | 312782566 | 0.230125482 | 0.000304001 | 12 | 0.001321023 | 52.14546393 | 8 mo | HOM |
| **1013978366** | 1 | VISa6a | 312782566 | 0.230125482 | 0 | 0 | 0 | 0 | 2 mo | HOM |
| **1014416155** | 2 | VISa6a | 312782566 | 0.230125482 | 0 | 0 | 0 | 0 | 2 mo | HOM |
| **1013978065** | 2 | VISa6a | 312782566 | 0.230125482 | 0 | 0 | 0 | 0 | 2 mo | HOM |
| **1009934102** | 2 | VISa6a | 312782566 | 0.230125482 | 0 | 0 | 0 | 0 | 4 mo | WILD |
| **1009322937** | 2 | VISa6a | 312782566 | 0.230125482 | 0 | 0 | 0 | 0 | 8 mo | WILD |
| **1011080637** | 2 | VISa6a | 312782566 | 0.230125482 | 0.000165635 | 7 | 0.00071976 | 30.41818729 | 8 mo | HOM |
| **1013609713** | 2 | VISa6a | 312782566 | 0.230125482 | 0 | 0 | 0 | 0 | 2 mo | WILD |
| **1010418842** | 1 | VISa6a | 312782566 | 0.230125482 | 0.000268741 | 16 | 0.001167803 | 69.52728524 | 8 mo | HOM |
| **1013288213** | 2 | VISa6a | 312782566 | 0.230125482 | 5.46E-06 | 0 | 2.37E-05 | 0 | 4 mo | HOM |
| **1008976060** | 2 | VISa6a | 312782566 | 0.230125482 | 0 | 0 | 0 | 0 | 8 mo | WILD |
| **1013446243** | 1 | VISa6a | 312782566 | 0.230125482 | 1.67E-05 | 3 | 7.26E-05 | 13.03636598 | 4 mo | HOM |
| **1013288951** | 2 | VISa6a | 312782566 | 0.230125482 | 4.32E-05 | 3 | 0.000187713 | 13.03636598 | 4 mo | HOM |
| **1013978665** | 1 | VISa6b | 312782570 | 0.040549656 | 0 | 0 | 0 | 0 | 2 mo | HOM |
| **1013445609** | 1 | VISa6b | 312782570 | 0.040549656 | 0 | 0 | 0 | 0 | 4 mo | HOM |
| **1013609015** | 1 | VISa6b | 312782570 | 0.040549656 | 1.53E-05 | 1 | 0.000377681 | 24.66112166 | 4 mo | HOM |
| **1013608106** | 2 | VISa6b | 312782570 | 0.040549656 | 0 | 0 | 0 | 0 | 4 mo | HOM |
| **1011937543** | 1 | VISa6b | 312782570 | 0.040549656 | 4.43E-05 | 1 | 0.001091809 | 24.66112166 | 8 mo | HOM |
| **1010419538** | 2 | VISa6b | 312782570 | 0.040549656 | 0 | 0 | 0 | 0 | 4 mo | WILD |
| **1010419187** | 1 | VISa6b | 312782570 | 0.040549656 | 2.06E-05 | 2 | 0.000508895 | 49.32224333 | 8 mo | HOM |
| **1013978366** | 1 | VISa6b | 312782570 | 0.040549656 | 0 | 0 | 0 | 0 | 2 mo | HOM |
| **1014416155** | 2 | VISa6b | 312782570 | 0.040549656 | 0 | 0 | 0 | 0 | 2 mo | HOM |
| **1013978065** | 2 | VISa6b | 312782570 | 0.040549656 | 0 | 0 | 0 | 0 | 2 mo | HOM |
| **1009934102** | 2 | VISa6b | 312782570 | 0.040549656 | 0 | 0 | 0 | 0 | 4 mo | WILD |
| **1009322937** | 2 | VISa6b | 312782570 | 0.040549656 | 0 | 0 | 0 | 0 | 8 mo | WILD |
| **1011080637** | 2 | VISa6b | 312782570 | 0.040549656 | 2.38E-05 | 1 | 0.000587836 | 24.66112166 | 8 mo | HOM |
| **1013609713** | 2 | VISa6b | 312782570 | 0.040549656 | 0 | 0 | 0 | 0 | 2 mo | WILD |
| **1010418842** | 1 | VISa6b | 312782570 | 0.040549656 | 1.12E-05 | 1 | 0.000274998 | 24.66112166 | 8 mo | HOM |
| **1013288213** | 2 | VISa6b | 312782570 | 0.040549656 | 7E-06 | 1 | 0.00017251 | 24.66112166 | 4 mo | HOM |
| **1008976060** | 2 | VISa6b | 312782570 | 0.040549656 | 0 | 0 | 0 | 0 | 8 mo | WILD |
| **1013446243** | 1 | VISa6b | 312782570 | 0.040549656 | 0 | 0 | 0 | 0 | 4 mo | HOM |
| **1013288951** | 2 | VISa6b | 312782570 | 0.040549656 | 7.32E-06 | 0 | 0.000180603 | 0 | 4 mo | HOM |
| **1013978665** | 1 | VISli | 312782574 | 0.506959292 | 2.66E-05 | 3 | 5.25E-05 | 5.917634902 | 2 mo | HOM |
| **1013445609** | 1 | VISli | 312782574 | 0.506959292 | 9.67E-05 | 6 | 0.000190671 | 11.8352698 | 4 mo | HOM |

| | | | | | | | | | | |
|---|---|---|---|---|---|---|---|---|---|---|
| **1013609015** | 1 | VISli | 312782574 | 0.506959292 | 0.000231412 | 12 | 0.000456471 | 23.67053961 | 4 mo | HOM |
| **1013608106** | 2 | VISli | 312782574 | 0.506959292 | 0 | 0 | 0 | 0 | 4 mo | HOM |
| **1011937543** | 1 | VISli | 312782574 | 0.506959292 | 0.001515947 | 23 | 0.002990273 | 45.36853425 | 8 mo | HOM |
| **1010419538** | 2 | VISli | 312782574 | 0.506959292 | 0 | 0 | 0 | 0 | 4 mo | WILD |
| **1010419187** | 1 | VISli | 312782574 | 0.506959292 | 0.001584996 | 30 | 0.003126476 | 59.17634902 | 8 mo | HOM |
| **1013978366** | 1 | VISli | 312782574 | 0.506959292 | 0 | 0 | 0 | 0 | 2 mo | HOM |
| **1014416155** | 2 | VISli | 312782574 | 0.506959292 | 0 | 0 | 0 | 0 | 2 mo | HOM |
| **1013978065** | 2 | VISli | 312782574 | 0.506959292 | 0 | 0 | 0 | 0 | 2 mo | HOM |
| **1009934102** | 2 | VISli | 312782574 | 0.506959292 | 0 | 0 | 0 | 0 | 4 mo | WILD |
| **1009322937** | 2 | VISli | 312782574 | 0.506959292 | 0 | 0 | 0 | 0 | 8 mo | WILD |
| **1011080637** | 2 | VISli | 312782574 | 0.506959292 | 0.000325315 | 17 | 0.000641699 | 33.53326444 | 8 mo | HOM |
| **1013609713** | 2 | VISli | 312782574 | 0.506959292 | 0 | 0 | 0 | 0 | 2 mo | WILD |
| **1010418842** | 1 | VISli | 312782574 | 0.506959292 | 0.000682964 | 25 | 0.001347178 | 49.31362418 | 8 mo | HOM |
| **1013288213** | 2 | VISli | 312782574 | 0.506959292 | 3.03E-05 | 3 | 5.98E-05 | 5.917634902 | 4 mo | HOM |
| **1008976060** | 2 | VISli | 312782574 | 0.506959292 | 4.75E-05 | 5 | 9.36E-05 | 9.862724836 | 8 mo | WILD |
| **1013446243** | 1 | VISli | 312782574 | 0.506959292 | 0.000158439 | 9 | 0.000312528 | 17.7529047 | 4 mo | HOM |
| **1013288951** | 2 | VISli | 312782574 | 0.506959292 | 0.00020487 | 7 | 0.000404114 | 13.80781477 | 4 mo | HOM |
| **1013978665** | 1 | VISli1 | 312782578 | 0.076393244 | 2.66E-05 | 3 | 0.00034859 | 39.27048827 | 2 mo | HOM |
| **1013445609** | 1 | VISli1 | 312782578 | 0.076393244 | 4.06E-09 | 0 | 5.32E-08 | 0 | 4 mo | HOM |
| **1013609015** | 1 | VISli1 | 312782578 | 0.076393244 | 0 | 0 | 0 | 0 | 4 mo | HOM |
| **1013608106** | 2 | VISli1 | 312782578 | 0.076393244 | 0 | 0 | 0 | 0 | 4 mo | HOM |
| **1011937543** | 1 | VISli1 | 312782578 | 0.076393244 | 0.000773715 | 2 | 0.010128061 | 26.18032551 | 8 mo | HOM |
| **1010419538** | 2 | VISli1 | 312782578 | 0.076393244 | 0 | 0 | 0 | 0 | 4 mo | WILD |
| **1010419187** | 1 | VISli1 | 312782578 | 0.076393244 | 0.001001223 | 3 | 0.013106172 | 39.27048827 | 8 mo | HOM |
| **1013978366** | 1 | VISli1 | 312782578 | 0.076393244 | 0 | 0 | 0 | 0 | 2 mo | HOM |
| **1014416155** | 2 | VISli1 | 312782578 | 0.076393244 | 0 | 0 | 0 | 0 | 2 mo | HOM |
| **1013978065** | 2 | VISli1 | 312782578 | 0.076393244 | 0 | 0 | 0 | 0 | 2 mo | HOM |
| **1009934102** | 2 | VISli1 | 312782578 | 0.076393244 | 0 | 0 | 0 | 0 | 4 mo | WILD |
| **1009322937** | 2 | VISli1 | 312782578 | 0.076393244 | 0 | 0 | 0 | 0 | 8 mo | WILD |
| **1011080637** | 2 | VISli1 | 312782578 | 0.076393244 | 7.1E-05 | 5 | 0.000929377 | 65.45081378 | 8 mo | HOM |
| **1013609713** | 2 | VISli1 | 312782578 | 0.076393244 | 0 | 0 | 0 | 0 | 2 mo | WILD |
| **1010418842** | 1 | VISli1 | 312782578 | 0.076393244 | 0.000134649 | 4 | 0.00176258 | 52.36065102 | 8 mo | HOM |
| **1013288213** | 2 | VISli1 | 312782578 | 0.076393244 | 1.33E-05 | 1 | 0.000174509 | 13.09016276 | 4 mo | HOM |
| **1008976060** | 2 | VISli1 | 312782578 | 0.076393244 | 4.3E-05 | 4 | 0.000562493 | 52.36065102 | 8 mo | WILD |
| **1013446243** | 1 | VISli1 | 312782578 | 0.076393244 | 4.94E-05 | 2 | 0.000646206 | 26.18032551 | 4 mo | HOM |
| **1013288951** | 2 | VISli1 | 312782578 | 0.076393244 | 1.12E-06 | 0 | 1.47E-05 | 0 | 4 mo | HOM |
| **1013978665** | 1 | VISli2/3 | 312782582 | 0.120795937 | 0 | 0 | 0 | 0 | 2 mo | HOM |
| **1013445609** | 1 | VISli2/3 | 312782582 | 0.120795937 | 1.83E-05 | 1 | 0.00015145 | 8.278424146 | 4 mo | HOM |
| **1013609015** | 1 | VISli2/3 | 312782582 | 0.120795937 | 2.2E-05 | 1 | 0.000181805 | 8.278424146 | 4 mo | HOM |
| **1013608106** | 2 | VISli2/3 | 312782582 | 0.120795937 | 0 | 0 | 0 | 0 | 4 mo | HOM |
| **1011937543** | 1 | VISli2/3 | 312782582 | 0.120795937 | 0.000115715 | 5 | 0.000957941 | 41.39212073 | 8 mo | HOM |
| **1010419538** | 2 | VISli2/3 | 312782582 | 0.120795937 | 0 | 0 | 0 | 0 | 4 mo | WILD |
| **1010419187** | 1 | VISli2/3 | 312782582 | 0.120795937 | 0.000119242 | 7 | 0.000987135 | 57.94896903 | 8 mo | HOM |
| **1013978366** | 1 | VISli2/3 | 312782582 | 0.120795937 | 0 | 0 | 0 | 0 | 2 mo | HOM |
| **1014416155** | 2 | VISli2/3 | 312782582 | 0.120795937 | 0 | 0 | 0 | 0 | 2 mo | HOM |
| **1013978065** | 2 | VISli2/3 | 312782582 | 0.120795937 | 0 | 0 | 0 | 0 | 2 mo | HOM |
| **1009934102** | 2 | VISli2/3 | 312782582 | 0.120795937 | 0 | 0 | 0 | 0 | 4 mo | WILD |
| **1009322937** | 2 | VISli2/3 | 312782582 | 0.120795937 | 0 | 0 | 0 | 0 | 8 mo | WILD |
| **1011080637** | 2 | VISli2/3 | 312782582 | 0.120795937 | 1.01E-05 | 0 | 8.34E-05 | 0 | 8 mo | HOM |
| **1013609713** | 2 | VISli2/3 | 312782582 | 0.120795937 | 0 | 0 | 0 | 0 | 2 mo | WILD |
| **1010418842** | 1 | VISli2/3 | 312782582 | 0.120795937 | 0.000157465 | 4 | 0.001303563 | 33.11369659 | 8 mo | HOM |
| **1013288213** | 2 | VISli2/3 | 312782582 | 0.120795937 | 0 | 0 | 0 | 0 | 4 mo | HOM |
| **1008976060** | 2 | VISli2/3 | 312782582 | 0.120795937 | 0 | 0 | 0 | 0 | 8 mo | WILD |
| **1013446243** | 1 | VISli2/3 | 312782582 | 0.120795937 | 2.95E-05 | 0 | 0.000244395 | 0 | 4 mo | HOM |
| **1013288951** | 2 | VISli2/3 | 312782582 | 0.120795937 | 2.37E-05 | 1 | 0.000196544 | 8.278424146 | 4 mo | HOM |
| **1013978665** | 1 | VISli4 | 312782586 | 0.058986563 | 0 | 0 | 0 | 0 | 2 mo | HOM |
| **1013445609** | 1 | VISli4 | 312782586 | 0.058986563 | 3.33E-05 | 2 | 0.000563834 | 33.90602727 | 4 mo | HOM |
| **1013609015** | 1 | VISli4 | 312782586 | 0.058986563 | 2.31E-08 | 0 | 3.92E-07 | 0 | 4 mo | HOM |
| **1013608106** | 2 | VISli4 | 312782586 | 0.058986563 | 0 | 0 | 0 | 0 | 4 mo | HOM |

| | | | | | | | | | | |
|---|---|---|---|---|---|---|---|---|---|---|
| **1011937543** | 1 | VISli4 | 312782586 | 0.058986563 | 1.1E-05 | 1 | 0.000185828 | 16.95301363 | 8 mo | HOM |
| **1010419538** | 2 | VISli4 | 312782586 | 0.058986563 | 0 | 0 | 0 | 0 | 4 mo | WILD |
| **1010419187** | 1 | VISli4 | 312782586 | 0.058986563 | 0.000101342 | 4 | 0.001718045 | 67.81205454 | 8 mo | HOM |
| **1013978366** | 1 | VISli4 | 312782586 | 0.058986563 | 0 | 0 | 0 | 0 | 2 mo | HOM |
| **1014416155** | 2 | VISli4 | 312782586 | 0.058986563 | 0 | 0 | 0 | 0 | 2 mo | HOM |
| **1013978065** | 2 | VISli4 | 312782586 | 0.058986563 | 0 | 0 | 0 | 0 | 2 mo | HOM |
| **1009934102** | 2 | VISli4 | 312782586 | 0.058986563 | 0 | 0 | 0 | 0 | 4 mo | WILD |
| **1009322937** | 2 | VISli4 | 312782586 | 0.058986563 | 0 | 0 | 0 | 0 | 8 mo | WILD |
| **1011080637** | 2 | VISli4 | 312782586 | 0.058986563 | 1.38E-05 | 1 | 0.000233115 | 16.95301363 | 8 mo | HOM |
| **1013609713** | 2 | VISli4 | 312782586 | 0.058986563 | 0 | 0 | 0 | 0 | 2 mo | WILD |
| **1010418842** | 1 | VISli4 | 312782586 | 0.058986563 | 0.000141084 | 6 | 0.002391802 | 101.7180818 | 8 mo | HOM |
| **1013288213** | 2 | VISli4 | 312782586 | 0.058986563 | 1.65E-07 | 0 | 2.8E-06 | 0 | 4 mo | HOM |
| **1008976060** | 2 | VISli4 | 312782586 | 0.058986563 | 4.24E-06 | 1 | 7.19E-05 | 16.95301363 | 8 mo | WILD |
| **1013446243** | 1 | VISli4 | 312782586 | 0.058986563 | 9.05E-06 | 1 | 0.000153498 | 16.95301363 | 4 mo | HOM |
| **1013288951** | 2 | VISli4 | 312782586 | 0.058986563 | 6.58E-05 | 2 | 0.001115918 | 33.90602727 | 4 mo | HOM |
| **1013978665** | 1 | VISli5 | 312782590 | 0.152417665 | 0 | 0 | 0 | 0 | 2 mo | HOM |
| **1013445609** | 1 | VISli5 | 312782590 | 0.152417665 | 4.51E-05 | 3 | 0.000295931 | 19.68275785 | 4 mo | HOM |
| **1013609015** | 1 | VISli5 | 312782590 | 0.152417665 | 0.000133851 | 6 | 0.000878189 | 39.3655157 | 4 mo | HOM |
| **1013608106** | 2 | VISli5 | 312782590 | 0.152417665 | 0 | 0 | 0 | 0 | 4 mo | HOM |
| **1011937543** | 1 | VISli5 | 312782590 | 0.152417665 | 0.000331441 | 8 | 0.002174555 | 52.48735426 | 8 mo | HOM |
| **1010419538** | 2 | VISli5 | 312782590 | 0.152417665 | 0 | 0 | 0 | 0 | 4 mo | WILD |
| **1010419187** | 1 | VISli5 | 312782590 | 0.152417665 | 0.000230003 | 11 | 0.001509033 | 72.17011211 | 8 mo | HOM |
| **1013978366** | 1 | VISli5 | 312782590 | 0.152417665 | 0 | 0 | 0 | 0 | 2 mo | HOM |
| **1014416155** | 2 | VISli5 | 312782590 | 0.152417665 | 0 | 0 | 0 | 0 | 2 mo | HOM |
| **1013978065** | 2 | VISli5 | 312782590 | 0.152417665 | 0 | 0 | 0 | 0 | 2 mo | HOM |
| **1009934102** | 2 | VISli5 | 312782590 | 0.152417665 | 0 | 0 | 0 | 0 | 4 mo | WILD |
| **1009322937** | 2 | VISli5 | 312782590 | 0.152417665 | 0 | 0 | 0 | 0 | 8 mo | WILD |
| **1011080637** | 2 | VISli5 | 312782590 | 0.152417665 | 0.000113347 | 6 | 0.000743658 | 39.3655157 | 8 mo | HOM |
| **1013609713** | 2 | VISli5 | 312782590 | 0.152417665 | 0 | 0 | 0 | 0 | 2 mo | WILD |
| **1010418842** | 1 | VISli5 | 312782590 | 0.152417665 | 0.000203533 | 9 | 0.001335362 | 59.04827355 | 8 mo | HOM |
| **1013288213** | 2 | VISli5 | 312782590 | 0.152417665 | 1.68E-05 | 2 | 0.000110367 | 13.12183857 | 4 mo | HOM |
| **1008976060** | 2 | VISli5 | 312782590 | 0.152417665 | 2.47E-07 | 0 | 1.62E-06 | 0 | 8 mo | WILD |
| **1013446243** | 1 | VISli5 | 312782590 | 0.152417665 | 5.51E-05 | 4 | 0.000361377 | 26.24367713 | 4 mo | HOM |
| **1013288951** | 2 | VISli5 | 312782590 | 0.152417665 | 6.62E-05 | 2 | 0.000434058 | 13.12183857 | 4 mo | HOM |
| **1013978665** | 1 | VISli6a | 312782594 | 0.081742172 | 0 | 0 | 0 | 0 | 2 mo | HOM |
| **1013445609** | 1 | VISli6a | 312782594 | 0.081742172 | 0 | 0 | 0 | 0 | 4 mo | HOM |
| **1013609015** | 1 | VISli6a | 312782594 | 0.081742172 | 7.56E-05 | 5 | 0.000924569 | 61.16793637 | 4 mo | HOM |
| **1013608106** | 2 | VISli6a | 312782594 | 0.081742172 | 0 | 0 | 0 | 0 | 4 mo | HOM |
| **1011937543** | 1 | VISli6a | 312782594 | 0.081742172 | 0.000255597 | 6 | 0.003126862 | 73.40152364 | 8 mo | HOM |
| **1010419538** | 2 | VISli6a | 312782594 | 0.081742172 | 0 | 0 | 0 | 0 | 4 mo | WILD |
| **1010419187** | 1 | VISli6a | 312782594 | 0.081742172 | 0.000130411 | 4 | 0.001595396 | 48.93434909 | 8 mo | HOM |
| **1013978366** | 1 | VISli6a | 312782594 | 0.081742172 | 0 | 0 | 0 | 0 | 2 mo | HOM |
| **1014416155** | 2 | VISli6a | 312782594 | 0.081742172 | 0 | 0 | 0 | 0 | 2 mo | HOM |
| **1013978065** | 2 | VISli6a | 312782594 | 0.081742172 | 0 | 0 | 0 | 0 | 2 mo | HOM |
| **1009934102** | 2 | VISli6a | 312782594 | 0.081742172 | 0 | 0 | 0 | 0 | 4 mo | WILD |
| **1009322937** | 2 | VISli6a | 312782594 | 0.081742172 | 0 | 0 | 0 | 0 | 8 mo | WILD |
| **1011080637** | 2 | VISli6a | 312782594 | 0.081742172 | 8.81E-05 | 4 | 0.00107787 | 48.93434909 | 8 mo | HOM |
| **1013609713** | 2 | VISli6a | 312782594 | 0.081742172 | 0 | 0 | 0 | 0 | 2 mo | WILD |
| **1010418842** | 1 | VISli6a | 312782594 | 0.081742172 | 3.11E-05 | 1 | 0.000380322 | 12.23358727 | 8 mo | HOM |
| **1013288213** | 2 | VISli6a | 312782594 | 0.081742172 | 0 | 0 | 0 | 0 | 4 mo | HOM |
| **1008976060** | 2 | VISli6a | 312782594 | 0.081742172 | 0 | 0 | 0 | 0 | 8 mo | WILD |
| **1013446243** | 1 | VISli6a | 312782594 | 0.081742172 | 9.42E-06 | 1 | 0.000115288 | 12.23358727 | 4 mo | HOM |
| **1013288951** | 2 | VISli6a | 312782594 | 0.081742172 | 4.54E-05 | 2 | 0.000555451 | 24.46717455 | 4 mo | HOM |
| **1013978665** | 1 | VISli6b | 312782598 | 0.016623711 | 0 | 0 | 0 | 0 | 2 mo | HOM |
| **1013445609** | 1 | VISli6b | 312782598 | 0.016623711 | 0 | 0 | 0 | 0 | 4 mo | HOM |
| **1013609015** | 1 | VISli6b | 312782598 | 0.016623711 | 0 | 0 | 0 | 0 | 4 mo | HOM |
| **1013608106** | 2 | VISli6b | 312782598 | 0.016623711 | 0 | 0 | 0 | 0 | 4 mo | HOM |
| **1011937543** | 1 | VISli6b | 312782598 | 0.016623711 | 2.85E-05 | 1 | 0.001715468 | 60.15504144 | 8 mo | HOM |
| **1010419538** | 2 | VISli6b | 312782598 | 0.016623711 | 0 | 0 | 0 | 0 | 4 mo | WILD |

| | | | | | | | | | | |
|---|---|---|---|---|---|---|---|---|---|---|
| **1010419187** | 1 | VISli6b | 312782598 | 0.016623711 | 2.78E-06 | 1 | 0.000166936 | 60.15504144 | 8 mo | HOM |
| **1013978366** | 1 | VISli6b | 312782598 | 0.016623711 | 0 | 0 | 0 | 0 | 2 mo | HOM |
| **1014416155** | 2 | VISli6b | 312782598 | 0.016623711 | 0 | 0 | 0 | 0 | 2 mo | HOM |
| **1013978065** | 2 | VISli6b | 312782598 | 0.016623711 | 0 | 0 | 0 | 0 | 2 mo | HOM |
| **1009934102** | 2 | VISli6b | 312782598 | 0.016623711 | 0 | 0 | 0 | 0 | 4 mo | WILD |
| **1009322937** | 2 | VISli6b | 312782598 | 0.016623711 | 0 | 0 | 0 | 0 | 8 mo | WILD |
| **1011080637** | 2 | VISli6b | 312782598 | 0.016623711 | 2.9E-05 | 1 | 0.001746738 | 60.15504144 | 8 mo | HOM |
| **1013609713** | 2 | VISli6b | 312782598 | 0.016623711 | 0 | 0 | 0 | 0 | 2 mo | WILD |
| **1010418842** | 1 | VISli6b | 312782598 | 0.016623711 | 1.51E-05 | 1 | 0.000911037 | 60.15504144 | 8 mo | HOM |
| **1013288213** | 2 | VISli6b | 312782598 | 0.016623711 | 0 | 0 | 0 | 0 | 4 mo | HOM |
| **1008976060** | 2 | VISli6b | 312782598 | 0.016623711 | 0 | 0 | 0 | 0 | 8 mo | WILD |
| **1013446243** | 1 | VISli6b | 312782598 | 0.016623711 | 5.99E-06 | 1 | 0.000360506 | 60.15504144 | 4 mo | HOM |
| **1013288951** | 2 | VISli6b | 312782598 | 0.016623711 | 2.62E-06 | 0 | 0.00015754 | 0 | 4 mo | HOM |
| **1013978665** | 1 | VISrl1 | 312782604 | 0.175900612 | 1.22E-05 | 1 | 6.95E-05 | 5.685028556 | 2 mo | HOM |
| **1013445609** | 1 | VISrl1 | 312782604 | 0.175900612 | 4.15E-05 | 3 | 0.000235754 | 17.05508567 | 4 mo | HOM |
| **1013609015** | 1 | VISrl1 | 312782604 | 0.175900612 | 9.9E-05 | 5 | 0.000562995 | 28.42514278 | 4 mo | HOM |
| **1013608106** | 2 | VISrl1 | 312782604 | 0.175900612 | 0 | 0 | 0 | 0 | 4 mo | HOM |
| **1011937543** | 1 | VISrl1 | 312782604 | 0.175900612 | 0.001213199 | 4 | 0.006897072 | 22.74011422 | 8 mo | HOM |
| **1010419538** | 2 | VISrl1 | 312782604 | 0.175900612 | 0 | 0 | 0 | 0 | 4 mo | WILD |
| **1010419187** | 1 | VISrl1 | 312782604 | 0.175900612 | 0.00058178 | 11 | 0.003307438 | 62.53531412 | 8 mo | HOM |
| **1013978366** | 1 | VISrl1 | 312782604 | 0.175900612 | 3.71E-06 | 1 | 2.11E-05 | 5.685028556 | 2 mo | HOM |
| **1014416155** | 2 | VISrl1 | 312782604 | 0.175900612 | 0 | 0 | 0 | 0 | 2 mo | HOM |
| **1013978065** | 2 | VISrl1 | 312782604 | 0.175900612 | 1.46E-06 | 0 | 8.31E-06 | 0 | 2 mo | HOM |
| **1009934102** | 2 | VISrl1 | 312782604 | 0.175900612 | 6.67E-08 | 0 | 3.79E-07 | 0 | 4 mo | WILD |
| **1009322937** | 2 | VISrl1 | 312782604 | 0.175900612 | 0 | 0 | 0 | 0 | 8 mo | WILD |
| **1011080637** | 2 | VISrl1 | 312782604 | 0.175900612 | 0.000427806 | 6 | 0.002432087 | 34.11017134 | 8 mo | HOM |
| **1013609713** | 2 | VISrl1 | 312782604 | 0.175900612 | 1.12E-05 | 1 | 6.34E-05 | 5.685028556 | 2 mo | WILD |
| **1010418842** | 1 | VISrl1 | 312782604 | 0.175900612 | 0.000697257 | 8 | 0.003963929 | 45.48022845 | 8 mo | HOM |
| **1013288213** | 2 | VISrl1 | 312782604 | 0.175900612 | 2.03E-05 | 2 | 0.00011565 | 11.37005711 | 4 mo | HOM |
| **1008976060** | 2 | VISrl1 | 312782604 | 0.175900612 | 5.1E-05 | 1 | 0.000290021 | 5.685028556 | 8 mo | WILD |
| **1013446243** | 1 | VISrl1 | 312782604 | 0.175900612 | 0.00021278 | 5 | 0.001209661 | 28.42514278 | 4 mo | HOM |
| **1013288951** | 2 | VISrl1 | 312782604 | 0.175900612 | 0.000101914 | 3 | 0.000579384 | 17.05508567 | 4 mo | HOM |
| **1013978665** | 1 | VISrl2/3 | 312782608 | 0.28474388 | 0 | 0 | 0 | 0 | 2 mo | HOM |
| **1013445609** | 1 | VISrl2/3 | 312782608 | 0.28474388 | 4.84E-05 | 2 | 0.000169956 | 7.023855955 | 4 mo | HOM |
| **1013609015** | 1 | VISrl2/3 | 312782608 | 0.28474388 | 0.000218324 | 5 | 0.000766738 | 17.55963989 | 4 mo | HOM |
| **1013608106** | 2 | VISrl2/3 | 312782608 | 0.28474388 | 0 | 0 | 0 | 0 | 4 mo | HOM |
| **1011937543** | 1 | VISrl2/3 | 312782608 | 0.28474388 | 0.000336144 | 11 | 0.001180514 | 38.63120775 | 8 mo | HOM |
| **1010419538** | 2 | VISrl2/3 | 312782608 | 0.28474388 | 0 | 0 | 0 | 0 | 4 mo | WILD |
| **1010419187** | 1 | VISrl2/3 | 312782608 | 0.28474388 | 0.000497657 | 13 | 0.001747735 | 45.65506371 | 8 mo | HOM |
| **1013978366** | 1 | VISrl2/3 | 312782608 | 0.28474388 | 0 | 0 | 0 | 0 | 2 mo | HOM |
| **1014416155** | 2 | VISrl2/3 | 312782608 | 0.28474388 | 0 | 0 | 0 | 0 | 2 mo | HOM |
| **1013978065** | 2 | VISrl2/3 | 312782608 | 0.28474388 | 0 | 0 | 0 | 0 | 2 mo | HOM |
| **1009934102** | 2 | VISrl2/3 | 312782608 | 0.28474388 | 0 | 0 | 0 | 0 | 4 mo | WILD |
| **1009322937** | 2 | VISrl2/3 | 312782608 | 0.28474388 | 0 | 0 | 0 | 0 | 8 mo | WILD |
| **1011080637** | 2 | VISrl2/3 | 312782608 | 0.28474388 | 0.000302104 | 8 | 0.001060967 | 28.09542382 | 8 mo | HOM |
| **1013609713** | 2 | VISrl2/3 | 312782608 | 0.28474388 | 0 | 0 | 0 | 0 | 2 mo | WILD |
| **1010418842** | 1 | VISrl2/3 | 312782608 | 0.28474388 | 0.00034751 | 12 | 0.00122043 | 42.14313573 | 8 mo | HOM |
| **1013288213** | 2 | VISrl2/3 | 312782608 | 0.28474388 | 3.68E-05 | 2 | 0.000129246 | 7.023855955 | 4 mo | HOM |
| **1008976060** | 2 | VISrl2/3 | 312782608 | 0.28474388 | 0 | 0 | 0 | 0 | 8 mo | WILD |
| **1013446243** | 1 | VISrl2/3 | 312782608 | 0.28474388 | 6.47E-05 | 5 | 0.000227239 | 17.55963989 | 4 mo | HOM |
| **1013288951** | 2 | VISrl2/3 | 312782608 | 0.28474388 | 3.88E-05 | 2 | 0.000136101 | 7.023855955 | 4 mo | HOM |
| **1013978665** | 1 | VISrl4 | 312782612 | 0.150715739 | 0 | 0 | 0 | 0 | 2 mo | HOM |
| **1013445609** | 1 | VISrl4 | 312782612 | 0.150715739 | 3.39E-05 | 2 | 0.000224954 | 13.27001426 | 4 mo | HOM |
| **1013609015** | 1 | VISrl4 | 312782612 | 0.150715739 | 0.000139348 | 4 | 0.000924576 | 26.54002851 | 4 mo | HOM |
| **1013608106** | 2 | VISrl4 | 312782612 | 0.150715739 | 0 | 0 | 0 | 0 | 4 mo | HOM |
| **1011937543** | 1 | VISrl4 | 312782612 | 0.150715739 | 0.000256571 | 4 | 0.001702353 | 26.54002851 | 8 mo | HOM |
| **1010419538** | 2 | VISrl4 | 312782612 | 0.150715739 | 0 | 0 | 0 | 0 | 4 mo | WILD |
| **1010419187** | 1 | VISrl4 | 312782612 | 0.150715739 | 0.000416472 | 10 | 0.002763295 | 66.35007129 | 8 mo | HOM |
| **1013978366** | 1 | VISrl4 | 312782612 | 0.150715739 | 0 | 0 | 0 | 0 | 2 mo | HOM |

| | | | | | | | | | | |
|---|---|---|---|---|---|---|---|---|---|---|
| **1014416155** | 2 | VlSrl4 | 312782612 | 0.150715739 | 0 | 0 | 0 | 0 | 2 mo | HOM |
| **1013978065** | 2 | VlSrl4 | 312782612 | 0.150715739 | 0 | 0 | 0 | 0 | 2 mo | HOM |
| **1009934102** | 2 | VlSrl4 | 312782612 | 0.150715739 | 0 | 0 | 0 | 0 | 4 mo | WILD |
| **1009322937** | 2 | VlSrl4 | 312782612 | 0.150715739 | 0 | 0 | 0 | 0 | 8 mo | WILD |
| **1011080637** | 2 | VlSrl4 | 312782612 | 0.150715739 | 0.000110198 | 4 | 0.000731162 | 26.54002851 | 8 mo | HOM |
| **1013609713** | 2 | VlSrl4 | 312782612 | 0.150715739 | 0 | 0 | 0 | 0 | 2 mo | WILD |
| **1010418842** | 1 | VlSrl4 | 312782612 | 0.150715739 | 0.000371292 | 7 | 0.002463528 | 46.4450499 | 8 mo | HOM |
| **1013288213** | 2 | VlSrl4 | 312782612 | 0.150715739 | 4.15E-05 | 3 | 0.000275042 | 19.90502139 | 4 mo | HOM |
| **1008976060** | 2 | VlSrl4 | 312782612 | 0.150715739 | 0 | 0 | 0 | 0 | 8 mo | WILD |
| **1013446243** | 1 | VlSrl4 | 312782612 | 0.150715739 | 1.6E-05 | 1 | 0.000106356 | 6.635007129 | 4 mo | HOM |
| **1013288951** | 2 | VlSrl4 | 312782612 | 0.150715739 | 1.34E-05 | 0 | 8.86E-05 | 0 | 4 mo | HOM |
| **1013978665** | 1 | VlSrl5 | 312782616 | 0.251677779 | 0 | 0 | 0 | 0 | 2 mo | HOM |
| **1013445609** | 1 | VlSrl5 | 312782616 | 0.251677779 | 1.58E-05 | 2 | 6.27E-05 | 7.946668989 | 4 mo | HOM |
| **1013609015** | 1 | VlSrl5 | 312782616 | 0.251677779 | 5.06E-05 | 2 | 0.000201204 | 7.946668989 | 4 mo | HOM |
| **1013608106** | 2 | VlSrl5 | 312782616 | 0.251677779 | 0 | 0 | 0 | 0 | 4 mo | HOM |
| **1011937543** | 1 | VlSrl5 | 312782616 | 0.251677779 | 0.000325288 | 6 | 0.001292478 | 23.84000697 | 8 mo | HOM |
| **1010419538** | 2 | VlSrl5 | 312782616 | 0.251677779 | 0 | 0 | 0 | 0 | 4 mo | WILD |
| **1010419187** | 1 | VlSrl5 | 312782616 | 0.251677779 | 0.000584694 | 13 | 0.002323185 | 51.65334843 | 8 mo | HOM |
| **1013978366** | 1 | VlSrl5 | 312782616 | 0.251677779 | 0 | 0 | 0 | 0 | 2 mo | HOM |
| **1014416155** | 2 | VlSrl5 | 312782616 | 0.251677779 | 0 | 0 | 0 | 0 | 2 mo | HOM |
| **1013978065** | 2 | VlSrl5 | 312782616 | 0.251677779 | 0 | 0 | 0 | 0 | 2 mo | HOM |
| **1009934102** | 2 | VlSrl5 | 312782616 | 0.251677779 | 0 | 0 | 0 | 0 | 4 mo | WILD |
| **1009322937** | 2 | VlSrl5 | 312782616 | 0.251677779 | 0 | 0 | 0 | 0 | 8 mo | WILD |
| **1011080637** | 2 | VlSrl5 | 312782616 | 0.251677779 | 0.000154298 | 7 | 0.000613077 | 27.81334146 | 8 mo | HOM |
| **1013609713** | 2 | VlSrl5 | 312782616 | 0.251677779 | 0 | 0 | 0 | 0 | 2 mo | WILD |
| **1010418842** | 1 | VlSrl5 | 312782616 | 0.251677779 | 0.000428357 | 8 | 0.001702004 | 31.78667596 | 8 mo | HOM |
| **1013288213** | 2 | VlSrl5 | 312782616 | 0.251677779 | 1.48E-05 | 3 | 5.88E-05 | 11.92000348 | 4 mo | HOM |
| **1008976060** | 2 | VlSrl5 | 312782616 | 0.251677779 | 0 | 0 | 0 | 0 | 8 mo | WILD |
| **1013446243** | 1 | VlSrl5 | 312782616 | 0.251677779 | 6.4E-05 | 4 | 0.000254446 | 15.89333798 | 4 mo | HOM |
| **1013288951** | 2 | VlSrl5 | 312782616 | 0.251677779 | 0.000172905 | 2 | 0.000687008 | 7.946668989 | 4 mo | HOM |
| **1013978665** | 1 | VlSrl6a | 312782620 | 0.156004906 | 0 | 0 | 0 | 0 | 2 mo | HOM |
| **1013445609** | 1 | VlSrl6a | 312782620 | 0.156004906 | 3.67E-05 | 5 | 0.00023554 | 32.05027403 | 4 mo | HOM |
| **1013609015** | 1 | VlSrl6a | 312782620 | 0.156004906 | 1.88E-05 | 3 | 0.00012047 | 19.23016442 | 4 mo | HOM |
| **1013608106** | 2 | VlSrl6a | 312782620 | 0.156004906 | 0 | 0 | 0 | 0 | 4 mo | HOM |
| **1011937543** | 1 | VlSrl6a | 312782620 | 0.156004906 | 0.000361377 | 6 | 0.002316443 | 38.46032883 | 8 mo | HOM |
| **1010419538** | 2 | VlSrl6a | 312782620 | 0.156004906 | 0 | 0 | 0 | 0 | 4 mo | WILD |
| **1010419187** | 1 | VlSrl6a | 312782620 | 0.156004906 | 0.000329121 | 13 | 0.002109686 | 83.33071248 | 8 mo | HOM |
| **1013978366** | 1 | VlSrl6a | 312782620 | 0.156004906 | 0 | 0 | 0 | 0 | 2 mo | HOM |
| **1014416155** | 2 | VlSrl6a | 312782620 | 0.156004906 | 0 | 0 | 0 | 0 | 2 mo | HOM |
| **1013978065** | 2 | VlSrl6a | 312782620 | 0.156004906 | 0 | 0 | 0 | 0 | 2 mo | HOM |
| **1009934102** | 2 | VlSrl6a | 312782620 | 0.156004906 | 0 | 0 | 0 | 0 | 4 mo | WILD |
| **1009322937** | 2 | VlSrl6a | 312782620 | 0.156004906 | 0 | 0 | 0 | 0 | 8 mo | WILD |
| **1011080637** | 2 | VlSrl6a | 312782620 | 0.156004906 | 6.7E-05 | 4 | 0.000429609 | 25.64021922 | 8 mo | HOM |
| **1013609713** | 2 | VlSrl6a | 312782620 | 0.156004906 | 0 | 0 | 0 | 0 | 2 mo | WILD |
| **1010418842** | 1 | VlSrl6a | 312782620 | 0.156004906 | 0.000181895 | 8 | 0.001165955 | 51.28043845 | 8 mo | HOM |
| **1013288213** | 2 | VlSrl6a | 312782620 | 0.156004906 | 1.63E-05 | 2 | 0.000104603 | 12.82010961 | 4 mo | HOM |
| **1008976060** | 2 | VlSrl6a | 312782620 | 0.156004906 | 0 | 0 | 0 | 0 | 8 mo | WILD |
| **1013446243** | 1 | VlSrl6a | 312782620 | 0.156004906 | 5.84E-05 | 2 | 0.000374644 | 12.82010961 | 4 mo | HOM |
| **1013288951** | 2 | VlSrl6a | 312782620 | 0.156004906 | 0.00010564 | 2 | 0.000677155 | 12.82010961 | 4 mo | HOM |
| **1013978665** | 1 | VlSrl6b | 312782624 | 0.029379957 | 0 | 0 | 0 | 0 | 2 mo | HOM |
| **1013445609** | 1 | VlSrl6b | 312782624 | 0.029379957 | 3.65E-06 | 1 | 0.000124353 | 34.03681004 | 4 mo | HOM |
| **1013609015** | 1 | VlSrl6b | 312782624 | 0.029379957 | 2.88E-06 | 0 | 9.79E-05 | 0 | 4 mo | HOM |
| **1013608106** | 2 | VlSrl6b | 312782624 | 0.029379957 | 0 | 0 | 0 | 0 | 4 mo | HOM |
| **1011937543** | 1 | VlSrl6b | 312782624 | 0.029379957 | 1.43E-05 | 1 | 0.000487177 | 34.03681004 | 8 mo | HOM |
| **1010419538** | 2 | VlSrl6b | 312782624 | 0.029379957 | 0 | 0 | 0 | 0 | 4 mo | WILD |
| **1010419187** | 1 | VlSrl6b | 312782624 | 0.029379957 | 3.86E-05 | 2 | 0.001314621 | 68.07362007 | 8 mo | HOM |
| **1013978366** | 1 | VlSrl6b | 312782624 | 0.029379957 | 0 | 0 | 0 | 0 | 2 mo | HOM |
| **1014416155** | 2 | VlSrl6b | 312782624 | 0.029379957 | 0 | 0 | 0 | 0 | 2 mo | HOM |
| **1013978065** | 2 | VlSrl6b | 312782624 | 0.029379957 | 0 | 0 | 0 | 0 | 2 mo | HOM |

| | | | | | | | | | | |
|---|---|---|---|---|---|---|---|---|---|---|
| **1009934102** | 2 | VISrl6b | 312782624 | 0.029379957 | 0 | 0 | 0 | 0 | 4 mo | WILD |
| **1009322937** | 2 | VISrl6b | 312782624 | 0.029379957 | 0 | 0 | 0 | 0 | 8 mo | WILD |
| **1011080637** | 2 | VISrl6b | 312782624 | 0.029379957 | 7.85E-06 | 2 | 0.00026717 | 68.07362007 | 8 mo | HOM |
| **1013609713** | 2 | VISrl6b | 312782624 | 0.029379957 | 0 | 0 | 0 | 0 | 2 mo | WILD |
| **1010418842** | 1 | VISrl6b | 312782624 | 0.029379957 | 3.05E-05 | 3 | 0.001039129 | 102.1104301 | 8 mo | HOM |
| **1013288213** | 2 | VISrl6b | 312782624 | 0.029379957 | 1.04E-06 | 0 | 3.55E-05 | 0 | 4 mo | HOM |
| **1008976060** | 2 | VISrl6b | 312782624 | 0.029379957 | 0 | 0 | 0 | 0 | 8 mo | WILD |
| **1013446243** | 1 | VISrl6b | 312782624 | 0.029379957 | 0 | 0 | 0 | 0 | 4 mo | HOM |
| **1013288951** | 2 | VISrl6b | 312782624 | 0.029379957 | 1.68E-07 | 0 | 5.71E-06 | 0 | 4 mo | HOM |
| **1013978665** | 1 | VISpor | 312782628 | 1.321164654 | 0 | 0 | 0 | 0 | 2 mo | HOM |
| **1013445609** | 1 | VISpor | 312782628 | 1.321164654 | 0.000105989 | 8 | 8.02E-05 | 6.055263421 | 4 mo | HOM |
| **1013609015** | 1 | VISpor | 312782628 | 1.321164654 | 0.000380741 | 10 | 0.000288186 | 7.569079276 | 4 mo | HOM |
| **1013608106** | 2 | VISpor | 312782628 | 1.321164654 | 0 | 0 | 0 | 0 | 4 mo | HOM |
| **1011937543** | 1 | VISpor | 312782628 | 1.321164654 | 0.005872524 | 25 | 0.00444496 | 18.92269819 | 8 mo | HOM |
| **1010419538** | 2 | VISpor | 312782628 | 1.321164654 | 5.2E-05 | 1 | 3.94E-05 | 0.756907928 | 4 mo | WILD |
| **1010419187** | 1 | VISpor | 312782628 | 1.321164654 | 0.004772774 | 82 | 0.003612551 | 62.06645007 | 8 mo | HOM |
| **1013978366** | 1 | VISpor | 312782628 | 1.321164654 | 0 | 0 | 0 | 0 | 2 mo | HOM |
| **1014416155** | 2 | VISpor | 312782628 | 1.321164654 | 0 | 0 | 0 | 0 | 2 mo | HOM |
| **1013978065** | 2 | VISpor | 312782628 | 1.321164654 | 0 | 0 | 0 | 0 | 2 mo | HOM |
| **1009934102** | 2 | VISpor | 312782628 | 1.321164654 | 3.29E-07 | 0 | 2.49E-07 | 0 | 4 mo | WILD |
| **1009322937** | 2 | VISpor | 312782628 | 1.321164654 | 6.66E-06 | 2 | 5.04E-06 | 1.513815855 | 8 mo | WILD |
| **1011080637** | 2 | VISpor | 312782628 | 1.321164654 | 0.001618132 | 55 | 0.001224777 | 41.62993602 | 8 mo | HOM |
| **1013609713** | 2 | VISpor | 312782628 | 1.321164654 | 3.05E-06 | 1 | 2.31E-06 | 0.756907928 | 2 mo | WILD |
| **1010418842** | 1 | VISpor | 312782628 | 1.321164654 | 0.001739563 | 65 | 0.001316689 | 49.1990153 | 8 mo | HOM |
| **1013288213** | 2 | VISpor | 312782628 | 1.321164654 | 9.02E-05 | 6 | 6.83E-05 | 4.541447566 | 4 mo | HOM |
| **1008976060** | 2 | VISpor | 312782628 | 1.321164654 | 2.1E-05 | 4 | 1.59E-05 | 3.027631711 | 8 mo | WILD |
| **1013446243** | 1 | VISpor | 312782628 | 1.321164654 | 0.000403027 | 19 | 0.000305054 | 14.38125063 | 4 mo | HOM |
| **1013288951** | 2 | VISpor | 312782628 | 1.321164654 | 0.000480061 | 25 | 0.000363362 | 18.92269819 | 4 mo | HOM |
| **1013978665** | 1 | VISpor1 | 312782632 | 0.257189495 | 0 | 0 | 0 | 0 | 2 mo | HOM |
| **1013445609** | 1 | VISpor1 | 312782632 | 0.257189495 | 3.48E-05 | 3 | 0.000135221 | 11.66455108 | 4 mo | HOM |
| **1013609015** | 1 | VISpor1 | 312782632 | 0.257189495 | 1.34E-05 | 1 | 5.21E-05 | 3.888183694 | 4 mo | HOM |
| **1013608106** | 2 | VISpor1 | 312782632 | 0.257189495 | 0 | 0 | 0 | 0 | 4 mo | HOM |
| **1011937543** | 1 | VISpor1 | 312782632 | 0.257189495 | 0.005021489 | 6 | 0.019524472 | 23.32910217 | 8 mo | HOM |
| **1010419538** | 2 | VISpor1 | 312782632 | 0.257189495 | 5.2E-05 | 1 | 0.000202226 | 3.888183694 | 4 mo | WILD |
| **1010419187** | 1 | VISpor1 | 312782632 | 0.257189495 | 0.002404538 | 21 | 0.009349287 | 81.65185758 | 8 mo | HOM |
| **1013978366** | 1 | VISpor1 | 312782632 | 0.257189495 | 0 | 0 | 0 | 0 | 2 mo | HOM |
| **1014416155** | 2 | VISpor1 | 312782632 | 0.257189495 | 0 | 0 | 0 | 0 | 2 mo | HOM |
| **1013978065** | 2 | VISpor1 | 312782632 | 0.257189495 | 0 | 0 | 0 | 0 | 2 mo | HOM |
| **1009934102** | 2 | VISpor1 | 312782632 | 0.257189495 | 3.29E-07 | 0 | 1.28E-06 | 0 | 4 mo | WILD |
| **1009322937** | 2 | VISpor1 | 312782632 | 0.257189495 | 6.66E-06 | 2 | 2.59E-05 | 7.776367388 | 8 mo | WILD |
| **1011080637** | 2 | VISpor1 | 312782632 | 0.257189495 | 0.000907121 | 18 | 0.003527053 | 69.9873065 | 8 mo | HOM |
| **1013609713** | 2 | VISpor1 | 312782632 | 0.257189495 | 3.05E-06 | 1 | 1.18E-05 | 3.888183694 | 2 mo | WILD |
| **1010418842** | 1 | VISpor1 | 312782632 | 0.257189495 | 0.000433225 | 19 | 0.001684457 | 73.87549019 | 8 mo | HOM |
| **1013288213** | 2 | VISpor1 | 312782632 | 0.257189495 | 4.26E-05 | 3 | 0.000165499 | 11.66455108 | 4 mo | HOM |
| **1008976060** | 2 | VISpor1 | 312782632 | 0.257189495 | 2.1E-05 | 4 | 8.16E-05 | 15.55273478 | 8 mo | WILD |
| **1013446243** | 1 | VISpor1 | 312782632 | 0.257189495 | 0.000182067 | 9 | 0.000707909 | 34.99365325 | 4 mo | HOM |
| **1013288951** | 2 | VISpor1 | 312782632 | 0.257189495 | 0.000299352 | 17 | 0.001163935 | 66.0991228 | 4 mo | HOM |
| **1013978665** | 1 | VISpor2/3 | 312782636 | 0.385275318 | 0 | 0 | 0 | 0 | 2 mo | HOM |
| **1013445609** | 1 | VISpor2/3 | 312782636 | 0.385275318 | 2.46E-05 | 1 | 6.39E-05 | 2.595546489 | 4 mo | HOM |
| **1013609015** | 1 | VISpor2/3 | 312782636 | 0.385275318 | 0.000111176 | 2 | 0.000288563 | 5.191092978 | 4 mo | HOM |
| **1013608106** | 2 | VISpor2/3 | 312782636 | 0.385275318 | 0 | 0 | 0 | 0 | 4 mo | HOM |
| **1011937543** | 1 | VISpor2/3 | 312782636 | 0.385275318 | 0.000287543 | 8 | 0.000746331 | 20.76437191 | 8 mo | HOM |
| **1010419538** | 2 | VISpor2/3 | 312782636 | 0.385275318 | 0 | 0 | 0 | 0 | 4 mo | WILD |
| **1010419187** | 1 | VISpor2/3 | 312782636 | 0.385275318 | 0.001072754 | 14 | 0.002784382 | 36.33765085 | 8 mo | HOM |
| **1013978366** | 1 | VISpor2/3 | 312782636 | 0.385275318 | 0 | 0 | 0 | 0 | 2 mo | HOM |
| **1014416155** | 2 | VISpor2/3 | 312782636 | 0.385275318 | 0 | 0 | 0 | 0 | 2 mo | HOM |
| **1013978065** | 2 | VISpor2/3 | 312782636 | 0.385275318 | 0 | 0 | 0 | 0 | 2 mo | HOM |
| **1009934102** | 2 | VISpor2/3 | 312782636 | 0.385275318 | 0 | 0 | 0 | 0 | 4 mo | WILD |
| **1009322937** | 2 | VISpor2/3 | 312782636 | 0.385275318 | 0 | 0 | 0 | 0 | 8 mo | WILD |

| | | | | | | | | | | |
|---|---|---|---|---|---|---|---|---|---|---|
| **1011080637** | 2 | VISpor2/3 | 312782636 | 0.385275318 | 0.00019387 | 11 | 0.000503198 | 28.55101138 | 8 mo | HOM |
| **1013609713** | 2 | VISpor2/3 | 312782636 | 0.385275318 | 0 | 0 | 0 | 0 | 2 mo | WILD |
| **1010418842** | 1 | VISpor2/3 | 312782636 | 0.385275318 | 0.000337426 | 10 | 0.000875804 | 25.95546489 | 8 mo | HOM |
| **1013288213** | 2 | VISpor2/3 | 312782636 | 0.385275318 | 1.38E-06 | 0 | 3.58E-06 | 0 | 4 mo | HOM |
| **1008976060** | 2 | VISpor2/3 | 312782636 | 0.385275318 | 0 | 0 | 0 | 0 | 8 mo | WILD |
| **1013446243** | 1 | VISpor2/3 | 312782636 | 0.385275318 | 4.48E-05 | 1 | 0.000116152 | 2.595546489 | 4 mo | HOM |
| **1013288951** | 2 | VISpor2/3 | 312782636 | 0.385275318 | 4.53E-05 | 2 | 0.00011748 | 5.191092978 | 4 mo | HOM |
| **1013978665** | 1 | VISpor4 | 312782640 | 0.062464602 | 0 | 0 | 0 | 0 | 2 mo | HOM |
| **1013445609** | 1 | VISpor4 | 312782640 | 0.062464602 | 1.5E-05 | 1 | 0.000240914 | 16.00906697 | 4 mo | HOM |
| **1013609015** | 1 | VISpor4 | 312782640 | 0.062464602 | 6.91E-05 | 1 | 0.00110572 | 16.00906697 | 4 mo | HOM |
| **1013608106** | 2 | VISpor4 | 312782640 | 0.062464602 | 0 | 0 | 0 | 0 | 4 mo | HOM |
| **1011937543** | 1 | VISpor4 | 312782640 | 0.062464602 | 0.000109116 | 2 | 0.001746838 | 32.01813394 | 8 mo | HOM |
| **1010419538** | 2 | VISpor4 | 312782640 | 0.062464602 | 0 | 0 | 0 | 0 | 4 mo | WILD |
| **1010419187** | 1 | VISpor4 | 312782640 | 0.062464602 | 0.000110617 | 5 | 0.001770876 | 80.04533486 | 8 mo | HOM |
| **1013978366** | 1 | VISpor4 | 312782640 | 0.062464602 | 0 | 0 | 0 | 0 | 2 mo | HOM |
| **1014416155** | 2 | VISpor4 | 312782640 | 0.062464602 | 0 | 0 | 0 | 0 | 2 mo | HOM |
| **1013978065** | 2 | VISpor4 | 312782640 | 0.062464602 | 0 | 0 | 0 | 0 | 2 mo | HOM |
| **1009934102** | 2 | VISpor4 | 312782640 | 0.062464602 | 0 | 0 | 0 | 0 | 4 mo | WILD |
| **1009322937** | 2 | VISpor4 | 312782640 | 0.062464602 | 0 | 0 | 0 | 0 | 8 mo | WILD |
| **1011080637** | 2 | VISpor4 | 312782640 | 0.062464602 | 4.51E-05 | 2 | 0.000722657 | 32.01813394 | 8 mo | HOM |
| **1013609713** | 2 | VISpor4 | 312782640 | 0.062464602 | 0 | 0 | 0 | 0 | 2 mo | WILD |
| **1010418842** | 1 | VISpor4 | 312782640 | 0.062464602 | 0.000138826 | 7 | 0.002222471 | 112.0634688 | 8 mo | HOM |
| **1013288213** | 2 | VISpor4 | 312782640 | 0.062464602 | 2.84E-05 | 2 | 0.000454423 | 32.01813394 | 4 mo | HOM |
| **1008976060** | 2 | VISpor4 | 312782640 | 0.062464602 | 0 | 0 | 0 | 0 | 8 mo | WILD |
| **1013446243** | 1 | VISpor4 | 312782640 | 0.062464602 | 7.32E-06 | 0 | 0.000117162 | 0 | 4 mo | HOM |
| **1013288951** | 2 | VISpor4 | 312782640 | 0.062464602 | 2.11E-05 | 1 | 0.000338066 | 16.00906697 | 4 mo | HOM |
| **1013978665** | 1 | VISpor5 | 312782644 | 0.396809683 | 0 | 0 | 0 | 0 | 2 mo | HOM |
| **1013445609** | 1 | VISpor5 | 312782644 | 0.396809683 | 2.49E-05 | 2 | 6.28E-05 | 5.040199584 | 4 mo | HOM |
| **1013609015** | 1 | VISpor5 | 312782644 | 0.396809683 | 0.000148277 | 5 | 0.000373673 | 12.60049896 | 4 mo | HOM |
| **1013608106** | 2 | VISpor5 | 312782644 | 0.396809683 | 0 | 0 | 0 | 0 | 4 mo | HOM |
| **1011937543** | 1 | VISpor5 | 312782644 | 0.396809683 | 0.000307035 | 6 | 0.00077376 | 15.12059875 | 8 mo | HOM |
| **1010419538** | 2 | VISpor5 | 312782644 | 0.396809683 | 0 | 0 | 0 | 0 | 4 mo | WILD |
| **1010419187** | 1 | VISpor5 | 312782644 | 0.396809683 | 0.000708284 | 25 | 0.001784947 | 63.00249479 | 8 mo | HOM |
| **1013978366** | 1 | VISpor5 | 312782644 | 0.396809683 | 0 | 0 | 0 | 0 | 2 mo | HOM |
| **1014416155** | 2 | VISpor5 | 312782644 | 0.396809683 | 0 | 0 | 0 | 0 | 2 mo | HOM |
| **1013978065** | 2 | VISpor5 | 312782644 | 0.396809683 | 0 | 0 | 0 | 0 | 2 mo | HOM |
| **1009934102** | 2 | VISpor5 | 312782644 | 0.396809683 | 0 | 0 | 0 | 0 | 4 mo | WILD |
| **1009322937** | 2 | VISpor5 | 312782644 | 0.396809683 | 0 | 0 | 0 | 0 | 8 mo | WILD |
| **1011080637** | 2 | VISpor5 | 312782644 | 0.396809683 | 0.000334314 | 18 | 0.000842505 | 45.36179625 | 8 mo | HOM |
| **1013609713** | 2 | VISpor5 | 312782644 | 0.396809683 | 0 | 0 | 0 | 0 | 2 mo | WILD |
| **1010418842** | 1 | VISpor5 | 312782644 | 0.396809683 | 0.000565075 | 21 | 0.001424045 | 52.92209563 | 8 mo | HOM |
| **1013288213** | 2 | VISpor5 | 312782644 | 0.396809683 | 1.16E-05 | 0 | 2.93E-05 | 0 | 4 mo | HOM |
| **1008976060** | 2 | VISpor5 | 312782644 | 0.396809683 | 0 | 0 | 0 | 0 | 8 mo | WILD |
| **1013446243** | 1 | VISpor5 | 312782644 | 0.396809683 | 0.000116505 | 7 | 0.000293604 | 17.64069854 | 4 mo | HOM |
| **1013288951** | 2 | VISpor5 | 312782644 | 0.396809683 | 9.61E-05 | 4 | 0.000242148 | 10.08039917 | 4 mo | HOM |
| **1013978665** | 1 | VISpor6a | 312782648 | 0.185710412 | 0 | 0 | 0 | 0 | 2 mo | HOM |
| **1013445609** | 1 | VISpor6a | 312782648 | 0.185710412 | 6.63E-06 | 1 | 3.57E-05 | 5.384727713 | 4 mo | HOM |
| **1013609015** | 1 | VISpor6a | 312782648 | 0.185710412 | 3.88E-05 | 1 | 0.000209016 | 5.384727713 | 4 mo | HOM |
| **1013608106** | 2 | VISpor6a | 312782648 | 0.185710412 | 0 | 0 | 0 | 0 | 4 mo | HOM |
| **1011937543** | 1 | VISpor6a | 312782648 | 0.185710412 | 0.000144163 | 3 | 0.000776281 | 16.15418314 | 8 mo | HOM |
| **1010419538** | 2 | VISpor6a | 312782648 | 0.185710412 | 0 | 0 | 0 | 0 | 4 mo | WILD |
| **1010419187** | 1 | VISpor6a | 312782648 | 0.185710412 | 0.000384403 | 15 | 0.002069908 | 80.7709157 | 8 mo | HOM |
| **1013978366** | 1 | VISpor6a | 312782648 | 0.185710412 | 0 | 0 | 0 | 0 | 2 mo | HOM |
| **1014416155** | 2 | VISpor6a | 312782648 | 0.185710412 | 0 | 0 | 0 | 0 | 2 mo | HOM |
| **1013978065** | 2 | VISpor6a | 312782648 | 0.185710412 | 0 | 0 | 0 | 0 | 2 mo | HOM |
| **1009934102** | 2 | VISpor6a | 312782648 | 0.185710412 | 0 | 0 | 0 | 0 | 4 mo | WILD |
| **1009322937** | 2 | VISpor6a | 312782648 | 0.185710412 | 0 | 0 | 0 | 0 | 8 mo | WILD |
| **1011080637** | 2 | VISpor6a | 312782648 | 0.185710412 | 0.000137027 | 6 | 0.000737854 | 32.30836628 | 8 mo | HOM |
| **1013609713** | 2 | VISpor6a | 312782648 | 0.185710412 | 0 | 0 | 0 | 0 | 2 mo | WILD |

| | | | | | | | | | | |
|---|---|---|---|---|---|---|---|---|---|---|
| 1010418842 | 1 | VISpor6a | 312782648 | 0.185710412 | 0.000175744 | 7 | 0.000946336 | 37.69309399 | 8 mo | HOM |
| 1013288213 | 2 | VISpor6a | 312782648 | 0.185710412 | 6.23E-06 | 1 | 3.36E-05 | 5.384727713 | 4 mo | HOM |
| 1008976060 | 2 | VISpor6a | 312782648 | 0.185710412 | 0 | 0 | 0 | 0 | 8 mo | WILD |
| 1013446243 | 1 | VISpor6a | 312782648 | 0.185710412 | 5.74E-06 | 1 | 3.09E-05 | 5.384727713 | 4 mo | HOM |
| 1013288951 | 2 | VISpor6a | 312782648 | 0.185710412 | 1.82E-05 | 1 | 9.82E-05 | 5.384727713 | 4 mo | HOM |
| 1013978665 | 1 | VISpor6b | 312782652 | 0.033715143 | 0 | 0 | 0 | 0 | 2 mo | HOM |
| 1013445609 | 1 | VISpor6b | 312782652 | 0.033715143 | 0 | 0 | 0 | 0 | 4 mo | HOM |
| 1013609015 | 1 | VISpor6b | 312782652 | 0.033715143 | 0 | 0 | 0 | 0 | 4 mo | HOM |
| 1013608106 | 2 | VISpor6b | 312782652 | 0.033715143 | 0 | 0 | 0 | 0 | 4 mo | HOM |
| 1011937543 | 1 | VISpor6b | 312782652 | 0.033715143 | 3.18E-06 | 0 | 9.43E-05 | 0 | 8 mo | HOM |
| 1010419538 | 2 | VISpor6b | 312782652 | 0.033715143 | 0 | 0 | 0 | 0 | 4 mo | WILD |
| 1010419187 | 1 | VISpor6b | 312782652 | 0.033715143 | 9.22E-05 | 2 | 0.002733995 | 59.32052488 | 8 mo | HOM |
| 1013978366 | 1 | VISpor6b | 312782652 | 0.033715143 | 0 | 0 | 0 | 0 | 2 mo | HOM |
| 1014416155 | 2 | VISpor6b | 312782652 | 0.033715143 | 0 | 0 | 0 | 0 | 2 mo | HOM |
| 1013978065 | 2 | VISpor6b | 312782652 | 0.033715143 | 0 | 0 | 0 | 0 | 2 mo | HOM |
| 1009934102 | 2 | VISpor6b | 312782652 | 0.033715143 | 0 | 0 | 0 | 0 | 4 mo | WILD |
| 1009322937 | 2 | VISpor6b | 312782652 | 0.033715143 | 0 | 0 | 0 | 0 | 8 mo | WILD |
| 1011080637 | 2 | VISpor6b | 312782652 | 0.033715143 | 6.59E-07 | 0 | 1.96E-05 | 0 | 8 mo | HOM |
| 1013609713 | 2 | VISpor6b | 312782652 | 0.033715143 | 0 | 0 | 0 | 0 | 2 mo | WILD |
| 1010418842 | 1 | VISpor6b | 312782652 | 0.033715143 | 8.93E-05 | 1 | 0.002647685 | 29.66026244 | 8 mo | HOM |
| 1013288213 | 2 | VISpor6b | 312782652 | 0.033715143 | 0 | 0 | 0 | 0 | 4 mo | HOM |
| 1008976060 | 2 | VISpor6b | 312782652 | 0.033715143 | 0 | 0 | 0 | 0 | 8 mo | WILD |
| 1013446243 | 1 | VISpor6b | 312782652 | 0.033715143 | 4.66E-05 | 1 | 0.001383532 | 29.66026244 | 4 mo | HOM |
| 1013288951 | 2 | VISpor6b | 312782652 | 0.033715143 | 0 | 0 | 0 | 0 | 4 mo | HOM |
| 1013978665 | 1 | ProS | 484682470 | 1.368200579 | 0 | 0 | 0 | 0 | 2 mo | HOM |
| 1013445609 | 1 | ProS | 484682470 | 1.368200579 | 3.66E-05 | 3 | 2.67E-05 | 2.192660963 | 4 mo | HOM |
| 1013609015 | 1 | ProS | 484682470 | 1.368200579 | 0.000201272 | 10 | 0.000147107 | 7.308869878 | 4 mo | HOM |
| 1013608106 | 2 | ProS | 484682470 | 1.368200579 | 0 | 0 | 0 | 0 | 4 mo | HOM |
| 1011937543 | 1 | ProS | 484682470 | 1.368200579 | 0.001942343 | 29 | 0.001419634 | 21.19572265 | 8 mo | HOM |
| 1010419538 | 2 | ProS | 484682470 | 1.368200579 | 0 | 0 | 0 | 0 | 4 mo | WILD |
| 1010419187 | 1 | ProS | 484682470 | 1.368200579 | 0.002112838 | 65 | 0.001544246 | 47.50765421 | 8 mo | HOM |
| 1013978366 | 1 | ProS | 484682470 | 1.368200579 | 0 | 0 | 0 | 0 | 2 mo | HOM |
| 1014416155 | 2 | ProS | 484682470 | 1.368200579 | 0 | 0 | 0 | 0 | 2 mo | HOM |
| 1013978065 | 2 | ProS | 484682470 | 1.368200579 | 0 | 0 | 0 | 0 | 2 mo | HOM |
| 1009934102 | 2 | ProS | 484682470 | 1.368200579 | 0 | 0 | 0 | 0 | 4 mo | WILD |
| 1009322937 | 2 | ProS | 484682470 | 1.368200579 | 0 | 0 | 0 | 0 | 8 mo | WILD |
| 1011080637 | 2 | ProS | 484682470 | 1.368200579 | 0.001262102 | 51 | 0.000922454 | 37.27523638 | 8 mo | HOM |
| 1013609713 | 2 | ProS | 484682470 | 1.368200579 | 0 | 0 | 0 | 0 | 2 mo | WILD |
| 1010418842 | 1 | ProS | 484682470 | 1.368200579 | 0.002801969 | 74 | 0.002047923 | 54.0856371 | 8 mo | HOM |
| 1013288213 | 2 | ProS | 484682470 | 1.368200579 | 3.02E-05 | 2 | 2.21E-05 | 1.461773976 | 4 mo | HOM |
| 1008976060 | 2 | ProS | 484682470 | 1.368200579 | 3.67E-06 | 1 | 2.69E-06 | 0.730886988 | 8 mo | WILD |
| 1013446243 | 1 | ProS | 484682470 | 1.368200579 | 0.000204219 | 16 | 0.000149261 | 11.6941918 | 4 mo | HOM |
| 1013288951 | 2 | ProS | 484682470 | 1.368200579 | 0.000105848 | 7 | 7.74E-05 | 5.116208914 | 4 mo | HOM |
| 1013978665 | 1 | APr | 484682508 | 0.33650652 | 0 | 0 | 0 | 0 | 2 mo | HOM |
| 1013445609 | 1 | APr | 484682508 | 0.33650652 | 1.9E-05 | 1 | 5.64E-05 | 2.971710622 | 4 mo | HOM |
| 1013609015 | 1 | APr | 484682508 | 0.33650652 | 6.98E-05 | 3 | 0.000207463 | 8.915131867 | 4 mo | HOM |
| 1013608106 | 2 | APr | 484682508 | 0.33650652 | 0 | 0 | 0 | 0 | 4 mo | HOM |
| 1011937543 | 1 | APr | 484682508 | 0.33650652 | 0.000402318 | 4 | 0.001195572 | 11.88684249 | 8 mo | HOM |
| 1010419538 | 2 | APr | 484682508 | 0.33650652 | 0.000135963 | 7 | 0.000404043 | 20.80197436 | 4 mo | WILD |
| 1010419187 | 1 | APr | 484682508 | 0.33650652 | 0.001720574 | 6 | 0.005113048 | 17.83026373 | 8 mo | HOM |
| 1013978366 | 1 | APr | 484682508 | 0.33650652 | 1.91E-06 | 0 | 5.68E-06 | 0 | 2 mo | HOM |
| 1014416155 | 2 | APr | 484682508 | 0.33650652 | 0 | 0 | 0 | 0 | 2 mo | HOM |
| 1013978065 | 2 | APr | 484682508 | 0.33650652 | 0 | 0 | 0 | 0 | 2 mo | HOM |
| 1009934102 | 2 | APr | 484682508 | 0.33650652 | 0.00016338 | 2 | 0.000485519 | 5.943421245 | 4 mo | WILD |
| 1009322937 | 2 | APr | 484682508 | 0.33650652 | 0 | 0 | 0 | 0 | 8 mo | WILD |
| 1011080637 | 2 | APr | 484682508 | 0.33650652 | 0.002404592 | 10 | 0.007145753 | 29.71710622 | 8 mo | HOM |
| 1013609713 | 2 | APr | 484682508 | 0.33650652 | 0.000110999 | 2 | 0.000329856 | 5.943421245 | 2 mo | WILD |
| 1010418842 | 1 | APr | 484682508 | 0.33650652 | 0.000420584 | 9 | 0.001249855 | 26.7453956 | 8 mo | HOM |
| 1013288213 | 2 | APr | 484682508 | 0.33650652 | 8.3E-05 | 3 | 0.000246601 | 8.915131867 | 4 mo | HOM |

| | | | | | | | | | | |
|---|---|---|---|---|---|---|---|---|---|---|
| **1008976060** | 2 | APr | 484682508 | 0.33650652 | 5.42E-06 | 1 | 1.61E-05 | 2.971710622 | 8 mo | WILD |
| **1013446243** | 1 | APr | 484682508 | 0.33650652 | 4.32E-05 | 2 | 0.000128471 | 5.943421245 | 4 mo | HOM |
| **1013288951** | 2 | APr | 484682508 | 0.33650652 | 0.000184712 | 3 | 0.00054891 | 8.915131867 | 4 mo | HOM |
| **1013978665** | 1 | scwm | 484682512 | 1.105604483 | 0 | 0 | 0 | 0 | 2 mo | HOM |
| **1013445609** | 1 | scwm | 484682512 | 1.105604483 | 1.07E-06 | 0 | 9.67E-07 | 0 | 4 mo | HOM |
| **1013609015** | 1 | scwm | 484682512 | 1.105604483 | 1.34E-05 | 1 | 1.21E-05 | 0.904482584 | 4 mo | HOM |
| **1013608106** | 2 | scwm | 484682512 | 1.105604483 | 0 | 0 | 0 | 0 | 4 mo | HOM |
| **1011937543** | 1 | scwm | 484682512 | 1.105604483 | 0.000697752 | 21 | 0.000631104 | 18.99413427 | 8 mo | HOM |
| **1010419538** | 2 | scwm | 484682512 | 1.105604483 | 0 | 0 | 0 | 0 | 4 mo | WILD |
| **1010419187** | 1 | scwm | 484682512 | 1.105604483 | 0.000185765 | 8 | 0.000168021 | 7.235860673 | 8 mo | HOM |
| **1013978366** | 1 | scwm | 484682512 | 1.105604483 | 0 | 0 | 0 | 0 | 2 mo | HOM |
| **1014416155** | 2 | scwm | 484682512 | 1.105604483 | 0 | 0 | 0 | 0 | 2 mo | HOM |
| **1013978065** | 2 | scwm | 484682512 | 1.105604483 | 0 | 0 | 0 | 0 | 2 mo | HOM |
| **1009934102** | 2 | scwm | 484682512 | 1.105604483 | 0 | 0 | 0 | 0 | 4 mo | WILD |
| **1009322937** | 2 | scwm | 484682512 | 1.105604483 | 1.95E-06 | 0 | 1.76E-06 | 0 | 8 mo | WILD |
| **1011080637** | 2 | scwm | 484682512 | 1.105604483 | 0.000188623 | 10 | 0.000170606 | 9.044825841 | 8 mo | HOM |
| **1013609713** | 2 | scwm | 484682512 | 1.105604483 | 0 | 0 | 0 | 0 | 2 mo | WILD |
| **1010418842** | 1 | scwm | 484682512 | 1.105604483 | 0.000531578 | 17 | 0.000480803 | 15.37620393 | 8 mo | HOM |
| **1013288213** | 2 | scwm | 484682512 | 1.105604483 | 6.82E-08 | 0 | 6.17E-08 | 0 | 4 mo | HOM |
| **1008976060** | 2 | scwm | 484682512 | 1.105604483 | 1.49E-06 | 0 | 1.35E-06 | 0 | 8 mo | WILD |
| **1013446243** | 1 | scwm | 484682512 | 1.105604483 | 1.68E-06 | 0 | 1.52E-06 | 0 | 4 mo | HOM |
| **1013288951** | 2 | scwm | 484682512 | 1.105604483 | 7.72E-07 | 0 | 6.99E-07 | 0 | 4 mo | HOM |
| **1013978665** | 1 | ccb | 484682516 | 3.13296153 | 0 | 0 | 0 | 0 | 2 mo | HOM |
| **1013445609** | 1 | ccb | 484682516 | 3.13296153 | 1.74E-05 | 0 | 5.56E-06 | 0 | 4 mo | HOM |
| **1013609015** | 1 | ccb | 484682516 | 3.13296153 | 5.8E-05 | 0 | 1.85E-05 | 0 | 4 mo | HOM |
| **1013608106** | 2 | ccb | 484682516 | 3.13296153 | 0 | 0 | 0 | 0 | 4 mo | HOM |
| **1011937543** | 1 | ccb | 484682516 | 3.13296153 | 0.001384103 | 25 | 0.000441788 | 7.979670278 | 8 mo | HOM |
| **1010419538** | 2 | ccb | 484682516 | 3.13296153 | 0 | 0 | 0 | 0 | 4 mo | WILD |
| **1010419187** | 1 | ccb | 484682516 | 3.13296153 | 0.000696537 | 29 | 0.000222325 | 9.256417523 | 8 mo | HOM |
| **1013978366** | 1 | ccb | 484682516 | 3.13296153 | 0 | 0 | 0 | 0 | 2 mo | HOM |
| **1014416155** | 2 | ccb | 484682516 | 3.13296153 | 0 | 0 | 0 | 0 | 2 mo | HOM |
| **1013978065** | 2 | ccb | 484682516 | 3.13296153 | 0 | 0 | 0 | 0 | 2 mo | HOM |
| **1009934102** | 2 | ccb | 484682516 | 3.13296153 | 1.21E-06 | 0 | 3.87E-07 | 0 | 4 mo | WILD |
| **1009322937** | 2 | ccb | 484682516 | 3.13296153 | 0 | 0 | 0 | 0 | 8 mo | WILD |
| **1011080637** | 2 | ccb | 484682516 | 3.13296153 | 0.000556773 | 22 | 0.000177715 | 7.022109845 | 8 mo | HOM |
| **1013609713** | 2 | ccb | 484682516 | 3.13296153 | 0 | 0 | 0 | 0 | 2 mo | WILD |
| **1010418842** | 1 | ccb | 484682516 | 3.13296153 | 0.000905683 | 38 | 0.000289082 | 12.12909882 | 8 mo | HOM |
| **1013288213** | 2 | ccb | 484682516 | 3.13296153 | 0.000192039 | 1 | 6.13E-05 | 0.319186811 | 4 mo | HOM |
| **1008976060** | 2 | ccb | 484682516 | 3.13296153 | 6.89E-05 | 5 | 2.2E-05 | 1.595934056 | 8 mo | WILD |
| **1013446243** | 1 | ccb | 484682516 | 3.13296153 | 2.05E-05 | 1 | 6.54E-06 | 0.319186811 | 4 mo | HOM |
| **1013288951** | 2 | ccb | 484682516 | 3.13296153 | 4.58E-05 | 1 | 1.46E-05 | 0.319186811 | 4 mo | HOM |
| **1013978665** | 1 | or | 484682520 | 1.906703366 | 0 | 0 | 0 | 0 | 2 mo | HOM |
| **1013445609** | 1 | or | 484682520 | 1.906703366 | 2.27E-05 | 2 | 1.19E-05 | 1.048930859 | 4 mo | HOM |
| **1013609015** | 1 | or | 484682520 | 1.906703366 | 2.93E-05 | 1 | 1.54E-05 | 0.52446543 | 4 mo | HOM |
| **1013608106** | 2 | or | 484682520 | 1.906703366 | 0 | 0 | 0 | 0 | 4 mo | HOM |
| **1011937543** | 1 | or | 484682520 | 1.906703366 | 0.001517852 | 26 | 0.000796061 | 13.63610117 | 8 mo | HOM |
| **1010419538** | 2 | or | 484682520 | 1.906703366 | 0 | 0 | 0 | 0 | 4 mo | WILD |
| **1010419187** | 1 | or | 484682520 | 1.906703366 | 0.000388625 | 21 | 0.000203821 | 11.01377402 | 8 mo | HOM |
| **1013978366** | 1 | or | 484682520 | 1.906703366 | 0 | 0 | 0 | 0 | 2 mo | HOM |
| **1014416155** | 2 | or | 484682520 | 1.906703366 | 0 | 0 | 0 | 0 | 2 mo | HOM |
| **1013978065** | 2 | or | 484682520 | 1.906703366 | 0 | 0 | 0 | 0 | 2 mo | HOM |
| **1009934102** | 2 | or | 484682520 | 1.906703366 | 0 | 0 | 0 | 0 | 4 mo | WILD |
| **1009322937** | 2 | or | 484682520 | 1.906703366 | 1.3E-06 | 0 | 6.83E-07 | 0 | 8 mo | WILD |
| **1011080637** | 2 | or | 484682520 | 1.906703366 | 0.000351352 | 19 | 0.000184272 | 9.964843161 | 8 mo | HOM |
| **1013609713** | 2 | or | 484682520 | 1.906703366 | 0 | 0 | 0 | 0 | 2 mo | WILD |
| **1010418842** | 1 | or | 484682520 | 1.906703366 | 0.000818828 | 31 | 0.000429447 | 16.25842831 | 8 mo | HOM |
| **1013288213** | 2 | or | 484682520 | 1.906703366 | 5.72E-06 | 1 | 3E-06 | 0.52446543 | 4 mo | HOM |
| **1008976060** | 2 | or | 484682520 | 1.906703366 | 2.75E-05 | 4 | 1.44E-05 | 2.097861718 | 8 mo | WILD |
| **1013446243** | 1 | or | 484682520 | 1.906703366 | 3.61E-05 | 2 | 1.89E-05 | 1.048930859 | 4 mo | HOM |

| | | | | | | | | | | |
|---|---|---|---|---|---|---|---|---|---|---|
| **1013288951** | 2 | or | 484682520 | 1.906703366 | 0.000138963 | 1 | 7.29E-05 | 0.52446543 | 4 mo | HOM |
| **1013978665** | 1 | ar | 484682524 | 0.41973839 | 0 | 0 | 0 | 0 | 2 mo | HOM |
| **1013445609** | 1 | ar | 484682524 | 0.41973839 | 7.69E-08 | 0 | 1.83E-07 | 0 | 4 mo | HOM |
| **1013609015** | 1 | ar | 484682524 | 0.41973839 | 0 | 0 | 0 | 0 | 4 mo | HOM |
| **1013608106** | 2 | ar | 484682524 | 0.41973839 | 0 | 0 | 0 | 0 | 4 mo | HOM |
| **1011937543** | 1 | ar | 484682524 | 0.41973839 | 0.000651631 | 6 | 0.001552469 | 14.29461812 | 8 mo | HOM |
| **1010419538** | 2 | ar | 484682524 | 0.41973839 | 0 | 0 | 0 | 0 | 4 mo | WILD |
| **1010419187** | 1 | ar | 484682524 | 0.41973839 | 0.000107861 | 6 | 0.000256972 | 14.29461812 | 8 mo | HOM |
| **1013978366** | 1 | ar | 484682524 | 0.41973839 | 0 | 0 | 0 | 0 | 2 mo | HOM |
| **1014416155** | 2 | ar | 484682524 | 0.41973839 | 0 | 0 | 0 | 0 | 2 mo | HOM |
| **1013978065** | 2 | ar | 484682524 | 0.41973839 | 0 | 0 | 0 | 0 | 2 mo | HOM |
| **1009934102** | 2 | ar | 484682524 | 0.41973839 | 0 | 0 | 0 | 0 | 4 mo | WILD |
| **1009322937** | 2 | ar | 484682524 | 0.41973839 | 0 | 0 | 0 | 0 | 8 mo | WILD |
| **1011080637** | 2 | ar | 484682524 | 0.41973839 | 3.32E-05 | 3 | 7.9E-05 | 7.147309059 | 8 mo | HOM |
| **1013609713** | 2 | ar | 484682524 | 0.41973839 | 0 | 0 | 0 | 0 | 2 mo | WILD |
| **1010418842** | 1 | ar | 484682524 | 0.41973839 | 0.000218415 | 10 | 0.000520359 | 23.82436353 | 8 mo | HOM |
| **1013288213** | 2 | ar | 484682524 | 0.41973839 | 5.33E-08 | 0 | 1.27E-07 | 0 | 4 mo | HOM |
| **1008976060** | 2 | ar | 484682524 | 0.41973839 | 2.43E-06 | 0 | 5.78E-06 | 0 | 8 mo | WILD |
| **1013446243** | 1 | ar | 484682524 | 0.41973839 | 2.3E-05 | 0 | 5.47E-05 | 0 | 4 mo | HOM |
| **1013288951** | 2 | ar | 484682524 | 0.41973839 | 8.77E-06 | 0 | 2.09E-05 | 0 | 4 mo | HOM |
| **1013978665** | 1 | stc | 484682528 | 0.036189744 | 0 | 0 | 0 | 0 | 2 mo | HOM |
| **1013445609** | 1 | stc | 484682528 | 0.036189744 | 0 | 0 | 0 | 0 | 4 mo | HOM |
| **1013609015** | 1 | stc | 484682528 | 0.036189744 | 0 | 0 | 0 | 0 | 4 mo | HOM |
| **1013608106** | 2 | stc | 484682528 | 0.036189744 | 0 | 0 | 0 | 0 | 4 mo | HOM |
| **1011937543** | 1 | stc | 484682528 | 0.036189744 | 3.92E-08 | 0 | 1.08E-06 | 0 | 8 mo | HOM |
| **1010419538** | 2 | stc | 484682528 | 0.036189744 | 0 | 0 | 0 | 0 | 4 mo | WILD |
| **1010419187** | 1 | stc | 484682528 | 0.036189744 | 1.4E-05 | 0 | 0.000387591 | 0 | 8 mo | HOM |
| **1013978366** | 1 | stc | 484682528 | 0.036189744 | 0 | 0 | 0 | 0 | 2 mo | HOM |
| **1014416155** | 2 | stc | 484682528 | 0.036189744 | 0 | 0 | 0 | 0 | 2 mo | HOM |
| **1013978065** | 2 | stc | 484682528 | 0.036189744 | 0 | 0 | 0 | 0 | 2 mo | HOM |
| **1009934102** | 2 | stc | 484682528 | 0.036189744 | 0 | 0 | 0 | 0 | 4 mo | WILD |
| **1009322937** | 2 | stc | 484682528 | 0.036189744 | 0 | 0 | 0 | 0 | 8 mo | WILD |
| **1011080637** | 2 | stc | 484682528 | 0.036189744 | 9.69E-06 | 0 | 0.000267782 | 0 | 8 mo | HOM |
| **1013609713** | 2 | stc | 484682528 | 0.036189744 | 0 | 0 | 0 | 0 | 2 mo | WILD |
| **1010418842** | 1 | stc | 484682528 | 0.036189744 | 3.02E-06 | 0 | 8.35E-05 | 0 | 8 mo | HOM |
| **1013288213** | 2 | stc | 484682528 | 0.036189744 | 2.34E-09 | 0 | 6.46E-08 | 0 | 4 mo | HOM |
| **1008976060** | 2 | stc | 484682528 | 0.036189744 | 0 | 0 | 0 | 0 | 8 mo | WILD |
| **1013446243** | 1 | stc | 484682528 | 0.036189744 | 4E-07 | 0 | 1.11E-05 | 0 | 4 mo | HOM |
| **1013288951** | 2 | stc | 484682528 | 0.036189744 | 0 | 0 | 0 | 0 | 4 mo | HOM |
| **1013978665** | 1 | LGd-sh | 496345664 | 0.213569754 | 0 | 0 | 0 | 0 | 2 mo | HOM |
| **1013445609** | 1 | LGd-sh | 496345664 | 0.213569754 | 0 | 0 | 0 | 0 | 4 mo | HOM |
| **1013609015** | 1 | LGd-sh | 496345664 | 0.213569754 | 0 | 0 | 0 | 0 | 4 mo | HOM |
| **1013608106** | 2 | LGd-sh | 496345664 | 0.213569754 | 0 | 0 | 0 | 0 | 4 mo | HOM |
| **1011937543** | 1 | LGd-sh | 496345664 | 0.213569754 | 0.000216637 | 2 | 0.001014361 | 9.364621939 | 8 mo | HOM |
| **1010419538** | 2 | LGd-sh | 496345664 | 0.213569754 | 0 | 0 | 0 | 0 | 4 mo | WILD |
| **1010419187** | 1 | LGd-sh | 496345664 | 0.213569754 | 8.41E-05 | 4 | 0.000393833 | 18.72924388 | 8 mo | HOM |
| **1013978366** | 1 | LGd-sh | 496345664 | 0.213569754 | 0 | 0 | 0 | 0 | 2 mo | HOM |
| **1014416155** | 2 | LGd-sh | 496345664 | 0.213569754 | 0 | 0 | 0 | 0 | 2 mo | HOM |
| **1013978065** | 2 | LGd-sh | 496345664 | 0.213569754 | 0 | 0 | 0 | 0 | 2 mo | HOM |
| **1009934102** | 2 | LGd-sh | 496345664 | 0.213569754 | 0 | 0 | 0 | 0 | 4 mo | WILD |
| **1009322937** | 2 | LGd-sh | 496345664 | 0.213569754 | 3.66E-11 | 0 | 1.72E-10 | 0 | 8 mo | WILD |
| **1011080637** | 2 | LGd-sh | 496345664 | 0.213569754 | 5.66E-06 | 0 | 2.65E-05 | 0 | 8 mo | HOM |
| **1013609713** | 2 | LGd-sh | 496345664 | 0.213569754 | 0 | 0 | 0 | 0 | 2 mo | WILD |
| **1010418842** | 1 | LGd-sh | 496345664 | 0.213569754 | 0.000269069 | 5 | 0.001259865 | 23.41155485 | 8 mo | HOM |
| **1013288213** | 2 | LGd-sh | 496345664 | 0.213569754 | 0 | 0 | 0 | 0 | 4 mo | HOM |
| **1008976060** | 2 | LGd-sh | 496345664 | 0.213569754 | 3.41E-06 | 0 | 1.6E-05 | 0 | 8 mo | WILD |
| **1013446243** | 1 | LGd-sh | 496345664 | 0.213569754 | 0 | 0 | 0 | 0 | 4 mo | HOM |
| **1013288951** | 2 | LGd-sh | 496345664 | 0.213569754 | 0 | 0 | 0 | 0 | 4 mo | HOM |
| **1013978665** | 1 | LGd-co | 496345668 | 0.431841469 | 0 | 0 | 0 | 0 | 2 mo | HOM |

| | | | | | | | | | | |
|---|---|---|---|---|---|---|---|---|---|---|
| **1013445609** | 1 | LGd-co | 496345668 | 0.431841469 | 2.33E-05 | 2 | 5.4E-05 | 4.63132919 | 4 mo | HOM |
| **1013609015** | 1 | LGd-co | 496345668 | 0.431841469 | 0.000180652 | 4 | 0.000418331 | 9.26265838 | 4 mo | HOM |
| **1013608106** | 2 | LGd-co | 496345668 | 0.431841469 | 0 | 0 | 0 | 0 | 4 mo | HOM |
| **1011937543** | 1 | LGd-co | 496345668 | 0.431841469 | 0.000879458 | 13 | 0.00203653 | 30.10363974 | 8 mo | HOM |
| **1010419538** | 2 | LGd-co | 496345668 | 0.431841469 | 0 | 0 | 0 | 0 | 4 mo | WILD |
| **1010419187** | 1 | LGd-co | 496345668 | 0.431841469 | 0.000198527 | 5 | 0.000459721 | 11.57832298 | 8 mo | HOM |
| **1013978366** | 1 | LGd-co | 496345668 | 0.431841469 | 0 | 0 | 0 | 0 | 2 mo | HOM |
| **1014416155** | 2 | LGd-co | 496345668 | 0.431841469 | 0 | 0 | 0 | 0 | 2 mo | HOM |
| **1013978065** | 2 | LGd-co | 496345668 | 0.431841469 | 0 | 0 | 0 | 0 | 2 mo | HOM |
| **1009934102** | 2 | LGd-co | 496345668 | 0.431841469 | 0 | 0 | 0 | 0 | 4 mo | WILD |
| **1009322937** | 2 | LGd-co | 496345668 | 0.431841469 | 1.48E-05 | 3 | 3.43E-05 | 6.946993785 | 8 mo | WILD |
| **1011080637** | 2 | LGd-co | 496345668 | 0.431841469 | 0.000331735 | 9 | 0.000768186 | 20.84098136 | 8 mo | HOM |
| **1013609713** | 2 | LGd-co | 496345668 | 0.431841469 | 0 | 0 | 0 | 0 | 2 mo | WILD |
| **1010418842** | 1 | LGd-co | 496345668 | 0.431841469 | 0.000715711 | 14 | 0.001657346 | 32.41930433 | 8 mo | HOM |
| **1013288213** | 2 | LGd-co | 496345668 | 0.431841469 | 0 | 0 | 0 | 0 | 4 mo | HOM |
| **1008976060** | 2 | LGd-co | 496345668 | 0.431841469 | 3.44E-05 | 2 | 7.96E-05 | 4.63132919 | 8 mo | WILD |
| **1013446243** | 1 | LGd-co | 496345668 | 0.431841469 | 3.87E-05 | 2 | 8.95E-05 | 4.63132919 | 4 mo | HOM |
| **1013288951** | 2 | LGd-co | 496345668 | 0.431841469 | 1.49E-05 | 2 | 3.45E-05 | 4.63132919 | 4 mo | HOM |
| **1013978665** | 1 | LGd-ip | 496345672 | 0.098747064 | 0 | 0 | 0 | 0 | 2 mo | HOM |
| **1013445609** | 1 | LGd-ip | 496345672 | 0.098747064 | 0 | 0 | 0 | 0 | 4 mo | HOM |
| **1013609015** | 1 | LGd-ip | 496345672 | 0.098747064 | 5.97E-06 | 0 | 6.04E-05 | 0 | 4 mo | HOM |
| **1013608106** | 2 | LGd-ip | 496345672 | 0.098747064 | 0 | 0 | 0 | 0 | 4 mo | HOM |
| **1011937543** | 1 | LGd-ip | 496345672 | 0.098747064 | 7.6E-05 | 1 | 0.000769251 | 10.12688339 | 8 mo | HOM |
| **1010419538** | 2 | LGd-ip | 496345672 | 0.098747064 | 0 | 0 | 0 | 0 | 4 mo | WILD |
| **1010419187** | 1 | LGd-ip | 496345672 | 0.098747064 | 4.59E-05 | 2 | 0.000465022 | 20.25376678 | 8 mo | HOM |
| **1013978366** | 1 | LGd-ip | 496345672 | 0.098747064 | 0 | 0 | 0 | 0 | 2 mo | HOM |
| **1014416155** | 2 | LGd-ip | 496345672 | 0.098747064 | 0 | 0 | 0 | 0 | 2 mo | HOM |
| **1013978065** | 2 | LGd-ip | 496345672 | 0.098747064 | 0 | 0 | 0 | 0 | 2 mo | HOM |
| **1009934102** | 2 | LGd-ip | 496345672 | 0.098747064 | 0 | 0 | 0 | 0 | 4 mo | WILD |
| **1009322937** | 2 | LGd-ip | 496345672 | 0.098747064 | 0 | 0 | 0 | 0 | 8 mo | WILD |
| **1011080637** | 2 | LGd-ip | 496345672 | 0.098747064 | 5.34E-05 | 1 | 0.000540753 | 10.12688339 | 8 mo | HOM |
| **1013609713** | 2 | LGd-ip | 496345672 | 0.098747064 | 0 | 0 | 0 | 0 | 2 mo | WILD |
| **1010418842** | 1 | LGd-ip | 496345672 | 0.098747064 | 0.000120464 | 3 | 0.00121993 | 30.38065016 | 8 mo | HOM |
| **1013288213** | 2 | LGd-ip | 496345672 | 0.098747064 | 0 | 0 | 0 | 0 | 4 mo | HOM |
| **1008976060** | 2 | LGd-ip | 496345672 | 0.098747064 | 2.19E-09 | 0 | 2.22E-08 | 0 | 8 mo | WILD |
| **1013446243** | 1 | LGd-ip | 496345672 | 0.098747064 | 0 | 0 | 0 | 0 | 4 mo | HOM |
| **1013288951** | 2 | LGd-ip | 496345672 | 0.098747064 | 5.33E-06 | 1 | 5.4E-05 | 10.12688339 | 4 mo | HOM |
| **1013978665** | 1 | FRP5 | 526157192 | 0.382815126 | 0 | 0 | 0 | 0 | 2 mo | HOM |
| **1013445609** | 1 | FRP5 | 526157192 | 0.382815126 | 0 | 0 | 0 | 0 | 4 mo | HOM |
| **1013609015** | 1 | FRP5 | 526157192 | 0.382815126 | 1.05E-05 | 1 | 2.74E-05 | 2.612226976 | 4 mo | HOM |
| **1013608106** | 2 | FRP5 | 526157192 | 0.382815126 | 0 | 0 | 0 | 0 | 4 mo | HOM |
| **1011937543** | 1 | FRP5 | 526157192 | 0.382815126 | 0.000293626 | 6 | 0.000767018 | 15.67336186 | 8 mo | HOM |
| **1010419538** | 2 | FRP5 | 526157192 | 0.382815126 | 0 | 0 | 0 | 0 | 4 mo | WILD |
| **1010419187** | 1 | FRP5 | 526157192 | 0.382815126 | 0.000560968 | 18 | 0.001465377 | 47.02008557 | 8 mo | HOM |
| **1013978366** | 1 | FRP5 | 526157192 | 0.382815126 | 0 | 0 | 0 | 0 | 2 mo | HOM |
| **1014416155** | 2 | FRP5 | 526157192 | 0.382815126 | 0 | 0 | 0 | 0 | 2 mo | HOM |
| **1013978065** | 2 | FRP5 | 526157192 | 0.382815126 | 0 | 0 | 0 | 0 | 2 mo | HOM |
| **1009934102** | 2 | FRP5 | 526157192 | 0.382815126 | 0 | 0 | 0 | 0 | 4 mo | WILD |
| **1009322937** | 2 | FRP5 | 526157192 | 0.382815126 | 1.59E-06 | 0 | 4.15E-06 | 0 | 8 mo | WILD |
| **1011080637** | 2 | FRP5 | 526157192 | 0.382815126 | 9.39E-05 | 4 | 0.000245335 | 10.4489079 | 8 mo | HOM |
| **1013609713** | 2 | FRP5 | 526157192 | 0.382815126 | 0 | 0 | 0 | 0 | 2 mo | WILD |
| **1010418842** | 1 | FRP5 | 526157192 | 0.382815126 | 0.000202494 | 12 | 0.000528961 | 31.34672371 | 8 mo | HOM |
| **1013288213** | 2 | FRP5 | 526157192 | 0.382815126 | 2.64E-05 | 2 | 6.9E-05 | 5.224453952 | 4 mo | HOM |
| **1008976060** | 2 | FRP5 | 526157192 | 0.382815126 | 8.15E-06 | 1 | 2.13E-05 | 2.612226976 | 8 mo | WILD |
| **1013446243** | 1 | FRP5 | 526157192 | 0.382815126 | 2.9E-05 | 2 | 7.57E-05 | 5.224453952 | 4 mo | HOM |
| **1013288951** | 2 | FRP5 | 526157192 | 0.382815126 | 3.79E-05 | 2 | 9.9E-05 | 5.224453952 | 4 mo | HOM |
| **1013978665** | 1 | FRP6a | 526157196 | 0.115613912 | 0 | 0 | 0 | 0 | 2 mo | HOM |
| **1013445609** | 1 | FRP6a | 526157196 | 0.115613912 | 0 | 0 | 0 | 0 | 4 mo | HOM |
| **1013609015** | 1 | FRP6a | 526157196 | 0.115613912 | 4.76E-05 | 1 | 0.000412141 | 8.649478071 | 4 mo | HOM |

| | | | | | | | | | | |
|---|---|---|---|---|---|---|---|---|---|---|
| 1013608106 | 2 | FRP6a | 526157196 | 0.115613912 | 0 | 0 | 0 | 0 | 4 mo | HOM |
| 1011937543 | 1 | FRP6a | 526157196 | 0.115613912 | 5.27E-05 | 1 | 0.000455964 | 8.649478071 | 8 mo | HOM |
| 1010419538 | 2 | FRP6a | 526157196 | 0.115613912 | 0 | 0 | 0 | 0 | 4 mo | WILD |
| 1010419187 | 1 | FRP6a | 526157196 | 0.115613912 | 0.000105441 | 4 | 0.000912007 | 34.59791228 | 8 mo | HOM |
| 1013978366 | 1 | FRP6a | 526157196 | 0.115613912 | 0 | 0 | 0 | 0 | 2 mo | HOM |
| 1014416155 | 2 | FRP6a | 526157196 | 0.115613912 | 0 | 0 | 0 | 0 | 2 mo | HOM |
| 1013978065 | 2 | FRP6a | 526157196 | 0.115613912 | 0 | 0 | 0 | 0 | 2 mo | HOM |
| 1009934102 | 2 | FRP6a | 526157196 | 0.115613912 | 0 | 0 | 0 | 0 | 4 mo | WILD |
| 1009322937 | 2 | FRP6a | 526157196 | 0.115613912 | 0 | 0 | 0 | 0 | 8 mo | WILD |
| 1011080637 | 2 | FRP6a | 526157196 | 0.115613912 | 8.57E-05 | 1 | 0.000741334 | 8.649478071 | 8 mo | HOM |
| 1013609713 | 2 | FRP6a | 526157196 | 0.115613912 | 0 | 0 | 0 | 0 | 2 mo | WILD |
| 1010418842 | 1 | FRP6a | 526157196 | 0.115613912 | 8.81E-05 | 4 | 0.000761743 | 34.59791228 | 8 mo | HOM |
| 1013288213 | 2 | FRP6a | 526157196 | 0.115613912 | 0 | 0 | 0 | 0 | 4 mo | HOM |
| 1008976060 | 2 | FRP6a | 526157196 | 0.115613912 | 0 | 0 | 0 | 0 | 8 mo | WILD |
| 1013446243 | 1 | FRP6a | 526157196 | 0.115613912 | 2.16E-05 | 1 | 0.000186483 | 8.649478071 | 4 mo | HOM |
| 1013288951 | 2 | FRP6a | 526157196 | 0.115613912 | 1.26E-07 | 0 | 1.09E-06 | 0 | 4 mo | HOM |
| 1013978665 | 1 | FRP6b | 526322264 | 0.000970472 | 0 | 0 | 0 | 0 | 2 mo | HOM |
| 1013445609 | 1 | FRP6b | 526322264 | 0.000970472 | 0 | 0 | 0 | 0 | 4 mo | HOM |
| 1013609015 | 1 | FRP6b | 526322264 | 0.000970472 | 0 | 0 | 0 | 0 | 4 mo | HOM |
| 1013608106 | 2 | FRP6b | 526322264 | 0.000970472 | 0 | 0 | 0 | 0 | 4 mo | HOM |
| 1011937543 | 1 | FRP6b | 526322264 | 0.000970472 | 0 | 0 | 0 | 0 | 8 mo | HOM |
| 1010419538 | 2 | FRP6b | 526322264 | 0.000970472 | 0 | 0 | 0 | 0 | 4 mo | WILD |
| 1010419187 | 1 | FRP6b | 526322264 | 0.000970472 | 0 | 0 | 0 | 0 | 8 mo | HOM |
| 1013978366 | 1 | FRP6b | 526322264 | 0.000970472 | 0 | 0 | 0 | 0 | 2 mo | HOM |
| 1014416155 | 2 | FRP6b | 526322264 | 0.000970472 | 0 | 0 | 0 | 0 | 2 mo | HOM |
| 1013978065 | 2 | FRP6b | 526322264 | 0.000970472 | 0 | 0 | 0 | 0 | 2 mo | HOM |
| 1009934102 | 2 | FRP6b | 526322264 | 0.000970472 | 0 | 0 | 0 | 0 | 4 mo | WILD |
| 1009322937 | 2 | FRP6b | 526322264 | 0.000970472 | 0 | 0 | 0 | 0 | 8 mo | WILD |
| 1011080637 | 2 | FRP6b | 526322264 | 0.000970472 | 0 | 0 | 0 | 0 | 8 mo | HOM |
| 1013609713 | 2 | FRP6b | 526322264 | 0.000970472 | 0 | 0 | 0 | 0 | 2 mo | WILD |
| 1010418842 | 1 | FRP6b | 526322264 | 0.000970472 | 0 | 0 | 0 | 0 | 8 mo | HOM |
| 1013288213 | 2 | FRP6b | 526322264 | 0.000970472 | 0 | 0 | 0 | 0 | 4 mo | HOM |
| 1008976060 | 2 | FRP6b | 526322264 | 0.000970472 | 0 | 0 | 0 | 0 | 8 mo | WILD |
| 1013446243 | 1 | FRP6b | 526322264 | 0.000970472 | 0 | 0 | 0 | 0 | 4 mo | HOM |
| 1013288951 | 2 | FRP6b | 526322264 | 0.000970472 | 0 | 0 | 0 | 0 | 4 mo | HOM |
| 1013978665 | 1 | ORBm6b | 527696977 | 0.003820074 | 0 | 0 | 0 | 0 | 2 mo | HOM |
| 1013445609 | 1 | ORBm6b | 527696977 | 0.003820074 | 0 | 0 | 0 | 0 | 4 mo | HOM |
| 1013609015 | 1 | ORBm6b | 527696977 | 0.003820074 | 0 | 0 | 0 | 0 | 4 mo | HOM |
| 1013608106 | 2 | ORBm6b | 527696977 | 0.003820074 | 0 | 0 | 0 | 0 | 4 mo | HOM |
| 1011937543 | 1 | ORBm6b | 527696977 | 0.003820074 | 0 | 0 | 0 | 0 | 8 mo | HOM |
| 1010419538 | 2 | ORBm6b | 527696977 | 0.003820074 | 0 | 0 | 0 | 0 | 4 mo | WILD |
| 1010419187 | 1 | ORBm6b | 527696977 | 0.003820074 | 0 | 0 | 0 | 0 | 8 mo | HOM |
| 1013978366 | 1 | ORBm6b | 527696977 | 0.003820074 | 0 | 0 | 0 | 0 | 2 mo | HOM |
| 1014416155 | 2 | ORBm6b | 527696977 | 0.003820074 | 0 | 0 | 0 | 0 | 2 mo | HOM |
| 1013978065 | 2 | ORBm6b | 527696977 | 0.003820074 | 0 | 0 | 0 | 0 | 2 mo | HOM |
| 1009934102 | 2 | ORBm6b | 527696977 | 0.003820074 | 0 | 0 | 0 | 0 | 4 mo | WILD |
| 1009322937 | 2 | ORBm6b | 527696977 | 0.003820074 | 0 | 0 | 0 | 0 | 8 mo | WILD |
| 1011080637 | 2 | ORBm6b | 527696977 | 0.003820074 | 0 | 0 | 0 | 0 | 8 mo | HOM |
| 1013609713 | 2 | ORBm6b | 527696977 | 0.003820074 | 0 | 0 | 0 | 0 | 2 mo | WILD |
| 1010418842 | 1 | ORBm6b | 527696977 | 0.003820074 | 1.68E-11 | 0 | 4.39E-09 | 0 | 8 mo | HOM |
| 1013288213 | 2 | ORBm6b | 527696977 | 0.003820074 | 0 | 0 | 0 | 0 | 4 mo | HOM |
| 1008976060 | 2 | ORBm6b | 527696977 | 0.003820074 | 0 | 0 | 0 | 0 | 8 mo | WILD |
| 1013446243 | 1 | ORBm6b | 527696977 | 0.003820074 | 0 | 0 | 0 | 0 | 4 mo | HOM |
| 1013288951 | 2 | ORBm6b | 527696977 | 0.003820074 | 0 | 0 | 0 | 0 | 4 mo | HOM |
| 1013978665 | 1 | RPF | 549009203 | 0.064047028 | 0 | 0 | 0 | 0 | 2 mo | HOM |
| 1013445609 | 1 | RPF | 549009203 | 0.064047028 | 0 | 0 | 0 | 0 | 4 mo | HOM |
| 1013609015 | 1 | RPF | 549009203 | 0.064047028 | 0 | 0 | 0 | 0 | 4 mo | HOM |
| 1013608106 | 2 | RPF | 549009203 | 0.064047028 | 0 | 0 | 0 | 0 | 4 mo | HOM |
| 1011937543 | 1 | RPF | 549009203 | 0.064047028 | 0 | 0 | 0 | 0 | 8 mo | HOM |

| | | | | | | | | | | |
|---|---|---|---|---|---|---|---|---|---|---|
| **1010419538** | 2 | RPF | 549009203 | 0.064047028 | 0 | 0 | 0 | 0 | 4 mo | WILD |
| **1010419187** | 1 | RPF | 549009203 | 0.064047028 | 0 | 0 | 0 | 0 | 8 mo | HOM |
| **1013978366** | 1 | RPF | 549009203 | 0.064047028 | 0 | 0 | 0 | 0 | 2 mo | HOM |
| **1014416155** | 2 | RPF | 549009203 | 0.064047028 | 0 | 0 | 0 | 0 | 2 mo | HOM |
| **1013978065** | 2 | RPF | 549009203 | 0.064047028 | 0 | 0 | 0 | 0 | 2 mo | HOM |
| **1009934102** | 2 | RPF | 549009203 | 0.064047028 | 0 | 0 | 0 | 0 | 4 mo | WILD |
| **1009322937** | 2 | RPF | 549009203 | 0.064047028 | 0 | 0 | 0 | 0 | 8 mo | WILD |
| **1011080637** | 2 | RPF | 549009203 | 0.064047028 | 0 | 0 | 0 | 0 | 8 mo | HOM |
| **1013609713** | 2 | RPF | 549009203 | 0.064047028 | 0 | 0 | 0 | 0 | 2 mo | WILD |
| **1010418842** | 1 | RPF | 549009203 | 0.064047028 | 4.71E-05 | 1 | 0.00073541 | 15.61352703 | 8 mo | HOM |
| **1013288213** | 2 | RPF | 549009203 | 0.064047028 | 0 | 0 | 0 | 0 | 4 mo | HOM |
| **1008976060** | 2 | RPF | 549009203 | 0.064047028 | 0 | 0 | 0 | 0 | 8 mo | WILD |
| **1013446243** | 1 | RPF | 549009203 | 0.064047028 | 0 | 0 | 0 | 0 | 4 mo | HOM |
| **1013288951** | 2 | RPF | 549009203 | 0.064047028 | 0 | 0 | 0 | 0 | 4 mo | HOM |
| **1013978665** | 1 | MA3 | 549009211 | 0.01685036 | 0 | 0 | 0 | 0 | 2 mo | HOM |
| **1013445609** | 1 | MA3 | 549009211 | 0.01685036 | 0 | 0 | 0 | 0 | 4 mo | HOM |
| **1013609015** | 1 | MA3 | 549009211 | 0.01685036 | 0 | 0 | 0 | 0 | 4 mo | HOM |
| **1013608106** | 2 | MA3 | 549009211 | 0.01685036 | 0 | 0 | 0 | 0 | 4 mo | HOM |
| **1011937543** | 1 | MA3 | 549009211 | 0.01685036 | 0 | 0 | 0 | 0 | 8 mo | HOM |
| **1010419538** | 2 | MA3 | 549009211 | 0.01685036 | 0 | 0 | 0 | 0 | 4 mo | WILD |
| **1010419187** | 1 | MA3 | 549009211 | 0.01685036 | 0 | 0 | 0 | 0 | 8 mo | HOM |
| **1013978366** | 1 | MA3 | 549009211 | 0.01685036 | 0 | 0 | 0 | 0 | 2 mo | HOM |
| **1014416155** | 2 | MA3 | 549009211 | 0.01685036 | 0 | 0 | 0 | 0 | 2 mo | HOM |
| **1013978065** | 2 | MA3 | 549009211 | 0.01685036 | 0 | 0 | 0 | 0 | 2 mo | HOM |
| **1009934102** | 2 | MA3 | 549009211 | 0.01685036 | 0 | 0 | 0 | 0 | 4 mo | WILD |
| **1009322937** | 2 | MA3 | 549009211 | 0.01685036 | 0 | 0 | 0 | 0 | 8 mo | WILD |
| **1011080637** | 2 | MA3 | 549009211 | 0.01685036 | 7.45E-06 | 1 | 0.000442322 | 59.34591273 | 8 mo | HOM |
| **1013609713** | 2 | MA3 | 549009211 | 0.01685036 | 0 | 0 | 0 | 0 | 2 mo | WILD |
| **1010418842** | 1 | MA3 | 549009211 | 0.01685036 | 2.87E-05 | 1 | 0.001701902 | 59.34591273 | 8 mo | HOM |
| **1013288213** | 2 | MA3 | 549009211 | 0.01685036 | 0 | 0 | 0 | 0 | 4 mo | HOM |
| **1008976060** | 2 | MA3 | 549009211 | 0.01685036 | 0 | 0 | 0 | 0 | 8 mo | WILD |
| **1013446243** | 1 | MA3 | 549009211 | 0.01685036 | 0 | 0 | 0 | 0 | 4 mo | HOM |
| **1013288951** | 2 | MA3 | 549009211 | 0.01685036 | 0 | 0 | 0 | 0 | 4 mo | HOM |
| **1013978665** | 1 | P5 | 549009215 | 0.33112875 | 0 | 0 | 0 | 0 | 2 mo | HOM |
| **1013445609** | 1 | P5 | 549009215 | 0.33112875 | 0 | 0 | 0 | 0 | 4 mo | HOM |
| **1013609015** | 1 | P5 | 549009215 | 0.33112875 | 0 | 0 | 0 | 0 | 4 mo | HOM |
| **1013608106** | 2 | P5 | 549009215 | 0.33112875 | 0 | 0 | 0 | 0 | 4 mo | HOM |
| **1011937543** | 1 | P5 | 549009215 | 0.33112875 | 0.00061347 | 3 | 0.001852663 | 9.059920035 | 8 mo | HOM |
| **1010419538** | 2 | P5 | 549009215 | 0.33112875 | 0 | 0 | 0 | 0 | 4 mo | WILD |
| **1010419187** | 1 | P5 | 549009215 | 0.33112875 | 0 | 0 | 0 | 0 | 8 mo | HOM |
| **1013978366** | 1 | P5 | 549009215 | 0.33112875 | 0 | 0 | 0 | 0 | 2 mo | HOM |
| **1014416155** | 2 | P5 | 549009215 | 0.33112875 | 0 | 0 | 0 | 0 | 2 mo | HOM |
| **1013978065** | 2 | P5 | 549009215 | 0.33112875 | 0 | 0 | 0 | 0 | 2 mo | HOM |
| **1009934102** | 2 | P5 | 549009215 | 0.33112875 | 0 | 0 | 0 | 0 | 4 mo | WILD |
| **1009322937** | 2 | P5 | 549009215 | 0.33112875 | 0 | 0 | 0 | 0 | 8 mo | WILD |
| **1011080637** | 2 | P5 | 549009215 | 0.33112875 | 0 | 0 | 0 | 0 | 8 mo | HOM |
| **1013609713** | 2 | P5 | 549009215 | 0.33112875 | 0 | 0 | 0 | 0 | 2 mo | WILD |
| **1010418842** | 1 | P5 | 549009215 | 0.33112875 | 4.54E-05 | 1 | 0.000137019 | 3.019973345 | 8 mo | HOM |
| **1013288213** | 2 | P5 | 549009215 | 0.33112875 | 0 | 0 | 0 | 0 | 4 mo | HOM |
| **1008976060** | 2 | P5 | 549009215 | 0.33112875 | 0 | 0 | 0 | 0 | 8 mo | WILD |
| **1013446243** | 1 | P5 | 549009215 | 0.33112875 | 0 | 0 | 0 | 0 | 4 mo | HOM |
| **1013288951** | 2 | P5 | 549009215 | 0.33112875 | 0 | 0 | 0 | 0 | 4 mo | HOM |
| **1013978665** | 1 | Acs5 | 549009219 | 0.01433249 | 0 | 0 | 0 | 0 | 2 mo | HOM |
| **1013445609** | 1 | Acs5 | 549009219 | 0.01433249 | 0 | 0 | 0 | 0 | 4 mo | HOM |
| **1013609015** | 1 | Acs5 | 549009219 | 0.01433249 | 0 | 0 | 0 | 0 | 4 mo | HOM |
| **1013608106** | 2 | Acs5 | 549009219 | 0.01433249 | 0 | 0 | 0 | 0 | 4 mo | HOM |
| **1011937543** | 1 | Acs5 | 549009219 | 0.01433249 | 0 | 0 | 0 | 0 | 8 mo | HOM |
| **1010419538** | 2 | Acs5 | 549009219 | 0.01433249 | 0 | 0 | 0 | 0 | 4 mo | WILD |
| **1010419187** | 1 | Acs5 | 549009219 | 0.01433249 | 0 | 0 | 0 | 0 | 8 mo | HOM |

| | | | | | | | | | | |
|---|---|---|---|---|---|---|---|---|---|---|
| **1013978366** | 1 | Acs5 | 549009219 | 0.01433249 | 0 | 0 | 0 | 0 | 2 mo | HOM |
| **1014416155** | 2 | Acs5 | 549009219 | 0.01433249 | 0 | 0 | 0 | 0 | 2 mo | HOM |
| **1013978065** | 2 | Acs5 | 549009219 | 0.01433249 | 0 | 0 | 0 | 0 | 2 mo | HOM |
| **1009934102** | 2 | Acs5 | 549009219 | 0.01433249 | 0 | 0 | 0 | 0 | 4 mo | WILD |
| **1009322937** | 2 | Acs5 | 549009219 | 0.01433249 | 0 | 0 | 0 | 0 | 8 mo | WILD |
| **1011080637** | 2 | Acs5 | 549009219 | 0.01433249 | 0 | 0 | 0 | 0 | 8 mo | HOM |
| **1013609713** | 2 | Acs5 | 549009219 | 0.01433249 | 0 | 0 | 0 | 0 | 2 mo | WILD |
| **1010418842** | 1 | Acs5 | 549009219 | 0.01433249 | 0 | 0 | 0 | 0 | 8 mo | HOM |
| **1013288213** | 2 | Acs5 | 549009219 | 0.01433249 | 0 | 0 | 0 | 0 | 4 mo | HOM |
| **1008976060** | 2 | Acs5 | 549009219 | 0.01433249 | 0 | 0 | 0 | 0 | 8 mo | WILD |
| **1013446243** | 1 | Acs5 | 549009219 | 0.01433249 | 0 | 0 | 0 | 0 | 4 mo | HOM |
| **1013288951** | 2 | Acs5 | 549009219 | 0.01433249 | 0 | 0 | 0 | 0 | 4 mo | HOM |
| **1013978665** | 1 | PC5 | 549009223 | 0.072845149 | 0 | 0 | 0 | 0 | 2 mo | HOM |
| **1013445609** | 1 | PC5 | 549009223 | 0.072845149 | 0 | 0 | 0 | 0 | 4 mo | HOM |
| **1013609015** | 1 | PC5 | 549009223 | 0.072845149 | 0 | 0 | 0 | 0 | 4 mo | HOM |
| **1013608106** | 2 | PC5 | 549009223 | 0.072845149 | 0 | 0 | 0 | 0 | 4 mo | HOM |
| **1011937543** | 1 | PC5 | 549009223 | 0.072845149 | 0 | 0 | 0 | 0 | 8 mo | HOM |
| **1010419538** | 2 | PC5 | 549009223 | 0.072845149 | 0 | 0 | 0 | 0 | 4 mo | WILD |
| **1010419187** | 1 | PC5 | 549009223 | 0.072845149 | 0 | 0 | 0 | 0 | 8 mo | HOM |
| **1013978366** | 1 | PC5 | 549009223 | 0.072845149 | 0 | 0 | 0 | 0 | 2 mo | HOM |
| **1014416155** | 2 | PC5 | 549009223 | 0.072845149 | 0 | 0 | 0 | 0 | 2 mo | HOM |
| **1013978065** | 2 | PC5 | 549009223 | 0.072845149 | 0 | 0 | 0 | 0 | 2 mo | HOM |
| **1009934102** | 2 | PC5 | 549009223 | 0.072845149 | 0 | 0 | 0 | 0 | 4 mo | WILD |
| **1009322937** | 2 | PC5 | 549009223 | 0.072845149 | 0 | 0 | 0 | 0 | 8 mo | WILD |
| **1011080637** | 2 | PC5 | 549009223 | 0.072845149 | 0 | 0 | 0 | 0 | 8 mo | HOM |
| **1013609713** | 2 | PC5 | 549009223 | 0.072845149 | 0 | 0 | 0 | 0 | 2 mo | WILD |
| **1010418842** | 1 | PC5 | 549009223 | 0.072845149 | 8.83E-05 | 5 | 0.001212211 | 68.63875012 | 8 mo | HOM |
| **1013288213** | 2 | PC5 | 549009223 | 0.072845149 | 0 | 0 | 0 | 0 | 4 mo | HOM |
| **1008976060** | 2 | PC5 | 549009223 | 0.072845149 | 2.08E-07 | 0 | 2.86E-06 | 0 | 8 mo | WILD |
| **1013446243** | 1 | PC5 | 549009223 | 0.072845149 | 0 | 0 | 0 | 0 | 4 mo | HOM |
| **1013288951** | 2 | PC5 | 549009223 | 0.072845149 | 0 | 0 | 0 | 0 | 4 mo | HOM |
| **1013978665** | 1 | I5 | 549009227 | 0.045849133 | 0 | 0 | 0 | 0 | 2 mo | HOM |
| **1013445609** | 1 | I5 | 549009227 | 0.045849133 | 0 | 0 | 0 | 0 | 4 mo | HOM |
| **1013609015** | 1 | I5 | 549009227 | 0.045849133 | 0 | 0 | 0 | 0 | 4 mo | HOM |
| **1013608106** | 2 | I5 | 549009227 | 0.045849133 | 0 | 0 | 0 | 0 | 4 mo | HOM |
| **1011937543** | 1 | I5 | 549009227 | 0.045849133 | 0.000737995 | 0 | 0.016096169 | 0 | 8 mo | HOM |
| **1010419538** | 2 | I5 | 549009227 | 0.045849133 | 0 | 0 | 0 | 0 | 4 mo | WILD |
| **1010419187** | 1 | I5 | 549009227 | 0.045849133 | 0 | 0 | 0 | 0 | 8 mo | HOM |
| **1013978366** | 1 | I5 | 549009227 | 0.045849133 | 0 | 0 | 0 | 0 | 2 mo | HOM |
| **1014416155** | 2 | I5 | 549009227 | 0.045849133 | 0 | 0 | 0 | 0 | 2 mo | HOM |
| **1013978065** | 2 | I5 | 549009227 | 0.045849133 | 0 | 0 | 0 | 0 | 2 mo | HOM |
| **1009934102** | 2 | I5 | 549009227 | 0.045849133 | 0 | 0 | 0 | 0 | 4 mo | WILD |
| **1009322937** | 2 | I5 | 549009227 | 0.045849133 | 0 | 0 | 0 | 0 | 8 mo | WILD |
| **1011080637** | 2 | I5 | 549009227 | 0.045849133 | 0 | 0 | 0 | 0 | 8 mo | HOM |
| **1013609713** | 2 | I5 | 549009227 | 0.045849133 | 0 | 0 | 0 | 0 | 2 mo | WILD |
| **1010418842** | 1 | I5 | 549009227 | 0.045849133 | 4.83E-07 | 0 | 1.05E-05 | 0 | 8 mo | HOM |
| **1013288213** | 2 | I5 | 549009227 | 0.045849133 | 0 | 0 | 0 | 0 | 4 mo | HOM |
| **1008976060** | 2 | I5 | 549009227 | 0.045849133 | 0 | 0 | 0 | 0 | 8 mo | WILD |
| **1013446243** | 1 | I5 | 549009227 | 0.045849133 | 0 | 0 | 0 | 0 | 4 mo | HOM |
| **1013288951** | 2 | I5 | 549009227 | 0.045849133 | 0 | 0 | 0 | 0 | 4 mo | HOM |
| **1013978665** | 1 | Eth | 560581551 | 0.242836395 | 0 | 0 | 0 | 0 | 2 mo | HOM |
| **1013445609** | 1 | Eth | 560581551 | 0.242836395 | 0 | 0 | 0 | 0 | 4 mo | HOM |
| **1013609015** | 1 | Eth | 560581551 | 0.242836395 | 0 | 0 | 0 | 0 | 4 mo | HOM |
| **1013608106** | 2 | Eth | 560581551 | 0.242836395 | 0 | 0 | 0 | 0 | 4 mo | HOM |
| **1011937543** | 1 | Eth | 560581551 | 0.242836395 | 0.000255055 | 5 | 0.001050316 | 20.58999433 | 8 mo | HOM |
| **1010419538** | 2 | Eth | 560581551 | 0.242836395 | 0 | 0 | 0 | 0 | 4 mo | WILD |
| **1010419187** | 1 | Eth | 560581551 | 0.242836395 | 7.85E-05 | 3 | 0.000323059 | 12.3539966 | 8 mo | HOM |
| **1013978366** | 1 | Eth | 560581551 | 0.242836395 | 0 | 0 | 0 | 0 | 2 mo | HOM |
| **1014416155** | 2 | Eth | 560581551 | 0.242836395 | 0 | 0 | 0 | 0 | 2 mo | HOM |

| | | | | | | | | | | |
|---|---|---|---|---|---|---|---|---|---|---|
| **1013978065** | 2 | Eth | 560581551 | 0.242836395 | 0 | 0 | 0 | 0 | 2 mo | HOM |
| **1009934102** | 2 | Eth | 560581551 | 0.242836395 | 0 | 0 | 0 | 0 | 4 mo | WILD |
| **1009322937** | 2 | Eth | 560581551 | 0.242836395 | 0 | 0 | 0 | 0 | 8 mo | WILD |
| **1011080637** | 2 | Eth | 560581551 | 0.242836395 | 4.4E-05 | 3 | 0.000181237 | 12.3539966 | 8 mo | HOM |
| **1013609713** | 2 | Eth | 560581551 | 0.242836395 | 0 | 0 | 0 | 0 | 2 mo | WILD |
| **1010418842** | 1 | Eth | 560581551 | 0.242836395 | 0.000205278 | 4 | 0.000845333 | 16.47199546 | 8 mo | HOM |
| **1013288213** | 2 | Eth | 560581551 | 0.242836395 | 2.51E-06 | 1 | 1.04E-05 | 4.117998866 | 4 mo | HOM |
| **1008976060** | 2 | Eth | 560581551 | 0.242836395 | 4.7E-09 | 0 | 1.93E-08 | 0 | 8 mo | WILD |
| **1013446243** | 1 | Eth | 560581551 | 0.242836395 | 0 | 0 | 0 | 0 | 4 mo | HOM |
| **1013288951** | 2 | Eth | 560581551 | 0.242836395 | 0 | 0 | 0 | 0 | 4 mo | HOM |
| **1013978665** | 1 | Xi | 560581559 | 0.090315702 | 0 | 0 | 0 | 0 | 2 mo | HOM |
| **1013445609** | 1 | Xi | 560581559 | 0.090315702 | 0 | 0 | 0 | 0 | 4 mo | HOM |
| **1013609015** | 1 | Xi | 560581559 | 0.090315702 | 0 | 0 | 0 | 0 | 4 mo | HOM |
| **1013608106** | 2 | Xi | 560581559 | 0.090315702 | 0 | 0 | 0 | 0 | 4 mo | HOM |
| **1011937543** | 1 | Xi | 560581559 | 0.090315702 | 5.8E-05 | 2 | 0.000642665 | 22.14454349 | 8 mo | HOM |
| **1010419538** | 2 | Xi | 560581559 | 0.090315702 | 0 | 0 | 0 | 0 | 4 mo | WILD |
| **1010419187** | 1 | Xi | 560581559 | 0.090315702 | 3.94E-06 | 0 | 4.37E-05 | 0 | 8 mo | HOM |
| **1013978366** | 1 | Xi | 560581559 | 0.090315702 | 0 | 0 | 0 | 0 | 2 mo | HOM |
| **1014416155** | 2 | Xi | 560581559 | 0.090315702 | 0 | 0 | 0 | 0 | 2 mo | HOM |
| **1013978065** | 2 | Xi | 560581559 | 0.090315702 | 0 | 0 | 0 | 0 | 2 mo | HOM |
| **1009934102** | 2 | Xi | 560581559 | 0.090315702 | 0 | 0 | 0 | 0 | 4 mo | WILD |
| **1009322937** | 2 | Xi | 560581559 | 0.090315702 | 0 | 0 | 0 | 0 | 8 mo | WILD |
| **1011080637** | 2 | Xi | 560581559 | 0.090315702 | 3.28E-05 | 1 | 0.000362983 | 11.07227175 | 8 mo | HOM |
| **1013609713** | 2 | Xi | 560581559 | 0.090315702 | 0 | 0 | 0 | 0 | 2 mo | WILD |
| **1010418842** | 1 | Xi | 560581559 | 0.090315702 | 4.91E-05 | 4 | 0.000543882 | 44.28908699 | 8 mo | HOM |
| **1013288213** | 2 | Xi | 560581559 | 0.090315702 | 0 | 0 | 0 | 0 | 4 mo | HOM |
| **1008976060** | 2 | Xi | 560581559 | 0.090315702 | 0 | 0 | 0 | 0 | 8 mo | WILD |
| **1013446243** | 1 | Xi | 560581559 | 0.090315702 | 0 | 0 | 0 | 0 | 4 mo | HOM |
| **1013288951** | 2 | Xi | 560581559 | 0.090315702 | 0 | 0 | 0 | 0 | 4 mo | HOM |
| **1013978665** | 1 | PIL | 560581563 | 0.180952845 | 0 | 0 | 0 | 0 | 2 mo | HOM |
| **1013445609** | 1 | PIL | 560581563 | 0.180952845 | 0 | 0 | 0 | 0 | 4 mo | HOM |
| **1013609015** | 1 | PIL | 560581563 | 0.180952845 | 0 | 0 | 0 | 0 | 4 mo | HOM |
| **1013608106** | 2 | PIL | 560581563 | 0.180952845 | 0 | 0 | 0 | 0 | 4 mo | HOM |
| **1011937543** | 1 | PIL | 560581563 | 0.180952845 | 0.000297319 | 7 | 0.001643075 | 38.68411136 | 8 mo | HOM |
| **1010419538** | 2 | PIL | 560581563 | 0.180952845 | 0 | 0 | 0 | 0 | 4 mo | WILD |
| **1010419187** | 1 | PIL | 560581563 | 0.180952845 | 1.65E-06 | 0 | 9.13E-06 | 0 | 8 mo | HOM |
| **1013978366** | 1 | PIL | 560581563 | 0.180952845 | 0 | 0 | 0 | 0 | 2 mo | HOM |
| **1014416155** | 2 | PIL | 560581563 | 0.180952845 | 0 | 0 | 0 | 0 | 2 mo | HOM |
| **1013978065** | 2 | PIL | 560581563 | 0.180952845 | 0 | 0 | 0 | 0 | 2 mo | HOM |
| **1009934102** | 2 | PIL | 560581563 | 0.180952845 | 0 | 0 | 0 | 0 | 4 mo | WILD |
| **1009322937** | 2 | PIL | 560581563 | 0.180952845 | 0 | 0 | 0 | 0 | 8 mo | WILD |
| **1011080637** | 2 | PIL | 560581563 | 0.180952845 | 1.14E-05 | 1 | 6.29E-05 | 5.526301624 | 8 mo | HOM |
| **1013609713** | 2 | PIL | 560581563 | 0.180952845 | 0 | 0 | 0 | 0 | 2 mo | WILD |
| **1010418842** | 1 | PIL | 560581563 | 0.180952845 | 1.26E-05 | 1 | 6.97E-05 | 5.526301624 | 8 mo | HOM |
| **1013288213** | 2 | PIL | 560581563 | 0.180952845 | 0 | 0 | 0 | 0 | 4 mo | HOM |
| **1008976060** | 2 | PIL | 560581563 | 0.180952845 | 0 | 0 | 0 | 0 | 8 mo | WILD |
| **1013446243** | 1 | PIL | 560581563 | 0.180952845 | 0 | 0 | 0 | 0 | 4 mo | HOM |
| **1013288951** | 2 | PIL | 560581563 | 0.180952845 | 0 | 0 | 0 | 0 | 4 mo | HOM |
| **1013978665** | 1 | PoT | 563807435 | 0.288052968 | 0 | 0 | 0 | 0 | 2 mo | HOM |
| **1013445609** | 1 | PoT | 563807435 | 0.288052968 | 0 | 0 | 0 | 0 | 4 mo | HOM |
| **1013609015** | 1 | PoT | 563807435 | 0.288052968 | 0 | 0 | 0 | 0 | 4 mo | HOM |
| **1013608106** | 2 | PoT | 563807435 | 0.288052968 | 0 | 0 | 0 | 0 | 4 mo | HOM |
| **1011937543** | 1 | PoT | 563807435 | 0.288052968 | 0.000446806 | 10 | 0.001551126 | 34.7158374 | 8 mo | HOM |
| **1010419538** | 2 | PoT | 563807435 | 0.288052968 | 0 | 0 | 0 | 0 | 4 mo | WILD |
| **1010419187** | 1 | PoT | 563807435 | 0.288052968 | 2.75E-06 | 0 | 9.53E-06 | 0 | 8 mo | HOM |
| **1013978366** | 1 | PoT | 563807435 | 0.288052968 | 0 | 0 | 0 | 0 | 2 mo | HOM |
| **1014416155** | 2 | PoT | 563807435 | 0.288052968 | 0 | 0 | 0 | 0 | 2 mo | HOM |
| **1013978065** | 2 | PoT | 563807435 | 0.288052968 | 0 | 0 | 0 | 0 | 2 mo | HOM |
| **1009934102** | 2 | PoT | 563807435 | 0.288052968 | 0 | 0 | 0 | 0 | 4 mo | WILD |

| | | | | | | | | | | |
|---|---|---|---|---|---|---|---|---|---|---|
| **1009322937** | 2 | PoT | 563807435 | 0.288052968 | 0 | 0 | 0 | 0 | 8 mo | WILD |
| **1011080637** | 2 | PoT | 563807435 | 0.288052968 | 2.58E-05 | 1 | 8.97E-05 | 3.47158374 | 8 mo | HOM |
| **1013609713** | 2 | PoT | 563807435 | 0.288052968 | 0 | 0 | 0 | 0 | 2 mo | WILD |
| **1010418842** | 1 | PoT | 563807435 | 0.288052968 | 0.000224287 | 3 | 0.000778632 | 10.41475122 | 8 mo | HOM |
| **1013288213** | 2 | PoT | 563807435 | 0.288052968 | 0 | 0 | 0 | 0 | 4 mo | HOM |
| **1008976060** | 2 | PoT | 563807435 | 0.288052968 | 1.85E-05 | 1 | 6.43E-05 | 3.47158374 | 8 mo | WILD |
| **1013446243** | 1 | PoT | 563807435 | 0.288052968 | 0 | 0 | 0 | 0 | 4 mo | HOM |
| **1013288951** | 2 | PoT | 563807435 | 0.288052968 | 0 | 0 | 0 | 0 | 4 mo | HOM |
| **1013978665** | 1 | IntG | 563807439 | 0.023180063 | 0 | 0 | 0 | 0 | 2 mo | HOM |
| **1013445609** | 1 | IntG | 563807439 | 0.023180063 | 0 | 0 | 0 | 0 | 4 mo | HOM |
| **1013609015** | 1 | IntG | 563807439 | 0.023180063 | 0 | 0 | 0 | 0 | 4 mo | HOM |
| **1013608106** | 2 | IntG | 563807439 | 0.023180063 | 0 | 0 | 0 | 0 | 4 mo | HOM |
| **1011937543** | 1 | IntG | 563807439 | 0.023180063 | 1.55E-05 | 1 | 0.000667058 | 43.14052216 | 8 mo | HOM |
| **1010419538** | 2 | IntG | 563807439 | 0.023180063 | 0 | 0 | 0 | 0 | 4 mo | WILD |
| **1010419187** | 1 | IntG | 563807439 | 0.023180063 | 0 | 0 | 0 | 0 | 8 mo | HOM |
| **1013978366** | 1 | IntG | 563807439 | 0.023180063 | 0 | 0 | 0 | 0 | 2 mo | HOM |
| **1014416155** | 2 | IntG | 563807439 | 0.023180063 | 0 | 0 | 0 | 0 | 2 mo | HOM |
| **1013978065** | 2 | IntG | 563807439 | 0.023180063 | 0 | 0 | 0 | 0 | 2 mo | HOM |
| **1009934102** | 2 | IntG | 563807439 | 0.023180063 | 0 | 0 | 0 | 0 | 4 mo | WILD |
| **1009322937** | 2 | IntG | 563807439 | 0.023180063 | 0 | 0 | 0 | 0 | 8 mo | WILD |
| **1011080637** | 2 | IntG | 563807439 | 0.023180063 | 0 | 0 | 0 | 0 | 8 mo | HOM |
| **1013609713** | 2 | IntG | 563807439 | 0.023180063 | 0 | 0 | 0 | 0 | 2 mo | WILD |
| **1010418842** | 1 | IntG | 563807439 | 0.023180063 | 2.02E-09 | 0 | 8.72E-08 | 0 | 8 mo | HOM |
| **1013288213** | 2 | IntG | 563807439 | 0.023180063 | 0 | 0 | 0 | 0 | 4 mo | HOM |
| **1008976060** | 2 | IntG | 563807439 | 0.023180063 | 0 | 0 | 0 | 0 | 8 mo | WILD |
| **1013446243** | 1 | IntG | 563807439 | 0.023180063 | 0 | 0 | 0 | 0 | 4 mo | HOM |
| **1013288951** | 2 | IntG | 563807439 | 0.023180063 | 0 | 0 | 0 | 0 | 4 mo | HOM |
| **1013978665** | 1 | VMPO | 576073699 | 0.051643119 | 0 | 0 | 0 | 0 | 2 mo | HOM |
| **1013445609** | 1 | VMPO | 576073699 | 0.051643119 | 0 | 0 | 0 | 0 | 4 mo | HOM |
| **1013609015** | 1 | VMPO | 576073699 | 0.051643119 | 0 | 0 | 0 | 0 | 4 mo | HOM |
| **1013608106** | 2 | VMPO | 576073699 | 0.051643119 | 0 | 0 | 0 | 0 | 4 mo | HOM |
| **1011937543** | 1 | VMPO | 576073699 | 0.051643119 | 0.000293325 | 0 | 0.00567984 | 0 | 8 mo | HOM |
| **1010419538** | 2 | VMPO | 576073699 | 0.051643119 | 0 | 0 | 0 | 0 | 4 mo | WILD |
| **1010419187** | 1 | VMPO | 576073699 | 0.051643119 | 0 | 0 | 0 | 0 | 8 mo | HOM |
| **1013978366** | 1 | VMPO | 576073699 | 0.051643119 | 0 | 0 | 0 | 0 | 2 mo | HOM |
| **1014416155** | 2 | VMPO | 576073699 | 0.051643119 | 0 | 0 | 0 | 0 | 2 mo | HOM |
| **1013978065** | 2 | VMPO | 576073699 | 0.051643119 | 0 | 0 | 0 | 0 | 2 mo | HOM |
| **1009934102** | 2 | VMPO | 576073699 | 0.051643119 | 0 | 0 | 0 | 0 | 4 mo | WILD |
| **1009322937** | 2 | VMPO | 576073699 | 0.051643119 | 0 | 0 | 0 | 0 | 8 mo | WILD |
| **1011080637** | 2 | VMPO | 576073699 | 0.051643119 | 1.18E-06 | 0 | 2.28E-05 | 0 | 8 mo | HOM |
| **1013609713** | 2 | VMPO | 576073699 | 0.051643119 | 0 | 0 | 0 | 0 | 2 mo | WILD |
| **1010418842** | 1 | VMPO | 576073699 | 0.051643119 | 1.79E-05 | 1 | 0.000346902 | 19.36366399 | 8 mo | HOM |
| **1013288213** | 2 | VMPO | 576073699 | 0.051643119 | 0 | 0 | 0 | 0 | 4 mo | HOM |
| **1008976060** | 2 | VMPO | 576073699 | 0.051643119 | 0 | 0 | 0 | 0 | 8 mo | WILD |
| **1013446243** | 1 | VMPO | 576073699 | 0.051643119 | 0 | 0 | 0 | 0 | 4 mo | HOM |
| **1013288951** | 2 | VMPO | 576073699 | 0.051643119 | 0 | 0 | 0 | 0 | 4 mo | HOM |
| **1013978665** | 1 | PeF | 576073704 | 0.22072983 | 0 | 0 | 0 | 0 | 2 mo | HOM |
| **1013445609** | 1 | PeF | 576073704 | 0.22072983 | 0 | 0 | 0 | 0 | 4 mo | HOM |
| **1013609015** | 1 | PeF | 576073704 | 0.22072983 | 0 | 0 | 0 | 0 | 4 mo | HOM |
| **1013608106** | 2 | PeF | 576073704 | 0.22072983 | 0 | 0 | 0 | 0 | 4 mo | HOM |
| **1011937543** | 1 | PeF | 576073704 | 0.22072983 | 0.000197537 | 3 | 0.000894928 | 13.59127584 | 8 mo | HOM |
| **1010419538** | 2 | PeF | 576073704 | 0.22072983 | 0 | 0 | 0 | 0 | 4 mo | WILD |
| **1010419187** | 1 | PeF | 576073704 | 0.22072983 | 4.99E-05 | 2 | 0.000225891 | 9.060850559 | 8 mo | HOM |
| **1013978366** | 1 | PeF | 576073704 | 0.22072983 | 0 | 0 | 0 | 0 | 2 mo | HOM |
| **1014416155** | 2 | PeF | 576073704 | 0.22072983 | 0 | 0 | 0 | 0 | 2 mo | HOM |
| **1013978065** | 2 | PeF | 576073704 | 0.22072983 | 0 | 0 | 0 | 0 | 2 mo | HOM |
| **1009934102** | 2 | PeF | 576073704 | 0.22072983 | 0 | 0 | 0 | 0 | 4 mo | WILD |
| **1009322937** | 2 | PeF | 576073704 | 0.22072983 | 0 | 0 | 0 | 0 | 8 mo | WILD |
| **1011080637** | 2 | PeF | 576073704 | 0.22072983 | 3.82E-05 | 2 | 0.000173161 | 9.060850559 | 8 mo | HOM |

| | | | | | | | | | | |
|---|---|---|---|---|---|---|---|---|---|---|
| **1013609713** | 2 | PeF | 576073704 | 0.22072983 | 0 | 0 | 0 | 0 | 2 mo | WILD |
| **1010418842** | 1 | PeF | 576073704 | 0.22072983 | 0.000188811 | 3 | 0.000855395 | 13.59127584 | 8 mo | HOM |
| **1013288213** | 2 | PeF | 576073704 | 0.22072983 | 0 | 0 | 0 | 0 | 4 mo | HOM |
| **1008976060** | 2 | PeF | 576073704 | 0.22072983 | 2.05E-06 | 0 | 9.31E-06 | 0 | 8 mo | WILD |
| **1013446243** | 1 | PeF | 576073704 | 0.22072983 | 0 | 0 | 0 | 0 | 4 mo | HOM |
| **1013288951** | 2 | PeF | 576073704 | 0.22072983 | 0 | 0 | 0 | 0 | 4 mo | HOM |
| **1013978665** | 1 | HATA | 589508447 | 0.434068813 | 0 | 0 | 0 | 0 | 2 mo | HOM |
| **1013445609** | 1 | HATA | 589508447 | 0.434068813 | 3.86E-05 | 4 | 8.89E-05 | 9.215128758 | 4 mo | HOM |
| **1013609015** | 1 | HATA | 589508447 | 0.434068813 | 7.96E-06 | 0 | 1.83E-05 | 0 | 4 mo | HOM |
| **1013608106** | 2 | HATA | 589508447 | 0.434068813 | 0 | 0 | 0 | 0 | 4 mo | HOM |
| **1011937543** | 1 | HATA | 589508447 | 0.434068813 | 0.000764165 | 9 | 0.00176047 | 20.73403971 | 8 mo | HOM |
| **1010419538** | 2 | HATA | 589508447 | 0.434068813 | 8.44E-06 | 2 | 1.95E-05 | 4.607564379 | 4 mo | WILD |
| **1010419187** | 1 | HATA | 589508447 | 0.434068813 | 0.000547644 | 24 | 0.001261652 | 55.29077255 | 8 mo | HOM |
| **1013978366** | 1 | HATA | 589508447 | 0.434068813 | 0 | 0 | 0 | 0 | 2 mo | HOM |
| **1014416155** | 2 | HATA | 589508447 | 0.434068813 | 0 | 0 | 0 | 0 | 2 mo | HOM |
| **1013978065** | 2 | HATA | 589508447 | 0.434068813 | 0 | 0 | 0 | 0 | 2 mo | HOM |
| **1009934102** | 2 | HATA | 589508447 | 0.434068813 | 0 | 0 | 0 | 0 | 4 mo | WILD |
| **1009322937** | 2 | HATA | 589508447 | 0.434068813 | 6.48E-06 | 1 | 1.49E-05 | 2.30378219 | 8 mo | WILD |
| **1011080637** | 2 | HATA | 589508447 | 0.434068813 | 0.00017957 | 8 | 0.00041369 | 18.43025752 | 8 mo | HOM |
| **1013609713** | 2 | HATA | 589508447 | 0.434068813 | 0 | 0 | 0 | 0 | 2 mo | WILD |
| **1010418842** | 1 | HATA | 589508447 | 0.434068813 | 0.000544116 | 22 | 0.001253525 | 50.68320817 | 8 mo | HOM |
| **1013288213** | 2 | HATA | 589508447 | 0.434068813 | 6.6E-05 | 4 | 0.000152164 | 9.215128758 | 4 mo | HOM |
| **1008976060** | 2 | HATA | 589508447 | 0.434068813 | 3.61E-06 | 1 | 8.31E-06 | 2.30378219 | 8 mo | WILD |
| **1013446243** | 1 | HATA | 589508447 | 0.434068813 | 2.23E-05 | 1 | 5.15E-05 | 2.30378219 | 4 mo | HOM |
| **1013288951** | 2 | HATA | 589508447 | 0.434068813 | 4.85E-05 | 4 | 0.000111791 | 9.215128758 | 4 mo | HOM |
| **1013978665** | 1 | Pa5 | 589508451 | 0.10573611 | 0 | 0 | 0 | 0 | 2 mo | HOM |
| **1013445609** | 1 | Pa5 | 589508451 | 0.10573611 | 0 | 0 | 0 | 0 | 4 mo | HOM |
| **1013609015** | 1 | Pa5 | 589508451 | 0.10573611 | 0 | 0 | 0 | 0 | 4 mo | HOM |
| **1013608106** | 2 | Pa5 | 589508451 | 0.10573611 | 0 | 0 | 0 | 0 | 4 mo | HOM |
| **1011937543** | 1 | Pa5 | 589508451 | 0.10573611 | 0 | 0 | 0 | 0 | 8 mo | HOM |
| **1010419538** | 2 | Pa5 | 589508451 | 0.10573611 | 0 | 0 | 0 | 0 | 4 mo | WILD |
| **1010419187** | 1 | Pa5 | 589508451 | 0.10573611 | 0 | 0 | 0 | 0 | 8 mo | HOM |
| **1013978366** | 1 | Pa5 | 589508451 | 0.10573611 | 0 | 0 | 0 | 0 | 2 mo | HOM |
| **1014416155** | 2 | Pa5 | 589508451 | 0.10573611 | 0 | 0 | 0 | 0 | 2 mo | HOM |
| **1013978065** | 2 | Pa5 | 589508451 | 0.10573611 | 0 | 0 | 0 | 0 | 2 mo | HOM |
| **1009934102** | 2 | Pa5 | 589508451 | 0.10573611 | 0 | 0 | 0 | 0 | 4 mo | WILD |
| **1009322937** | 2 | Pa5 | 589508451 | 0.10573611 | 0 | 0 | 0 | 0 | 8 mo | WILD |
| **1011080637** | 2 | Pa5 | 589508451 | 0.10573611 | 0 | 0 | 0 | 0 | 8 mo | HOM |
| **1013609713** | 2 | Pa5 | 589508451 | 0.10573611 | 0 | 0 | 0 | 0 | 2 mo | WILD |
| **1010418842** | 1 | Pa5 | 589508451 | 0.10573611 | 0 | 0 | 0 | 0 | 8 mo | HOM |
| **1013288213** | 2 | Pa5 | 589508451 | 0.10573611 | 0 | 0 | 0 | 0 | 4 mo | HOM |
| **1008976060** | 2 | Pa5 | 589508451 | 0.10573611 | 1.38E-06 | 0 | 1.3E-05 | 0 | 8 mo | WILD |
| **1013446243** | 1 | Pa5 | 589508451 | 0.10573611 | 0 | 0 | 0 | 0 | 4 mo | HOM |
| **1013288951** | 2 | Pa5 | 589508451 | 0.10573611 | 0 | 0 | 0 | 0 | 4 mo | HOM |
| **1013978665** | 1 | VeCB | 589508455 | 0.087719535 | 0 | 0 | 0 | 0 | 2 mo | HOM |
| **1013445609** | 1 | VeCB | 589508455 | 0.087719535 | 1.21E-07 | 0 | 1.38E-06 | 0 | 4 mo | HOM |
| **1013609015** | 1 | VeCB | 589508455 | 0.087719535 | 0 | 0 | 0 | 0 | 4 mo | HOM |
| **1013608106** | 2 | VeCB | 589508455 | 0.087719535 | 0 | 0 | 0 | 0 | 4 mo | HOM |
| **1011937543** | 1 | VeCB | 589508455 | 0.087719535 | 0 | 0 | 0 | 0 | 8 mo | HOM |
| **1010419538** | 2 | VeCB | 589508455 | 0.087719535 | 0 | 0 | 0 | 0 | 4 mo | WILD |
| **1010419187** | 1 | VeCB | 589508455 | 0.087719535 | 1.57E-05 | 0 | 0.000178581 | 0 | 8 mo | HOM |
| **1013978366** | 1 | VeCB | 589508455 | 0.087719535 | 1.99E-06 | 0 | 2.27E-05 | 0 | 2 mo | HOM |
| **1014416155** | 2 | VeCB | 589508455 | 0.087719535 | 0 | 0 | 0 | 0 | 2 mo | HOM |
| **1013978065** | 2 | VeCB | 589508455 | 0.087719535 | 0 | 0 | 0 | 0 | 2 mo | HOM |
| **1009934102** | 2 | VeCB | 589508455 | 0.087719535 | 0 | 0 | 0 | 0 | 4 mo | WILD |
| **1009322937** | 2 | VeCB | 589508455 | 0.087719535 | 0 | 0 | 0 | 0 | 8 mo | WILD |
| **1011080637** | 2 | VeCB | 589508455 | 0.087719535 | 0.000113822 | 3 | 0.001297567 | 34.19990754 | 8 mo | HOM |
| **1013609713** | 2 | VeCB | 589508455 | 0.087719535 | 0 | 0 | 0 | 0 | 2 mo | WILD |
| **1010418842** | 1 | VeCB | 589508455 | 0.087719535 | 4.05E-05 | 0 | 0.000461521 | 0 | 8 mo | HOM |

| | | | | | | | | | | |
|---|---|---|---|---|---|---|---|---|---|---|
| 1013288213 | 2 | VeCB | 589508455 | 0.087719535 | 1.06E-05 | 1 | 0.000120324 | 11.39996918 | 4 mo | HOM |
| 1008976060 | 2 | VeCB | 589508455 | 0.087719535 | 6.13E-05 | 0 | 0.000699069 | 0 | 8 mo | WILD |
| 1013446243 | 1 | VeCB | 589508455 | 0.087719535 | 0 | 0 | 0 | 0 | 4 mo | HOM |
| 1013288951 | 2 | VeCB | 589508455 | 0.087719535 | 3.3E-05 | 0 | 0.000376282 | 0 | 4 mo | HOM |
| 1013978665 | 1 | SCO | 599626923 | 0.014915598 | 0 | 0 | 0 | 0 | 2 mo | HOM |
| 1013445609 | 1 | SCO | 599626923 | 0.014915598 | 0 | 0 | 0 | 0 | 4 mo | HOM |
| 1013609015 | 1 | SCO | 599626923 | 0.014915598 | 0 | 0 | 0 | 0 | 4 mo | HOM |
| 1013608106 | 2 | SCO | 599626923 | 0.014915598 | 0 | 0 | 0 | 0 | 4 mo | HOM |
| 1011937543 | 1 | SCO | 599626923 | 0.014915598 | 2.05E-05 | 0 | 0.00137297 | 0 | 8 mo | HOM |
| 1010419538 | 2 | SCO | 599626923 | 0.014915598 | 0 | 0 | 0 | 0 | 4 mo | WILD |
| 1010419187 | 1 | SCO | 599626923 | 0.014915598 | 0 | 0 | 0 | 0 | 8 mo | HOM |
| 1013978366 | 1 | SCO | 599626923 | 0.014915598 | 0 | 0 | 0 | 0 | 2 mo | HOM |
| 1014416155 | 2 | SCO | 599626923 | 0.014915598 | 0 | 0 | 0 | 0 | 2 mo | HOM |
| 1013978065 | 2 | SCO | 599626923 | 0.014915598 | 0 | 0 | 0 | 0 | 2 mo | HOM |
| 1009934102 | 2 | SCO | 599626923 | 0.014915598 | 0 | 0 | 0 | 0 | 4 mo | WILD |
| 1009322937 | 2 | SCO | 599626923 | 0.014915598 | 0 | 0 | 0 | 0 | 8 mo | WILD |
| 1011080637 | 2 | SCO | 599626923 | 0.014915598 | 2.71E-06 | 1 | 0.000181575 | 67.04391136 | 8 mo | HOM |
| 1013609713 | 2 | SCO | 599626923 | 0.014915598 | 0 | 0 | 0 | 0 | 2 mo | WILD |
| 1010418842 | 1 | SCO | 599626923 | 0.014915598 | 0 | 0 | 0 | 0 | 8 mo | HOM |
| 1013288213 | 2 | SCO | 599626923 | 0.014915598 | 4.31E-06 | 0 | 0.000289191 | 0 | 4 mo | HOM |
| 1008976060 | 2 | SCO | 599626923 | 0.014915598 | 0 | 0 | 0 | 0 | 8 mo | WILD |
| 1013446243 | 1 | SCO | 599626923 | 0.014915598 | 0 | 0 | 0 | 0 | 4 mo | HOM |
| 1013288951 | 2 | SCO | 599626923 | 0.014915598 | 7.6E-06 | 1 | 0.000509322 | 67.04391136 | 4 mo | HOM |
| 1013978665 | 1 | PDTg | 599626927 | 0.049479646 | 0 | 0 | 0 | 0 | 2 mo | HOM |
| 1013445609 | 1 | PDTg | 599626927 | 0.049479646 | 0 | 0 | 0 | 0 | 4 mo | HOM |
| 1013609015 | 1 | PDTg | 599626927 | 0.049479646 | 1.18E-05 | 0 | 0.000238555 | 0 | 4 mo | HOM |
| 1013608106 | 2 | PDTg | 599626927 | 0.049479646 | 0 | 0 | 0 | 0 | 4 mo | HOM |
| 1011937543 | 1 | PDTg | 599626927 | 0.049479646 | 0 | 0 | 0 | 0 | 8 mo | HOM |
| 1010419538 | 2 | PDTg | 599626927 | 0.049479646 | 0 | 0 | 0 | 0 | 4 mo | WILD |
| 1010419187 | 1 | PDTg | 599626927 | 0.049479646 | 8.94E-06 | 0 | 0.000180683 | 0 | 8 mo | HOM |
| 1013978366 | 1 | PDTg | 599626927 | 0.049479646 | 8.59E-08 | 0 | 1.74E-06 | 0 | 2 mo | HOM |
| 1014416155 | 2 | PDTg | 599626927 | 0.049479646 | 0 | 0 | 0 | 0 | 2 mo | HOM |
| 1013978065 | 2 | PDTg | 599626927 | 0.049479646 | 0 | 0 | 0 | 0 | 2 mo | HOM |
| 1009934102 | 2 | PDTg | 599626927 | 0.049479646 | 0 | 0 | 0 | 0 | 4 mo | WILD |
| 1009322937 | 2 | PDTg | 599626927 | 0.049479646 | 0 | 0 | 0 | 0 | 8 mo | WILD |
| 1011080637 | 2 | PDTg | 599626927 | 0.049479646 | 6.18E-05 | 0 | 0.001249033 | 0 | 8 mo | HOM |
| 1013609713 | 2 | PDTg | 599626927 | 0.049479646 | 0 | 0 | 0 | 0 | 2 mo | WILD |
| 1010418842 | 1 | PDTg | 599626927 | 0.049479646 | 9.38E-06 | 1 | 0.0001896 | 20.2103304 | 8 mo | HOM |
| 1013288213 | 2 | PDTg | 599626927 | 0.049479646 | 0 | 0 | 0 | 0 | 4 mo | HOM |
| 1008976060 | 2 | PDTg | 599626927 | 0.049479646 | 4.5E-05 | 0 | 0.000909306 | 0 | 8 mo | WILD |
| 1013446243 | 1 | PDTg | 599626927 | 0.049479646 | 6.05E-07 | 0 | 1.22E-05 | 0 | 4 mo | HOM |
| 1013288951 | 2 | PDTg | 599626927 | 0.049479646 | 2.29E-06 | 0 | 4.62E-05 | 0 | 4 mo | HOM |
| 1013978665 | 1 | MMI | 606826647 | 0.219085586 | 1.41E-06 | 0 | 6.44E-06 | 0 | 2 mo | HOM |
| 1013445609 | 1 | MMI | 606826647 | 0.219085586 | 0 | 0 | 0 | 0 | 4 mo | HOM |
| 1013609015 | 1 | MMI | 606826647 | 0.219085586 | 0 | 0 | 0 | 0 | 4 mo | HOM |
| 1013608106 | 2 | MMI | 606826647 | 0.219085586 | 0 | 0 | 0 | 0 | 4 mo | HOM |
| 1011937543 | 1 | MMI | 606826647 | 0.219085586 | 0.001002101 | 4 | 0.004574015 | 18.25770504 | 8 mo | HOM |
| 1010419538 | 2 | MMI | 606826647 | 0.219085586 | 1.67E-06 | 0 | 7.6E-06 | 0 | 4 mo | WILD |
| 1010419187 | 1 | MMI | 606826647 | 0.219085586 | 0.001456773 | 1 | 0.006649333 | 4.56442626 | 8 mo | HOM |
| 1013978366 | 1 | MMI | 606826647 | 0.219085586 | 1.21E-05 | 1 | 5.5E-05 | 4.56442626 | 2 mo | HOM |
| 1014416155 | 2 | MMI | 606826647 | 0.219085586 | 0 | 0 | 0 | 0 | 2 mo | HOM |
| 1013978065 | 2 | MMI | 606826647 | 0.219085586 | 0 | 0 | 0 | 0 | 2 mo | HOM |
| 1009934102 | 2 | MMI | 606826647 | 0.219085586 | 1.47E-21 | 0 | 6.73E-21 | 0 | 4 mo | WILD |
| 1009322937 | 2 | MMI | 606826647 | 0.219085586 | 0 | 0 | 0 | 0 | 8 mo | WILD |
| 1011080637 | 2 | MMI | 606826647 | 0.219085586 | 1.94E-05 | 1 | 8.84E-05 | 4.56442626 | 8 mo | HOM |
| 1013609713 | 2 | MMI | 606826647 | 0.219085586 | 0 | 0 | 0 | 0 | 2 mo | WILD |
| 1010418842 | 1 | MMI | 606826647 | 0.219085586 | 4E-08 | 0 | 1.82E-07 | 0 | 8 mo | HOM |
| 1013288213 | 2 | MMI | 606826647 | 0.219085586 | 0 | 0 | 0 | 0 | 4 mo | HOM |
| 1008976060 | 2 | MMI | 606826647 | 0.219085586 | 1.4E-05 | 3 | 6.38E-05 | 13.69327878 | 8 mo | WILD |

| | | | | | | | | | | |
|---|---|---|---|---|---|---|---|---|---|---|
| **1013446243** | 1 | MMI | 606826647 | 0.219085586 | 0 | 0 | 0 | 0 | 4 mo | HOM |
| **1013288951** | 2 | MMI | 606826647 | 0.219085586 | 0 | 0 | 0 | 0 | 4 mo | HOM |
| **1013978665** | 1 | MMm | 606826651 | 0.139888071 | 0 | 0 | 0 | 0 | 2 mo | HOM |
| **1013445609** | 1 | MMm | 606826651 | 0.139888071 | 0 | 0 | 0 | 0 | 4 mo | HOM |
| **1013609015** | 1 | MMm | 606826651 | 0.139888071 | 0 | 0 | 0 | 0 | 4 mo | HOM |
| **1013608106** | 2 | MMm | 606826651 | 0.139888071 | 0 | 0 | 0 | 0 | 4 mo | HOM |
| **1011937543** | 1 | MMm | 606826651 | 0.139888071 | 0.000753404 | 4 | 0.005385761 | 28.59428942 | 8 mo | HOM |
| **1010419538** | 2 | MMm | 606826651 | 0.139888071 | 1.22E-07 | 0 | 8.7E-07 | 0 | 4 mo | WILD |
| **1010419187** | 1 | MMm | 606826651 | 0.139888071 | 4.73E-05 | 0 | 0.000338444 | 0 | 8 mo | HOM |
| **1013978366** | 1 | MMm | 606826651 | 0.139888071 | 2.32E-10 | 0 | 1.66E-09 | 0 | 2 mo | HOM |
| **1014416155** | 2 | MMm | 606826651 | 0.139888071 | 0 | 0 | 0 | 0 | 2 mo | HOM |
| **1013978065** | 2 | MMm | 606826651 | 0.139888071 | 0 | 0 | 0 | 0 | 2 mo | HOM |
| **1009934102** | 2 | MMm | 606826651 | 0.139888071 | 1.72E-06 | 0 | 1.23E-05 | 0 | 4 mo | WILD |
| **1009322937** | 2 | MMm | 606826651 | 0.139888071 | 0 | 0 | 0 | 0 | 8 mo | WILD |
| **1011080637** | 2 | MMm | 606826651 | 0.139888071 | 7.35E-06 | 1 | 5.25E-05 | 7.148572355 | 8 mo | HOM |
| **1013609713** | 2 | MMm | 606826651 | 0.139888071 | 0 | 0 | 0 | 0 | 2 mo | WILD |
| **1010418842** | 1 | MMm | 606826651 | 0.139888071 | 5.44E-05 | 5 | 0.000388822 | 35.74286177 | 8 mo | HOM |
| **1013288213** | 2 | MMm | 606826651 | 0.139888071 | 4.09E-06 | 0 | 2.92E-05 | 0 | 4 mo | HOM |
| **1008976060** | 2 | MMm | 606826651 | 0.139888071 | 5.49E-06 | 1 | 3.93E-05 | 7.148572355 | 8 mo | WILD |
| **1013446243** | 1 | MMm | 606826651 | 0.139888071 | 0 | 0 | 0 | 0 | 4 mo | HOM |
| **1013288951** | 2 | MMm | 606826651 | 0.139888071 | 0 | 0 | 0 | 0 | 4 mo | HOM |
| **1013978665** | 1 | MMp | 606826655 | 0.039022863 | 0 | 0 | 0 | 0 | 2 mo | HOM |
| **1013445609** | 1 | MMp | 606826655 | 0.039022863 | 0 | 0 | 0 | 0 | 4 mo | HOM |
| **1013609015** | 1 | MMp | 606826655 | 0.039022863 | 2.3E-06 | 0 | 5.91E-05 | 0 | 4 mo | HOM |
| **1013608106** | 2 | MMp | 606826655 | 0.039022863 | 0 | 0 | 0 | 0 | 4 mo | HOM |
| **1011937543** | 1 | MMp | 606826655 | 0.039022863 | 3.26E-07 | 0 | 8.36E-06 | 0 | 8 mo | HOM |
| **1010419538** | 2 | MMp | 606826655 | 0.039022863 | 0 | 0 | 0 | 0 | 4 mo | WILD |
| **1010419187** | 1 | MMp | 606826655 | 0.039022863 | 1.85E-05 | 1 | 0.000475261 | 25.62600319 | 8 mo | HOM |
| **1013978366** | 1 | MMp | 606826655 | 0.039022863 | 0 | 0 | 0 | 0 | 2 mo | HOM |
| **1014416155** | 2 | MMp | 606826655 | 0.039022863 | 0 | 0 | 0 | 0 | 2 mo | HOM |
| **1013978065** | 2 | MMp | 606826655 | 0.039022863 | 0 | 0 | 0 | 0 | 2 mo | HOM |
| **1009934102** | 2 | MMp | 606826655 | 0.039022863 | 0 | 0 | 0 | 0 | 4 mo | WILD |
| **1009322937** | 2 | MMp | 606826655 | 0.039022863 | 0 | 0 | 0 | 0 | 8 mo | WILD |
| **1011080637** | 2 | MMp | 606826655 | 0.039022863 | 9E-05 | 3 | 0.002307558 | 76.87800956 | 8 mo | HOM |
| **1013609713** | 2 | MMp | 606826655 | 0.039022863 | 0 | 0 | 0 | 0 | 2 mo | WILD |
| **1010418842** | 1 | MMp | 606826655 | 0.039022863 | 6.9E-06 | 2 | 0.000176833 | 51.25200637 | 8 mo | HOM |
| **1013288213** | 2 | MMp | 606826655 | 0.039022863 | 5.18E-06 | 2 | 0.000132819 | 51.25200637 | 4 mo | HOM |
| **1008976060** | 2 | MMp | 606826655 | 0.039022863 | 3.48E-05 | 1 | 0.000890772 | 25.62600319 | 8 mo | WILD |
| **1013446243** | 1 | MMp | 606826655 | 0.039022863 | 0 | 0 | 0 | 0 | 4 mo | HOM |
| **1013288951** | 2 | MMp | 606826655 | 0.039022863 | 0 | 0 | 0 | 0 | 4 mo | HOM |
| **1013978665** | 1 | MMd | 606826659 | 0.079181031 | 0 | 0 | 0 | 0 | 2 mo | HOM |
| **1013445609** | 1 | MMd | 606826659 | 0.079181031 | 0 | 0 | 0 | 0 | 4 mo | HOM |
| **1013609015** | 1 | MMd | 606826659 | 0.079181031 | 1.21E-05 | 1 | 0.000152507 | 12.6292875 | 4 mo | HOM |
| **1013608106** | 2 | MMd | 606826659 | 0.079181031 | 0 | 0 | 0 | 0 | 4 mo | HOM |
| **1011937543** | 1 | MMd | 606826659 | 0.079181031 | 0.000122985 | 1 | 0.00155321 | 12.6292875 | 8 mo | HOM |
| **1010419538** | 2 | MMd | 606826659 | 0.079181031 | 0 | 0 | 0 | 0 | 4 mo | WILD |
| **1010419187** | 1 | MMd | 606826659 | 0.079181031 | 4.93E-05 | 1 | 0.000622688 | 12.6292875 | 8 mo | HOM |
| **1013978366** | 1 | MMd | 606826659 | 0.079181031 | 0 | 0 | 0 | 0 | 2 mo | HOM |
| **1014416155** | 2 | MMd | 606826659 | 0.079181031 | 0 | 0 | 0 | 0 | 2 mo | HOM |
| **1013978065** | 2 | MMd | 606826659 | 0.079181031 | 4.08E-06 | 1 | 5.15E-05 | 12.6292875 | 2 mo | HOM |
| **1009934102** | 2 | MMd | 606826659 | 0.079181031 | 6.1E-06 | 1 | 7.7E-05 | 12.6292875 | 4 mo | WILD |
| **1009322937** | 2 | MMd | 606826659 | 0.079181031 | 0 | 0 | 0 | 0 | 8 mo | WILD |
| **1011080637** | 2 | MMd | 606826659 | 0.079181031 | 0 | 0 | 0 | 0 | 8 mo | HOM |
| **1013609713** | 2 | MMd | 606826659 | 0.079181031 | 0 | 0 | 0 | 0 | 2 mo | WILD |
| **1010418842** | 1 | MMd | 606826659 | 0.079181031 | 0 | 0 | 0 | 0 | 8 mo | HOM |
| **1013288213** | 2 | MMd | 606826659 | 0.079181031 | 0 | 0 | 0 | 0 | 4 mo | HOM |
| **1008976060** | 2 | MMd | 606826659 | 0.079181031 | 3.33E-05 | 1 | 0.000420816 | 12.6292875 | 8 mo | WILD |
| **1013446243** | 1 | MMd | 606826659 | 0.079181031 | 0 | 0 | 0 | 0 | 4 mo | HOM |
| **1013288951** | 2 | MMd | 606826659 | 0.079181031 | 3.97E-06 | 1 | 5.01E-05 | 12.6292875 | 4 mo | HOM |

| | | | | | | | | | | |
|---|---|---|---|---|---|---|---|---|---|---|
| **1013978665** | 1 | Pa4 | 606826663 | 0.016654617 | 0 | 0 | 0 | 0 | 2 mo | HOM |
| **1013445609** | 1 | Pa4 | 606826663 | 0.016654617 | 0 | 0 | 0 | 0 | 4 mo | HOM |
| **1013609015** | 1 | Pa4 | 606826663 | 0.016654617 | 0 | 0 | 0 | 0 | 4 mo | HOM |
| **1013608106** | 2 | Pa4 | 606826663 | 0.016654617 | 0 | 0 | 0 | 0 | 4 mo | HOM |
| **1011937543** | 1 | Pa4 | 606826663 | 0.016654617 | 0 | 0 | 0 | 0 | 8 mo | HOM |
| **1010419538** | 2 | Pa4 | 606826663 | 0.016654617 | 0 | 0 | 0 | 0 | 4 mo | WILD |
| **1010419187** | 1 | Pa4 | 606826663 | 0.016654617 | 0 | 0 | 0 | 0 | 8 mo | HOM |
| **1013978366** | 1 | Pa4 | 606826663 | 0.016654617 | 0 | 0 | 0 | 0 | 2 mo | HOM |
| **1014416155** | 2 | Pa4 | 606826663 | 0.016654617 | 0 | 0 | 0 | 0 | 2 mo | HOM |
| **1013978065** | 2 | Pa4 | 606826663 | 0.016654617 | 0 | 0 | 0 | 0 | 2 mo | HOM |
| **1009934102** | 2 | Pa4 | 606826663 | 0.016654617 | 0 | 0 | 0 | 0 | 4 mo | WILD |
| **1009322937** | 2 | Pa4 | 606826663 | 0.016654617 | 0 | 0 | 0 | 0 | 8 mo | WILD |
| **1011080637** | 2 | Pa4 | 606826663 | 0.016654617 | 0 | 0 | 0 | 0 | 8 mo | HOM |
| **1013609713** | 2 | Pa4 | 606826663 | 0.016654617 | 0 | 0 | 0 | 0 | 2 mo | WILD |
| **1010418842** | 1 | Pa4 | 606826663 | 0.016654617 | 2.26E-05 | 1 | 0.001356625 | 60.04340892 | 8 mo | HOM |
| **1013288213** | 2 | Pa4 | 606826663 | 0.016654617 | 0 | 0 | 0 | 0 | 4 mo | HOM |
| **1008976060** | 2 | Pa4 | 606826663 | 0.016654617 | 0 | 0 | 0 | 0 | 8 mo | WILD |
| **1013446243** | 1 | Pa4 | 606826663 | 0.016654617 | 0 | 0 | 0 | 0 | 4 mo | HOM |
| **1013288951** | 2 | Pa4 | 606826663 | 0.016654617 | 0 | 0 | 0 | 0 | 4 mo | HOM |
| **1013978665** | 1 | PN | 607344830 | 0.022392971 | 0 | 0 | 0 | 0 | 2 mo | HOM |
| **1013445609** | 1 | PN | 607344830 | 0.022392971 | 0 | 0 | 0 | 0 | 4 mo | HOM |
| **1013609015** | 1 | PN | 607344830 | 0.022392971 | 0 | 0 | 0 | 0 | 4 mo | HOM |
| **1013608106** | 2 | PN | 607344830 | 0.022392971 | 0 | 0 | 0 | 0 | 4 mo | HOM |
| **1011937543** | 1 | PN | 607344830 | 0.022392971 | 0 | 0 | 0 | 0 | 8 mo | HOM |
| **1010419538** | 2 | PN | 607344830 | 0.022392971 | 0 | 0 | 0 | 0 | 4 mo | WILD |
| **1010419187** | 1 | PN | 607344830 | 0.022392971 | 0 | 0 | 0 | 0 | 8 mo | HOM |
| **1013978366** | 1 | PN | 607344830 | 0.022392971 | 0 | 0 | 0 | 0 | 2 mo | HOM |
| **1014416155** | 2 | PN | 607344830 | 0.022392971 | 0 | 0 | 0 | 0 | 2 mo | HOM |
| **1013978065** | 2 | PN | 607344830 | 0.022392971 | 0 | 0 | 0 | 0 | 2 mo | HOM |
| **1009934102** | 2 | PN | 607344830 | 0.022392971 | 0 | 0 | 0 | 0 | 4 mo | WILD |
| **1009322937** | 2 | PN | 607344830 | 0.022392971 | 0 | 0 | 0 | 0 | 8 mo | WILD |
| **1011080637** | 2 | PN | 607344830 | 0.022392971 | 0 | 0 | 0 | 0 | 8 mo | HOM |
| **1013609713** | 2 | PN | 607344830 | 0.022392971 | 0 | 0 | 0 | 0 | 2 mo | WILD |
| **1010418842** | 1 | PN | 607344830 | 0.022392971 | 0 | 0 | 0 | 0 | 8 mo | HOM |
| **1013288213** | 2 | PN | 607344830 | 0.022392971 | 0 | 0 | 0 | 0 | 4 mo | HOM |
| **1008976060** | 2 | PN | 607344830 | 0.022392971 | 0 | 0 | 0 | 0 | 8 mo | WILD |
| **1013446243** | 1 | PN | 607344830 | 0.022392971 | 0 | 0 | 0 | 0 | 4 mo | HOM |
| **1013288951** | 2 | PN | 607344830 | 0.022392971 | 0 | 0 | 0 | 0 | 4 mo | HOM |
| **1013978665** | 1 | IPR | 607344834 | 0.082768272 | 0 | 0 | 0 | 0 | 2 mo | HOM |
| **1013445609** | 1 | IPR | 607344834 | 0.082768272 | 0 | 0 | 0 | 0 | 4 mo | HOM |
| **1013609015** | 1 | IPR | 607344834 | 0.082768272 | 0 | 0 | 0 | 0 | 4 mo | HOM |
| **1013608106** | 2 | IPR | 607344834 | 0.082768272 | 0 | 0 | 0 | 0 | 4 mo | HOM |
| **1011937543** | 1 | IPR | 607344834 | 0.082768272 | 3.38E-05 | 0 | 0.000408242 | 0 | 8 mo | HOM |
| **1010419538** | 2 | IPR | 607344834 | 0.082768272 | 1.83E-07 | 0 | 2.21E-06 | 0 | 4 mo | WILD |
| **1010419187** | 1 | IPR | 607344834 | 0.082768272 | 0.000133197 | 2 | 0.001609281 | 24.1638488 | 8 mo | HOM |
| **1013978366** | 1 | IPR | 607344834 | 0.082768272 | 0 | 0 | 0 | 0 | 2 mo | HOM |
| **1014416155** | 2 | IPR | 607344834 | 0.082768272 | 0 | 0 | 0 | 0 | 2 mo | HOM |
| **1013978065** | 2 | IPR | 607344834 | 0.082768272 | 0 | 0 | 0 | 0 | 2 mo | HOM |
| **1009934102** | 2 | IPR | 607344834 | 0.082768272 | 1.42E-08 | 0 | 1.72E-07 | 0 | 4 mo | WILD |
| **1009322937** | 2 | IPR | 607344834 | 0.082768272 | 0 | 0 | 0 | 0 | 8 mo | WILD |
| **1011080637** | 2 | IPR | 607344834 | 0.082768272 | 4.5E-06 | 1 | 5.43E-05 | 12.0819244 | 8 mo | HOM |
| **1013609713** | 2 | IPR | 607344834 | 0.082768272 | 0 | 0 | 0 | 0 | 2 mo | WILD |
| **1010418842** | 1 | IPR | 607344834 | 0.082768272 | 3E-05 | 3 | 0.000361998 | 36.24577319 | 8 mo | HOM |
| **1013288213** | 2 | IPR | 607344834 | 0.082768272 | 0 | 0 | 0 | 0 | 4 mo | HOM |
| **1008976060** | 2 | IPR | 607344834 | 0.082768272 | 7.36E-05 | 2 | 0.000888641 | 24.1638488 | 8 mo | WILD |
| **1013446243** | 1 | IPR | 607344834 | 0.082768272 | 1.11E-06 | 0 | 1.34E-05 | 0 | 4 mo | HOM |
| **1013288951** | 2 | IPR | 607344834 | 0.082768272 | 1.1E-05 | 1 | 0.000132492 | 12.0819244 | 4 mo | HOM |
| **1013978665** | 1 | IPC | 607344838 | 0.070539508 | 0 | 0 | 0 | 0 | 2 mo | HOM |
| **1013445609** | 1 | IPC | 607344838 | 0.070539508 | 0 | 0 | 0 | 0 | 4 mo | HOM |

| | | | | | | | | | | |
|---|---|---|---|---|---|---|---|---|---|---|
| **1013609015** | 1 | IPC | 607344838 | 0.070539508 | 0 | 0 | 0 | 0 | 4 mo | HOM |
| **1013608106** | 2 | IPC | 607344838 | 0.070539508 | 0 | 0 | 0 | 0 | 4 mo | HOM |
| **1011937543** | 1 | IPC | 607344838 | 0.070539508 | 0 | 0 | 0 | 0 | 8 mo | HOM |
| **1010419538** | 2 | IPC | 607344838 | 0.070539508 | 0 | 0 | 0 | 0 | 4 mo | WILD |
| **1010419187** | 1 | IPC | 607344838 | 0.070539508 | 0 | 0 | 0 | 0 | 8 mo | HOM |
| **1013978366** | 1 | IPC | 607344838 | 0.070539508 | 0 | 0 | 0 | 0 | 2 mo | HOM |
| **1014416155** | 2 | IPC | 607344838 | 0.070539508 | 0 | 0 | 0 | 0 | 2 mo | HOM |
| **1013978065** | 2 | IPC | 607344838 | 0.070539508 | 0 | 0 | 0 | 0 | 2 mo | HOM |
| **1009934102** | 2 | IPC | 607344838 | 0.070539508 | 1.1E-05 | 1 | 0.000155442 | 14.17645268 | 4 mo | WILD |
| **1009322937** | 2 | IPC | 607344838 | 0.070539508 | 1.19E-05 | 1 | 0.000168852 | 14.17645268 | 8 mo | WILD |
| **1011080637** | 2 | IPC | 607344838 | 0.070539508 | 0 | 0 | 0 | 0 | 8 mo | HOM |
| **1013609713** | 2 | IPC | 607344838 | 0.070539508 | 0 | 0 | 0 | 0 | 2 mo | WILD |
| **1010418842** | 1 | IPC | 607344838 | 0.070539508 | 3.87E-06 | 1 | 5.48E-05 | 14.17645268 | 8 mo | HOM |
| **1013288213** | 2 | IPC | 607344838 | 0.070539508 | 0 | 0 | 0 | 0 | 4 mo | HOM |
| **1008976060** | 2 | IPC | 607344838 | 0.070539508 | 0 | 0 | 0 | 0 | 8 mo | WILD |
| **1013446243** | 1 | IPC | 607344838 | 0.070539508 | 0 | 0 | 0 | 0 | 4 mo | HOM |
| **1013288951** | 2 | IPC | 607344838 | 0.070539508 | 0 | 0 | 0 | 0 | 4 mo | HOM |
| **1013978665** | 1 | IPA | 607344842 | 0.025866889 | 0 | 0 | 0 | 0 | 2 mo | HOM |
| **1013445609** | 1 | IPA | 607344842 | 0.025866889 | 0 | 0 | 0 | 0 | 4 mo | HOM |
| **1013609015** | 1 | IPA | 607344842 | 0.025866889 | 0 | 0 | 0 | 0 | 4 mo | HOM |
| **1013608106** | 2 | IPA | 607344842 | 0.025866889 | 0 | 0 | 0 | 0 | 4 mo | HOM |
| **1011937543** | 1 | IPA | 607344842 | 0.025866889 | 0 | 0 | 0 | 0 | 8 mo | HOM |
| **1010419538** | 2 | IPA | 607344842 | 0.025866889 | 0 | 0 | 0 | 0 | 4 mo | WILD |
| **1010419187** | 1 | IPA | 607344842 | 0.025866889 | 0 | 0 | 0 | 0 | 8 mo | HOM |
| **1013978366** | 1 | IPA | 607344842 | 0.025866889 | 0 | 0 | 0 | 0 | 2 mo | HOM |
| **1014416155** | 2 | IPA | 607344842 | 0.025866889 | 0 | 0 | 0 | 0 | 2 mo | HOM |
| **1013978065** | 2 | IPA | 607344842 | 0.025866889 | 0 | 0 | 0 | 0 | 2 mo | HOM |
| **1009934102** | 2 | IPA | 607344842 | 0.025866889 | 0 | 0 | 0 | 0 | 4 mo | WILD |
| **1009322937** | 2 | IPA | 607344842 | 0.025866889 | 0 | 0 | 0 | 0 | 8 mo | WILD |
| **1011080637** | 2 | IPA | 607344842 | 0.025866889 | 0 | 0 | 0 | 0 | 8 mo | HOM |
| **1013609713** | 2 | IPA | 607344842 | 0.025866889 | 0 | 0 | 0 | 0 | 2 mo | WILD |
| **1010418842** | 1 | IPA | 607344842 | 0.025866889 | 0 | 0 | 0 | 0 | 8 mo | HOM |
| **1013288213** | 2 | IPA | 607344842 | 0.025866889 | 0 | 0 | 0 | 0 | 4 mo | HOM |
| **1008976060** | 2 | IPA | 607344842 | 0.025866889 | 0 | 0 | 0 | 0 | 8 mo | WILD |
| **1013446243** | 1 | IPA | 607344842 | 0.025866889 | 0 | 0 | 0 | 0 | 4 mo | HOM |
| **1013288951** | 2 | IPA | 607344842 | 0.025866889 | 0 | 0 | 0 | 0 | 4 mo | HOM |
| **1013978665** | 1 | IPL | 607344846 | 0.063886313 | 0 | 0 | 0 | 0 | 2 mo | HOM |
| **1013445609** | 1 | IPL | 607344846 | 0.063886313 | 0 | 0 | 0 | 0 | 4 mo | HOM |
| **1013609015** | 1 | IPL | 607344846 | 0.063886313 | 0 | 0 | 0 | 0 | 4 mo | HOM |
| **1013608106** | 2 | IPL | 607344846 | 0.063886313 | 0 | 0 | 0 | 0 | 4 mo | HOM |
| **1011937543** | 1 | IPL | 607344846 | 0.063886313 | 0.000136293 | 0 | 0.002133369 | 0 | 8 mo | HOM |
| **1010419538** | 2 | IPL | 607344846 | 0.063886313 | 0 | 0 | 0 | 0 | 4 mo | WILD |
| **1010419187** | 1 | IPL | 607344846 | 0.063886313 | 1.37E-05 | 1 | 0.000214631 | 15.65280508 | 8 mo | HOM |
| **1013978366** | 1 | IPL | 607344846 | 0.063886313 | 0 | 0 | 0 | 0 | 2 mo | HOM |
| **1014416155** | 2 | IPL | 607344846 | 0.063886313 | 0 | 0 | 0 | 0 | 2 mo | HOM |
| **1013978065** | 2 | IPL | 607344846 | 0.063886313 | 0 | 0 | 0 | 0 | 2 mo | HOM |
| **1009934102** | 2 | IPL | 607344846 | 0.063886313 | 0 | 0 | 0 | 0 | 4 mo | WILD |
| **1009322937** | 2 | IPL | 607344846 | 0.063886313 | 5.22E-06 | 1 | 8.17E-05 | 15.65280508 | 8 mo | WILD |
| **1011080637** | 2 | IPL | 607344846 | 0.063886313 | 0 | 0 | 0 | 0 | 8 mo | HOM |
| **1013609713** | 2 | IPL | 607344846 | 0.063886313 | 0 | 0 | 0 | 0 | 2 mo | WILD |
| **1010418842** | 1 | IPL | 607344846 | 0.063886313 | 1.52E-05 | 2 | 0.000238698 | 31.30561016 | 8 mo | HOM |
| **1013288213** | 2 | IPL | 607344846 | 0.063886313 | 0 | 0 | 0 | 0 | 4 mo | HOM |
| **1008976060** | 2 | IPL | 607344846 | 0.063886313 | 3.51E-05 | 3 | 0.000549105 | 46.95841524 | 8 mo | WILD |
| **1013446243** | 1 | IPL | 607344846 | 0.063886313 | 0 | 0 | 0 | 0 | 4 mo | HOM |
| **1013288951** | 2 | IPL | 607344846 | 0.063886313 | 0 | 0 | 0 | 0 | 4 mo | HOM |
| **1013978665** | 1 | IPI | 607344850 | 0.032893024 | 0 | 0 | 0 | 0 | 2 mo | HOM |
| **1013445609** | 1 | IPI | 607344850 | 0.032893024 | 0 | 0 | 0 | 0 | 4 mo | HOM |
| **1013609015** | 1 | IPI | 607344850 | 0.032893024 | 0 | 0 | 0 | 0 | 4 mo | HOM |
| **1013608106** | 2 | IPI | 607344850 | 0.032893024 | 0 | 0 | 0 | 0 | 4 mo | HOM |

| | | | | | | | | | | |
|---|---|---|---|---|---|---|---|---|---|---|
| **1011937543** | 1 | IPI | 607344850 | 0.032893024 | 0 | 0 | 0 | 0 | 8 mo | HOM |
| **1010419538** | 2 | IPI | 607344850 | 0.032893024 | 0 | 0 | 0 | 0 | 4 mo | WILD |
| **1010419187** | 1 | IPI | 607344850 | 0.032893024 | 0 | 0 | 0 | 0 | 8 mo | HOM |
| **1013978366** | 1 | IPI | 607344850 | 0.032893024 | 0 | 0 | 0 | 0 | 2 mo | HOM |
| **1014416155** | 2 | IPI | 607344850 | 0.032893024 | 0 | 0 | 0 | 0 | 2 mo | HOM |
| **1013978065** | 2 | IPI | 607344850 | 0.032893024 | 0 | 0 | 0 | 0 | 2 mo | HOM |
| **1009934102** | 2 | IPI | 607344850 | 0.032893024 | 0 | 0 | 0 | 0 | 4 mo | WILD |
| **1009322937** | 2 | IPI | 607344850 | 0.032893024 | 0 | 0 | 0 | 0 | 8 mo | WILD |
| **1011080637** | 2 | IPI | 607344850 | 0.032893024 | 0 | 0 | 0 | 0 | 8 mo | HOM |
| **1013609713** | 2 | IPI | 607344850 | 0.032893024 | 0 | 0 | 0 | 0 | 2 mo | WILD |
| **1010418842** | 1 | IPI | 607344850 | 0.032893024 | 3.12E-06 | 0 | 9.49E-05 | 0 | 8 mo | HOM |
| **1013288213** | 2 | IPI | 607344850 | 0.032893024 | 0 | 0 | 0 | 0 | 4 mo | HOM |
| **1008976060** | 2 | IPI | 607344850 | 0.032893024 | 3.18E-06 | 1 | 9.68E-05 | 30.40158321 | 8 mo | WILD |
| **1013446243** | 1 | IPI | 607344850 | 0.032893024 | 0 | 0 | 0 | 0 | 4 mo | HOM |
| **1013288951** | 2 | IPI | 607344850 | 0.032893024 | 0 | 0 | 0 | 0 | 4 mo | HOM |
| **1013978665** | 1 | IPDM | 607344854 | 0.019646391 | 0 | 0 | 0 | 0 | 2 mo | HOM |
| **1013445609** | 1 | IPDM | 607344854 | 0.019646391 | 0 | 0 | 0 | 0 | 4 mo | HOM |
| **1013609015** | 1 | IPDM | 607344854 | 0.019646391 | 0 | 0 | 0 | 0 | 4 mo | HOM |
| **1013608106** | 2 | IPDM | 607344854 | 0.019646391 | 0 | 0 | 0 | 0 | 4 mo | HOM |
| **1011937543** | 1 | IPDM | 607344854 | 0.019646391 | 0 | 0 | 0 | 0 | 8 mo | HOM |
| **1010419538** | 2 | IPDM | 607344854 | 0.019646391 | 0 | 0 | 0 | 0 | 4 mo | WILD |
| **1010419187** | 1 | IPDM | 607344854 | 0.019646391 | 0 | 0 | 0 | 0 | 8 mo | HOM |
| **1013978366** | 1 | IPDM | 607344854 | 0.019646391 | 0 | 0 | 0 | 0 | 2 mo | HOM |
| **1014416155** | 2 | IPDM | 607344854 | 0.019646391 | 0 | 0 | 0 | 0 | 2 mo | HOM |
| **1013978065** | 2 | IPDM | 607344854 | 0.019646391 | 0 | 0 | 0 | 0 | 2 mo | HOM |
| **1009934102** | 2 | IPDM | 607344854 | 0.019646391 | 0 | 0 | 0 | 0 | 4 mo | WILD |
| **1009322937** | 2 | IPDM | 607344854 | 0.019646391 | 0 | 0 | 0 | 0 | 8 mo | WILD |
| **1011080637** | 2 | IPDM | 607344854 | 0.019646391 | 0 | 0 | 0 | 0 | 8 mo | HOM |
| **1013609713** | 2 | IPDM | 607344854 | 0.019646391 | 0 | 0 | 0 | 0 | 2 mo | WILD |
| **1010418842** | 1 | IPDM | 607344854 | 0.019646391 | 0 | 0 | 0 | 0 | 8 mo | HOM |
| **1013288213** | 2 | IPDM | 607344854 | 0.019646391 | 0 | 0 | 0 | 0 | 4 mo | HOM |
| **1008976060** | 2 | IPDM | 607344854 | 0.019646391 | 0 | 0 | 0 | 0 | 8 mo | WILD |
| **1013446243** | 1 | IPDM | 607344854 | 0.019646391 | 0 | 0 | 0 | 0 | 4 mo | HOM |
| **1013288951** | 2 | IPDM | 607344854 | 0.019646391 | 0 | 0 | 0 | 0 | 4 mo | HOM |
| **1013978665** | 1 | IPDL | 607344858 | 0.039066132 | 0 | 0 | 0 | 0 | 2 mo | HOM |
| **1013445609** | 1 | IPDL | 607344858 | 0.039066132 | 0 | 0 | 0 | 0 | 4 mo | HOM |
| **1013609015** | 1 | IPDL | 607344858 | 0.039066132 | 0 | 0 | 0 | 0 | 4 mo | HOM |
| **1013608106** | 2 | IPDL | 607344858 | 0.039066132 | 0 | 0 | 0 | 0 | 4 mo | HOM |
| **1011937543** | 1 | IPDL | 607344858 | 0.039066132 | 0 | 0 | 0 | 0 | 8 mo | HOM |
| **1010419538** | 2 | IPDL | 607344858 | 0.039066132 | 0 | 0 | 0 | 0 | 4 mo | WILD |
| **1010419187** | 1 | IPDL | 607344858 | 0.039066132 | 3.41E-06 | 0 | 8.74E-05 | 0 | 8 mo | HOM |
| **1013978366** | 1 | IPDL | 607344858 | 0.039066132 | 0 | 0 | 0 | 0 | 2 mo | HOM |
| **1014416155** | 2 | IPDL | 607344858 | 0.039066132 | 0 | 0 | 0 | 0 | 2 mo | HOM |
| **1013978065** | 2 | IPDL | 607344858 | 0.039066132 | 0 | 0 | 0 | 0 | 2 mo | HOM |
| **1009934102** | 2 | IPDL | 607344858 | 0.039066132 | 0 | 0 | 0 | 0 | 4 mo | WILD |
| **1009322937** | 2 | IPDL | 607344858 | 0.039066132 | 0 | 0 | 0 | 0 | 8 mo | WILD |
| **1011080637** | 2 | IPDL | 607344858 | 0.039066132 | 0 | 0 | 0 | 0 | 8 mo | HOM |
| **1013609713** | 2 | IPDL | 607344858 | 0.039066132 | 0 | 0 | 0 | 0 | 2 mo | WILD |
| **1010418842** | 1 | IPDL | 607344858 | 0.039066132 | 8.18E-06 | 1 | 0.000209513 | 25.59761995 | 8 mo | HOM |
| **1013288213** | 2 | IPDL | 607344858 | 0.039066132 | 0 | 0 | 0 | 0 | 4 mo | HOM |
| **1008976060** | 2 | IPDL | 607344858 | 0.039066132 | 4.61E-08 | 0 | 1.18E-06 | 0 | 8 mo | WILD |
| **1013446243** | 1 | IPDL | 607344858 | 0.039066132 | 0 | 0 | 0 | 0 | 4 mo | HOM |
| **1013288951** | 2 | IPDL | 607344858 | 0.039066132 | 0 | 0 | 0 | 0 | 4 mo | HOM |
| **1013978665** | 1 | IPRL | 607344862 | 0.019650512 | 0 | 0 | 0 | 0 | 2 mo | HOM |
| **1013445609** | 1 | IPRL | 607344862 | 0.019650512 | 0 | 0 | 0 | 0 | 4 mo | HOM |
| **1013609015** | 1 | IPRL | 607344862 | 0.019650512 | 0 | 0 | 0 | 0 | 4 mo | HOM |
| **1013608106** | 2 | IPRL | 607344862 | 0.019650512 | 0 | 0 | 0 | 0 | 4 mo | HOM |
| **1011937543** | 1 | IPRL | 607344862 | 0.019650512 | 5.29E-05 | 1 | 0.002693694 | 50.88926018 | 8 mo | HOM |
| **1010419538** | 2 | IPRL | 607344862 | 0.019650512 | 0 | 0 | 0 | 0 | 4 mo | WILD |

| | | | | | | | | | | |
|---|---|---|---|---|---|---|---|---|---|---|
| **1010419187** | 1 | IPRL | 607344862 | 0.019650512 | 0 | 0 | 0 | 0 | 8 mo | HOM |
| **1013978366** | 1 | IPRL | 607344862 | 0.019650512 | 0 | 0 | 0 | 0 | 2 mo | HOM |
| **1014416155** | 2 | IPRL | 607344862 | 0.019650512 | 0 | 0 | 0 | 0 | 2 mo | HOM |
| **1013978065** | 2 | IPRL | 607344862 | 0.019650512 | 0 | 0 | 0 | 0 | 2 mo | HOM |
| **1009934102** | 2 | IPRL | 607344862 | 0.019650512 | 0 | 0 | 0 | 0 | 4 mo | WILD |
| **1009322937** | 2 | IPRL | 607344862 | 0.019650512 | 0 | 0 | 0 | 0 | 8 mo | WILD |
| **1011080637** | 2 | IPRL | 607344862 | 0.019650512 | 9.53E-07 | 0 | 4.85E-05 | 0 | 8 mo | HOM |
| **1013609713** | 2 | IPRL | 607344862 | 0.019650512 | 0 | 0 | 0 | 0 | 2 mo | WILD |
| **1010418842** | 1 | IPRL | 607344862 | 0.019650512 | 0 | 0 | 0 | 0 | 8 mo | HOM |
| **1013288213** | 2 | IPRL | 607344862 | 0.019650512 | 0 | 0 | 0 | 0 | 4 mo | HOM |
| **1008976060** | 2 | IPRL | 607344862 | 0.019650512 | 6.24E-06 | 2 | 0.000317453 | 101.7785204 | 8 mo | WILD |
| **1013446243** | 1 | IPRL | 607344862 | 0.019650512 | 0 | 0 | 0 | 0 | 4 mo | HOM |
| **1013288951** | 2 | IPRL | 607344862 | 0.019650512 | 0 | 0 | 0 | 0 | 4 mo | HOM |
| **1013978665** | 1 | Su3 | 614454277 | 0.043384835 | 0 | 0 | 0 | 0 | 2 mo | HOM |
| **1013445609** | 1 | Su3 | 614454277 | 0.043384835 | 0 | 0 | 0 | 0 | 4 mo | HOM |
| **1013609015** | 1 | Su3 | 614454277 | 0.043384835 | 0 | 0 | 0 | 0 | 4 mo | HOM |
| **1013608106** | 2 | Su3 | 614454277 | 0.043384835 | 0 | 0 | 0 | 0 | 4 mo | HOM |
| **1011937543** | 1 | Su3 | 614454277 | 0.043384835 | 0 | 0 | 0 | 0 | 8 mo | HOM |
| **1010419538** | 2 | Su3 | 614454277 | 0.043384835 | 0 | 0 | 0 | 0 | 4 mo | WILD |
| **1010419187** | 1 | Su3 | 614454277 | 0.043384835 | 0 | 0 | 0 | 0 | 8 mo | HOM |
| **1013978366** | 1 | Su3 | 614454277 | 0.043384835 | 0 | 0 | 0 | 0 | 2 mo | HOM |
| **1014416155** | 2 | Su3 | 614454277 | 0.043384835 | 0 | 0 | 0 | 0 | 2 mo | HOM |
| **1013978065** | 2 | Su3 | 614454277 | 0.043384835 | 0 | 0 | 0 | 0 | 2 mo | HOM |
| **1009934102** | 2 | Su3 | 614454277 | 0.043384835 | 0 | 0 | 0 | 0 | 4 mo | WILD |
| **1009322937** | 2 | Su3 | 614454277 | 0.043384835 | 0 | 0 | 0 | 0 | 8 mo | WILD |
| **1011080637** | 2 | Su3 | 614454277 | 0.043384835 | 0 | 0 | 0 | 0 | 8 mo | HOM |
| **1013609713** | 2 | Su3 | 614454277 | 0.043384835 | 0 | 0 | 0 | 0 | 2 mo | WILD |
| **1010418842** | 1 | Su3 | 614454277 | 0.043384835 | 4.25E-07 | 0 | 9.79E-06 | 0 | 8 mo | HOM |
| **1013288213** | 2 | Su3 | 614454277 | 0.043384835 | 0 | 0 | 0 | 0 | 4 mo | HOM |
| **1008976060** | 2 | Su3 | 614454277 | 0.043384835 | 4.75E-06 | 0 | 0.000109394 | 0 | 8 mo | WILD |
| **1013446243** | 1 | Su3 | 614454277 | 0.043384835 | 0 | 0 | 0 | 0 | 4 mo | HOM |
| **1013288951** | 2 | Su3 | 614454277 | 0.043384835 | 0 | 0 | 0 | 0 | 4 mo | HOM |
