## Supplementary Table 5 for "Fibrillar Aβ causes profound microglial metabolic perturbations in a novel APP knock-in mouse model"

| Metabolite | Internal Stan | RT(QTRAP) | Q1 m/z | Q3 m/z | CE(QTRAP) | Ionization m |
|---|---|---|---|---|---|---|
| 1-Methylhist | Arginine-d4C | 6.55 | 170 | 124 | 20 | + |
| 1-Methylnico | 15N4-Inosine | 4.58 | 137 | 94 | 20 | + |
| 1-Methylxan | 13C5-Hypoxa | 2.9 | 167 | 110 | 30 | + |
| 1-Methylxan | 13C5-Hypoxa | 2.79 | 299 | 187.1 | 30 | + |
| 1,3-15N2-Uri | N/A | 5.22 | 171.1 | 143 | 20 | + |
| 13C10,15N5- | N/A | 3.97 | 283 | 146.1 | 27 | + |
| 13C3-Thiami | N/A | 5.15 | 268 | 122.07 | 25 | + |
| 13C5-Hypoxa | N/A | 3.82 | 142.2 | 124 | 20 | + |
| 13C5-Xantho | N/A | 4.62 | 290 | 153 | 20 | + |
| 15N4-Inosine | N/A | 4.42 | 273.1 | 141.01 | 20 | + |
| 2-Aminoben | Niacinamide | 1.05 | 138.1 | 92 | 20 | + |
| 3-Hydroxy-N | Arginine-d4C | 6.68 | 205.16 | 128.1 | 25 | + |
| 3-Hydroxybut | Methionine- | 1.3 | 105 | 87 | 20 | + |
| 3-Hydroxykyr | Niacinamide | 0.95 | 225.1 | 179.1 | 15 | + |
| 3-Hydroxytyr | 13C5-Hypoxa | 3.58 | 137.1 | 119 | 20 | + |
| 3-Methoxyty | Methionine- | 5.2 | 212.1 | 166.1 | 20 | + |
| 3-Methylxan | 13C5-Hypoxa | 3.1 | 167 | 69 | 30 | + |
| 3-Methylxan | 13C5-Hypoxa | 2.98 | 299.1 | 187.1 | 30 | + |
| 4-Hydroxypro | 6C13-Tyrosin | 5.9 | 132 | 68.1 | 20 | + |
| 4-Trimethyla | Arginine-d4C | 6.76 | 130.12 | 71.05 | 20 | + |
| 5-Glutamylal | 6C13-Phenyl | 7 | 219.1 | 90.1 | 12 | + |
| 5-Hydroxyind | Niacinamide | 1.6 | 192.07 | 146.1 | 20 | + |
| 5-Hydroxytry | Methionine- | 4.9 | 221.11 | 175 | 20 | + |
| 5'-Methylthi | 13C10,15N5- | 2.75 | 298 | 136 | 20 | + |
| 6-Aminourac | 15N4-Inosine | 4.3 | 128 | 85 | 19 | + |
| 6C13-Phenyl | N/A | 4.8 | 172.1 | 126.1 | 18 | + |
| 6C13-Tyrosin | N/A | 5.31 | 188.1 | 142.1 | 24 | + |
| 7-alpha-Hy | Niacinamide | 1.43 | 431.32 | 395.3 | 15 | + |
| 7-Methylgua | 15N4-Inosine | 4.15 | 166.11 | 124.1 | 25 | + |
| 7-Methylgua | 13C10,15N5- | 5.39 | 298 | 166 | 20 | + |
| 7-Methylxan | 13C5-Hypoxa | 3.3 | 167.1 | 69 | 30 | + |
| 7-Methylxan | 13C5-Hypoxa | 3.2 | 299.2 | 167.1 | 30 | + |
| 8-Hydroxy-de | 15N4-Inosine | 5.04 | 284.1 | 140 | 20 | + |
| 8-Hydroxygu | 15N4-Inosine | 4.8 | 300 | 168 | 20 | + |
| 8-Oxo-deoxy | N/A | 4.92 | 287.3 | 171 | 20 | + |
| 9-Hexadecen | Palmitoylcar | 3 | 398.3 | 85 | 35 | + |
| Acetylcarniti | Acetylcarniti | 4.2 | 204.1 | 85 | 25 | + |
| Acetylcarniti | N/A | 4.2 | 207.14 | 85 | 25 | + |
| Adenine | 13C5-Hypoxa | 3.92 | 136.1 | 119 | 30 | + |
| Adenosine | 13C10,15N5- | 3.98 | 268.15 | 136.1 | 27 | + |
| Adenosine di | Arginine-d4C | 7.35 | 560.1 | 136.1 | 30 | + |
| Adenosylcob | 6C13-Phenyl | 6 | 801.5 | 676.3 | 31 | + |
| AICA-ribosid | 15N4-Inosine | 4.25 | 259 | 110 | 20 | + |
| Alanine | Alanine-d4 | 5.78 | 90.1 | 44.2 | 20 | + |

| | | | | | | |
|---|---|---|---|---|---|---|
| Alanine-d4 | N/A | 5.78 | 94.08 | 48.1 | 25 | + |
| Alpha-Tocopl | 6C13-Phenyla | 0.93 | 431.4 | 165.1 | 20 | + |
| Arabitol | 6C13-Phenyla | 4.7 | 153.1 | 153.1 | 10 | + |
| Arginine | Arginine-d4C | 6.75 | 175.12 | 70 | 27 | + |
| Arginine-d4C | N/A | 6.74 | 180.1 | 75 | 27 | + |
| Ascorbate | 13C5-Hypoxa | 3.8 | 177 | 95 | 20 | + |
| Asparagine | N15C13-Glyc | 6.21 | 133.06 | 74.02 | 15 | + |
| Aspartic acid | N15C13-Glyc | 6.25 | 134.04 | 74.02 | 18 | + |
| Aspartic acid | IS | 6.25 | 137.04 | 75.02 | 18 | + |
| Asymmetric | N15C13-Glyc | 6.3 | 203.15 | 70.3 | 40 | + |
| Betaine | Betaine-d11 | 5 | 118.1 | 59.1 | 20 | + |
| Betaine-d11 | IS | 5.1 | 129.1 | 68.1 | 20 | + |
| Bilirubin | Niacinamide | 0.92 | 585.3 | 299.1 | 20 | + |
| Butyrobetain | Carnitine-d9 | 4.3 | 146.1 | 87 | 20 | + |
| Butyrylcarnit | Butyrylcarnit | 3.76 | 232.2 | 85 | 30 | + |
| Butyrylcarnit | NA | 3.93 | 235.16 | 85 | 30 | + |
| Cadaverine | 6C13-Phenyla | 7.21 | 102.9 | 86.1 | 14 | + |
| cADPRibose | Arginine-d4C | 7.4 | 542.1 | 136.1 | 30 | + |
| Caffeine | Niacinamide | 1.26 | 195.1 | 138 | 20 | + |
| Capsaicin | Niacinamide | 1.37 | 306.2 | 137.1 | 20 | + |
| Carnitine | Carnitine-d9 | 5.1 | 162.1 | 103.05 | 20 | + |
| Carnitine-d9 | N/A | 5.1 | 171.16 | 85 | 20 | + |
| Choline | Acetylcarniti | 4.04 | 104 | 60 | 21 | + |
| Citrulline | Citrulline-d2 | 6.24 | 176.1 | 113.1 | 12 | + |
| Citrulline-d2 | N/A | 6.24 | 178.11 | 115.1 | 12 | + |
| Cotinine | Niacinamide | 1.6 | 177.1 | 80 | 32 | + |
| Creatine | Alanine-d4 | 5.66 | 132 | 90 | 18 | + |
| Creatinine | 15N4-Inosine | 4.15 | 114 | 44.1 | 22 | + |
| Creatinine-d3 | N/A | 4.17 | 117 | 86 | 15 | + |
| Curcumin | Niacinamide | 1.05 | 369 | 177 | 30 | + |
| Cyanocobalai | 6C13-Phenyla | 6.09 | 678.5 | 997.5 | 28 | + |
| Cystathionine | 6C13-Phenyla | 7 | 223.1 | 134 | 15 | + |
| Cysteine | Alanine-d4 | 5.54 | 122.03 | 76.02 | 15 | + |
| Cysteinylglyc | 6C13-Phenyla | 7 | 179.1 | 162.2 | 15 | + |
| Cystine | Arginine-d4C | 7.25 | 241 | 74 | 25 | + |
| Cytosine | 6C13-Phenyla | 4.98 | 112.1 | 94.9 | 25 | + |
| Decanoylcarr | Octanoylcarr | 3.22 | 316.2 | 85 | 35 | + |
| Demethoxycu | 6C13-Phenyla | 1.41 | 339.1 | 255.2 | 25 | + |
| Deoxyadenos | 13C5-Hypoxa | 3.55 | 252.1 | 136 | 20 | + |
| Deoxyguanos | 15N4-Inosine | 4.62 | 268.1 | 152 | 20 | + |
| Deoxyinosine | 13C5-Hypoxa | 3.5 | 253 | 137 | 20 | + |
| Dimethyletha | 15N4-Inosine | 4.3 | 90 | 72 | 15 | + |
| Dimethylglyc | Alanine-d4 | 5.5 | 104.02 | 58 | 21 | + |
| DNL201 | Niacinamide | 1 | 340.3 | 313.2 | 20 | + |
| Dodecanoylca | Isovalerylcar | 3.8 | 343.3 | 85 | 30 | + |

| | | | | | | |
|---|---|---|---|---|---|---|
| Dopa | Alanine-d4 | 5.7 | 198 | 152 | 20 | + |
| Dopamine | 15N4-Inosine | 4.64 | 154 | 137 | 13 | + |
| Dopamine 3- | 13C5-Hypoxa | 3.7 | 234 | 137.1 | 15 | + |
| Dopamine 4- | 13C5-Hypoxa | 3.9 | 234.1 | 137.2 | 15 | + |
| Epinephrine | 6C13-Phenyla | 4.8 | 166 | 107 | 20 | + |
| Ergothionein | Methionine-( | 5.6 | 230.1 | 127 | 22 | + |
| Ethanolamin | Methionine-( | 5.14 | 62 | 44.2 | 12 | + |
| Formylanthra | Niacinamide | 1.3 | 166 | 120 | 16 | + |
| Gamma-Ami | 6C13-Phenyla | 5.08 | 104.1 | 87 | 13 | + |
| Gamma-Glu | 6C13-Phenyla | 6.5 | 251.1 | 122.02 | 15 | + |
| Glucosamine | Glutamic aci | 6.45 | 180 | 162 | 20 | + |
| Glucose | 6C13-Phenyla | 5.15 | 202.8 | 202.8 | 10 | + |
| Glutamic aci | Glutamic aci | 5.98 | 148.06 | 84.04 | 20 | + |
| Glutamic aci | N/A | 5.98 | 151 | 133.1 | 20 | + |
| Glutamine | Glutamic aci | 6.09 | 147.08 | 84 | 15 | + |
| Glutathione | Glutamic aci | 6.3 | 308.1 | 179.1 | 17 | + |
| Glutathione ( | Arginine-d4C | 7.6 | 613.2 | 355.1 | 25 | + |
| Glutathione- | IS | 6.3 | 311.3 | 182.1 | 16 | + |
| Glycerophosp | N15C13-Glyc | 6.12 | 258.1 | 104 | 16 | + |
| Glycerophosphorylcholine | | | 259.1 | 103.9 | 16 | + |
| Glycine | N15C13-Glyc | 5.96 | 76.04 | 30 | 25 | + |
| Guanine | 15N4-Inosine | 4.6 | 152.2 | 110 | 20 | + |
| Guanosine | 15N4-Inosine | 5 | 284.1 | 152 | 20 | + |
| Hexanoylcarn | Propionylcart | 3.5 | 260.2 | 85 | 35 | + |
| Histamine | N15C13-Glyc | 6.35 | 112.09 | 95 | 20 | + |
| Histidine | Arginine-d4C | 6.81 | 156.08 | 110.07 | 16 | + |
| Homocystein | Valine-d8 | 5.23 | 136.12 | 90.1 | 17 | + |
| Homoserine | N15C13-Glyc | 5.91 | 120.15 | 56.2 | 24 | + |
| Hydroxybutyr | Carnitine-d9 | 4.89 | 248.2 | 85 | 30 | + |
| Hydroxyhexan | Carnitine-d9 | 4.5 | 276.2 | 85 | 30 | + |
| Hydroxyisova | Carnitine-d9 | 4.6 | 262.2 | 85 | 30 | + |
| Hydroxypropi | Propionylcart | 4.9 | 234.1 | 85 | 30 | + |
| Hypoxanthine | 13C5-Hypoxa | 3.82 | 137.2 | 118.8 | 20 | + |
| Imidazoleace | Alanine-d4 | 5.3 | 127 | 81 | 20 | + |
| Indole | Niacinamide | 1 | 118 | 91 | 20 | + |
| Inosine | 15N4-Inosine | 4.41 | 269.1 | 137.01 | 20 | + |
| Isovalerylcarn | Propionylcart | 3.77 | 246.2 | 85 | 33 | + |
| Isovalerylcarn | NA | 3.8 | 255.2 | 85 | 30 | + |
| Kynurenic aci | 15N4-Inosine | 4.27 | 190.05 | 116 | 36 | + |
| Kynurenine | 6C13-Phenyla | 4.81 | 209 | 146 | 25 | + |
| Leucine | Leucine-d3 | 4.84 | 132.1 | 86.01 | 15 | + |
| Leucine-d3 | N/A | 4.84 | 135.1 | 89.1 | 15 | + |
| Linoleyl carni | Palmitoylcarn | 2.96 | 424.3 | 85 | 45 | + |
| LPC(18:1(d7) | N/A | 3.97 | 529.3 | 184.1 | 40 | + |
| Lysine | Arginine-d4C | 6.83 | 147.11 | 84 | 20 | + |

| | | | | | | |
|---|---|---|---|---|---|---|
| Mannitol | 6C13-Phenyla | 5.36 | 183 | 69 | 20 | + |
| Mannose | 6C13-Phenyla | 5.5 | 202.81 | 202.81 | 15 | + |
| Melatonin | 15N4-Inosine | 3.9 | 233.1 | 174.1 | 20 | + |
| Methionine | Methionine-c | 5.1 | 150.06 | 104.05 | 10 | + |
| Methionine S | N15C13-Glyc | 6.07 | 166.05 | 74.02 | 20 | + |
| Methionine-c | N/A | 5.1 | 153.07 | 107 | 20 | + |
| Methylcobala | 6C13-Phenyla | 6.04 | 673 | 971.5 | 42 | + |
| Myristoylcarn | Myristoylcarn | 3.3 | 372 | 85 | 40 | + |
| Myristoylcarn | NA | 3.3 | 381.36 | 85 | 35 | + |
| N-Acetylalan | Octanoylcarr | 1.8 | 160.1 | 72.1 | 18 | + |
| N-Acetylaspa | 13C5-Hypoxa | 3.56 | 176.06 | 176.06 | 10 | + |
| N-Acetylcyste | 6C13-Phenyla | 1.3 | 163 | 121.2 | 15 | + |
| N-Acetylgluta | Glutamic aci | 3.49 | 189 | 130 | 20 | + |
| N-Acetylneur | N15C13-Glyc | 6.5 | 310.1 | 274 | 15 | + |
| N-Acetylputr | Leucine-d3 | 4.89 | 131.1 | 114 | 12 | + |
| N-Acetylserin | 6C13-Phenyla | 3.55 | 148 | 106 | 14 | + |
| N-Alpha-acet | Alanine-d4 | 5.61 | 189 | 84 | 18 | + |
| N,N'-Bis(gan | 6C13-Phenyla | 7 | 499.1 | 453.1 | 15 | + |
| N1-Acetylspe | N15C13-Glyc | 6.5 | 188.2 | 171 | 19 | + |
| N1-Acetylspe | IS | 6.5 | 194.3 | 106.2 | 19 | + |
| N1-Acetylspe | Arginine-d4C | 7.7 | 245.2 | 100.1 | 22 | + |
| N1-N12-Diac | N15C13-Glyc | 6.1 | 287.2 | 100.1 | 22 | + |
| N1-N8-Diace | N15C13-Glyc | 4.87 | 230.5 | 100.2 | 19 | + |
| N1-N8-Diace | IS | 4.87 | 236.2 | 103.2 | 19 | + |
| N15C13-Glyc | IS | 5.98 | 78.04 | 32 | 25 | + |
| N6,N6,N6-Tr | Arginine-d4C | 6.53 | 189.16 | 84.08 | 20 | + |
| NAD | 6C13-Phenyla | 7 | 664.1 | 428.2 | 24 | + |
| Niacinamide | Niacinamide | 1.79 | 123.06 | 80 | 24 | + |
| Niacinamide | NA | 1.7 | 127 | 84 | 24 | + |
| Nicotinamide | Valine-d8 | 6.6 | 335.2 | 123.1 | 30 | + |
| Nicotinamide | Alanine-d4 | 5.3 | 255 | 123 | 30 | + |
| Nicotinic acid | Niacinamide | 1.7 | 124.1 | 78 | 30 | + |
| Nitrotyrosine | 6C13-Phenyla | 4.78 | 227.1 | 210 | 12 | + |
| Nonanoylcarn | Isovalerylcarn | 3.3 | 302.2 | 85 | 30 | + |
| Norepinephri | 6C13-Phenyla | 5.1 | 152 | 107 | 15 | + |
| Octanoylcarr | Octanoylcarr | 3.31 | 288.2 | 85 | 33 | + |
| Octanoylcarr | N/A | 3.31 | 291.2 | 85 | 33 | + |
| Ornithine | Ornithine-d2 | 6.89 | 133 | 70 | 15 | + |
| Ornithine-d2 | N/A | 6.89 | 135.1 | 117.2 | 15 | + |
| Palmitoylcarn | Palmitoylcarn | 3 | 400 | 85 | 40 | + |
| Palmitoylcarn | N/A | 3 | 403.35 | 85 | 40 | + |
| Paraxanthine | Niacinamide | 1.7 | 181.1 | 124 | 24 | + |
| Phenylalanin | 6C13-Phenyla | 4.8 | 166.09 | 120.08 | 18 | + |
| Piperanine | Niacinamide | 1.34 | 288.2 | 135.2 | 33 | + |
| Piperine | Niacinamide 1.1 | | 286.1 | 201.1 | 20 | + |

| | | | | | | |
|---|---|---|---|---|---|---|
| Pipernonalin | Niacinamide | 1.3 | 342.1 | 229.2 | 23 | + |
| Piperyline | Niacinamide | 1.39 | 272.2 | 201.1 | 28 | + |
| Proline | 6C13-Tyrosin | 5.4 | 116 | 70.1 | 20 | + |
| Propionylcarı | Propionylcarı | 3.9 | 218.1 | 85 | 20 | + |
| Propionylcarı | NA | 3.93 | 221.15 | 85 | 20 | + |
| Putrescine | N15C13-Glyc | 6.3 | 89 | 72 | 20 | + |
| Pyridoxamine | Alanine-d4 | 6.16 | 169 | 152 | 20 | + |
| Pyroglutamic | Octanoylcarr | 3.42 | 130 | 84 | 15 | + |
| S-Adenosylho | N15C13-Glyc | 6.22 | 385.1 | 136 | 21 | + |
| S-Adenosylm | Arginine-d4C | 6.98 | 399 | 250 | 20 | + |
| Sarcosine | Alanine-d4 | 5.65 | 90.04 | 44.1 | 20 | + |
| Serine | N15C13-Glyc | 6.19 | 106 | 60 | 16 | + |
| Serotonin | 15N4-Inosine | 4.3 | 177 | 160 | 15 | + |
| Spermidine | Arginine-d4C | 7.8 | 146.16 | 72 | 17 | + |
| Spermine | 6C13-Phenyla | 4.95 | 203.1 | 129.1 | 15 | + |
| Stearoylcarni | Isovalerylcarı | 2.98 | 428.4 | 85 | 35 | + |
| Sucrose | 6C13-Phenyla | 6.01 | 364.8 | 202.8 | 35 | + |
| Taurine | 6C13-Tyrosin | 5.3 | 126.02 | 108 | 15 | + |
| Tetrahydrocu | 6C13-Phenyla | 1.38 | 355 | 137.1 | 24 | + |
| Theobromine | Niacinamide | 2.1 | 181 | 138.3 | 25 | + |
| Theophylline | Niacinamide | 1.62 | 181.13 | 124 | 20 | + |
| Thiamine | 13C3-Thiami | 5.15 | 265 | 122 | 25 | + |
| Threonine | N15C13-Glyc | 5.9 | 120 | 74 | 13 | + |
| Trigonelline | Methionine-( | 5.2 | 138.1 | 94.1 | 20 | + |
| Trimethylam | IS | 4.1 | 85 | 66 | 20 | + |
| Trimethylam | Trimethylam | 4.1 | 76.08 | 58.07 | 20 | + |
| Tryptophan | 6C13-Phenyla | 4.81 | 205.01 | 146.06 | 20 | + |
| Tyrosine | 6C13-Phenyla | 5.31 | 182.1 | 136 | 15 | + |
| Uracil | Palmitoylcarı | 2.44 | 113 | 70 | 23 | + |
| Ureidopropic | Palmitoylcarı | 3.15 | 133 | 115 | 20 | + |
| Uric acid | 1,3-15N2-Uri | 5.22 | 169.1 | 141 | 20 | + |
| Uridine | 13C5-Hypoxa | 3.81 | 245 | 113 | 18 | + |
| Valerobetain | Valerobetain | 3.98 | 160 | 101.1 | 21 | + |
| Valerobetain | IS | 4.2 | 169.13 | 101.1 | 21 | + |
| Valine | 6C13-Phenyla | 5.3 | 118.09 | 55 | 20 | + |
| Valine-d8 | N/A | 5.3 | 126 | 80.1 | 20 | + |
| Xanthine | 15N4-Inosine | 4.15 | 153 | 110 | 20 | + |
| Xanthosine | 15N4-Inosine | 4.6 | 285 | 153 | 20 | + |

| Metabolite | Internal Std | RT(XEVO) | Q1 m/z | Q3 m/z | CE(XEVO) | Ionization mo |
|---|---|---|---|---|---|---|
| 13C3-Citric ac | N/A | 3.8 | 194.2 | 113 | 8 | - |
| 13C4-Fumaric | N/A | 3.3 | 119.2 | 74.3 | 11 | - |
| 13C4-Malic ac | N/A | 3.3 | 137.2 | 119.2 | 8 | - |
| 13C4-Succini | N/A | 3.3 | 121.3 | 76 | 8 | - |
| 13C5-Oxoglut | N/A | 3 | 150.2 | 105.3 | 8 | - |
| 13C6-Benzoic | N/A | 1 | 127 | 83 | 14 | - |
| 13C8-Octanic | N/A | 1.4 | 151.1 | 151.1 | 3 | - |
| 15N5-ADP | N/A | 3.6 | 431.1 | 79 | 40 | - |
| 2-Hydroxyglut | Glutamic acid | 3 | 145 | 101 | 10 | - |
| 2-Phosphogly | Glutamic acid | 3.6 | 185.5 | 97 | 25 | - |
| 3-Hydroxybut | Glutamic acid | 0.87 | 103.2 | 59.2 | 10 | - |
| 3-Phosphogly | Glutamic acid | 3.6 | 185.5 | 97.1 | 25 | - |
| 6-Phosphoglu | Glutamic acid | 3.8 | 275 | 97 | 15 | - |
| Acetoacetic a | Glutamic acid | tune 1 | 101 | 101 | 3 | - |
| Adenosine mo | Glutamic acid | 3.3 | 346 | 79 | 30 | - |
| Adenosine tri | 13C4-Succini | 3.7 | 506.1 | 158.9 | 23 | - |
| Adenosine tri | N/A | 3.7 | 510 | 158.9 | 23 | - |
| ADP | 13C4-Malic ac | 3.5 | 426.1 | 79 | 40 | - |
| Alanine | Alanine-d4 | 2.4 | 88 | 88 | 7 | - |
| Alanine-d4 | N/A | 2.4 | 92 | 92 | 7 | - |
| Arginine | Arginine-d4C | 4 | 173 | 131 | 10 | - |
| Arginine-d4C | N/A | 4 | 178 | 136 | 10 | - |
| Asparagine | Glutamic acid | 2.7 | 131 | 114 | 7 | - |
| Aspartic acid | Aspartic acid- | 3.1 | 132 | 88 | 13 | - |
| Aspartic acid- | N/A | 3.1 | 135 | 91 | 13 | - |
| cis-Aconitic ac | Glutamic acid | 3.4 | 173 | 129 | 12 | - |
| Citric acid | Glutamic acid | 3.7 | 191.2 | 111 | 8 | - |
| Dihydroxyace | Glutamic acid | 3.5 | 169.1 | 79 | 7 | - |
| Erythose 4-ph | Glutamic acid | 3.5 | 199 | 97 | 15 | - |
| Fructose 1,6-l | Glutamic acid | 4.1 | 339 | 97 | 15 | - |
| Fructose 6-ph | Glutamic acid | 3.7 | 259 | 97.1 | 15 | - |
| Fumaric acid | 13C4-Fumaric | 3.3 | 115.2 | 71.3 | 11 | - |
| Glucose 1-ph | 13C8-Octanic | 3.7 | 259 | 241 | 15 | - |
| Glucose 6-ph | 13C8-Octanic | 3.7 | 259 | 97 | 15 | - |
| Glutamic acid | Glutamic acid | 3.2 | 146 | 128 | 7 | - |
| Glutamic acid | Glutamic acid | 3.2 | 149 | 131 | 7 | - |
| Glutamine | | 2.7 | 145 | 127 | 17 | - |
| Glyceraldehy | Glutamic acid | 3.5 | 169.1 | 97 | 7 | - |
| Isocitric acid | Glutamic acid | 3.7 | 191.2 | 111.1 | 8 | - |
| Lactic acid | Glutamic acid | 1.2 | 89.2 | 43.2 | 13 | - |
| Malic acid | 13C4-Malic ac | 3.3 | 133.2 | 115.2 | 8 | - |
| NAD | | 3.2 | 662.1 | 540.3 | 12 | - |
| NADH | | 2.99 | 664.1 | 408.1 | 11 | - |
| NADP | | 3.92 | 742.1 | 620 | 20 | - |

| | | | | | | |
|---|---|---|---|---|---|---|
| NADPH | | 3.75 | 744.1 | 408 | 11 | - |
| Oxalacetic ac | 13C5-Oxoglut | 3.63 | 131 | 131 | 7 | - |
| Oxoglutaric a | Glutamic acid | 3 | 145.2 | 101.3 | 8 | - |
| Phosphocrea | Glutamic acid | 3.5 | 210 | 79 | 15 | - |
| Phosphoenol | Glutamic acid | 3.8 | 167 | 79 | 29 | - |
| Pyruvic acid | U-13C3-Pyruv | 0.7 | 87.2 | 43.2 | 14 | - |
| Ribose 5-pho | Glutamic acid | 3.4 | 229 | 97 | 15 | - |
| Ribulose 5-ph | Glutamic acid | 3.4 | 229 | 97.1 | 15 | - |
| Sedoheptulos | Glutamic acid | 3.6 | 289 | 97 | 15 | - |
| Serine | Glutamic acid | 2.7 | 104 | 74 | 8 | - |
| Succinic acid | 13C4-Succini | 3.3 | 117.2 | 73.2 | 8 | - |
| U-13C10,U-15 | N/A | 3.4 | 361 | 79 | 30 | - |
| U-13C3-Pyruv | N/A | 0.7 | 90.2 | 45.2 | 14 | - |
| U-13C6-Gluco | N/A | 3.7 | 265 | 97 | 15 | - |
| Xylulose 5-ph | Glutamic acid | 3.4 | 229.1 | 97 | 15 | - |

**ɔde**
